# *In silico* Identification and Functional Characterization of Conserved miRNAs in Fibre Biogenesis Crop *Corchorus capsularis*

**DOI:** 10.1101/2020.04.22.056176

**Authors:** Mahmudul Hasan, Milad Ahmed, Foeaz Ahmed, Jamil Ahmed, Mst Rubaiat Nazneen Akhand, Kazi Faizul Azim, Md. Abdus Shukur Imran, Syeda Farjana Hoque

## Abstract

Corchorus capsularis, commonly known as jute occupies the leading position in the production of natural fibre and fibre based products alongside lower environmental threat. Nowadays, the study of lignin biosynthesis pathways with other molecular basis of fibres formation are being more focused for its economic perspective. Small noncoding ∼21 to 24 nt nucleotides long microRNAs play significant roles in regulating the gene expression as well as different functions in cellular growth and development. Here, the study adopted a comprehensive in silico approach to identify and characterize the conserved miRNAs in the genome of C. capsularis including specific gene targets involved in the crucial cellular process. Expressed Sequence Tags (ESTs) based homology search of 3350 known miRNAs of dicotyledons were allowed against 763 non-redundant ESTs of jute genome resulted in the prediction of 5 potential miRNA candidates belonging five different miRNA families (miR1536, miR9567-3p, miR4391, miR11300, and miR8689). The putative miRNAs were 18 nucleotide length, within a range of -0.49 to -1.56 MFEI values and 55% to 61% of (A+U) content of their correspondence pre-miRNAs. A total of 1052 gene targets of putative miRNAs were identified and their functions were extensively analyzed. Most of the gene targets were involved in plant growth, cell cycle regulation, organelle synthesis, developmental process and environmental responses. The five gene targets, namely, NAC Domain Containing Protein, WRKY DNA binding protein, 3-dehydroquinate synthase, S-adenosyl-L-Met–dependent methyl transferase and Vascular-related NAC-Domain were found to be involved in the lignin biosynthesis, phenylpropanoid pathways and secondary wall formation which could play significant roles in the overall fibre biogenesis. The characterization of conserved miRNAs and their functional annotation of specific gene targets might enhance the more miRNA discovery, strengthening the complete understanding of miRNAs association in the cellular basis of lignin biosynthesis towards the production of high standard jute products.

## 1. Introduction

Jute, natural fibre is the blessing of future economy on account of raising awareness with environmental and SDG (Sustainable Development Goal) issues around all over the world [1,2,3].

It is now placed in the second and only after cotton in case of economic importance among the natural fibers and over 95% of the global jute production is served by India, Bangladesh, China, Nepal and Thailand [4]. Jute products are more environmentally sound and their disposal stage is also less harmful to environment than its competing synthetic materials [5]. Moreover, the natural fibre products are likely to be the favorable for combating against global worming rather [6]. Moreover, on an average farm value of ∼US$2.3 billion is being generated by the jute products annually [7]. Now-a-days, scientists are more likely to be interested in developing environment-friendly and cost-effective polymer matrix composites from natural resources as implementation of new laws and raising pressure from environmental activists [8,9,10,11]. Recently, genome sequencing of different bast (phloem) fibres plants such as Jute (*Corchorus sp*.) and kenaf (*Hibiscus cannabinus L*.) has been completed focusing on the analysis of key regulatory and structural genes involved in fibre biogenesis [12,13].

Jute (*Corchorus sp*.) covers about 80% of worldwide bast fibre production [12]. Among more than 100 of *Corchorus* species in the Malvaceae family, only *Corchorus olitorius* and *Corchorus capsularis* are commercially cultivated [12,14]. Lignocellulosic bast fibres are extracted from *Corchorus capsularis* for commercial purposes in the different industrial uses and textile sectors [15,16]. Usually jute fibre is composed of 61 % cellulose, 15.9 % hemicelluloses and 13.5 % lignin [17], but in some cases it could contains as high as 15–22 % of lignin [18]. The high content of lignin gives a unique xylan-type fibre properties of jute fibres which are distinguished from other gelatinous-type bast fibres of flax, hemp and ramie [19]. Jute fibre lignin seems the polymerization of three hydroxycinnamyl alcohols (monolignols), viz., p-coumaryl alcohol, coniferyl alcohol and sinapyl alcohol resulting in p-hydroxyphenyl, guaiacyl and syringyl units respectively [20]. However, this high level of lignin content causes several complexities in the formation of jute fibre and its valued product processing. For examples, photo-yellowing of jute products could be happened due to the α-carbonyl group of the lignin structure which reacts with UV radiation. Besides, high lignin content also cause complexity in jute fibre separation as well as maintaining standard quality [21]. The above mentioned features has made the jute fibres less suitable for making divers valued finer fabrics and other standard products, and it mostly happens in the fibres from *Corchorus capsularis*. So, the development of low-lignin fibres has become a priority needs in jute industries as well as global market [15, 22].

Different studies reported several metabolic pathways as well as genes involved in the key processes of fibres formation [23,24]. But, there are yet limited studies in the molecular basis and regulatory process of lignin biosynthesis in xylan type bast fibres [25]. However, the availability of genomic data and comparative genome based analysis could allow the study of gene regulation and molecular interaction of different pathways involved in complex fibre formation process. MicroRNAs (miRNAs), small regulatory RNAs consisting of about 18–22 nucleotides could play an significant roles in different cellular functions in plants and animals [26]. It was found that miRNA might regulate gene expression at the different stages of posttranscriptional levels by interacting to specific targets for initiating mRNA inhibition or cleavage of mRNA translation [27,28]. Though, there are enormous possibilities to work with miRNA for controlling any specific biological process but still limited number of miRNAs has been identified in plants by experimental and computational approaches [29,30]. As miRNAs are highly evolutionarily conserved in the plant and animal kingdom, and that is why the nature of evolutionary conservation in the miRNAs allows an effective approach to the identification and characterization of newly conserved miRNAs using comparative genomics approach [30,31,32,33].

Though the plethora of economic importance in the view of trading and commercial purposes, there has been no report on miRNAs in *Corchorus capsularis*. In this study, we employed an EST based homology search to identify miRNAs in jute using currently available 826 expressed sequence tags (ESTs) from the NCBI Genbank database. To avoid the difficulties of searching from genomic sequence surveys or unavailability of whole genome sequences, EST based miRNA identification by using specialized software has already been accepted successfully over the other approaches [34,35,36,37,38]. The present study was employed with the Identification and Characterization of conserved microRNAs in the genome of fibre biogenesis crop jute by using EST based comparative genomics approach.

## 2. Methods

### 2.1 Acquisition of EST Sequences and Reference miRNAs

The available EST sequences of *Corchorus capsularis* were retrieved from the recently deposited genome (GenBank assembly accessionGCA_001974805.1) in the GenBank databases of NCBI(http://www.ncbi.nlm.nih.gov/) [12,39]. Moreover, to search potential miRNAs in *C. capsularis*, all previously known mature miRNA sequences of plant dicotyledons were extracted from miRBase database (http://www.mirbase.org/). These miRNAs were employed as the reference set with the retrieved ESTs of *C. capsularis* genome for identifying conserved miRNAs family in Jute. To avoid redundant or overlapping of miRNAs and ESTs, the repeated sequences of miRNAs and ESTs were removed by CD-HIT [40] and only the single ones were retained for further study.

### 2.2 Search for potential miRNAs in the jute genome

The reference miRNAs retrieved from miRBase were used as the query templates for homology search against the ESTs of *Corchorus capsularis* by using BLASTn of NCBI database (https://blast.ncbi.nlm.nih.gov/Blast.cgi) considering all default parameters. Only the top EST hit for each miRNA search was taken. Again, all of ESTs from BALSTn search were screened by CD-HIT for redundancy check.

### 2.3 Screening for the non-coding miRNA candidates

As the miRNAs are non-protein coding genes, the precursor miRNAs should also be non-protein coding. After redundancy check, the top hits from BLASTn were subjected to BLASTx (https://blast.ncbi.nlm.nih.gov/Blast.cgi) analysis for removing the protein-coding sequence. A number of EST sequences were found that were non protein coding. These miRNA candidates were then used for further screening to identify pre-miRNA by evaluating the miRNA precursor prediction properties [41].

### 2.4 Identification of Novel miRNAs with Reported Criteria

The different criteria were used to screen precursor miRNAs based on statistical comparative genomics approaches reported by Zhang et al., 2006 [42], which had already been adopted in many ESTs targeted miRNAs identification and characterization studies [43,44,45]. Here, only candidate ESTs sequences adopted the following criteria were prioritized as considerable miRNAs in *C. capsularis*: (1) not more than 3 mismatches were allowed in the putative miRNAS with all previously known plant miRNAs; (2) the length of predicted mature miRNAs should be in the range of 18 nucleotides; (3) pre-miRNA sequence must fold into an appropriate hairpin secondary structure and mature miRNA should be located in one arm of the hairpin stem–loop secondary structure; (4) the mature miRNAs should allow less than 3 mismatches with the opposite miRNA strand in the other arm; (5) A+U content of pre-miRNA must be in range of 30% to 70%,; and (6) the secondary structure of mature miRNAs must have lower minimal (highly negative) folding free energy (MFE) and minimal free energy index (MFEI) value [46,47,48].

### 2.5 Prediction of pre-miRNA and Hair-loop Secondary Structure of mature miRNAs

The ESTs for putative miRNA prediction were subjected to mirEval (https://tools4mirs.org/) for determining the pre-miRN candidates [49]. The minimal length of the pre-miRNA was set as 85 nucleotide and these precursor sequences of potential miRNA homologs were assessed for secondary structures prediction using the Zuker folding algorithm [50] in the Mfold web server (http://unafold.rna.albany.edu/?q=mfold). As the predicted secondary structure should have the higher minimal negative value of minimal folding free energy index (MFEI) and minimal folding free energy (MFE), only the best fitted pre-miRNA candidates were obtained considering all parameters. The MFEI was calculated using the following equation;

Adjusted MFE (AMFE) represented the MFE of 100 nucleotides.

AMFE= (MFE ÷ length of RNA sequence) * 100 MFEI = AMFE/(G + C)%

MFEI = [(MFE/length of the RNA sequence)×100]/(G+C) %

### 2.6 Nomenclature and family annotation of predicted microRNAs

The putative miRNAs were used as query sequence for local blast in miRBase to reconfirm the family of the putative mature miRNAs. The predicted microRNAs were named by a nomenclature followed by miRBase [51]. Sequence alignment of homologous members of predicted miRNA family was investigated in the miRBase search output and rechecked by Clustal Omega [52,53]. The percentage of each nucleotide base of putative mature miRNAs and their respective homologue miRNAs from miRBase were also calculated and analyzed.

### 2.7 Prediction and functional analysis of putative miRNA targets

Due to the unavailability of *C. capsularis* resources regarding miRNA targeted genes, *Arabidopsis* was considered as the reference organism for prediction and functional annotation of the newly identified mature miRNAs in jute genome. The predicted mature miRNAs were employed as query against the *A. thaliana* by using psRNATarget database (http://plantgrn.noble.org/psRNATarget/) with following parameters: (1) maximum expectation value should be 5 ; (2) complementarity scoring (HSP) size should be kept within the length of 18; (2) target sites’ multiplicity must be allowed to 2; (3) range of central mismatch for translational inhibition should be in the range of 10–11 nucleotide; (4) highest level of mismatches at the complementary site 4 except any gaps. The psRNATarget ensures reverse complementary matching between targets transcripts and respective miRNAs, leading to the target site accessibility by calculating unpaired energy (UPE) needed for beginning the secondary structure around the miRNA target site [54]. The schematic outline of the total methodology was illustrated in Figure 1.

**Figure 1:**
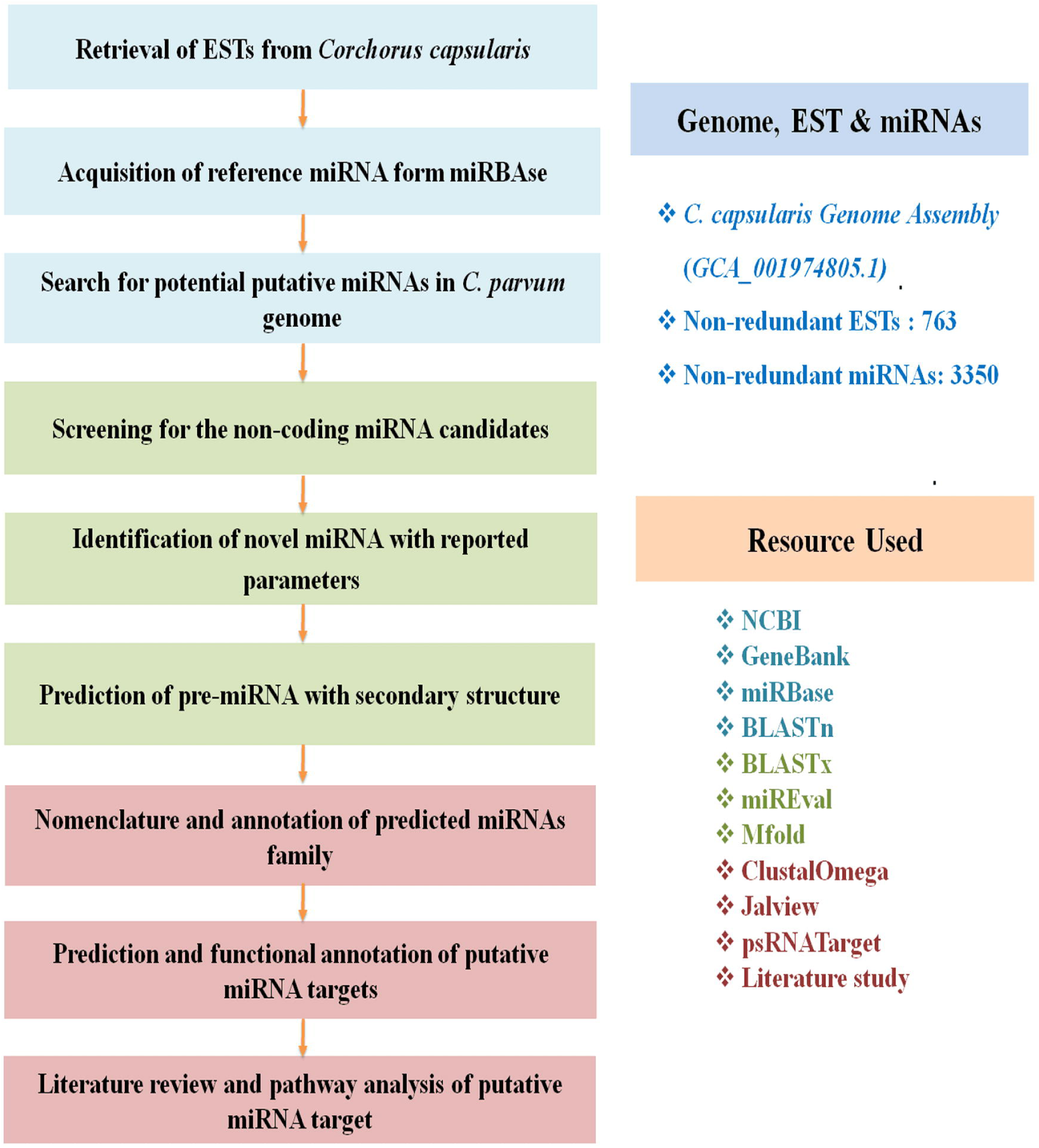
The schematic outline of the total methodology.

## 3. Results

### 3.1 Search for potential miRNAs in genome of *C. capsularis*

To identify and characterize the conserved microRNA of *C. capsularis*, a comprehensive EST based study was employed. There were different *in silico* steps where a huge volume of preliminary ESTs of *C. capsularis* were screened to finalize the putative miRNAs with specific gene targets involved in the biological process of jute alongside biogenesis of fibres. The summery of details process with the numerical data was described in the Table 1. The previously known mature miRNAs under the dicotyledons, belonging to twenty plants families (Amaranthaceae, Araliaceae, Asteraceae, Cucurbitaceae, Euphorbiaceae, Fabaceae, Caricaceae, Lamiales, Linaceae, Malvaceae, Myrtaceae, Paeoniaceae, Brassicaceae, Ranunculaceae, Rhizophoraceae, Rosaceae, Rutaceae Salicaceae, Solanaceae, Vitaceae) were extracted from the microRNA database. In this approach, a total of 6386 of plant miRNA (Supplementary File 1) were retrieved. About 826 ESTs of *Corchorus capsularis* (Supplementary File 2) were retrieved from the GenBank databases of NCBI. Redundant miRNAs and ESTs were removed, and only the unique ones were retained. About 3036 redundant miRNA sequences were removed, retaining 3350 miRNAs containing no repeated sequences were kept for the further investigation (Supplementary File 3). On the other hand, 763 non-redundant ESTs of jute were kept from 826 ESTs (Supplementary File 4). Here, 763 non-redundant Jute ESTs were searched against 3350 non-redundant mature miRNA sequences of all dicotyledons. After the removal of redundancy by CDHIT, around 621 ESTs were selected as potential miRNA sequences for *Corchorus capsularis* (Supplementary File 5). Moreover, It had been found that around 36 ESTs were non-coding which could be the potential candidates of miRNA homologs (Supplementary File 6).

**Table 1:**
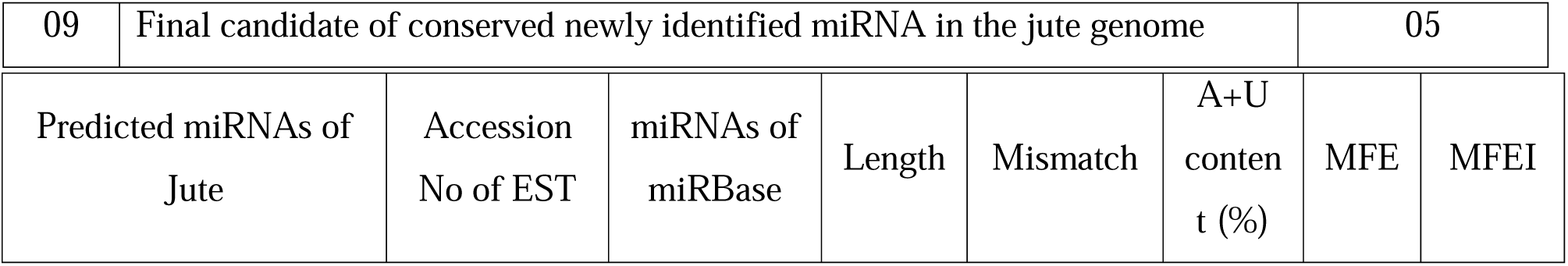
Steps involved in the characterization of putative miRNA from ESTs of jute genome

### 3.2 Prediction of pre-miRNAs and Hair-loop Secondary Structure of mature miRNAs

About the 36 non-coding ESTs were investigated to identify the potential miRNA precursors by evaluating the miRNA precursor prediction properties using mirEval (https://tools4mirs.org/). The different preset criteria were strictly followed in each steps to screen out the putative miRNAs in *Corchorus capsularis*. Here, five ESTs were considered as pre-miRNA by careful evaluation (Table 2 and Supplementary Table 1), while the other 31 ESTs (Supplementary File 7) were filtered out as they failed to fulfill the criteria mentioned in the Materials and Methods section. The final five precursor sequences of potential miRNAs were assessed for confirming their ability to form secondary structures using the Zuker folding algorithm of MFOLD software (Figure 2). It was observed that all five pre-miRNAs were capable of folding themselves into an appropriate hairpin structure as predicted by mirEval. There was also minimal loop involvement in the hairpin structures of the miRNAs. The four mature miRNAs were incorporated in only one loop of their respective hairpin structure while the other one was in a single-arm without involving any loop (Figure 2). The minimal folding free energy (MFE, *dG* in kcal/mol) of all the pre-miRNAs were ranged between -59.60 and -14.90 while the percentage of A+U contents of the precursors were ranged between 55% and 65% (Table 2).

**Table 2:**
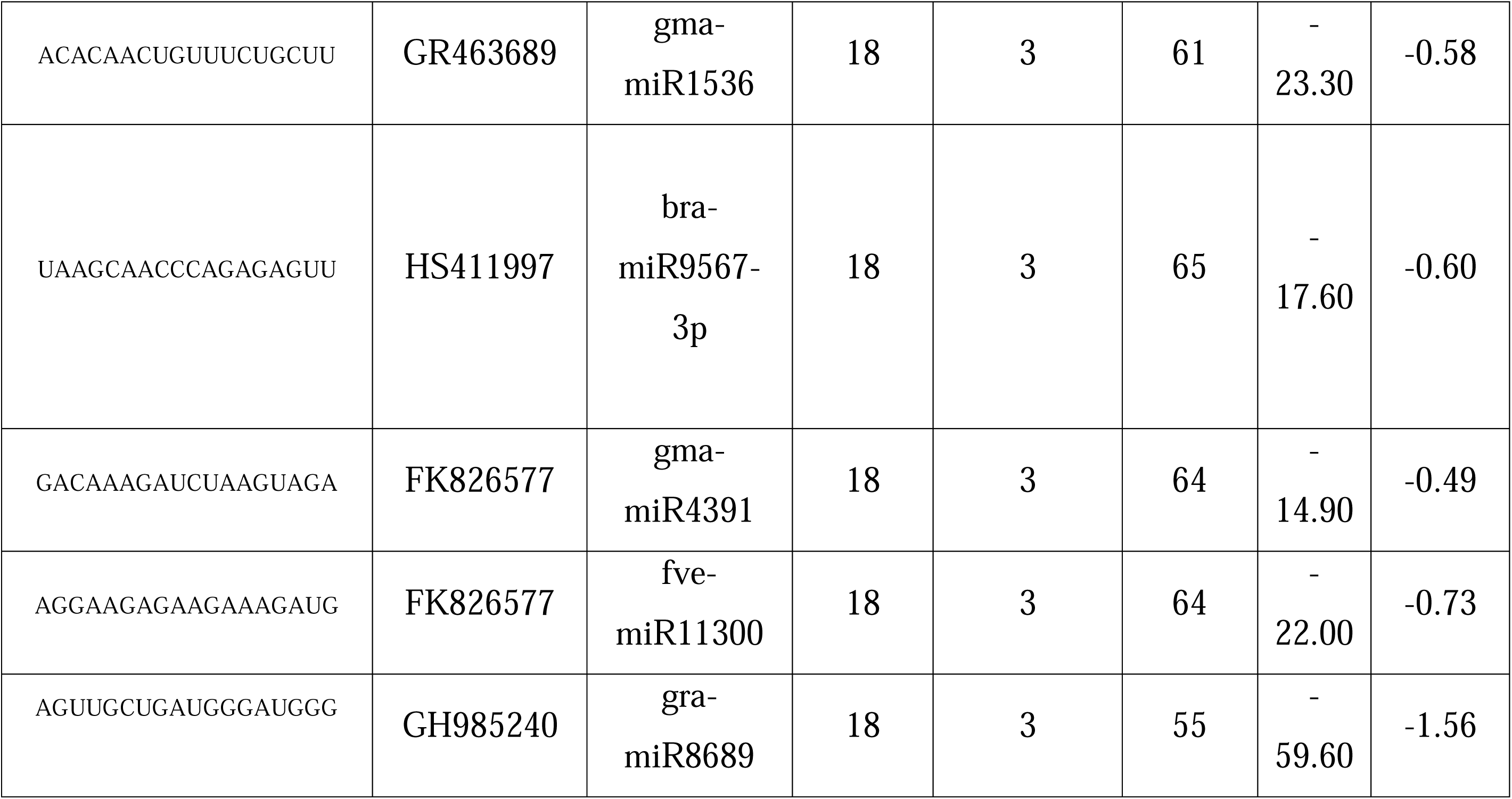
Putative miRNA in the jute genome with suggested criteria

**Figure 2:**
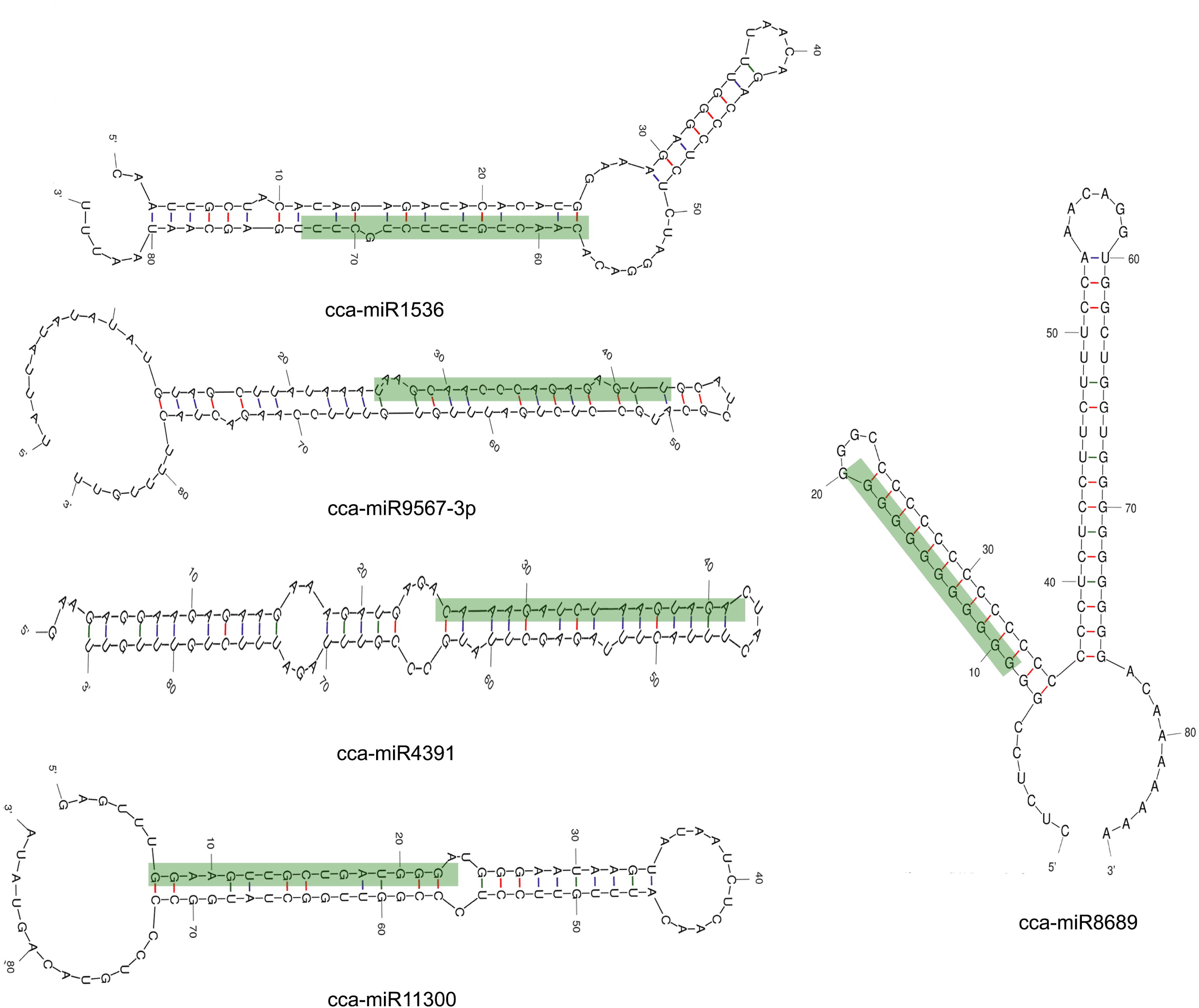
Secondary structures of final putative miRNA by using the Zuker folding algorithm of MFOLD software.

### 3.3 Nomenclature and family annotation of predicted microRNAs

The miRBase database was used to search out and address the microRNAs family of the predicted five putative miRNAs. It had been found that the five miRNAs of jute belong to five different miRNA families (miR1536, miR9567-3p, miR4391, miR11300 and miR8689) and the nomenclature of the predicted miRNAs were done as cca-miR1536, cca-miR9567-3p, cca-miR4391, cca-miR11300, and cca-miR8689. Sequence alignment of putative miRNAs of jute genome and its homologous member from miRBase were visualized in Figure 3. The average each nucleotide base percentage of cca-miR1536, cca-miR9567-3p, cca-miR4391, cca-miR11300, and cca-miR8689 were calculated, and compared with the each nucleotide base percentage of gma-miR1536, bra-miR9567-3p, gma-miR4391, fve-miR11300 and gra-miR8689 respectively. It had been investigated that the percentage of adenine, uracil, guanine and cytosine of both putative mature miRNA of jute and their correspondence reference miRNA were almost in the similar quantity. In addition, the adenine percentage was found higher in the miRNA family of miR9567-3p and miR4391 for whereas other two families, namely, miR1536 and miR8689 showed higher percentages in case of uracil and guanine respectively (Figure 4).

**Figure 3:**
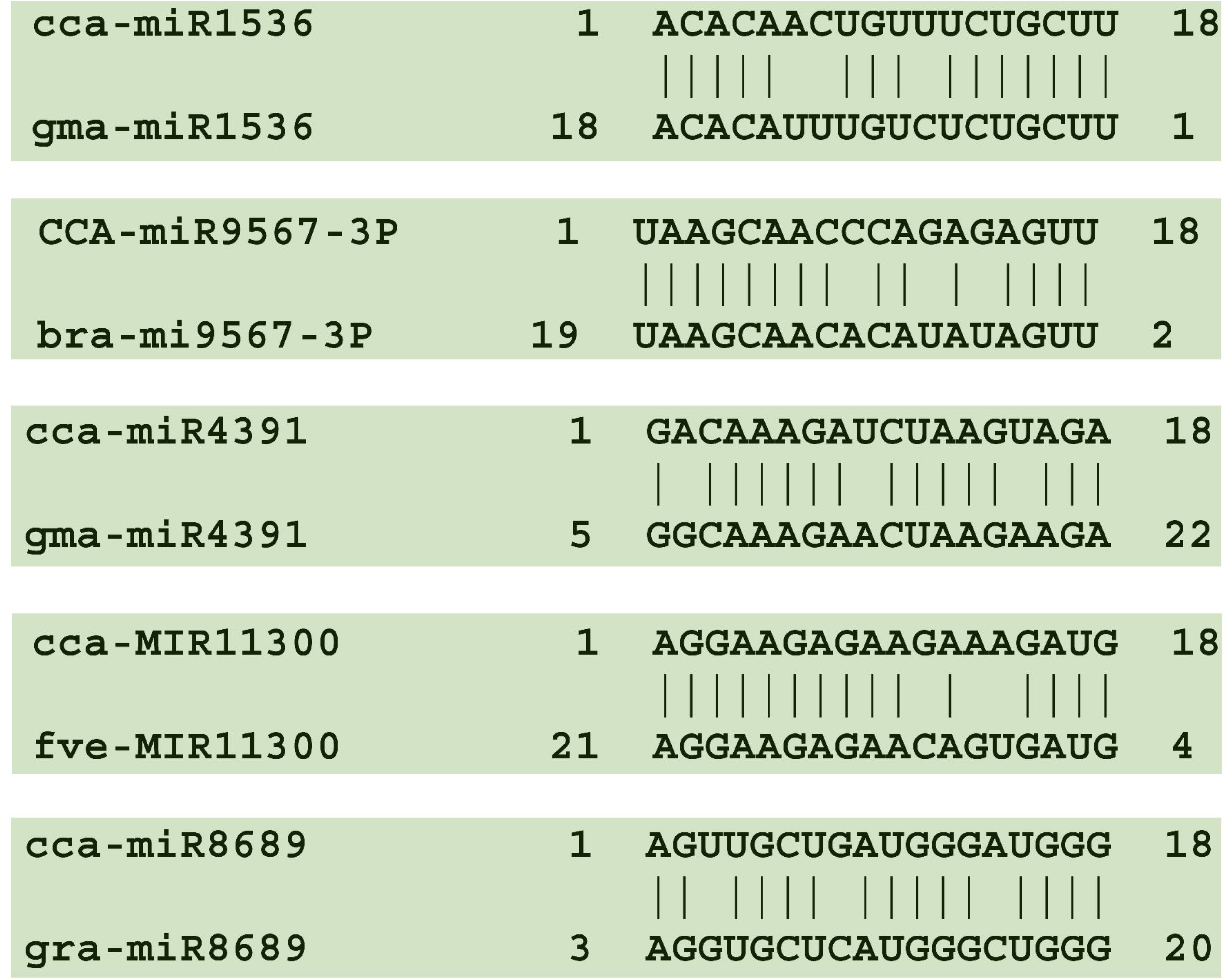
Alignments of the putative miRNAs of jute genome and its homologues from respective microRNA family.

**Figure 4:**
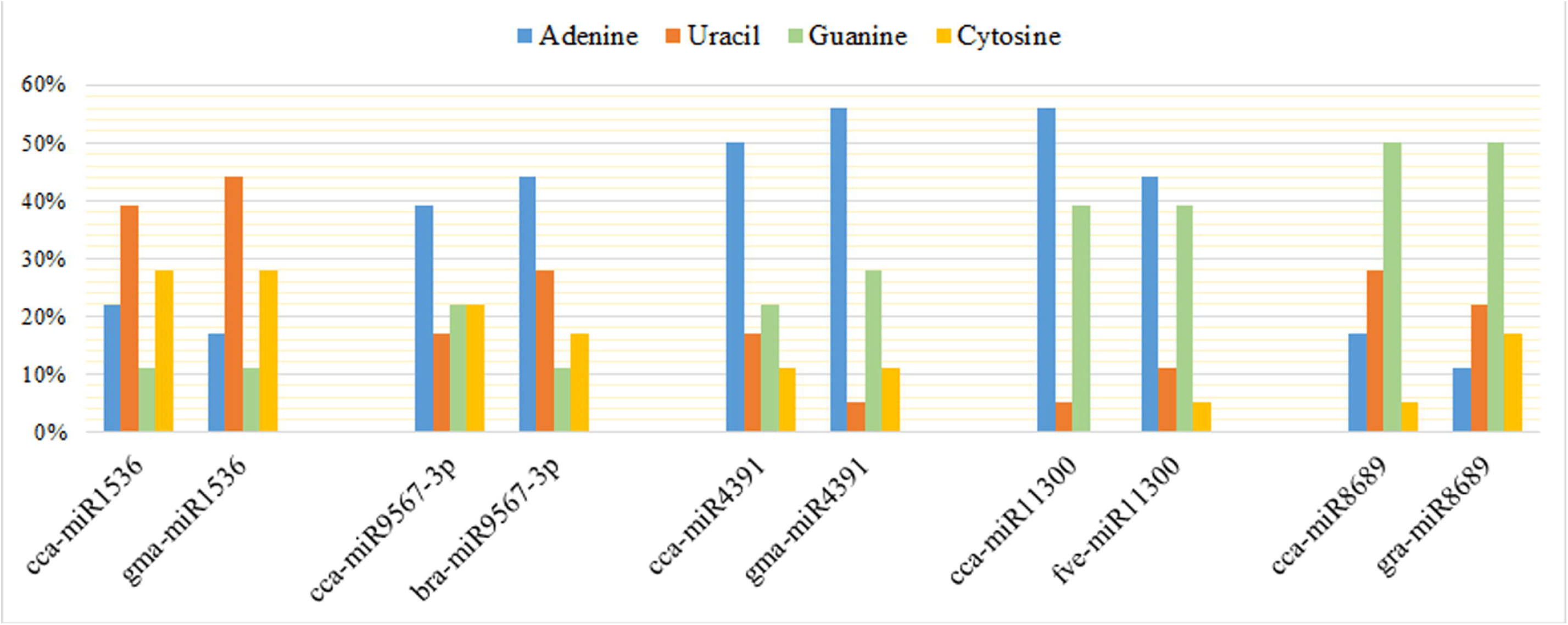
Overall nucleotide compositions (%) of putative miRNA from *Corchorus capsularis* compared with the homologues mRNA of miRBase.

### 3.4 Prediction and functional analysis of putative miRNA targets

The five novel jute miRNAs were found to be involved with a total of 1052 targets as predicted by the psRNATarget (Supplementary Table 2). Among the around 1052 gene targets, 147 was found to be uncharacterized. All of the gene targets were cross checked where four gene targets were found as the common target for the five putative miRNAs, and those targets were also highly abundant in the target pool (‘Leucine-rich repeat family protein; 13 times’, ‘Transposable element gene; 87 times’, ‘Protein kinase superfamily protein; 17 times’, ‘Pentatricopeptide repeat superfamily protein; 9 times’). Comprehensive literature studies were employed to find out the involvement of predicted gene targets in different biological process. There were few gene targets which were found to be directly or indirectly involved with the lignin biosynthesis and secondary wall biogenesis (Table 3) [55-68]. The functions of the common gene targets were also described in the Table 3 [69-76]. Moreover, there were three gene targets (alpha/beta-Hydrolases superfamily protein, RING/U-box superfamily protein and P-loop containing nucleoside triphosphate hydrolases superfamily protein coding genes) were found to be involved with four putative miRNAs of jute (Supplementary Table 2).

**Table 3:**
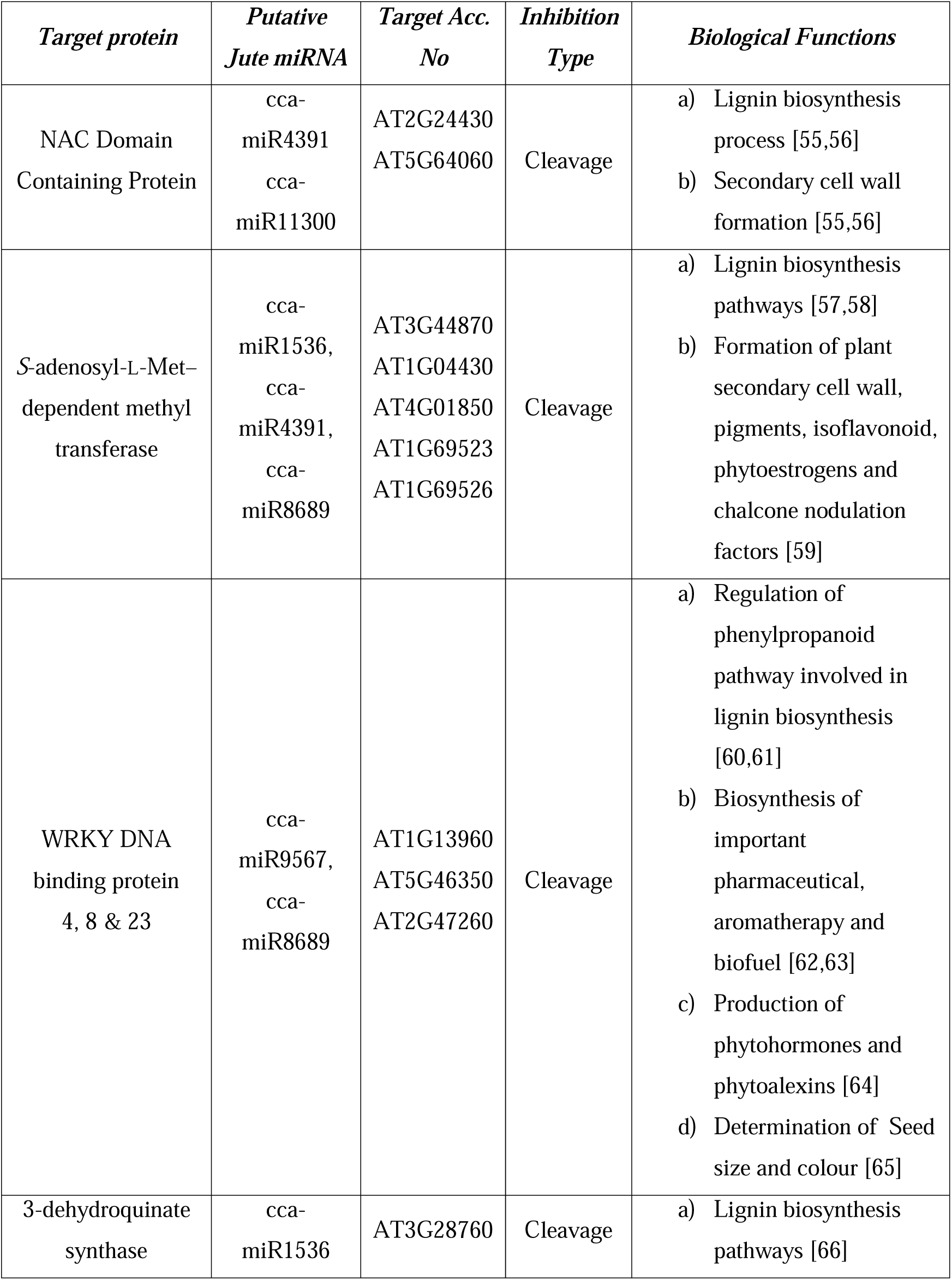
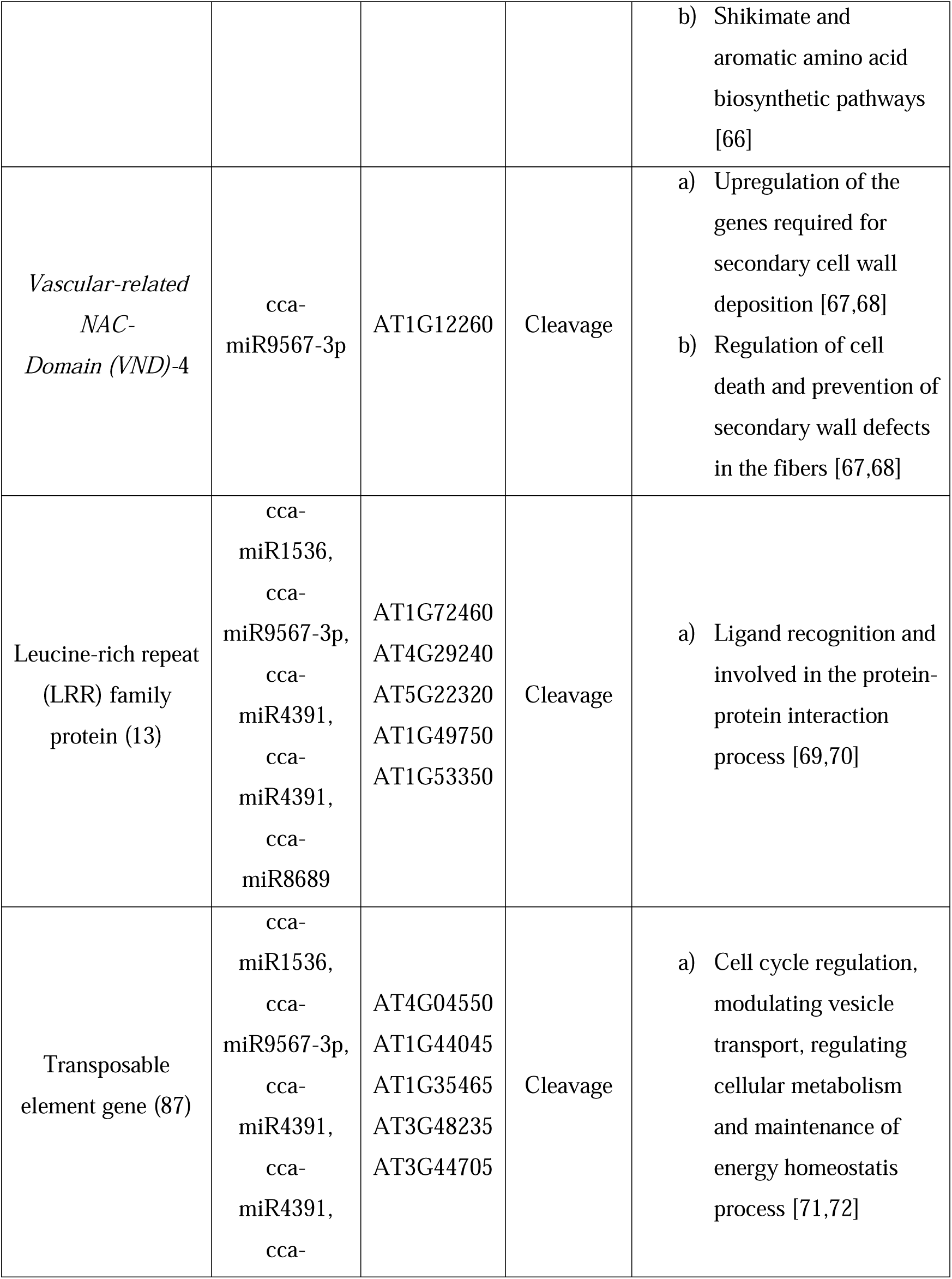
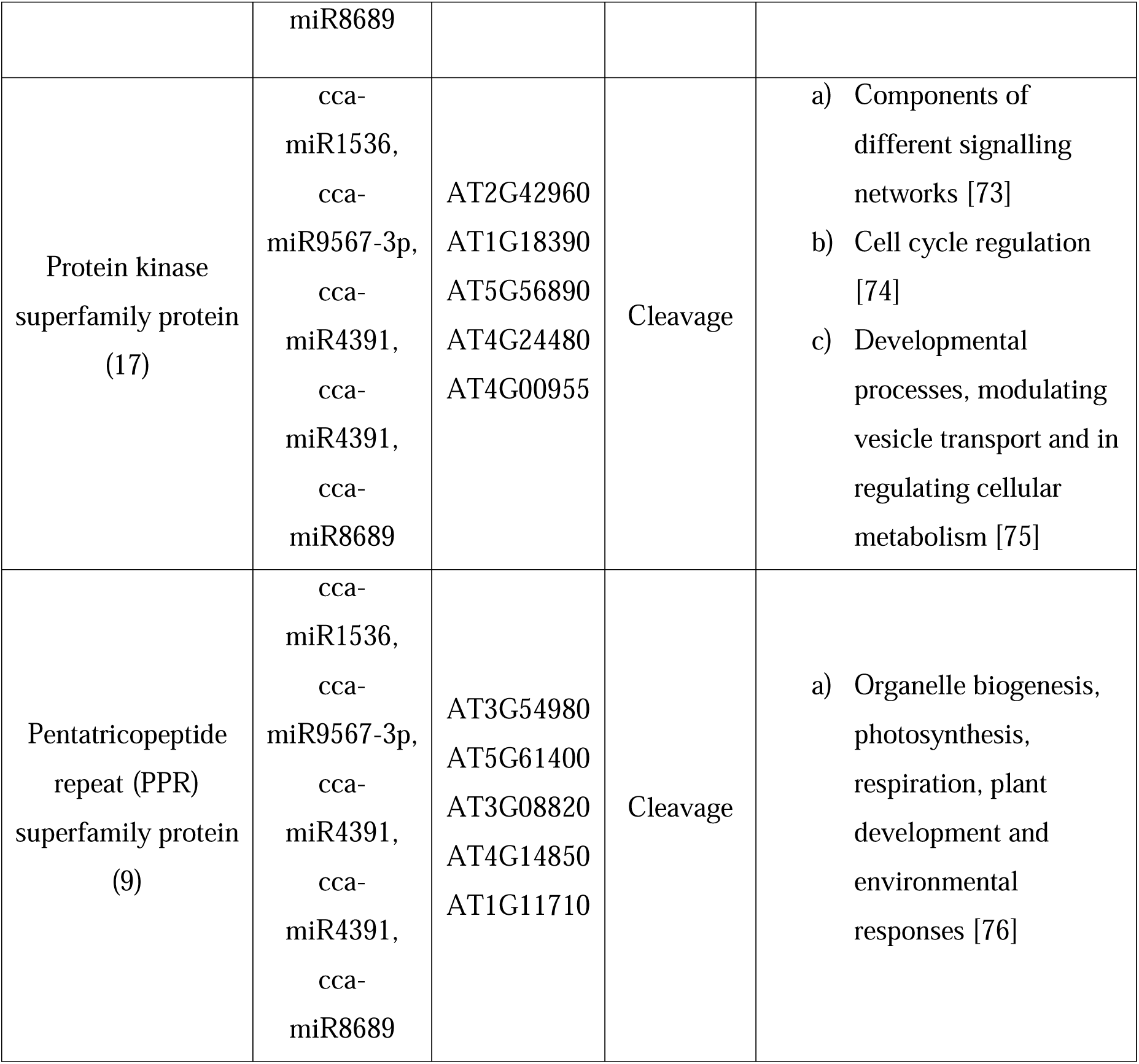
Putative miRNA targets in the crucial cellular process of *Corchorus capsularis*

## 4. Discussion

Identification of conserved microRNA in the genome of *C. capsularis* may have significant roles in the improvement of fibres quality including its mass production. Though xylan type bast fibres are traditionally used in the low quality household products because of its lower smoothness and higher percentage of lignin compounds, but now jute products are being emphasized for the sustainable application for its eco-friendly nature[77,78,79,80]. Hence, lignin biogenesis pathways and other molecular basis of fibres formation are now getting more importance for its commercial and economic perspective [81,82,83]. The regulation of lignin biosynthesis leading to fibres formation could initiate the lower percentage of lignin compound in the jute fibres which is the prerequisite for high quality fibres product. MicroRNAs are now being prioritized in the molecular research for their diverse cellular applications from the post-transcriptional regulation to gene expression [84,85]. Although miRNAs have been extensively studied for several years in many plant species [86,87,88], but no systematic study has been performed on the *C. capsularis*. The present study was carried out to identify the putative conserved microRNA candidates from the genome of *C. capsularis* by employing different screening process. Though there are several computational approaches for identifying the miRNAs in the higher plants, animals or even in the microorganisms, but EST based miRNA search has been reported more convenient than other methods [89,90]. ESTs, partially transcribed gene sequences could be used to detect the presence and expression of potential miRNAs in the higher plants [91,92]. Plant miRNAs are found to be conserved in the deferent evolutionary convergent species [93]. In the present study, previously known mature miRNAs of dicotyledons were used to conduct homology based search for identifying the putative miRNA homologs of *C. capsularis* in the available ESTs of jute genome. Though, few previous miRNA studies of higher plants used only the miRNAs of *Arabidopsis thaliana* for their homology search [94,95], but present study considered around 3350 non-redundant mature miRNAs of dicotyledons plants for enhancing the strength of broad spectrum homology search for conserved miRNAs in the jute genome.

Retrieved mature miRNAs of dicotyledons were allowed to homology search with 763 non-redundant ESTs of *C. capsularis* where 621 ESTs were identified for the further screening process. Non-coding ESTs were prioritized for being the putative miRNAs candidates [96,97,98] and only 36 non-coding ESTs were finalized in the present study. Different criteria were set up for screening the ESTs in order to identify the putative miRNAs of jute [99,100]. Though it had been reported that one miRNA exists per 10,000 ESTs in plants [101,102], but five miRNAs have been characterized in *C. capsularis* genome through scrutinizing a series of screening processes from initial 826 ESTs. The putative miRNAs were found with 18 nucleotide length which are also observed in the other studies [103,104]. The pre-miRNA were checked for the ability of folding into an appropriate hairpin secondary structure and mature miRNA should be placed in one arm of the hairpin structure (Figure 2). Minimal free energy index (MFEI) indicates the probable miRNA (average MFEI = -0.65) candidates with respect to other RNAs such as tRNAs (MFEI = -0.64) and rRNAs (MFEI = -0.59) [105,106,107,108]. The MFEI value of the newly identified five putative miRNAs of jute were found within the average range (−0.49, -0.58, -0.6, -0.73, and -1.56). Here, (A+U) content of pre-miRNA were found between the range of 55% to 61%, which was also observed in the other higher plants miRNAs [109,110,111]. Five conserved miRNA families (miR1536, miR9567-3p, miR4391, miR11300 and miR8689) were found in the jute genome which were also reported in the different plant families [112,113,114,115] and the predictive putative miRNAs were entitled as cca-miR1536, cca-miR9567-3p, cca-miR4391, cca-miR11300, and cca-miR8689. Multiple sequence alignment and comparative nucleotide composition studies exhibited that the putative miRNAs of jute were highly conserved in the other organisms which supported the evolutionary convergent relationships among the higher plant species [116,117,118].

In addition, a total of 1052 genes were identified which could be inhibited by the predicted mature miRNAs of jute. Among this huge gene pool, there were five genes which were somehow involved in the lignin biosynthesis or secondary cell wall formation process. ‘NAC Domain Containing Protein’, ‘WRKY DNA binding protein’, ‘3-dehydroquinate synthase’, ‘S-adenosyl-L-Met–dependent methyl transferase’ and ‘Vascular-related NAC-Domain’ coding genes associated with the putative miRNA of jute were studied exclusively for their important roles in the lignin biosynthesis or secondary cell wall formation. The study revealed that cca-miR4391 and cca-miR11300 might inhibit the ‘NAC Domain Containing Protein’ whereas cca-miR1536, cca-miR4391 and cca-miR8689 might cleave the ‘*S*-adenosyl-l-Met–dependent methyl transferase’.NAC are functionally redundant, being sufficient for all vessel secondary cell wall formation in fibres and vessels. Secondary cell wall formation associated NAC protein and its functional homologues are master switches that turn on a subset of transcription factors which also directly activate the expression of SCW biosynthetic genes [55,56]. The diverse variety of ‘*S*-adenosyl-l-Met–dependent methyl transferase’ has been observed in the plant. During the biogenesis of numerous plant secondary compounds alongside lignin molecules, ‘*S*-adenosyl-l-Met–dependent methyl transferase’ usually act on Phe-derived substrates. This enzymes has also been reported for its contribution in the synthesis of phytoestrogens, allelochemicals, flower pigments, antimicrobial compounds and isoflavonoid [57,58,59]. ‘WRKY DNA binding protein’ could be inhibited by the predicted cca-miR9567 and cca-miR8689 of Corchorus capsularis, and this protein could regulate the phenylpropanoid pathway and contribute in the biosynthesis of important pharmaceutical, aromatherapy and biofuel [60,61,62]. WRKYs are exclusively needed for proper expression of genes in the lignin biosynthetic pathway, and these are also found to be involved in the synthesis of phytohormones, phytoalexins, and other defense-related chemicals [63,64,65]. Again, the putative cca-miR1536 and cca-miR9567 were found to inhibit the ‘3-dehydroquinate synthase’ and ‘S-adenosyl-L-Met–dependent methyl transferase’ where ‘3-dehydroquinate synthase’ had significance contribution in the shikimate and aromatic amino acid biosynthetic pathways [66]. In addition, ‘*Vascular-related NAC-Domain* (*VND)’ could be inhibited by the predicted* cca-miR9567-3p. In Arabidopsis, secondary wall biosynthesis in the xylem conducting cells, vessels, has also been shown to be regulated by *VND* genes [67]. The expression of *VND4* gene was found mostly in the vessels and it was able to turn on transcription factors involved in the expression of secondary wall-associated functions and programmed cell death. VND4 was also found to prevent the the secondary wall defects in the fibers [68]. Though lignin provides mechanical strength to save fibres from biological or chemical degradation, it assumes to easy separation of lignin during fibres process for making quality products [119]. Here, the predicted miRNAs were found to be potent to target essential genes involved in the lignin or secondary wall synthesis alongside with different cellular process. The miRNA targeted regulation over the gene expression has already been reported in different plant species [120,121,122], hence, the putative miRNA of jute might exhibit regulatory roles over the important cellular expression through *in vivo* and *in vitro* validation.

## 5. Conclusion

As the screening and characterization of mature miRNAs has yet to be investigated in *C. capsularis*, it seems the predicted mature miRNAs and target genes might be larger and these are needed to be studied for complete understanding of miRNA derived regulation over the production of high quality fibres. The current study concluded with the suggestion of putative miRNAs in the jute genome with their correspondent gene targets, and also reflected some of the key regulatory proteins involved in the lignin biosynthesis that might be controlled by miRNA dependent cellular actions.

## Acknowledgements

Authors would like to acknowledge the Faculty of Biotechnology and Genetic Engineering, Sylhet Agricultural University for the technical support of the project.

---

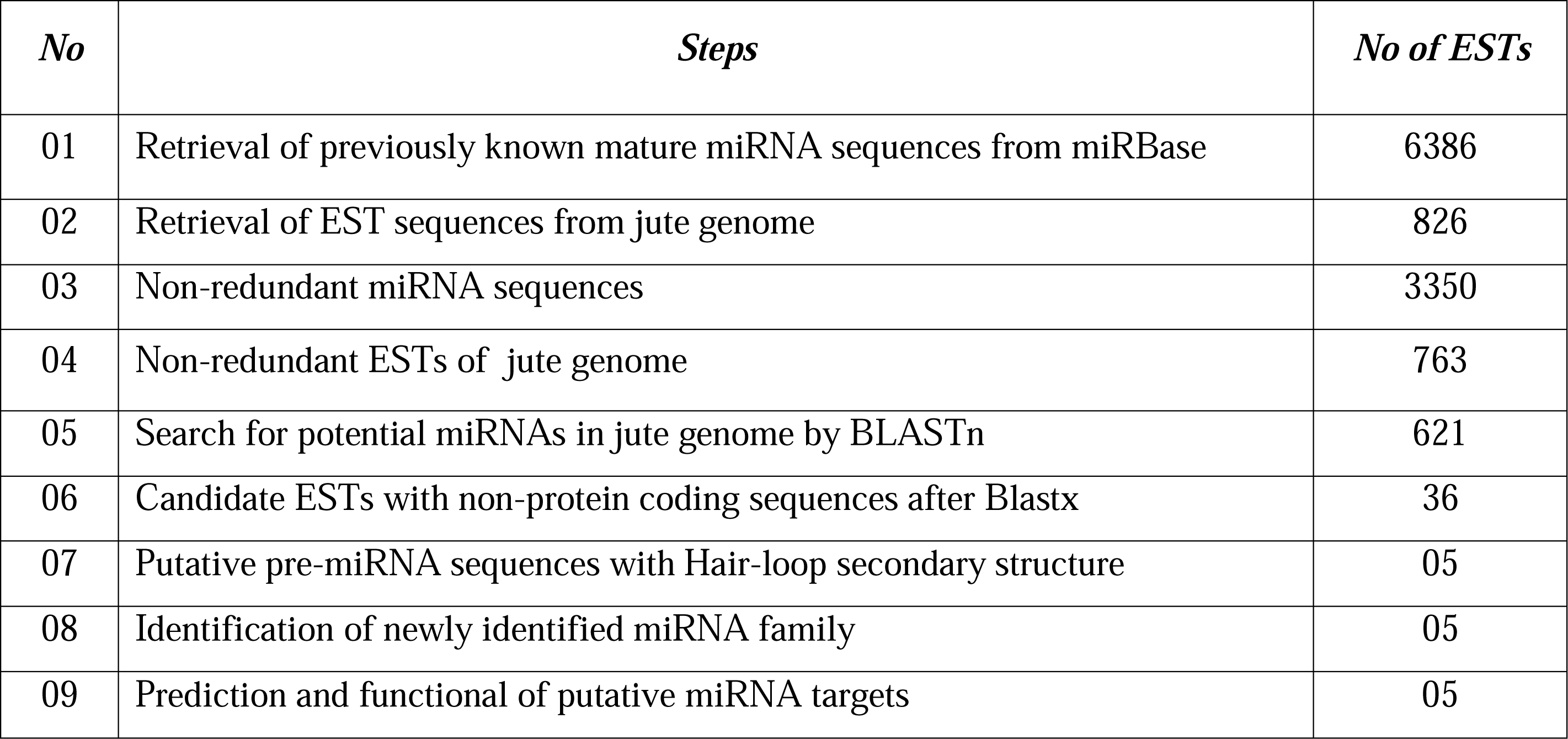

## All Legends

**Supplementary Table 1:** Non-coding EST sequence containing the predicted precursor miRNA

**Supplementary Table 2:** Around 1052 targets of five miRNAs by the psRNATarget

**Supplementary File 1:** A total of 6386 of plant miRNA from miRBase database. **Supplementary File 2:** Retrieved 826 EST sequences of Corchorus capsularis **Supplementary File 3:** Total of 3350 miRNA after redundancy screening **Supplementary File 4:** Total of 763 non-redundant EST sequences of jute

**Supplementary File 5:** Potential 621 ESTs of jute genome towards the prediction putative miRNA

**Supplementary File 6:** Total of 36 Putative non-coding EST sequences of jute genome

**Supplementary File 7:** Non-coding ESTs of jute genome failed to fulfil the suggested criteria

