## Supplementary Table 1 for "*In silico* Identification and Functional Characterization of Conserved miRNAs in Fibre Biogenesis Crop *Corchorus capsularis*"

**Supplementary Table 1:** Non-coding EST sequence containing the predicted precursor miRNA

| ***EST Accession No.*** | ***Non-coding EST sequence containing the predicted precursor miRNA (highlighted)*** |
| --- | --- |
| >GR463689.1 | AGCGTGGTCGCGGCCGAGGTACATACGGGGAAACTGAAGCCAGGAGAAGAAGAGGGAGAAACTGTCCAGGGCCTCACAGGGAGGCATTCATGGAGCTGGAGAAACCAGGGATCCTGACCGCAGGGCCCATCAATTGCTACATAGAGATACACATGGAAAGAGGGTTTAACAGACCCTCTCTAGGACACAACTGTTTCTGCTTTGAGCAATAATTTCTAACCTGAGGCCAACATGTCCCACTGCCCCTTGGGTTGGCTGGGGTTGTTTCCTGAGCCAAGCATCCAATATCATGCCCCACTCAATGGCCTAGAAGCTGCCTGTACCTGCCCGGGCGGCCGCTCGA |
| >HS411997.1 | TAACAACGATATTTACCCAGTTTCCCGACAACACCGAATTTGCAGTTCCCTTGAAGGAGTCTAGGCCTGAAGGTTCAGGAAGCTTCGACTGTACCTGATGTGGTGAAAGCTATATTAGGGGAAGGGTATCTATTATATATATGTAGCTTATAAATAAGCAACCCAGAGAGTTGCATCGCATGCCTCTGATTTGTGTTTCCAAGACTACTTTTGTTTAACTGCATTTGGTCGGAGCTCGCCATGGGCATTTAAGACTTTTAAATTACTTATTAGTGTGTGAACTATATACCTTCCATTAAATATATCACATGCTTAATGCTAATTCCGGTACTGTACCAAAAAAAAAAAGCTTCCTTTTAGGGACTTTTGAACGCCCTCTATGACCAGTTTCTGATGTGCTTCTTCGCTCATCACTTGGCTTTGCAAGGCTTCCTTGCTTGGCTTGCGTCGGGTACCCACTACCACCTCTGACATTAAGGGATGCATTCGTCCTAGGATGCTTTCAAGGAACCAGTGAGGTCATGGCCACATCTTTTGCCTTACAGCCAAAATCCTTGCCAAATAAATTAACGTCCTCTGTACATACCGTCATAAATGTCCCTTGATAAACGTAAGGGTTCCACAAGCTTCACAGAAACGAAACCATTCT |
| >FK826577.1 | ATTCACGCCGCTGCTTCTGTAGAAACAACATCTAACAAGTAGATTTGGTACTGCAGGAGACCCTTTTCGAAGACTTTTCACCATGAGAGCCAAGTGGAAGAAGAAGCGTATGAGAAGGTTGAAGAGGAAGAGAAGAAAGATGAGACAAAGATCTAAGTAGACTACTTTACTTTAGAGCTTATGCCCGTTTAGATTTCTGTTTGTTTGTTTTGATAAGATCCTTTTGTTAAAAGGATCGAACTTTATCTTCTTGTTTTCTGGGTTTAGTTTGTAACTTTTGTTGAACCAATTTTCATTTACTGGGAAAGTGTTATTTGAAGTTTAATATGCTTTTGGATCTAAAAAA |
| >GH985240.1 | GTTGAGGAACGAGTTTGGAAGTTGCTGATGGGATGGGAATAAGTATAATCTCAACATTTGTTCCTCCCGGTTGGCTATGGCCCCTGTACAGTATACTTTGCCCTGCCACGCAATGTAGTTGATGTAAATTACGGCTCACCTGCAACTAAAATTTCCCATCTTACCATTCCAGCTTCGATCCTGTGAGGCTGACCTTGTGATATAGAGGTGAGTTGGTAGTAGCAAAGAAGGTAAAAGTAATAGTGAAGCTCTTATGGCAATGTATACATCAAAGTTGTAGCATAGAGGTAGATATCATGCATATCCTTGTTTTGTAAAGCATCTTTCATGATGGACCCTTTGGTGGGTCAGTTGGGTCTAATCAAGCAATCCGACGAAATTTCGACATCAATTCTCATTTCCCTGTACAGAAAAATGCAGAACACATGTAGAAAAGAATAGCATTTTCCAGTTTCCGCATATTATCAATATTATGCCCTCA |
| >JK743800.1 | CCGGCCCCCCTTTTTTCCCTCTCTCCGGGGGGGGGGGGGGGGCCCCCCCCCCCCCCCCCTCTCCTTCTTTCCAAACAGGTGGCTGGTGGGGGGGGGACAAAAAAATTACCCCCCAAATTGGTTTCCCCCTTGCGTTGTTTAGGCGGGGTGGGTCTCCACCTTTTAGACGTAGGGTTCGCCTCCCATTCCGAGTTATTTCCCTGAAAAAAGGGGGGGTTGTGTGGGGGGGGGTGACCGGGAAACTAGGGAACCTAACAAAAAGGCGGGCGACTGGGGACAGCGGGGGTATCGGTCGTAAACGGGCCCCGTGGGTAAATGGCCGGCCCCCCCCCCCTTTTTTTCGGGGTTTTGGGGGGGGGGGGGGCGCCGAGAAACCCCCCCCCTGGAAACCCCCCCCCCCAAATACATTTTTTTTGGGGGGGAGGCGGGAGAACGGGGGCAATGTTAGGGGAAGCCACAGAGAGGGAAGAGCGCGCGTTGGGGGGGGGGGGGTGAGGATAGCTTGGACATATTTGTCATAAAACAGCGACACTATCCCCCCGCCAACCGGCGCCGGGGTTAAGGGGGGGGGAGTAGTGCGGCGGGTAACCGGCCGTATATTTCTCCCGCCACGCCCCCACGCTATTATTCCTGGTCGTTAGATCGTTTTATATGTGGGCTCACTTGTCGGGGGAGAATGCAGGCACGAAAGAACACACGAGATGATGATACTTTTCTTTTCTCTATAGATGACGAAAGGGAGGAGGGGTGCGGGACTGTCTAGTGAGTGTGCCTCCCCCCCGCCGACCTGTTGTCCCTCCCTCGTTTATCTGGGCCGTAGATATGGCCTTCATGAAG |
