## Supplementary Table 2 for "*In silico* Identification and Functional Characterization of Conserved miRNAs in Fibre Biogenesis Crop *Corchorus capsularis*"

**Supplementary Table 2:** Around 1052 targets of five miRNAs by the psRNATarget

| **cca-miR1536** | | |
| --- | --- | --- |
| Target_Acc. | Inhibition | Target_Desc. |
| AT3G53480.1 | Cleavage | \| Symbols: PIS1, PDR9, ATPDR9, ABCG37 \| pleiotropic drug resistance 9 \| chr3:19825307-19831797 FORWARD LENGTH=4565 |
| AT2G28950.1 | Cleavage | \| Symbols: ATEXPA6, ATEXP6, ATHEXP ALPHA 1.8, EXPA6 \| expansin A6 \| chr2:12431341-12433595 REVERSE LENGTH=1386 |
| AT5G43760.1 | Cleavage | \| Symbols: KCS20 \| 3-ketoacyl-CoA synthase 20 \| chr5:17585763-17588597 FORWARD LENGTH=1841 |
| AT5G10630.1 | Cleavage | \| Symbols: \| Translation elongation factor EF1A/initiation factor IF2gamma family protein \| chr5:3360173-3364596 FORWARD LENGTH=2280 |
| AT5G10630.2 | Cleavage | \| Symbols: \| Translation elongation factor EF1A/initiation factor IF2gamma family protein \| chr5:3360117-3364619 FORWARD LENGTH=2362 |
| AT1G52880.1 | Cleavage | \| Symbols: NAM, ANAC018, ATNAM, NARS2 \| NAC (No Apical Meristem) domain transcriptional regulator superfamily protein \| chr1:19688957-19690542 REVERSE LENGTH=1403 |
| AT3G21420.1 | Cleavage | \| Symbols: \| 2-oxoglutarate (2OG) and Fe(II)-dependent oxygenase superfamily protein \| chr3:7541477-7543518 FORWARD LENGTH=1494 |
| AT2G19110.1 | Cleavage | \| Symbols: HMA4, ATHMA4 \| heavy metal atpase 4 \| chr2:8278881-8286445 FORWARD LENGTH=3812 |
| AT4G10440.1 | Cleavage | \| Symbols: \| S-adenosyl-L-methionine-dependent methyltransferases superfamily protein \| chr4:6459728-6461932 REVERSE LENGTH=1902 |
| AT5G16270.1 | Cleavage | \| Symbols: ATRAD21.3, SYN4 \| sister chromatid cohesion 1 protein 4 \| chr5:5316299-5322696 FORWARD LENGTH=3689 |
| AT5G02310.1 | Cleavage | \| Symbols: PRT6 \| proteolysis 6 \| chr5:474279-482783 FORWARD LENGTH=6252 |
| AT2G40030.1 | Cleavage | \| Symbols: NRPD1B, DRD3, ATNRPD1B, DMS5, NRPE1 \| nuclear RNA polymerase D1B \| chr2:16714494-16723583 FORWARD LENGTH=6348 |
| AT2G14260.2 | Cleavage | \| Symbols: PIP \| proline iminopeptidase \| chr2:6041253-6043705 REVERSE LENGTH=1408 |
| AT2G14260.1 | Cleavage | \| Symbols: PIP \| proline iminopeptidase \| chr2:6041253-6043974 REVERSE LENGTH=1436 |
| AT1G60000.1 | Cleavage | \| Symbols: \| RNA-binding (RRM/RBD/RNP motifs) family protein \| chr1:22093569-22094626 REVERSE LENGTH=972 |
| AT1G75310.1 | Cleavage | \| Symbols: AUL1 \| auxin-like 1 protein \| chr1:28261004-28266124 FORWARD LENGTH=4347 |
| AT2G30080.1 | Cleavage | \| Symbols: ZIP6, ATZIP6 \| ZIP metal ion transporter family \| chr2:12838730-12840112 REVERSE LENGTH=1026 |
| AT1G52920.1 | Cleavage | \| Symbols: GCR2, GPCR \| G protein coupled receptor \| chr1:19709275-19711052 REVERSE LENGTH=1322 |
| AT3G25590.1 | Cleavage | \| Symbols: \| unknown protein; Has 149 Blast hits to 140 proteins in 44 species: Archae - 0; Bacteria - 6; Metazoa - 40; Fungi - 6; Plants - 39; Viruses - 0; Other Eukaryotes - 58 (source: NCBI BLink). \| chr3:9302010-9303615 FORWARD LENGTH=1606 |
| AT2G17510.2 | Cleavage | \| Symbols: EMB2763 \| ribonuclease II family protein \| chr2:7609338-7616342 REVERSE LENGTH=3196 |
| AT3G29515.1 | Cleavage | \| Symbols: \| transposable element gene \| chr3:11341766-11346095 REVERSE LENGTH=4330 |
| AT3G52250.1 | Cleavage | \| Symbols: \| Duplicated homeodomain-like superfamily protein \| chr3:19376138-19383100 FORWARD LENGTH=5366 |
| AT1G27440.1 | Cleavage | \| Symbols: GUT2, IRX10, ATGUT1 \| Exostosin family protein \| chr1:9529141-9531375 REVERSE LENGTH=1525 |
| AT1G32790.2 | Cleavage | \| Symbols: CID11 \| CTC-interacting domain 11 \| chr1:11874704-11877387 REVERSE LENGTH=1744 |
| AT5G58375.1 | Cleavage | \| Symbols: \| Methyltransferase-related protein \| chr5:23596272-23596960 FORWARD LENGTH=610 |
| AT1G78360.1 | Cleavage | \| Symbols: ATGSTU21, GSTU21 \| glutathione S-transferase TAU 21 \| chr1:29482070-29482966 REVERSE LENGTH=669 |
| AT2G34520.1 | Cleavage | \| Symbols: RPS14 \| mitochondrial ribosomal protein S14 \| chr2:14547784-14548758 REVERSE LENGTH=885 |
| AT5G23030.1 | Cleavage | \| Symbols: TET12 \| tetraspanin12 \| chr5:7726788-7727809 FORWARD LENGTH=926 |
| AT1G30920.1 | Cleavage | \| Symbols: \| F-box family protein \| chr1:11004242-11005444 REVERSE LENGTH=1203 |
| AT4G29150.1 | Cleavage | \| Symbols: IQD25 \| IQ-domain 25 \| chr4:14378767-14380511 FORWARD LENGTH=1374 |
| AT3G49080.1 | Cleavage | \| Symbols: \| Ribosomal protein S5 domain 2-like superfamily protein \| chr3:18194154-18196727 REVERSE LENGTH=1709 |
| AT3G01890.1 | Cleavage | \| Symbols: \| SWIB/MDM2 domain superfamily protein \| chr3:310015-311976 FORWARD LENGTH=1898 |
| AT5G35753.1 | Cleavage | \| Symbols: \| Domain of unknown function (DUF3444) \| chr5:13922569-13925272 REVERSE LENGTH=1927 |
| AT2G42370.1 | Cleavage | \| Symbols: \| unknown protein; BEST Arabidopsis thaliana protein match is: unknown protein (TAIR:AT3G58110.2); Has 205 Blast hits to 191 proteins in 60 species: Archae - 3; Bacteria - 23; Metazoa - 73; Fungi - 8; Plants - 34; Viruses - 0; Other Eukaryotes - 64 (source: NCBI BLink). \| chr2:17643334-17645533 FORWARD LENGTH=2148 |
| AT5G58350.1 | Cleavage | \| Symbols: WNK4, ZIK2 \| with no lysine (K) kinase 4 \| chr5:23584789-23587801 FORWARD LENGTH=2552 |
| AT3G54980.1 | Cleavage | \| Symbols: \| Pentatricopeptide repeat (PPR) superfamily protein \| chr3:20370234-20373069 FORWARD LENGTH=2836 |
| AT5G57160.1 | Cleavage | \| Symbols: ATLIG4, LIG4 \| DNA ligase IV \| chr5:23154965-23161746 REVERSE LENGTH=3903 |
| AT5G45116.1 | Cleavage | \| Symbols: \| transposable element gene \| chr5:18233906-18237928 FORWARD LENGTH=4023 |
| AT5G29708.1 | Cleavage | \| Symbols: \| transposable element gene \| chr5:11631511-11634456 REVERSE LENGTH=2946 |
| AT4G24990.1 | Cleavage | \| Symbols: ATGP4 \| Ubiquitin family protein \| chr4:12849709-12851633 REVERSE LENGTH=909 |
| AT1G30900.1 | Cleavage | \| Symbols: VSR6, VSR3;3, BP80-3;3 \| VACUOLAR SORTING RECEPTOR 6 \| chr1:10997275-11000543 FORWARD LENGTH=1896 |
| AT3G52170.1 | Cleavage | \| Symbols: \| DNA binding \| chr3:19346799-19349513 REVERSE LENGTH=1893 |
| AT3G52170.2 | Cleavage | \| Symbols: \| DNA binding \| chr3:19346752-19349533 REVERSE LENGTH=1974 |
| AT5G50510.1 | Translation | \| Symbols: \| Molecular chaperone Hsp40/DnaJ family protein \| chr5:20567386-20568075 REVERSE LENGTH=690 |
| AT5G50620.1 | Translation | \| Symbols: \| Molecular chaperone Hsp40/DnaJ family protein \| chr5:20600727-20601416 REVERSE LENGTH=690 |
| AT5G06470.1 | Cleavage | \| Symbols: \| Glutaredoxin family protein \| chr5:1974659-1975378 REVERSE LENGTH=720 |
| AT5G16660.2 | Cleavage | \| Symbols: \| unknown protein; FUNCTIONS IN: molecular_function unknown; INVOLVED IN: biological_process unknown; LOCATED IN: chloroplast, membrane; EXPRESSED IN: 23 plant structures; EXPRESSED DURING: 14 growth stages; BEST Arabidopsis thaliana protein match is: unknown protein (TAIR:AT3G02900.1); Has 106 Blast hits to 106 proteins in 32 species: Archae - 0; Bacteria - 28; Metazoa - 0; Fungi - 0; Plants - 78; Viruses - 0; Other Eukaryotes - 0 (source: NCBI BLink). \| chr5:5465484-5467063 REVERSE LENGTH=773 |
| AT5G16660.1 | Cleavage | \| Symbols: \| unknown protein; FUNCTIONS IN: molecular_function unknown; INVOLVED IN: biological_process unknown; LOCATED IN: chloroplast thylakoid membrane, chloroplast, membrane, chloroplast envelope; EXPRESSED IN: 23 plant structures; EXPRESSED DURING: 14 growth stages; BEST Arabidopsis thaliana protein match is: unknown protein (TAIR:AT3G02900.1); Has 30201 Blast hits to 17322 proteins in 780 species: Archae - 12; Bacteria - 1396; Metazoa - 17338; Fungi - 3422; Plants - 5037; Viruses - 0; Other Eukaryotes - 2996 (source: NCBI BLink). \| chr5:5465484-5467063 REVERSE LENGTH=779 |
| AT2G42960.1 | Cleavage | \| Symbols: \| Protein kinase superfamily protein \| chr2:17868597-17870630 REVERSE LENGTH=1485 |
| AT4G34320.1 | Cleavage | \| Symbols: \| Protein of unknown function (DUF677) \| chr4:16422166-16423948 FORWARD LENGTH=1519 |
| AT1G79610.1 | Cleavage | \| Symbols: ATNHX6, NHX6 \| Na+/H+ antiporter 6 \| chr1:29952873-29957189 REVERSE LENGTH=1943 |
| AT5G28928.1 | Cleavage | \| Symbols: \| transposable element gene \| chr5:10962068-10963114 REVERSE LENGTH=1047 |
| AT3G33076.1 | Cleavage | \| Symbols: \| transposable element gene \| chr3:13739731-13744449 REVERSE LENGTH=4719 |
| AT5G28335.1 | Cleavage | \| Symbols: \| transposable element gene \| chr5:10308601-10313586 REVERSE LENGTH=4986 |
| AT3G33084.1 | Cleavage | \| Symbols: \| transposable element gene \| chr3:13675144-13680170 FORWARD LENGTH=5027 |
| AT2G12770.1 | Cleavage | \| Symbols: \| transposable element gene \| chr2:5239769-5245033 REVERSE LENGTH=5265 |
| AT4G04300.1 | Cleavage | \| Symbols: \| transposable element gene \| chr4:2086200-2087758 REVERSE LENGTH=1559 |
| AT3G25890.1 | Translation | \| Symbols: \| Integrase-type DNA-binding superfamily protein \| chr3:9475575-9477455 FORWARD LENGTH=1781 |
| AT4G30960.1 | Cleavage | \| Symbols: CIPK6, SIP3, SNRK3.14, ATCIPK6 \| SOS3-interacting protein 3 \| chr4:15067053-15069010 FORWARD LENGTH=1958 |
| AT3G25890.2 | Translation | \| Symbols: \| Integrase-type DNA-binding superfamily protein \| chr3:9475350-9477460 FORWARD LENGTH=2111 |
| AT4G20850.1 | Cleavage | \| Symbols: TPP2 \| tripeptidyl peptidase ii \| chr4:11160721-11169889 REVERSE LENGTH=4357 |
| AT2G15920.1 | Cleavage | \| Symbols: \| transposable element gene \| chr2:6935939-6940405 FORWARD LENGTH=4467 |
| AT5G34965.1 | Cleavage | \| Symbols: \| transposable element gene \| chr5:13253349-13258229 REVERSE LENGTH=4881 |
| AT1G02580.1 | Cleavage | \| Symbols: MEA, EMB173, FIS1, SDG5 \| SET domain-containing protein \| chr1:544783-549202 FORWARD LENGTH=2291 |
| AT1G11710.1 | Cleavage | \| Symbols: \| Pentatricopeptide repeat (PPR) superfamily protein \| chr1:3948714-3951359 FORWARD LENGTH=2646 |
| AT5G27606.1 | Cleavage | \| Symbols: \| unknown protein; Has 30201 Blast hits to 17322 proteins in 780 species: Archae - 12; Bacteria - 1396; Metazoa - 17338; Fungi - 3422; Plants - 5037; Viruses - 0; Other Eukaryotes - 2996 (source: NCBI BLink). \| chr5:9752002-9753404 FORWARD LENGTH=813 |
| AT3G55180.1 | Cleavage | \| Symbols: \| alpha/beta-Hydrolases superfamily protein \| chr3:20454903-20456682 FORWARD LENGTH=939 |
| AT5G16380.1 | Cleavage | \| Symbols: \| Protein of unknown function, DUF538 \| chr5:5359363-5360710 REVERSE LENGTH=1052 |
| AT1G76170.1 | Cleavage | \| Symbols: \| 2-thiocytidine tRNA biosynthesis protein, TtcA \| chr1:28584039-28586225 REVERSE LENGTH=1139 |
| AT2G39400.1 | Cleavage | \| Symbols: \| alpha/beta-Hydrolases superfamily protein \| chr2:16452612-16454801 FORWARD LENGTH=1194 |
| AT3G57640.1 | Cleavage | \| Symbols: \| Protein kinase superfamily protein \| chr3:21344800-21346057 REVERSE LENGTH=1258 |
| AT5G50830.1 | Cleavage | \| Symbols: \| unknown protein; FUNCTIONS IN: molecular_function unknown; INVOLVED IN: N-terminal protein myristoylation; LOCATED IN: cellular_component unknown; EXPRESSED IN: 11 plant structures; EXPRESSED DURING: L mature pollen stage, M germinated pollen stage, 4 anthesis, C globular stage, petal differentiation and expansion stage; Has 4984 Blast hits to 3288 proteins in 342 species: Archae - 12; Bacteria - 257; Metazoa - 1366; Fungi - 452; Plants - 199; Viruses - 77; Other Eukaryotes - 2621 (source: NCBI BLink). \| chr5:20683282-20684827 FORWARD LENGTH=1294 |
| AT5G50830.2 | Cleavage | \| Symbols: \| unknown protein; Has 4750 Blast hits to 3160 proteins in 341 species: Archae - 14; Bacteria - 239; Metazoa - 1329; Fungi - 394; Plants - 197; Viruses - 79; Other Eukaryotes - 2498 (source: NCBI BLink). \| chr5:20683279-20684855 FORWARD LENGTH=1328 |
| AT1G60710.1 | Cleavage | \| Symbols: ATB2 \| NAD(P)-linked oxidoreductase superfamily protein \| chr1:22354753-22356761 REVERSE LENGTH=1492 |
| AT3G17290.1 | Cleavage | \| Symbols: \| transposable element gene \| chr3:5905052-5906716 REVERSE LENGTH=1665 |
| AT2G37930.1 | Cleavage | \| Symbols: \| Protein of unknown function (DUF3527) \| chr2:15873117-15874972 FORWARD LENGTH=1773 |
| AT3G44580.1 | Cleavage | \| Symbols: \| BEST Arabidopsis thaliana protein match is: Arabidopsis retrotransposon ORF-1 protein (TAIR:AT2G14000.1); Has 12 Blast hits to 12 proteins in 2 species: Archae - 0; Bacteria - 0; Metazoa - 0; Fungi - 0; Plants - 12; Viruses - 0; Other Eukaryotes - 0 (source: NCBI BLink). \| chr3:16160512-16161003 REVERSE LENGTH=492 |
| AT3G55670.1 | Cleavage | \| Symbols: \| CONTAINS InterPro DOMAIN/s: FBD (InterPro:IPR013596), FBD-like (InterPro:IPR006566); BEST Arabidopsis thaliana protein match is: F-box/RNI-like/FBD-like domains-containing protein (TAIR:AT3G52680.2); Has 400 Blast hits to 249 proteins in 3 species: Archae - 0; Bacteria - 0; Metazoa - 0; Fungi - 0; Plants - 400; Viruses - 0; Other Eukaryotes - 0 (source: NCBI BLink). \| chr3:20658013-20659503 REVERSE LENGTH=657 |
| AT2G19700.1 | Cleavage | \| Symbols: \| unknown protein; FUNCTIONS IN: molecular_function unknown; INVOLVED IN: biological_process unknown; LOCATED IN: endomembrane system; Has 5 Blast hits to 5 proteins in 2 species: Archae - 0; Bacteria - 0; Metazoa - 0; Fungi - 0; Plants - 5; Viruses - 0; Other Eukaryotes - 0 (source: NCBI BLink). \| chr2:8504967-8505639 REVERSE LENGTH=673 |
| AT2G20580.1 | Cleavage | \| Symbols: RPN1A, ATRPN1A \| 26S proteasome regulatory subunit S2 1A \| chr2:8858988-8864865 FORWARD LENGTH=3065 |
| AT5G26731.1 | Cleavage | \| Symbols: \| unknown protein; Has 30201 Blast hits to 17322 proteins in 780 species: Archae - 12; Bacteria - 1396; Metazoa - 17338; Fungi - 3422; Plants - 5037; Viruses - 0; Other Eukaryotes - 2996 (source: NCBI BLink). \| chr5:9295574-9296380 FORWARD LENGTH=807 |
| AT3G44570.1 | Cleavage | \| Symbols: \| Arabidopsis retrotransposon ORF-1 protein \| chr3:16158405-16159835 FORWARD LENGTH=922 |
| AT2G14000.1 | Cleavage | \| Symbols: \| Arabidopsis retrotransposon ORF-1 protein \| chr2:5884600-5885985 REVERSE LENGTH=1041 |
| AT1G09240.1 | Cleavage | \| Symbols: NAS3, ATNAS3 \| nicotianamine synthase 3 \| chr1:2984881-2986130 FORWARD LENGTH=1250 |
| AT4G04760.1 | Cleavage | \| Symbols: \| Major facilitator superfamily protein \| chr4:2424164-2427769 FORWARD LENGTH=1404 |
| AT1G43171.1 | Cleavage | \| Symbols: \| unknown protein; FUNCTIONS IN: molecular_function unknown; INVOLVED IN: biological_process unknown; LOCATED IN: cellular_component unknown; BEST Arabidopsis thaliana protein match is: unknown protein (TAIR:AT1G50220.1); Has 30201 Blast hits to 17322 proteins in 780 species: Archae - 12; Bacteria - 1396; Metazoa - 17338; Fungi - 3422; Plants - 5037; Viruses - 0; Other Eukaryotes - 2996 (source: NCBI BLink). \| chr1:16269075-16270513 FORWARD LENGTH=1439 |
| AT1G69260.1 | Cleavage | \| Symbols: AFP1 \| ABI five binding protein \| chr1:26039048-26040720 FORWARD LENGTH=1454 |
| AT1G59900.1 | Cleavage | \| Symbols: AT-E1 ALPHA, E1 ALPHA \| pyruvate dehydrogenase complex E1 alpha subunit \| chr1:22051295-22053885 FORWARD LENGTH=1468 |
| AT1G24180.1 | Cleavage | \| Symbols: IAR4 \| Thiamin diphosphate-binding fold (THDP-binding) superfamily protein \| chr1:8560525-8563495 REVERSE LENGTH=1547 |
| AT5G22320.1 | Cleavage | \| Symbols: \| Leucine-rich repeat (LRR) family protein \| chr5:7388004-7390485 REVERSE LENGTH=1589 |
| AT5G22320.2 | Cleavage | \| Symbols: \| Leucine-rich repeat (LRR) family protein \| chr5:7387985-7390520 REVERSE LENGTH=1595 |
| AT4G34020.1 | Cleavage | \| Symbols: \| Class I glutamine amidotransferase-like superfamily protein \| chr4:16298262-16300938 REVERSE LENGTH=1751 |
| AT1G04220.1 | Cleavage | \| Symbols: KCS2 \| 3-ketoacyl-CoA synthase 2 \| chr1:1119699-1122526 REVERSE LENGTH=1784 |
| AT1G50190.1 | Cleavage | \| Symbols: \| Cysteine/Histidine-rich C1 domain family protein \| chr1:18588229-18590799 REVERSE LENGTH=1860 |
| AT4G36270.1 | Cleavage | \| Symbols: \| ATP binding \| chr4:17161024-17165241 REVERSE LENGTH=2163 |
| AT2G25560.1 | Cleavage | \| Symbols: \| DNAJ heat shock N-terminal domain-containing protein \| chr2:10880671-10883868 FORWARD LENGTH=2512 |
| AT5G24320.1 | Cleavage | \| Symbols: \| Transducin/WD40 repeat-like superfamily protein \| chr5:8284436-8288039 REVERSE LENGTH=2570 |
| AT5G24320.2 | Cleavage | \| Symbols: \| Transducin/WD40 repeat-like superfamily protein \| chr5:8284436-8288039 REVERSE LENGTH=2582 |
| AT2G05250.1 | Cleavage | \| Symbols: \| DNAJ heat shock N-terminal domain-containing protein \| chr2:1913338-1916623 REVERSE LENGTH=2839 |
| AT2G05230.1 | Cleavage | \| Symbols: \| DNAJ heat shock N-terminal domain-containing protein \| chr2:1899507-1902802 REVERSE LENGTH=2849 |
| AT3G31317.1 | Cleavage | \| Symbols: \| transposable element gene \| chr3:12685821-12691103 REVERSE LENGTH=5283 |
| AT3G62290.2 | Translation | \| Symbols: ARFA1E \| ADP-ribosylation factor A1E \| chr3:23051544-23053817 FORWARD LENGTH=944 |
| AT3G62290.3 | Translation | \| Symbols: ARFA1E \| ADP-ribosylation factor A1E \| chr3:23051544-23053817 FORWARD LENGTH=949 |
| AT5G50360.1 | Cleavage | \| Symbols: \| unknown protein; BEST Arabidopsis thaliana protein match is: unknown protein (TAIR:AT3G48510.1); Has 1807 Blast hits to 1807 proteins in 277 species: Archae - 0; Bacteria - 0; Metazoa - 736; Fungi - 347; Plants - 385; Viruses - 0; Other Eukaryotes - 339 (source: NCBI BLink). \| chr5:20505180-20506338 REVERSE LENGTH=1159 |
| AT3G49080.1 | Cleavage | \| Symbols: \| Ribosomal protein S5 domain 2-like superfamily protein \| chr3:18194154-18196727 REVERSE LENGTH=1709 |
| AT3G49080.1 | Cleavage | \| Symbols: \| Ribosomal protein S5 domain 2-like superfamily protein \| chr3:18194154-18196727 REVERSE LENGTH=1709 |
| AT2G02147.1 | Cleavage | \| Symbols: LCR73 \| low-molecular-weight cysteine-rich 73 \| chr2:546664-547026 FORWARD LENGTH=243 |
| AT3G11640.1 | Cleavage | \| Symbols: \| unknown protein; FUNCTIONS IN: molecular_function unknown; INVOLVED IN: biological_process unknown; LOCATED IN: endomembrane system; BEST Arabidopsis thaliana protein match is: unknown protein (TAIR:AT3G52480.1); Has 36 Blast hits to 36 proteins in 7 species: Archae - 0; Bacteria - 0; Metazoa - 0; Fungi - 0; Plants - 36; Viruses - 0; Other Eukaryotes - 0 (source: NCBI BLink). \| chr3:3674499-3675198 FORWARD LENGTH=700 |
| AT1G69526.3 | Cleavage | \| Symbols: \| S-adenosyl-L-methionine-dependent methyltransferases superfamily protein \| chr1:26131381-26132388 FORWARD LENGTH=706 |
| AT1G69880.1 | Translation | \| Symbols: ATH8, TH8 \| thioredoxin H-type 8 \| chr1:26321479-26322993 FORWARD LENGTH=707 |
| AT1G11125.1 | Cleavage | \| Symbols: \| unknown protein; BEST Arabidopsis thaliana protein match is: unknown protein (TAIR:AT1G61170.1). \| chr1:3718783-3719711 FORWARD LENGTH=929 |
| AT1G07120.1 | Cleavage | \| Symbols: \| FUNCTIONS IN: molecular_function unknown; INVOLVED IN: biological_process unknown; LOCATED IN: chloroplast envelope; EXPRESSED IN: inflorescence meristem, petal, leaf whorl, flower; EXPRESSED DURING: 4 anthesis, petal differentiation and expansion stage; BEST Arabidopsis thaliana protein match is: Tetratricopeptide repeat (TPR)-like superfamily protein (TAIR:AT4G18570.1); Has 288 Blast hits to 260 proteins in 50 species: Archae - 0; Bacteria - 8; Metazoa - 27; Fungi - 15; Plants - 163; Viruses - 0; Other Eukaryotes - 75 (source: NCBI BLink). \| chr1:2184759-2186580 REVERSE LENGTH=1294 |
| AT1G69523.1 | Cleavage | \| Symbols: \| S-adenosyl-L-methionine-dependent methyltransferases superfamily protein \| chr1:26129551-26131292 FORWARD LENGTH=1314 |
| AT1G21600.1 | Cleavage | \| Symbols: PTAC6 \| plastid transcriptionally active 6 \| chr1:7571185-7573846 REVERSE LENGTH=1380 |
| AT1G69526.2 | Cleavage | \| Symbols: \| S-adenosyl-L-methionine-dependent methyltransferases superfamily protein \| chr1:26131294-26133196 FORWARD LENGTH=1385 |
| AT5G10745.1 | Translation | \| Symbols: \| unknown protein; FUNCTIONS IN: molecular_function unknown; INVOLVED IN: biological_process unknown; LOCATED IN: endomembrane system; BEST Arabidopsis thaliana protein match is: unknown protein (TAIR:AT5G24980.1); Has 35333 Blast hits to 34131 proteins in 2444 species: Archae - 798; Bacteria - 22429; Metazoa - 974; Fungi - 991; Plants - 531; Viruses - 0; Other Eukaryotes - 9610 (source: NCBI BLink). \| chr5:3396475-3398063 FORWARD LENGTH=1391 |
| AT1G69526.1 | Cleavage | \| Symbols: \| S-adenosyl-L-methionine-dependent methyltransferases superfamily protein \| chr1:26131421-26133196 FORWARD LENGTH=1429 |
| AT4G37840.1 | Cleavage | \| Symbols: HKL3 \| hexokinase-like 3 \| chr4:17790147-17792198 REVERSE LENGTH=1482 |
| AT4G11340.1 | Cleavage | \| Symbols: \| Disease resistance protein (TIR-NBS-LRR class) family \| chr4:6894208-6899130 REVERSE LENGTH=1488 |
| AT2G37930.1 | Cleavage | \| Symbols: \| Protein of unknown function (DUF3527) \| chr2:15873117-15874972 FORWARD LENGTH=1773 |
| AT3G17830.1 | Cleavage | \| Symbols: \| Molecular chaperone Hsp40/DnaJ family protein \| chr3:6101789-6104650 FORWARD LENGTH=1780 |
| AT1G29890.1 | Translation | \| Symbols: \| O-acetyltransferase family protein \| chr1:10463423-10467170 FORWARD LENGTH=1850 |
| AT1G29890.2 | Translation | \| Symbols: \| O-acetyltransferase family protein \| chr1:10463403-10467170 FORWARD LENGTH=1901 |
| AT5G19160.1 | Cleavage | \| Symbols: TBL11 \| TRICHOME BIREFRINGENCE-LIKE 11 \| chr5:6430486-6432728 FORWARD LENGTH=1906 |
| AT1G55870.2 | Cleavage | \| Symbols: ATPARN, AHG2 \| Polynucleotidyl transferase, ribonuclease H-like superfamily protein \| chr1:20895585-20898422 FORWARD LENGTH=2141 |
| AT1G55870.1 | Cleavage | \| Symbols: ATPARN, AHG2 \| Polynucleotidyl transferase, ribonuclease H-like superfamily protein \| chr1:20895605-20898422 FORWARD LENGTH=2191 |
| AT2G45810.1 | Cleavage | \| Symbols: \| DEA(D/H)-box RNA helicase family protein \| chr2:18859599-18862918 FORWARD LENGTH=2329 |
| AT1G32940.1 | Cleavage | \| Symbols: ATSBT3.5, SBT3.5 \| Subtilase family protein \| chr1:11937596-11940978 FORWARD LENGTH=2485 |
| AT5G04460.1 | Cleavage | \| Symbols: \| RING/U-box superfamily protein \| chr5:1260009-1263862 FORWARD LENGTH=2759 |
| AT5G04460.3 | Cleavage | \| Symbols: \| RING/U-box superfamily protein \| chr5:1259531-1264025 FORWARD LENGTH=3112 |
| AT1G48120.1 | Cleavage | \| Symbols: \| hydrolases;protein serine/threonine phosphatases \| chr1:17773688-17779955 REVERSE LENGTH=4346 |
| AT5G27250.1 | Cleavage | \| Symbols: \| transposable element gene \| chr5:9601673-9603014 FORWARD LENGTH=1047 |
| AT2G27310.1 | Cleavage | \| Symbols: \| F-box family protein \| chr2:11683862-11684967 REVERSE LENGTH=1106 |
| AT3G22790.1 | Cleavage | \| Symbols: \| Kinase interacting (KIP1-like) family protein \| chr3:8052318-8058764 REVERSE LENGTH=5537 |
| AT5G28225.1 | Cleavage | \| Symbols: \| transposable element gene \| chr5:10193379-10194578 REVERSE LENGTH=1200 |
| AT2G13370.1 | Cleavage | \| Symbols: CHR5 \| chromatin remodeling 5 \| chr2:5544181-5556464 REVERSE LENGTH=5938 |
| AT2G04450.1 | Cleavage | \| Symbols: ATNUDT6, NUDX6, ATNUDX6, NUDT6 \| nudix hydrolase homolog 6 \| chr2:1543320-1545462 FORWARD LENGTH=1311 |
| AT4G18070.4 | Cleavage | \| Symbols: \| unknown protein; BEST Arabidopsis thaliana protein match is: unknown protein (TAIR:AT1G29530.1); Has 30201 Blast hits to 17322 proteins in 780 species: Archae - 12; Bacteria - 1396; Metazoa - 17338; Fungi - 3422; Plants - 5037; Viruses - 0; Other Eukaryotes - 2996 (source: NCBI BLink). \| chr4:10030663-10032935 FORWARD LENGTH=1312 |
| AT4G04550.1 | Cleavage | \| Symbols: \| transposable element gene \| chr4:2281319-2282656 REVERSE LENGTH=1338 |
| AT1G21990.1 | Cleavage | \| Symbols: \| F-box/RNI-like/FBD-like domains-containing protein \| chr1:7740530-7742106 REVERSE LENGTH=1398 |
| AT2G26480.1 | Cleavage | \| Symbols: UGT76D1 \| UDP-glucosyl transferase 76D1 \| chr2:11263963-11265684 FORWARD LENGTH=1471 |
| AT5G65280.1 | Cleavage | \| Symbols: GCL1 \| GCR2-like 1 \| chr5:26085973-26088153 REVERSE LENGTH=1542 |
| AT1G34630.2 | Translation | \| Symbols: \| FUNCTIONS IN: molecular_function unknown; INVOLVED IN: biological_process unknown; LOCATED IN: cellular_component unknown; EXPRESSED IN: 25 plant structures; EXPRESSED DURING: 15 growth stages; BEST Arabidopsis thaliana protein match is: Mitochondrial import inner membrane translocase subunit Tim17/Tim22/Tim23 family protein (TAIR:AT5G51150.1); Has 30201 Blast hits to 17322 proteins in 780 species: Archae - 12; Bacteria - 1396; Metazoa - 17338; Fungi - 3422; Plants - 5037; Viruses - 0; Other Eukaryotes - 2996 (source: NCBI BLink). \| chr1:12685241-12687575 FORWARD LENGTH=1732 |
| AT1G28110.1 | Cleavage | \| Symbols: SCPL45 \| serine carboxypeptidase-like 45 \| chr1:9803865-9806896 REVERSE LENGTH=1738 |
| AT1G34630.1 | Translation | \| Symbols: \| BEST Arabidopsis thaliana protein match is: Mitochondrial import inner membrane translocase subunit Tim17/Tim22/Tim23 family protein (TAIR:AT5G51150.1); Has 323 Blast hits to 315 proteins in 124 species: Archae - 0; Bacteria - 0; Metazoa - 95; Fungi - 110; Plants - 73; Viruses - 0; Other Eukaryotes - 45 (source: NCBI BLink). \| chr1:12685200-12687641 FORWARD LENGTH=1769 |
| AT2G34960.1 | Translation | \| Symbols: CAT5 \| cationic amino acid transporter 5 \| chr2:14744037-14745961 REVERSE LENGTH=1925 |
| AT3G50160.1 | Translation | \| Symbols: \| Plant protein of unknown function (DUF247) \| chr3:18598429-18601009 REVERSE LENGTH=2015 |
| AT1G19370.1 | Cleavage | \| Symbols: \| unknown protein; LOCATED IN: endoplasmic reticulum; EXPRESSED IN: 22 plant structures; EXPRESSED DURING: 13 growth stages; BEST Arabidopsis thaliana protein match is: unknown protein (TAIR:AT1G75140.1); Has 45 Blast hits to 43 proteins in 15 species: Archae - 0; Bacteria - 0; Metazoa - 0; Fungi - 0; Plants - 44; Viruses - 0; Other Eukaryotes - 1 (source: NCBI BLink). \| chr1:6692764-6694934 REVERSE LENGTH=2171 |
| AT3G32990.1 | Cleavage | \| Symbols: \| pseudogene, ATP synthase C subunit, blastp match of 86% identity and 9.3e-27 P-value to SP\|P06286\|ATPH_TOBAC ATP synthase C chain (EC 3.6.3.14) (Lipid-binding protein) (Subunit III). (Common tobacco) {Nicotiana tabacum} \| chr3:13534183-13536711 REVERSE LENGTH=2529 |
| AT1G49490.1 | Cleavage | \| Symbols: \| Leucine-rich repeat (LRR) family protein \| chr1:18317563-18320106 REVERSE LENGTH=2544 |
| AT1G09720.1 | Cleavage | \| Symbols: \| Kinase interacting (KIP1-like) family protein \| chr1:3144438-3147303 REVERSE LENGTH=2787 |
| AT1G20400.1 | Cleavage | \| Symbols: \| Protein of unknown function (DUF1204) \| chr1:7072192-7075838 REVERSE LENGTH=2835 |
| AT4G16280.4 | Cleavage | \| Symbols: \| RNA binding;abscisic acid binding \| chr4:9206597-9214825 REVERSE LENGTH=2903 |
| AT5G35560.1 | Cleavage | \| Symbols: \| DENN (AEX-3) domain-containing protein \| chr5:13742109-13748312 REVERSE LENGTH=2923 |
| AT5G34960.1 | Cleavage | \| Symbols: \| transposable element gene \| chr5:13246527-13251225 REVERSE LENGTH=3102 |
| AT3G09750.1 | Cleavage | \| Symbols: \| Galactose oxidase/kelch repeat superfamily protein \| chr3:2991655-2992679 REVERSE LENGTH=936 |
| AT2G09994.1 | Cleavage | \| Symbols: \| pseudogene of U-box domain-containing protein / armadillo/beta-catenin repeat family protein \| chr2:3782956-3784075 FORWARD LENGTH=1120 |
| AT5G40480.1 | Cleavage | \| Symbols: EMB3012 \| embryo defective 3012 \| chr5:16213222-16224058 FORWARD LENGTH=5987 |
| AT1G27090.1 | Translation | \| Symbols: \| glycine-rich protein \| chr1:9403900-9406256 REVERSE LENGTH=1562 |
| AT1G11710.1 | Cleavage | \| Symbols: \| Pentatricopeptide repeat (PPR) superfamily protein \| chr1:3948714-3951359 FORWARD LENGTH=2646 |
| AT2G18090.1 | Cleavage | \| Symbols: \| PHD finger family protein / SWIB complex BAF60b domain-containing protein / GYF domain-containing protein \| chr2:7864314-7867766 FORWARD LENGTH=3111 |
| AT2G15330.1 | Cleavage | \| Symbols: \| transposable element gene \| chr2:6672892-6673989 REVERSE LENGTH=1098 |
| AT4G16310.1 | Cleavage | \| Symbols: LDL3 \| LSD1-like 3 \| chr4:9218636-9225264 FORWARD LENGTH=5387 |
| AT3G30838.1 | Cleavage | \| Symbols: \| transposable element gene \| chr3:12579885-12585786 REVERSE LENGTH=5902 |
| AT3G18060.1 | Cleavage | \| Symbols: \| transducin family protein / WD-40 repeat family protein \| chr3:6183773-6187074 FORWARD LENGTH=2223 |
| AT2G22241.1 | Cleavage | \| Symbols: \| unknown protein; LOCATED IN: mitochondrion; Has 30201 Blast hits to 17322 proteins in 780 species: Archae - 12; Bacteria - 1396; Metazoa - 17338; Fungi - 3422; Plants - 5037; Viruses - 0; Other Eukaryotes - 2996 (source: NCBI BLink). \| chr2:9454244-9454399 FORWARD LENGTH=156 |
| AT1G61065.1 | Cleavage | \| Symbols: \| Protein of unknown function (DUF1218) \| chr1:22490244-22491461 REVERSE LENGTH=860 |
| AT1G04120.1 | Cleavage | \| Symbols: ATMRP5, MRP5, ATABCC5, ABCC5 \| multidrug resistance-associated protein 5 \| chr1:1064454-1070927 REVERSE LENGTH=5277 |
| AT1G04120.2 | Cleavage | \| Symbols: MRP5 \| multidrug resistance-associated protein 5 \| chr1:1064450-1071047 REVERSE LENGTH=5386 |
| AT4G26910.3 | Cleavage | \| Symbols: \| Dihydrolipoamide succinyltransferase \| chr4:13519811-13522442 REVERSE LENGTH=1565 |
| AT4G26910.2 | Cleavage | \| Symbols: \| Dihydrolipoamide succinyltransferase \| chr4:13519811-13523225 REVERSE LENGTH=1815 |
| AT4G26910.1 | Cleavage | \| Symbols: \| Dihydrolipoamide succinyltransferase \| chr4:13519811-13523222 REVERSE LENGTH=1815 |
| AT1G16680.1 | Cleavage | \| Symbols: \| Chaperone DnaJ-domain superfamily protein \| chr1:5702753-5705838 FORWARD LENGTH=2143 |
| AT3G61947.1 | Cleavage | \| Symbols: \| unknown protein; Has 30201 Blast hits to 17322 proteins in 780 species: Archae - 12; Bacteria - 1396; Metazoa - 17338; Fungi - 3422; Plants - 5037; Viruses - 0; Other Eukaryotes - 2996 (source: NCBI BLink). \| chr3:22939271-22939387 REVERSE LENGTH=117 |
| AT2G28950.1 | Cleavage | \| Symbols: ATEXPA6, ATEXP6, ATHEXP ALPHA 1.8, EXPA6 \| expansin A6 \| chr2:12431341-12433595 REVERSE LENGTH=1386 |
| AT5G56860.1 | Cleavage | \| Symbols: GNC, GATA21 \| GATA type zinc finger transcription factor family protein \| chr5:22989265-22991467 REVERSE LENGTH=1678 |
| AT3G09620.1 | Cleavage | \| Symbols: \| P-loop containing nucleoside triphosphate hydrolases superfamily protein \| chr3:2949152-2952205 REVERSE LENGTH=2970 |
| AT5G40340.1 | Cleavage | \| Symbols: \| Tudor/PWWP/MBT superfamily protein \| chr5:16131654-16134916 REVERSE LENGTH=3096 |
| AT3G58640.2 | Cleavage | \| Symbols: \| Mitogen activated protein kinase kinase kinase-related \| chr3:21686741-21693841 REVERSE LENGTH=3182 |
| AT3G58640.1 | Cleavage | \| Symbols: \| Mitogen activated protein kinase kinase kinase-related \| chr3:21686743-21693831 REVERSE LENGTH=3268 |
| AT1G04470.1 | Cleavage | \| Symbols: \| Protein of unknown function (DUF810) \| chr1:1211049-1214662 REVERSE LENGTH=3307 |
| AT1G04470.1 | Cleavage | \| Symbols: \| Protein of unknown function (DUF810) \| chr1:1211049-1214662 REVERSE LENGTH=3307 |
| AT3G05415.1 | Cleavage | \| Symbols: \| transposable element gene \| chr3:1555537-1560588 REVERSE LENGTH=5052 |
| AT2G37930.1 | Cleavage | \| Symbols: \| Protein of unknown function (DUF3527) \| chr2:15873117-15874972 FORWARD LENGTH=1773 |
| AT5G29708.1 | Cleavage | \| Symbols: \| transposable element gene \| chr5:11631511-11634456 REVERSE LENGTH=2946 |
| AT3G07525.2 | Cleavage | \| Symbols: ATG10, ATATG10 \| autophagocytosis-associated family protein \| chr3:2399143-2400402 REVERSE LENGTH=837 |
| AT3G07525.1 | Cleavage | \| Symbols: ATG10, ATATG10 \| autophagocytosis-associated family protein \| chr3:2399123-2400437 REVERSE LENGTH=889 |
| AT1G80850.1 | Cleavage | \| Symbols: \| DNA glycosylase superfamily protein \| chr1:30385449-30387522 REVERSE LENGTH=1392 |
| AT3G28760.1 | Cleavage | \| Symbols: \| FUNCTIONS IN: molecular_function unknown; INVOLVED IN: biological_process unknown; LOCATED IN: chloroplast; EXPRESSED IN: 20 plant structures; EXPRESSED DURING: 13 growth stages; CONTAINS InterPro DOMAIN/s: 3-dehydroquinate synthase, prokaryotic-type (InterPro:IPR002812); Has 390 Blast hits to 390 proteins in 131 species: Archae - 144; Bacteria - 105; Metazoa - 0; Fungi - 0; Plants - 54; Viruses - 0; Other Eukaryotes - 87 (source: NCBI BLink). \| chr3:10792955-10795312 REVERSE LENGTH=1448 |
| AT2G44550.1 | Cleavage | \| Symbols: AtGH9B10, GH9B10 \| glycosyl hydrolase 9B10 \| chr2:18389381-18391142 REVERSE LENGTH=1473 |
| AT1G05370.1 | Cleavage | \| Symbols: \| Sec14p-like phosphatidylinositol transfer family protein \| chr1:1569192-1572280 REVERSE LENGTH=1480 |
| AT1G05370.1 | Cleavage | \| Symbols: \| Sec14p-like phosphatidylinositol transfer family protein \| chr1:1569192-1572280 REVERSE LENGTH=1480 |
| AT3G28760.2 | Cleavage | \| Symbols: \| CONTAINS InterPro DOMAIN/s: 3-dehydroquinate synthase, prokaryotic-type (InterPro:IPR002812); Has 35333 Blast hits to 34131 proteins in 2444 species: Archae - 798; Bacteria - 22429; Metazoa - 974; Fungi - 991; Plants - 531; Viruses - 0; Other Eukaryotes - 9610 (source: NCBI BLink). \| chr3:10792955-10795304 REVERSE LENGTH=1506 |
| AT4G28485.1 | Cleavage | \| Symbols: DMP7, AtDMP7 \| DUF679 domain membrane protein 7 \| chr4:14075235-14076011 REVERSE LENGTH=584 |
| AT1G36950.1 | Cleavage | \| Symbols: \| RING/U-box superfamily protein \| chr1:14009673-14010822 REVERSE LENGTH=681 |

| **cca-miR9567-3p** | | |
| --- | --- | --- |
| Target_Acc. | Inhibition | Target_Desc. |
| AT4G00630.2 | Cleavage | \| Symbols: KEA2 \| K+ efflux antiporter 2 \| chr4:261449-268031 REVERSE LENGTH=4006 |
| AT4G00630.1 | Cleavage | \| Symbols: KEA2, ATKEA2 \| K+ efflux antiporter 2 \| chr4:261531-268050 REVERSE LENGTH=3910 |
| AT2G45830.2 | Cleavage | \| Symbols: DTA2 \| downstream target of AGL15 2 \| chr2:18866061-18868542 FORWARD LENGTH=2136 |
| AT2G45830.1 | Cleavage | \| Symbols: DTA2 \| downstream target of AGL15 2 \| chr2:18866061-18868542 FORWARD LENGTH=2033 |
| AT1G01790.1 | Cleavage | \| Symbols: KEA1, ATKEA1 \| K+ efflux antiporter 1 \| chr1:284610-291094 FORWARD LENGTH=3892 |
| AT1G09190.1 | Cleavage | \| Symbols: \| Tetratricopeptide repeat (TPR)-like superfamily protein \| chr1:2966120-2967718 REVERSE LENGTH=1599 |
| AT3G03790.2 | Cleavage | \| Symbols: \| ankyrin repeat family protein / regulator of chromosome condensation (RCC1) family protein \| chr3:961997-968149 FORWARD LENGTH=4032 |
| AT3G03790.1 | Cleavage | \| Symbols: \| ankyrin repeat family protein / regulator of chromosome condensation (RCC1) family protein \| chr3:961997-968149 FORWARD LENGTH=4023 |
| AT3G03790.3 | Cleavage | \| Symbols: \| ankyrin repeat family protein / regulator of chromosome condensation (RCC1) family protein \| chr3:961997-968149 FORWARD LENGTH=3932 |
| AT5G67280.1 | Cleavage | \| Symbols: RLK \| receptor-like kinase \| chr5:26842270-26845190 REVERSE LENGTH=2480 |
| AT1G28310.1 | Cleavage | \| Symbols: \| Dof-type zinc finger DNA-binding family protein \| chr1:9912203-9913837 REVERSE LENGTH=1635 |
| AT1G28310.2 | Cleavage | \| Symbols: \| Dof-type zinc finger DNA-binding family protein \| chr1:9911898-9913698 REVERSE LENGTH=1627 |
| AT2G37100.1 | Cleavage | \| Symbols: \| protamine P1 family protein \| chr2:15591101-15592104 FORWARD LENGTH=1004 |
| AT4G32340.1 | Cleavage | \| Symbols: \| Tetratricopeptide repeat (TPR)-like superfamily protein \| chr4:15612584-15614305 REVERSE LENGTH=960 |
| AT5G66070.2 | Cleavage | \| Symbols: \| RING/U-box superfamily protein \| chr5:26421855-26423232 FORWARD LENGTH=1005 |
| AT5G66070.1 | Cleavage | \| Symbols: \| RING/U-box superfamily protein \| chr5:26421855-26423240 FORWARD LENGTH=941 |
| AT2G27402.1 | Cleavage | \| Symbols: \| BEST Arabidopsis thaliana protein match is: plastid transcriptionally active 18 (TAIR:AT2G32180.1); Has 35333 Blast hits to 34131 proteins in 2444 species: Archae - 798; Bacteria - 22429; Metazoa - 974; Fungi - 991; Plants - 531; Viruses - 0; Other Eukaryotes - 9610 (source: NCBI BLink). \| chr2:11723525-11724362 FORWARD LENGTH=838 |
| AT1G12260.1 | Cleavage | \| Symbols: VND4, EMB2749, ANAC007, NAC007 \| NAC 007 \| chr1:4162832-4165852 REVERSE LENGTH=1871 |
| AT3G22170.2 | Cleavage | \| Symbols: FHY3 \| far-red elongated hypocotyls 3 \| chr3:7822018-7826085 REVERSE LENGTH=3208 |
| AT3G52800.1 | Cleavage | \| Symbols: \| A20/AN1-like zinc finger family protein \| chr3:19569311-19570682 FORWARD LENGTH=1129 |
| AT1G44045.1 | Cleavage | \| Symbols: \| transposable element gene \| chr1:16730937-16735967 REVERSE LENGTH=5031 |
| AT5G43310.3 | Cleavage | \| Symbols: \| COP1-interacting protein-related \| chr5:17379486-17385535 REVERSE LENGTH=3832 |
| AT1G32090.1 | Cleavage | \| Symbols: \| early-responsive to dehydration stress protein (ERD4) \| chr1:11540025-11544131 REVERSE LENGTH=2730 |
| AT4G14400.1 | Cleavage | \| Symbols: ACD6 \| ankyrin repeat family protein \| chr4:8294446-8298602 FORWARD LENGTH=2354 |
| AT4G23850.1 | Cleavage | \| Symbols: LACS4 \| AMP-dependent synthetase and ligase family protein \| chr4:12403454-12408335 REVERSE LENGTH=2339 |
| AT4G14400.3 | Cleavage | \| Symbols: ACD6 \| ankyrin repeat family protein \| chr4:8294452-8298602 FORWARD LENGTH=2335 |
| AT4G14400.2 | Cleavage | \| Symbols: ACD6 \| ankyrin repeat family protein \| chr4:8294446-8298602 FORWARD LENGTH=2271 |
| AT2G47260.1 | Cleavage | \| Symbols: WRKY23, ATWRKY23 \| WRKY DNA-binding protein 23 \| chr2:19404820-19407084 REVERSE LENGTH=1877 |
| AT3G28150.1 | Cleavage | \| Symbols: TBL22 \| TRICHOME BIREFRINGENCE-LIKE 22 \| chr3:10471842-10473754 REVERSE LENGTH=1382 |
| AT4G09960.4 | Cleavage | \| Symbols: STK \| K-box region and MADS-box transcription factor family protein \| chr4:6236375-6240932 REVERSE LENGTH=1238 |
| AT1G61790.1 | Cleavage | \| Symbols: \| Oligosaccharyltransferase complex/magnesium transporter family protein \| chr1:22814390-22815430 FORWARD LENGTH=1041 |
| AT5G23370.1 | Cleavage | \| Symbols: \| GRAM domain-containing protein / ABA-responsive protein-related \| chr5:7863331-7864271 REVERSE LENGTH=941 |
| AT3G56730.1 | Cleavage | \| Symbols: \| Putative endonuclease or glycosyl hydrolase \| chr3:21014187-21015078 FORWARD LENGTH=690 |
| AT1G14390.1 | Cleavage | \| Symbols: \| Leucine-rich repeat protein kinase family protein \| chr1:4924277-4926794 FORWARD LENGTH=2244 |
| AT2G18500.1 | Cleavage | \| Symbols: ATOFP7, OFP7 \| ovate family protein 7 \| chr2:8027034-8028251 FORWARD LENGTH=1218 |
| AT5G35935.1 | Cleavage | \| Symbols: \| transposable element gene \| chr5:14083388-14089891 REVERSE LENGTH=6504 |
| AT2G18090.1 | Cleavage | \| Symbols: \| PHD finger family protein / SWIB complex BAF60b domain-containing protein / GYF domain-containing protein \| chr2:7864314-7867766 FORWARD LENGTH=3111 |
| AT4G34980.1 | Cleavage | \| Symbols: SLP2 \| subtilisin-like serine protease 2 \| chr4:16656691-16659339 REVERSE LENGTH=2649 |
| AT2G01190.1 | Cleavage | \| Symbols: \| Octicosapeptide/Phox/Bem1p family protein \| chr2:114975-117640 FORWARD LENGTH=2555 |
| AT2G44090.1 | Cleavage | \| Symbols: \| Ankyrin repeat family protein \| chr2:18238360-18241293 REVERSE LENGTH=2536 |
| AT5G41410.1 | Cleavage | \| Symbols: BEL1 \| POX (plant homeobox) family protein \| chr5:16580044-16584008 FORWARD LENGTH=2454 |
| AT2G44090.2 | Cleavage | \| Symbols: \| Ankyrin repeat family protein \| chr2:18238360-18241316 REVERSE LENGTH=2436 |
| AT5G13990.1 | Cleavage | \| Symbols: ATEXO70C2, EXO70C2 \| exocyst subunit exo70 family protein C2 \| chr5:4514568-4516892 REVERSE LENGTH=2325 |
| AT3G61850.2 | Cleavage | \| Symbols: DAG1 \| Dof-type zinc finger DNA-binding family protein \| chr3:22895519-22897558 FORWARD LENGTH=1956 |
| AT4G13260.1 | Cleavage | \| Symbols: YUC2 \| Flavin-binding monooxygenase family protein \| chr4:7721618-7723989 REVERSE LENGTH=1843 |
| AT3G61850.4 | Cleavage | \| Symbols: DAG1 \| Dof-type zinc finger DNA-binding family protein \| chr3:22895268-22897488 FORWARD LENGTH=1540 |
| AT3G61850.1 | Cleavage | \| Symbols: DAG1 \| Dof-type zinc finger DNA-binding family protein \| chr3:22895268-22897558 FORWARD LENGTH=1526 |
| AT4G31410.2 | Cleavage | \| Symbols: \| Protein of unknown function (DUF1644) \| chr4:15243732-15245835 FORWARD LENGTH=1495 |
| AT4G10640.1 | Cleavage | \| Symbols: IQD16 \| IQ-domain 16 \| chr4:6571899-6574427 FORWARD LENGTH=1487 |
| AT4G31410.1 | Cleavage | \| Symbols: \| Protein of unknown function (DUF1644) \| chr4:15243723-15245838 FORWARD LENGTH=1431 |
| AT3G61850.3 | Cleavage | \| Symbols: DAG1 \| Dof-type zinc finger DNA-binding family protein \| chr3:22895729-22897558 FORWARD LENGTH=1401 |
| AT5G38360.1 | Cleavage | \| Symbols: \| alpha/beta-Hydrolases superfamily protein \| chr5:15331711-15333625 REVERSE LENGTH=1282 |
| AT2G31570.1 | Cleavage | \| Symbols: ATGPX2, GPX2 \| glutathione peroxidase 2 \| chr2:13437974-13439881 REVERSE LENGTH=853 |
| AT1G29520.1 | Cleavage | \| Symbols: \| AWPM-19-like family protein \| chr1:10323655-10324743 FORWARD LENGTH=775 |
| AT1G05065.1 | Cleavage | \| Symbols: CLE20 \| CLAVATA3/ESR-RELATED 20 \| chr1:1454969-1455220 REVERSE LENGTH=252 |
| AT1G19370.1 | Cleavage | \| Symbols: \| unknown protein; LOCATED IN: endoplasmic reticulum; EXPRESSED IN: 22 plant structures; EXPRESSED DURING: 13 growth stages; BEST Arabidopsis thaliana protein match is: unknown protein (TAIR:AT1G75140.1); Has 45 Blast hits to 43 proteins in 15 species: Archae - 0; Bacteria - 0; Metazoa - 0; Fungi - 0; Plants - 44; Viruses - 0; Other Eukaryotes - 1 (source: NCBI BLink). \| chr1:6692764-6694934 REVERSE LENGTH=2171 |
| AT2G25880.1 | Cleavage | \| Symbols: AtAUR2, AUR2 \| ataurora2 \| chr2:11034652-11036910 REVERSE LENGTH=1185 |
| AT2G25880.2 | Cleavage | \| Symbols: AtAUR2, AUR2 \| ataurora2 \| chr2:11034664-11036909 REVERSE LENGTH=1076 |
| AT5G46510.1 | Cleavage | \| Symbols: \| Disease resistance protein (TIR-NBS-LRR class) family \| chr5:18860451-18867013 FORWARD LENGTH=4708 |
| AT4G11830.2 | Cleavage | \| Symbols: PLDGAMMA2 \| phospholipase D gamma 2 \| chr4:7115794-7121245 REVERSE LENGTH=3858 |
| AT4G11830.1 | Cleavage | \| Symbols: PLDGAMMA2 \| phospholipase D gamma 2 \| chr4:7115736-7121245 REVERSE LENGTH=3820 |
| AT5G35180.4 | Cleavage | \| Symbols: \| Protein of unknown function (DUF1336) \| chr5:13424418-13433018 FORWARD LENGTH=2787 |
| AT5G35180.2 | Cleavage | \| Symbols: \| Protein of unknown function (DUF1336) \| chr5:13424418-13433018 FORWARD LENGTH=2776 |
| AT5G35180.1 | Cleavage | \| Symbols: \| Protein of unknown function (DUF1336) \| chr5:13424418-13433018 FORWARD LENGTH=2688 |
| AT1G14390.1 | Cleavage | \| Symbols: \| Leucine-rich repeat protein kinase family protein \| chr1:4924277-4926794 FORWARD LENGTH=2244 |
| AT3G58030.1 | Cleavage | \| Symbols: \| RING/U-box superfamily protein \| chr3:21484467-21487042 FORWARD LENGTH=2172 |
| AT2G44280.3 | Cleavage | \| Symbols: \| Major facilitator superfamily protein \| chr2:18301350-18304284 REVERSE LENGTH=2073 |
| AT3G58030.4 | Cleavage | \| Symbols: \| RING/U-box superfamily protein \| chr3:21484156-21487054 FORWARD LENGTH=2036 |
| AT3G58030.2 | Cleavage | \| Symbols: \| RING/U-box superfamily protein \| chr3:21484479-21487051 FORWARD LENGTH=1994 |
| AT2G44280.1 | Cleavage | \| Symbols: \| Major facilitator superfamily protein \| chr2:18301350-18304284 REVERSE LENGTH=1959 |
| AT2G44280.2 | Cleavage | \| Symbols: \| Major facilitator superfamily protein \| chr2:18301350-18304201 REVERSE LENGTH=1791 |
| AT3G48710.1 | Cleavage | \| Symbols: \| DEK domain-containing chromatin associated protein \| chr3:18040937-18044226 FORWARD LENGTH=1707 |
| AT3G11220.2 | Cleavage | \| Symbols: ELO1 \| Paxneb protein-related \| chr3:3513531-3516408 REVERSE LENGTH=1635 |
| AT3G11220.1 | Cleavage | \| Symbols: ELO1 \| Paxneb protein-related \| chr3:3513531-3516408 REVERSE LENGTH=1581 |
| AT4G18197.1 | Cleavage | \| Symbols: ATPUP7, PEX17, PUP7 \| purine permease 7 \| chr4:10071634-10073236 FORWARD LENGTH=1521 |
| AT3G26250.1 | Cleavage | \| Symbols: \| Cysteine/Histidine-rich C1 domain family protein \| chr3:9611696-9613168 REVERSE LENGTH=1473 |
| AT5G55610.1 | Cleavage | \| Symbols: \| unknown protein; LOCATED IN: mitochondrion, chloroplast, plastid, membrane; EXPRESSED IN: 25 plant structures; EXPRESSED DURING: 13 growth stages; Has 1807 Blast hits to 1807 proteins in 277 species: Archae - 0; Bacteria - 0; Metazoa - 736; Fungi - 347; Plants - 385; Viruses - 0; Other Eukaryotes - 339 (source: NCBI BLink). \| chr5:22525807-22528110 FORWARD LENGTH=1416 |
| AT3G18773.1 | Cleavage | \| Symbols: \| RING/U-box superfamily protein \| chr3:6465825-6467175 FORWARD LENGTH=1351 |
| AT3G17980.1 | Cleavage | \| Symbols: \| Calcium-dependent lipid-binding (CaLB domain) family protein \| chr3:6152121-6153434 FORWARD LENGTH=1072 |
| AT3G52130.1 | Cleavage | \| Symbols: \| Bifunctional inhibitor/lipid-transfer protein/seed storage 2S albumin superfamily protein \| chr3:19331932-19332473 REVERSE LENGTH=542 |
| AT1G53635.1 | Cleavage | \| Symbols: \| unknown protein; FUNCTIONS IN: molecular_function unknown; INVOLVED IN: biological_process unknown; LOCATED IN: cellular_component unknown; Has 6 Blast hits to 6 proteins in 1 species: Archae - 0; Bacteria - 0; Metazoa - 0; Fungi - 0; Plants - 6; Viruses - 0; Other Eukaryotes - 0 (source: NCBI BLink). \| chr1:20019293-20019799 REVERSE LENGTH=507 |
| AT5G48280.1 | Cleavage | \| Symbols: \| unknown protein; FUNCTIONS IN: molecular_function unknown; INVOLVED IN: biological_process unknown; LOCATED IN: endomembrane system; Has 1807 Blast hits to 1807 proteins in 277 species: Archae - 0; Bacteria - 0; Metazoa - 736; Fungi - 347; Plants - 385; Viruses - 0; Other Eukaryotes - 339 (source: NCBI BLink). \| chr5:19567064-19567688 REVERSE LENGTH=438 |
| AT5G29015.1 | Cleavage | \| Symbols: \| transposable element gene \| chr5:11031699-11038817 FORWARD LENGTH=7119 |
| AT1G46552.1 | Cleavage | \| Symbols: \| transposable element gene \| chr1:17247859-17252452 REVERSE LENGTH=4594 |
| AT1G58889.1 | Cleavage | \| Symbols: \| transposable element gene \| chr1:21797094-21801662 REVERSE LENGTH=4569 |
| AT1G59265.1 | Cleavage | \| Symbols: \| transposable element gene \| chr1:21833352-21837779 REVERSE LENGTH=4428 |
| AT4G21100.1 | Cleavage | \| Symbols: DDB1B \| damaged DNA binding protein 1B \| chr4:11258761-11265463 REVERSE LENGTH=3576 |
| AT3G09560.1 | Cleavage | \| Symbols: ATPAH1, PAH1 \| Lipin family protein \| chr3:2934617-2939456 REVERSE LENGTH=3511 |
| AT3G09560.3 | Cleavage | \| Symbols: \| Lipin family protein \| chr3:2934617-2939456 REVERSE LENGTH=3472 |
| AT3G09560.2 | Cleavage | \| Symbols: \| Lipin family protein \| chr3:2934617-2939456 REVERSE LENGTH=3465 |
| AT4G06528.1 | Cleavage | \| Symbols: \| transposable element gene \| chr4:3327079-3330294 FORWARD LENGTH=3216 |
| AT3G07660.1 | Cleavage | \| Symbols: \| Kinase-related protein of unknown function (DUF1296) \| chr3:2444778-2450561 REVERSE LENGTH=3212 |
| AT3G51620.1 | Cleavage | \| Symbols: \| PAP/OAS1 substrate-binding domain superfamily \| chr3:19143652-19148242 FORWARD LENGTH=2826 |
| AT3G51620.2 | Cleavage | \| Symbols: \| PAP/OAS1 substrate-binding domain superfamily \| chr3:19143652-19148056 FORWARD LENGTH=2809 |
| AT2G18940.1 | Cleavage | \| Symbols: \| Tetratricopeptide repeat (TPR)-like superfamily protein \| chr2:8203730-8206395 REVERSE LENGTH=2666 |
| AT4G30200.2 | Cleavage | \| Symbols: VEL1, VIL2 \| vernalization5/VIN3-like \| chr4:14786633-14790501 REVERSE LENGTH=2662 |
| AT4G30200.3 | Cleavage | \| Symbols: VEL1, VIL2 \| vernalization5/VIN3-like \| chr4:14786633-14790501 REVERSE LENGTH=2626 |
| AT4G30200.4 | Cleavage | \| Symbols: VEL1, VIL2 \| vernalization5/VIN3-like \| chr4:14786633-14790501 REVERSE LENGTH=2585 |
| AT3G45090.1 | Translation | \| Symbols: \| P-loop containing nucleoside triphosphate hydrolases superfamily protein \| chr3:16490600-16494762 REVERSE LENGTH=2535 |
| AT4G30200.1 | Cleavage | \| Symbols: VEL1, VIL2 \| vernalization5/VIN3-like \| chr4:14786633-14790501 REVERSE LENGTH=2532 |
| AT3G45090.2 | Translation | \| Symbols: \| P-loop containing nucleoside triphosphate hydrolases superfamily protein \| chr3:16490600-16494761 REVERSE LENGTH=2529 |
| AT3G01040.1 | Cleavage | \| Symbols: GAUT13 \| galacturonosyltransferase 13 \| chr3:8957-12444 FORWARD LENGTH=2502 |
| AT3G01040.2 | Cleavage | \| Symbols: GAUT13 \| galacturonosyltransferase 13 \| chr3:8957-12444 FORWARD LENGTH=2499 |
| AT5G51230.1 | Cleavage | \| Symbols: EMF2, VEF2, CYR1, AtEMF2 \| VEFS-Box of polycomb protein \| chr5:20823736-20829564 FORWARD LENGTH=2427 |
| AT5G51230.2 | Cleavage | \| Symbols: EMF2, VEF2, CYR1, AtEMF2 \| VEFS-Box of polycomb protein \| chr5:20823736-20829564 FORWARD LENGTH=2412 |
| AT5G49720.1 | Cleavage | \| Symbols: ATGH9A1, TSD1, DEC, KOR, RSW2, IRX2, KOR1, GH9A1 \| glycosyl hydrolase 9A1 \| chr5:20197391-20200288 REVERSE LENGTH=2360 |
| AT1G18390.1 | Cleavage | \| Symbols: \| Protein kinase superfamily protein \| chr1:6325642-6330079 FORWARD LENGTH=2325 |
| AT1G18150.3 | Cleavage | \| Symbols: ATMPK8 \| Protein kinase superfamily protein \| chr1:6244298-6247812 REVERSE LENGTH=2269 |
| AT1G53210.1 | Cleavage | \| Symbols: \| sodium/calcium exchanger family protein / calcium-binding EF hand family protein \| chr1:19844632-19847836 FORWARD LENGTH=2219 |
| AT1G18150.1 | Cleavage | \| Symbols: ATMPK8 \| Protein kinase superfamily protein \| chr1:6244384-6247769 REVERSE LENGTH=2137 |
| AT4G23230.1 | Cleavage | \| Symbols: CRK15 \| cysteine-rich RLK (RECEPTOR-like protein kinase) 15 \| chr4:12157569-12160270 REVERSE LENGTH=2133 |
| AT4G36980.2 | Cleavage | \| Symbols: \| FUNCTIONS IN: molecular_function unknown; INVOLVED IN: biological_process unknown; EXPRESSED IN: 23 plant structures; EXPRESSED DURING: 13 growth stages; CONTAINS InterPro DOMAIN/s: Splicing factor, suppressor of white apricot (InterPro:IPR019147); Has 5391 Blast hits to 4388 proteins in 280 species: Archae - 1; Bacteria - 114; Metazoa - 3014; Fungi - 666; Plants - 308; Viruses - 14; Other Eukaryotes - 1274 (source: NCBI BLink). \| chr4:17433644-17437069 REVERSE LENGTH=2119 |
| AT5G25620.2 | Cleavage | \| Symbols: YUC6 \| Flavin-binding monooxygenase family protein \| chr5:8935040-8938758 REVERSE LENGTH=2111 |
| AT5G25620.1 | Cleavage | \| Symbols: YUC6 \| Flavin-binding monooxygenase family protein \| chr5:8935109-8938666 REVERSE LENGTH=2107 |
| AT1G18150.2 | Cleavage | \| Symbols: ATMPK8 \| Protein kinase superfamily protein \| chr1:6244384-6247655 REVERSE LENGTH=2100 |
| AT5G65540.1 | Cleavage | \| Symbols: \| unknown protein; Has 6825 Blast hits to 3811 proteins in 289 species: Archae - 0; Bacteria - 184; Metazoa - 2139; Fungi - 760; Plants - 816; Viruses - 52; Other Eukaryotes - 2874 (source: NCBI BLink). \| chr5:26195689-26198322 FORWARD LENGTH=2049 |
| AT4G36980.4 | Cleavage | \| Symbols: \| FUNCTIONS IN: molecular_function unknown; INVOLVED IN: biological_process unknown; EXPRESSED IN: 22 plant structures; EXPRESSED DURING: 13 growth stages; CONTAINS InterPro DOMAIN/s: Splicing factor, suppressor of white apricot (InterPro:IPR019147). \| chr4:17433601-17437065 REVERSE LENGTH=1984 |
| AT4G36980.3 | Cleavage | \| Symbols: \| FUNCTIONS IN: molecular_function unknown; INVOLVED IN: biological_process unknown; EXPRESSED IN: 22 plant structures; EXPRESSED DURING: 13 growth stages; CONTAINS InterPro DOMAIN/s: Splicing factor, suppressor of white apricot (InterPro:IPR019147). \| chr4:17433601-17437065 REVERSE LENGTH=1978 |
| AT5G61400.1 | Cleavage | \| Symbols: \| Pentatricopeptide repeat (PPR) superfamily protein \| chr5:24681550-24683514 FORWARD LENGTH=1965 |
| AT4G36980.1 | Cleavage | \| Symbols: \| FUNCTIONS IN: molecular_function unknown; INVOLVED IN: biological_process unknown; EXPRESSED IN: 23 plant structures; EXPRESSED DURING: 13 growth stages; CONTAINS InterPro DOMAIN/s: Splicing factor, suppressor of white apricot (InterPro:IPR019147); Has 7672 Blast hits to 5479 proteins in 321 species: Archae - 0; Bacteria - 89; Metazoa - 5155; Fungi - 712; Plants - 341; Viruses - 39; Other Eukaryotes - 1336 (source: NCBI BLink). \| chr4:17433643-17437069 REVERSE LENGTH=1943 |
| AT1G52290.1 | Cleavage | \| Symbols: \| Protein kinase superfamily protein \| chr1:19470033-19472449 REVERSE LENGTH=1835 |
| AT1G80420.4 | Cleavage | \| Symbols: ATXRCC1 \| BRCT domain-containing DNA repair protein \| chr1:30235164-30237771 REVERSE LENGTH=1774 |
| AT2G15480.1 | Cleavage | \| Symbols: UGT73B5 \| UDP-glucosyl transferase 73B5 \| chr2:6758681-6760633 FORWARD LENGTH=1772 |
| AT2G15480.2 | Cleavage | \| Symbols: UGT73B5 \| UDP-glucosyl transferase 73B5 \| chr2:6758681-6763501 FORWARD LENGTH=1724 |
| AT1G55530.1 | Cleavage | \| Symbols: \| RING/U-box superfamily protein \| chr1:20729182-20731283 REVERSE LENGTH=1650 |
| AT4G34160.1 | Cleavage | \| Symbols: CYCD3;1, CYCD3 \| CYCLIN D3;1 \| chr4:16357639-16359555 FORWARD LENGTH=1646 |
| AT1G34270.1 | Cleavage | \| Symbols: \| Exostosin family protein \| chr1:12492411-12494544 REVERSE LENGTH=1624 |
| AT1G63170.1 | Cleavage | \| Symbols: \| Zinc finger, C3HC4 type (RING finger) family protein \| chr1:23425352-23427311 FORWARD LENGTH=1606 |
| AT1G30740.1 | Cleavage | \| Symbols: \| FAD-binding Berberine family protein \| chr1:10903029-10904630 FORWARD LENGTH=1602 |
| AT3G28300.1 | Cleavage | \| Symbols: AT14A \| Protein of unknown function (DUF677) \| chr3:10565737-10567600 FORWARD LENGTH=1575 |
| AT3G28290.1 | Cleavage | \| Symbols: AT14A \| Protein of unknown function (DUF677) \| chr3:10547504-10549367 FORWARD LENGTH=1575 |
| AT3G61850.4 | Cleavage | \| Symbols: DAG1 \| Dof-type zinc finger DNA-binding family protein \| chr3:22895268-22897488 FORWARD LENGTH=1540 |
| AT3G61850.1 | Cleavage | \| Symbols: DAG1 \| Dof-type zinc finger DNA-binding family protein \| chr3:22895268-22897558 FORWARD LENGTH=1526 |
| AT3G26790.1 | Cleavage | \| Symbols: FUS3 \| AP2/B3-like transcriptional factor family protein \| chr3:9853828-9855989 REVERSE LENGTH=1380 |
| AT5G37660.2 | Cleavage | \| Symbols: PDLP7 \| plasmodesmata-located protein 7 \| chr5:14959797-14961585 FORWARD LENGTH=1362 |
| AT5G61920.1 | Cleavage | \| Symbols: \| unknown protein; INVOLVED IN: biological_process unknown; LOCATED IN: chloroplast; EXPRESSED IN: 6 plant structures; EXPRESSED DURING: 4 anthesis, F mature embryo stage, petal differentiation and expansion stage, E expanded cotyledon stage, D bilateral stage; BEST Arabidopsis thaliana protein match is: unknown protein (TAIR:AT1G67170.1); Has 1807 Blast hits to 1807 proteins in 277 species: Archae - 0; Bacteria - 0; Metazoa - 736; Fungi - 347; Plants - 385; Viruses - 0; Other Eukaryotes - 339 (source: NCBI BLink). \| chr5:24864302-24866019 FORWARD LENGTH=1348 |
| AT3G13445.2 | Cleavage | \| Symbols: TBP1, TFIID-1 \| TATA binding protein 1 \| chr3:4379796-4382224 FORWARD LENGTH=1326 |
| AT1G17160.1 | Cleavage | \| Symbols: \| pfkB-like carbohydrate kinase family protein \| chr1:5867635-5869356 FORWARD LENGTH=1324 |
| AT5G61920.2 | Cleavage | \| Symbols: \| unknown protein; INVOLVED IN: biological_process unknown; LOCATED IN: chloroplast; EXPRESSED IN: 6 plant structures; EXPRESSED DURING: 4 anthesis, F mature embryo stage, petal differentiation and expansion stage, E expanded cotyledon stage, D bilateral stage; BEST Arabidopsis thaliana protein match is: unknown protein (TAIR:AT1G67170.1); Has 30201 Blast hits to 17322 proteins in 780 species: Archae - 12; Bacteria - 1396; Metazoa - 17338; Fungi - 3422; Plants - 5037; Viruses - 0; Other Eukaryotes - 2996 (source: NCBI BLink). \| chr5:24864305-24866019 FORWARD LENGTH=1298 |
| AT1G17160.2 | Cleavage | \| Symbols: \| pfkB-like carbohydrate kinase family protein \| chr1:5867635-5869356 FORWARD LENGTH=1292 |
| AT1G07320.1 | Cleavage | \| Symbols: RPL4 \| ribosomal protein L4 \| chr1:2249133-2250529 FORWARD LENGTH=1246 |
| AT1G07320.2 | Cleavage | \| Symbols: RPL4 \| ribosomal protein L4 \| chr1:2249133-2250527 FORWARD LENGTH=1238 |
| AT3G13445.1 | Cleavage | \| Symbols: TBP1, TFIID-1 \| TATA binding protein 1 \| chr3:4379793-4382224 FORWARD LENGTH=1223 |
| AT1G07320.4 | Cleavage | \| Symbols: RPL4 \| ribosomal protein L4 \| chr1:2249134-2250491 FORWARD LENGTH=1211 |
| AT4G32890.1 | Cleavage | \| Symbols: GATA9 \| GATA transcription factor 9 \| chr4:15875470-15876762 FORWARD LENGTH=1202 |
| AT5G59662.1 | Cleavage | \| Symbols: \| other RNA \| chr5:24039142-24040411 REVERSE LENGTH=1193 |
| AT4G12920.1 | Cleavage | \| Symbols: \| Eukaryotic aspartyl protease family protein \| chr4:7568286-7569455 FORWARD LENGTH=1170 |
| AT5G47310.1 | Cleavage | \| Symbols: \| PPPDE putative thiol peptidase family protein \| chr5:19200782-19202921 FORWARD LENGTH=1167 |
| AT5G37660.1 | Cleavage | \| Symbols: PDLP7 \| plasmodesmata-located protein 7 \| chr5:14959797-14961594 FORWARD LENGTH=1127 |
| AT1G07320.3 | Cleavage | \| Symbols: RPL4 \| ribosomal protein L4 \| chr1:2249134-2250164 FORWARD LENGTH=1031 |
| AT4G08330.1 | Cleavage | \| Symbols: \| unknown protein; FUNCTIONS IN: molecular_function unknown; INVOLVED IN: biological_process unknown; LOCATED IN: plasma membrane; EXPRESSED IN: 24 plant structures; EXPRESSED DURING: 14 growth stages; BEST Arabidopsis thaliana protein match is: unknown protein (TAIR:AT2G17705.1); Has 98 Blast hits to 98 proteins in 13 species: Archae - 0; Bacteria - 0; Metazoa - 0; Fungi - 0; Plants - 98; Viruses - 0; Other Eukaryotes - 0 (source: NCBI BLink). \| chr4:5255120-5256704 REVERSE LENGTH=938 |
| AT4G35840.1 | Cleavage | \| Symbols: \| RING/U-box superfamily protein \| chr4:16980961-16982365 FORWARD LENGTH=932 |
| AT1G10990.2 | Cleavage | \| Symbols: \| unknown protein; Has 6 Blast hits to 6 proteins in 2 species: Archae - 0; Bacteria - 0; Metazoa - 0; Fungi - 0; Plants - 6; Viruses - 0; Other Eukaryotes - 0 (source: NCBI BLink). \| chr1:3670278-3671187 REVERSE LENGTH=910 |
| AT1G62422.1 | Cleavage | \| Symbols: \| unknown protein; EXPRESSED IN: 22 plant structures; EXPRESSED DURING: 13 growth stages; BEST Arabidopsis thaliana protein match is: unknown protein (TAIR:AT1G12020.1); Has 87 Blast hits to 86 proteins in 14 species: Archae - 0; Bacteria - 0; Metazoa - 0; Fungi - 0; Plants - 87; Viruses - 0; Other Eukaryotes - 0 (source: NCBI BLink). \| chr1:23100139-23101033 REVERSE LENGTH=895 |
| AT5G03230.1 | Cleavage | \| Symbols: \| Protein of unknown function, DUF584 \| chr5:769514-770385 FORWARD LENGTH=872 |
| AT4G00695.2 | Cleavage | \| Symbols: \| FUNCTIONS IN: molecular_function unknown; INVOLVED IN: microtubule cytoskeleton organization; LOCATED IN: spindle pole, microtubule organizing center; EXPRESSED IN: cotyledon; CONTAINS InterPro DOMAIN/s: Spc97/Spc98 (InterPro:IPR007259); BEST Arabidopsis thaliana protein match is: Spc97 / Spc98 family of spindle pole body (SBP) component (TAIR:AT1G80245.3); Has 30201 Blast hits to 17322 proteins in 780 species: Archae - 12; Bacteria - 1396; Metazoa - 17338; Fungi - 3422; Plants - 5037; Viruses - 0; Other Eukaryotes - 2996 (source: NCBI BLink). \| chr4:284031-285142 FORWARD LENGTH=841 |
| AT1G10990.1 | Cleavage | \| Symbols: \| unknown protein; Has 4 Blast hits to 4 proteins in 1 species: Archae - 0; Bacteria - 0; Metazoa - 0; Fungi - 0; Plants - 4; Viruses - 0; Other Eukaryotes - 0 (source: NCBI BLink). \| chr1:3670278-3671187 REVERSE LENGTH=832 |
| AT4G00695.1 | Cleavage | \| Symbols: \| FUNCTIONS IN: molecular_function unknown; INVOLVED IN: microtubule cytoskeleton organization; LOCATED IN: spindle pole, microtubule organizing center; EXPRESSED IN: cotyledon; CONTAINS InterPro DOMAIN/s: Spc97/Spc98 (InterPro:IPR007259); BEST Arabidopsis thaliana protein match is: Spc97 / Spc98 family of spindle pole body (SBP) component (TAIR:AT1G80245.3); Has 27 Blast hits to 27 proteins in 8 species: Archae - 0; Bacteria - 0; Metazoa - 0; Fungi - 0; Plants - 27; Viruses - 0; Other Eukaryotes - 0 (source: NCBI BLink). \| chr4:283918-285142 FORWARD LENGTH=829 |
| AT4G08330.2 | Cleavage | \| Symbols: \| unknown protein. \| chr4:5255201-5256705 REVERSE LENGTH=825 |
| AT5G47570.1 | Cleavage | \| Symbols: \| unknown protein; Has 30201 Blast hits to 17322 proteins in 780 species: Archae - 12; Bacteria - 1396; Metazoa - 17338; Fungi - 3422; Plants - 5037; Viruses - 0; Other Eukaryotes - 2996 (source: NCBI BLink). \| chr5:19292608-19294737 REVERSE LENGTH=710 |
| AT5G02420.1 | Cleavage | \| Symbols: \| unknown protein; Has 90 Blast hits to 90 proteins in 10 species: Archae - 0; Bacteria - 0; Metazoa - 0; Fungi - 0; Plants - 90; Viruses - 0; Other Eukaryotes - 0 (source: NCBI BLink). \| chr5:523389-524076 FORWARD LENGTH=688 |
| AT1G15400.2 | Cleavage | \| Symbols: \| unknown protein; FUNCTIONS IN: molecular_function unknown; INVOLVED IN: biological_process unknown; LOCATED IN: plasma membrane; EXPRESSED IN: 23 plant structures; EXPRESSED DURING: 13 growth stages; BEST Arabidopsis thaliana protein match is: unknown protein (TAIR:AT1G80180.1); Has 82 Blast hits to 82 proteins in 16 species: Archae - 0; Bacteria - 4; Metazoa - 0; Fungi - 0; Plants - 75; Viruses - 0; Other Eukaryotes - 3 (source: NCBI BLink). \| chr1:5296225-5296902 REVERSE LENGTH=678 |
| AT2G17300.1 | Cleavage | \| Symbols: \| unknown protein; FUNCTIONS IN: molecular_function unknown; INVOLVED IN: biological_process unknown; LOCATED IN: chloroplast; EXPRESSED IN: 20 plant structures; EXPRESSED DURING: 13 growth stages; BEST Arabidopsis thaliana protein match is: unknown protein (TAIR:AT4G35320.1); Has 42 Blast hits to 42 proteins in 10 species: Archae - 0; Bacteria - 0; Metazoa - 0; Fungi - 0; Plants - 42; Viruses - 0; Other Eukaryotes - 0 (source: NCBI BLink). \| chr2:7522401-7523033 REVERSE LENGTH=633 |
| AT5G51620.3 | Cleavage | \| Symbols: \| Uncharacterised protein family (UPF0172) \| chr5:20967046-20968043 FORWARD LENGTH=369 |
| AT1G69825.1 | Translation | \| Symbols: \| Encodes a defensin-like (DEFL) family protein. \| chr1:26286496-26286929 FORWARD LENGTH=222 |
| AT1G22610.1 | Cleavage | \| Symbols: \| C2 calcium/lipid-binding plant phosphoribosyltransferase family protein \| chr1:7994291-7997588 FORWARD LENGTH=3298 |
| AT5G45310.1 | Cleavage | \| Symbols: \| unknown protein; LOCATED IN: endomembrane system; EXPRESSED IN: stem, inflorescence meristem, root, leaf; EXPRESSED DURING: LP.04 four leaves visible; Has 30201 Blast hits to 17322 proteins in 780 species: Archae - 12; Bacteria - 1396; Metazoa - 17338; Fungi - 3422; Plants - 5037; Viruses - 0; Other Eukaryotes - 2996 (source: NCBI BLink). \| chr5:18359275-18361114 REVERSE LENGTH=1289 |
| AT5G55370.1 | Cleavage | \| Symbols: \| MBOAT (membrane bound O-acyl transferase) family protein \| chr5:22445085-22446116 REVERSE LENGTH=1032 |
| AT3G07660.1 | Cleavage | \| Symbols: \| Kinase-related protein of unknown function (DUF1296) \| chr3:2444778-2450561 REVERSE LENGTH=3212 |
| AT1G18150.3 | Cleavage | \| Symbols: ATMPK8 \| Protein kinase superfamily protein \| chr1:6244298-6247812 REVERSE LENGTH=2269 |
| AT1G53210.1 | Cleavage | \| Symbols: \| sodium/calcium exchanger family protein / calcium-binding EF hand family protein \| chr1:19844632-19847836 FORWARD LENGTH=2219 |
| AT1G18150.1 | Cleavage | \| Symbols: ATMPK8 \| Protein kinase superfamily protein \| chr1:6244384-6247769 REVERSE LENGTH=2137 |
| AT1G18150.2 | Cleavage | \| Symbols: ATMPK8 \| Protein kinase superfamily protein \| chr1:6244384-6247655 REVERSE LENGTH=2100 |
| AT1G52980.1 | Cleavage | \| Symbols: \| GTP-binding family protein \| chr1:19737432-19740348 FORWARD LENGTH=1939 |
| AT5G55970.2 | Cleavage | \| Symbols: \| RING/U-box superfamily protein \| chr5:22667235-22669545 FORWARD LENGTH=1811 |
| AT5G55970.1 | Cleavage | \| Symbols: \| RING/U-box superfamily protein \| chr5:22667906-22669545 FORWARD LENGTH=1378 |
| AT1G76185.1 | Cleavage | \| Symbols: \| unknown protein; BEST Arabidopsis thaliana protein match is: unknown protein (TAIR:AT1G20460.1); Has 37 Blast hits to 37 proteins in 11 species: Archae - 0; Bacteria - 0; Metazoa - 0; Fungi - 0; Plants - 37; Viruses - 0; Other Eukaryotes - 0 (source: NCBI BLink). \| chr1:28590325-28591518 FORWARD LENGTH=854 |
| AT5G24350.2 | Cleavage | \| Symbols: \| FUNCTIONS IN: molecular_function unknown; INVOLVED IN: biological_process unknown; LOCATED IN: cellular_component unknown; EXPRESSED IN: 23 plant structures; EXPRESSED DURING: 13 growth stages; CONTAINS InterPro DOMAIN/s: Secretory pathway Sec39 (InterPro:IPR013244). \| chr5:8301105-8310895 FORWARD LENGTH=7577 |
| AT5G24350.1 | Cleavage | \| Symbols: \| CONTAINS InterPro DOMAIN/s: Secretory pathway Sec39 (InterPro:IPR013244); Has 1807 Blast hits to 1807 proteins in 277 species: Archae - 0; Bacteria - 0; Metazoa - 736; Fungi - 347; Plants - 385; Viruses - 0; Other Eukaryotes - 339 (source: NCBI BLink). \| chr5:8301105-8310895 FORWARD LENGTH=7517 |
| AT3G18290.1 | Cleavage | \| Symbols: EMB2454, BTS \| zinc finger protein-related \| chr3:6273995-6280510 FORWARD LENGTH=4414 |
| AT2G05935.1 | Cleavage | \| Symbols: \| transposable element gene \| chr2:2278835-2282788 FORWARD LENGTH=3954 |
| AT1G74360.1 | Cleavage | \| Symbols: \| Leucine-rich repeat protein kinase family protein \| chr1:27954231-27958114 FORWARD LENGTH=3592 |
| AT2G31010.1 | Cleavage | \| Symbols: \| Protein kinase superfamily protein \| chr2:13194185-13200128 FORWARD LENGTH=3054 |
| AT2G31010.2 | Cleavage | \| Symbols: \| Protein kinase superfamily protein \| chr2:13194185-13200128 FORWARD LENGTH=3049 |
| AT1G75730.1 | Cleavage | \| Symbols: \| unknown protein; Has 327 Blast hits to 272 proteins in 89 species: Archae - 0; Bacteria - 129; Metazoa - 68; Fungi - 14; Plants - 20; Viruses - 0; Other Eukaryotes - 96 (source: NCBI BLink). \| chr1:28435706-28439667 REVERSE LENGTH=2739 |
| AT2G27228.1 | Cleavage | \| Symbols: CPuORF6 \| conserved peptide upstream open reading frame 6 \| chr2:11650358-11654143 FORWARD LENGTH=2706 |
| AT2G27230.2 | Cleavage | \| Symbols: LHW \| transcription factor-related \| chr2:11650359-11654143 FORWARD LENGTH=2705 |
| AT1G80280.1 | Cleavage | \| Symbols: \| alpha/beta-Hydrolases superfamily protein \| chr1:30183541-30186586 REVERSE LENGTH=2687 |
| AT2G27230.1 | Cleavage | \| Symbols: LHW \| transcription factor-related \| chr2:11650358-11654116 FORWARD LENGTH=2641 |
| AT5G17725.1 | Cleavage | \| Symbols: \| transposable element gene \| chr5:5848296-5850501 REVERSE LENGTH=2206 |
| AT5G25050.1 | Cleavage | \| Symbols: \| Major facilitator superfamily protein \| chr5:8631680-8634027 FORWARD LENGTH=2041 |
| AT5G28910.1 | Cleavage | \| Symbols: \| unknown protein; FUNCTIONS IN: molecular_function unknown; INVOLVED IN: biological_process unknown; LOCATED IN: mitochondrion; BEST Arabidopsis thaliana protein match is: unknown protein (TAIR:AT5G28960.1); Has 35333 Blast hits to 34131 proteins in 2444 species: Archae - 798; Bacteria - 22429; Metazoa - 974; Fungi - 991; Plants - 531; Viruses - 0; Other Eukaryotes - 9610 (source: NCBI BLink). \| chr5:10930569-10932999 REVERSE LENGTH=2004 |
| AT5G28910.2 | Cleavage | \| Symbols: \| unknown protein; FUNCTIONS IN: molecular_function unknown; INVOLVED IN: biological_process unknown; BEST Arabidopsis thaliana protein match is: unknown protein (TAIR:AT5G28960.1); Has 82 Blast hits to 80 proteins in 14 species: Archae - 0; Bacteria - 0; Metazoa - 1; Fungi - 0; Plants - 78; Viruses - 0; Other Eukaryotes - 3 (source: NCBI BLink). \| chr5:10930586-10932999 REVERSE LENGTH=1992 |
| AT1G17370.1 | Cleavage | \| Symbols: UBP1B \| oligouridylate binding protein 1B \| chr1:5951542-5955037 REVERSE LENGTH=1772 |
| AT1G17370.2 | Cleavage | \| Symbols: UBP1B \| oligouridylate binding protein 1B \| chr1:5951542-5955037 REVERSE LENGTH=1763 |
| AT5G43420.1 | Cleavage | \| Symbols: \| RING/U-box superfamily protein \| chr5:17451669-17453124 FORWARD LENGTH=1456 |
| AT2G41330.1 | Cleavage | \| Symbols: \| Glutaredoxin family protein \| chr2:17226948-17228378 FORWARD LENGTH=1431 |
| AT5G46350.1 | Cleavage | \| Symbols: WRKY8, ATWRKY8 \| WRKY DNA-binding protein 8 \| chr5:18801218-18804043 REVERSE LENGTH=1308 |
| AT4G19390.1 | Cleavage | \| Symbols: \| Uncharacterised protein family (UPF0114) \| chr4:10574768-10576428 REVERSE LENGTH=1030 |
| AT1G65420.1 | Cleavage | \| Symbols: NPQ7 \| Protein of unknown function (DUF565) \| chr1:24297480-24298570 REVERSE LENGTH=979 |
| AT1G63380.1 | Cleavage | \| Symbols: \| NAD(P)-binding Rossmann-fold superfamily protein \| chr1:23505582-23506504 FORWARD LENGTH=849 |
| AT1G70581.1 | Cleavage | \| Symbols: \| other RNA \| chr1:26616267-26616996 FORWARD LENGTH=730 |
| AT5G22270.1 | Cleavage | \| Symbols: \| unknown protein; BEST Arabidopsis thaliana protein match is: unknown protein (TAIR:AT3G11600.1); Has 136 Blast hits to 136 proteins in 15 species: Archae - 0; Bacteria - 0; Metazoa - 0; Fungi - 0; Plants - 136; Viruses - 0; Other Eukaryotes - 0 (source: NCBI BLink). \| chr5:7372233-7373102 REVERSE LENGTH=646 |
| AT3G52800.1 | Cleavage | \| Symbols: \| A20/AN1-like zinc finger family protein \| chr3:19569311-19570682 FORWARD LENGTH=1129 |
| AT3G01040.1 | Cleavage | \| Symbols: GAUT13 \| galacturonosyltransferase 13 \| chr3:8957-12444 FORWARD LENGTH=2502 |
| AT3G01040.2 | Cleavage | \| Symbols: GAUT13 \| galacturonosyltransferase 13 \| chr3:8957-12444 FORWARD LENGTH=2499 |
| AT5G51230.1 | Cleavage | \| Symbols: EMF2, VEF2, CYR1, AtEMF2 \| VEFS-Box of polycomb protein \| chr5:20823736-20829564 FORWARD LENGTH=2427 |
| AT4G23230.1 | Cleavage | \| Symbols: CRK15 \| cysteine-rich RLK (RECEPTOR-like protein kinase) 15 \| chr4:12157569-12160270 REVERSE LENGTH=2133 |
| AT5G25620.2 | Cleavage | \| Symbols: YUC6 \| Flavin-binding monooxygenase family protein \| chr5:8935040-8938758 REVERSE LENGTH=2111 |
| AT5G25620.1 | Cleavage | \| Symbols: YUC6 \| Flavin-binding monooxygenase family protein \| chr5:8935109-8938666 REVERSE LENGTH=2107 |
| AT5G09690.1 | Cleavage | \| Symbols: ATMGT7, MGT7, MRS2-7 \| magnesium transporter 7 \| chr5:3000830-3003343 REVERSE LENGTH=1495 |
| AT5G09690.3 | Cleavage | \| Symbols: ATMGT7, MGT7, MRS2-7 \| magnesium transporter 7 \| chr5:3000809-3003262 REVERSE LENGTH=1413 |
| AT5G37660.2 | Cleavage | \| Symbols: PDLP7 \| plasmodesmata-located protein 7 \| chr5:14959797-14961585 FORWARD LENGTH=1362 |
| AT5G09690.4 | Cleavage | \| Symbols: ATMGT7, MGT7, MRS2-7 \| magnesium transporter 7 \| chr5:3000990-3003262 REVERSE LENGTH=1209 |
| AT5G37660.1 | Cleavage | \| Symbols: PDLP7 \| plasmodesmata-located protein 7 \| chr5:14959797-14961594 FORWARD LENGTH=1127 |
| AT1G17370.1 | Cleavage | \| Symbols: UBP1B \| oligouridylate binding protein 1B \| chr1:5951542-5955037 REVERSE LENGTH=1772 |
| AT1G17370.2 | Cleavage | \| Symbols: UBP1B \| oligouridylate binding protein 1B \| chr1:5951542-5955037 REVERSE LENGTH=1763 |
| AT5G46350.1 | Cleavage | \| Symbols: WRKY8, ATWRKY8 \| WRKY DNA-binding protein 8 \| chr5:18801218-18804043 REVERSE LENGTH=1308 |
| AT5G41810.1 | Cleavage | \| Symbols: \| unknown protein; BEST Arabidopsis thaliana protein match is: unknown protein (TAIR:AT1G64340.1); Has 876 Blast hits to 690 proteins in 132 species: Archae - 0; Bacteria - 38; Metazoa - 180; Fungi - 112; Plants - 59; Viruses - 2; Other Eukaryotes - 485 (source: NCBI BLink). \| chr5:16738067-16739663 FORWARD LENGTH=1291 |
| AT5G41810.2 | Cleavage | \| Symbols: \| unknown protein; BEST Arabidopsis thaliana protein match is: unknown protein (TAIR:AT1G64340.1); Has 514 Blast hits to 437 proteins in 98 species: Archae - 0; Bacteria - 21; Metazoa - 115; Fungi - 53; Plants - 52; Viruses - 5; Other Eukaryotes - 268 (source: NCBI BLink). \| chr5:16738062-16739663 FORWARD LENGTH=1269 |
| AT4G23850.1 | Cleavage | \| Symbols: LACS4 \| AMP-dependent synthetase and ligase family protein \| chr4:12403454-12408335 REVERSE LENGTH=2339 |
| AT5G55610.1 | Cleavage | \| Symbols: \| unknown protein; LOCATED IN: mitochondrion, chloroplast, plastid, membrane; EXPRESSED IN: 25 plant structures; EXPRESSED DURING: 13 growth stages; Has 1807 Blast hits to 1807 proteins in 277 species: Archae - 0; Bacteria - 0; Metazoa - 736; Fungi - 347; Plants - 385; Viruses - 0; Other Eukaryotes - 339 (source: NCBI BLink). \| chr5:22525807-22528110 FORWARD LENGTH=1416 |
| AT1G07320.1 | Cleavage | \| Symbols: RPL4 \| ribosomal protein L4 \| chr1:2249133-2250529 FORWARD LENGTH=1246 |
| AT1G07320.2 | Cleavage | \| Symbols: RPL4 \| ribosomal protein L4 \| chr1:2249133-2250527 FORWARD LENGTH=1238 |
| AT1G07320.4 | Cleavage | \| Symbols: RPL4 \| ribosomal protein L4 \| chr1:2249134-2250491 FORWARD LENGTH=1211 |
| AT1G07320.3 | Cleavage | \| Symbols: RPL4 \| ribosomal protein L4 \| chr1:2249134-2250164 FORWARD LENGTH=1031 |
| AT1G28310.1 | Cleavage | \| Symbols: \| Dof-type zinc finger DNA-binding family protein \| chr1:9912203-9913837 REVERSE LENGTH=1635 |
| AT1G28310.1 | Cleavage | \| Symbols: \| Dof-type zinc finger DNA-binding family protein \| chr1:9912203-9913837 REVERSE LENGTH=1635 |
| AT3G28150.1 | Cleavage | \| Symbols: TBL22 \| TRICHOME BIREFRINGENCE-LIKE 22 \| chr3:10471842-10473754 REVERSE LENGTH=1382 |
| AT2G44280.3 | Cleavage | \| Symbols: \| Major facilitator superfamily protein \| chr2:18301350-18304284 REVERSE LENGTH=2073 |
| AT2G44280.1 | Cleavage | \| Symbols: \| Major facilitator superfamily protein \| chr2:18301350-18304284 REVERSE LENGTH=1959 |
| AT2G44090.1 | Cleavage | \| Symbols: \| Ankyrin repeat family protein \| chr2:18238360-18241293 REVERSE LENGTH=2536 |
| AT5G51230.2 | Cleavage | \| Symbols: EMF2, VEF2, CYR1, AtEMF2 \| VEFS-Box of polycomb protein \| chr5:20823736-20829564 FORWARD LENGTH=2412 |
| AT4G13260.1 | Cleavage | \| Symbols: YUC2 \| Flavin-binding monooxygenase family protein \| chr4:7721618-7723989 REVERSE LENGTH=1843 |
| AT1G29520.1 | Cleavage | \| Symbols: \| AWPM-19-like family protein \| chr1:10323655-10324743 FORWARD LENGTH=775 |
| AT5G02420.1 | Cleavage | \| Symbols: \| unknown protein; Has 90 Blast hits to 90 proteins in 10 species: Archae - 0; Bacteria - 0; Metazoa - 0; Fungi - 0; Plants - 90; Viruses - 0; Other Eukaryotes - 0 (source: NCBI BLink). \| chr5:523389-524076 FORWARD LENGTH=688 |
| AT1G17370.1 | Cleavage | \| Symbols: UBP1B \| oligouridylate binding protein 1B \| chr1:5951542-5955037 REVERSE LENGTH=1772 |
| AT1G17370.2 | Cleavage | \| Symbols: UBP1B \| oligouridylate binding protein 1B \| chr1:5951542-5955037 REVERSE LENGTH=1763 |
| AT5G03230.1 | Cleavage | \| Symbols: \| Protein of unknown function, DUF584 \| chr5:769514-770385 FORWARD LENGTH=872 |
| AT1G15400.2 | Cleavage | \| Symbols: \| unknown protein; FUNCTIONS IN: molecular_function unknown; INVOLVED IN: biological_process unknown; LOCATED IN: plasma membrane; EXPRESSED IN: 23 plant structures; EXPRESSED DURING: 13 growth stages; BEST Arabidopsis thaliana protein match is: unknown protein (TAIR:AT1G80180.1); Has 82 Blast hits to 82 proteins in 16 species: Archae - 0; Bacteria - 4; Metazoa - 0; Fungi - 0; Plants - 75; Viruses - 0; Other Eukaryotes - 3 (source: NCBI BLink). \| chr1:5296225-5296902 REVERSE LENGTH=678 |
| AT5G41810.1 | Cleavage | \| Symbols: \| unknown protein; BEST Arabidopsis thaliana protein match is: unknown protein (TAIR:AT1G64340.1); Has 876 Blast hits to 690 proteins in 132 species: Archae - 0; Bacteria - 38; Metazoa - 180; Fungi - 112; Plants - 59; Viruses - 2; Other Eukaryotes - 485 (source: NCBI BLink). \| chr5:16738067-16739663 FORWARD LENGTH=1291 |
| AT5G41810.2 | Cleavage | \| Symbols: \| unknown protein; BEST Arabidopsis thaliana protein match is: unknown protein (TAIR:AT1G64340.1); Has 514 Blast hits to 437 proteins in 98 species: Archae - 0; Bacteria - 21; Metazoa - 115; Fungi - 53; Plants - 52; Viruses - 5; Other Eukaryotes - 268 (source: NCBI BLink). \| chr5:16738062-16739663 FORWARD LENGTH=1269 |
| AT4G19390.1 | Cleavage | \| Symbols: \| Uncharacterised protein family (UPF0114) \| chr4:10574768-10576428 REVERSE LENGTH=1030 |

| **cca-miR4391** | | |
| --- | --- | --- |
| Target_Acc. | Inhibition | Target_Desc. |
| AT4G28200.1 | Cleavage | \| Symbols: \| FUNCTIONS IN: molecular_function unknown; INVOLVED IN: RNA processing; LOCATED IN: intracellular; EXPRESSED IN: 22 plant structures; EXPRESSED DURING: 13 growth stages; CONTAINS InterPro DOMAIN/s: RNA-processing protein, HAT helix (InterPro:IPR003107), U3 small nucleolar RNA-associated protein 6 (InterPro:IPR013949); Has 492 Blast hits to 480 proteins in 206 species: Archae - 0; Bacteria - 2; Metazoa - 128; Fungi - 191; Plants - 60; Viruses - 0; Other Eukaryotes - 111 (source: NCBI BLink). \| chr4:13987602-13990450 REVERSE LENGTH=2181 |
| AT1G72460.1 | Cleavage | \| Symbols: \| Leucine-rich repeat protein kinase family protein \| chr1:27279510-27281533 FORWARD LENGTH=1935 |
| AT4G37760.1 | Cleavage | \| Symbols: SQE3 \| squalene epoxidase 3 \| chr4:17743887-17746678 FORWARD LENGTH=2028 |
| AT5G51500.1 | Cleavage | \| Symbols: \| Plant invertase/pectin methylesterase inhibitor superfamily \| chr5:20917929-20919838 REVERSE LENGTH=1623 |
| AT5G51490.1 | Cleavage | \| Symbols: \| Plant invertase/pectin methylesterase inhibitor superfamily \| chr5:20913553-20915606 REVERSE LENGTH=1738 |
| AT3G08960.1 | Cleavage | \| Symbols: \| ARM repeat superfamily protein \| chr3:2729819-2736884 REVERSE LENGTH=3591 |
| AT5G63930.1 | Cleavage | \| Symbols: \| Leucine-rich repeat protein kinase family protein \| chr5:25583006-25586499 FORWARD LENGTH=3416 |
| AT5G04240.1 | Cleavage | \| Symbols: ELF6 \| Zinc finger (C2H2 type) family protein / transcription factor jumonji (jmj) family protein \| chr5:1169544-1174878 FORWARD LENGTH=4263 |
| AT3G55320.1 | Cleavage | \| Symbols: PGP20 \| P-glycoprotein 20 \| chr3:20506992-20513393 REVERSE LENGTH=4626 |
| AT2G39480.1 | Cleavage | \| Symbols: PGP6 \| P-glycoprotein 6 \| chr2:16477792-16485039 REVERSE LENGTH=4893 |
| AT2G46100.1 | Translation | \| Symbols: \| Nuclear transport factor 2 (NTF2) family protein \| chr2:18953280-18954619 FORWARD LENGTH=921 |
| AT5G53420.2 | Cleavage | \| Symbols: \| CCT motif family protein \| chr5:21674389-21675733 FORWARD LENGTH=1055 |
| AT5G53420.1 | Cleavage | \| Symbols: \| CCT motif family protein \| chr5:21673525-21675733 FORWARD LENGTH=1217 |
| AT3G13222.1 | Cleavage | \| Symbols: GIP1 \| GBF-interacting protein 1 \| chr3:4251014-4254267 REVERSE LENGTH=2038 |
| AT2G41700.1 | Cleavage | \| Symbols: ABCA1, AtABCA1 \| ATP-binding cassette A1 \| chr2:17383042-17396110 REVERSE LENGTH=5846 |
| AT2G41700.2 | Cleavage | \| Symbols: ABCA1, AtABCA1 \| ATP-binding cassette A1 \| chr2:17382890-17395942 REVERSE LENGTH=5900 |
| AT2G24430.1 | Cleavage | \| Symbols: ANAC038, NAC038 \| NAC domain containing protein 38 \| chr2:10383517-10386625 REVERSE LENGTH=1025 |
| AT2G24430.2 | Cleavage | \| Symbols: ANAC039 \| NAC domain containing protein 38 \| chr2:10383517-10386480 REVERSE LENGTH=1045 |
| AT3G22380.2 | Cleavage | \| Symbols: TIC \| time for coffee \| chr3:7912905-7919510 FORWARD LENGTH=5487 |
| AT3G22380.1 | Cleavage | \| Symbols: TIC \| time for coffee \| chr3:7912905-7919484 FORWARD LENGTH=5534 |
| AT3G08820.1 | Cleavage | \| Symbols: \| Pentatricopeptide repeat (PPR) superfamily protein \| chr3:2677118-2679179 REVERSE LENGTH=2062 |
| AT1G35465.1 | Cleavage | \| Symbols: \| transposable element gene \| chr1:13046927-13047661 FORWARD LENGTH=735 |
| AT1G31300.1 | Cleavage | \| Symbols: \| TRAM, LAG1 and CLN8 (TLC) lipid-sensing domain containing protein \| chr1:11193693-11196312 FORWARD LENGTH=1410 |
| AT2G42380.1 | Cleavage | \| Symbols: ATBZIP34, BZIP34 \| Basic-leucine zipper (bZIP) transcription factor family protein \| chr2:17646900-17648945 REVERSE LENGTH=1523 |
| AT2G42380.2 | Cleavage | \| Symbols: ATBZIP34, BZIP34 \| Basic-leucine zipper (bZIP) transcription factor family protein \| chr2:17646900-17648945 REVERSE LENGTH=1556 |
| AT1G31300.2 | Cleavage | \| Symbols: \| TRAM, LAG1 and CLN8 (TLC) lipid-sensing domain containing protein \| chr1:11193635-11196312 FORWARD LENGTH=1610 |
| AT1G28310.2 | Cleavage | \| Symbols: \| Dof-type zinc finger DNA-binding family protein \| chr1:9911898-9913698 REVERSE LENGTH=1627 |
| AT1G28310.1 | Cleavage | \| Symbols: \| Dof-type zinc finger DNA-binding family protein \| chr1:9912203-9913837 REVERSE LENGTH=1635 |
| AT1G57820.2 | Cleavage | \| Symbols: VIM1, ORTH2 \| Zinc finger (C3HC4-type RING finger) family protein \| chr1:21414170-21417937 REVERSE LENGTH=2136 |
| AT1G57820.1 | Cleavage | \| Symbols: VIM1, ORTH2 \| Zinc finger (C3HC4-type RING finger) family protein \| chr1:21414170-21417946 REVERSE LENGTH=2154 |
| AT5G41000.1 | Cleavage | \| Symbols: YSL4 \| YELLOW STRIPE like 4 \| chr5:16420780-16423830 FORWARD LENGTH=2276 |
| AT2G07784.1 | Cleavage | \| Symbols: \| transposable element gene \| chr2:3317798-3318634 REVERSE LENGTH=837 |
| AT1G06910.1 | Cleavage | \| Symbols: TRFL7 \| TRF-like 7 \| chr1:2120991-2123748 FORWARD LENGTH=1528 |
| AT1G65730.1 | Cleavage | \| Symbols: YSL7 \| YELLOW STRIPE like 7 \| chr1:24442504-24446291 FORWARD LENGTH=2371 |
| AT4G29580.1 | Cleavage | \| Symbols: \| Cytidine/deoxycytidylate deaminase family protein \| chr4:14510147-14511043 FORWARD LENGTH=897 |
| AT4G29580.2 | Cleavage | \| Symbols: \| Cytidine/deoxycytidylate deaminase family protein \| chr4:14510147-14512186 FORWARD LENGTH=1338 |
| AT2G12420.1 | Cleavage | \| Symbols: \| transposable element gene \| chr2:5027376-5029321 REVERSE LENGTH=1946 |
| AT5G35070.2 | Translation | \| Symbols: \| pseudogene, hypothetical protein \| chr5:13336981-13338990 REVERSE LENGTH=2010 |
| AT2G37160.1 | Cleavage | \| Symbols: \| Transducin/WD40 repeat-like superfamily protein \| chr2:15608827-15612934 FORWARD LENGTH=2260 |
| AT2G37160.2 | Cleavage | \| Symbols: \| Transducin/WD40 repeat-like superfamily protein \| chr2:15608828-15612934 FORWARD LENGTH=2346 |
| AT4G20940.1 | Translation | \| Symbols: \| Leucine-rich receptor-like protein kinase family protein \| chr4:11202728-11206280 FORWARD LENGTH=3176 |
| AT2G21380.1 | Cleavage | \| Symbols: \| Kinesin motor family protein \| chr2:9141631-9149309 FORWARD LENGTH=3614 |
| AT5G56890.1 | Cleavage | \| Symbols: \| Protein kinase superfamily protein \| chr5:23010523-23015671 REVERSE LENGTH=3732 |
| AT2G22830.1 | Cleavage | \| Symbols: SQE2 \| squalene epoxidase 2 \| chr2:9723680-9726270 REVERSE LENGTH=2025 |
| AT3G31357.1 | Cleavage | \| Symbols: \| transposable element gene \| chr3:12713010-12713628 FORWARD LENGTH=619 |
| AT2G29850.1 | Cleavage | \| Symbols: \| unknown protein; BEST Arabidopsis thaliana protein match is: Plant protein 1589 of unknown function (TAIR:AT2G29605.1); Has 35333 Blast hits to 34131 proteins in 2444 species: Archae - 798; Bacteria - 22429; Metazoa - 974; Fungi - 991; Plants - 531; Viruses - 0; Other Eukaryotes - 9610 (source: NCBI BLink). \| chr2:12734725-12736394 FORWARD LENGTH=855 |
| AT1G13050.2 | Cleavage | \| Symbols: \| unknown protein; FUNCTIONS IN: molecular_function unknown; INVOLVED IN: biological_process unknown; LOCATED IN: endomembrane system; EXPRESSED IN: 14 plant structures; EXPRESSED DURING: 9 growth stages; BEST Arabidopsis thaliana protein match is: unknown protein (TAIR:AT3G26350.1); Has 260 Blast hits to 259 proteins in 20 species: Archae - 0; Bacteria - 0; Metazoa - 0; Fungi - 0; Plants - 260; Viruses - 0; Other Eukaryotes - 0 (source: NCBI BLink). \| chr1:4450238-4451742 FORWARD LENGTH=1149 |
| AT2G29605.1 | Cleavage | \| Symbols: \| Plant protein 1589 of unknown function \| chr2:12657909-12660087 REVERSE LENGTH=1206 |
| AT1G13050.1 | Cleavage | \| Symbols: \| unknown protein; BEST Arabidopsis thaliana protein match is: unknown protein (TAIR:AT3G26350.1); Has 538 Blast hits to 510 proteins in 88 species: Archae - 0; Bacteria - 23; Metazoa - 81; Fungi - 36; Plants - 361; Viruses - 8; Other Eukaryotes - 29 (source: NCBI BLink). \| chr1:4450382-4451742 FORWARD LENGTH=1361 |
| AT3G48300.1 | Translation | \| Symbols: CYP71A23 \| cytochrome P450, family 71, subfamily A, polypeptide 23 \| chr3:17885524-17887118 FORWARD LENGTH=1452 |
| AT4G23420.2 | Cleavage | \| Symbols: \| NAD(P)-binding Rossmann-fold superfamily protein \| chr4:12225606-12228593 FORWARD LENGTH=1530 |
| AT3G30710.1 | Cleavage | \| Symbols: \| transposable element gene \| chr3:12291812-12294137 REVERSE LENGTH=1572 |
| AT4G23420.3 | Cleavage | \| Symbols: \| NAD(P)-binding Rossmann-fold superfamily protein \| chr4:12225515-12228687 FORWARD LENGTH=1672 |
| AT4G23420.1 | Cleavage | \| Symbols: \| NAD(P)-binding Rossmann-fold superfamily protein \| chr4:12225607-12228718 FORWARD LENGTH=1788 |
| AT5G65800.1 | Cleavage | \| Symbols: ACS5, CIN5, ETO2, ATACS5 \| ACC synthase 5 \| chr5:26330746-26332812 REVERSE LENGTH=1877 |
| AT5G45840.1 | Cleavage | \| Symbols: \| Leucine-rich repeat protein kinase family protein \| chr5:18594080-18597221 REVERSE LENGTH=2007 |
| AT2G10080.1 | Cleavage | \| Symbols: \| transposable element gene \| chr2:3820974-3822999 REVERSE LENGTH=2026 |
| AT5G65690.1 | Cleavage | \| Symbols: PCK2, PEPCK \| phosphoenolpyruvate carboxykinase 2 \| chr5:26266340-26269616 FORWARD LENGTH=2176 |
| AT4G06557.1 | Cleavage | \| Symbols: \| transposable element gene \| chr4:3476600-3479077 REVERSE LENGTH=2478 |
| AT5G45840.2 | Cleavage | \| Symbols: \| Leucine-rich repeat protein kinase family protein \| chr5:18593917-18597995 REVERSE LENGTH=2675 |
| AT2G12850.1 | Cleavage | \| Symbols: \| transposable element gene \| chr2:5275903-5278679 FORWARD LENGTH=2777 |
| AT5G49890.1 | Cleavage | \| Symbols: CLC-C, ATCLC-C \| chloride channel C \| chr5:20288119-20292350 REVERSE LENGTH=2836 |
| AT2G31981.1 | Cleavage | \| Symbols: \| unknown protein; Has 30201 Blast hits to 17322 proteins in 780 species: Archae - 12; Bacteria - 1396; Metazoa - 17338; Fungi - 3422; Plants - 5037; Viruses - 0; Other Eukaryotes - 2996 (source: NCBI BLink). \| chr2:13609981-13610094 FORWARD LENGTH=114 |
| AT1G35115.1 | Cleavage | \| Symbols: \| transposable element gene \| chr1:12842158-12845455 FORWARD LENGTH=3298 |
| AT1G75640.1 | Cleavage | \| Symbols: \| Leucine-rich receptor-like protein kinase family protein \| chr1:28403466-28407051 REVERSE LENGTH=3586 |
| AT5G19165.1 | Cleavage | \| Symbols: \| transposable element gene \| chr5:6433770-6438242 REVERSE LENGTH=4473 |
| AT4G08375.1 | Cleavage | \| Symbols: \| transposable element gene \| chr4:5303928-5308612 REVERSE LENGTH=4685 |
| AT1G21722.1 | Cleavage | \| Symbols: \| unknown protein; FUNCTIONS IN: molecular_function unknown; INVOLVED IN: biological_process unknown; LOCATED IN: cellular_component unknown; BEST Arabidopsis thaliana protein match is: unknown protein (TAIR:AT1G78922.1); Has 47 Blast hits to 47 proteins in 13 species: Archae - 0; Bacteria - 0; Metazoa - 1; Fungi - 1; Plants - 42; Viruses - 0; Other Eukaryotes - 3 (source: NCBI BLink). \| chr1:7628456-7629252 FORWARD LENGTH=702 |
| AT2G33710.2 | Cleavage | \| Symbols: \| Integrase-type DNA-binding superfamily protein \| chr2:14258809-14260739 REVERSE LENGTH=939 |
| AT1G34760.2 | Cleavage | \| Symbols: GRF11, GF14 OMICRON, RHS5 \| general regulatory factor 11 \| chr1:12743821-12745648 REVERSE LENGTH=964 |
| AT1G34760.1 | Cleavage | \| Symbols: GRF11, GF14 OMICRON, RHS5 \| general regulatory factor 11 \| chr1:12743848-12745648 REVERSE LENGTH=1027 |
| AT5G50130.2 | Cleavage | \| Symbols: \| NAD(P)-binding Rossmann-fold superfamily protein \| chr5:20389997-20393184 FORWARD LENGTH=1213 |
| AT5G50130.1 | Cleavage | \| Symbols: \| NAD(P)-binding Rossmann-fold superfamily protein \| chr5:20390015-20393263 FORWARD LENGTH=1281 |
| AT2G10940.1 | Cleavage | \| Symbols: \| Bifunctional inhibitor/lipid-transfer protein/seed storage 2S albumin superfamily protein \| chr2:4310412-4312103 REVERSE LENGTH=1338 |
| AT2G17570.1 | Cleavage | \| Symbols: \| Undecaprenyl pyrophosphate synthetase family protein \| chr2:7644053-7646254 FORWARD LENGTH=1413 |
| AT2G33710.1 | Cleavage | \| Symbols: \| Integrase-type DNA-binding superfamily protein \| chr2:14258509-14260733 REVERSE LENGTH=1500 |
| AT3G06380.1 | Cleavage | \| Symbols: ATTLP9, TLP9 \| tubby-like protein 9 \| chr3:1936124-1938221 FORWARD LENGTH=1596 |
| AT4G06598.1 | Cleavage | \| Symbols: \| BEST Arabidopsis thaliana protein match is: Basic-leucine zipper (bZIP) transcription factor family protein (TAIR:AT1G58110.2); Has 392 Blast hits to 388 proteins in 30 species: Archae - 0; Bacteria - 0; Metazoa - 15; Fungi - 8; Plants - 363; Viruses - 0; Other Eukaryotes - 6 (source: NCBI BLink). \| chr4:3663485-3665981 FORWARD LENGTH=1658 |
| AT2G10940.2 | Cleavage | \| Symbols: \| Bifunctional inhibitor/lipid-transfer protein/seed storage 2S albumin superfamily protein \| chr2:4310412-4312103 REVERSE LENGTH=1692 |
| AT3G20040.1 | Cleavage | \| Symbols: ATHXK4, HKL2 \| Hexokinase \| chr3:6994894-6998171 FORWARD LENGTH=2039 |
| AT5G05490.1 | Cleavage | \| Symbols: SYN1, ATREC8, REC8 \| Rad21/Rec8-like family protein \| chr5:1624206-1629234 FORWARD LENGTH=2093 |
| AT5G05490.2 | Cleavage | \| Symbols: SYN1, DIF1 \| Rad21/Rec8-like family protein \| chr5:1624563-1629269 FORWARD LENGTH=2128 |
| AT1G48370.1 | Cleavage | \| Symbols: YSL8 \| YELLOW STRIPE like 8 \| chr1:17874512-17877397 FORWARD LENGTH=2364 |
| AT3G17650.1 | Cleavage | \| Symbols: YSL5, PDE321 \| YELLOW STRIPE like 5 \| chr3:6034198-6037229 FORWARD LENGTH=2396 |
| AT5G12440.2 | Cleavage | \| Symbols: \| CCCH-type zinc fingerfamily protein with RNA-binding domain \| chr5:4035322-4041334 REVERSE LENGTH=2524 |
| AT5G12440.3 | Cleavage | \| Symbols: \| CCCH-type zinc fingerfamily protein with RNA-binding domain \| chr5:4035322-4041334 REVERSE LENGTH=2526 |
| AT5G12440.1 | Cleavage | \| Symbols: \| CCCH-type zinc fingerfamily protein with RNA-binding domain \| chr5:4035294-4038795 REVERSE LENGTH=2573 |
| AT4G19960.2 | Translation | \| Symbols: KUP9 \| K+ uptake permease 9 \| chr4:10813548-10817070 FORWARD LENGTH=2756 |
| AT4G19960.1 | Translation | \| Symbols: KUP9, ATKUP9, HAK9, KT9 \| K+ uptake permease 9 \| chr4:10813548-10817070 FORWARD LENGTH=2804 |
| AT1G79840.2 | Cleavage | \| Symbols: GL2 \| HD-ZIP IV family of homeobox-leucine zipper protein with lipid-binding START domain \| chr1:30036956-30041440 FORWARD LENGTH=2895 |
| AT3G42270.1 | Cleavage | \| Symbols: \| transposable element gene \| chr3:14441782-14444827 FORWARD LENGTH=3046 |
| AT5G33990.1 | Cleavage | \| Symbols: \| transposable element gene \| chr5:12791432-12794551 REVERSE LENGTH=3120 |
| AT4G38590.2 | Cleavage | \| Symbols: BGAL14 \| beta-galactosidase 14 \| chr4:18036116-18040928 FORWARD LENGTH=3159 |
| AT4G38590.1 | Cleavage | \| Symbols: BGAL14 \| beta-galactosidase 14 \| chr4:18036092-18040928 FORWARD LENGTH=3187 |
| AT5G61980.1 | Translation | \| Symbols: AGD1 \| ARF-GAP domain 1 \| chr5:24894472-24899973 FORWARD LENGTH=3282 |
| AT3G45775.1 | Cleavage | \| Symbols: \| transposable element gene \| chr3:16809432-16813871 REVERSE LENGTH=4440 |
| AT1G08600.2 | Translation | \| Symbols: ATRX \| P-loop containing nucleoside triphosphate hydrolases superfamily protein \| chr1:2723824-2733613 FORWARD LENGTH=4869 |
| AT4G30150.1 | Cleavage | \| Symbols: \| CONTAINS InterPro DOMAIN/s: Nucleolar 27S pre-rRNA processing, Urb2/Npa2 (InterPro:IPR018849); Has 58 Blast hits to 49 proteins in 21 species: Archae - 0; Bacteria - 2; Metazoa - 2; Fungi - 0; Plants - 44; Viruses - 3; Other Eukaryotes - 7 (source: NCBI BLink). \| chr4:14742452-14749987 FORWARD LENGTH=6253 |
| AT3G57920.1 | Cleavage | \| Symbols: SPL15 \| squamosa promoter binding protein-like 15 \| chr3:21444321-21446035 REVERSE LENGTH=1377 |
| AT1G70550.2 | Cleavage | \| Symbols: \| Protein of Unknown Function (DUF239) \| chr1:26597152-26600219 FORWARD LENGTH=1754 |
| AT4G33495.1 | Cleavage | \| Symbols: RPD1 \| Ubiquitin carboxyl-terminal hydrolase family protein \| chr4:16110712-16112597 REVERSE LENGTH=1786 |
| AT1G70550.1 | Cleavage | \| Symbols: \| Protein of Unknown Function (DUF239) \| chr1:26597313-26600219 FORWARD LENGTH=1926 |
| AT3G17040.2 | Translation | \| Symbols: HCF107 \| high chlorophyll fluorescent 107 \| chr3:5809266-5812678 REVERSE LENGTH=2042 |
| AT3G17040.1 | Translation | \| Symbols: HCF107 \| high chlorophyll fluorescent 107 \| chr3:5809266-5812678 REVERSE LENGTH=2144 |
| AT3G10550.1 | Cleavage | \| Symbols: MTM1, AtMTM1 \| Myotubularin-like phosphatases II superfamily \| chr3:3292899-3298522 REVERSE LENGTH=2904 |
| AT5G39500.1 | Cleavage | \| Symbols: GNL1, ERMO1 \| GNOM-like 1 \| chr5:15815274-15820045 FORWARD LENGTH=4467 |
| AT1G12890.1 | Cleavage | \| Symbols: \| Integrase-type DNA-binding superfamily protein \| chr1:4391734-4392393 FORWARD LENGTH=660 |
| AT2G38660.4 | Cleavage | \| Symbols: \| Amino acid dehydrogenase family protein \| chr2:16165958-16168386 FORWARD LENGTH=1187 |
| AT2G38660.1 | Cleavage | \| Symbols: \| Amino acid dehydrogenase family protein \| chr2:16165958-16168386 FORWARD LENGTH=1335 |
| AT2G38660.2 | Cleavage | \| Symbols: \| Amino acid dehydrogenase family protein \| chr2:16165952-16168386 FORWARD LENGTH=1354 |
| AT2G38660.3 | Cleavage | \| Symbols: \| Amino acid dehydrogenase family protein \| chr2:16165952-16168386 FORWARD LENGTH=1356 |
| AT5G27450.3 | Cleavage | \| Symbols: MK \| mevalonate kinase \| chr5:9690607-9693289 FORWARD LENGTH=1423 |
| AT5G27450.1 | Cleavage | \| Symbols: MVK, MK \| mevalonate kinase \| chr5:9690607-9693225 FORWARD LENGTH=1459 |
| AT3G24230.1 | Cleavage | \| Symbols: \| Pectate lyase family protein \| chr3:8774616-8777407 FORWARD LENGTH=1481 |
| AT5G64370.1 | Translation | \| Symbols: BETA-UP, PYD3 \| beta-ureidopropionase \| chr5:25739178-25741212 FORWARD LENGTH=1494 |
| AT2G31210.1 | Cleavage | \| Symbols: \| basic helix-loop-helix (bHLH) DNA-binding superfamily protein \| chr2:13296143-13298277 FORWARD LENGTH=1693 |
| AT5G27450.2 | Cleavage | \| Symbols: MVK, MK \| mevalonate kinase \| chr5:9690740-9693225 FORWARD LENGTH=1698 |
| AT2G41700.1 | Cleavage | \| Symbols: ABCA1, AtABCA1 \| ATP-binding cassette A1 \| chr2:17383042-17396110 REVERSE LENGTH=5846 |
| AT2G17470.1 | Cleavage | \| Symbols: \| Aluminium activated malate transporter family protein \| chr2:7584505-7588008 REVERSE LENGTH=1684 |
| AT1G64960.1 | Cleavage | \| Symbols: \| ARM repeat superfamily protein \| chr1:24129918-24134204 FORWARD LENGTH=3767 |
| AT4G19960.2 | Cleavage | \| Symbols: KUP9 \| K+ uptake permease 9 \| chr4:10813548-10817070 FORWARD LENGTH=2756 |
| AT4G19960.1 | Cleavage | \| Symbols: KUP9, ATKUP9, HAK9, KT9 \| K+ uptake permease 9 \| chr4:10813548-10817070 FORWARD LENGTH=2804 |
| AT1G22130.1 | Cleavage | \| Symbols: AGL104 \| AGAMOUS-like 104 \| chr1:7812387-7814264 REVERSE LENGTH=1013 |
| AT5G43280.1 | Cleavage | \| Symbols: ATDCI1, DCI1 \| delta(3,5),delta(2,4)-dienoyl-CoA isomerase 1 \| chr5:17367888-17369314 FORWARD LENGTH=1097 |
| AT4G30980.1 | Cleavage | \| Symbols: LRL2 \| LJRHL1-like 2 \| chr4:15079243-15081657 REVERSE LENGTH=1230 |
| AT1G28640.1 | Cleavage | \| Symbols: \| GDSL-like Lipase/Acylhydrolase superfamily protein \| chr1:10067563-10069177 REVERSE LENGTH=1238 |
| AT5G43280.2 | Cleavage | \| Symbols: ATDCI1, DCI1 \| delta(3,5),delta(2,4)-dienoyl-CoA isomerase 1 \| chr5:17367888-17369399 FORWARD LENGTH=1268 |
| AT1G23340.2 | Cleavage | \| Symbols: \| Protein of Unknown Function (DUF239) \| chr1:8283644-8286528 REVERSE LENGTH=1289 |
| AT1G28660.2 | Cleavage | \| Symbols: \| GDSL-like Lipase/Acylhydrolase superfamily protein \| chr1:10071712-10073388 REVERSE LENGTH=1310 |
| AT1G28660.1 | Cleavage | \| Symbols: \| GDSL-like Lipase/Acylhydrolase superfamily protein \| chr1:10071712-10073411 REVERSE LENGTH=1336 |
| AT1G28670.1 | Cleavage | \| Symbols: ARAB-1 \| GDSL-like Lipase/Acylhydrolase superfamily protein \| chr1:10074507-10076344 REVERSE LENGTH=1411 |
| AT1G04620.1 | Cleavage | \| Symbols: \| coenzyme F420 hydrogenase family / dehydrogenase, beta subunit family \| chr1:1282741-1286584 REVERSE LENGTH=1609 |
| AT4G29240.1 | Cleavage | \| Symbols: \| Leucine-rich repeat (LRR) family protein \| chr4:14418605-14420250 FORWARD LENGTH=1646 |
| AT2G21120.1 | Cleavage | \| Symbols: \| Protein of unknown function (DUF803) \| chr2:9051748-9054613 REVERSE LENGTH=1689 |
| AT1G23340.1 | Cleavage | \| Symbols: \| Protein of Unknown Function (DUF239) \| chr1:8283896-8286832 REVERSE LENGTH=1796 |
| AT4G00350.1 | Translation | \| Symbols: \| MATE efflux family protein \| chr4:151919-154130 FORWARD LENGTH=1830 |
| AT1G31970.1 | Cleavage | \| Symbols: STRS1 \| DEA(D/H)-box RNA helicase family protein \| chr1:11479866-11482890 FORWARD LENGTH=1852 |
| AT5G05010.2 | Cleavage | \| Symbols: \| clathrin adaptor complexes medium subunit family protein \| chr5:1476673-1480197 FORWARD LENGTH=2007 |
| AT1G24160.1 | Cleavage | \| Symbols: \| unknown protein; BEST Arabidopsis thaliana protein match is: unknown protein (TAIR:AT1G70100.3); Has 5230 Blast hits to 4162 proteins in 477 species: Archae - 30; Bacteria - 540; Metazoa - 2022; Fungi - 429; Plants - 280; Viruses - 32; Other Eukaryotes - 1897 (source: NCBI BLink). \| chr1:8553420-8556290 REVERSE LENGTH=2037 |
| AT5G05010.1 | Cleavage | \| Symbols: \| clathrin adaptor complexes medium subunit family protein \| chr5:1476583-1480197 FORWARD LENGTH=2059 |
| AT5G25620.1 | Cleavage | \| Symbols: YUC6 \| Flavin-binding monooxygenase family protein \| chr5:8935109-8938666 REVERSE LENGTH=2107 |
| AT5G25620.2 | Cleavage | \| Symbols: YUC6 \| Flavin-binding monooxygenase family protein \| chr5:8935040-8938758 REVERSE LENGTH=2111 |
| AT5G18550.1 | Cleavage | \| Symbols: \| Zinc finger C-x8-C-x5-C-x3-H type family protein \| chr5:6160178-6163130 FORWARD LENGTH=2136 |
| ATCG00350.1 | Cleavage | \| Symbols: PSAA \| Photosystem I, PsaA/PsaB protein \| chrC:39605-41857 REVERSE LENGTH=2253 |
| AT3G18230.1 | Cleavage | \| Symbols: \| Octicosapeptide/Phox/Bem1p family protein \| chr3:6251483-6254221 FORWARD LENGTH=2263 |
| AT1G24160.2 | Cleavage | \| Symbols: \| unknown protein; FUNCTIONS IN: molecular_function unknown; INVOLVED IN: biological_process unknown; LOCATED IN: cellular_component unknown; EXPRESSED IN: guard cell; BEST Arabidopsis thaliana protein match is: unknown protein (TAIR:AT1G70100.3). \| chr1:8553411-8556555 REVERSE LENGTH=2316 |
| AT1G72990.2 | Translation | \| Symbols: BGAL17 \| beta-galactosidase 17 \| chr1:27457282-27462392 REVERSE LENGTH=2529 |
| AT1G72990.1 | Translation | \| Symbols: BGAL17 \| beta-galactosidase 17 \| chr1:27457266-27462389 REVERSE LENGTH=2529 |
| AT5G62090.2 | Cleavage | \| Symbols: SLK2 \| SEUSS-like 2 \| chr5:24934963-24939109 REVERSE LENGTH=2828 |
| AT1G20780.1 | Translation | \| Symbols: PUB44, ATPUB44, SAUL1 \| senescence-associated E3 ubiquitin ligase 1 \| chr1:7217128-7220784 FORWARD LENGTH=2877 |
| AT4G07339.1 | Cleavage | \| Symbols: \| transposable element gene \| chr4:4157880-4160875 REVERSE LENGTH=2996 |
| AT3G29641.1 | Cleavage | \| Symbols: \| transposable element gene \| chr3:11490749-11493971 REVERSE LENGTH=3223 |
| AT5G62090.1 | Cleavage | \| Symbols: SLK2 \| SEUSS-like 2 \| chr5:24934963-24939060 REVERSE LENGTH=3229 |
| AT5G49030.2 | Cleavage | \| Symbols: OVA2 \| tRNA synthetase class I (I, L, M and V) family protein \| chr5:19876166-19882462 REVERSE LENGTH=3238 |
| AT2G21440.1 | Cleavage | \| Symbols: \| RNA-binding (RRM/RBD/RNP motifs) family protein \| chr2:9173560-9179166 REVERSE LENGTH=3265 |
| AT1G10130.1 | Cleavage | \| Symbols: ECA3, ATECA3 \| endoplasmic reticulum-type calcium-transporting ATPase 3 \| chr1:3310930-3322213 FORWARD LENGTH=3478 |
| AT5G49030.1 | Cleavage | \| Symbols: OVA2 \| tRNA synthetase class I (I, L, M and V) family protein \| chr5:19874944-19882462 REVERSE LENGTH=3600 |
| AT5G49030.3 | Cleavage | \| Symbols: OVA2 \| tRNA synthetase class I (I, L, M and V) family protein \| chr5:19874944-19883297 REVERSE LENGTH=4033 |
| AT1G04580.1 | Cleavage | \| Symbols: AAO4, ATAO-4, ATAO2, AO4 \| aldehyde oxidase 4 \| chr1:1252122-1257510 REVERSE LENGTH=4104 |
| AT5G24710.1 | Cleavage | \| Symbols: \| Transducin/WD40 repeat-like superfamily protein \| chr5:8458847-8468173 REVERSE LENGTH=4688 |
| AT1G08730.1 | Cleavage | \| Symbols: XIC, ATXIC \| Myosin family protein with Dil domain \| chr1:2779963-2788447 FORWARD LENGTH=4739 |
| AT5G33310.1 | Cleavage | \| Symbols: \| pseudogene, hypothetical protein, predicted proteins - Arabidopsis thaliana \| chr5:12577321-12582199 REVERSE LENGTH=4879 |
| AT5G28692.1 | Cleavage | \| Symbols: \| transposable element gene \| chr5:10728378-10733393 REVERSE LENGTH=5016 |
| AT2G01029.1 | Translation | \| Symbols: \| transposable element gene \| chr2:28465-38652 FORWARD LENGTH=10188 |
| AT4G33560.1 | Cleavage | \| Symbols: \| Wound-responsive family protein \| chr4:16135400-16135914 FORWARD LENGTH=515 |
| AT3G63160.1 | Cleavage | \| Symbols: \| FUNCTIONS IN: molecular_function unknown; INVOLVED IN: biological_process unknown; LOCATED IN: chloroplast outer membrane, thylakoid, chloroplast thylakoid membrane, chloroplast, chloroplast envelope; EXPRESSED IN: 21 plant structures; EXPRESSED DURING: 13 growth stages; BEST Arabidopsis thaliana protein match is: outer envelope membrane protein 7 (TAIR:AT3G52420.1); Has 26 Blast hits to 26 proteins in 8 species: Archae - 0; Bacteria - 0; Metazoa - 0; Fungi - 0; Plants - 26; Viruses - 0; Other Eukaryotes - 0 (source: NCBI BLink). \| chr3:23333513-23334034 REVERSE LENGTH=522 |
| AT4G04100.1 | Cleavage | \| Symbols: \| transposable element gene \| chr4:1966652-1967584 FORWARD LENGTH=933 |
| AT3G60176.1 | Cleavage | \| Symbols: \| other RNA \| chr3:22240155-22241164 REVERSE LENGTH=1010 |
| AT4G19150.1 | Cleavage | \| Symbols: \| Ankyrin repeat family protein \| chr4:10471331-10472742 REVERSE LENGTH=1044 |
| AT1G51530.1 | Cleavage | \| Symbols: \| RNA-binding (RRM/RBD/RNP motifs) family protein \| chr1:19109854-19112152 FORWARD LENGTH=1176 |
| AT3G18030.1 | Translation | \| Symbols: ATHAL3A, HAL3A, HAL3, ATHAL3 \| HAL3-like protein A \| chr3:6167640-6168948 REVERSE LENGTH=1192 |
| AT4G10920.2 | Translation | \| Symbols: KELP \| transcriptional coactivator p15 (PC4) family protein (KELP) \| chr4:6696712-6699199 REVERSE LENGTH=1208 |
| AT1G51530.2 | Cleavage | \| Symbols: \| RNA-binding (RRM/RBD/RNP motifs) family protein \| chr1:19109854-19112157 FORWARD LENGTH=1217 |
| AT3G42930.1 | Cleavage | \| Symbols: \| transposable element gene \| chr3:14998007-14999235 REVERSE LENGTH=1229 |
| AT5G30545.1 | Cleavage | \| Symbols: \| transposable element gene \| chr5:11641770-11643020 REVERSE LENGTH=1251 |
| AT1G28710.1 | Cleavage | \| Symbols: \| Nucleotide-diphospho-sugar transferase family protein \| chr1:10086637-10088071 REVERSE LENGTH=1282 |
| AT4G07420.1 | Cleavage | \| Symbols: \| transposable element gene \| chr4:4214468-4215758 FORWARD LENGTH=1291 |
| AT4G19150.2 | Cleavage | \| Symbols: \| Ankyrin repeat family protein \| chr4:10471399-10472726 REVERSE LENGTH=1328 |
| AT3G57040.1 | Cleavage | \| Symbols: ARR9, ATRR4 \| response regulator 9 \| chr3:21109614-21111432 FORWARD LENGTH=1354 |
| AT3G21100.1 | Cleavage | \| Symbols: \| RNA-binding (RRM/RBD/RNP motifs) family protein \| chr3:7400239-7402436 FORWARD LENGTH=1464 |
| AT1G28710.2 | Cleavage | \| Symbols: \| Nucleotide-diphospho-sugar transferase family protein \| chr1:10086637-10089535 REVERSE LENGTH=1466 |
| AT3G09540.1 | Translation | \| Symbols: \| Pectin lyase-like superfamily protein \| chr3:2928811-2931228 REVERSE LENGTH=1496 |
| AT5G65510.1 | Cleavage | \| Symbols: AIL7 \| AINTEGUMENTA-like 7 \| chr5:26185829-26189413 FORWARD LENGTH=1497 |
| AT5G60770.1 | Cleavage | \| Symbols: ATNRT2.4, NRT2.4 \| nitrate transporter 2.4 \| chr5:24444396-24447026 FORWARD LENGTH=1584 |
| AT1G78510.1 | Cleavage | \| Symbols: SPS1 \| solanesyl diphosphate synthase 1 \| chr1:29535285-29537417 REVERSE LENGTH=1600 |
| AT1G78510.2 | Cleavage | \| Symbols: SPS1 \| solanesyl diphosphate synthase 1 \| chr1:29535285-29537417 REVERSE LENGTH=1604 |
| AT1G75710.1 | Cleavage | \| Symbols: \| C2H2-like zinc finger protein \| chr1:28428706-28431249 FORWARD LENGTH=1610 |
| AT1G04500.1 | Cleavage | \| Symbols: \| CCT motif family protein \| chr1:1221433-1224366 REVERSE LENGTH=1616 |
| AT2G29710.1 | Cleavage | \| Symbols: \| UDP-Glycosyltransferase superfamily protein \| chr2:12698673-12700343 FORWARD LENGTH=1671 |
| AT1G28710.3 | Cleavage | \| Symbols: \| Nucleotide-diphospho-sugar transferase family protein \| chr1:10086600-10089772 REVERSE LENGTH=1718 |
| AT3G55320.1 | Cleavage | \| Symbols: PGP20 \| P-glycoprotein 20 \| chr3:20506992-20513393 REVERSE LENGTH=4626 |
| AT2G39480.1 | Cleavage | \| Symbols: PGP6 \| P-glycoprotein 6 \| chr2:16477792-16485039 REVERSE LENGTH=4893 |
| AT2G07784.1 | Cleavage | \| Symbols: \| transposable element gene \| chr2:3317798-3318634 REVERSE LENGTH=837 |
| AT3G22380.2 | Cleavage | \| Symbols: TIC \| time for coffee \| chr3:7912905-7919510 FORWARD LENGTH=5487 |
| AT3G22380.1 | Cleavage | \| Symbols: TIC \| time for coffee \| chr3:7912905-7919484 FORWARD LENGTH=5534 |
| AT5G23950.1 | Cleavage | \| Symbols: \| Calcium-dependent lipid-binding (CaLB domain) family protein \| chr5:8082744-8083604 FORWARD LENGTH=861 |
| AT5G61980.1 | Cleavage | \| Symbols: AGD1 \| ARF-GAP domain 1 \| chr5:24894472-24899973 FORWARD LENGTH=3282 |
| AT1G10840.2 | Cleavage | \| Symbols: TIF3H1 \| translation initiation factor 3 subunit H1 \| chr1:3607631-3610386 REVERSE LENGTH=1471 |
| AT1G10840.1 | Cleavage | \| Symbols: TIF3H1 \| translation initiation factor 3 subunit H1 \| chr1:3607631-3610520 REVERSE LENGTH=1489 |
| AT5G47550.1 | Cleavage | \| Symbols: \| Cystatin/monellin superfamily protein \| chr5:19286450-19286995 REVERSE LENGTH=546 |
| AT2G42500.2 | Cleavage | \| Symbols: PP2A-3 \| protein phosphatase 2A-3 \| chr2:17697670-17701417 REVERSE LENGTH=1421 |
| AT2G42500.1 | Cleavage | \| Symbols: PP2A-3 \| protein phosphatase 2A-3 \| chr2:17697667-17701417 REVERSE LENGTH=1565 |
| AT3G02690.1 | Cleavage | \| Symbols: \| nodulin MtN21 /EamA-like transporter family protein \| chr3:579586-581611 FORWARD LENGTH=1458 |
| AT1G13050.2 | Cleavage | \| Symbols: \| unknown protein; FUNCTIONS IN: molecular_function unknown; INVOLVED IN: biological_process unknown; LOCATED IN: endomembrane system; EXPRESSED IN: 14 plant structures; EXPRESSED DURING: 9 growth stages; BEST Arabidopsis thaliana protein match is: unknown protein (TAIR:AT3G26350.1); Has 260 Blast hits to 259 proteins in 20 species: Archae - 0; Bacteria - 0; Metazoa - 0; Fungi - 0; Plants - 260; Viruses - 0; Other Eukaryotes - 0 (source: NCBI BLink). \| chr1:4450238-4451742 FORWARD LENGTH=1149 |
| AT1G13050.1 | Cleavage | \| Symbols: \| unknown protein; BEST Arabidopsis thaliana protein match is: unknown protein (TAIR:AT3G26350.1); Has 538 Blast hits to 510 proteins in 88 species: Archae - 0; Bacteria - 23; Metazoa - 81; Fungi - 36; Plants - 361; Viruses - 8; Other Eukaryotes - 29 (source: NCBI BLink). \| chr1:4450382-4451742 FORWARD LENGTH=1361 |
| AT3G48300.1 | Cleavage | \| Symbols: CYP71A23 \| cytochrome P450, family 71, subfamily A, polypeptide 23 \| chr3:17885524-17887118 FORWARD LENGTH=1452 |
| AT1G64960.1 | Cleavage | \| Symbols: \| ARM repeat superfamily protein \| chr1:24129918-24134204 FORWARD LENGTH=3767 |
| AT2G10940.1 | Translation | \| Symbols: \| Bifunctional inhibitor/lipid-transfer protein/seed storage 2S albumin superfamily protein \| chr2:4310412-4312103 REVERSE LENGTH=1338 |
| AT2G10940.2 | Translation | \| Symbols: \| Bifunctional inhibitor/lipid-transfer protein/seed storage 2S albumin superfamily protein \| chr2:4310412-4312103 REVERSE LENGTH=1692 |
| AT2G21380.1 | Cleavage | \| Symbols: \| Kinesin motor family protein \| chr2:9141631-9149309 FORWARD LENGTH=3614 |
| AT3G57920.1 | Cleavage | \| Symbols: SPL15 \| squamosa promoter binding protein-like 15 \| chr3:21444321-21446035 REVERSE LENGTH=1377 |
| AT3G57920.1 | Cleavage | \| Symbols: SPL15 \| squamosa promoter binding protein-like 15 \| chr3:21444321-21446035 REVERSE LENGTH=1377 |
| AT5G62090.2 | Translation | \| Symbols: SLK2 \| SEUSS-like 2 \| chr5:24934963-24939109 REVERSE LENGTH=2828 |
| AT5G62090.1 | Translation | \| Symbols: SLK2 \| SEUSS-like 2 \| chr5:24934963-24939060 REVERSE LENGTH=3229 |
| AT5G28692.1 | Cleavage | \| Symbols: \| transposable element gene \| chr5:10728378-10733393 REVERSE LENGTH=5016 |
| AT5G60770.1 | Cleavage | \| Symbols: ATNRT2.4, NRT2.4 \| nitrate transporter 2.4 \| chr5:24444396-24447026 FORWARD LENGTH=1584 |

| **cca-miR11300** | | |
| --- | --- | --- |
| Target_Acc. | Inhibition | Target_Desc. |
| AT4G36090.3 | Cleavage | \| Symbols: \| oxidoreductase, 2OG-Fe(II) oxygenase family protein \| chr4:17078311-17080749 REVERSE LENGTH=1707 |
| AT4G36090.1 | Cleavage | \| Symbols: \| oxidoreductase, 2OG-Fe(II) oxygenase family protein \| chr4:17078311-17080776 REVERSE LENGTH=1830 |
| AT4G36090.2 | Cleavage | \| Symbols: \| oxidoreductase, 2OG-Fe(II) oxygenase family protein \| chr4:17078311-17080786 REVERSE LENGTH=2025 |
| AT3G62190.1 | Cleavage | \| Symbols: \| Chaperone DnaJ-domain superfamily protein \| chr3:23021121-23023332 FORWARD LENGTH=1160 |
| AT2G36830.1 | Translation | \| Symbols: GAMMA-TIP, TIP1;1, GAMMA-TIP1 \| gamma tonoplast intrinsic protein \| chr2:15445425-15446574 FORWARD LENGTH=1059 |
| AT5G24810.1 | Cleavage | \| Symbols: \| ABC1 family protein \| chr5:8516635-8522767 REVERSE LENGTH=3448 |
| AT3G44870.1 | Cleavage | \| Symbols: \| S-adenosyl-L-methionine-dependent methyltransferases superfamily protein \| chr3:16382219-16383605 FORWARD LENGTH=1198 |
| AT5G52060.1 | Translation | \| Symbols: ATBAG1, BAG1 \| BCL-2-associated athanogene 1 \| chr5:21152214-21154244 REVERSE LENGTH=1767 |
| AT5G09680.2 | Cleavage | \| Symbols: RLF \| reduced lateral root formation \| chr5:2999220-3000601 REVERSE LENGTH=1194 |
| AT2G20140.1 | Cleavage | \| Symbols: \| AAA-type ATPase family protein \| chr2:8692700-8695084 FORWARD LENGTH=1615 |
| AT1G80210.1 | Cleavage | \| Symbols: BRCC36A, AtBRCC36A \| Mov34/MPN/PAD-1 family protein \| chr1:30162835-30165755 REVERSE LENGTH=1870 |
| AT1G80210.2 | Cleavage | \| Symbols: BRCC36A, AtBRCC36A \| Mov34/MPN/PAD-1 family protein \| chr1:30162835-30165755 REVERSE LENGTH=1954 |
| AT4G01970.1 | Cleavage | \| Symbols: AtSTS, STS \| stachyose synthase \| chr4:853922-857008 REVERSE LENGTH=2837 |
| AT5G25060.1 | Translation | \| Symbols: \| RNA recognition motif (RRM)-containing protein \| chr5:8634074-8640217 REVERSE LENGTH=3219 |
| AT5G11320.2 | Cleavage | \| Symbols: YUC4 \| Flavin-binding monooxygenase family protein \| chr5:3611990-3613361 REVERSE LENGTH=1141 |
| AT5G11320.1 | Cleavage | \| Symbols: YUC4 \| Flavin-binding monooxygenase family protein \| chr5:3611247-3613361 REVERSE LENGTH=1418 |
| AT5G11840.1 | Cleavage | \| Symbols: \| Protein of unknown function (DUF1230) \| chr5:3813449-3814805 REVERSE LENGTH=959 |
| AT4G31060.1 | Cleavage | \| Symbols: \| Integrase-type DNA-binding superfamily protein \| chr4:15116148-15117756 FORWARD LENGTH=1372 |
| AT3G47600.1 | Translation | \| Symbols: MYB94, ATMYBCP70, ATMYB94 \| myb domain protein 94 \| chr3:17539338-17541230 REVERSE LENGTH=1486 |
| AT5G64060.1 | Cleavage | \| Symbols: anac103, NAC103 \| NAC domain containing protein 103 \| chr5:25633536-25635781 REVERSE LENGTH=1658 |
| AT3G48235.1 | Cleavage | \| Symbols: \| transposable element gene \| chr3:17863754-17865925 FORWARD LENGTH=2172 |
| AT5G04670.1 | Cleavage | \| Symbols: \| Enhancer of polycomb-like transcription factor protein \| chr5:1337839-1340884 REVERSE LENGTH=2460 |
| AT3G42255.1 | Cleavage | \| Symbols: \| transposable element gene \| chr3:14416062-14419442 FORWARD LENGTH=3381 |
| AT3G30211.1 | Cleavage | \| Symbols: \| transposable element gene \| chr3:11841552-11845610 REVERSE LENGTH=4059 |
| AT4G02600.2 | Cleavage | \| Symbols: MLO1, ATMLO1 \| Seven transmembrane MLO family protein \| chr4:1143962-1147422 FORWARD LENGTH=1942 |
| AT4G02600.1 | Cleavage | \| Symbols: MLO1, ATMLO1 \| Seven transmembrane MLO family protein \| chr4:1143999-1147409 FORWARD LENGTH=1976 |
| AT4G14970.1 | Cleavage | \| Symbols: \| unknown protein; FUNCTIONS IN: molecular_function unknown; INVOLVED IN: biological_process unknown; LOCATED IN: chloroplast; EXPRESSED IN: 14 plant structures; EXPRESSED DURING: 6 growth stages; Has 257 Blast hits to 164 proteins in 70 species: Archae - 0; Bacteria - 4; Metazoa - 189; Fungi - 0; Plants - 38; Viruses - 0; Other Eukaryotes - 26 (source: NCBI BLink). \| chr4:8553643-8561664 FORWARD LENGTH=4681 |
| AT3G21520.1 | Cleavage | \| Symbols: DMP1, AtDMP1 \| DUF679 domain membrane protein 1 \| chr3:7581959-7582793 FORWARD LENGTH=835 |
| AT2G35320.1 | Cleavage | \| Symbols: ATEYA, EYA \| EYES ABSENT homolog \| chr2:14866832-14868860 REVERSE LENGTH=1277 |
| AT3G29830.1 | Cleavage | \| Symbols: \| F-box/RNI-like superfamily protein \| chr3:11737049-11738847 FORWARD LENGTH=1395 |
| AT3G27473.1 | Cleavage | \| Symbols: \| Cysteine/Histidine-rich C1 domain family protein \| chr3:10169242-10171230 REVERSE LENGTH=1989 |
| AT4G14850.1 | Translation | \| Symbols: LOI1, MEF11 \| Pentatricopeptide repeat (PPR) superfamily protein \| chr4:8513859-8516360 FORWARD LENGTH=2228 |
| AT3G18770.1 | Cleavage | \| Symbols: \| Autophagy-related protein 13 \| chr3:6459884-6463059 REVERSE LENGTH=2343 |
| AT5G06460.1 | Cleavage | \| Symbols: ATUBA2, UBA 2 \| ubiquitin activating enzyme 2 \| chr5:1970029-1974527 FORWARD LENGTH=3438 |
| AT3G01460.1 | Cleavage | \| Symbols: MBD9, ATMBD9 \| methyl-CPG-binding domain 9 \| chr3:173316-182454 FORWARD LENGTH=6947 |
| AT2G07751.1 | Cleavage | \| Symbols: \| NADH:ubiquinone/plastoquinone oxidoreductase, chain 3 protein \| chr2:3270008-3270364 FORWARD LENGTH=357 |
| ATMG00990.1 | Cleavage | \| Symbols: NAD3 \| NADH dehydrogenase 3 \| chrM:260647-261006 REVERSE LENGTH=360 |
| AT1G03860.2 | Cleavage | \| Symbols: ATPHB2, PHB2 \| prohibitin 2 \| chr1:979308-981868 REVERSE LENGTH=1355 |
| AT5G43720.1 | Cleavage | \| Symbols: \| Protein of unknown function (DUF2361) \| chr5:17557166-17559354 REVERSE LENGTH=1371 |
| AT5G53750.1 | Translation | \| Symbols: \| CBS domain-containing protein \| chr5:21817366-21818843 FORWARD LENGTH=1397 |
| AT1G27370.4 | Translation | \| Symbols: SPL10 \| squamosa promoter binding protein-like 10 \| chr1:9505202-9508322 REVERSE LENGTH=1673 |
| AT3G50780.1 | Cleavage | \| Symbols: \| BEST Arabidopsis thaliana protein match is: BTB/POZ domain-containing protein (TAIR:AT1G63850.1); Has 298 Blast hits to 298 proteins in 22 species: Archae - 0; Bacteria - 0; Metazoa - 10; Fungi - 0; Plants - 287; Viruses - 0; Other Eukaryotes - 1 (source: NCBI BLink). \| chr3:18875399-18877534 REVERSE LENGTH=2019 |
| AT3G48500.1 | Translation | \| Symbols: PDE312, PTAC10 \| Nucleic acid-binding, OB-fold-like protein \| chr3:17962204-17965613 FORWARD LENGTH=2217 |
| AT3G48500.2 | Translation | \| Symbols: PDE312, PTAC10 \| Nucleic acid-binding, OB-fold-like protein \| chr3:17962163-17965613 FORWARD LENGTH=2345 |
| AT3G05970.1 | Cleavage | \| Symbols: LACS6, ATLACS6 \| long-chain acyl-CoA synthetase 6 \| chr3:1786318-1791802 REVERSE LENGTH=2354 |
| AT3G42255.1 | Translation | \| Symbols: \| transposable element gene \| chr3:14416062-14419442 FORWARD LENGTH=3381 |
| AT3G27700.2 | Cleavage | \| Symbols: \| zinc finger (CCCH-type) family protein / RNA recognition motif (RRM)-containing protein \| chr3:10257117-10261974 REVERSE LENGTH=3526 |
| AT3G27700.1 | Cleavage | \| Symbols: \| zinc finger (CCCH-type) family protein / RNA recognition motif (RRM)-containing protein \| chr3:10257117-10261974 REVERSE LENGTH=3689 |
| AT4G34131.1 | Cleavage | \| Symbols: UGT73B3 \| UDP-glucosyl transferase 73B3 \| chr4:16343057-16344818 REVERSE LENGTH=1762 |
| AT4G16500.1 | Cleavage | \| Symbols: \| Cystatin/monellin superfamily protein \| chr4:9301388-9302022 REVERSE LENGTH=635 |
| AT1G52880.1 | Cleavage | \| Symbols: NAM, ANAC018, ATNAM, NARS2 \| NAC (No Apical Meristem) domain transcriptional regulator superfamily protein \| chr1:19688957-19690542 REVERSE LENGTH=1403 |
| AT1G69828.1 | Cleavage | \| Symbols: \| Defensin-like (DEFL) family protein \| chr1:26287522-26287948 FORWARD LENGTH=225 |
| AT5G45010.1 | Cleavage | \| Symbols: ATDSS1(V), DSS1(V) \| DSS1 homolog on chromosome V \| chr5:18167244-18168374 REVERSE LENGTH=582 |
| AT1G09640.2 | Cleavage | \| Symbols: \| Translation elongation factor EF1B, gamma chain \| chr1:3119967-3122324 FORWARD LENGTH=1057 |
| AT5G42030.1 | Cleavage | \| Symbols: ABIL4 \| ABL interactor-like protein 4 \| chr5:16811384-16813183 REVERSE LENGTH=1070 |
| AT3G62620.2 | Cleavage | \| Symbols: \| sucrose-phosphatase-related \| chr3:23159783-23161607 FORWARD LENGTH=1260 |
| AT2G02470.1 | Cleavage | \| Symbols: AL6 \| alfin-like 6 \| chr2:652580-654915 FORWARD LENGTH=1322 |
| AT3G17060.1 | Cleavage | \| Symbols: \| Pectin lyase-like superfamily protein \| chr3:5816672-5818498 REVERSE LENGTH=1356 |
| AT2G02960.3 | Cleavage | \| Symbols: \| RING/FYVE/PHD zinc finger superfamily protein \| chr2:862087-864364 REVERSE LENGTH=1459 |
| AT2G02960.5 | Cleavage | \| Symbols: \| RING/FYVE/PHD zinc finger superfamily protein \| chr2:862087-864359 REVERSE LENGTH=1560 |
| AT3G50080.1 | Cleavage | \| Symbols: VFB2 \| VIER F-box proteine 2 \| chr3:18572736-18574514 FORWARD LENGTH=1779 |
| AT1G09640.1 | Cleavage | \| Symbols: \| Translation elongation factor EF1B, gamma chain \| chr1:3119878-3122526 FORWARD LENGTH=1795 |
| AT5G50150.1 | Cleavage | \| Symbols: \| Protein of Unknown Function (DUF239) \| chr5:20402257-20406731 REVERSE LENGTH=1806 |
| AT2G32990.1 | Cleavage | \| Symbols: AtGH9B8, GH9B8 \| glycosyl hydrolase 9B8 \| chr2:14003250-14006017 FORWARD LENGTH=1862 |
| AT2G42290.1 | Cleavage | \| Symbols: \| Leucine-rich repeat protein kinase family protein \| chr2:17616841-17619593 REVERSE LENGTH=2213 |
| AT2G44170.1 | Cleavage | \| Symbols: NMT2, ATNMT2 \| pseudogene, myristoyl-CoA:protein N-myristoyltransferase (NMT), putative, similar to N-myristoyltransferase 1 (NMT1) (Arabidopsis thaliana) GI:7339834; contains Pfam profiles PF01233: Myristoyl-CoA:protein N-myristoyltransferase N-terminal domain, PF02799: Myristoyl-CoA:protein N-myristoyltransferase C-terminal domain; gene contains a frameshift. This could be a pseudogene or a sequencing error.; blastp match of 62% identity and 5.1e-121 P-value to GP\|20804654\|dbj\|BAB92343.1\|\|AP003273 putative N-myristoyl transferase {Oryza sativa (japonica cultivar-group)} \| chr2:18265431-18267426 REVERSE LENGTH=1996 |
| AT2G43250.1 | Translation | \| Symbols: \| unknown protein; Has 32 Blast hits to 32 proteins in 11 species: Archae - 0; Bacteria - 0; Metazoa - 0; Fungi - 0; Plants - 32; Viruses - 0; Other Eukaryotes - 0 (source: NCBI BLink). \| chr2:17977228-17979608 FORWARD LENGTH=2381 |
| AT2G04270.5 | Cleavage | \| Symbols: RNEE/G \| RNAse E/G-like \| chr2:1474907-1480210 FORWARD LENGTH=3256 |
| AT1G14790.1 | Cleavage | \| Symbols: RDR1, ATRDRP1 \| RNA-dependent RNA polymerase 1 \| chr1:5093961-5098032 REVERSE LENGTH=3807 |
| AT3G13290.1 | Cleavage | \| Symbols: VCR \| varicose-related \| chr3:4297381-4303508 FORWARD LENGTH=4566 |
| AT3G18200.2 | Cleavage | \| Symbols: \| nodulin MtN21 /EamA-like transporter family protein \| chr3:6234306-6236091 REVERSE LENGTH=1317 |
| AT3G18200.1 | Cleavage | \| Symbols: \| nodulin MtN21 /EamA-like transporter family protein \| chr3:6234306-6236091 REVERSE LENGTH=1318 |
| AT3G27700.2 | Cleavage | \| Symbols: \| zinc finger (CCCH-type) family protein / RNA recognition motif (RRM)-containing protein \| chr3:10257117-10261974 REVERSE LENGTH=3526 |
| AT3G27700.1 | Cleavage | \| Symbols: \| zinc finger (CCCH-type) family protein / RNA recognition motif (RRM)-containing protein \| chr3:10257117-10261974 REVERSE LENGTH=3689 |
| AT3G22650.1 | Cleavage | \| Symbols: ATSFL61, SFL61, CEG \| F-box and associated interaction domains-containing protein \| chr3:8014809-8015960 FORWARD LENGTH=1119 |
| AT2G16040.1 | Cleavage | \| Symbols: \| hAT dimerisation domain-containing protein / transposase-related \| chr2:6976017-6977594 FORWARD LENGTH=1149 |
| AT3G30685.1 | Cleavage | \| Symbols: \| transposable element gene \| chr3:12234878-12236714 REVERSE LENGTH=1837 |
| AT2G07706.1 | Cleavage | \| Symbols: \| unknown protein; BEST Arabidopsis thaliana protein match is: unknown protein (TAIR:ATMG00470.1); Has 35333 Blast hits to 34131 proteins in 2444 species: Archae - 798; Bacteria - 22429; Metazoa - 974; Fungi - 991; Plants - 531; Viruses - 0; Other Eukaryotes - 9610 (source: NCBI BLink). \| chr2:3381901-3386102 REVERSE LENGTH=2080 |
| AT1G59870.1 | Cleavage | \| Symbols: PEN3, PDR8, ATPDR8, ABCG36, ATABCG36 \| ABC-2 and Plant PDR ABC-type transporter family protein \| chr1:22034506-22040038 FORWARD LENGTH=4759 |
| AT3G60540.1 | Cleavage | \| Symbols: \| Preprotein translocase Sec, Sec61-beta subunit protein \| chr3:22374963-22375865 REVERSE LENGTH=481 |
| AT5G39100.1 | Cleavage | \| Symbols: GLP6 \| germin-like protein 6 \| chr5:15653090-15654020 REVERSE LENGTH=835 |
| AT3G60540.2 | Cleavage | \| Symbols: \| Preprotein translocase Sec, Sec61-beta subunit protein \| chr3:22374963-22375865 REVERSE LENGTH=903 |
| AT2G14840.1 | Cleavage | \| Symbols: \| pseudogene, similar to phosphoenolpyruvate carboxykinase, similar to GB:L31899; frameshift at base 16259 possibly caused by a base a pair deletion.; blastp match of 72% identity and 3.7e-81 P-value to GP\|13785471\|dbj\|BAB43909.1\|\|AB050473 phosphoenolpyruvate carboxykinase {Flaveria pringlei} \| chr2:6371902-6373082 FORWARD LENGTH=1181 |
| AT2G14840.1 | Cleavage | \| Symbols: \| pseudogene, similar to phosphoenolpyruvate carboxykinase, similar to GB:L31899; frameshift at base 16259 possibly caused by a base a pair deletion.; blastp match of 72% identity and 3.7e-81 P-value to GP\|13785471\|dbj\|BAB43909.1\|\|AB050473 phosphoenolpyruvate carboxykinase {Flaveria pringlei} \| chr2:6371902-6373082 FORWARD LENGTH=1181 |
| AT4G01960.1 | Translation | \| Symbols: \| unknown protein; BEST Arabidopsis thaliana protein match is: unknown protein (TAIR:AT1G02380.1); Has 67 Blast hits to 67 proteins in 11 species: Archae - 0; Bacteria - 0; Metazoa - 0; Fungi - 0; Plants - 67; Viruses - 0; Other Eukaryotes - 0 (source: NCBI BLink). \| chr4:851210-853165 REVERSE LENGTH=1191 |
| AT4G12980.1 | Cleavage | \| Symbols: \| Auxin-responsive family protein \| chr4:7589395-7591102 REVERSE LENGTH=1488 |
| AT3G20880.1 | Cleavage | \| Symbols: WIP4 \| WIP domain protein 4 \| chr3:7313538-7315883 REVERSE LENGTH=1551 |
| AT1G11860.3 | Cleavage | \| Symbols: \| Glycine cleavage T-protein family \| chr1:4001113-4003267 FORWARD LENGTH=1666 |
| AT1G52040.1 | Cleavage | \| Symbols: MBP1, ATMBP \| myrosinase-binding protein 1 \| chr1:19350375-19352782 REVERSE LENGTH=1681 |
| AT1G01660.1 | Translation | \| Symbols: \| RING/U-box superfamily protein \| chr1:240057-242608 REVERSE LENGTH=1707 |
| AT4G36700.1 | Translation | \| Symbols: \| RmlC-like cupins superfamily protein \| chr4:17298200-17300367 REVERSE LENGTH=1842 |
| AT5G46420.1 | Cleavage | \| Symbols: \| 16S rRNA processing protein RimM family \| chr5:18829955-18832953 FORWARD LENGTH=2066 |
| AT1G04430.2 | Translation | \| Symbols: \| S-adenosyl-L-methionine-dependent methyltransferases superfamily protein \| chr1:1198508-1201495 FORWARD LENGTH=2268 |
| AT1G64250.1 | Cleavage | \| Symbols: \| transposable element gene \| chr1:23840888-23843251 FORWARD LENGTH=2364 |
| AT1G04430.1 | Translation | \| Symbols: \| S-adenosyl-L-methionine-dependent methyltransferases superfamily protein \| chr1:1198118-1201527 FORWARD LENGTH=2379 |
| AT3G13065.1 | Cleavage | \| Symbols: SRF4 \| STRUBBELIG-receptor family 4 \| chr3:4187371-4191298 FORWARD LENGTH=2638 |
| AT1G15300.1 | Cleavage | \| Symbols: \| transposable element gene \| chr1:5265726-5268602 REVERSE LENGTH=2668 |
| AT3G48195.1 | Cleavage | \| Symbols: \| Phox (PX) domain-containing protein \| chr3:17828572-17832731 REVERSE LENGTH=3294 |
| AT2G42270.1 | Cleavage | \| Symbols: \| U5 small nuclear ribonucleoprotein helicase \| chr2:17604330-17611333 FORWARD LENGTH=6906 |
| AT5G41450.1 | Cleavage | \| Symbols: \| RING/U-box superfamily protein \| chr5:16588600-16589094 REVERSE LENGTH=495 |
| AT5G25240.1 | Translation | \| Symbols: \| unknown protein; FUNCTIONS IN: molecular_function unknown; INVOLVED IN: biological_process unknown; LOCATED IN: cellular_component unknown; EXPRESSED IN: 23 plant structures; EXPRESSED DURING: 13 growth stages; Has 1807 Blast hits to 1807 proteins in 277 species: Archae - 0; Bacteria - 0; Metazoa - 736; Fungi - 347; Plants - 385; Viruses - 0; Other Eukaryotes - 339 (source: NCBI BLink). \| chr5:8746605-8747197 REVERSE LENGTH=593 |
| AT1G24260.1 | Translation | \| Symbols: SEP3, AGL9 \| K-box region and MADS-box transcription factor family protein \| chr1:8593642-8595909 REVERSE LENGTH=948 |
| AT1G24260.3 | Translation | \| Symbols: SEP3 \| K-box region and MADS-box transcription factor family protein \| chr1:8593637-8596105 REVERSE LENGTH=1110 |
| AT1G24260.2 | Translation | \| Symbols: SEP3, AGL9 \| K-box region and MADS-box transcription factor family protein \| chr1:8593642-8596098 REVERSE LENGTH=1140 |
| AT4G25310.1 | Translation | \| Symbols: \| 2-oxoglutarate (2OG) and Fe(II)-dependent oxygenase superfamily protein \| chr4:12949664-12951234 FORWARD LENGTH=1247 |
| AT4G11240.1 | Cleavage | \| Symbols: TOPP7 \| Calcineurin-like metallo-phosphoesterase superfamily protein \| chr4:6847112-6849300 FORWARD LENGTH=1458 |
| AT3G27220.1 | Cleavage | \| Symbols: \| Galactose oxidase/kelch repeat superfamily protein \| chr3:10051612-10053678 REVERSE LENGTH=1495 |
| AT1G44575.2 | Translation | \| Symbols: NPQ4, PSBS \| Chlorophyll A-B binding family protein \| chr1:16871696-16873378 FORWARD LENGTH=1520 |
| AT4G30993.2 | Cleavage | \| Symbols: \| Calcineurin-like metallo-phosphoesterase superfamily protein \| chr4:15098067-15100558 FORWARD LENGTH=1615 |
| AT5G17630.1 | Cleavage | \| Symbols: \| Nucleotide/sugar transporter family protein \| chr5:5809334-5810962 FORWARD LENGTH=1629 |
| AT4G06504.1 | Translation | \| Symbols: \| transposable element gene \| chr4:3171711-3173348 FORWARD LENGTH=1638 |
| AT3G58900.3 | Cleavage | \| Symbols: \| F-box/RNI-like superfamily protein \| chr3:21772838-21774743 FORWARD LENGTH=1702 |
| AT4G30993.1 | Cleavage | \| Symbols: \| Calcineurin-like metallo-phosphoesterase superfamily protein \| chr4:15098067-15100558 FORWARD LENGTH=1708 |
| AT3G58900.4 | Cleavage | \| Symbols: \| F-box/RNI-like superfamily protein \| chr3:21772838-21774743 FORWARD LENGTH=1713 |
| AT1G67110.1 | Cleavage | \| Symbols: CYP735A2 \| cytochrome P450, family 735, subfamily A, polypeptide 2 \| chr1:25061731-25065446 REVERSE LENGTH=1735 |
| AT3G58900.1 | Cleavage | \| Symbols: \| F-box/RNI-like superfamily protein \| chr3:21772857-21774743 FORWARD LENGTH=1783 |
| AT2G42720.1 | Cleavage | \| Symbols: \| FBD, F-box, Skp2-like and Leucine Rich Repeat domains containing protein \| chr2:17785409-17787421 FORWARD LENGTH=1786 |
| AT5G37590.1 | Translation | \| Symbols: \| Tetratricopeptide repeat (TPR)-like superfamily protein \| chr5:14927313-14932787 REVERSE LENGTH=1824 |
| AT5G28923.1 | Translation | \| Symbols: \| transposable element gene \| chr5:10946562-10948829 REVERSE LENGTH=2268 |
| AT4G13710.1 | Cleavage | \| Symbols: \| Pectin lyase-like superfamily protein \| chr4:7962432-7966461 FORWARD LENGTH=1980 |
| AT4G13710.2 | Cleavage | \| Symbols: \| Pectin lyase-like superfamily protein \| chr4:7962430-7966518 FORWARD LENGTH=1982 |
| AT1G23720.1 | Translation | \| Symbols: \| Proline-rich extensin-like family protein \| chr1:8388655-8391540 FORWARD LENGTH=2886 |
| AT4G13550.1 | Translation | \| Symbols: \| triglyceride lipases;triglyceride lipases \| chr4:7871057-7876963 REVERSE LENGTH=2971 |
| AT4G03970.1 | Translation | \| Symbols: \| transposable element gene \| chr4:1892890-1897940 FORWARD LENGTH=3132 |
| AT1G10290.1 | Translation | \| Symbols: ADL6, DRP2A \| dynamin-like protein 6 \| chr1:3370558-3377463 FORWARD LENGTH=3304 |
| AT4G24480.1 | Translation | \| Symbols: \| Protein kinase superfamily protein \| chr4:12649987-12654984 FORWARD LENGTH=3523 |
| AT2G37280.1 | Translation | \| Symbols: PDR5, ATPDR5 \| pleiotropic drug resistance 5 \| chr2:15650400-15656417 FORWARD LENGTH=4242 |
| AT1G21630.1 | Translation | \| Symbols: \| Calcium-binding EF hand family protein \| chr1:7581152-7588151 FORWARD LENGTH=4317 |
| AT1G21630.2 | Translation | \| Symbols: \| Calcium-binding EF hand family protein \| chr1:7581084-7588064 FORWARD LENGTH=4385 |
| AT1G47860.1 | Cleavage | \| Symbols: \| transposable element gene \| chr1:17620677-17630270 REVERSE LENGTH=9594 |
| AT3G57040.1 | Cleavage | \| Symbols: ARR9, ATRR4 \| response regulator 9 \| chr3:21109614-21111432 FORWARD LENGTH=1354 |
| AT2G25520.1 | Cleavage | \| Symbols: \| Drug/metabolite transporter superfamily protein \| chr2:10860828-10862357 FORWARD LENGTH=1530 |
| AT3G46080.1 | Cleavage | \| Symbols: \| C2H2-type zinc finger family protein \| chr3:16922753-16923247 REVERSE LENGTH=495 |
| AT1G17240.1 | Cleavage | \| Symbols: AtRLP2, RLP2 \| receptor like protein 2 \| chr1:5896416-5898717 REVERSE LENGTH=2302 |
| AT1G72300.1 | Cleavage | \| Symbols: \| Leucine-rich receptor-like protein kinase family protein \| chr1:27217477-27221123 REVERSE LENGTH=3647 |
| AT4G33420.1 | Translation | \| Symbols: \| Peroxidase superfamily protein \| chr4:16084828-16086296 FORWARD LENGTH=1197 |
| AT3G14550.1 | Cleavage | \| Symbols: GGPS3 \| geranylgeranyl pyrophosphate synthase 3 \| chr3:4884566-4885982 REVERSE LENGTH=1308 |
| AT1G72210.1 | Cleavage | \| Symbols: \| basic helix-loop-helix (bHLH) DNA-binding superfamily protein \| chr1:27179790-27182430 FORWARD LENGTH=1401 |
| AT2G26460.1 | Cleavage | \| Symbols: SMU2 \| RED family protein \| chr2:11254932-11258973 REVERSE LENGTH=1935 |
| AT1G76550.1 | Cleavage | \| Symbols: \| Phosphofructokinase family protein \| chr1:28722702-28727006 REVERSE LENGTH=2129 |
| AT5G24206.1 | Cleavage | \| Symbols: \| other RNA \| chr5:8213626-8214689 REVERSE LENGTH=467 |
| AT1G59720.1 | Cleavage | \| Symbols: CRR28 \| Tetratricopeptide repeat (TPR)-like superfamily protein \| chr1:21939752-21941784 REVERSE LENGTH=2033 |
| AT3G33565.1 | Cleavage | \| Symbols: \| transposable element gene \| chr3:14060883-14064624 FORWARD LENGTH=3742 |
| AT3G33565.1 | Cleavage | \| Symbols: \| transposable element gene \| chr3:14060883-14064624 FORWARD LENGTH=3742 |
| ATCG01110.1 | Cleavage | \| Symbols: NDHH \| NAD(P)H dehydrogenase subunit H \| chrC:122011-123192 REVERSE LENGTH=1182 |
| AT5G45890.1 | Translation | \| Symbols: SAG12 \| senescence-associated gene 12 \| chr5:18613259-18614930 FORWARD LENGTH=1253 |
| AT5G48340.2 | Cleavage | \| Symbols: \| unknown protein; FUNCTIONS IN: molecular_function unknown; INVOLVED IN: biological_process unknown; LOCATED IN: cellular_component unknown; EXPRESSED IN: 24 plant structures; EXPRESSED DURING: 14 growth stages. \| chr5:19590436-19592949 FORWARD LENGTH=2140 |
| AT5G48340.1 | Cleavage | \| Symbols: \| unknown protein; Has 1807 Blast hits to 1807 proteins in 277 species: Archae - 0; Bacteria - 0; Metazoa - 736; Fungi - 347; Plants - 385; Viruses - 0; Other Eukaryotes - 339 (source: NCBI BLink). \| chr5:19590473-19593000 FORWARD LENGTH=2156 |
| AT2G06880.1 | Translation | \| Symbols: \| transposable element gene \| chr2:2782435-2785689 FORWARD LENGTH=3255 |
| AT4G14695.2 | Cleavage | \| Symbols: \| Uncharacterised protein family (UPF0041) \| chr4:8419799-8421030 FORWARD LENGTH=495 |
| AT4G14695.1 | Cleavage | \| Symbols: \| Uncharacterised protein family (UPF0041) \| chr4:8419799-8420998 FORWARD LENGTH=517 |
| AT2G19960.1 | Cleavage | \| Symbols: \| hAT family dimerisation domain \| chr2:8622215-8622736 FORWARD LENGTH=522 |
| AT1G50320.1 | Cleavage | \| Symbols: ATHX, ATX, THX \| thioredoxin X \| chr1:18638384-18639536 REVERSE LENGTH=843 |
| AT3G12410.1 | Cleavage | \| Symbols: \| Polynucleotidyl transferase, ribonuclease H-like superfamily protein \| chr3:3946036-3946960 REVERSE LENGTH=925 |
| AT1G67100.1 | Cleavage | \| Symbols: LBD40 \| LOB domain-containing protein 40 \| chr1:25053734-25054847 FORWARD LENGTH=957 |
| AT1G34520.1 | Cleavage | \| Symbols: \| MBOAT (membrane bound O-acyl transferase) family protein \| chr1:12623477-12624487 FORWARD LENGTH=1011 |
| AT1G34490.1 | Cleavage | \| Symbols: \| MBOAT (membrane bound O-acyl transferase) family protein \| chr1:12609482-12610495 FORWARD LENGTH=1014 |
| AT5G44020.1 | Cleavage | \| Symbols: \| HAD superfamily, subfamily IIIB acid phosphatase \| chr5:17712409-17714336 FORWARD LENGTH=1133 |
| AT4G27450.1 | Cleavage | \| Symbols: \| Aluminium induced protein with YGL and LRDR motifs \| chr4:13727484-13728886 REVERSE LENGTH=1137 |
| AT3G15840.5 | Translation | \| Symbols: PIFI \| post-illumination chlorophyll fluorescence increase \| chr3:5356601-5358593 REVERSE LENGTH=1179 |
| AT3G02420.1 | Cleavage | \| Symbols: \| unknown protein; FUNCTIONS IN: molecular_function unknown; INVOLVED IN: biological_process unknown; LOCATED IN: membrane; EXPRESSED IN: 25 plant structures; EXPRESSED DURING: 15 growth stages; CONTAINS InterPro DOMAIN/s: Uncharacterised protein family UPF0121 (InterPro:IPR005344); Has 72 Blast hits to 71 proteins in 25 species: Archae - 0; Bacteria - 0; Metazoa - 2; Fungi - 2; Plants - 60; Viruses - 0; Other Eukaryotes - 8 (source: NCBI BLink). \| chr3:495885-498841 REVERSE LENGTH=1410 |
| AT4G24220.2 | Cleavage | \| Symbols: VEP1, AWI31 \| NAD(P)-binding Rossmann-fold superfamily protein \| chr4:12564945-12566755 FORWARD LENGTH=1719 |
| AT4G24220.1 | Cleavage | \| Symbols: VEP1, AWI31 \| NAD(P)-binding Rossmann-fold superfamily protein \| chr4:12564945-12566755 FORWARD LENGTH=1722 |
| AT4G27020.1 | Translation | \| Symbols: \| unknown protein; FUNCTIONS IN: molecular_function unknown; LOCATED IN: vacuole; BEST Arabidopsis thaliana protein match is: unknown protein (TAIR:AT5G54870.1); Has 1807 Blast hits to 1807 proteins in 277 species: Archae - 0; Bacteria - 0; Metazoa - 736; Fungi - 347; Plants - 385; Viruses - 0; Other Eukaryotes - 339 (source: NCBI BLink). \| chr4:13568432-13571474 REVERSE LENGTH=1837 |
| AT1G61960.1 | Cleavage | \| Symbols: \| Mitochondrial transcription termination factor family protein \| chr1:22902150-22903837 FORWARD LENGTH=1688 |
| AT2G16570.1 | Translation | \| Symbols: ATASE, ATASE1, ASE1 \| GLN phosphoribosyl pyrophosphate amidotransferase 1 \| chr2:7180145-7182193 REVERSE LENGTH=2049 |
| AT5G17460.1 | Cleavage | \| Symbols: \| unknown protein; FUNCTIONS IN: molecular_function unknown; INVOLVED IN: response to salt stress; LOCATED IN: mitochondrion; Has 30201 Blast hits to 17322 proteins in 780 species: Archae - 12; Bacteria - 1396; Metazoa - 17338; Fungi - 3422; Plants - 5037; Viruses - 0; Other Eukaryotes - 2996 (source: NCBI BLink). \| chr5:5757098-5759795 REVERSE LENGTH=2251 |
| AT2G01750.1 | Cleavage | \| Symbols: ATMAP70-3, MAP70-3 \| microtubule-associated proteins 70-3 \| chr2:328089-332113 FORWARD LENGTH=2336 |
| AT2G01750.2 | Cleavage | \| Symbols: MAP70-3 \| microtubule-associated proteins 70-3 \| chr2:328089-332113 FORWARD LENGTH=2339 |
| AT4G33620.1 | Cleavage | \| Symbols: \| Cysteine proteinases superfamily protein \| chr4:16147692-16152853 FORWARD LENGTH=2352 |
| AT1G32940.1 | Cleavage | \| Symbols: ATSBT3.5, SBT3.5 \| Subtilase family protein \| chr1:11937596-11940978 FORWARD LENGTH=2485 |
| AT1G73920.1 | Cleavage | \| Symbols: \| alpha/beta-Hydrolases superfamily protein \| chr1:27790875-27794914 FORWARD LENGTH=2650 |
| AT1G73920.2 | Cleavage | \| Symbols: \| alpha/beta-Hydrolases superfamily protein \| chr1:27790876-27794914 FORWARD LENGTH=2654 |
| AT4G32300.1 | Translation | \| Symbols: SD2-5 \| S-domain-2 5 \| chr4:15599475-15602595 FORWARD LENGTH=3121 |
| AT4G38140.1 | Cleavage | \| Symbols: \| RING/U-box superfamily protein \| chr4:17899868-17900305 REVERSE LENGTH=438 |
| AT4G11211.1 | Cleavage | \| Symbols: \| unknown protein; Has 35333 Blast hits to 34131 proteins in 2444 species: Archae - 798; Bacteria - 22429; Metazoa - 974; Fungi - 991; Plants - 531; Viruses - 0; Other Eukaryotes - 9610 (source: NCBI BLink). \| chr4:6836106-6836588 FORWARD LENGTH=483 |
| AT5G25060.1 | Cleavage | \| Symbols: \| RNA recognition motif (RRM)-containing protein \| chr5:8634074-8640217 REVERSE LENGTH=3219 |
| AT1G69295.1 | Cleavage | \| Symbols: PDCB4 \| plasmodesmata callose-binding protein 4 \| chr1:26050197-26052567 REVERSE LENGTH=1688 |
| AT3G60540.1 | Translation | \| Symbols: \| Preprotein translocase Sec, Sec61-beta subunit protein \| chr3:22374963-22375865 REVERSE LENGTH=481 |
| AT3G60540.2 | Translation | \| Symbols: \| Preprotein translocase Sec, Sec61-beta subunit protein \| chr3:22374963-22375865 REVERSE LENGTH=903 |
| AT4G01960.1 | Cleavage | \| Symbols: \| unknown protein; BEST Arabidopsis thaliana protein match is: unknown protein (TAIR:AT1G02380.1); Has 67 Blast hits to 67 proteins in 11 species: Archae - 0; Bacteria - 0; Metazoa - 0; Fungi - 0; Plants - 67; Viruses - 0; Other Eukaryotes - 0 (source: NCBI BLink). \| chr4:851210-853165 REVERSE LENGTH=1191 |
| AT5G20640.1 | Cleavage | \| Symbols: \| Protein of unknown function (DUF567) \| chr5:6984240-6985365 FORWARD LENGTH=947 |
| AT5G60700.1 | Cleavage | \| Symbols: \| glycosyltransferase family protein 2 \| chr5:24402129-24404948 REVERSE LENGTH=2426 |
| AT2G28360.1 | Cleavage | \| Symbols: \| SIT4 phosphatase-associated family protein \| chr2:12124415-12130069 REVERSE LENGTH=2746 |
| AT1G14790.1 | Cleavage | \| Symbols: RDR1, ATRDRP1 \| RNA-dependent RNA polymerase 1 \| chr1:5093961-5098032 REVERSE LENGTH=3807 |
| AT5G15750.1 | Cleavage | \| Symbols: \| Alpha-L RNA-binding motif/Ribosomal protein S4 family protein \| chr5:5141103-5142866 FORWARD LENGTH=901 |
| AT3G18430.2 | Cleavage | \| Symbols: \| Calcium-binding EF-hand family protein \| chr3:6325915-6327692 FORWARD LENGTH=914 |
| AT3G14530.1 | Cleavage | \| Symbols: \| Terpenoid synthases superfamily protein \| chr3:4879112-4880448 REVERSE LENGTH=1264 |
| AT2G37630.1 | Cleavage | \| Symbols: ATPHAN, AS1, ATMYB91, MYB91 \| myb-like HTH transcriptional regulator family protein \| chr2:15781615-15783433 REVERSE LENGTH=1490 |
| AT3G07550.1 | Cleavage | \| Symbols: \| RNI-like superfamily protein \| chr3:2409477-2411294 FORWARD LENGTH=1708 |
| AT3G07550.1 | Cleavage | \| Symbols: \| RNI-like superfamily protein \| chr3:2409477-2411294 FORWARD LENGTH=1708 |
| AT5G49100.1 | Translation | \| Symbols: \| unknown protein; BEST Arabidopsis thaliana protein match is: unknown protein (TAIR:AT3G06868.1); Has 30201 Blast hits to 17322 proteins in 780 species: Archae - 12; Bacteria - 1396; Metazoa - 17338; Fungi - 3422; Plants - 5037; Viruses - 0; Other Eukaryotes - 2996 (source: NCBI BLink). \| chr5:19897370-19899114 REVERSE LENGTH=1745 |
| AT1G13040.1 | Cleavage | \| Symbols: \| Pentatricopeptide repeat (PPR-like) superfamily protein \| chr1:4447647-4449200 FORWARD LENGTH=1554 |
| AT3G07550.2 | Cleavage | \| Symbols: \| RNI-like superfamily protein \| chr3:2409727-2411294 FORWARD LENGTH=1568 |
| AT3G07550.2 | Cleavage | \| Symbols: \| RNI-like superfamily protein \| chr3:2409727-2411294 FORWARD LENGTH=1568 |
| AT5G43820.1 | Cleavage | \| Symbols: \| Pentatricopeptide repeat (PPR) superfamily protein \| chr5:17618948-17620588 FORWARD LENGTH=1641 |
| AT5G18500.1 | Cleavage | \| Symbols: \| Protein kinase superfamily protein \| chr5:6138489-6141630 FORWARD LENGTH=2007 |
| AT3G43240.1 | Cleavage | \| Symbols: \| ARID/BRIGHT DNA-binding domain-containing protein \| chr3:15209936-15214742 REVERSE LENGTH=2642 |

| **cca-miR8689** | | |
| --- | --- | --- |
| Target_Acc. | Inhibition | Target_Desc. |
| AT1G18120.1 | Cleavage | \| Symbols: \| pseudogene, putative myrosinase-associated protein, blastp match of 46% identity and 1.7e-57 P-value to GP\|6522943\|emb\|CAB62165.1\|\|AJ223307 myrosinase-associated protein {Brassica napus} \| chr1:6232092-6233926 FORWARD LENGTH=1835 |
| AT3G53690.1 | Translation | \| Symbols: \| RING/U-box superfamily protein \| chr3:19898788-19900065 REVERSE LENGTH=1193 |
| AT5G10790.1 | Translation | \| Symbols: UBP22 \| ubiquitin-specific protease 22 \| chr5:3410539-3412633 FORWARD LENGTH=1847 |
| AT1G49750.1 | Translation | \| Symbols: \| Leucine-rich repeat (LRR) family protein \| chr1:18410929-18412801 REVERSE LENGTH=1755 |
| AT1G47380.1 | Cleavage | \| Symbols: \| Protein phosphatase 2C family protein \| chr1:17372537-17376075 REVERSE LENGTH=2046 |
| AT2G22060.1 | Cleavage | \| Symbols: \| BEST Arabidopsis thaliana protein match is: Galactose oxidase/kelch repeat superfamily protein (TAIR:AT2G22030.1); Has 148 Blast hits to 148 proteins in 2 species: Archae - 0; Bacteria - 0; Metazoa - 0; Fungi - 0; Plants - 148; Viruses - 0; Other Eukaryotes - 0 (source: NCBI BLink). \| chr2:9380992-9382145 FORWARD LENGTH=1154 |
| AT1G62085.1 | Translation | \| Symbols: \| Mitochondrial transcription termination factor family protein \| chr1:22948757-22950142 REVERSE LENGTH=1386 |
| AT5G28150.1 | Cleavage | \| Symbols: \| Plant protein of unknown function (DUF868) \| chr5:10135434-10136977 FORWARD LENGTH=1453 |
| AT4G24400.2 | Cleavage | \| Symbols: CIPK8, PKS11 \| CBL-interacting protein kinase 8 \| chr4:12617256-12620684 FORWARD LENGTH=1615 |
| AT4G24400.1 | Cleavage | \| Symbols: CIPK8, SnRK3.13, PKS11, ATCIPK8 \| CBL-interacting protein kinase 8 \| chr4:12617256-12620684 FORWARD LENGTH=1664 |
| AT1G74350.1 | Translation | \| Symbols: \| Intron maturase, type II family protein \| chr1:27949022-27951283 REVERSE LENGTH=2262 |
| AT1G11710.1 | Cleavage | \| Symbols: \| Pentatricopeptide repeat (PPR) superfamily protein \| chr1:3948714-3951359 FORWARD LENGTH=2646 |
| AT3G44705.1 | Cleavage | \| Symbols: \| transposable element gene \| chr3:16244952-16250205 FORWARD LENGTH=5254 |
| AT5G12860.2 | Cleavage | \| Symbols: DiT1 \| dicarboxylate transporter 1 \| chr5:4059686-4061950 REVERSE LENGTH=1866 |
| AT5G12860.1 | Cleavage | \| Symbols: DiT1 \| dicarboxylate transporter 1 \| chr5:4059686-4061950 REVERSE LENGTH=1946 |
| AT5G54480.1 | Cleavage | \| Symbols: \| Protein of unknown function (DUF630 and DUF632) \| chr5:22118004-22120166 FORWARD LENGTH=2163 |
| AT1G49910.1 | Cleavage | \| Symbols: BUB3.2 \| Transducin/WD40 repeat-like superfamily protein \| chr1:18479025-18481475 FORWARD LENGTH=1224 |
| AT5G27900.1 | Cleavage | \| Symbols: \| transposable element gene \| chr5:9913203-9918023 FORWARD LENGTH=4821 |
| AT1G49005.1 | Cleavage | \| Symbols: CLE11 \| CLAVATA3/ESR-RELATED 11 \| chr1:18128686-18131522 REVERSE LENGTH=582 |
| AT1G19970.1 | Cleavage | \| Symbols: \| ER lumen protein retaining receptor family protein \| chr1:6931012-6932689 REVERSE LENGTH=1093 |
| AT3G03890.2 | Cleavage | \| Symbols: \| FMN binding \| chr3:999467-1002083 REVERSE LENGTH=1273 |
| AT1G03340.1 | Cleavage | \| Symbols: \| unknown protein; BEST Arabidopsis thaliana protein match is: unknown protein (TAIR:AT4G02920.1); Has 44 Blast hits to 41 proteins in 13 species: Archae - 0; Bacteria - 1; Metazoa - 0; Fungi - 0; Plants - 43; Viruses - 0; Other Eukaryotes - 0 (source: NCBI BLink). \| chr1:819662-821317 FORWARD LENGTH=1298 |
| AT3G07550.2 | Cleavage | \| Symbols: \| RNI-like superfamily protein \| chr3:2409727-2411294 FORWARD LENGTH=1568 |
| AT3G03890.1 | Cleavage | \| Symbols: \| FMN binding \| chr3:999467-1002484 REVERSE LENGTH=1654 |
| AT3G07550.1 | Cleavage | \| Symbols: \| RNI-like superfamily protein \| chr3:2409477-2411294 FORWARD LENGTH=1708 |
| AT1G45170.2 | Translation | \| Symbols: \| unknown protein; FUNCTIONS IN: molecular_function unknown; INVOLVED IN: biological_process unknown; LOCATED IN: cellular_component unknown; EXPRESSED IN: 24 plant structures; EXPRESSED DURING: 13 growth stages; BEST Arabidopsis thaliana protein match is: unknown protein (TAIR:AT5G42960.1); Has 60 Blast hits to 60 proteins in 18 species: Archae - 0; Bacteria - 0; Metazoa - 0; Fungi - 0; Plants - 60; Viruses - 0; Other Eukaryotes - 0 (source: NCBI BLink). \| chr1:17095980-17098006 REVERSE LENGTH=853 |
| AT5G35918.1 | Translation | \| Symbols: \| transposable element gene \| chr5:14051737-14052722 FORWARD LENGTH=986 |
| AT1G45170.1 | Translation | \| Symbols: \| unknown protein; BEST Arabidopsis thaliana protein match is: unknown protein (TAIR:AT5G42960.1); Has 62 Blast hits to 62 proteins in 18 species: Archae - 0; Bacteria - 0; Metazoa - 0; Fungi - 0; Plants - 62; Viruses - 0; Other Eukaryotes - 0 (source: NCBI BLink). \| chr1:17095980-17098006 REVERSE LENGTH=991 |
| AT2G15555.1 | Cleavage | \| Symbols: \| other RNA \| chr2:6785904-6786976 REVERSE LENGTH=1073 |
| AT1G70505.1 | Cleavage | \| Symbols: \| unknown protein; BEST Arabidopsis thaliana protein match is: unknown protein (TAIR:AT1G10660.1); Has 141 Blast hits to 140 proteins in 16 species: Archae - 0; Bacteria - 0; Metazoa - 4; Fungi - 0; Plants - 135; Viruses - 0; Other Eukaryotes - 2 (source: NCBI BLink). \| chr1:26570647-26572463 FORWARD LENGTH=1077 |
| AT2G21190.1 | Cleavage | \| Symbols: \| ER lumen protein retaining receptor family protein \| chr2:9080865-9082904 FORWARD LENGTH=1123 |
| AT5G22250.1 | Translation | \| Symbols: \| Polynucleotidyl transferase, ribonuclease H-like superfamily protein \| chr5:7365532-7366778 REVERSE LENGTH=1247 |
| AT2G26480.1 | Translation | \| Symbols: UGT76D1 \| UDP-glucosyl transferase 76D1 \| chr2:11263963-11265684 FORWARD LENGTH=1471 |
| AT2G22690.2 | Cleavage | \| Symbols: \| zinc ion binding \| chr2:9649690-9651439 REVERSE LENGTH=1553 |
| AT3G12200.1 | Cleavage | \| Symbols: AtNek7, Nek7 \| NIMA-related kinase 7 \| chr3:3886992-3890826 REVERSE LENGTH=2173 |
| AT3G12200.2 | Cleavage | \| Symbols: Nek7 \| NIMA-related kinase 7 \| chr3:3887108-3891000 REVERSE LENGTH=2261 |
| AT4G37640.1 | Cleavage | \| Symbols: ACA2 \| calcium ATPase 2 \| chr4:17682977-17686941 REVERSE LENGTH=3426 |
| AT2G39580.1 | Cleavage | \| Symbols: \| CONTAINS InterPro DOMAIN/s: Putative zinc-finger domain (InterPro:IPR019607); Has 249 Blast hits to 219 proteins in 85 species: Archae - 0; Bacteria - 144; Metazoa - 29; Fungi - 8; Plants - 50; Viruses - 0; Other Eukaryotes - 18 (source: NCBI BLink). \| chr2:16510425-16517073 FORWARD LENGTH=4974 |
| AT1G36250.1 | Cleavage | \| Symbols: \| transposable element gene \| chr1:13617739-13623656 FORWARD LENGTH=5918 |
| AT3G10585.1 | Cleavage | \| Symbols: \| Homeodomain-like superfamily protein \| chr3:3308489-3309509 REVERSE LENGTH=656 |
| AT1G44920.1 | Cleavage | \| Symbols: \| unknown protein; FUNCTIONS IN: molecular_function unknown; INVOLVED IN: biological_process unknown; LOCATED IN: chloroplast; EXPRESSED IN: 22 plant structures; EXPRESSED DURING: 13 growth stages; CONTAINS InterPro DOMAIN/s: Protein of unknown function DUF3054 (InterPro:IPR021414); Has 246 Blast hits to 246 proteins in 119 species: Archae - 14; Bacteria - 181; Metazoa - 0; Fungi - 0; Plants - 45; Viruses - 0; Other Eukaryotes - 6 (source: NCBI BLink). \| chr1:16982904-16984419 REVERSE LENGTH=1113 |
| AT4G05130.1 | Cleavage | \| Symbols: ATENT4, ENT4 \| equilibrative nucleoside transporter 4 \| chr4:2636875-2638965 REVERSE LENGTH=1257 |
| AT4G02800.1 | Cleavage | \| Symbols: \| unknown protein; FUNCTIONS IN: molecular_function unknown; INVOLVED IN: biological_process unknown; LOCATED IN: chloroplast; EXPRESSED IN: 16 plant structures; EXPRESSED DURING: 9 growth stages; BEST Arabidopsis thaliana protein match is: unknown protein (TAIR:AT5G01970.1); Has 3209 Blast hits to 2720 proteins in 308 species: Archae - 13; Bacteria - 213; Metazoa - 1207; Fungi - 247; Plants - 183; Viruses - 21; Other Eukaryotes - 1325 (source: NCBI BLink). \| chr4:1250015-1251681 FORWARD LENGTH=1316 |
| AT2G39450.1 | Translation | \| Symbols: MTP11, ATMTP11 \| Cation efflux family protein \| chr2:16471632-16473818 REVERSE LENGTH=1380 |
| AT4G01850.1 | Cleavage | \| Symbols: SAM-2, MAT2, SAM2, AtSAM2 \| S-adenosylmethionine synthetase 2 \| chr4:796097-798286 REVERSE LENGTH=1480 |
| AT5G53430.1 | Cleavage | \| Symbols: SDG29, SET29, ATX5 \| SET domain group 29 \| chr5:21677146-21683494 FORWARD LENGTH=3937 |
| AT5G41755.1 | Cleavage | \| Symbols: \| transposable element gene \| chr5:16700028-16705892 FORWARD LENGTH=5865 |
| AT5G63870.1 | Cleavage | \| Symbols: PP7, ATPP7 \| serine/threonine phosphatase 7 \| chr5:25561161-25563248 REVERSE LENGTH=1523 |
| AT5G63870.2 | Cleavage | \| Symbols: PP7 \| serine/threonine phosphatase 7 \| chr5:25561160-25563274 REVERSE LENGTH=1539 |
| AT5G63870.3 | Cleavage | \| Symbols: PP7 \| serine/threonine phosphatase 7 \| chr5:25561172-25563274 REVERSE LENGTH=1615 |
| AT5G22170.1 | Translation | \| Symbols: \| unknown protein; BEST Arabidopsis thaliana protein match is: unknown protein (TAIR:AT5G22150.1); Has 30201 Blast hits to 17322 proteins in 780 species: Archae - 12; Bacteria - 1396; Metazoa - 17338; Fungi - 3422; Plants - 5037; Viruses - 0; Other Eukaryotes - 2996 (source: NCBI BLink). \| chr5:7350575-7351190 FORWARD LENGTH=540 |
| AT5G54480.1 | Translation | \| Symbols: \| Protein of unknown function (DUF630 and DUF632) \| chr5:22118004-22120166 FORWARD LENGTH=2163 |
| AT3G15358.1 | Cleavage | \| Symbols: \| unknown protein; FUNCTIONS IN: molecular_function unknown; INVOLVED IN: biological_process unknown; LOCATED IN: endomembrane system; BEST Arabidopsis thaliana protein match is: unknown protein (TAIR:AT1G53035.1); Has 35333 Blast hits to 34131 proteins in 2444 species: Archae - 798; Bacteria - 22429; Metazoa - 974; Fungi - 991; Plants - 531; Viruses - 0; Other Eukaryotes - 9610 (source: NCBI BLink). \| chr3:5178013-5178838 FORWARD LENGTH=729 |
| AT1G53035.1 | Cleavage | \| Symbols: \| unknown protein; FUNCTIONS IN: molecular_function unknown; INVOLVED IN: biological_process unknown; LOCATED IN: endomembrane system; EXPRESSED IN: 23 plant structures; EXPRESSED DURING: 13 growth stages; BEST Arabidopsis thaliana protein match is: unknown protein (TAIR:AT3G15358.1); Has 49 Blast hits to 49 proteins in 9 species: Archae - 0; Bacteria - 0; Metazoa - 0; Fungi - 0; Plants - 49; Viruses - 0; Other Eukaryotes - 0 (source: NCBI BLink). \| chr1:19761700-19762551 REVERSE LENGTH=852 |
| AT4G23010.2 | Cleavage | \| Symbols: ATUTR2, UTR2 \| UDP-galactose transporter 2 \| chr4:12060131-12062757 REVERSE LENGTH=1547 |
| AT4G23010.1 | Cleavage | \| Symbols: ATUTR2, UTR2 \| UDP-galactose transporter 2 \| chr4:12060131-12062816 REVERSE LENGTH=1555 |
| AT4G23010.3 | Cleavage | \| Symbols: UTR2 \| UDP-galactose transporter 2 \| chr4:12060184-12062858 REVERSE LENGTH=1685 |
| AT1G72820.1 | Cleavage | \| Symbols: \| Mitochondrial substrate carrier family protein \| chr1:27402878-27404673 FORWARD LENGTH=1796 |
| AT2G46500.2 | Cleavage | \| Symbols: ATPI4K GAMMA 4, UBDK GAMMA 4, PI4K GAMMA 4 \| phosphoinositide 4-kinase gamma 4 \| chr2:19086561-19088981 REVERSE LENGTH=2186 |
| AT5G41690.1 | Cleavage | \| Symbols: \| RNA-binding (RRM/RBD/RNP motifs) family protein \| chr5:16670001-16674966 REVERSE LENGTH=2268 |
| AT2G46500.1 | Cleavage | \| Symbols: ATPI4K GAMMA 4, UBDK GAMMA 4, PI4K GAMMA 4 \| phosphoinositide 4-kinase gamma 4 \| chr2:19086561-19088964 REVERSE LENGTH=2311 |
| AT5G20500.1 | Cleavage | \| Symbols: \| Glutaredoxin family protein \| chr5:6938598-6939848 FORWARD LENGTH=645 |
| AT5G44140.1 | Cleavage | \| Symbols: ATPHB7, PHB7 \| prohibitin 7 \| chr5:17762491-17763629 FORWARD LENGTH=837 |
| AT1G24148.2 | Cleavage | \| Symbols: \| unknown protein; FUNCTIONS IN: molecular_function unknown; INVOLVED IN: biological_process unknown; LOCATED IN: cellular_component unknown. \| chr1:8544420-8545551 FORWARD LENGTH=1132 |
| AT3G29638.1 | Cleavage | \| Symbols: \| General transcription factor 2-related zinc finger protein \| chr3:11473584-11475559 FORWARD LENGTH=1239 |
| AT3G05155.1 | Cleavage | \| Symbols: \| Major facilitator superfamily protein \| chr3:1448570-1451213 FORWARD LENGTH=1287 |
| AT4G11330.1 | Cleavage | \| Symbols: ATMPK5, MPK5 \| MAP kinase 5 \| chr4:6892055-6894149 FORWARD LENGTH=1523 |
| AT1G16180.1 | Translation | \| Symbols: \| Serinc-domain containing serine and sphingolipid biosynthesis protein \| chr1:5540427-5542902 FORWARD LENGTH=1618 |
| AT1G16180.2 | Translation | \| Symbols: \| Serinc-domain containing serine and sphingolipid biosynthesis protein \| chr1:5540427-5542902 FORWARD LENGTH=1621 |
| AT2G33835.1 | Cleavage | \| Symbols: FES1 \| Zinc finger C-x8-C-x5-C-x3-H type family protein \| chr2:14311787-14314700 REVERSE LENGTH=2256 |
| AT5G04480.2 | Translation | \| Symbols: \| UDP-Glycosyltransferase superfamily protein \| chr5:1271518-1277946 REVERSE LENGTH=3629 |
| AT3G28415.1 | Cleavage | \| Symbols: \| ABC transporter family protein \| chr3:10647123-10651540 REVERSE LENGTH=3666 |
| AT5G04480.1 | Translation | \| Symbols: \| UDP-Glycosyltransferase superfamily protein \| chr5:1271513-1277985 REVERSE LENGTH=3718 |
| AT2G23330.1 | Translation | \| Symbols: \| transposable element gene \| chr2:9927826-9932319 REVERSE LENGTH=4494 |
| AT3G54560.1 | Translation | \| Symbols: HTA11 \| histone H2A 11 \| chr3:20196266-20197650 FORWARD LENGTH=689 |
| AT5G54790.1 | Translation | \| Symbols: \| unknown protein; BEST Arabidopsis thaliana protein match is: unknown protein (TAIR:AT1G50930.1); Has 53 Blast hits to 53 proteins in 10 species: Archae - 0; Bacteria - 0; Metazoa - 0; Fungi - 0; Plants - 53; Viruses - 0; Other Eukaryotes - 0 (source: NCBI BLink). \| chr5:22253986-22255663 REVERSE LENGTH=881 |
| AT4G09960.2 | Cleavage | \| Symbols: STK, AGL11 \| K-box region and MADS-box transcription factor family protein \| chr4:6236485-6240820 REVERSE LENGTH=962 |
| AT4G09960.1 | Cleavage | \| Symbols: STK, AGL11 \| K-box region and MADS-box transcription factor family protein \| chr4:6236473-6240820 REVERSE LENGTH=1016 |
| AT2G28690.1 | Translation | \| Symbols: \| Protein of unknown function (DUF1635) \| chr2:12306890-12308628 REVERSE LENGTH=1097 |
| AT2G38740.1 | Translation | \| Symbols: \| Haloacid dehalogenase-like hydrolase (HAD) superfamily protein \| chr2:16194327-16196113 REVERSE LENGTH=1165 |
| AT4G09960.3 | Cleavage | \| Symbols: STK, AGL11 \| K-box region and MADS-box transcription factor family protein \| chr4:6236473-6240681 REVERSE LENGTH=1198 |
| AT4G09960.4 | Cleavage | \| Symbols: STK \| K-box region and MADS-box transcription factor family protein \| chr4:6236375-6240932 REVERSE LENGTH=1238 |
| AT5G19120.1 | Cleavage | \| Symbols: \| Eukaryotic aspartyl protease family protein \| chr5:6414488-6415907 FORWARD LENGTH=1420 |
| AT1G70090.2 | Cleavage | \| Symbols: GATL9, LGT8 \| glucosyl transferase family 8 \| chr1:26400790-26402727 FORWARD LENGTH=1540 |
| AT3G09580.1 | Cleavage | \| Symbols: \| FAD/NAD(P)-binding oxidoreductase family protein \| chr3:2942504-2944078 REVERSE LENGTH=1575 |
| AT1G70090.1 | Cleavage | \| Symbols: GATL9, LGT8 \| glucosyl transferase family 8 \| chr1:26400790-26402397 FORWARD LENGTH=1608 |
| AT1G49600.1 | Cleavage | \| Symbols: ATRBP47A, RBP47A \| RNA-binding protein 47A \| chr1:18356886-18360150 REVERSE LENGTH=1688 |
| AT2G18193.1 | Cleavage | \| Symbols: \| P-loop containing nucleoside triphosphate hydrolases superfamily protein \| chr2:7917505-7919277 REVERSE LENGTH=1697 |
| AT3G19130.1 | Cleavage | \| Symbols: ATRBP47B, RBP47B \| RNA-binding protein 47B \| chr3:6611198-6614044 REVERSE LENGTH=1729 |
| AT4G02005.2 | Translation | \| Symbols: \| Unknown gene \| chr4:876908-879741 REVERSE LENGTH=1826 |
| AT4G02005.1 | Translation | \| Symbols: \| other RNA \| chr4:876908-879181 REVERSE LENGTH=1894 |
| AT5G52410.1 | Cleavage | \| Symbols: \| CONTAINS InterPro DOMAIN/s: S-layer homology domain (InterPro:IPR001119); BEST Arabidopsis thaliana protein match is: unknown protein (TAIR:AT5G23890.1); Has 30201 Blast hits to 17322 proteins in 780 species: Archae - 12; Bacteria - 1396; Metazoa - 17338; Fungi - 3422; Plants - 5037; Viruses - 0; Other Eukaryotes - 2996 (source: NCBI BLink). \| chr5:21275956-21279444 FORWARD LENGTH=2440 |
| AT5G52410.2 | Cleavage | \| Symbols: \| INVOLVED IN: biological_process unknown; LOCATED IN: chloroplast; EXPRESSED IN: 13 plant structures; EXPRESSED DURING: 6 growth stages; CONTAINS InterPro DOMAIN/s: S-layer homology domain (InterPro:IPR001119); BEST Arabidopsis thaliana protein match is: unknown protein (TAIR:AT5G23890.1); Has 35333 Blast hits to 34131 proteins in 2444 species: Archae - 798; Bacteria - 22429; Metazoa - 974; Fungi - 991; Plants - 531; Viruses - 0; Other Eukaryotes - 9610 (source: NCBI BLink). \| chr5:21275956-21279443 FORWARD LENGTH=2461 |
| AT5G64570.1 | Cleavage | \| Symbols: XYL4, ATBXL4 \| beta-D-xylosidase 4 \| chr5:25810115-25813335 REVERSE LENGTH=2493 |
| AT3G45775.1 | Translation | \| Symbols: \| transposable element gene \| chr3:16809432-16813871 REVERSE LENGTH=4440 |
| AT5G01335.1 | Translation | \| Symbols: \| transposable element gene \| chr5:135831-141287 FORWARD LENGTH=5457 |
| AT5G04560.2 | Cleavage | \| Symbols: DME \| HhH-GPD base excision DNA repair family protein \| chr5:1309202-1318401 FORWARD LENGTH=6475 |
| AT5G04560.1 | Cleavage | \| Symbols: DME \| HhH-GPD base excision DNA repair family protein \| chr5:1309193-1318386 FORWARD LENGTH=6963 |
| AT5G22340.2 | Cleavage | \| Symbols: \| unknown protein; FUNCTIONS IN: molecular_function unknown; INVOLVED IN: biological_process unknown; LOCATED IN: chloroplast; EXPRESSED IN: 22 plant structures; EXPRESSED DURING: 13 growth stages; Has 43 Blast hits to 43 proteins in 15 species: Archae - 0; Bacteria - 0; Metazoa - 0; Fungi - 0; Plants - 42; Viruses - 0; Other Eukaryotes - 1 (source: NCBI BLink). \| chr5:7394473-7396875 FORWARD LENGTH=1325 |
| AT5G22340.1 | Cleavage | \| Symbols: \| unknown protein; FUNCTIONS IN: molecular_function unknown; INVOLVED IN: biological_process unknown; LOCATED IN: chloroplast; EXPRESSED IN: 22 plant structures; EXPRESSED DURING: 13 growth stages; Has 58 Blast hits to 58 proteins in 20 species: Archae - 0; Bacteria - 0; Metazoa - 0; Fungi - 0; Plants - 57; Viruses - 0; Other Eukaryotes - 1 (source: NCBI BLink). \| chr5:7394444-7396852 FORWARD LENGTH=1335 |
| AT3G21040.1 | Translation | \| Symbols: \| transposable element gene \| chr3:7372760-7374364 FORWARD LENGTH=1605 |
| AT2G24240.1 | Cleavage | \| Symbols: \| BTB/POZ domain with WD40/YVTN repeat-like protein \| chr2:10310648-10312328 FORWARD LENGTH=1681 |
| AT2G44210.1 | Cleavage | \| Symbols: \| Protein of Unknown Function (DUF239) \| chr2:18280623-18282876 FORWARD LENGTH=1719 |
| AT2G44210.2 | Cleavage | \| Symbols: \| Protein of Unknown Function (DUF239) \| chr2:18280623-18282876 FORWARD LENGTH=1809 |
| AT5G14880.1 | Translation | \| Symbols: \| Potassium transporter family protein \| chr5:4814244-4817667 FORWARD LENGTH=2346 |
| AT4G01030.1 | Cleavage | \| Symbols: \| pentatricopeptide (PPR) repeat-containing protein \| chr4:447941-450642 REVERSE LENGTH=2702 |
| AT3G24340.1 | Cleavage | \| Symbols: chr40 \| chromatin remodeling 40 \| chr3:8832085-8835722 REVERSE LENGTH=3399 |
| AT1G17360.1 | Cleavage | \| Symbols: \| BEST Arabidopsis thaliana protein match is: COP1-interacting protein-related (TAIR:AT1G72410.1); Has 9949 Blast hits to 7480 proteins in 576 species: Archae - 12; Bacteria - 1007; Metazoa - 3636; Fungi - 982; Plants - 444; Viruses - 50; Other Eukaryotes - 3818 (source: NCBI BLink). \| chr1:5946764-5951473 FORWARD LENGTH=3530 |
| AT1G32120.1 | Cleavage | \| Symbols: \| FUNCTIONS IN: molecular_function unknown; INVOLVED IN: biological_process unknown; LOCATED IN: membrane; EXPRESSED IN: 14 plant structures; EXPRESSED DURING: 4 anthesis, C globular stage, F mature embryo stage, petal differentiation and expansion stage, E expanded cotyledon stage; CONTAINS InterPro DOMAIN/s: Aminotransferase-like, plant mobile domain (InterPro:IPR019557), Protein of unknown function DUF716 (InterPro:IPR006904); BEST Arabidopsis thaliana protein match is: Aminotransferase-like, plant mobile domain family protein (TAIR:AT1G51538.1); Has 16736 Blast hits to 9656 proteins in 576 species: Archae - 4; Bacteria - 1182; Metazoa - 7098; Fungi - 2631; Plants - 1178; Viruses - 174; Other Eukaryotes - 4469 (source: NCBI BLink). \| chr1:11552926-11558777 FORWARD LENGTH=3790 |
| AT3G30838.1 | Cleavage | \| Symbols: \| transposable element gene \| chr3:12579885-12585786 REVERSE LENGTH=5902 |
| AT3G14270.1 | Cleavage | \| Symbols: FAB1B \| phosphatidylinositol-4-phosphate 5-kinase family protein \| chr3:4753925-4761456 FORWARD LENGTH=5987 |
| AT2G07000.1 | Translation | \| Symbols: \| unknown protein; Has 1 Blast hits to 1 proteins in 1 species: Archae - 0; Bacteria - 0; Metazoa - 0; Fungi - 0; Plants - 1; Viruses - 0; Other Eukaryotes - 0 (source: NCBI BLink). \| chr2:2901870-2902290 REVERSE LENGTH=322 |
| AT3G46090.1 | Cleavage | \| Symbols: ZAT7 \| C2H2 and C2HC zinc fingers superfamily protein \| chr3:16926216-16926850 REVERSE LENGTH=635 |
| AT3G43826.1 | Cleavage | \| Symbols: \| pseudogene, hypothetical protein \| chr3:15675393-15676148 REVERSE LENGTH=756 |
| AT3G45638.2 | Translation | \| Symbols: \| other RNA \| chr3:16755470-16756290 REVERSE LENGTH=821 |
| AT3G45638.1 | Translation | \| Symbols: \| other RNA \| chr3:16755215-16756263 REVERSE LENGTH=878 |
| AT5G13790.1 | Cleavage | \| Symbols: AGL15 \| AGAMOUS-like 15 \| chr5:4449014-4450843 REVERSE LENGTH=962 |
| AT2G26455.1 | Translation | \| Symbols: \| pseudogene, similar to Ubiquitin conjugating enzyme 7 interacting protein 4 (UbcM4- interacting protein 4) (RING finger protein 144). (Mouse), contains a Prosite:PS00518: Zinc finger, C3HC4 type (RING finger), signature and Pfam:PF01485 IBR domain; blastp match of 25% identity and 5.7e-18 P-value to SP\|Q925F3\|U7I4_MOUSE Ubiquitin conjugating enzyme 7 interacting protein 4 (UbcM4- interacting protein 4) (RING finger protein 144). (Mouse) {Mus musculus} \| chr2:11253700-11254930 REVERSE LENGTH=1231 |
| AT1G22750.4 | Cleavage | \| Symbols: \| unknown protein; CONTAINS InterPro DOMAIN/s: Protein of unknown function DUF1475 (InterPro:IPR009943); Has 186 Blast hits to 155 proteins in 21 species: Archae - 0; Bacteria - 8; Metazoa - 3; Fungi - 0; Plants - 65; Viruses - 0; Other Eukaryotes - 110 (source: NCBI BLink). \| chr1:8050860-8052904 FORWARD LENGTH=1015 |
| AT1G22750.3 | Cleavage | \| Symbols: \| unknown protein; FUNCTIONS IN: molecular_function unknown; INVOLVED IN: biological_process unknown; LOCATED IN: vacuole; EXPRESSED IN: 23 plant structures; EXPRESSED DURING: 15 growth stages; CONTAINS InterPro DOMAIN/s: Protein of unknown function DUF1475 (InterPro:IPR009943); Has 35333 Blast hits to 34131 proteins in 2444 species: Archae - 798; Bacteria - 22429; Metazoa - 974; Fungi - 991; Plants - 531; Viruses - 0; Other Eukaryotes - 9610 (source: NCBI BLink). \| chr1:8050860-8052904 FORWARD LENGTH=1063 |
| AT1G22750.1 | Cleavage | \| Symbols: \| unknown protein; FUNCTIONS IN: molecular_function unknown; INVOLVED IN: biological_process unknown; LOCATED IN: vacuole; EXPRESSED IN: 23 plant structures; EXPRESSED DURING: 15 growth stages; CONTAINS InterPro DOMAIN/s: Protein of unknown function DUF1475 (InterPro:IPR009943); Has 185 Blast hits to 155 proteins in 21 species: Archae - 0; Bacteria - 8; Metazoa - 3; Fungi - 0; Plants - 64; Viruses - 0; Other Eukaryotes - 110 (source: NCBI BLink). \| chr1:8050860-8052904 FORWARD LENGTH=1072 |
| AT1G22750.2 | Cleavage | \| Symbols: \| unknown protein; FUNCTIONS IN: molecular_function unknown; INVOLVED IN: biological_process unknown; LOCATED IN: vacuole; EXPRESSED IN: 23 plant structures; EXPRESSED DURING: 15 growth stages; CONTAINS InterPro DOMAIN/s: Protein of unknown function DUF1475 (InterPro:IPR009943); Has 185 Blast hits to 155 proteins in 21 species: Archae - 0; Bacteria - 8; Metazoa - 3; Fungi - 0; Plants - 64; Viruses - 0; Other Eukaryotes - 110 (source: NCBI BLink). \| chr1:8050860-8052930 FORWARD LENGTH=1094 |
| AT3G42436.1 | Cleavage | \| Symbols: \| transposable element gene \| chr3:14570230-14571625 FORWARD LENGTH=1208 |
| AT3G18780.2 | Cleavage | \| Symbols: ACT2, DER1, LSR2, ENL2 \| actin 2 \| chr3:6474842-6477204 FORWARD LENGTH=1756 |
| AT3G18780.1 | Cleavage | \| Symbols: ACT2, DER1, LSR2, ENL2 \| actin 2 \| chr3:6474871-6477204 FORWARD LENGTH=1813 |
| AT5G22550.1 | Translation | \| Symbols: \| Plant protein of unknown function (DUF247) \| chr5:7483870-7485482 REVERSE LENGTH=1466 |
| AT5G22550.2 | Translation | \| Symbols: \| Plant protein of unknown function (DUF247) \| chr5:7483873-7485458 REVERSE LENGTH=1586 |
| AT4G26100.1 | Cleavage | \| Symbols: CK1, CKL1 \| casein kinase 1 \| chr4:13227450-13230797 REVERSE LENGTH=2077 |
| AT1G26890.1 | Cleavage | \| Symbols: \| FBD, F-box and Leucine Rich Repeat domains containing protein \| chr1:9317589-9319246 FORWARD LENGTH=1428 |
| AT3G05940.1 | Cleavage | \| Symbols: \| Protein of unknown function (DUF300) \| chr3:1777364-1780254 REVERSE LENGTH=1858 |
| AT5G55330.1 | Translation | \| Symbols: \| MBOAT (membrane bound O-acyl transferase) family protein \| chr5:22437825-22438865 REVERSE LENGTH=1041 |
| AT4G10613.1 | Cleavage | \| Symbols: \| RNA-directed DNA polymerase (reverse transcriptase)-related family protein \| chr4:6560363-6561726 FORWARD LENGTH=1364 |
| AT4G31020.2 | Cleavage | \| Symbols: \| alpha/beta-Hydrolases superfamily protein \| chr4:15108101-15110147 REVERSE LENGTH=1491 |
| AT1G78560.1 | Cleavage | \| Symbols: \| Sodium Bile acid symporter family \| chr1:29546605-29548824 REVERSE LENGTH=1507 |
| AT4G28620.1 | Translation | \| Symbols: ATATM2, ATM2 \| ABC transporter of the mitochondrion 2 \| chr4:14135526-14137953 REVERSE LENGTH=2043 |
| AT1G69610.1 | Cleavage | \| Symbols: \| Protein of unknown function (DUF1666) \| chr1:26186954-26189489 FORWARD LENGTH=2051 |
| AT5G61690.1 | Cleavage | \| Symbols: ATATH15, ATH15 \| ABC2 homolog 15 \| chr5:24789495-24793487 REVERSE LENGTH=2760 |
| AT5G61730.1 | Cleavage | \| Symbols: ATATH11, ATH11 \| ABC2 homolog 11 \| chr5:24803583-24807898 REVERSE LENGTH=2823 |
| AT5G45480.1 | Cleavage | \| Symbols: \| Protein of unknown function (DUF594) \| chr5:18426104-18429090 REVERSE LENGTH=2987 |
| AT3G47730.1 | Cleavage | \| Symbols: ATATH1, ATH1, ABCA2 \| ATP-binding cassette A2 \| chr3:17594060-17598896 REVERSE LENGTH=3302 |
| AT3G43436.1 | Cleavage | \| Symbols: \| transposable element gene \| chr3:15361697-15365347 FORWARD LENGTH=3651 |
| AT1G35870.1 | Cleavage | \| Symbols: \| transposable element gene \| chr1:13335257-13339587 FORWARD LENGTH=4331 |
| AT4G34790.1 | Cleavage | \| Symbols: \| SAUR-like auxin-responsive protein family \| chr4:16594469-16595153 FORWARD LENGTH=685 |
| AT5G11690.1 | Cleavage | \| Symbols: ATTIM17-3, TIM17-3 \| translocase inner membrane subunit 17-3 \| chr5:3761244-3762346 FORWARD LENGTH=856 |
| AT3G49980.1 | Cleavage | \| Symbols: \| F-box and associated interaction domains-containing protein \| chr3:18530329-18531477 REVERSE LENGTH=1149 |
| AT4G35985.1 | Cleavage | \| Symbols: \| Senescence/dehydration-associated protein-related \| chr4:17032268-17033862 REVERSE LENGTH=1347 |
| AT5G17780.1 | Cleavage | \| Symbols: \| alpha/beta-Hydrolases superfamily protein \| chr5:5867283-5869050 REVERSE LENGTH=1474 |
| AT5G17780.1 | Cleavage | \| Symbols: \| alpha/beta-Hydrolases superfamily protein \| chr5:5867283-5869050 REVERSE LENGTH=1474 |
| AT5G17780.2 | Cleavage | \| Symbols: \| alpha/beta-Hydrolases superfamily protein \| chr5:5867283-5869050 REVERSE LENGTH=1480 |
| AT5G17780.2 | Cleavage | \| Symbols: \| alpha/beta-Hydrolases superfamily protein \| chr5:5867283-5869050 REVERSE LENGTH=1480 |
| AT2G20590.1 | Cleavage | \| Symbols: \| Reticulon family protein \| chr2:8867117-8869228 REVERSE LENGTH=1484 |
| AT2G20590.2 | Cleavage | \| Symbols: \| Reticulon family protein \| chr2:8867120-8869256 REVERSE LENGTH=1640 |
| AT4G11570.2 | Translation | \| Symbols: \| Haloacid dehalogenase-like hydrolase (HAD) superfamily protein \| chr4:7004335-7006081 FORWARD LENGTH=1747 |
| AT4G03745.1 | Cleavage | \| Symbols: \| transposable element gene \| chr4:1665242-1667033 REVERSE LENGTH=1792 |
| AT1G11780.1 | Translation | \| Symbols: \| oxidoreductase, 2OG-Fe(II) oxygenase family protein \| chr1:3976839-3979195 REVERSE LENGTH=1831 |
| AT1G22710.1 | Cleavage | \| Symbols: SUC2, SUT1, ATSUC2 \| sucrose-proton symporter 2 \| chr1:8030641-8033117 REVERSE LENGTH=1956 |
| AT1G74780.1 | Cleavage | \| Symbols: \| Nodulin-like / Major Facilitator Superfamily protein \| chr1:28095815-28098185 FORWARD LENGTH=1976 |
| AT2G45830.2 | Cleavage | \| Symbols: DTA2 \| downstream target of AGL15 2 \| chr2:18866061-18868542 FORWARD LENGTH=2136 |
| AT2G45830.1 | Cleavage | \| Symbols: DTA2 \| downstream target of AGL15 2 \| chr2:18866061-18868542 FORWARD LENGTH=2033 |
| AT3G57150.1 | Cleavage | \| Symbols: NAP57, AtNAP57, CBF5, AtCBF5 \| homologue of NAP57 \| chr3:21153973-21156009 REVERSE LENGTH=2037 |
| AT3G26782.1 | Translation | \| Symbols: \| Tetratricopeptide repeat (TPR)-like superfamily protein \| chr3:9850570-9852864 FORWARD LENGTH=2186 |
| AT4G03730.1 | Cleavage | \| Symbols: \| transposable element gene \| chr4:1653587-1655944 REVERSE LENGTH=2358 |
| AT2G21300.2 | Translation | \| Symbols: \| ATP binding microtubule motor family protein \| chr2:9114154-9118687 REVERSE LENGTH=2975 |
| AT2G21300.1 | Translation | \| Symbols: \| ATP binding microtubule motor family protein \| chr2:9114218-9119946 REVERSE LENGTH=3009 |
| AT2G22950.1 | Cleavage | \| Symbols: \| Cation transporter/ E1-E2 ATPase family protein \| chr2:9766127-9769766 FORWARD LENGTH=3048 |
| AT5G06810.1 | Translation | \| Symbols: \| Mitochondrial transcription termination factor family protein \| chr5:2108493-2112256 FORWARD LENGTH=3426 |
| AT5G47490.1 | Cleavage | \| Symbols: \| RGPR-related \| chr5:19264675-19270800 FORWARD LENGTH=4399 |
| AT1G60983.1 | Translation | \| Symbols: SCRL8 \| SCR-like 8 \| chr1:22455320-22455589 FORWARD LENGTH=270 |
| AT5G42445.1 | Translation | \| Symbols: \| 60S ribosomal protein L3 (RPL3C), pseudogene, temporary automated functional assignment; blastp match of 61% identity and 1.3e-25 P-value to SP\|P35684\|RL3_ORYSA 60S ribosomal protein L3. (Rice) {Oryza sativa} \| chr5:16976272-16976613 FORWARD LENGTH=342 |
| AT1G74000.1 | Cleavage | \| Symbols: SS3 \| strictosidine synthase 3 \| chr1:27829142-27831431 REVERSE LENGTH=1144 |
| AT1G08110.2 | Cleavage | \| Symbols: \| lactoylglutathione lyase family protein / glyoxalase I family protein \| chr1:2535409-2537930 FORWARD LENGTH=1151 |
| AT4G00955.1 | Cleavage | \| Symbols: \| FUNCTIONS IN: molecular_function unknown; INVOLVED IN: biological_process unknown; LOCATED IN: endomembrane system; EXPRESSED IN: 14 plant structures; EXPRESSED DURING: 9 growth stages; CONTAINS InterPro DOMAIN/s: EGF-like (InterPro:IPR006210); BEST Arabidopsis thaliana protein match is: Protein kinase superfamily protein (TAIR:AT2G23450.1); Has 94 Blast hits to 88 proteins in 13 species: Archae - 0; Bacteria - 0; Metazoa - 10; Fungi - 0; Plants - 84; Viruses - 0; Other Eukaryotes - 0 (source: NCBI BLink). \| chr4:412118-413327 FORWARD LENGTH=1210 |
| AT2G03915.1 | Cleavage | \| Symbols: \| transposable element gene \| chr2:1199311-1200432 REVERSE LENGTH=1122 |
| AT1G65820.1 | Cleavage | \| Symbols: \| microsomal glutathione s-transferase, putative \| chr1:24485101-24486892 FORWARD LENGTH=763 |
| AT1G65820.2 | Cleavage | \| Symbols: \| microsomal glutathione s-transferase, putative \| chr1:24485101-24486892 FORWARD LENGTH=811 |
| AT1G65820.3 | Cleavage | \| Symbols: \| microsomal glutathione s-transferase, putative \| chr1:24485101-24486892 FORWARD LENGTH=907 |
| AT5G13610.1 | Cleavage | \| Symbols: \| Protein of unknown function (DUF155) \| chr5:4383013-4385029 FORWARD LENGTH=1490 |
| AT5G19500.1 | Translation | \| Symbols: \| Tryptophan/tyrosine permease \| chr5:6578952-6581945 FORWARD LENGTH=1831 |
| AT1G13960.2 | Cleavage | \| Symbols: WRKY4 \| WRKY DNA-binding protein 4 \| chr1:4776463-4779350 FORWARD LENGTH=1930 |
| AT5G58520.1 | Translation | \| Symbols: \| Protein kinase superfamily protein \| chr5:23655234-23658162 FORWARD LENGTH=2112 |
| AT1G10240.1 | Cleavage | \| Symbols: FRS11 \| FAR1-related sequence 11 \| chr1:3356680-3359838 REVERSE LENGTH=2295 |
| AT5G55480.1 | Cleavage | \| Symbols: SVL1 \| SHV3-like 1 \| chr5:22474230-22478132 FORWARD LENGTH=2661 |
| AT1G53350.1 | Cleavage | \| Symbols: \| Disease resistance protein (CC-NBS-LRR class) family \| chr1:19903899-19907515 FORWARD LENGTH=2784 |
| AT2G16390.1 | Cleavage | \| Symbols: DRD1, CHR35, DMS1 \| SNF2 domain-containing protein / helicase domain-containing protein \| chr2:7097280-7101261 FORWARD LENGTH=3104 |
| AT2G15800.1 | Cleavage | \| Symbols: \| transposable element gene \| chr2:6880825-6882741 FORWARD LENGTH=1917 |
| AT3G26614.1 | Translation | \| Symbols: \| transposable element gene \| chr3:9784890-9787571 REVERSE LENGTH=2682 |
| AT4G01975.1 | Translation | \| Symbols: \| transposable element gene \| chr4:858636-862475 REVERSE LENGTH=3840 |
| AT1G79350.1 | Cleavage | \| Symbols: EMB1135 \| RING/FYVE/PHD zinc finger superfamily protein \| chr1:29844633-29853414 REVERSE LENGTH=4283 |
| AT5G57625.1 | Translation | \| Symbols: \| CAP (Cysteine-rich secretory proteins, Antigen 5, and Pathogenesis-related 1 protein) superfamily protein \| chr5:23337850-23338807 FORWARD LENGTH=869 |
| AT5G51140.2 | Cleavage | \| Symbols: \| Pseudouridine synthase family protein \| chr5:20784082-20786816 REVERSE LENGTH=1277 |
