## Supplementary File 1 for "*In silico* Identification and Functional Characterization of Conserved miRNAs in Fibre Biogenesis Crop *Corchorus capsularis*"

**Supplementary File 1:** A total of 6386 of plant miRNA from miRBase database.

>aqc-miR477b MIMAT0012586

CUCUCCCUCAAGGGCUUCUA

>aqc-miR477e MIMAT0012584

CUCUCCCUCAAGGGCUUCUA

>aqc-miR156a MIMAT0012551

UGACAGAAGAUAGAGAGCAC

>aqc-miR167 MIMAT0012560

UCAAGCUGCCAGCAUGAUCUA

>aqc-miR530 MIMAT0012593

UGCAUUUGCACCUGCAUCUC

>aqc-miR398a MIMAT0012578

UGUGUUCUCAGGUCACCCCUU

>aqc-miR169b MIMAT0012564

UAGCCAAGGAUGACUUGCCUG

>aqc-miR160b MIMAT0012554

UGCCUGGCUCCCUGUAUGCCA

>aqc-miR482a MIMAT0012589

UCUUGCCGACUCCUCCCAUACC

>aqc-miR166d MIMAT0012559

UCGGACCAGGCUUCAUUCCUC

>aqc-miR156b MIMAT0012550

UGACAGAAGAUAGAGAGCAC

>aqc-miR169c MIMAT0012562

CAGCCAAGGAUGACUUGCCGG

>aqc-miR160a MIMAT0012553

UGCCUGGCUCCCUGGAUGCCA

>aqc-miR166c MIMAT0012558

UCGGACCAGGCUUCAUUCCU

>aqc-miR482b MIMAT0012590

UCUUGCCGACUCCUCCCAUACC

>aqc-miR477c MIMAT0012587

CUCUCCCUCAAGUUCUUCUA

>aqc-miR166b MIMAT0012557

UCGGACCAGGCUUCAUUCCCC

>aqc-miR168 MIMAT0012561

UGGCUUAGUGCAGCUCGGGGA

>aqc-miR529 MIMAT0012592

AGAAGAGAGAGAGCACAACCC

>aqc-miR171d MIMAT0012569

UGAUUGAGCCGUGCCAAUAUC

>aqc-miR477d MIMAT0012588

CUCUUCUUCAAAGGCUUCUA

>aqc-miR477g MIMAT0012582

CUCUCCCUCAAGUUCUUCUA

>aqc-miR171c MIMAT0012568

UAAUUGAACCGCACUAAUAUC

>aqc-miR172b MIMAT0012572

GGAAUCUUGAUGAUGCUGCAU

>aqc-miR171b MIMAT0012567

UGAUUGAGCCGUGCCAAUAUC

>aqc-miR159 MIMAT0012552

UUUGGACUGAAGGGAGCUCUA

>aqc-miR171e MIMAT0012570

UGAAUGAACCGAGCCAACAUC

>aqc-miR166e MIMAT0012555

UCGGACCAGGCUUCAUUCCCC

>aqc-miR395a MIMAT0012575

CUGAAGGGUUUGGAGGAACUC

>aqc-miR171f MIMAT0012565

UAAUUGAGCCGUGCCAAUAUC

>aqc-miR395b MIMAT0012574

CUGAAGGGUUUGGAGGAACUC

>aqc-miR169a MIMAT0012563

UAGCCAAGGAUGACUUGCCUA

>aqc-miR396b MIMAT0012577

UUCCACAGCUUUCUUGAACUU

>aqc-miR398b MIMAT0012579

UGUGUUCUCAGGUCGCCCCUG

>aqc-miR399 MIMAT0012580

UGCCAAAGGAGAGUUGCCCUA

>aqc-miR396a MIMAT0012576

UUCCACAGCUUUCUUGAACUG

>aqc-miR166a MIMAT0012556

UCGGACCAGGCUUCAUUCCUC

>aqc-miR171a MIMAT0012566

UGAUUGAGCCGUGCCAAUAUC

>aqc-miR172a MIMAT0012571

AGAAUCUUGAUGAUGCUGCAU

>aqc-miR477a MIMAT0012585

CUCUCCCUCAAGGGCUUCUA

>aqc-miR482c MIMAT0012591

UCUUGCCGACUCCUCCCAUACC

>aqc-miR319 MIMAT0012573

UUGGACUGAAGGGAGCUCCCU

>aqc-miR408 MIMAT0012581

UGCUCUGCCUCAUCCUUGUCU

>aqc-miR535 MIMAT0012594

UGACAACGAGAGAGAGCACGCG

>aqc-miR477f MIMAT0012583

CUCUCCUUCAAAGGCUUCUA

>ama-miR396-3p MIMAT0031155

AAGCUCAAGAAAGCUGUGGGA

>ama-miR396-5p MIMAT0031154

UUCCACAGCUUUCUUGAACUU

>ama-miR156 MIMAT0031153

CUGACAGAAGAGAGUGAGCAC

>bcy-miR529 MIMAT0020967

GAAGAAGAGAGAUGGUAGAG

>bcy-miR396a MIMAT0020965

UUCCACAGCUUUCUUGAACUG

>bcy-miR396b MIMAT0020966

UUCCACAGCUUUCUUGAACUU

>bcy-miR156 MIMAT0020964

UUUGACAGAAGAUAGAGAGCAC

>bgy-miR396b MIMAT0020962

UUCCACAGCUUUCUUGAACUU

>bgy-miR396a MIMAT0020961

UUCCACAGCUUUCUUGAACUG

>bgy-miR529 MIMAT0020963

GAAGAAGAGAGAUGGUAGAG

>bgy-miR156 MIMAT0020960

UUUGACAGAAGAUAGAGAGCAC

>ccl-miR167b MIMAT0014083

UGAAGCUGCCAGCAUGAUCUGA

>ccl-miR396 MIMAT0014078

UUCCACAGCUUUCUUGAACUU

>ccl-miR171 MIMAT0014075

UGAUUGAGCCGCGCCAAUAUC

>ccl-miR168 MIMAT0014073

UCGCUUGGUGCAGGUCGGGAA

>ccl-miR167a MIMAT0014072

UGAAGCUGCCAGCAUGAUCUGA

>crt-miR166a MIMAT0014084

UCGGACCAGGCUUCAUUCCCGU

>crt-miR171 MIMAT0014087

UGAUUGAGCCGUGCCAAUAUC

>crt-miR168 MIMAT0014086

UCGCUUGGUGCAGGUCGGGAA

>crt-miR166b MIMAT0014085

UCGGACCAGGCUUCAUUCCCUU

>csi-miR395b-3p MIMAT0048925

CUGAAGUGUUUGGGGGAACUC

>csi-miR3952-5p MIMAT0037415

GCUGUAGAUAGGCCCUUCAAC

>csi-miR2275a-3p MIMAT0048951

UUUAGUUUCCUCCAAUAUCUUA

>csi-miR399f-5p MIMAT0048935

GGGCAAUUCUCCUUUGGCAGA

>csi-miR171f-5p MIMAT0048910

UAUUGGCCUGGUUCACUCAGA

>csi-miR3954 MIMAT0018497

UGGACAGAGAAAUCACGGUCA

>csi-miR169c-5p MIMAT0048889

UAGCCAAGGAUGACUUGCCU

>csi-miR396c MIMAT0018493

UUCAAGAAAUCUGUGGGAAG

>csi-miR403a-5p MIMAT0037405

CAUGUUUCGGGUUUGUGCGUG

>csi-miR3947-5p MIMAT0018487

UUAUUUCAGUAGACGACGUCACA

>csi-miR399e-3p MIMAT0048934

UGCCAAAGGAGAUUUGCCCGG

>csi-miR166k-3p MIMAT0048883

UCUUGGACCAGGCUUCAUUCC

>csi-miR9560-3p MIMAT0048980

UCAUAUUUGCUCCACCACCUGUGG

>csi-miR156c-5p MIMAT0048852

UGACAGAAGAGAGUGAGCAC

>csi-miR390a-3p MIMAT0037374

CGCUAUCCAUCCUGAGUUUCA

>csi-miR2275b-3p MIMAT0048953

UUUAGUUUCCUCCAAUAUCUUA

>csi-miR399c-5p MIMAT0037401

GUGCAGUCCUCCUUUGGCGUG

>csi-miR482b-5p MIMAT0037411

AAUGGGAGGCUUGGCAAGAAG

>csi-miR156e-3p MIMAT0048856

GCUCUCUGUGCUUCUGUCAUCA

>csi-miR396f-5p MIMAT0048930

UUCCACAGCUUUCUUGAACUU

>csi-miR169l-5p MIMAT0048901

UAGCCAAGGAUGACUUGCCU

>csi-miR536-5p MIMAT0048967

CGCACCCCAGCGUGGAACCAUC

>csi-miR482g-5p MIMAT0048987

AGUGGGAGCGUGGGGUAAGAAG

>csi-miR396d-3p MIMAT0048927

UUCAAGAAAUCUGUGGGAAG

>csi-miR166d-5p MIMAT0048874

CUGUUGUCUGGUUCAAGG

>csi-miR171b-5p MIMAT0037396

UGUGAUAUUGGUUCGGCUCAUC

>csi-miR535a MIMAT0018484

UGACAAUGAGAGAGAGCACAC

>csi-miR399e-5p MIMAT0048933

GGGCAAUUCUCUAUUGGCAGU

>csi-miR156a-3p MIMAT0037391

UGCUCGCUCCUCUUCUGUCAGC

>csi-miR3952b-3p MIMAT0048962

UGAAGGGCCUUUCUAGAGCAC

>csi-miR477c-3p MIMAT0037408

GAAGUCCUUGGGGUUGAGUGA

>csi-miR159a-5p MIMAT0037392

GAGCUCCUUGAAGUCCAAAAG

>csi-miR169a MIMAT0014074

GAGCCAAGAAUGACUUGCCGA

>csi-miR408-3p MIMAT0018472

AUGCACUGCCUCUUCCCUGGC

>csi-miR169p-5p MIMAT0048996

UAGCCAAGGAUGACUUGCCU

>csi-miR164b-5p MIMAT0048868

UGGAGAAGCAGGGCACGUGCA

>csi-miR156i-5p MIMAT0048990

UUGACAGAAGAUAGAUAGCAC

>csi-miR535d-5p MIMAT0048998

UGACAAUGAGAGAGAGCACAC

>csi-miR3627b-3p MIMAT0048955

UUGUCGCAGGAGCGUUGGCACC

>csi-miR167c-3p MIMAT0037394

AGGUCAUCUUGCAGCUUCAAU

>csi-miR482e-5p MIMAT0048973

GGUCAUGGGAGGAUUGGCGA

>csi-miR827 MIMAT0018485

UUAGAUGACCAUCAACAAACA

>csi-miR12108-3p MIMAT0048976

AUUUUAAAACGUGCGGAUGUCUGC

>csi-miR162-3p MIMAT0014088

UCGAUAAACCUCUGCAUCCAG

>csi-miR167d-3p MIMAT0048885

AUCGGAUCAUGUGGUAGCUUCACC

>csi-miR535c-5p MIMAT0048947

UGACAAUGAGAGAGAGCACAC

>csi-miR12109-5p MIMAT0048977

UGCUUGGUGCAUAAAACUACAGAA

>csi-miR395c-3p MIMAT0048926

CUGAAGUGUUUGGGGGAACUC

>csi-miR395a MIMAT0018462

CUGAAGUGUUUGGGGGAACUC

>csi-miR172a-5p MIMAT0017387

GCAGCGUCCUCAAGAUUCACA

>csi-miR858-3p MIMAT0048949

CUCGUUGUCUGUUCGACCUUG

>csi-miR390a-5p MIMAT0014092

AAGCUCAGGAGGGAUAGCGCC

>csi-miR2111-5p MIMAT0049002

UAAUCUGCAUCCUGAGGUUUG

>csi-miR168-3p MIMAT0048986

CCCGCCUUGCAUCAACUGAAU

>csi-miR482a-3p MIMAT0018480

UCUUCCCUAUGCCUCCCAUUCC

>csi-miR171c-3p MIMAT0048907

UGAUUGAGCCGUGCCAAUAUC

>csi-miR394a MIMAT0018461

UUGGCAUUCUGUCCACCUCC

>csi-miR166h-3p MIMAT0048880

UUUGGACCAGGCUUCCUUCCUC

>csi-miR164b-3p MIMAT0048869

CACGCGCUCCCCUUCUCCAAC

>csi-miR3951a-3p MIMAT0037414

UUUCUCUUAUCGUUAUCUGU

>csi-miR393b-5p MIMAT0048919

UUCCAAAGGGAUCGCAUUGAUC

>csi-miR408-5p MIMAT0037406

ACGGGGAACAGGCAGAGCAUG

>csi-miR3952b-5p MIMAT0048961

GCUGUAGAAAGGCCCCUCAAC

>csi-miR169b-5p MIMAT0048888

UAGCCAAGGAUGACUUGCCU

>csi-miR482d-3p MIMAT0048944

UCCCUACUCCACCCAUGCCAUA

>csi-miR171e-3p MIMAT0048909

UUGAGCCGCGUCAAUAUCUCC

>csi-miR482c-3p MIMAT0018495

UUCCCUAGUCCCCCUAUUCCUA

>csi-miR396e-5p MIMAT0048928

UUCCACGGCUUUCUUGAACU

>csi-miR166g-3p MIMAT0048879

UCUCGGACCAGGCUUCAUUCC

>csi-miR169e-5p MIMAT0048893

UAGCCAAGGAUGACUUGCCU

>csi-miR156e-5p MIMAT0048855

GUGACAGAAGAUAGAGAGCGC

>csi-miR3950 MIMAT0018491

UUUUUCGGCAACAUGAUUUCU

>csi-miR167b-3p MIMAT0037395

AGAUCAUGCGGCAGUUUCACC

>csi-miR477a-5p MIMAT0018475

ACCUCCCUCGAAGGCUUCCAA

>csi-miR12107-3p MIMAT0048972

UCAUUCGCGCUCUCAUCAUUA

>csi-miR3948 MIMAT0018488

UGGAGUGGGAGUGGGAGUAGGGUG

>csi-miR393b-3p MIMAT0048920

UCAUGCGAUCCCUUCGGAAUU

>csi-miR171i-3p MIMAT0048914

UGAUUGAGCCGUGCCAAUAUC

>csi-miR160b-5p MIMAT0048864

UGCCUGGCUCCCUGUAUGCCG

>csi-miR3952-3p MIMAT0018494

UGAAGGGCCUUUCUAGAGCAC

>csi-miR159c-3p MIMAT0048863

CUUGGACUGAAGGGAGCUCCC

>csi-miR12110-5p MIMAT0048981

CUGGGGAGUUGCACCCGGAGU

>csi-miR172c-5p MIMAT0037397

GGAGCACCAUCAAGAUUCACA

>csi-miR166f-3p MIMAT0048877

UCGGACCAGGCUUCAUUCCCU

>csi-miR12111-5p MIMAT0048983

GAAAAACUUAGGUGGGAUAGUGUG

>csi-miR169e-3p MIMAT0048894

CAGGCAGUCUCCUUGGCUAAG

>csi-miR530a-3p MIMAT0037412

AGGUGCAGCUGUAUAUGCAGG

>csi-miR166c-5p MIMAT0037393

GGGAAUGUUGUCUGGUUCGAG

>csi-miR535f-5p MIMAT0049001

UGACAAUGAGAGAGAGCACAC

>csi-miR172b-5p MIMAT0048915

GCAGCAUCAUCAAGAUUCACA

>csi-miR395b-5p MIMAT0048924

GUUCCCCUGAACACUUCAUUG

>csi-miR156f-3p MIMAT0048858

GCUCUCUCUUCUCCUGUCAUUG

>csi-miR1515b-5p MIMAT0048959

UCAUUUUUGCGUGCAAUGAUCC

>csi-miR156g-3p MIMAT0048860

GCUCUCUAUUCUUCUGUCAUC

>csi-miR159c-5p MIMAT0048862

GAGCUUUCUUCGGUCCACUU

>csi-miR164d-3p MIMAT0048994

CACGCGCUCCCCUUCUCCAAC

>csi-miR3627a-5p MIMAT0048954

UUGUCGCCGGAGAGAUAGCACC

>csi-miR172c-3p MIMAT0018458

UGGAAUCUUGAUGAUGCUGCAG

>csi-miR2275b-5p MIMAT0048952

AGAAUUGGAUGGAACUAAACA

>csi-miR164a-3p MIMAT0037371

CAUGUGCCCUUCUUCCCCAUC

>csi-miR169j-5p MIMAT0048899

UAGCCAAGGAUGACUUGCCU

>csi-miR156h-5p MIMAT0048989

ACAGAAGAUAGAGAACACAUA

>csi-miR160b-3p MIMAT0048865

GCGUACGAGGAGCCAAGCAUA

>csi-miR164d-5p MIMAT0048993

UGGAGAAGCAGGGCACGUGCA

>csi-miR482f-3p MIMAT0018473

UUUUCCCACACCUCCCAUCCC

>csi-miR169d-5p MIMAT0048891

UAGCCAAGGAUGACUUGCCU

>csi-miR166b-5p MIMAT0048872

GGAAUGUUGUUUGGCUCGAGGG

>csi-miR399d-3p MIMAT0018468

UGCCAAAGGAGAGUUGCCCUG

>csi-miR156c-3p MIMAT0048853

GCUCACUCUCUAUCUGUCACC

>csi-miR857 MIMAT0018486

UUUUGAAUGUUGAAUGGUGGCUAU

>csi-miR167c-5p MIMAT0018453

UGAAGCUGCCAGCAUGAUCUG

>csi-miR160a-3p MIMAT0037370

GCGUAUGAGGAGCCAUGCAUA

>csi-miR535d-3p MIMAT0048999

GUGCUCUCUACCAUUGUCAUA

>csi-miR160c-3p MIMAT0048867

GCGUGCGAGGAGCCAUGCAUG

>csi-miR391-5p MIMAT0048957

UGCAGGUGAGAUGAUACCGUCA

>csi-miR403b-5p MIMAT0048937

AGUUUGUGCGUGAAUCUAACC

>csi-miR169n-3p MIMAT0048905

GGCAAGUUGCCCUUGGCUACA

>csi-miR169i-5p MIMAT0048898

UAGCCAAGGAUGACUUGCCU

>csi-miR171d-3p MIMAT0048908

UGAUUGAGCCGUGCCAAUAUC

>csi-miR171b-3p MIMAT0018455

CGAGCCGAAUCAAUAUCACUC

>csi-miR167e-3p MIMAT0048887

GGUCAUGCUCUGACAGCCUCACU

>csi-miR169n-5p MIMAT0048904

CAGCCAAGGAUGACUUGCCGG

>csi-miR477e-3p MIMAT0048942

UGAGGUUCUUGGGGAGAGUAG

>csi-miR393c-3p MIMAT0048922

AUCAUGCUAUCCCUUUGGAUU

>csi-miR172b-3p MIMAT0048916

AGAAUCUUGAUGAUGCUGCAU

>csi-miR156b-5p MIMAT0048850

UGACAGAAGAGAGUGAGCAC

>csi-miR403b-3p MIMAT0048938

UUAGAUUCACGCACAAACUCG

>csi-miR482c-5p MIMAT0037416

GGAAUUGGGUGCUAGGGAAGG

>csi-miR167b-5p MIMAT0018454

UGAAGCUGCCAGCAUGAUCUU

>csi-miR3951b-5p MIMAT0049003

CAGGUAAAGAUGAGAGAAAAA

>csi-miR156b-3p MIMAT0048851

GCUCACUGCUCUUUCUGUCAGCU

>csi-miR166i-3p MIMAT0048881

UCUUGGACCAGGCUUCAUUCC

>csi-miR166e-5p MIMAT0017386

GGAAUGUUGUCUGGCUCGAGG

>csi-miR171a MIMAT0018456

UUGAGCCGUGCCAAUAUCAC

>csi-miR166a-5p MIMAT0037373

GGAAUGUUGUCUGGCUCGAGG

>csi-miR535c-3p MIMAT0048948

GUGCUCUCUACCAUUGUCAUA

>csi-miR390b-5p MIMAT0048918

AGCUCAGGAGGGAUAGCGCC

>csi-miR164a-5p MIMAT0014069

UGGAGAAGCAGGGCACGUGCA

>csi-miR3954b-3p MIMAT0048964

ACCGUGUUUCUCUGCCCAAUC

>csi-miR399b-5p MIMAT0037404

AGGGCUUCUCUCCUUUGGCAG

>csi-miR156d-5p MIMAT0048854

UGACAGAAGAUAGAGAGCGC

>csi-miR398a-3p MIMAT0014079

UGUGUUCUCAGGUCACCCCUU

>csi-miR12110-3p MIMAT0048982

CCCGAGUCACAGCUCCCUGGAC

>csi-miR156a-5p MIMAT0018448

UGACAGAAGAGAGUGAGCAC

>csi-miR12108-5p MIMAT0048975

AGACGUUCGCACUUUUUAAAAUGC

>csi-miR3954b-5p MIMAT0048963

UUGGACAGAGAAAUCACGGUCA

>csi-miR169h-5p MIMAT0048897

UAGCCAAGGAUGACUUGCCU

>csi-miR399b-3p MIMAT0018470

UGCCAAAGGAGAGUUGCCCUA

>csi-miR477c-5p MIMAT0018474

ACUCUCCCUCAAGGGCUUCGC

>csi-miR477b-3p MIMAT0037410

UGAAGCCCUUAGGGCAGAGUGA

>csi-miR12111-3p MIMAT0048984

GACUGUCCCAUCUAAGUUUUUCUC

>csi-miR12109-3p MIMAT0048978

CUGCAGAUUUGUGCAUUAAGCUGC

>csi-miR3627c-5p MIMAT0048956

UCGCAGGGGAGAUGGGACCAAC

>csi-miR397-5p MIMAT0018465

UCAUUGAGUGCAGCGUUGAUG

>csi-miR168-5p MIMAT0048985

UCGCUUGGUGCAGGUCGGGAA

>csi-miR12106-5p MIMAT0048969

GUUUAGGACAGCUGCUGCCAAA

>csi-miR169q-5p MIMAT0048997

UAGCCAAGGAUGACUUGCCU

>csi-miR3951a-5p MIMAT0018492

UAGAUAAAGAUGAGAGAAAAA

>csi-miR12106-3p MIMAT0048970

UGGGGCAGCUGUCCUAAACGG

>csi-miR169r-5p MIMAT0049005

CAGCCAAGGAUGACUUGCCGG

>csi-miR2275a-5p MIMAT0048950

AGAAUUGGAUGGAACUAAACA

>csi-miR393c-5p MIMAT0048921

UCCAAAGGGAUCGCAUUGAUC

>csi-miR482e-3p MIMAT0048974

UUGCCAACUCCUCCCAUGCCGA

>csi-miR160c-5p MIMAT0048866

UGCCUGGCUCCCUGUAUGCUU

>csi-miR167a-5p MIMAT0018498

UGAAGCUGCCAGCAUGAUCUG

>csi-miR399c-3p MIMAT0018467

UGCCAAAGGAGAAUUGCCCUG

>csi-miR530b-5p MIMAT0018483

UGCAUUUGCACCUGCAUCUUG

>csi-miR398a-5p MIMAT0037372

AGAACAGAGGGUGGCGUUGGCU

>csi-miR482a-5p MIMAT0018479

AGUGGGAGCGUGGGGUAAGAAG

>csi-miR530a-5p MIMAT0018482

UGCAUUUGCACCUGCACCUUG

>csi-miR397-3p MIMAT0037400

ACCAGCGCUGCACUCGAUCAU

>csi-miR166a-3p MIMAT0014090

UCGGACCAGGCUUCAUUCCCCC

>csi-miR393a MIMAT0018460

AUCCAAAGGGAUCGCAUUGAUC

>csi-miR159d MIMAT0018459

UUUGGACUGAAGGGAGCUCCU

>csi-miR3951b-3p MIMAT0049004

UUUCUCUUAUCUUUUAUCUGUG

>csi-miR536-3p MIMAT0048968

UGGUGCCACGCUGUGUGCGUC

>csi-miR399d-5p MIMAT0037402

GGGCACCUCUUACUUGGCAUG

>csi-miR171g-3p MIMAT0048912

UUGAGCCGCGUCAAUAUCUCC

>csi-miR171f-3p MIMAT0048911

UGAUUGAGCCGUGCCAAUAUC

>csi-miR398b-3p MIMAT0048932

UGUGUUCUCAGGUCGCCCCUG

>csi-miR477b-5p MIMAT0018476

CUCUCCCUCAAGGGCUUCUCU

>csi-miR396a-3p MIMAT0037398

AUUCAAGAAAGCUGUGGAAAA

>csi-miR162-5p MIMAT0017388

UGGAGGCAGCGGUUCAUCGAUC

>csi-miR12105-5p MIMAT0048965

UUUCUUGAGACAAAACAUGCAUC

>csi-miR156j-3p MIMAT0048992

AUCCAACGAAGCAGGAGCUGC

>csi-miR399f-3p MIMAT0048936

CGCCAAAGGAGAAUUGCCCUG

>csi-miR396b-3p MIMAT0037399

GUUCAAUAAAGCUGUGGGAAG

>csi-miR396a-5p MIMAT0018463

UUCCACAGCUUUCUUGAACUG

>csi-miR164c-5p MIMAT0048870

UGGAGAAGCAGGGCACGUGCA

>csi-miR169l-3p MIMAT0048902

AGGCAGUCUCCUUGGCUAAC

>csi-miR399a-3p MIMAT0018469

UGCCAAAGGAGAUUUGCCCGG

>csi-miR477e-5p MIMAT0048941

ACUCUCCCUCAAGGGCUUCUGA

>csi-miR394b-5p MIMAT0048923

UUGGCAUUCUGUCCACCUCC

>csi-miR172d-5p MIMAT0048917

GCGGCAUCAUCAAGAUUCACA

>csi-miR482g-3p MIMAT0048988

UCUUCCCUAUGCCUCCCAUUCC

>csi-miR9560-5p MIMAT0048979

ACAGGAGGUGGAACAAAUAUGAAA

>csi-miR477d-3p MIMAT0048940

UGAGGCCGUUGGGGAGAGUGG

>csi-miR166e-3p MIMAT0014070

UCGGACCAGGCUUCAUUCCCC

>csi-miR477d-5p MIMAT0048939

ACUCUCCCUCAAGGGCUUCUGG

>csi-miR396f-3p MIMAT0048931

GCUCAAGAAAGCUGUGGGAGA

>csi-miR169g-5p MIMAT0048896

CAGCCAAGGAUGACUUGCCGG

>csi-miR169f-5p MIMAT0048895

UAGCCAAGGAUGACUUGCCU

>csi-miR169k-5p MIMAT0048900

CAGCCAAGGAUGACUUGCCGG

>csi-miR530b-3p MIMAT0037413

AGGUGCAGUUGCAAGUGCAGA

>csi-miR169o-5p MIMAT0048995

CAGCCAAGGAUGACUUGCCGG

>csi-miR535b-5p MIMAT0048945

UGACAAUGAGAGAGAGCACAC

>csi-miR169c-3p MIMAT0048890

CAGGCAGUCUCCUUGGCUAAG

>csi-miR477a-3p MIMAT0037409

GGAAACCCUAGGGGGAGGUCG

>csi-miR535e-5p MIMAT0049000

UGACAAUGAGAGAGAGCACAC

>csi-miR403a-3p MIMAT0018471

UUAGAUUCACGCACAAACUCG

>csi-miR1515b-3p MIMAT0048960

AUCAUUCACGCAAAAAUGAUU

>csi-miR159b-5p MIMAT0048861

AGCUGCCGACUCAUUCAUUCA

>csi-miR535b-3p MIMAT0048946

GUGCUCUCUACCAUUGUCAUA

>csi-miR399a-5p MIMAT0037403

GGGCAACAUCUCCAUUGGCAGG

>csi-miR171h-3p MIMAT0048913

UGAUUGAGCCGUGCCAAUAUC

>csi-miR166f-5p MIMAT0048876

GGACUGUUGUCUGGCUCGAUG

>csi-miR167d-5p MIMAT0048884

UGAAGCUGCCAGCAUGAUCUA

>csi-miR166d-3p MIMAT0048875

UCUCGGACCAGGCUUCAUUCC

>csi-miR12107-5p MIMAT0048971

CUGAUGAGAGAGCGAAUGAUA

>csi-miR482b-3p MIMAT0018481

UCUUGCCCACCCCUCCCAUUCC

>csi-miR156f-5p MIMAT0048857

AUGACAGAAGAGAGAGAGUAC

>csi-miR166g-5p MIMAT0048878

GAAUGCUGUCUGGUUCGAGAC

>csi-miR1515a MIMAT0018489

UCAUUUUUGCGUGCAAUGAUCC

>csi-miR482d-5p MIMAT0048943

UGGUAUGGGUGAGUAGGGAAG

>csi-miR3949 MIMAT0018490

UGAUGUUGAGGCAAAAAUGUAG

>csi-miR164c-3p MIMAT0048871

CACGCGCUCCCCUUCUCCAAC

>csi-miR160a-5p MIMAT0014068

GCCUGGCUCCCUGUAUGCCAU

>csi-miR167a-3p MIMAT0037417

AGAUCAUCUGGCAGUUUCACC

>csi-miR12105-3p MIMAT0048966

UUGUUUUGGGUGAAACGGGUGU

>csi-miR396e-3p MIMAT0048929

GCUCAAGAAUGCCGUGGGAAA

>csi-miR171c-5p MIMAT0048906

UGUUGGAACGGCUCAAUCAAA

>csi-miR396b-5p MIMAT0018464

UUCCACAGCUUUCUUGAACUG

>csi-miR159a-3p MIMAT0018450

UUUGGAUUGAAGGGAGCUCUA

>csi-miR156g-5p MIMAT0048859

UUGACGGAAGAUAGAGAGCAC

>csi-miR156j-5p MIMAT0048991

AGCUGCUGACUCGUUGGUUCA

>csi-miR172a-3p MIMAT0014076

AGAAUCUUGAUGAUGCUGCA

>csi-miR167e-5p MIMAT0048886

UGAAGCUGCCAGCAUGAUCUA

>csi-miR169d-3p MIMAT0048892

CAGGCAGUCUCCUUGGCUAAG

>csi-miR166c-3p MIMAT0018451

UCGGACCAGGCUUCAUUCCC

>csi-miR169m-5p MIMAT0048903

CAGCCAAGGAUGACUUGCCGG

>csi-miR166j-3p MIMAT0048882

UUGGACCAGGCUUCAUUCCUC

>csi-miR391-3p MIMAT0048958

CCGGAAUCAUUUCUCCCGCGUG

>csi-miR3946 MIMAT0018449

UUGUAGAGAAAGAGAAGAGAGCAC

>csi-miR166b-3p MIMAT0048873

UCUCGGACCAGGCUUCAUUCC

>csi-miR482f-5p MIMAT0037407

GAUGGGUGAGUUGGGAAAAUA

>csi-miR3953 MIMAT0018496

UUGAGUUCUGCAAGCCGUCGA

>ctr-miR164 MIMAT0014080

UGGAGAAGCAGGGCACGUGCA

>ctr-miR156 MIMAT0014067

UGACAGAAGAGAGUGAGCAC

>ctr-miR171 MIMAT0014082

UUGAGCCGCGUCAAUAUCUCC

>ctr-miR167 MIMAT0014081

UGAAGCUGCCAGCAUGAUCUGA

>ctr-miR319 MIMAT0014077

UUGGACUGAAGGGAGCUCCC

>ctr-miR166 MIMAT0014071

UCGGACCAGGCUUCAUUCCCCC

>dpr-miR172b MIMAT0023535

AGAAUCUUGAUGAUGCUGCAU

>dpr-miR397 MIMAT0023537

CCAUUGAGUGCAGCGUUGAUG

>dpr-miR408 MIMAT0023538

CUGCACUGCCUCUUCCCUGGC

>dpr-miR156a MIMAT0023526

UGACAGAAGAGAGGGAGCAC

>dpr-miR396 MIMAT0023536

UUCCACAGCUUUCUUGAACUG

>dpr-miR172a MIMAT0023534

AGAAUCUUGAUGAUGCUGCAU

>dpr-miR166b MIMAT0023530

UCGGACCAGGCUUCAUUCCCC

>dpr-miR160 MIMAT0023528

UGCCUGGCUCCUUGUAUGCCA

>dpr-miR166a MIMAT0023529

UCGGACCAGGCUUCAUUCCUC

>dpr-miR167a MIMAT0023531

UGAAGCUGCCAGCAUGAUCUA

>dpr-miR167b MIMAT0023532

UGAAGCUGCCAGCAUGAUCUA

>dpr-miR156b MIMAT0023527

UGACAGAAGAGAGUGAGCAC

>dpr-miR167c MIMAT0023533

UGAAGCUGCCAGCAUGAUCUG

>eun-miR482a-5p MIMAT0041082

CAUGGGUUGUUUGGUGAGAGG

>eun-miR530-5p MIMAT0041088

UCUGCAUUUGCACCUGCACCU

>eun-miR10214a-3p MIMAT0041105

UAGAUAUAGUGGAUUUUCGAU

>eun-miR397a-5p MIMAT0041078

UCAUUGAGUGCAGCGUUGAU

>eun-miR397a-3p MIMAT0041079

CGGUUUCGACAGCGCUGCACU

>eun-miR10220-3p MIMAT0041117

UCGUGAAGGAAGAAUGUGCAAU

>eun-miR535b-5p MIMAT0041092

UGACAACGAGAGAGAGCACGC

>eun-miR395-5p MIMAT0041072

UCCCCUAGAGUUCUCCUGAACA

>eun-miR159-3p MIMAT0041052

CUUGCAUAUGCCAGGAGCUUC

>eun-miR10214b-3p MIMAT0041106

CAGAGAUAUAGUGGAUUUACG

>eun-miR530-3p MIMAT0041089

AGGUGCGGGUGCAGGUGCAGA

>eun-miR167a-5p MIMAT0041059

UGAAGCUGCCAGCAUGAUCUGA

>eun-miR10216-5p MIMAT0041108

UCUCUGUUGAUCUGAUAAAUA

>eun-miR396b-3p MIMAT0041077

GUUCAAGCUAGCUGUGGGAAG

>eun-miR167b-3p MIMAT0041062

UCAGGUCAUCUUGCAGCUUCA

>eun-miR10217-3p MIMAT0041111

UAAGUGGUGAUCUGACUCUAA

>eun-miR397b-3p MIMAT0041081

UCAACGCUGCACUCAAUGAUG

>eun-miR167d-5p MIMAT0041065

UGAAGCUGCCACAUGAUCUGA

>eun-miR535a-3p MIMAT0041091

GUGCUCUCUAUCGCUGUCAUA

>eun-miR10219-5p MIMAT0041114

UUCAAGUCUAACAACCUCAGCU

>eun-miR167c-3p MIMAT0041064

AUCAGAUCAUGUGGCAGCUUCACC

>eun-miR10218-3p MIMAT0041113

AUCUGUGGAAGAGACUCGACU

>eun-miR10222-3p MIMAT0041121

UUCAUGGACAGCCAGCUUAUU

>eun-miR10223-3p MIMAT0041123

UCUCGUUCCGCUUCAUCUGAA

>eun-miR10211a-3p MIMAT0041097

AUCCAGAAAUUGGCAGCCGUU

>eun-miR10213-3p MIMAT0041103

UGGAUCAAUAGAACGAGCAGGUGA

>eun-miR166-5p MIMAT0041057

GGAAUGUUGUCUGGCUCGAGG

>eun-miR482a-3p MIMAT0041083

UCUUGCCAAUACCACCCAUGCC

>eun-miR172a-3p MIMAT0041067

AGAAUCUUGAUGAUGCUGCAU

>eun-miR395-3p MIMAT0041073

AUGAAGUGUUUGGGGGAACUC

>eun-miR482b-3p MIMAT0041085

UUUCCUAUUCCUCCCAUUCCAU

>eun-miR10215-3p MIMAT0041107

GAGAAUGAUGAGUUAAAUGGA

>eun-miR396a-3p MIMAT0041075

GUUCAAUAAAGCUGUGGGAAG

>eun-miR10221-5p MIMAT0041118

GCUCGAGGUCAGUUUGUCGCC

>eun-miR167c-5p MIMAT0041063

UGAAGCUGCCAGCGUGAUCUCA

>eun-miR10223-5p MIMAT0041122

UGAAGCAGAUCAAGAACCCAG

>eun-miR482c-3p MIMAT0041087

UUCCCAAGGCCGCCCAUUCCGA

>eun-miR10213-5p MIMAT0041102

UACUCGUUCCGUUGAUCCAUC

>eun-miR172c-5p MIMAT0041070

GUAGCAUCAUCAAGAUUCACA

>eun-miR10214a-5p MIMAT0041104

UCGUAAAUCCACUAUAUCUCU

>eun-miR397b-5p MIMAT0041080

UGCAGCGCUGUCGAAACCGAU

>eun-miR10211b-5p MIMAT0041098

CAAUUUCUGGAUUUCAGUUCG

>eun-miR10212-3p MIMAT0041101

CGAUCUAAUCAAUCAUUUUUCGGG

>eun-miR10220-5p MIMAT0041116

UGCUGUUCUUCCGUUCACGAAU

>eun-miR10218-5p MIMAT0041112

UCGAGCCCCUCCCACAGAUUG

>eun-miR10212-5p MIMAT0041100

CGAAAAAUGAUUGGUUGUAUCGCU

>eun-miR482b-5p MIMAT0041084

GAAAUGGGAGGGUGGGAAAGA

>eun-miR10216-3p MIMAT0041109

UUUGUCGGAUGAACAGGGAAU

>eun-miR162-3p MIMAT0041056

UCGAUAAACCUCUGCAUCCAG

>eun-miR167b-5p MIMAT0041061

UGAAGCUGCCAGCAUGAUCUGG

>eun-miR396a-5p MIMAT0041074

UUCCACGGCUUUCUUGAACUG

>eun-miR827-5p MIMAT0041094

CUUUGUUGAUGGCCAUCUAAUC

>eun-miR482c-5p MIMAT0041086

GAGAUUCGAGCUACCGGAAGUUGUG

>eun-miR166-3p MIMAT0041058

UCGGACCAGGCUUCAUUCCCC

>eun-miR10222-5p MIMAT0041120

CAUGUAACAAGUUGGCUGUCA

>eun-miR10224-3p MIMAT0041124

CAAUGAACGCAUUUGCAGGUG

>eun-miR535a-5p MIMAT0041090

UGACAACGAGAGAGAGCACGC

>eun-miR535b-3p MIMAT0041093

UGCUCUCUACCGUUGUCAUG

>eun-miR159-5p MIMAT0041051

AGCUGCUGGUCUAUGGAUCCC

>eun-miR160-5p MIMAT0041053

UGCCUGGCUCCCUGUAUGCCA

>eun-miR10211a-5p MIMAT0041096

UCGGCUGUCAAUUUCUGGAUU

>eun-miR162-5p MIMAT0041055

GGAGGCAGCGGUUCAUCGAUC

>eun-miR10217-5p MIMAT0041110

UAGGGUCAGAUCGCUACUUAG

>eun-miR396b-5p MIMAT0041076

UUCCACAGCUUUCUUGAACUG

>eun-miR172b-3p MIMAT0041069

AGAAUCUUGAUGAUGCUGCAU

>eun-miR827-3p MIMAT0041095

UUAGAUGACCAUCAGCGAACA

>eun-miR172b-5p MIMAT0041068

GCAGCAUCAUCAAGAUUCACA

>eun-miR10221-3p MIMAT0041119

CGGCAAACUGGACCUCGAGAUC

>eun-miR172c-3p MIMAT0041071

AGAAUCUUGAUGAUGCUGCAU

>eun-miR167a-3p MIMAT0041060

AGAUCAUCUGGCAGUUUCAAC

>eun-miR10219-3p MIMAT0041115

UGGAGGUUGUUUGGCUUGAGCU

>eun-miR160-3p MIMAT0041054

GCGUAUGAGGAGCCAAGCAUA

>eun-miR10211b-3p MIMAT0041099

UCGAACUGAAAUCCAGAAAUU

>eun-miR172a-5p MIMAT0041066

CAGGUGUAGCAUCAUCAAGAU

>fve-miR159a-3p MIMAT0044488

UUUGGAUUGAAGGGAGCUCUA

>fve-miR11287 MIMAT0044568

UCAGGGAUUGUUUCAUAGACC

>fve-miR11299 MIMAT0044582

CAAAUAGGGUUGGCUGAUACU

>fve-miR11314 MIMAT0044598

AGAGUUGUGGAUGCUAUGAAU

>fve-miR171e MIMAT0044520

UGAUUGAGCCGUGCCAAUAUC

>fve-miR11289 MIMAT0044571

UGCUUCAAGUCUGGCCAAUACU

>fve-miR482a MIMAT0044557

UCUUUCCAAUUCCUCCCAUGCC

>fve-miR169b MIMAT0044513

UAGCCAAGGAUGACUUGCCU

>fve-miR11284 MIMAT0044473

UUCGUGAUCUGCGAAAGGCUC

>fve-miR11297 MIMAT0044580

CAGACAAGAUCGAUCUCGCCU

>fve-miR11300 MIMAT0044583

CAACAUCACUGUUCUCUUCCU

>fve-miR166a MIMAT0044500

UCGGACCAGGCUUCAUUCCCC

>fve-miR477a MIMAT0044556

ACUCUCCCUCAAGGGCUUCUC

>fve-miR167c MIMAT0044508

UGAAGCUGCCAGCAUGAUCU

>fve-miR408 MIMAT0044555

UGCACUGCCUCUUCCCUGGCU

>fve-miR396c-3p MIMAT0044549

GUUCAAUAAAGCUGUGGGAAG

>fve-miR535a MIMAT0044563

UGACGAUGAGAGAGAGCACGC

>fve-miR160a MIMAT0044492

UGCCUGGCUCCCUGUAUGCCA

>fve-miR845 MIMAT0044566

AACCGGCUCUGAUACCAAUUG

>fve-miR171h MIMAT0044524

UUGAGCCGCGUCAAUAUCUCC

>fve-miR393a MIMAT0044542

UCCAAAGGGAUCGCAUUGAUC

>fve-miR399a MIMAT0044553

UGCCAAAGGAGAGUUGCCCUG

>fve-miR482b MIMAT0044558

UCUUUCCUAGUCCUGCCAUUCC

>fve-miR11294 MIMAT0044577

UUCACCUGGACCAUAACUGACC

>fve-miR156c MIMAT0044478

UUGACAGAAGAGAGUGAGCAC

>fve-miR11291 MIMAT0044574

UUGCGGUCUUGUCUCUUCCAAU

>fve-miR156g-5p MIMAT0044482

UUGACAGAAGAUAGAGAGCAC

>fve-miR394 MIMAT0044543

UUGGCAUUCUGUCCACCUCC

>fve-miR390b MIMAT0044540

AAGCUCAGGAGGGAUAGCGCC

>fve-miR164a-5p MIMAT0044495

UGGAGAAGCAGGGCACGUGCA

>fve-miR166c MIMAT0044504

UCGGACCAGGCUUCAUUCCCC

>fve-miR827 MIMAT0044565

UUAGAUGACCAUCAACAAACA

>fve-miR11295 MIMAT0044578

CUCAUUCAAUUUCGGUAUUCAG

>fve-miR11305 MIMAT0044588

UUUUGGUCCGAAUCCGAGCUCC

>fve-miR159c MIMAT0044490

AUUGGAUUGAAGGGAGCUCCC

>fve-miR172b MIMAT0044528

AGAAUCUUGAUGAUGCUGCAU

>fve-miR11301 MIMAT0044584

UCAGAGUUGUAAUAUAUUGAU

>fve-miR164b MIMAT0044497

UGGAGAAGCAGGGCACGUGCA

>fve-miR11286 MIMAT0044567

UUGGAGAGAGAGUAGACAAUG

>fve-miR2109 MIMAT0044531

UGCGAGUGUCUUCACCUCUGAA

>fve-miR166e MIMAT0044499

UCGGACCAGGCUUCAUUCCCC

>fve-miR11309 MIMAT0044592

UUUGUUUGGCAUGCAGUUGGC

>fve-miR156j MIMAT0044486

UUGACGGAAGAGAGCGAGCAC

>fve-miR390a MIMAT0044539

AAGCUCAGGAGGGAUAGCGCC

>fve-miR172c MIMAT0044529

AGAAUCUUGAUGAUGCUGCAU

>fve-miR2111c MIMAT0044532

UAAUCUGCAUCCUGAGGUUU

>fve-miR166d-5p MIMAT0044502

GGGAAUGUCGUCUGGUUCGA

>fve-miR11312 MIMAT0044595

UGGAGCUGUUGGGGAGAGUUA

>fve-miR11288e MIMAT0044601

UGAAGUGGGAUUUGGCGAAUU

>fve-miR162-3p MIMAT0044494

UCGAUAAACCUCUGCAUCCAG

>fve-miR164a-3p MIMAT0044496

CACGUGCUCCCCUUCUCCAAC

>fve-miR396c-5p MIMAT0044548

UUCCACAGCUUUCUUGAACUG

>fve-miR396b-3p MIMAT0044545

GCUCAAGAAAGCUGUGGGACA

>fve-miR11304 MIMAT0044587

UAGUGUUUCAGCUUGACAAG

>fve-miR168-3p MIMAT0044511

CCCGCCUUGCAUCAACUGAAU

>fve-miR156i MIMAT0044485

UGACAGAAGAUAGAGAGCAC

>fve-miR535b MIMAT0044564

UGACGAUGAGAGAGAGCACGC

>fve-miR166b MIMAT0044501

UCGGACCAGGCUUCAUUCCCC

>fve-miR2111a MIMAT0044533

UAAUCUGCAUCCUGAGGUUU

>fve-miR319 MIMAT0044536

UUUGGACUGAAGGGAGCUCCU

>fve-miR156a MIMAT0044477

UUGACAGAAGAGAGUGAGCAC

>fve-miR168-5p MIMAT0044510

UCGCUUGGUGCAGGUCGGGAA

>fve-miR11308 MIMAT0044591

UAAGUUAGGAUUCUAGUUACC

>fve-miR171b MIMAT0044525

CGAGCCGAACCAAUAUCACUC

>fve-miR11288a MIMAT0044573

UGGGAUUUGGCGAAUUGUGGU

>fve-miR11288c-5p MIMAT0044569

UUCGUUCGGAUUCCAAUUCAAA

>fve-miR11310 MIMAT0044593

AGGGAUCACAACUCCAGUGCU

>fve-miR482c MIMAT0044559

UCUUUCCUAUUCCUCCCAUCCC

>fve-miR167b MIMAT0044507

UGAAGCUGCCAGCAUGAUCU

>fve-miR171g MIMAT0044521

UGAUUGAGCCGUGCCAAUAUC

>fve-miR166d-3p MIMAT0044503

UCGGACCAGGCUUCAUUCCCC

>fve-miR159a-5p MIMAT0044487

GAGCUCCUUGAAGUCCAAUAG

>fve-miR171f-5p MIMAT0044526

CGAUGUUGGUGAGGUUCAAUC

>fve-miR11288c-3p MIMAT0044570

UGAAUUGGGAUUUGUCGAAUU

>fve-miR156b MIMAT0044476

UUGACAGAAGAGAGUGAGCAC

>fve-miR3627b MIMAT0044538

UCGCAGGAGAGAUGGCACUACC

>fve-miR156e MIMAT0044479

UUGACAGAAGAGAGUGAGCAC

>fve-miR171c-3p MIMAT0044523

UGAUUGAGCCGUGCCAAUAUC

>fve-miR11292 MIMAT0044575

UUGUAGUUCAGCGCCUCCGCC

>fve-miR11298 MIMAT0044581

UUGAGGGGCUUAACGAUUACC

>fve-miR169f MIMAT0044517

UGAGCCAAGAAUGACUUGCUG

>fve-miR396d MIMAT0044550

UUCCACAGCUUUCUUGAACUG

>fve-miR11293 MIMAT0044576

UUCUUCCUCAGGAACCUCCACC

>fve-miR160b MIMAT0044491

UGCCUGGCUCCCUGUAUGCCA

>fve-miR164c MIMAT0044498

UGGAGAAGCAGGGCACAUGCU

>fve-miR11290 MIMAT0044572

UCGUCGACAUCAAAGGGCACC

>fve-miR156d MIMAT0044480

UGACAGAAGAGAGUGAGCAC

>fve-miR172a MIMAT0044530

AGAAUCUUGAUGAUGCUGCAU

>fve-miR11302 MIMAT0044585

AGGACCGCCAUCACGUUUUGG

>fve-miR482d MIMAT0044560

UUCCCUAUUCCACCUAUUCCCC

>fve-miR11285 MIMAT0044474

UCUAUUCAAAGAGAUGACUGUU

>fve-miR397 MIMAT0044552

UCAUUGAGUGCAGCGUUGAUG

>fve-miR11313 MIMAT0044597

GAAAAGAAUGGACUCUCCGGGG

>fve-miR169c MIMAT0044514

UAGCCAAGGAUGACUUGCCU

>fve-miR11311 MIMAT0044594

UGAGAAUGAUGUGGAUCUCAGC

>fve-miR171f-3p MIMAT0044527

UUGAGCCGCGCCAAUAUCACU

>fve-miR171d MIMAT0044519

UGAUUGAGCCGUGCCAAUAUC

>fve-miR11315 MIMAT0044602

CUAGUCAUUGGUCAUAGCAUC

>fve-miR399b MIMAT0044554

AGCCAAAGGAGAAUUGCCCUG

>fve-miR393b MIMAT0044541

UCCAAAGGGAUCGCAUUGAUCU

>fve-miR2111b-5p MIMAT0044534

UAAUCUGCAUCCUGAGGUUU

>fve-miR396a-3p MIMAT0044547

GUUCAAUAAAGCUGUGGGAAG

>fve-miR11296 MIMAT0044579

UUUUUGAUGGCUGGAAUCCAGU

>fve-miR169a MIMAT0044512

UAGCCAAGGAUGACUUGCCU

>fve-miR396a-5p MIMAT0044546

UUCCACAGCUUUCUUGAACUG

>fve-miR162-5p MIMAT0044493

GGAGGCAGCGGUUCAUCGAUC

>fve-miR11306 MIMAT0044589

CUACCGAAGAACUUUGCAAAAG

>fve-miR11303 MIMAT0044586

UCAAACAUCACUGCAGCUGUA

>fve-miR169d MIMAT0044515

UAGCCAAGGAUGACUUGCCU

>fve-miR156h MIMAT0044484

UGACAGAAGAGAGUGAGCUC

>fve-miR3627a MIMAT0044537

UCGCAGGAGAGAUGGCACUACC

>fve-miR169e MIMAT0044516

UGAGCCAAGGAUGACUUGCCU

>fve-miR11288b MIMAT0044599

UGGGAUUGGGCGAAUUUUGGU

>fve-miR166f MIMAT0044505

UCGGACCAGGCUUCAUUCCCC

>fve-miR171c-5p MIMAT0044522

AGAUAUUGGUGCGGUUCAAUC

>fve-miR167d MIMAT0044506

UGAAGCUGCCAGCAUGAUCUCA

>fve-miR1511 MIMAT0044475

ACCUAGCUCUGAUACCAUGUG

>fve-miR156f MIMAT0044481

UUGACAGAAGAUAGAGAGCAC

>fve-miR530 MIMAT0044562

UGCAUUUGCACCUGCACCUCU

>fve-miR156g-3p MIMAT0044483

GCUCUCUAUGCUUCUGUCAUC

>fve-miR11283 MIMAT0044472

AGGCUUUGUAGAGGAUGGAAU

>fve-miR11307 MIMAT0044590

UAAGCGACGGACUCCAAUCGC

>fve-miR5225 MIMAT0044561

CUGUCGUAGGAGAGAUGGCGCC

>fve-miR477b MIMAT0044596

CGCGCACCCGUUCAUCUUCGC

>fve-miR396b-5p MIMAT0044544

UUCCACAGCUUUCUUGAACUU

>fve-miR2111b-3p MIMAT0044535

GCCCUUGGGAUGCGGAUUACC

>fve-miR171a MIMAT0044518

UGAUUGAGCCGUGCCAAUAUC

>fve-miR11288d MIMAT0044600

UCCAUCGUUUUGAGACACAGG

>fve-miR396e MIMAT0044551

UUCCACAGGCUUUCUUGAACU

>fve-miR167a MIMAT0044509

UGAAGCUGCCAGCAUGAUCU

>fve-miR159b MIMAT0044489

AUUGGAUUGAAGGGAGCUCUC

>gar-miR2947 MIMAT0014330

UAUACCGUGCCCAUGACUGUAG

>ghb-miR169a MIMAT0005814

UAGCCAAGGAUGACUUGCCUG

>ghr-miR7492b MIMAT0029135

CUAUAGAACAUGAUCUUUAGCGG

>ghr-miR7507 MIMAT0029154

AAGGUAGUGAAGUAGGCAAUUGGG

>ghr-miR399e MIMAT0025840

UGCCAAAGGAGAUUUGCCCCG

>ghr-miR482b MIMAT0014340

UCUUGCCUACUCCACCCAUGCC

>ghr-miR7508 MIMAT0029155

CAAGAAAAGAAGUCGGGAGAG

>ghr-miR7498 MIMAT0029145

AUGGUGACACAUGGUAGUCUCACA

>ghr-miR7497 MIMAT0029144

ACAUGUGGACUGUCAUAUGGGUU

>ghr-miR827b MIMAT0014342

UUAGAUGACCAUCAACAAACA

>ghr-miR399d MIMAT0014350

UGCCAAAGGAGAUUUGCCCUG

>ghr-miR156d MIMAT0005809

UGACAGAAGAGAGUGAGCAC

>ghr-miR2950 MIMAT0014348

UGGUGUGCAGGGGGUGGAAUA

>ghr-miR399a MIMAT0005820

CGCCAAUGGAGAUUUGUCCGG

>ghr-miR7490 MIMAT0029132

AGUCUAGAAAACUUCACUGACGGU

>ghr-miR3476-3p MIMAT0015654

AGCCAACAACAUCAGUUCUAA

>ghr-miR2949a-5p MIMAT0014345

ACUUUUGAACUGGAUUUGCCGA

>ghr-miR167b MIMAT0025838

UGAAGCUGCCAGCAUGAUCUA

>ghr-miR7495b MIMAT0029141

UUACUUUAGAUGUCUCCUUCA

>ghr-miR393 MIMAT0014334

UCCAAAGGGAUCGCAUUGAUCU

>ghr-miR7494 MIMAT0029139

AGCUUGUGGACUAGUUUUAACAA

>ghr-miR3476-5p MIMAT0015653

UGAACUGGGUUUGUUGGCUGC

>ghr-miR169b MIMAT0029157

CAGCCAAGGAUGAUUUGCCGG

>ghr-miR827c MIMAT0014343

UUAGAUGACCAUCAACAAACA

>ghr-miR172 MIMAT0014333

AGAAUCCUGAUGAUGCUGCAG

>ghr-miR7504a MIMAT0029151

UAUGAAACUGUGAUUCCACGUCAU

>ghr-miR827a MIMAT0014341

UUAGAUGACCAUCAACAAACA

>ghr-miR7504b MIMAT0029159

AGGAGGAAAAAUCUGAUUUGUCAU

>ghr-miR7505 MIMAT0029152

UUCAGAAACCAUCCCUUCCUU

>ghr-miR7510b MIMAT0029163

AAGAACAUGAUCUUUAGCGGCGU

>ghr-miR396b MIMAT0005819

UUCCACAGCUUUCUUGAACUG

>ghr-miR156b MIMAT0005807

UGACAGAAGAGAGUGAGCAC

>ghr-miR7513 MIMAT0029162

AAUCAGCCAGGAAUCGUUUGA

>ghr-miR7484a MIMAT0029124

UUUGUAUAUUAGAUCAAAGAGCAA

>ghr-miR169a MIMAT0025839

UAGCCAAGGAUGACUUGCCUG

>ghr-miR396a MIMAT0005818

UUCCACAGCUUUCUUGAACUG

>ghr-miR162a MIMAT0005812

UCGAUAAACCUCUGCAUCCAG

>ghr-miR479 MIMAT0014338

CGUGAUAUUGGUUCGGCUCAUC

>ghr-miR167a MIMAT0014332

UGAAGCUGCCAGCAUGAUCUA

>ghr-miR2949b MIMAT0014346

UCUUUUGAACUGGAUUUGCCGA

>ghr-miR7500 MIMAT0029147

AUCGAGUUAUUCGAGUUAAUCGAG

>ghr-miR7486b MIMAT0029128

AAGGAAGCGCUUUGUCCACGUGGA

>ghr-miR398 MIMAT0014337

UGUGUUCUCAGGUCACCCCUU

>ghr-miR7510a MIMAT0029158

AAGGUCAUGAUCUUUAGCGGCGUU

>ghr-miR7496a MIMAT0029142

AUGACCAAAUUGAUAGAAUGUGUA

>ghr-miR7506 MIMAT0029153

AUGUCUGGGACAUGGCGUUGGCAC

>ghr-miR7501 MIMAT0029148

AUAUCUGAUUCUGACACGAAAAAA

>ghr-miR7493 MIMAT0029138

AAUAUUUUAAUAAUUCAAUCGUCA

>ghr-miR7503 MIMAT0029150

AGAUCGAUGGCUGAACAAGUUAGA

>ghr-miR7492c MIMAT0029136

CUAUAGAACAUGAUCUUUAGCGG

>ghr-miR156c MIMAT0005808

UGUCAGAAGAGAGUGAGCAC

>ghr-miR7485 MIMAT0029126

AAAGACAUCUUUGAAUUCUUGGAG

>ghr-miR7487 MIMAT0029129

AUACUCUUAUAGGACACUUGUUAA

>ghr-miR482a MIMAT0014339

UCUUUCCUACUCCUCCCAUACC

>ghr-miR2948-5p MIMAT0014344

UGUGGGAGAGUUGGGCAAGAAU

>ghr-miR7499 MIMAT0029146

AUAUAAUUUUCGGUUAAUUCGGUU

>ghr-miR7502 MIMAT0029149

UUUUUAACAGUAGAAAUGAAUGAA

>ghr-miR7492a MIMAT0029134

CUAUAGAACAUGAUCUUUAGCGG

>ghr-miR7514 MIMAT0029164

AUAAAGUGAUAAGUGAGAUCGUCU

>ghr-miR7486a MIMAT0029127

AAGGAAGCGCUUUGUCCACGUGGA

>ghr-miR390b MIMAT0005816

AAGCUCAGGAGGGAUAGCGCC

>ghr-miR399c MIMAT0014349

UGCCAAAGGAGAGUUGGCCUU

>ghr-miR2949a-3p MIMAT0015374

UGCAAAUCCAGUCAAAAGUUA

>ghr-miR7511 MIMAT0029160

AGAAGUUUUGCAUGUGUAGCUGAG

>ghr-miR156a MIMAT0005806

UGACAGAAGAGAGUGAGCAC

>ghr-miR7491 MIMAT0029133

UGGGAUCUUCGAGAGGAUUGAGCC

>ghr-miR399b MIMAT0005821

CGCCAAUGGAGAUUUGUCCGG

>ghr-miR7489 MIMAT0029131

AUUGUUGCCAAUACAGGAGAACGU

>ghr-miR7509 MIMAT0029156

UCAAAAGCACUUUUUGACAGCAAU

>ghr-miR394a MIMAT0014336

UUGGCAUUCUGUCCACCUCC

>ghr-miR7488 MIMAT0029130

UUUUGAGUACAGGGGACAAAA

>ghr-miR164 MIMAT0014331

UGGAGAAGCAGGGCACGUGCA

>ghr-miR160 MIMAT0029137

UAUGAGGAGCCAUGCAUGUAU

>ghr-miR2949c MIMAT0014347

UCUUUUGAACUGGAUUUGCCGA

>ghr-miR390c MIMAT0005817

AAGCUCAGGAGGGAUAGCGCC

>ghr-miR390a MIMAT0005815

AAGCUCAGGAGGGAUAGCGCC

>ghr-miR394b MIMAT0014335

UUGGCAUUCUGUCCACCUCC

>ghr-miR7495a MIMAT0029140

UUACUUUAGAUGUCUCCUUCA

>ghr-miR7484b MIMAT0029125

UUUGUAUAUUAGAUCAAAGAGCAA

>ghr-miR166b MIMAT0005813

UCGGACCAGGCUUCAUUCCCC

>ghr-miR7512 MIMAT0029161

UGCUACUUGUAGUUAUGCAUG

>ghr-miR7496b MIMAT0029143

AUGACCAAAUUGAUAGAAUGUGUA

>gra-miR8787 MIMAT0034241

UUUUCUUUUAAUUGGACGAGAUA

>gra-miR8659a MIMAT0034028

AAUUUUUUAAGGUUGUUUGUGGAA

>gra-miR8639h MIMAT0033994

UAAACAUAAGUAGAAUUAAACAAG

>gra-miR8721a MIMAT0034111

CAUCGAUAGUUUGAGGAUGUA

>gra-miR8710b MIMAT0034098

AUACAUGAACUUUGGUCCAA

>gra-miR8738a MIMAT0034150

GUUAACUUUAACGGUCAACGGUU

>gra-miR8772 MIMAT0034210

UUGGACUGUGGCUACAUAUAG

>gra-miR8760 MIMAT0034190

UGUGACAUCGUCAAAUUCGGCCAU

>gra-miR7494c MIMAT0034104

AUGGAGGAAAACAGAGGGAGAAGC

>gra-miR8663a MIMAT0034001

UUUGAACUUGGCAACUUUUUUCAC

>gra-miR3476 MIMAT0033982

UCGGACUGGAUUUGUUGACAA

>gra-miR7504i MIMAT0034101

AUAUGAAACUGAGAUACCAUG

>gra-miR8728 MIMAT0034259

CGGGCUUGGGCAAAAUUUUAGGCU

>gra-miR7504e MIMAT0034067

AGGAGAAAAAAAUCUGAUUUGUCA

>gra-miR8740 MIMAT0034152

UAAUGAUGUGGCACAAUAUUA

>gra-miR8716 MIMAT0034106

AUGUGUCAAAAUGUGAGGCUGUCA

>gra-miR8670b MIMAT0034245

AAAGUUGGGCCCCUGUUGGUGCGG

>gra-miR8765 MIMAT0034199

UUAGUGGAGCUCAUGCUAGAA

>gra-miR8771d MIMAT0034182

UGCCAUGUAGGAUUGUCGUUA

>gra-miR8660 MIMAT0034030

ACGAAAAAUUAAAUGGAGGGCUA

>gra-miR8642 MIMAT0033998

UGAUCAAAACAGGAACGAAUUCAA

>gra-miR8639b MIMAT0033990

UAAACAUAAGUAGAAUUAAACAAG

>gra-miR8757a MIMAT0034186

UGGGCCGAGCUUGGACAAGCAUA

>gra-miR8661 MIMAT0034031

CAUUACUUUUUCAUUCAUUAA

>gra-miR8776c MIMAT0034221

UUUCAAAGUCCUUGCAUACUAUUU

>gra-miR8664 MIMAT0034034

UGGAGUCAACGGUGGGACGGAGA

>gra-miR7486h MIMAT0034255

CAGCAACUCGCGCUGACGUGGACA

>gra-miR482c MIMAT0034205

UUCCCAAAACCUCCAAUUCCAA

>gra-miR172a MIMAT0033988

GCGGCAUUAUCAAGAUUCACA

>gra-miR8703c MIMAT0034090

AGUAGUCUAAUUGGUAUAGCUGAA

>gra-miR8707 MIMAT0034094

AUAAAAACUUUUGAAUAAUUCAGU

>gra-miR8638 MIMAT0034266

UGAAUACAGGAAUGGCUCUCU

>gra-miR7486j MIMAT0034110

CAUCAGCGCGAUGUCAGGGGACGU

>gra-miR8694a MIMAT0034074

AGGAUGCACUGUCAGCAAAAGUAU

>gra-miR8657a MIMAT0034022

UGUAGUAAUUGUAGAAGUUCAGGG

>gra-miR164a MIMAT0034113

CAUGUGCCUUGGCUCUCCAUC

>gra-miR7484h MIMAT0034215

UUUAACCGUAGAAAUGGAUGA

>gra-miR8637 MIMAT0033987

AGGCCCCUGUAUUGAGAGUCGGAU

>gra-miR8654c MIMAT0034019

AAGGAUACUACUUUGAUGGAGAAA

>gra-miR8674b MIMAT0034197

UUAGAAAUGGACCAUGGAUGAUGU

>gra-miR8724 MIMAT0034117

CCUGACAUGUCAGUAGAAAGCUC

>gra-miR8713 MIMAT0034102

AUGACUGUUUUAAUGAAGAUUGCG

>gra-miR8678 MIMAT0034049

AAUUUGGACUGUCACGUAGGA

>gra-miR8646 MIMAT0034004

UAGUGAGGAUGGGAAAUUUGU

>gra-miR8703a MIMAT0034088

AGUAGUCUAAUUGGUAUAGCUGAA

>gra-miR7484e MIMAT0034107

AUUUAAUGUAUAGGGACUAAU

>gra-miR8701 MIMAT0034082

AGGGGCUUAGAAAGAUGGGAC

>gra-miR7492o MIMAT0034251

CUAAAGAUCUGAGCAUUAGUGGCG

>gra-miR7492g MIMAT0034124

CGUGAUCUUUAGCGGCGUUUG

>gra-miR8670a MIMAT0034244

AAAGUUGGGCCCCUGUUGGUGCGG

>gra-miR8639i MIMAT0033995

UAAACAUAAGUAGAAUUAAACAAG

>gra-miR8665 MIMAT0034036

UUAAUUAUUAUAUAGAUCAAGGAU

>gra-miR7486d MIMAT0034035

UGGGGGCGCGCUUUGCUUACG

>gra-miR166c MIMAT0034156

UAUAUGAUCUCGGACCAGGCU

>gra-miR8639f MIMAT0033992

UAAACAUAAGUAGAAUUAAACAAG

>gra-miR398 MIMAT0014328

UGUGUUCUCAGGUCACCCCUU

>gra-miR8706b MIMAT0034093

AGUUUUAGGAUUGAUUUGAUGAAA

>gra-miR8699 MIMAT0034080

AGGGCCAUUUUGACAAAACAUGCA

>gra-miR157b MIMAT0005811

UUGACAGAAGAUAGAGAGCAC

>gra-miR8688 MIMAT0034061

AGAGGAUGCUUUAUAAAACUCAUA

>gra-miR8692 MIMAT0034065

AGCAGGUCGCGCUGACGUGGACA

>gra-miR8641 MIMAT0034262

UUUUUACUUUGGGACACUGAUGGC

>gra-miR167c MIMAT0034167

UCAGAUGAAGCUGCCAGCAUGA

>gra-miR8743a MIMAT0034163

UAUGAAAAGUUAUAAAAUGGUCAU

>gra-miR7502c MIMAT0034014

UUGUUAAAAGUUUCAUCCAUU

>gra-miR7504f MIMAT0034068

AGGAGAAAAAAAUCUGAUUUGUCA

>gra-miR8762a MIMAT0034192

UUAACAUUUGUUAACUUUGCUGAC

>gra-miR8730 MIMAT0034130

CUAAGAGAUUGGGAUUUGGUAGGA

>gra-miR8685 MIMAT0034058

AGACGACACUUCAAAGAUUUCUCC

>gra-miR8757b MIMAT0034188

UGGGUCGAGCUUAGGCAAGCAUA

>gra-miR8736 MIMAT0034148

GUGGAACGUGUUAAGAGAGGAAUA

>gra-miR8764 MIMAT0034198

UUAGAUUGCAUUUUACCCCUU

>gra-miR8683 MIMAT0034054

AGAACUCAUAACACAUUUAGAUAA

>gra-miR8762e MIMAT0034242

UUUUUAACUUUACUGACAUGGCAU

>gra-miR7492k MIMAT0034128

CGUGAUCUUUAGCGGCGUUUG

>gra-miR166d MIMAT0034180

UGAUGGGAAUGUUGUUUGGCU

>gra-miR8653b MIMAT0034201

UUCAAACUUAUUUUACGGCCA

>gra-miR8771c MIMAT0034154

UAGGGAGGUAACGAAGCUUAACGG

>gra-miR8719 MIMAT0034108

CAAAUUGGAUAACUGGACAGGUAA

>gra-miR7504j MIMAT0034247

AUAUGAAACUGAGAUACCAUG

>gra-miR8711 MIMAT0034099

AUACUACACUAAGGGCACUAGAUC

>gra-miR8777 MIMAT0034223

UUUCCAAUAGAAGAAUGACA

>gra-miR8674a MIMAT0034196

UUAGAAAGGGACCAUGGAUGAUGU

>gra-miR8657e MIMAT0034025

UGUAGUAAUUGUAGAAGUUCAGGG

>gra-miR8639e MIMAT0034264

UAAACAUAAGUAGAAUUAAACAAG

>gra-miR8743b MIMAT0034231

UUUGGAAAGUUAUAAAAUGGUCAU

>gra-miR8703b MIMAT0034089

AGUAGUCUAAUUGGUAUAGCUGAA

>gra-miR8681 MIMAT0034052

ACAUUGUUGAGGGUCUAAUCGG

>gra-miR8738b MIMAT0034151

GUUAACUUUAACGGUCAACGGUU

>gra-miR7494b MIMAT0034062

AGAGGGAGAAGCAGAAGAGAAUA

>gra-miR8735 MIMAT0034147

GGGGACAAUACCUUCGAUUGUUGG

>gra-miR7486g MIMAT0034122

CGCUUUGCUGACGUGGAAACAAAU

>gra-miR8676 MIMAT0034047

AAGGUUGAUGGUUAAAUUUGACU

>gra-miR8691 MIMAT0034064

AGAUGAUGAGAAAGGAAAGUCAAG

>gra-miR8697 MIMAT0034078

AGGGAAAUUGAUUCAAGUGAGCAU

>gra-miR8746 MIMAT0034170

UCCAUAUUUCACUAUCUCUUA

>gra-miR7486i MIMAT0034142

GCUGACGUGGAAGGAAAUCGCUA

>gra-miR8712 MIMAT0034100

AUAUCAUUGGUGAUGUAUCGUCUU

>gra-miR7492h MIMAT0034125

CGUGAUCUUUAGCGGCGUUUG

>gra-miR8720 MIMAT0034109

CAGUGUAGUUCUAAACCCGUCGGG

>gra-miR8743e MIMAT0034162

UAUGAAAAGUUACAAAAUGGUCAU

>gra-miR7492m MIMAT0034133

CUAUAGAACAUGACCUUUAGCAGC

>gra-miR8761 MIMAT0034191

UGUUGACGUUGCAUACAUGUGGAU

>gra-miR172b MIMAT0034141

GCAGCAUUAUCAAGAUUCACA

>gra-miR8782 MIMAT0034232

UUUGGUGUUGAAGGGGAAUAA

>gra-miR8654b MIMAT0034018

AAGGAUACUACUUUGAUGGAGAAA

>gra-miR8702 MIMAT0034083

AGGUAUUGUCUCUGGGGAAGGGUU

>gra-miR399 MIMAT0034146

GGGCACCUCUCACUUAGGCAGG

>gra-miR7502e MIMAT0034193

UUAACGGUAGAAAUGGAUGAA

>gra-miR7484d MIMAT0034056

AGAAUGACCGGUUUGCUCUUU

>gra-miR8721b MIMAT0034112

CAUCGAUAGUUUGAGGAUGUA

>gra-miR157a MIMAT0005810

UUGACAGAAGAUAGAGAGCAC

>gra-miR7505b MIMAT0033977

ACAGCUUUAGAAAUCAUCCCU

>gra-miR1446 MIMAT0034042

AACUCUCUCCCUCAAAGGCUA

>gra-miR7484m MIMAT0034234

UUUGUAUGUUAGAUCGAAGAG

>gra-miR8725 MIMAT0034118

CCUGACUAGAGACAAUACCAACCU

>gra-miR8658 MIMAT0034026

UUUGAAAAGUACAGGGACUAU

>gra-miR8737 MIMAT0034149

GUGUAUCUCCUGAAAACGACGACA

>gra-miR7486c MIMAT0034235

UUUGUCCACGUGAACAGAAAACGC

>gra-miR8767a MIMAT0034202

UUCAACUCUGCCAAGCAAUUG

>gra-miR8667 MIMAT0034038

UUUCCAAUAGUUUUAUGCCACAUC

>gra-miR7504l MIMAT0034226

UUUCUCCUGAUUUUUAGCAUUUUU

>gra-miR8750 MIMAT0034176

UCUUAGUUGGCAUAUACUCAAGGA

>gra-miR8698 MIMAT0034079

AGGGACAAUUAACUUUAACGGUCA

>gra-miR8773 MIMAT0034211

UUGGAUGAACGGUGCGUUUACUU

>gra-miR8756 MIMAT0034185

UGGACUGUUAAAAUUUUAAUGGCA

>gra-miR8645 MIMAT0034003

UUGGUACCUGAACUUGACGUCU

>gra-miR8643a MIMAT0033999

ACAUAUUAAGAAGUCGAAUUG

>gra-miR8653a MIMAT0034012

UAAAUCUACGUAUGGAAAUUC

>gra-miR8639a MIMAT0033989

UAAACAUAAGUAGAAUUAAACAAG

>gra-miR8673 MIMAT0034045

AAGAUAUGUAAACCUUGAGGGCUA

>gra-miR8705 MIMAT0034091

AGUUCAUACUAGUUCGUGGGUCA

>gra-miR8656 MIMAT0034021

UGGAAAAGUGGUAAUAAGGGG

>gra-miR167b MIMAT0033984

UCAGAUCAUCUUGCAGCUUCA

>gra-miR8684 MIMAT0034057

AGAAUGACUGGUUUACUCUUU

>gra-miR8762d MIMAT0034214

UUGUUAACUUUGAUGAUGUGGCAU

>gra-miR8649 MIMAT0034007

ACACUGUGGAAGUGGAUCUCUCUC

>gra-miR8634 MIMAT0033981

UUGGUAUGGAGGAUGGAAAAG

>gra-miR8652 MIMAT0034011

AUUAUGUGUAUGAAACUUUAGUU

>gra-miR8640 MIMAT0033996

UCUGACAGUGCACUGAAAACG

>gra-miR8695 MIMAT0034076

AGGAUGUAAAAGAAUAGGUGA

>gra-miR477 MIMAT0034119

CGAAGUCUUGGAAGAGAGUAA

>gra-miR8662 MIMAT0034032

GGCGUACUUUCAGAGAACAUUUUG

>gra-miR8709a MIMAT0034015

AUAUUAAAACUAUACAUGAACUUU

>gra-miR8689 MIMAT0034265

AGAGGUGCUCAUGGGCUGGGUCGG

>gra-miR8666 MIMAT0034037

UUGGUGAUCAUAAGUACAUAA

>gra-miR8723a MIMAT0034250

UGUAACAGUAAGCUGACGUGACA

>gra-miR8709c MIMAT0034096

AUACAUGAACUUCGAUUUAA

>gra-miR8753 MIMAT0034181

UGCCAAAUCAGGGAAGCGAAA

>gra-miR8654a MIMAT0034017

AAGGAUACUACUUUGAUGGAGAAA

>gra-miR8751b MIMAT0034178

UGAAAUUUGUAGAGAGAUAACGCU

>gra-miR8770 MIMAT0034208

UUGAUGGUGGUAAGAAAUGUGCAU

>gra-miR8675b MIMAT0034084

AGGUGAUGAUGUGGUACAAUCUCA

>gra-miR7504k MIMAT0034225

UUUCUCCUGAUUUUUAGCAUUUUU

>gra-miR8693 MIMAT0034073

AGGAUGAAAAUAUUGAUGUAGCAU

>gra-miR7492i MIMAT0034126

CGUGAUCUUUAGCGGCGUUUG

>gra-miR8677 MIMAT0034048

AAUGAAUCUAGGUUCUCUCUU

>gra-miR8690 MIMAT0034063

AGAGUGACUACUUCGUAACAAAAC

>gra-miR8778 MIMAT0034260

UUUCCAUAUUAGGGUUUGAACUUU

>gra-miR8680 MIMAT0034051

ACAGUGGAGGUAUUGUGCCUG

>gra-miR8784 MIMAT0034236

UUUGUCGACAUGUCAGGAAAGCGC

>gra-miR7492l MIMAT0034132

CUAUAGAACAUGACCUUUAGCAGC

>gra-miR8742a MIMAT0034158

UAUCUUAUUCAUCUUGGACUG

>gra-miR8639c MIMAT0033991

UAAACAUAAGUAGAAUUAAACAAG

>gra-miR7504g MIMAT0034069

AGGAGGAAAUCUGAUUUGUCAUUC

>gra-miR8785 MIMAT0034237

UUUUACAGCAGCUACAUCCAU

>gra-miR8657d MIMAT0034024

UGUAGUAAUUGUAGAAGUUCAGGG

>gra-miR7484f MIMAT0034157

UAUCUAACGUGUAGGGACUAA

>gra-miR8771b MIMAT0034144

GGAUUGUCGUUAGGGAGGUAA

>gra-miR7484i MIMAT0034216

UUUAACCGUAGAAAUGGAUGA

>gra-miR8657c MIMAT0034023

UGUAGUAAUUGUAGAAGUUCAGGG

>gra-miR8669 MIMAT0034040

AAAGACGAAAGACAAAAUCUCAAU

>gra-miR8747 MIMAT0034171

UCCCAGUUGUAGUUGGUCGUUCGG

>gra-miR8786b MIMAT0034239

UUUUAGUGAUGUGGCAGAAAGAUG

>gra-miR8745 MIMAT0034166

UCAACGGAGUUGGGAGACAAA

>gra-miR8776a MIMAT0034219

UUUCAAAGUCCUUGCAUACUAUUU

>gra-miR7492d MIMAT0034016

UGGGCUUAGAUUUUUUGCGGCGUU

>gra-miR8743d MIMAT0034161

UAUGAAAAGUUACAAAAUGGUCAU

>gra-miR7493b MIMAT0034256

UUGGAUUGUUAAAAGGUUAAUUGU

>gra-miR8694b MIMAT0034075

AGGAUGCACUGUCAGCAAAAGUAU

>gra-miR8774 MIMAT0034212

UUGGAUUUUGAUUCAUAGAUUCGU

>gra-miR7492e MIMAT0034115

CCAUGAUCUUUAGCGGCGUUU

>gra-miR8726 MIMAT0034120

CGAUGGAGUCUGGAGACAAAA

>gra-miR8648 MIMAT0034006

UUUCAGCGCAGUAGAAGGAUU

>gra-miR8733 MIMAT0034138

GAGCUUGGAAGUGCAUCCGGC

>gra-miR8644 MIMAT0034002

UUAAACUCCAGAAGUAGGGCU

>gra-miR8732 MIMAT0034137

GAAGAGUAUAGGGACUUAUGGCAU

>gra-miR8639d MIMAT0034263

UAAACAUAAGUAGAAUUAAACAAG

>gra-miR8748 MIMAT0034172

UCGGUGGAGAUGGAUAAAAUGAAU

>gra-miR8755 MIMAT0034184

UGGACGCGCUUUGCUGACGUGGCA

>gra-miR8708 MIMAT0034095

AGGAGGAGUUAUGGAUAGUUUUA

>gra-miR7484g MIMAT0034206

UUCUUAUAUGUUAGAUCAAAGAGC

>gra-miR7494d MIMAT0034168

UCAUGGACUUUAGCGGCGUU

>gra-miR8767b MIMAT0034203

UUCAACUCUGCCAAGCAAUUG

>gra-miR8714 MIMAT0034103

AUGGACGAAAUGAAAGUAGAG

>gra-miR8723b MIMAT0034116

CCAUUAACGGUGUAACAGUAAGCU

>gra-miR7484l MIMAT0034229

UUUGAUCUAAUGUAUGGAGACUA

>gra-miR7484c MIMAT0034055

AGAAGUCGAAUUGCAUUUUG

>gra-miR8635 MIMAT0033985

UCGGAUCUUCAAACGGUGGAG

>gra-miR8643b MIMAT0034000

ACAUAUUAAGAAGUCGAAUUG

>gra-miR8639g MIMAT0033993

UAAACAUAAGUAGAAUUAAACAAG

>gra-miR3267 MIMAT0033980

UCUGUCGCAGGGGAGAUGGCUG

>gra-miR482d MIMAT0034224

UUUCCUAUGCCCCCCAUUCCAC

>gra-miR7502f MIMAT0034267

UUUAGCAGUAGAAAUAGAUGA

>gra-miR8776d MIMAT0034222

UUUCAAAGUCCUUGCAUACUAUUU

>gra-miR7494f MIMAT0034174

UCGUGAUCUUUAGCGGUGUUU

>gra-miR8663b MIMAT0034033

UGAGGCAUGAACUUGGCAAUUUU

>gra-miR8771a MIMAT0034143

GGAUUGUCGUUAGGGAGGUAA

>gra-miR8771e MIMAT0034145

GGCAUUUAAAACACAUUUGGACUG

>gra-miR8766 MIMAT0034200

UUAUUUUGGAAUUAGAAAAGUCGU

>gra-miR167a MIMAT0033983

UCAGAUCAUCUUGCAGCUUCA

>gra-miR482 MIMAT0014329

UCUUUCCAAUUCCUCCCAUUCC

>gra-miR530a MIMAT0034086

AGGUGCAGAUGCAGUUGCAGG

>gra-miR8769 MIMAT0034207

UUGAACUUUGACCGGAUCUAGGGA

>gra-miR8672 MIMAT0034044

AAGAGAAAUGAUUGUAUGAAACAG

>gra-miR7486f MIMAT0034121

CGCUUUGCUGACGUGGAAACAAAU

>gra-miR8709b MIMAT0034169

UCAUGUAUAAUUUUGAGAUUUGUC

>gra-miR7504d MIMAT0034066

AGGAAAAAAAAUCUGAUUUGUCAU

>gra-miR8675a MIMAT0034046

AAGGUGAUGACCUGCUACAAUUUU

>gra-miR8659b MIMAT0034029

AAUUUUUUAAGGUUGUUUGUGGAA

>gra-miR8751a MIMAT0034177

UGAAAUUUGUAGAGACAAAACGCU

>gra-miR8759 MIMAT0034189

UGGUGGAAGUAUUGUGCCCGG

>gra-miR8710a MIMAT0034097

AUACAUGAACUUUGGUCCAA

>gra-miR7484j MIMAT0034217

UUUACUCUUUGAUCUAACGUACA

>gra-miR8729 MIMAT0034129

CUAAAAUCGAGCAUAGACAUC

>gra-miR7484k MIMAT0034218

UUUAUACAUUAGAUCAAAGAGCAA

>gra-miR8767c MIMAT0034240

UUUUCAACUCUGCCAAGCAAU

>gra-miR827b MIMAT0036968

UUUGUUUAUGGUCAUCUAAGC

>gra-miR8632 MIMAT0033978

AUGAGCUAGAAGUUGGAACUC

>gra-miR8762c MIMAT0034243

UUAACGUUUGUUAACUUUGUUGAU

>gra-miR7502b MIMAT0034013

UUGUUAAAAGUUUCAUCCAUU

>gra-miR8741 MIMAT0034153

UAGCACUGAAGAUGAUGAUGG

>gra-miR8651 MIMAT0034010

UUAAUGGAUCAUAACAGCAGGUAU

>gra-miR8657b MIMAT0034246

UGUAGUAAUUGUAGAAGUUCAGGG

>gra-miR7492f MIMAT0034249

CCAUGAUCUUUAGCGGCGUUU

>gra-miR7504h MIMAT0034070

AGGAGGAAUAAGUCUGAUUUGUCA

>gra-miR7504c MIMAT0034027

AAGAGAAAAAAUCGGAUUUAUCAU

>gra-miR8743c MIMAT0034160

UAUGAAAAGUUACAAAAUGGUCAU

>gra-miR8786a MIMAT0034238

UUUUAGUGAUGUGGCAGAAAGAUG

>gra-miR8783 MIMAT0034233

UUUGUACGUGGCGGGAGAUAU

>gra-miR8696 MIMAT0034077

AGGCUCUAAUGUGGGAACGACUGC

>gra-miR8633 MIMAT0033979

UAAGUGAAGAAAGAGGUAGGUU

>gra-miR8781a MIMAT0034164

UAUUGAAUUAUUUAGAACUAGGAU

>gra-miR7484n MIMAT0034258

CUAGUUUGCUCUUUGAUCUAAUGU

>gra-miR8739 MIMAT0034257

UAACUAAAUAGUGACACGUGGCAU

>gra-miR8754 MIMAT0034183

UGCCGCCUGACACACUGACGACAC

>gra-miR8717 MIMAT0034248

AUUGGUUGUUCUGAUUCGAGGCUA

>gra-miR8655 MIMAT0034020

AUAUGUUCUGAUUAUAUCUGU

>gra-miR7492j MIMAT0034127

CGUGAUCUUUAGCGGCGUUUG

>gra-miR7506b MIMAT0034134

CUGGGACAUGGCGUUGGCAA

>gra-miR7504m MIMAT0034227

UUUCUCCUGAUUUUUAGCAUUUUU

>gra-miR8775 MIMAT0034213

UUGUGAGAUUGAAGCUGAUGG

>gra-miR8668 MIMAT0034039

UUUUGGAAUCAAGACUACAU

>gra-miR8706a MIMAT0034092

AGUUUUAGGAUUGAUUUGAUGAAA

>gra-miR7502d MIMAT0034136

CUUUUAACAGUAGAAUUUGAUGGA

>gra-miR8752 MIMAT0034179

UGAUGGAGAUAGGUAUCUGCA

>gra-miR8776b MIMAT0034220

UUUCAAAGUCCUUGCAUACUAUUU

>gra-miR8704 MIMAT0034252

AGUCAGUUCAGGCAGUCAGACCGU

>gra-miR8700 MIMAT0034081

AGGGCGACAACGGUUACUGUGAUU

>gra-miR8687 MIMAT0034060

AGAGACAAAAGAAACACGUUCUAC

>gra-miR7492n MIMAT0034155

UAUAGAUCAGGGUCUUUAGCGGCG

>gra-miR7504n MIMAT0034072

AGGAUAAAAUUACUGAUGUGGCAU

>gra-miR8731 MIMAT0034131

CUAUAAACAGUCGAUGGUAUC

>gra-miR8715 MIMAT0034105

AUGGGCUGGCACAUGGGCGUGUGG

>gra-miR5207 MIMAT0034135

CUUAAGGUGGGUUUGGAUGGGCGA

>gra-miR8674c MIMAT0034253

AAGGGAUACAGGAUGAUGUGGCAU

>gra-miR7494e MIMAT0034173

UCGUGAUCUUUAGCGGUGUUU

>gra-miR8734 MIMAT0034139

GAUCAUAUCUCGUACGUUAGGACA

>gra-miR8686 MIMAT0034059

AGAGAAUGUCAGUACUAGAGGCAG

>gra-miR7484o MIMAT0034140

GAUUUAACGUAUAGGGACUAA

>gra-miR8762b MIMAT0034194

UUAACGUUUGUUAACUUUGUUGAU

>gra-miR8763 MIMAT0034195

UUAAUACUGUUAAAUUUGUUGGU

>gra-miR8675c MIMAT0034085

AGGUGAUGAUGUGGUACAAUCUUA

>gra-miR8781b MIMAT0034254

UUUGAUUAGGAAGUUUGAGGAUCA

>gra-miR8647 MIMAT0034005

ACCGAUUUGCUCUUUGAUCUA

>gra-miR8768 MIMAT0034204

UUCCAUGUCACAGAGAUGUUG

>gra-miR8780 MIMAT0034230

UUUGAUGAUGUGGCAACAAAGCGC

>gra-miR8744 MIMAT0034165

UCAAAAAAUGGGCAAAGUAGUCAU

>gra-miR8722 MIMAT0034114

CAUGUUUUUCCUGUUCAUCUUC

>gra-miR8636 MIMAT0033986

CGGACCCUCUAACAGUGGAGG

>gra-miR8679 MIMAT0034050

ACAAGCGAGUUAAGGAUGUGACAA

>gra-miR7486e MIMAT0033997

CGACUUGCUGACGUGGAAGGAAAU

>gra-miR8682 MIMAT0034053

ACUAGCUCAGGCUCUCGGCAG

>gra-miR5741 MIMAT0034009

UAGUAUAAGGACUAAAUUGGU

>gra-miR8749 MIMAT0034175

UCUCGGACCCUCAACGGAAGG

>gra-miR7584e MIMAT0034071

AGGAGUUAGAUUGUAUUUUAU

>gra-miR8718 MIMAT0034261

AUUUUGGAAGAAUUUCAGCUG

>gra-miR8727 MIMAT0034123

CGGACUCUCAAACAGUGGAGGUA

>gra-miR530b MIMAT0034087

AGGUGCAGGUGCAGGCGCAGC

>gra-miR8650 MIMAT0034008

AGUGGAAGGACAAAAUGCAAUCUG

>gra-miR8671 MIMAT0034043

AACUGUUCGUGAUAAGAUGUCGGU

>gra-miR8771f MIMAT0034209

UUGGACAUCCAAGUUAGCAUUUA

>gra-miR7503b MIMAT0034041

AACCGAUGGUUGAACAUGUUAGAC

>gra-miR8758 MIMAT0034187

UGGGCUUCUUGCAAGAUGAAGGUA

>gra-miR8779 MIMAT0034228

UUUGAUCUAACGUACAGAGACUA

>gra-miR8742b MIMAT0034159

UAUCUUAUUCAUCUUGGACUG

>lus-miR166e MIMAT0027217

UCGGACCAGGCUUCAUUCCUC

>lus-miR160f MIMAT0027195

UGCCUGGCUCCCUGUAUGCCA

>lus-miR171e MIMAT0027149

UGAUUGAGCCGUGCCAAUAUC

>lus-miR159a MIMAT0027182

AUUGGAGUGAAGGGAGCUCGA

>lus-miR172h MIMAT0027228

AGAAUCUUGAUGAUGCUGCAU

>lus-miR172d MIMAT0027164

AGAAUCUUGAUGAUGCUGCAU

>lus-miR172a MIMAT0027135

AGAAUCUUGAUGAUGCUGCAU

>lus-miR398d MIMAT0031859

UGUGUUCUCAGGUCACCCCUC

>lus-miR156d MIMAT0027180

UGACAGAAGAAAGAGAGCAC

>lus-miR164b MIMAT0027138

UGGAGAAGCAGGGCACGUGCA

>lus-miR171h MIMAT0027219

AUGAGCCGAACCAAUAUCACU

>lus-miR399c MIMAT0027154

UGCCAAAGGAGAUUUGCCCGG

>lus-miR398a MIMAT0027147

UGUGUUCUCAGGUCGCCCCUG

>lus-miR168a MIMAT0027185

UCGCUUGGUGCAGGUCGGGAA

>lus-miR399e MIMAT0027156

UGCCAAAGGAGAUUUGCCCGG

>lus-miR390d MIMAT0027203

AAGCUCAGGAGGGAUAGCGCC

>lus-miR156e MIMAT0027184

UUGACAGAAGAUAGAGAGCAC

>lus-miR390a MIMAT0027167

AAGCUCAGGAGGGAUAGCGCC

>lus-miR171b MIMAT0027134

UGAUUGAGCCGUGCCAAUAUC

>lus-miR160g MIMAT0027202

UGCCUGGCUCCCUGGAUGCCA

>lus-miR399d MIMAT0027155

UGCCAAAGGAGAAUUGCCCUG

>lus-miR169k MIMAT0027209

UAGCCAAGGAUGACUUGCCUG

>lus-miR397b MIMAT0027181

UCAUUGAGUGCAGCGUUGAUG

>lus-miR395c MIMAT0027136

CUGAAGUGUUUGGGGGAACUC

>lus-miR171i MIMAT0027224

UUGAGCCGUGCCAAUAUCACG

>lus-miR171d MIMAT0027146

AGAUAUUGGUGCGGUUCAAUC

>lus-miR169d MIMAT0027128

UAGCCAAGGAUGACUUGCCCA

>lus-miR166h MIMAT0027231

UCGGACCAGGCUUCAUUCCCC

>lus-miR166d MIMAT0027204

UCGGACCAGGCUUCAUUCCCC

>lus-miR395e MIMAT0027208

CUGAAGUGUUUGGAGGAACUC

>lus-miR156i MIMAT0027178

UUGACAGAAGAUAGAGAGCAC

>lus-miR159b MIMAT0027233

UUUGGAUUGAAGGGAGCUCUC

>lus-miR319b MIMAT0027130

UUGGACUGAAGGGAGCUCCC

>lus-miR169l MIMAT0027215

CAGCCAAGGAUGACUUGCCGA

>lus-miR396b MIMAT0027212

UUCCACAGCUUUCUUGAACUU

>lus-miR394a MIMAT0027141

UUGGCAUUCUGUCCACCUCC

>lus-miR396e MIMAT0031858

UUCCACAGCUUUCUUGAACUU

>lus-miR159c MIMAT0027235

UUUGGAUUGAAGGGAGCUCUU

>lus-miR395d MIMAT0027145

CUGAAGUGUUUGGGGGAACUC

>lus-miR171g MIMAT0027193

UGAUUGAGCCGCGUCAAUAUC

>lus-miR168b MIMAT0027206

UCGCUUGGUGCAGGUCGGGAA

>lus-miR156f MIMAT0027186

UUGACAGAAGAUAGAGAGCAC

>lus-miR530a MIMAT0027162

UGCAUUUGCACCUGCACCUU

>lus-miR395b MIMAT0027133

CUGAAGUGUUUGGGGGAACUC

>lus-miR171j MIMAT0031857

UGAUUGAGCCGCGUCAAUAUC

>lus-miR172i MIMAT0027230

GGAAUCUUGAUGAUGCUGCAG

>lus-miR169a MIMAT0027125

UAGCCAAGGAUGACUUGCCCA

>lus-miR396a MIMAT0027207

UUCCACAGCUUUCUUGAACUG

>lus-miR169g MIMAT0027196

CAGCCAAGGAUGACUUGCCGA

>lus-miR172f MIMAT0027205

AGAAUCUUGAUGAUGCUGCAU

>lus-miR399a MIMAT0027152

UGCCAAAGGAGAUUUGCCCGG

>lus-miR167e MIMAT0027165

UGAAGCUGCCAGCAUGAUCUA

>lus-miR398f MIMAT0031861

GGUGUUCUCAGGUCGCCCCUG

>lus-miR395a MIMAT0027132

CUGAAGUGUUUGGGGGAACUC

>lus-miR393b MIMAT0027218

UCCAAAGGGAUCGCAUUGAUC

>lus-miR172c MIMAT0027144

AGAAUCUUGAUGAUGCUGCAU

>lus-miR166k MIMAT0031856

UCGGACCAGGCUUCAUUCCCCC

>lus-miR164e MIMAT0027227

UGGAGAAGCAGGGCACGUGCA

>lus-miR156b MIMAT0027161

UUGACAGAAGAUAGAGAGCAC

>lus-miR397a MIMAT0027170

AUUGAGUGCAGCGUUGAUGAA

>lus-miR169c MIMAT0027127

UAGCCAAGGAUGACUUGCCUG

>lus-miR393c MIMAT0027229

UCCAAAGGGAUCGCAUUGAUC

>lus-miR156a MIMAT0027216

UGACAGAAGAGAGUGAGCAC

>lus-miR390b MIMAT0027191

AAGCUCAGGAGGGAUAGCGCC

>lus-miR167f MIMAT0027175

UGAAGCUGCCAGCAUGAUCUG

>lus-miR156c MIMAT0027179

UUGACAGAAGAUAGAGAGCAC

>lus-miR166i MIMAT0027236

UCGGACCAGGCUUCAUUCCCCC

>lus-miR166c MIMAT0027192

UCGGACCAGGCUUCAUUCCCC

>lus-miR171a MIMAT0027122

AGAUUGAGCCGCGCCAAUAUC

>lus-miR167b MIMAT0027131

UGAAGCUGCCAGCAUGAUCUU

>lus-miR160d MIMAT0027171

UGCCUGGCUCCCUGUAUGCCA

>lus-miR156h MIMAT0031854

UUGACAGAAGAUAGAGAGCAC

>lus-miR169e MIMAT0027139

UGAGCCAAGGAUGACUUGCCG

>lus-miR160e MIMAT0027189

UGCCUGGCUCCCUGUAUGCCA

>lus-miR160i MIMAT0027222

UGCCUGGCUCCCUGUAUGCCA

>lus-miR167i MIMAT0027190

UGAAGCUGCCAGCAUGAUCUG

>lus-miR530b MIMAT0027169

UGCAUUUGCACCUGCACCUU

>lus-miR398b MIMAT0027160

UGUGUUCUCAGGUCACCCCU

>lus-miR166f MIMAT0027221

UCGGACCAGGCUUCAUUCCUU

>lus-miR169i MIMAT0027199

UGAGCCAAGGAUGACUUGCCG

>lus-miR166b MIMAT0027172

UCGGACCAGGCUUCAUCCCCC

>lus-miR160j MIMAT0027225

UGCCUGGCUCCCUGUAUGCCA

>lus-miR167d MIMAT0027159

UGAAGCUGCCAGCAUGAUCUA

>lus-miR399g MIMAT0027188

UGCCAAAGGAGAUUUGCCCAG

>lus-miR393a MIMAT0027173

UCCAAAGGGAUCGCAUUGAUC

>lus-miR390c MIMAT0027197

AAGCUCAGGAGGGAUAGCGCC

>lus-miR164d MIMAT0027183

UGGAGAAGCAGGGCACGUGCA

>lus-miR169h MIMAT0027198

CAGCCAAGGAUGACUUGCCGG

>lus-miR167c MIMAT0027158

UGAAGCUGCCAGCAUGAUCUA

>lus-miR828a MIMAT0027220

UCUUGCUCAAAUGAGUGUUCCA

>lus-miR319a MIMAT0027123

UUGGACUGAAGGGAGCUCCCU

>lus-miR166j MIMAT0031855

UCGGACCAGGCUUCAUUCCCC

>lus-miR396c MIMAT0027213

UUCCACAGCUUUCUUGAACUG

>lus-miR166a MIMAT0027168

UCGGACCAGGCUUCAUUCCCC

>lus-miR399f MIMAT0027157

UGCCAAAGGAGAUUUGCCCAG

>lus-miR162b MIMAT0027187

UCGAUAAACCUCUGCAUCCAG

>lus-miR160c MIMAT0027151

UGCCUGGCUCCCUGGAUGCCA

>lus-miR169b MIMAT0027126

UAGCCAAGGAUGACUUGCCUG

>lus-miR398e MIMAT0031860

UGUGUUCUCAGGUCACCCCUC

>lus-miR172g MIMAT0027226

GGAAUCUUGAUGAUGCUGCAU

>lus-miR393d MIMAT0027232

UCCAAAGGGAUCGCAUUGAUC

>lus-miR164c MIMAT0027142

UGGAGAAGCAGGGCACGUGCA

>lus-miR167g MIMAT0027176

UGAAGCUGCCAGCAUGAUCUG

>lus-miR167a MIMAT0027124

UGAAGCUGCCAGCAUGAUCUC

>lus-miR171f MIMAT0027163

AGAUUGAGCCGCGCCAAUAUC

>lus-miR394b MIMAT0027194

UUGGCAUUCUGUCCACCUCC

>lus-miR162a MIMAT0027166

UCGAUAAACCUCUGCAUCCAG

>lus-miR156g MIMAT0027201

UGACAGAAGAGAGUGAGCAC

>lus-miR172b MIMAT0027143

AGAAUCUUGAUGAUGCUGCAU

>lus-miR160a MIMAT0027137

UGCCUGGCUCCCUGUAUGCCA

>lus-miR172j MIMAT0027234

GCAGCAUCAUCAAGAUUCCCA

>lus-miR166g MIMAT0027223

UCGGACCAGGCUUCAUUCCCC

>lus-miR171c MIMAT0027140

UGAUUGAGCCGUGCCAAUAUC

>lus-miR167h MIMAT0027177

UGAAGCUGCCAGCAUGAUCUA

>lus-miR160b MIMAT0027150

UGCCUGGCUCCCUGUAUGCCA

>lus-miR169f MIMAT0027148

CAGCCAAGGAUGACUUGCCGG

>lus-miR160h MIMAT0027211

UGCCUGGCUCCCUGUAUGCCA

>lus-miR399b MIMAT0027153

UGCCAAAGGAGAAUUGCCCUG

>lus-miR398c MIMAT0027214

UGUGUUCUCAGGUCACCCCU

>lus-miR164a MIMAT0027129

UGGAGAAGCAGGGCACGUGCA

>lus-miR169j MIMAT0027200

CAGCCAAGGAUGACUUGCCGG

>lus-miR172e MIMAT0027174

GGAAUCUUGAUGAUGCUGCAG

>lus-miR396d MIMAT0027237

UCCCACAGCUUUAUUGAACUG

>lus-miR408a MIMAT0027210

AUGCACUGCCUCUUCCCUGGC

>mdm-miR2118b MIMAT0026035

CUACCGAUGCCACUAAGUCCCA

>mdm-miR159c MIMAT0026053

GAAUUCCUUCUCCUCUCCUUU

>mdm-miR390e MIMAT0025973

AAGCUCAGGAGGGAUAGCGCC

>mdm-miR156e MIMAT0025871

UGACAGAAGAGAGUGAGCAC

>mdm-miR156ad MIMAT0025896

UGACAGAAGAAAGUGAGCAC

>mdm-miR398c MIMAT0026000

UGUGUUCUCAGGUCGCCCCUG

>mdm-miR7121e MIMAT0026044

UCCUCUUGGUGAUCGCCCUGC

>mdm-miR395i-5p MIMAT0037471

GUUCCCUUGACCACUUCAUUG

>mdm-miR171i MIMAT0025946

UGAGCCGAACCAAUAUCACUC

>mdm-miR828b MIMAT0026030

UCUUGCUCAAAUGAGUAUUCCA

>mdm-miR11012b MIMAT0043615

CCGCUGCCUAUGCAAUGCACCC

>mdm-miR399c MIMAT0026003

UGCCAAAGGAGAAUUGCCCUG

>mdm-miR156m MIMAT0025879

UGACAGAAGAGAGUGAGCAC

>mdm-miR390a MIMAT0025969

AAGCUCAGGAGGGAUAGCGCC

>mdm-miR171m MIMAT0025950

UUGAGCCGUGCCAAUAUCACA

>mdm-miR10999b MIMAT0043592

GGGCGUGAUAUUCACACACCU

>mdm-miR169f MIMAT0026068

UGAAGAGAAGAGCGUUGUUUGG

>mdm-miR482a-3p MIMAT0011165

UUCCCAAGCCCGCCCAUUCCUA

>mdm-miR395k MIMAT0043586

GUUUCCUCAAACACUUCAUU

>mdm-miR3627d MIMAT0043534

UCCAUCCUCCUGUGACAUGAA

>mdm-miR319e MIMAT0043567

GAGCUUUCUUCAGUCCACUC

>mdm-miR408c MIMAT0026032

ACAGGGAAGAGGUAGAGCAUG

>mdm-miR159a MIMAT0025898

CUUGGAUUGAAGGGAGCUCC

>mdm-miR399d MIMAT0026004

UGCCAAAGGAGAGUUGCCCUA

>mdm-miR319d MIMAT0043552

AACUGCCGACUCAUUCACUCA

>mdm-miR156i MIMAT0025875

UGACAGAAGAGAGUGAGCAC

>mdm-miR10989c MIMAT0043557

CAAAGCUUUUAAUAUCAGUCGA

>mdm-miR10980b MIMAT0043531

CACCUGGGACUUGCAGCCAUG

>mdm-miR393f MIMAT0026063

AUCAUGCGAUCCCUUCGGACG

>mdm-miR166d MIMAT0025916

UCGGACCAGGCUUCAUUCCCC

>mdm-miR393d MIMAT0026061

AUCAUGCGAUCCCUUCGGACG

>mdm-miR166b MIMAT0025914

UCGGACCAGGCUUCAUUCCCC

>mdm-miR403a MIMAT0026011

UUAGAUUCACGCACAAACUCG

>mdm-miR399g MIMAT0026007

UGCCAAAGGAGAUUUGCUCGG

>mdm-miR156b MIMAT0025868

UGACAGAAGAGAGUGAGCAC

>mdm-miR172g MIMAT0025958

AGAAUCUUGAUGAUGCUGCAU

>mdm-miR319a MIMAT0025967

UUGGACUGAAGGGAGCUCCCU

>mdm-miR156h MIMAT0025874

UGACAGAAGAGAGUGAGCAC

>mdm-miR169k MIMAT0043616

UAGCCAAGGAUGACUUGCCUG

>mdm-miR2118c MIMAT0026036

CUACCGAUGCCACUAAGUCCCA

>mdm-miR156l MIMAT0025878

UGACAGAAGAGAGUGAGCAC

>mdm-miR395f MIMAT0025985

CUGAAGUGUUUGGGGGAACUC

>mdm-miR10993f MIMAT0043590

ACAUGUGGUGUACCAUCCUGU

>mdm-miR11001 MIMAT0043599

UACAAAGAUGGACUCCACCCUA

>mdm-miR10991c MIMAT0043563

CGAGCCAUUGAAAUUCGAUCC

>mdm-miR408d MIMAT0026033

ACAGGGAAGAGGUAGAGCAUG

>mdm-miR160d MIMAT0025903

UGCCUGGCUCCCUGUAUGCCA

>mdm-miR396d MIMAT0025992

UUCCACAGCUUUCUUGAACUU

>mdm-miR10984a-5p MIMAT0043547

GGUAAUUGACUGUGAAAUCGU

>mdm-miR390d MIMAT0025972

AAGCUCAGGAGGGAUAGCGCC

>mdm-miR172l MIMAT0025963

GGAAUCUUGAUGAUGCUGCAG

>mdm-miR167g MIMAT0025928

UGAAGCUGCCAGCAUGAUCUA

>mdm-miR171f-3p MIMAT0025943

UUGAGCCGUGCCAAUAUCACG

>mdm-miR164c MIMAT0025909

UGGAGAAGCAGGGCACGUGCA

>mdm-miR395c MIMAT0025982

CUGAAGUGUUUGGGGGAACUC

>mdm-miR169c MIMAT0025936

UAGCCAAGGAUGACUUGCCCG

>mdm-miR172d MIMAT0025955

AGAAUCUUGAUGAUGCUGCAU

>mdm-miR156s MIMAT0025885

CUGACAGAAGAUAGAGAGCAC

>mdm-miR10991e MIMAT0043565

CGAGCCAUUGAAAUUCGAUCC

>mdm-miR7124a MIMAT0026054

CACCAAUAUCAACUUUAUUUG

>mdm-miR535a MIMAT0026024

UGACAACGAGAGAGAGCACGC

>mdm-miR167c MIMAT0025924

UGAAGCUGCCAGCAUGAUCUA

>mdm-miR171e MIMAT0025942

UGAUUGAGCCGCGCCAAUAUC

>mdm-miR11002b MIMAT0043601

GAGGAUGAGCUUCGGCGGUGA

>mdm-miR396g MIMAT0025995

UUCCACGGCUUUCUUGAACUG

>mdm-miR10986 MIMAT0043551

UGGCACCAAAGUCACCACCCG

>mdm-miR156ae MIMAT0025897

UGACAGAAGAAAGUGAGCAC

>mdm-miR530a MIMAT0043542

UGCAUUUGCACCUGCACUUGU

>mdm-miR399k MIMAT0043550

UGCCAAAGGAGAGUUGCCCUU

>mdm-miR10978b MIMAT0043528

GUUGGGAAUCGAAGCAUCACGA

>mdm-miR172p MIMAT0043620

AGAAUCUUGAUGAUGCUGCAU

>mdm-miR7121d MIMAT0026043

UCCUCUUGGUGAUCGCCCUGC

>mdm-miR159d MIMAT0043539

UUUGGAUUGAAGGGAGCUCUA

>mdm-miR10997 MIMAT0043581

UAACCUUAUUUGAUUUCACGA

>mdm-miR156ac MIMAT0025895

UUGACAGAAGAUAGAGAGCAC

>mdm-miR395b MIMAT0025981

CUGAAGUGUUUGGGGGAACUC

>mdm-miR535b MIMAT0026025

UGACAAGGAGAGAGAGCACGC

>mdm-miR10990 MIMAT0043560

CCAAGGAAAAUUUUAUGACGA

>mdm-miR396a MIMAT0025989

UUCCACAGCUUUCUUGAACAG

>mdm-miR11002a MIMAT0043600

GAGGAUGAGCUUCGGCGGUGA

>mdm-miR10982c MIMAT0043537

CGGAAUGAAGCUUACGAGAAUG

>mdm-miR164d MIMAT0025910

UGGAGAAGCAGGGCACGUGCA

>mdm-miR10981a MIMAT0043532

UGACCAACAUAUAUGGGCCGU

>mdm-miR169b MIMAT0025935

UAGCCAAGGAUGAUUUGCCUGC

>mdm-miR11007 MIMAT0043607

CCGUCAAUUAUUUUGGUACGU

>mdm-miR156r MIMAT0025884

CUGACAGAAGAUAGAGAGCAC

>mdm-miR167b MIMAT0025923

UGAAGCUGCCAGCAUGAUCUA

>mdm-miR827 MIMAT0026028

UUAGAUGACCAUCAACGAACA

>mdm-miR10994-3p MIMAT0043573

UGCUUUUUUCUUGACCAUAGC

>mdm-miR11000 MIMAT0043598

GUGUUCCAAAGAAAUCCGGAGU

>mdm-miR393g MIMAT0043583

AUCAUGCUAUCCCUUUGGAUU

>mdm-miR3627b MIMAT0026017

UCGCAGGAGAGAUGGCACUA

>mdm-miR395j MIMAT0043574

GUUCCCUUGACCACUUCAUUG

>mdm-miR530c MIMAT0043544

UGCAUUUGCACCUGCACUUGU

>mdm-miR169o MIMAT0043632

UAGCCAGGGAUGACUUGCCU

>mdm-miR169g MIMAT0043575

CAGCCAAGGAUGACUUGCCGG

>mdm-miR10993b MIMAT0043571

AUCCCACCAUUUAUAUAGCGA

>mdm-miR10989d MIMAT0043558

CAAAGCUUUUAAUAUCAGUCGA

>mdm-miR7127a MIMAT0026064

AUACUCAUCGAAUUUGUCAUA

>mdm-miR11012a MIMAT0043614

CCGCUGCCUAUGCAAUGCACCC

>mdm-miR11009 MIMAT0043611

AUGCACAACAAUAUGAGGGUGU

>mdm-miR10993e MIMAT0043589

ACAUGUGGUGUACCAUCCUGU

>mdm-miR10984b-3p MIMAT0043610

CUCACGUACGCUGUCCCGAGAA

>mdm-miR11014 MIMAT0043623

CGACCAUUCAUGAAAACUGCC

>mdm-miR156n MIMAT0025880

UGACAGAAGAGAGUGAGCAC

>mdm-miR167f MIMAT0025927

UGAAGCUGCCAGCAUGAUCUA

>mdm-miR172i MIMAT0025960

GGAAUCUUGAUGAUGCUGCAU

>mdm-miR11011a MIMAT0043613

AAGUUCAUUCAAACACCAUGU

>mdm-miR7124b MIMAT0026055

CACCAAUAUCAACUUUAUUUG

>mdm-miR156p MIMAT0025882

CUGACAGAAGAUAGAGAGCAC

>mdm-miR171n MIMAT0025951

UUGAGCCGUGCCAAUAUCACA

>mdm-miR395d-5p MIMAT0037469

GUUCCCUUGACCACUUCAUUG

>mdm-miR171q MIMAT0043605

GGAUAUUGGUCCGGUUCAAUA

>mdm-miR396e MIMAT0025993

UUCCACAGCUUUCUUGAACUU

>mdm-miR156t MIMAT0025886

UUGACAGAAGAGAGAGAGCAC

>mdm-miR11020 MIMAT0043631

GACAUUACAACGGUUACACGG

>mdm-miR10999a MIMAT0043591

GGGCGUGAUAUUCACACACCU

>mdm-miR390c MIMAT0025971

AAGCUCAGGAGGGAUAGCGCC

>mdm-miR7121h MIMAT0026047

UCCUCUUGGUGAUCGCCCUGC

>mdm-miR172f MIMAT0025957

AGAAUCUUGAUGAUGCUGCAU

>mdm-miR10996b MIMAT0043622

UCACCAUUGCAUCUCAUGUUCC

>mdm-miR5225a MIMAT0026056

UCUGUCGAAGGUGAGAUGGUGC

>mdm-miR169j MIMAT0043578

CAGCCAAGGAUGACUUGCCGG

>mdm-miR156f MIMAT0025872

UGACAGAAGAGAGUGAGCAC

>mdm-miR156x MIMAT0025890

UGACAGAAGAUAGAGAGCAC

>mdm-miR399a MIMAT0026001

UGCCAAAGGAGAAUUGCCCUG

>mdm-miR156j MIMAT0025876

UGACAGAAGAGAGUGAGCAC

>mdm-miR408b MIMAT0026031

ACAGGGAAGAGGUAGAGCAUG

>mdm-miR171j MIMAT0025947

UUGAGCCGCGCCAAUAUCACU

>mdm-miR397b MIMAT0025997

UUGAGUGCAGCGUUGAUGAAA

>mdm-miR172c MIMAT0025954

AGAAUCUUGAUGAUGCUGCA

>mdm-miR156q MIMAT0025883

CUGACAGAAGAUAGAGAGCAC

>mdm-miR166h MIMAT0025920

UCGGACCAGGCUUCAUUCCCC

>mdm-miR390f MIMAT0025974

AAGCUCAGGAGGGAUAGCGCC

>mdm-miR10979 MIMAT0043529

CUUGCCGAUAGAUUUGGGGAG

>mdm-miR319h MIMAT0043595

GAGCUCUUCUUCAGUCCAGUCC

>mdm-miR168b MIMAT0025933

UCGCUUGGUGCAGGUCGGGAA

>mdm-miR530b MIMAT0043543

UGCAUUUGCACCUGCACUUGU

>mdm-miR166e MIMAT0025917

UCGGACCAGGCUUCAUUCCCC

>mdm-miR482b MIMAT0026022

UCUUUCCUAUCCCUCCCAUUCC

>mdm-miR11017 MIMAT0043627

CUCUAAUCUGCCCAACAACUU

>mdm-miR403b MIMAT0026012

UUAGAUUCACGCACAAACUCG

>mdm-miR171b MIMAT0025939

UUGAGCCGCGUCAAUAUCUCC

>mdm-miR10989a MIMAT0043555

CAAAGCUUUUAAUAUCAGUCGA

>mdm-miR10983 MIMAT0043546

CAGAGCAAAACAGUCGUGGAA

>mdm-miR11018 MIMAT0043628

CCAACAUCAAAAGUAAAAGGAA

>mdm-miR11015 MIMAT0043624

UGGGUAAAAUGACCCAACCUG

>mdm-miR399e MIMAT0026005

UGCCAAAGGAGAUUUGCUCGG

>mdm-miR10978a MIMAT0043527

GUUGGGAAUCGAAGCAUCACGA

>mdm-miR319g MIMAT0043569

GAGCUUUCUUCAGUCCACUC

>mdm-miR10981d MIMAT0043594

AGCCGUUUAAUCAAAAUCCAA

>mdm-miR11011b MIMAT0043630

UAAGUUCAUCCAAACACCAUA

>mdm-miR319c-3p MIMAT0026058

AUCCAACGAAGCAGGAGCUGA

>mdm-miR319b-5p MIMAT0037468

GAGCUUUCUUCAGUCCACUC

>mdm-miR396b MIMAT0025990

UUCCACAGCUUUCUUGAACUG

>mdm-miR482d MIMAT0026039

AAUGGAAGGGUAGGAAAGAAG

>mdm-miR169a MIMAT0025934

CAGCCAAGGAUGACUUGCCGG

>mdm-miR393h MIMAT0043584

AUCAUGCUAUCCCUUUGGAUU

>mdm-miR167a MIMAT0025922

AGAUCAUCUGGCAGUUUCACC

>mdm-miR169n MIMAT0043619

UAGCCAAGGAUGACUUGCCUG

>mdm-miR399i MIMAT0026009

UGCCAAAGGAGAGUUGCCCUG

>mdm-miR11006 MIMAT0043606

CAAUGGGGAGGAGUCAUUCGUA

>mdm-miR10987 MIMAT0043553

CCAUAUGUCCCUCCAUAUACU

>mdm-miR10984b-5p MIMAT0043609

GGUAAUUGACUGUGAAAUCGU

>mdm-miR477b MIMAT0026020

ACUCUCCCUCAAGGGCUUCGAC

>mdm-miR159e MIMAT0043540

UUUGGAUUGAAGGGAGCUCUA

>mdm-miR156c MIMAT0025869

UGACAGAAGAGAGUGAGCAC

>mdm-miR164a MIMAT0025907

UGGAGAAGCAGGGCACAUGCC

>mdm-miR535c MIMAT0026026

UGACAAGGAGAGAGAGCACGC

>mdm-miR395g-3p MIMAT0025986

CUGAAGUGUUUGGGGGAACUC

>mdm-miR3627c MIMAT0026018

UCGCAGGAGAGAUGGCACUA

>mdm-miR11005 MIMAT0043604

CUACUAUAUGGUCGUACACAUC

>mdm-miR858 MIMAT0026070

UUCGUUGUCUGUUCGACCUGA

>mdm-miR7121g MIMAT0026046

UCCUCUUGGUGAUCGCCCUGC

>mdm-miR10993a MIMAT0043570

AUCCCACCAUUUAUAUAGCGA

>mdm-miR156g MIMAT0025873

UGACAGAAGAGAGUGAGCAC

>mdm-miR172m MIMAT0025964

AGAAUCUUGAUGAUGCUGCAG

>mdm-miR166a MIMAT0025913

UCGGACCAGGCUUCAUUCCCC

>mdm-miR395e MIMAT0025984

CUGAAGUGUUUGGGGGAACUC

>mdm-miR166i MIMAT0025921

UCGGACCAGGCUUCAUUCCCC

>mdm-miR7121c MIMAT0026042

UCCUCUUGGUGAUCGCCCUGU

>mdm-miR11004 MIMAT0043603

GUAUUCUUUCAUCUUCUACUA

>mdm-miR395a MIMAT0025980

CUGAAGUGUUUGGGGGAACUC

>mdm-miR167j MIMAT0025931

UGAAGCUGCCAGCAUGAUCUUA

>mdm-miR172o MIMAT0025966

AGAAUCUUGAUGAUGCUGCAG

>mdm-miR7125 MIMAT0026059

CGAACUUAUUGCAACUAGCUU

>mdm-miR171o MIMAT0026066

UGGGAUGUUGGUAUGGUUCAA

>mdm-miR169i MIMAT0043577

CAGCCAAGGAUGACUUGCCGG

>mdm-miR160a MIMAT0025900

UGCCUGGCUCCCUGUAUGCCA

>mdm-miR2111b MIMAT0026015

UAAUCUGCAUCCUGAGGUUUA

>mdm-miR10981b MIMAT0043533

UGACCAACAUAUAUGGGCCGU

>mdm-miR10996a MIMAT0043580

ACACCAUCGCAUCUCAUGUUCC

>mdm-miR167e MIMAT0025926

UGAAGCUGCCAGCAUGAUCUA

>mdm-miR10989e MIMAT0043559

CAAAGCUUUUAAUAUCAGUCGA

>mdm-miR10991a MIMAT0043561

CGAGCCAUUGAAAUUCGAUCC

>mdm-miR319c-5p MIMAT0037474

GAGCUCUUCUUCAGUCCAGUCC

>mdm-miR482a-5p MIMAT0011164

AGGAAUGGGCUGUUUGGGAAGA

>mdm-miR398a MIMAT0025998

UGUGUUCUCAGGUCACCCCUU

>mdm-miR11013 MIMAT0043621

CAUGGGAAUUUUAAAGUCACCU

>mdm-miR7120a-3p MIMAT0037472

CAGUCUGACAAUAUAACGUGC

>mdm-miR172j MIMAT0025961

GGAAUCUUGAUGAUGCUGCAU

>mdm-miR171k MIMAT0025948

UUGAGCCGCGCCAAUAUCACU

>mdm-miR394a MIMAT0025978

UUGGCAUUCUGUCCACCUCC

>mdm-miR10991d MIMAT0043564

CGAGCCAUUGAAAUUCGAUCC

>mdm-miR164e MIMAT0025911

UGGAGAAGCAGGGCACGUGCA

>mdm-miR396f MIMAT0025994

UUCCACGGCUUUCUUGAACUG

>mdm-miR395l MIMAT0043626

CUGAAGUGUUUGGGGGAACCC

>mdm-miR172b MIMAT0025953

AGAAUCUUGAUGAUGCUGCA

>mdm-miR393c MIMAT0025977

UCCAAAGGGAUCGCAUUGAUCU

>mdm-miR166f MIMAT0025918

UCGGACCAGGCUUCAUUCCCC

>mdm-miR156ab MIMAT0025894

UUGACAGAAGAUAGAGAGCAC

>mdm-miR399h MIMAT0026008

UGCCAAAGGAGAUUUGCUCGG

>mdm-miR156u MIMAT0025887

UUGACAGAAGAGAGAGAGCAC

>mdm-miR397a MIMAT0025996

UUGAGUGCAGCGUUGAUGAAA

>mdm-miR156a MIMAT0025867

UGACAGAAGAGAGUGAGCAC

>mdm-miR535d MIMAT0026027

UGACGACGAGAGAGAGCACGC

>mdm-miR7120b-5p MIMAT0026038

UGUUAUAUUGUCAGAUUGUCA

>mdm-miR156y MIMAT0025891

UGACAGAAGAUAGAGAGCAC

>mdm-miR7126-5p MIMAT0026060

AAAGUAUCAAGGAGCGCAAAG

>mdm-miR7127b MIMAT0026065

AUACUCAUCGAAUUUGUCAUA

>mdm-miR167i MIMAT0025930

UGAAGCUGCCAGCAUGAUCUUA

>mdm-miR7128 MIMAT0026069

AUCAUUAACACUUAAUAACGA

>mdm-miR408a MIMAT0026013

AUGCACUGCCUCUUCCCUGGC

>mdm-miR11010 MIMAT0043612

AGUAACUAUAGCUGUUUUCUA

>mdm-miR168a MIMAT0025932

UCGCUUGGUGCAGGUCGGGAA

>mdm-miR167d MIMAT0025925

UGAAGCUGCCAGCAUGAUCUA

>mdm-miR162a MIMAT0025905

UCGAUAAACCUCUGCAUCCAG

>mdm-miR171c MIMAT0025940

UGAUUGAGCCGCGCCAAUAUC

>mdm-miR1511 MIMAT0026188

ACCUAGCUCUGAUACCAUGAA

>mdm-miR477a MIMAT0026021

ACUCUCCCUCAAGAGCUUCUC

>mdm-miR11003 MIMAT0043602

ACACAAUAUACGAUGAACAGA

>mdm-miR172a MIMAT0025952

AGAAUCUUGAUGAUGCUGCA

>mdm-miR7121b MIMAT0026041

UCCUCUUGGUGAUCGCCCUGU

>mdm-miR10989b MIMAT0043556

CAAAGCUUUUAAUAUCAGUCGA

>mdm-miR156v MIMAT0025888

UUGACAGAAGAGAGAGAGCAC

>mdm-miR166g MIMAT0025919

UCGGACCAGGCUUCAUUCCCC

>mdm-miR399b MIMAT0026002

UGCCAAAGGAGAAUUGCCCUG

>mdm-miR7122a MIMAT0026048

UUAUACAGAGAAAUCACGGUCG

>mdm-miR2111a MIMAT0026014

UAAUCUGCAUCCUGAGGUUUA

>mdm-miR10981c MIMAT0043593

AGCCGUUUAAUCAAAAUCCAA

>mdm-miR319f MIMAT0043568

GAGCUUUCUUCAGUCCACUC

>mdm-miR10991b MIMAT0043562

CGAGCCAUUGAAAUUCGAUCC

>mdm-miR160c MIMAT0025902

UGCCUGGCUCCCUGUAUGCCA

>mdm-miR10982a MIMAT0043535

CGGAAUGAAGCUUACGAGAAUG

>mdm-miR11002c-3p MIMAT0043597

GAGGAUGAGCUUCGGCGGUGA

>mdm-miR159f MIMAT0043541

UUUGGAUUGAAGGGAGCUCUA

>mdm-miR166c MIMAT0025915

UCGGACCAGGCUUCAUUCCCC

>mdm-miR164b MIMAT0025908

UGGAGAAGCAGGGCACGUGCA

>mdm-miR319b-3p MIMAT0025968

UUGGACUGAAGGGAGCUCCCU

>mdm-miR396c MIMAT0025991

UUCCACAGCUUUCUUGAACUU

>mdm-miR156d MIMAT0025870

UGACAGAAGAGAGUGAGCAC

>mdm-miR156w MIMAT0025889

UUGACAGAAGAGAGAGAGCAC

>mdm-miR395i-3p MIMAT0025988

CUGAAGUGUUUGGGGGAACUC

>mdm-miR171g MIMAT0025944

UGAUUGAGCCGUGCCAAUAUC

>mdm-miR171a MIMAT0025938

UUGAGCCGCGUCAAUAUCUCC

>mdm-miR399f MIMAT0026006

UGCCAAAGGAGAUUUGCUCGG

>mdm-miR828a MIMAT0026029

UCUUGCUCAAAUGAGUAUUCCA

>mdm-miR166j MIMAT0043545

UCGGACCAGGCUUCAUUCCCC

>mdm-miR171p MIMAT0043582

UUGAGCCGCGUCAAUAUCUCC

>mdm-miR10980a MIMAT0043530

CACCUGGGACUUGCAGCCAUG

>mdm-miR7121f MIMAT0026045

UCCUCUUGGUGAUCGCCCUGC

>mdm-miR169h MIMAT0043576

CAGCCAAGGAUGACUUGCCGG

>mdm-miR10993c MIMAT0043587

ACAUGUGGUGUACCAUCCUGU

>mdm-miR2118a MIMAT0026034

CUACCGAUGCCACUAAGUCCCA

>mdm-miR10988 MIMAT0043554

AGAGAAAAUUCAUUCCAACGC

>mdm-miR156z MIMAT0025892

UGACAGAAGAUAGAGAGCAC

>mdm-miR391 MIMAT0026019

UACGCAGGAGAGAUGACGCCG

>mdm-miR171l MIMAT0025949

UUGAGCCGCGCCAAUAUCACU

>mdm-miR7126-3p MIMAT0037475

UUGCGUUCCACUGAUUCUUUCG

>mdm-miR167h MIMAT0025929

UGAAGCUGCCAGCAUGAUCUUA

>mdm-miR11016 MIMAT0043625

UCCAUAAUUUUUCCAGAUCAA

>mdm-miR10992 MIMAT0043566

CAUAACAAAUUAUUACUCAGU

>mdm-miR394b MIMAT0025979

UUGGCAUUCUGUCCACCUCC

>mdm-miR5225c MIMAT0026052

UCUGUCGUGGGUGAGAUGGUGC

>mdm-miR7123a MIMAT0026050

AAGAGCGGGAUGUGUAAAAGG

>mdm-miR11002c-5p MIMAT0043596

CCCAGUCACCUCCGAAGCUCA

>mdm-miR393b MIMAT0025976

UCCAAAGGGAUCGCAUUGAUCU

>mdm-miR395g-5p MIMAT0037470

GUUCCCUUGACCACUUCAUUG

>mdm-miR482c MIMAT0026023

UCUUUCCUAACCCUCCCAUUCC

>mdm-miR172n MIMAT0025965

AGAAUCUUGAUGAUGCUGCAG

>mdm-miR164f MIMAT0025912

UGGAGAAGCAGGGCACGUGCA

>mdm-miR172h MIMAT0025959

AGAAUCUUGAUGAUGCUGCAU

>mdm-miR10995 MIMAT0043579

CAAGCUUCCUCUUCAUACUCGU

>mdm-miR156aa MIMAT0025893

UGACAGAAGAUAGAGAGCAC

>mdm-miR10993d MIMAT0043588

ACAUGUGGUGUACCAUCCUGU

>mdm-miR171f-5p MIMAT0037467

GGAUAUUGGUCCGGUUCAAUA

>mdm-miR156o MIMAT0025881

UGACAGAAGAGAGUGAGCAC

>mdm-miR160e MIMAT0025904

UGCCUGGCUCCCUGUAUGCCA

>mdm-miR10994-5p MIMAT0043572

UAUGGUUAAGAAACAAGCAGA

>mdm-miR393e MIMAT0026062

AUCAUGCGAUCCCUUCGGACG

>mdm-miR160b MIMAT0025901

UGCCUGGCUCCCUGUAUGCCA

>mdm-miR162b MIMAT0025906

UCGAUAAACCUCUGCAUCCAG

>mdm-miR398b MIMAT0025999

UGUGUUCUCAGGUCGCCCCUG

>mdm-miR169l MIMAT0043617

UAGCCAAGGAUGACUUGCCUG

>mdm-miR156k MIMAT0025877

UGACAGAAGAGAGUGAGCAC

>mdm-miR11008 MIMAT0043608

GUGACCGCACAAAAUAGAAGA

>mdm-miR10982b MIMAT0043536

CGGAAUGAAGCUUACGAGAAUG

>mdm-miR172k MIMAT0025962

GGAAUCUUGAUGAUGCUGCAU

>mdm-miR7122b MIMAT0026049

UUAUACAGAGAAAUCACGGUCG

>mdm-miR169e MIMAT0026067

UGAAGAGAAGAGCGUUGUUUGG

>mdm-miR171d MIMAT0025941

UGAUUGAGCCGCGCCAAUAUC

>mdm-miR10984a-3p MIMAT0043548

AGUCAAUUACCUCAUAAACUC

>mdm-miR159b MIMAT0025899

CUUGGAUUGAAGGGAGCUCC

>mdm-miR5225b MIMAT0026057

UCUGUCGAAGGUGAGAUGGUGC

>mdm-miR3627a MIMAT0026016

UCGCAGGAGAGAUGGCACUA

>mdm-miR171h MIMAT0025945

UGAUUGAGCCGUGCCAAUAUC

>mdm-miR169m MIMAT0043618

UAGCCAAGGAUGACUUGCCUG

>mdm-miR395h MIMAT0025987

CUGAAGUGUUUGGGGGAACUC

>mdm-miR7121a MIMAT0026040

UCCUCUUGGUGAUCGCCCUGU

>mdm-miR10985 MIMAT0043549

CCACUCGUAGUGAAACAGUUG

>mdm-miR390b MIMAT0025970

AAGCUCAGGAGGGAUAGCGCC

>mdm-miR172e MIMAT0025956

AGAAUCUUGAUGAUGCUGCAU

>mdm-miR11019 MIMAT0043629

CCAGAUGAUCAUAAUCUCCUGA

>mdm-miR10982d MIMAT0043538

CGGAAUGAAGCUUACGAGAAUG

>mdm-miR7120b-3p MIMAT0037473

CAGUCUGACAAUAUAACGUGC

>mdm-miR395d-3p MIMAT0025983

CUGAAGUGUUUGGGGGAACUC

>mdm-miR399j MIMAT0026010

UGCCAAAGGAGAGUUGCCCUG

>mdm-miR169d MIMAT0025937

UAGCCAAGGAUGACUUGCCCG

>mdm-miR7123b MIMAT0026051

AAGAGCGGGAUGUGUAAAAGG

>mdm-miR393a MIMAT0025975

UCCAAAGGGAUCGCAUUGAUCU

>mdm-miR10998 MIMAT0043585

CUUGGGAUUCAGUCUAGGACUU

>mdm-miR7120a-5p MIMAT0026037

UGUUAUAUUGUCAGAUUGUCA

>nta-miR159 MIMAT0024639

UUUGGAUUGAAGGGAGCUCUA

>nta-miR477a MIMAT0024721

ACUCUCCCUCAAGGGCUUCUG

>nta-miR319b MIMAT0024700

UUGGACUGAAGGGAGCUCCCU

>nta-miR156f MIMAT0024634

UGACAGAAGAGAAUGAGCAC

>nta-miR169c MIMAT0024669

CAGCCAAGGAUGACUUGCCGA

>nta-miR408 MIMAT0024720

UGCACUGCCUCUUCCCUGGCU

>nta-miR6021 MIMAT0023588

UUGGAAGAGGCUGCUAUUGGA

>nta-miR168d MIMAT0024665

UCGCUUGGUGCAGGUCGGGAA

>nta-miR169d MIMAT0024670

CAGCCAAGGAUGACUUGCCGA

>nta-miR167b MIMAT0024658

UGAAGCUGCCAGCAUGAUCUGG

>nta-miR390b MIMAT0024702

AAGCUCAGGAGGGAUAGCGCC

>nta-miR6151c MIMAT0024744

UGAAUGUGAGGCAUUGGAUUGA

>nta-miR1446 MIMAT0024763

UGAACUCUCUCCCUCAAUGGCU

>nta-miR482c MIMAT0024727

UUUCCAAUUCCACCCAUUCCUA

>nta-miR396b MIMAT0024709

UUCCACAGCUUUCUUGAACUU

>nta-miR167d MIMAT0024660

UGAAGCUGCCAGCAUGAUCUA

>nta-miR172g MIMAT0024695

AGAAUCUUGAUGAUGCUGCAU

>nta-miR168b MIMAT0024663

UCGCUUGGUGCAGGUCGGGAC

>nta-miR399e MIMAT0024717

CGCCAAAGGAGAGCUGCCCUG

>nta-miR6151i MIMAT0024750

UGAGUGUGAGGCGUUGGAUUGA

>nta-miR162b MIMAT0024645

UCGAUAAACCUCUGCAUCCAG

>nta-miR171a MIMAT0024686

UAUUGGUGCGGUUCAAUGAGA

>nta-miR6158a MIMAT0024766

AAGUUCGAUUUGUACGAAGGGC

>nta-miR6155 MIMAT0024757

UAAGGUUGCCUUGCUCUUGCA

>nta-miR172i MIMAT0024697

AGAAUCUUGAUGAUGCUGCAU

>nta-miR172e MIMAT0024693

AGAAUCUUGAUGAUGCUGCAU

>nta-miR6148b MIMAT0024738

UGUGUUAAUCGUUUGUUCUCA

>nta-miR6154b MIMAT0024756

UGGGUCUCCUGGAGAAAGGUC

>nta-miR166c MIMAT0024651

UCGGACCAGGCUUCAUUCCCC

>nta-miR6147 MIMAT0024736

UGACAUCUUCAAAACCCACUA

>nta-miR6025c MIMAT0024730

UCAAUUGAGAUGACAUCUAGU

>nta-miR394 MIMAT0024704

UUGGCAUUCUGUCCACCUCC

>nta-miR169f MIMAT0024672

CAGCCAAGGAUGACUUGCCGA

>nta-miR156a MIMAT0024629

UGACAGAAGAGAGUGAGCAC

>nta-miR169h MIMAT0024674

CAGCCAAGGAUGACUUGCCGA

>nta-miR6157 MIMAT0024760

UGGUAGACGUAGGAUUUGAAGA

>nta-miR6161b MIMAT0024775

GCUGGACCGGUAUACUUUGCUGAC

>nta-miR169a MIMAT0024667

CAGCCAAGGAUGACUUGCCGA

>nta-miR6152a MIMAT0024751

UAUUGUAUUCGACUGUAUUCACGG

>nta-miR6020a-3p MIMAT0023585

AGAUACUCAGCAAAACAUUUAC

>nta-miR6024 MIMAT0023595

UUUUAGCCAGAGUUGUUUUCCC

>nta-miR397 MIMAT0024711

AUUGAGUGCAGCGUUGAUGU

>nta-miR395a MIMAT0024705

CUGAAGUGUUUGGGGGAACUC

>nta-miR6145c MIMAT0024758

CAGUGCACAUAUAACAGUAA

>nta-miR168a MIMAT0024662

UCGCUUGGUGCAGGUCGGGAC

>nta-miR399b MIMAT0024714

CGCCAAAGGAGAGCUGCCCUG

>nta-miR6158c MIMAT0024768

AAGUUCGAUUUGUACGAAGGGC

>nta-miR169j MIMAT0024676

CAGCCAAGGAUGACUUGCCGA

>nta-miR395c MIMAT0024707

CUGAAGUGUUUGGGGGAACUC

>nta-miR156b MIMAT0024630

UGACAGAAGAGAGUGAGCAC

>nta-miR172f MIMAT0024694

AGAAUCUUGAUGAUGCUGCAU

>nta-miR6025a MIMAT0023606

UACCAACAAUUGAGAUAACAUC

>nta-miR6025e MIMAT0024733

UGCCAAUUAUAGAGAUGACAUC

>nta-miR169t MIMAT0024685

UAGCCAAGGAUGACUUGCCUU

>nta-miR169r MIMAT0024683

CAGCCAAGGAUGACUUGCCGG

>nta-miR6151g MIMAT0024748

UGAGUGUGAGGCGUUGGAUUGA

>nta-miR6145d MIMAT0024764

AUUGUUACAUGUAACACUGGC

>nta-miR6146b MIMAT0024735

UUUGUCCAAUGAAAUACUUAUC

>nta-miR399g MIMAT0024719

CGCCAAAGGAGAGCUGCCCUG

>nta-miR6148a MIMAT0024737

UACGUCGAUCGAUUGUUCUUA

>nta-miR164a MIMAT0024646

UGGAGAAGCAGGGCACGUGCA

>nta-miR166g MIMAT0024655

UCGGACCAGGCUUCAUUCCCC

>nta-miR396c MIMAT0024710

UUCCACAGCUUUCUUGAACUU

>nta-miR6163 MIMAT0024779

UGGAAGUACUGCCUAAGUUUGA

>nta-miR6019a MIMAT0023583

UACAGGUGACUUGUAAAUGUUU

>nta-miR399a MIMAT0024713

CGCCAAAGGAGAGCUGCCCUG

>nta-miR482a MIMAT0023599

UUUCCAAUUCCACCCAUUCCUA

>nta-miR156e MIMAT0024633

UGACAGAAGAGAGUGAGCAC

>nta-miR156h MIMAT0024636

UGACAGAAGAUAGAGAGCAC

>nta-miR160a MIMAT0024640

UGCCUGGCUCCCUGUAUGCCA

>nta-miR482d MIMAT0024776

UUCCCGACUCCCCCCAUACCAC

>nta-miR6161d MIMAT0024773

UGAACUCCAGCAUAUUAUACU

>nta-miR6149b MIMAT0024740

UUGAUACGCACCUGAAUCGGC

>nta-miR479a MIMAT0024723

CGUGAUAUUGGUUUGGCUCAUC

>nta-miR169m MIMAT0024679

CAGCCAAGGAUGACUUGCCGA

>nta-miR6150 MIMAT0024741

AGAUUUGUUUGAUCGUCUUGGC

>nta-miR6144 MIMAT0024729

UGGCAACUUCUUCAUCAUGCC

>nta-miR166e MIMAT0024653

UCGGACCAGGCUUCAUUCCCC

>nta-miR6152b MIMAT0024752

UAUUGUAUUCGACUGUAUUCACGG

>nta-miR399d MIMAT0024716

CGCCAAAGGAGAGCUGCCCUG

>nta-miR6160 MIMAT0024772

GCAUAUAUGGGCCAACUGUGUAAC

>nta-miR156d MIMAT0024632

UGACAGAAGAGAGUGAGCAC

>nta-miR166a MIMAT0024649

UCGGACCAGGCUUCAUUCCCC

>nta-miR162a MIMAT0024644

UCGAUAAACCUCUGCAUCCAG

>nta-miR172h MIMAT0024696

AGAAUCUUGAUGAUGCUGCAU

>nta-miR169o MIMAT0024680

CAGCCAAGGAUGACUUGCCGA

>nta-miR390c MIMAT0024703

AAGCUCAGGAGGGAUAGCGCC

>nta-miR6020b MIMAT0023587

AAAUGUUCUUCGAGUAUCUUC

>nta-miR5303c MIMAT0024770

ACGGGUGCGGCUACAUUUUGG

>nta-miR6151e MIMAT0024746

UGAAUGUGAGGCAUUGGAUUGA

>nta-miR169p MIMAT0024681

CAGCCAAGGAUGACUUGCCGA

>nta-miR169b MIMAT0024668

CAGCCAAGGAUGACUUGCCGA

>nta-miR6161a MIMAT0024774

GCUGGACCGGUAUACUUUGCUGAC

>nta-miR172b MIMAT0024690

AGAAUCAUGAUGAUGCUGCAU

>nta-miR168e MIMAT0024666

UCGCUUGGUGCAGGUCGGGAA

>nta-miR319a MIMAT0024699

UUGGACUGAAGGGAGCUCCCU

>nta-miR156g MIMAT0024635

UGACAGAAGAUAGAGAGCAC

>nta-miR6145f MIMAT0024771

AUCGUAACAUAUAGCACUAGC

>nta-miR6156 MIMAT0024759

UUGAAGAUGUUCUAUUUCUGU

>nta-miR169e MIMAT0024671

CAGCCAAGGAUGACUUGCCGA

>nta-miR6162 MIMAT0024777

AGAAAAAUGGUAGCCAUUGGA

>nta-miR171b MIMAT0024687

UUGAGCCGCGCCAAUAUCACU

>nta-miR6164a MIMAT0024780

UCACAUAAAUUGAAACGGAGG

>nta-miR167c MIMAT0024659

UGAAGCUGCCAGCAUGAUCUGG

>nta-miR6158b MIMAT0024767

AAGUUCGAUUUGUACGAAGGGC

>nta-miR6145b MIMAT0024753

UUAUCAUACGUAGCACUAGCC

>nta-miR166f MIMAT0024654

UCGGACCAGGCUUCAUUCCCC

>nta-miR169g MIMAT0024673

CAGCCAAGGAUGACUUGCCGA

>nta-miR398 MIMAT0024712

UGUGUUCUCAGGUCGCCCCUG

>nta-miR168c MIMAT0024664

UCGCUUGGUGCAGGUCGGGAC

>nta-miR5303a MIMAT0024761

AAAAUGUGGCCGGAUACGUGU

>nta-miR399f MIMAT0024718

CGCCAAAGGAGAGCUGCCCUG

>nta-miR395b MIMAT0024706

CUGAAGUGUUUGGGGGAACUC

>nta-miR482b-3p MIMAT0023605

UCUUGCCAAUGCCAUCCAUUCC

>nta-miR169k MIMAT0024677

CAGCCAAGGAUGACUUGCCGA

>nta-miR6145a MIMAT0024731

CAUUUUCACAUGUAGCACUGAC

>nta-miR169l MIMAT0024678

CAGCCAAGGAUGACUUGCCGA

>nta-miR160d MIMAT0024643

UGCCUGGCUCCCUGCAUGCCA

>nta-miR169i MIMAT0024675

CAGCCAAGGAUGACUUGCCGA

>nta-miR6151b MIMAT0024743

UGAAUGUGAGGCAUUGGAUUGA

>nta-miR164b MIMAT0024647

UGGAGAAGCAGGGCACGUGCA

>nta-miR167a MIMAT0024657

UGAAGCUGCCAGCAUGAUCUGG

>nta-miR169s MIMAT0024684

CAGCCAAGGAUGACUUGCCGG

>nta-miR166d MIMAT0024652

UCGGACCAGGCUUCAUUCCCC

>nta-miR156j MIMAT0024638

UGACAGAAGAUAGAGAGCAC

>nta-miR396a MIMAT0024708

UUCCACAGCUUUCUUGAACUG

>nta-miR477b MIMAT0024722

UCUCUCCCUCAAGGGCUUCUC

>nta-miR6025b MIMAT0023607

UGCCAACUAUUGAGAUGACAUC

>nta-miR479b MIMAT0024724

CGUGAUAUUUGUUUGGCUCAUC

>nta-miR6151d MIMAT0024745

UGAAUGUGAGGCAUUGGAUUGA

>nta-miR166b MIMAT0024650

UCGGACCAGGCUUCAUUCCCC

>nta-miR6151a MIMAT0024742

UGAAUGUGAGGCAUUGGAUUGA

>nta-miR172c MIMAT0024691

AGAAUCUUGAUGAUGCUGCAU

>nta-miR6161c MIMAT0024778

AAUAUACUGGAGUUCGGUGCACCU

>nta-miR6154a MIMAT0024755

UGGGUCUCCUGGAGAAAGGUC

>nta-miR6153 MIMAT0024754

UAGGACCAUAUUCACUAUUUG

>nta-miR169q MIMAT0024682

CAGCCAAGGAUGACUUGCCGG

>nta-miR6146a MIMAT0024734

UUUGUCCAAUGAAACACUUAUC

>nta-miR482b-5p MIMAT0023604

AGUGGGUGGAGUGGUAAGAUA

>nta-miR160c MIMAT0024642

UGCCUGGCUCCCUGUAUGCCA

>nta-miR172d MIMAT0024692

AGAAUCUUGAUGAUGCUGCAU

>nta-miR6151f MIMAT0024747

UGAGUGUGAGGCAUUGGAUUGA

>nta-miR167e MIMAT0024661

UGAAGCUGCCAGCAUGAUCUA

>nta-miR160b MIMAT0024641

UGCCUGGCUCCCUGUAUGCCA

>nta-miR399c MIMAT0024715

CGCCAAAGGAGAGCUGCCCUG

>nta-miR827 MIMAT0024725

UUAGAUGAACAUCAACAAACA

>nta-miR6159 MIMAT0024769

UAGCAUAGAAUUCUCGCACCUA

>nta-miR6151h MIMAT0024749

UGAGUGUGAGGCGUUGGAUUGA

>nta-miR6019b MIMAT0023586

UACAGGUGACUUGUAAAUGUUU

>nta-miR5303b MIMAT0024762

AAAAUGUGGCCGGAUACGUGU

>nta-miR156i MIMAT0024637

UGACAGAAGAUAGAGAGCAC

>nta-miR6149a MIMAT0024739

UUGAUACGCACCUGAAUCGGC

>nta-miR171c MIMAT0024688

UGAUUGAGCCGUGCCAAUAUC

>nta-miR390a MIMAT0024701

AAGCUCAGGAGGGAUAGCACC

>nta-miR6025d MIMAT0024732

AACAAUUGAGAUAACAUCUAGG

>nta-miR6020a-5p MIMAT0023584

AAAUGUUUUUCGAGUAUCUUC

>nta-miR6164b MIMAT0024781

UCACAUAAAUUGAAACGGAGG

>nta-miR164c MIMAT0024648

UGGAGAAGCAGGGCACAUGCU

>nta-miR166h MIMAT0024656

UCGGACCAGGCUUCAUUCCCC

>nta-miR1919 MIMAT0024726

GAGCGAGUCAUCUGUGACAGG

>nta-miR172j MIMAT0024698

GGAAUCUUGAUGAUGCUGCAU

>nta-miR172a MIMAT0024689

AGAAUCUUGAUGAUGCUGCAG

>nta-miR6145e MIMAT0024765

AUUGUUACAUGUAGCACUGGC

>nta-miR156c MIMAT0024631

UGACAGAAGAGAGUGAGCAC

>pla-miR11609 MIMAT0045462

UAGCUUGGUGUGAGGUCAACUU

>pla-miR11603 MIMAT0045456

UUAUAAUUAGGUUGAGCGGAC

>pla-miR11600 MIMAT0045453

AACGUUCCUCGAUUUCGCGAU

>pla-miR11612 MIMAT0045465

AAGACGGUCCAAAACGCCCAC

>pla-miR11598 MIMAT0045451

UCGUUCAAAGUAGGUUGUCAA

>pla-miR11608 MIMAT0045461

UCGCUUAGGGGUUGUUGAAGCGC

>pla-miR11601 MIMAT0045454

UGCUCUAAAAGAUCGUAGUUC

>pla-miR11610 MIMAT0045463

ACAGAUAUGGUAGGGGGCACA

>pla-miR11606 MIMAT0045459

UGGUGGACUCCAAUUCGCAUA

>pla-miR11599 MIMAT0045452

UUCAACCGUGGUAGAUGUUAA

>pla-miR11607 MIMAT0045460

GAUCACUCGGUUGUCUGACACAC

>pla-miR11604 MIMAT0045457

UCCAGAGGGAGAACGUGGCGA

>pla-miR11611 MIMAT0045464

UAGGCAACCGUGGUAAAAUGUC

>pla-miR11602 MIMAT0045455

UCUAACGGAACGCUAUUGGAUC

>pla-miR11605 MIMAT0045458

UUGAGGCGGCAUAUUCUCAAU

>peu-miR2916 MIMAT0012884

UGGGGACUCGAAGACGAUCAUAU

>peu-miR2912b MIMAT0012880

UCUAGAACUCGAGAUAUGGGC

>peu-miR2912a MIMAT0012879

UCUAGAACUCGAGAUAUGGGC

>peu-miR2913 MIMAT0012881

GAGGUCGGGGAUUGCAAGGAG

>ptc-miR475d-3p MIMAT0002070

UUACAGAGUCCAUUGAUUAAG

>ptc-miR156a MIMAT0001890

UGACAGAAGAGAGUGAGCAC

>ptc-miR1447 MIMAT0006009

CAGAAUUGCAGUGCCUUGAUU

>ptc-miR408-3p MIMAT0002058

AUGCACUGCCUCUUCCCUGGC

>ptc-miR482b-3p MIMAT0025271

UUACCAAUACCUCUCAUGCCAA

>ptc-miR169o MIMAT0001971

AAGCCAAGGAUGACUUGCCUG

>ptc-miR169b-3p MIMAT0022898

GGCAGGUUGUUCUUGGCUAC

>ptc-miR7834 MIMAT0030406

UAAUAAAAUCUCGACUAUUAU

>ptc-miR397c MIMAT0002040

UCAAUGAGUGGAGCUUUGAUG

>ptc-miR6434 MIMAT0025237

CUAUGUAUGCUUGACCAAUCA

>ptc-miR396d MIMAT0002034

UUCCACAGCUUUCUUGAACUU

>ptc-miR393c MIMAT0002017

UCCAAAGGGAUCGCAUUGAUC

>ptc-miR156l MIMAT0025268

UUGACAGAAGAUGGAGAGCAC

>ptc-miR172a MIMAT0001993

AGAAUCUUGAUGAUGCUGCAU

>ptc-miR167d MIMAT0001944

UGAAGCUGCCAGCAUGAUCUA

>ptc-miR481b MIMAT0002099

AGGACCUCACUUAACAGCUUAAGC

>ptc-miR169u-3p MIMAT0022900

GGCAGUCUCCUUUGGCUAUCC

>ptc-miR167h-3p MIMAT0022895

AGAUCAUGUGGCAGUUUCACC

>ptc-miR6467 MIMAT0025196

AUAAGCUGUCGAGCGUUUUGU

>ptc-miR482a.2 MIMAT0006785

UCUUGCCUACUCCUCCCAUU

>ptc-miR477e-5p MIMAT0002062

ACUCUCCCUCAAGGCUUCCA

>ptc-miR395a MIMAT0002021

CUGAAGGGUUUGGAGGAACUC

>ptc-miR6442 MIMAT0025245

CUGAACGGCUUGAAGGGCACG

>ptc-miR160e-3p MIMAT0022892

GCAUGAGGGGAGUCGAGCAGG

>ptc-miR169v MIMAT0001978

UAGCCAAGGAUGACUUGCCCA

>ptc-miR160h MIMAT0001914

UGCCUGGCUCCCUGCAUGCCA

>ptc-miR478l MIMAT0002086

UAACGUGUCUCCUAUUUUUAGGGA

>ptc-miR478f MIMAT0002081

UGACAUGUCUUCUAUUUUUAGGGA

>ptc-miR6430 MIMAT0025231

UGAUGAUUAAUUGACUGCAAA

>ptc-miR156g MIMAT0001896

UUGACAGAAGAUAGAGAGCAC

>ptc-miR162b MIMAT0001916

UCGAUAAACCUCUGCAUCCAG

>ptc-miR171l-5p MIMAT0002095

UGUGAUAUUGGUCCGGCUCAUC

>ptc-miR395e MIMAT0002025

CUGAAGUGUUUGGGGGAACUC

>ptc-miR160c-5p MIMAT0001909

UGCCUGGCUCCCUGUAUGCCA

>ptc-miR6437b MIMAT0025225

UCACGGACGGCGGCUCCAAGCA

>ptc-miR398c-3p MIMAT0002043

UGUGUUCUCAGGUCGCCCCUG

>ptc-miR164d MIMAT0001921

UGGAGAAGCAGGGCACGUGCA

>ptc-miR319a MIMAT0002002

UUGGACUGAAGGGAGCUCCC

>ptc-miR6428 MIMAT0025229

UCUGGCAACUCAUUAGACUCAU

>ptc-miR399b MIMAT0002045

UGCCAAAGGAGAUUUGCCCGG

>ptc-miR160c-3p MIMAT0022891

GCGUAUGAGGAGCCAUGCAUA

>ptc-miR171l-3p MIMAT0005994

CGAGCCGAAUCAAUAUCACU

>ptc-miR156b MIMAT0001891

UGACAGAAGAGAGUGAGCAC

>ptc-miR6478 MIMAT0025220

CCGACCUUAGCUCAGUUGGUG

>ptc-miR6421-3p MIMAT0025181

UAGAGCAGAUUGUAAGGGAAG

>ptc-miR169a MIMAT0001951

CAGCCAAGGAUGACUUGCCGA

>ptc-miR6422 MIMAT0025236

UGUGAUAAUGAAGGCUAUGGU

>ptc-miR172h-3p MIMAT0002000

GGAAUCUUGAUGAUGCUGCAG

>ptc-miR482d-3p MIMAT0025235

UUGCCGACCCCACCCAUGCCAA

>ptc-miR399e MIMAT0002055

CGCCAAAGGAGAGUUGCCCUC

>ptc-miR1444c MIMAT0006001

UUCACAUUCGGUCAACGUUC

>ptc-miR403c-5p MIMAT0022932

UUUGUGCGUGGAUCUGAGGCC

>ptc-miR166i MIMAT0001932

UCGGACCAGGCUUCAUUCCCC

>ptc-miR160b-5p MIMAT0001908

UGCCUGGCUCCCUGUAUGCCA

>ptc-miR171i MIMAT0001991

UGAUUGAGCCGUGCCAAUAUC

>ptc-miR6440c MIMAT0025200

GAGUUUGAUCGAUUUCGAGUU

>ptc-miR6459b MIMAT0030396

UCUCAAGCCCAAAUUCGAUC

>ptc-miR7817b MIMAT0030379

UCUCUUCUGUUCCUGAACGGU

>ptc-miR6429 MIMAT0025230

UAGUAGAAAUGCAUUGACUAG

>ptc-miR6438b MIMAT0025197

UUGUACACAGAAUAGGUGAAAU

>ptc-miR6461 MIMAT0025187

UAGCUAGCAAGUUCAUGGAUC

>ptc-miR319d MIMAT0002005

UUGGACUGAAGGGAGCUCCC

>ptc-miR169n-3p MIMAT0022899

GCAAGCAUCCUUGGUUCUCC

>ptc-miR172e MIMAT0001997

GGAAUCUUGAUGAUGCUGCAU

>ptc-miR169ac MIMAT0001954

UAGCCAAGGACGACUUGCCCA

>ptc-miR6474 MIMAT0025213

UGUUCAGAUCAGUAGAUAGCA

>ptc-miR393b-5p MIMAT0002016

UCCAAAGGGAUCGCAUUGAUC

>ptc-miR7821 MIMAT0030402

AGAUGGGCAUCGGCAUUGUGA

>ptc-miR169l MIMAT0001968

UAGCCAAGGAUGACUUGCCUG

>ptc-miR6423 MIMAT0025252

CCGCUGUCGCCACUAUCUUCCU

>ptc-miR398b MIMAT0002042

UGUGUUCUCAGGUCGCCCCUG

>ptc-miR478m MIMAT0002087

UAACGUGUCUCCUAUUUUUAGGGA

>ptc-miR398c-5p MIMAT0022912

GGAGCGACCUGAAAUCACAUG

>ptc-miR390a MIMAT0002011

AAGCUCAGGAGGGAUAGCGCC

>ptc-miR6476a MIMAT0025217

UCAGUGGAGAUGAAACAUGA

>ptc-miR6440d MIMAT0025226

GAGUUUGAUCGAUUUCGAGUU

>ptc-miR477d-3p MIMAT0025276

UGGACUCCUUUGGGGAGAUGG

>ptc-miR319e MIMAT0002006

UUGGACUGAAGGGAGCUCCU

>ptc-miR6425e MIMAT0025219

UUGUCUUCCAUGGAAUAGGCAG

>ptc-miR7815 MIMAT0030373

CUCUUCAAAUAAAUCGUGGGA

>ptc-miR7812 MIMAT0030371

CUGUUAUGAAUUGAUGGAGUG

>ptc-miR171m MIMAT0005995

CGAGCCGAAUCAAUAUCACU

>ptc-miR6447 MIMAT0025250

UUGACGAAAUGUGACGACUAC

>ptc-miR399i MIMAT0002052

UGCCAAAGGAGAGUUGCCCUA

>ptc-miR169ad MIMAT0001955

UAGCCAAGGACGACUUGCCCA

>ptc-miR1448 MIMAT0006010

CUUUCCAACGCCUCCCAUAC

>ptc-miR169h MIMAT0001964

CAGCCAAGGAUGACUUGCCGG

>ptc-miR1446d MIMAT0006007

UUCUGAACUCUCUCCCUCAA

>ptc-miR478s MIMAT0002093

UAACGUGUCUCCUAUUUUUAGGGA

>ptc-miR319g MIMAT0002008

UUGGACUGAAGGGAGCUCCU

>ptc-miR7829 MIMAT0030404

ACACAGAAACUCCAAGCCCAC

>ptc-miR171a-3p MIMAT0001983

UUGAGCCGUGCCAAUAUCACG

>ptc-miR478a MIMAT0002076

UGACGUGUCUUCUAUUUUUAGGGA

>ptc-miR159c MIMAT0001906

AUUGGAGUGAAGGGAGCUCGA

>ptc-miR7819 MIMAT0030401

UCUUGAGAACAUGAUGAAUCG

>ptc-miR477a-3p MIMAT0022918

GGAUGCCUUUGGGGGAGAUUG

>ptc-miR166d MIMAT0001927

UCGGACCAGGCUUCAUUCCCC

>ptc-miR171h-3p MIMAT0001990

UGAUUGAGCCGUGCCAAUAUC

>ptc-miR481d MIMAT0002102

AGGACCUCACCUAACAGCUUAAGC

>ptc-miR475a-3p MIMAT0002067

UUACAGUGCCCAUUGAUUAAG

>ptc-miR7835 MIMAT0030407

GAUGGGAUUUUUCGGGAAGUG

>ptc-miR6463 MIMAT0025189

UGGAUGAUCAUGUUGGCAACC

>ptc-miR1450 MIMAT0006012

UUCAAUGGCUCGGUCAGGUUAC

>ptc-miR403a MIMAT0002056

UUAGAUUCACGCACAAACUCG

>ptc-miR474b MIMAT0002065

CAAAAGUUGUUGGGUUUGGCUGGG

>ptc-miR166n MIMAT0001937

UCGGACCAGGCUUCAUUCCUU

>ptc-miR394a-5p MIMAT0002019

UUGGCAUUCUGUCCACCUCC

>ptc-miR169r MIMAT0001974

UAGCCAAGGAUGACUUGCCUA

>ptc-miR482c-5p MIMAT0025277

UAUGGGAGAGGCGGGAAUGACU

>ptc-miR477f MIMAT0002063

GCUCUCCCUCAGGGCUUCCA

>ptc-miR480 MIMAT0002096

ACUACUACAUCAUUGACGUUGAAC

>ptc-miR169d MIMAT0001960

CAGCCAAGGAUGACUUGCCGG

>ptc-miR6457b MIMAT0025203

UUAGUUUGGCAGCCUCUUCUC

>ptc-miR394b-3p MIMAT0006784

CUGUUGGUCUCUCUUUGUAA

>ptc-miR169n-5p MIMAT0001970

UGAGCCAAGGAUGACUUGCCG

>ptc-miR6436 MIMAT0025239

CCAGACUCAAUAGCAGGACCCA

>ptc-miR156h MIMAT0001897

UUGACAGAAGAUAGAGAGCAC

>ptc-miR7817a MIMAT0030377

UUUGGUUAUUGUCUCGAGACA

>ptc-miR156k MIMAT0001900

UGACAGAAGAGAGGGAGCAC

>ptc-miR6427-3p MIMAT0025228

GUGGGAAUGAACAUUAUGAGA

>ptc-miR1449 MIMAT0006011

UGAGGUGCACGUAAGAUAACUC

>ptc-miR396a MIMAT0002031

UUCCACAGCUUUCUUGAACUG

>ptc-miR408-5p MIMAT0022913

CGGGGAACAGGCAGAGCAUGG

>ptc-miR160a MIMAT0001907

UGCCUGGCUCCCUGUAUGCCA

>ptc-miR403c-3p MIMAT0003944

UUAGAUUCACGCACAAACUCG

>ptc-miR172g-5p MIMAT0022905

GGAGCAUCAUCAAGAUUCACA

>ptc-miR6462c-3p MIMAT0025206

UCUCUUAUGCAUUUUUGUCCC

>ptc-miR164c MIMAT0001920

UGGAGAAGCAGGGCACGUGCA

>ptc-miR390c MIMAT0002013

AAGCUCAGGAGGGAUAGCGCC

>ptc-miR169z MIMAT0001982

CAGCCAAGAAUGAUUUGCCGG

>ptc-miR1444e MIMAT0030386

CGAACGUUGACCGAAUGUGAA

>ptc-miR7830 MIMAT0030389

UGAUCUAGAGAACCGUUGCU

>ptc-miR6468-5p MIMAT0027337

GUUUUCCCUGAAUCACUCCCA

>ptc-miR6421-5p MIMAT0025180

UCCCUUACAAUCUACUCUUUC

>ptc-miR166o MIMAT0001938

UCGGACCAGGCUUCAUUCCUU

>ptc-miR167e MIMAT0001945

UGAAGCUGCCAGCAUGAUCUG

>ptc-miR1444b MIMAT0006000

UUCACAUUCGGUCAACGUUC

>ptc-miR6472 MIMAT0025211

UAGUGAAUUCUAGGUCUCAAUC

>ptc-miR475d-5p MIMAT0022917

AAUGGCCAUUGUAAGAGUAGA

>ptc-miR475b-5p MIMAT0022916

AAUGGCCAUUGUAAGAGUAGA

>ptc-miR390d-5p MIMAT0002014

AAGCUCAGGAGGGAUAGCGCC

>ptc-miR477c MIMAT0025257

GGAAACCUUUUGUGGGGGUUUG

>ptc-miR481c MIMAT0002100

AGGACCUCACUUAACAGCUUAAGC

>ptc-miR395d MIMAT0002024

CUGAAGUGUUUGGGGGAACUC

>ptc-miR6457a MIMAT0025267

UAAUCUCUCUGCAGAAUGCUG

>ptc-miR6451 MIMAT0025255

AUGUCAGAUCAUGUUAGGUAU

>ptc-miR476b MIMAT0002072

UAGUAAUUCUUCUUUGCAAAA

>ptc-miR166j MIMAT0001933

UCGGACCAGGCUUCAUUCCCC

>ptc-miR390d-3p MIMAT0022907

CGCUAUCCAUCCUGAGUUUUA

>ptc-miR172b-5p MIMAT0022904

GGAGCAUCAUCAAGAUUCACA

>ptc-miR1446a MIMAT0006004

UUCUGAACUCUCUCCCUCAA

>ptc-miR7828 MIMAT0030388

GAUGACAUGGACACCAAAAUC

>ptc-miR171j MIMAT0005996

CGAGCCGAAUCAAUAUCACU

>ptc-miR7839 MIMAT0030395

AGUGGCAUUGGAGGUAUCCC

>ptc-miR156c MIMAT0001892

UGACAGAAGAGAGUGAGCAC

>ptc-miR169p MIMAT0001972

CAGCCAAGGAUGACUUGCCGG

>ptc-miR395h MIMAT0002028

CUGAAGUGUUUGGGGGAACUC

>ptc-miR395i MIMAT0002029

CUGAAGUGUUUGGGGGAACUC

>ptc-miR6456 MIMAT0025266

UUGAGUCCUUCCAUUAGAUCC

>ptc-miR398a MIMAT0002041

UGUGUUCUCAGGUCACCCCUU

>ptc-miR394b-5p MIMAT0002020

UUGGCAUUCUGUCCACCUCC

>ptc-miR393b-3p MIMAT0022909

AUCAUGCUAUCCCUUUGGAUU

>ptc-miR6462d MIMAT0025224

UCUCUUAUGCAUUUUUGUCCC

>ptc-miR399a MIMAT0002044

UGCCAAAGGAGAUUUGCCCCG

>ptc-miR159d MIMAT0001904

CUUGGAUUGAAGGGAGCUCCU

>ptc-miR7824 MIMAT0030383

UUGAGAAAAGUCAAUCGGACC

>ptc-miR482b-5p MIMAT0027338

GGCAUGAGGUGUUUGGCAAGA

>ptc-miR6445b MIMAT0025256

UUCAUUCCUCUUCCUAAAAUGG

>ptc-miR474a MIMAT0002064

CAAAAGUUGCUGGGUUUGGCUGGG

>ptc-miR169f MIMAT0001962

CAGCCAAGGAUGACUUGCCGG

>ptc-miR6470 MIMAT0025209

CUCUGAUAUCAUAUUAAAAAA

>ptc-miR530a MIMAT0005997

UGCAUUUGCACCUGCACCUU

>ptc-miR478r MIMAT0002092

UAACGUGUCUCCUAUUUUUAGGGA

>ptc-miR167a MIMAT0001941

UGAAGCUGCCAGCAUGAUCUA

>ptc-miR403b MIMAT0002057

UUAGAUUCACGCACAAACUCG

>ptc-miR319h MIMAT0002009

UUGGACUGAAGGGAGCUCCU

>ptc-miR7822 MIMAT0030381

UUUGAAAUUGAACAAAUGGUA

>ptc-miR477b MIMAT0002075

AUCUCCCUCAGAGGCUUCCAA

>ptc-miR478h MIMAT0002082

UAACGUGUCUCCUAUUUUUAGGGA

>ptc-miR399j MIMAT0002053

UGCCAAAGGAGAUUUGUCCGG

>ptc-miR472a MIMAT0002060

UUUUCCCUACUCCACCCAUCCC

>ptc-miR399h MIMAT0002051

UGCCAAAGGAGAGUUUCCCUG

>ptc-miR160d MIMAT0001910

UGCCUGGCUCCCUGUAUGCCA

>ptc-miR169c MIMAT0001959

CAGCCAAGGAUGACUUGCCGA

>ptc-miR7814 MIMAT0030372

UAGAUUGUUUUUAUGCUUUGA

>ptc-miR172f MIMAT0001998

AGAAUCUUGAUGAUGCUGCAU

>ptc-miR396e-5p MIMAT0002035

UUCCACAGCUUUCUUGAACUU

>ptc-miR6448 MIMAT0025251

UAGGCACAGAAUUAACAAGGC

>ptc-miR169s MIMAT0001975

UCAGCCAAGGAUGACUUGCCG

>ptc-miR475c MIMAT0002069

UUACAAUGUCCAUUGAUUAAG

>ptc-miR472b MIMAT0002061

UUUUCCCAACUCCACCCAUCCC

>ptc-miR171f MIMAT0001988

UGAUUGAGCCGUGCCAAUAUC

>ptc-miR7818 MIMAT0030378

UUUCUUAUCGAUCACUAGACG

>ptc-miR6425d-5p MIMAT0025214

UUGUCUUCCAUGGAAUAGGCAG

>ptc-miR7840 MIMAT0030397

CAAGGAGUAAUUAGUGACAUC

>ptc-miR166q MIMAT0001940

UCGGACCAGGCUUCAUUCCUU

>ptc-miR166e MIMAT0001928

UCGGACCAGGCUUCAUUCCCC

>ptc-miR167b MIMAT0001942

UGAAGCUGCCAGCAUGAUCUA

>ptc-miR171c MIMAT0001985

AGAUUGAGCCGCGCCAAUAUC

>ptc-miR1446e MIMAT0006008

UUCUGAACUCUCUCCCUCAA

>ptc-miR6475 MIMAT0025216

UCUUGAGAAGUAAAGAACGAC

>ptc-miR6449 MIMAT0025253

CAUGAUUCUGAAUAACGGUUU

>ptc-miR6431 MIMAT0025232

UUAUGUGGCAUAAAAGAAUCAA

>ptc-miR481a MIMAT0002098

AGGACCUCACUUAACAGCUUAAGC

>ptc-miR6425a-5p MIMAT0025190

UUGUCUUCCAUGGAAUAGGCAG

>ptc-miR169i MIMAT0001965

UAGCCAAGGAUGACUUGCCUG

>ptc-miR6460 MIMAT0025186

UGAUAUGUGGCAUUCAAUCGA

>ptc-miR478e MIMAT0002080

UGACGAGUCUUCUAUUUUUAGGGA

>ptc-miR478k MIMAT0002085

UAACGUGUCUCCUAUUUUUAGGGA

>ptc-miR390b MIMAT0002012

AAGCUCAGGAGGGAUAGCGCC

>ptc-miR171k MIMAT0003943

GGAUUGAGCCGCGCCAAUAUC

>ptc-miR6473 MIMAT0025212

UCCACAAUCCCAUCAAGACUU

>ptc-miR6425b-5p MIMAT0025198

UUGUCUUCCAUGGAAUAGGCAG

>ptc-miR171g-3p MIMAT0001989

UGAUUGAGCCGUGCCAAUAUC

>ptc-miR172g-3p MIMAT0001999

GGAAUCUUGAUGAUGCUGCAG

>ptc-miR7836 MIMAT0030408

UGGGUGGGAGGUGUGGUAGCU

>ptc-miR169q MIMAT0001973

UAGCCAAGGACGACUUGCCUG

>ptc-miR482c-3p MIMAT0025278

UCUUUCCGAGUCCUCCCAUACC

>ptc-miR6440a MIMAT0025243

GAGUUUGAUCGAUUUCGAGUU

>ptc-miR166b MIMAT0001925

UCGGACCAGGCUUCAUUCCCC

>ptc-miR478p MIMAT0002090

UAACGUGUCUUCUAUUUUUAGGGA

>ptc-miR395c MIMAT0002023

CUGAAGUGUUUGGGGGAACUC

>ptc-miR396b MIMAT0002032

UUCCACAGCUUUCUUGAACUG

>ptc-miR828a MIMAT0025279

UCUUGCUCAAAUGAGUAUUCCA

>ptc-miR156d MIMAT0001893

UGACAGAAGAGAGUGAGCAC

>ptc-miR6444 MIMAT0025247

UAGGAAAAUUAGAGAUUAUGU

>ptc-miR172i MIMAT0002001

AGAAUCCUGAUGAUGCUGCAA

>ptc-miR7831 MIMAT0030390

UACAUGUAGAGACCACCAAAC

>ptc-miR6458 MIMAT0025182

ACGCUCAAAUAUAUAAGGUGUU

>ptc-miR164f MIMAT0001923

UGGAGAAGCAGGGCACAUGCU

>ptc-miR7838 MIMAT0030394

ACAUGUGCUGGGUAGGAGGAA

>ptc-miR169b-5p MIMAT0001958

CAGCCAAGGAUGACUUGCCGA

>ptc-miR399d MIMAT0002047

UGCCAAAGAAGAUUUGCCCCG

>ptc-miR169j MIMAT0001966

UAGCCAAGGAUGACUUGCCUG

>ptc-miR156i MIMAT0001898

UUGACAGAAGAUAGAGAGCAC

>ptc-miR6476c MIMAT0030369

UCAGUGGAGAUGAAACAUGA

>ptc-miR6465 MIMAT0025193

UUAAAGCAGGGGGUCUAGUAGU

>ptc-miR395j MIMAT0002030

CUGAAGUGUUUGGGGGAACUC

>ptc-miR7827 MIMAT0030403

GCUAGGACCAAGUUUUUUGGA

>ptc-miR478n MIMAT0002094

UAACGUGUCUCCUAUUUUUAGGGA

>ptc-miR6462c-5p MIMAT0025205

AAGGGACAAAAAUGGCAUAAGA

>ptc-miR6480 MIMAT0025223

UUGCUGAAACGAUUGAACUAU

>ptc-miR1444d MIMAT0030385

CGAACGUUGACCGAAUGUGAA

>ptc-miR6455 MIMAT0025265

UCAAAUAGCAUCCUCAACAUU

>ptc-miR394a-3p MIMAT0006783

CUGUUGGUCUCUCUUUGUAA

>ptc-miR478j MIMAT0002084

UAACGUGUCUCCUAUUUUUAGGGA

>ptc-miR7823 MIMAT0030382

UUGCAUGCAUGAACUUGAAAU

>ptc-miR6450b MIMAT0025261

CGAACACAGGACUCAAGGCUA

>ptc-miR6432 MIMAT0025233

CGGCUCUAGAGAAAAAGAUGG

>ptc-miR7842 MIMAT0030399

UAAAUCAAGCCCGGUACUUUU

>ptc-miR6462e MIMAT0030375

UCUUAUGCGUUUUUGUCUCU

>ptc-miR477d-5p MIMAT0025275

AUCUCCCUCAAAGGCUUCCUCU

>ptc-miR319c MIMAT0002004

UUGGACUGAAGGGAGCUCCC

>ptc-miR478d MIMAT0002079

UGACAUGUCUUCUAUUUUUAGUAA

>ptc-miR536 MIMAT0025260

CGCUCGCCAGCGUUGCACCACC

>ptc-miR169ae MIMAT0001956

UAGCCAAGGACGACUUGCCCA

>ptc-miR395g MIMAT0002027

CUGAAGUGUUUGGGGGAACUC

>ptc-miR475b-3p MIMAT0002068

UUACAGUGCCCAUUGAUUAAG

>ptc-miR397a MIMAT0002038

UCAUUGAGUGCAGCGUUGAUG

>ptc-miR6479 MIMAT0025222

AUCAAUGAGAAUACUACUGCA

>ptc-miR160f MIMAT0001912

UGCCUGGCUCCCUGAAUGCCA

>ptc-miR164e MIMAT0001922

UGGAGAAGCAGGGCACGUGCA

>ptc-miR7826 MIMAT0030387

UUACCAAGUUUCAAAUUCUCA

>ptc-miR169y MIMAT0001981

UAGCCAUGGAUGAAUUGCCUG

>ptc-miR6439b MIMAT0025258

AGCAGAAGCCAUCACUAGCGC

>ptc-miR171a-5p MIMAT0022901

GGAUAUUGGUACGGUUCAAUC

>ptc-miR166p MIMAT0001939

UCGGACCAGGCUCCAUUCCUU

>ptc-miR478q MIMAT0002091

UAACGUGUCUCCUAUUUUUAGGGA

>ptc-miR169m MIMAT0001969

UAGCCAAGGAUGACUUGCCUG

>ptc-miR3627a MIMAT0025263

UCUGUCGCUGGAAAGAUGGUAC

>ptc-miR6440b MIMAT0025183

GAGUUUGAUCGAUUUCGAGUU

>ptc-miR167g-5p MIMAT0001947

UGAAGCUGCCAGCAUGAUCUU

>ptc-miR167h-5p MIMAT0001948

UGAAGCUGCCAACAUGAUCUG

>ptc-miR1444a MIMAT0005999

UCCACAUUCGGUCAAUGUUC

>ptc-miR6452 MIMAT0025262

UGCAAAGGUAACUGAGACAAU

>ptc-miR6425b-3p MIMAT0025199

UCCAUGGAAGAUAAUGACUCG

>ptc-miR171d MIMAT0001986

AGAUUGAGCCGCGCCAAUAUC

>ptc-miR828b-3p MIMAT0025281

UCAUUCGAGCAAGAAAUAUUA

>ptc-miR395k MIMAT0025273

CUGAAGUGUUUGGGGGAACUC

>ptc-miR828b-5p MIMAT0025280

UCUUGCUCAAAUGAGUAUUCCA

>ptc-miR396f MIMAT0002036

UUCCACGGCUUUCUUGAACUG

>ptc-miR169g MIMAT0001963

CAGCCAAGGAUGACUUGCCGG

>ptc-miR169t MIMAT0001976

GAGCCAAGAAUGACUUGCCGG

>ptc-miR172c MIMAT0001995

AGAAUCUUGAUGAUGCUGCAU

>ptc-miR166l MIMAT0001935

UCGGACCAGGCUUCAUUCCCC

>ptc-miR169aa MIMAT0001952

GAGCCAAGAAUGACUUGUCGG

>ptc-miR7813 MIMAT0030400

UGGUAAUGCAAGUGUUGCUAA

>ptc-miR6459a-5p MIMAT0025184

AGCUCAAGCACAAAUUCGAUC

>ptc-miR164b MIMAT0001919

UGGAGAAGCAGGGCACGUGCA

>ptc-miR6441 MIMAT0025244

AAGUUGACGGAAGGACUACAU

>ptc-miR171g-5p MIMAT0022902

UGUUGGGAUGGCUCAAUCAUG

>ptc-miR166k MIMAT0001934

UCGGACCAGGCUUCAUUCCCC

>ptc-miR169u-5p MIMAT0001977

UAGCCAAGGACGACUUGCCUA

>ptc-miR169af MIMAT0001957

UAGCCAAGGACGACUUGCCCA

>ptc-miR172b-3p MIMAT0001994

AGAAUCUUGAUGAUGCUGCAU

>ptc-miR2111b MIMAT0025272

UAAUCUGCAUCCUGAGGUUUG

>ptc-miR393a-5p MIMAT0002015

UCCAAAGGGAUCGCAUUGAUC

>ptc-miR6426b MIMAT0025259

GUGGAGACAUGGAAGUGAAGA

>ptc-miR166f MIMAT0001929

UCGGACCAGGCUUCAUUCCCC

>ptc-miR478o MIMAT0002089

UAACGUGUCUCCUAUUUUUAGGGA

>ptc-miR530b MIMAT0005998

UGCAUUUGCACCUGCAUCUU

>ptc-miR1446b MIMAT0006005

UUCUGAACUCUCUCCCUCAA

>ptc-miR399g MIMAT0002050

UGCCAAAGGAGAAUUGCCCUG

>ptc-miR167g-3p MIMAT0022894

AGAUCAUGUGGCAGUUUCACC

>ptc-miR159a MIMAT0001901

UUUGGAUUGAAGGGAGCUCUA

>ptc-miR6469 MIMAT0025202

UGGCAGAAAAGGAUUCGUUUA

>ptc-miR396c MIMAT0002033

UUCCACAGCUUUCUUGAACUU

>ptc-miR403d MIMAT0030370

UUAGAUUCACGCACAAACUCG

>ptc-miR6450a MIMAT0025254

CUUUGUCAGGACUCAAGGCUA

>ptc-miR171b MIMAT0001984

UUGAGCCGUGCCAAUAUCACG

>ptc-miR169w MIMAT0001979

UAGCCAAGGAUGACUUGCCCA

>ptc-miR6443 MIMAT0025246

GUAUGAUCAUAGAUGAUGGAG

>ptc-miR169k MIMAT0001967

UAGCCAAGGAUGACUUGCCUG

>ptc-miR478i MIMAT0002083

UAACGUGUCUCCUAUUUUUAGGGA

>ptc-miR156e MIMAT0001894

UGACAGAAGAGAGUGAGCAC

>ptc-miR6425d-3p MIMAT0025215

UCCAUGGAAGAUAAUGACUCG

>ptc-miR166c MIMAT0001926

UCGGACCAGGCUUCAUUCCCC

>ptc-miR475a-5p MIMAT0022915

AAUGGCCAUUGUAAGAGUAGA

>ptc-miR171h-5p MIMAT0022903

UGUUGGGAUGGCUCAAUCAUA

>ptc-miR477a-5p MIMAT0002074

AUCUCCCUCAGAGGCUUCCAA

>ptc-miR393a-3p MIMAT0022908

AUCAUGCUAUCCCUUUGGAUU

>ptc-miR172h-5p MIMAT0022906

GGAGCACCAUCAAGAUUCACA

>ptc-miR7833 MIMAT0030392

UAAUUAGAACUCAUACUAGAC

>ptc-miR478c MIMAT0002078

UGACGUGUCUUCUAUUUUUAGGGA

>ptc-miR6438a MIMAT0025241

UUGUACACAGAAUAGGUGAAAU

>ptc-miR6464 MIMAT0025192

UGAUUGCUUGUUGGAUAUUAU

>ptc-miR6425c-5p MIMAT0025207

UUGUCUUCCAUGGAAUAGGCAG

>ptc-miR6437a MIMAT0025240

UCACGGACGGCGGCUCCAAGCA

>ptc-miR167c MIMAT0001943

UGAAGCUGCCAGCAUGAUCUA

>ptc-miR156f MIMAT0001895

UGACAGAAGAGAGUGAGCAC

>ptc-miR6427-5p MIMAT0025227

UCGUAAUGCUUCAUUCUCACAA

>ptc-miR160e-5p MIMAT0001911

UGCCUGGCUCCCUGAAUGCCA

>ptc-miR169e MIMAT0001961

CAGCCAAGGAUGACUUGCCGG

>ptc-miR3627b MIMAT0030391

UGUCGCAGGAGAGAUGGCGCUA

>ptc-miR162a MIMAT0001915

UCGAUAAACCUCUGCAUCCAG

>ptc-miR7832 MIMAT0030405

UAGUUCCCAACCUACACCACA

>ptc-miR319i MIMAT0002010

UUGGGCUGAAGGGAGCUCCC

>ptc-miR399f MIMAT0002049

UGCCAAAGGAGAAUUGCCCUG

>ptc-miR7841 MIMAT0030398

GGGGGUUGCUGUCAAGCAUAA

>ptc-miR482a.1 MIMAT0002103

CCUACUCCUCCCAUUCC

>ptc-miR396g-5p MIMAT0002037

UUCCACGGCUUUCUUGAACUU

>ptc-miR6476b MIMAT0030368

UCAGUGGAGAUGAAACAUGA

>ptc-miR477e-3p MIMAT0022914

UGAGGCCUUUGGGGGAGAGUGG

>ptc-miR169ag MIMAT0025269

AAGCCAAGGGUGACUUGCCUGA

>ptc-miR7816 MIMAT0030374

AAUGUUGUUAUUAACACUGUA

>ptc-miR166h MIMAT0001931

UCGGACCAGGCUUCAUUCCCC

>ptc-miR6477 MIMAT0025218

UGAACAGUAGACGUGAAUUAU

>ptc-miR6454 MIMAT0025264

CUUGUAACCUGAGUAGAGGCA

>ptc-miR319b MIMAT0002003

UUGGACUGAAGGGAGCUCCC

>ptc-miR159e MIMAT0001905

CUUGGGGUGAAGGGAGCUCCU

>ptc-miR166a MIMAT0001924

UCGGACCAGGCUUCAUUCCCC

>ptc-miR478b MIMAT0002077

UGACGUGUCUUCUAUUUUUAGGGA

>ptc-miR6462b MIMAT0025204

UCUCUUAUGCAUUUUUGUCCC

>ptc-miR397b MIMAT0002039

CCAUUGAGUGCAGCGUUGAUG

>ptc-miR6445a MIMAT0025248

UUCAUUCCUCUUCCUAAAAUGG

>ptc-miR482d-5p MIMAT0025234

GGACAUGGGUUGGUUUGCAAGA

>ptc-miR168b-5p MIMAT0001950

UCGCUUGGUGCAGGUCGGGAA

>ptc-miR166m MIMAT0001936

UCGGACCAGGCUUCAUUCCCC

>ptc-miR167f-5p MIMAT0001946

UGAAGCUGCCAGCAUGAUCUU

>ptc-miR474c MIMAT0002066

CAAAAGCUGUUGGGUUUGGCUGGG

>ptc-miR168a-5p MIMAT0001949

UCGCUUGGUGCAGGUCGGGAA

>ptc-miR6462f MIMAT0030376

UCUUAUGCGUUUUUGUCUCU

>ptc-miR6466-3p MIMAT0025195

UAUCAAUCAUCAAAUGUUCGU

>ptc-miR7825 MIMAT0030384

UUGAAGAAAGGUAGACAGAUAG

>ptc-miR6446 MIMAT0025249

UUGCUGGGUCCUGAUGAUGGA

>ptc-miR168a-3p MIMAT0022896

CCCGCCUUGCAUCAACUGAAU

>ptc-miR171e MIMAT0001987

UGAUUGAGCCGUGCCAAUAUC

>ptc-miR169ab MIMAT0001953

CAGCCAAGGAAGACUUGCCC

>ptc-miR164a MIMAT0001918

UGGAGAAGCAGGGCACGUGCA

>ptc-miR6439a MIMAT0025242

CCAAAUGAAGAAGCAGAAGCC

>ptc-miR6471 MIMAT0025210

UUUGGGAUCAUCAGGACAGCC

>ptc-miR159b MIMAT0001902

UUUGGAUUGAAGGGAGCUCUA

>ptc-miR6435 MIMAT0025238

UGAAUAAUGGAGACACUCUAG

>ptc-miR172d MIMAT0001996

GGAAUCUUGAUGAUGCUGCAU

>ptc-miR2111a MIMAT0025270

UAAUCUGCAUCCUGAGGUUUG

>ptc-miR1445 MIMAT0006002

UCCCUUGUAGACUAGAAAAA

>ptc-miR1446c MIMAT0006006

UUCUGAACUCUCUCCCUCAA

>ptc-miR160g MIMAT0001913

UGCCUGGCUCCCUGGAUGCCA

>ptc-miR6459a-3p MIMAT0025185

UCGAAUUUGGGCUUGAGAUUG

>ptc-miR827 MIMAT0006003

UUAGAUGACCAUCAACGAAAA

>ptc-miR7837 MIMAT0030393

UGGGUGGGAGGUGUGGUAGCU

>ptc-miR396g-3p MIMAT0022911

CUCAAGAAAGCCGUGGGAAAA

>ptc-miR6425a-3p MIMAT0025191

UCCAUGGAAGAUAAUGACUCG

>ptc-miR395f MIMAT0002026

CUGAAGUGUUUGGGGGAACUC

>ptc-miR168b-3p MIMAT0022897

CCCGCCUUGCAUCAACUGAAU

>ptc-miR160b-3p MIMAT0022890

GCGUAUGAGGAGCCAUGCAUA

>ptc-miR6468-3p MIMAT0025201

GGAGUGAUUCAGGGAACCCAU

>ptc-miR6462a MIMAT0025188

UCUCUUAUGCAUUUUUGUCCC

>ptc-miR319f MIMAT0002007

UUGGACUGAAGGGAGCUCCU

>ptc-miR395b MIMAT0002022

CUGAAGUGUUUGGGGGAACUC

>ptc-miR6425c-3p MIMAT0025208

UCCAUGGAAGAUAAUGACUCG

>ptc-miR156j MIMAT0001899

UUGACAGAAGAUAGAGAGCAC

>ptc-miR476a MIMAT0002071

UAGUAAUCCUUCUUUGCAAAG

>ptc-miR169x MIMAT0001980

UAGCCAAGGAUGACUUGCUCG

>ptc-miR399c MIMAT0002054

UGCCAAAGGAGAUUUGCUCAC

>ptc-miR6426a MIMAT0025221

GUGGAGACAUGGAAGUGAAGA

>ptc-miR167f-3p MIMAT0022893

AGAUCAUGUGGCAGUUUCACC

>ptc-miR396e-3p MIMAT0022910

CUCAAGAAAGCUGUGGGAGA

>ptc-miR6466-5p MIMAT0025194

UCUGGUAUGAGCAUUUGAUGA

>ptc-miR7820 MIMAT0030380

AUCAUAUAGGUUGAUCCUCGU

>ptc-miR166g MIMAT0001930

UCGGACCAGGCUUCAUUCCCC

>ppe-miR169f MIMAT0031459

UAGCCAAGGAUGACUUGCCUGC

>ppe-miR8122-5p MIMAT0031523

UUCCACAGAUCUUUCCUCAUU

>ppe-miR399h MIMAT0031504

UGCCAAAGAAGAGUUGCCCUA

>ppe-miR160a MIMAT0031438

UGCCUGGCUCCCUGUAUGCCA

>ppe-miR171g MIMAT0027304

UGAUUGAGCCGUGCCAAUAUC

>ppe-miR8130-3p MIMAT0031557

CCCUUCCAGUAAGGCACCCCC

>ppe-miR3627-5p MIMAT0031534

UCGCAGGAGAGAUGGCACUGUC

>ppe-miR6295 MIMAT0027331

GAGGACAGAAGAUGAUUCAGC

>ppe-miR399g MIMAT0031503

UGCCAAAGAAGAGUUGCCCUA

>ppe-miR6287 MIMAT0027319

CAAGAAGUGGAAGUUUUGGGC

>ppe-miR6273 MIMAT0027299

AAUGCAGCAUGAUUUUUUUUU

>ppe-miR394b MIMAT0031476

UUGGCAUUCUGUCCACCUCC

>ppe-miR396a MIMAT0031494

UUCCACAGCUUUCUUGAACGU

>ppe-miR2111a MIMAT0031517

UAAUCUGCAUCCUGAGGUUUA

>ppe-miR399j MIMAT0031506

UGCCAAAGAAGAGUUGCCCUA

>ppe-miR169i MIMAT0031462

UAGCCAAGGAUGACUUGCCUGC

>ppe-miR169l MIMAT0031465

GAGCCAAGGAUGAAUUGCCGG

>ppe-miR171c MIMAT0027323

UGAUUGAGCCGUGCCAAUAUC

>ppe-miR395m MIMAT0031491

CUGAAGUGUUUGGGGGAACUC

>ppe-miR2111d MIMAT0031520

UAAUCUGCAUCCUGAGGUUUA

>ppe-miR399e MIMAT0031501

UGCCAAAGAAGAGUUGCCCUA

>ppe-miR167c MIMAT0031452

UGAAGCUGCCAGCAUGAUCUGA

>ppe-miR319b MIMAT0027303

UAGCUGCCGAGUCAUUCAUCCA

>ppe-miR156g MIMAT0031434

UUGACAGAAGAUAGAGAGCAC

>ppe-miR6288c-5p MIMAT0031565

CUUGUUAUUUUUUAAUUGAUU

>ppe-miR390 MIMAT0031473

AAGCUCAGGAGGGAUAGCGCC

>ppe-miR169c MIMAT0031457

CAGCCAAGGAUGACUUGCCGG

>ppe-miR393a MIMAT0031474

CAUCCAAAGGGAUCGCAUUGA

>ppe-miR172b MIMAT0031470

AGAAUCUUGAUGAUGCUGCAU

>ppe-miR398b MIMAT0027328

CGUGUUCUCAGGUCGCCCCUG

>ppe-miR6288b-3p MIMAT0031561

UCAAUUAGAAAAUGAUAAGUG

>ppe-miR8123-3p MIMAT0031526

UUGUGCCAUUGCUCAAGC

>ppe-miR6267c-5p MIMAT0031542

AUUGCUGAUCACCUCUCUAAU

>ppe-miR398a-3p MIMAT0027294

UGUGUUCUCAGGUCGCCCCUG

>ppe-miR6265 MIMAT0027284

UUGAACUUUGACCCGAUUCGCAU

>ppe-miR8129-5p MIMAT0031554

GAUAUCCGCACAUUAUUAUUG

>ppe-miR160b MIMAT0031439

UGCCUGGCUCCCUGUAUGCCA

>ppe-miR6292 MIMAT0027327

UAUCUUUUAAUCGUUAGAUCA

>ppe-miR8124-5p MIMAT0031531

ACUUGGUAUCUUGGUGCCGGU

>ppe-miR6293 MIMAT0027329

UAAGAGGCUGAUGACUAAAAC

>ppe-miR156e MIMAT0031432

UGACAGAAGAGAGUGAGCAC

>ppe-miR156c MIMAT0031430

UGACAGAAGAGAGUGAGCAC

>ppe-miR482d-3p MIMAT0027315

CCUCCCAUGCCACGCAUUUCUA

>ppe-miR828-5p MIMAT0031168

UCUUGCUCAAAUGAGUAUUCCA

>ppe-miR827 MIMAT0031516

UUAGAUGACCAUCAACAAACA

>ppe-miR482a-5p MIMAT0031165

GGGUGAGAGGUUGCCGGAAAGA

>ppe-miR7125-3p MIMAT0031537

CGAACUUAUUGCAACUAGCUU

>ppe-miR8126-3p MIMAT0031539

UUCAGUAUUUUGACUCAGAA

>ppe-miR167b MIMAT0031451

UGAAGCUGCCAGCAUGAUCUA

>ppe-miR8127-5p MIMAT0031546

CAACUGUGGACAUACCCUUUG

>ppe-miR395a-3p MIMAT0031478

CUGAAGUGUUUGGGGGGACCC

>ppe-miR6276 MIMAT0027305

AAAGGCUCAUACAAAUAUUCC

>ppe-miR6261 MIMAT0027277

AAGUGAUUAUAUGGAGAAGCAC

>ppe-miR395f MIMAT0031484

CUGAAGUGUUUGGGGGAACUC

>ppe-miR6269 MIMAT0027293

UGUGAAUAGUGAUUGCCAUGG

>ppe-miR8124-3p MIMAT0031532

UGGCACCAAUGAUACCAAGUUU

>ppe-miR482c-3p MIMAT0031169

UUCCCAAGCCCGCCCAUUCCAA

>ppe-miR166b MIMAT0031446

UCGGACCAGGCUUCAUUCCCC

>ppe-miR6282 MIMAT0027312

GUUGAUCGAUGUGGGAUGUUACA

>ppe-miR395d MIMAT0031482

CUGAAGUGUUUGGGGGAACUC

>ppe-miR8130-5p MIMAT0031556

GGGUUCCUUGUUGGAAGGACU

>ppe-miR3627-3p MIMAT0031535

UGGUGUCAUCCCUCCUGUGACC

>ppe-miR399n MIMAT0031510

UGCCAAAGGAGAUUUGCUCGG

>ppe-miR164a MIMAT0031441

UGGAGAAGCAGGGCACGUGCA

>ppe-miR2111c MIMAT0031519

UAAUCUGCAUCCUGAGGUUUA

>ppe-miR169b MIMAT0031456

CAGCCAAGGAUGACUUGCCGG

>ppe-miR171h MIMAT0027285

UUGAGCCGCGUCAAUAUCUCC

>ppe-miR6274b-5p MIMAT0031552

AUUUCGACUAAUAACACAAUG

>ppe-miR6272 MIMAT0027298

UAGCUGUAAAUGAGUGUUUUU

>ppe-miR6257 MIMAT0027270

UCUUAACUGUUGGAUUAGGCU

>ppe-miR166d MIMAT0031448

UCGGACCAGGCUUCAUUCCCC

>ppe-miR397 MIMAT0031496

UCAUUGAGUGCAGCGUUGAUG

>ppe-miR6259 MIMAT0027275

UAGAAAAAUACGGGCGAUAAA

>ppe-miR156i MIMAT0031436

UUGACAGAAGAUAGAGAGCAC

>ppe-miR6270 MIMAT0027295

UUCUGGUAUUGGAAUUUCAUU

>ppe-miR477b-3p MIMAT0027288

GUUGGGGGCUCUUUUGGGACG

>ppe-miR1511-3p MIMAT0031522

ACCUGGCUCUGAUACCAUAAC

>ppe-miR8133-5p MIMAT0031563

UCCUGUGCGAACGUCCAGAAG

>ppe-miR477-3p MIMAT0031541

CGAAGCCUUUGGGGAGAGUAA

>ppe-miR8126-5p MIMAT0031538

UCUGAGUCAGAUUACUGAAUA

>ppe-miR530 MIMAT0031513

UCUGCAUUUGCACCUGCACCU

>ppe-miR6288b-5p MIMAT0031560

CUUGUUAUUUUUUAAUUGAUU

>ppe-miR7122a-5p MIMAT0031527

UUAUACAAUGAAAUCACGGCCG

>ppe-miR395l MIMAT0031490

CUGAAGUGUUUGGGGGAACUC

>ppe-miR6284 MIMAT0027314

UUUGGACCAUGGAUGAAGAUU

>ppe-miR477b-5p MIMAT0031171

UCCCUCAAGGGCUCCCAAUAUU

>ppe-miR482d-5p MIMAT0031174

GAGAUGGGUGGCUGGGAAGGA

>ppe-miR6297b MIMAT0027336

GAUGUAUUGUCGUCGCGCAAAGU

>ppe-miR399l MIMAT0031508

UGCCAAAGAAGAGUUGCCCUA

>ppe-miR168 MIMAT0031454

UCGCUUGGUGCAGGUCGGGAA

>ppe-miR399d MIMAT0031500

UGCCAAAGAAGAGUUGCCCUA

>ppe-miR403 MIMAT0031511

UUAGAUUCACGCACAAACUCG

>ppe-miR156h MIMAT0031435

UUGACAGAAGAUAGAGAGCAC

>ppe-miR156a MIMAT0031428

UGACAGAAGAAAGAGAGCAC

>ppe-miR169h MIMAT0031461

UAGCCAAGGAUGACUUGCCUGC

>ppe-miR396b MIMAT0031495

UUCCACAGCUUUCUUGAACUU

>ppe-miR169k MIMAT0031464

GAGCCAAGGAUGAAUUGCCGG

>ppe-miR159 MIMAT0031437

UUUGGAUUGAAGGGAGCUCUA

>ppe-miR399c MIMAT0031499

UGCCAAAGAAGAGUUGCCCUA

>ppe-miR6267b MIMAT0027297

AUUAGAGAGGCGGUAAACAAU

>ppe-miR172a-5p MIMAT0031468

GUAGCAUCAUCAAGAUUCACG

>ppe-miR8128-3p MIMAT0031551

UCGUGGGGAGAGAUCUAAUCG

>ppe-miR399i MIMAT0031505

UGCCAAAGAAGAGUUGCCCUA

>ppe-miR8128-5p MIMAT0031550

AUUAGACCUCUCCCGACGAAA

>ppe-miR5225-3p MIMAT0031549

UCAUCUCUCCUCGACUGAA

>ppe-miR6260 MIMAT0027276

UGGAGUGAGAGAAUGGGAGGU

>ppe-miR164d MIMAT0031444

UGGAGAAGCAGGGCACAUGCU

>ppe-miR8123-5p MIMAT0031525

UGAGCAAUGGCACACAGCCCU

>ppe-miR6266a MIMAT0027290

UAAAUGCAGGGGCAAAAUGAU

>ppe-miR172a-3p MIMAT0031469

AGAAUCUUGAUGAUGCUGCAU

>ppe-miR395g MIMAT0031485

CUGAAGUGUUUGGGGGAACUC

>ppe-miR6274a MIMAT0027301

UAUUUUGCUAUCUUCGGGCAAUA

>ppe-miR482e MIMAT0027318

UUGCCUAUUCCUCCCAUGCCAA

>ppe-miR8132 MIMAT0031562

UCCAACGAUGGGUGACCACAA

>ppe-miR171b MIMAT0027309

UUGAGCCGCGCCAAUAUCACU

>ppe-miR6288a MIMAT0027320

GAAAAUGACAAGUGGCUAGUU

>ppe-miR169g MIMAT0031460

UAGCCAAGGAUGACUUGCCUGC

>ppe-miR6263 MIMAT0027280

AAGUGGACAAAAGGGGAGUGG

>ppe-miR6291c-5p MIMAT0031544

CCACAUUUAUAGAUUACCUUG

>ppe-miR395b-3p MIMAT0031480

CUGAAGUGUUUGGGGGGACCC

>ppe-miR395n MIMAT0031492

CUGAAGUGUUUGGGGGAACUC

>ppe-miR6294 MIMAT0027330

UGGUGUAGGCUAAUCACAAUC

>ppe-miR395a-5p MIMAT0031477

GUUCCCUCAAACACUUCAUU

>ppe-miR171d-3p MIMAT0031467

CGAGCCGAAUCAAUAUCACUC

>ppe-miR7125-5p MIMAT0031536

GCUAGGUGCAACAAGUUCAAU

>ppe-miR535a MIMAT0031514

UGACAACGAGAGAGAGCACGC

>ppe-miR399a MIMAT0031497

CGCCAAAGGAGAGUUGCCCUU

>ppe-miR482a-3p MIMAT0027272

UUUCCGAAACCUCCCAUUCCAA

>ppe-miR399f MIMAT0031502

UGCCAAAGAAGAGUUGCCCUA

>ppe-miR6280 MIMAT0027310

UUGGCAGUAAGAUUUUUGGUG

>ppe-miR156f MIMAT0031433

UGACAGAAGAUAGAGAGCAC

>ppe-miR171f MIMAT0027274

UGAUUGAGCCGUGCCAAUAUC

>ppe-miR6296 MIMAT0027332

UAAGGCCCUUAGAUGAGACCC

>ppe-miR8122-3p MIMAT0031524

UGAAGGAAGAUUUGUGGAAAG

>ppe-miR6275 MIMAT0027302

AGUGGAAGUAGCAAGGGGAAGC

>ppe-miR6285 MIMAT0027316

UAGUGAAGUUUGAAUUAGGGCU

>ppe-miR395i MIMAT0031487

CUGAAGUGUUUGGGGGAACUC

>ppe-miR6271 MIMAT0027296

UCAAGAUUGAGAGAUAUAAUG

>ppe-miR319a MIMAT0027300

UUGGACUGAAGGGAGCUCCC

>ppe-miR393b MIMAT0031475

UCCAAAGGGAUCGCAUUGAUC

>ppe-miR169j MIMAT0031463

UAGCCAAGGAUGACUUGCCUGC

>ppe-miR6297a MIMAT0027335

AAUAAUUUUUCGUCGCGCAAAAU

>ppe-miR6281 MIMAT0027311

GUUAGAGAUAGAGAGAGUGAG

>ppe-miR482b-3p MIMAT0027273

CUUCCCAAACCUCCCAUUCCUA

>ppe-miR6258 MIMAT0027271

UUCCAGCUGUAAAGAUCAAGA

>ppe-miR6278 MIMAT0027307

UGAACCUUGUGUACAAAUUGGC

>ppe-miR167d MIMAT0031453

UGAAGCUGCCAGCAUGAUCUUA

>ppe-miR858 MIMAT0031567

CUCGUUGUCUGUUCGACCUUG

>ppe-miR5225-5p MIMAT0031548

UCUGUCGUAGGAGAGAUGGCGC

>ppe-miR477a-5p MIMAT0031170

UCCCUCAAGGGCUCCCAAUAUU

>ppe-miR167a MIMAT0031450

UGAAGCUGCCAGCAUGAUCUA

>ppe-miR169d MIMAT0031458

UGAGCCAAGGAUGACUUGCCA

>ppe-miR477-5p MIMAT0031540

ACUCUCCCUCAAAGGCUUCUAG

>ppe-miR8133-3p MIMAT0031564

UAACUUCCGAACGUCCGCAUA

>ppe-miR6291c-3p MIMAT0031545

CAAGGUAGUUUAUAAAUGUGG

>ppe-miR6288c-3p MIMAT0031566

AACCAAUUAGAAAAUAACAAGUGG

>ppe-miR477a-3p MIMAT0027287

GUUGGGGGCUCUUUUGGGACG

>ppe-miR169e-5p MIMAT0031172

UGAGCCAAGGAUGACUUGCCA

>ppe-miR171a MIMAT0027286

UGAUUGAGCCGUGCCAAUAUC

>ppe-miR156d MIMAT0031431

UGACAGAAGAGAGUGAGCAC

>ppe-miR7122a-3p MIMAT0031528

GCCGUGUUUCUUUGUAUAAAG

>ppe-miR6286 MIMAT0027317

UUUGAACCAUUGGAUCGUAGUUA

>ppe-miR394a MIMAT0031167

UUGGCAUUCUGUCCACCUCC

>ppe-miR8127-3p MIMAT0031547

UUCAAAGGGUACAUCCACAGU

>ppe-miR6262 MIMAT0027278

UCUUUAGAAAGUUAGAAUUGU

>ppe-miR6266c MIMAT0027334

UAAAUGCAGGGGCAAAAUGAU

>ppe-miR164b MIMAT0031442

UGGAGAAGCAGGGCACGUGCA

>ppe-miR482f MIMAT0031512

UCUUUCCUACUCCACCCAUUCC

>ppe-miR6290 MIMAT0027325

UGAAUGAGUUCAGAGAUCGUGUA

>ppe-miR6291a MIMAT0027326

CUUACCACAUUUUUAUACCAU

>ppe-miR6283 MIMAT0027313

CAAAAGGGGAGUGGGAAAAUC

>ppe-miR828-3p MIMAT0027282

UCAUUUCAGCAAGCAGCGUUA

>ppe-miR6266b MIMAT0027324

UAAAUGCAGGGGCAAAAUGAU

>ppe-miR395j MIMAT0031488

CUGAAGUGUUUGGGGGAACUC

>ppe-miR166a MIMAT0031445

UCGGACCAGGCUUCAUUCCCC

>ppe-miR395b-5p MIMAT0031479

GUUCCCUCAAACACUUCAUU

>ppe-miR6264 MIMAT0027281

AUGCCUAUGGACACGUGUCAA

>ppe-miR172c MIMAT0031471

GGAAUCUUGAUGAUGCUGCAU

>ppe-miR166c MIMAT0031447

UCGGACCAGGCUUCAUUCCCC

>ppe-miR395e MIMAT0031483

CUGAAGUGUUUGGGGGAACUC

>ppe-miR1511-5p MIMAT0031521

CGUGGUAUCAGAGUCAUGUUA

>ppe-miR6279 MIMAT0027308

UAGACAAGAAUUCCAGAGACC

>ppe-miR164c MIMAT0031443

UGGAGAAGCAGGGCACGUGCA

>ppe-miR8131-3p MIMAT0031559

AAUCAACUCAGCUUAGCUGAACUG

>ppe-miR171d-5p MIMAT0031466

UGUGAUAUUGGUUCGGUUCAUA

>ppe-miR169a MIMAT0031455

CAGCCAAGGAUGACUUGCCGG

>ppe-miR171e MIMAT0027289

UUAUUGAACCGGACCAAUAUC

>ppe-miR395o MIMAT0031493

GUUCCCUCAAACACUUCAUU

>ppe-miR6277 MIMAT0027306

UGUGUGUGGAAAGAGCGAGAC

>ppe-miR7122b-3p MIMAT0031530

CCGUGUUUCCUUGUAUAAAG

>ppe-miR399m MIMAT0031509

UGCCAAAGGAGAUUUGCUCGG

>ppe-miR395k MIMAT0031489

CUGAAGUGUUUGGGGGAACUC

>ppe-miR482c-5p MIMAT0027283

GGAAUGGGCUGUUUGGGAUG

>ppe-miR8125 MIMAT0031533

CAGGAAAGAAUGUGAUGAGUA

>ppe-miR6291b MIMAT0027333

CUUACCACAUUUUUAUACCAU

>ppe-miR162 MIMAT0031440

UCGAUAAACCUCUGCAUCCAG

>ppe-miR8129-3p MIMAT0031555

AUAAUAAUGUCCGGAUGUCAA

>ppe-miR2111b MIMAT0031518

UAAUCUGCAUCCUGAGGUUUA

>ppe-miR395c MIMAT0031481

CUGAAGUGUUUGGGGGAACUC

>ppe-miR7122b-5p MIMAT0031529

UUAUACAAUGAAAUCACGGUCG

>ppe-miR8131-5p MIMAT0031558

AUUUCAGCUAAGUUGAGUUGU

>ppe-miR398a-5p MIMAT0031173

GGAGCGACCUGGGAUCACAUG

>ppe-miR6274b-3p MIMAT0031553

UUGUGUUAUUGGCCGAAAAUAG

>ppe-miR399k MIMAT0031507

UGCCAAAGAAGAGUUGCCCUA

>ppe-miR535b MIMAT0031515

UGACGACGAGAGAGAGCACGC

>ppe-miR6289 MIMAT0027322

UCCUUUGAAUGGUUAGGCUCA

>ppe-miR482b-5p MIMAT0031166

GGAAUGGGAGGAUUGGGAAAA

>ppe-miR156b MIMAT0031429

UGACAGAAGAAAGAGAGCAC

>ppe-miR399b MIMAT0031498

UCUGCCAAAGGAGAAUUGCCC

>ppe-miR6267c-3p MIMAT0031543

UAGAGAGAUGGUCAGCAAUGU

>ppe-miR172d MIMAT0031472

GGAAUCUUGAUGAUGCUGCAG

>ppe-miR6267a MIMAT0027291

UAGAGAGGUGGUACAAUUGUG

>ppe-miR395h MIMAT0031486

CUGAAGUGUUUGGGGGAACUC

>ppe-miR166e MIMAT0031449

UCGGACCAGGCUUCAUUCCCC

>rgl-miR7801 MIMAT0032242

UACGAGAUGAAACACAGUUUG

>rgl-miR5142 MIMAT0020653

AUAUUGAUUGAUAAGUGAU

>rgl-miR5141 MIMAT0020652

AGACCCGACGCGACUGACAGAUAA

>rgl-miR7803b-3p MIMAT0032253

UACACGUGUCAAUCAUCUAU

>rgl-miR7797a MIMAT0032238

AAGACGGAAUCAAACCUCAA

>rgl-miR5140 MIMAT0020651

GCUGGUGAAGAUUUGGUG

>rgl-miR5577 MIMAT0023002

AGAAGCUGAGAAUCACUUUUU

>rgl-miR5574 MIMAT0022999

UUUAAUCAGAAUCUAAAAGCA

>rgl-miR7807b-5p MIMAT0032258

UAACUAUAUGAAAAUCUCAAUU

>rgl-miR7808 MIMAT0032254

AAGGAUGCUCGAUUCAGAAGAA

>rgl-miR7798 MIMAT0032239

AGGGAGUGUUUGCAAAAACU

>rgl-miR7797b MIMAT0032260

UUUGAUUUCGUCUUACAUUUUUC

>rgl-miR7804-5p MIMAT0032245

AGGGGUGUUCAUCGAAUCGAAUU

>rgl-miR5575 MIMAT0023000

UGGAUUUUGAACGUUUCGGUG

>rgl-miR7811 MIMAT0032257

UGAAUGGAGAUACGGAAUGAAGC

>rgl-miR5573 MIMAT0022998

GAGUAGUCGACCUGAGAUGGA

>rgl-miR7802 MIMAT0032243

AGGGAGUGUUUGCAAUCACUAAA

>rgl-miR7972 MIMAT0032261

UUGUCAGGCUUGUUAUUCUCC

>rgl-miR5578 MIMAT0023003

CAUGUGGCAUGAUGGCAGAAAGU

>rgl-miR7809 MIMAT0032255

UCCCAUUGCAUCAGCGGACACA

>rgl-miR7805-3p MIMAT0032248

UAUUCAUUUACACCAAAUUUGG

>rgl-miR7803a MIMAT0032244

UACGGAUAAUUGACACGUGUAUA

>rgl-miR7804-3p MIMAT0032246

UUUAAUCGAAUGAACAUUUUAAA

>rgl-miR5139 MIMAT0020650

AAACCUGGCUCUGAUACCA

>rgl-miR7807a-3p MIMAT0032251

UUGGGAUUUGCAUACAGUUAC

>rgl-miR7803b-5p MIMAT0032252

GGAUGAUUGCCACGUGUAUA

>rgl-miR7799 MIMAT0032240

AGUGGAAUAGGAGAUCUCAA

>rgl-miR164 MIMAT0023004

UGGGGAAGAAGAGCACAUGAA

>rgl-miR7807b-3p MIMAT0032259

UUGAGAUUUUCAUAUAGUUACU

>rgl-miR7810 MIMAT0032256

AGAGGAAGAGUUUUCUGGCUC

>rgl-miR7800 MIMAT0032241

UAUUUUUGUGUCGUUAUGGUC

>rgl-miR5576 MIMAT0023001

AGAAGUUGGCAUUUGCAAACACU

>rgl-miR7807a-5p MIMAT0032250

AACUAUAUGAAAAUCUCAAUU

>rgl-miR5138 MIMAT0020649

AAAAAUCGUUAGGCGCUA

>rgl-miR7806 MIMAT0032249

UAGAAGAUGUCCACAUGAGCA

>rgl-miR5137 MIMAT0020648

AGCGGAGAAGACGAUGGGCU

>rgl-miR7805-5p MIMAT0032247

AAAUUUGGUGUAGUGAAUAGU

>smi-miR12112 MIMAT0049006

CGAUCUUGAUACCACCAAUGG

>ssl-miR397 MIMAT0022518

UCAUUGAGUGCAGCGUUGAUG

>ssl-miR398 MIMAT0022519

UGUGUUCUCAGGUCACCCCUC

>ssl-miR399 MIMAT0022520

UGCCAAAGGAGAAUUGCCCGG

>ssl-miR828 MIMAT0022521

UCUUGCUCAAAUGAGUAUUCCA

>ssl-miR156 MIMAT0022506

UGACAGAAGAGAGUGAGCACA

>ssl-miR166b MIMAT0022510

UCGGACCAGGCUUCAUUCCCC

>ssl-miR169 MIMAT0022511

UAGCCAAGGAUGACUUGCCUA

>ssl-miR396 MIMAT0022517

UUCCACAGCUUUCUUGAACUG

>ssl-miR164a MIMAT0022507

UGGAGAAGCAGGGCACGUGCA

>ssl-miR166a MIMAT0022509

UCGGACCAGGCUUCAUUCCUC

>ssl-miR171a MIMAT0022512

UUGAGCCGUGCCAAUAUCACG

>ssl-miR395 MIMAT0022516

GGGAAAUGUUUGGGGAAACUU

>ssl-miR948 MIMAT0022522

UGUGGCUGUGUGGGUUCCGG

>ssl-miR164b MIMAT0022508

UGGAGAAGCAGGGCACGUGCA

>ssl-miR171b MIMAT0022513

UUGAGCCGCGCCAAUAUCACU

>ssl-miR172 MIMAT0022514

AGAAUCUUGAUGAUGCUGCAU

>ssl-miR394 MIMAT0022515

UUGGCAUUCUGUCCACCUCC

>ssl-miR1078 MIMAT0022523

CUUGAUUGAUUCAAUUGUGAU

>sly-miR6026 MIMAT0023610

UUCUUGGCUAGAGUUGUAUUGC

>sly-miR9472-5p MIMAT0035449

UUUCAGUAGACGUUGUGAAUA

>sly-miR169b MIMAT0007919

UAGCCAAGGAUGACUUGCCUG

>sly-miR164a-5p MIMAT0033973

UGGAGAAGCAGGGCACGUGCA

>sly-miR7981e MIMAT0042015

AAGUGUGUCUCUGAGAUUUCGGAU

>sly-miR9477-5p MIMAT0035471

UAUCCGUUGUUCCCUUUUCCUACC

>sly-miR9473-3p MIMAT0035462

AAACGAGUUCAGAUUUACAGC

>sly-miR9470-5p MIMAT0035439

UGAAAUCCAUGAGCCUAAACU

>sly-miR9469-5p MIMAT0035435

CCACAUAAGAAGACCGAAUUC

>sly-miR169e-5p MIMAT0035475

UAGCCAAGGAUGACUUGCCUUU

>sly-miR482e-5p MIMAT0020765

UGUGGGUGGGGUGGAAAGAUU

>sly-miR10539 MIMAT0042033

CUUGGAACCACAGUUACCACC

>sly-miR9475-3p MIMAT0035466

CUACAAUGUAGAGAUCGUUUU

>sly-miR482b MIMAT0023602

UCUUGCCUACACCGCCCAUGCC

>sly-miR7981b MIMAT0042030

ACCCCUUUUUAGCUUACGUGGCAC

>sly-miR396a-5p MIMAT0035455

UUCCACAGCUUUCUUGAACUG

>sly-miR394-3p MIMAT0035438

AGGUGGGCAUACUGUCAACA

>sly-miR10537 MIMAT0042025

AUUUACCCCAAGUUCGUUGUC

>sly-miR482e-3p MIMAT0032124

UCUUUCCUACUCCUCCCAUACC

>sly-miR5300 MIMAT0020764

UCCCCAGUCCAGGCAUUCCAAC

>sly-miR166c-3p MIMAT0035444

UCGGACCAGGCUUCAUUCCUC

>sly-miR391 MIMAT0022688

ACGCAGGAGAGAUGAUGCUGGA

>sly-miR390b-3p MIMAT0035480

CGCUAUCCAUCCUGAGUUUCA

>sly-miR9471a-5p MIMAT0035447

CAGGUGCUCACUCAGCUAAUA

>sly-miR1916 MIMAT0007908

AUUUCACUUAGACACCUCAA

>sly-miR398a MIMAT0042014

UAUGUUCUCAGGUCGCCCCUG

>sly-miR167b-5p MIMAT0035457

UAAAGCUGCCAGCAUGAUCUGG

>sly-miR5304 MIMAT0020768

UCAAUGCUACAUACUCAUCCC

>sly-miR164b-3p MIMAT0033976

CACGUGUUCUCCUUCUCCAAC

>sly-miR1919a MIMAT0007911

ACGAGAGUCAUCUGUGACAGG

>sly-miR156a MIMAT0009138

UUGACAGAAGAUAGAGAGCAC

>sly-miR9469-3p MIMAT0035436

AUUCGGUCUUCUUAUGUGGAC

>sly-miR10534 MIMAT0042020

CAAAAUACCCUUGUCAUCCAA

>sly-miR10540 MIMAT0042034

AUAAUAACUAUUAGUUGAAUG

>sly-miR156d-3p MIMAT0035452

GCUCACUGCUCUAUCUGUCACC

>sly-miR168a-3p MIMAT0031130

CCUGCCUUGCAUCAACUGAAU

>sly-miR6023 MIMAT0023592

UUCCAUGAAAGAGUUUUUGGAU

>sly-miR9477-3p MIMAT0035472

UUGGGAAAGGGAACAACUGAUAGU

>sly-miR9479-3p MIMAT0035478

GAGAAUGGUAGAGGGUCGGACC

>sly-miR9476-5p MIMAT0035469

UCUAGUCCUGCAUCUUUUUUU

>sly-miR171b-3p MIMAT0007923

UUGAGCCGUGCCAAUAUCACG

>sly-miR1918 MIMAT0007910

UGUUGGUGAGAGUUCGAUUCUC

>sly-miR156e-3p MIMAT0035454

GCUUACUCUCUAUCUGUCACC

>sly-miR396b MIMAT0035481

UUCCACAGCUUUCUUGAACUU

>sly-miR164b-5p MIMAT0033975

UGGAGAAGCAGGGCACGUGCA

>sly-miR482d-5p MIMAT0035459

GGAGUGGGUGGGAUGGAAAAA

>sly-miR319c-3p MIMAT0035432

UUGGACUGAAGGGAGCUCCUU

>sly-miR10529 MIMAT0042009

AACGAGUGAGACUUGCUCAGUUGG

>sly-miR403-3p MIMAT0035434

CUAGAUUCACGCACAAGCUCG

>sly-miR172b MIMAT0009144

AGAAUCUUGAUGAUGCUGCAU

>sly-miR390a-3p MIMAT0035468

CGCUAUCCAUCCUGAGUUUUA

>sly-miR172c MIMAT0042004

AGAAUCUUGAUGAUGCUGCAG

>sly-miR171e MIMAT0035482

UUGAGCCGCGUCAAUAUCUCU

>sly-miR5302b-5p MIMAT0035445

UGAAAUGCUAUAGUUGGAAAGU

>sly-miR395b MIMAT0007927

CUGAAGUGUUUGGGGGAACUCC

>sly-miR10533 MIMAT0042019

UCUUAUGAAUUCUAGGUCUUCU

>sly-miR171d MIMAT0007925

UUGAGCCGCGCCAAUAUCAC

>sly-miR9476-3p MIMAT0035470

AAAAAGAUGCAGGACUAGACC

>sly-miR156e-5p MIMAT0035453

UGAUAGAAGAGAGUGAGCAC

>sly-miR168b-3p MIMAT0031132

CCCGCCUUGCAUCAACUGAAU

>sly-miR319a MIMAT0009145

CUUGGACUGAAGGGAGCUCC

>sly-miR156c MIMAT0009140

UUGACAGAAGAUAGAGAGCAC

>sly-miR530 MIMAT0042011

AGGUGUAGGUGUUCAUGCAGA

>sly-miR390a-5p MIMAT0035467

AAGCUCAGGAGGGAUAGCACC

>sly-miR6025 MIMAT0042023

UACCAAUAAUUGAGAUAACAUC

>sly-miR167b-3p MIMAT0035458

AGGUCAUCUAGCAGCUUCAAU

>sly-miR10535a MIMAT0042022

UUGGCAUAAGUUUGUGAAAGCCGG

>sly-miR166b MIMAT0007916

UCGGACCAGGCUUCAUUCCCC

>sly-miR1919c-3p MIMAT0007913

ACGAGAGUCAUCUGUGACAGG

>sly-miR10542 MIMAT0042037

UAGAAGAAUCAUAUAUACCCCUA

>sly-miR397-3p MIMAT0037335

UCAACGCUAAACUCGAUCAUG

>sly-miR482a MIMAT0020769

UUUCCAAUUCCACCCAUUCCUA

>sly-miR6027-5p MIMAT0032133

AUGGGUAGCACAAGGAUUAAUG

>sly-miR5303 MIMAT0020767

UUUUUGAAGAGUUCGAGCAAC

>sly-miR159b MIMAT0042036

UUGGAAAGAAGGGAGCUCUAC

>sly-miR9473-5p MIMAT0035461

UGGCUGUAAAUCUAAACUCGU

>sly-miR156b MIMAT0009139

UUGACAGAAGAUAGAGAGCAC

>sly-miR482c MIMAT0023603

UCUUGCCAAUACCGCCCAUUCC

>sly-miR169d MIMAT0007921

UAGCCAAGGAUGACUUGCCUA

>sly-miR10536 MIMAT0042024

AGACAUGUUCUAAUCGUCAGCUUC

>sly-miR162 MIMAT0009142

UCGAUAAACCUCUGCAUCCAG

>sly-miR166c-5p MIMAT0035443

GGGAUGUUGUCUGGCUCGACA

>sly-miR397-5p MIMAT0007928

AUUGAGUGCAGCGUUGAUGA

>sly-miR403-5p MIMAT0035433

CGUUUGUGCGUGAAUCUAACA

>sly-miR399 MIMAT0009146

UGCCAAAGGAGAGUUGCCCUA

>sly-miR482d-3p MIMAT0035460

UUUCCUAUUCCACCCAUGCCAA

>sly-miR171c MIMAT0007924

UAUUGGUGCGGUUCAAUGAGA

>sly-miR7981c MIMAT0042021

UCGAAAUCUCAGAGACACACUUAU

>sly-miR319b MIMAT0035430

UUGGACUGAAGGGAGCUCCCU

>sly-miR393 MIMAT0042012

AUCAUGCGAUCUCUUCGGAAU

>sly-miR319d MIMAT0042007

AGGAAACUGUUUAGUCCAACC

>sly-miR164a-3p MIMAT0033974

CAUGUGCCUGUUUUCCCCAUC

>sly-miR7981a MIMAT0042029

ACCCCUUUUCGGCCUACGUGGCAC

>sly-miR9471b-3p MIMAT0035442

UUGGCUGAGUGAGCAUCACUG

>sly-miR319c-5p MIMAT0035431

AGAGCUUCCUUCAGCCCACUC

>sly-miR10530 MIMAT0042013

ACGUCCCUUCCCCAUCGUUCAACA

>sly-miR827 MIMAT0042010

UUAGAUGAACAUCAACAAACA

>sly-miR1919b MIMAT0007912

ACGAGAGUCAUCUGUGACAGG

>sly-miR10528 MIMAT0042006

AAUGCAAUGUCAUAUACCAUC

>sly-miR156d-5p MIMAT0035451

UGACAGAAGAGAGUGAGCAC

>sly-miR1919c-5p MIMAT0032040

UGUCGCAGAUGACUUUCGCCC

>sly-miR10541 MIMAT0042035

AGUCACUUUGAUGAUUGUCAAACA

>sly-miR169c MIMAT0007920

CAGCCAAGGAUGACUUGCCGA

>sly-miR5302b-3p MIMAT0035446

UUUUCAACUAUAGCAUUAUUUU

>sly-miR171a MIMAT0007922

UGAUUGAGCCGUGCCAAUAUC

>sly-miR6024 MIMAT0023594

UUUUAGCAAGAGUUGUUUUACC

>sly-miR9472-3p MIMAT0035450

UUCACAAUCUCUGCUGAAAAA

>sly-miR9479-5p MIMAT0035477

UCCAGUCCUCUACCCUUCUCCA

>sly-miR6027-3p MIMAT0023611

UGAAUCCUUCGGCUAUCCAUAA

>sly-miR159 MIMAT0009141

UUUGGAUUGAAGGGAGCUCUA

>sly-miR160a MIMAT0007914

UGCCUGGCUCCCUGUAUGCCA

>sly-miR168a-5p MIMAT0031129

UCGCUUGGUGCAGGUCGGGAC

>sly-miR9471b-5p MIMAT0035441

GAGGUGCUCACUCAGCUAAUA

>sly-miR9474-5p MIMAT0035463

UGUAGAAGUCAUGAAUAAAAUG

>sly-miR390b-5p MIMAT0035479

AAGCUCAGGAGGGAUAGCGCC

>sly-miR9474-3p MIMAT0035464

UUUUGUUCGCAGAUACUACAGU

>sly-miR10538 MIMAT0042032

AGAUUGAUAUACGUUACUCACAGU

>sly-miR169e-3p MIMAT0035476

UGGCAAGCAUCUUUGGCGACU

>sly-miR477-3p MIMAT0035484

AGUUCUUGUAGGGUGAGACAAC

>sly-miR1917 MIMAT0007909

AUUAAUAAAGAGUGCUAAAGU

>sly-miR408 MIMAT0042005

ACGGGGACGAGCCAGAGCAUG

>sly-miR399b MIMAT0042016

GGGCUACUCUCUAUUGGCAUG

>sly-miR7981d MIMAT0042031

UCGAAAUCUCAGAGACACACUUAU

>sly-miR394-5p MIMAT0035437

UUGGCAUUCUGUCCACCUCC

>sly-miR171f MIMAT0042038

UAUUGGCCUGGUUCACUCAGA

>sly-miR9478-5p MIMAT0035473

GCUUAAAUAUGUAGAUCGAACU

>sly-miR10532 MIMAT0042018

AAGUGUGUCUCUGAGAUUUCGGGC

>sly-miR172a MIMAT0009143

AGAAUCUUGAUGAUGCUGCAU

>sly-miR9471a-3p MIMAT0035448

UUGGCUGAGUGAGCAUCACGG

>sly-miR9475-5p MIMAT0035465

AACGAUCUCUACAUUGUAGGC

>sly-miR477-5p MIMAT0035483

UGUCUCUCCCUCAAGGGCUCC

>sly-miR10535b MIMAT0042027

UUGGCAUGAGUUUGUGAAAGCCGG

>sly-miR171b-5p MIMAT0037334

AUAUUGGUGCGGUUCAAUUAG

>sly-miR7981f MIMAT0042026

AAGUGUGUCUCUGGAAUUUCGGGC

>sly-miR172d MIMAT0042008

GGAAUCUUGAUGAUGCUGCAG

>sly-miR167a MIMAT0007917

UGAAGCUGCCAGCAUGAUCUA

>sly-miR9478-3p MIMAT0035474

UUCGAUGACAUAUUUGAGCCU

>sly-miR168b-5p MIMAT0031131

UCGCUUGGUGCAGGUCGGGAC

>sly-miR166a MIMAT0007915

UCGGACCAGGCUUCAUUCCCC

>sly-miR169f MIMAT0042028

UAGGCGUUGUCUGAGGCUAAC

>sly-miR6022 MIMAT0023590

UGGAAGGGAGAAUAUCCAGGA

>sly-miR395a MIMAT0007926

CUGAAGUGUUUGGGGGAACUCC

>sly-miR5302a MIMAT0020766

AAACGAGGUUUGUUACUUUGG

>sly-miR9470-3p MIMAT0035440

UUUGGCUCAUGGAUUUUAGC

>sly-miR396a-3p MIMAT0035456

GUUCAAUAAAGCUGUGGGAAG

>sly-miR10531 MIMAT0042017

UGGGGUCCUAGUAGAGUCGGUUC

>sly-miR169a MIMAT0007918

CAGCCAAGGAUGACUUGCCGG

>stu-miR391-3p MIMAT0031275

GCAUCAUACUCCUGCAUAUU

>stu-miR8032a-5p MIMAT0030936

UGGUCGGCAUGACUCCCGAGGU

>stu-miR8008a MIMAT0030901

AUUUCCAGAAAAGCGACGGACAGU

>stu-miR1919-3p MIMAT0031239

ACGAGAGUCAUCUGUGACAGG

>stu-miR8005b-5p MIMAT0031191

ACUCUAAAUUUUAAAUUCUAAAUC

>stu-miR167d-3p MIMAT0031310

GAUCAUGUGGUUGCUUCACC

>stu-miR395b MIMAT0031263

CUGAAGUGUUUGGGGGAACUC

>stu-miR166d-5p MIMAT0031254

AGAAUGUCGUCUGGUUCGAGA

>stu-miR8032a-3p MIMAT0031208

AGUGUGAGUCGGUGUGAUUAGG

>stu-miR1886b MIMAT0030870

AUGGUAUCGUGAGAUGAAAUCAGC

>stu-miR1886i-5p MIMAT0030925

AUGAGAUGAAAUUAGCGUUUGGAU

>stu-miR7999-3p MIMAT0030889

ACGACCCGUAGAACUGCCCACGAC

>stu-miR390-5p MIMAT0031343

AAGCUCAGGAGGGAUAGCACC

>stu-miR8030-3p MIMAT0031207

UUAAAACCAAAUCAACCCAAAU

>stu-miR1886i-3p MIMAT0031202

UUUACGUUGAUUUCAUCUCAUGA

>stu-miR167b-3p MIMAT0031306

GAUCAUGUGGCAGCAUCACC

>stu-miR482e-5p MIMAT0031163

AGUGGGUGGUGUGGUAAGAUU

>stu-miR395i MIMAT0031270

CUGAAGUGUUUGGGGGAACUC

>stu-miR6026-5p MIMAT0031164

AAUACAACUAUUGCCAAGACAA

>stu-miR399h MIMAT0030971

GGGCUACUCUCUAUUGGCAUG

>stu-miR7983-5p MIMAT0031183

UAAAGUCUUUAGCGACAUUGGUUC

>stu-miR479 MIMAT0031311

UGAGCCGAACCAAUAUCACUC

>stu-miR169e-5p MIMAT0031367

UAGCCAAGGAUGACUUGCCU

>stu-miR8038b-3p MIMAT0031217

GUUCAACUUGCUCACUUGGAG

>stu-miR7997c MIMAT0030887

AUAUUGCUCGGACUCUUCAAAAAU

>stu-miR8000 MIMAT0030890

ACACCGAAGAACUGACACCGAAGA

>stu-miR8019-3p MIMAT0030916

AAAAGAAUGACCUGGUUUGACUUG

>stu-miR8006-5p MIMAT0030898

UAGUUUUUGGACGACAGGGGCACC

>stu-miR169e-3p MIMAT0031368

GCAAGUUAUCCUGGCUAUC

>stu-miR8032e-5p MIMAT0030940

UGGUCGGCAUGACUCCCGAGGU

>stu-miR169c-3p MIMAT0031364

GCAGUCUCCUUGGCUACC

>stu-miR169b-3p MIMAT0031362

GCAGUCUCCUUGGCUACU

>stu-miR7984c-5p MIMAT0030861

ACGAUACCAAACUUUAUGAAGGAC

>stu-miR6025 MIMAT0023608

UACCAACAAUUGAGAUAACAUC

>stu-miR162b-3p MIMAT0031282

UCGAUAAACCUCUGCAUCCAG

>stu-miR8004 MIMAT0030894

AGGGGUUGUGUAUGUGUUUGGCCU

>stu-miR482a-5p MIMAT0031161

GGAAUUGGUGGAUUGGAAAGC

>stu-miR390-3p MIMAT0031344

CGCUAUCCAUCUUGAGUUUUA

>stu-miR7992-5p MIMAT0030868

UUUGACAAUGCACAUCUAGACACU

>stu-miR171a-3p MIMAT0031257

UGAUUGAGCCGUGCCAAUAUC

>stu-miR399k-3p MIMAT0031315

CGCCAAAGGAGAGCUGCCCUG

>stu-miR8040-3p MIMAT0030953

CUUAUAAUUGUAAUUAUGAUC

>stu-miR408a-3p MIMAT0031338

UGCACAGCCUCUUCCCUGGUU

>stu-miR8032c MIMAT0030938

UGGUCGGCAUGACUCCCGAGGU

>stu-miR162a-3p MIMAT0031280

UCGAUAAACCUCUGCAUCCAG

>stu-miR8032g-3p MIMAT0031213

AGUGUGAGUCGGUGCGAUUAGG

>stu-miR7990a MIMAT0030864

UUCAAAUGAUCGUAACUUUGGCCU

>stu-miR399i-5p MIMAT0031260

GGGCUACACUCUAUUGGCAUG

>stu-miR164-3p MIMAT0030973

CAUGUGCUCUAGCUCUCCAGC

>stu-miR399g-3p MIMAT0031233

CGCCAAAGGGGAGCUGCCCUA

>stu-miR7980b-5p MIMAT0031205

GUCCAAACACUGAUUCCAUCUCAU

>stu-miR8045 MIMAT0030959

AUUGAUAGUUGAGGUGUGUUU

>stu-miR172e-3p MIMAT0031295

AGAAUCUUGAUGAUGCUGCAU

>stu-miR8005c MIMAT0030897

UUUAGAGUUUAAGGUUUAGAGUUU

>stu-miR8022 MIMAT0030921

UUUAAAUGAGAAUUUUGGACUAUU

>stu-miR7987 MIMAT0030860

ACACUAGUUGAUCUUAUUGAUGAC

>stu-miR8032b-5p MIMAT0030937

UGGUCGGCAUGACUCCCGAGGU

>stu-miR5303f MIMAT0030904

AUUUUUGGAGAAUCUGACACGGGU

>stu-miR7980b-3p MIMAT0030930

GAGAUGGAAUCAGUGUUUGGACAU

>stu-miR477a-5p MIMAT0031326

CCUCUCCCUCAAGGGCUUCUC

>stu-miR8033-5p MIMAT0030943

UUCCAAAGCUGCAGAAAUGAGU

>stu-miR399l-3p MIMAT0031317

CGCCAAAGGAGAGCUGCCCUG

>stu-miR8002-3p MIMAT0031190

AUUCCAUUAUUAUCAAGAAAAAAG

>stu-miR169g MIMAT0031371

UAGCCAAGGAUGACUUGCCU

>stu-miR395g MIMAT0031268

CUGAAGUGUUUGGGGGAACUC

>stu-miR7993d MIMAT0030878

AUAUUUUAUGUGGUUAACUUAACU

>stu-miR172a-3p MIMAT0031291

AGAAUCUUGAUGAUGCUGCAU

>stu-miR8041b-3p MIMAT0030955

AUGAUGUAUAGCAAAGAGCCU

>stu-miR169a-3p MIMAT0031360

GCAGUCUCCUUGGCUACU

>stu-miR477b-5p MIMAT0031328

ACUCUCCCUCAAAGGCUUCUG

>stu-miR8048-3p MIMAT0030972

AGAUGGACAUGCUAAUGAACA

>stu-miR399d-3p MIMAT0031230

UGCCAAAGGAGAGCUGCCCUG

>stu-miR399n-5p MIMAT0031320

GGGCUACUCUCUAUUGGCAUG

>stu-miR399j-5p MIMAT0031312

GGGCUACUCUCUAUUGGCAUA

>stu-miR8051-5p MIMAT0030978

UAGUAUGGUAGAAAGAUUCA

>stu-miR169d-5p MIMAT0031365

UAGCCAAGGAUGACUUGCCU

>stu-miR167a-5p MIMAT0031307

UGAAGCUGCCAGCAUGAUCUA

>stu-miR164-5p MIMAT0031235

UGGAGAAGCAGGGCACAUGCU

>stu-miR8032b-3p MIMAT0031209

AGUGUGAGUCGGUGCGAUUAGG

>stu-miR399f-3p MIMAT0031232

UGCCAAAGGAGAGCUGCCCUG

>stu-miR827-3p MIMAT0031273

UUAGAUGAACAUCAACAAACA

>stu-miR8024a-5p MIMAT0031201

UUGAAGAAUUUAAAGACUUCAACU

>stu-miR399g-5p MIMAT0030970

GGGCUACUCUCUAUUGGCAUG

>stu-miR1886f MIMAT0030874

AUGGUAUCGUGAGAUGAAAUCAGC

>stu-miR156k-5p MIMAT0031357

UGACAGAAGAGAGUGAGCAC

>stu-miR8041a-5p MIMAT0031221

GUGCUUUGCUAUUUUCAUUG

>stu-miR167c-5p MIMAT0031303

UGAAGCUGCCAGCAUGAUCUA

>stu-miR5303b MIMAT0030857

UUUUGGAGAAUCCGACACGCACCC

>stu-miR482b-5p MIMAT0031160

GGAGUGGGUGGCAUGGUAAGA

>stu-miR6024-5p MIMAT0031159

AGAAACAACACUUGCUAAAAGA

>stu-miR172e-5p MIMAT0031294

GCAACAUCAUCAAGAUUCACA

>stu-miR399c-3p MIMAT0031229

UGCCAAAGGAGAGCUGCCCUG

>stu-miR399c-5p MIMAT0030966

GGGCUACUCUCUAUUGGCAUG

>stu-miR172b-3p MIMAT0031287

AGAAUCUUGAUGAUGCUGCAU

>stu-miR8001b-3p MIMAT0031200

GGAUUUUCAUACUAAUUCCUAGAA

>stu-miR399m-3p MIMAT0031319

CGCCAAAGGAGAGCUGCCCUG

>stu-miR156i-5p MIMAT0031353

UGACAGAAGAGAGUGAGCAC

>stu-miR398a-3p MIMAT0031332

UAUGUUCUCAGGUCGCCCCUG

>stu-miR399b-5p MIMAT0030965

GGGCUACUCUCUAUUGGCAUG

>stu-miR7984c-3p MIMAT0031185

CCUUCAUAAAGUUUGGUAUCGUAA

>stu-miR399a-5p MIMAT0030964

GGGCUACUCUCUAUUGGCAUG

>stu-miR395a MIMAT0031262

CUGAAGUGUUUGGGGGAACUC

>stu-miR167b-5p MIMAT0031305

UGAAGCUGCCAGCAUGAUCUA

>stu-miR8032f-3p MIMAT0031212

AGUGUGAGUCGGUGCGAUUAGG

>stu-miR399e-3p MIMAT0031231

UGCCAAAGGAGAGCUGCCCUG

>stu-miR8046-3p MIMAT0031225

CGCUGAAAUUUCGAUCAUAAU

>stu-miR7994b-3p MIMAT0030880

AUAUUAUACUUGGGCAUAAACUCC

>stu-miR166c-3p MIMAT0031253

UCGGACCAGGCUUCAUUCCCC

>stu-miR7122-5p MIMAT0031224

UUAUACAGAGAAACCGCUGUCG

>stu-miR391-5p MIMAT0031274

UACGCAGGAGAGAUGAUGCUG

>stu-miR5303d MIMAT0030858

UUUUGGAGAAUCCGACACGCACCC

>stu-miR8018 MIMAT0030915

ACGAACCGUAGAUCCCAUCCGUGG

>stu-miR8007a-5p MIMAT0030899

AUGUGGCACUUUUCGGAUUUUGAG

>stu-miR172d-5p MIMAT0031292

GGAGCAUCAUCAAGAUUCACA

>stu-miR8005a MIMAT0030895

UUUAGAGUUUAAGGUUUAGAGUUU

>stu-miR1886a MIMAT0030869

AUGGUAUCGUGAGAUGAAAUCAGC

>stu-miR160a-5p MIMAT0031283

UGCCUGGCUCCCUGUAUGCCA

>stu-miR6149-3p MIMAT0031226

UGAUUCAGGUUUGUAUGCAAAC

>stu-miR7996c MIMAT0030884

AUGUGGUACAUAUGAAAUUUGAAA

>stu-miR395j MIMAT0031271

CUGAAGUGUUUGGGGGAACUC

>stu-miR8021 MIMAT0030920

AUUCAAGGCUCAAACUCGAGACCU

>stu-miR399l-5p MIMAT0031316

GGGCUACUCUCUAUUGGCAUG

>stu-miR171d-3p MIMAT0031302

UUGAGCCGUGCCAAUAUCACG

>stu-miR6022 MIMAT0023589

UGGAAGGGAGAAUAUCCAGGA

>stu-miR171e MIMAT0031345

UGAUUGAGCCGUGCCAAUAUC

>stu-miR482d-5p MIMAT0031162

CGUGAGUGGUGGGGUAAGAUA

>stu-miR8033-3p MIMAT0031214

UCAAUUCUGCAGCUUUAGGAGU

>stu-miR7995 MIMAT0030881

UUACACGUAGACAAGUUGACCAUU

>stu-miR7984d-5p MIMAT0030928

AUCCGAACUUUGACCGAAAUUGCU

>stu-miR156c MIMAT0031298

UUGACAGAAGAUAGAGAGCAC

>stu-miR7992-3p MIMAT0031186

UGUCUAGAUGUGCAUUUCAAAGU

>stu-miR8011a-5p MIMAT0031195

UUGUGUGAGGUUUCUUUUUGUUUC

>stu-miR156d-3p MIMAT0031300

GCUCUCUAUGCUUCUGUCAUCA

>stu-miR7991a MIMAT0030865

AGGAGGUCGGAAUUUUUAAUGAAU

>stu-miR8001b-5p MIMAT0030917

AUGGGGAUUAGUAUGAAAAUUUGC

>stu-miR156h-3p MIMAT0031352

GCUCACUGCUCUAUCUGUCACC

>stu-miR7982b MIMAT0030850

AAGUUGGAUGAUAAUAAUAUAUAU

>stu-miR7988 MIMAT0030862

AACGGAAAAGGGCCAAAAAUACCC

>stu-miR397-5p MIMAT0031341

AUUGAGUGCAGCGUUGAUGAC

>stu-miR160a-3p MIMAT0031284

GCGUAUGAGGAGCCAAGCAUA

>stu-miR399a-3p MIMAT0031227

UGCCAAAGGAGAGCUGCCCUG

>stu-miR319a-3p MIMAT0031278

UUGGACUGAAGGGAGCUCCCU

>stu-miR7122-3p MIMAT0030960

ACAGCGUUUCUCUGUAUAACC

>stu-miR530 MIMAT0031330

UCUGCAUUUGCACCUGCACCU

>stu-miR7980a MIMAT0030847

AUGAGAUGAAGUCAAUGUUUGGAC

>stu-miR1886c MIMAT0030871

AUGGUAUCGUGAGAUGAAAUCAGC

>stu-miR5303a MIMAT0030856

UUUUGGAGAAUCCGACACGCACCC

>stu-miR399j-3p MIMAT0031313

CGCCAAAGGAGAGCUGCCCUG

>stu-miR8032d-3p MIMAT0031210

AGUGUGAGUUGGUGCGAUUAGG

>stu-miR482a-3p MIMAT0023598

UUUCCAAUUCCACCCAUUCCUA

>stu-miR7998 MIMAT0030888

ACGGACCGUAGAUCAAUCCACAGU

>stu-miR408a-5p MIMAT0031337

ACAGGGACGAGGCAGCGCAUG

>stu-miR8011a-3p MIMAT0030907

AAUAAAAAGAAGCCUCACACAACU

>stu-miR169a-5p MIMAT0031359

UAGCCAAGGAUGACUUGCCU

>stu-miR7999-5p MIMAT0031189

CUGGGUCACUUCUACGGGUCCUUC

>stu-miR8008b MIMAT0030931

AAACCCAGAAAAGCGACGGACAGU

>stu-miR8048-5p MIMAT0031234

CUCAUUAGCAUCUCCAUCUUG

>stu-miR8019-5p MIMAT0031199

AGGGAAGCAGGUCAUUCUUUAUG

>stu-miR8025-3p MIMAT0031203

UUUAAUUGCAUGCCAAGUGUGUGG

>stu-miR827-5p MIMAT0031272

UUUGUUGAUGGUCAUCUAUUC

>stu-miR5303j MIMAT0031241

AAUAUUUUUGAAGAGUCUGAGCAA

>stu-miR399i-3p MIMAT0031261

UGCCAAAGGAGAGUUGCCCUA

>stu-miR319b MIMAT0031276

UUGGACUGAAGGGAGCUCCU

>stu-miR1886g-5p MIMAT0030900

GAGAUGAGAUCAAUGUUUGGACAU

>stu-miR8050-3p MIMAT0030976

UGACUUGAGAUUCCUACUUGG

>stu-miR5303g MIMAT0031242

AUAUUUUUGAAGAGUCUGAGCAAC

>stu-miR395f MIMAT0031267

CUGAAGUGUUUGGGGGAACUC

>stu-miR8006-3p MIMAT0031192

UGCCCUGCCGUCCAAAAAAUAGA

>stu-miR7982a MIMAT0030849

AAGUUGGAUGAUAAUAAUAUAUAU

>stu-miR8050-5p MIMAT0031238

AAGUAGGAAUCAAGGUCAAU

>stu-miR7993b-3p MIMAT0030876

AUAUUUUAUGUGGUUAACUUAACU

>stu-miR399o-3p MIMAT0031323

CGCCAAAGGAGAGCUGCCCUG

>stu-miR477a-3p MIMAT0031327

GAAGCUCUAGCAGGGAGAGCCA

>stu-miR6026-3p MIMAT0023609

UUCUUGGCUAGAGUUGUAUUGC

>stu-miR8015-5p MIMAT0030912

UAUUGGAUAUUGAAAAUGAAACUU

>stu-miR166a-3p MIMAT0031250

UCGGACCAGGCUUCAUUCCCC

>stu-miR7981-5p MIMAT0031182

GUUAAUUAAACUAUGGUCCUAUUA

>stu-miR8014-5p MIMAT0031196

AUUGUUUCAUAUUGUAUUGUAUUU

>stu-miR8038b-5p MIMAT0030949

CCUUGUGAGUAAGUUGAAUCUC

>stu-miR482e-3p MIMAT0023601

UCUUGCCAAUACCGCCCAUUCC

>stu-miR8005b-3p MIMAT0030896

UUUAGAGUUUAAGGUUUAGAGUUU

>stu-miR8044-5p MIMAT0031223

UUUCAAAUAUGGUUGGAGAUG

>stu-miR8037 MIMAT0030947

AUAAUUUGGAGGAAUAGGAACC

>stu-miR8049-3p MIMAT0031237

CAUGUCUACAUGAGCCUGAUA

>stu-miR6149-5p MIMAT0030963

UUGCAACACACCUGAAUCGUC

>stu-miR7986 MIMAT0030859

AGUUUAAAACGUUACUGUCGGUAA

>stu-miR7984d-3p MIMAT0031204

CAAUUUCGGUCAAAGUUCGGAUAU

>stu-miR8028-5p MIMAT0031206

UCCUUAUGCUACAAUUGUGAACAA

>stu-miR7984b-3p MIMAT0031184

GGUCUUUCAUAAAAUUUGGUAUCG

>stu-miR3627-3p MIMAT0031246

AAGUGCCUCUGUCUUUCGACA

>stu-miR8007a-3p MIMAT0031193

CGAAAAAUGAAAAGUGCCACAUAA

>stu-miR8024a-3p MIMAT0030923

UUGGAGGAUUUGAAGAUUUCAACU

>stu-miR397-3p MIMAT0031342

CAUCAACGCUACACUCAAUCA

>stu-miR166d-3p MIMAT0031255

UCGGACCAGGCUUCAUUCCCC

>stu-miR6023 MIMAT0023591

UUCCAUGAAAGUGUUUUUGGAU

>stu-miR8007b-3p MIMAT0031197

CGAAAAAUGAAAAGUACCACAUAA

>stu-miR171d-5p MIMAT0031301

AGAUAUUGGUGCGGUUCAAUU

>stu-miR7993a MIMAT0030875

AUAUUUUAUGUGGUUAACUUAACU

>stu-miR8047 MIMAT0030962

CCAUUUUUUCGAAAUUAGACC

>stu-miR156a MIMAT0031296

UUGACAGAAGAUAGAGAGCAC

>stu-miR8024b MIMAT0030924

UUGGAGGAUUUGAAGAUUUCAACU

>stu-miR8001a MIMAT0030891

UCCUGGGGAUUAGUAUGAAAAUUC

>stu-miR169h MIMAT0031372

UAGCCAAGGAUGACUUGCCU

>stu-miR7981-3p MIMAT0030848

AUAGGACUUUAGUUUAGUUAAGGU

>stu-miR5303i MIMAT0031244

AUAUUUUUGAAGAGUCUGAGCAAC

>stu-miR171c-5p MIMAT0031258

UAUUGGCCUGGUUCACUCAGA

>stu-miR8032e-3p MIMAT0031211

AGUGUGAGUCGGUGUGAUUAGG

>stu-miR399e-5p MIMAT0030968

GGGCUACUCUCUAUUGGCAUG

>stu-miR7985 MIMAT0030854

CGGGCUUGCCUAGAACGGGUUACC

>stu-miR8015-3p MIMAT0031198

GUUUCAUUUUCAAGGUCCAAUAGC

>stu-miR1886e MIMAT0030873

AUGGUAUCGUGAGAUGAAAUCAGC

>stu-miR7984a MIMAT0030852

AUACCGAACUUUGGAAAUGACCUU

>stu-miR7993b-5p MIMAT0031187

UUAAGUUAACCACAUAAAAUAUGU

>stu-miR8041b-5p MIMAT0031222

GUGCUUUGCUAUUUUCAUUG

>stu-miR5303e MIMAT0030903

UUUUGGAGAAUCUGACACGGGUGU

>stu-miR8020 MIMAT0030919

AAUUUCAUUGAGUAUGUUGUUGUU

>stu-miR8039 MIMAT0030951

UUUCCUAUCUGAACUAUCACC

>stu-miR156d-5p MIMAT0031299

UUGACAGAAGAUAGAGAGCAC

>stu-miR8023 MIMAT0030922

UUUGGCACAAUUUCAUUGGCAACC

>stu-miR156i-3p MIMAT0031354

GCUCACUGCUCUAUCUGUCACC

>stu-miR7997a MIMAT0030885

AUGCUGCUCGGACUCUUCAAA

>stu-miR8017 MIMAT0030914

AUCCAAGUGAAGUGUAUCGUCUCA

>stu-miR7994b-5p MIMAT0031188

AGUUUAUGCCCAAGUAUAUAAUAUAU

>stu-miR171a-5p MIMAT0031256

UAUUGGCCUGGUUCACUCAGA

>stu-miR156f-3p MIMAT0031348

CUCACUUCUCUUUCUGUCAAUC

>stu-miR7990b MIMAT0030918

GAAUUUUCAAAUGAUCGUAACUUU

>stu-miR169f-5p MIMAT0031369

UAGCCAAGGAUGACUUGCCU

>stu-miR167d-5p MIMAT0031309

UGAAGCUGCCAGCAUGAUCUA

>stu-miR7989 MIMAT0030863

ACAAAUAAGUCCAUUACCUGAACC

>stu-miR171b-3p MIMAT0031248

UUGAGCCGCGUCAAUAUCUCU

>stu-miR1886h MIMAT0030905

AUUUUACGUUGAUUUCAUCUCAUG

>stu-miR482c MIMAT0023596

UUUCCUAUUCCACCCAUGCCAA

>stu-miR8025-5p MIMAT0030926

ACAUACUCGACAUGCAAUUAAAUU

>stu-miR395h MIMAT0031269

CUGAAGUGUUUGGGGGAACUC

>stu-miR169d-3p MIMAT0031366

GCAGGUCAUCUUUAGCUAACU

>stu-miR8038a-5p MIMAT0030948

CCUUGUGAGUAAGUUGAAUCUC

>stu-miR8014-3p MIMAT0030910

AUGAAUACAAUGUUUGGAUAAAUU

>stu-miR166b MIMAT0031251

UCGGACCAGGCUUCAUUCCUC

>stu-miR169c-5p MIMAT0031363

UAGCCAAGGAUGACUUGCCU

>stu-miR395e MIMAT0031266

CUGAAGUGUUUGGGGGAACUC

>stu-miR7991b MIMAT0030866

AGGAGGUCGGAAUUUUUAAUGAAU

>stu-miR384-3p MIMAT0031374

AGGGGGCCAAAGUGCCAAAC

>stu-miR8028-3p MIMAT0030932

GUUCAUAAUUAUAGUAUAAGGAUG

>stu-miR399o-5p MIMAT0031322

GGGCUACUCUCUAUUGGCAUG

>stu-miR7996b MIMAT0030883

AUGUGGUACAUAUGAAAUUUGAAA

>stu-miR166a-5p MIMAT0031249

GGAAUGUUGUCUGGCUCGAGG

>stu-miR395d MIMAT0031265

CUGAAGUGUUUGGGGGAACUC

>stu-miR167c-3p MIMAT0031304

GGUCAUGCUCGGACAGCCUCACU

>stu-miR1886g-3p MIMAT0031194

UUUCAUAUUGAUUUCAUCUCAU

>stu-miR8034 MIMAT0030944

UAUGACAAACACUGCAAAAACU

>stu-miR398b-5p MIMAT0031333

GAGUGUGCCUUAGAACACAGGU

>stu-miR5303h MIMAT0031243

AACAUUUUUGAAGAGUCUGAGCAA

>stu-miR6027 MIMAT0023612

UGAAUCCUUCGGCUAUCCAUAA

>stu-miR172c-3p MIMAT0031289

AGAAUCUUGAUGAUGCUGC

>stu-miR7979 MIMAT0030846

AGGUACAUGAACUCUAACGAGGCA

>stu-miR8032g-5p MIMAT0030942

UGGUCGGCAUGACUCCCGAGGU

>stu-miR408b-5p MIMAT0031339

ACGGGGACGAGACAGAGCAUG

>stu-miR482d-3p MIMAT0023600

UCUUGCCUACACCGCCCAUGCC

>stu-miR172b-5p MIMAT0031286

GCAGCACCAUCAAGAUUCACA

>stu-miR8029 MIMAT0030933

AGCCAUUUUUCUUUGUUUUGGAGC

>stu-miR156g-5p MIMAT0031349

UGACAGAAGAGAGUGAGCAC

>stu-miR8040-5p MIMAT0031220

UCAUAAUUACAAUUAUAAGCC

>stu-miR7994a MIMAT0030879

AUAUUAUACUUGGGCAUAAACUCC

>stu-miR393-3p MIMAT0031325

AUCAUGCGAUCUCUUCGGAAU

>stu-miR172d-3p MIMAT0031293

GGAAUCUUGAUGAUGCUGCAG

>stu-miR3627-5p MIMAT0031245

UCGCAGGAGAGAUGGCACUUAG

>stu-miR398b-3p MIMAT0031334

UUGUGUUCUCAGGUCACCCCU

>stu-miR156h-5p MIMAT0031351

UGACAGAAGAGAGUGAGCAC

>stu-miR8046-5p MIMAT0030961

UAUGAUCGAAGUUUCAAUGAC

>stu-miR1919-5p MIMAT0030977

UGUCGCAGAUGACUUUCGCCC

>stu-miR160b MIMAT0031285

UGCCUGGCUCCCUGUAUGCCA

>stu-miR8043 MIMAT0030957

UGAUAUAAUUGGACUUUGGCC

>stu-miR7984b-5p MIMAT0030853

AUACCGAACUUUGGAAAUGACCUU

>stu-miR8044-3p MIMAT0030958

UCUCCAGCGAUAUUUGAAACU

>stu-miR395c MIMAT0031264

CUGAAGUGUUUGGGGGAACUC

>stu-miR398a-5p MIMAT0031331

GGGUUGAUUUGAGAACAUAUG

>stu-miR399m-5p MIMAT0031318

GGGCUACUCUCUAUUGGCAUG

>stu-miR399k-5p MIMAT0031314

GGGCUACUCUCUAUUGGCAUG

>stu-miR399b-3p MIMAT0031228

UGCCAAAGGAGAGCUGCCCUG

>stu-miR156j-5p MIMAT0031355

UGACAGAAGAGAGUGAGCAC

>stu-miR156b MIMAT0031297

UUGACAGAAGAUAGAGAGCAC

>stu-miR8027 MIMAT0030929

AUCUCGAGAUAAGUUAUUCUGGAC

>stu-miR482b-3p MIMAT0023597

UUACCGAUUCCCCCCAUUCCAA

>stu-miR8038a-3p MIMAT0031216

GUUCAACUUGCUCACUUGGAG

>stu-miR6024-3p MIMAT0023593

UUUUAGCAAGAGUUGUUUUCCC

>stu-miR7983-3p MIMAT0030851

ACUAAUGCCGGUAAAGACUUUAAC

>stu-miR8049-5p MIMAT0030975

CAAGGCUCAUGCAGACAUGCA

>stu-miR399n-3p MIMAT0031321

CGCCAAAGGAGAGCUGCCCUG

>stu-miR171c-3p MIMAT0031259

UGAUUGAGCCGUGUCAAUAUC

>stu-miR319a-5p MIMAT0031277

AGAGCUUUCUUCGGUCCACAC

>stu-miR156g-3p MIMAT0031350

GCUUACUCUCUAUCUGUCACC

>stu-miR172a-5p MIMAT0031290

GUAGCAUAAUCAAGAUUCACA

>stu-miR8009 MIMAT0030902

AUUUCCAGAAAAGCGACGGACAGU

>stu-miR8036-3p MIMAT0030946

UAUGUCUUUCCGAUGCCUCCCA

>stu-miR319-5p MIMAT0031219

AGGAAACUGUUUAGUCCAACC

>stu-miR156j-3p MIMAT0031356

GCUCACUGCUCUAUCUGUCACC

>stu-miR477b-3p MIMAT0031329

GAGGUCUUUCGAGUGAGAGUGA

>stu-miR172c-5p MIMAT0031288

AGCAUCUUCAAGAUUCACA

>stu-miR8010 MIMAT0030906

AUAGGACCCUAGUUAAAUUUAGGU

>stu-miR169f-3p MIMAT0031370

GCAAGCAUCCUUGGCGACU

>stu-miR8041a-3p MIMAT0030954

AUGAUGUAUAGCAAAGAGCCU

>stu-miR393-5p MIMAT0031324

UCCAAAGGGAUCGCAUUGAUCC

>stu-miR8011b-3p MIMAT0030950

UUCGUGAGACAAAAAGAAGCCU

>stu-miR8036-5p MIMAT0031215

GGAGGAAUCGAAAGAUAUAAG

>stu-miR319-3p MIMAT0030952

UUGGACUGAAGGGUUCCCUUC

>stu-miR8051-3p MIMAT0031240

UAUUUCUUCUACCAUACUAUU

>stu-miR162b-5p MIMAT0031281

GGAGGCAGCGGUUCAUCGAUC

>stu-miR8003 MIMAT0030893

AUUUCGGUAUACAAAUGGGAUGAC

>stu-miR156e MIMAT0031346

UGACAGAAGAGAGUGAGCAC

>stu-miR8012 MIMAT0030908

AUGACUUUAAGUCGCGUCUGGCCC

>stu-miR156f-5p MIMAT0031347

CUGACAGAAGAGAGUGAGCA

>stu-miR8042 MIMAT0030956

AUUAGACUGAAGUGCUGAUCU

>stu-miR8011b-5p MIMAT0031218

ACUCAUUUUUGUCUCACAAAAA

>stu-miR156k-3p MIMAT0031358

GCUCACUGCUCUAUCUGUCACC

>stu-miR396-5p MIMAT0031335

UUCCACAGCUUUCUUGAACUU

>stu-miR8013 MIMAT0030909

AGAAGAAAAUCGCUCCGUCAGAAG

>stu-miR384-5p MIMAT0031373

UUGGCAUUCUGUCCACCUCC

>stu-miR167a-3p MIMAT0031308

GAUCAUGUGGCAGCCUCACC

>stu-miR399f-5p MIMAT0030969

GGGCUACUCUCUAUUGGCAUG

>stu-miR8002-5p MIMAT0030892

UUUUUCGUGAUAAUAAUGGAAUCA

>stu-miR8007b-5p MIMAT0030911

AUGUGACACUUUUUGAAUUUCGAG

>stu-miR171b-5p MIMAT0031247

AGAUAUUGAUGUGGCUCAAUC

>stu-miR1886d MIMAT0030872

AUGGUAUCGUGAGAUGAAAUCAGC

>stu-miR7997b MIMAT0030886

AUGCUGCUCGGACUCUUCAAA

>stu-miR8035 MIMAT0030945

UCCAUCUUCAAUAUCACUUUCU

>stu-miR399d-5p MIMAT0030967

GGGCUACUCUCUAUUGGCAUG

>stu-miR166c-5p MIMAT0031252

GGAAUGUUGUUUGGCUCGAGG

>stu-miR169b-5p MIMAT0031361

UAGCCAAGGAUGACUUGCCU

>stu-miR5304-3p MIMAT0031236

AGAUGAGUAUGGUGCAUUGGA

>stu-miR5303c MIMAT0030855

UUUUGGAGAAUCCGACACGCACCC

>stu-miR396-3p MIMAT0031336

GUCCAAGAAAGCUGUGGGAAA

>stu-miR7991c MIMAT0030867

AGGAGGUCGGAAUUUUUAAUGAAU

>stu-miR8031 MIMAT0030935

UUAGACACCUCAACUAAGACUUG

>stu-miR7996a MIMAT0030882

AUGUGGUACAUAUGAAAUUUGAAA

>stu-miR8030-5p MIMAT0030934

UUGGGUUGGUUUGGUCUCGGGUU

>stu-miR8016 MIMAT0030913

AUUUUUGAAUGGAAGGCCCAUGUG

>stu-miR7993c MIMAT0030877

AUAUUUUAUGUGGUUAACUUAACU

>stu-miR162a-5p MIMAT0031279

GGAGGCAGCGGUUCAUCGAUC

>stu-miR8026 MIMAT0030927

AUGUAGAGAAUAUGUGGUAACCCU

>stu-miR8032f-5p MIMAT0030941

UGGUCGGCAUGACUCCCGAGGU

>stu-miR5304-5p MIMAT0030974

CAAUGCAACAUACUCAUCACC

>stu-miR408b-3p MIMAT0031340

UGCACUGCCUCUUCCCUGGCU

>stu-miR8032d-5p MIMAT0030939

UGGUCGGCAUGACUCCCGAGGU

>tcc-miR171e MIMAT0020409

UGAUUGAGCCGUGCCAAUAUC

>tcc-miR395a MIMAT0020425

CUGAAGUGUUUGGGGGAACUC

>tcc-miR399a MIMAT0020435

CGCCAAAGGAGAGUUGCCCUG

>tcc-miR396a MIMAT0020427

UUCCACAGCUUUCUUGAACUG

>tcc-miR169n MIMAT0020404

UGAGUCAAGAAUGACUUGCCG

>tcc-miR319 MIMAT0020418

UUUGGACUGAAGGGAGCUCCU

>tcc-miR169e MIMAT0020395

CAGCCAAGGAUGACUUGCCGA

>tcc-miR399b MIMAT0020436

UGCCAAAGGAGAUUUGCCCGG

>tcc-miR168 MIMAT0020390

UCGCUUGGUGCAGGUCGGGAA

>tcc-miR169k MIMAT0020401

CAGCCAAGGAUGACUUGCCGG

>tcc-miR399e MIMAT0020439

CGCCAAAGGAGAAUUGCCCUG

>tcc-miR397 MIMAT0020432

UCAUUGAGUGCAGCGUUGAUG

>tcc-miR169l MIMAT0020402

CAGCCAAGGAUGACUUGCCGG

>tcc-miR530b MIMAT0020447

UGCAUUUGCACCUGCACCUU

>tcc-miR394a MIMAT0020423

UUGGCAUUCUGUCCACCUCC

>tcc-miR171g MIMAT0020411

UGAUUGAGCCGUGCCAAUAUC

>tcc-miR399f MIMAT0020440

UGCCAGAGGAGAUUUGCCCUG

>tcc-miR167c MIMAT0020389

UGAAGCUGCCAGCAUGAUCUU

>tcc-miR403a MIMAT0020444

UUAGAUUCACGCACAAACUCG

>tcc-miR164c MIMAT0020382

UGGAGAAGCAGGGCACAUGCU

>tcc-miR171b MIMAT0020406

AGAUUGAGCCGCGCCAAUAUC

>tcc-miR169m MIMAT0020403

UGAGCCAAGGAUGACUUGCCG

>tcc-miR398b MIMAT0020434

UGUGUUCUCAGGUCACCCCUU

>tcc-miR171d MIMAT0020408

UGAUUGAGCCGUGCCAAUAUC

>tcc-miR156f MIMAT0020374

UUGACAGAAGAUAGAGAGCAC

>tcc-miR156e MIMAT0020373

UUGACAGAAGAUAGAGAGCAC

>tcc-miR171f MIMAT0020410

UGAUUGAGCCGUGCCAAUAUC

>tcc-miR156g MIMAT0020375

UGACAGAAGAGAGUGAGCAC

>tcc-miR172b MIMAT0020414

AGAAUCUUGAUGAUGCUGCAU

>tcc-miR396e MIMAT0020431

UUCCACAGCUUUCUUGAACUU

>tcc-miR169i MIMAT0020399

UAGCCAAGGAUGAGUUGCCUG

>tcc-miR169f MIMAT0020396

AAGCCAAGAAUGACUUGCCUG

>tcc-miR535 MIMAT0020448

UGACAACGAUAGAGAGCACGC

>tcc-miR393b MIMAT0020422

UCCAAAGGGAUCGCAUUGAUC

>tcc-miR164b MIMAT0020381

UGGAGAAGCAGGGCACGUGCA

>tcc-miR166a MIMAT0020383

UCGGACCAGGCUUCAUUCCCC

>tcc-miR169d MIMAT0020394

UAGCCAAGGAUGACUUGCCUA

>tcc-miR156c MIMAT0020371

UGACAGAAGAGAGUGAGCAC

>tcc-miR169b MIMAT0020392

CAGCCAAGGAUGACUUGCCGG

>tcc-miR169a MIMAT0020391

CAGCCAAGGAUGACUUGCCGA

>tcc-miR169g MIMAT0020397

UAGCCAGGGAUGACUUGCCUA

>tcc-miR169j MIMAT0020400

UAGCCAAGGAUGACUUGCCUG

>tcc-miR396b MIMAT0020428

UUCCACAGCUUUCUUGAACUG

>tcc-miR398a MIMAT0020433

UGUGUUCUCAGGUCGCCCCUG

>tcc-miR166c MIMAT0020385

UCGGACCAGGCUUCAUUCCUC

>tcc-miR166b MIMAT0020384

UCGGACCAGGCUUCAUUCCC

>tcc-miR171c MIMAT0020407

AGAUUGAGCCGCGCCAAUAUC

>tcc-miR403b MIMAT0020445

UUAGAUUCACGCACAAACUCG

>tcc-miR399i MIMAT0020443

UGCCAAAGGAGAGUUGCCCUG

>tcc-miR169c MIMAT0020393

CAGCCAAGGAUGACUUGCCGA

>tcc-miR162 MIMAT0020379

UCGAUAAACCUCUGCAUCCAG

>tcc-miR164a MIMAT0020380

UGGAGAAGCAGGGCACGUGCA

>tcc-miR156d MIMAT0020372

UGACAGAAGAGAGUGAGCAC

>tcc-miR390b MIMAT0020420

AAGCUCAGGAGGGAUAGCGCC

>tcc-miR393a MIMAT0020421

UCCAAAGGGAUCGCAUUGAUCC

>tcc-miR160c MIMAT0020378

UGCCUGGCUCCCUGGAUGCCA

>tcc-miR156b MIMAT0020370

UGACAGAAGAGAGUGAGCAC

>tcc-miR160a MIMAT0020376

UGCCUGGCUCCCUGAAUGCCA

>tcc-miR171h MIMAT0020412

UGAUUGAGCCGUGCCAAUAUC

>tcc-miR390a MIMAT0020419

AAGCUCAGGAGGGAUAGCGCC

>tcc-miR172c MIMAT0020415

GGAAUCUUGAUGAUGCUGCAU

>tcc-miR399d MIMAT0020438

UGCCAAAGGAGAUUUGCCCGG

>tcc-miR169h MIMAT0020398

UAGCCAAGGAUGACUUGCCUG

>tcc-miR2111 MIMAT0020450

UAAUCUGCAUCCUGAGGUUUA

>tcc-miR399g MIMAT0020441

UGCCAAAGGAGAAUUGCCCUG

>tcc-miR156a MIMAT0020369

UGACAGAAGAGAGAGAGCACA

>tcc-miR394b MIMAT0020424

UUGGCAUUCUGUCCACCUCC

>tcc-miR395b MIMAT0020426

CUGAAGUGUUUGGGGGAACUC

>tcc-miR530a MIMAT0020446

UGCAUUUGCACCUGCACCUC

>tcc-miR171a MIMAT0020405

UGAUUGAGCCGCGCCAAUAUC

>tcc-miR172a MIMAT0020413

GGAAUCUUGAUGAUGCUGCA

>tcc-miR827 MIMAT0020449

UUAGAUGACCAUCAACAAACA

>tcc-miR399c MIMAT0020437

UGCCAAUGGAGAUUUGCCCAG

>tcc-miR167a MIMAT0020387

UGAAGCUGCCAGCAUGAUCUA

>tcc-miR396d MIMAT0020430

UUCCACGGCUUUCUUGAACUU

>tcc-miR172d MIMAT0020416

AGAAUCCUGAUGAUGCUGCAU

>tcc-miR166d MIMAT0020386

UCGGACCAGGCUUCAUUCCCC

>tcc-miR399h MIMAT0020442

UGCCAAAGGAGAUUUGCCCCG

>tcc-miR172e MIMAT0020417

AGAAUCUUGAUGAUGCUGCAU

>tcc-miR167b MIMAT0020388

UGAAGCUGCCAGCAUGAUCUA

>tcc-miR396c MIMAT0020429

UUCCACAGCUUUCUUGAACUU

>tcc-miR160b MIMAT0020377

UGCCUGGCUCCCUGUAUGCCA

>vvi-miR395c MIMAT0005713

CUGAAGUGUUUGGGGGAACUC

>vvi-miR3631c MIMAT0018033

CAUGUUGACAUCAUCCAAUAUA

>vvi-miR169r MIMAT0005687

UGAGUCAAGGAUGACUUGCCG

>vvi-miR169y MIMAT0005677

UAGCGAAGGAUGACUUGCCUA

>vvi-miR164d MIMAT0005661

UGGAGAAGCAGGGCACGUGCA

>vvi-miR171g MIMAT0006555

UUGAGCCGAACCAAUAUCACC

>vvi-miR3629a-3p MIMAT0018022

GGCUGCUGAGAAAAUGUAGGA

>vvi-miR395j MIMAT0005720

CUGAAGUGUUUGGGGGAACUC

>vvi-miR3637-3p MIMAT0018048

UUUCGACAAGACACAAUGCAUAAA

>vvi-miR3636-5p MIMAT0018045

UCGGUUUGCUUCUUUGAUAGAUUC

>vvi-miR394b MIMAT0005710

UUGGCAUUCUGUCCACCUCC

>vvi-miR159a MIMAT0005648

CUUGGAGUGAAGGGAGCUCUC

>vvi-miR166d MIMAT0005665

UCGGACCAGGCUUCAUUCCCC

>vvi-miR408 MIMAT0005733

AUGCACUGCCUCUUCCCUGGC

>vvi-miR3640-3p MIMAT0018054

AUCGAAAAGGCAUCAUCAAUCAGG

>vvi-miR3633a-5p MIMAT0018037

GGAAUGGAUGGUUAGGAGAG

>vvi-miR3632-3p MIMAT0018036

UUUCCCAGACCCCCAAUACCAA

>vvi-miR319e MIMAT0006556

UUUGGACUGAAGGGAGCUCCU

>vvi-miR3624-5p MIMAT0018011

UAGUAUGCUGCUGUCUUUAGA

>vvi-miR3634-3p MIMAT0018040

UUUCCGACUCGCACUCAUGCCGU

>vvi-miR2950-3p MIMAT0018010

UGGUGUGCACGGGAUGGAAUA

>vvi-miR3631a-3p MIMAT0018030

UAUAUUGGAUGAUGUCAACAA

>vvi-miR159c MIMAT0005650

UUUGGAUUGAAGGGAGCUCUA

>vvi-miR3632-5p MIMAT0018035

GGAUUGGGGGCCGAUGGAAAGG

>vvi-miR535a MIMAT0005735

UGACAACGAGAGAGAGCACGC

>vvi-miR169s MIMAT0005688

CAGCCAAGGAUGACUUGCCGG

>vvi-miR3638-3p MIMAT0018050

GCAACAAGCAUGAAAAGGCACACC

>vvi-miR166f MIMAT0005667

UCGGACCAGGCUUCAUUCCCC

>vvi-miR535c MIMAT0005738

UGACAACGAGAGAGAGCACGC

>vvi-miR3624-3p MIMAT0018012

UCAGGGCAGCAGCAUACUACU

>vvi-miR3629c MIMAT0018024

GGCUGCUGAGAAAAUGUAGGA

>vvi-miR3625-5p MIMAT0018013

UUCCAGCAGUCAUCUCCAAGG

>vvi-miR845c MIMAT0006581

AGGCUCUGAUACCAAUUGAUG

>vvi-miR393a MIMAT0006557

UCCAAAGGGAUCGCAUUGAUC

>vvi-miR156b MIMAT0005641

UGACAGAAGAGAGUGAGCAC

>vvi-miR399f MIMAT0006567

UGCCGAAGGAGAUUUGUCCUG

>vvi-miR169b MIMAT0006545

UGAGCCAAGGAUGGCUUGCCG

>vvi-miR395k MIMAT0005721

CUGAAGUGUUUGGGGGAACUC

>vvi-miR3626-5p MIMAT0018015

GGUAGUCGCUGUGAAAUUGAA

>vvi-miR169a MIMAT0005676

CAGCCAAGGAUGACUUGCCGG

>vvi-miR167d MIMAT0005673

UGAAGCUGCCAGCAUGAUCUA

>vvi-miR398b MIMAT0006563

UGUGUUCUCAGGUCGCCCCUG

>vvi-miR828a MIMAT0006577

UCUUGCUCAAAUGAGUAUUCCA

>vvi-miR395e MIMAT0005715

CUGAAGUGUUUGGGGGAACUC

>vvi-miR403a MIMAT0006569

UUAGAUUCACGCACAAACUCG

>vvi-miR399b MIMAT0005729

UGCCAAAGGAGAGUUGCCCUG

>vvi-miR3633b-3p MIMAT0018042

GUUCCCAUGCCAUCCAUUCCUA

>vvi-miR172a MIMAT0005699

UGAAUCUUGAUGAUGCUACAU

>vvi-miR166b MIMAT0005663

UCGGACCAGGCUUCAUUCC

>vvi-miR3636-3p MIMAT0018046

GUCUGUCGGAGAAGCAAGUCGGAG

>vvi-miR160c MIMAT0005653

UGCCUGGCUCCCUGUAUGCCA

>vvi-miR169i MIMAT0006547

GAGCCAAGGAUGACUGGCCGU

>vvi-miR3631b-3p MIMAT0018032

UGUUGGAUGAUGUCAAUAAGU

>vvi-miR403c MIMAT0006571

UUAGAUUCACGCACAAACUCG

>vvi-miR156a MIMAT0005640

UGACAGAAGAGAGGGAGCAC

>vvi-miR166e MIMAT0005666

UCGGACCAGGCUUCAUUCCCC

>vvi-miR169p MIMAT0005686

GAGCCAAGGAUGACUUGCCGG

>vvi-miR171j MIMAT0037305

UGAUUGAGCCGUGCCAAUAUC

>vvi-miR3630-3p MIMAT0018028

UUUGGGAAUCUCUCUGAUGCAC

>vvi-miR159b MIMAT0005649

CUUGGAGUGAAGGGAGCUCUC

>vvi-miR166g MIMAT0005668

UCGGACCAGGCUUCAUUCCCC

>vvi-miR167a MIMAT0005670

UGAAGCUGCCAGCAUGAUCUG

>vvi-miR156i MIMAT0005647

UUGACAGAAGAUAGAGAGCAC

>vvi-miR3633b-5p MIMAT0018041

GGAAUGGGUGGCUGGGAUCUA

>vvi-miR169e MIMAT0005680

UAGCCAAGGAUGACUUGCCUGC

>vvi-miR171a MIMAT0005691

UGAUUGAGCCGUGCCAAUAUC

>vvi-miR477b-3p MIMAT0018026

CGAAGUCUUUGGGGAGAGUGG

>vvi-miR171c MIMAT0005693

UGAUUGAGCCGUGCCAAUAUC

>vvi-miR3627-5p MIMAT0018017

UUGUCGCAGGAGAGACGGCACU

>vvi-miR156g MIMAT0005646

UUGACAGAAGAUAGAGAGCAC

>vvi-miR3639-3p MIMAT0018052

GAGCUUUUGGCUUCUCAGAAGUCA

>vvi-miR399d MIMAT0006566

UGCCAAAGGAGAUUUGCUCGU

>vvi-miR479 MIMAT0005734

UGUGGUAUUGGUUCGGCUCAUC

>vvi-miR3629a-5p MIMAT0018021

CGCAUUUUCUCAGCAGCCAAG

>vvi-miR171h MIMAT0005697

UGGUUGAGCCGCGCCAAUAUC

>vvi-miR2111-5p MIMAT0016364

UAAUCUGCAUCCUGAGGUCUA

>vvi-miR845e MIMAT0006583

UGGCUCUGAUACCAAUUGAUG

>vvi-miR845b MIMAT0006580

UAGCUCUGAUACCAAUUGAUA

>vvi-miR395a MIMAT0005711

CUGAAGUGUUUGGGGGAACUC

>vvi-miR172c MIMAT0005701

GGAAUCUUGAUGAUGCUGCAG

>vvi-miR169j MIMAT0005683

CAGCCAAGGAUGACUUGCCGG

>vvi-miR3627-3p MIMAT0018018

UCGCCGCUCUCCUGUGACAAG

>vvi-miR172d MIMAT0005702

UGAGAAUCUUGAUGAUGCUGCAU

>vvi-miR397a MIMAT0006561

UCAUUGAGUGCAGCGUUGAUG

>vvi-miR2950-5p MIMAT0018009

UUCCAUCUCUUGCACACUGGA

>vvi-miR403f MIMAT0006574

UUAGAUUCACGCACAAACUCG

>vvi-miR395h MIMAT0005718

CUGAAGUGUUUGGGGGAACUC

>vvi-miR396a MIMAT0005724

UUCCACAGCUUUCUUGAACUA

>vvi-miR171d MIMAT0005694

UGAUUGAGCCGUGCCAAUAUC

>vvi-miR160b MIMAT0005652

UGCCUGGCUCCCUGAAUGCCAUC

>vvi-miR156e MIMAT0005644

UGACAGAGGAGAGUGAGCAC

>vvi-miR167e MIMAT0005674

UGAAGCUGCCAGCAUGAUCUA

>vvi-miR164b MIMAT0005659

UGGAGAAGCAGGGCACAUGCU

>vvi-miR169u MIMAT0005690

UGAGUCAAGGAUGACUUGCCG

>vvi-miR169l MIMAT0006548

GAGCCAAGGAUGACUUGCCGU

>vvi-miR169c MIMAT0005678

CAGCCAAGGAUGACUUGCCGG

>vvi-miR166c MIMAT0005664

UCGGACCAGGCUUCAUUCCCC

>vvi-miR3631d MIMAT0018034

CAUGUUGACAUCAUCCAAUAUA

>vvi-miR3623-5p MIMAT0018007

UCACAAGUUCAUCCAAGCACCA

>vvi-miR395g MIMAT0005717

CUGAAGUGUUUGGGGGAACUC

>vvi-miR164c MIMAT0005660

UGGAGAAGCAGGGCACGUGCA

>vvi-miR390 MIMAT0005707

AAGCUCAGGAGGGAUAGCGCC

>vvi-miR477a MIMAT0006575

AUCUCCCUCAAAGGCUUCCAA

>vvi-miR169g MIMAT0005682

CAGCCAAGGAUGACUUGCCGA

>vvi-miR160e MIMAT0005656

UGCCUGGCUCCCUGUAUGCCA

>vvi-miR399a MIMAT0005728

UGCCAAAGGAGAAUUGCCCUG

>vvi-miR169f MIMAT0005681

CAGCCAAGGAUGACUUGCCGA

>vvi-miR393b MIMAT0005708

UCCAAAGGGAUCGCAUUGAUC

>vvi-miR3631a-5p MIMAT0018029

CAUGUUGACAUCAUCCAAUAUA

>vvi-miR169n MIMAT0006549

GAGCCAAGGAUGACUUGCCGG

>vvi-miR394c MIMAT0006558

UUGGCAUUCUGUCCACCUCCAU

>vvi-miR395n MIMAT0006559

CUGAAGAGUCUGGAGGAACUC

>vvi-miR162 MIMAT0005657

UCGAUAAACCUCUGCAUCCAG

>vvi-miR396d MIMAT0005726

UUCCACAGCUUUCUUGAACUG

>vvi-miR169t MIMAT0005689

CGAGUCAAGGAUGACUUGCCG

>vvi-miR399e MIMAT0005730

UGCCAAAGGAGAUUUGCCCGG

>vvi-miR3638-5p MIMAT0018049

UGUGCCUUUUCGCGCUUGUUGCUA

>vvi-miR168 MIMAT0005675

UCGCUUGGUGCAGGUCGGGAA

>vvi-miR171b MIMAT0005692

UGAUUGAGCCGCGUCAAUAUC

>vvi-miR319f MIMAT0005705

UUGGACUGAAGGGAGCUCCCU

>vvi-miR394a MIMAT0005709

UUGGCAUUCUGUCCACCUCCAU

>vvi-miR2111-3p MIMAT0016365

GUCCUCUGGUUGCAGAUUACU

>vvi-miR156c MIMAT0005642

UGACAGAAGAGAGUGAGCAC

>vvi-miR3628-5p MIMAT0018019

AUGCGAGAGCCGUGCUUAGUA

>vvi-miR160a MIMAT0005651

UGCCUGGCUCCCUGAAUGCCAUC

>vvi-miR395l MIMAT0005722

CUGAAGUGUUUGGGGGAACUC

>vvi-miR396b MIMAT0005725

UUCCACAGCUUUCUUGAACU

>vvi-miR319b MIMAT0005703

UUGGACUGAAGGGAGCUCCCU

>vvi-miR535b MIMAT0005736

UGACAACGAGAGAGAGCACGC

>vvi-miR169o MIMAT0006550

GAGCCAAGGAUGACUUGCCGC

>vvi-miR171f MIMAT0005696

UUGAGCCGCGCCAAUAUCACU

>vvi-miR398a MIMAT0005727

UGUGUUCUCAGGUCACCCCUU

>vvi-miR169x MIMAT0006554

UAGCCAAGGAUGACUUGCCUA

>vvi-miR399i MIMAT0006568

CGCCAAAGGAGAGUUGCCCUG

>vvi-miR3630-5p MIMAT0018027

UGCAAGUGACGAUAUCAGACA

>vvi-miR3635-3p MIMAT0018044

AUUAUGUCCCACACAUGCCUC

>vvi-miR403b MIMAT0006570

UUAGAUUCACGCACAAACUCG

>vvi-miR398c MIMAT0006564

UGUGUUCUCAGGUCGCCCCUG

>vvi-miR319g MIMAT0005706

UUGGACUGAAGGGAGCUCCCA

>vvi-miR399g MIMAT0005731

UGCCAAAGGAGAUUUGCCCCU

>vvi-miR3629b MIMAT0018023

GGCUGCUGAGAAAAUGUAGGA

>vvi-miR3631b-5p MIMAT0018031

CAUGUUGACAUCAUCCAAUAUA

>vvi-miR3623-3p MIMAT0018008

UGGUGCUUGGACGAAUUUGCUA

>vvi-miR166h MIMAT0005669

UCGGACCAGGCUUCAUUCCCC

>vvi-miR395f MIMAT0005716

CUGAAGUGUUUGGGGGAACUC

>vvi-miR319c MIMAT0005704

UUGGACUGAAGGGAGCUCCCU

>vvi-miR169d MIMAT0005679

CAGCCAAGAAUGAUUUGCCGG

>vvi-miR172b MIMAT0005700

UGAAUCUUGAUGAUGCUACAC

>vvi-miR169h MIMAT0006546

UGAGCCAAGGAUGGCUUGCCG

>vvi-miR3633a-3p MIMAT0018038

UUCCUAUACCACCCAUUCCCUA

>vvi-miR156h MIMAT0006544

UGACAGAAGAGAGAGAGCAU

>vvi-miR3639-5p MIMAT0018051

AUUGACUUCUGAAAGGCUAAAAGC

>vvi-miR845d MIMAT0006582

UGGCUCUGAUACCAAUUGAUG

>vvi-miR395i MIMAT0005719

CUGAAGUGUUUGGGGGAACUC

>vvi-miR403d MIMAT0006572

UUAGAUUCACGCACAAACUCG

>vvi-miR169w MIMAT0006553

CAGCCAAGGAUGACUUGCCGG

>vvi-miR3640-5p MIMAT0018053

ACCUGAUUGGUGAUGCUUUUUUGG

>vvi-miR399h MIMAT0005732

UGCCAAAGGAGAAUUGCCCUG

>vvi-miR169q MIMAT0006551

GAGCCAAGGAUGACUUGCCGG

>vvi-miR3634-5p MIMAT0018039

GGCAUAUGUGUGACGGAAAGA

>vvi-miR395m MIMAT0005723

CUGAAGUGUUUGGGGGAACUC

>vvi-miR169k MIMAT0005684

CAGCCAAGGAUGACUUGCCGG

>vvi-miR482 MIMAT0006576

UCUUUCCUACUCCUCCCAUUCC

>vvi-miR167b MIMAT0005671

UGAAGCUGCCAGCAUGAUCUA

>vvi-miR3637-5p MIMAT0018047

AUUUAUGUAUUGUGUUUUGUCGGA

>vvi-miR3635-5p MIMAT0018043

GGCAUGUGUGGGGCAUAAUAG

>vvi-miR3625-3p MIMAT0018014

CGGGAGAUGACUACUGGAAGC

>vvi-miR169m MIMAT0005685

GAGCCAAGGAUGACUUGCCGG

>vvi-miR403e MIMAT0006573

UUAGAUUCACGCACAAACUCG

>vvi-miR171e MIMAT0005695

UGAUUGAGCCGCGCCAAUAUC

>vvi-miR156d MIMAT0005643

UGACAGAAGAGAGUGAGCAC

>vvi-miR156f MIMAT0005645

UUGACAGAAGAUAGAGAGCAC

>vvi-miR169v MIMAT0006552

AAGCCAAGGAUGAAUUGCCGG

>vvi-miR160d MIMAT0005654

UGCCUGGCUCCCUGUAUGCCA

>vvi-miR396c MIMAT0006560

UUCCACAGCUUUCUUGAACUG

>vvi-miR3628-3p MIMAT0018020

CUAAGCACAGCUCUCGCAUCC

>vvi-miR395b MIMAT0005712

CUGAAGUGUUUGGGGGAACUC

>vvi-miR166a MIMAT0005662

UCGGACCAGGCUUCAUUCC

>vvi-miR845a MIMAT0006579

UAGCUCUGAUACCAAUUGAUA

>vvi-miR395d MIMAT0005714

CUGAAGUGUUUGGGGGAACUC

>vvi-miR3626-3p MIMAT0018016

CUUCAAUUUCACAGCGACCAC

>vvi-miR164a MIMAT0005658

UGGAGAAGCAGGGCACGUGCA

>vvi-miR167c MIMAT0005672

UGAAGCUGCCAGCAUGAUCUC

>vvi-miR171i MIMAT0005698

UGAUUGAGCCGUGCCAAUAUC

>vvi-miR399c MIMAT0006565

UGCCAAAGGAGAGUUGCCCUG

>vvi-miR477b-5p MIMAT0018025

ACUCUUUCUCAAGGGCUUCUAG>seu-miR11047 MIMAT0043661

CACUGAUGAUGAGGGGAGACCG

>seu-miR11035 MIMAT0043649

CGUUUUACCGAUCGUUUGGU

>seu-miR11025 MIMAT0043637

UGAAGGGCUUUCAUGAUGGGCA

>seu-miR11032 MIMAT0043645

UGACGUUGUUGCAGGCAUGGAU

>seu-miR11040 MIMAT0043654

AACAAUUGUCAUGACACGGAC

>seu-miR11049 MIMAT0043663

UAAGCUGAUAGUUGUGGCC

>seu-miR11033 MIMAT0043646

CCGGUACCGGGGCCACGUCUGC

>seu-miR11048 MIMAT0043662

UGAAACUUGAGGAACAGGCAG

>seu-miR11042 MIMAT0043656

CUUUCCUUUUCAGUCGUCUGACA

>seu-miR11026 MIMAT0043638

UGGGCAAUGAUGUAGGCCGGC

>seu-miR11034 MIMAT0043648

UUUUGGAUUUGUAUGAUGCAUC

>seu-miR319 MIMAT0043647

CGUGAAUGAUGCGGGAGAUAG

>seu-miR11030b MIMAT0043643

AAAUUGAUUUUGGCACUCCUC

>seu-miR11027 MIMAT0043639

UGGUAGAACAUUCUAGAGUAGAA

>seu-miR11044 MIMAT0043658

UAGUAUUUGUUAGUUAGUGGGUC

>seu-miR11030a MIMAT0043642

AAAUUGAUUUUGGCACUCCUC

>seu-miR11021 MIMAT0043633

UAGCAAGCGGAACAUUUGCACA

>seu-miR11045 MIMAT0043659

UUUGAAAUUCUGGAAUUGAGACA

>seu-miR11036 MIMAT0043650

UUUGGAUGAAUAUGAGGAUGAA

>seu-miR11024 MIMAT0043636

UAAUAAUUGCAACACUUUGAC

>seu-miR11038 MIMAT0043652

GGGUUGGCCAACCCGAUCGGGUC

>seu-miR11043 MIMAT0043657

UUGCACAUAAUAAGGACGGUA

>seu-miR11029 MIMAT0043641

UGGACCAGGUCUAUAGUGGAGCA

>seu-miR11022 MIMAT0043634

CGAGGCUGUGGGUGGAGAGGU

>seu-miR11041 MIMAT0043655

UAAGCAUUUAGAACAUUCAUA

>seu-miR11028 MIMAT0043640

AGAAUAUGUUAUAUACUCUAUCA

>seu-miR11039 MIMAT0043653

UGGAAAUUGUUGCUAGUAAUUCA

>seu-miR11023 MIMAT0043635

UUUUACAUCGUUUACUUUGGG

>seu-miR11037 MIMAT0043651

ACAUAUUAAUGGCAACGAUUU

>seu-miR11031 MIMAT0043644

UUCUGGUGCUGGUUGUUCGAU

>seu-miR11046 MIMAT0043660

AUGUGUUUGGGCGAAAAUAGG

>pgi-miR6140d MIMAT0027264

CGUUGAUGUGGCAUACUUCACC

>pgi-miR482a MIMAT0027238

UCUUGCCAAUUCCUCCCAUUCC

>pgi-miR6136b MIMAT0027255

UCAUACAACCGUCGUCUAUAC

>pgi-miR6137a MIMAT0027257

AUGAAAAUUGUCGCUAUAGAUC

>pgi-miR6140a MIMAT0027261

AAUGUUUGUAGAAUAGUUUGUGUC

>pgi-miR6140b MIMAT0027262

AAUGUUUGUAGAAUAAUUUGUGUA

>pgi-miR6136a.2 MIMAT0027254

ACGGGUGAGUAAGAUAAGGGGUAU

>pgi-miR6135d MIMAT0027244

GGGUAAGUUGGUCAAUUGAC

>pgi-miR6135h MIMAT0027249

ACGUAAGUUGGUCAAUUGGC

>pgi-miR6141 MIMAT0027265

UAACUAAAUCUGGCCUGUAGCGGA

>pgi-miR6135c MIMAT0027243

GGGUAAGUUGGUCAAUUGAC

>pgi-miR6135j MIMAT0027251

AAUUGACUAAUAGAAUACUGACAC

>pgi-miR6137b MIMAT0027258

AUGAAAAUUGUCGCUAUAGAUC

>pgi-miR6140c MIMAT0027263

GCUGAGGUGGAGUAUGCCACAUC

>pgi-miR482b MIMAT0027239

UCUUGCCAAUUCCUCCCAUUCC

>pgi-miR6135a MIMAT0027241

UGGUAAGUUGGUCAAUUGGC

>pgi-miR6135k MIMAT0027256

CGUGUCGAUACUGUAUUGGU

>pgi-miR6142 MIMAT0027266

GACGAUUUUUUGGGCUAUGACGAC

>pgi-miR6135e-5p MIMAT0027245

GGGUAAGUUGGUCAAUUGAC

>pgi-miR6135f MIMAT0027247

GAGUAAGUUGGUCAAUUGGC

>pgi-miR6135i MIMAT0027250

AAUUGGCCAAUAGAAUACUGACAC

>pgi-miR6135b MIMAT0027242

UGGUAAGUUGGUCAAUUGGC

>pgi-miR6138 MIMAT0027259

UACGUUUGGAUUGAAGGAAUGAAA

>pgi-miR6143b-3p MIMAT0027269

CAGCACUGUAUUGAACAUGA

>pgi-miR6135g MIMAT0027248

GAGUAAGUUGGUCAAUUGGC

>pgi-miR6143b-5p MIMAT0027268

ACAAUGUCGACACGCAGGCGGAGA

>pgi-miR6139 MIMAT0027260

AAGAAUCAUUGGGAAGGGAAGAAA

>pgi-miR6136a.1 MIMAT0027253

UAGACGACGGUUGUAUGACCG

>pgi-miR6135e.2-5p MIMAT0027246

AAUUGACUAAUAGAAUACUGACAC

>pgi-miR6143a MIMAT0027267

AGUACUGUAUUGGGCAUGAAG

>pgi-miR391 MIMAT0027252

UGCAGGAGAGAUGACGCCCAUC

>pgi-miR2118 MIMAT0027240

UUUCCUAUUCCACCCAUCCCAU

>cca-miR6108d MIMAT0024556

AUAAGGAGAGUUAAGCUGAGAAGC

>cca-miR6106-3p MIMAT0024540

UUAAAUCCGGAUACUUGCAAC

>cca-miR171 MIMAT0024520

UGAUUGAGCCGCACCAAUAUC

>cca-miR6108e-3p MIMAT0024558

AUCAAGGGUUCAGAUCUCUUCUU

>cca-miR6108f MIMAT0024559

UAUGGUGAGAAGGGUAAGAAG

>cca-miR6108b MIMAT0024545

UGAGAAGCGUAAGAAGGGAUC

>cca-miR393 MIMAT0024524

UCCAAAGGAAUCGCAUUGAUCC

>cca-miR168a MIMAT0024517

UCGCUUGGUGCAGGUCGGGAA

>cca-miR396a-5p MIMAT0024529

UUCCACAGCUUUCUUGAACUU

>cca-miR390 MIMAT0024523

AAGCUCAGGAGGGAUAGCGUC

>cca-miR398 MIMAT0024532

UGUGUUCUCAGGUCGCCCCUG

>cca-miR395c MIMAT0024528

CUGAAGUGUUUGGAGGAACUC

>cca-miR6110-5p MIMAT0024547

AUCUUGUAACAUUUGAUGAUGUGG

>cca-miR156b MIMAT0024511

UGACAGAAGAGAGUGAGCAUA

>cca-miR394 MIMAT0024525

UUGGCAUUCUGUCCACCUCC

>cca-miR6105b MIMAT0024542

CACGAAAACAGACUGGUCUCACA

>cca-miR164 MIMAT0024515

UGGAGAAGCAGGGUACGUGCA

>cca-miR6112 MIMAT0024551

GAAGUUUCAAGUGUAAAAAAGUGG

>cca-miR395b MIMAT0024527

CUGAAGUGUUUGGAGGAACUC

>cca-miR6118-5p MIMAT0024565

UGGAAUUGGGUGCUUCGGAAGA

>cca-miR6105a MIMAT0024538

ACGAAAACAUGUUGGUCUCACGUG

>cca-miR395a MIMAT0024526

UUGAAGUGUUUGGGGGGACUC

>cca-miR167 MIMAT0024516

UGAAGCUGCCAGCAUGAUCUGG

>cca-miR6108c MIMAT0024555

AAGCGUAAGAAGAGAUCUGAACC

>cca-miR6106-5p MIMAT0024539

UUGCAAGUAUCCGGAUUUAAA

>cca-miR156c MIMAT0024512

UUGACAGAAGAUAGAGAGCAC

>cca-miR6116-5p MIMAT0024562

CAUGCUUGUGAUCAAAUGAUG

>cca-miR6109 MIMAT0024546

AUGGACGUGUUAUUCAUCAUGAAU

>cca-miR399 MIMAT0024533

UGCCAAAGGAGAUUUGCCCUG

>cca-miR6108e-5p MIMAT0024557

GUAAGAAGAGAUCUCCACCCUUGG

>cca-miR6104 MIMAT0024537

AUACGACAAAUAGAACAAAUAAAC

>cca-miR6114-5p MIMAT0024553

UGAAAGGAAUCAUGAACGUGA

>cca-miR6111-3p MIMAT0024550

CAUGCAUGGUGAUAUAAAUAGC

>cca-miR408 MIMAT0024534

UGCACUGCCUCUUCCCUGGCU

>cca-miR6114-3p MIMAT0024554

UCACGUCCAUUGUCCCUUUCA

>cca-miR6110-3p MIMAT0024548

AUCCACGUCAUCAAAUGUUACAAG

>cca-miR319 MIMAT0024522

UUGGACUGAAGGGAGCUCCCU

>cca-miR156a MIMAT0024510

UGACAGAAGAGAGUGAGCAC

>cca-miR6113 MIMAT0024552

UCUGAAACUCAAGAACACGUUG

>cca-miR6108a MIMAT0024543

UGAGAAGCGUAAGAAGGGAUC

>cca-miR169b MIMAT0024519

UAGCCAAGGAUGACUUGCUAC

>cca-miR396c MIMAT0024561

UUCAAGAAAGCUGUGGGAAAA

>cca-miR6103-3p MIMAT0024536

CAAGAAGUUGUCUUAGGCAUG

>cca-miR160b MIMAT0024514

UGCCUGGCUCCCUGUAUGCCA

>cca-miR6118-3p MIMAT0024566

UUCCGAGGCCACCCAUUCCAAC

>cca-miR6108g MIMAT0024544

UGAGAAGCGUAAGAAGGGAUC

>cca-miR6115 MIMAT0024560

UCUGGACGGUAUGCACAUGUGCAU

>cca-miR160a MIMAT0024513

UGCCUGGCUCCCUGGAUGCCA

>cca-miR6107 MIMAT0024541

AAAGGGGACAAUAUCUGGUACGGU

>cca-miR6117 MIMAT0024564

GGUUAGGUUGAUCGGGUUGAAGAC

>cca-miR6111-5p MIMAT0024549

UCUUUAUGUCACGAUGUAUGAC

>cca-miR396a-3p MIMAT0024530

GUUCAAGAAAGCUGUGGGAAA

>cca-miR6116-3p MIMAT0024563

UCAUUUGAUCACAAGCAUGAG

>cca-miR172 MIMAT0024521

AGAAUCUUGAUGAUGCUGCAU

>cca-miR396b MIMAT0024531

UCCCACAGCUUUAUUGAACUG

>cca-miR6103-5p MIMAT0024535

UGUCUAAGACAACUCCUUGGA

>cca-miR169a MIMAT0024518

UAGCCAAGGAUGACUUGCCUU

>han-miR160a MIMAT0025498

UGCUCGGCUCCCUGUAUGCCA

>han-miR156a MIMAT0025488

UGACAGAAGAGAGUGAGCAC

>han-miR156c MIMAT0025496

UUGACAGAAGAUAGAGAGCAC

>han-miR3440 MIMAT0025524

UGGGUUGGUCAAGGGAAACGC

>han-miR156b MIMAT0025489

UGACAGAAGAGAGUGAGCAC

>han-miR3630-3p MIMAT0025526

UGUGGGAAUCUCUCUGAUGCUU

>han-miR3630-5p MIMAT0025525

GCAAGUGAUGAUAGACCAGACA

>har-miR403a MIMAT0025519

UUAGAUUCACGCACAAACUCG

>har-miR156c MIMAT0025495

UGACAGAAGAGAGGGAGCA

>har-miR156a MIMAT0025493

UGACAGAAGAGAGUGAGCAC

>hci-miR156a MIMAT0025490

UGACAGAAGAGAGUGAGUAC

>hci-miR164a MIMAT0025502

UGGAGAAGCAGGGCACGUGAA

>hci-miR156b MIMAT0025491

UGACAGAAGAGAGGGAGCAC

>hex-miR390a MIMAT0025509

AAGCUCAGGAGGGAUAGCGCC

>hex-miR390b MIMAT0025510

AAGCUCAGGAGGGAUAGCGCC

>hpa-miR171a MIMAT0025505

UGAUUGAGCCGUGCCAAUAUC

>hpa-miR166a MIMAT0025503

UCGGACCAGGCUUCAUUCCCC

>hpa-miR156a MIMAT0025494

UGACAGAAGAGAGAGAGCAC

>hpe-miR162a MIMAT0025501

UCGAUAAACCUCUGCAUCCAG

>hpe-miR403a MIMAT0025520

UUAGAUUCACGCACAAACUCG

>hpe-miR166a MIMAT0025504

UCGGACCAGGCUUCAUUCCCC

>htu-miR403b MIMAT0025515

UUAGAUUCACGCACAAACUCG

>htu-miR162a MIMAT0025500

UCGAUAAACCUCUGCAUCCAG

>htu-miR393b MIMAT0025512

UCCAAAGGGAUCGCAUUGAUCC

>htu-miR403a MIMAT0025514

UUAGAUUCACGCACAAACUCG

>htu-miR171a MIMAT0025506

UGAUUGAGCCGUGCCAAUAU

>htu-miR171c MIMAT0025508

UGAUUGAGCCGUGCCAAUAUC

>htu-miR403d MIMAT0025517

UUAGAUUCACGCACAAACUCG

>htu-miR159a MIMAT0025497

UUUGGAUUGAAGGGAGCUCUA

>htu-miR530 MIMAT0025521

UGCAUUUGCACCUGCACCUU

>htu-miR393a MIMAT0025511

UCCAAAGGGAUCGCAUUGAUCC

>htu-miR171b MIMAT0025507

UGAUUGAGCCGUGCCAAUAUC

>htu-miR403e MIMAT0025518

UUAGAUUCACGCACAAACUCG

>htu-miR393c MIMAT0025513

UCCAAAGGGAUCGCAUUGAUCC

>htu-miR156a MIMAT0025492

UGACAGAAGAGAGUGAGCAC

>htu-miR403c MIMAT0025516

UUAGAUUCACGCACAAACUCG

>htu-miR160a MIMAT0025499

UGCCUGGCUCCCUGUAUGCC

>aly-miR166h-3p MIMAT0031383

UCGGACCAGGCUUCAUUCCCC

>aly-miR4225 MIMAT0017913

GAAUCGAUGGUUAAACAACAC

>aly-miR169n-5p MIMAT0017511

UAGCCAAAGAUGACUUGCCUG

>aly-miR846-5p MIMAT0017635

UUCAGGGACUUCAAUUCAGAA

>aly-miR170-5p MIMAT0017513

UAUUGGCCUGGUUCACUCAGA

>aly-miR172a-3p MIMAT0017522

AGAAUCUUGAUGAUGCUGCAU

>aly-miR2112-3p MIMAT0017662

CUUUAUAUCCGCAUUUGCGCA

>aly-miR402-5p MIMAT0017681

UUGGCCUAUUGAACCUCUGUUU

>aly-miR157b-3p MIMAT0017416

GCUCUCUAAGCUUCUGUCAUCA

>aly-miR156e-5p MIMAT0017405

UGACAGAAGAGAGUGAGCAC

>aly-miR169m-5p MIMAT0017509

UAGCCAAGGAUGACUUGCCUG

>aly-miR4250 MIMAT0017941

UCCAAAGGCACAAGAACAUCA

>aly-miR859-3p MIMAT0017648

UGAUUUUACAAUAGAUAGAUA

>aly-miR3440-5p MIMAT0017711

UGGUUUCCCUGGCCAAUCCACU

>aly-miR166d-5p MIMAT0017465

GGAAUGUUGUCUGGCUCGAGG

>aly-miR825-5p MIMAT0017605

UCAAGCACCAGCUCGAAGAAGC

>aly-miR844-3p MIMAT0017630

CCAUCUUACUAGUCUUUCUUU

>aly-miR823-3p MIMAT0017602

UAGGUUGGUGAUCAUAUAGGAU

>aly-miR4230 MIMAT0017918

UGUCUCGGAAAUUUGCACCCU

>aly-miR841 MIMAT0017907

UACGACCCACUGGAAACUAAA

>aly-miR4242 MIMAT0017931

AAAGUUAACUAUGGCAUUCCC

>aly-miR166f-3p MIMAT0017470

UCGGACCAGGCUUCAUUCCCC

>aly-miR319c-5p MIMAT0017433

GGAGAUUCUUUCAGUCCAGUC

>aly-miR169l-3p MIMAT0017508

GGCAGUCUCCUUGGCUAUC

>ath-miR173-5p MIMAT0000206

UUCGCUUGCAGAGAGAAAUCAC

>ath-miR395a MIMAT0000938

CUGAAGUGUUUGGGGGAACUC

>ath-miR5021 MIMAT0020525

UGAGAAGAAGAAGAAGAAAA

>ath-miR157c-3p MIMAT0031872

GCUCUCUAUACUUCUGUCACC

>ath-miR167a-3p MIMAT0031883

GAUCAUGUUCGCAGUUUCACC

>ath-miR1888b MIMAT0022436

UUAGGCUAAGAUUUGUGAAGA

>ath-miR156a-3p MIMAT0031865

GCUCACUGCUCUUUCUGUCAGA

>ath-miR5638a MIMAT0022398

AUACCAAAACUCUCUCACUUU

>ath-miR779.2 MIMAT0004329

UGAUUGGAAAUUUCGUUGACU

>ath-miR5642a MIMAT0022403

UCUCGCGCUUGUACGGCUUU

>ath-miR5631 MIMAT0022390

UGGCAGGAAAGACAUAAUUUU

>ath-miR2111b-3p MIMAT0011153

AUCCUCGGGAUACAGUUUACC

>ath-miR169g-3p MIMAT0000912

UCCGGCAAGUUGACCUUGGCU

>ath-miR870-5p MIMAT0032025

AAGAACAUCAAAUUAGAAUGU

>ath-miR169b-3p MIMAT0031897

GGCAAGUUGUCCUUCGGCUACA

>ath-miR158a-3p MIMAT0000176

UCCCAAAUGUAGACAAAGCA

>ath-miR5665 MIMAT0022443

UUGGUGGACAAGAUCUGGGAU

>ath-miR5014a-5p MIMAT0021047

ACACUUAGUUUUGUACAACAU

>ath-miR397b MIMAT0000947

UCAUUGAGUGCAUCGUUGAUG

>ath-miR830-5p MIMAT0004247

UCUUCUCCAAAUAGUUUAGGUU

>ath-miR776 MIMAT0003935

UCUAAGUCUUCUAUUGAUGUU

>ath-miR5658 MIMAT0022431

AUGAUGAUGAUGAUGAUGAAA

>ath-miR5020a MIMAT0020524

UGGAAGAAGGUGAGACUUGCA

>ath-miR8168 MIMAT0032766

AGGUGCUGAGUGUGCUAGUGC

>ath-miR5999 MIMAT0023525

UCUUCACUAUUAGACGGACAA

>ath-miR396b-5p MIMAT0000945

UUCCACAGCUUUCUUGAACUU

>ath-miR166e-3p MIMAT0000193

UCGGACCAGGCUUCAUUCCCC

>ath-miR4239 MIMAT0017947

UUUGUUAUUUUCGCAUGCUCC

>ath-miR771 MIMAT0003930

UGAGCCUCUGUGGUAGCCCUCA

>ath-miR5654-3p MIMAT0023005

UGGAAGAUGCUUUGGGAUUUAUU

>ath-miR395d MIMAT0000941

CUGAAGUGUUUGGGGGAACUC

>ath-miR5632-5p MIMAT0032127

UUGAUUCUCUUAUCCAACUGU

>ath-miR5643b MIMAT0022445

AGGCUUUUAAGAUCUGGUUGC

>ath-miR847 MIMAT0004278

UCACUCCUCUUCUUCUUGAUG

>ath-miR171c-5p MIMAT0031900

AGAUAUUGGUGCGGUUCAAUC

>ath-miR161.2 MIMAT0006779

UCAAUGCAUUGAAAGUGACUA

>ath-miR156d-3p MIMAT0031868

GCUCACUCUCUUUUUGUCAUAAC

>ath-miR5630a MIMAT0022389

GCUAAGAGCGGUUCUGAUGGA

>ath-miR399c-5p MIMAT0031913

GGGCAUCUUUCUAUUGGCAGG

>ath-miR156f-5p MIMAT0000171

UGACAGAAGAGAGUGAGCAC

>ath-miR166f MIMAT0000194

UCGGACCAGGCUUCAUUCCCC

>ath-miR5641 MIMAT0022402

UGGAAGAAGAUGAUAGAAUUA

>ath-miR3440b-3p MIMAT0017946

UGGAUUGGUCAAGGGAAGCGU

>ath-miR416 MIMAT0001324

GGUUCGUACGUACACUGUUCA

>ath-miR160c-3p MIMAT0031875

CGUACAAGGAGUCAAGCAUGA

>ath-miR5595a MIMAT0023517

ACAUAUGAUCUGCAUCUUUGC

>ath-miR5024-3p MIMAT0021048

CCGUAUCUUGGCCUUGUCAUU

>ath-miR168b-5p MIMAT0000199

UCGCUUGGUGCAGGUCGGGAA

>ath-miR166b-3p MIMAT0000190

UCGGACCAGGCUUCAUUCCCC

>ath-miR167d MIMAT0000905

UGAAGCUGCCAGCAUGAUCUGG

>ath-miR8172 MIMAT0032771

AUGGAUCAUCUAGAUGGAGAU

>ath-miR860 MIMAT0004304

UCAAUAGAUUGGACUAUGUAU

>ath-miR472-3p MIMAT0003931

UUUUUCCUACUCCGCCCAUACC

>ath-miR2939 MIMAT0014143

UAACGCACAACACUAAGCCAU

>ath-miR834 MIMAT0004254

UGGUAGCAGUAGCGGUGGUAA

>ath-miR168a-5p MIMAT0000198

UCGCUUGGUGCAGGUCGGGAA

>ath-miR863-5p MIMAT0004309

UUAUGUCUUGUUGAUCUCAAU

>ath-miR841a-5p MIMAT0004263

UACGAGCCACUUGAAACUGAA

>ath-miR172e-3p MIMAT0001019

GGAAUCUUGAUGAUGCUGCAU

>ath-miR156i MIMAT0022421

UGACAGAAGAGAGAGAGCAG

>ath-miR824-5p MIMAT0004277

UAGACCAUUUGUGAGAAGGGA

>ath-miR5998a MIMAT0023521

ACAGUUUGUGUUUUGUUUUGU

>ath-miR833b MIMAT0022419

UGUUUGUUGACAUCGGUCUAG

>ath-miR5995b MIMAT0023518

ACAUAUGAUCUGCAUCUUUGC

>ath-miR399a MIMAT0000951

UGCCAAAGGAGAUUUGCCCUG

>ath-miR843 MIMAT0004265

UUUAGGUCGAGCUUCAUUGGA

>ath-miR447c-3p MIMAT0002115

UUGGGGACGACAUCUUUUGUUG

>ath-miR5998b MIMAT0023522

ACAGUUUGUGUUUUGUUUUGU

>ath-miR5648-5p MIMAT0022412

UUUGGAAAUAUUUGGCUUGACU

>ath-miR827 MIMAT0004243

UUAGAUGACCAUCAACAAACU

>ath-miR167b MIMAT0000197

UGAAGCUGCCAGCAUGAUCUA

>ath-miR5017-3p MIMAT0020521

UUAUACCAAAUUAAUAGCAAA

>ath-miR405d MIMAT0001008

AUGAGUUGGGUCUAACCCAUAACU

>ath-miR163 MIMAT0000184

UUGAAGAGGACUUGGAACUUCGAU

>ath-miR823 MIMAT0004240

UGGGUGGUGAUCAUAUAAGAU

>ath-miR447a.2-3p MIMAT0020345

UAUGGAAGAAAUUGUAGUAUU

>ath-miR5660 MIMAT0022434

CAGGUGGUUAGUGCAAUGGAA

>ath-miR156c-5p MIMAT0000168

UGACAGAAGAGAGUGAGCAC

>ath-miR167a-5p MIMAT0000196

UGAAGCUGCCAGCAUGAUCUA

>ath-miR390a-3p MIMAT0031902

CGCUAUCCAUCCUGAGUUUCA

>ath-miR840-3p MIMAT0032021

UUGUUUAGGUCCCUUAGUUUC

>ath-miR8183 MIMAT0032782

UUUAGUUGACGGAAUUGUGGC

>ath-miR169a-5p MIMAT0000200

CAGCCAAGGAUGACUUGCCGA

>ath-miR854d MIMAT0004283

GAUGAGGAUAGGGAGGAGGAG

>ath-miR393a-5p MIMAT0000934

UCCAAAGGGAUCGCAUUGAUCC

>ath-miR5654-5p MIMAT0022426

AUAAAUCCCAACAUCUUCCA

>ath-miR840-5p MIMAT0004262

ACACUGAAGGACCUAAACUAAC

>ath-miR398c-5p MIMAT0031912

AGGGUUGAUAUGAGAACACAC

>ath-miR3932b-5p MIMAT0032113

UUUGACGUGCUCGAUCUGCUC

>ath-miR402 MIMAT0001003

UUCGAGGCCUAUUAAACCUCUG

>ath-miR5629 MIMAT0022388

UUAGGGUAGUUAACGGAAGUUA

>ath-miR319b MIMAT0000512

UUGGACUGAAGGGAGCUCCCU

>ath-miR5012 MIMAT0020516

UUUUACUGCUACUUGUGUUCC

>ath-miR832-3p MIMAT0004251

UUGAUUCCCAAUCCAAGCAAG

>ath-miR319a MIMAT0000511

UUGGACUGAAGGGAGCUCCCU

>ath-miR156e MIMAT0000170

UGACAGAAGAGAGUGAGCAC

>ath-miR5029 MIMAT0020535

AAUGAGAGAGAACACUGCAAA

>ath-miR166a-3p MIMAT0000189

UCGGACCAGGCUUCAUUCCCC

>ath-miR8167d MIMAT0037259

AGAUGUGGAGAUCGUGGGGAUG

>ath-miR399b MIMAT0000952

UGCCAAAGGAGAGUUGCCCUG

>ath-miR8176 MIMAT0032775

GGCCGGUGGUCGCGAGAGGGA

>ath-miR158a-5p MIMAT0031873

CUUUGUCUACAAUUUUGGAAA

>ath-miR162b-3p MIMAT0000183

UCGAUAAACCUCUGCAUCCAG

>ath-miR169h MIMAT0000913

UAGCCAAGGAUGACUUGCCUG

>ath-miR171b-5p MIMAT0031899

AGAUAUUAGUGCGGUUCAAUC

>ath-miR781b MIMAT0022420

UUAGAGUUUUCUGGAUACUUA

>ath-miR8121 MIMAT0032262

AAAGUAUAAUGGUUUAGUGGUUUG

>ath-miR830-3p MIMAT0004248

UAACUAUUUUGAGAAGAAGUG

>ath-miR171a-3p MIMAT0000202

UGAUUGAGCCGCGCCAAUAUC

>ath-miR2111a-5p MIMAT0011150

UAAUCUGCAUCCUGAGGUUUA

>ath-miR8181 MIMAT0032780

UGGGGGUGGGGGGGUGACAG

>ath-miR157a-3p MIMAT0031870

GCUCUCUAGCCUUCUGUCAUC

>ath-miR167c-3p MIMAT0031917

UAGGUCAUGCUGGUAGUUUCACC

>ath-miR171b-3p MIMAT0000920

UUGAGCCGUGCCAAUAUCACG

>ath-miR396b-3p MIMAT0031909

GCUCAAGAAAGCUGUGGGAAA

>ath-miR3933 MIMAT0018348

AGAAGCAAAAUGACGACUCGG

>ath-miR8166 MIMAT0032762

AGAGAGUGUAGAAAGUUUCUCA

>ath-miR859 MIMAT0004303

UCUCUCUGUUGUGAAGUCAAA

>ath-miR842 MIMAT0004264

UCAUGGUCAGAUCCGUCAUCC

>ath-miR4240 MIMAT0017948

UGACUAGACCCGUAACAUUAC

>ath-miR394b-5p MIMAT0000937

UUGGCAUUCUGUCCACCUCC

>ath-miR5664 MIMAT0022442

AUAGUCAAUUUUAUCGGUCUG

>ath-miR851-5p MIMAT0004273

UCUCGGUUCGCGAUCCACAAG

>ath-miR829-3p.1 MIMAT0004245

AGCUCUGAUACCAAAUGAUGGAAU

>ath-miR172b-5p MIMAT0000204

GCAGCACCAUUAAGAUUCAC

>ath-miR5643a MIMAT0022404

AGGCUUUUAAGAUCUGGUUGC

>ath-miR8178 MIMAT0032777

UAACAGAGUAAUUGUACAGUG

>ath-miR2936 MIMAT0014140

CUUGAGAGAGAGAACACAGACG

>ath-miR5635d MIMAT0022405

UGUUAAGGAGUGUUAACGGUG

>ath-miR405a MIMAT0001006

AUGAGUUGGGUCUAACCCAUAACU

>ath-miR169k MIMAT0000916

UAGCCAAGGAUGACUUGCCUG

>ath-miR2938 MIMAT0014142

GAUCUUUUGAGAGGGUUCCAG

>ath-miR5630b MIMAT0022399

GCUAAGAGCGGUUCUGAUGGA

>ath-miR779.1 MIMAT0003938

UUCUGCUAUGUUGCUGCUCAU

>ath-miR406 MIMAT0001009

UAGAAUGCUAUUGUAAUCCAG

>ath-miR170-5p MIMAT0031887

UAUUGGCCUGGUUCACUCAGA

>ath-miR8169 MIMAT0032767

AUAGACAGAGUCACUCACAGA

>ath-miR5016 MIMAT0020520

UUCUUGUGGAUUCCUUGGAAA

>ath-miR162a-3p MIMAT0000182

UCGAUAAACCUCUGCAUCCAG

>ath-miR775 MIMAT0003934

UUCGAUGUCUAGCAGUGCCA

>ath-miR865-3p MIMAT0004314

UUUUUCCUCAAAUUUAUCCAA

>ath-miR5025 MIMAT0020531

ACUGUAUAUAUGUAAGUGACA

>ath-miR169b-5p MIMAT0000906

CAGCCAAGGAUGACUUGCCGG

>ath-miR5635a MIMAT0022395

UGUUAAGGAGUGUUAACGGUG

>ath-miR852 MIMAT0004275

AAGAUAAGCGCCUUAGUUCUG

>ath-miR837-3p MIMAT0004259

AAACGAACAAAAAACUGAUGG

>ath-miR169a-3p MIMAT0031886

GGCAAGUUGUCCUUGGCUAC

>ath-miR864-5p MIMAT0004311

UCAGGUAUGAUUGACUUCAAA

>ath-miR419 MIMAT0001327

UUAUGAAUGCUGAGGAUGUUG

>ath-miR773b-5p MIMAT0017735

GGCAAUAACUUGAGCAAACA

>ath-miR166g MIMAT0000195

UCGGACCAGGCUUCAUUCCCC

>ath-miR8180 MIMAT0032779

UGCGGUGCGGGAGAAGUGC

>ath-miR156a-5p MIMAT0000166

UGACAGAAGAGAGUGAGCAC

>ath-miR8167c MIMAT0032765

AGAUGUGGAGAUCGUGGGGAUG

>ath-miR5648-3p MIMAT0022413

AUCUGAAGAAAAUAGCGGCAU

>ath-miR156b-5p MIMAT0000167

UGACAGAAGAGAGUGAGCAC

>ath-miR2934-3p MIMAT0020364

CAUCCAAGGUGUUUGUAGAAA

>ath-miR166c MIMAT0000191

UCGGACCAGGCUUCAUUCCCC

>ath-miR394a MIMAT0000936

UUGGCAUUCUGUCCACCUCC

>ath-miR774a MIMAT0003933

UUGGUUACCCAUAUGGCCAUC

>ath-miR167c-5p MIMAT0001018

UAAGCUGCCAGCAUGAUCUUG

>ath-miR156j MIMAT0022423

UGACAGAAGAGAGAGAGCAC

>ath-miR395e MIMAT0000942

CUGAAGUGUUUGGGGGAACUC

>ath-miR5642b MIMAT0022435

UCUCGCGCUUGUACGGCUUU

>ath-miR168b-3p MIMAT0031885

CCCGUCUUGUAUCAACUGAAU

>ath-miR824-3p MIMAT0032024

CCUUCUCAUCGAUGGUCUAGA

>ath-miR396a-3p MIMAT0031908

GUUCAAUAAAGCUGUGGGAAG

>ath-miR165a-5p MIMAT0031879

GGAAUGUUGUCUGGAUCGAGG

>ath-miR836 MIMAT0004257

UCCUGUGUUUCCUUUGAUGCGUGG

>ath-miR8184 MIMAT0032783

UUUGGUCUGAUUACGAAUGUA

>ath-miR5645c MIMAT0022414

AACCUAUUUAACGACAUGACU

>ath-miR1886.1 MIMAT0007853

UGAGAGAAGUGAGAUGAAAUC

>ath-miR5019 MIMAT0020523

UGUUGGGAAAGAAAAACUCUU

>ath-miR854e MIMAT0018504

GAUGAGGAUAGGGAGGAGGAG

>ath-miR774b-3p MIMAT0017740

CAUCCAUAUUUUCAUCUCGAA

>ath-miR169l MIMAT0000917

UAGCCAAGGAUGACUUGCCUG

>ath-miR426 MIMAT0001337

UUUUGGAAAUUUGUCCUUACG

>ath-miR5015 MIMAT0020519

UUGGUGUUAUGUGUAGUCUUC

>ath-miR398a-3p MIMAT0000948

UGUGUUCUCAGGUCACCCCUU

>ath-miR8165 MIMAT0032761

AAUGGAGGCAAGUGUGAAGGA

>ath-miR826b MIMAT0035543

UGGUUUUGGACACGUGAAAAU

>ath-miR394b-3p MIMAT0031907

AGGUGGGCAUACUGCCAAUAG

>ath-miR8171 MIMAT0032770

AUAGGUGGGCCAGUGGUAGGA

>ath-miR5022 MIMAT0020527

GUCAUGGGGUAUGAUCGAAUG

>ath-miR161.1 MIMAT0000181

UGAAAGUGACUACAUCGGGGU

>ath-miR172a MIMAT0000203

AGAAUCUUGAUGAUGCUGCAU

>ath-miR4221 MIMAT0017942

UUUUCCUCUGUUGAAUUCUUGC

>ath-miR845b MIMAT0004317

UCGCUCUGAUACCAAAUUGAUG

>ath-miR165b MIMAT0000188

UCGGACCAGGCUUCAUCCCCC

>ath-miR168a-3p MIMAT0031884

CCCGCCUUGCAUCAACUGAAU

>ath-miR164c-3p MIMAT0031916

CACGUGUUCUACUACUCCAAC

>ath-miR415 MIMAT0001323

AACAGAGCAGAAACAGAACAU

>ath-miR393b-3p MIMAT0031906

AUCAUGCGAUCUCUUUGGAUU

>ath-miR5650 MIMAT0022416

UUGUUUUGGAUCUUAGAUACA

>ath-miR159b-5p MIMAT0031889

GAGCUCCUUGAAGUUCAAUGG

>ath-miR1887 MIMAT0007854

UACUAAGUAGAGUCUAAGAGA

>ath-miR5636 MIMAT0022396

CGUAGUUGCAGAGCUUGACGG

>ath-miR870-3p MIMAT0004322

UAAUUUGGUGUUUCUUCGAUC

>ath-miR8175 MIMAT0032774

GAUCCCCGGCAACGGCGCCA

>ath-miR864-3p MIMAT0004312

UAAAGUCAAUAAUACCUUGAAG

>ath-miR2112-3p MIMAT0011155

CUUUAUAUCCGCAUUUGCGCA

>ath-miR841a-3p MIMAT0032022

AUUUCUAGUGGGUCGUAUUCA

>ath-miR5639-5p MIMAT0022400

UAGUCCACUGUGGUCUAAGGC

>ath-miR159c MIMAT0001015

UUUGGAUUGAAGGGAGCUCCU

>ath-miR8179 MIMAT0032778

UGACUGCAUUAACUUGAUCGU

>ath-miR782 MIMAT0003941

ACAAACACCUUGGAUGUUCUU

>ath-miR8173 MIMAT0032772

AUGUGCUGAUUCGAGGUGGGA

>ath-miR156b-3p MIMAT0031866

UGCUCACCUCUCUUUCUGUCAGU

>ath-miR407 MIMAT0001010

UUUAAAUCAUAUACUUUUGGU

>ath-miR826a MIMAT0004242

UAGUCCGGUUUUGGAUACGUG

>ath-miR391-3p MIMAT0031904

ACGGUAUCUCUCCUACGUAGC

>ath-miR418 MIMAT0001326

UAAUGUGAUGAUGAACUGACC

>ath-miR169c MIMAT0000907

CAGCCAAGGAUGACUUGCCGG

>ath-miR863-3p MIMAT0004310

UUGAGAGCAACAAGACAUAAU

>ath-miR3440b-5p MIMAT0017945

UUUUCUUGGCCCAUCCACUUC

>ath-miR869.1 MIMAT0004320

AUUGGUUCAAUUCUGGUGUUG

>ath-miR5644 MIMAT0022406

GUGGGUUGCGGAUAACGGUA

>ath-miR5020b MIMAT0020529

AUGGCAUGAAAGAAGGUGAGA

>ath-miR832-5p MIMAT0004250

UGCUGGGAUCGGGAAUCGAAA

>ath-miR164b-3p MIMAT0031878

CAUGUGCCCAUCUUCACCAUC

>ath-miR156f-3p MIMAT0031869

GCUCACUCUCUAUCCGUCACC

>ath-miR838 MIMAT0004260

UUUUCUUCUACUUCUUGCACA

>ath-miR854b MIMAT0004281

GAUGAGGAUAGGGAGGAGGAG

>ath-miR781a MIMAT0003940

UUAGAGUUUUCUGGAUACUUA

>ath-miR835-3p MIMAT0004256

UGGAGAAGAUACGCAAGAAAG

>ath-miR171c-3p MIMAT0000921

UUGAGCCGUGCCAAUAUCACG

>ath-miR156g MIMAT0001012

CGACAGAAGAGAGUGAGCAC

>ath-miR162b-5p MIMAT0031877

UGGAGGCAGCGGUUCAUCGAUC

>ath-miR171a-5p MIMAT0031888

UAUUGGCCUGGUUCACUCAGA

>ath-miR414 MIMAT0001322

UCAUCUUCAUCAUCAUCGUCA

>ath-miR8182 MIMAT0032781

UUGUGUUGCGUUUCUGUUGAUU

>ath-miR8167a MIMAT0032763

AGAUGUGGAGAUCGUGGGGAUG

>ath-miR398b-3p MIMAT0000949

UGUGUUCUCAGGUCACCCCUG

>ath-miR405b MIMAT0001007

AUGAGUUGGGUCUAACCCAUAACU

>ath-miR5646 MIMAT0022410

GUUCGAGGCACGUUGGGAGG

>ath-miR865-5p MIMAT0004313

AUGAAUUUGGAUCUAAUUGAG

>ath-miR160c-5p MIMAT0000180

UGCCUGGCUCCCUGUAUGCCA

>ath-miR398a-5p MIMAT0031910

AAGGAGUGGCAUGUGAACACA

>ath-miR5027 MIMAT0020533

ACCGGUUGGAACUUGCCUUAA

>ath-miR2937 MIMAT0014141

AUAAGAGCUGUUGAAGGAGUC

>ath-miR778 MIMAT0003937

UGGCUUGGUUUAUGUACACCG

>ath-miR2111a-3p MIMAT0011151

GUCCUCGGGAUGCGGAUUACC

>ath-miR157b-5p MIMAT0000173

UUGACAGAAGAUAGAGAGCAC

>ath-miR166d MIMAT0000192

UCGGACCAGGCUUCAUUCCCC

>ath-miR170-3p MIMAT0000201

UGAUUGAGCCGUGUCAAUAUC

>ath-miR5651 MIMAT0022422

UUGUGCGGUUCAAAUAGUAAC

>ath-miR408-5p MIMAT0031915

ACAGGGAACAAGCAGAGCAUG

>ath-miR166e-5p MIMAT0031882

GGAAUGUUGUCUGGCACGAGG

>ath-miR395f MIMAT0000943

CUGAAGUGUUUGGGGGGACUC

>ath-miR169f-5p MIMAT0000910

UGAGCCAAGGAUGACUUGCCG

>ath-miR773a MIMAT0003932

UUUGCUUCCAGCUUUUGUCUC

>ath-miR158b MIMAT0001014

CCCCAAAUGUAGACAAAGCA

>ath-miR5635b MIMAT0022418

UGUUAAGGAGUGUUAACGGUG

>ath-miR866-5p MIMAT0004315

UCAAGGAACGGAUUUUGUUAA

>ath-miR5637 MIMAT0022397

AAUGCGCAACUCUAUAUUUCC

>ath-miR1886.2 MIMAT0012129

UGAGAUGAAAUCUUUGAUUGG

>ath-miR829-3p.2 MIMAT0004246

CAAAUUAAAGCUUCAAGGUAG

>ath-miR853 MIMAT0004276

UCCCCUCUUUAGCUUGGAGAAG

>ath-miR5633 MIMAT0022393

UAUGAUCAUCAGAAAACAGUG

>ath-miR164b-5p MIMAT0000186

UGGAGAAGCAGGGCACGUGCA

>ath-miR862-3p MIMAT0004308

AUAUGCUGGAUCUACUUGAAG

>ath-miR400 MIMAT0001001

UAUGAGAGUAUUAUAAGUCAC

>ath-miR866-3p MIMAT0004316

ACAAAAUCCGUCUUUGAAGA

>ath-miR5652 MIMAT0022424

UUGAAUGUGAAUGAAUCGGGC

>ath-miR169g-5p MIMAT0000911

UGAGCCAAGGAUGACUUGCCG

>ath-miR172c MIMAT0000922

AGAAUCUUGAUGAUGCUGCAG

>ath-miR165a-3p MIMAT0000187

UCGGACCAGGCUUCAUCCCCC

>ath-miR399c-3p MIMAT0000953

UGCCAAAGGAGAGUUGCCCUG

>ath-miR399f MIMAT0000956

UGCCAAAGGAGAUUUGCCCGG

>ath-miR160a-5p MIMAT0000178

UGCCUGGCUCCCUGUAUGCCA

>ath-miR10515 MIMAT0041984

ACCCCGAUGGUUAUCCUCACC

>ath-miR5645b MIMAT0022409

AUUUGAGUCAUGUCGUUAAG

>ath-miR397a MIMAT0000946

UCAUUGAGUGCAGCGUUGAUG

>ath-miR5026 MIMAT0020532

ACUCAUAAGAUCGUGACACGU

>ath-miR159a MIMAT0000177

UUUGGAUUGAAGGGAGCUCUA

>ath-miR169d MIMAT0000908

UGAGCCAAGGAUGACUUGCCG

>ath-miR408-3p MIMAT0001011

AUGCACUGCCUCUUCCCUGGC

>ath-miR5663-3p MIMAT0032129

UGAGAAUGCAAAUCCUUAGCU

>ath-miR5645f MIMAT0022441

AUUUGAGUCAUGUCGUUAAG

>ath-miR390b-5p MIMAT0000932

AAGCUCAGGAGGGAUAGCGCC

>ath-miR862-5p MIMAT0004307

UCCAAUAGGUCGAGCAUGUGC

>ath-miR169f-3p MIMAT0031898

GCAAGUUGACCUUGGCUCUGC

>ath-miR395c MIMAT0000940

CUGAAGUGUUUGGGGGGACUC

>ath-miR4245 MIMAT0023523

ACAAAGUUUUAUACUGACAAU

>ath-miR173-3p MIMAT0022843

UGAUUCUCUGUGUAAGCGAAA

>ath-miR1888a MIMAT0007855

UAAGUUAAGAUUUGUGAAGAA

>ath-miR854a MIMAT0004280

GAUGAGGAUAGGGAGGAGGAG

>ath-miR4243 MIMAT0017949

UUGAAAUUGUAGAUUUCGUAC

>ath-miR169m MIMAT0000918

UAGCCAAGGAUGACUUGCCUG

>ath-miR5023 MIMAT0020528

AUUGGUAGUGGAUAAGGGGGC

>ath-miR777 MIMAT0003936

UACGCAUUGAGUUUCGUUGCUU

>ath-miR855 MIMAT0004279

AGCAAAAGCUAAGGAAAAGGAA

>ath-miR5655 MIMAT0022427

AAGUAGACACAUAAGAAGGAG

>ath-miR399d MIMAT0000954

UGCCAAAGGAGAUUUGCCCCG

>ath-miR5018 MIMAT0020522

UUAAAGCUCCACCAUGAGUCCAAU

>ath-miR390a-5p MIMAT0000931

AAGCUCAGGAGGGAUAGCGCC

>ath-miR780.1 MIMAT0004218

UCUAGCAGCUGUUGAGCAGGU

>ath-miR3434-3p MIMAT0017738

UCAGAGUAUCAGCCAUGUGA

>ath-miR868-5p MIMAT0020355

UCAUGUCGUAAUAGUAGUCAC

>ath-miR157a-5p MIMAT0000172

UUGACAGAAGAUAGAGAGCAC

>ath-miR5656 MIMAT0022429

ACUGAAGUAGAGAUUGGGUUU

>ath-miR403-3p MIMAT0001004

UUAGAUUCACGCACAAACUCG

>ath-miR846-3p MIMAT0004269

UUGAAUUGAAGUGCUUGAAUU

>ath-miR2934-5p MIMAT0014138

UCUUUCUGCAAACGCCUUGGA

>ath-miR157d MIMAT0000175

UGACAGAAGAUAGAGAGCAC

>ath-miR166a-5p MIMAT0031880

GGACUGUUGUCUGGCUCGAGG

>ath-miR5666 MIMAT0022444

AUGGGACAUCGAGCAUUUAAU

>ath-miR413 MIMAT0001321

AUAGUUUCUCUUGUUCUGCAC

>ath-miR447b MIMAT0002114

UUGGGGACGAGAUGUUUUGUUG

>ath-miR2933b MIMAT0014137

GAAAUCGGAGAGGAAAUUCGCC

>ath-miR5014b MIMAT0023524

AUUUGUACACCUAGAUCUGUA

>ath-miR157b-3p MIMAT0031871

GCUCUCUAGCCUUCUGUCAUC

>ath-miR5997 MIMAT0023520

UGAAACCAAGUAGCUAAAUAG

>ath-miR157c-5p MIMAT0000174

UUGACAGAAGAUAGAGAGCAC

>ath-miR159b-3p MIMAT0000207

UUUGGAUUGAAGGGAGCUCUU

>ath-miR8174 MIMAT0032773

AUGUGUAUAGGGAAGCUAAUC

>ath-miR172d-3p MIMAT0000923

AGAAUCUUGAUGAUGCUGCAG

>ath-miR869.2 MIMAT0004321

UCUGGUGUUGAGAUAGUUGAC

>ath-miR868-3p MIMAT0004319

CUUCUUAAGUGCUGAUAAUGC

>ath-miR3932a MIMAT0018346

AACUUUGUGAUGACAACGAAG

>ath-miR858b MIMAT0022417

UUCGUUGUCUGUUCGACCUUG

>ath-miR5647 MIMAT0022411

UCAAGUUUGAUGACGAUUCCA

>ath-miR3434-5p MIMAT0017737

ACUUGGCUGAUUCUAUUAUU

>ath-miR5662 MIMAT0022439

AGAGGUGACCAUUGGAGAUG

>ath-miR841b-3p MIMAT0017742

CAAUUUCUAGUGGGUCGUAUU

>ath-miR5638b MIMAT0022407

ACAGUGGUCAUCUGGUGGGCU

>ath-miR156h MIMAT0001013

UGACAGAAGAAAGAGAGCAC

>ath-miR172e-5p MIMAT0031918

GCAGCACCAUUAAGAUUCAC

>ath-miR5017-5p MIMAT0032120

AUUUGUUACUAAUUUGGAAUG

>ath-miR829-5p MIMAT0032019

ACUUUGAAGCUUUGAUUUGAA

>ath-miR861-3p MIMAT0004306

GAUGGAUAUGUCUUCAAGGAC

>ath-miR5634 MIMAT0022394

AGGGACUUUGUGAAUUUAGGG

>ath-miR845a MIMAT0004268

CGGCUCUGAUACCAAUUGAUG

>ath-miR172d-5p MIMAT0031901

GCAACAUCUUCAAGAUUCAGA

>ath-miR4228-5p MIMAT0017944

AUAGCCUUGAACGCCGUCGUU

>ath-miR5645d MIMAT0022433

AUUUGAGUCAUGUCGUUAAG

>ath-miR5653 MIMAT0022425

UGGGUUGAGUUGAGUUGAGUUGGC

>ath-miR841b-5p MIMAT0017741

UACGAGCCACUGGAAACUGAA

>ath-miR391-5p MIMAT0000933

UUCGCAGGAGAGAUAGCGCCA

>ath-miR850 MIMAT0004272

UAAGAUCCGGACUACAACAAAG

>ath-miR396a-5p MIMAT0000944

UUCCACAGCUUUCUUGAACUG

>ath-miR403-5p MIMAT0031914

UGUUUUGUGCUUGAAUCUAAUU

>ath-miR169n MIMAT0000919

UAGCCAAGGAUGACUUGCCUG

>ath-miR401 MIMAT0001002

CGAAACUGGUGUCGACCGACA

>ath-miR861-5p MIMAT0004305

CCUUGGAGAAAUAUGCGUCAA

>ath-miR5663-5p MIMAT0022440

AGCUAAGGAUUUGCAUUCUCA

>ath-miR5649a MIMAT0022415

AUUGAAUAUGUUGGUUACUAU

>ath-miR8170-5p MIMAT0032768

AUAGCAAAUCGAUAAGCAAUG

>ath-miR447a-3p MIMAT0002113

UUGGGGACGAGAUGUUUUGUUG

>ath-miR162a-5p MIMAT0031876

UGGAGGCAGCGGUUCAUCGAUC

>ath-miR169i MIMAT0000914

UAGCCAAGGAUGACUUGCCUG

>ath-miR5657 MIMAT0022430

UGGACAAGGUUAGAUUUGGUG

>ath-miR831-3p MIMAT0004249

UGAUCUCUUCGUACUCUUCUUG

>ath-miR5628 MIMAT0022387

GAAAUAGCGAAGAUAUGAUUA

>ath-miR393a-3p MIMAT0031905

AUCAUGCUAUCUCUUUGGAUU

>ath-miR398b-5p MIMAT0031911

AGGGUUGAUAUGAGAACACAC

>ath-miR858a MIMAT0004302

UUUCGUUGUCUGUUCGACCUU

>ath-miR5659 MIMAT0022432

CGAUGAAGGUCUUUGGAACGGUA

>ath-miR2112-5p MIMAT0011154

CGCAAAUGCGGAUAUCAAUGU

>ath-miR780.2 MIMAT0003939

UUCUUCGUGAAUAUCUGGCAU

>ath-miR833a-3p MIMAT0004253

UAGACCGAUGUCAACAAACAAG

>ath-miR393b-5p MIMAT0000935

UCCAAAGGGAUCGCAUUGAUCC

>ath-miR417 MIMAT0001325

GAAGGUAGUGAAUUUGUUCGA

>ath-miR857 MIMAT0004301

UUUUGUAUGUUGAAGGUGUAU

>ath-miR5020c MIMAT0022391

UGGCAUGGAAGAAGGUGAGAC

>ath-miR395b MIMAT0000939

CUGAAGUGUUUGGGGGGACUC

>ath-miR420 MIMAT0001328

UAAACUAAUCACGGAAAUGCA

>ath-miR773b-3p MIMAT0017736

UUUGAUUCCAGCUUUUGUCUC

>ath-miR1886.3 MIMAT0013773

AAUUAAAGAUUUCAUCUUACU

>ath-miR8167f MIMAT0037261

AGAUGUGGAGAUCGUGGGGAUG

>ath-miR2111b-5p MIMAT0011152

UAAUCUGCAUCCUGAGGUUUA

>ath-miR166b-5p MIMAT0031881

GGACUGUUGUCUGGCUCGAGG

>ath-miR831-5p MIMAT0032020

AGAAGCGUACAAGGAGAUGAGG

>ath-miR844-5p MIMAT0004266

UGGUAAGAUUGCUUAUAAGCU

>ath-miR854c MIMAT0004282

GAUGAGGAUAGGGAGGAGGAG

>ath-miR169e MIMAT0000909

UGAGCCAAGGAUGACUUGCCG

>ath-miR5014a-3p MIMAT0020518

UUGUACAAAUUUAAGUGUACG

>ath-miR856 MIMAT0004300

UAAUCCUACCAAUAACUUCAGC

>ath-miR5996 MIMAT0023519

UGACAUCCAGAUAGAAGCUUUG

>ath-miR848 MIMAT0004270

UGACAUGGGACUGCCUAAGCUA

>ath-miR8177 MIMAT0032776

GUGUGAUGAUGUGUCAUUUAUA

>ath-miR8167b MIMAT0032764

AGAUGUGGAGAUCGUGGGGAUG

>ath-miR5640 MIMAT0022401

UGAGAGAAGGAAUUAGAUUCA

>ath-miR846-5p MIMAT0032023

CAUUCAAGGACUUCUAUUCAG

>ath-miR5639-3p MIMAT0032128

UUUAGCCUCAGACCACGGUGGACU

>ath-miR825 MIMAT0004241

UUCUCAAGAAGGUGCAUGAAC

>ath-miR828 MIMAT0004244

UCUUGCUUAAAUGAGUAUUCCA

>ath-miR822-3p MIMAT0032018

UGUGCAAAUGCUUUCUACAGG

>ath-miR839-5p MIMAT0004261

UACCAACCUUUCAUCGUUCCC

>ath-miR5024-5p MIMAT0020530

AUGACAAGGCCAAGAUAUAACA

>ath-miR399e MIMAT0000955

UGCCAAAGGAGAUUUGCCUCG

>ath-miR447c-5p MIMAT0020346

CCCCUUACAAUGUCGAGUAAA

>ath-miR4227 MIMAT0017943

UCACUGGUACCAAUCAUUCCA

>ath-miR160b MIMAT0000179

UGCCUGGCUCCCUGUAUGCCA

>ath-miR833a-5p MIMAT0004252

UGUUUGUUGUACUCGGUCUAGU

>ath-miR398c-3p MIMAT0000950

UGUGUUCUCAGGUCACCCCUG

>ath-miR2933a MIMAT0014136

GAAAUCGGAGAGGAAAUUCGCC

>ath-miR8170-3p MIMAT0032769

UUGCUUAAAGAUUUUCUAUGU

>ath-miR160a-3p MIMAT0031874

GCGUAUGAGGAGCCAUGCAUA

>ath-miR172b-3p MIMAT0000205

AGAAUCUUGAUGAUGCUGCAU

>ath-miR822-5p MIMAT0004239

UGCGGGAAGCAUUUGCACAUG

>ath-miR390b-3p MIMAT0031903

CGCUAUCCAUCCUGAGUUCC

>ath-miR164a MIMAT0000185

UGGAGAAGCAGGGCACGUGCA

>ath-miR851-3p MIMAT0004274

UGGGUGGCAAACAAAGACGAC

>ath-miR837-5p MIMAT0004258

AUCAGUUUCUUGUUCGUUUCA

>ath-miR844-3p MIMAT0004267

UUAUAAGCCAUCUUACUAGUU

>ath-miR5028 MIMAT0020534

AAUUGGGUUUAUGCUAGAGUU

>ath-miR156d-5p MIMAT0000169

UGACAGAAGAGAGUGAGCAC

>ath-miR169j MIMAT0000915

UAGCCAAGGAUGACUUGCCUG

>ath-miR319c MIMAT0001016

UUGGACUGAAGGGAGCUCCUU

>ath-miR835-5p MIMAT0004255

UUCUUGCAUAUGUUCUUUAUC

>ath-miR164c-5p MIMAT0001017

UGGAGAAGCAGGGCACGUGCG

>ath-miR5645e MIMAT0022446

AUUUGAGUCAUGUCGUUAAG

>ath-miR156c-3p MIMAT0031867

GCUCACUGCUCUAUCUGUCAGA

>ath-miR5649b MIMAT0022437

AUUGAAUAUGUUGGUUACUAU

>ath-miR5632-3p MIMAT0022392

UUGGAUUUAUAGUUGGAUAAG

>ath-miR4228-3p MIMAT0032109

UCGGAUGCGAAACGGUGGUGU

>ath-miR5013 MIMAT0020517

UUUGUGACAUCUAGGUGCUUU

>ath-miR774b-5p MIMAT0017739

UGAGAUGAAGAUAUGGGUGAU

>ath-miR404 MIMAT0001005

AUUAACGCUGGCGGUUGCGGCAGC

>ath-miR5661 MIMAT0022438

AGAGGUACAUCAUGUAGUCUG

>ath-miR867 MIMAT0004318

UUGAACAUGGUUUAUUAGGAA

>ath-miR472-5p MIMAT0032014

AUGGUCGAAGUAGGCAAAAUC

>ath-miR849 MIMAT0004271

UAACUAAACAUUGGUGUAGUA

>ath-miR5635c MIMAT0022428

UGUUAAGGAGUGUUAACGGUG

>ath-miR8167e MIMAT0037260

AGAUGUGGAGAUCGUGGGGAUG

>ath-miR3932b-3p MIMAT0018347

AACUUUGUGAUGACAACGAAG

>ath-miR5645a MIMAT0022408

AUUUGAGUCAUGUCGUUAAG

>bna-miR160b MIMAT0023618

UGCCUGGCUCCCUGUAUGCCA

>bna-miR390a MIMAT0005602

AAGCUCAGGAGGGAUAGCGCC

>bna-miR393 MIMAT0004447

UCCAAAGGGAUCGCAUUGAUC

>bna-miR169l MIMAT0005623

UAGCCAAGGAUGACUUGCCUGC

>bna-miR397b MIMAT0005601

UCAUUGAGUGCAGCGUUGAUGU

>bna-miR167b MIMAT0005627

UGAAGCUGCCAGCAUGAUCUAA

>bna-miR6030 MIMAT0023650

UCCACCCAUACCAUACAGACCC

>bna-miR166d MIMAT0005632

UCGGACCAGGCUUCAUUCCCC

>bna-miR824 MIMAT0005599

UAGACCAUUUGUGAGAAGGGA

>bna-miR156g MIMAT0023616

UUGACAGAAGAUAGAGAGCAC

>bna-miR390b MIMAT0005603

AAGCUCAGGAGGGAUAGCGCC

>bna-miR168b MIMAT0023628

UCGCUUGGUGCAGGUCGAGAA

>bna-miR164a MIMAT0005633

UGGAGAAGCAGGGCACGUGCA

>bna-miR172a MIMAT0023633

AGAAUCUUGAUGAUGCUGCAU

>bna-miR166e MIMAT0023626

UCGGACCAGGCUUCAUUCCCC

>bna-miR164b MIMAT0023622

UGGAGAAGCAGGGCACGUGCG

>bna-miR1140 MIMAT0005637

ACAGCCUAAACCAAUCGGAGC

>bna-miR169k MIMAT0005622

UAGCCAAGGAUGACUUGCCUGC

>bna-miR161 MIMAT0005634

UCAAUGCACUGAAAGUGACUA

>bna-miR171a MIMAT0005605

UUGAGCCGUGCCAAUAUCACG

>bna-miR395b MIMAT0023637

CUGAAGUGUUUGGGGGAACUC

>bna-miR167a MIMAT0005626

UGAAGCUGCCAGCAUGAUCUAA

>bna-miR2111b-3p MIMAT0011781

AUCCUCGGGAUACAGAUUACC

>bna-miR169a MIMAT0005612

CAGCCAAGGAUGACUUGCCGA

>bna-miR160c MIMAT0023619

UGCCUGGCUCCCUGUAUGCCA

>bna-miR166b MIMAT0005630

UCGGACCAGGCUUCAUUCCCC

>bna-miR156d MIMAT0023613

UGACAGAAGAGAGUGAGCAC

>bna-miR169g MIMAT0005618

UAGCCAAGGAUGACUUGCCUGC

>bna-miR159 MIMAT0005635

UUUGGAUUGAAGGGAGCUCUA

>bna-miR171g MIMAT0004446

UGAUUGAGCCGCGCCAAUAUCU

>bna-miR156e MIMAT0023614

UGACAGAAGAGAGUGAGCAC

>bna-miR169d MIMAT0005615

UAGCCAAGGAUGACUUGCCUA

>bna-miR6028 MIMAT0023648

UGGAGAGUAAGGACAUUCAGA

>bna-miR171d MIMAT0005608

UUGAGCCGUGCCAAUAUCACG

>bna-miR169b MIMAT0005613

CAGCCAAGGAUGACUUGCCGA

>bna-miR166f MIMAT0023625

UCGGACCAGGCUUCAUCCCCC

>bna-miR6036 MIMAT0023656

AUAGUACUAGUACUUGCAUGAUCA

>bna-miR397a MIMAT0005600

UCAUUGAGUGCAGCGUUGAUGU

>bna-miR171f MIMAT0005610

UGAUUGAGCCGCGCCAAUAUC

>bna-miR6035 MIMAT0023655

UGGAGUAGAAAAUGCAGUCGU

>bna-miR160a MIMAT0023617

UGCCUGGCUCCCUGUAUGCCA

>bna-miR169f MIMAT0005617

UAGCCAAGGAUGACUUGCCUA

>bna-miR395a MIMAT0023636

CUGAAGUGUUUGGGGGAACUC

>bna-miR172b MIMAT0023631

GGAAUCUUGAUGAUGCUGCAU

>bna-miR169h MIMAT0005619

UAGCCAAGGAUGACUUGCCUGC

>bna-miR399a MIMAT0004449

UGCCAAAGGAGAUUUGCCCGG

>bna-miR399b MIMAT0023642

UGCCAAAGGAGAUUUGCCCGG

>bna-miR394b MIMAT0023635

UUGGCAUUCUGUCCACCUCC

>bna-miR169n MIMAT0023629

CAGCCAAGGAUGACUUGCCGG

>bna-miR167c MIMAT0005628

UGAAGCUGCCAGCAUGAUCUA

>bna-miR166a MIMAT0005629

UCGGACCAGGCUUCAUUCCCC

>bna-miR395f MIMAT0023641

CUGAAGUGUUUGGGGGGACUC

>bna-miR396a MIMAT0004448

UUCCACAGCUUUCUUGAACUU

>bna-miR171b MIMAT0005606

UUGAGCCGUGCCAAUAUCACG

>bna-miR395d MIMAT0023639

CUGAAGUGUUUGGGGGGACUC

>bna-miR162a MIMAT0023621

UCGAUAAACCUGUGCAUCCAG

>bna-miR399c MIMAT0023643

UGCCAAAGGAGAUUUGUCCGG

>bna-miR169e MIMAT0005616

UAGCCAAGGAUGACUUGCCUA

>bna-miR172c MIMAT0023632

GGAAUCUUGAUGAUGCUGCAU

>bna-miR2111b-5p MIMAT0011780

UAAUCUGCAUCCUGAGGUUUA

>bna-miR160d MIMAT0023620

UGCCUGGCUCCCUGUAUGCCA

>bna-miR394a MIMAT0023634

UUGGCAUUCUGUCCACCUCC

>bna-miR2111a-3p MIMAT0011783

GUCCUCGGGAUGCGGAUUACC

>bna-miR156a MIMAT0004445

UGACAGAAGAGAGUGAGCACA

>bna-miR169i MIMAT0005620

UAGCCAAGGAUGACUUGCCUGC

>bna-miR171e MIMAT0005609

UUGAGCCGUGCCAAUAUCACG

>bna-miR169m MIMAT0005624

UGAGCCAAAGAUGACUUGCCG

>bna-miR403 MIMAT0023644

UUAGAUUCACGCACAAACUCG

>bna-miR156c MIMAT0005639

UUGACAGAAGAUAGAGAGCAC

>bna-miR171c MIMAT0005607

UUGAGCCGUGCCAAUAUCACG

>bna-miR390c MIMAT0005604

AAGCUCAGGAGGGAUAGCGCC

>bna-miR860 MIMAT0023645

UCAAUACAUUGGACUACAUAU

>bna-miR6029 MIMAT0023649

UGGGGUUGUGAUUUCAGGCUU

>bna-miR395e MIMAT0023640

CUGAAGUGUUUGGGGGGACUC

>bna-miR6033 MIMAT0023653

UGAACCAGAUAGAGUGGGACU

>bna-miR6034 MIMAT0023654

UCUGAUGUAUAUAGCUUUGGG

>bna-miR2111c MIMAT0023647

UAAUCUGCAUCCUGGGGUUUA

>bna-miR169c MIMAT0005614

UAGCCAAGGAUGACUUGCCUA

>bna-miR164c MIMAT0023623

UGGAGAAGCAGGGCACGUGCG

>bna-miR169j MIMAT0005621

UAGCCAAGGAUGACUUGCCUGC

>bna-miR2111d MIMAT0023646

UAAUCUGCAUCCUGAGGUUUA

>bna-miR2111a-5p MIMAT0011782

UAAUCUGCAUCCUGAGGUUUA

>bna-miR156b MIMAT0005636

UUGACAGAAGAUAGAGAGCAC

>bna-miR164d MIMAT0023624

UGGAGAAGCAGGGCACGUGCG

>bna-miR395c MIMAT0023638

CUGAAGUGUUUGGGGGAACUC

>bna-miR168a MIMAT0005625

UCGCUUGGUGCAGGUCGGGAA

>bna-miR172d MIMAT0023630

AGAAUCUUGAUGAUGCUGCAG

>bna-miR6032 MIMAT0023652

UGGAGCAUCAACAGAUCUCGG

>bna-miR6031 MIMAT0023651

AAGAGGUUCGGAGCGGUUUGAAGC

>bna-miR156f MIMAT0023615

UGACAGAAGAGAGUGAGCAC

>bna-miR167d MIMAT0023627

UGAAGCUGCCAGCAUGAUCU

>bna-miR166c MIMAT0005631

UCGGACCAGGCUUCAUUCCCC

>bol-miR9408 MIMAT0035363

GUUUCAUCUUAGAGAAUGUUGUC

>bol-miR9409 MIMAT0035364

UUUUGUUCAUGACUGCAUUUUC

>bol-miR172b MIMAT0010172

AGAAUCUUGAUGAUGCUGCAU

>bol-miR824 MIMAT0005597

UAGACCAUUUGUGAGAAGGGA

>bol-miR9411 MIMAT0035366

UACUGGACGACUUACACGGAAG

>bol-miR9410 MIMAT0035365

UACUUAAUUAUAAGUCGUCUGG

>bol-miR398a-5p MIMAT0010173

GGAGUGUCAUGAGAACACGGA

>bol-miR171a MIMAT0010170

UUGAGCCGUGCCAAUAUCACG

>bol-miR172a MIMAT0010171

AGAAUCUUGAUGAUGCUGCAU

>bol-miR398a-3p MIMAT0010174

UGUGUUCUCAGGUCACCCCUU

>bol-miR157a MIMAT0010169

UUGACAGAAGAUAGAGAGCAC

>bra-miR9565-3p MIMAT0035690

CUGAAGCUAGUGAAAGAGAGA

>bra-miR167b MIMAT0010158

UGAAGCUGCCAGCAUGAUCUA

>bra-miR9553-5p MIMAT0035659

UACAAAGCUGAAGCUAAUUAUG

>bra-miR5717 MIMAT0023014

GUUUGGAUUGUUUGCCUUGGC

>bra-miR396-3p MIMAT0035853

GCUCAAGAAAGCUGUGGGAAA

>bra-miR395b-3p MIMAT0035847

CUGAAGUGUUUGGGGGAACUC

>bra-miR860-3p MIMAT0035861

UCAAUACAUUGGACUACAUAU

>bra-miR167d MIMAT0010160

UGAAGCUGCCAGCAUGAUCUA

>bra-miR9567-3p MIMAT0035694

AAACUAUAUGUGUUGCUUAGA

>bra-miR9563b-5p MIMAT0035695

AAGAACUCGUCUCUUAACUUUUAA

>bra-miR171e MIMAT0010165

UGAUUGAGCCGCGCCAAUAUC

>bra-miR1140 MIMAT0023021

ACAGCCUAAACCAAUCGGAGC

>bra-miR172d-3p MIMAT0035877

GGAAUCUUGAUGAUGCUGCAU

>bra-miR156a-5p MIMAT0035816

UGACAGAAGAGAGUGAGCAC

>bra-miR9563b-3p MIMAT0035696

AAAUUAAGAGAUGAAUUCUUAC

>bra-miR164d-3p MIMAT0035867

CACGUGUUCUACUACUCCAAC

>bra-miR171b MIMAT0010162

UUGAGCCGUGCCAAUAUCACG

>bra-miR9564-3p MIMAT0035688

CGAGCUGUGUAAUCGUUUUGUU

>bra-miR172a MIMAT0010166

AGAAUCUUGAUGAUGCUGCAU

>bra-miR400-3p MIMAT0035857

GACUUAUAAUGAUCUCAUGAA

>bra-miR168c-5p MIMAT0035834

UCGCUUGGUGCAGGUCGGGAC

>bra-miR5718 MIMAT0023016

UCAGAACCAAACACAGAACAAG

>bra-miR168b-5p MIMAT0035832

UCGCUUGGUGCAGGUCGGGAC

>bra-miR9555b-5p MIMAT0035679

UGUAAUUGCGGGGUUCUAAGC

>bra-miR5716 MIMAT0023013

UUGGAUAAUUGAAGAUAUAAA

>bra-miR156a-3p MIMAT0035817

GCUUACUCUCUCUCUGUCACC

>bra-miR408-5p MIMAT0035683

GGGAGCCAGGGAAGAGGCAGU

>bra-miR2111a-3p MIMAT0016369

GUCCUCGGGAUGCGGAUUACC

>bra-miR2111b-5p MIMAT0016370

UAAUCUGCAUCCUGAGGUUUA

>bra-miR172c-5p MIMAT0035872

GCAUCAUCAUCAAGAUUCAGA

>bra-miR156b-5p MIMAT0035818

UGACAGAAGAGAGUGAGCAC

>bra-miR9564-5p MIMAT0035687

ACAAAACGAUUACACAGCUCGGUC

>bra-miR162-5p MIMAT0035830

GGAGGCAGCGGUUCAUCGAUC

>bra-miR9558-5p MIMAT0035669

AGAGAUGUCUGGCUUGCAACA

>bra-miR9554-5p MIMAT0035661

GAAUGAUACUUGGAUAUAAUC

>bra-miR2111-5p MIMAT0035874

UAAUCUGCAUCCUGGGGUUUA

>bra-miR9558-3p MIMAT0035670

UUGCAAGCCAGACAUUUCCUUU

>bra-miR1885a MIMAT0009213

CAUCAAUGAAAGGUAUGAUUCC

>bra-miR9552b-3p MIMAT0035656

UGACCGAGUAGACCGAUAGUC

>bra-miR5714 MIMAT0023011

AGACUCUACGACAUCAAGAAAC

>bra-miR395a-3p MIMAT0035851

CUGAAGUGUUUGGGGGAACUC

>bra-miR9568-3p MIMAT0035698

UCAUCGUAAGAGAUCUGCAUU

>bra-miR9563a-5p MIMAT0035685

ACCCGUCUCUUAAUUUUUAAC

>bra-miR9562-5p MIMAT0035681

ACUAUGCAAUUGUGAACAAAC

>bra-miR164c-3p MIMAT0035869

CACGUGUUCUACUACUCCAAC

>bra-miR9556-5p MIMAT0035665

GUCAAUUGGUGAUAGUAGUUC

>bra-miR9560a-5p MIMAT0035673

ACAGGUGGUGGAACAAAUAUGAGU

>bra-miR9565-5p MIMAT0035689

UCUCGUUCUCUCGUUUCAGCU

>bra-miR9567-5p MIMAT0035693

UAAACAACACAUAUAGUUUGC

>bra-miR5721 MIMAT0023019

AAAAAUGGAGUGAGAAAUGGA

>bra-miR161-3p MIMAT0035879

GUCACUUUCAAUGCGUUGAUC

>bra-miR319-3p MIMAT0035839

UUGGACUGAAGGGAGCUCCCU

>bra-miR9556-3p MIMAT0035666

UCUACUUUCACCAAUUGGCCU

>bra-miR156c-5p MIMAT0035820

UGACAGAAGAGAGUGAGCAC

>bra-miR164b-5p MIMAT0035864

UGGAGAAGCAGGGCACGUGCG

>bra-miR395d-3p MIMAT0035845

CUGAAGUGUUUGGGGGAACUC

>bra-miR172b-3p MIMAT0010168

AGAAUCUUGAUGAUGCUGCAU

>bra-miR9408-5p MIMAT0035657

CAACAGUCUCAGGAUGGAAAA

>bra-miR172b-5p MIMAT0010167

GCAGCACCAUUAAGAUUCACA

>bra-miR400-5p MIMAT0035856

UAUGAGAGUAUUAUAAGUCAC

>bra-miR9568-5p MIMAT0035697

UGCGGAUAUCUUAGGAUGAGGU

>bra-miR2111b-3p MIMAT0016371

AUCCUCGGGAUACGGAUUACC

>bra-miR164e-5p MIMAT0035870

UGGAGAAGCAGGGCACGUGCAA

>bra-miR161-5p MIMAT0035878

UCAAUGCACUGAAAGUGACUA

>bra-miR158-5p MIMAT0035701

CUUUGUCUAUCGUUUGGAAAAG

>bra-miR9552b-5p MIMAT0035655

CUAUCGGUCUACUUGGUCAGC

>bra-miR390-3p MIMAT0035841

CGCUGUCCAUCCUGAGUUUCA

>bra-miR9555b-3p MIMAT0035680

UUAGAAACUUGCAAUUAUAUA

>bra-miR156b-3p MIMAT0035819

GCUCACUCUCUAUCUGUCACC

>bra-miR156e-5p MIMAT0035824

UGACAGAAGAGAGUGAGCAC

>bra-miR408-3p MIMAT0035684

UGCUUGUUCCCUGUCUCUCUC

>bra-miR9566-5p MIMAT0035691

UUGUUGACAAAUACUUAGGCUC

>bra-miR167c MIMAT0010159

UGAAGCUGCCAGCAUGAUCUA

>bra-miR9562-3p MIMAT0035682

UUAUUCACAACUGCAUAAUUC

>bra-miR5654b MIMAT0023008

AUAAAUCCCAAGCAUCAUCCA

>bra-miR5723 MIMAT0023022

AAUGUGCUGCAAUAUCUCUGC

>bra-miR391-3p MIMAT0035843

ACGGUAUCUCUCCUACGUAGC

>bra-miR9552a-5p MIMAT0035653

CUAUCGGUCUACUCGGUCAGC

>bra-miR156c-3p MIMAT0035821

GCUCACUGCUCUAUCUGUCAGA

>bra-miR5654a MIMAT0023007

AUAAAUCCCAAGCAUCAUCCA

>bra-miR5711 MIMAT0023006

UGUUUUGUGGGUUUCUACCGA

>bra-miR2111a-5p MIMAT0016368

UAAUCUGCAUCCUGAGGUUUA

>bra-miR9555a-5p MIMAT0035663

UUCUAAGCUUUACGGGAAACC

>bra-miR390-5p MIMAT0035840

AAGCUCAGGAGGGAUAGCGCC

>bra-miR9569-3p MIMAT0035700

ACACAGGAACAAUACUAACUCAUU

>bra-miR159a MIMAT0010153

UUUGGAUUGAAGGGAGCUCUA

>bra-miR156d-3p MIMAT0035823

GCUCACUCUCUAUCUGUCACC

>bra-miR395a-5p MIMAT0035850

GUUCCUCUGAGCACUUCAUUG

>bra-miR391-5p MIMAT0035842

UUCGCAGGAGAGAUAGCGCCA

>bra-miR9553-3p MIMAT0035660

UAAUCAGCUCCAGCUAUGUACA

>bra-miR5719 MIMAT0023017

UUGUGAUGAUAAUACGACUUC

>bra-miR162-3p MIMAT0035831

UCGAUAAACCUCUGCAUCCAG

>bra-miR9552a-3p MIMAT0035654

UGACCAAGUAGACCGAUAGUC

>bra-miR395b-5p MIMAT0035846

GUUCCUCUGAGCACUUCAUUG

>bra-miR9561-5p MIMAT0035677

UGAGUCUCUCACCAGUCUUUCAC

>bra-miR860-5p MIMAT0035860

AUGUAGUCCAAUCUAUUGAAG

>bra-miR5713 MIMAT0023010

AGGCUUAGAAGAACGUUUGUU

>bra-miR6032-3p MIMAT0035863

UCUGCUGGUCGUUCCAUGUUAA

>bra-miR398-3p MIMAT0035855

UGUGUUCUCAGGUCACCCCUG

>bra-miR9408-3p MIMAT0035658

UUUCAUCUUAGAGAAUGUUGUU

>bra-miR167a MIMAT0010157

UGAAGCUGCCAGCAUGAUCUA

>bra-miR5725 MIMAT0023024

AUUUGGCACAAUCUGAUCUGC

>bra-miR824 MIMAT0005598

UAGACCAUUUGUGAGAAGGGA

>bra-miR9559-3p MIMAT0035672

ACAAUGAACGAAAUCCAAAUC

>bra-miR9560b-3p MIMAT0035676

UCAUAUUAGUUCUACCUCCUGCUG

>bra-miR5726 MIMAT0023025

CAAAGGUUGCUUGAAUAAGGU

>bra-miR172c-3p MIMAT0035873

AGAAUCUUGAUGAUGCUGCAG

>bra-miR396-5p MIMAT0035852

UUCCACAGCUUUCUUGAACUU

>bra-miR156f-3p MIMAT0035827

UGCUCACUGCUCUUUCUGUCAGA

>bra-miR171d MIMAT0010164

UUGAGCCGUGCCAAUAUCACG

>bra-miR160a-5p MIMAT0010154

UGCCUGGCUCCCUGUAUGCCA

>bra-miR156f-5p MIMAT0035826

UGACAGAAGAGAGUGAGCAC

>bra-miR156g-3p MIMAT0035829

GCUCACUGCUCUAUCUGUCAGA

>bra-miR395c-5p MIMAT0035848

GUUCCUCUGAGCACUUCAUUG

>bra-miR9561-3p MIMAT0035678

GAGAGACUCUGAAAGACUCACC

>bra-miR319-5p MIMAT0035838

AGAGCUUCCUUGAGUCCAUUC

>bra-miR403-5p MIMAT0035858

UGUUUUGUGCGUGAAUCUAAUU

>bra-miR168a-5p MIMAT0035836

UCGCUUGGUGCAGGUCGGGAA

>bra-miR157a MIMAT0010152

UUGACAGAAGAUAGAGAGCAC

>bra-miR9560a-3p MIMAT0035674

UCAUAUUAGUUCUACCUCCUGUUG

>bra-miR156d-5p MIMAT0035822

UGACAGAAGAGAGUGAGCAC

>bra-miR2111-3p MIMAT0035875

GACCUCAGGAUGCGGAUUACC

>bra-miR164b-3p MIMAT0035865

CACGUGUUCUACUACUCCAAC

>bra-miR5715 MIMAT0023012

ACGUGAUAAGCCUCUGAAGAA

>bra-miR164a MIMAT0010156

UGGAGAAGCAGGGCACGUGCA

>bra-miR172d-5p MIMAT0035876

GCAGCAUCAUUAAGAUUCACA

>bra-miR5712 MIMAT0023009

AAUAUUAAUAUAAUUGGUGAG

>bra-miR168c-3p MIMAT0035835

CCCGCCUUGCAUCAACUGAAU

>bra-miR395d-5p MIMAT0035844

GUUCCUCUGAGCACUUCAUUG

>bra-miR156g-5p MIMAT0035828

UGACAGAAGAGAGUGAGCAC

>bra-miR164d-5p MIMAT0035866

UGGAGAAGCAGGGCACGUGCG

>bra-miR9557-5p MIMAT0035667

UUUUGCGUUUCAACUCGGUCC

>bra-miR9555a-3p MIMAT0035664

UUUCCCGUAAAGCUUAGAACC

>bra-miR9557-3p MIMAT0035668

GCUGAGUUGGAACGCAAAAUC

>bra-miR171a MIMAT0010161

UUGAGCCGUGCCAAUAUCACG

>bra-miR1885b MIMAT0023015

UACAUCUUCUCCGCGGAAGCUC

>bra-miR398-5p MIMAT0035854

GGGUCGACAUGAGAACACAUG

>bra-miR5720 MIMAT0023018

UUGUGAUUUGGUUGGAAUAUC

>bra-miR164c-5p MIMAT0035868

UGGAGAAGCAGGGCACGUGCG

>bra-miR9560b-5p MIMAT0035675

ACAGGUGGUGGAACAAAUAUGAGU

>bra-miR9554-3p MIMAT0035662

UCAUAUCCAAGUAUCAUUCCU

>bra-miR9559-5p MIMAT0035671

UUUGGAUUUUGGUCAUUGUUG

>bra-miR9563a-3p MIMAT0035686

UAAAAGUUAAGAGACAAGUUA

>bra-miR395c-3p MIMAT0035849

CUGAAGUGUUUGGGGGAACUC

>bra-miR5724 MIMAT0023023

AACCGCCGGUUUGAUAAUAGC

>bra-miR9569-5p MIMAT0035699

UGAGUUAUCAUUGGUCUUGUG

>bra-miR164e-3p MIMAT0035871

CACGUGCUCCCCUCCUCCAAC

>bra-miR5722 MIMAT0023020

UGAAAUAGAGUCAUGUGGAACG

>bra-miR403-3p MIMAT0035859

UUAGAUUCACGCACAAACUCG

>bra-miR158-3p MIMAT0035702

UUUCCAAAUGUAGACAAAGCA

>bra-miR156e-3p MIMAT0035825

UGCUCACCUCUCUUUCUGUCAGU

>bra-miR168a-3p MIMAT0035837

CCCGCCUUGUAUCAAGUGAAU

>bra-miR171c MIMAT0010163

UUGAGCCGUGCCAAUAUCACG

>bra-miR168b-3p MIMAT0035833

CCCGCCUUGCAUCAACUGAAU

>bra-miR6032-5p MIMAT0035862

AACAUGGAGCAUCAACAGAUC

>bra-miR9566-3p MIMAT0035692

GAGCCUAAGUAUUUGUCAACAAUG

>bra-miR160a-3p MIMAT0010155

GCGUAUGAGGAGCCAUGCAUA

>cas-miR162b MIMAT0045372

AUCGAUGAACCGCUGCCUCC

>cas-miR172b MIMAT0045327

AGAAUCUUGAUGAUGCUGCAU

>cas-miR156e-5p MIMAT0045272

UGACAGAAGAGAGUGAGCAC

>cas-miR8171 MIMAT0045369

AUAGGUGGGCCAGUGGUAGGA

>cas-miR828 MIMAT0045360

UCUUGCUUAAAUGAGUAUUCC

>cas-miR399b MIMAT0045353

GGGCAAGAUCACCAUUGGCAGA

>cas-miR169b MIMAT0045311

CAGCCAAGGAUGACUUGCCGG

>cas-miR166a MIMAT0045298

GGAAUGUUGUCUGGCUCGAGG

>cas-miR167a MIMAT0045306

GAUCAUGUUCGCAGUUUCACC

>cas-miR156e-3p MIMAT0045273

GCUUACUCUCUCUCUGUCACC

>cas-miR167b MIMAT0045307

GGUCAUGCUGUGACAGCCUCACU

>cas-miR171b MIMAT0045322

UUGAGCCGUGCCAAUAUCACG

>cas-miR171a-5p MIMAT0045320

UAUUGGCCUGGUUCACUCAGA

>cas-miR156c-5p MIMAT0045268

UGACAGAAGAGAGUGAGCAC

>cas-miR156b-3p MIMAT0045267

UGCUCACCUCUCUUUCUGUCAGU

>cas-miR171c-5p MIMAT0045323

AGAUAUUGGUGCGGUUCAAUC

>cas-miR827a MIMAT0045359

UUAGAUGACCAUCAACAAACG

>cas-miR157c MIMAT0045284

GCUCUCUAUACUUCUGUCACC

>cas-miR172a-3p MIMAT0045326

AGAAUCUUGAUGAUGCUGCAU

>cas-miR165a MIMAT0045296

UCGGACCAGGCUUCAUCCCCC

>cas-miR169a MIMAT0045310

CAGCCAAGGAUGACUUGCCGA

>cas-miR408 MIMAT0045356

AUGCACUGCCUCUUCCCUGGC

>cas-miR396b MIMAT0045347

UUCCACAGCUUUCUUGAACUU

>cas-miR156f-5p MIMAT0045274

UGACAGAAGAGAGUGAGCAC

>cas-miR395c-3p MIMAT0045343

CUGAAGUGUUUGGGGGGACUC

>cas-miR161 MIMAT0045293

UCAAUGCAUUGAAAGUGACUA

>cas-miR160b-3p MIMAT0045292

GCGUACAGAGUAGUCAAGCAUG

>cas-miR164 MIMAT0045295

UGGAGAAGCAGGGCACGUGCA

>cas-miR156k-3p MIMAT0045281

UGCUUACUCUCUAUCUGUCACC

>cas-miR169e-5p MIMAT0045314

UAGCCAAGGAUGACUUGCCUG

>cas-miR390b MIMAT0045336

AAGCUCAGGAGGGAUAGCGCC

>cas-miR395c-5p MIMAT0045342

GUUCCCUUUAACGCUUCAUUG

>cas-miR398a-5p MIMAT0045349

GGGUCGACAUGAGAACACAUG

>cas-miR159a MIMAT0045289

UUUGGAUUGAAGGGAGCUCCU

>cas-miR159c-5 MIMAT0045263

AGAAGAAGAGGAAGAGCUCCU

>cas-miR159b-5p MIMAT0045287

AGCUGCUAAGCUAUGGAUCCC

>cas-miR157d MIMAT0045285

CAGAAGAUAGAGAGCACUGA

>cas-miR860 MIMAT0045364

UCAAUAGAUUGGACUAUAUAU

>cas-miR166c-5p MIMAT0045300

GGAAUGUUGUCUGGCACGAGG

>cas-miR398b MIMAT0045351

UGUGUUCUCAGGUCACCCCU

>cas-miR319a MIMAT0045331

AGAGCUUCCUUGAGUCCAUUC

>cas-miR159b-3p MIMAT0045288

UUUGGAUUGAAGGGAGCUCUU

>cas-miR390a-3p MIMAT0045335

CGCUAUCCAUCCUGAGUUUCA

>cas-miR156f-3p MIMAT0045275

GCUCACUCUCUAUCCGUCACC

>cas-miR393-3p MIMAT0045338

AUCAUGCUAUCUCUUUGGAUU

>cas-miR166d MIMAT0045302

UCGGACCAGGCUUCAUUCCCCU

>cas-miR156h MIMAT0045277

UGACAGAAGAAAGAGAGCAC

>cas-miR395d-5p MIMAT0045344

GUUCUCCCGAACACUUCAUUG

>cas-miR172a-5p MIMAT0045325

GUGGCAUCAUCAAGAUUCACA

>cas-miR159c-3p MIMAT0045264

UUUGGAUUGAAGGGAGCUCUU

>cas-miR168 MIMAT0045309

CCCGCCUUGCAUCAACUGAAU

>cas-miR173 MIMAT0045330

UUCGCUUGCAGAGAGAAAUCAC

>cas-miR403 MIMAT0045355

UUAGAUUCACGCACAAACUCGU

>cas-miR156k-5p MIMAT0045280

UGACAGAAGAGAGUGAGCAC

>cas-miR9560 MIMAT0045370

ACAGGUGGUGGAACAAAUAUGAGU

>cas-miR858 MIMAT0045363

UUUCGUUGUCUGUUCGACCUU

>cas-miR169e-3p MIMAT0045315

GGCAGUCUCCUUGGCUAUC

>cas-miR390a-5p MIMAT0045334

AAGCUCAGGAGGGAUAGCGCC

>cas-miR160a MIMAT0045290

UGCCUGGCUCCCUGUAUGCC

>cas-miR158a MIMAT0045286

CUGUGCUUCUUUGUCUACAAU

>cas-miR172c MIMAT0045328

GGAGCAUCAUCAAGAUUCACA

>cas-miR157b MIMAT0045283

GCUCUCUAUACUUCUGUCACC

>cas-miR166f-5p MIMAT0045304

GGAAUGUUGUUUGGCUCGAGG

>cas-miR395b MIMAT0045341

CUGAAGUGUUUGGGGGGACUC

>cas-miR166f-3p MIMAT0045305

UCGGACCAGGCUUCAUUCCCCU

>cas-miR156a MIMAT0045265

UGACAGAAGAGAGUGAGCAC

>cas-miR319b MIMAT0045332

AGAGCUUUCUUCGGUCCACUC

>cas-miR156d-5p MIMAT0045270

UGACAGAAGAGAGUGAGCAC

>cas-miR171a-3p MIMAT0045321

UGAUUGAGCCGCGCCAAUAUCU

>cas-miR160b-5p MIMAT0045291

UGCCUGGCUCCCUGUAUGCC

>cas-miR397 MIMAT0045348

UCAUUGAGUGCAGCGUUGAUG

>cas-miR857 MIMAT0045362

AAACUUUCACCAUACAAAAUAA

>cas-miR827b MIMAT0045262

UAAGAUGACCAUCAACAAAC

>cas-miR845 MIMAT0045361

CGGCUCUGAUACCAAUUGAUG

>cas-miR158b MIMAT0045261

UCCGAAAUGUAGACAAAGCA

>cas-miR169d MIMAT0045313

GCAAGUUGACCUUGGCUCUGU

>cas-miR162a MIMAT0045294

UCGAUAAACCUCUGCAUCCA

>cas-miR166e MIMAT0045303

UCGGACCAGGCUUCAUUCCCU

>cas-miR472 MIMAT0045357

AUGGUCGAAGUAGGCAAA

>cas-miR156g MIMAT0045276

GACAGAAGAGAGUGAGCAC

>cas-miR2111a MIMAT0045365

GUCCUCGGGAUGCGGAUUACC

>cas-miR3435 MIMAT0045367

GGCAUAAUUCUGUCAAUUCC

>cas-miR2111b MIMAT0045366

AUCCUCGGGAUACGGAUUACC

>cas-miR394 MIMAT0045339

UUUGGCAUUCUGUCCACCUCC

>cas-miR157a MIMAT0045282

GCUCUCUAGCCUUCUGUCAUCA

>cas-miR169g-3p MIMAT0045318

GGCAGUCUCCUUGGCUAUU

>cas-miR400 MIMAT0045354

UAUGAGAGUAUUAUAAGUCAC

>cas-miR395a MIMAT0045340

CUGAAGUGUUUGGGGGAACUC

>cas-miR156c-3p MIMAT0045269

GCUCACUGCUCUAUCUGUCAGA

>cas-miR156b-5p MIMAT0045266

UGACAGAAGAGAGUGAGCAC

>cas-miR319c MIMAT0045333

UUUGGACUGAAGGGAGCUCCU

>cas-miR166c-3p MIMAT0045301

UCGGACCAGGCUUCAUUCCCCU

>cas-miR156j MIMAT0045279

UGACAGAAGAGAGAGAGCAC

>cas-miR393-5p MIMAT0045337

UCCAAAGGGAUCGCAUUGAUCC

>cas-miR825 MIMAT0045358

UCAAGCACCAGCUCGAAGAAGC

>cas-miR170 MIMAT0045319

UAUUGGCCUGGUUCACUCAGA

>cas-miR165b MIMAT0045297

GAAUGUUGUUUGGAUCGAGG

>cas-miR171c-3p MIMAT0045324

UUGAGCCGUGCCAAUAUCACG

>cas-miR166b MIMAT0045299

GGACUGUUGUCUGGCUCGAGG

>cas-miR156d-3p MIMAT0045271

GCUCACUCUCUUUCUGUCAUA

>cas-miR169c MIMAT0045312

UAGCCAAGGAUGACUUGCCUG

>cas-miR396a MIMAT0045346

UUCCACAGCUUUCUUGAACUG

>cas-miR156i MIMAT0045278

CAGAAGAGAGAGAGCACAAU

>cas-miR172d MIMAT0045329

GCAACAUCUUCAAGAUUCAGA

>cas-miR11592 MIMAT0045371

GAACCGAACCGAACCGAAA

>cas-miR395d-3p MIMAT0045345

CUGAAGUGUUUGGGGGAACUC

>cas-miR169g-5p MIMAT0045317

UAGCCAAGGAUGACUUGCCUG

>cas-miR398a-3p MIMAT0045350

UGUGUUCUCAGGUCACCCCU

>cas-miR399a MIMAT0045352

GGGCAUCUUUCUAUUGGCAGG

>cas-miR169f MIMAT0045316

GCAAGUUGACCUUGGCUCUGC

>cas-miR5139 MIMAT0045368

AAACCUGGCUCUGAUACCA

>cas-miR167c MIMAT0045308

AGGUCAUGCUGGUAGCUUCAC

>cpa-miR8152 MIMAT0031849

UUGGACUGCUAGGUGGCCCAU

>cpa-miR166d MIMAT0031799

UCGGACCAGGCUUCAUUCCCG

>cpa-miR164b MIMAT0031792

UGGAGAAGCAGGGCACGUGCA

>cpa-miR8140 MIMAT0031837

CUUUUCAAGACUUCAGCUUCA

>cpa-miR8144 MIMAT0031841

AACAGUAGAACGAGUUAGAAAGGA

>cpa-miR5211 MIMAT0031851

GUGCCGUCGCACUGUGACAAG

>cpa-miR160b MIMAT0031784

UGCCUGGCUCCCUGUAUGCCA

>cpa-miR8146 MIMAT0031843

AGGAAGACGGUGAGUAGAAGCCAA

>cpa-miR160f-5p MIMAT0031789

UGCCUGGCUCCCUGUAUGCCA

>cpa-miR8139b MIMAT0031833

UCCUGGCUGAGGACGGGUGUUGAA

>cpa-miR319 MIMAT0031782

AUUGGACUGAAGGGAGCUCC

>cpa-miR160c-3p MIMAT0031786

GCGUAUGAGGAGCCAUGCAUA

>cpa-miR8153 MIMAT0031850

UGCACUGUAGAGCCGUAUUCGGAC

>cpa-miR395e MIMAT0031821

UGAAGUGUUUGGGGGAACUC

>cpa-miR172b MIMAT0031811

GGGAAUCUUGAUGAUGCUGCA

>cpa-miR8139a MIMAT0031832

UCCUGGCUGAGGACGGGUGUUGAA

>cpa-miR8137 MIMAT0031830

UUCGCCAGCCAUUCACAAAAU

>cpa-miR8139c MIMAT0031834

UCCUGGCUGAGGACGGGUGUUGAA

>cpa-miR169 MIMAT0031805

CAGCCAAGAAUGACUUGCCG

>cpa-miR8138 MIMAT0031831

UAAAGAUGGUAACAAAGGAUAA

>cpa-miR160d MIMAT0031787

UGCCUGGCUCCCUGAAUGCCA

>cpa-miR8147 MIMAT0031844

ACUGAUACUUGAUGAAUUUGCAUG

>cpa-miR172a MIMAT0031810

GGGAAUCUUGAUGAUGCUGCA

>cpa-miR164e MIMAT0031795

UGGAGAAGGGGAGCACGUGCA

>cpa-miR8139d MIMAT0031835

UCCUGGCUGAGGACGGGUGUUGAA

>cpa-miR171c MIMAT0031808

UGAUUGAGCCGUGCCAAUAUC

>cpa-miR160f-3p MIMAT0031790

GCGUAUGAGGAGCCAUGCAUA

>cpa-miR395c MIMAT0031819

UGAAGUGUUUGGGGGAACUC

>cpa-miR171a MIMAT0031806

UGAUUGAGCCGUGCCAAUAUC

>cpa-miR8139e MIMAT0031836

UCCUGGCUGAGGACGGGUGUUGAA

>cpa-miR535 MIMAT0031824

UGACAACGAGAGAGAGCACGC

>cpa-miR408 MIMAT0031823

CUGCACUGCCUCUUCCCUGGC

>cpa-miR164c MIMAT0031793

UGGAGAAGCAGGGCACGUGCA

>cpa-miR171d MIMAT0031809

UGAUUGAGCCGUGCCAAUAUC

>cpa-miR8155 MIMAT0031853

UAACCUGGCUCUGAUACCA

>cpa-miR8151 MIMAT0031848

UGGAUACCAGUAGACAGAUA

>cpa-miR159a MIMAT0031780

UUUGGAUUGAAGGGAGCUCUA

>cpa-miR167a MIMAT0031801

UGAAGCUGCCAGCAUGAUCUA

>cpa-miR156a MIMAT0031774

UGACAGAAGAGAGUGAGCAC

>cpa-miR8150 MIMAT0031847

AAAACCUGAGUCAGAUGAUGAGCG

>cpa-miR166e MIMAT0031800

GGACCAGGCUUCAUUCCCC

>cpa-miR396 MIMAT0031822

UUCCACAGCUUUCUUGAACUG

>cpa-miR394b MIMAT0031816

UUGGCAUUCUGUCCACCUCC

>cpa-miR156d MIMAT0031777

UGACAGAAGAGAGUGAGCAC

>cpa-miR395d MIMAT0031820

UGAAGUGUUUGGGGGAACUC

>cpa-miR156c MIMAT0031776

UGACAGAAGAGAGUGAGCAC

>cpa-miR164d MIMAT0031794

UGGAGAAGGGGAGCACGUGCA

>cpa-miR390b MIMAT0031813

AAGCUCAGGAGGGAUAGCGCC

>cpa-miR159b MIMAT0031781

CUUGGAUUGAAGGGAGCUCC

>cpa-miR390a MIMAT0031812

AAGCUCAGGAGGGAUAGCGCC

>cpa-miR395a MIMAT0031817

UGAAGUGUUUGGGGGAACUC

>cpa-miR162a MIMAT0005567

UCGAUAAACCUCUGCAUCCAG

>cpa-miR156f MIMAT0031779

UUGACAGAAGAUAGAGAGCAC

>cpa-miR171b MIMAT0031807

UGAUUGAGCCGUGCCAAUAUC

>cpa-miR156b MIMAT0031775

UGACAGAAGAGAGUGAGCAC

>cpa-miR398 MIMAT0031826

GGGACGACAUGAGAUCACACG

>cpa-miR164a MIMAT0031791

UGGAGAAGCAGGGCACGUGCA

>cpa-miR8148 MIMAT0031845

UGACUGGGUCUGCUGACGUGGCAU

>cpa-miR8154 MIMAT0031852

CAGAGGAGGAGAUGAAGAGGGA

>cpa-miR393 MIMAT0031814

UCCAAAGGGAUCGCAUUGAUCC

>cpa-miR8142 MIMAT0031839

UGAGGUAAGUAGACAGUAAAGGUU

>cpa-miR160a MIMAT0031783

UGCCUGGCUCCCUGUAUGCCA

>cpa-miR167d MIMAT0031804

UGAAGCUGCCAGCAUGAUCUGA

>cpa-miR8136 MIMAT0031829

UAAAGUGGAAUUGGGAUAAUA

>cpa-miR167c MIMAT0031803

UGAAGCUGCCAGCAUGAUCUU

>cpa-miR8145 MIMAT0031842

AUAAACAGAUAGAAUGACAGCCUU

>cpa-miR477 MIMAT0031828

AUUGGAGGACUUUGGGGGAGC

>cpa-miR167b MIMAT0031802

UGAAGCUGCCAGCAUGAUCUA

>cpa-miR8134 MIMAT0031825

UGAGAAUUAUGCGGAGGAUGU

>cpa-miR166a MIMAT0031796

UCGGACCAGGCUUCAUUCCCC

>cpa-miR160e MIMAT0031788

UGCCUGGCUCCCUGUAUGCCA

>cpa-miR395b MIMAT0031818

UGAAGUGUUUGGGGGAACUC

>cpa-miR8135 MIMAT0031827

AGGAUUUUGCAGGGUUGAU

>cpa-miR8141 MIMAT0031838

UGGAUACUAGUAGGCUGGUU

>cpa-miR160c-5p MIMAT0031785

UGCCUGGCUCCCUGUAUGCCA

>cpa-miR166c MIMAT0031798

UCGGACCAGGCUUCAUUCCCC

>cpa-miR166b MIMAT0031797

UCGGACCAGGCUUCAUUCCCC

>cpa-miR8149 MIMAT0031846

AGCGAAGGGGACGCCUGAAGACUC

>cpa-miR8143 MIMAT0031840

GAGAGAUGGUGGACAGAUCAGGUA

>cpa-miR156e MIMAT0031778

UUGACAGAAGAUAGAGAGCAC

>cpa-miR394a MIMAT0031815

UUGGCAUUCUGUCCACCUCC

>cme-miR390b MIMAT0022750

AAGCUCAGGAGGGAUAGCGCC

>cme-miR172f MIMAT0026099

UGAAUCUUGAUGAUGCCGCAC

>cme-miR160d MIMAT0026178

UGCCUGGCUCCCUGAAUGCCA

>cme-miR169c MIMAT0026081

UAGCCAAAGAUGACUUGCCUG

>cme-miR399d MIMAT0026078

UGCCAAAGGAGAGUUGCCCUU

>cme-miR167d MIMAT0026163

UGAAGCUGCCAGCAUGAUCUG

>cme-miR169b MIMAT0026094

UAGCCAAAAAUGACUUGCCUG

>cme-miR156d MIMAT0026111

UGACAGAAGAGAGUGAGCAC

>cme-miR399a MIMAT0026150

UGCCAAAGGAGAUUUGCCCCG

>cme-miR319c MIMAT0026130

UUGGACUGAAGGGAGCUCCU

>cme-miR172e MIMAT0026126

AGAAUCUUGAUGAUGCUGCAG

>cme-miR162 MIMAT0026080

UCGAUAAACCUCUGCAUCCAG

>cme-miR166e MIMAT0026179

UCGGACCAGGCUUCAUUCCUC

>cme-miR168 MIMAT0022751

UCGCUUGGUGCAGGUCGGGA

>cme-miR7129 MIMAT0026183

AGUCAAAUCUAAACGAUCGUGUAU

>cme-miR395e MIMAT0026142

CUGAAGUGUUUGGGGGAACUC

>cme-miR530a MIMAT0026182

UGCAUUUGCACCUGCACCUU

>cme-miR156b MIMAT0026131

UUGACAGAAGAUAGAGAGCAC

>cme-miR164a MIMAT0022749

UGGAGAAGCAGGGCACGUGCU

>cme-miR395d MIMAT0026143

CUGAAGUGUUUGGGGGAACUC

>cme-miR172b MIMAT0026124

AGAAUCUUGAUGAUGCUGCAU

>cme-miR166g MIMAT0026165

UCGGACCAGGCUUCAUUCCC

>cme-miR166c MIMAT0026097

UCGGACCAGGCUUCAUUCCCC

>cme-miR399e MIMAT0026082

UGCCAAAGGAGAGUUGCCCUU

>cme-miR156i MIMAT0026105

UGACAGAAGAGAGUGAGCAC

>cme-miR169m MIMAT0026092

UAGCCAAAAAUGACUUGCCUGC

>cme-miR393b MIMAT0026137

UCCAAAGGGAUCGCAUUGAUC

>cme-miR172a MIMAT0026181

GGAAUCUUGAUGAUGCUGCAG

>cme-miR399b MIMAT0026167

UGCCAAAGGAGAGUUGCCCUA

>cme-miR396e MIMAT0026162

UUCCACGGCUUUCUUGAACUG

>cme-miR166i MIMAT0026091

UCGGACCAGGCUUCAUUCUC

>cme-miR828 MIMAT0026161

UCUUGCUCAAAUGAGUAUUCCA

>cme-miR167e MIMAT0026177

UCAAGCUGCCAGCAUGAUCUA

>cme-miR166b MIMAT0026103

UCGGACCAGGCUUCAUUCCCC

>cme-miR160a MIMAT0026072

UGCCUGGCUCCCUGUAUGCCA

>cme-miR169d MIMAT0026077

UAGCCAAAGAUGACUUGCCUG

>cme-miR390d MIMAT0026133

AAGCUCAGGAGGGAUAGCGCC

>cme-miR164d MIMAT0026090

UGGAGAAGCAGGGCACGUGCA

>cme-miR390c MIMAT0026135

AAGCUCAGGAGGGAUAGCGCC

>cme-miR319b MIMAT0026128

UUGGACUGAAGGGAGCUCCC

>cme-miR169t MIMAT0026187

UGAGCCAAGAAUGACUUGCCGGC

>cme-miR169a MIMAT0026095

UAGCCAAAAAUGACUUGCCUG

>cme-miR169l MIMAT0026098

UAGCCAAAAAUGACUUGCCUGC

>cme-miR395a MIMAT0026141

CUGAAGUGUUUGGGGGAACUC

>cme-miR160b MIMAT0026171

UGCCUGGCUCCCUGUAUGCCA

>cme-miR166f MIMAT0026109

UCGGACCAGGCUUCAUUCCCC

>cme-miR167a MIMAT0026112

UGAAGCUGCCAGCAUGAUCUA

>cme-miR169k MIMAT0026079

UGAGCCAAGGAUGACUUGCCU

>cme-miR156j MIMAT0026186

GUUGACAGAAGAGAGUGAGCAC

>cme-miR166h MIMAT0026108

UCGGACCAGGCUUCAUUCCCC

>cme-miR396a MIMAT0026106

UUCCACAGCUUUCUUGAACUU

>cme-miR156g MIMAT0026159

UGACAGAAGAGAGGGAGCAC

>cme-miR2111b MIMAT0026169

UAAUCUGCAUCCUGAGGUUUA

>cme-miR171a MIMAT0026119

UGAUUGAGCCGCGCCAAUAUC

>cme-miR854 MIMAT0026100

GAUGAGGAUAGUGAGGAGGAG

>cme-miR171e MIMAT0026118

UGAUUGAGCCGCGCCAAUAUC

>cme-miR166a MIMAT0026110

UCGGACCAGGCUUCAUUCCCC

>cme-miR167f MIMAT0026164

UGAAGCUGCCAGCAUGAUCUG

>cme-miR408 MIMAT0026153

AUGCACUGCCUCUUCCCUGGC

>cme-miR171b MIMAT0026123

UUGAGCCGUGCCAAUAUCACG

>cme-miR477a MIMAT0026089

ACCUCCCUCAAAGGCUUCCAA

>cme-miR169o MIMAT0026076

UAGCCAAAGAUGACUUGCCUG

>cme-miR319a MIMAT0026129

UUGGACUGAAGGGAGCUCCC

>cme-miR156f MIMAT0026151

UGACAGAAGAUAGAGAGCAC

>cme-miR171h MIMAT0026173

UGAUUGAGCCGCGUCAAUAUC

>cme-miR319d MIMAT0026132

UUGGACUGAAGGGAGCUCCU

>cme-miR171f MIMAT0026166

UGAUUGAGCCGUGCCAAUAUC

>cme-miR169q MIMAT0026073

UGAGCCAAAGAUGACUUGCCU

>cme-miR169p MIMAT0026075

UGAGCCAAAGAUGACUUGCCU

>cme-miR156e MIMAT0026156

UUGACAGAAGAUAGAGGGCAC

>cme-miR160c MIMAT0026180

UGCCUGGCUCCCUGUAUGCCA

>cme-miR169r MIMAT0026175

GAGCCAAGAAUGACUUGCCGG

>cme-miR156a MIMAT0026071

UGACAGAAGAGAGUGAGCAC

>cme-miR399g MIMAT0026185

AGGGCUUCUCUCCAUUGGCAGG

>cme-miR845 MIMAT0026084

UCGCUCUGAUACCAAUAUGAUG

>cme-miR171g MIMAT0026120

UGAUUGAGCCGCGCCAAUAUC

>cme-miR7130 MIMAT0026184

GUUUGGAAUGUGCGAGAUGUGUGC

>cme-miR171i MIMAT0026085

UUGAGCCGUUCCAAUAUCACA

>cme-miR169j MIMAT0026093

UAGCCAAAAAUGACUUGCCUGC

>cme-miR156c MIMAT0026121

UGACAGAAGAGAGUGAGCAC

>cme-miR1863 MIMAT0026101

AGCUCUGAUACCAUGUUAGAUUUG

>cme-miR394b MIMAT0026139

UUGGCAUUCUGUCCACCUCC

>cme-miR156h MIMAT0026158

UGACAGAAGAGAGGGAGCAC

>cme-miR393a MIMAT0026136

UCCAAAGGGAUCGCAUUGAUC

>cme-miR171d MIMAT0026122

UUGAGCCGUGCCAAUAUCACG

>cme-miR166d MIMAT0026107

UCGGACCAGGCUUCAUUCCCC

>cme-miR171c MIMAT0026154

UGAUUGAGCCGUGCCAAUAUC

>cme-miR169s MIMAT0026074

UGAGCCAAAGAUGACUUGCCU

>cme-miR399f MIMAT0026087

UGCCAAAGGAGAAUUGCAC

>cme-miR396c MIMAT0026146

UUCCACAGCUUUCUUGAACUU

>cme-miR172c MIMAT0026125

AGAAUCUUGAUGAUGCUGCAU

>cme-miR393c MIMAT0026138

UCCAAAGGGAUCGCAUUGAUC

>cme-miR399c MIMAT0026152

UGCCAAAGGAGAUUUGCCCGG

>cme-miR164b MIMAT0026088

UGGAGAGGCAGGGCACAUGCU

>cme-miR395b MIMAT0026157

UUGAAGUGUUUGGGGGAACUC

>cme-miR167b MIMAT0026113

UGAAGCUGCCAGCAUGAUCUA

>cme-miR167c MIMAT0026176

UGAAGCUGCCAGCAUGAUCUU

>cme-miR169n MIMAT0026116

UAGCCAAGGAUGACUUGCCUG

>cme-miR396d MIMAT0026147

UUCCACAGCUUUCUUGAACUU

>cme-miR395c MIMAT0026170

UUGAAGUGUUUGGGGGAACUC

>cme-miR397 MIMAT0026148

UCAUUGAGUGCAGCGUUGAUG

>cme-miR169g MIMAT0026172

AAGCCAAGGAUGAAUUGCCGG

>cme-miR169f MIMAT0026115

CAGCCAAGGAUGACUUGCCGG

>cme-miR390a MIMAT0026134

AAGCUCAGGAGGGAUAGCGCC

>cme-miR858 MIMAT0026083

UCUCGUUGUCUGUUCGACCUU

>cme-miR172d MIMAT0026127

GGAAUCUUGAUGAUGCUGCAU

>cme-miR394a MIMAT0026140

UUGGCAUUCUGUCCACCUCC

>cme-miR530b MIMAT0026104

UGCAUUUGCACCUACACCUU

>cme-miR169i MIMAT0026096

UAGCCAAAAAUGACUUGCCUGC

>cme-miR159a MIMAT0026160

UUUGGAUUGAAGGGAGCUCUA

>cme-miR396b MIMAT0026145

UUCCACAGCUUUCUUGAACUG

>cme-miR395f MIMAT0026144

CUGAAGUGUUUGGGGGAACUC

>cme-miR159b MIMAT0026168

AUUGGAUUGAAGGGAGCUCCU

>cme-miR398b MIMAT0026149

UGUGUUCUCAGGUCACCCCUU

>cme-miR398a MIMAT0026174

UGUGUUCUCAGGUCGCCCCUG

>cme-miR164c MIMAT0026086

UGGAGAAGCAGGGCACGUGCA

>cme-miR169h MIMAT0026114

CAGCCAAGGAUGACUUGCCGG

>cme-miR169e MIMAT0026117

UAGCCAAGGAUGACUUGCCUG

>cme-miR2111a MIMAT0026155

UAAUCUGCAUCCUGAGGUUUA

>cme-miR477b MIMAT0026102

CUCUCCCUCAAAGGCUUCUG

>cst-miR11334 MIMAT0044624

AAUUACUAUAAUAACACCUUCACAU

>cst-miR11332 MIMAT0044622

CUUGUGGGUAAUAGGCUUUCCUUCCUUG

>cst-miR11333 MIMAT0044623

ACUUCGCGGUAAGUCUAACCCUAAUU

>cst-miR11331 MIMAT0044621

CGUUGAGUAUUAUCUUUCAGUUC

>cst-miR11335 MIMAT0044625

GUUGGGAGGAUGGAGCGGUU

>hbr-miR6485 MIMAT0025295

UAGGAUGUAGAAGAGCAUAA

>hbr-miR396b MIMAT0025287

UUCCACAGCUUUCUUGAACUG

>hbr-miR166b MIMAT0025285

UCGGACCAGGCUUCAUUCCCCC

>hbr-miR159a MIMAT0025283

UUUGGAUUGAAGGGAGCUCUA

>hbr-miR6168 MIMAT0024786

UCUUGGUGCGGUGGACUGCUG

>hbr-miR2118 MIMAT0024784

GAAAUGGGUGGAUGGGAGUGA

>hbr-miR6175 MIMAT0024796

UGGUGGCUUUUAGGGCUCAAG

>hbr-miR9386 MIMAT0035235

UUUGCAGUUCGAAAGUGGAAGC

>hbr-miR6172 MIMAT0024792

UGGACCGUGUUUAAGUAAGAA

>hbr-miR482a MIMAT0024787

AGAUGGGUGGCUGGGCAAGAAG

>hbr-miR476 MIMAT0025290

UAAUCCUUCUUUGCAAAGUC

>hbr-miR9387 MIMAT0035236

GAACACAAUUAUAGGAAUC

>hbr-miR6174 MIMAT0024795

AUGCUAAGCUGUCAGACUUUG

>hbr-miR6483 MIMAT0025292

UAUUGUAGAAAUUUUCAGGAUC

>hbr-miR6482 MIMAT0025291

CAGGAACUGGUAUCAACCCAGC

>hbr-miR408b MIMAT0025289

ACUGGGAACAGGCAGAGCAUGA

>hbr-miR156 MIMAT0025282

UUGACAGAAGAUAGAGAGC

>hbr-miR396a MIMAT0024791

CACAGCUUUCUUGAACUUUCU

>hbr-miR408a MIMAT0025288

AAGACUGGGAACAGGCAGAGCA

>hbr-miR166a MIMAT0025284

UCGGACCAGGCUUCAUUCC

>hbr-miR482b MIMAT0035237

GAAUGGGCGGUUUGGGAAAGA

>hbr-miR6169 MIMAT0024788

UAGUAUUUCUCUUUUCUCUCU

>hbr-miR6173 MIMAT0024793

AGCCGUAAACGAUGGAUACU

>hbr-miR398 MIMAT0024794

GGAGCGACCUGAGAUCACAUG

>hbr-miR6170 MIMAT0024789

CAAGAAACAGAAGAGAGGGAU

>hbr-miR319 MIMAT0025286

UUGGACUGAAGGGAGCUCCCU

>hbr-miR6166 MIMAT0024783

AUUGGAUAACGGGAUUACAUGGU

>hbr-miR6171 MIMAT0024790

AUGUGGAUUGCUGAAGGCUUU

>hbr-miR6484 MIMAT0025293

UAAUGGGCUCUGCAUAGAUGG

>hbr-miR6167 MIMAT0024785

UACCCAGGUGGAAGCUUUGA

>mes-miR1446 MIMAT0029305

UUCUGAACUCUCUCCCUCAU

>mes-miR9386 MIMAT0045985

UUUGCAGUUCGAAAGUGGAAGC

>mes-miR166j MIMAT0029201

UCGGACCAGGCUUCAUUCC

>mes-miR11891 MIMAT0045987

CAUAAAUUGAACUAUAGACC

>mes-miR159b MIMAT0029176

UUUGGAUUGAAGGGAGCUCUA

>mes-miR171i MIMAT0029245

UGAUUGAGCCGUGCCAAUAUC

>mes-miR169j MIMAT0029218

GAGCCAAGAAUGACUUGCCGG

>mes-miR393c MIMAT0029264

UCCAAAGGGAUCGCAUUGAUCU

>mes-miR172d MIMAT0029250

AGAAUCUUGAUGAUGCUGCAU

>mes-miR395d MIMAT0029271

CUGAAGUGUUUGGGGGAACUC

>mes-miR482d MIMAT0045978

UUCCCGACACCACCCAUUCCAU

>mes-miR169ac MIMAT0029237

UAGCCAAGGAUGACUUGCCU

>mes-miR167g MIMAT0029207

UGAAGCUGCCAGCAUGAUCUU

>mes-miR399h MIMAT0045972

GGGCACCUCUCGCUUGGCAGG

>mes-miR396a MIMAT0029273

UUCCACAGCUUUCUUGAACUG

>mes-miR828a MIMAT0029303

UCUUGCUCAAAUGAGUAUUCCA

>mes-miR2111b MIMAT0029307

UAAUCUGCAUCCUGAGGUUUA

>mes-miR399d MIMAT0029282

UGCCAAAGGAGAUUUGCCCGG

>mes-miR164a MIMAT0029188

UGGAGAAGCAGGGCACGUGCA

>mes-miR160a MIMAT0029179

UGCCUGGCUCCCUGUAUGCCA

>mes-miR169t MIMAT0029228

UAGCCAAGGAUGACUUGCCCG

>mes-miR169b MIMAT0029210

CAGCCAAGGAUGACUUGCCGG

>mes-miR156a MIMAT0029165

UGACAGAAGAGAGUGAGCAC

>mes-miR1446b MIMAT0045982

UGAACUCUCCCCCUCAACGGCU

>mes-miR167b MIMAT0029202

UGAAGCUGCCAGCAUGAUCU

>mes-miR159a-5p MIMAT0037459

AGCUGCUGAGCUAUGGAUCCC

>mes-miR319b MIMAT0029254

UUGGACUGAAGGGAGCUCCCU

>mes-miR164c MIMAT0029190

UGGAGAAGCAGGGCACGUGCA

>mes-miR477g MIMAT0029296

AUCUCCCUCAAAGGCUUCCA

>mes-miR160e MIMAT0029183

UGCCUGGCUCCCUGAAUGCCAUC

>mes-miR482b MIMAT0045976

UUCCCAAUGUCGCCCAUUCCGA

>mes-miR169l MIMAT0029220

CAGCCAAGAAUGACUUGCCGG

>mes-miR166a MIMAT0029192

UCGGACCAGGCUUCAUUCCCC

>mes-miR167c MIMAT0029203

UGAAGCUGCCAGCAUGAUCUA

>mes-miR395e MIMAT0029272

CUGAAGGGUUUGGAGGAACUC

>mes-miR399b MIMAT0029280

UGCCAAAGGAGAUUUGCCCGG

>mes-miR167f MIMAT0029206

UGAAGCUGCCAGCAUGAUCUGA

>mes-miR160g MIMAT0029185

UGCCUGGCUCCCUGUAUGCCAUC

>mes-miR169p MIMAT0029224

UAGCCAAGGAUGACUUGCCUG

>mes-miR396b MIMAT0029274

UUCCACAGCUUUCUUGAACUG

>mes-miR171f MIMAT0029242

AGAUUGAGCCGCGCCAAUAUC

>mes-miR166c MIMAT0029194

UCGGACCAGGCUUCAUUCCCC

>mes-miR319c MIMAT0029255

UUGGACUGAAGGGAGCUCCCU

>mes-miR156j MIMAT0029174

UUGACAGAAGAUAGAGAGCAC

>mes-miR160c MIMAT0029181

UGCCUGGCUCCCUGUAUGCCG

>mes-miR2950 MIMAT0029309

UUCCAUCUCUUGCACACUGGA

>mes-miR169h MIMAT0029216

UGAGCCAAGGAUGACUUGCCG

>mes-miR169q MIMAT0029225

UAGCCAAGGAUGACUUGCCUG

>mes-miR2275 MIMAT0029308

UUUGGUUUCCUCCAAUAUCUUA

>mes-miR171a MIMAT0024415

UUGAGCCGCGUCAAUAUCUCC

>mes-miR477i MIMAT0029289

ACUCUCCCUCAAGGGCUUCCG

>mes-miR169k MIMAT0029219

GAGCCAAGAAUGACUUGCCGG

>mes-miR535b MIMAT0029301

UGACAACGAGAGAGAGCACGG

>mes-miR172a MIMAT0024416

AGAAUCUUGAUGAUGCUGCAU

>mes-miR477b MIMAT0029291

CUCUCCCUCAAGGGCUUCUG

>mes-miR319h MIMAT0029260

CUUGGACUGAAGGGAGCUCCU

>mes-miR399e MIMAT0029283

UGCCAAAGGAGAUUUGCUCGG

>mes-miR156f MIMAT0029170

UGACAGAAGAGAGUGAGCAC

>mes-miR169z MIMAT0029234

UAGCCAAGGAUGACUUGCCCG

>mes-miR169v MIMAT0029230

UAGCCAAGGAUGACUUGCCCG

>mes-miR164b MIMAT0029189

UGGAGAAGCAGGGCACGUGCA

>mes-miR535d MIMAT0045981

UUGACGACGAGAGAGAGCACG

>mes-miR393a MIMAT0029262

UCCAAAGGGAUCGCAUUGAUC

>mes-miR156c MIMAT0029167

UGACAGAAGAGAGUGAGCAC

>mes-miR166g MIMAT0029198

UCGGACCAGGCUUCAUUCCCC

>mes-miR167h MIMAT0029208

UGAAGCUGCCAGCAUGAUCUU

>mes-miR319f MIMAT0029258

AUUGGACUGAAGGGAGCUCC

>mes-miR169f MIMAT0029214

UAGCCAAGGAUGACUUGCCGG

>mes-miR827 MIMAT0029302

UUAGAUGACCAUCAACAAACA

>mes-miR530a MIMAT0029298

UGCAUUUGCACCUGCACCUU

>mes-miR395b MIMAT0029269

CUGAAGUGUUUGGGGGAACUC

>mes-miR477k MIMAT0045974

ACUCUCCCUCAAGAGCUUCUC

>mes-miR398 MIMAT0045971

UGUGUUCUCAGGUCGCCCCUG

>mes-miR397b MIMAT0045970

UCAUUGAGUGCAGCGUUGAUG

>mes-miR169d MIMAT0029212

CAGCCAAGGAUGACUUGCCGG

>mes-miR390 MIMAT0029261

CGCUAUCCAUCCUGAGUUUC

>mes-miR169o MIMAT0029223

UAGCCAAGGAUGACUUGCCUG

>mes-miR394b MIMAT0029266

UUGGCAUUCUGUCCACCUCC

>mes-miR172b MIMAT0029248

AGAAUCUUGAUGAUGCUGCAU

>mes-miR482 MIMAT0029297

UCUUCCCUACUCCACCCAUUCC

>mes-miR399c MIMAT0029281

UGCCAAAGGAGAUUUGCCCGG

>mes-miR168a MIMAT0024414

UCGCUUGGUGCAGGUCGGGAA

>mes-miR159d MIMAT0029178

AUUGGAGUGAAGGGAGCUCUG

>mes-miR396f MIMAT0029278

UUCCACAGCUUUCUUGAACUU

>mes-miR166e MIMAT0029196

UCGGACCAGGCUUCAUUCCCC

>mes-miR166i MIMAT0029200

UUGGACCAGGCUUCAUUCCCC

>mes-miR169ad MIMAT0045986

UCACAGGCUCUUAUUUUUCAUG

>mes-miR3627 MIMAT0045983

UUGUCGCAGGAGCGGUGGCACC

>mes-miR397 MIMAT0029279

UUUGAGUGCAGCGUUGAUGA

>mes-miR159c MIMAT0029177

AUUGGAGUGAAGGGAGCUCUG

>mes-miR164d MIMAT0029191

UGGAGAAGCAGGGCACAUGCU

>mes-miR167d MIMAT0029204

UGAAGCUGCCAGCAUGAUCUGA

>mes-miR171e MIMAT0029241

UUGAGCCGCGCCAAUAUCACU

>mes-miR393d MIMAT0029265

UCCAAAGGGAUCGCAUUGAUCU

>mes-miR477a MIMAT0029290

CUCUCCCUCAAGGGCUUCUG

>mes-miR169g MIMAT0029215

CAGCCAAGGAUGACUUGCCGA

>mes-miR172c MIMAT0029249

UGAAUCUUGAUGAUGCUACGC

>mes-miR403b MIMAT0029287

UUAGAUUCACGCACAAACUCG

>mes-miR160b MIMAT0029180

UGCCUGGCUCCCUGUAUGCCA

>mes-miR166d MIMAT0029195

UCGGACCAGGCUUCAUUCCCC

>mes-miR171l MIMAT0045968

CGAGCCGAACCAAUAUCACUC

>mes-miR530b MIMAT0029299

UGCAUUUGCACCUGCACCUUA

>mes-miR169x MIMAT0029232

UAGCCAAGGAUGACUUGCCCG

>mes-miR156i MIMAT0029173

UUGACAGAAGAUAGAGAGCAC

>mes-miR172e MIMAT0029251

GGAAUCUUGAUGAUGCUGCAG

>mes-miR169ab MIMAT0029236

UAGCCAAGGAUGACUUGCCUA

>mes-miR319a MIMAT0029253

UUGGACUGAAGGGAGCUCCCU

>mes-miR408 MIMAT0024419

AUGCACUGCCUCUUCCCUGGC

>mes-miR2118 MIMAT0045975

GUUCCCAUGCCACCCAUUUCUA

>mes-miR156e MIMAT0029169

UGACAGAAGAGAGUGAGCAC

>mes-miR167a MIMAT0024413

UGAAGCUGCCAGCAUGAUCUG

>mes-miR11892 MIMAT0045988

UUGUCAUCUCAACCUUGUGUC

>mes-miR2111a MIMAT0029306

UAAUCUGCAUCCUGAGGUUUA

>mes-miR399a MIMAT0024418

UGCCAAAGGAGAAUUGCCCUG

>mes-miR169e MIMAT0029213

CAGCCAAGGAUGACUUGCCGG

>mes-miR396e MIMAT0029277

UUCCACAGCUUUCUUGAACUU

>mes-miR828b MIMAT0029304

UCUUGCUCAAAUGAGUAUUCCA

>mes-miR171g MIMAT0029243

UGAUUGAGCCGUGCCAAUAUC

>mes-miR477d MIMAT0029293

CUCUCCCUCAAGGGCUUCUC

>mes-miR160f MIMAT0029184

UGCCUGGCUCCCUGAAUGCCA

>mes-miR166h MIMAT0029199

UCGGACCAGGCUUCAUUCCCGU

>mes-miR482c MIMAT0045977

UUUUCCCAAGACCUCCCAUACC

>mes-miR169i MIMAT0029217

GAGCCAAGAAUGACUUGCCGG

>mes-miR169n MIMAT0029222

GAGCCAAGAAUGACUUGCCGA

>mes-miR396c MIMAT0029275

UUCCACAGCUUUCUUGAACUU

>mes-miR535c MIMAT0045980

UUGACGACGAGAGAGAGCACA

>mes-miR156k MIMAT0029175

UUGACAGAAGAGAGAGAGCAC

>mes-miR169a MIMAT0029209

CAGCCAAGGAUGACUUGCCGG

>mes-miR171h MIMAT0029244

UGAUUGAGCCGUGCCAAUAUC

>mes-miR171k MIMAT0029247

UGAUUGAGCCGUGCCAAUAUC

>mes-miR319d MIMAT0029256

UUGGACUGAAGGGAGCUCCCU

>mes-miR160d MIMAT0029182

UGCCUGGCUCCCUGUAUGCCA

>mes-miR399g MIMAT0029285

UGCCAAAGGAGAUUUGCCCGG

>mes-miR159a-3p MIMAT0024410

UUUGGAUUGAAGGGAGCUCUA

>mes-miR169u MIMAT0029229

UAGCCAAGGAUGACUUGCCUG

>mes-miR160h MIMAT0029186

UGCCUGGCUCCCUGUAUGCCAUU

>mes-miR169c MIMAT0029211

CAGCCAAGGAUGACUUGCCGG

>mes-miR6445 MIMAT0045984

UUCAUUCCUCUUCCUAAAAUGG

>mes-miR477f MIMAT0029295

AUCUCCCUCAAAGGCUUCCA

>mes-miR172f MIMAT0029252

GGAAUCUUGAUGAUGCUGCAG

>mes-miR166b MIMAT0029193

UCGGACCAGGCUUCAUUCCCC

>mes-miR169m MIMAT0029221

CAGCCAAGAAUGACUUGCCGG

>mes-miR319e MIMAT0029257

UUGGACUGAAGGGAGCUCCCU

>mes-miR169w MIMAT0029231

UAGCCAAGGAUGACUUGCCUG

>mes-miR156b MIMAT0029166

UGACAGAAGAGAGUGAGCAC

>mes-miR171d MIMAT0029240

AUGAGCCGUGCCAAUAUCACG

>mes-miR167e MIMAT0029205

UGAAGCUGCCAGCAUGAUCUGA

>mes-miR171b MIMAT0029238

UUGAGCCGUGCCAAUAUCACG

>mes-miR403a MIMAT0029286

UUAGAUUCACGCACAAACUCG

>mes-miR393b MIMAT0029263

UCCAAAGGGAUCGCAUUGAUCC

>mes-miR477e MIMAT0029294

CUCUCCCUCAAGGGCUUCUG

>mes-miR394a MIMAT0024417

UUGGCAUUCUGUCCACCUCC

>mes-miR396d MIMAT0029276

UUCCACAGCUUUCUUGAACUU

>mes-miR169s MIMAT0029227

UAGCCAAGGAUGACUUGCCUG

>mes-miR399f MIMAT0029284

UGCCAAAGGAGAGUUGCCCUG

>mes-miR319g MIMAT0029259

UUGGACUGAAGGGAGCUCCUU

>mes-miR156g MIMAT0029171

UGACAGAAGAGAGUGAGCAC

>mes-miR171c MIMAT0029239

UUGAGCCGUGCCAAUAUCACG

>mes-miR394c MIMAT0029267

UUGGCAUUCUGUCCACCUCCAU

>mes-miR395c MIMAT0029270

CUGAAGUGUUUGGGGGAACUC

>mes-miR477j MIMAT0045973

ACUCUCCCUAAAGGCUUCAAC

>mes-miR169r MIMAT0029226

UAGCCAAGGAUGACUUGCCCG

>mes-miR156d MIMAT0029168

UGACAGAAGAGAGUGAGCAC

>mes-miR162 MIMAT0029187

UCGAUAAACCUCUGCAUCCAG

>mes-miR166f MIMAT0029197

UCGGACCAGGCUUCAUUCCCC

>mes-miR477c MIMAT0029292

CUCUCCCUCAAGGGCUUCUG

>mes-miR169y MIMAT0029233

UAGCCAAGGAUGACUUGCCUG

>mes-miR482e MIMAT0045979

UCUUACCUACACCGCCCAUGCC

>mes-miR156h MIMAT0029172

UUGACAGAAGAUAGAGAGCAC

>mes-miR535a MIMAT0029300

UGACAACGAGAGAGAGCACGU

>mes-miR169aa MIMAT0029235

UAGCCAAGGAUGACUUGCCUG

>mes-miR477h MIMAT0029288

ACUCUCCCUCAAGGGCUUCAG

>mes-miR390b MIMAT0045969

AAGCUCAGGAGGGAUAGCGCC

>mes-miR395a MIMAT0029268

CUGAAGUGUUUGGGGGAACUC

>mes-miR171j MIMAT0029246

UGAUUGAGCCGUGCCAAUAUC

>rco-miR166d MIMAT0014163

UCGGACCAGGCUUCAUUCCCC

>rco-miR164a MIMAT0014156

UGGAGAAGCAGGGCACGUGCA

>rco-miR166a MIMAT0014160

UCGGACCAGGCUUCAUUCCCC

>rco-miR171c MIMAT0014174

UGAUUGAGCCGUGCCAAUAUC

>rco-miR169c MIMAT0014171

UGAGCCAAGGAUGACUUGCCG

>rco-miR160c MIMAT0014206

UGCCUGGCUCCCUGAAUGCCA

>rco-miR319c MIMAT0014182

UUGGACUGAAGGGAGCUCCCU

>rco-miR156c MIMAT0014146

UGACAGAAGAGAGUGAGCACA

>rco-miR399b MIMAT0014197

UGCCAAAGGAGAUUUGCCCGG

>rco-miR390a MIMAT0014184

AAGCUCAGGAGGGAUAGCGCC

>rco-miR319d MIMAT0014183

UUGGACUGAAGGGAGCUCCUU

>rco-miR171b MIMAT0014173

UUGAGCCGUGCCAAUAUCACG

>rco-miR167c MIMAT0014167

UGAAGCUGCCAGCAUGAUCUGG

>rco-miR164c MIMAT0014158

UGGAGAAGCAGGGCACGUGCA

>rco-miR398a MIMAT0014194

UGUGUUCUCAGGUCACCCCUU

>rco-miR166c MIMAT0014162

UCGGACCAGGCUUCAUUCCCC

>rco-miR166b MIMAT0014161

UCGGACCAGGCUUCAUUCCCC

>rco-miR171a MIMAT0014172

UUGAGCCGUGCCAAUAUCACG

>rco-miR160a MIMAT0014153

UGCCUGGCUCCCUGUAUGCCA

>rco-miR156b MIMAT0014145

UGACAGAAGAGAGUGAGCACA

>rco-miR167a MIMAT0014165

UGAAGCUGCCAGCAUGAUCUA

>rco-miR395e MIMAT0014191

CUGAAGUGUUUGGGGGAACUC

>rco-miR156g MIMAT0014150

UUGACAGAAGAUAGAGAGCAC

>rco-miR171g MIMAT0014178

AGAUUGAGCCGCGCCAAUAUC

>rco-miR156h MIMAT0014151

UUGACAGAAGAUAGAGAGCAC

>rco-miR395d MIMAT0014190

CUGAAGUGUUUGGGGGAACUC

>rco-miR398b MIMAT0014195

UGUGUUCUCAGGUCGCCCCUG

>rco-miR169b MIMAT0014170

CAGCCAAGGAUGACUUGCCGG

>rco-miR156e MIMAT0014148

UGACAGAAGAGAGAGAGCACA

>rco-miR535 MIMAT0014205

UGACAACGAGAGAGAGCACGC

>rco-miR172 MIMAT0014179

GGAAUCUUGAUGAUGCUGCAG

>rco-miR169a MIMAT0014169

CAGCCAAGGAUGACUUGCCGG

>rco-miR156d MIMAT0014147

UGACAGAAGAGAGUGAGCACA

>rco-miR162 MIMAT0014155

UCGAUAAACCUCUGCAUCCAG

>rco-miR395b MIMAT0014188

CUGAAGUGUUUGGGGGAACUC

>rco-miR160b MIMAT0014154

UGCCUGGCUCCCUGUAUGCCA

>rco-miR167b MIMAT0014166

UGAAGCUGCCAGCAUGAUCUA

>rco-miR395c MIMAT0014189

CUGAAGUGUUUGGGGGAACUC

>rco-miR156f MIMAT0014149

UUGACAGAAGAUAGAGAGCAC

>rco-miR408 MIMAT0014204

CUGCACUGCCUCUUCCCUGGC

>rco-miR399f MIMAT0014201

UGCCAAAGGAGAUUUGCUCAC

>rco-miR399d MIMAT0014199

UGCCAAAGGAGAGCUGCCCUG

>rco-miR399a MIMAT0014196

UGCCAAAGGAGAGUUGCCCUG

>rco-miR171d MIMAT0014175

UGAUUGAGCCGUGCCAAUAUC

>rco-miR159 MIMAT0014152

UUUGGAUUGAAGGGAGCUCUA

>rco-miR403a MIMAT0014202

UUAGAUUCACGCACAAACUCG

>rco-miR390b MIMAT0014185

AAGCUCAGGAGGGAUAGCGCC

>rco-miR171e MIMAT0014176

UGAUUGAGCCGUGCCAAUAUC

>rco-miR164d MIMAT0014159

UGGAGAAGCAGGGCACAUGCU

>rco-miR168 MIMAT0014168

UCGCUUGGUGCAGGUCGGGAA

>rco-miR395a MIMAT0014187

CUGAAGUGUUUGGGGGAACUC

>rco-miR166e MIMAT0014164

UCGGACCAGGCUUCAUUCCCC

>rco-miR164b MIMAT0014157

UGGAGAAGCAGGGCACGUGCA

>rco-miR399e MIMAT0014200

UGCCAAAGGAGAUUUGCCCAG

>rco-miR171f MIMAT0014177

UGAUUGAGCCGUGCCAAUAUC

>rco-miR319a MIMAT0014180

UUGGACUGAAGGGAGCUCCCU

>rco-miR156a MIMAT0014144

UGACAGAAGAGAGUGAGCACA

>rco-miR399c MIMAT0014198

UGCCAAAGGAGAUUUGCCCGG

>rco-miR319b MIMAT0014181

UUGGACUGAAGGGAGCUCCCU

>rco-miR397 MIMAT0014193

UCAUUGAGUGCAGCGUUGAUG

>rco-miR393 MIMAT0014186

UCCAAAGGGAUCGCAUUGAUC

>rco-miR403b MIMAT0014203

UUAGAUUCACGCACAAACUCG

>rco-miR396 MIMAT0014192

UUCCACAGCUUUCUUGAACUU

>aau-miR172 MIMAT0022673

UGAGAAUCUUGAUGAUGCUGCAU

>aau-miR160 MIMAT0022672

UGGCAUACAGGGAGCCAGGCA

>aau-miR168 MIMAT0022678

AUUCAGUUGAUGCAAGGCGGGAUC

>aau-miR2086 MIMAT0022676

GACAUGAAUGCAGAACUGGAA

>aau-miR162 MIMAT0022677

UCGAUAAACCUCUGCAUCCAG

>aau-miR319 MIMAT0022675

UUGGACUGAAGGGAGCUCCCU

>aau-miR396 MIMAT0022674

UUCCACAGCUUUCUUGAACUG

>ahy-miR408-3p MIMAT0016329

AUGCACUGCCUCUUCCCUGGC

>ahy-miR3511-3p MIMAT0016335

UGUUACUAUGGCAUCUGGUAA

>ahy-miR408-5p MIMAT0016328

CUGGGAACAGGCAGAGCAUGA

>ahy-miR3517 MIMAT0016343

CUGACCACUGUGAUCCCGGAA

>ahy-miR3511-5p MIMAT0016334

GCCAGGGCCAUGAAUGCAGA

>ahy-miR3510 MIMAT0016333

UUAUACCAUCUUGCGAGACUGA

>ahy-miR3518 MIMAT0016344

UGACCUUUGGGGAUAUUCGUG

>ahy-miR3519 MIMAT0016345

UCAAUCAAUGACAGCAUUUCA

>ahy-miR3513-5p MIMAT0016337

UUAAUUUCUGAGUUUGUCAUC

>ahy-miR3520-3p MIMAT0016347

AAGGGAGACGUUUGAAUUAUC

>ahy-miR3514-5p MIMAT0016339

AGGAUUCUGUAUUAACGGUGGA

>ahy-miR3513-3p MIMAT0016338

UUGAUAAGAUAGAAAUUGUAU

>ahy-miR167-5p MIMAT0016324

UGAAGCUGCCAGCAUGAUCUU

>ahy-miR156a MIMAT0016317

UGACAGAAGAGAGAGAGCAC

>ahy-miR167-3p MIMAT0016325

AGAUCAUGUGGCAGUUUCACC

>ahy-miR156b-3p MIMAT0016319

GCUCUCUAAGCUUCUGUCAUC

>ahy-miR159 MIMAT0016321

UUUGGAUUGAAGGGAGCUCUA

>ahy-miR3512 MIMAT0016336

CGCAAAUGAUGACAAAUAGA

>ahy-miR3514-3p MIMAT0016340

UCACCGUUAAUACAGAAUCCUU

>ahy-miR3515 MIMAT0016341

AAUGUAGAAAAUGAACGGUAU

>ahy-miR3509-3p MIMAT0016332

AUCUAACGACUCUCAGAUAUCA

>ahy-miR156c MIMAT0016320

UUGACAGAAGAGAGAGAGCAC

>ahy-miR160-5p MIMAT0016322

UGCCUGGCUCCCUGAAUGCCA

>ahy-miR3508 MIMAT0016330

UAGAGGGUCCCCAUGUUCUCA

>ahy-miR394 MIMAT0016326

UUGGCAUUCUGUCCACCUCC

>ahy-miR3520-5p MIMAT0016346

AGGUGAUGGUGAAUAUCUUAUC

>ahy-miR3509-5p MIMAT0016331

AUACUUGAGAGCCGUUAGAUGA

>ahy-miR3516 MIMAT0016342

GCUGGGUGAUAUUGACAGAAG

>ahy-miR398 MIMAT0016327

UGUGUUCUCAGGUCACCCCU

>ahy-miR3521 MIMAT0016348

UGGUGAGUCGUAUACAUACUG

>ahy-miR160-3p MIMAT0016323

GCAUGAAGGGAGUCACGCAGG

>ahy-miR156b-5p MIMAT0016318

UUGACAGAAGAUAGAGAGCAC

>amg-miR396 MIMAT0022680

UUCCACAGCUUUCUUGAACUG

>amg-miR319 MIMAT0022679

UUGGACUGAAGGGAGCUCCCU

>amg-miR2086 MIMAT0022681

GACAUGAAUGCAGAACUGGAA

>gma-miR10447 MIMAT0041727

UGUUUCCAUGUUGUUGAGUGAC

>gma-miR9732 MIMAT0036351

CAAGGGUAUGAUGUGCAAUCU

>gma-miR5038a MIMAT0021063

UGAGAAUUUGGCCUCUGUCCA

>gma-miR5672 MIMAT0022454

CAUGGUAGUGGAAGAAAUGGA

>gma-miR10406b MIMAT0041685

CAGUCGAGUUAUUGUAUAAUUCAC

>gma-miR10186d MIMAT0041653

UUUUGGGAAUUAAAAUGUAAACUU

>gma-miR5030b MIMAT0040957

UUCCGGAAGAACAAAGCUACC

>gma-miR393c-5p MIMAT0024910

UCCAAAGGGAUCGCAUUGAUCC

>gma-miR10419 MIMAT0041665

CAAAAAACACAUAUGAAACUG

>gma-miR1520f-3p MIMAT0018244

CAAUCAGAACAUGACACAUGACAA

>gma-miR9739 MIMAT0036359

UUUGAAUGUCCAGAUACGUAC

>gma-miR319p MIMAT0036379

UUUUGGACUGAAGGGAGCUCC

>gma-miR156v MIMAT0024881

UGACAGAAGAGAGUGAGCAC

>gma-miR862b MIMAT0021604

GCUGGAUGUCUUUGAAGGA

>gma-miR1520l MIMAT0018292

AAUCAGAACAUGACACGUGAUAGU

>gma-miR169a MIMAT0001693

CAGCCAAGGAUGACUUGCCGG

>gma-miR2606a MIMAT0023226

AAAAGCACUUAAGGAACGGUA

>gma-miR403b MIMAT0021614

UUAGAUUCACGCACAAACUUG

>gma-miR5371-5p MIMAT0021608

UAGGAAUUAGUCACUCAGAUC

>gma-miR482b-5p MIMAT0018286

UAUGGGGGGAUUGGGAAGGAAU

>gma-miR169w MIMAT0041676

CAAGGAUGACUUGCCGGCAUU

>gma-miR10430 MIMAT0041695

UCGGUCUGACCUUUUUAAAAGCCU

>gma-miR171i-3p MIMAT0021057

UUGAGCCGUGCCAAUAUCACG

>gma-miR1532 MIMAT0007394

AACACGCUAAGCGAGAGGAGCUC

>gma-miR408c-5p MIMAT0022993

CAGGGGAACAGGCAGAGCAUG

>gma-miR4994-3p MIMAT0032119

UGAUAUCCUUGAGCUAAUACA

>gma-miR391-5p MIMAT0018280

UACGCAGGAGAGAUGACGCUGU

>gma-miR1524 MIMAT0007385

CGAGUCCGAGGAAGGAACUCC

>gma-miR10424d MIMAT0041722

UUUUCUAAUUUAUUAGGGACU

>gma-miR156m MIMAT0021637

UUGACAGAAGAUAGAGAGCAC

>gma-miR408d MIMAT0020998

UGCACUGCCUCUUCCCUGGC

>gma-miR9760 MIMAT0036395

UGGAUGAUGUAGUUUUGAUUG

>gma-miR4358 MIMAT0018250

CAGUGCAUGACUAUAUCGCCAG

>gma-miR160f MIMAT0024888

UGCCUGGCUCCCUGUAUGCCA

>gma-miR4398 MIMAT0018314

UGUCAGCGGAGUGAGAAGACGAAA

>gma-miR393b MIMAT0023170

UUUGGGAUCAUGCUAUCCCUU

>gma-miR166d MIMAT0020979

UCGGACCAGGCUUCAUUCCCC

>gma-miR1520h MIMAT0018255

AACGUCCAAUCAGAACGUGACAUG

>gma-miR171b-5p MIMAT0017336

ACGGCGUGAUAUUGGUACGGCUC

>gma-miR395e MIMAT0023244

UGAAGUGUUUGGGGGAACUUU

>gma-miR4353 MIMAT0018242

CAAGUCGUAGCCGGUGUUAUUACU

>gma-miR4996 MIMAT0021014

UAGAAGCUCCCCAUGUUCUC

>gma-miR10405d MIMAT0041704

UUGUUUCUUAUAAAAAGGACC

>gma-miR169l-5p MIMAT0021652

CAGCCAAGAAUGACUUGCCGG

>gma-miR4379 MIMAT0018285

UAGAGUGUAUACUGUGAGAGGCCU

>gma-miR10426 MIMAT0041682

UAGAAAAGAAUCAAUGUAGAAAGU

>gma-miR169g MIMAT0020987

CAGCCAAGGAUGACUUGCCGG

>gma-miR10441 MIMAT0041714

CAACCCUGAGAACAAUGAAAUCGU

>gma-miR399m MIMAT0041681

GGGCUCCUCUCUCCUGGCAUG

>gma-miR2118b-5p MIMAT0022982

GGAGAUGGGAGGGUCGGUAAAG

>gma-miR156b MIMAT0001692

UGACAGAAGAGAGAGAGCACA

>gma-miR394a-5p MIMAT0022974

UUGGCAUUCUGUCCACCUCC

>gma-miR1513c MIMAT0021674

UAUGAGAGAAAGCCAUGAC

>gma-miR2108a MIMAT0010084

UUAAUGUGUUGUGUUUGUCGG

>gma-miR393i MIMAT0024916

UUCCAAAGGGAUCGCAUUGAUC

>gma-miR166t MIMAT0024902

UCGGACCAGGCUUCAUUCCC

>gma-miR4366 MIMAT0018265

CUACUUAGUAGAGAUUUGUUGG

>gma-miR1512a-5p MIMAT0007370

UAACUGAAAAUUCUUAAAGUAU

>gma-miR5679 MIMAT0022467

UUGGUGACCCAGAAGAAGUUGA

>gma-miR4389 MIMAT0018302

UCGGUCGGACCGAUCCAAUCGGAA

>gma-miR159d MIMAT0018340

AGCUGCUUAGCUAUGGAUCCC

>gma-miR9748 MIMAT0036375

GAAGGAAGUGUAGAGGGAUGAC

>gma-miR5784 MIMAT0023190

AAUUAGCUAAUGGUUAGCUAA

>gma-miR4411 MIMAT0018331

UUAUUGUAACUAAUUUGUCGGU

>gma-miR4401a MIMAT0018319

ACAACGUCUUUGAAAGUAGGCAUU

>gma-miR10438 MIMAT0041710

UUGGACUAAGGUUUUUUGGCAC

>gma-miR4997 MIMAT0021015

GAUCGUCAAGCGCGAAGAUGAGG

>gma-miR4412-3p MIMAT0018337

AGUGGCGUAGAUCCCCACAAC

>gma-miR4404 MIMAT0018323

AUUCGUGGAAGACUGGCGGAUCAA

>gma-miR160a-5p MIMAT0001676

UGCCUGGCUCCCUGUAUGCCA

>gma-miR530c MIMAT0023197

UGCAUUUGCACCUGCACUUUA

>gma-miR171i-5p MIMAT0021056

AUAAGAAAGCAAUGCUCAAA

>gma-miR4415b-3p MIMAT0021676

UUGAUUCUCAUCACAACAUGG

>gma-miR1521a MIMAT0007382

CUGUUAAUGGAAAAUGUUGA

>gma-miR5035-3p MIMAT0032121

UGUUUAGAAGCUCAUAGAAUAGAU

>gma-miR4390 MIMAT0018303

UCGUACUCGUCGGGUAUCGGGUAU

>gma-miR396k-5p MIMAT0032132

UUCCACAGCUUUCUUGAACUU

>gma-miR164a MIMAT0007354

UGGAGAAGCAGGGCACGUGCA

>gma-miR9723 MIMAT0036338

CAAAGGAGAUUUGGACAACUC

>gma-miR10405e MIMAT0041657

UUGUUUCUUAUAAAAAGGACC

>gma-miR5043 MIMAT0021074

UGUCCCCUUCUCUGCACCACC

>gma-miR10422 MIMAT0041669

UUCUGAUUAGAGAGCAACACC

>gma-miR1516d MIMAT0024927

GUACUUGUGGCUUGUAUCCAA

>gma-miR166m MIMAT0023247

CGGACCAGGCUUCAUUCCCC

>gma-miR1508c MIMAT0021007

UAGAAAGGGAAAUAGCAGUUG

>gma-miR4344 MIMAT0018232

AAGUAGACAUUCUAAGACGUUGCU

>gma-miR172b-5p MIMAT0020921

GUAGCAUCAUCAAGAUUCAC

>gma-miR319i MIMAT0021663

UUGGACUGAAGGGGAGCUCCUUC

>gma-miR3522 MIMAT0021633

AGACCAAAUGAGCAGCUGA

>gma-miR1520m MIMAT0018293

AAUCAGAACAUGACAUGUGACAAU

>gma-miR4363 MIMAT0018259

CGAUUACCAGAAGGCUUAUUAG

>gma-miR160a-3p MIMAT0022889

GCGUAUGAGGAGCCAAGCAUA

>gma-miR396a-5p MIMAT0001687

UUCCACAGCUUUCUUGAACUG

>gma-miR399e MIMAT0023232

UGCCAAAGGAGAUUUGCCCAG

>gma-miR2111a MIMAT0022457

GUCCUUGGGAUGCAGAUUACG

>gma-miR4383 MIMAT0018290

UAUUGGAUCUCAGUUGAACCGGUC

>gma-miR4370 MIMAT0018271

AGUAGACUCGUCCGAUUUUGCGUA

>gma-miR10186l MIMAT0041720

UUUUGGGAAUUAAAAUGUAAACUU

>gma-miR166n MIMAT0024896

UCGGACCAGGCUUCAUUCCCC

>gma-miR393d MIMAT0024911

UCCAAAGGGAUCGCAUUGAUCC

>gma-miR9752 MIMAT0036381

UGCUUCUUCUUUUCCCUGUUU

>gma-miR5773 MIMAT0023175

UUUUUAAAAGGUUCAGUUAGGU

>gma-miR5671a MIMAT0022453

CAUGGAAGUGAAUCGGGUGAC

>gma-miR4357 MIMAT0018249

CAGUCGUGUGAUUGUACGGUUCAU

>gma-miR167a MIMAT0001679

UGAAGCUGCCAGCAUGAUCUA

>gma-miR5770a MIMAT0023167

UUAGGACUAUGGUUUGGACGA

>gma-miR10186c MIMAT0041645

UUUUGGGAAUUAAAAUGUAAACUU

>gma-miR2606b MIMAT0023250

AAAAGCACUUAAGGAACGGUA

>gma-miR159e-3p MIMAT0021641

UUUGGAUUGAAGGGAGCUCUA

>gma-miR319o MIMAT0036377

UGGACUGAAGGGGAGCUCCUUC

>gma-miR319b MIMAT0001685

UUGGACUGAAGGGAGCUCCC

>gma-miR5764 MIMAT0023161

UCCAUUCGCGGACAUGAUGGAU

>gma-miR159f-5p MIMAT0021642

GAGUUCCCUGCACUCCAAGUC

>gma-miR10409 MIMAT0041646

CCUCUGAAGAUAUUAAAAGCCU

>gma-miR398a MIMAT0001689

UGUGUUCUCAGGUCACCCCUU

>gma-miR4414a-3p MIMAT0032112

UCCAACGAUGCGGGAGCUGC

>gma-miR167g MIMAT0018307

UGAAGCUGCCAGCAUGAUCUGA

>gma-miR166h-3p MIMAT0020984

UCUCGGACCAGGCUUCAUUCC

>gma-miR1520c MIMAT0007395

UUCAAUAAGAACGUGACACGUGA

>gma-miR4414b MIMAT0041688

AGCUGCUGACUCGUUGGUUCG

>gma-miR10418 MIMAT0041661

GAAGUAAUCCUAGGAACUCCCU

>gma-miR5044 MIMAT0021075

GUAGUGGAUGCCUAGAGGUCCA

>gma-miR9745 MIMAT0036365

AGAAUUAAAUUUGGACCGUAUAAC

>gma-miR166s MIMAT0024901

UCGGACCAGGCUUCAUUCCC

>gma-miR408a-5p MIMAT0022992

CAGGGGAACAGGCAGAGCAUG

>gma-miR171c-5p MIMAT0018334

AGAUAUUGGUGCGGUUCAAUC

>gma-miR9751 MIMAT0036380

GUAAUUUUAAACCUAAACCCUAAA

>gma-miR395f MIMAT0023245

UGAAGUGUUUGGGGGAACUUU

>gma-miR9740 MIMAT0036360

UGUAGGUUCCAGUGAGGGAAA

>gma-miR5780b MIMAT0036341

UCUGAGUCCAUGAUAUAUUAAA

>gma-miR1516c MIMAT0023178

AAUGUCUGGGCUUAGCGAGGCGGU

>gma-miR4415b-5p MIMAT0022997

AAGUUGUGAUGGGAAUCAAUGGCA

>gma-miR171r MIMAT0023229

CGAGCCGAAUCAAUACCACUC

>gma-miR4377 MIMAT0018281

UACGUCAUCGCUGAAUGGAAGACG

>gma-miR10406a MIMAT0041641

CAGUCGAGUUAUUGUAUAAUUCAC

>gma-miR4410 MIMAT0018329

UAUGUUGAUCCGUAUGAGUCGUAC

>gma-miR166h-5p MIMAT0020983

GGAAUGUUGUUUGGCUCGAGG

>gma-miR9738 MIMAT0036358

UGAAACAUGAUGUGGACUCUUC

>gma-miR169v MIMAT0024905

CAGCCAAGGAUGACUUGCC

>gma-miR156n MIMAT0021638

UUGACAGAAGAGAGUGAGCAC

>gma-miR169i-3p MIMAT0021068

CCGGUGCCAUCCCGUCUCAUA

>gma-miR4374c MIMAT0041680

CAACACCGUCUUUGAAGCCUGG

>gma-miR10405c MIMAT0041674

UUGUUUCUUAUAAAAAGGACC

>gma-miR169l-3p MIMAT0022994

CGGGCAAGUUGUUUUUGGCUAC

>gma-miR159c MIMAT0007352

AUUGGAGUGAAGGGAGCUCCG

>gma-miR10424b MIMAT0041689

UUUUCUAAUUUAUUAGGGACU

>gma-miR1525 MIMAT0007386

UGGGUUAAUUAAGUUUUUAGU

>gma-miR160e MIMAT0020973

UGCCUGGCUCCCUGUAUGCC

>gma-miR159f-3p MIMAT0021643

AUUGGAGUGAAGGGAGCUCCA

>gma-miR4368b MIMAT0018269

AAGGACGGUACUUACGUAAGCAAC

>gma-miR4365 MIMAT0018264

AAGAACUUCUUCCGCGAGAUCGCA

>gma-miR10186a MIMAT0040951

UUGGGAAUUAAAAUGUAAACUU

>gma-miR10198 MIMAT0040965

AAGGAACUCGAAAUUCAUAGU

>gma-miR156d MIMAT0001672

UUGACAGAAGAUAGAGAGCAC

>gma-miR171s MIMAT0023234

CGAGCCGAAUCAAUACCACUC

>gma-miR169f MIMAT0020986

CAGCCAAGGAUGACUUGCCGG

>gma-miR160d MIMAT0020972

UGCCUGGCUCCCUGUAUGCC

>gma-miR156s MIMAT0023216

UGACAGAAGAGAGUGAGCACU

>gma-miR5369 MIMAT0021603

UGAGAAAAGGAGGAUGUCA

>gma-miR10437 MIMAT0041709

UAUUGUUCAAACAUAUACUGUU

>gma-miR530d MIMAT0023199

UGCAUUUGCACCUGCACUUUA

>gma-miR5786 MIMAT0023222

UGUCGCAGGAUAGAGGGCACU

>gma-miR164h MIMAT0024892

UGGAGAAGCAGGGCACGUGCA

>gma-miR1507c-5p MIMAT0021005

GAGGUGUUUGGGAUGAGAGAA

>gma-miR9750-5p MIMAT0039317

AGGCGCGUAUGGACUUACGACC

>gma-miR162c MIMAT0021644

UCGAUAAACCUCUGCAUCCAG

>gma-miR5042-3p MIMAT0032123

UGGGGCUUGAUCCAAGAUAGG

>gma-miR4354 MIMAT0018246

CAAUUGGAUCGGUCCAACCGGC

>gma-miR9750-3p MIMAT0036378

UGUAAGUCCAUAUGUGCUCUCU

>gma-miR1521b MIMAT0021065

GACUGUCACGUGUCAUAAUCAUA

>gma-miR1515a MIMAT0007374

UCAUUUUGCGUGCAAUGAUCUG

>gma-miR156a MIMAT0001686

UGACAGAAGAGAGUGAGCAC

>gma-miR9746f MIMAT0036371

AAAGUGUUUGAAUCUCAAUUAGAU

>gma-miR9746c MIMAT0036368

AAAGUGUUUGAAUCUCAAUUAGAU

>gma-miR2111b MIMAT0023220

UAAUCUGCAUCCUGAGGUUUA

>gma-miR10407b MIMAT0041706

AGUUAACGGAUGAAUGAAUUUGUC

>gma-miR399f MIMAT0023241

UGCCAAAGGAGAUUUGCCCAG

>gma-miR5370 MIMAT0021607

CUAAAGAUUGUCCAAAAGGAA

>gma-miR9724 MIMAT0036339

UAGAGAUAGUGUCAAAAUAGAA

>gma-miR319k MIMAT0021665

UUGGACUGAAGGGAGCUCCCU

>gma-miR9758 MIMAT0036392

UGUUUAGUCAUGCAAGUUUAG

>gma-miR4362 MIMAT0018258

CCUUAGGACAGACGUCAUGUAG

>gma-miR5378 MIMAT0021619

CAUCUGAAGGAUAGAACACAUA

>gma-miR482d-3p MIMAT0021672

UCUUCCCUACACCUCCCAUACC

>gma-miR164b MIMAT0020975

UGGAGAAGCAGGGCACGUGC

>gma-miR4416a MIMAT0018344

ACGGGUCGCUCUCACCUAGG

>gma-miR10431 MIMAT0041697

UUAGAAUAAACAUGUGUAGAAAUU

>gma-miR4396 MIMAT0018311

UGUAGUUUCUAAGACGAUGCUGAC

>gma-miR4351 MIMAT0018240

AUUGGGAUUCAGUUGGAGUUGG

>gma-miR9766 MIMAT0036403

UUUGAAGGGAAGGAAUGAAAC

>gma-miR4405b MIMAT0041662

UUCCGAAUAACCGACUUAGAAAUC

>gma-miR4392b MIMAT0040970

UCUGCGAAAAUGUGAUUUCGGA

>gma-miR395l MIMAT0024924

AUGAAGUGUUUGGGGGAACUC

>gma-miR5762 MIMAT0023158

UCAUAGGAGGAAUCAACUGGC

>gma-miR10435 MIMAT0041703

UUCCAUUUCAACGUCGGGUUAGAA

>gma-miR10411 MIMAT0041648

UAAUUUUAGACUGAUAUUUUGCCU

>gma-miR9756 MIMAT0036387

UGAGAACUUUAUCCAAACAGAG

>gma-miR530a MIMAT0021003

UGCAUUUGCACCUGCACUUU

>gma-miR5037b MIMAT0021605

AACCCUCAAAGGCUUCCUAG

>gma-miR169h MIMAT0021062

GGCGAGACAUCUUGGCUCAUU

>gma-miR1522 MIMAT0007383

UUUAUUGCUUAAAAUGAAAU

>gma-miR10190 MIMAT0040955

UCCUGAUGACUAUUAUGAGCU

>gma-miR5041-3p MIMAT0032122

GUUGAGCAAGUUGAAGAUGAA

>gma-miR4368a MIMAT0018267

AAGACGGUACUUACCUCAGUAACA

>gma-miR10405b MIMAT0041673

UUGUUUCUUAUAAAAAGGACC

>gma-miR1528 MIMAT0007390

AUAGAUUAGAUCAAUAUAUUAGU

>gma-miR4378b MIMAT0018284

UAGAACUGUCUUAGAAUGUGCUAC

>gma-miR319h MIMAT0021662

UUGGACUGAAGGGAGCUCCCU

>gma-miR9767 MIMAT0036404

AUGGAAUGGUUACUUAUGAAAAGA

>gma-miR166b MIMAT0001678

UCGGACCAGGCUUCAUUCCCC

>gma-miR5674b MIMAT0023248

UAAUUGUGUUGUACAUUAUCA

>gma-miR1509b MIMAT0011201

UUAAUCAAGGAAAUCACGGUU

>gma-miR4399 MIMAT0018315

UUAACGAAAAAGGACUAACGAC

>gma-miR166j-5p MIMAT0021647

GGAAUGUUGUUUGGCUCGAGG

>gma-miR394b-5p MIMAT0022973

UUGGCAUUCUGUCCACCUCC

>gma-miR395g MIMAT0023246

UGAAGUGUUUGGGGGAACUUU

>gma-miR5676 MIMAT0022462

UCGACACCAUAUGUAGAGGCAG

>gma-miR171p MIMAT0023208

UUGAGCCGCGUCAAUAUCUUA

>gma-miR1520r MIMAT0018313

UGUCACAUCCUGGUUGGACAUGAA

>gma-miR171j-3p MIMAT0022990

UGAUUGAGCCGUGCCAAUAUC

>gma-miR4397-3p MIMAT0018312

UGUCAAAGAUGUGGCGAAUACU

>gma-miR10186k MIMAT0041719

UUUUGGGAAUUAAAAUGUAAACUU

>gma-miR171c-3p MIMAT0022972

UUGAGCCGUGCCAAUAUCACA

>gma-miR5559 MIMAT0023168

UACUUGGUGAAUUGUUGGAUC

>gma-miR156o MIMAT0021639

UUGACAGAAGAGAGUGAGCAC

>gma-miR167i MIMAT0021076

UCAUGCUGGCAGCUUCAACUGGU

>gma-miR4342 MIMAT0018230

AAUCGACUUAGAAUGUAGGAUGGU

>gma-miR9749 MIMAT0036376

UUAGCUUCUUUCACCUUUCCC

>gma-miR9728 MIMAT0036346

CGCAGAACUGAAACAAGUUGA

>gma-miR1520o MIMAT0018296

UCAUCGUCCAAUCAGAAUGUGACA

>gma-miR166k MIMAT0023223

UCUCGGACCAGGCUUCAUUCC

>gma-miR1519 MIMAT0007378

UAAGUGUUGCAAAAUAGUCAUU

>gma-miR1508a MIMAT0007366

UCUAGAAAGGGAAAUAGCAGUUG

>gma-miR169c MIMAT0007357

AAGCCAAGGAUGACUUGCCGA

>gma-miR1520i MIMAT0018257

AACGUGACACGUGACGGUCAACAU

>gma-miR160c MIMAT0020971

UGCCUGGCUCCCUGUAUGCC

>gma-miR530e MIMAT0023209

UGCAUUUGCACCUGCACUUUA

>gma-miR1514a-3p MIMAT0032031

AUGCCUAUUUUAAAAUGAAAA

>gma-miR171h MIMAT0021054

AUUGAGACGAGCCGAAUCAAU

>gma-miR5774b MIMAT0023193

GCUGGCGUCGACACGUGGCAU

>gma-miR5379 MIMAT0021620

AUGAAAAUCAUUCAUUAUGAUAUC

>gma-miR10201 MIMAT0040969

UACCGGUAGUAAAUGGAUGCCU

>gma-miR395i MIMAT0024921

AUGAAGUGUUUGGGGGAACUC

>gma-miR394g MIMAT0024919

UUGGCAUUCUGUCCACCUCC

>gma-miR10448 MIMAT0041729

UUAGGCUUUGUGGGACGUGAUU

>gma-miR10445 MIMAT0041724

AAGGACCAAAUUUAUGAAUUAAAU

>gma-miR9724b MIMAT0041687

UAUCUUGGCACUAUCUCUAAC

>gma-miR169u MIMAT0023249

CAGCCAAGGAUGACUUGCCGU

>gma-miR5368 MIMAT0021602

GGACAGUCUCAGGUAGACA

>gma-miR4997b MIMAT0041691

UCGCGGCAGAAGAAAUCCUGAU

>gma-miR5380b MIMAT0021622

GAAAAUGAAUGAUGAGGAUGGGGA

>gma-miR4348b MIMAT0036344

UCUUUUGAAUUUGACUAUUAG

>gma-miR156j MIMAT0021634

UUGACAGAAGAUAGAGAGCAC

>gma-miR172h-5p MIMAT0021657

GCAGCAGCAUCAAGAUUCACA

>gma-miR4346 MIMAT0018235

GAAAGACCAAACGAGAAGCUGCAU

>gma-miR4403 MIMAT0018321

ACGGACACCGAACACGACACGGAC

>gma-miR408c-3p MIMAT0021632

AUGCACUGCCUCUUCCCUGGC

>gma-miR5376 MIMAT0021617

UGAAGAUUUGAAGAAUUUGGGA

>gma-miR10191 MIMAT0040956

GGCGACUGCGGCUCCUCCGCCG

>gma-miR393g MIMAT0024914

UCCAAAGGGAUCGCAUUGAUCC

>gma-miR169r MIMAT0023217

UGAGCCAGGAUGGCUUGCCGGC

>gma-miR828b MIMAT0023198

UCUUGCUCAAAUGAGUAUUCCA

>gma-miR319l MIMAT0021666

UUGGACUGAAGGGAGCUCCUUC

>gma-miR5034 MIMAT0021055

GGUACCCUUUCAGAUAGUCUCA

>gma-miR862a MIMAT0021004

UGCUGGAUGUCUUUGAAGGAAU

>gma-miR10407a MIMAT0041643

AGUUAACGGAUGAAUGAAUUUGUC

>gma-miR9762 MIMAT0036399

UACAGAUCUUUGGAAACAGGC

>gma-miR169s-5p MIMAT0023218

AAGCCAAGGAUGACUUGCCGG

>gma-miR9730 MIMAT0036349

CGAUUGCUGUCAUAACUGCUGC

>gma-miR166i-5p MIMAT0021645

GGAAUGUCGUCUGGUUCGAG

>gma-miR5371-3p MIMAT0021609

UCUCAGUGACUAAUUUCUAGA

>gma-miR9736 MIMAT0036356

UGAAAGACAAACAAAGGUGGG

>gma-miR5038b MIMAT0021612

UGAGAAUUUGGCCUCUGUCCA

>gma-miR1513a-5p MIMAT0007371

UGAGAGAAAGCCAUGACUUAC

>gma-miR4387d MIMAT0018301

AUGUCACUGAUUAGGCAUGAUGAU

>gma-miR4348a-5p MIMAT0018237

AAACUUGUAAGAUGGUGACAUU

>gma-miR4349-3p MIMAT0037390

UCACUUAUAUACUCUUUCUUGGCC

>gma-miR10425 MIMAT0041675

UUACAACCGUCUUUGAAAUCGCCU

>gma-miR5380c MIMAT0036391

AUGAAUGGUGAAGAUGAAGAG

>gma-miR390g MIMAT0024909

AAGCUCAGGAGGGAUAGCGCC

>gma-miR1520f-5p MIMAT0032111

AUUGUCACGUGUCAUGUUCUGAUU

>gma-miR156u MIMAT0024880

UGACAGAAGAGAGUGAGCAC

>gma-miR10186b MIMAT0041642

UUUUGGGAAUUAAAAUGUAAACUU

>gma-miR396b-3p MIMAT0020923

GCUCAAGAAAGCUGUGGGAGA

>gma-miR395k MIMAT0024923

AUGAAGUGUUUGGGGGAACUC

>gma-miR399c MIMAT0023228

UGCCAAAGGAGAGUUGCCCUG

>gma-miR319g MIMAT0021661

UUGGACUGAAGGGAGCUCCUUC

>gma-miR4402 MIMAT0018320

ACAUAUUAUGGGUCUCAGACGGAC

>gma-miR10413b MIMAT0041659

UUUUGGUAGAGAACGAAACCCU

>gma-miR395h MIMAT0024920

AUGAAGUGUUUGGGAGAACUC

>gma-miR166i-3p MIMAT0021646

UCGGACCAGGCUUCAUUCCCC

>gma-miR10424a MIMAT0041671

UUUUCUAAUUUAUUAGGGACU

>gma-miR10405a MIMAT0041639

UUGUUUCUUAUAAAAAGGACC

>gma-miR4348c MIMAT0036389

UGUUAAACUUGCAAGAUGACA

>gma-miR1516a-5p MIMAT0020937

CAAGUUAUAAGCUCUUUUGAGAG

>gma-miR5669 MIMAT0022451

CAAUGUAGUGUGGUAAGUGGUC

>gma-miR9761 MIMAT0036397

UGAAGGUCUAGGAUAUUUUGU

>gma-miR5677 MIMAT0022465

UUUGGUCUUUAAUCAAGCUGA

>gma-miR393a MIMAT0007362

UCCAAAGGGAUCGCAUUGAUC

>gma-miR530b MIMAT0022463

UGCAUUUGCACCUGCACUUUA

>gma-miR1523b MIMAT0021051

UCAUCGCUCCUGAGCUCACA

>gma-miR5779 MIMAT0023184

CAAGUCCAAAGUAGGAAUGUUGCA

>gma-miR398b-5p MIMAT0037314

GAGUGGAUCUGAGAACACAAGG

>gma-miR4995 MIMAT0021013

AGGCAGUGGCUUGGUUAAGGG

>gma-miR396j MIMAT0023214

AUUCAAGAUAGCUGUGGAAAA

>gma-miR4378a MIMAT0018283

AUAGGACUGUCUUAGAAUGGUGUA

>gma-miR169j-5p MIMAT0021649

UAGCCAAGAAUGACUUGCCGG

>gma-miR10193f MIMAT0041725

UAAAAAAACCAAUGUUAACUGU

>gma-miR4994-5p MIMAT0021012

GGUUAGCUCAAGGAUCUCAC

>gma-miR171a MIMAT0007358

UGAGCCGUGCCAAUAUCACGA

>gma-miR399o MIMAT0041728

GGGCUCCUCUCUCCUGGCAUG

>gma-miR9755 MIMAT0036386

UGAUCCAGGAACUUUUCAUCU

>gma-miR172i-3p MIMAT0022996

GGAAUCUUGAUGAUGCUGCAU

>gma-miR1527 MIMAT0007389

UAACUCAACCUUACAAAACC

>gma-miR10424c MIMAT0041712

UUUUCUAAUUUAUUAGGGACU

>gma-miR1533 MIMAT0007396

AUAAUAAAAAUAAUAAUGA

>gma-miR4341 MIMAT0018229

UGUGUUGAAAGUUUAACAUGACGG

>gma-miR2118b-3p MIMAT0021000

UUGCCGAUUCCACCCAUUCCU

>gma-miR10412 MIMAT0041649

GAACGUUCUGGAGAACUCUAGAGU

>gma-miR1513b MIMAT0021008

UGAGAGAAAGCCAUGACUUAC

>gma-miR4409 MIMAT0018328

UAACAAGUGGGUUUGUUGACUG

>gma-miR10199 MIMAT0040966

AGCAAUGUUGAGCUUGGGCCU

>gma-miR482a-3p MIMAT0007364

UCUUCCCAAUUCCGCCCAUUCCUA

>gma-miR390a-3p MIMAT0007360

CGCUAUCCAUCCUGAGUUUC

>gma-miR156i MIMAT0020969

UUGACAGAAGAUAGAGAGCAC

>gma-miR9737 MIMAT0036357

UUGUGGCUGAAAUCACUGUUGC

>gma-miR167h MIMAT0021064

AUCAUGCUGGCAGCUUCAACUGGU

>gma-miR5040 MIMAT0021067

AUGAUAUAUAACAAGCAUGAG

>gma-miR4393a MIMAT0018308

UGAGAAAAGGACGGCAGAAAAGCC

>gma-miR156p MIMAT0023196

UUGACAGAAGAAAGGGAGCAC

>gma-miR10442 MIMAT0041715

CUACAUGCUGCACUUGGAUCC

>gma-miR396a-3p MIMAT0020922

UUCAAUAAAGCUGUGGGAAG

>gma-miR166l MIMAT0023235

GGAAUGUUGUCUGGCUCGAGG

>gma-miR319m MIMAT0021667

UUGGACUGAAGGGAGCUCCCU

>gma-miR171b-3p MIMAT0007363

CGAGCCGAAUCAAUAUCACUC

>gma-miR172a MIMAT0001682

AGAAUCUUGAUGAUGCUGCAU

>gma-miR164i MIMAT0024893

UGGAGAAGCAGGGCACGUGCA

>gma-miR169t MIMAT0023233

UAGCCAAGGAUGGACUUGCCUA

>gma-miR9729 MIMAT0036347

GUAAUGAGUAGAAACAUUUAGAAG

>gma-miR171q MIMAT0023215

UUGAGCCGUGCCAAUAUCACA

>gma-miR5785 MIMAT0023191

UAGUGUUGUCCUGUCGAACACGGA

>gma-miR169n-5p MIMAT0021654

CAGCCAAGGGUGAUUUGCCGG

>gma-miR1514b-3p MIMAT0032032

AUGCCUAUUUUAAAAUGAAAA

>gma-miR4345 MIMAT0018233

UAAGACGGAACUUACAAAGAUU

>gma-miR5678 MIMAT0022466

UUCCAUGAUAAGAUCUUUGAC

>gma-miR4374a MIMAT0018278

UAAGACGGUCGUGAUGUCAGCA

>gma-miR166u MIMAT0024903

UCUCGGACCAGGCUUCAUUC

>gma-miR391a-3p MIMAT0020951

AGCAUCAUAUCUCCUGCAUAG

>gma-miR5761b MIMAT0023173

UUUUGUGUCGUGAAGCUUUUG

>gma-miR9747 MIMAT0036374

CAUGCGAUGAUUUAAAUACUUU

>gma-miR395j MIMAT0024922

AUGAAGUGUUUGGGGGAACUC

>gma-miR169j-3p MIMAT0021650

UUUCGACGAGUUGUUCUUGGC

>gma-miR5380a MIMAT0021621

GAAAAUGAAUGAUGAGGAUGGGGA

>gma-miR5033 MIMAT0021053

GGCUGUACAAAAGGAAACUAC

>gma-miR9735 MIMAT0036355

UACGGCUUAAGUUCAACUUUGGAG

>gma-miR5765 MIMAT0023162

CGAAACGUUGAGGUAUAUGUGGAC

>gma-miR4393b MIMAT0018317

UUGAAAAGGGACAGCAGAGAAGCC

>gma-miR10192 MIMAT0040958

UAAACGUUUGAUCCCUUGUAU

>gma-miR5772 MIMAT0023174

AGAAUGUGAGUUAGAGUGAGCAUC

>gma-miR10189 MIMAT0040954

AAGAACACUAGAACCAUCUCC

>gma-miR4371a MIMAT0018273

AAGUGAUGACAUGACAAGCGAAGU

>gma-miR5771 MIMAT0023171

AUCUCAAGUGGAUUGCUUAAGGAC

>gma-miR396g MIMAT0021071

UUCUUGAACUUCUUAUGCAUC

>gma-miR1520a MIMAT0007381

UAGAACAUGAUACAUGACAGUCA

>gma-miR10407d MIMAT0041655

UUAACGGCUGGAUAUAUUUGUUGC

>gma-miR393j MIMAT0024917

UUCCAAAGGGAUCGCAUUGAUC

>gma-miR396b-5p MIMAT0001688

UUCCACAGCUUUCUUGAACUU

>gma-miR4416c-3p MIMAT0037453

ACGGGUCGCUCUCACCUGGAG

>gma-miR9734 MIMAT0036354

UCGUGAAUGAGAUUUGUGUUGCUU

>gma-miR10407c MIMAT0041650

UUAACAGUCGGAUAUAUUUAUCGC

>gma-miR4387b MIMAT0018272

AAGGUGUGAUGGCAUGACACUCUG

>gma-miR1520q MIMAT0018332

AUUGACCAAUCAGAACAUGACACA

>gma-miR4350 MIMAT0018239

UCAAAUGAUUUUGUGUCGUUGG

>gma-miR397b-3p MIMAT0022991

UAUUGACGCUGCACUCAAUCA

>gma-miR2111e MIMAT0023237

UAAUCUGCAUCCUGAGGUUUA

>gma-miR1510a-5p MIMAT0017338

AGGGAUAGGUAAAACAAUGACUGC

>gma-miR403a MIMAT0021613

UUAGAUUCACGCACAAACUUG

>gma-miR4352a MIMAT0018241

AUUUCUAGGACAUACUACGACGGU

>gma-miR4391 MIMAT0018304

UCUCGGCAAAGAACUAAGAAGAAG

>gma-miR395a MIMAT0021624

CUGAAGUGUUUGGGGGAACUC

>gma-miR2111d MIMAT0023224

GUCCUUGGGAUGCAGAUUACG

>gma-miR10413a MIMAT0041651

UUUUGGUAGAGAACGAAACCCU

>gma-miR4400 MIMAT0018316

UUCGGAAAAAUUCUGGAAGACGUC

>gma-miR167k MIMAT0036388

UGAAGCUGCCAGCCUGAUCUUA

>gma-miR4372b MIMAT0021073

UAAUAAAAUCGUGACAUGUAAC

>gma-miR5037d MIMAT0022455

CGGGAGCCUAUGAAGGUUAAC

>gma-miR398d MIMAT0036343

UGUGUUCUCAGGUCGCCCCUG

>gma-miR171f MIMAT0020990

UGAUUGAGCCGUGCCAAUAUC

>gma-miR156q MIMAT0023203

UGACAGAAGAGAGUGAGCACU

>gma-miR5375 MIMAT0021616

ACUAUAGAAGUACUUGUGGAGC

>gma-miR5781 MIMAT0023186

CUGAAACUGAGACUGCAUCUGG

>gma-miR399d MIMAT0023231

UGCCAAAGGAGAUUUGCCCAG

>gma-miR5032 MIMAT0021052

AGAGCCACUUUUGGGUUCCCUAU

>gma-miR5667-5p MIMAT0032130

AAUCCAUUCAGAUCUGUUUCG

>gma-miR396h MIMAT0021668

UCCACAGCUUUCUUGAACUG

>gma-miR5768 MIMAT0023165

AAGUGCAAUACUGAUCUUCGGAAC

>gma-miR1508b MIMAT0010081

UAGAAAGGGAAAUAGCAGUUG

>gma-miR4382 MIMAT0018289

UAUGUUAACUGAUUUCAUGGAU

>gma-miR408b-5p MIMAT0021630

CUGGGAACAGGCAGGGCACG

>gma-miR10186i MIMAT0041699

UUUUGGGAAUUAAAAUGUAAACUU

>gma-miR399k MIMAT0041672

UGCCAAAGGAGAUUUGCCCUG

>gma-miR167e MIMAT0011196

UGAAGCUGCCAGCAUGAUCUU

>gma-miR10186j MIMAT0041718

UUUUGGGAAUUAAAAUGUAAACUU

>gma-miR9733 MIMAT0036353

UAACAGACUUAGAUUAACAGA

>gma-miR390d MIMAT0024906

AAGCUCAGGAGGGAUAGCACC

>gma-miR319f MIMAT0020994

UUGGACUGAAGGGGCCUCUU

>gma-miR166a-3p MIMAT0001677

UCGGACCAGGCUUCAUUCCCC

>gma-miR167d MIMAT0010078

UGAAGCUGCCAGCAUGAUCUA

>gma-miR396i-3p MIMAT0021670

GUUCAAUAAAGCUGUGGGAAG

>gma-miR156z MIMAT0024885

AUUGGAGUGAAGGGAGCU

>gma-miR4371c MIMAT0018268

GACGUGACAGACGGAAUAUCACAU

>gma-miR4375 MIMAT0018279

UACCACUAGUGGUCGCGCCUGGCA

>gma-miR4380b MIMAT0018287

UAUGGUCAUACGGAUUGUUGAU

>gma-miR4388 MIMAT0018300

AAUCUUAGGGACCAAAUUGACAGC

>gma-miR828a MIMAT0023195

UCUUGCUCAAAUGAGUAUUCCA

>gma-miR4364b MIMAT0018234

UAACAACAGCGGAAGAACCUUCUU

>gma-miR156e MIMAT0001673

CUGACAGAAGAUAGAGAGCAC

>gma-miR10188 MIMAT0040953

UCUCUGAUUUUGCAGAUAAGGACU

>gma-miR159b-3p MIMAT0007351

AUUGGAGUGAAGGGAGCUCCA

>gma-miR5675 MIMAT0022460

UAGAGACGACAACAAUGGAAA

>gma-miR10427a MIMAT0041684

AAGAGAAUUGGGCUAGAGCUGC

>gma-miR172f MIMAT0018333

AGAAUCUUGAUGAUGCUGCA

>gma-miR162b MIMAT0020974

UCGAUAAACCUCUGCAUCCAG

>gma-miR2118a-5p MIMAT0022981

GGAGAUGGGAGGGUCGGUAAAG

>gma-miR10193a MIMAT0040960

UCUGAAACCGAUGUUAACUACU

>gma-miR1520d MIMAT0007379

AUCAGAACAUGACACGUGACAA

>gma-miR9754 MIMAT0036385

UGACCACAUUAGACUGAAAGUA

>gma-miR164j MIMAT0024894

UGGAGAAGCAGGGCACGUGCA

>gma-miR4408 MIMAT0018327

UAACAACAUUGGAUGAGGGUUGGA

>gma-miR10443 MIMAT0041721

UCGACUGAGAACAACAUAAUUCAU

>gma-miR1536 MIMAT0007380

AAGCAGAGACAAAUGUGUUUA

>gma-miR4394 MIMAT0018309

AAUGGACUAAAGAGAAAGGGGCCG

>gma-miR9764 MIMAT0036401

UGGAUACACUCUUACUUUCUC

>gma-miR4359b MIMAT0018253

AACGCGUGAUAUGUUAACAUCGGU

>gma-miR1520k MIMAT0018291

AAUCAGAACAUGACACAUGACAGU

>gma-miR9741 MIMAT0036361

AAUGUGUUGAACUGAGUGAAGACA

>gma-miR168a MIMAT0001681

UCGCUUGGUGCAGGUCGGGAA

>gma-miR4343b MIMAT0018306

UCUUACAGAUCAAGUUGAUUCGGA

>gma-miR4348d MIMAT0041640

UUUCGGUGUCGGUGAAUUGCC

>gma-miR1515b MIMAT0024926

UCAUUUUGCGUGCAAUGAUCUG

>gma-miR156l MIMAT0021636

UUGACAGAAGAUAGAGAGCAC

>gma-miR10410a MIMAT0041647

UUUCCACAUCGUUGCUGACAUAAG

>gma-miR10195 MIMAT0040962

UAUGAUUUUGUGGAUCAAAGGA

>gma-miR5775 MIMAT0023179

AUAAGCUCUUUUGAGAGCUUC

>gma-miR5763a MIMAT0023159

UGAACUAUACAAAGACGGUUA

>gma-miR169p MIMAT0023211

UGAGCCAAGGAUGACUUGCCG

>gma-miR10416 MIMAT0041658

AAGAUUUGACUGUUGGAACUGCCU

>gma-miR5776 MIMAT0023181

AACUUGGGCUGAGCUUAGGUG

>gma-miR166c-5p MIMAT0022980

GGAAUGUUGUCUGGCUCGAGG

>gma-miR397b-5p MIMAT0021628

UCAUUGAGUGCAGCGUUGAUG

>gma-miR1520g MIMAT0018245

CAAUCAGAACAUGACACGUGACAA

>gma-miR171o-3p MIMAT0023204

UUGAGCCGUGCCAAUAUCACA

>gma-miR394d MIMAT0023210

UUGGCAUUCUGUCCACCUCC

>gma-miR9746g MIMAT0036372

AAAGUGUUUGAAUCUCAAUUAGAU

>gma-miR164e MIMAT0024889

UGGAGAAGCAGGGCACGUGCA

>gma-miR156h MIMAT0020968

UGACAGAAGAGAGUGAGCAC

>gma-miR5037a MIMAT0021060

GCCUCAAAGGCUUCCACUACUG

>gma-miR156ab MIMAT0024887

AUUUAAGUGAUGGGAGCUCCG

>gma-miR4352b MIMAT0018276

UAAAAUGUAGACAUUCUAAGACGG

>gma-miR169k MIMAT0021651

CAGCCAAGAAUGACUUGCCGG

>gma-miR394e MIMAT0023230

UUGGCAUUCUGUCCACCUCC

>gma-miR393e MIMAT0024912

UCCAAAGGGAUCGCAUUGAUCC

>gma-miR5668 MIMAT0022450

AGCAAUGGAAUUAUAGACUGC

>gma-miR4415a-3p MIMAT0020955

UUGAUUCUCAUCACAACAUGG

>gma-miR172d MIMAT0011199

GGAAUCUUGAUGAUGCUGCAGCAG

>gma-miR10436 MIMAT0041707

UCUUGGCAACACAACUUUUAACAU

>gma-miR169e MIMAT0018335

AGCCAAGGAUGACUUGCCGG

>gma-miR167j MIMAT0021677

UGAAGCUGCCAGCAUGAUCUG

>gma-miR5674a MIMAT0022459

UAAUUGUGUUGUACAUUAUCA

>gma-miR4387e MIMAT0022464

UGUUAGUGAUAAGGCGUGAUG

>gma-miR390a-5p MIMAT0007359

AAGCUCAGGAGGGAUAGCGCC

>gma-miR5031 MIMAT0021050

UUAAUGAUUAACAUCUAAUUU

>gma-miR164k MIMAT0024895

UGGAGAAGCAGGGCACGUGCA

>gma-miR4385 MIMAT0018297

AAUCGAUGUAGAAAAGUGAUUGGU

>gma-miR2118a-3p MIMAT0020999

UUGCCGAUUCCACCCAUUCCU

>gma-miR166c-3p MIMAT0020978

UCGGACCAGGCUUCAUUCCCC

>gma-miR397a MIMAT0021627

UCAUUGAGUGCAGCGUUGAUG

>gma-miR5782 MIMAT0023187

UAGCUGGUAGGAGAAGUUCAG

>gma-miR10186h MIMAT0041696

UUUUGGGAAUUAAAAUGUAAACUU

>gma-miR10410b MIMAT0041679

UUUCCACAUCGUUGCUGACAUAAG

>gma-miR4348a-3p MIMAT0037389

UAUCACAAUGUCAUCAUCUUG

>gma-miR2111f MIMAT0023238

UAAUCUGCAUCCUGAGGUUUA

>gma-miR10414 MIMAT0041652

UCUGAGAAGAACCAUUGGUACC

>gma-miR393h MIMAT0024915

UUCCAAAGGGAUCGCAUUGAUC

>gma-miR395b MIMAT0021625

CUGAAGUGUUUGGGGGAACUC

>gma-miR1512c MIMAT0022458

UAACUGAACAUUCUUAGAGCAU

>gma-miR156r MIMAT0023212

CUGACAGAAGAUAGAGAGCAU

>gma-miR1535a MIMAT0007398

CUUGUUUGUGGUGAUGUCU

>gma-miR1509a MIMAT0007367

UUAAUCAAGGAAAUCACGGUCG

>gma-miR172l MIMAT0023205

GGAAUCUUGAUGAUGCUGCAU

>gma-miR9765 MIMAT0036402

UAAUACAGAAUUCGGAGACAAC

>gma-miR1530 MIMAT0007392

UUUUCACAUAAAUUAAAAUAU

>gma-miR396k-3p MIMAT0023219

GCUCAAGAAAGCUGUGGGAGA

>gma-miR1507a MIMAT0007365

UCUCAUUCCAUACAUCGUCUGA

>gma-miR4364a MIMAT0018260

CGCGAGAUCGCACGGAAGAAGGUU

>gma-miR171g MIMAT0020991

UGAUUGAGCCGUGCCAAUAUC

>gma-miR399l MIMAT0041678

UGCCAAAGGAGAGCUGCCCUG

>gma-miR156y MIMAT0024884

UGACAGAAGAGAGUGAGCAC

>gma-miR1512a-3p MIMAT0032030

GCUUUAAGAAUUUCAGUUAUG

>gma-miR10440 MIMAT0041713

UUGGGACAAUACUUUAGAUAU

>gma-miR390b-3p MIMAT0020935

UACUUGGCGCUAUCUAUCUUGA

>gma-miR10423 MIMAT0041670

AUGAAAGCACCAAAUUGAUAGGACU

>gma-miR9759 MIMAT0036394

UAUAAGCAAGUAGAAUUUAAU

>gma-miR5667-3p MIMAT0022449

AAACAGAUCUAAAUGGAUUCC

>gma-miR10200 MIMAT0040967

AGGUUUUAAAGAAAUUAAAUG

>gma-miR398b-3p MIMAT0001690

UGUGUUCUCAGGUCACCCCUU

>gma-miR171n MIMAT0023202

UUGAGCCGCGUCAAUAUCUUA

>gma-miR319n MIMAT0023207

UUUGGACCGAAGGGAGCCCCU

>gma-miR166g MIMAT0020982

UCGGACCAGGCUUCAUUCCCC

>gma-miR1446 MIMAT0040968

UUCUGAACUCUCUCCCUCAAU

>gma-miR10427b MIMAT0041694

CAAGAGAAUUGGGCUAGAGCUG

>gma-miR1520n MIMAT0018295

UCAAUCAGAACAUGACACGUGACA

>gma-miR4401b MIMAT0022461

UCAAAGACGUUGCUGAGGUAA

>gma-miR171d MIMAT0020988

UGAUUGAGUCGUGUCAAUAUC

>gma-miR319e MIMAT0020993

UUGGACUGAAGGGAGCUCCC

>gma-miR9726 MIMAT0036342

UAUAGGCAUUAUUUUUUUCUUC

>gma-miR408b-3p MIMAT0021631

AUGCACUGCCUCUUCCCUGGC

>gma-miR156g MIMAT0018339

ACAGAAGAUAGAGAGCACAG

>gma-miR5225 MIMAT0023169

CCUGUCGUAGGAGAGAUGACGC

>gma-miR2107 MIMAT0010083

CAAACCUCCGUAGCCUGUAUC

>gma-miR164f MIMAT0024890

UGGAGAAGCAGGGCACGUGCA

>gma-miR156aa MIMAT0024886

AUUGGAGUGAAGGGAGCU

>gma-miR319a MIMAT0001684

UUGGACUGAAGGGAGCUCCC

>gma-miR4993 MIMAT0021011

GAGCGGCGGCGGUGGAGGAUG

>gma-miR10408 MIMAT0041644

CGAGGACUCCUUUAAAUAGACU

>gma-miR169b MIMAT0007356

CAGCCAAGGAUGACUUGCCGA

>gma-miR394f MIMAT0023239

UUGGCAUUCUGUCCACCUCC

>gma-miR167f MIMAT0011197

UGAAGCUGCCAGCAUGAUCUU

>gma-miR9727 MIMAT0036345

UGAAGUUACUCUGAGCACUGAG

>gma-miR10444 MIMAT0041723

GACAUACAUGUGAGUUCGGAUGUA

>gma-miR10415 MIMAT0041656

AUCUUAGAAUGUGAGUUUGGAUCU

>gma-miR5039 MIMAT0021066

CCCUUUUUUAAUCGUUGCAUG

>gma-miR399a MIMAT0023194

UGCCAAAGGAGAGUUGCCCUG

>gma-miR9763 MIMAT0036400

ACCCAUCCAACUCUGAAGAUA

>gma-miR4413b MIMAT0021675

UAAGAGAAUUGUAAGUCACU

>gma-miR9742 MIMAT0036362

UGUGUUGUUUGUUUUGUAGCA

>gma-miR171k-5p MIMAT0021655

CGAUGUUGGUGAGGUUCAAUC

>gma-miR171l MIMAT0021678

CGAUGUUGGUGAGGUUCAAUC

>gma-miR172g MIMAT0021656

GCAGCACCAUCAAGAUUCAC

>gma-miR1513a-3p MIMAT0020936

UUUAAAUGUGUAUAAGUCAUGGU

>gma-miR4359a MIMAT0018251

AACGAAGUGACUCUAACAUCGGUU

>gma-miR1520j MIMAT0018263

AAGAACGUGACACAUGACAAUCAA

>gma-miR9722 MIMAT0036337

UAAUAGAGGGAAGAAGAUGAA

>gma-miR390c MIMAT0020995

CGCUAUCCAUCCUGAGUUUC

>gma-miR4369 MIMAT0018270

GGAUCAAGCUGAUCCGGAAGUGGA

>gma-miR5780c MIMAT0036352

UUUAAUAUAUCAGGGACUUGGA

>gma-miR1529 MIMAT0007391

UUAAAGGAAACAAUUAAUCGUUA

>gma-miR4413a MIMAT0018338

AAGAGAAUUGUAAGUCACUG

>gma-miR394c-5p MIMAT0020996

UUGGCAUUCUGUCCACCUCC

>gma-miR390f MIMAT0024908

AAGCUCAGGAGGGAUAGCGCC

>gma-miR396e MIMAT0018345

UUCCACAGCUUUCUUGAACUGU

>gma-miR9753 MIMAT0036384

ACAAUUUGGGACUUAGGGCUACAA

>gma-miR5774a MIMAT0023176

GCUGGCGUCGACACGUGGCAU

>gma-miR9746i MIMAT0036383

AAAGUGUUUGAAUUUCAAUUAGAU

>gma-miR10446 MIMAT0041726

UCAACUCAAGUCGAGCUCAAGCCU

>gma-miR10194 MIMAT0040961

UGUUGAAGGUGUGUAUGACAAG

>gma-miR4349-5p MIMAT0018238

UAUUGGCUAGAGAUAAGACAAAGA

>gma-miR164c MIMAT0020976

UGGAGAAGCAGGGCACGUGC

>gma-miR5770c MIMAT0040959

UAGGACUAUGGUUUGGAUUU

>gma-miR169o MIMAT0023206

UGAGCCAGGAUGGCUUGCCGGC

>gma-miR5764b MIMAT0041701

UGAUGGAUCUUUGCUUGGUACC

>gma-miR167l MIMAT0041654

UGAAGCUGCCAGCAUGAUCU

>gma-miR5770b MIMAT0023189

UUAGGACUAUGGUUUGGACAA

>gma-miR10420 MIMAT0041666

CCUGAACCAUCAUUUUUUU

>gma-miR399i MIMAT0036382

UGCCAAAGGAGAAUUGCCCUG

>gma-miR482a-5p MIMAT0017337

AGAAUUUGUGGGAAUGGGCUGA

>gma-miR5767 MIMAT0023164

UGGAGGACCUUUGAAGGUGCA

>gma-miR172e MIMAT0011200

GGAAUCUUGAUGAUGCUGCAGCAG

>gma-miR5035-5p MIMAT0021058

CUUCUAAACAUUUUUUCCCUUA

>gma-miR396f MIMAT0021069

AGCUUUCUUGAACUUCUUAUGCCUA

>gma-miR10433 MIMAT0041700

CUGCGGAUCAAGUUGAUUGUUACG

>gma-miR166o MIMAT0024897

UCGGACCAGGCUUCAUUCCCC

>gma-miR9743 MIMAT0036363

AGAGAGUGCUUUGAAGAAAAUGCC

>gma-miR171j-5p MIMAT0021623

UAUUGGCCUGGUUCACUCAGA

>gma-miR166f MIMAT0020981

UCGGACCAGGCUUCAUUCCCC

>gma-miR5783 MIMAT0023188

GACGACGACGGGGAGGACGCGC

>gma-miR5670b MIMAT0036393

CACAUCAUACCAUAUUUGCUUC

>gma-miR156x MIMAT0024883

UGACAGAAGAGAGUGAGCAC

>gma-miR6299 MIMAT0024928

AUUUAAAAUUAUUGAUUUGUCA

>gma-miR10417 MIMAT0041660

GAGGCUGUUUCCAGAUUUGAACCC

>gma-miR1518 MIMAT0007377

UGUGUUGUAAAGUGAAUAUCA

>gma-miR4412-5p MIMAT0020954

UGUUGCGGGUAUCUUUGCCUC

>gma-miR5377 MIMAT0021618

CUGAAGGAUCGAUGUAGAAUGCU

>gma-miR169m MIMAT0021653

CAGCCAAGGAUGACUUGCCGG

>gma-miR171u MIMAT0023240

UGAUUGAGCCGUGCCAAUAUC

>gma-miR9746e MIMAT0036370

AAAGUGUUUGAAUCUCAAUUAGAU

>gma-miR156c MIMAT0001674

UUGACAGAAGAUAGAGAGCAC

>gma-miR167b MIMAT0001680

UGAAGCUGCCAGCAUGAUCUA

>gma-miR4998 MIMAT0021016

AGUUUCGUGACUACAACUUCUG

>gma-miR399b MIMAT0023213

UGCCAAAGGAGAGUUGCCCUG

>gma-miR395c MIMAT0021626

CUGAAGUGUUUGGGGGAACUC

>gma-miR10193d MIMAT0041717

UCUGAAACCGAUGUUAACUACU

>gma-miR4356 MIMAT0018248

CAGGACUGUCUUAGAAAGCCAGGC

>gma-miR4416b MIMAT0023177

UGGGUGAGAGAAACGCGUAUC

>gma-miR10410c MIMAT0041705

UUUCCACAUCGUUGCUGACAUAAG

>gma-miR4407 MIMAT0018326

CAGAGGAAGCAGCACUUGUACC

>gma-miR4397-5p MIMAT0020953

CAUCGUUGACGCUGACUGUACG

>gma-miR5373 MIMAT0021611

UCUCUUGAUUCUAGAUGAUGU

>gma-miR1516b MIMAT0021061

AGCUUCUCUACAGAAAAUAUA

>gma-miR10187 MIMAT0040952

CGAGGAUAAUUGUUGGAACCA

>gma-miR2108b MIMAT0010085

UUAAUGUGUUGUGUUUGUGAG

>gma-miR6300 MIMAT0024929

GUCGUUGUAGUAUAGUGG

>gma-miR9746b MIMAT0036367

AAAGUGUUUGAAUCUCAAUUAGAU

>gma-miR2119 MIMAT0011167

UCAAAGGGAGUUGUAGGGGAA

>gma-miR1523a MIMAT0007384

AUGGGAUAAAUGUGAGCUCA

>gma-miR172j MIMAT0021660

GCAGCAGCAUCAAGAUUCACA

>gma-miR2109-5p MIMAT0010086

UGCGAGUGUCUUCGCCUCUG

>gma-miR10193c MIMAT0041716

UCUGAAACCGAUGUUAACUACU

>gma-miR10186g MIMAT0041686

UUUUGGGAAUUAAAAUGUAAACUU

>gma-miR9746a MIMAT0036366

AAAGUGUUUGAAUCUCAAUUAGAU

>gma-miR156t MIMAT0023225

UUGACAGAAGAAAGGGAGCAC

>gma-miR166r MIMAT0024900

UCGGACCAGGCUUCAUUCCC

>gma-miR5778 MIMAT0023183

CGACGAACUCUUCGUCGGCAUC

>gma-miR4395 MIMAT0018310

UGGAUAGGAGUAUGGGCUUGAG

>gma-miR172h-3p MIMAT0021658

AGAAUCUUGAUGAUGCUGCAU

>gma-miR5761a MIMAT0023157

UUUUGUGUCGUGAAGCUUUUG

>gma-miR171m MIMAT0023200

UUGAGCCGCGUCAAUAUCUCA

>gma-miR4360 MIMAT0018252

CAGUUGACGUACGUACGGAUUGAC

>gma-miR4347 MIMAT0018236

AAGCUUCUUACGGAUCAAGUUGAU

>gma-miR5374-3p MIMAT0032125

UUCAAAUGUCAGAUUAUAAAA

>gma-miR482b-3p MIMAT0020952

UCUUCCCUACACCUCCCAUACC

>gma-miR396c MIMAT0010079

UUCCACAGCUUUCUUGAACUU

>gma-miR1520b MIMAT0007387

GUGACAGUCAUCAUUUAAUAAGA

>gma-miR5377b MIMAT0041693

GACCGAUGUAGAAUGUUCAACAUU

>gma-miR4414a-5p MIMAT0018342

AGCUGCUGACUCGUUGGCUC

>gma-miR4373 MIMAT0018277

AAGUUGACGUACGUACGGAUUGAC

>gma-miR395m MIMAT0024925

AUGAAGUGUUUGGGGGAACUC

>gma-miR10193e MIMAT0041663

UAAAAAAACCAAUGUUAACUGU

>gma-miR4392 MIMAT0018305

UCUGCGAAAAUGUGAUUUCGGA

>gma-miR408a-3p MIMAT0021629

AUGCACUGCCUCUUCCCUGGC

>gma-miR10439 MIMAT0041711

CAGACAGAUAAUGCUAGAGCC

>gma-miR4387a MIMAT0018299

AACAAGACGUGAUGACGUGACACU

>gma-miR394b-3p MIMAT0018336

AGGUGGGCAUACUGUCAACU

>gma-miR5037c MIMAT0021606

AGUGGAACUUUGAGGCCUGC

>gma-miR160b MIMAT0020970

UGCCUGGCUCCCUGUAUGCC

>gma-miR159b-5p MIMAT0020934

GAGUUCCCUGCACUCCAAGUC

>gma-miR5030 MIMAT0021049

AGAACAAUUUGUGUUUUACCGG

>gma-miR5670a MIMAT0022452

CAUCAUACCAUAUUUGCUUCAU

>gma-miR1526 MIMAT0007388

CCGGAAGAGGAAAAUUAAGCAA

>gma-miR10197 MIMAT0040964

AUUUUAGAACGACUUUUUCCU

>gma-miR4355 MIMAT0018247

CACUGUUGUGCUGGGUGUACCA

>gma-miR1534 MIMAT0007397

UAUUUUGGGUAAAUAGUCAU

>gma-miR171k-3p MIMAT0022995

UUGAGCCGCGCCAAUAUCACU

>gma-miR5036 MIMAT0021059

AGAGGCCCUUGGGGAGGAGUAA

>gma-miR169i-5p MIMAT0022983

UGAGCCGGGAUGGCUUGCCGGCA

>gma-miR2111c MIMAT0023221

UAAUCUGCAUCCUGAGGUUUA

>gma-miR4381 MIMAT0018288

UAUGUGACGGUAAACGGUGACAAG

>gma-miR319d MIMAT0020992

UGGACUGAAGGGGAGCUCCUUC

>gma-miR1531-5p MIMAT0032033

AUAUGGACGAAGAGAUAGGUAAAU

>gma-miR5780d MIMAT0036396

UGUUUUGAGUUUCUGAUAAAUU

>gma-miR5042-5p MIMAT0021072

UAUCUUGGAUCACAGCCCCAUU

>gma-miR4361 MIMAT0018256

CCGGAAGAGACUUACGGAUCAACU

>gma-miR399g MIMAT0023242

UGCCAAAGGAGAUUUGCCCAG

>gma-miR5673 MIMAT0022456

CGUGGAAUCUCGCGGAAGACAU

>gma-miR5763c MIMAT0023192

UGAACUAUACAAAGACGGUUA

>gma-miR172b-3p MIMAT0001683

AGAAUCUUGAUGAUGCUGCAU

>gma-miR1516a-3p MIMAT0007375

CAAAAGAGCUUAUGGCUUGUA

>gma-miR4372a MIMAT0018275

UAAAAUCGUGACAUGUGACGGUCA

>gma-miR159a-5p MIMAT0020919

GAGCUCCUUGAAGUCCAAUUG

>gma-miR399j MIMAT0041664

UGCCAAAGGAGAUUUGCCCUG

>gma-miR10432 MIMAT0041698

UUUUCCAGAUCAUAUCGCCGGUUC

>gma-miR166p MIMAT0024898

UCGGACCAGGCUUCAUUCCC

>gma-miR1520p MIMAT0018322

AUGUUGUUAUUGGAUGAUGACGGU

>gma-miR393f MIMAT0024913

UCCAAAGGGAUCGCAUUGAUCC

>gma-miR393c-3p MIMAT0032134

AUCAUGCUAUCCCUUUGGAUU

>gma-miR9757 MIMAT0036390

CAACCCUCCUCAGUUAGAUCUC

>gma-miR1514a-5p MIMAT0007372

UUCAUUUUUAAAAUAGGCAUU

>gma-miR396i-5p MIMAT0021669

UUCCACAGCUUUCUUGAACUG

>gma-miR5769 MIMAT0023166

UGAGGGAAAUGAAGACGACGA

>gma-miR156w MIMAT0024882

UGACAGAAGAGAGUGAGCAC

>gma-miR1511 MIMAT0007369

AACCAGGCUCUGAUACCAUG

>gma-miR398c MIMAT0020997

UGUGUUCUCAGGUCGCCCCUG

>gma-miR10186e MIMAT0041668

UUGGGAAUUAAAAUGUAAACUU

>gma-miR171e MIMAT0020989

UGAUUGAGCCGUGCCAAUAUC

>gma-miR390e MIMAT0024907

AGCUCAGGAGGGAUAGCGCC

>gma-miR4343a MIMAT0018231

AAAAAACUUACGGAUCAAGUUGAU

>gma-miR4416c-5p MIMAT0023172

CUGGGUGAGAGAAACACGUAU

>gma-miR4340 MIMAT0018228

UGCAGAGAUAGGGACGCGCUUA

>gma-miR482e MIMAT0023227

UAUGGGGGGAUUGGGAAGGAA

>gma-miR166e MIMAT0020980

UCGGACCAGGCUUCAUUCCCC

>gma-miR10196 MIMAT0040963

UGAUUGUGGGAGAGCAUUUCAU

>gma-miR4406 MIMAT0018325

AUUGAUUCUGAGAGAACCGGUGUA

>gma-miR1507c-3p MIMAT0021006

CCUCAUUCCAAACAUCAUCU

>gma-miR10434 MIMAT0041702

CUACUGAUUAUAUAUCUGAUACCAG

>gma-miR156k MIMAT0021635

UUGACAGAAGAGAGUGAGCAC

>gma-miR164g MIMAT0024891

UGGAGAAGCAGGGCACGUGCA

>gma-miR4371b MIMAT0018274

AAGUGAUGACGUGGUAGACGGAGU

>gma-miR1517 MIMAT0007376

AGUCUUGGUCAAUGUCGUUCGAAA

>gma-miR169n-3p MIMAT0032126

UGCCGGCAAGUUUCUCUUGGC

>gma-miR5041-5p MIMAT0021070

UUUCAUCUUCAACUUGCUCAA

>gma-miR4380a MIMAT0018261

CGGAUUGUUGAUCCGUAUGUGCAU

>gma-miR1510b-5p MIMAT0020939

AGGGAUAGGUAAAACAACUACU

>gma-miR162a MIMAT0007353

UCGAUAAACCUCUGCAUCCA

>gma-miR5374-5p MIMAT0021615

UUAUAGUCUGACAUCUGGAAU

>gma-miR9746h MIMAT0036373

AAAGUGUUUGAAUCUCAAUUAGAU

>gma-miR1507b MIMAT0010080

UCUCAUUCCAUACAUCGUCUG

>gma-miR9746d MIMAT0036369

AAAGUGUUUGAAUCUCAAUUAGAU

>gma-miR4367 MIMAT0018266

CUGAACCCUAGCGAAGUAAAUC

>gma-miR171t MIMAT0023236

UUGAGCCGCGUCAAUAUCUCA

>gma-miR4374b MIMAT0018282

UACUUUCAAAGACGUUGUUGAG

>gma-miR10186f MIMAT0041677

UUUUGGGAAUUAAAAUGUAAACUU

>gma-miR4415a-5p MIMAT0018343

AAGUUGUGAUGAGAAUCAAUG

>gma-miR5766 MIMAT0023163

UUGAGGCUGAGAAGAGGCAAG

>gma-miR9731 MIMAT0036350

AUACAUAUCGUGUUGCCAAGC

>gma-miR169s-3p MIMAT0037454

CGGCAAGUAAUCUUUGGCUGC

>gma-miR10428 MIMAT0041690

UGAGGACAAAACACGUUGAAAAGU

>gma-miR9725 MIMAT0036340

UUAAUUUUUUUGGAUCAGCAU

>gma-miR172c MIMAT0011198

GGAAUCUUGAUGAUGCUGCAG

>gma-miR10193b MIMAT0041683

UCUGAAACCGAUGUUAACUACU

>gma-miR5372 MIMAT0021610

UUGUUCGAUAAAACUGUUGUG

>gma-miR1510a-3p MIMAT0007368

UUGUUGUUUUACCUAUUCCACCC

>gma-miR171o-5p MIMAT0032131

AGAUAUUGGUACGGUUCAAUC

>gma-miR166j-3p MIMAT0021648

UCGGACCAGGCUUCAUUCCCG

>gma-miR159e-5p MIMAT0021640

GAGCUCCUUGAAGUCCAAUU

>gma-miR5777 MIMAT0023182

CUAGCAAUAAUGUUGGAUGCAC

>gma-miR1535b MIMAT0021009

CUUGUUUGUGGUGAUGUCUAG

>gma-miR167c MIMAT0007355

UGAAGCUGCCAGCAUGAUCUG

>gma-miR1510b-3p MIMAT0010082

UGUUGUUUUACCUAUUCCACC

>gma-miR10429 MIMAT0041692

UGAGGACAAAACAUGUUGAAAAGU

>gma-miR156f MIMAT0018318

UUGACAGAAGAGAGAGAGCACA

>gma-miR172k MIMAT0023201

UGAAUCUUGAUGAUGCUGCAU

>gma-miR394a-3p MIMAT0018341

AGCUCUGUUGGCUACACUUU

>gma-miR4405 MIMAT0018324

AUUCUAAGACGGUUAUCUGGGACC

>gma-miR1531-3p MIMAT0007393

UCGUCCAUAUGGGAAGACUUGUC

>gma-miR4384 MIMAT0018294

AAUCAGACACUGCAUUCAAAGACG

>gma-miR159a-3p MIMAT0001675

UUUGGAUUGAAGGGAGCUCUA

>gma-miR5780a MIMAT0023185

AUCACUUAGCUGACGGUAGGGAC

>gma-miR399n MIMAT0041708

UGCCAAAGGAGAGCUGCCCUG

>gma-miR2109-3p MIMAT0032101

GGAGGCGUAGAUACUCACACC

>gma-miR10421 MIMAT0041667

CAGAAUGAGACCUUGAGCGUGG

>gma-miR395d MIMAT0023243

UGAAGUGUUUGGGGGAACUUU

>gma-miR164d MIMAT0020977

UGGAGAAGCAGGGCACGUGC

>gma-miR1514b-5p MIMAT0007373

UUCAUUUUUAAAAUAGACAUU

>gma-miR1512b MIMAT0021673

UAACUGGAAAUUCUUAAAGCAU

>gma-miR4387c MIMAT0018254

AGCGUGAUGACGUGACACUCCGUC

>gma-miR319q MIMAT0036398

UGGACUGAAGGGAGCUCCUUC

>gma-miR172i-5p MIMAT0021659

GCAGCAGCAUCAAGAUUCACA

>gma-miR1520e MIMAT0018243

CAAUAAGAACGUGACAUAUGACAG

>gma-miR166q MIMAT0024899

UCGGACCAGGCUUCAUUCCC

>gma-miR390b-5p MIMAT0007361

AAGCUCAGGAGGGAUAGCACC

>gma-miR9744 MIMAT0036364

AAUGGAUAUGAGCUGCAUACA

>gma-miR393k MIMAT0024918

UUCCAAAGGGAUCGCAUUGAUC

>gma-miR482c-3p MIMAT0021002

UUCCCAAUUCCGCCCAUUCCU

>gma-miR482d-5p MIMAT0021671

UAUGGGGGGAUUGGGAAGGAAU

>gma-miR4386 MIMAT0018298

UCGAAGGUUCUGGAGAGGACUGCA

>gma-miR5671b MIMAT0036348

UACCCGAAUUUGCUUCCAUGAU

>gma-miR168b MIMAT0020985

UCGCUUGGUGCAGGUCGGG

>gma-miR399h MIMAT0023251

UGCCAAAGGAGAGUUGCCCUG

>gma-miR4992 MIMAT0021010

AUUCUAAGAUGGUUUUUGUUAG

>gma-miR169d MIMAT0018330

UGAGCCAAGGAUGACUUGCCGGU

>gma-miR319c MIMAT0001691

UUGGACUGAAGGGAGCUCCU

>gma-miR482c-5p MIMAT0021001

AUUUGUGGGAAUGGGCUGAUUGG

>gma-miR396d MIMAT0018262

AAGAAAGCUGUGGGAGAAUAUGGC

>gma-miR166a-5p MIMAT0020920

GGAAUGUUGUCUGGCUCGAGG

>gma-miR319j MIMAT0021664

UUGGACUGAAGGGAGCUCCCU

>gma-miR5763b MIMAT0023160

UGAACUAUACAAAGACGGUUA

>gso-miR1510b MIMAT0016355

UGUUGUUUUACCUAUUCCACC

>gso-miR167a MIMAT0016351

UGAAGCUGCCAGCAUGAUCUG

>gso-miR2218 MIMAT0016361

UUGCCGAUUCCACCCAUUCCUA

>gso-miR1508a MIMAT0016352

UAGAAAGGGAAAUAGCAGUUG

>gso-miR482b MIMAT0016354

UCUUCCCUACACCUCCCAUAC

>gso-miR2109 MIMAT0016363

UGCGAGUGUCUUCGCCUCUGA

>gso-miR3522b MIMAT0016360

UGAGACCAAAUGAGCAGCUGAC

>gso-miR1507b MIMAT0016358

UCUCAUUCCAUACAUCGUCUGA

>gso-miR482a MIMAT0016353

UCUUCCCUACACCUCCCAUAC

>gso-miR1509a MIMAT0016362

UUAAUCAAGGAAAUCACGGUCG

>gso-miR1510a MIMAT0016356

UGUUGUUUUACCUAUUCCACC

>gso-miR1507a MIMAT0016357

UCUCAUUCCAUACAUCGUCUGA

>gso-miR3522a MIMAT0016359

UGAGACCAAAUGAGCAGCUGA

>lja-miR7541 MIMAT0029368

UGCAUUCUCUUUUGGUGGCCC

>lja-miR11078g-3p MIMAT0043981

UAAAAUCUGAUUGGCUGACGGUGU

>lja-miR11068a-5p MIMAT0043703

AUCUCUAAUGCAAGGUUCU

>lja-miR7526d MIMAT0029345

AUCAAGGUAGCUGUAACUUCC

>lja-miR11114-5p MIMAT0043836

UGUUCCUAUAUAUCUGUCCACUU

>lja-miR11069-5p MIMAT0043708

GACACCAAGAGAUCUCGAC

>lja-miR11108n-3p MIMAT0043901

UUGGAUGAUGAAUUAUUUAACC

>lja-miR11087-3p MIMAT0043768

AAAAUAUGAACCCAACCCGGC

>lja-miR7536b MIMAT0029362

UAAGACAUGCUCAAGAGUG

>lja-miR11108e-5p MIMAT0043790

CCCAUCAUCCAAUCACAUUGGAG

>lja-miR7546 MIMAT0029373

UUGGUGACCGACAGGCGCGUGC

>lja-miR7526h MIMAT0029349

AUCAAGGUAGCUGUAACUUCC

>lja-miR171b MIMAT0029315

UGAUUGAGCCGCGUCAAUAUC

>lja-miR167 MIMAT0010087

UGAAGCUGCCAGCAUGAUCUG

>lja-miR11145b-3p MIMAT0043927

GAUAAGUGAGGGACCAAAAGAGC

>lja-miR477-3p MIMAT0043829

UGAAGUUCUUAGGGAGAGUGG

>lja-miR11095-5p MIMAT0043784

UCUAUAAUUGAUUCUGAAGCC

>lja-miR7532b MIMAT0029356

GAAGCUGCCUCUGGUCGUGGU

>lja-miR11108s-5p MIMAT0043875

CAUUGAAGAUAAGUGAGUUGGA

>lja-miR7536a MIMAT0029361

UAAGACAUGCUCAAGAGUG

>lja-miR11134-5p MIMAT0043897

UUUACUCUUGAUGGUCAGAGAA

>lja-miR11179-3p MIMAT0043991

UUUAAAGUUUAGGACCACGUUUGC

>lja-miR7540a MIMAT0029366

UGAUAUGAUAAGUGAUGUGA

>lja-miR11108e-3p MIMAT0043791

CAAUGUGAUUGGAUGAUGGGU

>lja-miR11095-3p MIMAT0043785

AUUCAGAAUCAAUUGUAGUAG

>lja-miR11180-3p MIMAT0043995

UUUUACUUUCGAUUUUGUAACGGC

>lja-miR11075b-3p MIMAT0043758

GUCGUACUAGGAUUGCAAGC

>lja-miR11155d-5p MIMAT0043963

AGUUUGAUUUUAGUUCCUCUAAGG

>lja-miR11082g-3p MIMAT0043741

CACAGAGAGUUGGUUGGACU

>lja-miR11129-3p MIMAT0043884

UAUUAUUCUGUUUAUGGAAGGC

>lja-miR11108o-3p MIMAT0043976

GAUUGGAUGAUGAGUUAUUUAACC

>lja-miR11143-5p MIMAT0043920

CAAACGGACGUUACGGCAACCGC

>lja-miR11108j-3p MIMAT0043839

UGGAUGAUGAGUUAUUUAACC

>lja-miR11085-3p MIMAT0043764

AUUGCAUUUCAUCUUUGAUA

>lja-miR11080-3p MIMAT0043727

AGAUGGUAAAACUUGACACU

>lja-miR477-5p MIMAT0043828

ACUCUCCCUCAAGGGCUUCUGA

>lja-miR1507a MIMAT0029330

UCUUCCAUCCAUACAUCAUCU

>lja-miR11108p-3p MIMAT0043978

GAUUGGAUGAUGAGUUAUUUAACC

>lja-miR11107a-3p MIMAT0043814

UAGUUCAGUGUAUUAUUUGGC

>lja-miR11067-5p MIMAT0043701

ACGGGUUGGCUAAAUGAGC

>lja-miR11155a-5p MIMAT0043949

AAAUUAGUCUCUGAGUUUGUAAGC

>lja-miR171d-5p MIMAT0029318

UUGAGCCGCGCCAAUAUCACU

>lja-miR11108b-3p MIMAT0043823

UCAUGGGUCAUUUAGACAACC

>lja-miR7526f MIMAT0029347

AUCAAGGUAGCUGUAACUUCC

>lja-miR11130-3p MIMAT0043890

UCUGGUGUCUCAUUUAGCCACA

>lja-miR7543 MIMAT0029370

UUAAUGAUACAUGUUUGACU

>lja-miR11111-3p MIMAT0043830

UGAGAUGAUGAAACUUGACAC

>lja-miR11101b-3p MIMAT0043797

CGAGAAACAACAUUUUGGGGG

>lja-miR7533a MIMAT0029357

GAGGGGAUGGAGAGAAGCUGG

>lja-miR11108r-5p MIMAT0043893

AAUAAUCCAUCAUCCAAUCACAUU

>lja-miR11083d-3p MIMAT0043931

GGAUUUAAGCAACUCAUAAGCAA

>lja-miR11128-3p MIMAT0043883

GUUGCUGACCACUCAAAGACAGUA

>lja-miR11078f-3p MIMAT0043835

UGAUUGGUUGACGGUGUAAAA

>lja-miR11082l-3p MIMAT0043820

UCACAGAGAGUUGGUUGGACU

>lja-miR396 MIMAT0010088

UUCCACAGCUUUCUUGAACUG

>lja-miR11117b-3p MIMAT0043849

UUAUGGAGGAAGAGGAAGAAG

>lja-miR11121-3p MIMAT0043860

UUUAAGGGGGGUUUGUCACGC

>lja-miR398-3p MIMAT0043859

UUGUGUUCUCAGGUCACCCCU

>lja-miR11126-5p MIMAT0043877

UGUUCUGGGGACCCUGACCGGG

>lja-miR11108p-5p MIMAT0043977

UUAAAUAACUCAUCAUUCAAUCAC

>lja-miR11152-5p MIMAT0043946

GUUGAAAUUUAGUGUCUAUCUCUC

>lja-miR11108x-5p MIMAT0043915

ACAUUGAAGAUAAGUGAGUUGGA

>lja-miR11101a-3p MIMAT0043796

CGAGAAACAACAUUUUGGGGG

>lja-miR11078c-3p MIMAT0043722

AAAUCUGAUUGGUUGACGGU

>lja-miR11075c-3p MIMAT0043925

GAAGUCGUACUAGGAUUGCAAGC

>lja-miR11130-5p MIMAT0043889

UGGUAAAUGAUACAUCUCCAAG

>lja-miR11072g-3p MIMAT0043733

AGGGACUAAUUUCAGACGGC

>lja-miR11144-5p MIMAT0043923

UUGUCCAUUUUACCCCUGUUAAA

>lja-miR11078b-3p MIMAT0043720

AAAUCUGAUUGGUUGACGGU

>lja-miR7533c-5p MIMAT0043700

AAGUUGGAUGGAGAGGAUC

>lja-miR11098-3p MIMAT0043792

CACUUUGUGGAUCUAAUGUAG

>lja-miR11109-5p MIMAT0043824

UCAUUGAUUGGUUGAAAUUCA

>lja-miR7516-3p MIMAT0029329

AGAGACGUGACUCCCGCUACG

>lja-miR11108q-3p MIMAT0043917

UCAACUCACUUAUCUUAAAUGUGA

>lja-miR172c MIMAT0029321

AGAAUCUUGAUGAUGCUGCA

>lja-miR5287d-3p MIMAT0043993

UUUGCUUAUAAUAGUGACCGGAGG

>lja-miR11153-5p MIMAT0043947

UCCUAUCGGUAAGUUUCUGACAGC

>lja-miR11131-5p MIMAT0043891

UCUUUUGGUUCGAUUCUAAGAC

>lja-miR7527 MIMAT0029350

CAUGGCGUGCAAAACCCCACGC

>lja-miR390a-5p MIMAT0029322

AAGCUCAGGAGGGAUAGCGCC

>lja-miR11108d-5p MIMAT0043885

CAUCAUCCAAUCAUAUUGGAGA

>lja-miR11093-3p MIMAT0043781

AUAAAUCACGACCGUUAGAUC

>lja-miR11115-5p MIMAT0043840

UGGUAUGUAACUUAGAAGUUU

>lja-miR11090d-3p MIMAT0043778

GCGUCGAUAUCUCUUAGUGUUU

>lja-miR11166-3p MIMAT0043969

CAUUGUAUGGAUAGUAGAGGGGCU

>lja-miR11113-5p MIMAT0043833

UGAUAUGUAAUUAGGAAACUA

>lja-miR11107c-3p MIMAT0043817

UAGUUCAGUGUAUUAUUUGGC

>lja-miR11077d-3p MIMAT0043804

AAAUUCACUCCUCAAUUUCCGUG

>lja-miR2111-3p MIMAT0016367

GUCCUUAGGAUGCAGAUUACC

>lja-miR7518 MIMAT0029333

UUGCGCACUGAGCAAGGACAGG

>lja-miR7539 MIMAT0029365

UCGAGAGAGAGAGCGACGAGG

>lja-miR11108h-5p MIMAT0043869

ACAUUGGAGAUAAAUGAGUUGG

>lja-miR11091-3p MIMAT0043779

AGAAGAGAGAAGUUGAGUAGA

>lja-miR7526g MIMAT0029348

AUCAAGGUAGCUGUAACUUCC

>lja-miR172a MIMAT0029319

AGAAUCUUGAUGAUGCUGCAG

>lja-miR11108r-3p MIMAT0043894

UGAUUGGAUGAUGAGUUAUUUA

>lja-miR11118-5p MIMAT0043850

CAUACUAAUAGAAUACGGAAAAU

>lja-miR11097a-3p MIMAT0043789

CACCGUCGGUGCAUCAAACUUU

>lja-miR11092-3p MIMAT0043780

AGUUUGAAACUGACACACGGU

>lja-miR7540b MIMAT0029367

UGAUAUGAUAAGUGAUGUGA

>lja-miR11076-3p MIMAT0043716

UUCCCUUCACUGCCAUCGG

>lja-miR11068e-3p MIMAT0043772

AAGAACCUUGCAUUGAAGAUG

>lja-miR11085-5p MIMAT0043763

UCAAAUAUGAAGAUGCAAUAA

>lja-miR7526j-3p MIMAT0043760

UGAUAGCGGACGUUACAGCU

>lja-miR11127-5p MIMAT0043880

CGGUUCGGACCCGCUGUAGAGC

>lja-miR11105c-5p MIMAT0043809

GUAAGAUUGGAUACUCUCACC

>lja-miR7520 MIMAT0029335

GAGGGGGAAGGUGAUGACAUC

>lja-miR7542 MIMAT0029369

UGCUUGCUUAUAGAUGGUG

>lja-miR11067-3p MIMAT0043702

UCAAGCGAGCCAACCCGUCG

>lja-miR11072c-3p MIMAT0043728

AGGGACUAAUUUCAGACGGC

>lja-miR11172-5p MIMAT0043982

UAGCGGUGAAUAGUGGAGUAGCGG

>lja-miR11128-5p MIMAT0043882

CUGUCUUUGAGUGGUCAGCGGC

>lja-miR11155c-3p MIMAT0043984

UCAGAAGGACUAAAAUCAAACUCG

>lja-miR11068a-3p MIMAT0043704

ACUUUACCAUUGGAGGUGC

>lja-miR11079-3p MIMAT0043725

GUGGUGUCCAAUAUCUCCUUGG

>lja-miR2111-5p MIMAT0016366

UAAUCUGCAUCCUGAGGUUUA

>lja-miR11140-3p MIMAT0043914

UUUCCGAGCUUAACGCAUAAUUGC

>lja-miR7533b MIMAT0029358

GAGGGGAUGGAGAGAAGCUGG

>lja-miR11073-5p MIMAT0043712

GGGUUGGAUGGAUUUGAAC

>lja-miR11132-5p MIMAT0043895

UUAGCAACUUUGAGUACUUUGA

>lja-miR11125-5p MIMAT0043873

AGAGGUCACCUGUUGUCACGGU

>lja-miR11162-3p MIMAT0043960

AGCUAGAUACCUUCGUCACUUUGU

>lja-miR11118-3p MIMAT0043851

UUCCGUAUUCUAUUAAAGGGG

>lja-miR11156-5p MIMAT0043951

AAGAUGGGAUGAUUGAAUUUUGAC

>lja-miR11083c-3p MIMAT0043750

CAUAGGAUUUAAGCAACUCA

>lja-miR11074-5p MIMAT0043713

GGGUUGGAUGGAUUUGAAC

>lja-miR11137-5p MIMAT0043909

UUUUUGGCAUCAUGAUACUGUG

>lja-miR11108w-5p MIMAT0043966

AUUAAGUGAGGUUGUCUGAAUGAC

>lja-miR11072b-5p MIMAT0043752

CGUAAAAUCUAAGUCCUCGUG

>lja-miR11083a-3p MIMAT0043747

CAUAGGAUUUAAGCAACUCA

>lja-miR11075b-5p MIMAT0043757

CUUUGGUCAUUGGUUGAUCUU

>lja-miR7523b MIMAT0029339

ACCACCGGGCUCGAGGAUCAGC

>lja-miR11105d-5p MIMAT0043810

GUAAGAUUGGAUACUCUCACC

>lja-miR171a MIMAT0029314

UGAUUGAGCCGUGCCAAUAUC

>lja-miR11140-5p MIMAT0043913

AAUUAUGCGUUGGCUCAACAAGC

>lja-miR11068f-3p MIMAT0043881

CUAAGAACUUUACCAUUGGAGG

>lja-miR11175-5p MIMAT0043987

UUGAAUUGUUCGAUCAAGAUCGGG

>lja-miR11108v-3p MIMAT0043939

UUGGAUGAUGAGUUAUUUAACCC

>lja-miR11155b-5p MIMAT0043952

AAGCGAGUUUGAUUUUAGUCCCUC

>lja-miR7534 MIMAT0029359

GCAACUUGACUACAGUUUGAC

>lja-miR7529 MIMAT0029352

CCGUAGCAUCAAUUUAUCCGA

>lja-miR11112-3p MIMAT0043831

UGAGAUGAUGAAACUUGACAC

>lja-miR11089c-5p MIMAT0043964

AUGAUUUUAGAGGGACCUAAGACC

>lja-miR11124-5p MIMAT0043865

UCUCGUCUGCAUUUUGUGGAUG

>lja-miR11077a-5p MIMAT0043717

AAAUCGAGGAGUGAAUUUGG

>lja-miR11169-5p MIMAT0043974

GAGUUGGGCUCCUGAAAAGAAGUU

>lja-miR11084-5p MIMAT0043762

UUUCUGGAUCCACGUUUGGC

>lja-miR11174-3p MIMAT0043986

UGUUUCUGUUAGAGUACAGCUGGC

>lja-miR11107b-3p MIMAT0043815

UAGUUCAGUGUAUUAUUUGGC

>lja-miR11123-3p MIMAT0043864

UAUUAGGCACCAUUGUUUCUUG

>lja-miR11159-3p MIMAT0043956

ACUGAAAACUUGUUUGGUAAGACC

>lja-miR11177-5p MIMAT0043989

UUGGAUACACAAUGGAACCACGGU

>lja-miR11108k-3p MIMAT0043871

UCUACUCACUUAUCUUCAAUGUGA

>lja-miR11178-3p MIMAT0043990

UUGGUGGUUAGUAGAAUUAAGACU

>lja-miR11108s-3p MIMAT0043876

UCCAACUCACUGAUCUUCAAUGUG

>lja-miR171d-3p MIMAT0029317

CGAUGUUGGUGAGGUUCAAUC

>lja-miR398-5p MIMAT0043858

GAGUGAGCUUAAGAACACAAGC

>lja-miR11103a-3p MIMAT0043800

CUUGAGAGGAUUUCACUGGAG

>lja-miR11086-3p MIMAT0043766

UAGAUGAAGCAACUCGACACU

>lja-miR11157-3p MIMAT0043954

ACGAGAUAAAAUUAGGGUUGUUUG

>lja-miR7523a MIMAT0029338

ACCACCGGGCUCGAGGAUCAGC

>lja-miR11170-3p MIMAT0043975

GAUGAUAUUAGAGUGAUUCGGAGU

>lja-miR11124-3p MIMAT0043866

UCCCAUGAUGACAGAUGAGAUA

>lja-miR11097e-5p MIMAT0043832

UGAGUUUAAUGGAUAUGCACU

>lja-miR11176-3p MIMAT0043988

UUGAGAUCUGCUGCGGCUAGGACC

>lja-miR11143-3p MIMAT0043921

GGUUGUUGUUGCGUCGUUUUUAAG

>lja-miR11108n-5p MIMAT0043900

UUAAAUAAUCCAUCAUCCAAUC

>lja-miR390b-3p MIMAT0029325

CGCUAUCCAUCCUGAGUUUCA

>lja-miR11148-5p MIMAT0043934

UAAAACUUAUUUACACCGUCGGU

>lja-miR11090e-5p MIMAT0043922

GAACACUUUAUAAGUGAGAGGAC

>lja-miR11107c-5p MIMAT0043816

CCAAAUAAUACACUGAACUAACC

>lja-miR7519 MIMAT0029334

CAAAUUUUCUAAGUGGGCUAGC

>lja-miR11082m-3p MIMAT0043912

AAUCACAGAGAGUUGGUUGGACU

>lja-miR11110b-3p MIMAT0043827

CUCAUUCAAAGACAUAAAAUC

>lja-miR11089a-5p MIMAT0043771

AAGAACCAUGAUUUUAGAGGG

>lja-miR11082i-3p MIMAT0043744

CACAGAGAGUUGGUUGGACU

>lja-miR11154-5p MIMAT0043948

UGAUACACGGUAGAAAAUCACAGC

>lja-miR7530 MIMAT0029353

CCUUCCUCUCUUCACUAUCUUC

>lja-miR11097b-3p MIMAT0043879

CGGUGCAUAGAAAUUAAUCUCU

>lja-miR7545 MIMAT0029372

UUGGGAAAGCUAGAGUGCU

>lja-miR11108j-5p MIMAT0043838

GUUAAAUAACUCAUUAUCCAAUC

>lja-miR7516-5p MIMAT0029328

UAGCGGGUGUCUUCGCCUCUGA

>lja-miR11078a-3p MIMAT0043719

AAAUCUGAUUGGUUGACGGU

>lja-miR11090b-5p MIMAT0043775

ACUUUAUAAGAGAUGUGACUC

>lja-miR11135d-3p MIMAT0043970

CAUUGUAUUGUAUAAGCUUAUACA

>lja-miR7526l-3p MIMAT0043819

UAUCUUGAUAGCGGAAGUUAC

>lja-miR11108f-3p MIMAT0043941

UUGGAUGGUGGGUUAUUUAACCC

>lja-miR11082h-3p MIMAT0043742

CACAGAGAGUUGGUUGGACU

>lja-miR11155c-5p MIMAT0043983

AGUUUGAUUUGAGUCCCUCUGACG

>lja-miR7526k-3p MIMAT0043761

UGAUAGCGGACGUUACAGCU

>lja-miR11090c-5p MIMAT0043776

ACUUUAUAAGAGAUGUGACUC

>lja-miR11117a-5p MIMAT0043847

UCUUCCUCUUCCUCCAUAACU

>lja-miR11120-3p MIMAT0043857

UCACGGAUUCUUCCCGCCAAAA

>lja-miR11083e-3p MIMAT0043932

GGAUUUAAGCAACUCAUAAGCAA

>lja-miR11135b-3p MIMAT0043905

UUGUAUUGUAUAAGCUUAUCCA

>lja-miR11097h-5p MIMAT0043846

UUAAUGGAUAUGCACUGACAG

>lja-miR11136-5p MIMAT0043907

UUUGGAUCUUGAUAAUUGUUGG

>lja-miR11173-3p MIMAT0043985

UCUAUUGGUGGGAUAAAGAAGAGA

>lja-miR11139-3p MIMAT0043911

AAGAUGUGUUGACCUGUUAUAGC

>lja-miR11097c-3p MIMAT0043767

AAAAAACUUUACACCGUCGGU

>lja-miR11168-5p MIMAT0043972

GAACCGUGGCAAUAGAUGACCUCU

>lja-miR11144-3p MIMAT0043924

GAACAGGGAUAAAAUGGACAAUU

>lja-miR11145d-3p MIMAT0043962

AGUGCAAAAUCGUGAUAAGUGAGG

>lja-miR11146-3p MIMAT0043929

GCAACGACAUGUAUAGUUGGAGG

>lja-miR7526i-3p MIMAT0043759

UGAUAGCGGACGUUACAGCU

>lja-miR11082b-3p MIMAT0043736

CACAGAGAGUUGGUUGGACU

>lja-miR11072d-3p MIMAT0043729

AGGGACUAAUUUCAGACGGC

>lja-miR11082c-3p MIMAT0043737

CACAGAGAGUUGGUUGGACU

>lja-miR5287c-3p MIMAT0043979

GCUUAUAUUUGUGACCGGAGGGUG

>lja-miR166-3p MIMAT0043967

AUUUCGGACCAGGCUUCAUUCCCC

>lja-miR11072i-3p MIMAT0043756

GAGGGACCUAGAUCAGACGG

>lja-miR7537 MIMAT0029363

UAGGAAAUACGCCUGCGGUUCC

>lja-miR1120-3p MIMAT0043973

GAGUCUUAUAAUAAGGGACGGAGG

>lja-miR11086-5p MIMAT0043765

UGUCGAGUUGCUCCAUCUAAC

>lja-miR7524 MIMAT0029340

ACCAGUGAGUCAUUGGGCGGA

>lja-miR11120-5p MIMAT0043856

UUGGUUGGAAAGUCCUCGAAC

>lja-miR5287a-3p MIMAT0043862

UUUGCUUAUAAUAGUGACCGG

>lja-miR11105b-5p MIMAT0043808

GUAAGAUUGGAUACUCUCACC

>lja-miR11116-3p MIMAT0043843

UGUUUUUUGCUUAUAUAUGUG

>lja-miR11108q-5p MIMAT0043916

ACAUUGGAGAUAAGUGAGUUGGA

>lja-miR11136-3p MIMAT0043908

GAUAAUUGUUGGGAUUUUGGUU

>lja-miR11106-3p MIMAT0043812

UAAGAACAGGAUAUGAUAAGG

>lja-miR11110b-5p MIMAT0043826

UCUUUAUGUCUUUGAAUGAGG

>lja-miR11088a-5p MIMAT0043769

AAAUGCGGGUUGGGUUGGGCU

>lja-miR11126-3p MIMAT0043878

CGGACAGGGGACCUAGGACAGG

>lja-miR11068b-3p MIMAT0043705

ACUUUACCAUUGGAGGUGC

>lja-miR11135a-3p MIMAT0043904

UUGUAUUGUAUAAGCUUAUCCA

>lja-miR11089b-3p MIMAT0043782

AUGGUUCUUUGGUUUAGGUCC

>lja-miR11097d-5p MIMAT0043874

AGAGUUUAAUGGAUAUGCACUG

>lja-miR172b MIMAT0029320

AGAAUCUUGAUGAUGCUGCA

>lja-miR11113-3p MIMAT0043834

GUUUCCUGAUUACAUAUCAGA

>lja-miR11131-3p MIMAT0043892

CUUAGAUUGAACAAAUGAUC

>lja-miR11077a-3p MIMAT0043718

AAAUUCACUCCUCGAUUUCC

>lja-miR168-5p MIMAT0043887

UCGCUUGGUGCAGGUCGGGAAC

>lja-miR7526e MIMAT0029346

AUCAAGGUAGCUGUAACUUCC

>lja-miR11108t-3p MIMAT0043902

UUGGAUGAUGGGUUAUUUAACC

>lja-miR7517 MIMAT0029332

AUAUGGUAAAGGUUAGGGACC

>lja-miR11108i-3p MIMAT0043935

UAUCAUGGGUCAUUUAGACAACC

>lja-miR11142-3p MIMAT0043919

AUAAGAUCAGGGUCAACAAUCGC

>lja-miR167a MIMAT0029311

UGAAGCUGCCAGCAUGAUCU

>lja-miR7528 MIMAT0029351

CCGAAAUGCUAAUCUGAAGCUU

>lja-miR11103b-5p MIMAT0043853

UUCUCCAGUGGACAUUUGUGG

>lja-miR11088a-3p MIMAT0043770

UCUCAACCCAACCCAUAUUUUGGC

>lja-miR7526a MIMAT0029342

AUCAAGGUAGCUGUAACUUCC

>lja-miR11133-5p MIMAT0043896

UUCCUUAAGGCUAACAGCUUGG

>lja-miR11075a-3p MIMAT0043714

UCGUACUAGGAUUGCAAGC

>lja-miR11102-3p MIMAT0043799

CUAUAUGAAUUUGAUGAAGGC

>lja-miR11077b-5p MIMAT0043798

CGAUGGAUGAAUUAAGACUAC

>lja-miR11083b-3p MIMAT0043748

CAUAGGAUUUAAGCAACUCA

>lja-miR11114-3p MIMAT0043837

UGGACAGAUAAAUAGGAACGG

>lja-miR5287b-3p MIMAT0043938

UUAUAUAUGUGAUCGGAGGGUGU

>lja-miR11072f-3p MIMAT0043732

AGGGACUAAUUUCAGACGGC

>lja-miR11138-3p MIMAT0043910

AAAGCUUAAGCUGUUGGGUAAGG

>lja-miR11088b-5p MIMAT0043867

AAUAUGGGUUGGGUUGAGACUU

>lja-miR11077c-5p MIMAT0043802

GGAAAUCGAGGAGUGAACUUG

>lja-miR11123-5p MIMAT0043863

AGAAAUAGUUGGCCUAAUAAU

>lja-miR11090a-5p MIMAT0043774

ACUUUAUAAGAGAUGUGACUC

>lja-miR11082f-3p MIMAT0043740

CACAGAGAGUUGGUUGGACU

>lja-miR11106-5p MIMAT0043811

UUAUCAUAUCCUGUGUUUGAG

>lja-miR7531 MIMAT0029354

CGUGUUUUCUUUCAUUCCCCA

>lja-miR11149a-5p MIMAT0043936

UUUUACACCAUCAUUCAAUCACAU

>lja-miR11082d-3p MIMAT0043738

CACAGAGAGUUGGUUGGACU

>lja-miR11082k-3p MIMAT0043746

CACAGAGAGUUGGUUGGACU

>lja-miR11147-5p MIMAT0043930

GGAUUCCCGGGAAUGUUUGGGCC

>lja-miR11096b-3p MIMAT0043787

AUUUUGAAGUUUGGGGACGAU

>lja-miR11122-3p MIMAT0043861

UUUAAGUUGGGGAAGUUUGAC

>lja-miR11149a-3p MIMAT0043937

UGAUUGAAUGAUGGUGUAAAACU

>lja-miR7533d-3p MIMAT0043801

GAAGCUGGAUGGAGAGGAUCC

>lja-miR7535 MIMAT0029360

GGGAAAAUGUGGGUGUGGU

>lja-miR11077d-5p MIMAT0043803

GGAAAUCGAGGAGUGAACUUG

>lja-miR11104-3p MIMAT0043806

GGGUUGGGUUGGGAUUUUGAA

>lja-miR390a-3p MIMAT0029323

CGCUAUCCAUCCUGAGUUUCA

>lja-miR11160-5p MIMAT0043958

AGACUGAUUUGAUAGGAGAACUCC

>lja-miR11078d-3p MIMAT0043723

AAAUCUGAUUGGUUGACGGU

>lja-miR390b-5p MIMAT0029324

AAGCUCAGGAGGGAUAGCGCC

>lja-miR7532a MIMAT0029355

GAAGCUGCCUCUGGUCGUGGU

>lja-miR11108k-5p MIMAT0043870

ACAUUGGAGAUAAGUGAGUUGG

>lja-miR11081-3p MIMAT0043734

AGGGCCUGUAAAAUUUUCGC

>lja-miR11090d-5p MIMAT0043777

ACUUUAUAAGAGAUGUGACUC

>lja-miR7522 MIMAT0029337

AACUGCGGACAGGUUUUAUGAC

>lja-miR11072j-3p MIMAT0043842

CACACAACCACAGGGACCAAU

>lja-miR11070-5p MIMAT0043709

GGAGGCCCUGGGCUGUGGG

>lja-miR168-3p MIMAT0043888

UCCCGCCUUGCAUCAACUGAAU

>lja-miR7525 MIMAT0029341

AGGGCGUUUUGGUACAUGACU

>lja-miR11103c-3p MIMAT0043855

ACAAAAUUCCACUGGAGACAA

>lja-miR11103d-5p MIMAT0043942

UUUCUCUAGUGGACAUUUGUGGC

>lja-miR11103e-3p MIMAT0043957

AGAAAUCUACUGGAGAGGAUCUGG

>lja-miR11161-5p MIMAT0043959

AGCCAAAACCAUCGUGGACAGAGG

>lja-miR7538 MIMAT0029364

UCAACGGAGAGCUUGCUGUC

>lja-miR11097f-5p MIMAT0043992

UUUAUACAGACGUCCAAUGAUAGG

>lja-miR11107a-5p MIMAT0043813

CCAAAUAAUACACUGAACUAACC

>lja-miR11068d-3p MIMAT0043754

GAACCUUGCAUUGGAGAUGC

>lja-miR11097g-5p MIMAT0043845

UUAAUGGAUAUGCACUGACAG

>lja-miR7544 MIMAT0029371

UUAGAAAGAAAAUGUUGUUAGC

>lja-miR11072b-3p MIMAT0043753

CGAGGACUUAGAUUUUACGG

>lja-miR7526l-5p MIMAT0043818

UAACUUCCGCUAUCUUAGUAGC

>lja-miR11078c-5p MIMAT0043721

CGUCAACCAAUCAGAUUUCA

>lja-miR11171-5p MIMAT0043980

GCUUGACGAUCGGGCCUGGAGAGC

>lja-miR164-5p MIMAT0043726

ACGCUGGAGAAGCAGGGCAC

>lja-miR397 MIMAT0029326

UAUUGAGUGCAGCGUUGAUGA

>lja-miR11151-3p MIMAT0043945

CGUGAGUAAUUCCUGAGUGUCAGA

>lja-miR11103c-5p MIMAT0043854

UUCUCCAGUGGACAUUUGUGG

>lja-miR11082e-3p MIMAT0043739

CACAGAGAGUUGGUUGGACU

>lja-miR11066-5p MIMAT0043698

UCAAGAUGCUCUAUGUUUG

>lja-miR11100-3p MIMAT0043795

GGGUGCCAAACCCAAUAGGUGGG

>lja-miR11167-5p MIMAT0043971

GAAAAUGGUCUCUAAACUAUGAGC

>lja-miR7526c MIMAT0029344

AUCAAGGUAGCUGUAACUUCC

>lja-miR11079-5p MIMAT0043724

AAGGAGAUCUUGGACACCAC

>lja-miR11165-3p MIMAT0043968

CAUAAAAGGAAACCUCAAUGUAGC

>lja-miR11068c-5p MIMAT0043706

AUCUCUAAUGCAAGGUUCU

>lja-miR11083f-3p MIMAT0043933

GGAUUUAAGCAACUCAUAAGCAA

>lja-miR1507b MIMAT0029331

UCUUCCAUCCAUACAUCAUCU

>lja-miR11117a-3p MIMAT0043848

UUAUGGAGGAAGAGGAAGAAG

>lja-miR11135c-3p MIMAT0043906

UUGUAUUGUAUAAGCUUAUCCA

>lja-miR11099-5p MIMAT0043793

CAUAGGACCUGCAUAGAAGCC

>lja-miR11100-5p MIMAT0043794

CCUAUUGGGUUUCGCACCCCA

>lja-miR11134-3p MIMAT0043898

UUCUGAUCAUCUAGGGUAAACU

>lja-miR11082j-3p MIMAT0043745

CACAGAGAGUUGGUUGGACU

>lja-miR11083c-5p MIMAT0043749

UGGGUUGCUUAAAUACUAUUCU

>lja-miR11082a-3p MIMAT0043735

CACAGAGAGUUGGUUGGACU

>lja-miR11108g-5p MIMAT0043868

ACAUUGGAGAUAAAUGAGUUGG

>lja-miR11108u-3p MIMAT0043903

UUGGAUGAUGGGUUAUUUAACC

>lja-miR11068c-3p MIMAT0043707

ACUUUACCAUUGGAGGUGC

>lja-miR7526b MIMAT0029343

AUCAAGGUAGCUGUAACUUCC

>lja-miR11108d-3p MIMAT0043886

UCCAAUGUGAUUGGAUGAUGGG

>lja-miR167b MIMAT0029312

UGAAGCUGCCAGCAUGAUCU

>lja-miR11108f-5p MIMAT0043940

GUUAAAUAACCCACCAUCCAAUC

>lja-miR171c MIMAT0029316

UGAGCCGAAUCAAUAUCACUC

>lja-miR11119-5p MIMAT0043852

UUCGUAUCGUGACCUGUAUCG

>lja-miR167c MIMAT0029313

UGAAGCUGCCAGCAUGAUCU

>lja-miR11108l-5p MIMAT0043872

ACAUUGGAGAUAAGUGAGUUGG

>lja-miR11158-3p MIMAT0043955

ACUCAACCUUAGGCGUGGGACGCA

>lja-miR408 MIMAT0029327

CAGGGAAGAGGCAGAGCAUGG

>lja-miR11078e-3p MIMAT0043751

CAUCCUUGAAAUCUGAUUGG

>lja-miR11072a-3p MIMAT0043711

GGGAGGGACUAAUUUCAGA

>lja-miR11082i-5p MIMAT0043743

GCCUAUAGCUUUUUGUGUG

>lja-miR11108a-5p MIMAT0043821

UUGUCUAAAUGACCCAUGAUA

>lja-miR11066-3p MIMAT0043699

AACAUAGGGGAUCUUGAAU

>lja-miR11104-5p MIMAT0043805

UCAAAACUCUUAGCAUCAAACGUG

>lja-miR11145c-3p MIMAT0043928

GAUAAGUGAGGGACCAAAAGAGC

>lja-miR11108c-3p MIMAT0043773

AAUGUGAUUGGAUGAUGGGUU

>lja-miR1511-3p MIMAT0043950

AACCAGGCUCUGAUACCAUGAAGC

>lja-miR11150-3p MIMAT0043944

ACUUUCGAGGUGAGUAGUUAACGC

>lja-miR11108m-3p MIMAT0043899

UUGGAUGAUGAAUUAUUUAACC

>lja-miR11072h-3p MIMAT0043731

AGGGACUAAUUUCAGACGGC

>lja-miR11076-5p MIMAT0043715

GAUGGCAGUGAAGGGAACA

>lja-miR11072e-3p MIMAT0043730

AGGGACUAAUUUCAGACGGC

>lja-miR11096a-3p MIMAT0043786

AUUUUGAAGUUUGGGGACGAU

>lja-miR11072j-5p MIMAT0043841

UGGUCCCUGUGGUUGUGUGGC

>lja-miR395-3p MIMAT0043943

ACUGUUGAGGGGAACUCUAGGAGC

>lja-miR11072i-5p MIMAT0043755

ACUCUGGCGAGGAUCCGCUCCA

>lja-miR11097a-5p MIMAT0043788

AUUUUGAUGCACCGACGGUGU

>lja-miR11071-3p MIMAT0043710

GGAUUAAGAGAAGUAGGAC

>lja-miR11094-3p MIMAT0043783

AUUAGGCCUGAUAGUAUAAGC

>lja-miR11110a-5p MIMAT0043825

UCUUUAUGUCUUUGAAUGAGG

>lja-miR11164-5p MIMAT0043965

AUGUUUGGAGACUCAUUUGAAAAC

>lja-miR11149b-3p MIMAT0043953

AAUAGAAGGUUGUGAUUGAAUGAU

>lja-miR11141-3p MIMAT0043918

ACGUCAGCUGGCGGAGCUCCGGC

>lja-miR11155e-5p MIMAT0043994

UUUGUAAGCGAGUUUGAUUUUAGU

>lja-miR11163-5p MIMAT0043961

AGGUCUCUAUUGCCACGGUUAAGC

>lja-miR11097i-3p MIMAT0043844

UUAAUGGAUAUGCACUGACAG

>lja-miR11145a-3p MIMAT0043926

GAUAAGUGAGGGACCAAAAGAGC

>lja-miR11108a-3p MIMAT0043822

UCAUGGGUCAUUUAGACAACC

>lja-miR11105a-5p MIMAT0043807

GUAAGAUUGGAUACUCUCACC

>lja-miR7521 MIMAT0029336

UCAUGGGUGGGGUGUUAAACC

>mtr-miR5758 MIMAT0023144

UAAGUUGGAAGAAUGUAUUUG

>mtr-miR2618b MIMAT0013338

GUGAAUUCAGUUUACGUACGUU

>mtr-miR5261 MIMAT0021261

UCAUUGUAGAUGGCUUUGGCU

>mtr-miR2632b MIMAT0013397

CCUGAAGUUACUAAUCCUUCCA

>mtr-miR395e MIMAT0003858

AUGAAGUGUUUGGGGGAACUC

>mtr-miR2592bk MIMAT0021231

UGGAACAUUGGGAAUGCCGGU

>mtr-miR2592ab MIMAT0021158

CAACAGGACUCAAGCAUUUCGC

>mtr-miR5554a-3p MIMAT0022200

ACCAUCGUUGCAGAUGCUCAUC

>mtr-miR2086-5p MIMAT0010026

CCAGUUCUGCGUUCAUGUCCC

>mtr-miR399s-5p MIMAT0030003

GGGUGAGUUCUCCAUUGGCAGG

>mtr-miR159b MIMAT0021267

AUUGGAGUGAAGGGAGCUCCA

>mtr-miR5223 MIMAT0021180

CGUGGAAUUUACUUGAAGAUGC

>mtr-miR2677 MIMAT0013505

UUUAUUGAUAUUGCUAAUAGAU

>mtr-miR5272a MIMAT0021302

GAAUUGAUUUAUGUUUGGAUACAC

>mtr-miR168c-5p MIMAT0022235

UCGCUUGGUGCAGGUCGGGAA

>mtr-miR2609b MIMAT0013316

UGGAAGUAAUAGGUUCUCACU

>mtr-miR5281b MIMAT0021321

UCUUAUAAAUAGGACCGGAGGGAG

>mtr-miR2586b MIMAT0021172

CGGUGUCGUAUCGGUGUUGGAC

>mtr-miR2111d-3p MIMAT0026926

AUCCUUAGAAUGCAGAUUAUC

>mtr-miR5236a MIMAT0021222

UGAAUUUCGGGCAGAUUUGGU

>mtr-miR2111m-5p MIMAT0013297

UAAUCUGCAUCCUGAGGUUUA

>mtr-miR5037a MIMAT0023150

AACCCUCAAAGGCUUCCACGG

>mtr-miR5267e MIMAT0021280

AGGCAUUUGCUAGAAUACACCCAC

>mtr-miR2592an MIMAT0021170

AAAUGCUUGAGUCCUGUUGUU

>mtr-miR2592bq-5p MIMAT0030011

CGACCAUGACUCAAGUAUUAA

>mtr-miR399r MIMAT0021271

UGCCAAAGAAGAUUUGCCCCG

>mtr-miR2593c MIMAT0013282

UUAAAUGAAUGAACCUAGAAU

>mtr-miR5262 MIMAT0021262

UCUGUCAGUAGACUCAAUUUC

>mtr-miR2592w MIMAT0021153

CAACAGGACUCAAGCAUUUCGC

>mtr-miR2592g MIMAT0013350

AAAUGCUUGAGUCCUGUUGUU

>mtr-miR2592ar MIMAT0021189

AGGCUGGUUUAGAUGAAGGUA

>mtr-miR5274b-3p MIMAT0022210

CAUUUACACUCCGUCAUAUUG

>mtr-miR5257 MIMAT0021257

ACAAGUAGAACCUUUUUUCUG

>mtr-miR164d MIMAT0011103

UGGAGAAGCAGGGCACAUGCU

>mtr-miR7696a-5p MIMAT0029986

UCAAGUUCUCAUAAUUCAAAA

>mtr-miR5749 MIMAT0023129

UUCGGGUUGAUAAUAUUUUCUU

>mtr-miR2664a MIMAT0013471

AAUUGUGGUGGGUUGACAGUC

>mtr-miR1507-5p MIMAT0010030

AGAGUUGUAUGGAACGAAAGAU

>mtr-miR399d MIMAT0001646

UGCCAAAGGAGAGCUGCCCUA

>mtr-miR2629f MIMAT0013368

AGUUUUCCUCGGUAGUUAACU

>mtr-miR172a MIMAT0011086

AGAAUCCUGAUGAUGCUGCAG

>mtr-miR2593d MIMAT0023154

AUACAUCAUUGAUUGAAUGAACCU

>mtr-miR5556-5p MIMAT0022211

UGGAAUUCUUCCGCCAUCCAA

>mtr-miR2592i MIMAT0013273

AAAUGCUUGAGUCCUGUUGUU

>mtr-miR5236c MIMAT0021224

UGAAUUUCGGGCAGAUUUGGU

>mtr-miR399q MIMAT0013477

UGCCAAAGGAGAGCUGCUCUU

>mtr-miR2630q MIMAT0013389

UGGUUUUGGUCCUUGGUAUUU

>mtr-miR156h-3p MIMAT0026729

GCUCUUUAUUCUUCUGUCAUC

>mtr-miR2653a MIMAT0013436

UCACGCUGCUGUGAACAUGAU

>mtr-miR169g MIMAT0011090

CAGCCAAGGAUGACUUGCCGG

>mtr-miR1510b-5p MIMAT0010028

CCAUGGAUCCCUACCAUGUGG

>mtr-miR2594 MIMAT0013283

CCAUGGCCAAGGAUGCCAGAG

>mtr-miR5558-5p MIMAT0022219

UUUUCCAAUUCUAAGUCUAUC

>mtr-miR2654 MIMAT0013439

AUUCAGGGACAAAGUGUGCG

>mtr-miR1509a-3p MIMAT0010035

ACCGGAUUUCCUUGAUUAAAG

>mtr-miR393b-5p MIMAT0011087

UCCAAAGGGAUCGCAUUGAUC

>mtr-miR2663 MIMAT0013470

UUAGAGAGGGCGUUACAAUU

>mtr-miR2652b MIMAT0013425

UAUGCAGGGUGCAUAAGGAUU

>mtr-miR2619b-5p MIMAT0022203

AUAUGUUUUGAUUCUUUGGCA

>mtr-miR5267l MIMAT0021287

AGGCAUUUGCUAGGAUACACCCAC

>mtr-miR1509b MIMAT0013333

UUAAUCUAGGAAAUUACACUCG

>mtr-miR5214-5p MIMAT0027101

UAGCUCUAUUAACAAUUAAAU

>mtr-miR2592h MIMAT0013351

AAAUGCUUGAGUCCUGUUGUU

>mtr-miR5241c MIMAT0021237

UGACUGAAUGGAAGAGUGCAU

>mtr-miR172c-3p MIMAT0021266

AGAAUCUUGAUGAUGCUGCAU

>mtr-miR2629g MIMAT0013369

AGUUUUCCUCGGUAGUUAACU

>mtr-miR5289b MIMAT0021351

CGAGGAAAACUGAAAACUUCGGCA

>mtr-miR5282 MIMAT0021326

GACGGAAUUAGAGAGGGAUUUCAU

>mtr-miR1510a-3p MIMAT0010033

CGGAGGAUUAGGUAAAACAAC

>mtr-miR2680a MIMAT0013510

UCCUCGGUACCUAUGUUGAU

>mtr-miR2592z MIMAT0021156

CAACAGGACUCAAGCAUUUCGC

>mtr-miR2111d-5p MIMAT0013291

UAAUCUGCAUCCUGAGGUUUA

>mtr-miR2655l MIMAT0013451

CGUUUAGGUCCCUUAACUUUA

>mtr-miR397-5p MIMAT0029984

UCAUUGAGUGCAGCGUUGAUG

>mtr-miR171b MIMAT0011066

UGAUUGAGCCGCGUCAAUAUC

>mtr-miR5299 MIMAT0021373

UUCAUUGGUAUUGUAAAGCGACAU

>mtr-miR2605 MIMAT0013309

ACUUAGUUUAUAUGACCUAC

>mtr-miR393b-3p MIMAT0022936

UUUGGGAUCAUGCUAUCCCUU

>mtr-miR166d MIMAT0011068

UCGGGCCAGGCUUCAUCCCCC

>mtr-miR399k MIMAT0011078

UGCCAAAGAAGAUUUGCCCUG

>mtr-miR2592ax MIMAT0021195

AGGCUGGUUUAGAUGAAGGUA

>mtr-miR2656b MIMAT0013456

AAGUUGCAUAAUCGAGUUGG

>mtr-miR2629b MIMAT0013366

AGUUUUCCUCGGUAGUUAACU

>mtr-miR398a-3p MIMAT0011085

UGUGUUCUCAGGUCACCCCUU

>mtr-miR5291c MIMAT0021356

GUUUGAUGGAUGGAUUGGAUGGAU

>mtr-miR5215 MIMAT0021150

AGGAGGAUGAGCUACCUGCUU

>mtr-miR5232 MIMAT0021217

UACAUGUCGCUCUCACCUGAA

>mtr-miR2636 MIMAT0013402

UUUGGUUAGUGUGCUGAAUAU

>mtr-miR2602b MIMAT0013306

UGGCAGUGAUUGCCACGUCAU

>mtr-miR2655d MIMAT0013443

CGUUUAGGUCCCUUAACUUUA

>mtr-miR2111b MIMAT0013358

UAAUCUGCAUCCUGAGGUUUA

>mtr-miR1507-3p MIMAT0010031

CCUCGUUCCAUACAUCAUCUAG

>mtr-miR399a MIMAT0001651

UGCCAAAGGAGAUUUGCCCAG

>mtr-miR2592v MIMAT0021152

CAACAGGACUCAAGCAUUUCGC

>mtr-miR2617a MIMAT0013334

UGUAGUGUAGCAUGCCCGUU

>mtr-miR2647c MIMAT0013419

AUUCACGGGGACGAACCUCCU

>mtr-miR156c-3p MIMAT0026724

UGCUUACUCUCUAUCUGUCACC

>mtr-miR5212-5p MIMAT0021145

UGGAUUUCGUAUUUCUUUGGUA

>mtr-miR2592bo-3p MIMAT0030008

AAAUGCUUGAGUCCUGUUGUU

>mtr-miR2592bj MIMAT0021230

UGGAACAUUGGGAAUGCCGGU

>mtr-miR2632a MIMAT0013396

CCUGAAGUUACUAAUCCUUCCA

>mtr-miR156i-5p MIMAT0011112

UGACAGAAGAGAGUGAGCAC

>mtr-miR2669b MIMAT0029977

AAAGUUCAGUCUUCAUAGUAUC

>mtr-miR5263 MIMAT0021263

UGACUAAAACUAGUAACGGGG

>mtr-miR2644 MIMAT0013412

CACUUCAGAUUGAUGGUGUGU

>mtr-miR2646b MIMAT0013416

CAUGACAUUUAGUGAUGAUGU

>mtr-miR2614 MIMAT0013328

CGGUUCGACUCGUUAGGUUC

>mtr-miR5225c MIMAT0023143

UCGCAGGAGAGAUGACACCUUC

>mtr-miR5283 MIMAT0021327

CGUGCGUAUCGGGAUGUAUCGGAA

>mtr-miR160a MIMAT0001641

UGCCUGGCUCCCUGUAUGCCA

>mtr-miR2590e MIMAT0013265

AUCUAAAGGUGAUUAUUGUGCC

>mtr-miR395h MIMAT0003861

AUGAAGUGUUUGGGGGAACUU

>mtr-miR396a-5p MIMAT0011107

UUCCACAGCUUUCUUGAACUU

>mtr-miR5298c MIMAT0021371

UGAUGGAGAUGAUAUGAAGAUGAA

>mtr-miR2679a MIMAT0013507

CUUUUCACUUUCGAACGGGUG

>mtr-miR2592bm-3p MIMAT0022214

GGAAAACAUGAAUGUCGGGUG

>mtr-miR5295c MIMAT0021365

UCGGCUCUGGGAAUGAAAAGAGGC

>mtr-miR5237 MIMAT0021227

UUCAAAAGAUUUAGUUGGGAU

>mtr-miR7696a-3p MIMAT0029987

UUGAAUUAUGAGAACUUGAAG

>mtr-miR399t-5p MIMAT0030005

GGGUGAGUUCUCCAUUGGCAGGU

>mtr-miR399s-3p MIMAT0030004

UGCCAAAGGAGAUUUGCCCAG

>mtr-miR398a-5p MIMAT0022935

GGAGUGACACUGAGAACACAAG

>mtr-miR5294a MIMAT0021360

GCUAAACGGAAUGAGGGUAGUCAU

>mtr-miR2653b MIMAT0013437

UCACGCUGCUGUGAACAUGAU

>mtr-miR5740 MIMAT0023117

UGAACAGAAAGAACAUUUGGC

>mtr-miR2652i MIMAT0013432

UAUGCAGGGUGCAUAAGGAUU

>mtr-miR168b MIMAT0021270

UCGCUUGGUGCAGGUCGGGAA

>mtr-miR2592ae MIMAT0021161

CAACAGGACUCAAGCAUUUCG

>mtr-miR5563-3p MIMAT0022232

ACUGAGUUGCCUAAUGUCGUU

>mtr-miR166e-5p MIMAT0026726

GGAAUGUUGGCUGGCUCGAGG

>mtr-miR5236b MIMAT0021223

UGAAUUUCGGGCAGAUUUGGU

>mtr-miR399p MIMAT0011120

UGCCAAAGGAGAGUUGCCCUG

>mtr-miR2592bn-3p MIMAT0022218

GGAAAACAUGAAUGUCGGGUG

>mtr-miR399i MIMAT0011076

UGCCAAAGGAGAUUUGCCCUG

>mtr-miR2592au MIMAT0021192

AGGCUGGUUUAGAUGAAGGUA

>mtr-miR2600a MIMAT0013303

ACAUUAGCCAAUCACAAUGCC

>mtr-miR5298d MIMAT0021372

UGGAGAUGAUAUGAAGAUGAAAAA

>mtr-miR2629h MIMAT0021310

GCAGAAGAUCCUCGGCAGUUAACU

>mtr-miR5249 MIMAT0021245

ACUUAGGGGGCAGUUUUGUAG

>mtr-miR2655j MIMAT0013449

CGUUUAGGUCCCUUAACUUUA

>mtr-miR2635 MIMAT0013401

AUUAUUGUCAACGUGACUAG

>mtr-miR319c-3p MIMAT0029981

UUGGACUGAAGGGAGCUCCCA

>mtr-miR5233 MIMAT0021218

GAGGAGGAUGGCCGUCUGGAC

>mtr-miR2655c MIMAT0013442

CGUUUAGGUCCCUUAACUUUA

>mtr-miR5562-5p MIMAT0022229

UGUGGAGUCUUUUGCAUGAAG

>mtr-miR2592be MIMAT0021202

AGGCUGGUUUAGAUGAAGGUA

>mtr-miR7699-3p MIMAT0029999

UGUAAUCAAUGCAUUAAAUGC

>mtr-miR5267k MIMAT0021286

AGGCAUUUGCUAGAAUACACCCAC

>mtr-miR2588b MIMAT0013259

UAACACUGUGCAACUAAGUCC

>mtr-miR2111i MIMAT0013296

UAAUCUGCAUCCUGAGGUUUA

>mtr-miR5284a MIMAT0021335

GAGGGACCAAAAGUGGAAGAAUCU

>mtr-miR2592s-5p MIMAT0026925

ACAACAGGACUCAAGCAUUUC

>mtr-miR2629a MIMAT0013363

AGUUUUCCUCGGUAGUUAACU

>mtr-miR399c MIMAT0001650

UGCCAAAGGAGAUUUGCCCUG

>mtr-miR5241b MIMAT0021236

UGACUGAAUGGAAGAGUGCAU

>mtr-miR2659g MIMAT0029970

CCAUGGGUGCGACUUGGUAAG

>mtr-miR5280 MIMAT0021319

UAAUUAGAAACGGGCCGUGAUGGG

>mtr-miR2592ai MIMAT0021165

CAACAGGACUCAAGCAUUUCGC

>mtr-miR2111g-3p MIMAT0013288

AGCCUCGGAGUGCGGAUUAUC

>mtr-miR2680b MIMAT0013511

UCCUCGGUACCUAUGUUGAU

>mtr-miR2593b MIMAT0013281

UUAAAUGAAUGAACCUAGAAU

>mtr-miR5559-5p MIMAT0022221

UACUUGGUGAAUUGUUGGAUC

>mtr-miR5291b MIMAT0021355

GUUUGAUGGAUGGAUUGGAUGGAU

>mtr-miR2610a MIMAT0013318

AGAUUGAGACUUGUAUGGCUU

>mtr-miR2592y MIMAT0021155

CAACAGGACUCAAGCAUUUCGC

>mtr-miR5759 MIMAT0023146

AAGGGGGUGAAAAGAUUCAAA

>mtr-miR2592bb MIMAT0021199

AGGCUGGUUUAGAUGAAGGUA

>mtr-miR2615a MIMAT0013329

CCUGAUCGCAUUUUAAAAGGC

>mtr-miR2592q-3p MIMAT0013277

AAAUGCUUGAGUCCUGUUGUU

>mtr-miR5271a MIMAT0021297

CGGAUAAUUGUGGUUACUAACGGU

>mtr-miR396a-3p MIMAT0026456

GCUCAAGAAAGCUGUGGGAGA

>mtr-miR5224a MIMAT0021182

UCGAGGACAUGAGGGACGUUAU

>mtr-miR171h MIMAT0021269

CGAGCCGAAUCAAUAUCACUC

>mtr-miR399j MIMAT0011077

CGCCAAAGAAGAUUUGCCCCG

>mtr-miR2670c MIMAT0013480

CAAGAAGGUUGCUCACUAUUU

>mtr-miR5206a MIMAT0021137

AUGGGAUCCUGUUGGUGGGUUAC

>mtr-miR2645 MIMAT0013414

UUUCUAGAGAUGAGCAUAUAU

>mtr-miR171c MIMAT0011083

UGAUUGAGCCGUGCCAAUAUU

>mtr-miR5560-5p MIMAT0022223

CUCAUUCACUCAGCCGGUACA

>mtr-miR408-5p MIMAT0022237

ACAGGGAACAUGCAGAGCAUG

>mtr-miR5295d MIMAT0021366

UCGGCUCUGGGAAUGAAAAGAGGC

>mtr-miR5221 MIMAT0021178

AGGAGAGAUGGUGUUUUGACUU

>mtr-miR5741a MIMAT0023118

UAGGGACUAAAUUGAUGGUUU

>mtr-miR2088-3p MIMAT0010039

UCCAAUGUAAUCUAGGUCUA

>mtr-miR5284h MIMAT0021342

GAGGGAUCAAAAGUGGAGGAAUCU

>mtr-miR2592ad MIMAT0021160

CAACAGGACUCAAGCAUUUCGC

>mtr-miR2592b-5p MIMAT0026924

ACAACAGGACUCAAGCAUUUC

>mtr-miR2676f MIMAT0013503

CAUUGUUUGGAUAAUAAUUUG

>mtr-miR399g MIMAT0011074

UGCCAAAGGAGAUUUGCCCAG

>mtr-miR2592al MIMAT0021168

CAACAGGACUCAAGCAUUUCGC

>mtr-miR2590c MIMAT0013263

AUCUAAAGGUGAUUAUUGUGCC

>mtr-miR2592u MIMAT0021151

CAACAGGACUCAAGCAUUUCGC

>mtr-miR2634 MIMAT0013400

UUUAUUCUCAGUUUGUUGCUC

>mtr-miR5556-3p MIMAT0022212

UGAUGACGGAAGAAAUCCAAA

>mtr-miR5250 MIMAT0021246

UGAGAAUGUUAGAUACGGAAC

>mtr-miR169a MIMAT0001643

CAGCCAAGGAUGACUUGCCGA

>mtr-miR5297 MIMAT0021368

AUCGGGAAGUAUCGGAUAAUUAUU

>mtr-miR2671j MIMAT0013482

UUAAAAGUUUCGUUUCGGUCC

>mtr-miR2655f MIMAT0013445

CGUUUAGGUCCCUUAACUUUA

>mtr-miR2593a MIMAT0013280

UUAAAUGAAUGAACCUAGAAU

>mtr-miR5254 MIMAT0021252

AGGAGGUGGAAGCAUUUGUGA

>mtr-miR5269b MIMAT0021294

AAAGUGGUGGGACAUACAUUGAUU

>mtr-miR2670d MIMAT0013481

CAAGAAGGUUGCUCACUAUUU

>mtr-miR5745a MIMAT0023125

CGUGACAACACAUCAUUGGAUGCA

>mtr-miR5270b MIMAT0021296

GAGGAGGAGUAGUUUUAGGUCAUU

>mtr-miR5286a MIMAT0021346

CAGGACAAACUGGAGGCAAGGGAC

>mtr-miR398c MIMAT0011097

UGUGUUCUCAGGUCGCCCCUG

>mtr-miR319d-3p MIMAT0029983

UUGGACUGAAGGGAGCUCCCU

>mtr-miR2643a MIMAT0013411

UUUGGGAUCAGAAAUUAGAGA

>mtr-miR5747 MIMAT0023127

AAAAGAAUACUCAUACAUAACAUU

>mtr-miR2592at MIMAT0021191

AGGCUGGUUUAGAUGAAGGUA

>mtr-miR2679b MIMAT0013508

CUUUUCACUUUCGAACGGGUG

>mtr-miR5222 MIMAT0021179

UUACAGGAGAAGAAUGUAUGGC

>mtr-miR2630v MIMAT0013394

UGGUUUUGGUCCUUGGUAUUU

>mtr-miR2590j MIMAT0021334

AGAAUGACAUGGCAGAAUAAUCAC

>mtr-miR5559-3p MIMAT0022222

UCUAAUUAUUCACCAAGUAAA

>mtr-miR482-3p MIMAT0027102

CUUACCUACACCUCCCAUGCC

>mtr-miR2599 MIMAT0013302

UGGGUACAAGGAAUCUACUUU

>mtr-miR2670g MIMAT0021210

AGUGGUCUGUUAGGUUGGGGA

>mtr-miR160d MIMAT0011101

UGCCUGGCUCCCUGUAUGCCA

>mtr-miR395b MIMAT0001649

AUGAAGUAUUUGGGGGAACUC

>mtr-miR5272e MIMAT0021306

GAAUUGAUUUAUGUUUGGAUACAC

>mtr-miR2592az MIMAT0021197

AGGCUGGUUUAGAUGAAGGUA

>mtr-miR7696c-3p MIMAT0029991

UUUUGAAUUAUGAGAACUUGA

>mtr-miR172d-3p MIMAT0029979

AGAAUCUUGAUGAUGCUGCA

>mtr-miR2592bl-5p MIMAT0022205

UGGCAAGUUUGAAUUUACCUCA

>mtr-miR2607 MIMAT0013313

AUGUGAUUAUGUGAUAAGUGU

>mtr-miR2630c MIMAT0013372

UGGUUUUGGUCCUUGGUAUUU

>mtr-miR2592bm-5p MIMAT0022213

CUCGGCAUUCAUGUUUUUCCUU

>mtr-miR5288 MIMAT0021349

CAGCAUUGAAGAACAUAGGGAUUA

>mtr-miR5267d MIMAT0021279

AGGCAUUUGCUAGAAUACACCCAC

>mtr-miR2659j MIMAT0029973

CCAUGGGUGCGACUUGGUAAG

>mtr-miR5208a MIMAT0021139

AACAUGGAUGUUGUGAGUUUGUU

>mtr-miR2613 MIMAT0013327

CGGUCGCCGGUGGUCAAUGGU

>mtr-miR395j MIMAT0003863

AUGAAGUGUUUGGGGGAACUC

>mtr-miR2656d MIMAT0029968

AAGUUGCAUAAUCGAGUUGG

>mtr-miR5246 MIMAT0021242

UUGCAGACAGCUUUGAAGGUU

>mtr-miR169d-3p MIMAT0022933

GGCAGGUCAUCCUUCGGCUAUA

>mtr-miR5216a MIMAT0021173

UUAGGAGUGAAAAACGGUGGAA

>mtr-miR2087-3p MIMAT0010037

CUGCAGUCGGUUUCUUACUUC

>mtr-miR2111a-5p MIMAT0022239

UAAUCUGCAUCCUGAGGUUUA

>mtr-miR5563-5p MIMAT0022231

UGAUAUCAGGCAACUCGGUCC

>mtr-miR5287b MIMAT0021348

UGCUUAUAAUAGUGAUCGGAGGGU

>mtr-miR396c MIMAT0023119

AUUCAAGAAGGUCGUGGAAAA

>mtr-miR2111h MIMAT0013295

UAAUCUGCAUCCUGAGGUUUA

>mtr-miR2639 MIMAT0013406

UAGUCGGCUUACGUCACCUUG

>mtr-miR2606c MIMAT0021312

AGUUAAGAACCAUACAAAAAACAC

>mtr-miR5267j MIMAT0021285

AGGCAUUUGCUAGAAUACACCCAC

>mtr-miR168a MIMAT0011089

UUGCUUGGUGCUGGUCGGGAA

>mtr-miR2675 MIMAT0013496

CGAGGCAUAUUUGCAGGGAUU

>mtr-miR169d-5p MIMAT0011064

AAGCCAAGGAUGACUUGCCGG

>mtr-miR5743a MIMAT0023121

UGAGAACUGUUUUCCGCACCUU

>mtr-miR2199 MIMAT0011319

UGAUACACUAGCACGGAUCAC

>mtr-miR5756 MIMAT0023140

CCGACCGGAUUCUCAGACGGG

>mtr-miR2625 MIMAT0013346

CCAUCGUGCCACGUUACGAUCC

>mtr-miR2630b MIMAT0013371

UGGUUUUGGUCCUUGGUAUUU

>mtr-miR2592bp-3p MIMAT0030010

CGGCAGAACUCCAGCAUUCGA

>mtr-miR159a MIMAT0011166

UUUGGAUUGAAGGGAGCUCUA

>mtr-miR2630o MIMAT0013387

UGGUUUUGGUCCUUGGUAUUU

>mtr-miR399o MIMAT0011096

UGCCAAAGGAGAGCUGCCCUG

>mtr-miR2592n MIMAT0013355

AAAUGCUUGAGUCCUGUUGUU

>mtr-miR5256 MIMAT0021254

UAAUGGAUUAUGUAAGAUUAA

>mtr-miR2653c MIMAT0013438

UCACGCUGCUGUGAACAUGAU

>mtr-miR171e-3p MIMAT0011106

AGAUUGAGCCGCGCCAAUAUC

>mtr-miR5230 MIMAT0021215

CAAAUCUUGAAUCGAUUGGCA

>mtr-miR5270a MIMAT0021295

GAGGAGGAGUAGUUUUAGGUCAUU

>mtr-miR2630j MIMAT0013382

UGGUUUUGGUCCUUGGUAUUU

>mtr-miR5272d MIMAT0021305

GAAUUGAUUUAUGUUUGGAUACAC

>mtr-miR2671i MIMAT0013491

UUAAAAGUUUCGUUUCGGUCC

>mtr-miR5269a MIMAT0021293

AAAGUGGUGGAACAUACAUUGAUU

>mtr-miR2592bi MIMAT0021229

UGGAACAUUGGGAAUGCCGGU

>mtr-miR399b MIMAT0001645

UGCCAAAGGAGAGCUGCCCUG

>mtr-miR2587e MIMAT0013256

UUGACCGUUCAUAUGAACCCUG

>mtr-miR5244 MIMAT0021240

UAUCUCAUGAAGAUUGUUGGU

>mtr-miR2592ao MIMAT0021171

AAAUGCUUGAGUCCUGUUGUU

>mtr-miR399n MIMAT0011095

UGCCAAAGGAGAGCUGCCCUA

>mtr-miR2592a.2-3p MIMAT0013349

AAAUGCUUGAGUCCUGUUGUU

>mtr-miR2655g MIMAT0013446

CGUUUAGGUCCCUUAACUUUA

>mtr-miR2630w MIMAT0013373

UGGUUUUGGUCCUUGGUAUUU

>mtr-miR169i MIMAT0029963

UGAGCCAAAGAUGACUUGCCGG

>mtr-miR5281e MIMAT0021324

UCUUAUAAAUAGGACCGGAGGGAG

>mtr-miR5225a MIMAT0021183

UCAGUCGCAGGAGAGAUGACAC

>mtr-miR156j MIMAT0021272

UGACAGAAGAGGGUGAGCAC

>mtr-miR2679c MIMAT0013509

CUUUUCACUUUCGAACGGGUG

>mtr-miR2620 MIMAT0013340

UUCUGAUAGACACCGGCUCUGC

>mtr-miR2592x MIMAT0021154

CAACAGGACUCAAGCAUUUCGC

>mtr-miR2601 MIMAT0013304

UAUUUGGUAUCGCUUUGGUCCC

>mtr-miR2592o-3p MIMAT0013275

AAAUGCUUGAGUCCUGUUGUU

>mtr-miR2642 MIMAT0013410

AUGAGUUUCAUCAAAUCAUGU

>mtr-miR2590d MIMAT0013264

AUCUAAAGGUGAUUAUUGUGCC

>mtr-miR5286b MIMAT0021352

ACAAACUGGAGGCAAGGGACAGGA

>mtr-miR396b-3p MIMAT0026732

GUUCAAUAAAGCUGUGGGAAG

>mtr-miR5213-3p MIMAT0021147

CAGAGUGCAGAUACACGCAUC

>mtr-miR5296 MIMAT0021367

AUUUUGUGUGGGUGUAAGAGGUGU

>mtr-miR5757 MIMAT0023141

UAGAGAUUUGUUUAACAGCCA

>mtr-miR390 MIMAT0011072

AAGCUCAGGAGGGAUAGCGCC

>mtr-miR5267c MIMAT0021278

AGGCAUUUGCUAGAAUACACCCAC

>mtr-miR5214-3p MIMAT0021148

UGAUAGAGCUAGACCAUCGGAG

>mtr-miR7699-5p MIMAT0029998

AUUUAAUGCAUUGAUUACACA

>mtr-miR156a MIMAT0001654

UGACAGAAGAGAGAGAGCACA

>mtr-miR7700-5p MIMAT0030000

GUGGAGUGUGGGACAGCUUGC

>mtr-miR5284g MIMAT0021341

GAGGGACCAAAAGUGGAAGAAUCU

>mtr-miR2592aw MIMAT0021194

AGGCUGGUUUAGAUGAAGGUA

>mtr-miR2670e MIMAT0021208

UCUCAACAGGACGGAUCACUA

>mtr-miR2671b MIMAT0013484

UUAAAAGUUUCGUUUCGGUCC

>mtr-miR171a MIMAT0001655

UGAUUGAGUCGUGCCAAUAUC

>mtr-miR5255 MIMAT0021253

UGACUUGAUAGAGGACAUGGG

>mtr-miR5260 MIMAT0021260

UUUGUAUUGUUGACAUGGCUU

>mtr-miR5206b MIMAT0023152

GUGGGAUCCGUUGAUGGGUUAC

>mtr-miR5284b MIMAT0021336

GAGGGAUCAAAAGUGGAAGAAUCU

>mtr-miR2609a MIMAT0013315

UGGAAGUAAUAGGUUCUCACU

>mtr-miR399h MIMAT0011075

UGCCAAAGGAGAUUUGCCCUG

>mtr-miR5272f MIMAT0021307

GAAUUGAUUAUGUUUGGAUACACU

>mtr-miR5287a MIMAT0021347

UGCUUAUAUUAGUGACCGGAGGAU

>mtr-miR2669a MIMAT0013476

AAAGUUCAGUCUUCAUAGUAUC

>mtr-miR2111f MIMAT0021256

UAAUCUGCAUCCUGAGGUUUA

>mtr-miR5247 MIMAT0021243

GCAGGAGCAAGCAUCUGAUGA

>mtr-miR5289a MIMAT0021350

CGAGGAAAACUGAAAACUUCGGCA

>mtr-miR160e MIMAT0011115

UGCCUGGCUCCCUGUAUGCCA

>mtr-miR5293 MIMAT0021359

GAUGAAGAAGUGGAAGGAAGAAGA

>mtr-miR169f MIMAT0011069

AAGCCAAGGAUGACUUGCCUA

>mtr-miR7696b-3p MIMAT0029989

UUGAAUUAUGAGAACUUGAAG

>mtr-miR2660 MIMAT0013466

UAAGACAUCAGCUAUAAGCUA

>mtr-miR5241a MIMAT0021235

UGACUGAAUGGAAGAGUGCAU

>mtr-miR5231 MIMAT0021216

UUAUGCAAGUAGAUAAGCUCA

>mtr-miR2608 MIMAT0013314

GUUGUACAUAUAUCACUACUCU

>mtr-miR7698-5p MIMAT0029996

UUUUCAUCAAAGUUUUCUGGA

>mtr-miR167b-5p MIMAT0022233

UGAAGCUGCCAGCAUGAUCUG

>mtr-miR2626 MIMAT0013347

AACGUCGGGAUUUAGGGUGUU

>mtr-miR2652f MIMAT0013429

UAUGCAGGGUGCAUAAGGAUU

>mtr-miR2633 MIMAT0013399

UGACAUUUUGCUCCAGAUUCA

>mtr-miR2655m MIMAT0013452

CGUUUAGGUCCCUUAACUUUA

>mtr-miR2676a MIMAT0013498

CAUUGUUUGGAUAAUAAUUUG

>mtr-miR397-3p MIMAT0029985

UCUACGCUACACUCAAUUAUG

>mtr-miR7697-5p MIMAT0029994

UCCGAGAUAUUUGAUCGGAGG

>mtr-miR5290 MIMAT0021353

AAUUUGGAGAGAGAUAGACACAUA

>mtr-miR5275 MIMAT0021313

AGCUGGAGUCACAUGCUUGAAUUU

>mtr-miR2670a MIMAT0013478

CAAGAAGGUUGCUCACUAUUU

>mtr-miR2653d MIMAT0029966

UCACGCUGCUGUGAACAUGAU

>mtr-miR395g MIMAT0003860

UUGAAGUGUUUGGGGGAACUC

>mtr-miR5240 MIMAT0021234

UUGAAAAAAUUGUGGAUUUGA

>mtr-miR2630n MIMAT0013386

UGGUUUUGGUCCUUGGUAUUU

>mtr-miR2592q-5p MIMAT0020944

CAACAGGACUCAAGCAUUUCGC

>mtr-miR2652l MIMAT0013435

UAUGCAGGGUGCAUAAGGAUU

>mtr-miR2592c MIMAT0013269

AAAUGCUUGAGUCCUGUUGUU

>mtr-miR5236d MIMAT0021225

UGAAUUUCGGGCAGAUUUGGU

>mtr-miR2655e MIMAT0013444

CGUUUAGGUCCCUUAACUUUA

>mtr-miR5562-3p MIMAT0022230

AUGUGGAGAAGGCUGCAAC

>mtr-miR5743b MIMAT0023122

UGAGAACUGUUUUCCGCACCUU

>mtr-miR5554a-5p MIMAT0022199

UGUGCAUCUUGAACAAUGGUAU

>mtr-miR2618a MIMAT0013337

GUGAAUUCAGUUUACGUACGUU

>mtr-miR2631 MIMAT0013395

UGACACGCCACGUGGCACACU

>mtr-miR5208b MIMAT0021140

AACAUGGAUGUUGUGAGUUUGUU

>mtr-miR5272c MIMAT0021304

GAAUUGAUUUAUGUUUGGAUACAC

>mtr-miR2592bl-3p MIMAT0022206

GAGUAAUUCAAACUUGUUAAA

>mtr-miR169l-3p MIMAT0030002

GGCAAGUUUUUCCUUGGCUAUA

>mtr-miR2587f MIMAT0013257

UUGACCGUUCAUAUGAACCCUG

>mtr-miR7698-3p MIMAT0029997

CAGACAACUUUGAUGAAAAGC

>mtr-miR5274b-5p MIMAT0022209

AUAUGACGGAGUGUAAAUGCC

>mtr-miR7701-3p MIMAT0030016

UUAGGUUCAUUCAAUUAAUGA

>mtr-miR5554c-3p MIMAT0022226

ACCAUCGUUGCAGAUGCUCAUC

>mtr-miR160f MIMAT0021268

GCGUGAAGGGAGUCAAGCAGG

>mtr-miR2592aj MIMAT0021166

CAACAGGACUCAAGCAUUUCGC

>mtr-miR2592bo-5p MIMAT0030007

CGGCCAGGACUCAAGCAUUUCG

>mtr-miR2592av MIMAT0021193

AGGCUGGUUUAGAUGAAGGUA

>mtr-miR172d-5p MIMAT0029978

AGUGGAGCAUCAUCAAGAUUCACA

>mtr-miR2592l MIMAT0013353

AAAUGCUUGAGUCCUGUUGUU

>mtr-miR2629c MIMAT0013367

AGUUUUCCUCGGUAGUUAACU

>mtr-miR5267h MIMAT0021283

AGGCAUUUGCUAGAAUACACCCAC

>mtr-miR319b-5p MIMAT0026727

GAGCUUUCUUUAGUCCACUCA

>mtr-miR2592br-5p MIMAT0030013

GACUAGGACUCAAGUAUUUCG

>mtr-miR5210 MIMAT0021143

UAAAUGUGUUGGAAUUAAGGUU

>mtr-miR5554b-5p MIMAT0022207

UGUGCAUCUUGAACAAUGGUAU

>mtr-miR2592bf MIMAT0021203

AGGCUGGUUUAGAUGAAGGUA

>mtr-miR2671c MIMAT0013485

UUAAAAGUUUCGUUUCGGUCC

>mtr-miR2627 MIMAT0013348

UUUCGGUAGUUAACUGCUGAGG

>mtr-miR5561-5p MIMAT0022227

CAUUUGGAGAGACAUAGACAA

>mtr-miR2597 MIMAT0013287

UUUGGUACUUCGUCGAUUUGA

>mtr-miR2592af MIMAT0021162

CAACAGGACUCAAGCAUUUCGC

>mtr-miR2671h MIMAT0013490

UUAAAAGUUUCGUUUCGGUCC

>mtr-miR319b-3p MIMAT0011082

UUGGACUGAAGGGAGCUCCC

>mtr-miR2658 MIMAT0013459

AUGUGACCUUGUAUAUGAUC

>mtr-miR2600b MIMAT0021328

AAGCAUUGUGGCAUUGUGAUUGGU

>mtr-miR5281f MIMAT0021325

UCUUAUAAAUAGGACCGGAGGGAG

>mtr-miR395i MIMAT0003862

AUGAAGUGUUUGGGGGAACUC

>mtr-miR168c-3p MIMAT0022236

CCCGCCUUGCAUCAACUGAAU

>mtr-miR156d-5p MIMAT0011071

UGACAGAAGAGAGUGAGCAC

>mtr-miR2592d-5p MIMAT0020941

CAACAGGACUCAAGCAUUUCGC

>mtr-miR4414a-3p MIMAT0021207

AUCCAACGAUGCGGGAGCUGC

>mtr-miR5267b MIMAT0021277

AGGCAUUUGCUAGAAUACACCCAC

>mtr-miR2638b MIMAT0013405

AUGAUUAAUAUUUGCAGUGGC

>mtr-miR5258 MIMAT0021258

UCAAGUGACAAGGAAGAUCUU

>mtr-miR2680e MIMAT0013514

UCCUCGGUACCUAUGUUGAU

>mtr-miR5267i MIMAT0021284

AGGCAUUUGCUAGAAUACACCCAC

>mtr-miR166a MIMAT0001642

UCGGACCAGGCUUCAUUCCCC

>mtr-miR2641 MIMAT0013409

GUUUGAUCCUUUACGUUUAU

>mtr-miR2612 MIMAT0013325

UGAUAGUGUCAACUAGUACAG

>mtr-miR2656a MIMAT0013455

AAGUUGCAUAAUCGAGUUGG

>mtr-miR2640 MIMAT0013407

UUCCUUGCCGGAGCUGGACUAC

>mtr-miR2590a MIMAT0013261

AUCUAAAGGUGAUUAUUGUGCC

>mtr-miR2659i MIMAT0029972

CCAUGGGUGCGACUUGGUAAG

>mtr-miR2587c MIMAT0013254

UUGACCGUUCAUAUGAACCCUG

>mtr-miR5274a MIMAT0021311

CGUUCUACAAUAUGACGGAGUGUA

>mtr-miR2671g MIMAT0013489

UUAAAAGUUUCGUUUCGGUCC

>mtr-miR5284c MIMAT0021337

GAGGGACCAAAAGUGGAAGAAUCU

>mtr-miR2087-5p MIMAT0010036

GAAGUAAAGAACCGGCUGCAG

>mtr-miR5226 MIMAT0021184

UUUGUACAACUUGGAGGAUUCA

>mtr-miR393a MIMAT0001647

UCCAAAGGGAUCGCAUUGAUC

>mtr-miR2656e MIMAT0029969

AAGUUGCAUAAUCGAGUUGG

>mtr-miR319a-5p MIMAT0026556

AGAGCUUCCUUCAGUCCACUC

>mtr-miR5741c MIMAT0023139

UAGGGACUAAAUUGAUGGUUU

>mtr-miR2637 MIMAT0013403

AAAUACUUCCUCUGAUCACUG

>mtr-miR156c-5p MIMAT0011070

UGACAGAAGAGAGUGAGCAC

>mtr-miR5252 MIMAT0021248

UGAGAGCUCACUGAAGUCUGC

>mtr-miR2659e MIMAT0013464

CCAUGGGUGCGACUUGGUAAG

>mtr-miR5229a MIMAT0021213

UUAGCAGGAAGAGUGACUAUG

>mtr-miR5760 MIMAT0023149

UGCUUUAAGGAUAUUUGUUAAGGA

>mtr-miR5285b MIMAT0021344

UGGGACUUUGGGUAGAAUUAGGCG

>mtr-miR2670f MIMAT0021209

AGUGGUCUGUUAGGUUGGGGA

>mtr-miR2630e MIMAT0013377

UGGUUUUGGUCCUUGGUAUUU

>mtr-miR5284f MIMAT0021340

GAGGGACCAAAAGUGGAAGAAUCU

>mtr-miR2086-3p MIMAT0010027

GACAUGAAUGCAGAACUGGAA

>mtr-miR395m MIMAT0003866

AUGAAGUGUUUGGGGGAACUC

>mtr-miR2592br-3p MIMAT0030014

AAAUGCUUGAGUCCUGUUGUU

>mtr-miR2610b MIMAT0013319

AGAUUGAGACUUGUAUGGCUU

>mtr-miR2606a MIMAT0013310

UACAAUUCCUUAGGUGCUUUU

>mtr-miR2590g MIMAT0021309

AAAUGAGACUGAAAUCUAAAGGUG

>mtr-miR2676c MIMAT0013500

CAUUGUUUGGAUAAUAAUUUG

>mtr-miR5277 MIMAT0021316

AGGUUGUUUCUUGAAGUGCAAGGC

>mtr-miR5271e MIMAT0021301

CGGAUAAUUGUGGUUACUAACGGU

>mtr-miR2619b-3p MIMAT0022204

CCAAAGAAUCAAUACAUAGGG

>mtr-miR396b-5p MIMAT0011108

UUCCACAGCUUUCUUGAACUG

>mtr-miR2621 MIMAT0013341

AGCUUGGGCUAGGAAUUUGUGC

>mtr-miR2673a MIMAT0013493

CCUCUUCCUCUUCCUCUUCCAC

>mtr-miR2089-5p MIMAT0010040

UUACCUAUUCCACCAAUUCCAU

>mtr-miR2592bp-5p MIMAT0030009

AAAUGCUUGAGUCCUGUUGUU

>mtr-miR399m MIMAT0011094

UGCCAAAGGAGAGCUGCCCUA

>mtr-miR2118 MIMAT0011318

UUACCGAUUCCACCCAUUCCUA

>mtr-miR169l-5p MIMAT0030001

CAGCCAAGGAUGACUUGCCGG

>mtr-miR2672 MIMAT0013492

UUAAUCGACCAAGUGGGUACUA

>mtr-miR5294c MIMAT0021362

GCUAAACGGAAUGAGGGUAGUCAU

>mtr-miR156g-3p MIMAT0026728

GCUCUCUAGACUUCUGUCAUC

>mtr-miR5219 MIMAT0021177

UCAUGGAAUCUCAGCUGCUGCA

>mtr-miR319a-3p MIMAT0001653

UUGGACUGAAGGGAGCUCCC

>mtr-miR395d MIMAT0003857

AUGAAGUGUUUGGGGGAACUC

>mtr-miR169c MIMAT0011063

CAGCCAAGGGUGAUUUGCCGG

>mtr-miR2585c MIMAT0013322

CAGGAUUAGCGAUUACAGGGAC

>mtr-miR2652m MIMAT0029965

UAUGCAGGGUGCAUAAGGAUU

>mtr-miR2676d MIMAT0013501

CAUUGUUUGGAUAAUAAUUUG

>mtr-miR5557-3p MIMAT0022216

UGCUUCCUUAGUACUUGUUGA

>mtr-miR162 MIMAT0001640

UCGAUAAACCUCUGCAUCCAG

>mtr-miR2665 MIMAT0013472

UGAUUUCAGGUCAAGAAUUGA

>mtr-miR2630t MIMAT0013392

UGGUUUUGGUCCUUGGUAUUU

>mtr-miR2655n MIMAT0013453

CGUUUAGGUCCCUUAACUUUA

>mtr-miR2667a MIMAT0013474

UCCUUGAUCUGACGGCUACC

>mtr-miR395l MIMAT0003865

AUGAAGUGUUUGGGGGAACUC

>mtr-miR5236e MIMAT0021226

UGAAUUUCGGGCAGAUUUGGU

>mtr-miR2592bg MIMAT0021204

AGGCUGGUUUAGAUGAAGGUA

>mtr-miR5265 MIMAT0021274

AAGUGAUGUUGGAAUGGUUA

>mtr-miR166f MIMAT0011100

UCGGACCAGGCUUCAUUCCUC

>mtr-miR5225b MIMAT0023142

UCGCAGGAGAGAUGACACCUUC

>mtr-miR2592b-3p MIMAT0013268

AAAUGCUUGAGUCCUGUUGUU

>mtr-miR2630r MIMAT0013390

UGGUUUUGGUCCUUGGUAUUU

>mtr-miR5037c MIMAT0023153

AACCCUCAAAGGCUUCCACGG

>mtr-miR5294b MIMAT0021361

GCUAAACGGAAUGAGGGUAGUCAU

>mtr-miR5561-3p MIMAT0022228

GUCUAUCUCUCUCUAAAUGGA

>mtr-miR5205d MIMAT0021136

CUUAUAAUUAGGGACGGAGGUAGU

>mtr-miR5752b MIMAT0023134

CAUUGUUUGGUUUAGUACAAA

>mtr-miR5211 MIMAT0021144

UCGCAGGAGUGAUGGGACCGGC

>mtr-miR2600c MIMAT0021329

AAGCAUUGUGGCAUUGUGAUUGGU

>mtr-miR482-5p MIMAT0021232

GGCAUGGGAUAGUAGGGAAGA

>mtr-miR2630d MIMAT0013376

UGGUUUUGGUCCUUGGUAUUU

>mtr-miR5266 MIMAT0021275

CUGGGGGACUGUCUGGGGCG

>mtr-miR156e MIMAT0011081

UUGACAGAAGAUAGAGAGCAC

>mtr-miR2598 MIMAT0013301

CUAAGGGUGAUUAUUCUGCCA

>mtr-miR5284d MIMAT0021338

GAGGGACCAAAAGUGGAAGAAUCU

>mtr-miR166g-3p MIMAT0011104

UCGGACCAGGCUUCAUUCCCC

>mtr-miR2590h MIMAT0021332

AGAAUGACAUGGCAGAAUAAUCAC

>mtr-miR2587a MIMAT0013252

UUGACCGUUCAUAUGAACCCUG

>mtr-miR2611 MIMAT0013320

UAUUUGUCAGUGUUUGAUGAA

>mtr-miR5216b MIMAT0021174

UUAGGAGUGAAAAACGGUGGAA

>mtr-miR5217 MIMAT0021175

AGGUCAUUUUGAACGGUCGGAU

>mtr-miR2591 MIMAT0013267

GGAACUUCUACGGUACACCUGC

>mtr-miR2592j MIMAT0013274

AAAUGCUUGAGUCCUGUUGUU

>mtr-miR2590b MIMAT0013262

AUCUAAAGGUGAUUAUUGUGCC

>mtr-miR2590i MIMAT0021333

AGAAUGACAUGGCAGAAUAAUCAC

>mtr-miR2629d MIMAT0013364

AGUUUUCCUCGGUAGUUAACU

>mtr-miR2676e MIMAT0013502

CAUUGUUUGGAUAAUAAUUUG

>mtr-miR5271b MIMAT0021298

CGGAUAAUUGUGGUUACUAACGGU

>mtr-miR2587d MIMAT0013255

UUGACCGUUCAUAUGAACCCUG

>mtr-miR2602a MIMAT0013305

UGGCAGUGAUUGCCACGUCAU

>mtr-miR2592e-3p MIMAT0013271

AAAUGCUUGAGUCCUGUUGUU

>mtr-miR5748 MIMAT0023128

ACAAAGACAUUGGAAGGCUUA

>mtr-miR5284e MIMAT0021339

GAGGGACCAAAAGUGGAAGAAUCU

>mtr-miR2659h MIMAT0029971

CCAUGGGUGCGACUUGGUAAG

>mtr-miR2652a MIMAT0013424

UAUGCAGGGUGCAUAAGGAUU

>mtr-miR2592t MIMAT0021149

CAACAGGACUCAAGCAUUUCGC

>mtr-miR2592am MIMAT0021169

CAACAGGACUCAAGCAUUUCGC

>mtr-miR319d-5p MIMAT0029982

AGAGCUCUCUUCAGUCCACUC

>mtr-miR2111e-3p MIMAT0026927

AGCCUUGGGAUGCUGAUUAUC

>mtr-miR2592aq MIMAT0021188

AGGCUGGUUUAGAUGAAGGUA

>mtr-miR156i-3p MIMAT0026733

UGCUCACUUCUCUUUCUGUCAUC

>mtr-miR156b-5p MIMAT0011057

UGACAGAAGAGAGUGAGCAC

>mtr-miR5281c MIMAT0021322

UCUUAUAAAUAGGACCGGAGGGAG

>mtr-miR2592m MIMAT0013354

AAAUGCUUGAGUCCUGUUGUU

>mtr-miR7696c-5p MIMAT0029990

AAGUUCUCAUAAUUCAAAAAG

>mtr-miR2592ba MIMAT0021198

AGGCUGGUUUAGAUGAAGGUA

>mtr-miR2643b-5p MIMAT0021249

UCUAAUCUCUGUUCCCAAUUA

>mtr-miR2655i MIMAT0013448

CGUUUAGGUCCCUUAACUUUA

>mtr-miR5741b MIMAT0023123

UAGGGACUAAAUUGAUGGUUU

>mtr-miR5259 MIMAT0021259

CAAGGGGUAUUGCGGAGGAUA

>mtr-miR2676b MIMAT0013499

CAUUGUUUGGAUAAUAAUUUG

>mtr-miR172c-5p MIMAT0021265

GUAGCAUCAUCAAGAUUCACA

>mtr-miR7701-5p MIMAT0030015

AUUAAAUGAAUGAAUCUAAAA

>mtr-miR166g-5p MIMAT0026730

GGAAUGUUGUCUGGCUCGAGG

>mtr-miR5267a MIMAT0021276

AGGCAUUUGCUAGAAUACACCCAC

>mtr-miR319c-5p MIMAT0029980

GGAGUUCCUUGCAGCCCAAAG

>mtr-miR2603 MIMAT0013307

UUUGGUAUUGGUCCCUGCACUU

>mtr-miR2674 MIMAT0013495

CACUCGCUUUGGAAGUCAUGG

>mtr-miR5285c MIMAT0021345

UGGGACUUUGGGUAGAAUUAGGCG

>mtr-miR2622 MIMAT0013342

UUUGUGUGCCAUCGUGAACUUA

>mtr-miR2088-5p MIMAT0010038

AGGCCUAGAUUACAUUGGAC

>mtr-miR5267m MIMAT0021288

AGGCAUUUGCUAGAAUACACCCAC

>mtr-miR2592e-5p MIMAT0020942

CAACAGGACUCAAGCAUUUCGC

>mtr-miR2650 MIMAT0013422

AACUUAAAUAUGUUUUCAGUCC

>mtr-miR2630u MIMAT0013393

UGGUUUUGGUCCUUGGUAUUU

>mtr-miR5213-5p MIMAT0021146

UACGUGUGUCUUCACCUCUGAA

>mtr-miR2659d MIMAT0013463

CCAUGGGUGCGACUUGGUAAG

>mtr-miR5245 MIMAT0021241

CAUCGUAGAACACAGGCAGUA

>mtr-miR169e-5p MIMAT0011065

GAGCCAAGGAUGACUUGCCGG

>mtr-miR2656c MIMAT0029967

AAGUUGCAUAAUCGAGUUGG

>mtr-miR169b MIMAT0011111

CAGCCAAGGAUGACUUGCCGG

>mtr-miR5229b MIMAT0021214

UUAGCAGGAAGAGUGACUAUG

>mtr-miR7696d-5p MIMAT0029992

AAGUUCUCAUAAUUCAAAACG

>mtr-miR2662 MIMAT0013469

GAGUAAAAAUGUGAACCGAAU

>mtr-miR5755 MIMAT0023137

CUUUAACGCGGGAAAGACACA

>mtr-miR169e-3p MIMAT0022934

GGCAGGUCAUCCUUCGGCUAUA

>mtr-miR2111o MIMAT0013300

UAAUCUGCAUCCUGAGGUUUA

>mtr-miR5292a MIMAT0021357

AUUCAGAUGAUAGCAACAAAGAGC

>mtr-miR2630l MIMAT0013384

UGGUUUUGGUCCUUGGUAUUU

>mtr-miR2659k MIMAT0029974

CCAUGGGUGCGACUUGGUAAG

>mtr-miR2111n MIMAT0013299

UAAUCUGCAUCCUGAGGUUUA

>mtr-miR5276 MIMAT0021314

AGGGGGAGCACCUUGCUGGGGCAU

>mtr-miR5253 MIMAT0021251

GAUGAAAAUGAUUAUGUUGGA

>mtr-miR5285a MIMAT0021343

UGGGACUUUGGGUAGAAUUAGGCG

>mtr-miR2585d MIMAT0013323

CAGGAUUAGCGAUUACAGGGAC

>mtr-miR395a MIMAT0001648

AUGAAGUGUUUGGGGGAACUC

>mtr-miR169h MIMAT0011109

UGAGCCAAAGAUGACUUGCCGG

>mtr-miR5745b MIMAT0023145

UUUAAUUUAUAUACAUCGUCA

>mtr-miR399e MIMAT0001652

UGCCAAAGGAGAUUUGCCCAG

>mtr-miR2592r MIMAT0013278

AAAUGCUUGAGUCCUGUUGUU

>mtr-miR5204 MIMAT0021132

GCUGGAAGGUUUUGUAGGAAC

>mtr-miR398b MIMAT0011088

UGUGUUCUCAGGUCGCCCCUG

>mtr-miR156d-3p MIMAT0026725

UGCUCACUCAUCUUUCUGUCAAA

>mtr-miR5751 MIMAT0023131

UUGAUUUGAUCAGAUGGUUUU

>mtr-miR2652j MIMAT0013433

UAUGCAGGGUGCAUAAGGAUU

>mtr-miR2655o MIMAT0013454

CGUUUAGGUCCCUUAACUUUA

>mtr-miR2119 MIMAT0011168

UCAAAGGGAGGUGUGGAGUAG

>mtr-miR2111g-5p MIMAT0013294

UAAUCUGCAUCCUGAGGUUUA

>mtr-miR2628 MIMAT0013362

CAUGAAAGAAUGAUGAGUAA

>mtr-miR2673b MIMAT0013494

CCUCUUCCUCUUCCUCUUCCAC

>mtr-miR5298b MIMAT0021370

UGAUGGAGAUGAUAUGAAGAUGAA

>mtr-miR5271d MIMAT0021300

CGGAUAAUUGUGGUUACUAACGGU

>mtr-miR2666 MIMAT0013473

CGAAAGUGAGGAUAUCAAGGA

>mtr-miR5281d MIMAT0021323

UCUUAUAAAUAGGACCGGAGGGAG

>mtr-miR2646a MIMAT0013415

CAUGACAUUUAGUGAUGAUGU

>mtr-miR2652d MIMAT0013427

UAUGCAGGGUGCAUAAGGAUU

>mtr-miR2589 MIMAT0013260

GGCAUCCACGUGUGCUUCACCG

>mtr-miR2680c MIMAT0013512

UCCUCGGUACCUAUGUUGAU

>mtr-miR172b MIMAT0021264

AGAAUCUUGAUGAUGCUGCAU

>mtr-miR5037b MIMAT0023151

AACCCUCAAAGGCUUCCACGG

>mtr-miR5264 MIMAT0021273

UUGAUCAAGGACUUUGCAUC

>mtr-miR2587b MIMAT0013253

UUGACCGUUCAUAUGAACCCUG

>mtr-miR5205b MIMAT0021134

CUUAUAAUUAGGGACGGAGGGAGU

>mtr-miR2648 MIMAT0013420

UAGCCAAUGGGAAUAACAGAU

>mtr-miR5218 MIMAT0021176

UGAGACUUGGUAGUAAGAUGAU

>mtr-miR5752a MIMAT0023133

CAUUGUUUGGUUUAGUACAAA

>mtr-miR2659a MIMAT0013460

CCAUGGGUGCGACUUGGUAAG

>mtr-miR399l MIMAT0011092

UGCCAAAGGAGAGUUGCCCUG

>mtr-miR395k MIMAT0003869

UUGAAGCGUUUGGGGGAACUC

>mtr-miR5208d MIMAT0021315

CAUAUUAGUCAUAUUUGUAGGCAU

>mtr-miR2619a MIMAT0013339

ACAUAGGAGGCUGUUUUGUAU

>mtr-miR5205c MIMAT0021135

CUUAUAAUUAGGGACGGAGGUAGU

>mtr-miR171d MIMAT0011105

UGAUUGAGCCGUGCCAAUAUC

>mtr-miR2592ah MIMAT0021164

CAACAGGACUCAAGCAUUUCGC

>mtr-miR2647b MIMAT0013418

AUUCACGGGGACGAACCUCCU

>mtr-miR2592ap MIMAT0021187

AGGCUGGUUUAGAUGAAGGUA

>mtr-miR5207 MIMAT0021138

CAUUAAUGUGGGUUUGGACGGUU

>mtr-miR1510a-5p MIMAT0010032

UUGUCUUACCCAUUCCUCCCA

>mtr-miR7696b-5p MIMAT0029988

UCAAGUUCUCAUAAUUCAAAA

>mtr-miR5273 MIMAT0021308

UAGGGGCUGUAGUUUGAGAAGAGG

>mtr-miR5267g MIMAT0021282

AGGCAUUUGCUAGAAUACACCCAC

>mtr-miR2595 MIMAT0013285

UACAUUUUCUUCUUUAUGUCU

>mtr-miR2592bd MIMAT0021201

AGGCUGGUUUAGAUGAAGGUA

>mtr-miR5279 MIMAT0021318

CGGAACCACUCGGAUGACUCGGUU

>mtr-miR2617b MIMAT0013336

UGUAGUGUAGCAUGCCCGUU

>mtr-miR2630i MIMAT0013381

UGGUUUUGGUCCUUGGUAUUU

>mtr-miR2623 MIMAT0013343

UCGGCUGUACUGUCCUUCAUG

>mtr-miR171f MIMAT0011113

UUGAGCCGUGCCAAUAUCACG

>mtr-miR2600d MIMAT0021330

AAGCAUUGUGGCAUUGUGAUUGGU

>mtr-miR408-3p MIMAT0022238

AUGCACUGCCUCUUCCCUGGC

>mtr-miR164b MIMAT0011098

UGGAGAAGCAGGGCACGUGCA

>mtr-miR166c MIMAT0011067

UCGGACCAGGCUUCAUUCCUC

>mtr-miR2651 MIMAT0013423

UUUGAUUGGUAUGCCUGCAUU

>mtr-miR5234 MIMAT0021219

UUUUGUUGUGGAUGGCAGAAG

>mtr-miR5292b MIMAT0021358

GAUUCAGAUGAUAGCAACAAAGAG

>mtr-miR2671e MIMAT0013487

UUAAAAGUUUCGUUUCGGUCC

>mtr-miR5268b MIMAT0021292

CCAGAGUGGAAUGAAGAUAUGGUU

>mtr-miR2659c MIMAT0013462

CCAUGGGUGCGACUUGGUAAG

>mtr-miR5235b MIMAT0021221

AUAAGGUCAAUGAUUGGCGUG

>mtr-miR399f MIMAT0011073

UGCCAAAGGAGAUUUGCCCAG

>mtr-miR5741e MIMAT0023155

UAGGGACUAAAUUGAUGGUUU

>mtr-miR171e-5p MIMAT0026731

CGAUGUUGGUGAGGUUCAAUC

>mtr-miR160c MIMAT0011062

UGCCUGGCUCCCUGAAUGCCA

>mtr-miR171g MIMAT0023132

CGAGCCGAAUCAAUAUCACUC

>mtr-miR2630g MIMAT0013379

UGGUUUUGGUCCUUGGUAUUU

>mtr-miR2655k MIMAT0013450

CGUUUAGGUCCCUUAACUUUA

>mtr-miR5742 MIMAT0023120

CCACAUCAAUGGUCGUUGGAU

>mtr-miR2671a MIMAT0013483

UUAAAAGUUUCGUUUCGGUCC

>mtr-miR5238 MIMAT0021228

UGUAGAAAAAACAAAGGGCAA

>mtr-miR5295a MIMAT0021363

UCGGCUCUGGGAAUGAAAAGAGGC

>mtr-miR2661 MIMAT0013468

UAGGUUUGAGAAAAUGGGCAG

>mtr-miR5228 MIMAT0021212

UCUGGUGUACAACUUGAUGGA

>mtr-miR5298a MIMAT0021369

UGGAUAUGAUAUGAAGAUGAAGAA

>mtr-miR2629e MIMAT0013365

AGUUUUCCUCGGUAGUUAACU

>mtr-miR2592f MIMAT0013272

AAAUGCUUGAGUCCUGUUGUU

>mtr-miR2615b MIMAT0013330

CCUGAUCGCAUUUUAAAAGGC

>mtr-miR2652k MIMAT0013434

UAUGCAGGGUGCAUAAGGAUU

>mtr-miR2111c MIMAT0013289

UAAUCUGCAUCCUGAGGUUUA

>mtr-miR2592o-5p MIMAT0020943

CAACAGGACUCAAGCAUUUCGC

>mtr-miR167a MIMAT0011058

UGAAGCUGCCAGCAUGAUCUA

>mtr-miR5224b MIMAT0021185

CGGAAGAGGAUUGUCGAGGACA

>mtr-miR2671d MIMAT0013486

UUAAAAGUUUCGUUUCGGUCC

>mtr-miR5212-3p MIMAT0027100

CCAAGGAAAUAGAUAUCCAGC

>mtr-miR2586a MIMAT0013251

CGAGGAGUGUCCGUGCUUCAU

>mtr-miR2111k MIMAT0013361

UAAUCUGCAUCCUGAGGUUUA

>mtr-miR2652g MIMAT0013430

UAUGCAGGGUGCAUAAGGAUU

>mtr-miR2592bc MIMAT0021200

AGGCUGGUUUAGAUGAAGGUA

>mtr-miR5555-3p MIMAT0022202

AAGUCGUAUUACACUCUUAGA

>mtr-miR169k MIMAT0013248

UGAGCCAGGAUGGCUUGCCGG

>mtr-miR156h-5p MIMAT0011099

UUGACAGAAGAUAGAGAGCAC

>mtr-miR5754 MIMAT0023136

UAUUGCACUCAUCUUCCAUGGC

>mtr-miR5291a MIMAT0021354

GUUUGAUGGAUGGAUUGGAUGGAU

>mtr-miR5560-3p MIMAT0022224

UGCCGGCUCAAUGAAUGCGGAG

>mtr-miR5267n MIMAT0021289

AGGCAUUUGCUAGAAUACACCCAC

>mtr-miR395o MIMAT0003868

AUGAAGUGUUUGGGGGAACUC

>mtr-miR2592as MIMAT0021190

AGGCUGGUUUAGAUGAAGGUA

>mtr-miR5558-3p MIMAT0022220

UAGAUUUAGAAUUAGAAAAGC

>mtr-miR2585a MIMAT0013249

CAGGAUUAGCGAUUACAGGGAC

>mtr-miR5242 MIMAT0021238

UUGUAGAAACAAGCGAUGUCA

>mtr-miR164a MIMAT0011059

UGGAGAAGCAGGGCACGUGCA

>mtr-miR2652e MIMAT0013428

UAUGCAGGGUGCAUAAGGAUU

>mtr-miR2630f MIMAT0013378

UGGUUUUGGUCCUUGGUAUUU

>mtr-miR2630s MIMAT0013391

UGGUUUUGGUCCUUGGUAUUU

>mtr-miR2592ac MIMAT0021159

CAACAGGACUCAAGCAUUUCGC

>mtr-miR5750 MIMAT0023130

AAGAGAGAUAGAUCAGAAUUGACA

>mtr-miR2587g MIMAT0029964

UUGACCGUUCAUAUGAACCCUG

>mtr-miR5744 MIMAT0023124

UAGGUAUUUUAAGGAGCACGUU

>mtr-miR5251 MIMAT0021247

AGUAGAUCUAGUUGGUGUCUU

>mtr-miR5272b MIMAT0021303

GAAUUGAUUUAUGUUUGGAUACAC

>mtr-miR5281a MIMAT0021320

CUCUUGUAAAUAGGAUCGGAGGGA

>mtr-miR7697-3p MIMAT0029995

UCCGAUCAAAUAGAUCGGAUA

>mtr-miR5248 MIMAT0021244

UUUUUAGUUGGCAUGCAUUCA

>mtr-miR2624 MIMAT0013344

CGAAAGACGAGGUUGCCGGCU

>mtr-miR2616 MIMAT0013332

AUUGGGUUUGGUUCGGGCGGAU

>mtr-miR5555-5p MIMAT0022201

UAAGAGUAUAAUAUGACUUUG

>mtr-miR2089-3p MIMAT0010041

AGGAUUGGUGUAAUAGGUAAAA

>mtr-miR2630m MIMAT0013385

UGGUUUUGGUCCUUGGUAUUU

>mtr-miR5554b-3p MIMAT0022208

ACCAUCGUUGCAGAUGCUCAUC

>mtr-miR2592a-5p MIMAT0020945

CCCGGCAUUCAUGUUUUCCU

>mtr-miR2592s-3p MIMAT0013279

AAAUGCUUGAGUCAUGUUGUU

>mtr-miR2670b MIMAT0013479

CAAGAAGGUUGCUCACUAUUU

>mtr-miR2647a MIMAT0013417

AUUCACGGGGACGAACCUCCU

>mtr-miR2592d-3p MIMAT0013270

AAAUGCUUGAGUCCUGUUGUU

>mtr-miR1509a-5p MIMAT0010034

UUAAUCUAGGAAAAUACGGUG

>mtr-miR2649 MIMAT0013421

AAAGGUGCCAAUUAUGAGUGU

>mtr-miR2592a-3p MIMAT0020946

GAAAAACAUGAAUGUCGAGCG

>mtr-miR4414a-5p MIMAT0021206

AGCUGCUGACUCGUUGGUUCA

>mtr-miR2630h MIMAT0013380

UGGUUUUGGUCCUUGGUAUUU

>mtr-miR2592bn-5p MIMAT0022217

CUCGGCAUUCAUGUUUUUCCUU

>mtr-miR2111e-5p MIMAT0013292

UAAUCUGCAUCCUGAGGUUUA

>mtr-miR2659f MIMAT0013465

CCAUGGGUGCGACUUGGUAAG

>mtr-miR2643b-3p MIMAT0021250

UUUGGGAUCAGAAAUUAGAGA

>mtr-miR2630x MIMAT0013374

UGGUUUUGGUCCUUGGUAUUU

>mtr-miR399t-3p MIMAT0030006

UGCCAAAGGAGAUUUGCCCAG

>mtr-miR2604 MIMAT0013308

UAAUUUUUAUGUGGGAGUGUU

>mtr-miR530 MIMAT0023156

UGCAUUUGCACCUGCACUUUC

>mtr-miR5268a MIMAT0021291

CCAGAGUGGAAUGAAGAUAUGGUU

>mtr-miR2678 MIMAT0013506

UGAAAUUGUUGCGAGUGUCUU

>mtr-miR4414b MIMAT0021211

UGUGAAUGAUGCGGGAGCUAA

>mtr-miR2617c MIMAT0013335

UGUAGUGUAGCAUGCCCGUU

>mtr-miR5235a MIMAT0021220

AUAAGGUCAAUGAUUGGCGUG

>mtr-miR5205a MIMAT0021133

CAUACAAUUUGGGACGGAGGGAG

>mtr-miR2664b MIMAT0029975

AAUUGUGGUGGGUUGACAGUC

>mtr-miR5267f MIMAT0021281

AGGCAUUUGCUAGAAUACACCCAC

>mtr-miR2111j MIMAT0013360

UAAUCUGCAUCCUGAGGUUUA

>mtr-miR5278 MIMAT0021317

GAAAUUAUCUGCAGGAAAUGUGAA

>mtr-miR2606b MIMAT0013311

UACAAUUCCUUAGGUGCUUUU

>mtr-miR2655b MIMAT0013441

CGUUUUGGUCCCUUAACUUUA

>mtr-miR2630p MIMAT0013388

UGGUUUUGGUCCUUGGUAUUU

>mtr-miR2630k MIMAT0013383

UGGUUUUGGUCCUUGGUAUUU

>mtr-miR2592p MIMAT0013276

AAAUGCUUGAGUCCUGUUGUU

>mtr-miR2111m-3p MIMAT0026928

GUCCUCGGGAUACAGAUUACU

>mtr-miR2671f MIMAT0013488

UUAAAAGUUUCGUUUCGGUCC

>mtr-miR2592k MIMAT0013352

AAAUGCUUGAGUCCUGUUGUU

>mtr-miR2585b MIMAT0013250

CAGGAUUAGCGAUUACAGGGAC

>mtr-miR2590f MIMAT0013266

AUCUAAAGGUGAUUAUUGUGCC

>mtr-miR5243 MIMAT0021239

UGGGCAGAGAAAUCGUGAGGC

>mtr-miR5208c MIMAT0021141

AACAUGGAUGUUGUGAGUUUGUU

>mtr-miR5741d MIMAT0023147

UAGGGACUAAAUUGAUGGUUU

>mtr-miR2592ay MIMAT0021196

AGGCUGGUUUAGAUGAAGGUA

>mtr-miR164c MIMAT0011102

UGGAGAAGCAGGGCACGUGCA

>mtr-miR2659b MIMAT0013461

CCAUGGGUGCGACUUGGUAAG

>mtr-miR2593e MIMAT0023148

AUACAUCAUUGAUUGAAUGAACCU

>mtr-miR2630a MIMAT0013370

UGGUUUUGGUCCUUGGUAUUU

>mtr-miR160b MIMAT0011060

UGCCUGGCUCCCUGUAUGCCA

>mtr-miR2680d MIMAT0013513

UCCUCGGUACCUAUGUUGAU

>mtr-miR2630y MIMAT0013375

UGGUUUUGGUCCUUGGUAUUU

>mtr-miR5746 MIMAT0023126

UGGAUUCAACUCUAAGUGUCGGUU

>mtr-miR7696d-3p MIMAT0029993

UUUUGAAUUAUGAGAACUUGA

>mtr-miR166e-3p MIMAT0011080

UCGGACCAGGCUUCAUUCCCC

>mtr-miR2655h MIMAT0013447

CGUUUAGGUCCCUUAACUUUA

>mtr-miR2652h MIMAT0013431

UAUGCAGGGUGCAUAAGGAUU

>mtr-miR2652c MIMAT0013426

UAUGCAGGGUGCAUAAGGAUU

>mtr-miR2588a MIMAT0013258

UAACACUGUGCAACUAAGUCC

>mtr-miR5295b MIMAT0021364

UCGGCUCUGGGAAUGAAAAGAGGC

>mtr-miR2111a-3p MIMAT0022240

AGCCUUGGAAUGCAGAUUAUC

>mtr-miR395n MIMAT0003867

AUGAAGUGUUUGGGGGAACUC

>mtr-miR5227 MIMAT0021186

UGAAGAGAAGAAGAUUGAUGAA

>mtr-miR169j MIMAT0013321

UGAGCCAGGAUGACUUGCCGG

>mtr-miR156g-5p MIMAT0011093

UUGACAGAAGAUAGAGGGCAC

>mtr-miR5557-5p MIMAT0022215

AACAAGUACUAAGGAAGCACA

>mtr-miR5271c MIMAT0021299

CGGAUAAUUGUGGUUACUAACGGU

>mtr-miR395f MIMAT0003859

AUGAAGUGUUUGGGGGAACUC

>mtr-miR2638a MIMAT0013404

AUGAUUAAUAUUUGCAGUGGC

>mtr-miR5209 MIMAT0021142

CGAGGAGGCGGUAUUGUUUGAA

>mtr-miR2615c MIMAT0013331

CCUGAUCGCAUUUUAAAAGGC

>mtr-miR5239 MIMAT0021233

UGGGAGAAAAGAUAGAAUGUG

>mtr-miR5554c-5p MIMAT0022225

UGUGCAUCUUGAACAAUGGUAU

>mtr-miR166b MIMAT0011061

UCGGACCAGGCUUCAUUCCUA

>mtr-miR2592bq-3p MIMAT0030012

AAAUGCUUGAGUCAUGUUGUU

>mtr-miR167b-3p MIMAT0022234

GAUCAUGUUGGAGCUUCACC

>mtr-miR2655a MIMAT0013440

CGUUUAGGUCCCUUAACUUUA

>mtr-miR2657 MIMAT0013457

UGUUAUUUCAUCGAUUUUGUUG

>mtr-miR156f MIMAT0011091

UUGACAGAAGAUAGAGAGCAC

>mtr-miR5267o MIMAT0021290

AGGCAUUUGCUAGGAUACACCCAC

>mtr-miR2596 MIMAT0013286

UCUAUUUCAUUGUUCCACACA

>mtr-miR1510b-3p MIMAT0010029

ACAUGGUCGGUAUCCCUGGAA

>mtr-miR156b-3p MIMAT0026723

UGCUCACUCUCUAUCUGUCACC

>mtr-miR2668 MIMAT0013475

UUCAUCCUUGCAAUUAGGGGUC

>mtr-miR2667b MIMAT0029976

UCCUUGAUCUGACGGCUACC

>mtr-miR5753 MIMAT0023135

AUUUUGAUUGGUGCCAACUAA

>mtr-miR2111l MIMAT0021255

AUCCUUGGAAUGCAGAUUAUC

>mtr-miR395c MIMAT0003856

AUGAAGUGUUUGGGGGAACUC

>mtr-miR2632c MIMAT0013398

CCUGAAGUUACUAAUCCUUCCA

>mtr-miR2600e MIMAT0021331

AAGCAUUGUGGCAUUGUGAUUGGC

>pvu-miR159a.1 MIMAT0015301

UUUGGAUUGAAGGGAGCUCUA

>pvu-miR482-3p MIMAT0011173

UCUUCCCAAUUCCGCCCAUUCC

>pvu-miR482-5p MIMAT0011172

GGAAUGGGCUGAUUGGGAAGCA

>pvu-miR399a MIMAT0011177

UGCCAAAGGAGAGUUGCCCUG

>pvu-miR319c MIMAT0011176

UUGGACUGAAGGGAGCUCCUU

>pvu-miR159a.2 MIMAT0011170

CUUCCAUAUCUGGGGAGCUUC

>pvu-miR166a MIMAT0011175

UCGGACCAGGCUUCAUUCCCC

>pvu-miR2119 MIMAT0011174

UCAAAGGGAGUUGUAGGGGAA

>pvu-miR2118 MIMAT0011169

UUGCCGAUUCCACCCAUUCCUA

>pvu-miR1514a MIMAT0011171

UUCAUUUUGAAAAUAGGCAUUG

>vun-miR319b MIMAT0022754

CUUGGACUGAAGGGAGCUCCU

>vun-miR395 MIMAT0022762

CUGAAGUGUUUGGGGGAACUC

>vun-miR399a MIMAT0022763

UGCCAAAGGAGAGUUGCCCUG

>vun-miR408 MIMAT0022765

AUGCACUGCCUCUUCCCUGGC

>vun-miR172 MIMAT0022760

AGAAUCUUGAUGAUGCUGCAU

>vun-miR156a MIMAT0022752

UGACAGAAGAGAGUGAGCAC

>vun-miR1507a MIMAT0010089

UCUCAUUCCAUACAUCGUCUG

>vun-miR164 MIMAT0022757

UGGAGAAGGGGAGCACGUGCA

>vun-miR2118 MIMAT0022767

UUGCCGAUUCCACCCAUUCCUA

>vun-miR162 MIMAT0022756

UCGAUAAACCUCUGCAUCCAG

>vun-miR169 MIMAT0022759

CAGCCAAGGAUGACUUGCCGG

>vun-miR156b MIMAT0022753

UUGACAGAAGAUAGAGAGCAC

>vun-miR1507b MIMAT0011178

UCUCAUUCCAUACAUCGUCUG

>vun-miR160 MIMAT0022755

UGCCUGGCUCCCUGUAUGCCA

>vun-miR482 MIMAT0022766

UCUUCCCAAUUCCGCCCAUUCCUA

>vun-miR168 MIMAT0022758

UCGCUUGGUGCAGGUCGGGAA

>vun-miR399b MIMAT0022764

UGCCAAAGGAGAAUUGCCCUG

>vun-miR319a MIMAT0022761

UUGGACUGAAGGGAGCUCCCU
