## Supplementary File 2 for "*In silico* Identification and Functional Characterization of Conserved miRNAs in Fibre Biogenesis Crop *Corchorus capsularis*"

**Supplementary File 2:** Retrieved 826 EST sequences of Corchorus capsularis

>GH985272.1 JutEST5278 Jute Plasmid library 20-45 days after sowing Corchorus capsularis cDNA similar to MLP-like protein 34, mRNA sequence

GACCGATGTAGAGATCAAGGCTTCTGCCAAGGATTTCCATGATATGTTCTTTAAAAAACCCCACCACGTC

TCCAAAACCTGCGGTGACAAGATCCACGGATGTGAATTGCATGAGGGTGAATTTGGCAAAGTCGGTTCCA

TTGTCGTCTGGCATTACACGCATGATGGAGTGAGGAAGGTGGCAAAGGAGGTAATTGAAGCAGCAGACGA

GGCGAAGAACTCAATCACTTTTAGTGTGATCGAAGGGGATGTGATGAACGAGTACAAGAACTTCATAATC

CATATTCAAGCTACACCCAAGAGCCATGGCAGCGGCTGTATAGTTCACTGGGTTATGGAATACGAAAAGC

TGCATGAGGGAGTTGCACATCCTGAGACACTGCTGCAATTTGCCGTGGATGTCTCCAAGGACATCGACGC

TCACCTTTGCAAATAATTGAAGATTATGAAATATACCATGTTTCTTATGAGTGTGCTAAGTTATAATAAT

GCAACAACTACTCTACTAAGAAAAAATT

>GH985271.1 JutEST527 Jute Plasmid library 20-45 days after sowing Corchorus capsularis cDNA, mRNA sequence

GCAAGGAAGAAGGAACAGTTAACTTACTTCCTATAGATTGTTGTCGAATGGAGGCTTCTTATGTAATCAT

CAACAGAATTAATCTCTATGAAGAGATGTGAAAAACTTTGAGCCAAAAGCCCCTTCAACGAGTAATTCAA

TCGTCCTGTTCCAGGCATGCACGGGGTAGCATCAAGACGGAAAGCTCTGGCACGGGTCCTTGTACAAAGA

AATGCTGACCAATACCACAGGGCTCCTTGGTAGAGTGTGTGTGAAAAATAGGTATTAAAAGTATATATCT

GGCATTTGTACATCATGCTTAAGAGAGGGTTCTTTGGCAAAGGGTACTCCATTCCATGAACCATTTGCCA

CCAATCTTTCGGTTACTTTTAGTCTGTGTTCTGTGAATGTGTGGTATTTGGAAATGCCTAGTTTTGACTT

TCAAAACATGGGAGGAGTTTGTATTGGATTTGTTTAGAGATTCTGTAGTTCAGTCTATAATTTATTTATA

ATATAATATTTTTAGTTACA

>GH985270.1 JutEST5133 Jute Plasmid library 20-45 days after sowing Corchorus capsularis cDNA similar to unknown, mRNA sequence

GGAATCTCATTCTCAAGTCCTAATTCGATCCTAGTCTTCATCGCAGCTTGGTTCAGGTTTACTATAAACC

CGAGACTTCAAATGGCTGGTGGTAGGGTTGCTCATGCTACCTTGAAAGGACCAAGTGTTGTTAAGGAGAT

AATTATTGGATTATCACTTGGTCTTTTTGCTGGTGGGCTCTGGAAGATGCATCACTGGAATGAGCAAAGG

AAAGTGAGGGCATTCTATGACATGCTTGAGAAAGGTGATATCTCTGTTGTCTTAGAAGAATAAATGTTCA

TGTCTTGTGCAATTGTGGACAGTCTTGGCAGTATTTTGATATTTTGTGTTTCCTTCATTTGGCTCTTAGA

TTCAAAATGAGCAAACTCAGTTTGGAGTTATCAAGAATAATTTTATTGTATTAAGATCCAGATCACATAT

CTGCTTAACAAATCTCAATGGTTTTGTGATATTGTGTCCTTAAAACTTTTTTTCCTTCTGTTTTTTGTGG

CATCTCTTGAGATGTTTTGTAAAATGTGAATGCAGTTCTCTTCCAGACCACTGAGTTGGC

>GH985238.1 JutEST5241 Jute Plasmid library 20-45 days after sowing Corchorus capsularis cDNA similar to ATP synthase protein 9, mRNA sequence

GAAGAAAAATAAGGGTCTAATTTCTGGTTTTTTTTTATATAAAATAAAATGAAAAAAGAAAATCTTAAAG

CGTGAGAAATTCAATTATAAATAATTTAATAAATGCTAGAAGGAGCTAAATTAATTGGTGCAGGAGCAGC

TACAATTGCTTTGGCGGGCGCAGCAGTAGGAATCGGAAATGTATTTAGTTCATTAATACATTCCGTGGCA

AATCCATCCCTAGCAAAACAATTATTCGGATATGCAATTTTAGGCTTTGCTCTAACCGAAGCAATTGCCC

TCTTTGCATTAATGATGGCATTTTTGATCTTATTCGTATTTTGAGAGTATTGGCTCAACATTACA

>GH985237.1 JutEST5240 Jute Plasmid library 20-45 days after sowing Corchorus capsularis cDNA similar to Ribosomal protein S3, mRNA sequence

GAAGTTCCTTTTCCCAAAGAAGTCCCGCTCCTACGGCCAACGAGTCATCTACTTAAAAGAACCCCCCGCC

CCTCCTTGAATTATTTGGTGATGCAATACTTATTAAATAGAAAAAACCAAATTCATTTCGACCCCGTCGT

AGTTGTCAAACATTTCGTGGCACGAGGCGTGTCTGAACCATCTACGATGGGGGGAGCGAATGCATGTGGA

AGAAGCTTAGATAAGAGAATACGTTCTCGCATCGCTTTTTTTGTAGAAAGCTTGACCAGCGAGAAAAAGT

GTTTGGACGAAGCCAAAAAGAGGTTGACCCACTTCATCCGCTTGGCGAATGATCTTCGCTTCGCGGGAAC

AACAAAAACCACCATCTCGCTCTTTCCTTTCTTCGGTGCTACCTTTTTCTTTCTAAGGGATGGGGTTGGG

GGTGAAAAAAAACCTTTTTTTGTGAGAATGACCGAAAAAAACTCCTAGAAAAATTAAGGAAAAAATGTTG

TAACCTCATGGGTAGTAAGGGTAAGGTAATAGAATTGATAGAGAAATTCATAAACTTAGGTGGGGATAGA

AGAAGTGATAAAGGGAATAGAG

>GH985236.1 JutEST5239 Jute Plasmid library 20-45 days after sowing Corchorus capsularis cDNA similar to Putative uncharacterized protein precursor, mRNA sequence

GAGAAAATGTCGTGGGCAGCCAGAATTCGTATGCAGGCCAAAGGGTAGACAATCGCAGCATTAATGGAGA

AAAATAACAAACAGTACTGATTAACATAGTCAAAGAACATATTGATACTTGTTTTCAACGTTTCACTCTT

TCCTTGTTTTATACATATCTTCAAAAAGATAGATTCTCTAGATTTCTGGCTCTCTTGGCGTCAAGAAAGT

GGCAACATCTGTAGCTGGTGTTGATTATCTTTACGA

>GH985235.1 JutEST5238 Jute Plasmid library 20-45 days after sowing Corchorus capsularis cDNA similar to Ole e 1 allergen and extensin family protein, mRNA sequence

GGCTTTGGGTATTGTCTCAAGTTCTCAACTGAAAGAAGAAAATCCCAACATCCTATGGAATCAAAGAACA

AGAATTTGGTTCTGAGTTTTGTTTTCTTAGCTGTCTTCTTAACCCCAGGAGTCAAATCTTGGACTGGAGA

GATTCATGGTAGAGTTGTGTGTGATGTCTGTGCTGATTCTTCAATTGGACCTGAAGATCATGTTTTAGAA

GGTGATGAGGTTGCCGTTCTTTGCATTACAAAATCTGGGGAAGTAGTAAACTATCAAGCTTTTACAAATG

CCAAGGGGATTTACACCGTAGCTGAGACAATGCCTGAGAGTGACCGTTGGGATGCATGCTTAGCACGACC

TATTAGTAGCTTCCATGATCGCTGCAACCATCTTGGGGGAGGGAACACCGGAGTTAAGTTCACCTATAAT

CGTCCCTCGGGTCATTTTCACACTGTCAGACCTTTCGTGTATCGACCCTCCAGCTCTCCTACCTACTGCA

TCTAAAATGTCAGTCAATTTGTAGACTTTTGTTGCCATCATGAATGTCTTAAATCAGAGCTAATGAAAAT

GAATGCTATTTGATGTTGTAATATACATGTGTTCTTATAGTTACAACTTATAATATCCGAAAACGTGTAT

GTTAATTA

>GH985234.1 JutEST5237 Jute Plasmid library 20-45 days after sowing Corchorus capsularis cDNA similar to Peroxidase precursor, mRNA sequence

CCATGGTTTTGGATAACAATTACTACAGGAACATACTGGACAACAAGGGTTTGTTGATTGTGGATCACCA

ACTGGCTTACGACAAAAGGACCAGACCTTATGTGAAGAAAATGGCCAAGAGCCAAGACTATTTCTTCAAG

GAGTTCTCAAGGGCCATTACTAAACTTTCTGAGAACAATCCTCTCACTGGTAGCAAGGGTGAGATCAGGA

AGCAATGCAATCTTGCAAACAAGCTTCACTAAAATGCTATAATCTGAGTGGCAGAATAAAAATGGTTGTT

TGGATCTAGTCTTGAGACAACTTTGTTAATATGCAGTAGTGACCAGTTATATCTGGTTATATGATGACTT

TGCTCATCAAATTCCCCTGTTTTCTTTAGCATATAGCTAGGTTTCTCTTCTCTTGTTTGCCCAGGTTGAT

GAATGATGATGGTCCTGGCTGGTGGTTGGATGTTAGGAT

>GH985233.1 JutEST5236 Jute Plasmid library 20-45 days after sowing Corchorus capsularis cDNA similar to Putative uncharacterized protein, mRNA sequence

ACGCCTCTTCTTCTTTGGCTTTGAACAACTCCAAATCCAACCAAGTTGCCTTGAGAGCGAAATTTAAAAT

TACAATCAAGAGATCTTGAACTTTAGTATCCTAGAAGATGGCGTACGTCGACCATGCCTTCTCCATCTCC

GACGAGGATATCATGATGGAGACATCCTACGCCGTCAACAATCGCCCGCCGTTCAAGGAGATCGGCCTCG

CCATCGCCCTTCTCGTCTTTGGAATCGCTTGGGATCGTCTTGGGCATCTTCATGGCCTACAACAAAGTCG

GCGGTGACAGAGCCCATGGGCTTTTCTTTGCAATATTAGGAGTGCTTTTGTTTATACCGGGCTTCTATTA

CACCAGGATTGCCTATTATGCTTATAAGGGTTATAAAGGCTTCTCTTTCTCCAACATTCCCCCTGTTTAA

CTGATGTTCAACTTTCCTGCAACGCCCGACTATTGTACAGCATGCTTCTTCATATCTTTCCCCATTTTCG

TGTTCCCTTCCCTGTACTTTCTCTTGTTTACTGTATTCAGGATGATTTGTATTCTGTAGTTATATTCTGT

TCCATTGTAGAATTCTTACACATATTGTCTTGTTTCTTGTGGTTGTTTATGGCTTATACACTACTTAATC

TCAATACTTATAC

>GH985232.1 JutEST5235 Jute Plasmid library 20-45 days after sowing Corchorus capsularis cDNA similar to Putative uncharacterized protein, mRNA sequence

GACTCTTAAAAAAGATAATCAATCTAAAAAAGGGAAGAATCAGAGAGCAAAGTAAAGCAAATATAATGCG

TAAATTCATTATACTTTCCGTTTTGGTTCTCTCTGTTTCTTTTTCTGGAAAGTTAGTACAAGCCGGCTAC

ATCACCTTCCCCACTCCCCCAAACCCTCCCCGAAGGCGCCCCTCTATCTCCGGCGAAACCTCCGACTACT

TCTTCGACCCCTCCTGGGCTTCCCGACGCAATCCCCTCCACCTGCCAGTCCCCAACAACAACATCATCCC

CACCCCTGCCAACTACTCCCGACTGCTACGCCGAATCACCGGTTCAATCCCCGGTTGCATCCACTCCGCC

GGCTTCTTCAAGCCCCTCCGGTTCAATCCCCACGCGAGATACTGCTGCACCAACTGGCTCACCCCCGGCT

TGATCACCCACAGTTTCCCCCTGCATTGACACCGACTACTTCTCCTCCACAATGGGCTCCGGTCGGGTCT

CCGCCTATTGCAGAGAGTCCAGTTGGAACTCCTGAGTCGTCTGCTAATATTCCTTCGAGTTCAGCGACGC

CAGCCGAGGGTCCTGCAATTTTTCCGTCCACCAGTAGCCCACCGAATCCCACTCCAGCGGCTTCATCGCC

G

>GH985231.1 JutEST5134 Jute Plasmid library 20-45 days after sowing Corchorus capsularis cDNA similar to HOMEOBOX 7 (ARABIDOPSIS THALIANA), mRNA sequence

ACTCAGCTTAAGTTTGTCTATGAAGACAAAGATTATAGCAGAGATGTTCGACGGAGAGAATAAGCAAGAG

ATGGAGAGCCGTTTTCAAGCAGTAGAATAAGTTACAACAAAAAGAAGAAGAACAAGAACAAACAGAGAAG

GTTTAGCGACGAACAAATCAAGTCGTTGGAATCCATGTTTGAATCAGAATCCAGGCTTGAGCCACGGAAA

AAGTTGCAGGTTGCTAGGGAATTGGGTTTGCAGCCACGGCAAGTTGCGATATGGTTTCAGAACAAAAGAG

CCAGGTGGAAATCAAAGCAGCTTGAACGCGATTACAGTATTTTCAATCCAATCTACAATAATCTTGCTTC

CAAGTTCGAGTCCTTAAAGAAAGAAAAGCAGGCTTTGGTAATTCAGTTGCAGAAGCTGAATGATATGCTG

AAGAAGCCGAAAGAGGAAGGCGAGTGTTGCGGACAAGTGACGGCGACGAATAGCATTGAAGGAGACTCTG

AAAAGGGAGAGGGTGTCAATAAGTCTGATTCCATGGAAAGATCGGAACAT

>GH985230.1 JutEST5233 Jute Plasmid library 20-45 days after sowing Corchorus capsularis cDNA similar to Putative uncharacterized protein, mRNA sequence

GTCTCAGCCCAAGCTACTCTCCAGCCCAAAGAAGACTTTCTTTACTCCCCTGACATGGCGAAAACTTCCC

TTCTTTTCCTGCTACTCTCTCTTCTCTACATCGCTTCCCATGCAACGGCAGATAGCTCCGGTGGCCCAAG

CTCTCGCGCCACAAATTTCATCAAGTCAAAATGCAGCGCCACCACATACCCAGCCTTGTGCTTCCAATCC

CTGGCCCATTACGCCCCTTCAATCCAACAAAGCCCACGCCAGTTGGCTCAGACCGCCCTGTCCGTGAGCC

TAGCTAGGGCTCAGTCGGCCAAGGGATTCGTGGCCAAGATGAAGAAGTTCAAGGGACTTAAGAAGAGGGA

ATACGAAGCCATCAAGGATTGCATCGAGGAAATGGGCGATTCCGTAGACCGGCTCAGTAAATCGGTCCAG

GAGCTCAAGCGCATGGGGTCAACCAAGGGTCAAGACTTTCATTGGCACCTTAGTAATGTGGACACATGGG

TTAGTGCCGCACTGACCGATGAGAACACTTGCCTTGATGGGTTCGCTGGGCGGGCCTTGGATGGTAAAAT

TAAGTCCTCGATTCGAGCCCGTATTTGTAAATGTGGCTCAGGTCACTAGCAATGCTCTTGCTTTGGT

>GH985229.1 JutEST5232 Jute Plasmid library 20-45 days after sowing Corchorus capsularis cDNA similar to Unknown, mRNA sequence

GTCATGGCTGCTTTAATGCTTACGGTGCTCATTCTGGCCATGACTGGCCACTCAAGTGCCACATGGTGTG

TTTGCAAAGATGTGGGTGATGCAAATATGCAGAAAGCTTTGGACTATGCTTGTGGAGCTGGGGCTGACTG

TAACCCTATCCATTCAAACGGGCCTTGTTTCAACCCCAACACCGTTAAGGCTCATTGTAATTACGCCGTC

AACAGCTATTTCCAAAGAAAAGGCCAAGCTCCAGGCTCATGTGAATTTGCTGGCGCTGCCACTGTCACTG

CTTCTGACCCTAGCACTGCTGGCTGTGTTTACCCATCTAGTGCCAGCAGCACTGGAACCGCAACACCAAC

CACTCCGGTCACAACAACCACTCCATCGACCACCACACCCACAACCATGAACCCAGGCACCAACACCCCC

ACAGGCACAACACCCTACAGTTCAACAACACCTGGAGTGCTAGGCGGAGTTGGTACCGGCTTAGGCCCTT

CCGGAGCTGGGATCAACACAGACTACAGTGAAGGATATCGGCTCCAAGGCTTCTCTTTCGTCACAACCCT

TCTGTTTTCAGGCTTGATGCTACTGAGGGCTTGAAAAGAAGATATTGTGGGGT

>GH985228.1 JutEST5230 Jute Plasmid library 20-45 days after sowing Corchorus capsularis cDNA similar to Cyc07, mRNA sequence

GCTTGCAGGCCTAACCTTCCCTTCCCAACATGGCCGTCGGGAAGAACAAGAGGATTTCCAAGGGAAAGAA

GGGAGGAAAGAAGAAGGCGGCGGATCCTTATGCAAAGAAGGATTGGTATGACATCAAGGCACCTTCAATG

TTCAATGAGAGGAACATTGGCAAAACACTTGTTACTAGAACCCAGGGCACTAAGATTGCTTCAGAAGGAC

TTAAGCATAGAGTGTTTGAGGTTTGTTTGGCTGACCTTCAAAAGGATGAAGATCAAGCTTACAGGAAGAT

TCGACTCAGAGCTGAGGATGTCCAAGGGAGGAATGTTCTCACAAACTTCTGGGGAATGGACTTCACCACA

GACAAGCTGCGATCTTTGGTGCGCAAGTGGCAGACATTGATTGAAGCCCATGTTGATGTCAAGACTACTG

ACAATTACAGCTTGAGGTTGTTCTGCATTGCTTATACCAAGAGACGTGCATATCAGGTCAAGCGAACCTG

TTATGCTCAGTCCAGCCAGATCCGTCAGATTCGCCGGAAGATGAGGGAAATCATGGTTAATCAGGCCTCG

TCCTGTGATCTGAAAGAACTTGTTGCGAAGTTCATCCCTGAGGTGATTGGCAGAGAGATTGAGAAAGCTA

>GH985227.1 JutEST5229 Jute Plasmid library 20-45 days after sowing Corchorus capsularis cDNA similar to Putative H-type thioredoxin, mRNA sequence

GGTTGAACTTAAAACAGAGAAGCGTTTCTTTTTGTCTTTCAAAAGAACAAAGTTTTTTCTTTTTAGTGAA

GAAAACAGGCTAAGCAAAAATGGCAGCAGAGGAAGGACAAGTGATAGGTTGCCACACTCGTGAGTCATGG

ACGGAGCAGTTAAACAAGGGCAAAGAATCCAAGAAGCTGGTGGTTGTTGATTTCACCGCTTCATGGTGTG

GACCCTGCCGCTTCATTGCACCCTTCCTGGCTGAACTGGCCAAGAAGCTCCCTAATGTCACGTTTCTTAA

GGTTGATGTCGATGAGTTGAAGGAAGTTGCTGCCGAATATTCTGTGGAGGCTATGCCAACTTTCCTGTTT

CTCAAGGATGGGAAGGAGGTAGACAAGGTGGTGGGAGCAAAGAAAGATGAGCTTCAGAACACTGTGGCTA

AGCATATGGCTACTGCTTCTGCTTGATAATCATATATTTGGAACTACCTTCATATTTGGAGGAATTTATA

TCATATATATAGACCTTAAGTTATGAGTTTGTGCTGCAACTAATAGACAGATCTATGTAACTATGTTAGA

CCTCTCAGTTTGTTCCAGTGTCATTTGAGGATTGTTGACCTTTTCTATGATGAAAACTAAGTGAAGTTT

>GH985226.1 JutEST5228 Jute Plasmid library 20-45 days after sowing Corchorus capsularis cDNA similar to Unknown, mRNA sequence

GTGATCAAGACTAGAGTTATGAACATGAAGGTTGAGCCCGGGCAAAAGCCACCGTATGCCGGCGCGTTGG

ACTGTGCTCTGAAAACGGTGAAGGCAGAAGGGCCTATGGCTTTGTATAAGGGATTTATTCCGACGATTTC

GAGACAGGGACCTTTCACGGTGGTCCTTTTCGTTACGTTGGAACAGGTTCGGAAGCTGCTCAAGGATTTT

TGATCGATGATGATGATGATGAAAAATAATCTTAGGAATTCTATGCTTCAAGTTACGTAATAAGTTTCAG

TTTTATTTTATGTGGTCAAAAAAAAAACTAGAAAAATG

>GH985225.1 JutEST5227 Jute Plasmid library 20-45 days after sowing Corchorus capsularis cDNA similar to Malate dehydrogenase, mRNA sequence

GTTATGTCAAAGAATGTGTCTATTTACAAGTCTCAGGCTTCTGCCTTGGAACAGCATGCAGCTCCTAACT

GCAAGGTTTTGGTTGTTGCTAACCCTGCCAACACCAATGCATTGATCCTGAAGGAATTTGCACCTTCAAT

CCCTGCAAAGAACATTACCGCTTTGACAAGGCTGGACCATAACAGGGCTTTGGGTCAAATCTCAGAGAGA

CTAAATGTCCAAGTTTGTGATGTTAAAAATGTGATTATTTGGGGTAATCATTCATCAACTCAGTATCCTG

ATGTCAATCATGCAACTGTTAAAACTCCATCTGGAGAAAAGCCTGTTCGGGAGCTTGTGAAGGATGATGC

ATGGCTGAATGGAGAATTCATAACCACTGTCCAACAACGGGGTGCTGCAATCATCAAAGCTAGGAAACTC

TCAAGTGCACTATCAGCTGCCAGTTCTGCTTGTGACCACATTCGTGATTGGGTCCTTGGAACCCCAGAGG

GCACATGGGTATCCATGGGAGTTTACTCTGATGGTTCATACAATGTACCTGCTGGTGTCATTTACTCCTT

CCCCTGTAAACTTGCAAGAATGGAGAATGGACAATTGTCCAAGGGGCTGGCTATTGAATGACTTCGTCAA

G

>GH985224.1 JutEST5225 Jute Plasmid library 20-45 days after sowing Corchorus capsularis cDNA similar to 60S ribosomal protein L11, mRNA sequence

GACCCTCCATTTTCATTCTCAGAAGCGGCTGCGGCGGATTCAGCTTCAGCATCCAAAAATGGCGTCGGAG

AAGAAGCTATCAAACCCAATGAGGGAAATCAAGGTCCAGAAGCTTGTCCTTAACATCTCCGTCGGTGAAA

GTGGAGATCGTCTCACCAGGGCTGCCAAGGTGTTAGAGCAACTCAGCGGCCAAACACCGGTCTTCTCTAA

AGCTAGGTACACTGTCCGGTCCTTTGGAATCAGACGTAATGAGAAGATCGCGTGCTATGTCACTGTTAGA

GGGGAGAAGGCAATGCAACTTTTGGAAAGTGGGTTGAAGGTTAAGGAGTATGAGTTGTTGAGAAGGAACT

TTAGTGATACTGGTTGCTTTGGATTTGGTATTCAGGAGCATATTGATCTTGGAATTAAATATGACCCATC

TACTGGTATTTATGGTATGGACTTCTTTGTTGTCCTGGAGCGTGCTGGTTATCGTGTGGGTCGTCGCCGT

AGGTGCAAGTCCCGTGTTGGGATCCAGCACAGGGTTACAAAGGAGGATGCTATGAAATGGTTCCAGGTGA

AATATGAAGGTGTGATCCTCAACAAGTCCCAGCAATTAACTTC

>GH985223.1 JutEST5222 Jute Plasmid library 20-45 days after sowing Corchorus capsularis cDNA similar to Lipid transfer protein 4, mRNA sequence

GATTCAAAAGCACTAATTAGTATGTTTTTCTCTCGCTTAATCTCTATATTAATCGAAATGGCTCCCTATT

AAGCTTGCATGTGTGGCGGTGTTGTGCATGGTTTGGTGGGTGCACCCCTGGCTCAAGGAGCCGTAACCTG

TGGTCAAGTCACAAGCTCCCTCGCGGCCTGCATTGGTTACTTGACAGGGAATGGTCCTTGTGACGTCCTC

CAGGCTGCTGCAATGGGGTGAGATCTCTCAACAGCGCCGCCAAAACCACCCCGGACCGCCAAGCCGCTTG

CAACTGCCTTAAAAGCGCCGCCGGTTCCATTCCTGGCATCAACTTTGGTCTCGCTGGAGGACTCCCTGGC

AAGTGCCGCGTCAGCATTCCTTACAAGATTAGCCCCTCCACTGACTGCAGCAAGGTTAAGTGAAATGCCA

AGATTTTGGAAGGGTTTTATTTTAATTTAAGCTCCAAGGAGATGAAGAGAGTTTACTTTTAGATGGAAAA

ATAAAATGGATGTAAAAGTCCATGTGGGATAGCACTCTTTCGTAGGAGAGTTGTCCTAATTTTAGATTTG

TTATTTGTACTTTTTATCGTGTGTACAAGGTCGCTTTTATATATGCGACTATAACTTTCATATATGAAAA

ATTAAT

>GH985222.1 JutEST5221 Jute Plasmid library 20-45 days after sowing Corchorus capsularis cDNA similar to Elongation factor EF-2, mRNA sequence

GGCGAGGCTATTGATGATGGACGTATTGGTCCAAGAGATGATCCAAAGGTTCGTTCCAAGATCTTGGCTG

AGGAGTTTGGTTGGGACAAGGATCTTGCAAAGAAAATCTGGTGCTTTGGTCCTGAGACCACCGGGCCAAA

CATGGTTGTGGATATGTGTAAGGGAGTTCAGTACCTTAATGAAATCAAGGATTCTGTTGTTGCTGGTTTC

CAGTGGGCTTCAAAGGAAGGTGCGATGTGTGAAGAAAACATGCGAGGTATCTGCTTTGAAGTCTGTGATG

TGGTTCTTCATTCCGATGCCATTCACAGAGGTGGTGGGCAGATCATTCCTACTGCCAGGAGGGTCTTCTA

TGCCTCCCAGCTGACTGCCAAGCCCAGGCTTCTTGAGCCCGTGTACATGGTGGAGATCCAAGCCCCCGAG

CAGGCCCTCGGTGGCATCTACAGTGTGCTTAACCAGAAACGTGGACACGTGTTTGAAGAAATTCAGAGGC

CCGGTACCCCCCTGTACAATATCAAGGCGTATCTTCCTGTTGTCGAGTCTTTTGGATTTTCCGGCACTTT

GAGGGCTGCAACTTCAGGGCAGGCCTTCCCACAATGTGTCTTTGATCATTGGGATATGATGACTCAAG

>GH985221.1 JutEST5220 Jute Plasmid library 20-45 days after sowing Corchorus capsularis cDNA similar to Lipoxygenase LOX2, mRNA sequence

GGGGGGGGGGAAATAGAGTCATCATGGGGTGAAGTTCCAATTATAAAGGCAGCTTTTGAGAAATTCAATG

GGAAATTGAAGGAGCTAGAAGGGATAATTGATGAAAGGAATGCTAATCAACACTTGCAAAATAGAAGTGG

AGCTGGATTGTACCATATGAGCTTTTGAAGCCATTTTCTGAGCCTGGTGTTACAGGAAAAGGAGTTCCAT

ATAGTATTTCCATATGAAATCCACAGACTGCTAATAGATCGTTCTTGATAAGACATTTGAATAAATTAAG

CATAAATTTATTTGAGTGGTTC

>GH985220.1 JutEST5218 Jute Plasmid library 20-45 days after sowing Corchorus capsularis cDNA similar to Putative uncharacterized protein, mRNA sequence

GATTGAAACATTTAACAATGGCGTCAGAGAAAGACCACAAGCCAACTTCGTCCGACCTTTTTTTCCAGTG

CTAAGGTGGTTGCGGAGGCCGCAAAATCAGCATTCGGCGGCAATGAAGGCGGCGACAAGTTCGATAAGGC

GAAGGTTGCCGAAGCCGCTTCTGATCTTTTAGATGCAGCCAAAGACTATGGCAAGCTTGAGCAGGATAAG

GGAATTGGTCAGTATGTTGACAAGGCACAGAGTTACCTTCACCAATTCGAGTCTTCCCAATCAACCACTA

AGGATACTCCGGCACCGGCGCCGGCTAAAGATGGTGAGGCAAAGCCAGAGTCGCATCCAAGCGGTGGCCA

TGAAGGCAAGGAGTCTGGAAGCGGTGGCGGTGTTGGGGATTACATGAAGATGGCTCAAAGTTTCTTGGGC

AAGTAATTAAGCTTAATTAGCTATGTTGTTAAATGTTTAGGGTTTTTATGTACCATAATTTCAATATTTC

CTCTCTTCTATGCTCTGTATGGCTTATAATTCTGTACCCAATCTTCTGTCTAATAATATGCAGCAATAAT

TTGGTACCCAAAAAA

>GH985219.1 JutEST5217 Jute Plasmid library 20-45 days after sowing Corchorus capsularis cDNA similar to Chlorophyll A/B binding protein precursor, mRNA sequence

GGACTCACAACCACAAACTCCCATCAGCAAACAAAAAATCTCTCAATATTTCACAATGGCCGCCGCAACC

ATGGCTCTTTCCTCCCCAACCTTCGCCGGGAAAGCTGTCAAGCTTTACCCATCCACCTCTGAAATCCAGG

GCAATGGCCGTGTTTCCATGAGGAAGGCCACCAAGCCCTCAGCTCCCTCCGGCAGCCCATGGTACGGCCC

AGACAGAGTCTTGTACTTGGGCCCACTCTCCGGCGAGCCACCATCCTACTTGACCGGTGAGTTCCCCGGT

GACTACGGTTGGGACACCGCCGGTCTCTCAGCCGACCCCGAGACTTTCGCTAAGAACCGTGAGCTGGAGG

TCATCCACTGCAGATGGGCCATGTTGGGAGCTCTCGGCTGCGTTTTCCCGGAGCTTTTGGCCAGGAACGG

CGTCAAGTTCGGCGAAGCCGTGTGGTTCAAGGCCGGAGCCCAGATCTTCAGCGAAGGAGGACTCGACTAC

TTGGGAAACCCAAGCTTGATCCACGCTCAGAGCATCTTGGCCATCTGGGCTTGCCAAGTTATCTTGATGG

GAGCCGTCGAGGGTTACCGTATTGCCGGTGGACCCTCGGGTGAGGT

>GH985218.1 JutEST5213 Jute Plasmid library 20-45 days after sowing Corchorus capsularis cDNA similar to Ethylene-responsive element binding protein, mRNA sequence

GGGTAGCACCATTGACGAATCCTCCAGCCCAACGGTGGAGCTCACTTTGGTGGCAACACTACGCAGTGGG

GTTGTAGGGAGATTCCCTTTCGCGTGCCACCAGCAGCTGGCTCTGGGTGGTGGAGTCGCTCGTGGTGGGA

TGGGGTGGGTCACACGGTCGCGGCCGGTTCTATTATTTGAAGCGTTGGGAAGAGCTGAGCCGATTGGAGT

GGAGTTGAGTATGGGATTTGGGAGGTAGGGTTCAAAGTGAATCAAACTCTTTAACTATGGTTTACCTTTC

GATGTAAATAATTTAGGAATAGGCGATCGCTGTTTTTTACAACTATTTAAAGTTCAAACTATTATT

>GH985217.1 JutEST5212 Jute Plasmid library 20-45 days after sowing Corchorus capsularis cDNA similar to Elongation factor 1-alpha, mRNA sequence

GGGGTTATCAAGAGTGTTGAGAAGAAGGACCCATCTGGTGCTAAGGTCACCAAGTCTGCTGCCAAGAAGA

AATGAGGTTGCTTATTGAGAGAGAGTATTATTACAGCTAGGATGATAGTTGGTCAATTATCTTAATTTTG

GTCTTTAATTTATTCTGAGTATTTTACATTGCGGTTTTGTGGATGACTTAATACTGCTTTTCTATCTGTC

CTGTTTTTGTCAATGTCTGCGTGTTAAAATTGTAGTTGGATTGCTTTTAAAAGCAATCTATAATTTACAG

AATTTACTGGA

>GH985216.1 JutEST5211 Jute Plasmid library 20-45 days after sowing Corchorus capsularis cDNA, mRNA sequence

GCCTCTGCTCCTGCATTTACCAACTAGCAGCTTCATAGACCCTGCTGAGGGAAATGATGAGTCCATAGTT

GGTAATGTTGGCTCAGATGCATCGGCTAAATCGTTCTCTCATGTGGAGGTTGCAGCTGCTAAATAACTTA

AAAGGATTTTCTCTGCAGCAGGGAGATTTCATTTCTCATGTCATTTTGCTACTAAGAGTATGGTTAACTG

ATTACAGATATAAATGTGTTACCATGGTAGGAAGATCACTTGTGAAATTTGAAACATTTAAAAGCTGCAA

TGAATCTGAAGGGCCAAGAATCCATGATCTGTTATATGAGGTTCAGCAGATCGACTAAAACTGAATTGAG

GGTAGACTCAACAACAAATTACAGTTCCAGGTTCCTATTCCTGTAATTGATGGATTTTGAATCAATTCTG

AAGTGCGTGTAATAATCTAAGATGTTCTGAAAAATTACCTCTTTCTGTTGTAAATTGGTATCTATGAGTG

TTTGTTTCCGGTTTAGAATCAGTCATCATATTTGTTGCTCCAAGCAAGGGTTTTATTGGATCTTCTAGTC

AGTTGAAAGAAAATAAAACTCTGTAATTACAGAGCCTTAAAAGGC

>GH985215.1 JutEST528 Jute Plasmid library 20-45 days after sowing Corchorus capsularis cDNA similar to Heat shock protein 81-3, mRNA sequence

GAACAAGATCTTGAAGGTTATCCGCAAGAACTTGGTGAAGAAGTGCATTGAGCTCTTCTTTGAAATTGCT

GAGAACAAGGAGGATTACAACAAGTTCTACGAGGCTTTCTCCAAGAACCTCAAGCTTGGAATTCACGAGG

ATTCTCAGAACAGAACCAAGATTGCTGAATTGCTCAGGTACCACTCCACCAAGAGCGGTGATGAGATGAC

TAGCTTGAAGGACTATGTGACCAGGATGAAGGAGGGCCAGAGTGACATTTACTACATCACTGGTGAAAGC

AAGAAGGCTGTGGAGAACTCCCCATTCCTTGAGAAGCTGAAGAAGAAGGGCTATGAGGTGCTCTTCATGG

TTGATGCCATTGATGAGTATGCTGTTGGTCAGCTTAAGGAATTTGAGGGCAAGAAGCTTGTGTCTGCTAC

CAAGGAAGGTCTGAAGCTTGATGAGAGTGAGGATGAGAAGGCTAAGAAGGAAGCTTTGAAGGAGAAGTTT

GAGGGTCTGTGCAAGGTTATCAAGGATGTGTTGGGTGACAAGGTTGAGAAAGGTGGTTGTCTCTGACCGT

GTGGTTGACTCCCCCTGCTGTTTGGTGACTGGTGAATATGGCTGGACTGCCAACATGGGAGAGAA

>GH985214.1 JutEST525 Jute Plasmid library 20-45 days after sowing Corchorus capsularis cDNA similar to ubiquitin-protein ligase, mRNA sequence

GGGGAGATGCGAATGAGAATTCTGAGGTTTCAGCTCATCTTACATCTTACAGCAGTGATCATTCAGGAGA

AGTGAGATCCAACCCACATGCTGCAAGCTAGCTCAGCCACTCCTCACAGAGAGATGCCTGAGTTTTCTCC

TCGTTTGATGGATGCTCGATCTCGAAGCCAAACAATTGGCGCCGGCCATCAGAGAGATTCATTCCAAGGA

TAGTTTCTTCTCCTGGTATTGAGAACAGGGCGTGATCTTTCTGAGGTCGAAACTCAAGTTAAGAAGTTGG

TGGAGGAGTTGAAAAGCAGCTCAGTTGATACTCAGAGACAAGCAACCGCAGAATTGAGGTTGCTAGCCAA

GCACAACATGGATAATCGGATTATCATTGCAAACTCTGGGGCAATTAACTTATTGGTTGATTTGCTTCGC

TCGACAGATACAAAGATCCAGGAAAATGCTGTTACTGCACTTCTTAACTTATCGATCAATGATAACAACA

AATCGGCTATTGCTAATGCTAATGCAATTGAGCCTCTGATTTATGTCCTCGAGACAGGAAGTCCTGAAGC

TAAAGAGAATTCAGCTGCCACGCTCTTCAGCCTTTCAGTCATCGAGGATAATAAAGTCAAGATTGGAAGG

TCA

>GH985213.1 JutEST524 Jute Plasmid library 20-45 days after sowing Corchorus capsularis cDNA similar to Putative Major latex-like protein, mRNA sequence

GACCCACATCTCAAAACCTAAAAAAGTCTCATTAGCAAAAAAAAAAAAATGTCTTCGTCCATAAGTGGCA

AGCTTGAGGGATCTGTCGAGATCGATTCTTCTCCCGAAAAGTTCCATGATATGTTCTTTCACAGGCCGCA

CCATGTCCAACACACCTGCTCCGAAAAAGTACAAGGATGCGATTTACACGAAGGTGATGTTGGGCAACAC

GGTGCCGTCGTCTGCTTCCGTTATACTCATGATGGGAAGGCCAAAACTGCAAAACATATAGTGGAAGTGG

ATGATCCAAAGACCCACTCGGTCAGATATAAGATGCTGGAAGGAGACTTGCTGGAGGAGTATAAGAGCTT

CGTGATTACAATACGGCAACTCCGAAGAGCGACGGTAAGGGGAGTATAGTGAAGTGGGGTTTTGATTATG

AGAAGCTGCATGAGGGCATTCCTCATCCTCAGTCTCTTCTTGACTTCTTCTTGGATCTCACCAAAGATTT

GGGAACTCACCTCACCCAAGGCCCAAAACATAATTAATTACAAAATCATGGCATGAAATTATAATATTCC

ATGGTGGAATATTACTCTATCTGCCTGTACTGTTCTAATTATTGCTACCAAA

>GH985212.1 JutEST522 Jute Plasmid library 20-45 days after sowing Corchorus capsularis cDNA similar to Putative uncharacterized protein, mRNA sequence

AGTCTGGAAGCGGTGGTGGGTATGGTGACTACGTGAAGATGGCTGAAGGGTTTCTTGGGCAAGTACTGAA

GCTAATTAGCTTTGCTTGTTAAACCTGTGTGAGGTGTTGTTATGTACCATAATTTACAATATTTCCTCTC

TTCTATGCTCTGTATGTTGCTTATAATTCTGTACCCAATCTTCTGTCTAAGTAATTGCAGCAAATTTGGT

ACATTACTGTTAACTTTATGTATCAATTAATCCAAGATAGTCGCTAGTTGC

>GH985211.1 JutEST521 Jute Plasmid library 20-45 days after sowing Corchorus capsularis cDNA similar to Glutamine synthetase nodule isozyme, mRNA sequence

GGACCCCTCTCCATCCTAACTTCTCTCTGTCTCCTTTACTGTCGAAGCAGCAACAGCAGCCATGTCTCTT

CTCACTGATCTCATCAACCTTAACCTCACTGAAACCACTGAAAAGATCATCGCCGAATATGTCTGGATCG

GCGGTTCTGGTTTGGACTTGAGGAGCAAAGCAAGGACTTTGTCTGCACCAGTTTCCGACCCTGCCAAACT

CCCTAAATGGAACTACGACGGTTCCAGCACCAACCAAGCCCCCGGAGACGACAGTGAAGTGATTCTTTAT

CCTCAAGCGATATTCAAGGATCCATTTAGAAGGGGAAACAACATCTTGGTCATGTGCGACGCTTACACGC

CAGCCGGCGAACCCATCCCCGGCAACAATCGATTCAATGCCGCTAAGATCTTTAGCCACCCTGAGGTGGC

TGCAGAGGTGCCATGGTATGGAATTGAGCAAGAATACACTCTCCTCCAGAAGGACGTTAAGTGGCCTCTG

GGGTGGCCTGTTGGAGGGTTCCCCGGACCACAGGGCCCTTACTACTGTGGGAGTAGGGGCTGACAAGTCC

TTTGGCCGTGACATCGTGGATGCTCATTACAAGGCATGTCTCTATGCTGGCATTAACATCAGT

>GH985210.1 JutEST51100 Jute Plasmid library 20-45 days after sowing Corchorus capsularis cDNA similar to Unknown, mRNA sequence

GGGATCCATCGAATCAAAGCTGAGAGTGACAGAGAGTGTGAGTCTGAGATTGAGAAAATGAAGAACGCCG

GCAAGAGTGGCAAAGGCAACGGGAAGCCGTCGGTGGAGGTGAAGAAAGACAGGAAATCCGGGACCGGAAT

GAACGGATCGCCTAAGAAAGGAGGTCACGGTGGCAAATTCACCTGGGCCGGAGATGGATTCTCGCCGGCG

GAGATCGGATTCGAGAAAGAGGTCGTTGATGTTAAGGATCCTAACTTTGAAGATCCCGATGAGATCGTCA

CCGTTTGATTATTATTATTAGGTTGGCGATTATGATGATGATCATCATTATCTTTTCTGCTTTTTTCTAG

TCTGATGATCGAGATGTAAATATTTTATGACTGAATAAATAAATTATAGTTATTGCCATTTTCC

>GH985209.1 JutEST5196 Jute Plasmid library 20-45 days after sowing Corchorus capsularis cDNA similar to Metallothionein-like), mRNA sequence

GATAGCAATCCCATAGCAAGCAATCTTCAATAACCCAAAAGCTCTTCCAACTTCTTTCAAAAAACCTTCC

TCAATTCACCATGTCGGACAAGTGCGGAAACTGCGACTGCGCTGACAAGAGCCAGTGCACCAAGAAGGGA

AACACCATGATCATTGAGACTGAGAAGAGCTACGTTGCCACCGTGGTGATGGAGGCTCCTCCAGCAGAGA

ACGATGGAAAGTGCAAGTGTGGATCTAGCTGCAGCTGCACAAACTGCACATGCGGTGGTCATTAAGCAAA

CATATATATAGAAATGAAACAAATGATCCCATGTTGCAGAAACATGGGCAATATTTGGGACCTAAAATCT

TATTTTGGGTTTGAATAATTTGTGTCTTTCCTGTTTCTGTTGGTGTCTGTGTTTTCATGTCCTGTCTGTC

ATGTTTGTGTCTCCTTATATACTGTGATGTAATGGCCATGGCCATGCCTTTGGCCTTGTCTCATCTTAAT

GAACTATTATTAG

>GH985208.1 JutEST5194 Jute Plasmid library 20-45 days after sowing Corchorus capsularis cDNA similar to Blue copper-like protein, mRNA sequence

GGTTATCACAAAAAAACTCTCATTGTCATTCTTTTCACTCTCATTTCAAAGTGTTTCTCCACCTTTTCTC

TTTCACAAGAAAATGGCAAGCTCAAGTGTGGGAATGGCCTGTCTTTTGCTAGTTTTATGCATGGCGGTTG

TGCCAATTTTGGCCACGGTTTACAATGTTGGTGACGCCAACGGCTGGGCAACCGGTGTCGATTTTAGCTC

TTGGGCTAAAGGCAAGACCTTCAAAGTTGGGGATAACCTTGTTTTCAATTACCCAACAAGCCATACGGTG

GAGGAGGTAAGTTCTAGCGATTACAGTTCTTGCTCAGTGGGAAATTCAATCTCAACAGATAGCAGTGGAG

CCACCACCATTGCTCTCAAGACAGCCGGCACTCATTACTTCATTTGTGGTGTGATGG

>GH985207.1 JutEST5191 Jute Plasmid library 20-45 days after sowing Corchorus capsularis cDNA similar to Dehydrin, mRNA sequence

GAGAGACTATTTGCTTCAGCAGCGGCGCTGCCAGAGCTTTTTGAGAAGGGTCAATTCTCAGGTTTCATTT

CTAGTTGAAAGATGGGTAGTCGGGAGCTTGCTTGCTCTTACGCTGCTATGATTCTCTATGATGATGGCAT

CGCCATCACTGCAGAGAAGATTGCAGAATTCCGCAAAGCAGCCAATGTGCACTTTGAATCTTTCTGGCCA

AGCTTGTTTGCCAAGCTTCTTGAGAAGCGCAACATTGAGGATCTCATAACCAATGTTGGAGCTGTGGTGG

CGGTGGCGCTGCTCCTGTTGCTGCAGCTGTTGGTGGTGAGGCTGGTGCTGCTGCTCCAGCTGCTTGCTTT

CGAGGGGAAGAAAGAAGAAGAGAAAGCTGAGCCAGAAGAAGAAAGTGA

>GH985206.1 JutEST5189 Jute Plasmid library 20-45 days after sowing Corchorus capsularis cDNA similar to Unknown, mRNA sequence

CGATGTGATTCGAATAGTACACTCAAATGGACGCGTGGAAGAAATCAGTGGCACCGTCAAAGCCAGCGAA

ATCATGAAGGCTTACCCTAAACATGTCTTGAAGAAACCTTCGTCGCCTTCCGACGACGGAATGGTCCCGA

AAATCGTCGTCGTCCCACCGGACGCCGAGCTCCAACGTGGCAAGATTTATTTCCTCATGCCAGTCCCTTC

GGCGCCCGAAAAAACCCGATCGAAATCTTCCTCAACGACGTCGTCCAAGAAGAAAAGAAGAGATAGCAAC

CTTGACGGAAACAACAACAATATTGATAAAAATAGCGACGGAAACAGCCATAGCCGACATCATCGTCGTC

AAAGCAGAAGCATGAGCAACGCCAGCAAAAACAACAACGCCGCCGCCGCCAACAACAACACCATTTCAAT

GACGAATCTTTTGATATCGGATCAGTATTTGAGCGAGATACTTTCGAGAGAAGTTGTCGACGCAAAGAGA

TCGACGAAGAGGCCGAGTTGGGGTTTGGAGGCCACACTTAGAGAGCATCTCCGAGACACCCAACGATGGT

TGATGATCAGTTTA

>GH985205.1 JutEST5186 Jute Plasmid library 20-45 days after sowing Corchorus capsularis cDNA similar to Putative uncharacterized protein, mRNA sequence

GATTGAAACATTTAACAATGGCGTCAGAGAAAGACCACAAGCCAACTTCGTCCGACCTTTTTTCCAGTGC

TAAGGTGGTTGCGGAGGCCGCAAAATCAGCATTCGGCGGCAATGAAGGCGGCGACAAGTTCGATAAGGCG

AAGGTTGCCGAAGCCGCTTCTGATCTTTTAGATGCAGCCAAAGACTATGGCAAGCTTGAGCAGGATAAGG

GAATTGGTCAGTATGTTGACAAGGCACAGAGTTACCTTCACCAATTCGAGTCTTCCCAATCAACCACTAA

GGATACTCCGGCACCGGCGCCGGCTAAAGATGGTGAGGCAAAGCCAGAGTCGCATCCAAGCGGTGGCCAT

GAAGGCAAGGAGTCTGGAAGCGGTGGCGGTGTTGGGGATTACGTGAAGATGGCTCAAAGTTTCTTGGGCA

AGTAATTAAGCTTAATTAGCTATGTTGTTAAATGTTTAGGGTTTTTATGTACCATAATTTCAATATTTCC

TCTCTTCTATGCTCTGTATGGCTTATAATTCTGTACCCAATCTTCTGTCTAATAATATGCAGCAATAATT

TGG

>GH985204.1 JutEST5184 Jute Plasmid library 20-45 days after sowing Corchorus capsularis cDNA similar to Unknown, mRNA sequence

GACTCAGTTCCATTGTAGGTTCCTGTTTCCCTCTTCTCTCCCCCTCTGAAATTAAGAACCCTAGACAGAA

AGGAAGAGAAAAGCTACGACCCCGATGGCTACTCTGGCAGCGATTGCAGCTCGTAGAGCAGCGACTTATG

CTCGTACACCGGCTTCTTCTCAAGCCGCTTCTCTCATTTCTCGCCGTGGTTTTGCCGGCGCCGCGGATCA

CCATGGACCCCCCAAAGTTAATTGCTGGCAAGACCCAATGAGCCCGTCCAAATGGAAGGAAGAACACTTC

GTTATTGTCTCTTTATCTGGTGGGGTCTGGCTATC

>GH985203.1 JutEST5183 Jute Plasmid library 20-45 days after sowing Corchorus capsularis cDNA similar to ATP binding, mRNA sequence

AACACATTGAGGCCGCCTGTACCAGCCTACTGTGACCCCGAATGGAAGTTACTAATGGAGCAGTGTTGGG

CTCCCGATCCTGTGGTTCGTCCATCATTCACTGAGATTGCAAGGCGTTTACGTATTATGTCCACAGCATC

GGTCCAGACAAAACCACATGGTATCAGCTGCCACCAAGTATCCAAGTGATGGCTCTCTCGATGATTGCTC

TTACTTAAATTATTTTAAATGAAATATGTATAGTTTATTTTCTTTCTACTCCATAAACGTTTGTGTATTA

TATTATGAGTAAGGGGAGAAGGAGCTCCCATTGGCATTGAGAGGGTGAGGTGACTCAGAAAATTTGTTTA

TATATGTAAAGCTATTATTTCTGTCTATAGTGGATACAGACTTTGTTGGTGCTTGTGGGCAAAGTTAATA

CAAAAAAAAATGCTTTTCATTGT

>GH985202.1 JutEST5182 Jute Plasmid library 20-45 days after sowing Corchorus capsularis cDNA similar to Low molecular weight heat shock protein, mRNA sequence

TGCTTCAGATCAGCGGTGAGAGGAACATGGAGAAGGAAGACAAGAATGACACGTGGCATCGGGTGGAACG

CAGTAGTGGCAAGTTCATGAGGAGGTTCAGGTTGCCTGAGAATGCCAAAATGGACCAGATCAAGGCTTCC

ATGGAGAATGGTGTGCTTACTGTGACTGTTCCAAAGGCGGAAATGAAAAAGCCCGATGTCAAGGCCATTG

ACATTTCTGGCTGAGTAATGGAACATTGAGTTCATAAATTCATGCCCTCTCTGTCTCCTGCGACGGATTC

CTCTAAATATAACTACTTTTTTTGTGTCTTTTGTTTCTGTCAGTTCAGCCGAGAGTTTTGTTCAAATTGT

GTCTGAGAAAATCTGTGTGCTACTATGTTATCTTGTATAAAGTGTGTGTAATTTTCCATTTCCATATGTA

GAAATAAAATCACTAGTTTCTTTATAA

>GH985201.1 JutEST5181 Jute Plasmid library 20-45 days after sowing Corchorus capsularis cDNA similar to S-adenosylmethionine synthetase 2, mRNA sequence

GGGGTATCAGTGTCGGTGATGTGAAAAATGTATGAGAATTTTGCTTTAATAGATGGCACTGAAAGTAACA

AGTGCCTCTTTAGATTTCTTGTGAAATTAGCTTTGCAAAGGCAGCAGCAATATACTTATGTGTAATTGTA

CTTTGGATTTGGGATAATCAAATTTATTTTAC

>GH985200.1 JutEST5179 Jute Plasmid library 20-45 days after sowing Corchorus capsularis cDNA similar to Actin-depolymerizing factor, mRNA sequence

GGGTAGATGGCATTCAGGTAGAGCTGCAGGCAACTGATCCAACTGAAATGGGTCTTGATGTCATTAAGAG

CCGAGCCAGCTAAATGGAAACCTGGAAATTTCCAAGAGGCGTGTGATGGAGTCTATATCAAAGCTTTCTG

CTTTTTGCTTAATATGAAATGTTAATAGGTTGTGAGGGGGGCGGGTATCGACCTGACTGAATGATAAGAT

TGGTTTGATCGTTCAAGTTTTATATTGAAGTCTTTATCTAGAACAAGTTTAGTGATTTTAAATTTATATT

TATGATTTGTTATGCGTGGAGATTAACC

>GH985199.1 JutEST5180 Jute Plasmid library 20-45 days after sowing Corchorus capsularis cDNA similar to Metallothionein 2a, mRNA sequence

GGTTCAAGTTTCCAGTGTTAGCTCTTTGATTTCCCAGCAAAAAAATCTCTTCAATTCTCTCAATTAGAAA

ACTGAAAATGTCTTGCTGCGGAGGAAACTGTGGCTGCGGCTCTGGCTGCAAGTGCGGCAATGGCTGTGGA

GGATGCAAGATGTACCCTGACATGAGCTTCGCCGAGTCAACCACAACCGAGACTCTTGTTATGGGAGTTG

CACCCCAGAAAGTGCACTTCGAGGGAGCTGCTGAGATGGAAGTTGGGGCCGAGAATGGCTGCAAGTGCGG

AGCTGACTGCAAATGCAACCCTTGCACTTGTAAATGAGAGAGAAAAAATGGCTTAATTGATCCCCGAAAA

CAGCAGAGAGTCCTTAATTTCCCAACAAATAATATCTCAGCTACTAGTCTTTCGGAAAGATTATCTTGTT

CCTGTGTTTGTTTAAATAAAGTAGCCAGGTTTTATCTGTGGCAAAATCTCACCAGATTTTCTTGGTGCAG

CAGCCATGGTGCTTGCTTGGTGGTATAAGTATAGAATAGTATTAGTTTCTTATCGTCTAATGTAAGCATC

TCTACTGTCATCAGGATGATACTCTATCTATGCTTCTTATTGTAAGAAAGTGAGTCGGTTTTTCTAATGA

AATGAACGGGAAAAC

>GH985198.1 JutEST5176 Jute Plasmid library 20-45 days after sowing Corchorus capsularis cDNA similar to axi 1-like protein, mRNA sequence

GACTAATGGTTATCTTTTAGTACACGCGAATGGCGGATTGAATCAGATGAGAACTGGGATATGTGACATG

GTTGCTGTTGCAAAGATTATGAATGCGACTCTTGTTCTTCCTCTCTCGACCATGAATCTTTCTGGACTGA

CCCAAGTACTTTCAAAGATATTTTTGACTGGAGACATTTCATGAATGTACTGAAAGATGATATTGACATT

GTTGAGTACTTACCTGCTAAGTACGCAGCGAAAAAGCCACTTGACAAAGCCCCTGTTTCCTGGTCAAAGG

CTAGTTACTATAGAAGTGAGGTGCTCCCACAACTAAAGCGGCATAAGGTGATCAAGTTCACACACAGTGA

TTCACGCCTTGCTAATAATGGCTTGCCACCATCCATTCAAAAGCTTAGGTGCCGAGCCAATTATGAGGCT

CTCCGATATACAAAAGAAATTGAAGATCTTGGCAGAGATCTGGTTGATAGATTAAAGAACAACAATGAAC

CATACATTGCTTTACACTTAAGGTATGAGAAAGATATGCTCGCATTTACAGGCTGCATTCATAACCTTAC

TGTTGAAGAAGGTGAGGAACTCACGGCTATGAGATACAGTGTAAAACACTGGGAAGAAAA

>GH985197.1 JutEST5175 Jute Plasmid library 20-45 days after sowing Corchorus capsularis cDNA similar to Aminoimidazolecarboximide ribonucleotide transformylase/inosine monophosphate cyclohydrolase, mRNA sequence

GCTGCTGCCACGCCCAAATCCAATCATTCTCGACAAAAGCAGCAACTTGAAGAGAAACAGAAACAAACAC

AAGAGACGTTTTTGAGTTTTGGAACAGGGCTTAAGGTTAAAACAGAGGAATTGGACAACAGGGAAGACAT

TTTCCCTTCGTTTACTTTCCCTGTGGGATCAGAAGAAGTGGACAATACAGGCATGTTCGTGAATTCCCTG

ATGGAAAATGATATCATGGGGAATCTGTCTCGGGCAACATCTGAATCCAACTACTTCTCAATGTCACCAT

GCCATATGAACAGCTTTGGAATAGGTTAGGCCAAAATGTCCAGACTTCAGAGTCTGAACTCACCACGGAG

ATAATATCAGCACCAACTTCGGTAACTAATTCGCCTATCGGGGATTTTGATATCTCATCATTGATAAGTT

GGAATTTGATACCACTTTATAATCCTGAAATATTCTTCTCTTAATGTGGTTTTCAAATCCCCACCCCATT

AGAAGCAGAAGGAAAGAGGAAAATACAATATTGGAGATAAGTGATGTTTAGTGCACTAATTTAATTTGTA

GAGAAGCTGAGCCAGCTTTAGAAAAGAGATTGGAGGAGTGTTCATATAAGCGCTGCTGTGACCAGACAGC

AAA

>GH985196.1 JutEST5174 Jute Plasmid library 20-45 days after sowing Corchorus capsularis cDNA similar to Auxin-repressed protein-like protein ARP1, mRNA sequence

GATTACTTGAAAAGAGAAACCGGAGAGGAAAAGAGAAAGAAAAAGAAAAGAAAAATGGTTTTACTAGACA

AGATGTGGGATGATGTTGTTGCCGGACCGCAGCCGGAACGTGGCCTTGGCAGTCTTAGAAAGCTGGTCAC

CAGCCCTCTTAACATCAAAGGTGAAGGAGAAGGGAGCAGGTTCCAGAGATCGTTGTCGATGCCGTCGAGT

CCAGGGACACCGACGACTCCGATAACACCAACGACGCCGGTGTCGGCGCGTAGGGATAACGTGTGGCGGA

GCGTGTTCCATCCGGGTAGCAACCTTGCCGCCAAGGGAATCGGCGCTGCGGTTTTCGACAAGCCTTCACA

AGCCAACTCTCCTACTGTCTATGACTGGCTTTACAGTGGGGAGACAAGGAGCAAGCATCATCGCTAATTG

GGTGGTGGTGACTGTGACTGACAGACGGGGGAAGCTTAGCAGTATAAGCAGGGGGGATTTGGTTGAAGCA

TGTAAATAGGCGTCGTTCTTTCAATCTTCGATGTGCCAACGTTGTCCTTGTTTTGATGGGTATTTGTGAT

TAAGTGTCCTAAGTAGTAGAACGTTTAAGACTTGACTAAGTTCTTAATTAAGCCCTTTCCTTCCTCTTCC

TTTTTTTGTAAGTATAAGA

>GH985195.1 JutEST5172 Jute Plasmid library 20-45 days after sowing Corchorus capsularis cDNA similar to 60S ribosomal protein L27, mRNA sequence

GTCGACCTAGGGTTTTTAGCGAGGAGAGAAACAGAGCGCTGCAGCCGCCGTCGCCAATGGTGAAGTTCTT

GAAGCCCAACAAGGCCGTCATTGTCCTCCAAGGCAGATATGCCGGACGCAAAGCCGTGATCGTCAGGTCC

TTCGACGATGGTACACGTGATCGCCCGTACGGCCACTGTTTGGTTGCAGGGATTAAGAAGTACCCAAGCA

AGGTTATCCGCAAGGACTCCGCCAAAAAGACTGCCAAGAAATCCCGGGTCAAGTGCTTCATCAAGCTAGT

GAATTACCAGCACCTGATGCCCACACGTTACACGCTGGATGTGGACTTGAAAGATGTGGTGACCGCTGAT

TGTTCTTCAACAAAGGACAAGAAAGTTGCAGCTTGTAAGGCTACTAAGGAACGTTTTGAGGAGAGGTTCA

AGACCGGGAAGAACAGGTGGTTTTTCACCAAGCTTAGGTTCTGAGCTTTTGTTGGTTTGTCAGTTGTTGA

GGTATTATAAGTCGTTTTGAATAGGAAAAATTTGAGCTGTCATGTAGTGTCTTTTAACAGTTTTGATTTT

AAGATAATCGAACTGCTTCCATGAGATATTACTTCTGGGCATGATAGTAATGCTGATTTGCTCATTTCTG

TTTGAGAGTTATGTTC

>GH985194.1 JutEST5170 Jute Plasmid library 20-45 days after sowing Corchorus capsularis cDNA similar to Actin depolymerizing factor 6, mRNA sequence

GGGGATGACAGCAATTTCCAAGTTTTGCCATAAAGACTTTTATCAGTCTCGATGATGATAAACTTGTACT

TTCGGGTATTTGTGATTTGTTATCTATAGTACCTATGTGTTGAGTGCCTTTCTGTAGTCTTTTATCTCCT

TCAACGCTAATATGCAACTGTTTGTGAGGGTTAGACAACACCTTTAGTTCATTTTCAAGTCTTTTTTCTT

TTTTGAAATATTGACCCTCGTCAATGTTTATTTGGGAGTGTTAAAAAGGGCATGCTTTCATAG

>GH985193.1 JutEST5168 Jute Plasmid library 20-45 days after sowing Corchorus capsularis cDNA similar to GTP-binding protein, mRNA sequence

GGGGCACGCTTGACATACAAGAATGTCCCTACATGGCACAGGGATGTTTGCAGGGTCTGTGAAAACATTC

CTATTGTTCTTTGTGGAAACAAGGTGGATGTGAAGAACAGGCAGGTGAAGGCGAAGCAGGTTACATTCCA

CAGGAAGAAGAATCTTCAGTACTATGATATTTCTGCTCAAGAGCAATTACAACTTTGAGAAGCCCTTCTT

GTACCTTGCCAGGAAGCTTGCCGGTGATCCCAACTTGCACTTTGTGGAGTCTC

>GH985192.1 JutEST5167 Jute Plasmid library 20-45 days after sowing Corchorus capsularis cDNA similar to RAN, mRNA sequence

GACGCTACTCATTTTGCTCCTTCCTTCTCTGTGCTTTCACACAAAAAACGCCAGCTCTCGCTAGATTCTT

CCTGCCTTTTCCTTTTCGAATAATATGGCTCTACCAAACCAACAAACTGTAGATTATCCAAGCTTCAAGC

TTGTAATCGTGGGTGATGGAGGAACTGGGAAAACAACATTTGTGAAGAGGCATCTGACTGGAGAGTTCGA

AAAGAAATATGAGCCCACTATTGGTGTGGAGGTTCATCCATTAGACTTCTTCACAAATTGTGGGAAAATT

CGCTTTTATTGCTGGGATACTGCTGGACAAGAGAAGTTTGGAGGTCTACGGGATGGTTACTACATTCATG

GGCAATGTGCAATTATCATGTTTGATGTTACA

>GH985191.1 JutEST5165 Jute Plasmid library 20-45 days after sowing Corchorus capsularis cDNA similar to Enolase, mRNA sequence

GGAAACTCTTCCTGTGACTAATCTCTAAGCTCTGTTTCTAGTTCTCTCAGTCTCTCTTCAGATCTCAAAA

AGAGCTTTGAACTCATGGCTGCACTATCACCGCCGTCAAAGCTAGGCAGATCTTCGACAGCCGTGGAAAT

CCAACGGTTGAGGTTGATGTGGAATTATCCAACGGTATAAAGTGTAGGGCCGCAGTTCCTAGTGGTGCAT

CCACTGGAATTTACGAGGCTCTTGAGTTGAGAGATGGAGGTTCCGATTACCTCGGTAAAGGTGTATCAAA

GGCTGTTGCTAATGTTAACGCCATCATCGGCCCTGCATTGATTGGAAAGGACCCAACCGAGCAGACTGCA

ATTGACAATTTTATGGTTCAACAACTGGATGGAACCCAAAATGAGTGGGGTTGGTGCAAACAAAAGCTCG

GTGCAAATGCCATTCTTGCCGTGTCCCTTGCTGTTTGCAAAGCTGGAGCTGAAGTCAAGAAACTTCCCCT

TTACAAGCACATTGCTAACCTTGCCGGTAACAAGAAGTTGGTTCTGCCAGTCCCTGCTTTCAATGTCATC

AATGGTGGTTCACATGCAGGAAACAAACTTGCTATGCAAGAGTTCATGATTCTTCCTGTAGGAGCATCAT

CCTTT

>GH985190.1 JutEST5164 Jute Plasmid library 20-45 days after sowing Corchorus capsularis cDNA similar to Unknown, mRNA sequence

GGTCCCCGCAATTTAAACTCACTTTCACTTAGATTCCCTCTCAAAAGCAAACTCACCACCAAACTGGATA

AAACTTTCGCCTTTAACTGTTAAAAAAAAGGAAAAAAACAGATTCAGGAAGATCCAAAAGGAATCATGGT

TCAGCTTTTGCCTTCTCCATCTTTTTCCCTCTCTGAGACAGAGCAAGAACATCTCTTGGAGAAGCTCGAT

GTCTTCAAGGTTCAGGGTAGGGACAAGGTCGGCCGGAAGGTTCTTCTCATCATCGGCAAGCACTTTCCTG

CTCGGATGGTTAGCAGTGAGGTGATCAAGAACTATTTGGAGGAGAAAATTTACCCCAAGTTAGGGGAAAA

GCAATTTTCTGTTGTGTATGTGAACACTGATGTGCAGAGGAGTGAGAATTTCCCCGGAATCTCGGCCCTC

CGGTCAATCTATGACGCTATTCCGACCAACGTGAAGGACAACATCGAGGCTGTTTATTTCTTGCACCCTG

GCTTACAGGCCAGGCTCTTCTTCGCCACCTTTGGCCGTCTGTTTTTCAACGGAGGGTTATACTCGAAGCT

TAAGTACGTCAGCAGGGTTGAATTTCTGTGGGAGCATGTGAGGAGAAAGGAGATAGAACTGCCGGGAGTT

TGTGTTCGAACACGACGA

>GH985189.1 JutEST5162 Jute Plasmid library 20-45 days after sowing Corchorus capsularis cDNA similar to Unknown, mRNA sequence

TTCCACTTGTGATTTCCGATTCCGCTGAGAGTGTTGAGAAGACTTCGGGTGCTATTAAGGTCTTGAAGCG

GATCGGGGCTTTTGCTGATGTCGAGAAGGCCAAAGATAGCCAAGCTATTCGTGCTGGAAAAGGAAAAATG

AGGAACCGCCGCTACATTTCTCGAAAAGGTCCATTGATTGTGTATGGAACCGAAGGAGCTAAGCTCGTTA

AGGCATTCAGGAACCTTCCTGGTGTTGAGATTGTCAATGTGGACAGGCTGAATCTTCTCAAGCTTGCTCC

TGGTGGCCATTTGGGGCGTTTTGTGATCTGGACAAAATCGGCTTTTGAGAAGCTTGATTCCATCTATGGA

ACTTTTGACAAGGCTTCTGAGAAGAAGAAGGGATATGTGCTTCCTCGTCCCAAGATGGTTAACGCTGATG

TGGCTAGGATTATTAACTCTGATGAGGTCCAATCAGTTGTGAAGCCAATTAAGAAGGATGTAAAGAGAGC

GACATTGAAGAAGAATCCTCTTAAGAACTTGAATGCTATGTTTAAGCTCAACCCTTATGCTAAGACTGCT

AAGAGAATGTCTCTTTTGGGCTGAAGCTCAACGAGTCAAGGCCAAGAAGGAGAAGCTTGACAAGATTAGA

AAACCA

>GH985188.1 JutEST5159 Jute Plasmid library 20-45 days after sowing Corchorus capsularis cDNA similar to Tonoplast intrinsic protein bobTIP26-1, mRNA sequence

GGGCCTTCTCTGGAGCCTCAATGAACCCAGCTGTTTCATTTGGACCAGCTGTGGTTAGCTGGACATGGGC

CAACCACTGGATCTACTGGGTCGGACCCCTTGTTGGCGGTGGGATTGCTGGAGTCCTGTACGAGTTCATC

TTCATTAGCCACACCCATGAGCAACTCCCCACAACCGATTATTAAACATGATTTGAGTTAAATCAAATAA

AATGTCACATGAAAAGCGGATTTCTTCCGGTTGTATTTAAGTTTGTTGAGTCTTTCTTTGTCTATTATGT

TTCCTTTGTAATTTGTGTTTCTTGTGGCCGTTGTTTTCATTATCCATCCTCTATCATGATCATCATGGAA

GGTAGTTGTGATTGAAAATTGCATGCCCTGGTAAAATATTGTGTGGGAATATATTTATGACCAAAAATAC

CAGGGATTGATGTAGATGTGAAATGTTAGCTTTTTTTAATTTCTTTGTTAATTTCTATGGTTTCATTTTC

CTTCATA

>GH985187.1 JutEST5158 Jute Plasmid library 20-45 days after sowing Corchorus capsularis cDNA similar to 10 kDa photosystem II polypeptide, mRNA sequence

GTCTCCTAAAGTTTTCAAGGTTCTAGCCTCCGGTGGCAAGAAGATCAAGACTGACAAGCCTTATGGAATC

AATGGAGGCATGGCCTTGAGGTATGGTCTTGATGCCTCTGGAAGGAAGGGCAAGGGAAAGGGTGTTTACC

AATTTGTTGACAAATATGGTGCTAACGTCGATGGCTACAGCCCTATCTACGACCAAAAGGACTGGTCCCC

ATCCGGTGATGTTTATGCTGGCGGTGTAACTGGTTTGGCAATCTGGGCAGTTACCCTAGCCGGTATTCTT

GCAGGTGGTGCTCTTCTTGTTTACAACACAAGTGCTTTGGCACAGTAGATTTCTACTAATGTAATGTACC

GTGTTGTATAATATAGCCACTCCAAGTTATTGTTCTCATACATAATCACTTCAAGCTACTTTGCCTATCT

ATATGTCATTTTATTGTTTAAGCCTGGACCAA

>GH985186.1 JutEST5154 Jute Plasmid library 20-45 days after sowing Corchorus capsularis cDNA similar to Endo-xyloglucan transferase, mRNA sequence

ACATGGAAAGGGTGACAGGGAACAAAGGGTTAACCTTTGGTTCGATCCATCTCCAGATTTCCATACTTAC

CAAATTAGATGGAACCATAAGGAAATTGTGTTTTTGGTTGATGATACGCCAATTAGGGTTTACAAGAACA

ATGAAGCAAAAGGTATCCCATACCCAAAACTCCAACCAATGGGAGTCTATTCAACACTTTGGGAAGCCGA

TGATTGGGCGACAAGGGGTGGTTTAGAGAAAATTGATTGGACCAAAGCTCCTTTCTTAGCTTATTACAAG

GATTTCGACATCGAAGGTTGTGCTGT

>GH985185.1 JutEST5153 Jute Plasmid library 20-45 days after sowing Corchorus capsularis cDNA similar to Beta-glucosidase G2, mRNA sequence

TTATTGGCATAAACTATTACTCTTCTTGCTATGCTTCTGATGCACCTCAATTAAGCAATGGCAAACCTAG

CTACCTAACTGATTCTCAACGTCGCACCTCAACTGGAAAAGATGGAGTTCCAATTGGCACACCTACTGTT

GTGAGTTGGCTTTACCTTGTTCCAAGAGGGTTTCGAGAACTTGTGATACACATAAAGGACAAATACAACA

ATCCTGTTGTTTACATTAATGAAAATGGTGTAGCTGAGCAAGACAATGCTTCGTTGTCGGTAACTGAAGC

AATCAAGGATACCTATAGGATTGACAGCCCTTGACGCCACCTAAACGCTCTTAGAGAGGAAATCAAGGCG

GGTGCTAATGTCAAGGGTTATTACATGTGGTCATTTTTGGATAATTACGAATGGAATTCTGGATATACTG

TTAGATTTGGGTTGGTCTATATTGATTTCAAGCATAAATTGCAACGACACCTAAATATCTCAGCTTACTG

GTTCAAGAATTTTCTCAGACCATTAATTGATCTGCCGCTGCCGAACCAAACCAAGAGATGGACGTAATTA

ACAGTTTGTCGTCGTCGTTTGTTGTTGTTGTTGTTGGCTTTCTTTAATTTTAATTT

>GH985184.1 JutEST5152 Jute Plasmid library 20-45 days after sowing Corchorus capsularis cDNA similar to MipC, mRNA sequence

GATCCAAAGCATAGAAAACCCAAACTCAAACAGCTCCTTTTGCTTTTCCTAGCAGAAAAAAACAGTCTTT

ATTTAGCAGCAAAGAGAGGAAAACAATGACTAAGGATCTTGAGACCGCAGAGCAACATGCAGGGGAATAC

TCAGCAAAAGACTACCATGATCCACCACCTGCTCCATTGATTGATTTCGAAGAGTTAACCAAATGGTCGT

TTTACAGGGCAGTGATTGCCGAGTTCATTGCTACACTTCTCTTCCTGTATGTCACTGTGTTGACCGTGAT

TGGTTACAAGGTCCAAACCGACCCTACCAAGAACACTGTTGATGCTGACTGCGGTGGCGTTGGTATTCTT

GGTATTGCTTGGGCTTTCGGTGGCATGATCTTTATTCTTGTTTACTGCACCGCTGGTATCTCTGGAGGGC

ATATCAACCCGGCTGTGACATTCGGGCTATTCTTGGGACGCAAGGTGTCACTCATCCGAGCCCTGATGTA

CATGGTGGCTCAGTGCTTGGGTGCCATATGTGGGTGTGGATTGGTCAAGGCTTTCCAGAAGACTTACTAC

AACAACTACGGGGGTGGTGCGAACGAGCTCGCCGCTGGATTCAACAAGGGAACCGGT

>GH985183.1 JutEST5150 Jute Plasmid library 20-45 days after sowing Corchorus capsularis cDNA similar to Arabinogalactan proteinr, mRNA sequence

GATCGCTTGTCTCAATTTGCTTAGTACATTTTCCTTTTTCACCACAAAACTCAATTTCCTCTCTTTTGCA

ACAATTTCCACCCCCTTACTACACCAACAAATGGAGGCCCTGAGGATAAGACTATTCCTGGCTATGATGG

TTGCCTTGATGGCCATCTCAGCGGTCCAATATGTCGCCGCAGCTGATGCCCCCGCAGCCGCCC

>GH985182.1 JutEST5149 Jute Plasmid library 20-45 days after sowing Corchorus capsularis cDNA similar to oxysterol binding protein, mRNA sequence

GAACACCTTCAATGATCATCAATATTATCTCAATTGAAACTAAAATAAATAAAGAGTCTCTCATTCTTCC

AGCTTGGCCTTTCTCTCACCAAACCATCTTGGAGGATCGAACCCAGAGCCACCAAGAGGATTGCAGCGAC

AAAGACGCCAAGTTGTGAGAACAGTGCCTTTAAGAACCCCATATTTCTTGTAAGCTTCCATTGAATACTC

ACTACAAGTCGGAACATAACGGCAGCTTTTTGGTAACAAAGGCGATATCTCTCTCTTGTAGAATCTCAGC

ATTGCTAAAGCTGCTTTGACT

>GH985181.1 JutEST5147 Jute Plasmid library 20-45 days after sowing Corchorus capsularis cDNA, mRNA sequence

GGAAAAAACATCTACGGTTAAGCATCTTTTAACTTTGCCGCAGTTCAACAGCAGTGTGACTGCACTCTCT

TGGGCTGGTCTTGATCGAAAGAGAAATCATGGGCTTCTCGCAGTTGGAATGGAGAGCGGACTCCTTGAAC

TTTGGAGCCTTGATGTTGAAAGAACACCAGATAAAAGCACCCTGACACCAGCTGTTTCTGCTGCTCTAAT

CGTGCGGCTAGATCCTTTCATGTGCCATGTTTCTGCAATTAACCGGTTGGCCTGGCAGAACCGAGAGAAT

GATGAAAATTGCACAAGCCTCAAGCTTGCTTCTTGTGGAGCTGATCATTGTGTGAGGTTGTATGAGGTAA

TTGTTGACTAAGTCTAATCTGATAATGATTTTGTATTATAGATACATCAGTATGTCATTCTTCAAATCAA

ATACTACAACATATGTGATTCATATATATATATCTAGATTTTAACTTTCATCATTC

>GH985180.1 JutEST5146 Jute Plasmid library 20-45 days after sowing Corchorus capsularis cDNA similar to Aux/IAA protein, mRNA sequence

AGGTTGGCCTCCTATTAGATCGTTTAGAAAGAACTCATTGGCCACCTCTTCCAAGAACACTGATGAAGTA

GATGGAAAGACAGGGCCTGATGCTCTTTTTGTTAAGGTCAGCATGGATGGTGCTCCGTATTTGAGGAAAG

TAGATTTGAAAAACTACTCCAAATATCAGGAGCTATCTTCTGCCCTTGAGAAGATGTTCAGTTGCTTTAC

CATAGGTCAATATGGATCTCACGGAACCCTGGCTAGGGAGCTGCTAAGCGAAAGCAAGCTGAAGGATCTG

CTCCATGGATCAGAATATGTTCTTACTTATGAGGATAAAGATGGTGACTGGATGCTTGTGGGTGATGTCC

CATGGGAGATGTTTATTGAGACCTGCAAAAGGCTGAGGATTATGAAAAGCTCTGATGCCATTGGACTTGC

CCCAAGGGCCGTGGAAAAGTGCAGGAACCGGAACTAGACACTTTCAGACATGATTTTTGCTACCCATCCA

GCAGCTCTTTCTGGAGGAGAAAGAGGATGTTGGAGCGGGCATTCAAGATTCTGTCTAAAGGGATGAGAGG

GATGTTGGAGTGGCTAGGAGAACATGGAGTATCTGCTATCATAAGTTGCTACGTTTATCTGCTCTTTGCG

GGGGA

>GH985179.1 JutEST5145 Jute Plasmid library 20-45 days after sowing Corchorus capsularis cDNA similar to Putative uncharacterized protein, mRNA sequence

GACTCTTCAAACCTCCAGACGTAGCAGCAGCAGCCAAAGCAGCCGCTCTCAAGTTTCCAGCCATGGGACG

TGTACGTACCAAGACAGTGAAGAAATCCTCTCGCCAAGTCATCGAGAGGTACTACTCTCGCATGACCCTC

GACTTCCACACCAACAAGAAGATCATCGAGGAGGTTGCCATCATCCCATCGAAGAGGCTCCGCAACAAGA

TCGCCGGCTTCTCCACCCATCTCATGAAGCGGATCCAGAAGGGCCCCGTCCGTGGCATCTCCCTCAAGCT

CCAGGAGGAAGAACGTGAGCGCCGCATGGACTTTGTCCCCGATGAGTCCGCTATCAAGGTCGACCAAATC

GAAGTCGACAAGGAGACCCTTGATATGCTTTCTGTTCTTGGGATGTCTGACATTCCTGGCCTCGTTAAGG

TTGACCCAGTTGCTGTTTCCGCTCCTCAAGTTGGATTCGGCCGTGGCAGTGGTGGACCTGGAAGGAGGTT

TTAAGTGGTGGGGCTCCATGGACATCTGATGGGTTCTTTAGATTTTTGCTATGATTTTACATTTTAAGTA

GTGTTTTAAATTTGTTTTTTTTGTTGGAACAATATTGGTAGTTGCTATTATCTTAAATGGTACTGA

>GH985178.1 JutEST5141 Jute Plasmid library 20-45 days after sowing Corchorus capsularis cDNA similar to Unknown, mRNA sequence

GGATGCTGTACCAAATATCTTAATCTATATCCTCCTTCTCTCCAGCGACGTGAAGCAAGCGGCAAGTGTT

TACACTTTCCGAAACCTCTTCCGTCAAAATTACACCAAATAATTCTAAAGATAATATGGAGGGAGGGAAA

GAGCGAGGAATGGTGTTTCGTATGCCGCAAAATACTTTCAGACAGTTGCAAGTCAGGTATAAGGAATTGG

AGGAGGGCTTTAAGTCTTGGCTAGCCAAACAGTCTTTGCTTGTCGAGGCGGCTGTCGTGACGGCCGCCAC

CGCTATTCAAGGTGCTGCTATCGGAGGTTTTATGGGTACCCTCACCAACGATTTCTCCTCCACTCTTCCA

ACGCCTCCCCAGGGTTCCCTCAATCCCCAAGCCATGGCTTCTCTCCAGCAAGCCCAGGCTCTAGTCAGGA

GGTCCCTTGGTTCAAGCTCGCAATTTTGCGTGAAGTGACAGGTGTTAATGCTGGCATATCTTGTGTTATG

AAAAGATTGAGAGGAAAGGAGGATGGTCCAATCTAGCATGGTTGCAGCTTTTGGTTCTGGGGCTATGTTT

TCATTAGTGAGTGGCTTCGGTGGCCCAAATCAGGCTGCAAATGCGGTCTCTTCCGGGCTCTTCTTCGCA

>GH985177.1 JutEST5140 Jute Plasmid library 20-45 days after sowing Corchorus capsularis cDNA similar to Alpha-tubulin, mRNA sequence

GGAAAGGCGTCTTGATAAACGCCCCATATTCTCTTTAGATTACGTTCCTTTTTTCCTCCTAAGAATTTCC

TTCGCGTTTTTTCGAATTTTAAGAAAAAAATGAGAGAGTGCATTTCAGTTCACATCGGTCAGGCCGGTAT

TCAGGTCGGAAATGCTTGCTGGGAGCTTTACTGTCTCGAGCATGGCATTCAGCCTGATGGTCAAATGCCA

AGTGATAAGACTGTTGGTGGAGGAGACGATGCTTTCAACACCTTTTTCAGCGAAACTGGTGCCGGGAAGC

ACGTCCCTCGCGCTGTCTTTGTTGATCTGGAGCCTACTGTTATCGATGAAGTGAGGACTGGTACGTACCG

CCAGTTATTCCACCCTGAGCAACTCATCAGTGGCAAAGAAGATGCTGCCAACAATTTCGCTCGTGGCCAT

TATACAATTGGCAAAGAGATTGTTGATCTCTGCTTGGATCGTATCCGAAAGCTTGCTGATAACTGTACTG

GGCTCCAAGGCTTCTTGGTTTTCAATGCTGTTGGTGGTGGTACTGGTTCTGGTCTTGGATCTCTTCTCTT

GGAGCGTCTCTCTGTTGACTATGGAAAGAAGTCGAAGCTTGGTTTCACCGTCTATCCATCACC

>GH985176.1 JutEST5138 Jute Plasmid library 20-45 days after sowing Corchorus capsularis cDNA, mRNA sequence

GGCATGCATTACACATTCTAAAACTCTCTTAACTCTCTGCTCTAAGAGATGGTAGCCATGAAAGTTGTTT

CTATGATACTTCTTGCTGTGTTCATATTGACGCTTGTTGAACACCAAGCATATTGTGAAATATCAGCAGA

CACACAAAGATTAATTCACAAAGCCAATAACAAAGGGCCTTACCTAGGCTTGGTGATTCCTAACCTTTTT

GAAATGAACCCTCTTCTTGAACACCCAAACTATACTTCCACTGGCTTAACCATTGATGTTTCTGGAAGGC

GATTCCGATTCGGGAAAGTTTTTGAGAAGAAGGTTGTATCAGTCATGACAGGACTTGGCACGATTAATGC

AGGCATCACCACCCAGCTACTGTTAAGTTTGTTCAAGATAGAAGGGGTAGTCCATTATGGAATAGCTGGA

AATGCCAACCCCTCTTTACACGTTGGAGATGTCACAATTCCTCAGTATTGGTCACATTTAGCTCTTTGGA

GTTGGCAGAGGTACGGAAATGGACCCCAAGACGAGCTACCTCTTGAAGGACAAGGAGACTATACAAGGGG

AAATTGGTTACATACAATTTGCAGATTATGCCACAAATGTCTCCAACTGTAATTCACACGACAA

>GH985175.1 JutEST5137 Jute Plasmid library 20-45 days after sowing Corchorus capsularis cDNA similar to Ubiquitin extensive-like protein, mRNA sequence

GCTCGCCGTCCTCCAGTTCTACAAGGTGGATGATTCTGGCAAAGTTCAGAGGCTGAGGAAGGAGTGCCCT

AACGCCGAGTGTGGTGCGGGGACTTTCATGGCAAACCACTTTGATAGGCATTACTGCGGCAAGTGTGGCT

TGACTTATGTTTATCAGAAGGCTGGTGGTGATTAGGTTCCTTTCTAGTGGGATTTTCATGTTTTTCTTAT

GAGCTTTTATATCTATGTTTAATGGTACTTTATTTGGTTTAATTTAGTACTTTTGAAATGGATGGTAATT

GTCAGATTTTGGAATTTTGGATGGTTAATTAGATATTGGTATATGCTGATTTTACCG

>GH985174.1 JutEST5136 Jute Plasmid library 20-45 days after sowing Corchorus capsularis cDNA similar to specefic tissue protein 2, mRNA sequence

CCCTACATTGCAGTTCTGCAACTATGAAATCTTTCCCTTCTTTCTTAGGTTTCCTGTCACTTCTCATGCT

TGCTGACACCATAGCAGCAGCAAGAAAAGATGGTGGGGAATACTGGAGAGTAGCTGTAATGAAAGATGAA

TCCATTCCAGAAGCCATCGAAAGCGCCCTTGTTCCGGTTAATGCCGCAGCTGCTTCCTCCTCCGGTGACA

AAACCAACTGTCACGATTTACCTAGCAACATCGAGATTAAGGAAGAGAAGATTTTTGTTGAGGATTTTGA

ATTACCAAGGCCTAATAATTATAACAGTGTTGAAGGAGGGGTGACGAAGGAACGTTCATTTGCTAAAGAT

TTCGAACCCAGGCCCAACCTTTCGGCTTATGGTGATGATGGTGATCTTAAAGAAGAAAAAAAATCATTTG

CTAAAGATTTCGAACCGAGGCCCAACCTTTCGGCTTATGGTGGTGATGGTGATCTTGAAGAAGAGGAAAA

ATCATTTGCTAAAGATTTCGAACCCAGGCCTAACCTTTCGGCTTATGGTGATGATGGTTATCTTTAAAGA

GGGGAAAAAAT

>GH985173.1 JutEST5135 Jute Plasmid library 20-45 days after sowing Corchorus capsularis cDNA similar to Ripening-related protein, mRNA sequence

GGTCTCAGCCCAATCTTACTCCCAGTCCCAAAGTAAACTTTCTTTAGCTCCCCATTGACAATGGCAGCAA

AAACTTCCCTTCTTTTCCTCTCACTCTCTCTTCTCTACATCGCTTCCCATGCCGGCAGATCTCCGGTGGC

CGCAAGCTCTCGCGCCACAAATTTCATCAAGGCCAAATGCAGTGCCACCACATACCCAGCATTGTGCGTC

CAATCGCTGGCCGCATTCGCCCCTTCAATCCAACAAAGCCCACGCCAGTTGGCTCAGACCGCCCTGTCCG

TGAGCCTAGCTAGGGCTCAGTCGGCCCAAGGGTTCGTGGCCAAGATGAAGAAGTTCAAGGGACTTAAGAA

GAGGGAATACGAAGCCATCAAAGATTGCATCGAGGAAATGGGCGATTCCGTAGACCGGCTCAGTAAATCG

GTCCAGGAGCTCAAGCGCATGGGTCGAGCCAAGGGTCAAGACTTCATTTGGCACCTTAGTAATGTGGACA

CATGGGTTAGTGCCGCACTGACCGATGAGAACACTTGCCTTGATGGGTTCGCTGGGCGGGCCTTGGATGG

TAAAATTAAGTCCTCGATTCGAGCCCGTATTTTAAATGTGGCTCAGGTCACTAGCAATGCTTT

>GH985172.1 JutEST5132 Jute Plasmid library 20-45 days after sowing Corchorus capsularis cDNA similar to Inositol-1,4,5-triphosphate-5-phosphatase, mRNA sequence

GGATCTGTTTCTGTTAGCATGACTCTTTACCAATCCAGGCTGTGCTTTGTGTGTTCTCATTTGACATCAG

GCCAGAAAGATGGAGCAAGGCGTAACTCGGACGTGTGTGAAATTATAAGGCGTACCCAATTCTCATCTGC

CCTTGACACAGATCAACCATTGACAATTCCTTCTCATGATCAGAGATTTTGGTTTGGGGATTTGAACTAT

CGTCTTGATATGTTGGATTCGGAGGTGAGAAAGCTTGTTGCTCTTAAACGCTGGGATGAACTAATCAACC

ATGATCAGCTACATAAATAACTGCGTACCGGGTATGCATTTGATGGATGGAAAGAGGGAATCATAAACTT

TCCTCCTACATACAAATATGAAATAAATTCTGATAGATATGTTGGTGAGGTCCATAGGGAAGGGGAGAAG

AAGAGATCTCCAGCATGGTGTGATCGTATACTTTGGTCAGGAAAAGGTATAAAACAGCTTTGTTACCAAC

GAGCAGAATTAAAACTATCAGATCATAGACCAGTAAGTTCAATGTTTTTAGTCGAAGTTGAAGTTTTAGA

CCCCAGGAACTGCAAAGGGCTCTTAATGTTAGCACTGCTGCTGTCCATCCAGAGATTTTCTTT

>GH985171.1 JutEST5130 Jute Plasmid library 20-45 days after sowing Corchorus capsularis cDNA similar to Unknown, mRNA sequence

GTGATGGCTATGGCCACTGGTGAGGCTCCAGCGGAAGTTGCCACGGTTGAGACGCCTGAGATTGTCAAGA

CTATTCAAGAAGCGTGGGATAAAGTTGAAGACAAGTATGCTGTGAGTTCACTTGCTGTAGCAGGAGTGGT

TGCCCTTTGGGGATCCGCTGGAATGATCTCGGCAATTGATAGGCTTCCTCTGATTCCTGGCCTTCTGGAG

GTCGTGGGTATTGGCTACTCAGGGTACTTTGCATACAAAAACTTCTTAAAAGGGTCAGACAGGGAAGCTC

TGATTAAAAAAGTAAAAGAAACATACAAGGAAATAATCGGAAGCAGCAGCTAGGGGGAATCGGTTTCAAA

TGCTGCTGAGTGCTGAGATGAGGTGAACACTAGAAATCTGAAGCCATATATACTTTGCAGCATCAACAAC

ATATGACCTTGCTTCATTTTGTTAGCAAAAGTTCTAACCAGATGAAAACCAATGTAATGTGGTTTTGTCG

TGCGCAATTGAGATACATGTATATTTCTATTTTCTACTCTGTAGCACACTATCAAGTATCATCATCATAT

GCTAATAAACACCTTCCTTTTGTCAC

>GH985170.1 JutEST5129 Jute Plasmid library 20-45 days after sowing Corchorus capsularis cDNA similar to cyc07-like protein, mRNA sequence

GATGAAGATCAAGCTTACAGGAAGATTCGACTCAGAGCTGAGGATGTCCAAGGGAGGAATGTTCTCACAA

ACTTCTGGGGAATGGACTTCACCACAGACAAGTTGCGATCTTTGGTGCGGAAGTGGCAGACATTGATTGA

AGCCCATGTTGATGTCAAGACTACTGACAATTACAGCTTGAGGTTGTTCTGCATTGCTTTTACCAAGAGA

CGTGCAAATCAGGTCAAGCGTACCTGTTATGCTCAGTCCAGCCAGATCCGTCAGATTCGCCGGAAGATGA

GGGAAATCATGGTTAATCAGGCCTCGTCCTGTGATCTGAAAGAACTTGTTGCGAAGTTCATCCCTGAGGT

GATTGGCAGAGAGATTGAGAAAGCTACCTCAAGCATCTATCCCCTACAAAATGTTTTCATTCGTAAAGTG

AAGATCTTGAAGTCTCCCAAGTTCGATCTTGGCAAGTTGATGGAGGTTCATGGTGACTATTCAGAGGATG

TTGGTGTGAAGTTGGAGAGACCCGCTGAAGAGACGGCAGCAGAGGCGGAAACCGAGGTTGTTGGAGCCTG

AAGAGAGCTTTATTTTATAAGTATTAGTATAGTGATACTTCCA

>GH985169.1 JutEST5128 Jute Plasmid library 20-45 days after sowing Corchorus capsularis cDNA similar to Unknown, mRNA sequence

GAACCAAAATCAAAATTTAGTCAAATTGAAAGAAGGAAGAAAAAATGGCGTCGTCGTCTTGGAAGGCGAT

GTCTAGGATATCGTCTCGGTTACAGTCTTCACCACTCAAGTTCAAAAAAACTTCACTCTCTGCTCTGCAA

TCTACTTCGGATTCTTCAGTTACACGCGTTTCTCGTATCTCAAGGTTACCATTGGAGCTGAGTAGTGTAG

GGCCGATGATGCCTTTCCACAGTGCTATTGCTTCGGCTCGATTGGTATCGAGCTTGGCTTCCGAGTCTCG

GACTTGGAGTTTAGTTCCTCAAGGTATCTCAATGCCTTTATGACAGCCTCTTTTTGCAGTAAGTTTTTGG

AGAGGAACATGCATAGAAATATGGAGTTTTCATTGAGACCTTTGTTTTTTGGCTTCGCCGGGTTTCAGAG

GCAAAACATTTTAAAGTTGACCCAGTATCTTTGAGGAACACTGGTTGCAATTGACATCATTTATGTTTTG

GTTCCTTTCGTGGATACTTTGCTTTCTTTTCAGAATGTTTCTCAACCTGGATTAACTTTGCATAATCATG

TTGAAGTGTGTTATTCATGAACTTTAATGAGAGGTGATCAGCTTTTAG

>GH985168.1 JutEST5127 Jute Plasmid library 20-45 days after sowing Corchorus capsularis cDNA similar to Dehydration-induced protein RD22-like protein 2, mRNA sequence

GAACAAACCCTAACTTGCAAACTCTCTTCTTTCCATAGCTAAGCACACAGTTAGTTTTTCAGCAATGGAG

TTTCGTCTCTGTTGGGTCCTTGCTTTCCTTGCTCTTGCGGTGGTAGCAAGCCAAGCTGCTCTATCACCGG

AGCAATATTGGAAGTATGCGCTGCCAAATACTCCAATGCCGAAGGTTGTCAAAGATATGCTATATCCTGA

CATGGTGGAGGAGAAAAGTACCTCTGTTATGTAGGGGGTGGTGGCGTAAACGTGAACACAGGAAAAGGGA

AGCCCAGCGGCACTTCGGTGAACGTTGGAAAAGGTGGAGTGAATGTAAACACCGGGAAAGGAAAGCCCAG

CGGCACTTCTGTTAATGTTGGGAAAGGCGGTGTAAATGTAAATACTGGGAAAGGAAAGCCCGGGGGAACT

TCTGTTAATGTTGGAAAAGGCGGAGTTGGAGTGAACACTAGGAAACCAGGTGGTGGCACCTCCGTAAATG

TGGGTGGCAAAGGCGGCGGAGTATCTGTAAACACAGGACATAAGAAAAAGCCAGTGATAGTAAATGTTGG

TTATTCACCATTTATATACAATTATGCAGCCACTGAGACCCAAATTCATGATGATCCA

>GH985167.1 JutEST5125 Jute Plasmid library 20-45 days after sowing Corchorus capsularis cDNA similar to Putative ABC transporter, mRNA sequence

ATAGTGGGAGAACGTGGTTTGAGGCTTTAGGAGGCCAGAAACAAAGAGTGGCAATTGCAAGAGCACTATT

AATGAATCCAAAAGTTTTGCTGTTGGATGAAGCAACAAGTGCTCTGGATGCTGAGAGTGAATACCTTGTC

CAGGATGCTATGGATTCTTTGATGAGGGGTAGGACTGTTCTGGTCATAGCACACAGGCTTTCAACAGTCA

AAAGTGCAAACAGTGTAGCTGTTATATCTGATGGTCAGATAGCTGAGAGTGGCACCCATGAAGAGCTTCT

CCGAGAGGATGGTATCTACACTGCGCTTGTGAGGAGGCAACTACAAGGTTCAAACACCAACTAAATCTAG

TTCCAAATGGTCAAATACTGTATGCTCCTCTTTCTGAAAATTTTGAACTGTCCTCCCTCCCTTGTTTCTT

CCATTTGGCCTAAAAAAATTTTGGGTTGGAATGAAATTACACTTTAATTGTATCTGTTGATAGGACGACA

AGTCAAATGTTTGACTAGTACGTAGAATGAAATTTCTTGTCTATCTCAGGCAGCA

>GH985166.1 JutEST5124 Jute Plasmid library 20-45 days after sowing Corchorus capsularis cDNA, mRNA sequence

GGAGAAATGGAAATGTGCATTGAATGGCGCAGACTACCCTTCTAAATGAGCAGATAAATTGGGCCGTAAT

TATTAAAGAAAAAAAAAAAAGGTTCATATCTAACCAGAAAAGTTGGTAAGAGTTTCTTCGCTAGCTCCCT

GCTAGTGCTCTCCTTTTTCTTTTTGCTGCAAAGCAAACAATTCCAATCTCTTTAGAGATGGAAGCAACGC

TTTCTGCTGCAGCGCCCGCATTCACTCTCTGCTCCATTTCCAAAGCTCTCCATCGTCATTTGTCTAACAA

CAACTCCCCGGCTCTCTTCGCTCGATGCTTCCACTCTCGTCTCTCCGCTCGGGGCCGACTCGTTACTCCT

CTCTCATCCCTCTCTTATTCCTCTCCACTAAGGACCTTGTGCCCGCCCCCTGTCCCCAAACGCTTTCACC

GCCCTTTGGAATGCTCCGCTAGTTCGATCGCCTCCTTTGGGACTGGTGGAGGCGACGATGGGGAGTGGTG

CTGGCGGCGGCGGCGGCGGCGGCGGCGGAAGTGGTTGGTGGAAAATTCGGAGGCAAAATTGGGAGCGGCA

GGGGCTAATGAGCTCTCTCCCTTGTCTCCTGATGTTATAATTCTTGATGT

>GH985165.1 JutEST5123 Jute Plasmid library 20-45 days after sowing Corchorus capsularis cDNA similar to Metallothionein 2b, mRNA sequence

GCAACCCTTGCACTTGTAAATGGGAGAGAAAAAATCTGAAGATCCCCTGAAAACAGCAGAGAGTCCTTAA

TTTCCCTTAACAAATAATATCTAGTATGTCTTTCGGACAAGATTATCTTGTTCCTGTGTTTGTTTAGAAT

AAAGTAGCCATGGGTTTTATCTGTCGGCAAAATCCTCACCAGGATTTGTCTGGTGCAGCAGCCATGGTTG

CTTGCTTGGTGTGTATAAGCATAGTAATAGTAATTTAGTTTTCTTATGTCTAATGTAAGCATCTCTGCTG

TCTCAGGATGATGACTCTATCTATGCTTCTTATTGTAAGAAAGTGAGTCGGTTTTTCTAATGAAATGAAC

GGAAAACTCCTTTCTCTTT

>GH985164.1 JutEST5122 Jute Plasmid library 20-45 days after sowing Corchorus capsularis cDNA similar to Metallothionein-like, mRNA sequence

GATAGCGATCCCATAGCAAGCAATCTTCAATAACCCAAAAGCTCTTCCAACTTCTTTCAAAAAACCTTCC

TCAATTCACCATGTCGGACAAGTGCGGAAACTGCGACTGCGCTGACAAGAGCCAGTGCACCAAGAAGGGA

AACACCATGATCATTGAGACTGAGAAGAGCTACGTTGCCACCGTGGTGATGGAGGCTCCTCCAGCAGAGA

ACGATGGAAAGTGCAAGTGTGGATCTAGCTGCAGCTGCACAAACTGCACATGCGGTGGTCATTAAGCAAA

CATATATATAGAAATGAAACAAATGATCCCATGTTGCAGAAACATGGGCAATATTTGGGACCTAAAATCT

TATTTTGGGTTTGAATAATTTGCGTCTTTCCTGTTTCTGTTGGTGTCTGTGTTTTCATGTCCTGTCTGTC

ATGTTTGTGTCTCCTTATATACTGTGATGTAATGGCCATGGCCATGCCTTTGGCCTTGTCTCATCTTAAT

GAACTATTATTAG

>GH985163.1 JutEST5121 Jute Plasmid library 20-45 days after sowing Corchorus capsularis cDNA similar to Metallothionein 2b, mRNA sequence

GGGGCAGCCATGGTTGCTTGCTTGGTGTGTATAAGTATAGTAATAGTAATTTAGTTTTCTTATGTCTAAT

GTAAGCATCTCTGCTGTCTCAGGATGATGACTCTATCTATGCTTCTTATTGTAAGAAAGTGAGTCGGTTT

TTCTAATGAAATGAACGGAAAACTCCTTTCTCTTGG

>GH985162.1 JutEST5120 Jute Plasmid library 20-45 days after sowing Corchorus capsularis cDNA, mRNA sequence

GAAAGAAAGAAGACGGAGAAAGTAAGTGAAAAGGCGTGCCTTGCTCATAGAGTGGAAGACTCAACATCGT

GAAGTGGTGAATTCTCTCCCCAGACTGATTTATGGGTTTAACACAATTCTTATTGCGTCTAACAAAATTC

CAAAAAGATATGTCATGAATAAGTACAAGATTATTCCAAAGGATAGATGGAAAGTGATGGAGCTAGAATA

GCTAAAATTGGGTAATTTTTAAAGGAAAGAAATTTAATGCCTGTAATCCCAGCACTTCAGGAGTCCAAGG

TGGGTGGATCACGAGGTCAGGAGATCGAGACCATCCTGGCTAACACAGTGAAACCCTGTCTCTACTAAAA

ATACAAAAAATTAGCTGGGCATGGTGGTGGGTGCCTGTAGTCCCAGCTACTCGGGAGGCTGAGGCAGGAG

AATGGCGTGAACCTGGGAGGCGGGGCTTGCAGTGAGCCGAGACCGCACCACTGCACTCCAGCCTGGGTGA

TAGAGCGAGACTCTGACAAAAA

>GH985161.1 JEST5119 Jute Plasmid library 20-45 days after sowing Corchorus capsularis cDNA similar to Temperature-induced lipocalin (TIL) mRNA, mRNA sequence

GAGAATCACGAAAAGCAGATTACCCTTAGCTATCTTAAACCCTAACCAAAATCCCTTCAGAATCTAGCAA

AGAAAAAAAGAAAAACCCAGAAACAATGTCTGAGAAAACAATGGAAGTGGTGAAGAATTTGGACATCAAG

AGATACATGGGAAGGTGGTACGAGATTGCTTCATTCCCATCAAGGTTTCAGCCTAAGGATGGGAAGGACA

CCAGGGCCACCTATACCTTGAATGAAGATGGAACCGTTAAAGTTCTTAATGAGACTTTCGTCGATGGGAA

ACGTTCCTATATTGAAGGCACTGCTTACAAGGCCGATCCCAAGAGTGATGAGGCAAAGCTCAAAGTGAAA

TTCTATGTTCCTCCATTTTTGCCCATCATTCCTGTTGTTGGGGATTACTGGGTTTTGTACATTGATGATG

ATTATCGGTATGCTCTCATTGGTCAGCCCAGCAGGAAATATCTTTGGATATTGAGCAGGACGCCACACTT

GGATGAAGAAATCTACAATCAGCTGGTGGAGAAGGCCAAAGAGGAAGGCTATGATGTGAGCAAATTGCAC

AAGACCCCACAAAGTGATCTCCACCAGAAGG

>GH985160.1 JEST5117 Jute Plasmid library 20-45 days after sowing Corchorus capsularis cDNA similar to EMBRYO DEFECTIVE 2769 (Arabidopsis thaliana), mRNA sequence

GATGACAGAGATCGCAGAAGAGGGAGTCAACTTCTTTTAGAAGGGGCAAAGAGGGATGCTGAGGACCGGA

TTGTTCCACGCAGTATTGATGCTGATGATTCTGATGTTGAAGTTGCTAATGACGATGAAGCGATGATGAT

GATGAGGATGATGATGAGGAGGATGATGAGGCTCTGCTTGCTGAGCTTGAACGAATAAAGAAAGAGAGAG

CTGAGGAGAAGCTCAGCAAGGAAGCACG

>GH985159.1 JEST5116 Jute Plasmid library 20-45 days after sowing Corchorus capsularis cDNA similar to P-rich protein NtEIG, mRNA sequence

TTGTTGGGTTTAGTTAACGTTGTTGTTGGCTCACCCCCAAAGACTCCTTGCTGCACTCTCATCTAAGGTC

TTGCTGATCTCGAAGCTGCTGCTTGCCTTTGCACTGCTATTAAGGCTAATGTTTTGGGCATTAATCTTAA

TGTACCCGTTTCACTGTCTTTGCTTCTTAATTACTGCTGGAAAGAGGTTCCTTCTGGCTACCAGTGTGCT

TAATTATTATTCAAGCTCTTTAATTTCGTCTTACTTTATTATCAATTACTGTCTTGGTACTTTCGGGCTT

ATGTTTGGTCGTTTTATATAGTCGTTTTCGAGATGGTGTGATATGGAATTTGGAGATCTGGGATACGTAC

GTATAGTATTATGTCATACTGCAATTTACTTACTAATTTTTACTTCCTTAATTTGATCAAGGAATTAGGA

AGGCTTCTCACCTTCTAATCCTTTTATATATGCGAAACAAGAATGTATTATTTACATCACTCTGTATCTA

ATAAAATGAGAAGTTTTACTTTCAAA

>GH985158.1 JEST5114 Jute Plasmid library 20-45 days after sowing Corchorus capsularis cDNA similar to Actin 7, mRNA sequence

ATTGTGCTGGATTCCGGTGATGGAGTCAGTCATACAGTTCCCATCTATGAAGGTTATGCACTTCCCCATG

CCATCCTCAGGCTTGACCTTGCTGGACGAGATTTAACAGATGCCTTGATGAAAATCCTGACAGAGAGGGG

GTACACATTCACCACAACGGCTGAACGAGAAATTGTTCGTGACATCAAGGAAAAACTTGCTTATGTTGCA

CTTGACTATGAGCAGGAGCTTGAAACTGCTAAAAGCAGCTCTTCAATTGAGAAAAGCTATGAGCTACCTG

ATGGTCAGGTAATAACAATTGGAGCAGAGCGTTTCCGTTGTCCTGAGGTCTTATTTCAACCATCACTTAT

TGGAATGGAAGCTCCTGGCATTCACGAGACCACCTATAACTCGATCATGAAATGTGATGTCGATATTAGA

AAGGACTTGTATGGTAACATAGTGCTTAGCGGTGGGTCTACCATGTTTCCTGGTATTGCTGATCGTATGA

GCAAAGAAATCACAGCTCTTGCACCAAGCAGCATGAAAATTAAGGTGGTAGCACCTCCAGAGAGAAAATA

CAGTGTCTGGGATTGGTGGATCTATTTTGGCTTCCCTTAGTACATTCCAACCAGATGTGGAT

>GH985157.1 JEST5112 Jute Plasmid library 20-45 days after sowing Corchorus capsularis cDNA similar to 40S ribosomal protein S17, mRNA sequence

AGGAGGAAGAACGTGAGCGCCGCATGGACTTTGTCCCCGATGAGTCCGCTATCAAGGTCGACCAAATCGA

AGTCGACAAGGAGAGCCTTGATATGCTTTCTGTTCTTGGGATGTCTGACATTCCTGGCCTCGTTAAGGTT

GACCCAGTTGCTGTTTCCGCTCCTCAAGTTGGATTCGGCCGTGGCAGTGGTGGACCTGGAAGGAGGTTTT

AAGTGGTGGGGCTCCATGGACTTCTGATGGGTTCTTTAAGATTTTTGCTATGATTTTACATTTTAAGTAG

TGTTTAAAATTTGTGTTTTTTGTTGGAACAATATTGGTAGTTGCTATTATCTTAAATGGTACTGAGATGC

TAAGTTTATGTTTAC

>GH985156.1 JEST5110 Jute Plasmid library 20-45 days after sowing Corchorus capsularis cDNA similar to Epsin, mRNA sequence

GGGGCCTCTTCCTTATCCATAACATTTTAGTACACGGTCCGTTCATCCTGAAAGATCAGCTCTCTGTATA

TCCCTGTTTTGGTGGTCGGAATTATCTGAAACTTTCTGCTTTTCGAGATGATTCTGATTCAGAAACGCTT

GAATTATCAACGTGGGTTCGATGGTACGCAGGATATCTTGAACAGAATTTATCGTCTTCAAGAGTCCTAG

GTTATCACCTTTATTCCTCATCGTGTGCTAATAAGAACAAGATTTGGTCATTGT

>GH985155.1 JEST517 Jute Plasmid library 20-45 days after sowing Corchorus capsularis cDNA similar to Tonoplast intrinsic protein 1, mRNA sequence

AGTGTACGCCACAGCCGTTGATCCCAAGAAGGGAAGTTTGGGAACCATCGCACCAATTGCCATCGGTTTC

ATCGTTGGGGCCAACATTTTGGCTGGTGGGCCATTCTCTGGAGCCTCAATGAACCCAGCCGTTGCTTTCG

GACCAGCTGTCGTTGGCTGGAGCTGGGCCAACCACTGGATCTACTGGGCTGGACCTCTTGTTGGCGGTGG

GATTGCTGGAGTCCTGTACGAGTTCATCTTCATTAGCCACACCCATGAGCAACTCCCCACAACCGAATAT

AATTAAACAGTGGGATTTGATTAAATCAAATAAAATCGTCACATGAAAAGCGGATTTCTTCCGGTTGTAT

TTAATTTGTTGAGTTTTCTTTTCTATTATTTTCCTTTGTAATTTGTGTTTCATTGTGGCCGTTGATCTTT

CATTATCATCCATCATCATGATCATCATGGAAGGTGTTGTGATTGAAAAATTTGCATGCCCCCTGGTAAA

ATATTGTGTGGGAATATATTTATGACCAAAAATAC

>GH985154.1 JEST515 Jute Plasmid library 20-45 days after sowing Corchorus capsularis cDNA similar to Metallothionein 2b, mRNA sequence

GGGGGTTCAAGTTTCCAGTGTTAGCTCTTTGATTTCCCAGCAAAAAAAATCTCTTCAATTCTCTCAATTA

GAAAACTGAAAATGTCTTGCTGCGGAGGAAACTGTGGCTGCGGCTCTGGCTGCAAGTGCGGCAATGGCTG

TGGAGGATGCAAGATGTACCCTGACATGAGCTTCGCCGAGTCAA

>GH985153.1 JEST514 Jute Plasmid library 20-45 days after sowing Corchorus capsularis cDNA similar to Metallothionein-like protein, mRNA sequence

GGTTCAAGTTTCCAGTGTTAGCTCTTTGATTTCCCAGCAAAAAAAATCTCTTCAATTCTCTCAATTAGAA

AACTGAAAATGTCTTGCTGCGGAGGAAACTGTGGCTGCGGCTCTGGCTGCAAGTGCGGCAATGGCTGTGG

AGGATGCAAGATGTACCCTGACATGAGCTTCGCCGAGTCAACCACAACCGAGACTCTTGTTATGGGAGTT

GCACCCCAGAAAGTGCACTTCGAGGGAGCTGAGATGGAAGTTGGGGCCGAGAATGGCTGCAAGTGCGGAG

CTGACTGCAAATGCAACCCTTGCACTTGTAAATGAGAGAGAAAAAAATCTGAAGATCCCCTGAAAACAGC

AGAGAGTCCTTAATTTCCCTTAACAAATAATATCTAGTATGTCTTTCGGACAAGATTATCTTGTTCCTGT

GTTTGTTTAGAATAAAGTAGCCATGGGTTTTATCTGTCGGCAAAATCCTCACCAGGATTTGTCTGGTGCA

GCAGCCATGGTTGCTTGCTTGGTGTGTATAAGTATAGTAATAGGTAATTTAGTTTTCTTTATGTCTAATG

TAAGCATCTCTGCTGTCTCAGGATGATGACTCTATCTATGCTTCTTATTGTAA

>GH985152.1 JEST513 Jute Plasmid library 20-45 days after sowing Corchorus capsularis cDNA similar to Translationally-controlled tumor protein homolog, mRNA sequence

GGCTCTCTCGTCGCCTTTTGCGTCTCTCTCTTCATAAACACAATAAAGTCTTATTCTTTGCGTCGATTTC

TCAAAGAGAGAGACTGCAATCATGTTGGTTTACCAGGATCTCCTTACCGGCGATGAGCTTCTTTCTGACT

CATTCCCATACAAGGAAATTGAGAATGGGATTCTGTGGGAAGTTGAAGGAAAGTGGGTTGTGCAAGGAGC

TGTAGAAGTTAACATTGGTGCCAATCCTTCTGCTGAAGGTGCAGACGAGGATGAGGGTGTTGATGACCAA

GCTGTTAAGGTTGTCGACATTGTCGACACCTTTAGGCTTCAGGAGCAACCTCCTTTCGACAAGAAGCAGT

TTGTTTCTTACATCAAGAAGTATATCAAGTTGTTGACACCCAAGCTGGATGCAGATAAGCAGGAAGAGTT

TAAGAAGAACATTGAGGGAGCAACCAAGTTTTTGCTTGGAAAGCTCAAGGAC

>GH985151.1 JEST512 Jute Plasmid library 20-45 days after sowing Corchorus capsularis cDNA similar to Disease resistance response protein-like, mRNA sequence

GCCAGTGATCGGAGGAAGTGGACTTTTCCGGTTTGCTAGAGGTTATGTTCAGGCGAGAACTCGCATGGTT

CACCAGAAGACAGGAGATGCTACAGTTCAGTATACTTGCTATGTGTTTCATTATTGAAAATTAGACCCTT

TTCTTGCCTCTCTTTATTAAGTTATTGAAACTTTTGGCGAATGCTTGCTTTCATCATTGTGTTGGCCACA

TTAAGTAACGAGTATATTTTTTAGGTGGGGTGGGGTTACTTTCTCGTTACTATGTAAT

>GH985150.1 JutEST511 Jute Plasmid library 20-45 days after sowing Corchorus capsularis cDNA similar to 6-phosphofructokinase, mRNA sequence

GATGGAATTCTATGCACTGTTCTAGGACAAAATGCAGTTCATGGTGCATTTGCTGGATATAGCGGCATTA

CAGTTGGCATAATCAACACTCACTATGCTTACTTCCCCATTACTGAGGTAATTGCTTATGCCCGGGAAGT

GGACCCTAACAGCCGCATGTGGCATCGATGCTTGACTTCTACAGGCCAGCCCGACTTCATCTAAATTACA

AAGGGTTAGTTTAATCAAAAGATCAAAGAAATTTGAGTTTCCAAATTGTTGATTAAAGAAC

>GH985269.1 JutEST52100 Jute Plasmid library 20-45 days after sowing Corchorus capsularis cDNA similar to glycine-rich protein, mRNA sequence

GGGGGCTGGGGAAACGGGATCATAGGTGGTGGATACGGTGGAGGGTTTGGTGGTCCTTCTGGAGGCTACT

CCAAAGGAGGTGTGGTGAGACCAACGGTTGTTTGCAAAGAGAAAGGACCTTGTTTCAACAAAAAGCTGAC

TTGCCCTGCTAAGTGTTTCACCTCTTACAGCAGGTCCGGGAAAGGATATGGCGGCGGAGGTGGCGGTGGA

GGATGCACCATGGATTGTAAGAAGAAATGTGTTGCTTATTGTTAAAAGGGTAACATAAATTATTATGCTA

CTAGAATTTACTATCATTAGTGGTTTGATTACTACTTAAAACTATATATTATGTATGACTATATAAACTA

GATGCATGCAGTTTTATCTAAAAATCATATCATATGGAGTATATACATAAAAATGGAATATTTGGTGTAG

TATAGCTCCATGATTGAGCTTTCTCTTTGTTTTTCTCTTCCGTTTCTTCTCTCCATCTTTATAAATTTTA

CTGCTCGGATGAGTTTAAACTATTC

>GH985268.1 JutEST5299 Jute Plasmid library 20-45 days after sowing Corchorus capsularis cDNA similar to Plasma membrane associated protein-like, mRNA sequence

GGCTGAACAATTGCTCAGCTTTTCTTCTTTCTTGATCTGTCAGCCAAAGAAAATTAGCTTATAAGCATGG

CTTCTGGAGGGTCAAAGTCTGCGGCTTTCATGCTTCTGTTTCTCAATCTTGGTCTCTATTTTATTGTCAC

CATCATAGCAGGGTGGGCTGTTAATCATGGCATAGAGAGAACTCATGAAACTGCTTCTGTTTTGTCACTT

CCGGCGACGATATTTCCCATATACTTTCCAATGGGGAACATGGCCACCGGTTTCTTCATCATCTTCTCCC

TCATTGCCGGCGTCGTCGGCGTGGCGACCTCGGTTACTGGACTCACCAATGTTATTCAATGGGATGAACC

AAATTTACACTCCGCCGCTGCTTCCTCTCTCCTTACCTGGTCTCTCACTCTCCCAGCCATGGGATTGGCC

TGCAAGGAGATCAACATAGGCTGGACCGAAGCCAATCTGCGAACTTTGGAAGTGATGACTATAATTCTGA

GCGCTACGCAGTTATTTTGCACGGGTGCTATCCATGCCGGGGTTGGAGATTTAGTCTCCTAAGGAGACCA

GCCTGTTGCCGGAACAGTTTGATGATCATTATAAATCTCAAGCGCTGAATTCCCATTGCTTGAGTGTGTT

TCATTTGTCTGTTAATCCATTA

>GH985267.1 JutEST5295 Jute Plasmid library 20-45 days after sowing Corchorus capsularis cDNA similar to Heme oxygenase 2 precursor, mRNA sequence

GATCTACTTTTCACACATTGCTGGCGGTCAAGTGATTTGTAAGACAGGTATCTGAAATGCTCCTTGAAGG

AAAGGAGCTGGAGTTCTACAAATGGGAGGGGGATGAACAAGAATTATTGAAAGGTGTACGTGAGAATCTC

AATATTCTTGGAGAGCACTGGTCTCGTGATGATAGTAACAAATGCTTAAAGGAAGCTGCAAAGTCATTCA

AGTTTTTGGGGCAGATAATTCGCTTGATCATCTTATAGACCAGATTGGTTTGCATTACAGCTCTGCTTAT

CTGTAAGGCTACAACCTTTTCCACCTAATAGCTCTCCCCTACCCCCTTCCTACTGTCCATGATATTGGAA

GATCAGCTCAAATCTATTCATATCTCCTGGTAACAAAGCAACAATCCTCTTGCTCAGACTTAAGCTCTGC

TTCCAAATCTCATTGTCTTAGCTAAATATCTTCTTGAGGTTTCTTTTCTTTCTGTTCACCAGGTAAGACA

TTCCAATTGTAGACGTTAACACCACTTTTATGAGTCTTCTATTCACCGAGTTGTCAGTAATCAGGATAGC

CAGCATGAGCGACTGGAGTTATATGTCATGTTGAGGAATATCAACGACATGGTAATGTCATAATTTTACT

TAGGAAAATAAG

>GH985266.1 JutEST6293 Jute Plasmid library 20-45 days after sowing Corchorus capsularis cDNA similar to Omega-3 fatty acid desaturase, mRNA sequence

GAGGGCAGGCAGCAAAACGGAAAACCTCAACTTCCTCCCATTCACCCAATATAAATTTTCAGTTGTTTTT

GCAGAGAAGAAATAGAATCTTTTACGAAAGGGTCCTGAGGTTTTCCTTTAGACCGTATTGATAGCGGCAA

AGAACCCTTTTATTTCTCTCAATCTTAAAGTTGGCAGCTTGGGTTTTAGTAATAGTAGAATTTGATTTCT

CTTGGGATACTTTTGTTTTTGGTTTGTTTTTTTAAGTTTTTTTTTTTTTCTCAAACCCTGAGTTTGACTT

TCTCCAGTTCGGCTTTTAATGGCGAGTTTTGTAATATCAGAATGTGGTTTAAAGCCTCTTCCTCAAATTT

ACGCTAGACCCAGAGCTGGTTTTGTCAACCCTTCAAAGTCCAGATTTTTACATGTAAGCAAGACTTTTAC

AGATCTAAAAGTACAACCCTTTAATAAGTTCTCAACTGGGATTTTTAGGGAGAAAAGAACTGGGGGCTTA

AAGTGAGTGTTCCATTGAGAGTTGCAACCGTTGAAGAAAAA

>GH985265.1 JutEST5292 Jute Plasmid library 20-45 days after sowing Corchorus capsularis cDNA similar to 14-3-3 protein, mRNA sequence

GATCTGTGATGGCATTTTGAGCCTTCTTGAGTCTCACCTCATTCCTTCAGCCTCATCTGCCGAGTCCAAG

GTTTTTTACCTTAAGATGAAGGGTGATTATCACAGGTACCTTGCTGAGTTTAAGACTGGTGCTGAGAGGA

AAGAAGCTGCTGAAAGCACATTGTTGGCTTATAAGTCTGCCCAGGATATTGCTCTTGCTGAGCTTGCTCC

TACTCACCCAATCAGGCTGGGACTTGCACTTAACTTCTCTGTCTTCTACTATGAAATCCTCAACTCACCT

GACCGTGCTTGCAATCTTGCTAAAAATGCCTTTGATGAGGCTATTTCTGAGTTGGATACATTGGGTGAGG

AATCTTACAAGGATAGTACATTGATTATGCAACTTCTCCGGGATAATTTGACCCTTTGGACATCTGACAT

TACGGATGAAACTGGAGATGAGATCAAGGAAGCTTCAAAACGTGAGTCAGGCGAGGGACAGCAGCAGTGA

TGAGTTATAAGTCGATAGCATATGTACTCTGTACTTTTATCTTGCAGGTTTACATAGGTGCAATGTGGGT

TGTTTTAGAAGGTTTTTTCTTATGCATTCATCAACGCTTGGAGTTGAAATGTTCCCAGTGGGATTATATA

TGTGCTTTATTATTGAACAAGAA

>GH985264.1 JutEST5291 Jute Plasmid library 20-45 days after sowing Corchorus capsularis cDNA similar to Putative uncharacterized protein T5K6_60, mRNA sequence

GGTCTTCCATCACAACAAAAAATCGTTCACACAAAAGAAAAAAAGAAGAAACAAAAATGCACAACGAATG

CTGCAACTTCCCATCGCCAAGAGAGAGAGTCAAGGAGAAGTTGAAATCTTCCATCTGCTGCTTCCTACCG

CCGCATCTCAACGACGGCGCCGATATGGAGCCGTTGGAGGATCAGATGAAGTCAAGGACGCCGAGATCCC

CTCACGTGATAAAGGACAAGTGCAGACACTTAATCGCAAGAATCGGGAAACGAGGGCGCAGGCAATACCA

CTCGTCGGATTTCAGTTACGATCCAATGAGCTACGCGCTGAATTTCGAGAACGAGTCCAGCCGAGCCGAT

GAGCTTCCCATTATGAACTTCATGTCGCGGCTTCCCGCCACGCCGGAGCGAACTCCAGAGCCGATGCCGG

TGTCGGTGAGGAGGGG

>GH985263.1 JutEST5290 Jute Plasmid library 20-45 days after sowing Corchorus capsularis cDNA similar to Putative uncharacterized protein, mRNA sequence

GCAAGCCACAAATCCTTGTTCCAGCTGATATACCGGGCACATCTGCCCCCATCACCAATTACCTCTCTCC

TTATGTTGCAAAGAATAACCACTTGGCTTATTCATGTTTTTGCTAACCCTCCATTTCAAAGAAGCCTTAG

AAGAAAAAGGTGTCATTTGAAACATCCACTTGTGAAAGCCTTGATAAAAAAAGGTTCTTTTGAGCCTCCT

CATTTACCTCAGTCTTAAGTCTTTTTGAGTAAGTAGTTGGACAGATCAACTTTGCTGTTCATACTTCATT

TGGGTGTGAAGAATTACCCTGGAAAAAAAAAGTTATGTTAAGATAAGCTTTTTCTTTTTTTCTTTTTAAT

TTATTTAATATATGTTTTTCAATAGGGTAGAAATGTTACAAGCTTTGAGTCAATATGTCGTTTGATGTGA

TGCAAAACAGAAATTTCATTTCTATAATTTATTCATTGTTATAAGGGCGCAAA

>GH985262.1 JutEST5289 Jute Plasmid library 20-45 days after sowing Corchorus capsularis cDNA similar to Fructose-bisphosphate aldolase, mRNA sequence

GAAGCCTAACATGGTTACCCCTGGTTCTGATTCACCAAAGGTTGCCCCTGAGGTTATTGCCGAGTACACT

GTCCGTGCCTTGCAGAGAACCGTCCCTCCTGCAGTCCCAGCTGTTGTTTTCTTGTCCGGTGGACAAAGTG

AGGAGGAGGCCACTGTTAACTTGAATGCCATGAACAAGCTCCAGACCAAGAAGCCATGGGCACTCTCATT

CTCATTCGGCCGTGCCCTTCAACAGAGCACACTCAAGGCCTGGGGTGGAAAGGCAGAGAATGTGAAGAAA

GCCCAAGAGGCATTCCTTGTTAGGTGCAAGGCTAACTCTGAGGCCACTCTTGGTACCTACAAGGGTGATG

CTAAGCTTGGTGAGGGTGCCGCTGAAAGCCTCCATGTCAAGGACTACAAATACTAAGAAGATAGGCTTTT

GTGGTCTCTTTTTAATGTCTATTTTGCAGAGTACATTGCGGTATGAATATTTTTAGGCTTTTAGAACTTT

TTTCCTTTTTGGGTTTGTGTCGTTTCTCATGGAATTTAGAGGGTGAAAATCAGACTTAATAAGCTTGGGG

AGCCCTTAGTGGCTCGTCTGAGTTTGGACATGTTTTGTTTATTTTCAGTTTAAAAACAATATTCTCTGCT

TTTTTT

>GH985261.1 JutEST5283 Jute Plasmid library 20-45 days after sowing Corchorus capsularis cDNA similar to Unknown, mRNA sequence

GAGAGTCGGAGGCCACAGCCACAGCAACAGCGCTCCTCAAAAACCCTAGTTTCCTATCTCTCTTTCGTTT

GCCTGAAGCCATGTCGAGGAGAAAGGTTAGAGAGCCCAAGGAGGAGAATACTAATCTCGGACCAGCTGTA

CGAGAGGGCGAGTATGTCTTCGGTGTTGCTCACATCTTTGCCTCATTTAATGACACATTTATTCATGTGA

CTGATCTGTCTGGAAGGGAAACAATGGTTCGCATTACTGGTGGTATGAAGGTTAAAGCCGACAGAGATGA

ATCATCCCCTTATGCTGCTATGCTTGCAGCACAGGATGTTGCTCAACGTTGCAAGGAGTTGGGTATTACT

GCCCTTCATATTAAACTCAGAGCTACTGGAGGAAACAAGACCAAGACCCCTGGTCCTGGTGCTCAGTCAG

CACTTAGAGCTCTTGCTCGTTCAGGCATGAAAATTGGTCGCATAGAGGATGTGACCCCCATTCCCACTGA

CAGTACCCGTAGAAAGGGTGGTAGGCGTGGA

>GH985260.1 JutEST5281 Jute Plasmid library 20-45 days after sowing Corchorus capsularis cDNA similar to Eukaryotic translation initiation factor 5A isoform IVr, mRNA sequence

GCTCCTCCTTATCTCTCGCATCTTCCTCGAAAAGCTTTGAAATTTCTTAGTGTCTGGTAATCGCCATGTC

TGACGAGGAGCACCACTTCGAGTCCAAGGCCGACGCCGGAGCTTCCAAGACTTACCCTCAGCAGGCCGGT

ACCATCCGCAAGAACGGTTATATTGTCATCAAGAATCGTCCTTGCAAGGTTGTGGAAGTGTCAACTTCCA

AGACTGGCAAGCACGGTCACGCTAAGTGTCACTTTGTTGGAATTGATATCTTCAATGCTAAAAAGCTTGA

AGATATTGTTCCCTCTTCTCACAACTGTGATGTTCCTCATGTCAACCGTGTTGATTACCAGCTGATTGAT

ATCTCTGAGGATGGATTTGTGAGTCTGCTCACTGAGGATGGTAACACTAAGGATGACCTCAGGCTTCCAA

CCGATGAGGCTCTTCTTAAGACGATTAAGGATGGATTTGCTGAAGGGAAAGATCTGATTGTGTCTGTCAT

GTCTTCCATGGGAGAGGAGCAGATCTGTGCTGTGAAGGACATTGGCCCCAAATAAATGTTCCAGCTTCTA

TCATCCTAGAGATTTCTTATTTGCCTAAGTGCTTGAAAACTCAGATTAAATATTAGTTTTTTACTGTTTT

ACTGAGGGGTTCTATATGG

>GH985259.1 JutEST5279 Jute Plasmid library 20-45 days after sowing Corchorus capsularis cDNA similar to Ankyrin, mRNA sequence

GACGAGGGTTCATTGAGATTGAAGATCATGGTCTTAGTAGCAGTTTTTTTCCTTTCATAGTTGTCGAGGA

GGATGTTTGTTCAGAAATCTGTATGCTTGAGAGCATACTTGAATTTGCTGACGCTGATCCAGATGTTGCA

AGACCTGAAATAATGGAAGCCAAAAATCAAGCCATGGACTTCATTCATGAAGTTGGTTGGCTTCTTCATA

GAAGTCAATTGAAGTCTAGATTGGGCCACTTAATCCTAACCCGGATCCGTTTCCTTTAAGGCGGTTTAAA

TGGATTATGGAGTTCTCTATGGACCGCGAATGGTGTGCAGTAGTGAAAAAACTCTTAAATATTCTTCTAG

ATGGGAGAGTGGGTTCAGGGGAGCACCCTTCCCTGTATCTTGCATTAACAGAGATGGGTCTTCTTCATAG

AGCTGTTAGGAAAAATTGCAGACCTCTTGTGGAGCTCCTTTTAAGATTTGTCCCTGAGAAGCATTCGGAT

GCATTAGAATTTGGACACAAGACAGTAGCTGGTGGTGTTCATGACAGCTTCTTGTTTAAACCTGATGTCC

TAGGCCCTGCAGGTTTGACTCCTCTCCACATTGCACTGGTAAAGATGGTTCTGAAGAAGTGTTGGATGCA

TTA

>GH985258.1 JutEST5277 Jute Plasmid library 20-45 days after sowing Corchorus capsularis cDNA similar to ATP-dependent protease catalytic subunit, mRNA sequence

GATCTTAACTGTCGTGGGCCCATTTCAGCGTCAAAAACTTTTTTCCTTTTCACACCAGAAACTATTAACA

ATATTTATGTTATGAAAGAGTACAAAAAAAAGAATCTTTTTTCTTTCTTTTTTTGAAAAAAAAAAACTTT

GGGATTGCTGAATAAATATATAAAGCAACGGAGCCATCATAGTATTTTTTATCCGAAAAAAAAGAAGGGT

GGTAATTGGATTATTTATTGATCTGGGTCTAATAGACTCTATATTTTCTCTTTTTTCGATGAAAGAAAAA

AAGAGAATTCATTCCATTGTAAAGCCGTATGCAATGCGCAAAAAATGCCTGTACGGTTGTTCATCAGGGT

AATGATCCATCAACCTACTAGTTCTTTTTATGAGGACCGAACGGGAGAGTTTTCGCTGGACACGGGAGAA

CTCATGAGAATTCGCCAAAAGATCACAAAAGTTTATGCACAAAGAACGGGCAAGCC

>GH985257.1 JutEST5276 Jute Plasmid library 20-45 days after sowing Corchorus capsularis cDNA similar to Metallothionein 3a, mRNA sequence

GAGACTGAGAAGAGCTACGTTGCCACCGTGGTGATGGAGGCTCCTCCAGCAGAGAACGATGGAAAGTGCA

AGTGTGGATCTAGCTGCAGCTGCACAAACTGCACATGCGGTGGTCATTAAGCAAACATATATATAGAAAT

GAAACAAATGATCCCATGTTGCAGAAACATGGGCAATATTTGGGACCTAAAATCTTATTTTGGGTTTGAA

TAATTTGTGTCTTTCCTGTTTCTGTTGGTGTCTGTGTTTTCATGTCCTGTCTGTCATGTTTGTGTCTCCT

TATATACTGTGATGTAATGGCCATGGCCATGCCTTTGGCCTTGTCTCATCTTAATGAACTATTATTAGTA

CCA

>GH985256.1 JutEST5275 Jute Plasmid library 20-45 days after sowing Corchorus capsularis cDNA similar to Ubiquitin extension protein, mRNA sequence

GCTAGCCGAAAATCCATTTCCCCTCTGCATCTTCTGAGAACACCAAGATGCAGATCTTCGTGAAAACCCT

CACGGGTAAGACCATTACCCTTGAGGTTGAGTCCTCCGACACCATCGACAATGTCAAAGCCAAGATCCAG

GATAAGGAAGGAATCCCACCGGACCAGCAGCGTCTCATCTTCGCCGGCAAGCAGCTCGAAGACGGCCGCA

CCTTAGCAGACTATAACATCCAGAAGGAATCCACTCTCCACCTTGTCCTCCGTCTCAGGGGTGGCGCCAA

AAAGAGAAAGAAGAAGACCTACACTAAGCCCAAGAAGATCAAGCATAAGAAGAAGAAGGTCAAGCTCGCC

GTCCTCCAGTTTTACAAGGTTGATGATTCTGGAAAGGTTCAGAGGCTGAGGAAAGAGTGCCCCAATGCTG

AGTGTGGAGCCGGAACTTTCATGGCTAACCATTTTGATAGGCACTATTGTGGTAAGTGTGGGCTCACTTA

TGTTTAC

>GH985255.1 JutEST5274 Jute Plasmid library 20-45 days after sowing Corchorus capsularis cDNA similar to Unknown, mRNA sequence

GGGGGAAATCTACAGTTCATCTAAAACCCTTGCTCCAACAAAAATCCCTGAAACTCAACGCCGCCTTTAG

TTGACAACTTCAACATGCCTCGCCGAAGCTCTGGAAGGTCTGCTTCCCGCCCAGCTCCTCGCCCTGCTGC

CGCACGCAGTCCACCACCTCAGCCAGTTCAACATGCTCCTCCTCCAGCCCCTGCTCAGAGCAGTGGTGGT

GGATCCATTCTTGGAGGGATAGGCTCTACCATAGCTCAGGGCATGGCTTTTGGCACTGGAAGTGCTGTGG

CTCACAGGGCTGTTGATGCTGTTATGGGTCCCCGTACTATTCAACACGAAACTGTGGTTACTGAGGGTGC

AGCTGCCCCTGCACCAGCAGCCACCACAAATAGTTTTGGCGGCTCTGATGCATGCGGTGTCCACTACAAG

GCATTCCAAGATTGCCTGAATGGCTCCGGAAACGACATCAGCAAGTGTCAATTCTACATGGGATATGTTG

TCCGAGTGCAGGAAGAACTCAGGATCCATGGATGAGTTCCTAAGCTCATTCCCCTATTTGTTGCCATGCG

ATGGAATATCCGCTATTGGCTTTGCTTTTATTTATTCCTGTTTCTTGTCTAATGATGATGAAG

>GH985254.1 JutEST5273 Jute Plasmid library 20-45 days after sowing Corchorus capsularis cDNA similar to Unknown, mRNA sequence

GCCCCGCAATTTAAACTCACTTTCACTTAGATTCCCTCTCAAAAGCAAACTCACCACCAAACTGGATAAA

ACTTTCGCCTTTAACTGTTAAAAAAAGGAAAAAAACAGATTCAGGAAGATCCAAAAGGAATCATGGTTCA

GCTTTTGCCTTCTCCATCTTTTTCCCTCTCTGAGACAGAGCAAGAACATCTCTTGGAGAAGCTCGATGTC

TTCAAGGTTCAGGGTAGGGACAAGGTCGGCCGGAAGGTTCTTCTCATCATCGGCAAGCACCTTCCTGCTC

GGATGGTTAGCAGTGAGGTGATCAAGAACTATTTGGAGGAGAAAATTTACCCCAAGTTAGGGGAAAAGCA

ATTTTCTGTTGTGTATGTGAACACTGATGTGCAGAGGAGTGAGAATTTCCCCGGAATCTCGGCCCTCCGG

TCAATCTATGACGCTATTCCGACCAACGTGAAGGACAACATCGAGGCTGTTTATTTCTTGCACCCTGGCT

TACAGGCCAGGCTCTTCTTCGCCACCTTTGGCCGTCTGTTTTTCAACGGAGGGTTATACTCGAAGCTTAA

GTACGTCAGCAGGGTTGAATTTCTGTGGGAGCATGTGAGGAGAAAGGGAGATAGAACTGCCGGAGTTTGT

GTTCGAACACGACGAGGAGCTGGG

>GH985253.1 JutEST5272 Jute Plasmid library 20-45 days after sowing Corchorus capsularis cDNA similar to Ribosomal protein L29, mRNA sequence

GGACCCAAAGTTCTTGAGGAACCAGAGGTATGCAAGGAAGCACAACAAGAAGAGTGGAGAGTCTGCCGAT

GAAGAGTAAATTTTCAATGGGGGAAATCTAGCTTTGATTAGGTCTCTCTGATTGGATCTTTTGTAGTCTG

TTTTAATGTTTTGAGATTATGGGAATGCTTGTTAGTACTATTATTATTTTTGGAAAACAATTTGAAGTTT

TTAATATGAAGAATATGTTTTGATTTG

>GH985252.1 JutEST5268 Jute Plasmid library 20-45 days after sowing Corchorus capsularis cDNA similar to Lipid transfer protein precursor, mRNA sequence

GCGGACCGCCAAGCCGCTTGCAACTGCCTTAAAAGCGCCGCCGGTTCCATTCCTGGCATCAACTTTGGTC

TCGCTGGAGGACTCCCTGGCAAGTGCCGCGTCAGCATTCCTTACAAGATTACTTTTATTTTAATTTAAGC

TCCAAGGAGATGAAGAGAGTTTACTTTTAGATGGAAAAATAAAATGGGATGTAAAAGTCCATGTGGATAG

CACTCTTTCGTAGGAGAGTTGTCCTAATTTTAGATTTGTTATTTGTACTTTTATCGTGTGTACAAGGTCG

CTTTTATATATGCGACTATAACTTTCATATATGAAAAATTAATGGTTGCTAT

>GH985251.1 JutEST5266 Jute Plasmid library 20-45 days after sowing Corchorus capsularis cDNA similar to Ribosomal protein L15, mRNA sequence

GCTCTCTCTTCTCTAAAACGAAAACCCTAGCCACTCTCTCCGCAGCTTGCCAATGGGGGCTTACACTTAC

GTTTCGGAGCTATGGAGGAAGAAGCAGTCCGATGTAATGAGGTTTTTGCAGAGGGTTAGATGCTGGGAAT

ACCGTCAGCATCCATCAATTGTCAGGGTCAACCACCCTACTCGTCCTGACAAGGCACGCCGCTTGGGTTA

CAAGGCCAAGCAGGGTTATGTTGTTTATCGTGTTCGTGTTAGGAGGGGTGGTCGTAAGAGGCCTGTTCCC

AAGGGTATTGTGTATGGTAAGCCCACAAACCAGGGTGTTACCCAATTGAAGTTCCAGCGCAGCAAGCGAT

CTGTTGCAGAGGAGCGAGCTGGTCGGAAATTAGGAGCTCTCAGGGTTGTTAACTCATACTGGATCAATGA

GGATTCCACTTACAAGTACTTTGAGGTTATCCTGGTTGATCCTCAACACAAGGCCATTCAAAATGACCCA

AGGATTAACTGGATCTGCAATCCTGTTCATAAGCACAGGGGAGCTCCGAGGACTCACTTCTGCTGGTAAA

AAGAACAGGGGTCTGCACGGGGGAGGGTCACTTGTACCACAAGAATCGTCCTTCT

>GH985250.1 JutEST5265 Jute Plasmid library 20-45 days after sowing Corchorus capsularis cDNA similar to Putative uncharacterized protein, mRNA sequence

GATCCTTTCATTGCTTATCCTTAGATTTTCGGATTTATAGCTAAGTCATCAAGTTTATTTAATCCAACTC

CCAAAATTAAGTTGAATCAGAGAGGGAAGAAAGAGAAACAGAGAGATGAAAATAACAGGATCCATTAGAT

TTTTTCCTATCTTCTTGATCACTTTAGCTACTTTGCTGTTTATCAGCACAAGTGTGTCCAGAGCAACTGC

TTCTGAAAGCGAAGGAAATGCTATATTTATTTCAGAATACATAGAAGGTGAAGAAGAAGAAGGGATGTCG

TTGGGGGATCCGGAGCCAAGCAGAGGATTCTTCAAGGATTTTACTTTTCTGCAAGGTATATTTCTTATCA

GAGCTTAGAAAAACAACCAATTTGCAATGAAAACATCTACGGAAACTGCATAAAACCGATCAATAAGAAT

CTTCGGCCTTGCACTTTATACAATCGGTGCAAACGGGGTAGCTGATTTCTGGGGAGCGATGTCACAGATT

TGAGTGATTCCCATGCCGTTCTTTGTAAGTTAGTGTGATCAGAAAGAATGTTTCAGTTTGTTCTGTTTGT

ACGAGAGAGTTTAAGAAATGTTAGAATTTTGTGAATGATAATTGTCATTTGTACCAAGCTATATATA

>GH985249.1 JutEST5262 Jute Plasmid library 20-45 days after sowing Corchorus capsularis cDNA similar to 60s RIBOSOMAL PROTEIN L27A, mRNA sequence

GTTTCTAAGACAGCTGAGAAGAAAATTAAGGAGGCCGGTGGGGCTGTTGTTCTCACTGCTTAGAGGTTTT

AAGGTTTTAAGTTTTAGGGTGGATTTTTAGTTTTGTTTTCTAAATGTTTATTATTAGCACTTAAATTGGT

ATTTTGATTTGAGTTTTGTTTTTGGTATGAACTTGGATACTGATATATGAGTTCGTTTCTTTTTATTAAG

AAATGGAATGGAGTTATCTTTTTTTCCGA

>GH985248.1 JutEST5256 Jute Plasmid library 20-45 days after sowing Corchorus capsularis cDNA similar to Fiber protein Fb19, mRNA sequence

ATTGAATTCAAAGCCCGAGTCAAATTAATCTCAAGTCCAAAAGCAAACTTTCCAGGAAGAATTAGGGTTG

AGTGAGAGCTGCAAATTAATGGCGACGTCCGAGAAGGAAGTGATGATCGTTGCAATCGACGACAGCGAGC

ATAGCGTCTACGCCTTGGAATGGACGCTTAATCATTTCTTTACTCCTTTCGCTTCCAATCCTCCTTTCAA

GCTCGTCATCGTTCACGCCAAGCCTACTGCTACTTCTGCTGTTGGTCTCGCTGGACCTGGCGCTGCTGAG

GTATTGCCGTATGTGGATGCAGATTTGAGGAAGATTGCTGCTAGGGTTGTTGAGAAAGCCAAGGATCTTT

GCATTACTAAATCGGTGAATGATGCTGTTGTTGAAGTGATCGAAGGTGATGCTAGAAATGTTCTTTGCGA

GGCCGTAGAAAAACACCATGCCTCAATGCTTGTTGTTGGCAGCCATGGTTATGGAGCTATAAAGAGGGCT

GTTCTTGGCAGTGTGAGTGATTACTGTGCTCATCATGCTCACTGCTCTGTGATGATTGTGAAGAAGCCTA

AGATCAAACATTAATTAATTGTTCCATGAAACGAGCCTCCCAATAATGTATTTGTTTCACTTTGGTTATG

CAATTAATAATGATGGTGC

>GH985247.1 JutEST5254 Jute Plasmid library 20-45 days after sowing Corchorus capsularis cDNA similar to Eukaryotic translation initiation factor 5A isoform VI, mRNA sequence

GCTCCTCCTTATCCTCTCGCTATCTTCCTCGAAAAGCTTTGAAATTTCTTAGTGTCTGGTCAATCGCGGC

CATGTCTGACGAGGAGCAGCACTTCGAGTCCAAAGCTGACGCGGGAGCTTCGAAAACTTACCCTCAACAA

GCCGGTACCATCCGCAAGAGCGGTTACATTGCCATCAAGAATCGTCCTTGCAAGGTTGTGGAAGTGTCAA

CTTCCAAAACTGGCAAGCACGGTCACGCTAAGTGTCACTTTGTTGGAATTGATATCTTCAATGCTAAAAA

GCTTGAAGATATTGTTCCCTCTTCTCACAACTGTGATGTTCCTCATGTCAACCGTGTTGATTACCAGCTG

ATTGATATCTCTGAGGATGGATTTGTGAGTCTGCTCACTGAGGATGGTAACACTAAGGATGACCTCAGGC

TTCCAACCGATGAGGCTCTTCTTAAGACGATTAAGGGATGGATTTGCTGA

>GH985246.1 JutEST5253 Jute Plasmid library 20-45 days after sowing Corchorus capsularis cDNA similar to Unknown, mRNA sequence

GCACAAAACAGGGTTTCCTGGACATGGACTCTGAATCAAGCCTGGAGAATATCATCAAGAAAAATGGCTG

GTGGACAAGAAGCAACTGGGCATTCCTGAACGAACCACCTGTTCTTGAACGCCCTACCAACAGTTACAAA

GCCCAGTTCCACATAGCCAACCTTGCTGCTTCAAAGCTTAATAGGGATTCTGGAATCAGTGCTTAAATCC

TCAGCTATCTTAATCTTAATCCTACCTTGTGCTAGAAATATATAGCATAAAACTTGTTCTGCCTAAAATC

TGTATATATAATGATGTTCTACAAAGTTTGTAAGAACATTACCCTGATGTGTCGGCTGAGTGTGTTTAGT

CATGAGTGTTGTCCTGATCTGGCGTCTGTACAAAACTAGTAGTAAAATGAGCTGTGTAATTTGCAGGTGG

ATGATATTTCGTAAGCTGTCTGCATTATATAAAGATTTGTTTGAGATTTTTAT

>GH985245.1 JutEST5252 Jute Plasmid library 20-45 days after sowing Corchorus capsularis cDNA similar to Gip1-like protein, mRNA sequence

GAACCAAACCCCTCTTTCCCTTCTTCACTCACTTTCACTTCTCTTCTCTTTTCTTTGTCTTATTTTGACA

GTTTCAAGAAGAAGGGTTTTGCTTTGTAGCAAAAGGCCATGGCTAAACTTGTTGCTGCCTTCTTCTTGGC

TCTCATTGCCATCTCCATGCTTCAAACCATGGTCATGGCTTCACATGGACATGGAGGTCACCACTATGAT

CAAAAACCATTATGGTCCTGGAAGCCTCAAGAGCTACCAATGCCCATCACAATGCACGAGGAGATGCAGC

AAGACCCAATACCACAAGCCCTGCATGTTCTTCTGTCAGAAATGCTGCAGGACATGCCTGTGTGTGCCCC

CAGGGTATTATGGGAACAAAGCTGTGTGCCCTTGCTACAACAACTGGAAGACCAAGGAAGGAGGACCCAA

GTGCCCTTGAAGACCCCTTCCTTTAACATCCCTATTTTCCTACTTATCAATTCCTTGAATCATAAGTTAA

GCTAATGTTCTACTAACTACTCTATACTATATACT

>GH985244.1 JutEST5251 Jute Plasmid library 20-45 days after sowing Corchorus capsularis cDNA similar to COP1-interacting protein 4, mRNA sequence

GTCCACTGTTTACAACACTGTCTTTGTAGAAACCAGCCTTGACACTCACCTCGCCATGCATGTCTCCGAC

TCCGACACTGTTTCCGATTTCAAAGACAGATTATGCAAGGAGCACCCATTGTGCTTTCCTAGAAATTGGA

GAAATCAAGATTAATGCCTTGAAGGTAAAGCGACAAAGGATATCTTATCACCTTTCAGATTCCATGTTTG

TAAATAAGGCTTTTGATGGAGTGAGCAAAAGTTGGTTTCTTTCTGTTGATGCCATTAGAGGCGACGAGTG

TCAGAAATATGCTATAATTCCCGCAAACCTGATACTGGCAACACTATAACCTGTTTGGGTATCACAACTA

ATAATTCTTCAGCTGATGTGGTCGATCTTCTACCTGACGGTCCTTCAAAAAGATTGCGCAACATCAAGGA

TTCCTCATTGCGCCAAGATGAAAACAACCACCTTGCAGAACAGACGTCTGCGCCTCAACCATTAGGTTTT

GGTAATTCTGCTGAAGCAAATTCAGAAGATTTGCCTATGGAAGTTGATCAGACGGCTGACTGTAATTCTA

AGGTTTTATTCCCTGTTACCAATATTGGGTCAAGGCCAAAGGTACAGGGATCAAGATGTTGCAAATAAGA

AAATTTCTGATGATCTA

>GH985243.1 JutEST5249 Jute Plasmid library 20-45 days after sowing Corchorus capsularis cDNA similar to Unknown, mRNA sequence

GTCGGCAACCGATATAGCGGTCTCGCTGCGTATGATTTGTCAACGTGGAAGCGACTTTTTCTTACCCACC

TCCCTAGTGATGAAAAGTCCTTTGCGGATGATGTTGCAGTAGATGCAGATGGTACTGCATATGTTACCGA

TACCAAAGGCAGTAAAATTTGGAAGGTTGGTGTCAATGGTGAACTCTTGTCTGTCGTTAGGAACCCACTC

TTCAATCCCAAAGAGTGGTACAAAACCTTGTTCGGATTGAATGGAATCGTTTACCACCCAGATGGATACT

TGATTGTAATCCATACTTTCTCTGGTAATTTGCTGAAGATTGACCTCACCAAGGGTGAGGAAGTTAAGTT

GATTCAAGTTACAGGAGGTTCACTTCAATTTGGTGATGGTATAGAGCTAGTTTCTCCAACAAAGCTTGTG

GTTGCTGGGAACCCTTCTGGGAGATTGGTTGAGAGCTCTGATGGGATGGGAGACGGCTTCCGT

>GH985242.1 JutEST5247 Jute Plasmid library 20-45 days after sowing Corchorus capsularis cDNA similar to Metallothionein-like protein class II, mRNA sequence

GGTTCAAGTTTCCAGTGTTAGCTCTTTGATTTCCCAGCAAAAAAAATCTCTTCAATTCTCTCAATTAGAA

AACTGAAAATGTCTTGCTGCGGAGGAAACTGTGGCTGCGGCTCTGGCTGCAAGTGCGGCAATGGCTGTGG

AGGATGCAAGATGTACCCTGACATGAGCTTCGCCGAGTCAACCACAACCGAGACTCTTGTTATGGGAGTT

GCACCCCAGAAAGTGCACTTCGAGGGAGCTGAGATGGAAGTTGGGGCCGAGAATGGCTGCAAGTGCGGAG

CTGACTGCAAATGCAACCCTTGCACTTGTAAATGAGAGAGAAAAAATCTGAAGATCCCCTGAAAACAGCA

GAGAGTCCTTAATTTCCCTTAACAAATAATATCTAGTATGTCTTTCGGACAAGATTATCTTGTTCCTGTG

TTTGTTTAGAATAAAGTAGCCATGGGTTTTATCTGTCGGCAAAATCCTCACCAGGATTTGTCTGGTGCAG

CAGCCATGGTTGCTTGCTTGGTGTGTATAAGTATAGTAATAGTAATTTTAGTTTTCTTATGTCTAATGTA

AGCATCTCTGCTGTCTCAGGATGATGACTCTATCTATGCTTCTTATTGTAAGAAAGTGAGTCGGTTTTTC

T

>GH985241.1 JutEST5245 Jute Plasmid library 20-45 days after sowing Corchorus capsularis cDNA similar to Putative uncharacterized protein, mRNA sequence

AGGGTTTAGTTTTAGTGCAAAACCCCCGAACTCTGAACTCTGAACTCTGAAGTAGAAACTAGAACGAAAC

GAAAAGATCTTTGAGATTTTCTCATCTCGCAAAGATCCCAATGGCATCGTTTGGCGCCGCCGCAAGATCA

GTTCTCCGATCAGGCTCTGCTCGCAACGCTGCAGCTCGGCTCGCACCCCAAGCCAAAGCAGCACCTGCGC

CGTTTCGTGTCTCCTCTAGGACCCCTCTCTCCAACCGCATTTTCAGGTGTCCAGTTGAAGCGAGCTTTTG

CGTGGAATCGATGATGCCGTATCACACTGCGACTGCTTCGGCTTTGATGACTTCGATGCTCTCCATTTCT

CGCCAGAGCTGTGGTTGGTTCCCTGAAGCTTGCAATGATGATGTGTGACTAATGCTCAGAGGATTCAAGC

AAGTTTATTCCACACAATTCTCCATTTTATGAGAGTGACTGCAGAAAACATAGAAGTTTTAAGTGAGCTG

ATGAGGAGCCAAGTTGAATTTCATAAACTTTTAAACCTAGATGGGACCAGTTTGCATTAATCCTAAGCTC

CTAAAAAGTTGCTAATACTCTATTTCAATCTTTTTAATGCCTTTTGTAGCACATACATTGGATGCAA

>GH985240.1 JutEST5244 Jute Plasmid library 20-45 days after sowing Corchorus capsularis cDNA, mRNA sequence

GTTGAGGAACGAGTTTGGAAGTTGCTGATGGGATGGGAATAAGTATAATCTCAACATTTGTTCCTCCCGG

TTGGCTATGGCCCCTGTACAGTATACTTTGCCCTGCCACGCAATGTAGTTGATGTAAATTACGGCTCACC

TGCAACTAAAATTTCCCATCTTACCATTCCAGCTTCGATCCTGTGAGGCTGACCTTGTGATATAGAGGTG

AGTTGGTAGTAGCAAAGAAGGTAAAAGTAATAGTGAAGCTCTTATGGCAATGTATACATCAAAGTTGTAG

CATAGAGGTAGATATCATGCATATCCTTGTTTTGTAAAGCATCTTTCATGATGGACCCTTTGGTGGGTCA

GTTGGGTCTAATCAAGCAATCCGACGAAATTTCGACATCAATTCTCATTTCCCTGTACAGAAAAATGCAG

AACACATGTAGAAAAGAATAGCATTTTCCAGTTTCCGCATATTATCAATATTATGCCCTCA

>GH985239.1 JutEST5242 Jute Plasmid library 20-45 days after sowing Corchorus capsularis cDNA similar to T-complex protein 1, alpha subunit, mRNA sequence

GCAGGCGGAGGTGCTGTTGAAGCTGCTTTGTCTGTGTACTTGGAGTATCTAGCCACGACTCTAGGATCAC

GAGAACAGTGGGCCATTGCGGAGTTTGCGGAATCTCTATTGATTATACCAAAGGTGCTTGCGGTCAATGC

TGCCAAGGATGCCACTGAATTAGTTGCAAAACTTCGGGCTTATCACCACACCGCTCAAACCAAGGCTGAT

AAGAAACATTTTTCAAGCATGGGTCTAGACCTTTCCAACGGTACTGTTCGTAACAACTTAGAAGCTGGAG

TCATTGAACCCGCAATGAGCAAAGTGAAGATCATTCAGTTTGCTACTGAAGCAGCCATAACAATTCTTCG

CATTGATGATATGATCAAGCTTGTCAAAGATGAAAGTCAGAATGAAGAAAGGGATAAGGGGGGTTTGGTT

TCAACGTTAGTGATGTTATCTCTGTATCTTAATTTTGGGGAAGGGATTGGGATTCCATTTGGGTTAGATT

>FK826618.1 JutEST4019 Jute Plasmid library 20-45 days after sowing Corchorus capsularis cDNA similar to superoxide dismutase,complete, mRNA sequence

TAATTAGGATCCAGAAAAGGACCATTCATTGTAAGCAATGCATACAATTGCAAATATCTCTGATTCACTG

ATGGTAAATCTGTTTCGGAGTGCCATAAGATTTTACTTAAAAGAAGTCTTCAAGAAAACATCCGACATTC

AATTCATAAGAAGAGGTCTTTCACCAACTCAACCAATCACAACGAAGAAGGTCTCTTGAAATATGGAGAG

ACATGGAAATCATAAATGAAAACACGAAACATAATCAACCCAAAAAATTTTCCATTTTCATTTAAAATGC

CTAGAACTGAACAGAGAGAGATAAGCAAACTTGTTTAAATGCTTGATATTGTATTCCCATCCCTTGTGAT

TAATTGTAGTTGTCTTCATGCATAGTTCAACCTGAACATGTCATTCAACACATAGTAGCTCCCCTGTGGT

GTTGGCATCAAATGGAACATCTGGCTGAACTTGAGAGCATGCTGTTCGCCGGCGAGCTGGAGATTACCAC

TGACGAAGACGAGCATGCCACCGGCGGCGCCGGAAGGCA

>FK826617.1 JutEST4018 Jute Plasmid library 20-45 days after sowing Corchorus capsularis cDNA similar to Solanesyl diphosphate synthase 2, mRNA sequence

GATCATTGAGTCTGAGTTTTGTGAGCCTGGATCGCTTGAAGAAGCGATTGAAATAGTTAGAAATCGCGGT

GGGATCAAGAAAGCTCAAGAATTAGCAAAGGAGAAAGCTGAACTTGCGTTAAAGAATTTGAATTGTCTTC

CTAGGAGTGGTTTCAGATCGGCTCTTGAGGATATGGTAATGTTTAATCTTGAAAGGATTGATTAGTTTCA

AAGGGTATTTTCAAGAGTCTACAATGTCTCCCTCAGAGCATGTTTCGGTTAGCTCTCGAAGACATGGTGA

TGTTTAATCTTGAAAGGATCGATTAGTTAGACAACATTTTCACAGTACAAGCAAATTCCTTGAAATGGTG

GACCATTATTCTTAAATCTTAAACATACCCAAAAGCATAAACAGAGGATGCATGAAGTTGAAGCCTCTTA

CTCATATAGAAAATCTTTTTCTTAGCACAATAGTTGGTTAGTCCAGCTGAATCCAGGTTCCTTGTGACTG

GTTTTTTTTTTTTACACCATAGGACCACAACAATAACTCTAAAAATAAATTAAGAAAAACGGAATTTAAT

GAAAAAAAAATAAAAGCCAAAAAAGAAATTTTTGGGAGGTGGAAATTAAAAAAAAAAAAAAGAGCAAAGG

GGGCCAACCATGTGATTGATGATGAAGACTGGAATATCAATCCTCAACCGGATA

>FK826616.1 JutEST4017 Jute Plasmid library 20-45 days after sowing Corchorus capsularis cDNA similar to Glycolate oxidase 2, mRNA sequence

TAGAATATACAATGAAGGCAGCAGCAGCCATAAGATAATCATCATACAAAAGTATTAACTCATTGTTGAA

ACATAAAATTTTCAAAAGATTTGAAGATCCAGGTCTAGTTGAACGACCAAGTAAAATCAACCCAACATGC

TATGATTTGCCAAAAGTAGATAATTAAGCATGCAGATACTAGATACTCTGAGATCATGCATTTACAACCT

TGGGGCCGGAAGACCTTTAGGAGTGTCCCAATCTGTCACAATGTGATTGCGGGTGATTTCTTTCAGTGAA

CGGCACCTACTCAGCGCCATGGTTAGCTCAAATTCCTCCCTCAGCATTGACAGTACTTTCCTCACCCCAG

CTTCTCCCTCAGCAGCCAATGAGAACACCACAGGTCGCCCAATAAATATTCAGAGGCTCCCAGGGCTAAG

A

>FK826615.1 JutEST4015 Jute Plasmid library 20-45 days after sowing Corchorus capsularis cDNA similar to Hybrid proline-rich protein, mRNA sequence

TTTTCTCTTCAAAAACAGAGAAAAAAGAGGACCATTCTAAGCACTTAGTTGTGTCACTTGAAAAATGGCT

TCAAAATCTGTAGCATCCATTGCTCTTCTTCTCACTCTCAACCTGTCTCTTCTTCACTTTGGTCAGTTCA

ACCTACACCCCTTGCCCACCACACCAAGAGAGCCACAAGCCTACCCACAAAAGCCCACCCTCGGCACCTG

AACAGCCAGGCTCACCCAAAAAGCCACACCCCGACCCCCCCAAGCCAAAACCCAACACCCAAGCCGCCAG

CCTCTTGCCCTAAAGACACTCTCAAATTGGGAGTTTGTGCTGATTTGTTGGGTTTGGTTAACGTTGTTGT

TGGTCCCTCCAAGACTCCATCCTGCACTCTCCTCGAAGGTTTGGCTGATCTTGAAGCTGCTGCTTGCCTT

TGCACTGCCATTAAGGCTAATGTTTTGGGAATTAACCTTAATGTTCCCGTTTCATTGTCTTTGCTCCTTA

ATTACTGTGGAAAGAAGGTTCCTGCTGGCTACCAGTGTGCTTAATCTAAAAGCTACTTTAATTTCGTCTA

TACTTGTAATAATCAATTACGTGTTCTTGGTATTTCTGGGCTTGTTTGGTTTTTTATAGTCGTTTGTCAT

GGGTGATATTGGAAGTTTAAGGAATCTGGGTACGTTAACGTATAGTCATTATGTCATCTATTTATACTAT

ATTTTTACTTTTAATTTGATCGAAGGAATTAGGAAGGCTNCTCACCTTCTANTCCTTTTATATATGCGAA

ANAAGAATGTATTATTTACATCACTCTGTATCTAANAAAATGAAANGTTTTACTTTTATCAGCGGGCCGC

GAAATNANTTCACCAAGNGGTGGAATGNTNCNCCCNCNAAANGGGANCGNGANCTGGGNATNAACCNCNC

NNGAAAANG

>FK826614.1 JutEST4014 Jute Plasmid library 20-45 days after sowing Corchorus capsularis cDNA similar to Caffeoyl-CoA-O-methyltransferase, mRNA sequence

ATGCCAAGATCTTGGCCATGGACATTAACAGGGAAAACTACGAGTTGGGTCTGCCTGTAATCCAAAAAGC

AGGCGTTGCACACAAAATTGAATTCAAAGAGGGCCCTGCTTTGCCTGTTCTTGACAAACTAGTCGAAGAT

GAAAAGAATCATGAATCATATGACTTCATCTTCGTGGATGCTGACAAAGACAACTACATAAACTACATAA

ACTACCACAAGAGGTTAATAGACCTTGTGAAAGTTGGAGGCTTAATCGGCTACGACAACACTCTATGGAA

TGGTTCGGTCGTCGCACCACCTGACGCACCAATGAGGAAGTATGTTCGGTACTATCGGGATTTCGTCTTG

GAAATAAATAAAGCACTCGCTGCCGACCCAAGGATCGAGATTTGCATGCTACCTGTCGAAGATGGTATCA

CAGTATGTAGGCGGATCCAATGAGGGATTCTGGCTATCCAGCTAG

>FK826613.1 JutEST4013 Jute Plasmid library 20-45 days after sowing Corchorus capsularis cDNA similar to Fatty acid desaturase 2, mRNA sequence

AACGCTCCATCCCACGCTCCTTCGTTCTTTATGACCTCTATCATCTCCTCCTCGGCTCCTCCCTTGCAAC

TAACTATTTCCACAGCCTACCTTACAATCTCCCCTACGTCGCCTGGCCGCTCTACTGGGCCTGCCAAGGC

TGCGTCCTAACGGGCCTCTGGGTCATAGCCCACGAATGCGGTCATCATGCCTTCAGCGACCACTAATGGC

TTGACGACGCCGTCGGCCTTGTCCTACACTCATCTCTCCTCGTTCCCTATTTCTCCTGGAAGTACAGCCA

GATCTCGC

>FK826612.1 JutEST4012 Jute Plasmid library 20-45 days after sowing Corchorus capsularis cDNA similar to DELLA protein GAI1 (Gibberellic acid-insensitive mutant protein 1), mRNA sequence

AAACAGGGGATGCAGTGGCCCGCGCTGATGCAAGCCCTTGCCCTGAGAGAAGGCGGTCCGCCCTCGTTCC

GGCTAACCGGAATTGGGCCTCCCGCGGCGGACAACACCGATCATCTGCATGAAGTGGGTTGGAAATTGGC

CCAGCTGGCGGAGGCGATTCATGTGGAGTTCGAGTACAGAGGTTTCGTGGCCAACAGTTTGGCAGATCTC

GACGCATCCATGCTTGAGCTTCGACGAGCA

>FK826611.1 JutEST4011 Jute Plasmid library 20-45 days after sowing Corchorus capsularis cDNA similar to Digalactosyldiacylglycerol synthase, mRNA sequence

AAATTGCCGGACTTACCAAGACGGACAAGGTTTTGTGAAAGCCACGCTGAAAGCTCTAAGTGAAGAGCCT

GCCCCATTGACTGAGCAACAACGGCATGAACTGTCATGGGAAGCTGCTACAGAGCGGTTTCTAAGTGTAT

CTGAGCTCAAACAGGTATTGATAAAAAATTCGAAAATTTCTCAGAGAAGAAAGTTTGCATCAACGTCTTT

AAACCTACGAAAGAATATGGATGATACATCAGCATATTTTCATCATGTGGCCTTAGGATTT

>FK826610.1 JutEST4010 Jute Plasmid library 20-45 days after sowing Corchorus capsularis cDNA similar to Delta-12 fatty acid desaturase, mRNA sequence

ATGACCGACAAGGAGCGGGAGAAGCAGGAGCAGCTCGCCCGAGCTACCGGTGGCGCCGCGATGCAGCGGT

CGCCGGTGGAGAAGCCGCCGTTCACTCTCGGTCAGATCAAGAAGGCGATCCCGCCTCACTGCTTCGCGCT

GGCCATCATCGCGTCCTTCTCCTACGTGGTTCATGACCTGGTCCTCATCGCCGCCCTGCTGTACTC

>FK826609.1 JutEST4009 Jute Plasmid library 20-45 days after sowing Corchorus capsularis cDNA similar to Delta-12 oleate desaturase, mRNA sequence

ATGGGAGCTGGTGGCAGAATGCAAGTCTCTCCTCCCTCCAGCTCCCCCGGAACCAACACCCTCAAACGCG

TCCCTGCTGAGACACCACCATTCACTCTCGGAGACCTCAAGAAAGCAATCCCACCTCACTGCTTCAAACG

CTACACCCTTTCTCCTTCGTTCTTTATCACCTCATCATAGCCTTTCTCCTTTCCTCCCTTGCAACCACCT

ATTTCCACAGCCTCCCTTACAATCTATCGTACCTCGCCTGACCCCTCTACTGGGCCTGCCAAGGCTGCGT

CCTAACAGGCCTCTGGGTCATAGCCCACGAATGTGGTCATCACGCCTTCAGTGACTATCAATGGCTTGAT

GACGCTGTTGGCCTTGTCCTACACACATCTCTACTTGTTCCCTATTTCTCTTGGAAATATAGTCA

>FK826608.1 JutEST4008 Jute Plasmid library 20-45 days after sowing Corchorus capsularis cDNA similar to Fiber protein Fb9, mRNA sequence

CCCATCTTCTGCCCTCGGCATCAACATCCTCCAATGGTGGCATGTCAGCATCAGCTTCAGCTTCACCAGC

ATCCTCATCAATGCTCATCCCAATTTCAACATTCTGTGGATTCTGTTGCCGAAGTGTTGGGGTCATCCAG

GCTGANGCCAGAGGTGAGAATTTCAGTCTCAAAGAGCAGGAGAACAAGGTCCTTGACAGACTTGTCATTC

TTGTCAGCATCAGCTTCTTCTGCTACCTCTAGTTGGGTCTCTTGGGGTAGTACCTTCTCAACTCCTTCTC

TCGACTACGACTGTTCTTACTGTTCAGACAGTTCCTGGAACAGAGGACGAGAAACTCTGACGGGAAGAGT

GGAGACNNNAGTCGGACCTACTGGGGTTGTGGAAGCCGTTGTCTTAGGTGTCTTACAACTTTAACCCTAA

CTCGTAACTACTCCTACGACCACTTCGACTTCGACTACGACTGTACGGTGGTAACCTCCTACAACTACGG

CTCCCAGTCGTTCTACCTCTCGCCAACCTTAGCTCGAGCC

>FK826607.1 JutEST4007 Jute Plasmid library 20-45 days after sowing Corchorus capsularis cDNA similar to Unknown protein, mRNA sequence

TCCCCGTAGGAGTCTGGGCCGTGTCTCAATCCCAGTGTGGCTGATCATCCTCTCGGACCAGACTACTGAT

CATCGCCTTGGTAAGCTATTGCCTCACCAACTAGCTAATCAGACGCGAGCCCCTCCTCGGGCGGATTCCT

CCTTTTGCTCCTCAGGCTACGGGGTATTAGCAGCCGTTTCCAGCTGTTGGTCCCCTCCCAAGGGCAGGTT

CTTACGCGTTACTCACCCGACCGCCACTGGAAACACCACTTCCCGTCCGACTTGCATGTGTTAAGCATGC

CGCCAGCGTTCATCCTGAGCCAGGAT

>FK826606.1 JutEST4006 Jute Plasmid library 20-45 days after sowing Corchorus capsularis cDNA similar to Bimodular protein, mRNA sequence

GCAATTCATGCAAACCCAAACCTCCTCACAGCTTGCGGTTCATGCAAACCCAAGCCCAAACCCACCCCTA

GTCCTCCTCCAGCTCCAAAAGCTTGCCCTCCACCACCAAAACCTGCGTCTTGCCCTAAGGACACATTGAA

GCTCGGTGTTTGTGCTGACGTCCTTGGACTTGTCAACGTCATTGTCGGAACACCTCCTTCAAGCAAATGC

TGCGCTTTGCTTCAAGGCTTGGCTGATGCCGAAGCTGCCCTTTGCCTTTGCACTGCCATTAAAGCCAATG

TACTTGGAATCACCCTCAACATTCCTGTTACTCTTAGCTTACTCCTTAGTGCATGCCAAAAGGAAATTCC

TCCAGGCTTCAAGT

>FK826605.1 JutEST4005 Jute Plasmid library 20-45 days after sowing Corchorus capsularis cDNA similar to Solanesyl diphosphate synthase 2, complete, mRNA sequence

GAAGCGATTGAAATAGTTAGAAATCGCGGTGGGATCAAGAAAGCTCAAGAATTAGCAAAGGAGAAAGCTG

AACTTGCGTTTCAAGAATCTGAATTGTCTCCCTCAGGCATGGTTTCGGTTAGGCTCTCGAAGACATGGTG

ATGTTTAATCTTGAAAGGATCGATTAGTTAGACAACATATTTCACAGTACAAGCAAATTCCTTGAAATGG

TGGACCATTATTCTTAAATCTTAAACATACCCAAAAGCATAAACAGAGGATGCATGAAGTTGAAGCCTCT

TACTCATATAGAAAA

>FK826604.1 JutEST4004 Jute Plasmid library 20-45 days after sowing Corchorus capsularis cDNA similar to Peptidyl-prolyl cis-trans isomerase, mRNA sequence

TCTCAAATAACCTTTATTTTCTCTGTTCATCAGGTTATTTTTCTGTCTCCCAAATTTCCCTACAGAATTT

TATCGCAAATCCTCAACCATGTCGTCCAAGCAAGGTGGCAAATTGAAGCCTTTGAAGCAACCCAAGGCTG

AGAAGAAAGAACTCGACGAGAATGATAAGGCTTTGCTTCAGAAGAAGAAGGAAGAAGAGAAGGCTCTTAA

AGAGCTGAGAGCAAAGGCACAACAAAAGGGATCATTTGGTGGAGCTGGACTCAAGAAGAGTGGGAAGAAA

TGAGGGTCTGACAAACCCCAACTAAACAGAACTTATTGAATATTGTCTTGTGATTATTACTATTTATCAT

TATGATGACTCTTTTGTTTAGTGTGATGCTCTGGCATGACTTGATGCACTCGTTTAATTCTGAATATGCT

ATGCTAAACATTGTTTCATATTTCATTATTTAAAACTTAATAAATTTAAGTTCTACGTTTGTGGCTTTTG

TCTGTGTTTTAAAAAAAAAAAAAAA

>FK826603.1 JutEST4003 Jute Plasmid library 20-45 days after sowing Corchorus capsularis cDNA similar to Acyl-CoA-binding protein, mRNA sequence

GATGAGCACGCTGAGAAACCAAAGACCCTACCAGAGACCACAACAAACGAAAACAAGCTTATTCTCTATG

GACTGTTCAAGCAAGCAACCGTTGGACCTGTGAATACGAGCCGTCCTGGAATGTTCAACATGAGAGACAG

GTACAAGTGGGATGCATGGAAGGCTGTTGAAGGGAAATCAAAGGAGGAAGCTATGAGTGATTATATCACC

AAGGTGAAACAGTTGCTAGAGGAGGCTGCTGCATCTTCTTGATGTCTGACATAACATTGTGTGTTCCCAG

CCTTGCTACATGTTTCTGAAAATGTAAAACTAAAAATAAGTGTTTGTACTTTTATTGTTGCTACCATGTT

ATTCCCGAATTATGCAGGTTTAAACCAGATTTGATTTATCTATCAATAGTATAAACAAGCCCAAATAAGA

TGTTTA

>FK826602.1 JutEST4002 Jute Plasmid library 20-45 days after sowing Corchorus capsularis cDNA similar to Hybrid proline-rich protein, mRNA sequence

TTTTCTCTTCAAAAACAGAGAAAAAAGAGGACCATTCTAAGCACTTTTGCATTAAGTCTCAAAGATATCA

CCAGACATGGCTTCAAAATCTGTTGCATCCATTGCTCTTCTTCTTACTCTGAACTTGTCTCTACTTCACT

TTGGTTAGTTCAACCTACACCCCTTGCCCACCATCACCCAAAAAGCCACACCCCGACCCCACCGAAGCCA

AAACCCAAACCCAAGCCGCCGCCTCTTGCCCTAAAGACACTCTCAAATTGGGAGTTTGTGCTGATTTGTT

GGGCTCGGTTAACGTTGTTGTTGGTCCCTCCAAGACTCCATCCTGCACTCTCCTCGAAGGTTTGGCTGAT

CTTGAAGCTGCTGCTTGCCTTTGCACTGCCATTAAGGCTAATGTTTTGGGAATTAACCTTAATGTTCCCG

TTTCATTGTCTTTGCTCCTTAATTACTGTGGAAAGAAGGTTCCTGCTGGCTACCAGTGTGCTTAATCTAA

AAGCTACTTTAATTTCGTCTATACTTGTAATAATCAATTACGTGTTCTTGGTATTTCTGGGCTTGTTTGG

TTTTTTATAGTCGTTTGTCATGGGTGATATTGGAAGTTTAAGGAATCTGGGTACGTTAACGTATAGTCAT

TTGTCATCTATTTATACTATATTTTTACTTTTAATTTGATCAAGGAATTAGGAAGGCTTCTCACCTTCTA

ATCCTTTTATATATGCGAAACAAGAATGTATTATTTACATCACTCTGTATCTAANAAANGGAAAAGTTTT

ANTTTTATCA

>FK826601.1 JutEST4001 Jute Plasmid library 20-45 days after sowing Corchorus capsularis cDNA similar to Ribulose bisphosphate carboxylase small chain, mRNA sequence

TGGAGCACGGATTCGTCTACCGTAAGAACCACAGCTCCCCAGGGTACTACGATGGACGCTACTGGACCAT

GTGGAAACTTCCCATGTTCGGGTGCACCGACTCATCCCAGGTGTTGAAGGAGCTTGAGGAGGCCAAGAAG

GAATACCCCGAGTCCTTCATCAGAATCATTGGATTCGACAACAAGCGTCAAGTGCAATGCATCAGTTTCA

TTGCCTACAAGCCCCCAAGCTTCACCTCAGCTTAAATGAACAGAGTTGTTTTCCCCACATTGTAGGGATT

CGGCTTGTTTAAATTGTACTTAGCATTTGTCGAAACCTTTTTACTTTTCCTTTCTTTTTGAGGTATTGTC

CTTTGTTTTTTT

>FK826600.1 JutEST4363 Jute Plasmid library 20-45 days after sowing Corchorus capsularis cDNA similar to AAF02153.1, mRNA sequence

GGGACTGTAGTCTATAGTTATCCCTAAAAAAGAACAACCTTAAATTAAAAGTTTTAAAACCCTTGTGAGA

AACCTCAGGGAAACCCCAAGTTTCTGTCTTTGTCTTTCGCAAGATCCTCTGCTCCAAACAATGTCTCTAT

CTCTGAAAACTCTGACACCCATATCTTTCTTAACTCTCTTTTTCATCCTACCATTCACAGTCACCTCTTC

CATTCAAGATCTCTTAAAGAGTCAAGGCCTTCCCGGTGGTCTTTTCCCCGACAATGTCAAATTCTACGAA

CTCGACCGTGTAGGCCGCCTTGAAGTCCACCTTGAACAGCCGTGCATGGCGAAGTTTGATGGCAGAGTCC

ATTTTGACCGTGTGGTGAGAGCTAATCTCAGTTATGGTGGCCTTATGGGTCTGGAAGGCCTTTCTCAGGA

AGAGTTATTTTTGTGGTTGCCTGTTAAAGGCATCATTGTGAATGATCCTTCTTCTGGGCTTATTTTGTTC

GATATTGGTGTTGCTCATAAACAACTCTCTCTTTCTCTCTTTGAAAACCCTCCTGTTTGCAAGCCTCAAG

AAGTTCTGGCACAGAAGGTTGGGGAGGAAGAACAAGGGGATTTCAAGTTCTGCGAT

>FK826599.1 JutEST46142 Jute Plasmid library 20-45 days after sowing Corchorus capsularis cDNA similar to F21J9.22, mRNA sequence

AAGAATTCTTGTTGGGAAAACACAGGATTTGGTGGAAGAGAAGAGCCATGAACTGAAGGGTTTGAATGAA

ACTGTTAGTCTATTAGTTAAGGAGAAAGAACATATTGGTACTTTACTGAGGAGCGCATTGTCCAAGAGGA

TGGTATCGGAAAATAATTCTAAAACAAATGAGTTGTTTCAAACTGCAGAGAATGGTTTAAGAGAGGCTGG

AATTGATTCTAAATTTGCTAAACTCTTAGGGGATGGGAATAAATCTGAAGCTCAGACTCAAGAGCATGAT

GAAATATATAATTTGGCCAGCACATTGGAGAATATTGTCAAAACATCTCAGCTCGAGATCGTCGAACTGC

AGCACTTGTTGGGACAATCAAGGCAAGAACCTAGTGTACTTAAAGAACATGTGGAGGCCCAAGCCAAGGA

GCTCAACCAGAGACTGCGTCGTATAGAGGAGCTTGAAGAGAAGGAGAAGAAAGAAAAACAGAGAGATGAA

AATAACAGGATCCATTAGATTTTTTCCTATCTTCTTGATCACTTTAGCTACTTTGCTGTTTATCAGCACA

AGTGTGTCCAGAGCAACTGCTTCTGAAAGCGAAGGAAATGCTACATTTATTTCAGAATACATA

>FK826598.1 JutEST44120 Jute Plasmid library 20-45 days after sowing Corchorus capsularis cDNA similar to Hypothetical protein T14E10_70, mRNA sequence

GGCATCCAGTTCCTTGTTCCAACCACTCTGTCCTTCAAAAATCTTCTCAAGAAAATCAATTTCAGCATGT

TGAGGGAACTGATGTTGAAGACCTTAGCACTCTTGCCTCTTTTGACCCAAGTTCGCCTATAGAGCAAGAT

GATTCTAGCACAGTATCAGTTGCGATTGATGATTGTTCTATGGAGGACACAGTACTTTACCGCCTACAAG

ATGTAATTGCGAAGTTGGATGTCAGAATTAGACTGTGCATAGGAGATAGTTTGTACCGTTTGGCTCAAAG

TGCAATGCAGAGACATTGTGCCAGTGATACAAACAGCACTAACAAAAGTAGCAGGGATGAAAATGAAGTG

GCAAAAGAGGACAATAAGAACCATAACAGGATGTCTGATGCAGAAACAGGGACTAACCCCATAGACCGTA

CTGTTGCTCATTTACTCTTCCATCGGCCTCTGGAGTTGCCAGGGAAAGCATCCTGAGACGCCTGAATCAC

CTGGTTCCGCAAAATTTCAATCTGAGCGCAAATCAGCAGGTTTAGTAGGTTTGCCAATGGGTTCCATATC

AGACAATCCACAAGTCAAACAAAACGTAACTCACCAGGTTTCAAAAGGTCCTTCCCCATTTATTGGATAC

CCAACATGTGGAGCAATTCAAGA

>FK826597.1 JutEST4442 Jute Plasmid library 20-45 days after sowing Corchorus capsularis cDNA similar to unknown protein, mRNA sequence

GGGGATTGAAACATTTAACAATGGCGTCAGAGAAAGACCACACAGACCAACTTCGTACGCTGACCTTTTT

TCCAGTGCTAAGGTGGTTGCGGAGGCCGCAAAATCAGCTTCGCGGCAAGAGGCGGACAAGTTCGATAAGG

CGAAGGTTGCCGAAGCCGCTTCTGATCTTTTAGATGCAGCCAATAGATATGGCAAGCTTGAGCAGGATAA

GGGAATTGGTCAGTATGTTGACAAGGCACAGAGTTACCTTCACCAATTCGAGTCTTTCCAATCAACCACT

AAGGATACTCCGGCACCGGCGCCGGCTAAAGATGGTGAGGCAAAGCCAGAGTCGCATCCAAGCGGTGGCC

ATGAAGGCAAGGAGTCTGGAAGCGGTGGCGGTGTTGGGGATTACATGAAGATGGCTCAAAGTTTCTTGGG

CAAGTAATTAAGCTTAATTAGCTATGTTGTTAAATGTTTAGGGTTTTTTATGTACCATAATTTCAATATT

TCCTCTCTTCTATGCTCTGTATGGCTTATAATTCTGTACCCAATCTTCTGTCTAATAATATGCAGCAATA

ATTTGGTACCAAAAACC

>FK826596.1 JutEST4493 Jute Plasmid library 20-45 days after sowing Corchorus capsularis cDNA similar to Putative uncharacterized protein, mRNA sequence

GGGACAACATGTACAAGGAATACCGTGACACTACTTTGAATGGAGCTGTTGAACAGATGTACACTGAGAT

GGCTTCTCGACACAGAGTCAGGTTCCCTTGCATTCAGATCATCAAGACTGCAACCATCCCTGCTAAACTC

TGCAAGAGGGATAGCACCAAGCAGTTCCACAACTCCAAAATCAAGTTCCCCTTGGTGTTCAAGAAGGTCA

GGCCACCAACCAGGAAGCTCAAGACAACCTTCAAGGCTACAAGGCCCAACTTGTTCATGTAATTTCTCTC

ATTTGATATAAAAGGAGAGTTAAAAAAGTGTTTCTAATTCTGAAATTTTTGGTAGCGAACTTATATCTGT

TGGTTGTTGATCGGATATTTGTTTTTATTTGAGATGCTTAGGCTGAACTTGGGATCTTTTGGGTTTCAAT

ATCTCATTTTGTTTTTTAAAA

>FK826595.1 JutEST4532 Jute Plasmid library 20-45 days after sowing Corchorus capsularis cDNA similar to Unknown protein, mRNA sequence

GGGGAAAAACTTGATGATGCTAAAGTTGAGTCTGCACATTCAGTTATTGTGGACTCCTCAGAAGTAGAGC

AGAAGCCACTGAAAAACACATCTGATACGCAGAGAGAACAGGACTCTGTGACTGTTAATCTCTCATACAG

GTCATCCCCAGAAGCCTCTCCAAGGAGCCATTTGACTGTTCCAGAATCCCAGGGAACACCTTCTAGTCAG

GTATCAGTGAAACCC

>FK826594.1 JutEST4747 Jute Plasmid library 20-45 days after sowing Corchorus capsularis cDNA similar to F6A14.17 protein, mRNA sequence

GGAGCAATCCATATGTTATTTGTTTGGCCGATTTTCTTCACAGCTATTCTCCTTCTTGACTATACTCCTT

CTCTGTTTAATCTCCCCCAGATCGATTTTTCTCTTTTTGGAAGTCACGTGGCTTTGGTTTTCAACTTTGG

GTTCTTGGTTACTCTAATCTATGCCGTGTTTTACGTGTGTTTGGATGTCAAAGCTGGAACCTTGGCGGCT

CTGCTTTGTTGGGTCTGTTGGATTGGTAGCAGTTTTCTTGCCACTCGACTTGGGTTTTCCTTTGCTTGGA

AAATTGTCCTGGTGGTTCAAATAGTGTGTTGGACTGGACAGTTCATTGGCCTTGGGGTCTTTGAGGGACG

AGCACCGGCTTTGTTGGACAACCTTGTTCAAGCCTTGATAATGGCACCATTTTTTTGTGCTCTTTGAGGC

ACTTCAATTCTTTGGATATGAACCATACCCTGGGTTTCATGCAAGTGTTCAATCAAAGATTGAGGCTGAA

ATCAATGAATGGAAAGAAAGGAAGCAGAAGTTAATCACTTAGATTATGCTAACATTGTCTTATATATACA

CCCACACATTTATCTGTCATCATGAAATGTGACTCAAGGCATTTGAAGTGACAGTATATGTTTACAGCAA

ACAGTTA

>FK826593.1 JutEST4794 Jute Plasmid library 20-45 days after sowing Corchorus capsularis cDNA, mRNA sequence

ATCTTCCCTTTAACTAATCATTAAAAAGTGCATTTCTCACAACCACTTTCACCAAATATCTCCCCACTGC

CAACACCTCACTGCCACCCATGTCCGCCGCCTCTGCCGCCACAATTTCCACCCCTACCCTCGACTCCGAG

TCCCCTTCCCTCCTCCAATCCATCACCTCCCATGGCGGCTACGCCTATGCCCGCATGGCCACTTTAGCCG

CTTCCGGCGACCAACGCGCTGCCGAGGCGGCGCGTGAGATGGCCTGGGAGCAGCTTCATTCTGGCCCTTG

GCACTCCGTTCTCCCCGTCTGGCGTGACGCTTACTCCATGGCTTGCCTCTACGTTGCCAAGTTCCATTTC

TCCAATGGTGAGTTTAGAGATGCTCTTAGGGCTTTGGATATGGGTATCATCATGGGCGGTCCTTTGCTTA

GGAAAGACTTGGACTCGGCGATCGAAGTCGTTTCGGCCGCTAAAGCAAGAGTTGATCAGAACGGTGGAGC

TGCTGCTGGTAATGAGGGTTCTCAAAAAGCTGAGTCTCGATTGCTTGTTTCTCCGGAGGAGATTGATAAG

TCCGAGTTACTACGGCTTTTACCAATAAGGTCCTTGTCTTCTAAGATTGTGGGGAAGAGGTCTGCCCTGT

CTTTGGAGGGATTTCTA

>FK826592.1 JutEST4766 Jute Plasmid library 20-45 days after sowing Corchorus capsularis cDNA similar to Expressed protein, mRNA sequence

GAAGCTTGTGGGCATGGTTTGGGAGTCGGTAATGTTGCATTTTCCGAATCATGCTCTATAGAGGGTGGAG

ATTTCCAGAAACTTGCAAACTCGCATTCAATCCATGATCATCTAGTTTCAAGAATGGAAAAGAGGGAAAG

GGGGCAGACTGATTTGGTTGTCTTATGCCATGGAGGATCTCATTCTGTGGAAGAGCTTGACCAGCCTCGC

CCTGAGAGTGAGATTTTGTCTGAAGTTATCAGTTCAGTGGAGCAGTCTGGGACAAAATATGCTGCTCTTT

ATGTTTCTGATCCTTTGAAGCCGGTTCACTATCCTGCTTATAGAGAGCTTGAAAGGCTTCTCGCTGAAAG

TGCAGCACGAAATGGTTCAGCACCTTCCACAGGATGTGATGAGGTTTGCAAGATAAAATCATCACTCCTA

GAGGGAGTTTTAGTTGGAATCGTGTTGCTTTTAATTTTAATATCAGGCCTTTGTTGTATGATGGGCATCG

ACACTCCAACAAGATTTGAGACACCCCAAGAGAATTGATTGGCTGTTTGTGCCCCAGCATCTTTATTTGT

ATTAGGCTTCTGTCCTATGGACCGTTGGGAAGGTATTACTAGTTATTAGATAGTTGCAATGGGCAGGAAT

GTT

>FK826591.1 JutEST47137 Jute Plasmid library 20-45 days after sowing Corchorus capsularis cDNA similar to Unknown Protein, mRNA sequence

ACTGTAGTCTATAGTTATCCCTAAAAAAGAACAACCTTAAATTAAAAGTTTTAAAACCCTTGTGAGAAAA

CCTCAGGGAAACCCCAAGTTTCTGTCTTTGTCTTTCGCAAGATCCTCTGCTCCAAACAATGTCTCTATCT

CTGAAAACTCTGACACCCATATCTTTCTTAATCTCTTTTTCATCACTCCATTCACGCACTCTTCCATTCA

AGATCTCTTAAAAGCTCAAGGCCTTCCCGGTGGTCTTTTCCCCGACAATGTCAAGAGTTCTAGTCGAGAT

GATAAGATGGCCGCGCTTGAAGTCCACCTTGAACAGCCGTGCATGGCGAAGTTTGATGGCAGAGTCCATT

TTGACCGTGTGGTGAGAGCTAATCTCAGTTATGGTGGCCTTATGGGTCTGGAAGGCCTTTCTCAGGAAGA

GTTATTTTTGTGGTTGCCTGTTAAAGGCATCATTGTGAATGATCCTTCTTCTGGGCTTATTTTGTTCGAT

ATTGGTGTTGCTCATAAACAACTCTCTCTTTCTCTCTTTGAAAACCCTCCTGTTTGCAAGCCTCAAGAAG

TTCTGGCACAGAAGTTGGGAGGAAGCAAGGTGGATATTCAGTTCTAGCGATGAATGCCATTGCTACATTT

TTCATGGAGAT

>FK826590.1 JutEST47123 Jute Plasmid library 20-45 days after sowing Corchorus capsularis cDNA similar to Unknown Protein, mRNA sequence

AATATTGCCTCGGATGGAACTAATGGTGAAGAGCAGGAAAAGACATGTGTTGTTCCAGATCGCGAACAGT

ATAACAGAGAAGAATTTCCTGTGGATTACACCGATGCCAAGTTCTTTGTGATTAAATCATACAGTGAAGA

TGATGTCCATAAGAGCATTAAATACAATGTTTGGGCCAGCACACCCAATGGTAACAAGAAGCTTGATGCA

GCTTACCAGGAAGCTCAGCAGAAACCTGGTGGCTGTCCTGTATATCTTTTCTTCTCGGTCAACACAAGTG

GGCAATTTGTGGGTGTTGCTGAGATGGTAGGGCCTGTTGATTTTCAGAAGAATGTAGAGTACTGGCAACA

GGACAAGTGGACTGGCTGTTTCCCAGTGAAGTGGCACATTGTAAAGGATGTACCAAACAATTCTTTGAAG

CACATCACCTTAGAGAACAATGAGAATAAACCTGTCACGAATAGCACGGATACACAAGAGATCAAGCTTG

AGCAAGGACTGAAGATTGATCAAAATTTTCAAGGAACATTCCAGCAAAACTTGCATTTTGGATGATTTCT

GATTTTATGAGGTCCGCCAGAAGGCAATCCAGGATAAAAAGGCTAAACAGACTCAAAAGCAGGTATGGGA

GGAAAGCCTG

>FK826589.1 JutEST471 Jute Plasmid library 20-45 days after sowing Corchorus capsularis cDNA similar to AT5g57120/MUL3_6, mRNA sequence

GGGCCTGCGCTGATAGCTTTGAAGCCTCGCCAAGTTGTGCTTTCTTCCAACTCTGCCAAAACCATGAAAA

ACAAGAGCTTCCCCTCTGCCTTAGATTCCCACCAAAAGTCTCTTCTCCTTCTCTCAGTCGCTCAATTCCT

CCAACGCAATTCCTTCTCCAAAACCCTCAAAAAGTTCCTTTCTGAAGCCCAAATCCAGAAAAACGATTTG

AACAGGTTCTTCCTTGATCTGGAAGAAATGTGCTGCAAGTATTTGGACAAGAGTGATAATTCAGCCAAAA

TTAATGCAGCCAGCGAGTCCAAGATAGTGGAAATGGCAATAGTGAATGGACCAAGTAGGAGGACTTGTAG

CTGTTGAACAATGCTACTAAGAAGAAGGTAAAGGAGGTAGAGAAAGTAAAAGTGATAAATGCTGCGAATC

AAGAAAGAGGGAAGGAGCAGTCTCTTATGAAGCATACAAAGGCGATCTTAACTGATGATATTATGGCCAG

TCACACTGATGATCTGATTGAAGAAACAGTTAAGGAGAAAAAGAAGAAGAAAAGCAAGGTGGATTCAGAT

GATGCTCATGTTGATAAGCAGAATGTTTCAAGGTGGGGGAATCGGGAAAAATCTAAAGTTGCAGTATCTA

>FK826588.1 JutEST4670 Jute Plasmid library 20-45 days after sowing Corchorus capsularis cDNA similar to Hypothetical protein, mRNA sequence

TGCCTGAAGGTCAAGCTACCAAAAGACAGAAGCTGGAAGTTGGTCTACTTAGCAGGGTTGCTCACTTGAC

GCATCAAGCTAGTTTTCTGCACAGGAAACCTAAGATGGATCCAACCCTTGATGGTAACTCAATACATGCC

AAGCCAAAGATTACCGTTCCACAGGAACCAGAGCTACAAACGGCAATAAGGGCTGAAAGACTACGGTACT

AGAGTTACGGTAAGTCGGATGAAGTTCAACATCACGGGCGCATTCCTTTAAAGCACTCCCCTTAAATAGA

AAAATTTTTGACTCTCCATCATGGCCCCTTCCTAAGAAGAGCACTCCTCAGCTACCAGAGTTTCAATTAT

TTCACCTGAGGACATCAGAGAGAGCCAGGCAGCATGCATCAACAATGACAATGAGACTACCTAATTACGT

TTCAACTTC

>FK826587.1 JutEST4652 Jute Plasmid library 20-45 days after sowing Corchorus capsularis cDNA similar to Predicted protein, mRNA sequence

AAGAGAATGGTACTGGCAAGGAGAGCCCTCATTGTGAAGATGGTGATCTGCCAGAAAGTTTGGAGCACAC

CCTTCCTGATAGTCCTTTTGCATCTCCTACAAAAGCCCCTGGAGAGTACTTGTCTGCTTTCTCAAATGCT

GAAGCAGACAAGGCTGCTTCAGTTAACAATAATTGGGTTTCTTCAACCTTACTTCCTGCGCATAATGCAT

TAGACATGGCATCCTTGAACTCCTTCAACTGCCAAATGCCCAGGTTCTCGTCTGGACATGGGACCATTGG

CATGTATGCGGGCACTGGCGGGCCGACTTGCCCTGTTGGAATTTAGCTCCTATGCTTAGGAAGATTGAAA

GTATAACCAAAGAAAAGAAAATTTCATATCAGAGATCTGTAGAGAAGCGCCACTACAAAGAGAAGCACGT

CAACCATTGTGATGAATACACCAGTATACACTACTTACTACCATACAAAAATGCATACGTAAAAGGCAGT

GAAGGAAGTGGGTCAGCCATTCCTTTCTTTTGTTGTAATATTGTTATCTATTGTTTCAAGTAGTTGAAGT

TTGGACCACAGGTAAAAAAGCACGG

>FK826586.1 JutEST4651 Jute Plasmid library 20-45 days after sowing Corchorus capsularis cDNA similar to Unknown Protein, mRNA sequence

GACGGCTGCTGACAGAGAGTTGGCTGGCTTTGTTGAATTATTGAGATTTTGGAGATCTGAAACGAAAGAG

TGCTGAAACTCATCGGAGGAGAAAAGGGGAACATAGAGAAAACACAGGAGCGAAGATGATTCACCGGAAA

TGGAGTTTGCTAACAGGCCCAGTCGTCATCATCGGCGGAGTCGTGGGAGCCGCTATTGCCGCTCACCTCC

TCTTCGTTGACGATCCATATCTTAAACCTAGGGTGAAGAAGCCGGATACACAACCATTGACCAAGTAGAA

AGACATTTACAGTTGCTCGGTTTTCATAATCTCTCTTGGAGCAGGCAAATGCCTAGGTTGCTGTTTGAAA

CATGAGTTGTTTGTTCAATAAAATCCAATCTGATTGTTGTGTAGTTTGGGGGTATCTTTCTTTTCTCTTT

TTAATAACAACTCAACTGAAATGTTCCTAAACTTTGAATGTCAAAACAGTTTATAATGTTTCACCTGGAA

GTTTTGCATTCGAAAAA

>FK826585.1 JutEST454 Jute Plasmid library 20-45 days after sowing Corchorus capsularis cDNA similar to Hypothetical protein, mRNA sequence

GGCCTGTTAATGCCTTCCTTCCAAGCATTCTTGGCAACAATTTGGTGGAAAAAACTTGCAAAAAGACACC

CTTTTACAACCTCTGCCTTCTCCCTCTACAATCAGACCCTCGGAGTTCCAAATGCTGACGTGGCTGGGCT

AGCCCACATCAGGAGCCATAAAGTTAAGGCCAGAGCCACTCACACACTAAGGCAGATTAACAGGCTGCTT

AAAGCGAGTCAAGACCCAAAGTTGAAACGAGCCTTGCACTGATGTGCCTGTTGATACAACACCATTATTG

TAGTTGACATTCCGGTAGCCATTGATGCTTTGTCAAAGGGAATCCACAAGTTTGGGGTGGCAGGTGCAGC

TGATGCAGCCAATGAGGCTAGATGATGTGAAAGAGAATTCAGGAGATGCAAGTCGCCTATTTCTGGTAGC

AACAAGGCTTGTGCATGATCTTTCTATCATGGTTGCATCCATTGTTCAGTTATTACTGTGAGGCTGCCTG

CCTCATCCTTTCTAAATCTGTATTTATCATGTTTAGTATATTTGATAGATCAATAAAGCTACATG

>FK826584.1 JutEST4519 Jute Plasmid library 20-45 days after sowing Corchorus capsularis cDNA similar to Putative uncharacterized protein, mRNA sequence

GGGAGGCTGGATAAGGATCGGAAGAGTTTGTTGGATCGTAAGGCCAAAGGAAGAGCTGCCGCCGATAAGG

ATAAGGGTACTAAATTTACTGCTGAGGATATTATGCAGAGTGTTGACTGAGTGAAAAGTTTTTAGGGTTT

AGTTGTCTTTGCTGGAATTCCGAAAGTGGTCTTTTTTATTTTTGTTAAATTTACAATGTCATTTGGGATA

TTTAGACTTTGTTGTCAATCTTATGATGGCTTAATGTTATGTTCTAAATTTTGT

>FK826583.1 JutEST4511 Jute Plasmid library 20-45 days after sowing Corchorus capsularis cDNA similar to Hypothetical protein, mRNA sequence

GGGCTGACCAGACTGTTTCAGAAGTCTCTGGGATCGTCTTCTTCATTTCTCACAAGCTGCCAACATGTTT

GACGATCAAGACTTGGGCTTCTTCGCGAATTTCCTCGGAATCTTCATTTTCGTTCTGGTGATTGCGTATC

ATTACGTGATGGCTGACCCAAAATATGCAGGCAATTGAGTATTTTGATCTTTTTGTTTAGGTGTCTGCAA

TGGACATTTTGATGATAGTTTTGTAGACTGAGTTGTTTAAGAAATGAAGACAGAATCAATGTTGTTAAAA

ACCTTAGATATACTATTTCTATAAATTTCATCTTATTGCC

>FK826582.1 JutEST4467 Jute Plasmid library 20-45 days after sowing Corchorus capsularis cDNA similar to unknown protein, mRNA sequence

GGTGAGACAACTTTGTTAATATGCAGTAGTGACCCAGTTATTCTGGTGATATGATGACTTTGCTCATCAA

ATTCCCCTGTTTTCTTTAGCATATAGCTAGGTTTCTCTTCCTTGTTTGCCCTTCGGTTATCTCCACGCCA

ATGGATTCGTTTAGTACTCAGAAACCTAAATCAAGAAAGATGAAGAAGAGGCTTGCGAAAAAGTCTCGAG

TTGAGATAACAGGTTGATGAATGATGATGGTCCTGGCTGGTGGTTGGATGTTAGGATGTATG

>FK826581.1 JutEST4418 Jute Plasmid library 20-45 days after sowing Corchorus capsularis cDNA similar to hypothetical protein, mRNA sequence

GGATTTGCATGGAGCAAATGCAACATTGTATGGCTAAAGCAAGAATGCTGTATGCTGCAAAAGATCAGGC

TTATGCAGATGCAAGTATGCATCAACACATGTTGCTCGACCAGTTGCAACGAAAGGATAGCATTATTGGA

TATTTGCAAAAGGCCAAGTACGAAGAGCAGACTAAGAGACAGGTGGAGGTTCATAGGCTTGAGCATGAGC

TCTTTATGATGGAAAATCTACTAAATGGATACAGAAAGGCTTTAAAGGAAACAAACAAAGCTTTTACTGA

GTACAGGGCACGCTTTCGCCTGTCTGATGAACCACTCTACAAGGATGTTAGCGGGTCAGGGGGGTTAGTT

CTGAGCACTATGGAACTAGAAAAGCAGCTTCTGAAACAAGAAGAAGAAGACAGGCTTAATAAATCACTGA

TTGAGAAAAGGATTAAGGACTTTGAAGCTGGGTGGATTGGTAAATTTGAGGCTCATAAGGATGCAATTTC

TCTATTTAGTATCAGATTGATTGATGCTGAGAAGGATGTTAATCTTCTTAAAGAACTTTTTGCAAATCGC

AAGTCACCAGATACCCCAGAATGTGTTCCATAGAAACTGTAAATGTTTTAGTCTGTAAGTTATGCCTTT

>FK826580.1 JutEST4482 Jute Plasmid library 20-45 days after sowing Corchorus capsularis cDNA similar to 40S ribosomal protein S9, mRNA sequence

GGGCTCACTGACGAGAAGCCTACGCAGCCATGGTTCACGTCAGTTTCTACCGAAATTATGGAAAGACCTT

TAAGAAGCCAAGGCGTCCATATGAGAAGGAGCGTTTGGATGCTGAGTTGAGGCTTGTAGGAGAGTATGGT

CTCCGATGTAAGAGGGAGTTGTGGAGGGTTCAGTATGCATTGAGCCGTATTAGAAATGCTGCGAGGGAAC

TTCTCACTCTGGATGAAAAGAATCCCCGTCGTATTTTTGAAGGTGAAGCACTTCTTCGCAGGATGAATAG

GTATGGACTTCTGGATGAGAGCCAAAACAAGCTCGATTATGTCTTGGCTTTGACTGTTGAGAACTTCCTT

GAACGCCGTTTGCAAACACTTGTGTTCAAGTCTGGTATGGCCAAGTCTATTCACCATGCTCGTGTGCTGA

TTAGGCAAAGGCATATAAGGGTTGGGGAGGCAGGTGGTCAACATCCCATCCTTCATGGTGAGGATGGACT

CCCCAGAAGCACATAGACTTCTCTCTCACAAGTCCATTTGGGAGGTGGCCGCCCCTGGAAGAGTGAAGAG

AAAGAACCAGAAGGCTGCTGCCAAGAAGGCCTCTGGTGGAGACGGTGATGAAAGAGGAAGACGACGAGTA

AACCCTTAGGCTGATATCA

>FK826579.1 JutEST48117 Jute Plasmid library 20-45 days after sowing Corchorus capsularis cDNA similar to Ribosomal protein S21e, mRNA sequence

CGACTCACTATAGGGCGCTAGCTCGCCGCAGCCGAACGACCGAGCGCAGCGAGGACGAAGCATTTTTCTT

TCCTGCCGCCACTGTCTTCAAGGTGATTTGAGCTAAGTGTTTGAGTGTTCGGTTAAGAAAATCAGCAAAG

ATGCAGAACGAAGAAGGACACAACATGGATCTTTATATCCCCAGGAAATGCTCCGCCACCAACAGGCTCA

TCACCTCGAAGGATCATGCTGCTGTCCAGATCAATATTGGACATTTGGATGAGACTGGCAGATACACTAG

CTCATTCTCCACTTTCGCCCTTTGCGGCAATGTCCGTGCTCAGGGTGATGCCGACAGCGCTCTTGACAGG

CTTTGGCAAAAGAAGAAAGCTGAAATCAAACAGCAGTAGTAGGATACTTTCCATTACTAGTCTTGTCTAT

GAACTTGGAATGAAAAGTGTTTCGGCAATGACTATCATGTTTTAGTTTGTTTCAGTGGATTATGTTGAAG

TT

>FK826578.1 JutEST4745 Jute Plasmid library 20-45 days after sowing Corchorus capsularis cDNA similar to ribosomal protein S12, mRNA sequence

GGGTCCGGGTCGAAATATGGCTTTCAAATTAAGTTCCGAATTAGTGGATGCTGCCAAAGGGAGTGGCGAT

GCCATACGCAAAAAGGAAGAGACTCATAGAATGGCAGAGGCAAATAGAGCTTTTGCACATTTTCGTTAAT

CCATGAACAGGATCGATATAGACACATAGATTCATGGATCCATACATCTCGATCGGAAAAGAATCAATAG

AAAAAGAAAGAATCGGAATTGATCAATATATTTCTCGAAACAAACGAAAAGGAAACGAAAGATGAAACAT

AAATCATGGATCAACTAAGCCCTCTGTGGGGGACTTGCTTAAGAATAAGAAAGAGGAATCTCATGTAAAT

ACCATGGAATAAGGTTTGATCCTATTCATGGGGATTCCGTAAATATTCCATTTC

>FK826577.1 JutEST4892 Jute Plasmid library 20-45 days after sowing Corchorus capsularis cDNA similar to 60S ribosomal protein L41, mRNA sequence

ATTCACGCCGCTGCTTCTGTAGAAACAACATCTAACAAGTAGATTTGGTACTGCAGGAGACCCTTTTCGA

AGACTTTTCACCATGAGAGCCAAGTGGAAGAAGAAGCGTATGAGAAGGTTGAAGAGGAAGAGAAGAAAGA

TGAGACAAAGATCTAAGTAGACTACTTTACTTTAGAGCTTATGCCCGTTTAGATTTCTGTTTGTTTGTTT

TGATAAGATCCTTTTGTTAAAAGGATCGAACTTTATCTTCTTGTTTTCTGGGTTTAGTTTGTAACTTTTG

TTGAACCAATTTTCATTTACTGGGAAAGTGTTATTTGAAGTTTAATATGCTTTTGGATCTAAAAAA

>FK826576.1 JutEST4874 Jute Plasmid library 20-45 days after sowing Corchorus capsularis cDNA similar to 40S ribosomal protein S23, mRNA sequence

GGGGAACAACCTCTCTCAGCCAACATGGGCAAGACACGTGGTATGGGAGCTGGTCGCAAGCTGAGGACCC

ACCGCAGGAACCAGAGGTGGGCTGATAAGTCATACAAGAAATCTAACCTTGGAAATGAATGGAAGAAGCC

ATTTGCCGGTTCTTCCCACGCCAAGGGCATTGTCCTTGAGAAGATCGGTATTGAGGCTAAGCAGCCTAAC

TCTGCTATCCGTAAGTGTGCTAGAGTTCAATTGATCAAGAACGGAAAGAAGATCGCCGCTTTCGTTCCAA

ATGATGGTTGCTTAAACTTCATTGAAGAGAATGATGAGGTGTTGATTGCTGGGATTGGACGTAAGGGTCA

CGCCGTCGGAGATATTCCCGGAGTTCGATTCAAGGTCGTCAAGGTCTCTGGTGTTTCACTCTTGGCACTC

TTCAAGGAGAAGAAGGAGAAGCCTAGGTCGTAAGTTTTAGAAGAGGGATGTTCAGTTTTCTTATGGTCCT

TTTATAGTTCTACTTAAACCTGAGCTTTTTATTGCTTCTTAGCTATTATGGACAATGTTTTGTTTATTTT

CTGGTTATTGGATTGAATTTGTCATGTCATCTTGATCAAGAAAGATTTTGGTGCTTTATTGCAAAAAA

>FK826575.1 JutEST48138 Jute Plasmid library 20-45 days after sowing Corchorus capsularis cDNA similar to 30S ribosomal protein S7, mRNA sequence

AGGAGGGAACGCAAAGAGTGACATAGTGTTCGTGACCGAGAAAGAGAGAAAGAGGGGAGTTGGCTGATAA

AGATGGACAGTAACGATTGCGTAATATCAATTTTTCGGCCTCGTCATCGAAAGCGGCTTCCAATTGCTCG

GAAATTCTTAGCTATATGGGGGACTTGGATGGTGAGCAAAAACAATTGATCAAGAAGTTGGTCAACTTTC

GCATGAAAGAAGGTAAAAAAACGAGAGTTCGTGCTATTGTTTATCAAACTTTTCATCGCCTGGCTCGAAC

TGAACGCGATGTAATCAAACTTATGGTTGACGCCGTAGAGAATATAAAGCCCATATGCGAAGTGAAAAAA

GTAAGAAGAGCGGGTACTATTTATGATGTCCCTGGGATTGTAGCCAAGAATCGTCAACAAACCTTAGCTA

TTCGTTGGATCCTTGAAGCAGCTTTCAAACGACGTATAAGCTACAGGATAAGCTTAGAGAAATGTTTATT

TGATGAGATACTGGATGCTTACCGAAAGAGGGGAATTGCACGTAAGAAAAGGGAGAATCTTCATGGACTG

TCTTCCACCAATCGAAGTTTCGCGCGTTTCAGATGGTGGGTAAAGTGAGATCACATAAAGAGCTCTTCCT

CAGACTCCTCCCTT

>FK826574.1 JutEST45141 Jute Plasmid library 20-45 days after sowing Corchorus capsularis cDNA similar to 40S ribosomal protein, mRNA sequence

GCTCATCAAGCAAAACAGAGGCGGCGGCCATTTTTGCTTCAGCAACTTGAGTTTGAAAGGATGGTGAGAG

TCAGCGTTTTGAATGACGCCCTCAAGAGCATGTATAATGCAGAGAAGAGGGGAAAACGACAGGTCATGAT

TAGGCCTTCCTCAAAAGTGATCATCAAGTTCCTTTTGGTCATGCAGAAGCATGGATACATTGGAGAATTT

GAGTATGTTGATGACCACAGAGCTGGAAAAATTGTAGTTGAACTGAATGGTAGGCTGAACAAGTGTGGCG

TGATTAGTCCTCGTTTCGATGTTGGAGTCAAGGAGATCGAGGGCTGGACTGCTAGGTTGCTTCCTTCCAG

ACAGTTTGGGTACATTGTGCTCACCACATCTGCTGGCATCATGGACCATGAAGAAGCCAGAAGAAAGAGC

GTCGGTGGAAAAGTACTTGGTTTCTTCTATTGAGACAAGCTTCTGAGATTTTTATCTACTCTTCATGTAG

AACTTGGTTATTATCCTCTTTTGGTGCAATTCTAAATTAGTAAATTAGGGATATTATAATTTT

>FK826573.1 JutEST45123 Jute Plasmid library 20-45 days after sowing Corchorus capsularis cDNA similar to ribosomal protein 41, mRNA sequence

ATTCTCCAAGCGAATCGTCTCCTTTCTAGCGAAATCGCCTGAAGAATCTCGAGAAAGAAAGCGCTAACCC

AAGATGAGAGCCAAGTGGAAGAAGAAGCGTATGAGGAGGCTGAAGAGGAAGCGCCGAAAGATGAGACAGA

GATCCAAGTAGGCGCATGGACACAGTCAAGATGAGCTTTACTGTGACATTCCCTTGTCTTTGCCACCGAT

CCAGCGTTTGTTGCATGAAGTTGTGAAGATTTGAGGATGTTTTTTCGGGGAAGCTTATCTTTTATGTCTT

CTAAATCTAGGTTTAAGTTTCAAGACTATATGAGATGTTCTCTAGTACACTACTATAAATCTTGTTGGTA

TTTCTGTTTTTGAGATGTGTAGACTATTATCCTTAACATTGCATCCTAAGAGTTTGATATTGGAAACCTT

ATGGTCTTTCTAGT

>FK826572.1 JutEST4461 Jute Plasmid library 20-45 days after sowing Corchorus capsularis cDNA similar to 60S ribosomal protein L39, complete, mRNA sequence

GGGGCTCCAGCAGTTGTTCAAGCAAGAAAACTCCTAGCGCCGGCAGCCGCTCTCCCAACTGTCAAAATGC

CTTCACACAAGACTTTCATGATCAAGAAGAAGCTGGCTAAGAAGATGAGGCAGAACAGGCCCATCCCTTA

CTGGATCCGCATGCGCACTGACAACACCATCAGGTACAATGCAA

>FK826571.1 JutEST4457 Jute Plasmid library 20-45 days after sowing Corchorus capsularis cDNA similar to 40S ribosomal protein-like protein, mRNA sequence

GGGGTTCTCGGCGAGAAAGGTAGGAGGATTAGGGAGCTGACCTCTGTGGTTCAGAAGCGATTCAAGTTCC

CAGAAAACAGCGTTGAGCTTTACGCTGAGAAGGTCAACAACAGAGGCCTTTGCGCCATTGCCCAGGCTGA

GTCCCTTCGCTACAAGCTCCTTGGAGGCCTTGCTGTCCGAAGGGCCTGCTATGGTGTCCTCAGATTCATC

ATGGAGAGTGGCGCCAAGGGATGCGAGGTTATTGTTAGTGGAAAGCTAAGGGCACAGCGTGCCAAATCCA

TGAAGTTCAAGGATGGTTATATGATTTCTTCCGGTCATCCAGTCAATGAATATATTGACTCTGCTGTCAG

ACATGTTCTCTTGAGACAGGGGGTACTTGGTATCAAGGTGAAGATCATGCTCGACTGGGATCCTAAGGGT

AAGCAGGGCCCTATGACTCCTCTACCTGATCTTGTCACAATCCACCCACCAAAGGAGGACGAAGAGATCT

ACAAACCAGTACCTGAGCCAACTGAAATTGAGGTTCCAGTGATGGCCGCTTAAACATAGTTTGTCTATTT

AATTTTGGAGTATGATGTTGAGTTGTCTTTTAGTTTTTGTAGCCAAAATTTGGCCCCAAAGACCTC

>FK826570.1 JutEST43102 Jute Plasmid library 20-45 days after sowing Corchorus capsularis cDNA similar to 60S ribosomal protein L22-2, mRNA sequence

GGGGCTAAAACATTTTGCTATCTTCAAAACGTAGAAGAGGCAGCGAAGAAGCAGTCAGGTATTGCGAAGA

TGAGTCGCGGAGCAGCAGCTGCACCCAAGGGGAAGAAGAAGGGAGCGACGTTCACCATCGACTGCGCGAA

GCCGGTGGAGGATAAGATCATGGACATCGCCTCATTGGAAAAGTTCCTCCAAGAAAGAATTAAGGTCGGT

GGTAAAGCCGGTGCTCTCGGTGATTCCGTCACTATCTCCCGCGACAAGAACAAGATCACCGTCACTTCTG

ACTCCAACTTCTCCAAGCGGTATCTTAAATACTTGACTAAGAAGTACTTGAAGAAGCACAATGTCCGGGA

TTGGCTTCGGGTGATTGCTTCCAACAAGGACCGTTCCGTTTATGAACTCCGCTACTTCAACATTGCTGAG

AATGAGGGGGGAGGAGGAAGAGTAAAGAATCAGTTCAGTATTTGCTCCAGTTACCAATCTGTTTTATTAG

TTGTGAGGACGTTGTTAATCTATTTCAGGTGTCAAATATTTGGACGTGTTTGGGTTGTTTTCTGACCCTT

CTTACTTAATGTTCAATGGAAATTTGTTGCG

>FK826569.1 JutEST4291 Jute Plasmid library 20-45 days after sowing Corchorus capsularis cDNA similar to Acidic ribosomal protein P1a-like, mRNA sequence

GGCTAGAGACATTTGCTTCAGAGCGGCGCTGCAGAGCTTTTTGAGAAGGGTCAATTCCAGGTTTCATTTC

TGAGTTGAAGCGATGGCGTCAGTCGGGGAAGCTGCTTGCTCTTACGCTGCTATGATTCTCCATGATGATG

GCATCCCCATCACCGCTGAGAAGATTGCAGCACTCTGCAAAGCTGCAAATGTCACTGTTGAATCTTACTG

GCCAAGCTTGTTTGCTAAGCTCTTCGAGAAGTGCAACATCGAGGATCTCATAACCAATGTTGGAGCTGGT

GGTGGCGGTGGCGCTGCTCCTGTTGCTGCAGCTGTTGGTGGAGGTGGTGCTGCTGCTCCAGCTGCTGCCG

AGGAGAAGAAAGAGGAGAAGAAAGCTGAGCCAGAAGAAGAAAGTGATGATGATATGGGATTCAGTTTGTT

CAATTAGGAGCTCTTTTTCGGTGTGTGATCTGGTCAATTACTGGTTTAGTGCATTAAGCAATGATAGTAG

TTACTATTTTGAGGTTTATTGGTATTAGTATCTGTATCTGTCTGTTACTAAGGGGAATTCAATCAGAAAT

ATATGTAGTTTTGAATGAAGTACTGTTTGAATTTTGCTAT

>FK826568.1 JutEST41130 Jute Plasmid library 20-45 days after sowing Corchorus capsularis cDNA similar to Ribosomal protein L33, complete, mRNA sequence

GGGCGAATGGCATACATTTACAAGGCCAAAGTGAAGCAGAATGGCTCTCATTACCGTTGTATTTGGGGTA

AAGTAACAAGGGCCCATGGTAACAGTGGCGTTGTTCGTGCCAAGTTCAAGTCCAACCTCCCTCCCAAGTC

CATGGGTGATAAAGTGAGAGTTTTCATGTATCCCAGCAACATTTGAGAGAAATGGTGTAGATGCTAGATA

GGCATTGCAAGTGGATAGGCAGAGTCGGGAATGAATGAGTATCATTTTAGAGTTAAAAATTTTGCTGTTT

TGTACTTTGAGAGCTATTCTACCTAGTTTGCTTTTCTTTATATTTCTATGGTGTTTTCAAAACTA

>FK826567.1 JutEST4129 Jute Plasmid library 20-45 days after sowing Corchorus capsularis cDNA similar to DNA repair protein RadC, mRNA sequence

GGGAACCCAATTCCAGAAAAATTACTCTCGCAATCTCAAGCTCAAAACTTCATCTTCAAAAATCAAAATA

ATCCCAATTTTCTTCCAAAAATCACCAAATTCATTTCCCTCTCGAAACCGTTAAATCAATGGCGAAAACC

CAGCAGCAACTCACAGCGACTCTGTTCTTCTTAATCATCTTCACGGCGACTTCGCTGTCTTCCGCAAGGC

TTCTAAACGACTTCGTTTCGCATGGCCTGGCAGCGACGGAGCCAACCTTGAACCTGGCTCTTCCCCTCGT

CGACGTCAACGACGAACCGGTCACGACGGAGGAGCAGCGAGATCGTAAGCCGGCGCTTCTGCCATGTGAC

CATATGGTTCCGATCAAGAACTCCCACGTCGGCCTCCGGAGGCCTCCAAGACTCGCCGGAAAGTATGGAC

CCATGATTCTGAGCATGCTTCCGAAGGGACCCGTGCCGTCTTCTGGACCTAGCAAGGGAACTAACGACGT

CAAAAACTAGCTCTGCCTCTGCTCTGTTCTCTCATAGGAATTTCTTGGTTCATTTTTTTGTTCTTTTTG

>FK826566.1 JutEST4123 Jute Plasmid library 20-45 days after sowing Corchorus capsularis cDNA similar to drought-induced protein SDi-6(fragment), mRNA sequence

ACTCCATCTGCTACTAACCTCACTACTCTTAAGCCACAAAAGATTTTGCTACCGGCACCTAAGCTCCAAT

GGAGGGTGAGAAGAACTCGTCCGGGGACCTGATGTCCAGCAGCAAGCTGGTCGCGGAGGCCGCCAAGACA

GCCTACGAGCAGAAGAGCGTCGAGGGCTTCGACAAGGAAAAGGTAGCCGGCGCCTCCGCCGAGATGCTGG

ATTCCGCCGCCAAGTACGGCAAGCTGGAGGATAAGCCGGTGGGACAGTACCTCGACAAGGCCGAGGGGTA

CCTGAAACAGTACAGCTCCGGCGGCACCGACAAGGAGAAAACCGACGCACCGGCTGCCGCTAAAGATGGT

GAGGCAAAGCCAGAGTCGCATCCAAGCGGTGGCCATGAAGGCAAGGAGTCTGGAAGCGGTG

>FK826565.1 JutEST4114 Jute Plasmid library 20-45 days after sowing Corchorus capsularis cDNA similar to Diacylglycerol kinase variant A, mRNA sequence

GGGATACAGTTGTGGTTGAAATTTCTCATCTTGGACAAGTCAATATGCTTGCAACCCAAAATTGCCGTTC

GAAAAGCATTCATGATCCCTCATCAACTCATTATAATGGTGATGGCGACGAAGATGATAGTGATGAAGCA

GACCCCAAAACTGAGGATGAGTTTAGAAAATTTGGAGCAGCAGACACGTTCAGAATCCCAGATGACGTTG

ATATTTCTCGCCTTAGTTAAAGATATAACATTTCATTTATTCTAGTAATTATTTTCTCTATCCTGTGTAA

TAGCAATTCTCGTGGTTCAGTATTAATTTTCGGTTTAATATTGCTATATAGGCTTGCCATTCATATTTGA

AGTGCTGTTCATCTTATAGCAAAAAATCTCATCTTAGCGAAATCAATAATCTTGTTGTAAATTTAGATAT

CTTACAGTAATTTGTATTTGCTGCGTAATTAACAATTATGTGGTGGCCCTCGTTCATGAGCATTAAAATT

GTTTATGTTGGTTTG

>FK826564.1 JutEST4117 Jute Plasmid library 20-45 days after sowing Corchorus capsularis cDNA similar to Gip1-like protein, mRNA sequence

GGATTTCACTTTCTTCTCTGTTTTCTTTGCTCTTATTTTGACAGTTTCAAGAGGGGTTTGCCTTGTAGCA

AAAGGCCATGGCTAAACTTGTTGCTGCCTTCTTCTTGGCTCTCATTGCCATCTCCATGCTTCAAACCATG

GTCATGGCTTCACATGGACATGGAGGTCACCACTATGATCAAAAACATTATGGTCCTGGAAGCCTCAAGA

GCTACCAATGCCCATCACAATGCACGAGGAGATGCAGCAAGACCCAATACCACAAGCCCTGCATGTTCTT

CTGTCAGAAATGCTGCAGGACATGCCTGTGTGTGCCCCCAGGGTATTATGGGAACAAAGCTGTGTGCCCT

TGCTACAACAACTGGAAGACCAAGGAAGGAGGACCCAAGTGCCCTTGAAGACCCCCCTTCCTTTAATCAT

CCCTATATTTCCTACTTAATCAAGTTCCTTGAATCATAATTAAGCATAATGTTCTACTACTACTCCTAAC

TATATACTACTAGTACTTCATGTTACTTTATGTTTTGGAAATCAAAATGTGGTGGGATTTTGTAATCACT

CTGTCACTCCATAGGGTACCCAATGTCCCAATCCCTTTAAATGAGAGTGGCTAATCTTTGTTCTATTTAT

CAATGAAGGTGT

>FK826563.1 JutEST4113 Jute Plasmid library 20-45 days after sowing Corchorus capsularis cDNA similar to Pectate lyase, mRNA sequence

GGGCCTGGCACTTTAAGGCACGCCGTGATTCAAGATGAGCCGTTGTGGATCATATTCGCTCGTGACATGA

CCATTCAATTGAAGGAAGAACTCATCATGAACTCCTTCAAGACCATCGACGGCAGAGGCGCTAGCGTCCA

CATTGCCGGCGGTCCATGCATTACCATACAGTTCGTGACCAACATTATCATTCATGGACTCAACATTCAT

GACTGCAAGCAAGGAGGGAATGCTATGGTGAGGGACTCCCCACGGCACTACGGGTGGAGGACTCTATCGG

ACGGTGATGGTGTGTCCATCTTTGGGTCTAGCCATGTTTGGGTGGACCATAACTCTTTATCAAACTGCAG

AGATGGGCTGATTGATGCCATCCATGGGTCCACTGCAATCACCATTTCAAACAACTACATGACTCACCAT

GATAAGGTTATGCTATTGGGGCACAGTGATTCCTACACTCAAGACAAGAACATGCAAGTCACCATTGCCT

TTAATCACTTTGGTGAAGGGCTTGTCCAAAGAATGCCAAGATGTAGACATGGATATTTCCACGTGGTGAA

CAATGACTACACCCATTGGGAAATGTATGCTATTGGAGGAAGTGCTAACCCAACCATCAACAGCCAAGGG

AACA

>FK826562.1 JutEST418 Jute Plasmid library 20-45 days after sowing Corchorus capsularis cDNA similar to Probable ubiquitin-fold modifier 1 precursor, mRNA sequence

GGGAGAAATTTCAGAAATCTGAAAAACGGAAAAAGAAGGGAGAAATGACGACTGGTGGAAAAGTGTCTTT

CAAAGTCACTCTTACCTCCGATCCAAAGCTCCCTTTCAAAGTGTTTAGCGTTCCTGAAGCAGCGCCGTTT

ACAGCTGTTTTGAAATTCGCCGCCGAGGAATTCAAAGTCCCTCCTCAAACCAGTGCTATCATCACCAACG

ATGGTGTAGGAATCAATCCTCAGCAAAGTGCAGGAAATGTCTTCCTGAAACATGGTTCTGAACTTCGCCT

GATTCCCCGCGACAGAGTTGGAGCCTTTTGAGTTAAATGGTGAAGACTTGATTATCATCATAGCGAGAAG

ACTAGTTATGATCTTGGTTATCGATTAATTGTCTTTGTATCTTTGTACTTTTTCTCCGTGTAGAACAGAG

CTGATAATATTTTATTTTAGCCCATATTTAAGAGCTATACTACATAAACAATTAATTATAAGTATGGTGT

GCATGTAACAGTTTGCCTTTGCCTTGTGCTGTTTTGGTATTTTATCACCACCACCACCAACTAGTTACTG

CTGTACACAGCAAATAAATAAATATACCAGAGTTAGAAGCC

>FK826561.1 JutEST41125 Jute Plasmid library 20-45 days after sowing Corchorus capsularis cDNA similar to Beta-1,2-N-acetylglucosaminyltransferase II, mRNA sequence

GGGCCAGATTGCTATGCTGCCAATCTGGCACCTTGTGACGTGAGTTCAAGAGGAGAAGGATGGAATAGTT

TGATTGCAGAAAGAATGGGAAATGTGGGTTATGCCTTTAACCGGACTGTTTGGAGAAAAATCCATAGGAA

GGCTAAAGAGTTTTGCTTCTTTGACGATTACAACTGGGATATAACAATGTGGGCAACAGTTTATCCTTCA

TTTGGCAGTCCGGTGTACACATTACGAGGTCCTAGGACTAGTGCAGTTCACTTTGGGAAGTGTGGTTTGC

ATCAGGGCCAAGGGCAAAGTGCTGCTTGCATTGATAATGGATCAGTGAACATTCAAGTAGATGACACTGA

TAAGGTGGCTAATATCAGATCAGAATGGGATGTGCGTGTCTATGACAATCAACCAGGGTATAAAGC

>FK826560.1 JutEST41123 Jute Plasmid library 20-45 days after sowing Corchorus capsularis cDNA similar to Light-inducible protein ATLS1, complete, mRNA sequence

GGGAGAAAGTGAAGAGTGAAGTCAAAATCAAAAGCTGAGAAAGATGCCTTGCCTCAACCTTTCCACCAAC

GTCAACCTGGAGGGCGTCGACACCTCCTCCATCCTCTCTGAAGCCACCTCTACGGTTGCCAAACTCATCG

GCAAACCCGAGTCCTATGTGATGGTCGTGTTGAAGGGTTCGGTGCCCATGTCATTTGGTGGTACTGAGCA

GCCAGCGGCCTATGGTGAGTTGGTGTCCATTGGTGACCTTAACCCAGATGTGAACAAGAAATTGAGTGCT

GCAATTGCTGCAATTCTTGAAACTAAGCTTCAAGTGCCCAAGTCTCGGTACTTCCTCAAATTTTATGACA

CTAAGGGTTCCAACTTTGGATGGAATGGATCCACCTTCTGAGTCTCGCATGCTCCCAATTTATGAAGGAA

ACATCTGTGTTGGTTCTTAACGCTGTGTTTCTATGTTGATTTGGTGAACTTTGGCATGGTATTTAGTTTA

TCTGTGTCTTCGTAGGAACATAATATGCAAAACATTAATGCATACTAAGAACTTTATGTTATGTTGGCAA

TAGCCTATATAGTTGTTATGGGCACATCATTCAGTCCCTAGGGG

>FK826559.1 JutEST41120 Jute Plasmid library 20-45 days after sowing Corchorus capsularis cDNA similar to T30E16.31, mRNA sequence

GGAACGGATCCTCCAAAAACGCCAGATGGTTCGTCGAGTTCGTTCCCGGACACCAAAACATCATCCGCCT

CAAAAGCTCTTACAACAAGTATCTCACCGCCTCCAATCAACCCTTTTTACTTGGGATGACCGGTCGCAAG

GTGCTTCAGACGCTGCCTCGGAGGCTCGATTCTTCGGTTGAGTGGGAACCCATTAGAGAAGGGTCTCAGG

TTAAGCTCAAAACTCGGTACGGTAACTTCATGAGAGCAAATGGTGGGTTGCCTCCTTGGAGGAACTCGGT

CACCCATGATATTCCTCATAGAACTTCAACTCAAGATTGGGTTTTGTGGGATGTGGATATTGTGGAGATT

CAAGT

>FK826558.1 JutEST41107 Jute Plasmid library 20-45 days after sowing Corchorus capsularis cDNA similar to Ferredoxin, mRNA sequence

GGGGAAACAAAGGCAAATTCTCTAAACGCCTCTGATTCTCTCTCTTCCATTCCCAAAAATGGCGATTTCA

ACACTGCAAAAACTTTCCTCTCAAATCCACCGTCTCCCGTCACTATCAGCCCACATCCTCACCCGATCCT

CCGCCACCTCTGCCTCCCCCTCCGCCTCCACGTCCTCCAAGAAAGTTGCCGACAGGATCGTCAAGCTCTT

CGCCATTGACCCAGAAGGGCAGAAGCGTGAGGTAGTGGGCTTAACCGGCCAAACCTTACTTCGGGCCTTG

ACTAACAGTGGGTTGATTGACCCTGCCTCTCACCGCCTTGAGGAGATCG

>FK826557.1 JutEST4193 Jute Plasmid library 20-45 days after sowing Corchorus capsularis cDNA similar to Thiazole biosynthetic enzyme, mRNA sequence

TCATCATCCATGGCTGTAACTCTCACTTCCTCAACCAAACTCTACAAGAACCCTTCTGTTTTTGAGTCTT

CCTTCCATGGGGTTCCCATAACTCCACTTTCTTTCAATTTCAAAACAAAATCCTCTCCATGCAATGCATC

GATTTCCATGTCTGCTGCCTCATCATCACCTCCACCCTATGATCTGAATAATTTCAGGTTTGACCCCATC

AAGGAGTCCATCGTGTCTCGTGAGATGACACGTAGGTACATGATGGACATGATCACTTATGCTGACACTG

ATGTGGTCGTGGTTGGTGCTGGCTCCGCCGGTCTTTCTTGTGCCTATGAACTCAGCAAGAACCCTTCTGT

CCAGGTTGCCATTATTGAACAATCTGTTAGCCCTGGAGGTGGTGCCTGGCTTGGTGGACAACTTTTCTCT

GCCATGGATAGTTCGCAAACCAGCTCATCGCTTCCTT

>FK826556.1 JutEST44130 Jute Plasmid library 20-45 days after sowing Corchorus capsularis cDNA similar to Lipid transfer protein precursor, mRNA sequence

GGATGGCTAGCGCTATTAAGATTGCATGTGTTGTGGTTGTGTGCATGGCGTTGGTGAGTGCACCAGTAGC

TCAGGGAGCCATTTCATGTGGTCAAGTCGCCAAAGCCCTGGGTCCATGCATTCCTCACTTGACTGGTAAG

GGTCCTTTGGCTCCTGGGTGCTGCAGTGGGGTGAAATCTCTCAACGGCGCCGCCAAAACCACCCCTGACC

GCCAAGCTGCTTGCAACTGCCTTAAAAGTAACGCTGGCTCCATCTCTGGCATCAACTATGGCCTCGCTGC

TGGACTCCCAGGCAAATGTGGTGTCAGCATTCCTTACCAGATCAGCCCCTCCACTGACTGCAGCAAGGTT

AAGTGAAAATGCGGAGATTGGAACGGTTCTCAATGTGAAGGATATAAAGAGAGTTTTAGTCTTTTAGATG

ATACATAATAAAATGGATGTAAAAATCCATGTGGAAGCACCCCTTCGTAGGAGAGTTATCCTATAAGTTT

TAGAGTTGTACAAGGTCGCTTTATATGCGACTATGCAAAATTTCATTTCATATATAAAAAATTTATTACA

ATTATAATACCCC

>FK826555.1 JutEST4274 Jute Plasmid library 20-45 days after sowing Corchorus capsularis cDNA similar to Lipid transfer protein isoform 1, mRNA sequence

GGGGGACACCAAGAATCAACTTTCCCTTGTTATTCAAAAGCACTAATTAATATTTTTGCTCTCGCTTACT

CTCTGATATTAATCGAAATGGCTTCCGCTATTAAGTTTGCATGTGTGGCGATTTTGTGCATGGCTTTGGT

GAGTGCACCACTGGCTGAGGGAGCCATAACATGTGGTCAAGTCACCAGCTCCCTGGCGGCATGCATTGCT

TACTTGAGAGGGGGTCCTTTGACTCCTGGCTGCTGCAATGGGGGTGAGATCTCTCAACAGCGCCGCCAAA

ACCACCCCGGACCGCCAAGCCGCTTGCAACTGCCTTAAAAGCGCCGCCGGTTCCATTCCTGGCATCAACT

TTGGTCTCGCTGGAGGACTCCCTGGCAAGTGCCGCGTCAGCATTCCTTACAAGATTAGCCCCTCCACTGA

CTGCAGCAAGGTTAAGTGAAATGCCAAGATTTTGGAAGGGTTTTATTTTAATTTAAGCTCCAAGGAGATG

AAGAGAGTTTACTTTTAGATGGAAAAATAAAATGGGATGTAAAAGTCCATGTGGATAGCACTCTTTCGTA

GGAGAGTTGTCCTAATTTTAGATTTGTTATTTGTACTTTTATCGTGTGTACAAGGTCGCTTTTATATATG

CGACTATTACTTTCATATATG

>FK826554.1 JutEST4316 Jute Plasmid library 20-45 days after sowing Corchorus capsularis cDNA similar to Chlorophyll A/B binding protein, mRNA sequence

GGGGGGACTCACAACCACAAACTCCCATCAGCAAACAAAAAATCTCTCAATATTTCACAATGGCCGCCGC

AACCATGGCTCTTTCCTCCCCAACCTTCGCCGGGAAAGCTGTCAAGCTTTACCCATCCACCTCTGAAATC

CAGGGCAATGGCCGTGTTTCCATGAGGAAGGCCACCAAGCCCTCAGCTCCCTCCGGCAGCCCATGGTACG

GCCCAGACAGAGTCTTGTACTTGGGCCCACTCTCCGGCGAGCCACCATCCTACTTGACCGGTGAGTTCCC

CGGTGACTACGGTTGGGACACCGCCGGTCTCTCAGCTGACCCCGAGACTTTCGCTAAGAACCGTGAGCTG

GAGGTCATCCACTGCAGATGGGCCATGTTGGGAGCTCTCGGCTGCGTTTTCCCGGAGCTTTTGGCCAGGA

ACGGCGTCAAGTTCGGCGAAGCCGTGTGGTTCAAGGCCGGAGCCCAGATCTTCAGCGAAGGAGGACTCGA

CTACTTGGGAAACCCAAGCTTGATCCACGCTCAGAGCATCTTGGCCATCTGGGCTTGCCAAGTTATCTTG

ATGGGAGCCGTCGAGGGTTACCGTATTGCCGGTGGACCCCTCGGNTGAGGTGGTTGGACCCATTGTACCC

AGGTGGCAGCTTCGACCCATTGGGTCTTGCCGATGACCCAGAGGCCTTCGCTGAGCTCAAGGTTAAGGAA

ATCAAGAACGGCCGATTGGCTATGTTCTCCATGTTTGGATTCTTCGTCCAAGCCATTGTCACCGGAAAGG

GACCATTGGAGAACTTGGCCGACCACCTTGCTGACCCAGTCAACAACAACGCCTGGGCCTACGCCACCAA

CTTT

>FK826553.1 JutEST45139 Jute Plasmid library 20-45 days after sowing Corchorus capsularis cDNA similar to Ubiquitin ligase SINAT2, mRNA sequence

GATGGATGTACCTTCAATCATCGATATGTCAAATCAAATCCACATGAAGTTGAAAACGCTACATGGATGC

TCACGGTGTTCAATTGTTTTGGGAGACAATTCTGCTTGCATTTTGAGGCTTTCCAGCTCGGCATGGCACC

CGTGTATATGGCCTTTTTACGTTTTATGGGTGACGATAATGAAGCAAAAAAAGTTCAGTTACAGTTTAGA

AGTTGGTGCCAATGGCCGTAAACTAATATGGCAGGGAATTCCCAGGAGCATCCGGGACAGTCATAGGAAA

GTTCGTGACAGCCAGGATGGGCTTATTATCCAAAGGAATCTGGCTCTATACTTTTCCGGTGGGGATAGGC

AAGAGCTGAAGTTGAGGGTAACCGGCCGTATATGGAAAGAAGAATGAGATAGAAAACGATGGAGAATTGA

GGAGAGTTTGTACAGTTTCAAAAAGTTGAGTCTCATTGGCAAGCAGATGAAGCTTTTGTTTTTCTGATCA

GGGCTTTCTTTTGTATTACATGTTGTATTTTGACTGGAATCTGTCTTCTACCATCTAATCTGTCTGCTAA

CCGCGCCCTTTGGCCGAATGTTTATGTAGAAAGTCATGATTATGAGTCAACTTGGCTGCTCCAACATCTG

TGTATGCT

>FK826552.1 JutEST4287 Jute Plasmid library 20-45 days after sowing Corchorus capsularis cDNA similar to ubiquitin-protein ligase, mRNA sequence

GGAGCGAATTCCTACTCCGAAGCAACCTGCCGAGCCTACCGTTGTTCGGTCCTTGCTGAATCAAGCTTCC

TCTGTTTCCACTCCGTTTCAGTCTCGTCTTTCTGCTCTGCGGCGGCTCAAGGGACTCGCCCGTGACTCCG

ACAAGAACCGGTCCCTTATTTCCTCCCTTAACGCACGTGAAATACTCGCAAACATCGTGTTTTCCGAGAC

GGGTTCGGGTTCTTCCGAGTTGAACCATGAGGCTCTCGCGGTTTTAGTAATGTTTCAGCTGACGGAATCT

GAGTGCGCGTTAATCGCTTCGGATTCAATTAAAGTTGTCTACTTGTCGCAACTCTTGTTTCATTCCTCGA

TTGAAGTCCGAGTTAACTCAGCTTCGCTCATCGAAATCGTTCTCGCCGGAACGAGGTCGTCAGATCTCCG

AGCTGAGATAAGCAATGTGGATGGAATATTCGAAGGCGTCACCGACATATTGAAGAACCTCAACTCCTAT

CCACGCGCTCTTTAAATCAGCATTCGAGCTCTATTCGCCTTGTGCTTAGTAAAACAAACCAGGCACAAAG

CGGTCCAAGCCGGCGCCGCCGAAACTTTAGTCG

>FK826551.1 JutEST48119 Jute Plasmid library 20-45 days after sowing Corchorus capsularis cDNA similar to Specific tissue protein, mRNA sequence

GGGGCAGTGCAAAAGTTATTGCATTTTTTAAACCATGAAATCTTCTATTGCATTGTTTCTCACTTTTCTT

GCTTCTGCAGCATGCAGACAGCAAGAAAAGAGGTGGGGAATATGCTGGAGGGCACTATAGAGCTGAAACG

ATGAATCAACCTATGCCGGAAGCAATCCAAGCGCTTGTTCGGGTTAATGCCGCAGCTGCCCTCCCCGGTG

ACAAAACCGACTGTCATACACCAAGTAACGTGGAGACTAAAGAAGAGAAGATTATTGTGAATGAGGAATT

TGAACCAAGGCCAAACATATCACCTTGAGTTGAAGAGGGTAAGATCATTGTTGCTGATGATTTTGAACCA

AGGCCCAATATTTCTGCTTACCATGATGATGGTGATGTAAAGGGGGAGAAGAAATCATCTTGTGTTGAAG

ATTTTGAACCAAGGCCCAACATTTCTGCTTACGGTGGCGGATGATGCTGATCTTAAGGGGGGGAAAGAAT

CATTTGCTAATGATTTTGAACCAAGGCCCAACATTTCTACTTACCATGATTAATTGGAGGTGAAATTAAA

ACAGAAATAAATAAAAGAGAGAAACCAATTTCTTTGATTTCTCTTAATTAGAAAAAATCATTATGGCCTC

GTGGCCTCTATGTATGGCATGGTTTGTTTT

>FK826550.1 JutEST4466 Jute Plasmid library 20-45 days after sowing Corchorus capsularis cDNA similar to Specific tissue protein 2, mRNA sequence

GGGGGACAAGTTGATATATTTTATTATCAATATAATATAATATAATATTCTATTATATCCCTACATTGCA

GTTCTGCAACTATGAAATCTTTCCCTTCTTTCTTAGGTTTCCTGTCACTTCTCATGCTTGCTGACACCAT

AGCAGCAGCAAGAAAAGATGGTGGGGAATACTGGAGAGTAGCTGTAATGAAAGATGAATCCATTCCAGAA

GCCATCGAAAGCGCCCTTGTTCCGGTTAATGCCGCAGCTGCTTCCTCCTCCGGTGACAAAACCAACTGTC

ACGATTTACCTAGCAGCATCGAGATTAAGGAAGAGAAGATTTTTGTTGAGGATTTTGAATTACCAAGGCC

TAATAATTATAACAGTGTTGAAGGAGGGGTGACGAAGGAACGTTCATTTGCTAAAGATTTCGAACCCAGG

CCCAACCTTTCGGCTTATGGTGATGATGGTGATCTTAAAGAAGAAAAAAAAATCATTTGCTAAAGATTTC

GAACCGAGGCCCAACCTTTCGGCTTATGGTGGTGATGGTGATCTTGAAGAAGAGGAAAAATCATTTGCTA

AAGATTTCGAACCCAGGCCTAACCTTTCGGCTTATGGTGATGATGGTTATCTTAAAAGAGGGGGAAAATC

ATTACTAGTGATTGGAAC

>FK826549.1 JutEST4185 Jute Plasmid library 20-45 days after sowing Corchorus capsularis cDNA similar to sucrose synthase, mRNA sequence

GGGTTCTACATTCTCAAGTTCCGTGATTTGGTAAAGACTGTTCCTTTGGCCAGTGATGATTAGGCTTTTC

TATCCCTTGCTATTGGGCATAACCTATGGAATAGCCAATAGTGACTTCAAACAATAATGATAGTTTATAA

ATAAGGTTCCAGTTGTGGTTGAGGGTGCTGAGTATTATATGTATCTTCTGGTTGTGACTTGTGAGGCTGT

ATGTAATGTACCGAGTCTAATACCTGTGGAGCTTTTGTGACTCAACTTCTTTTCGAAGAAAACTTGTTAT

CATGGCTTTTGTTTTGTATGTAGTAAATTGCAATAATAAGACTTTCTCACCCTG

>FK826548.1 JutEST4284 Jute Plasmid library 20-45 days after sowing Corchorus capsularis cDNA similar to Acyl carrier protein, mRNA sequence

CGGATACTTCTCTTTCTTTTTCGCTCTTTATTCCTCTCACCATCACTAGAAGACAATCAAGAATCAGAAA

ACATATGAACAAAAACAACAATGAAGCAAAAAATAGATGTGATAGTAAAAAAGAACATTGACTGAGAAAG

TAGCAAAGACAAATTTTTAATACTGCATATTCAGACAAAAGCCTCTTACCAGATACACTTAAGAACAAAA

GTACTATACCTAGGTTGATACATTGGGGATCCGGCTTCTTCCTGTTTTTGATTCTCTCTTCC

>FK826547.1 JutEST4184 Jute Plasmid library 20-45 days after sowing Corchorus capsularis cDNA similar to Ubiquitin OUB2, mRNA sequence

GGCTGATTTTTGCTGGTAAGCAGCTTGAAGATGGCCGCACCCTTGCAGACTACAACATCCAAAAAGAATC

CACCCTCCACTTGGTGCTCCGTCTTCGTGGTGGGATGCAAATTTTTGTGAAGACCTTGACAGGAAAGACC

ATCACTCTGGAAGTTGAGAGTTCAGACACTATTGATAATGTTAAAGCCAAGATCCAGGATAAGGAAGGAA

TTCCCCCAGACCAGCAGAGGCTGATCTTTGCAGGGAAGCAACTTGAAGATGGGAGGACCCTGGCAGACTA

CAACATTCAGAAGGAATCAACTCTGCACCTTGTCCTCCGTCTCCGTGGTGGTTTTTAAGTGATGAACTTT

CAAGTATGATTTGCAATGTGGTCCTGCATTAGAAGTCCCAATTGTGTGTTGAGCATTGAATGGTTATGGA

TATCTCTGGTTGTGTGAGCTTGCTGCTATGCAGTGGCTTTTAAATAATTTCTTTAAACTTATCTAGTATT

ATGCTATGGTTAATGTTGCCATGCTCTGGCATTGAATAAATTTCAATTTCCAG

>FK826546.1 JutEST4182 Jute Plasmid library 20-45 days after sowing Corchorus capsularis cDNA similar to Senescence-associated protein, mRNA sequence

GGGACCAAACAACTTTTCAGTTTTCTCTCAGCATTTAACTTTCTCCTCATCCAAACAATCACAAAGAGAA

TCAACAGAATCCCGGTAACAATGGCGTTTCGTTTGAGCAACAATCTCGTTGGGATCCTCAACTTTGTAAC

GTTCCTCCTTTCCATCCCCATCCTCGGCGCCGGCATCTGGCTGAGCCGTGAAGGCGTTACTGAATGCGAA

CGGTTTCTAGACAAACCCGTTATCGTTATCGGAGTCTTCCTCATGCTTGTCTCACTCGCGGGACTAATCG

GCGCGTGCTGTCGCGTGACGTGGCTACTCTGGGTTTACCTTGTCGTCATGTTCCTCCTC

>FK826545.1 JutEST4181 Jute Plasmid library 20-45 days after sowing Corchorus capsularis cDNA similar to 36kDA porin II, mRNA sequence

GGGGGACTAAGAAAGGTGAATTGCTTTTGGCTGATGTAAGCACTCAGCTGAAGAACAAGAACATCACCAC

TGATGTGAAAGTTGACACTAATTCGAAAGTTTTTACCACCGTCACTGTTGATGAGCCTGCACCTGGATTG

AAGGCTATCTTTAGTTTTGTTGTCCCTGATCAAAGGTCTGGAAAGGTGGAACTGCAATACCAGCATGAAT

ATGCTGGAATTAGCACTAG

>FK826544.1 JutEST4384 Jute Plasmid library 20-45 days after sowing Corchorus capsularis cDNA similar to cytochrome c oxidase, Vc subunit, mRNA sequence

GGATTGGATTATCACTTGGTCTTTTTGCTGGTGGGCTCTGGAAGATGCATCACTGGAATGAGCAAAGGAA

AGTGAGGGCATTCTATGACATGCTTGAGAAAGGTGATATCTCTGTTGTCGTAGAAGAATAGATGTTCATG

TCTTGTGCAATTGTGGACAGTCTTGGCAGTATTTTGATATTTTGTGTTTCCTTCATTTGGCTCTTAGATT

CAAAATGAGCAAACTCAGTTTGGAGTTATTAAGAATAATTTTATTGTATTAAGATCCAGATCACATATCT

GCTTAACAATCTCAATGGTTTTGTGATATTGTGTCCTTAAACTTTTTTCCTTCTGTTTTTGTGGCATCTC

TTGAGATG

>FK826543.1 JutEST43105 Jute Plasmid library 20-45 days after sowing Corchorus capsularis cDNA similar to Cytochrome B6-F complex iron-sulfur subunit, mRNA sequence

GGGACAACTCCAACAGCACACTTTCCATTCCTTTCCATGGCTTCCTCTACTCCCTCCCCTGCAGCTTCTT

CTCAGCTATGCTCGAGCAAGAGTGGCATGTTCTGCCCATCACAAGCTCTGCTAATGAAACCAACAAGGAC

CCAGATGGTTAAAAGGGAAAAGGGAATGAAGTTGACTTGCCAGGCCACTAGCATCCCTGCTGATAGAGTT

CCAGACATGGGCAAGAGGCAGCTCATGAATCTGCTTCTGTTGGGGGCTATTTCACTCCCCACAGGTTACA

TGTTGTATCCTTATACATACTTCTTTGTCCCACCTGGTGGGAGAGGTTCCGGTGGTGGTACTGTTGCTAA

GGATGCAATCGGAAATGATATCCTTGCTGCTGAATGGCTTAAGAACCATGGCCCCGGGGACCGAACCCTG

ACTCAAGGATTGAAGGGAGATCCAACCTACCTTGTGGTGGAGAAGGACAGAACTCTTGCAACATATGGTA

TAAATGCAGTCTGCACACATCTTGGATGTGTTGTTCCATGGAATTCTGCTGAGAACAAATTCATATGCCC

CTGCCACGGATCTCAGTACAATGACCAAGGAAGAGTTGTGAGAGGA

>FK826542.1 JutEST4294 Jute Plasmid library 20-45 days after sowing Corchorus capsularis cDNA similar to Cytochrome b5 domain-containing protein-like, mRNA sequence

GGCGGCGCCTACGCCATGTTCGCTGGCAAAGACGCCAGCAGAGCTCTTGCAAAGATGAGCAAGAACGAAG

AAGACGTCACCCCCTCCCTTGATGGCCTCGACGAAAAACAGATCGGCGTTCTTAACGACTGGGTCCAGAA

ATTCCAAGCTAAGTACCCTGTTGTTGGCCGCGTTGTCAATTTGTCTTATGCATCTCGTCCCTCTATGTTC

TAATATATCATGTAACATGCTTGGTCTCTTTATGTTATGGGAATCTATCTATGTGGTTAATATGTTTCTG

GCTTTCCATAAACTTGTTGGATTATGGTAATCTATCTGTGCTACTTCCAACTTGTCTTTGACTAAATGCA

GAACATTCGATTGTAACTAATAAAGTTTAAAGCTGT

>FK826541.1 JutEST4180 Jute Plasmid library 20-45 days after sowing Corchorus capsularis cDNA similar to Cytochrome b561,complete, mRNA sequence

GGGGAAGAACGAAAATCTGGTTTCCATGGAAGGAGAAGGAGAATCAGCAGAAGAAGTCAAGCCAAACGTC

GTCGTTAAAGAGAACGGCGATGTTGTCGGTGACTACGATGCAGGAAAGCATGGGATCCATAGTTGTCAAG

CCCCCGGTTTTCTTGGGAAGTAAAATTTTCCAGAAACGGTTCCCCTGCAATGCCGCCGTCTGGACTGTTG

AATACCACGAGTCCCGCCCCATCCCAGTTGACCATCTTCTTTGGTGGACAAGTTTGTGTGTTTGATGGAA

TTCCTATG

>FK826540.1 JutEST4271 Jute Plasmid library 20-45 days after sowing Corchorus capsularis cDNA similar to CLAVATA1 receptor kinase, mRNA sequence

GGGGTCCTAAAAAACTTTCTTAACAATTACATATTTCACCATTACCTCTTGTTTTCTTGTTCTGTTCTCT

TGCTGCGCATAGCTAGCAAAAAAGACCAGAGCTTTAGTGGTCCGATTCTGGTTCATGGATATTGATCCCT

TCTGGATTACTGGGGGGTGGTTCATTATTCTGGGTTTTCTGTCATCATAAAAATTTCATGGCAATCACCA

AAACTAAATCATCACATTTTCAAACTAGTCTCAATCTCACACTACCCTTGACTCCTCCTCTTCCTAATTG

TTCTTCTCTTAGATTTCCCCTCTCTTTGCTCGCCCAGCTCATTATCTCCAAGAGTCCCCGTATGTTACTT

CTTACAGGCAACTTCTCTCATCATCTTCCACCACAAATTCTGCACCCTTTTCCCACTTCTCCATCCTTCC

CAAGCTGCTACTTCCAGTTCCACCACTGCCACACCAGCAACTGCTAAAAGTCAACAGTTCCAGGCGGCTG

CTCACGAAGTTCCCAGTGGGACCAAATCCTGAATCCAACAAGTAAGCAAAAAAAGCAGCATCAAAATTTC

AGTTTTCACCACCCATTATTATCGTCTTCGATTGTGTCCTTAATTCTCTTCTTGTTTTATTGATCT

>FK826539.1 JutEST4323 Jute Plasmid library 20-45 days after sowing Corchorus capsularis cDNA similar to Metallothionein 3b, mRNA sequence

GGGGGGCACATTTTCACTGTAAACAGAAAGCTCAATACAGGAACATATCCACCATCAACGAGCAACAATG

GCAACCTTATCTAGCCTTAACCTTGTAACTCCAAGAGTCGTCGCCAACGTGCCAGACTCCACGAGGGTTG

GCCCTGTGAAAGTCCCATGCCTTAACCAGCCCTGGAAAGGGTATCTCAGTTGGGGTCAAGGCGGATGCAG

GTGTTCAGACCTGTGAGAGCCGCTCCCGACCAAATATCGGAGAAGGTGGAAAAGAGCATCAAGGAAGCCG

AGGAGGCATGCTCAGGTGACCCAACGAGCTCAGAGTGCGTGGCGGCTTGGGACGAGGTGGAGGAGCTCAG

CGCCGCCGCCAGCCACGCCAGGGACAGGCAGAAAGCAAACAACGACCCTTTGGAGAACTACTGCAAAGAG

AACCCTGAGACAGATGAGTGCCGCACTTATGATAATTGAAAATCGCTTTCTTTTTCTTTAATTAGTTTAT

GATGATAAAGGTTTTAGAACCTTCGATGATGAATATGGCGGGGGATTTATGTGGGGGGG

>FK826538.1 JutEST4124 Jute Plasmid library 20-45 days after sowing Corchorus capsularis cDNA similar to Metallothionein 2a, mRNA sequence

GGGGGTTCAAGTTTCCAGTGTTAGCTCTTTGATTTCCCAGCAAAAAAAAATCTCTTCAATTCTCTCAATT

AGAAAACTGAAAATGTCTTGCTGCGGAGGAAACTGTGGCTGCGGCTCTGGCTGCAAGTGCGGCAATGGCT

GTGGAGGATGCAAGATGTACCCTGACATGAGCTTCGCCGAGTCAACCACAACCGAGACTCTTGTTATGGG

AGTTGCACCCCAGAAAGTGCACTTCGAGGGAGCTGAGATGGAAGTGTGGGGCCGAGAATGGCTGCAAGTG

CGGAGCTGACTGCAAATGCAACCCTTGCACTTGTAAATGAGAGAGAAAAAAATCTGAAGATCCCCTGAAA

ACAGCAGAGAGTCCTTAATTTCCCTTAACAAATAATATCTAGTNATGTCTTTCGNGNACNAAGATNTATC

TTGTTCCTGTGTTNTGTTTAGAATAAAGTAGCCATGGGTTTTATCTGTCGGCAAAATCCTCACCAGGATT

TGTCTGGTGCAGCAGCCATGGTTGCTTGCTTNGGTGTGTATAAGTATAGTNAANTANGTAATNTTTAGTT

TTTCTTTATGTCTAATGTAAGCATCTCTGCTGTCTCAGGATGATGACTCTATCTATGCTTCTTATTGTAA

GAAAGTGAGTCGGTTTTTCTAATGAAATGAACGGAAAACTCCTTTCTCTTTG

>FK826537.1 JutEST4429 Jute Plasmid library 20-45 days after sowing Corchorus capsularis cDNA similar to Metallothionein-like, mRNA sequence

GGGATGGGATAGCAATCCACATAGCAAGCAATCTTCAATAACCCAAAAAGCTCTTCCAAACTTCTTTCAA

AAAACCTTCCTCAAATTCACCATGTCGGACAAGTGCGGAAACTGCGACTGCGCTGACAAGAGCCAGTGCA

CCAAGAAGGGAAACACCATGATCATTGAGACTGAGAAGAGCTACGTTGCCACCGTGGTGATGGAGGCTCC

TCCAGCAGAGAACGATGGAAAGTGCAAGTGTGGGATCTAGCTGCAGCTGCACAAACTGCACATGCGGTGG

GTCATTAAGCAAACATATATATAGAAATGAAACAAATGATCCCATGTTGCAGAAACATGGGGCTGAATAA

TTTGGGGACCTAAAATCTTATTTTGGGTTTGAATAAATTGTGTGTCTTTCCCTGTTTCTGTTGGGTGTCT

GTGTTTTCATGTCCTGTCTGTCATGTTTGTGTCTCCTTATATACTGTGATGTAATGGCCATGGCCATGCC

TTTGGCCTTGTCTCATCTTAATGAACTATTATTAGGTTGAAAA

>FK826536.1 JutEST4016 Jute Plasmid library 20-45 days after sowing Corchorus capsularis cDNA similar to Bimodular protein, mRNA sequence

CTGTAGTTTCAAATATTAATGTATTTTTGTTTAGTGTTTGGTCGATTTGAAGTTGGGATCTAGGGTTTCC

AGTTGGTTTTACATATTTTATAGGAGTCTTTTTACAAAGACTTCCGACAATTGTCGTTGTAATATTTGTT

CATGTTTAATAACTAGGCTTGTGGGTTAATTTACCTCGGCCTAATGTAATGTTTAATTAATGTTGTTTGT

TCTCAATTAATGTCAAGTTTGATTTTGCTTGA

>FK826535.1 JutEST4896 Jute Plasmid library 20-45 days after sowing Corchorus capsularis cDNA similar to Pectin methylesterase, mRNA sequence

CGGCACTGTTGCTGCCATGGGTACTGGATTTATTGCGAGAGACATAACATTCATGAACAATGCTGGTCCA

GCCAAGCACCAAGCTGTAGCATTACGAGTTGGAGCAGATAATGCAGTTGTTTTCAGATGTTTGGTAATTG

GATACCAAGATACTCTCTGCACCCATTCCAAACGCCAATTCTACAGGGAAACTGACATTTACGGAACTGT

GGATTTCATCTTCGGGAACTCAGCTGTTGTGTTACAAAACTGCAACATTTATGTGAGGAAATCTGTGTCA

TATCAGGACAACTTCGTGACGGCACAAGGTCGGTCAGCCCATAATCAAAACACAGGCATTTCGATTCATA

ACTGCAAAATAGGAGCAGCTTCAGACTTGGTTCCTTTCAAAGTCGAAAGTGAAAACGTTTCTTGGGAGGC

CATGGAAGCAGTATTCTCGGACGGTGTGATGCAATCGTTCCTGGATGATTGTATAAATCCTTCAGGATGG

TCTCCTTGGTCTGGGGTCTGGGTCTGGGAATTCGGGCTTGAAAACGGTTTACTATGGGGAATACATGAAT

TCAGGGCCTGGAGCTTCAACATCAGGCAGGGTTAAATGGGCTGGTTACCATCCTTCTCTCACATCCCTTG

AGCCCAAGCTTTCACTGTT

>FK826534.1 JutEST4890 Jute Plasmid library 20-45 days after sowing Corchorus capsularis cDNA similar to Glycosyl transferase family 2, mRNA sequence

AGTGAATTGATTCCTAATCATCTGAAGATCATAATCTCAGTGTCAAATTGATGGCACAATTATTTTCTCC

CTATACAAATTGATGGCACAATTGGGAATTTAATCTTGTCTACAGAGCCTTCCTGACGTATTTATCTAGT

CTAAGTTGTATAAATAGGCCTGTTAATATAGGGCAACATTAGCTAGGATCAGAAATGCCATTGCAGCAAC

AGATTACCAGAAGACAAAAGTTTTGTATCTAAATATTTATATTTACTGTCATTTTTAACTGTTATAAAAG

GTATATATATAAGTAAATTTTCAT

>FK826533.1 JutEST4888 Jute Plasmid library 20-45 days after sowing Corchorus capsularis cDNA similar to Putative SAH7 protein, mRNA sequence

GGGTGCCCAGTCTAATTGATTAGTTGGCGAATGATATGATGTTGTCTGCTACTATTTGTTACGTGAATTG

TTTTGTTATTTCTCTGTCTAGAGGATTTATATAACTTATTCTCTTTGAATCTATTACAATTCTATAAAAA

TATGCTAATGCTACTACGGCTTCTGTCTCAGAAGTATTTTGGC

>FK826532.1 JutEST4873 Jute Plasmid library 20-45 days after sowing Corchorus capsularis cDNA similar to putative prolylcarboxypeptidase, mRNA sequence

GCCCTATATTACAGTTTGATAAGATAATTCCTTGGTCAAGCTTTTATGATGCTGTTTCTCAGGATTTCAA

GGATGTAAGCTTGAATTGCTTTGAAGTGATAAAGGGGAGCTGGGATGAACTGGATGCAATGTCAACTCAG

AAAGAAGCCTTAGCCGAATTAAGCAAAGCTTTCAGAACCTGCAAGAGCCTCCATTCAACAGCCTCTGCCC

AAAACTGGCTATGGACTGCTTTTGTTTACACAGCCATGGTTAACTACCCAACTGAAGCAAATTTTTTGAA

GCCTTTACCTGCTTATCCTGTGGAGCAGATGTGCAGGATAATTGATAAACTTCCAGCAGGAGCAACAAAG

CTGAGTCGCGCTTTTGCAGCTGCAAGCTTATATTACAACTATTCTCGGGCAGAAAGTTGCTTTGAGATAG

AACATGAAGTTGATGCTCATGGCCTTCATGGTTGGGAATGGCAGTCATGCACAGAGATGGTTATGCCCAT

GACTTGTTCTAAGGAGAGCATGTTCCCTCCATCTGGCTTTGATTATGAAGAGTTCTCAGAACAATGCCAG

ATAAAATATGGAGTTTTGCCTCGGCCACACTGGATTACTACTGAATTTGGTGGCCAAAGTAT

>FK826531.1 JutEST4870 Jute Plasmid library 20-45 days after sowing Corchorus capsularis cDNA similar to Auxin-repressed protein, mRNA sequence

ATTACTTGAAAAGAGAAACCGGAGAGGAAAAGAGAAAGAAAAAGAAAAGAAAAATGGTTCTCCTTGATAA

GCTTTGGGATGACGTAGTAGCCGGGCCTCAACCGGAACGTGGCCTCGGCAGTCTTAGGAAGCTGGTCACC

GCCCAGCCCTTGACCATCAAAGATGAAGGAGAAGGGAGCAGGTTCCAGAGATCGTTGTCGATGCCGTCGA

GTCCAGGGACACCGACGACTCCGATAACACCAACGACGCCGGTGTCGGCGCGTAGGGATAACGTGTGGCG

GAGCGTGTTCCATCCGGGTAGCAACCTTGCCACCAAGGGAATCGGCGCTGCGGTTTTCGACAAGCCTCAA

ACCAACTCTCCTACTGTCTATGACTGGCTTTACAGTGGGGAGACAAGGAGCAAGCATCATCGCTGATTGG

GTGGTGGCTGACTGTGACTGACAGACGGGGGAAGCTTAGCAGTATAAGCAGGGGGGATTTTGGTTGAAGC

ATGTAAATAGGCGTCGTTCTTTCAATCTTCGATGTGCCAACGTTGTCCTTGTTTTGATGGGTATTTTGTG

ATTAAGTGTCCTAAGTAGTAGAACGTTTAAGACTTGACTAAGTTCTTAATTAAGCCCTTTCCTTCCT

>FK826530.1 JutEST4869 Jute Plasmid library 20-45 days after sowing Corchorus capsularis cDNA similar to Actin depolymerizing factor, mRNA sequence

AAGAAGCCCTAGATTTTCTCGGCATTTCTCGCTCTTTCTTTCGCTTTCTGGGGAAAAAGAAAAAAATGGC

TAACGCAGCTTCAGGAATGGCTGTGCATGATGATTGCAAGCTGAAGTTTTTAGAACTGAAGGCTAAAAGG

ACATACCGCTTCATAGTTTTCAAGATTGAGGAAAAACAAAAGCAAGTAATAGTGGAAAAGCTCGGTGAAC

CTACTCAGAGCTATGAGGATTTCACTGCTAGCCTTCCAGCTGATGAGTGTCGATATGCTGTCTATGACTT

TGATTTCGTGACCGCAGAGAACTGCCAGAAGAGCAGAATCTTTTTCATTGCCTGGTCTCCTGATACCTCA

AGAGTAAGAAGCAAGATGATCTATGCCAGCTCTAAGGACAGGTTCAAGAGGGAATTGGATGGTATCCAGG

TAGAATTGCAAGCTACTGATCCTACTGAGATGGGTCTCGATGTCATAAGGGAGCCGTGCCAATTAATTAA

TTGTATTACCTACTCTGCAATCAAAGTTCATACTATCTGGTTCACTTAAATGTAATGTTTTATAACCAAC

TTGGGAACTTAGAACTTGCTTATGATATTTGGGTTGTTGTTTCAGGAGTTCATACTCTTCACTCCTTGTC

TTGATTT

>FK826529.1 JutEST4856 Jute Plasmid library 20-45 days after sowing Corchorus capsularis cDNA similar to Thioredoxin H-type, mRNA sequence

TTGAACTTAAAACAGAGAAGCGTTTCTTTTTGTCTTTCAAAAGAACAAAGTTTTTTCTTTTTAGTGAAGA

AAACAGCGCTAAGCAAAAATGGCAGCAGCAGGAGAGGGACAAGTGATAGGTTGCCACACCGTGAGTCATG

GACGGAGCAGTTAAACAAGGGCAAAGAATCCAAGAGGCTGGTGGTTGTTGATTTCACCGCTTCATGGTGT

GGACCCTGCCGCTTCATTGCACCCTTCCTGGCTGAACTGGCCAAGAAGCTCCCTAATGTCACGTTTCTTA

AGGTTGATGTCGATGAGTTGAAGGAAGTTGCTGCCGAATATTCTGTGGAGGCTATGCCAACTTTCCTGTT

TCTCAAGGATGGGAAGGAGGTAGACAAGGTGGTGGGAGCAAAGAAAGATGAGCTTCAGAACACTGTGGCT

AAGCATATGGCTACTGCTTCTGCTTGATAAGTCATATATTTGGAACTACCTTCATACTTTGGAGGATATT

TATATCAATATATACTAGCACTCTGTAAGTTATGAGTTTGTGTGCATAATAGACAGTCTATGTAACTATG

TTAGACCTCTCAGTTTGTTCCAGTTCTTTGAGGATCATGCTGTGACCTTTTCTATGATGAAAACTAAGTG

AAGTTTTGTTTGGGTTATATATATC

>FK826528.1 JutEST4850 Jute Plasmid library 20-45 days after sowing Corchorus capsularis cDNA similar to Leucine-rich repeat protein, mRNA sequence

GAGTACCAACCAAAGTGGGAGTCTCAATATAGTCAGTGTGAGTATAAGGCTATAAGCCAGAGGTATTTTA

CAGACAACATTAAACCATACAAGGGTACATACCCGAAGTAATCTTGTTACTACCATTACATCAAACACTG

GTTTGTCTTAAAACATCATAGGAGTTTTTGGTATATATGCTAATAAATTCTGATTTACAGATACTTATAC

GAGATTGCATTTCCCCACACTGACTATATTGAGACTCCCACTTTGGTTGGTACTCTTTAAGGACACTGAC

TATATTGAGACTCCCACTTTGGTTGGTACTCTTTAAGG

>FK826527.1 JutEST48136 Jute Plasmid library 20-45 days after sowing Corchorus capsularis cDNA similar to Hydroxyproline-rich glycoprotein DZ-HRGP precursor, mRNA sequence

GTGAGACGAGGTGAGTCGTTGGATTCGAAGATTAAGAGCAGTAACGTGGAAATGAAGGCGGGTTGGTTGT

TACGGGTACGGATAGATTGGGTCCGGACCCCAAAATGGTTCCAAAACAGATCAAGATCGTGGATATTAGG

TCTCCGGTCACCGGGAATTCCGACGTTTACGCCGGATCGGCCTTTTCAGTATCTCCGCCCGCCGAGTTCT

CTTCCTTTGCCGTCGTTCTCAAAGAAGAAGCAGGTTTCGATCGATGATTCAGCTACTAGGGATCTCAGGC

GTCTACTCCGGCTCGACTTCTGAGACCCGAATAGAATCTTTCGGGTCCGGGTTGCCTTTTTCTCTCCATG

TTAATCTCTCTCCTGATCCGGATTTATATTCGGATCCCAGAAAAAGTCTCCGTCTCTTATCTCCTCTCTC

CCTCTCTATCTATCGATCGATCAGTTGTCTCTAATTAGAACTCTCCGGAAATTTTCGAAATCCAGTTTCC

TCCGTCATATTTTTTTCAAAACGGTG

>FK826526.1 JutEST48123 Jute Plasmid library 20-45 days after sowing Corchorus capsularis cDNA similar to Low temperature and salt responsive protein, mRNA sequence

GCCTCAAGTATGGTTGCAAGGTGGAGTTCTGGATCTGCCTTATACTGACCATCTTTGGGTACATCCCTGG

CATCATTTATGCTGTCTATGCTATCACCAAATAAGATGATGATCATCTTATAAATATTATATTACTCCAT

TTCTAATTTGTGGAGGACACAAATAATATTGCAGGTTAAGGTGGTGATTATTTTTGGATTCCTTGTTGTG

TTCGCTTGTCCTATCATTCTACTTGCTGGCCTCTGTTTTTCTTAGTAATTTTCAGTATAGCTATGTAGAT

ATTGTGTTGTTTTTTGATTCTCATGTAATTATATTGTCTGTTGGTGTTTGGTTTCATGATTATGATGAAC

TACTTGATCGATGATGATAATTATGGGGTTCGTCTATTTGATTTGATTTGATGTAAGAAAA

>FK826525.1 JutEST48113 Jute Plasmid library 20-45 days after sowing Corchorus capsularis cDNA similar to Formate dehydrogenase, mRNA sequence

CAGAGAGGATCAAGAAAGCCAAAAACTTGCAACTGCTTCTCACAGCAGGGATTGGATCTGACCATGTTGA

TCTGAAGGCAGCTGCAGAAGCTGGTTCGACAGTTGCTGAGGTTACAGGAAGCAATGTTGTCTCTGTTGCT

GAAGACGAGTTGATGAGAATTCTCATTCTTGTCCGGAACTTCTTACCTGGCTACCACCAAGTTATCACTG

GAGATTGGAATGTTGCCGGTATAGCTTATAGAGCTTATGATCTTGAAGGGAAGACAGTAGGAACTTTTGG

TGCAGGGCGCATTGGCAGGCTCTTACTCCAACGACTGAAACCATTCAACTGTAACCTCCTTTACCATGAT

CGGCTGAAGATGGATCCAGAGTTGGAGAAGCAGACTGGTGCAAAATTTGAGGAGGATCTTGATGCAATGC

TTCCAAAATGTGACATAATTGTCATAAACACGCCTCTTACTGAGAAGACAAGAGGGGATGTTCGACAAAG

AAAGGATTGCAAAACTGAAAAAGGGAGTCCTGATTGTTAACAATGCTCGAGGAGCTATCATGGATGCACA

GGCAGTTGCAGATGCTTGTGCAAGTGGACACATTGGAGGTTACAGTGGAGACGTTTGGGTTCCACAACCA

GCTCCAAAAGACATCCATG

>FK826524.1 JutEST48110 Jute Plasmid library 20-45 days after sowing Corchorus capsularis cDNA similar to ADIPOR-like receptor, mRNA sequence

AGATCAGAGTGGCCCTTGAAGCAAACTTTGCTCAGCATCTTCACAATTCACAACGAGACTTTAAATGTTT

GGACGCATTTGATTGGTTTCTTCATTTTCCTCTCCCTTACCATATACACTGCAATGAAGGTTCCAAAGGT

TGTTGATCTTCACTCTCTGCAGCATATTCCTGATGTGCTGAGAAGGGCCGATTTACACAAATTGCATTCA

GAATTGATGACCTGTCCTTCCCGATTTGCATAAACTTAGGGAGGAGTTGAATAACTCATTCCCTTCAATG

GATTTACTTCCATCAATCGCAGGCTGGCATGTCATGGATCTTCTCGTAAATTGTTTACCTGAGCGGTTTT

CCCACCGGAATCACACTGATGTTTGTGTTCTGCGAAGTGTAAAGGAGGATGTGGAAAACATAATAGCACC

GCTGATGGTGAGACCTATAACACGGTGGCCATTTTTTGCCTTCATGGGTGGTGCAATGTTTTGCTTGCTA

GCTAGCAGTACCTGCCACCTCCTCTCGTGCCATTCTCAGCGCCTATCATACATCATGCTCAGGCTAGATT

ACGCTGGGATTGCAGCTCTCATATCTACTTCTTTCTATCCTCCTGTGTATTACTCCTTTTATGTGTGACC

CATTCTTCTGTAACCTCT

>FK826523.1 JutEST48101 Jute Plasmid library 20-45 days after sowing Corchorus capsularis cDNA similar to Uridine monophosphate kinase, mRNA sequence

GGGGAGAAGCGTCCTCAGCGGCAGACCGGGCCCAAGTCCCCTGGAAGGGGGCGCCGGAGAGGGTGAGAGC

CCCGTCGTGCCCGGACCCTGTCGCACCACGAGGCGCTGTCTACGAGTCGGGTTGTTTGGGAATGCAGCCC

CAATCGGGCGGTAAATTCCGTCCAAGGCTAAATACGGGCGAGAGACCGATAGCAAACAAGTACCGCGAGG

C

>FK826522.1 JutEST4837 Jute Plasmid library 20-45 days after sowing Corchorus capsularis cDNA similar to Multidrug resistance-associated protein MRP1, mRNA sequence

AGAGTTATGGAAAAGTCTTGAGCGTTGCCAACTTAAAGATTCGGTAGCTTCAAAACCTGAAAAACTCGAT

TCTCCAGTGGTCGGTAATGGTGACAATTGGAGTGTGGGACAAAGGCAGCTTTTATGTTTAGCAAGAGTTA

TGCTAAAGCGCAGCAGGCTATTGTTCATGGACGAGGCAACTGCATCAGTCGATTCTCAAACCGATGCTAC

CATCCAAAAGATCATTAGAGAGGACTTTGCAGCCTGCACAATCATCAGCATTGCTCACAGAATACCCACA

GTCATGGACTGTGACCGGATTTTAGTCGTAGATGCAGGAAGGGCAAAAGAATTTGACACACCGACCGACT

TGCTGGAGAGCCCAACGCTCTTTTCAGCCTTGGTTCAGGAATACGCCAACCGCTCTGCTGGACTATAAGA

AACAAGACAAAGACAAATGGTCTCTGCTGTTCCGCTTGCATTTTGTTTCTGCTAATTCGGTCAATAATCT

TCTGCAGACTTCTCAATTAATAATCGTCTCATGAATTCTATGGTGAAATATAGTTTTCATGTACCAAATG

TACAATGAACTCTCGCACTCAAGAGCTATCTATCTAACCTTCATAATATATATTGTGCGATTTTGTTTCT

GAACAAAAAAAAAAAA

>FK826521.1 JutEST4816 Jute Plasmid library 20-45 days after sowing Corchorus capsularis cDNA similar to Peroxidase,complete, mRNA sequence

ACTACACAAATCTCCAGAACAACCGTGGCCTTCTCCAAACCGATCAGGATTTGTTTTCAAGCACCGGTGC

AAGCACCGTGGCCATTGTGAACAGTTTTGCTGGTAGCCAAACTGAATTCTTCGATGCCTTTGTTCAATCC

ATGATCAAAATGGGAAACTTAAGCCCTTTGACAGGAACCAATGGAGAAATCAGGACTAATTGCAGAAGGA

TCAATTAGAAACGTTGCTATATATATATATAGCTTCGGCTGAAACTTGTTTGATATATATACTTTTTTTG

TGTGAAGGATATACTAATGATATCCCAATATATATATACTTTCTTTGGAAATTGTATAATAAGGGCAATG

AAATCTTTTGTCCATTTTCATGTTAGT

>FK826520.1 JutEST489 Jute Plasmid library 20-45 days after sowing Corchorus capsularis cDNA similar to Ubiquitin Carboxyl-terminal Hydrolase-like zinc finger protein, mRNA sequence

TATTAAAAATAATTATTATTAATAGATCGGTAATCCAATATTTGTATTACATCTCAATTTACCGGTAGAA

TATTTTATGATAAACAATAATTTCAGAACAGTACAATAGTATAGAGTAAGATCAGAAAATGTACTATATT

AATACACAAACTACTCCATATGCTAATATTATGGAAAACATAATTACAAGTAGATCATCAGATGCATAAT

TATGCCAAATATATAGCCTTAAAGCAGCTGACTGGCGGTGGGGATTACCGATGCCGCGATTTGTAATTAA

TGTGGATGATGATGATGGTGAGGACCTGGTGGAGGAGCGTGGGGAGCTGGGGCAGGATGAGGATGCTGCG

GTGGACCTGGATGCGGTGCATGATGTGGTGGGGGGTGGTGGGGATCATGGTGGGGCTGTGGTGGTGGTGG

TGGATTGTGCGGTGGTGGGTGGTGGTGGTCATGGTGGTGTGGTGGTGGTGGATTATGCATCTTCAATTTT

CTTTTTTTTGCTATGTTAGGCTTTGTCTAGATATGGTGTGATTTAGCGAAGAT

>FK826519.1 JutEST484 Jute Plasmid library 20-45 days after sowing Corchorus capsularis cDNA similar to Serine/threonine kinase, mRNA sequence

ACCTCTTGATGTTGATGCAGTTTTCTGGTTCATGATCAGGCGTAATGTCACCGCATTTGATGCGAAGAAA

GATATGGCTAAGCTCACCAATTTGAATGCCTTTGACATTATTTCTCTTTCAAGTGGGTTCGATTTGTCTG

GACTATTTGCAGATAATGATGAGAAGAAGGAACTGCAGTTCACTTCCATGCATACTGCCTCAACCATCAC

ATCTAAACTAGAGGACATTGTCAGCGTTTGAAGTTGAAAGTTAAGAAGGAAGGAGGTGCTATGCTGAAAC

TGGAAGGATCAAATCGAGGTAGAAAAGGCTCATTGGCTATTGATGCAGAAATATTTGAGTTCACCCCTTC

TTTCCATTTAGTAGAACTGAAGAAGTCCAATGGTGATACTTTGGAGTTCCAACAAAACCAGAAAAAGGAT

GTTAGACCAGGTCTCAAGGACATTGTTTGGGCTTGGCAAGGTGAACGCCAGCAACAACAACATGCATATC

ACGGTCATCCTAACTACAGGTTCAATGTCTAAGCATCTCAAGTTTGTTTTCTAGGTTGTGTTCAATTTAT

ATGTTTTAATAAATGGTAGCAAAACAAGAAGTTGCCCGTCTAGAACTCTAATTAAAATGACCAATTCCTA

CCAAAACTTTGAC

>FK826518.1 JutEST47139 Jute Plasmid library 20-45 days after sowing Corchorus capsularis cDNA similar to Aux/IAA protein, mRNA sequence

AAATTATAAAACCATCAAGGCCTAGTGTTTGTGAGTAACCTTTTTACTATTGAATCAAGTTTTAGAGTCT

CTGGACAACTCTCATGTTCCCACGCAGGATCTCTAGTTTGTAAAGTGCATTGTTTGTTAAAAACTTTCAT

CGGTGTGTGTTTGTGTGATGAAAGACCTCGACATTATCTGTAACCCCGACATGATTGTATTAACTCTATT

GTGCGGCTGCTTGTGAGTGTTGTATTTCTTCTGCTTAAGTATTATTCTGCAAATCATTTTAGTTCCTATT

CTGCTAACTATAACCCCAATTCCGAAAACCGGTCGGGGGCTTTGAAAATGCATATGTACTGAATCATATT

AGTAGTAGTTTTATAATGTGTTTATGAGTTCCAAAAAAA

>FK826517.1 JutEST47118 Jute Plasmid library 20-45 days after sowing Corchorus capsularis cDNA similar to Protein phosphatase 2C-like, mRNA sequence

ATCAATTGAACACAAAATTCATCAATACAGTTTGATCATTTGGAAACAATCTTACTTGAAAACTTATTCT

GATATTGACCTTGAGCTTAAGCAGCATCGCTGGATAGATTCTTTCCGTAGTGGGACAACAGCCTTGACAA

TCATCAAACAGGGTGAACTCATTGTTATAGCCAATGTTGGTGATTCTCGGGCGGTACTTGCAACCACATC

GGATGATGGCAATTTGGTTCCACTTCAGCTTACAGTTGACTTCAAGCCTAACATCCCTGAGGAGGCTGAA

AGAATAACGCAGTCAAAAGGGCGAGTGTTTTGTTTAGAGGATGAACCAGGGGTTTACAGGGTGTGGATGC

CTAATGGTGACTCGCCAGGGTTAGCCATATCAAGAGCCTTTGGTGATTACTGCGTCAAAGAGTTTGGCCT

CATTTCTGTGCCTGATGTCACCCAAAGAAACATCACAACCAAAGACCATTTCGTCATTTTAGCAACAGAT

GGGGTTTGGGATGTGATTTCCAATGAAGAAGCAGTGGAGATCGTTTCTTCGACATCAGAAAGGGAAAAAT

CAGCCAAAAGATTGGTTCAATGTGCCATCCGCGCTTGGAAATACAAGAAACAGAGGCATTGCAATGGGAT

GATATGTCGGCTATTTGC

>FK826516.1 JutEST47113 Jute Plasmid library 20-45 days after sowing Corchorus capsularis cDNA similar to Peptide methionine sulfoxide reductase, mRNA sequence

GGGAGAGTCAGGTCCAATACACCCAAAAACCCAACCTCCAACTCATAATAACAATGCTTCGAAGTCTTAC

GCTTGCCACCTCTTCATCTCTTTCTTCTTCCCTCATCTCCAAGCCTTTCTTTTCTCTCTCTCCAAAACTC

CCTCTCTCTTTCTCTCCCATTTCCCTCCCTTTCCCTCAAACCAAACGCTCCATTTCTCTCTACAGGGCCC

CCATGAACATCCTCAAGAACCTCGGCTTCGGAGCCAAGCCGGCCCAGTCAGCTGAAAACACTGCCATCGC

TCAGGGTCCGGACGACGACGTCCCGGCTCCCGGTCAGCAATTTGCTCAGTTCGGCGCGGGTTGCTTTTGG

GGCGTTGAATTGGCTTTCCAGAGAGTTCCGGGGGTCACCAAAACAGAGGTTGGGTATACCCAAGGGTTCA

TACATAATCCGACTTATGAAGACGTGTGTTCCGGGACCACGAACCATAGTGAAGTTGTGAGGGTTCAGTA

CGACCCTAAGGGAGTGTAGCTATGAGAGTTTGCTAGATGCCTTCTGGGCTCGCCATGATCCTACTACACT

GAATCGCCAGGGGGAATGATGTTGGAACACAGTACAGATCCGGTATATACTTTTACACTCCTGAGC

>FK826515.1 JutEST47111 Jute Plasmid library 20-45 days after sowing Corchorus capsularis cDNA similar to Acidic elicitin A1 precursor, mRNA sequence

AAAAACGGAGTGAAAATTCTGTGTTTGAGGCAGCGCGCGGTTGACGGCGATCGGAGAATTGCTCGGTGAG

AAGAAAGCTATATCGTTCTTGAAGGGAAAGGCTTTTGTTACCAAGATTGCGCCGATAGCTACGTACGCAG

TACTTCCGGCTCCTATGTTTGCCGCTCTTTTTTACTCTCCTCCCGATTATGGTTCTGCTTACAAGGGCTT

AGAAAAACCCCCCAAATCTTCATAGTTTGTCCTTTTGTTTTTCTCCTGGAGTACTCTTTGACTTGCTGAA

GGTTTGCATGTCATGTAGCCTCTATTCATAATAGATGGGGTGTCTCCTAATAATTGCTGCATATGTTCTT

AGGTATGTAGAGAGTTGAATGGCTTTTGTTGCCTTTGAAGCACCTTTGAGCTATAAATCTTCTGTTTTTC

TGAGAGTTTGTCAAAGACTAAAACATTATGTAATGAAATCACAATTCCCCCATTAGTTATACTGTTGAAG

TGTTGTGTTATAAAATTATATCAATGCTCTTTGTGCTGTCTAAAAAAAAAAAAAAGTAAAAAAA

>FK826514.1 JutEST47101 Jute Plasmid library 20-45 days after sowing Corchorus capsularis cDNA similar to BLE1 protein, mRNA sequence

GGGGGTGATTTTTAGGGTTTGAGTGACAGAGGAGAGAGAGAGATGGGAGGAGGGCATGGAGAAAGCACAA

CTTACAAAGGTGTAACTTTGCATCACCCAAAGAGATGGCATACCGTCACCGGCAAGGGCTTGTGCGCTGT

CATGTGGTTTTGGGTCTTGTACAGGGCTAAACAAGATGGTCCTGTAGTATTGGGTTGGCGTCACCCTTGG

GAAGGCCATGATGGTCATTCTCACGGTCATGGAGACAAGCATTAGTGATTACATGATGCATGGCCTGGAA

TGACAGATCTTGTTCAGGGGTTGCTGGTGAGGTCTGTTCTCGAATCGTGGATGTGTTCTTTTCAAAAAAA

>FK826513.1 JutEST4786 Jute Plasmid library 20-45 days after sowing Corchorus capsularis cDNA similar to importin like protein, mRNA sequence

GCATTTTGGAGATGTATGAATACTGAAGAAGCTGACGATGAAGCTGATGATCCTGGTGCTCTAGCTGCAG

TCGGCTGTTTGCGTGCCATAAGCACAATACTTGAATCAGCTAGCAGGCTTCCACACCTTTTTGTTCAGAT

TGAGCCAACTTTGCTCCCAATTATGCGGAGAATGTTGACAACTGATGGGCAGAGGTATTTGAAGAAGTTT

GGGAGATTGTTTCATACATGACCTTCTTTTCTCCAACTATTTCCTTGGATATATGGAGTCTCTGGCCTTT

AATGGTTGAAGCATTGGCTGATTGGGCTATCGACTTCTTTCCAAATATTTTGGTTCCTTTGGACAACTAT

ATATCCAGGGGAACTGCATCATTTCCTTACTTGCAAGGAACCAGACTACCAGCAAAGTCTATGGACTATG

ATTTCATCAATTATGACCGACAGAAATTTAGAAGACAACGATATTGAGCCTGCTCCAAAGCTGATAGAAG

TAGTATTTCAAAACTGTAGAGGACAGGTGGATCATTGGGCTGAGCCATATCTGAGAATTGTAGTTGATCG

ATTGCGTCGAACAGAGAAATCACGCTTGAAATGTCTTCTTGTACAAGTGATTGCGAATGCAGTTTA

>FK826512.1 JutEST4774 Jute Plasmid library 20-45 days after sowing Corchorus capsularis cDNA similar to ADIPOR-like receptor, mRNA sequence

AGATCAGAGTGGCCCTTGAAGCAAACTTTGCTCAGCATCTTCACAATTCACAACGAGACTTTAAATGTTT

GGACGCATTTGATTGGTTTCTTCATTTTCCTCTCCCTTACCATATACACTGCAATGAAGGTTCCAAAGGT

TGTTGATCTTCACTGCTTGCAGCATATTCCTGATGTGCTGAGAAGGGCCGATTTACACAAGATTCATTCA

GAACTTGTGACCTGTCCTTCCCGATTTGCATAAACTTAGGGAGGAGTTGAATAACTCATTCCCTTCAATG

GATTTACTTCCATCAATCGCAGGCTGGCATGTCATGGATCTTCTCGTAAATTGTTTACCTGAGCGGTTTT

CCCACCGGAATCACACTGATGTTTGTGTTCTGCGAAGTGTAAAGGAGGATGTGGAAAACATAATAGCACC

GCTGATGGTGAGACCTATAACACGGTGGCCATTTTTTGCCTTCATGGGTGGTGCAATGTTTTGCTTGCTA

GCTAGCAGTACCTGCCACCTCCTCTCGTGCCATTCTCAGCGCCTATCATACATCATGCTCAGGCTAGATT

ACGCTGGGATTGCAGCTCTCATATCTACTTCTTTCTATCCTCCTGTGTATTACTCCTTTATGTGTGACCC

ATTC

>FK826511.1 JutEST4771 Jute Plasmid library 20-45 days after sowing Corchorus capsularis cDNA similar to Xyloglucan endotransglycosylase precursor, mRNA sequence

GGGCACACACAGAAAAATGGCTTCCACATCTTCTGCATTCGCTACTTTCGGCTCTTCCGCTGCTGGTTTT

ATGGGCTCCAAAGTTTCCGTCAAGGAACCGACTTCCATGCCAACACGTGTCCAGTTCCGGCCGCTCCGTG

TCTCCGCTGCCTGTAGTTCGACCGCCGAGAGAACGGTTTCCCACATTGCTTTTCCATCTTCACTCTACGA

GGTCTTGGGGATCCAGATGGGCGCCACGTGTCAAGAGATCAAGTCCGCTTACCGGAGATTGGCGCGAGTT

TTACACCCGGACGTCTCGGCCAACGGTCAGAATAACGCCACGGCACACGAGTTCATCAAAGTACATGAGG

CTTACGCGACTCTGTCTGATCCAGAGAAACGCGCCGATTATGACCGGACGTTGTTGTTCAGGACAAGGCG

GTCCTTTGCGATGTCCATCGCATCTATGGCCAGTTCCGCGGCTTCTGGATTTTCTGGGTACACAAGGCGA

AGCTGGGAAACTGATCAATGTTGGTGATACTAAATATAAATTGTACAAAATTATGAAATTGAGGGATTGG

ACTGTCTAGTTGTGTAGGCTACAGAGT

>FK826510.1 JutEST4755 Jute Plasmid library 20-45 days after sowing Corchorus capsularis cDNA similar to Ozone-responsive stress-related protein-like, mRNA sequence

GTTCACAGACACGATCTTTTTCATCTTCCAGGGCAGCTTAATCTCATTTCTTCTTTCTCGAGCTATCTGC

TTCTCCTTGACCTGGGACCGAGCAAGCCAGCTTTGAAAATGGCAGCAAAGTACATAATTTCTGCTGTAGT

TGGATCATTTGCCGTTGCTTATGTGTGTGATCAGGTTTTTTCTGACAAGAAGATTTTTGGAGGGACTACA

CCTTCGACTGTTTCAAACAAGGAATGGTGGGCAGAAACTGACAAGAAATTTCAGGCATGGCCCCGCACTG

CTGGTCCACCGGTGGTCATGAATCCCATCAGCCGCCAAAATTTCATTGTTAAGGCTGGTGAATCTTGAAG

CCAATTTTTTCTTCTGCTTTGTCAATGGTAATTCATTTGATTGATATTACTGTGAATGAAACTGTATTGC

GATGTTTCAGTGCTGGCACTTAGCCATTTAGAATCACTCTTATCAATGGAAATTGTGCTGTGCCTTTCTT

ATTTTAAATCTCAATTCCTGGATCTGAGGATCATTTGAGGAATGATGTTAATCTACAATCTGAATAAATT

TCTGTGAAGCAAATGCCAAACTTATTTTTTTTC

>FK826509.1 JutEST4748 Jute Plasmid library 20-45 days after sowing Corchorus capsularis cDNA similar to Eukaryotic initiation factor 4A-2, mRNA sequence

CATTAGGTGATTACCTTGGTGTTAAGGTTCATGCTTGTGTAGGTGGGACCAGTGTCCGTGAAGATCAAAG

GATTCTCTCCAGTGGAGTTCATGTGGTTGTTGGTACACCTGGTCGTGTGTTTGACATGTTGAGGAGACAA

TCTCTCCGCCCTGATCACATAAAGATGTTTGTGTTGGATGAAGCAGATGAAATGCTCTCCCGAGGTTTCA

AGGATCAGATCTATGATATTTTCCAATTACTGCCACCAAAAATTCAAGTAGGGGTTTTCTCTGCTACAAT

GCCTCCTGAAGCCCTTGAGATCACAAGGAAGTTTATGAATAAACCTGTGAGAATTCTGGTGAAGCGTGAT

GAACTTACCCTTGAGGGTATTAAGCAATTTCATGTCAATGTTGAGAAGGAGGAATGGAAGCTTGAGACGC

TATGTGATCTCTATGAAACCTTGGCAATCACCCAAAGTGTCATTTTTGTGAACACTAGGCGTAAAGTGGA

TTGGCTTACAGACAAGATGAGAGGTCGTGACCATACAGTGTCTGCCACCCATGGAGATATGGACCAAAAT

ACAAGGGACATCATCATGCGGGAATTCCGATCTGGATCCTCTCGTGTGCTGATCACTACTGACCTCT

>FK826508.1 JutEST4742 Jute Plasmid library 20-45 days after sowing Corchorus capsularis cDNA similar to Wound-induced protein 1, mRNA sequence

ACCAGTTCTTTCTGGTTTTCGTTTATCCAAATCTGAAACAACTCTCCTTAATTAAGGAACCCAAGAACTC

TTTTCATCTTTCTTTTGTGCCTATCGATCGGTTTTCTTGGTTCGTTGTCTCTTTCAAGAACTGGCTAACT

CGCAGGAAAAAATTCTGGAGGAAACCAACAAGAACCAAGAGGTGGTTCTTGCCCTCTATGAGGCCTTAAA

GGCTCGAGACGTAGAAACCGTTCACCAAATCCTCGCACCAGACCTAGAGTGGTGGTTCCATGGTCCGCCC

ACCCACCAGTTTTTGATGCGCCTTCTCACTGGAGCTTCAGCCGACGATTCCTTCCGTTTCGACGTTGATC

CTAACTCCGTCACCACCTTTGGGTCCACCGTAATCGTCGAAGGCTGCGATCACAGCCGCTCAATCTCCTG

GGTTCACGCCTGGGACTGTTTCTGATGGGATAATCACACAAGTGAGGGAGTACTTCAACACTTCTCTCAC

CGTCACTCGCCT

>FK826507.1 JutEST4741 Jute Plasmid library 20-45 days after sowing Corchorus capsularis cDNA similar to Wound-induced protein 1, mRNA sequence

AGACCAGTTCTTTCTGGTTTTCGTTTATCCAAATCTGAAACAACTCTCCTTAATTAAGGAACCCAAGAAC

TCTTTTCATCTTTCTTTTGTGCCTATCGATCGGTTTTCTTGGTCCGTTGTCTCTTTCAAGAACTGGCTAA

CTCGCAGGAAAAAATTCTGGAGGAAACCAACAAGAACCAAGAGGTGGTTCTTGCCCTCTATGAGGCCTTA

AAGGCTCGAGACGTAGAAACCGTTCACCAAATCCTCGCACCAGACCTAGAGTGGTGGTTCCATGGTCCGC

CCACCCACCAGTTTTTGATGCGCCTTCTCACTGGAGCTTCAGCCGACGATTCCTTCCGTTTCGACGTTGA

TCCTAACTCCGTCACCACCTTTGGGTCCACCGTAATCGTCGAAGGCTGCGATCACAGCCGCTCAATCTCC

TGGGTTCACGCCTGGACTGTTTCTGATGGGATAATCACACAAGTGAGGGAGTACTTCAACACTTCTCTCA

CCGTCACTCGCCTTGGCAACTCCACCCAATCTCCGCCGTCTAACTACGCATCCTCGACGGCCGAGATTAC

CTCCGTGCATTGCCCTTCCGTTTGGGAGAGCAGCTTTTCCCACAGGGTTGGAAAGTCCGTGCCGGGGCAT

GTC

>FK826506.1 JutEST4738 Jute Plasmid library 20-45 days after sowing Corchorus capsularis cDNA similar to CCR protein, mRNA sequence

GACTCACCACTTTTTTGATCAACAAAATTTATTAAACAACCATCATCATCCGATCACAAATATTCAAAAT

GCAGGCTGCAGCAAGCATTTCACCAATAATGCACCAACCTCTAACAACCTCCAAGAACTTTTCCCCAAAA

TCTATTTCTACTCAGCCGCTAGTAGGCAGCTCACTTGCTTCCAGACGTGTTTCAATCAAAGTAGCTGCTA

CAACAAGCAATGACCCTTCCACATTTGATTACAACTCCATCTTCTCAGTGTTTCCAGCAGAAGCATTTGA

GACAGTGGGTGGAGATGCTTGCTCGGGCCGACATGTACCCGGAAGTCAGGCTACAACCGGAGGCCAGAAA

TGTCACATCCGCCGCCGCAGCTGAGTTGATCGACAGAGAATATCTCCAATACAATGAAGCCAAAACGGTT

TTCTGTGCTGAGGCTTGTGATGATCTGGGGGGAGAATTCTGCAGCCGAGAGTATCAAAGGGGAGTCTATT

AGAAATCGAAAGCCAATTCCAGGAGATTTGCGCAAGGATAAATATATATGTTATATATAGGAATTTAGAA

TTTTCAGTTTGGTTGTGTGTAGCTACTAAATTTCCATAATAATGATATACATATATATG

>FK826505.1 JutEST4737 Jute Plasmid library 20-45 days after sowing Corchorus capsularis cDNA similar to ATP binding, mRNA sequence

GGGTTCAGTCCGTTCTGGACTCTTTTTTCATGAAATAAATTAATTTCTGTCTCAAGCTGCTTCTTCTTCT

TCTTCTCCCCGGTTTTCCTTCTCTTAAATCTGCCGATTTGAATCTCGCAACGACGCCGTTTTAGTTATAA

TCTCCAGAAAGATTTTATAGATTCCGTCAATCAAAAACCATTTCCGACGAGAA

>FK826504.1 JutEST4728 Jute Plasmid library 20-45 days after sowing Corchorus capsularis cDNA similar to Photosystem II reaction center W protein, mRNA sequence

GGAAATAAAGACTATTTTCACATTTTTGCACTTAACAATTCGTGTCACATAAAATTAAATCTCCTCCTTC

TTCCATACACTAATTTCTTTCCAATTTTGGGTGTTTGATCCAATTTGGGTTCAAAGAAAATGGCAACCAT

TACTGCCAGCTCCTCAGCAACTTTGCTTGCACGTCCTAGCCTAAGGCAACTTTGAAAGGATCAAGCTTCA

ATCATCTCCAAGTCTTGGACTTCCTGTGATTGCAACAAATGGAAGGGTTAAGTGCTCCCTAGAGCAGAAG

CCATCTGTCAACGGCAAGAGCTCGAACACAAGCATGGTTGCAACACTGATGGCGGCGGCAGCGGCTGCAA

CCACGACGAGCGCGGCCATGGCATTGGTGGAAGAGAGGATGTCAACAGAAGGAACAGGACTTCCTTTTGG

TTTGAGTAACAACCTTTTTGGGTGGATCCTTTTTG

>FK826503.1 JutEST4720 Jute Plasmid library 20-45 days after sowing Corchorus capsularis cDNA similar to Glycerol-3-phosphate acyltransferase, mRNA sequence

GGCTATGGAAGTTTTGCTCCTGTCATTTTAATCTCGGTTGCTTGTTTCATTAATTGGTCAAGTAAGTGAC

TTTTGTAGTCAGGGCCATATATCATATATGTTGTTTAGCATTGACTTTAAGAATTGTAATCGATGCAAAT

GGATCGAGTGTCTATCACTCCATGAAGTTCTAATTGATAATGCAACTTTGTTTTTAAGGTGC

>FK826502.1 JutEST4715 Jute Plasmid library 20-45 days after sowing Corchorus capsularis cDNA similar to FH protein interacting protein FIP1, mRNA sequence

GCTACCAGCGCTGTCACTACCATGCCGGCTGAGTCCAATCCCTACGTGTCTCCTGCTCCCGCTCCATCTT

CTTCCTCTAAGAATACAGTGGAGGCGGTGAAGGATGTGTTAGGGAAATGGGGTAAGAAGGCAGCAGAGGA

TACTAAGAAGGCGAAAGAGATTGCTGGGAACATGTGGCAACACTTGAAAACTGGGCCAAGTCTTGCTGAT

GTTGCTGTAGGAAGAATTGCACAGGGTACTAAAGTTCTTGCAGAAGGTGGTTATGAGAAGATCTTTCGAG

GAACATTTGACACTGTTCCTGAGGAAAAGCTTCTGAAAACATATGCGTGCTACCTGTCCACCTCGGCTGG

TCCTGTTATGGGAGTGATGTATCTATCAACTGCAAAGCTCGCTTTTTGTAGTGACAATCCACTGTCTTAC

CAAGTTGGTGGTCAGACTCAGTGGAGCTATTATAAGGTGGTTCTACCATTACATCAGCTCAGGGCGGTGA

ATCAATCGGCTAGCAAAGCCAACCCAG

>FK826501.1 JutEST46137 Jute Plasmid library 20-45 days after sowing Corchorus capsularis cDNA similar to 5-methyltetrahydropteroyltriglutamate--homocysteine methyltransferase, mRNA sequence

GCAACAGAAGAGATTGCCGACAGAATCAACAAGATGCTTGCTGTGCTCGAGACCAACATCTTGTGGGTTA

ATCCTGACTGTGGACTCAAGACTCGTAAGTATGCTGAGGTGAAGCCAGCCCTCCAGAACATGGTTGCTGC

CGCCAAGCTCCTCCGCACCCAACTTGCCAGCGCCAAGTGAAAACCTCATGATGAGAAACCGGAGATCCTT

CCTTCAGTGATTTATTATACCCAAAAGATGAGAGATGAAAGAGAAAAAAGAAAAAAAGAGAAAAATTTCT

GTCTTATCATTTGAATAACTGTGTTTTCTTGATGCAATATTTGTTAGGCATAATATGCCCATGGGGCCAA

TTGATTTCCTTCTTTTTTGGTTGTTTTCCTTTTTACTAAATATTTTGTATTTTTGGAATATATATCCAGC

CTTCAATTTCTTTGAAAGA

>FK826500.1 JutEST46126 Jute Plasmid library 20-45 days after sowing Corchorus capsularis cDNA similar to RAB1Y, mRNA sequence

GGAACTTCGCAAGAGATAATGGTTGCCTTTTCATTGAATGCAGTGCTAAAACACGTGTTAACGTACAGCA

GTGCTTTGAAGAGCTTGTTCTGAAGATTCTAGACACTCCCAGCCTTTTAGCTGAGGGCTCCAAAGGTGTC

AAAAAGAACATCTTCAAACAGAAGCCGCCACAACCGGACGCATCTACTAGCAGTTGTTGCTGATATATTG

ATTCTTTTACTTATACTCCCAAATCTCTAACCAGCGTGTGGTGGTTCTCTCCCTGTTCTGACAAAGACTA

TTCATGGTTGATCTCTTTATTATATGAAACTTACATTCTCCTTGTAATATAGCATCGTTGCGATTTCTAT

TTCTTTTCCAGGC

>FK826499.1 JutEST46124 Jute Plasmid library 20-45 days after sowing Corchorus capsularis cDNA similar to transporter, mRNA sequence

GAAGGATTTGCAAACAATCCTACTGCAGTTGTGAACATGTCTGCTATGTGGCTTGTGCCACAATTTGTCT

TGAATGGCCTAGCTGAAGCTTTCACTGCAATAGGGCAGACAGAGTTCTTCTATTCTGAATTGCCCAAGAG

TATGTCAACTATTGCTGCCGCTCTCTTTGGACTAGGATTGGCAGTGGCAAATTTGCTAGCAAGTGTTGTA

GTGAGCATTATCAATGACATTACCTCGAAAGGCGGCAAAGAGAGTTGGGTTTCAAATAACATAAACAAAG

GTCAGGATTACAAGTACTATTGGGTTCTTGCCATATTGAGTTTCATCAATCTATTCTATTACTTTGTTTG

TAGCTGGGCTTATGGACCTTTGGTCGAACAAGCAACAAAAGGTAAGGGTGATGCTGGGGCAGAGGAACAG

GAGTTGAATGACAGTGAAGAAGAACAATCTAATATGGAAGATCAAGGTTAGGGATGAGAGTAATGGCTTG

AAGAATGGACAAATTCAGTAATCCAAAAAAAGGCCTTCCTATTTGCTAGCTTGATGATGAGTGTTTCAAG

TCTTGGAATGACTATTTCAGTTTAAGATTTGCCTACATAGTCTGCTGTTAGTAACTAGCTAGGAATATAG

T

>FK826498.1 JutEST46116 Jute Plasmid library 20-45 days after sowing Corchorus capsularis cDNA similar to Inhibitor of apoptosis IAP, mRNA sequence

GCATGGACTCTGAATCAAGCCTGGAGAATATCATCAAGAAAAATGGCTGGTGGACAAGAAGCAACTGGGC

ATTCCTGAACGAACCACCTGTTCTTGAACGCCCTACCAACAGTTACAAAGCCCAGTTCCACATAGCCAAC

CTTGCTGCTTCAAAGCTTAATAGTGGATTCAATGCGAAGTCAGTGCTTATCTTCAGCTATCTTAATCTTA

ATCCTACCTTGTGCTAGAAATATATAGCATAAAACTTGTTCTGCCTAAAATCTGTATATATAATGATGTT

CTACAAAGTTTGTAAGAACATTACCCTGATGTGTCGGCTGAGTGTGTTTAGTCATGAGTGTTGTCCTGAT

CTGGCGTCTGTACAAAACTAGTAGTAAAATGAGCTGTGTAATTTGCAGGTGGATGATATTTCGTAAGCTG

TCTGCATTATATAAAGATTTGTTTGAGATTTTTATCTAATGATAAGACTTCTTTCTTTG

>FK826497.1 JutEST46114 Jute Plasmid library 20-45 days after sowing Corchorus capsularis cDNA similar to Peptidyl-prolyl cis-trans isomerase, mRNA sequence

GCTACCCATCCAAACAGAAAAAATCCCCAATTTTACCAGTTCTTTAAAAAATCTCTTTAACCATGCCAAA

CCCCAAGGTTTTCTTTGACATGACCATCGGCGGCCAACCTGCCGGCCGGATTGTCATGGAGCTCTTTGCT

GATGTCACTCCTCGCACCGCCGAGAACTTCCGTGCCCTCTGCACTGGTGAGAAGGGTGTTGGCCGCAGCG

GCAAACCCCTCCACTACAAAGACTCAACCTTCCACCGTGTGATCCCCAATTTCATGTGCCAGGGAGGTGA

CTTCACCGCCGGAAACGGTACCGGAGGTGAATCCATCTACGGTTCCAAGTTCGCTGACGAGAACTTCGTC

AAGAAGCACACCGGACCCGGTATCCTCTCCATGGCAAATGCTGGACCCGGAACCAACGGATCTCAGTTCT

TCATTTGCACCGCCAAAACCGAGTGGCTCGATGGAAAGCACGTCGTGTTTGGACAAGTTGTTGAAGGCAT

GGGACGTGGTTCAAGCCATTGAGAAGGTTGGGTCAAGCACTGGAAGGACTTCCAAGGCCGTTAAGATTGC

TGACTGTGGGACAGCTCTGTTAAATTGGCAAACCTGGAGTCGTTTATGTTGACTTTGGGTGGTCT

>FK826496.1 JutEST46113 Jute Plasmid library 20-45 days after sowing Corchorus capsularis cDNA similar to RAB1C, mRNA sequence

GACTTCTTTGAATTCTCCTTTGATCTCACGGTAATAGAACTTCTTCTCTCCGTCTCTCTTTGTCTCCGGC

GACGATCTCAAATCATCGTTAAGTCTAGCCTTAAGTCTAATTTCCGGTTCGTTCTCCGTCATCTCCGCCG

ACTTCTCGTGATTCAGTTTATTCAAACTCGTTGCCATCATGAATCCCGAATATGATTATCTGTTTAAGCT

TTTGCTTATTGGAGATTCTGGTGTCGGAAAATCATGTCTACTTTTGAGGTTTGCTGATGACTCCTATCTT

GAGAGTTACATCAGTACCATTGGTGTTGACTTTAAAATCCGCACTGTGGAACAAGATGGAAAGACCATCA

AACTTCAGATTTGGGGACACTGCTGGCCAAGAACGTTTCAGGACAATCACTAGTAGCTGCTATCGTGGGG

C

>FK826495.1 JutEST4695 Jute Plasmid library 20-45 days after sowing Corchorus capsularis cDNA similar to Oxygen evolving complex 33 kDa photosystem II protein, complete, mRNA sequence

TTCATCTTTCTTGGACCCAAAGGGCAGAGGTGGTTCGACCGGTTATGACAATGCAGTTGCCTTGCCAGCT

GGAGGAAGAGGAGATGAGGAAGAACTTGCTAAGGAGAACAACAAGAGTGCTGCTTCTTCATCAGGAAAAA

TCACCTTGAGTGTCACCGGGACTAAACCCGACACCGGAGAAGTAATTGGTGTGTTTGAGAGTGTTAAGCC

TTCTGACACTGACTCGGGTGCCAAGACACCAAAGGATGTTAAGATCACTGGTATCTGGTATGCTCAGCTT

GAACAATAGAGAGGGTCTTGTGTCTCAAATGTGCATTTATTCTTGCTGCAGATTTTGCTACATGTTGAAG

GTGTTGCTAAACTTTGCTGTAACTCCAATGAGAAACACTGAGATATATTATCCTTCTCTTTGTATATTAT

TATGTTGCCTTC

>FK826494.1 JutEST4691 Jute Plasmid library 20-45 days after sowing Corchorus capsularis cDNA similar to Small GTP-binding protein, mRNA sequence

GAAGCTATTAGACTGAAAAAAGACTAGAACTTGAAAGTAAAAAAAAGAGAAAAAGAACAAGAGAAGAGAA

AAATGGCTAGTGGAGGGTATGGAGACCCAAGCCAGAAGATTGACTATGTCTTCAAGGTAGTTTTGATTGG

TGACTCTGCCGTGGGTAAGTCTCAGATACTTGCTCGCTTTGCTAGAAACGAGTTTAGTTTAGATTCAAAG

GCTACCATTGGAGTTGAGTTTCAGACAAGAACTCTTGTTATTGAACATAAGAGTGTCAAGGCTCAGATCT

GGGATACTGCTGGTCAAGAACGATATAGAGCTGTTACAAGTGCATACTACCGAGGTGCTGTTGGAGCAAT

GCTGGTTTATGATATAACGAAACGCCAGACTTTTGATCACATCCCTCGTTGGCTTGAAGAGTTGCGTGGC

CATGCCGATAAGAACATTGTTATTATCCTGATTGGAAACAAATGTGACCTTGAAAACCAGCGGGCAGTTC

CAACAGAGGATGCCAAAGAATTTGCTCAGAAGGAGGGACTATTTTTCTTGGAGACCTCAGCTCTTGAATC

AACAAATGTGGGAGACTGCCTTCACAACTGTGCTTTATGGAGATCTTCAACATTGTGAACAAGAA

>FK826493.1 JutEST4689 Jute Plasmid library 20-45 days after sowing Corchorus capsularis cDNA similar to Alpha-tubulin, mRNA sequence

GAGAGAAGGAAAATTGTGGATTTGTTTGTTTCCCTTTACTATTTCCCACCTTCCTATTTTCCCCATAAAA

AATCTTCTGATTAAAAAGTGAAGTAGCAGGAGGAGGAGAATAAGAGGAAAGAATCCTTCTGTATTTGGTA

TTGTGATCAGCCATTGTTTCTTAGAATATTTATTTTAACACTTTAGACATTGAAAGAGATCATGAGGGAG

ATTATAAGCATCCATATAGGCCAAGCTGGAATTCAGGTTGGAAATGCTTGCTGGGAGCTCTACTGTCTTG

AACATGACATTCACCCTGATGGAATGATGCCCGGTGACACTTCTGTGGGTGTTGCACATGATGCATTCAA

CACTTTCTTCAGTGAAACTGGTTCTGGAAAGCATGTTCCTAGGGCTGTCTTTGTTGATCTGGAACCAACT

GTCATTGATGAAGTTAGGACTGGTCCTTACCGACAACTTTTCCACCCTGAACAGCTTATTTCTGGCAAGG

AAGATGCTGCCAACAATTTTGCTAGAGGCCATTACACAATTGGGAAGGAGATTGTAGATCTCTGCCTTGA

TCGAGTTAGGAAGTTGGCTGATAACTGCACTGGTTTGCAAGGCTTTTTGGTGTTTAATGCTG

>FK826492.1 JutEST4686 Jute Plasmid library 20-45 days after sowing Corchorus capsularis cDNA similar to Peroxidase precursor, mRNA sequence

AGGTCAGTGACCTTTGAGTGAGGAAATAATGGTGCCAAAGTTGTCTTCTTCTTTGCTTTGCTCTGTTTTG

CAGCTGTGTCTGCATTTCAGGATGAGGAAGACCAAGGCCTTGTTATGAACTTTTACAAGGATTCATGTCC

TCAGGCTGAAGAAATTATCAGAGAGCAAGTTAGGCTTCTTTACAAGCGCCACAAGAACACTGCATTTTCT

TGGCTCAGAAACATTTTCCATGATTGTGGTGTTCAGTCATGTGATGCTTCACTGCTGCTGGACTCGACCA

GAAGGACCTTGTCTGAGAAGGAAACAGACAGGAGCTTTGGCCTTAGGAATTTCAGATACATTGAAGACAT

CAAAGAAGCTGTCGAAAGGGAGTGTCCTGGAGTTGTTTCTTGCGCTGATATCCTTGTGCTCTCTGCAAGA

GAAGGAATTGTTTCGTTGGGAGGCCCTTACATCCCTCTAAAAACTGGAAGAAGAGATGGCAGAAAAAGCA

GAGCAGATGTTCTAGAAGAGTATCTTCCAGACCACAATGAGTCCATGTCTGTCGTCCTTGACAGATTTGC

TGCAATCGGAATCGACACCCCTGGGAGTCGTTGCCCTGCTAGGATCTCACAGTGTCGGTCGAACCCACTG

TGT

>FK826491.1 JutEST4683 Jute Plasmid library 20-45 days after sowing Corchorus capsularis cDNA similar to Acidic elicitin A1 precursor, mRNA sequence

GAAAAACGGAGTGAAAATTCTGTGTTTAGGCAGCGCGCGGTTGACGGCGATCGGAAGAATTAGCGTCGGT

GGAGCAAGAAAGCAATATGTCGCTGTTGAAGGGAGAGCCTTTGGTGACCAAGTTGGTAGCTTTGGCCAAG

TACGTGGTACTGCCGGCTCCTATGTTTGCCGCTCTTTTTTACTCTCCTCCCGATTACGCTCAAAAACAAA

CCTCCGCCACCACCAATTAGGTGCACGAGGGTAGCATGAAGGGTTTGCTTACAAGGGCTTAGAAAAACCC

CCCAAATCTTCATAGTTTGTCCTTTTTGTCAGCTTTTCTCCTGGAGTATCTTTATTGCTAAGGTTTGCAT

GTCATGTGCCTCTTTCATATAATGGGGTGTCTCCAATAATTGCTGCATATGTTCTTAGGTATGTAGAGAG

TTGAATGGCTTTTGTTGCCTTTGAAGCACCTTTGAGCTATAAATCTTCTGTTTTTCTGAGAGTTTGTCAA

AGACTAAAACATTATGTAATGAAATCACAATTCCCCCATTAGTTATACTGTTGAAGTGTTGTGTTATAAA

ATTATATCAATGCTCTTTGTGCTGTCT

>FK826490.1 JutEST4682 Jute Plasmid library 20-45 days after sowing Corchorus capsularis cDNA similar to Glycosyl transferase family 2, mRNA sequence

GGTGAATTGATTCCAATCATCTGAGATGATCATAATCTCAGTGCATCAAATGATGGCACAATTATTTTCT

CCCTATACAAATTGATGGCACAATTGGGAATTTATCTTGTCTACATGAGCCTTTGCTATTTACTAGTTAT

AGTTGTATAAATAGGCCTGTTAATATAGGGCAACATTAGCTTAGGTAGAATCAGAAATGCCATTGCAGTC

AACCAGATTACCAGAAGACAAAATTTGTATCTAAATATTTATATTACTGTCATTTTTAACTGTTATAGAA

AGGTATATATATAAGTAAATTTTCAT

>FK826489.1 JutEST4671 Jute Plasmid library 20-45 days after sowing Corchorus capsularis cDNA similar to Oxygen evolving complex 33 kDa photosystem II protein, complete, mRNA sequence

TTCATCTTTCTTGGACCCAAAGGGCAGAGGTGGTTCGACCGGTTATGACAATGCAGTTGCCTTGCCAGCT

GGAGGAAGAGGAGATGAGGAAGAACTTGCTAAGGAGAACAACAAGAGTGCTGCTTCTTCATCAGGAAAAA

TCACCTTGAGTGTCACCGGGACTAAACCCGACACCGGAGAAGTAATTGGTGTGTTTGAGAGTGTTAAGCC

TTCTGACACTGACTCGGGTGCCAAGACACCAAAGGATGTTAAGATCACTGGTATCTGGTATGCTCAGCTT

GAACAATAGAGAGGGTCTTGTGTCTCAAATGTGCATTTATTCTTGCTGCAGATTTTGCTACATGTTGAAG

GTGTTGCTAAACTTTGCTGTAACTCCAATGAGAAACACTGAGATATATTATCTTCTTTGTATATTATTAT

GTTGCCTTC

>FK826488.1 JutEST4669 Jute Plasmid library 20-45 days after sowing Corchorus capsularis cDNA similar to Expansin11 precursor, mRNA sequence

CAGCGCCGTGCTTACTGTTTTACTGATAGCAGCCCGCCGCCTTCGTCTGTCTTCTTTTATTCGATATTCG

AAATATATTGGCTCGATGTTAGCTGTAGTATATTGCGGCCATAGTAGTACGCCTGTGATCAAAAAGTTTG

TACATCTCAAACTTATTATATCCATGTATTCAAGTGTCAAATCTGAATCTTATACATATACTGTATTAAA

TGTCCCAATATACTCCCGTCTTTTATACTAAGCTTGTATTGTAATAAGGTCATATTATACATATATTTTA

TCAAATGTTCCTATGTTTTGCTTCAAAAG

>FK826487.1 JutEST4664 Jute Plasmid library 20-45 days after sowing Corchorus capsularis cDNA similar to Superoxide dismutase, mRNA sequence

ATTGTCATTTCTCAGTCATTTCTTTCTTTACTTTACTCTTTACTCTTGAAGGGGTGCACTGAGATCACAT

CGAAAATGGTGAAGGCCGTTGCCGTCCTTAGCAGCAGTGAAGGTGTTAGTGGAACCATTTTCTTCACTCA

AGAGGGAGATGGTCCAACAACTGTGACCGGGAATGTTTCTGGTCTTAAGCCTGGACTCCATGGCTTCCAT

GTTCATGCCCTTGGGGACACTACAAATGGTTGCATGTCAACAGGACCCCACTTTAATCCTGCTGGGAAAG

AGCATGGTGCCCCAGAAGATGAGAATCGCCATGCTGGTGATCTAGGAAATATCACTGTTGGTGATGATGG

CACTGCAAGTTTCTGTAAAACTGACAGTCAGATTCCTCTCACTGGGCCAAACTCTATTATTGGAAGAGCT

GTTGTTGTCCATGGAGATCCTGATGACCTTGGCAAGGGTGGCCATGAACTTAGCAAAAGCACAGGAAATG

CCGGTGGCAGAGTAGCTTGCGGTATCATTGGTCTGCAAGGCTGAAGTCCTTATCCCAGTGATCAGAATAA

TATGTGCAGTCGGAGATGCAATGATCTTGAGAATAACTCTTGTTATGTGAACCAGAGAGACTTTGTGAAT

AAAATGT

>FK826486.1 JutEST4657 Jute Plasmid library 20-45 days after sowing Corchorus capsularis cDNA similar to Beta-glucosidase precursor, mRNA sequence

GGGAGATGTAGCTTCGGATTTCTATCATAAATACAAGGAGGATATGTATTGAATGAAAGAAGATTGGGTT

GGATTCCTTCAGATTCTCCATTTCATGGTCTCGAGTTTTACCAAAGGGGAGGTTAAGTGAAGGAGTAAAC

GACCTAGGAGTCAAATTCTACAACGACCTCATCGATGAACTCATTAAGAATGACATCAAGCCTTTTGTGA

CTTTGCTTCATTTTGATCATCCACAAGCTATTGAAGAAGAATACGGTGCATTCGGAAATCGCAAGTATTG

TAGATGATTTTGCTGAATATGCGGAATTCTGCTTCAAAACATTTGGTGACCGAGTCAAGTACTGGGCAAC

CTTGAATGAACCATGGGTGGTCAGTATCCATCATCAAGACAGGTGGACTATGGCATCGGGTTGAACAAGG

CGGTAATGCCACTGAGTCGGGGCCCTTTCCCTTGGGCCATCATTTGATTCTTGCCCATGCAGCTGCTGCC

AGAATCTACATGGGACAAGTACCAGCCAACTCAAAAGGGAAAGATTGGAATTACCATAAATACCACTTGG

TACAAACCACTGAATGCCACAAAAGAACAGGAAGAGGCAGCGGGATAGAGCTCTTGGATTCGTTTTTGGG

ATGGTTTTGTGGAGCCCA

>FK826485.1 JutEST4647 Jute Plasmid library 20-45 days after sowing Corchorus capsularis cDNA similar to Peroxidase precursor, mRNA sequence

AACCAGTAGGTCAGTGACCTTTGAGTGAGGAAATAATGGGTTCCAAAGTTGTCTTCTTCTTTGCTTTGCT

CTGTTTTGCAGCTGTGTCTGCATTTCAGGATGAGGAAGACCAAGGCCTTGTTATGAACTTTTACAAGGAT

TCATGTCCTCAGGCTGAAGAAATTATCAGAGAGCAAGTTAGGCTTCTTTACAAGCGCCACAAGAACACTG

CATTTTCTTGGCTCAGAAACATTTTCCACGATTGTGGTGTTCAGTCATGTGATGCTTCACTGCTGCTGGA

CTCGACCAGAAGGACCTTGTCTGAGAAGGAAACAGACAGGAGCTTTGGCCTTAGGAATTTCAGATACATT

GAAGACATCAAAGAAGCTGTCGAAAGGGTGTGTCCTGGAGTTGTTTCTTGCGCTGATATCCTTGTGCTCT

CTGCAAGAGAAGGAATTGTTTCGTTGGGGAGGCCCTTACATCCCTCTAAAAACTGGAAGAAGAGATGGCA

GAAAAAGCAGAGCAGATGTTCTAGAAGAGTATCTTCCAGACCACAATGAGTCCATGTCTGTCGTCCTTGA

CAGATTTGCTGCAATCGGAATCGACACCCCTGGAGTCGTTGCCCCTGCTAGGATCTCACAGTGTCGGTCG

AACCCACTGTGTCAAAC

>FK826484.1 JutEST4634 Jute Plasmid library 20-45 days after sowing Corchorus capsularis cDNA similar to catalytic/ protein phosphatase type 2C, mRNA sequence

GATCAATTGAACAAAAATCATCATCAGTTTGAATATTTGGAAACAATCTTACTTGAAAACTTATTCTGAT

ATTGACCTTGAGCTTAAGCAGCATCCTGGAATAGATTCTTTCCGTAGTGGGACAACAGCCTTGAACAATC

ATCAAACAGGGTGAACATCTTGTTATAGCCAATGTTGGTGATTCTCGGGCGGTACTTGCAACCACATCGG

ATGATGGCAATTTGGTTCCACTTCAGCTTACAGTTGACTTCAAGCCTAACATCCCTGAGGAGGCTGAAAG

AATAACGCAGTCAAAAGGGCGAGTGTTTTGTTTAGATGATGAACCAGGGGTTTACAGGGTGTGGATGCCT

AATGGTGACTCGCCAGGGTTAGCCATATCAAGAGCCTTTGGTGATTACTGCGTAAAGAGTTTCGGCCTCA

TTTCTGTGCCTGATGTCACCCAAAGAAACATCACAACCAAAGACCATTTCGTCATTTTAGCAACAGATGG

GGTTTGGGATGTGATTTCCAATGAAGAAGCAGTGGAGATCGTTTCTTCGACATCAGAAAGGGAAAAATCA

GCCAAAAGATTGGTTCAATGTGCCATCCGCGCTTGGAAATACAAGAAAAGAGGCATTGCAATGGGATGAT

ATGTCGGCTATTTGCCTCTTCGTTCACTCCTCCCCTCAATCTCAACAAGTCCATGCTGTTAGTACACCAA

AATAGTGATCATAAAATGTAAAT

>FK826483.1 JutEST4626 Jute Plasmid library 20-45 days after sowing Corchorus capsularis cDNA similar to nucleic acid binding, mRNA sequence

TTTTGCAGCCTGCCTTCAACCCCAACTCGCACAATCCCAACTCGGTCTGGCCTTGCTCAGTTCGATCTCT

GGGGAGATAGCAATGCTTTTGAAGAGGAACCTGCAATGGAGAGGGTGGAATCTGGGAGAGACCTAAGGGC

AAGGATGTATGCCAAGCTGAGCAAGGAAAATTCTCTGAACCATGATCGAGTTGATCCTACCGGGTCGGGT

CCTGATGTTGGGTGGGTATCTGAGTTGGTGAAGTGAACTGGATGGAGGGGTAATAACTCACTCACTGCTG

TTTTGTTTATTTCTGTGGAATATCTGATTAATCGGACTTCGGTGGCTATCAATTTTCTGACTTTGTGTGA

ATATTTAATTGATGAAATTTCTGATTTGGGTTATCTGCACTGTAAATAGACGACGGGGAAAGGGAGATCG

GAAAAGTCTCTATTTTGTGGTGCTGCTGCTTCTTTCTTCATTTTGTCAATGGAAAAAAAAATGAAAATTG

CAAGTTTTGTATAGATAAGAAGGATTTATTAGGGCTGCCGGAAGATTGGGTAGCGGTCTGTTTTTCATTT

TGTCTACTATGAATCGTAGTAATTCCATTTCATTTTCTTCCTTTATCTTTGTAATTTCTTATCATGGCTA

ATCAATGTAATATATTTATTTCTTCA

>FK826482.1 JutEST45142 Jute Plasmid library 20-45 days after sowing Corchorus capsularis cDNA similar to Transporter, mRNA sequence

GGGAAGGATTTGCAAACAATCCTACTGCAGTTGTGAACATGTCTGCTATGTGGCTTGTGCCACAATTTGT

CTTGAATGGCCTAGCTGAAGCTTTCACTGCAATAGGGCAGACAGAGTTCTTCTATTCTGAATTGCCCAAG

AGTATGTCAACTATTGCTGCCGCTCTCTTTGGACTAGGATTGGCAGTGGCAAATTTGCTAGCAAGTGTTG

TAGTGAGCATTATCAATGACATTACCTCGAAAGGCGGCAAAGAGAGTTGGGTTTCAAATAACATAAACAA

AGGTCAGGATTGACAAGTACTATTGGGTTCTTGCCATATTGAGTTTCATCAATCTATTCTATTACTTTGT

TTGTAGCTGGGCTTATGGACCTTTGGTCGAACAAGCAACAAAAGGTAAGGTTGATGCTGGGGCAGAGGAA

CAGGAGTTGAAGAAGGAAGAAGAACAATCTAATATGGAAATCAAGGTTAGGGATGAGAGTAATGGCCTTG

AAGAATGGACAAATTCAGTAATCCAAAAAAAGGCCTTCCTATTTGCTAGCTTGATGATGAGTGTTTCAAG

TCTTGGAATGACTATTTCAGTTTAAGAATTTGCCTACATAGTCTGCTGTTAGTAACTAGCTAGGAATATA

GTTTTTGAAAA

>FK826481.1 JutEST45126 Jute Plasmid library 20-45 days after sowing Corchorus capsularis cDNA similar to Glyceraldehyde-3-phosphate dehydrogenase, mRNA sequence

CAACTGCCTTGCTCCATTGGCTAAGGTTATCAACGACAAGTTTGGCATTGTTGAGGGCCTTATGACCACT

GTTCATTCTATTACCGCTACCCAGAAGACTGTTGATGGTCCATCTATGAAGGACTGGAGAGGTGGAAGAG

CTGCTTGATGGAATCTCATTCCATGAAGGACTGGAGAGGTGGTAGAGCTGCTTCCTTCAATATCATTCCT

AGCAGCACTGGAGCTGCCAAGGCCGTCGGCAAAGTTTTGCCAGCACTGAATGGCAAGCTGA

>FK826480.1 JutEST45124 Jute Plasmid library 20-45 days after sowing Corchorus capsularis cDNA similar to Cytosolic ascorbate peroxidase 1, mRNA sequence

GGCTCAACTCGTGTCCATCGCGTGGTGGTTTAGGGTTTAAATTCATTTCCTGCAAACTCAATCACTTCAT

TTTCAAAAAGAAACCAGAGAGAAATGACGAAGTGCTACCCAACTGTGAGCGAGGAATACCAAAAGGCCGT

TGAAAGGGCCAAGAGGAAGCTCAGGGGTCTCATCGCCGAGAAGAACTGTGCTCCTCTCATGCTCCGTCTA

GCGTGGCACTCAGCTGGAACTTTTGATGTTAAGACCAGGACCGGAGGTCCATTCGGAACCATGAAGCAAC

CTGCTGAGCTGGCACATGGTGCTAACAACGGTCTCGATATTGCCGTCAGGCTTCTCGAGCCGATCAAGGA

GCAGGTCCCTATCCTCTCTTACGCTGACTTCTATCAGCTTGCCGGTGTCGTTGCCGTTGAGGTCACCGGT

GGACCTGAAGTTCCCTTCCACCCTGGAAGAGAGGACAAGCCTCACCCACCACCCGAGGGACGTCTCCCTG

ATGCTACCCAGGGAGCTGATCACTTGAGGCAGGTCTTTACAACTCAAATGGGTCTTAGCGACAAGGACAT

TGTTGCTCTTTCTGGTGGCCACACCCCTGGGAAGGTGCCACAAGGAGAGGTCTGGATTTGAGGGGACCAT

GGGACTACCAAC

>FK826479.1 JutEST45119 Jute Plasmid library 20-45 days after sowing Corchorus capsularis cDNA similar to Oxygen-evolving enhancer protein 2-1, mRNA sequence

AATTATTTCCCATACACAAAAACAAAACCCCCCAAAGCTAACCAAGACCACCACCACCACCACCACCAGC

TAGCAATGGCCTCAACCTCATGTTTCTTGCACCAGCATGCACTCACCACCGCCGCCGCCGCCAGATCATC

TTCTTCATCTTCACAGCGCCAATTTGTAAACCTCAAGCCTAGCCAGCTAGTTTGCAGGGCTCAGAAGCAG

GCCTGACCTCAAGAAGAGGATGGGACCAGCTTCGTTGTCTCTCGTCGGTTGGCTCTCACCGTCCTCATTG

GCGCCGCCGCCGTCGGCTCCAAGGTTTCCCCTGCTGATGCCGCCTACGGTGAAGCTGCAAATGTATTTGG

AAAGCCAAAAGAAAACACTGATTTTTTGACATACAATGGGGATGGATTCAAGCTGCGATTCCCATCAAAA

TGGAACCCAAGCAAAGAGGTTGAGTTCCCAGGTCAGGTTCTGAGATACGAGGACAACTTTGATTCAACCA

GCAATCTTACTGTCATGGTTACTCCCACTGACAAGAAGTCCATTACTGACTTCGGTTCCCCTGAGGAATT

CCTCTCTCAAGTGGACTATTTGCTTGGCAAACAAGCATACAATGGTAATACTGATGCTGAGGGTGGATTC

ACAACAGATGCATATAATATATAACATATGTAATAGCTATCATAGACTTCATTTTTGGTACCTCCCATTC

CCTTGTTTTTCGAATTTAAAGCAACCATTTACATACGTGGTTTGAGTACATACATGTGTGTGCGCGCGCG

CATATGCAGGGTGGTTTTGACTCCGGTGCGGT

>FK826478.1 JutEST45118 Jute Plasmid library 20-45 days after sowing Corchorus capsularis cDNA similar to Glutamine synthetase, mRNA sequence

GGGATTACAAGGCATGTCTCTATGCTGGCATTAACATCAGTGGAATCAATGGGGAGGTTATGCCTGGTCA

GTGGGAATTCCAAGTTGGACCAACTGTTGGAATTGCTGCAGGTGACCAAATATGGATGGCTCGTTACATT

CTTGAGCGAATCACTGAAATTGCTGGAGTGGTACTCACCTTTGACCCCAAACCAATTCAGGGTGACTGGA

ATGGCGCAGGCGCCCACACCAATTACAGTACCAAGTCCATGAGAAATGATGGTGGCATCAATGTGATCAA

AAAGGCCATTGAGAACTTGGGCAAGCGCCACAAGGAACACATTGCTGCTTATGGAGAGGGTAACGAGAGG

CGTCTAACAGGTCGTCATGAGACGGCAGATATCAATACATTCTCTTGGGGTGTGGCAAACAGGGGAGCGT

CCATCCGTGTTGGACGCGACACTGAGAAAGAAGGAAAAGGCTACTTCGAAGACAGAAGGCCAGCATCCAA

CATGGATCCCTACGTGGTGACATCAATGATTGCTGAAACCACCATTCTGTGGAAGCCTTGAAGGGTAGAT

ATTGGAGGTTTGCAGAGGATGAGAGGAGGCAAAGCAACATCCAGATCTAATTGAGTTCATACA

>FK826477.1 JutEST45113 Jute Plasmid library 20-45 days after sowing Corchorus capsularis cDNA similar to Calmodulin-binding protein-like, mRNA sequence

ACCCATCTCGAAGAAATACACGTTTGAAACCTCAAGTGGCAATGCAGGTGACGGTAGCAAATCAGAAAAG

AAAGAGAAGGAGAAGGAGAAGGAGAAAAAGATAGAGGCTAAGGAGAAGAAACAGAGAAGAAGGAGAAGAA

GAATCATAAATAGAAAAAGATAGAGGCTAAGGAGAAGAATGGACACTGCGTCATACGTCTTGTTGAAATC

GAAGAGCCGTCTGATCATGGGGCTGTTGTTTTGAGACAGGCTTTTGCCAAAAGAGCTGGTGTTATTGAGA

ATAAGAGGGGCAAGAAGAAGGAATTGTCTCCTCAAGATGCTGCTGCGATGATCCAAATTACATTCAGAGC

TTATCTGATCCGTAGGTCCCAGGCTCTTCGTGCCCTTAGGGACTTGGCAGTTGCCAAGACTAAGCTGAAA

GAGATCAGAGCATATTTCAATAACTTTAACTATCGTCGTCGAGTAGCCCAAGATGCAGCAGAACGTCAAA

AGTTCTCAGAGAAAATCATTGTCCTACTCCTCACAGTCGATGCCATTGAGGGAGTTGATGTAATGGTTCG

AGCTGCAAAGCGGTCAATGGTGGATGAGCTGGAGGCAATGCTTGATGTGGTGGACCCCCAGCCCCTTGGT

AAATCACTTTCAATGAGGAGAAGGACATTTGACATGCCGGATAGTG

>FK826476.1 JutEST4542 Jute Plasmid library 20-45 days after sowing Corchorus capsularis cDNA similar to Oxygen-evolving enhancer protein 2-2, mRNA sequence

GGGAATTATTTCCCATACACAAAAACAAAACCCCCCAAAGCTAACCAAGACCACCACCACCACCACCACC

AGCTAGCATGGCCTCCACCTCATGCTTCTTGCACCACCATGCACTCACCACCGCCGCCGCCGCCAGATCA

TCTTCTTCATCTTCACAGCGCCAATTTGTAAACCTCAAGCCTAGCCAGCTAGTTTGCAGGGCTCAGAAGC

AGGCTGCTCAAGAAGAGGATGGGACCCTTGTTTCCCGTCGGTTGGCTCTCACCGTCCTCATTGGCGCCGC

CGCCGTCGGCTCCAAGGTTTCCCCTGCTGATGCCGCCTACGGTGAAGCTGCAAATGTATTTGGAAAGCCA

AAAGAAAACACTGATTTTTTGACATACAATGGGGATGGATTCAAGCTGTCGATTCCATCAAAATGGAACC

CAAGCAAAGAGGTTGAGTTCCCAGGTCAGGTTCTGAGATACGAGGACAACTTTGATTCAACCAGCAATCT

TACTGTCATGGTTACTCCCACTGACAAGAAGTCCATTACTGACTTCGGTTCCCCTGAGGAATTCCTCTCC

AAGGTGGGACTATTTGCTTGGCAAACAAGCATACAATGGTAATACTGATGCTGAGGGTGGTTTTGAATC

>FK826475.1 JutEST4534 Jute Plasmid library 20-45 days after sowing Corchorus capsularis cDNA similar to beta-glucosidase, mRNA sequence

GCTGATTGACGGAAGGGGATTTAGCATATGGGATAAATTCACGAGGGAACATCCAGAAAAGATTTGGGAT

CGCAGCAATGGGAGATGTAGCTTCGGATTTCTATCATAAATACAAGGAGGATATAGTATTGATGAAGAAG

ATTGGGTTGGATTCCTTCAGATTCTCCATTTCATGGTCTCGAGTTTTACCAAAGGGGAGAGTTAGTGAAG

GAGTAAACGACCTAGGAGTCAAATTCTACAACGACCTCATCGATGAACTCATTAAGAATGACATCAAGCC

TTTTGTGACTTTGCTTCATTTTGATCATCCACAAGCTATTGAAGAAGAATACGGTGCATTCGGAAATCGC

AAGTTTGTAGATGATTTTGCTGAATATGCGGAATTCTGCTTCAAAACATTTGGTGACCGAGTCAAGTACT

GGGCAACCTTGAATGAACCAAATGGGTGGATCGTATCCATCATCCAAGCAGGTGGCTATGGATGGGTTGA

ACAAGGCGGTAATGCCACTGAGTCGGGGCCCTTTTCCTTGGGCCATCATTGATCTTGCCCATGCAGCTGC

TGCCAGAATCTACATGGACAAGTACCAGCCAACTCAAAAGGGA

>FK826474.1 JutEST4531 Jute Plasmid library 20-45 days after sowing Corchorus capsularis cDNA similar to MLP-like protein 34, complete, mRNA sequence

GGGATAGCACTAAAACCAGTACTCCTCAAAGCAAAAACTATACTATTCACTTAATCAAGAAATCTTGTTG

AAAATGGCGTCAAGGGGTCTAACTGGTAAGCTTGAGACCGATGTCGAGATCAAGGCTTCTGCCAAGGACT

TCCACGATATGTTCTGGAGCAAACCCCACCACGTCTCCAACACCTGCAATGACAAGATCCATGCATGTGA

TTTACATGAGGGTGAATACGGCAAAGTCGGTTCCATTGTCCTCTGGCATTACACACATGATGGAGTGAGG

AAAGTAGCAAAGGAGGTAATTGAAGCAGTAGATGAGGCGAACAACGCGATCACTTTCAGAGTGATCGAAG

GGGATCTGATGAAGGAGTACAAGAACTTTGTAATCCATATTCAAGCTACACCCAAGAGCGATGGTAACGG

CTGCATAGTTCACTGGATTCTGGGAATACGAAAAGCTGCACAAGGGAGTTGATCATCCTGAGACACTGCT

GCAGTTTGTCGCCGATGTCTCCAGAGACATCGATGCTCACCTCACCTCTGCTGCAAAGGCGTCATCGGGT

CTAACTGGTAAGCTTGAGACCGATGTAGAGATCAAGCTTCT

>FK826473.1 JutEST4528 Jute Plasmid library 20-45 days after sowing Corchorus capsularis cDNA similar to Hydroquinone glucosyltransferase, mRNA sequence

GCTCGTTGGATCAGATAACTGAATTGGCTCTGGGTTTGGAAATGAGCGAGCAAAGATTTTTGTGGGTTGC

GAGGGTTCCAAATGATAAAGTAGCTAACGCCACATGAGCGAGCAACGCTTTTTGTGGGTCGTGAGAAGCC

CAAACGATAAGGTTGCAAACGCAACGTTTTTCACTGTACAGAGCCAGAAGGACCCTTTTGATTTTCTACC

AAAGGGATTCCTTGAAAGGACCAAAGGCCGGGGTCTCATCGTGCCATCCTGGGCTCCACAGGCTCAAGTC

TTGAGCCACGGCTCCACGGGTGGGTTCTTGACCCATTGTGGATGGAATTCAATCCTCGAAAGTGTCGTTA

ACGGCGTCCCTTTGATCGCCTGGCCGCTTTACGCCGAGCAGAAAATGAACGCCGTGATGCTGACGGAGGA

TATCAAAGTTGCTCTCAGACCAAAACCCAGTGAAAATGGGTTAGTTTGCAGAGACGAGATTGCACGAGTC

GTTAAGGGTTTAATGGAAGGAGAAGAAGGGGAAAGGTGTCCGAAACAGAATGAAAGACCTAAAAGACGCA

GCAGCAAAAGTTCTAAGTGAAGAAGGGTCTTCCACCAAGGCACTCTCCGAGGTTGCCCAAAAGTGGAGGA

CTCAGGCTCAGGCTCAGGTTCGCCAGCATCTAGAACTACTTGTAGAAATTTGCATCTTTCCCTACAGTTT

AAGCTAGTGTATTTAATGTTACAACCTAGTATATATATT

>FK826472.1 JutEST4521 Jute Plasmid library 20-45 days after sowing Corchorus capsularis cDNA similar to Alpha-tubulin, mRNA sequence

GGGAGAGAAGGAAAATTGTGGATTTGTTTGTTTCCCTTTACTATTTCCCACCTTCCTATTTTCCCCATAA

AAAATCTTCTGATTAAAAAGTGAAGTAGCAGGAGGAGGAGAATAAGAGGAAAGAATCCTTCTGTATTTGG

TATTGTGATCAGCCATTGTTTCTTAGAATATTTCTTTTAACACTTTAGACATTGAAAGAGATCATGAGGG

AGATTATAAGCATCCATATAGGCCAAGCTGGAATTCAGGTTGGAAATGCTTGCTGGGAGCTCTACTGTCT

TGAACATGACATTCACCCTGATGGAATGATGCCCGGTGACACTTCTGTGGGTGTTGCACATGATGCATTC

AACACTTTCTTCAGTGAAACTGGTTCTGGAAAGCATGTTCCTAGGGCTGTCTTTGTTGATCTGGAACCAA

CTGTCATTGATGAAGTTAGGACTGGTCCTTACCGACAACTTTTCCACCCTGAACAGCTTATTTCTGGCAA

GGAAGATGCTGCCAACAATTTTGCTAGAGGCCATTACACAATTGGGAAGGAGATTGTAGATCTCTGCCTT

GATCGAGTTAGGAAGTTGGCTGATAACTGCACTGGTTTGCAAGGCTTTTTGGT

>FK826471.1 JutEST4516 Jute Plasmid library 20-45 days after sowing Corchorus capsularis cDNA similar to Purple acid phosphatase-like protein, complete, mRNA sequence

GGGTTGATATTGTATTTGCCGGTCATGTGCATGCTTATGAACGCTCTAAACGCGTCAACAATGGAAAATC

AGATCCTTGTGGTGCTGTCCATATTACTATTGGGGATGGAGGCAACAAAGAAGGCTTAGCTCAAAAGTAC

CGAAACCCACAACCTGAGTGGTCTTTATTCCGCGAAGCAAGCTTTGGTCATGGTGAGTTCAAGATAGTGA

ATTCAACACATGCATTTTGGAGCTGGCACCGGAATGATGATGACGAACCTGTGAAGTCTGATGAGGTCTG

GATAACTTCTCTTGTCAGTTCAGGATGCCTCGCGGAAAAGAACTATGAATCAATGAAAATTCTCATGTCC

CCCTAAAATTACAGAGTTTGATGTACATAAATATTATATATCTTAGTCATAGTTTATGTTTTTAAAATGC

TTGTTTCCATGTAATTTAGTTTAATGAGTGGCTACTAGATTAATACTATCTGAAGTTCTTACTTGCTGTT

TGTTATATGTAAAGGCTTTAGAGTAAATCCAGCTACAGCTTGCAGCTTATCAATTACTAAATTAAATTCC

CCATCCATCAAA

>FK826470.1 JutEST4514 Jute Plasmid library 20-45 days after sowing Corchorus capsularis cDNA similar to Caffeic acid 3-O-methyltransferase, mRNA sequence

GGTGATTGTATCATGTTGGCTCACAACCCTGGTGGGAAAGAAAGGACAGAGAAGGAATTTGAAGCCTTGG

CAAAGGGGGCAGGTTTTCAAGGTTTCCAAGTAAAATGTTGTGCTTTTGGCACTTACATCATGGAGTTCCT

CAAAACTGTTTAAATGATCTCTGGCTAAAAACTTCATGCTTGACCATTTCTTTTGGATTCTGAGCTTTGT

GATTGTTTCAAAGGTGGCTTTTCCGGAAAATATATAGTTCTTTCTTTCTTGTCACAGGAAACAATAAGAA

TATGAAATTTCATGCTCCTTGTGTAAAATAAGTAACCTGTTTTCTATATTGATAATTGCCCTTTCTTAAT

GTAACTATTACAGAATATACATATGTTGATTGGTTTCTCTACAAGGCCTAATGCCTTAATCATGGAAGAA

GCTGATTGGTATAAATACAAATTTCATTTTCTGCAAA

>FK826469.1 JutEST4510 Jute Plasmid library 20-45 days after sowing Corchorus capsularis cDNA similar to Translationally-controlled tumor protein homolog, mRNA sequence

GGGGCTTCAGGAGCAACCTCCTTTTGACAAGAAGCAGTTTGTTTCTTACATCAAGAAGTATATCAAGTTG

TTGACACCCAAACTGGATGCAGATAAGCAGGAAGAGTTTAAGAAGAACATTGAGGGAGCAACCAAGTTTT

TGCTTGGAAAGCTCAAGGACTTGCAATTCTTTGTAGGAGAGAGCATGGCTAATGATAGCACTGTTGTGTT

TGCCTACTACAAGGATGGTGCTACCGACCCGACATTTTTGTACTTTGCTCATGGATTGAAGGAGGTCAAG

TGCTAAGCTTTGAAGCAGAAGTGCTGGCACTATAGTTTGCTTTCTAGTATTGATTATTATTCGCTCTTCT

TCCTTATTTTCTGTACTGCTAGACTTAAGCTACTGTTTTGGTTTTTAATGCTTTCTATTTTATGAGTTGA

CTAAATAACTTATATTGATATGTACTTTTGTCATAATTTAATTGGT

>FK826468.1 JutEST4487 Jute Plasmid library 20-45 days after sowing Corchorus capsularis cDNA similar to GMFP7 (Fragment), complete, mRNA sequence

GGGACAGGATAACAAATAAAGAGAATAGTTTCACATAACGACACAGGCGGAGGGGTTTTCGTCGCTGAAA

GCAGTGGTGTAGCGGACCTCAGTGGAGCTGGCACGTGCGAGATGGGAATAGTGAGGATCATCGGTGTTGC

GGACATAACCGGAGTCAAACAACAAAGAAAAAATACAACCTGTTCTGAAAAACAGGTTAAAGAAAGGAAA

TGGAAGAGTCATCGCTTGATGGAAGAACGGAGGAAGAC

>FK826467.1 JutEST44144 Jute Plasmid library 20-45 days after sowing Corchorus capsularis cDNA similar to Low temperature and salt responsive protein, mRNA sequence

GGCCTCAAGTATGGTTGCAAGGTGGAGTTCTGGATCTGCCTTATACTGACCATCTTTGGGTACATCCCTG

GCATCATTTATGCTGTCTATGCTATCACCAAATAAGATGATGATCATCCTTATAAATATTATATTACTCC

ATTTTAATTTGTGGAGGACACAAATAATATTGCAGGTGTAAGGTGGTGATTATTTTTGGTTCCTTGTTGT

GTTCGCTTGTCCTATCATTCTATTGCTGGCCTCTGTTTTTCTTAGTAATTTTCAGTATAGCTTGTAGTAT

TTGTTCGTTTATTTGATTCCATGTAATTAATTGTCTGTTGGTGTTTGGTTTCATGATTATGATGAACTAC

TTGATCGATGATGATAATTATGGGGTTCGTCTATTTGATTTGATTTGATGTAAGAAAAAAA

>FK826466.1 JutEST44139 Jute Plasmid library 20-45 days after sowing Corchorus capsularis cDNA similar to Gip1-like protein, complete, mRNA sequence

GGAACCAAACCCCTCTTTCCCTTCTTCACTTCACATTTCACTTTCTTCTCTGTTTTCTTTGCTCTTATTT

TGACAGTTTCAAGAGGGGTTTGCTTTGTAGCAAAAGGCCATGGCTAAACTTGTTGCTGCCTTCTTCTTGG

CTCTCACTGCCATCTCCATGCTTCAAACCATGGTCATGGCTTCACATGGACATGGAGGTCACCACTATGA

TCAAAAACATTATGGTCCTGGAAGCCTCAAGAGCTACCAATGCCCATCACAATGCACGAGGAGATGCAGC

AAGACCCAATACCACAAGCCCTGCATGTTCTTCTGTCAGAAATGCTGCAGGACATGCCTGTGTGTGCCCC

CAGGGTATTATGGGAACAAAGCTGTGTGCCCTTGCTACAACAACTGGAAGACCAAGGAAGGAGGACCCAA

GTGCCCTTGAAGACCCCCCTTCCTTTAATCATCCCTATATTTCCTACTTAATCAAGTTCCTTGAATCATA

ATTAAGCATAATGTTCTACTACTACTCCTAACTATATACTACTAGTACTTCATGTTACTTTATGTTTTGG

AAATCAAAATGTGGTGGGATTTTGTAATCACTCTGTCACTCCATAGGGTACCCAA

>FK826465.1 JutEST44134 Jute Plasmid library 20-45 days after sowing Corchorus capsularis cDNA similar to Putative SAH7 protein, mRNA sequence

GGGCTCATCTCATTTCAAGAGAAACCCAGAAAAAGTAGAGGCCAAAAAGATGGCGAAAATCTTGTTGCTC

ATTGCACTATGTGTGCTCCCAGCTCTGGTTAGTGCAACACGCGGCTTGAACAAACCATTGACGGTGCAAG

GATTCGTGCACTGCGACCGTTGCCGTGCTGGTTTCGAAACCCCCAAAACTAGAACCATGGCTGGAGCTAA

GGTAAAGTTGGAATGCCGTGACAGGGTGTCTAGCCACGTTGTGTACAGGAAGGAGGGATACACAGATTCA

ACTGGACAATACAATATTCCTGTGAATGAGGATCATTTGGATCAGGTTTGTGATGCCATTCTTGTGAAGA

GCTCCCAGCCTGACTGTGCCGAAATGTCCCCTGGCCGTGAACGTGCCCGTGTTGTCTTGACCAACTACAA

TGGCCTTGCATCCACCACCCGTTATGCCAATGCCTTGCACTTCTTGGCTGCTCAGCCCGACTCTGGCTGC

ACCGAGATCATGAAGCTTTACCAGGAGGATGAGGAAAATCTGTAGAAATGTG

>FK826464.1 JutEST44132 Jute Plasmid library 20-45 days after sowing Corchorus capsularis cDNA similar to Glycosyl transferase family 2, mRNA sequence

GGTGAATTGATTCCAATCATCTGAGATGATCATAATCTCAGTGCATCAAATTGATGGCACAATTATTTTC

TCCCTATACAAATGATGGCACAATTGGGAATTTATCTTGTCTACATGAGCCTTCCTGACGTATTTATCTA

GTTATAGTTGTATAAATAGGCCTGTTAATATAGGGCAACATTAGCTTAGGTAGAATCAGAAATGCCATTG

CAGTCAACCAGATTACCAGAAGACAAAAGTTTTGTATCTAAATATTTATATTACTGTCATTTTTAACTGT

TATAGAAAGGTATATATATAAGTAAATTTT

>FK826463.1 JutEST44127 Jute Plasmid library 20-45 days after sowing Corchorus capsularis cDNA similar to Actin depolymerizing factor, mRNA sequence

GGAAGAAGCCCTAGATTTTCTCGGCATTTCTCGCTCTTTCTTTCGCTTTCTGGGGAAAAAGAAAAAAATG

GCTAACGCAGCTTCAGGAATGGCTGTGCATGATGATTGCAAGCTGAAGTTTTTAGAACTGAAGGCTAAAA

GGACATACCGCTTCATAGTTTTCAAGATTGAGGAAAAACAAAAGCAAGTAATAGTGGAAAAGCTCGGTGA

ACCTACTCAGAGCTATGAGGATTTCACTGCTAGCCTTCCAGCTGATGAGTGTCGATATGCTGTCTATGAC

TTTGATTTCGTGACCGCAGAGAACTGCCAGAAGAGCAGAATCTTTTTCATTGCCTGGTCTCCTGATACCT

CAAGAGTAAGAAGCAAGATGATCTATGCCAGCTCTAAGGACAGGTTCAAGAGGGAATTGGATGGTATCCA

GGTAGAATTGCAAGCTACTGATCCTACTGAGATGGGTCTCGATGTCATAAGGAGCCGTGCCAATTAATTA

ATTGTATTACCTACTCTGCAATCAAAGTTCATACTATCTGGTTCACTTAAATGTAATGTTTATAACCAAC

TTGGAAACTTAGAACTTGCTTATGATATTTGGGTTGTTGTTCAGGGAGTTCATACTCTTCACTCCTTGTC

TGGATTTGTTATCTGTGGAT

>FK826462.1 JutEST44122 Jute Plasmid library 20-45 days after sowing Corchorus capsularis cDNA similar to Putative calmodulin-binding protein (Fragment), mRNA sequence

GGACCCATCTCGAAGAAATACACGTTCACAGCCTCGACCGGCAATGCAGGTGATTGTAGCAAACCGGAGA

AGAAGGAGAAGAAGAATCATAAAAGTGAAAAGAAAGGAGAGAATGACACGCGTATAGTGGAGATCGAAGA

GCCGTCTGATCATAGGGTTGTCGTTTTGAGACAGGCTTTTGCCAAAAGAGCTGGTGTTATTAGAAATAAG

AGGGGCAAGAAGAAGGAATTGTCTCCTCAAGATGCTGCTGCGATGATCCAAATTACATTCAGAGCTTATC

TGATCCGTAGGTCCCAGGCTCTTCGTGCCCTTAGGGACTTGGCAGTTGCCAAGACTAAGCTGAAAGAGAT

CAGAGCATATTTCAATAACTTTAACTATCGTCGTCGAGTAGCCCAAGATGCAGCAGAACGTCAAAAGTTC

TCAGAGAAAATCATTGTCCTACTCCTCACAGTCGATGCCATTGAGGGAGTTGATGTAATGGTGCGAGCTG

CAAAGAGATCAATGGTTGATGAACTGGAAGCAATGCTCGACGTGGTTGATCCTCAGCCCCTTGGGAAATC

ACTTTCAATGAGGAGAAGGACATTTGACATGCCGGATAGTGTTATCCAGAAGGAAATTGCTGAGGGTGTG

GCGCAGGTTGTTCAATGCTTGAAA

>FK826461.1 JutEST44116 Jute Plasmid library 20-45 days after sowing Corchorus capsularis cDNA similar to 14-3-3-like protein B (VFA-1433B), complete, mRNA sequence

GGGGCAACCAAACCAAATGAGAAGCAAATAAATAAATTCGTCTGCAAAAAAAAGAGAAAAAAGGGGAGAG

AATCTCTCTTTTAGATTTCCAGCAGCAAAATTCTCTCACTATAAATCCCTAAAGGAGAAGTCTGTTTTTG

TCAGAGATCAGCTATGGCTTCTTCTAAAGAACGCGAAACTTTCGTCTACGTTGCCAAACTCGCTGAACAA

GCTGAACGCTACGATGAAATGGTGGAGTCCATGAAGAATGTGGCAAAGCTTGATGTTGAGTTGACTGTGA

GGAGAGGAACTTGTTGTCCGTCGGTTATAAGAATGTGATTGGTGCTCGGAGAGCTTCATGGAGGATTCTT

TCCTCTATAGAGCAAAAGGAAGAAGGTAAAGGAAATGAGGTTAATGCAAAGCGGATCAAGGAGTACAGGC

ACAAAGTTGAATCTGAGCTTTCAAACATATGCAATGATATAATGACTGTGATTGATGAACATCTTATCCC

TTCAGCATCTGCTGGGGAATTGACTGTCTTCTTCTATAAAATGAAAGGAGATTATTATCGATACCTTGCT

GAATTCAAATCTGGCAATGAGAAAAAAGAGGCTGCTGATCATTCTATGAAGGCATATGAGACTGCTACCA

CTACTGCAGAGGCTGAGTTGCCTC

>FK826460.1 JutEST44113 Jute Plasmid library 20-45 days after sowing Corchorus capsularis cDNA similar to ATP binding, mRNA sequence

GGCATCTTCTTCCCCCAAAACAGGGTCAAACCTCCGAGACCAACAAAAAAATGACAAACTACGGCACAAT

TCCGACATCGTCATCGCCGGGACCGTCGGCGAATCTCGAATACTTATCGCGAGCCAAGGCACGAATCAAG

GAAGGCCTGGGAGCTCGACGCCCATGGAAAATCATGTTCAATATCCACTCCCTCAACTGCCCTGCAAGCT

TCTTCGAAGCCATGTCCAGAATCAAGACTAATTTAGCTTACTTCCGCATGAACTACGCCATAATCGTGCT

CATCATCCTTTTCCTCAGTCTTCTATGGCACCCAATCTCCCTCATCGTCTTCATAGTCATGATGGCCGCT

TGGTTGTTCTTGTACTTCCTCCGCGATGAACCCTTGGTTATTTTCAACCGTACGATCGACGATCGTTGGG

TGCTGGCCGTTTTGGGGGTTTTGACCATCGTCTTCTTGCTGTTGACTGGTGCTTTCCTCAATATTCTGGT

TTCCCTCTTGGTTGCCGCCGCTGTCGTTTTGATTCACGCTTCCTTTAGGAGGACTGATGATTTATATGAT

GAAGAATCTGCTGGTTTGATGGCTCCTGCTTCATCTTCTTCTTAATTGTGGGATCTTTAATTTGAGTTAA

TTGGTATTAAGGTGTATGATCTC

>FK826459.1 JutEST44107 Jute Plasmid library 20-45 days after sowing Corchorus capsularis cDNA similar to AXS2 (UDP-D-APIOSE/UDP-D-XYLOSE SYNTHASE 2), mRNA sequence

GGCTGTTGCCTCTTAAGAAGCTGAAGACGTTCATGACTACTCGTAAGAAAGGGATATATCAAGATGTGCA

CCTTTTTAGAAAAGTGTTTCAATTCTTTTTAAGGCAGTCTTAAATATATCTTAATTGTTGATTCCTTATA

CACTAAAATTTTTGTCTTCCCTGTGTCGGTAGTCAATAGCTCGATGTAGCTCAGGATGATGTGTTCTCAG

TATGGATATTATGTGGAAACAAATAGATTCTACTTAAGGGTGGTATTTGTGATAGTATTATAAATATTCC

AATGGAGCTTTTGGGCTACAGCATGTTGGGGGTATATTATATAGAGGAAAGGAAAGCTGTTA

>FK826458.1 JutEST44105 Jute Plasmid library 20-45 days after sowing Corchorus capsularis cDNA similar to Putative prolylcarboxypeptidase, mRNA sequence

GGCCCTATATTACAGTTTGATAAGATAATTCCTTGGTCAAGCTTTTATGATGCTGTTTCTCAGGATTTCA

AGGATGTAAGCTTGAATTGCTTTGAAGTGATAAAGGGGAGCTGGGATGAACTGGATGCAATGTCAACTCA

GAAAGAAGCCTTAGCCGAATTAAGCAAAGCTTTCAGAACCTGCAAGAGCCTCCATTCAACAGCCTCTGCC

CAAAACTGGCTATGGACTGCTTTTGTTTACACAGCCATGGTTAACTACCCAACTGAAGCAAATTTTTTGA

AGCCTTTACCTGCTTATCCTGTGGAGCAGATGTGCAGGATAATTGATAAACTTCCAGCAGGAGCAACAAA

GCTGAGTCGCGCTTTTGCAGCTGCAAGCTTATATTACAACTATTCTCGGGCAGAAAGTTGCTTTGAGATA

GAACATGAAGTTGATGCTCATGGCCTTCATGGTTGGGAATGGCAGTCATGCACAGAGATGGTTATGCCCA

TGACTTGTTCTAAGGAGAGCATGTTCCCTCCATCTGGCTTTGATTATGGAAGAGTTCTCAGAACAATGCC

AGATAAAATATGGAGTTTTGCCTCGGCCACACTGGATTACTACTGAATTTGGTGGCCCAAGTATCCACAA

AGTATTGAAGAGATTTGGTAGC

>FK826457.1 JutEST44104 Jute Plasmid library 20-45 days after sowing Corchorus capsularis cDNA similar to Cyclophilin, mRNA sequence

GGGCATCTCATTTCAAGAGAAACCCAGAAAAAGTAGAGGCCAAAAAGATGGCGAAAATCTTGTTGCTCAT

TGCACTATGTGTGCTCCCAGCTCTGGTTAGTGCAACACGCGGCTTGAACAAACCATTGACGGTGCAAGGA

TTCGTGCACTGCGACCGTTGCCGTGCTGGTTTCGAAACCCCCAAAACTAGAACCATGGCTGGAGCTAAGG

TAAAGTTGGAATGCCGTGACAGGGTGTCTAGCCACGTTGTGTACAGGAAGGAGGGATACACAGATTCAAC

TGGACAATACAATATTCCTGTGAATGAGGATCATTTGGATCAGGTTTGTGATGCCATTCTTGTGAAGAGC

TCCCAGCCTGACTGTGCCGAAATGTCCCCTGGCCGTGAACGTGCCCGTGTTGTCTTGACCAACTACAATG

GCCTTGCATCCACCACCCGTTATGCCAATGCCTTGCACTTCTTGGCTGCTCAGCCCGACTCTGGCTGCAC

CGAGATCATGAAGCTTTACCAGGAGGATGAGGAAAATCTGTAGAAATGTGGTTTTGTGGGAGTATCTTGC

CCAGTCTAATTGATTAGTTGGCGAATGATATGATGTTGTCTGCTACTATTTGTTACGTGAATTGTTTT

>FK826456.1 JutEST44101 Jute Plasmid library 20-45 days after sowing Corchorus capsularis cDNA similar to Caffeic acid 3-O-methyltransferase (S-adenosysl-L-methionine:caffeic acid 3-O-methyltransferase) (COMT), mRNA sequence

GGTGATTGTATCATGTTGGCTCACAACCCTGGTGGGAAAGAAAGGACAGAGAAGGAATTTGAAGCCTTGG

CAAAGGGGGCAGGTTTTCAAGGTTTCCAAGTAAAATGTTGTGCTTTTGGCACTTACATCATGGAGTTCCT

CAAAACTGTTTAAATGATCTCTGGCTAAAAACTTCATGCTTGACCATTTCTTTTGGATTCTGAGCTTTGT

GATTGTTTCAAAGGTGGCTTTTCCGGAAAATATATAGTTCTTTCTTTCTTGTCACAGGAAACAATAAGAA

TATGAAATTTCATGCTCCTTGTGTAAAATAAGTAACCTGTTTTCTATATTGATAATTGCCCTTTCTTAAT

GTAACTATTACAGAATATACATATGTTGATTGGTTTCTCTACAAGGCCTAATGCCTTAATCATGGAAGAA

GCTGATTGGTATAAATACAAATTTCATTTTCTGCAAA

>FK826455.1 JutEST4488 Jute Plasmid library 20-45 days after sowing Corchorus capsularis cDNA similar to Oxygen evolving complex 33 kDa photosystem II protein, complete, mRNA sequence

GGGGTTCATCTTTCTTGGACCCAAAGGGCAGAGGTGGTTCGACCGGTTATGACAATGCAGTTGCCTTGCC

AGCTGGAGGAAGAGGAGATGAGGAAGAACTTGCTAAGGAGAACAACAAGAGTGCTGCTTCTTCATCAGGA

AAAATCACCTTGAGTGTCACCGGGACTAAACCCGACACCGGAGAAGTAATTGGTGTGTTTGAGAGTGTTA

AGCCTTCTGACACTGACTCGGGTGCCAAGACACCAAAGGATGTTAAGATCACTGGTATCTGGTATGCTCA

GCTTGAACAATAGAGAGGGTCTTGTGTCTCAAATGTGTTTATTCTTGCTGCAGATTTTGTAATTTGAAGG

TGTTGCTAAACTTTGCTGTAACTCAATGAGAAACACTGAGATTATATTATCCTTCTCTTTGTATATTATT

ATGTTGCCTTCCTCAAAAAAAA

>FK826454.1 JutEST4486 Jute Plasmid library 20-45 days after sowing Corchorus capsularis cDNA similar to Purple acid phosphatase-like protein, complete, mRNA sequence

GGGTTGATATTGTATTTGCCGGTCATGTGCATGCTTATGAACGCTCTAAACGCGTCAACAATGGAAAATC

AGATCCTTGTGGTGCTGTCCATATTACTATTGGGGATGGAGGCAACAAAGAAGGCTTAGCTCAAAAGTAC

CGAAACCCACAACCTGAGTGGTCTTTATTCCGCGAAGCAAGCTTTGGTCATGGTGAGTTCAAGATAGTGA

ATTCAACACATGCATTTTGGAGCTGGCACCGGAATGATGATGACGAACCTGTGAAGTCTGATGAGGTCTG

GATAACTTCTCTTGTCAGTTCAGGATGCCTCGCGGAAAAGAACTATGAATCAATGAAAATTCTCATGTCC

CCCTAAATTACAGAGTTTTGATGTACATTAAATATTATATATCTTAGTCATAGTTTTATGTTTTTTAAAT

GCTTGTTTCCATGTAATTTAGTTAATGAGTGGCTACTAGATAATACTATCTGAAGTTCTTACTTGCTGTT

TGTTATATGTAAAGGCTTTTAGAGTAAATCCAGCTACAGCTTGCAGCTTATCATTACTAATTTAATTTTC

CC

>FK826453.1 JutEST4483 Jute Plasmid library 20-45 days after sowing Corchorus capsularis cDNA similar to NOI protein, mRNA sequence

GGGATTTTTTTTTCTCTCTGAACCCTCCCCTTCTCTTTCTCTCTCTTCAAACAAAAGTTTCTCTCAAGAA

TTCATGTCGGACAAGGGTCGACCATTGCCAAAATTTGGTGAATGGGATGTCAATGATCCTGCATCAGCTG

AGGGATTTACTGTGATCTTTAACAAGGCAAGGGACGAGAAGAAAACAGGTGGCAAACCTGAGTCACCACC

AAAGGCTGATCATCATGTTAAGCCTGGAGTGGATCCAGGCAAACCCCAGCCTAAAAAATGGCTTTGCTGC

ATTCAAGCCCCACCAGCGGAATCTTGATGAAGTCCTGCCTGTATAAGAGAGCTTCAACAAGTATGTTCAA

AGCCCCACCATATATTCCTAAACAAGATAGGATCTATAGGGCTGTAATAAGCCCCTGTGGAATTTGTTTT

TTTGTGGTGTAAAATTTATTTCTGGTGGGGAGGAAAGAGCAAAGAAGACAAAACCCTTACCTGTATCTAT

ATGTACTGATTTGGTGGGTTATTATTAGAAGCTGAGTTATTCTTCTTGCCTTTGCCTTCTCTATATGTTC

TAAGCTGTAGAGCTTCTGTGTGACATATTCTTTTTTTTTTCATGTTATACGA

>FK826452.1 JutEST4479 Jute Plasmid library 20-45 days after sowing Corchorus capsularis cDNA similar to DNA binding, mRNA sequence

GGGACCATCATCAGTGCTTGAGTTGGATACTTCTGCATCTGTTTCGGTCTCAAAATCAAATGACAATGTT

TACTGTGAAGATGAAGATGCCAACAAAGAAGGGTTTGATGCAAACTTCGCCAATCTCCCAATACCCGATT

TCTGTTGCTTCATTGATGATCCATTGCTGTCTGGTTCAGTTGGACAAGAACTCAACTTAGATGCAGAGCA

CTGCTATTATCTGTTAGTCAATGACTTCGGGTTAGATGATCTGGAGATTATTGGTCATGCATGTGAAGAC

CTCAGCGATTTGCGGGACGAAGGTGACTGAGCCAAGGAGCTTCCTGATTGTGATTTCGGCACTGATTTTG

ACCATTTTGGCACCGGTGATCCTCTTTGGAGAAGAAGTAGCCCCCCTCTTAACATTGCATGCCCCTAAGT

TTTGCAGTTAGGAAATATCATTAGTATTAATATATTAATTTGTTAAGTTATATATTATATATAAGTATAA

GCCTTGTTAGGTGTGAATTATTTAAGGTTTTAATCAATGTTGTTGGAAGGAAGAAGAGATCCCCAGGGGA

CTCTCTAAGATGAGTTCTTGGGATAGTACTTTTTTGTGAGTGGAGTT

>FK826451.1 JutEST4476 Jute Plasmid library 20-45 days after sowing Corchorus capsularis cDNA similar to Superoxide dismutase [Cu-Zn], mRNA sequence

GGGGATTGTCATTATCTCAGTCATTTCTGTTTTATCTTGTACTCTCTACATTCATTCAAGGGGTGCACTG

AGATCACATACGAAAATGGTGAAGGCCGTTGCCGTCCTTAGCAGCAGTGAAGGTGTTAGCGGCACCATCT

TCTTCACTCAAGATGGAGACGCACCAACCACAGTTACTGGAAATGTCTCTGGCCTAAAACCCGGACTTCA

TGGCTTCCATGTCCACGCCCTTGGAG

>FK826450.1 JutEST4458 Jute Plasmid library 20-45 days after sowing Corchorus capsularis cDNA similar to Spermatid-specific protein T2, mRNA sequence

GGATTGTCTTCCATCACAACAAAAAATCGTTCACACAAAAGAAAAAAGAAGAAACAAAAATGCACAACGA

ATGCTGCAACTTCCCATCGCCAAGAGAGAGAGTCAAGGAGAAGTTGAAATCTTCCATCTGCTGCTTCCTA

CCGCCGCATCTCAACGACGGCGCCGATATGGAGCCGTTGGAGGATCAGATGAAGTCAAGGACGCCGAGAT

CCCCTCACGTGATAAAGGACAAGTGCAGACACTTAATCGCAAGAATCGGGAAACGAGGGCGCAGGCAATA

CCACTCGTCGGATTTCAGTTACGATCCAATGAGCTACGCGCTGAATTTCGAGAACGAGTCCAGCCGAGCC

GATGAGCTTCCCATTATGAACTTCATGTCGCGGCTTCCCGCCACGCCGGAGCGAACTCCAGAGCCGATGC

CGGTGTCGGTGAGGAGGGGAAGTGTTTTGCCTATAGTTGACAAATGGCTCGGCTCCATTATTGATTAGGT

TTATTTTGGGGGCTGAGGCTCGATCGGCTCCATTATTAACGGCAACTTGTCCGAGTTGAGTGTTTGGTTA

GGGTGACCAATTGATCCCCTACAACAACTTTTATGTACAGAGATTTAACACTGTAAAGTTTCTGATCACC

ATTCT

>FK826449.1 JutEST4453 Jute Plasmid library 20-45 days after sowing Corchorus capsularis cDNA similar to Acyl-[acyl-carrier protein] desaturase, mRNA sequence

GGATTCCAAGAAAGGGCAACTTTTATCTCCCATGGTAACACTGCCAGGCTTGCCAAGGAGCATGGGGACA

TTAACTTGGCTCAGATATGTGGAATAATTGCCTCAGACGAAAAACGCCATGAGACAGCCTATACTAAAAT

TGTGGAGAAGCTCTTTGAGATTGATCCTGATGGTACTGTCCTGGCCTTTGCTGACCTGATGAAGAAGAAA

ATCTCCATGCCAGCCCACTTGATGTATGATGGCAAGGATGACAACCTTTTCGACCACTTCTCAGCTGTTG

CACAAAGGCTTGGGGTGTACACTGCCAAGGACTATGCTGATATCCTAGAGTTCCTTGTGGAAAGATGGAA

GGTGAAGGACCTAACTGGACTTTCTGGAGATGGGCGTAAAGCTCAGGATTATGTGTGTGGACTTCCTGCA

AGAATTAGGAGGCTGGAGGAGAGAGCTCAAGGAAGGGCCAAGCAAGGACCCAGTATTCCTTTCAGTTGGA

TTTATGACAGAGAAGTGATGCTGTAGGTGTTTAAAGTAAAGAAGTGCTATCAATTTTCGTCTCTTGCAAG

CGTCTTATTTGATTCAAGGAAAAACAGAGATATAGAGAAAC

>FK826448.1 JutEST4452 Jute Plasmid library 20-45 days after sowing Corchorus capsularis cDNA similar to Dynein light chain, mRNA sequence

TTTCCCAAAAATGAGCACTGAAGAAGCAAAGAGAAGCCTCAACGGAGCTCTGACGGTGAAACCCAATTCA

GATGACCGGAAGCCTTCTATGGCGCCAATCGCGGCGTCGGCTGCTCCAACCGGCAAGAAAGTCATCATCA

AAAGCGCCGACATGAAAGATGACATGCAAAAAGAAGCTATCGATATAGCCATCTCTGCATTTGAGAAGAA

CAATGTAGAGAAAGATGTAGCAGAGCACATAAAGAAAGAATTCGATAAGAGACATGGACCCACTTGGCAT

TGCATCGTTGGTCGCAATTTTGGATCATATGTAACTCATGAGACAAACCACTTTGTTTACTTCTATTTAG

ATCAGAAAGCAGTTTTGCTCTTCAAATCTGGTTGATCATGTTGTTATGTTGATGAAGACTGCTGATCATG

AGGATTATGGAAGGCTTTTCACTTTTATTAATTATGCTACTATAAATAGCAACTGTATGTATTATCAATT

GAAAAACCTGAGTACGTATGTGGAATTCTCTCCTAAATTTCTTGTTT

>FK826447.1 JutEST4450 Jute Plasmid library 20-45 days after sowing Corchorus capsularis cDNA similar to NADH-ubiquinone oxidoreductase chain 1, mRNA sequence

GGGCAAGGCGGTGCATTCTTGTCCAACATTTTGGAACAAAACAGTTTCTCATAAGGAATTGTTGAGGATC

TTGAGTGACAAGGCAGGGTTGAAATGGTCTAAACGACCACCTGCTCTGTACTCTTTGTGGTCTCCAGGCT

TTGTATTGTTCTGTTCCTGCATATGGTTGCGTTGGTCCCGCGAGTGGCCACATGAGTAACTCAAGACTGA

TGGGACACCAGGGTAGGGTTTATGGTTCCAATTTTATATTGATGATGATGCACTTGTTAATATTTAAGTT

CAAACATAAAAGTTGAGTTATTTATTGTGTTTTTTATTTAGCCTTTTGTTTGTGTTCTAAATACATCGTG

TCCACACACATATGTCATGAGTTGCGCAAAGGAGCTTATGACCAAGGCACAC

>FK826446.1 JutEST4447 Jute Plasmid library 20-45 days after sowing Corchorus capsularis cDNA similar to specific tissue protein 2, mRNA sequence

GCATTGAAAATAAGGAAAAGAAAGGTTTTGTTAAAGATTTTGAGCCACGGCCAACAATTAAACATATGGA

GAGGAGGGGATAGACGTTAAGGAACAGTTCATTTGCGTCAAAGATTTCGAACCCAGGCCCAACCTTTCGG

CTTATGGTGATGATGTCGTTGATCTTAAAGAAAAGAAGAAATTTGCTAAAGATTTCGAACCGAGGCCCAA

CCTTTCGGCTTATGGTGGTGATGACAGTGATCTTGAAGAAGAGAAGCAAATCATTAAAGATTTCGAACCC

AGGCCTAACCTTTCGGCTTATGGTGATGATGGTATCTAAAAGAGGGGAAAAAATGCATTTACTAGTGATT

TGGAACCGAGGCCCAATACCTCTCCTTCTCTACACTACATCATAGATTTTAAGGAAAAGGAGAAACTAAA

CCAAATAAATAAAATGAAACTATTTGGTTTAATTTCTCTTGATTAGGAACTAATATGGCATCGTGGTCTG

TGTTGTATGGTTTGTCTTAATTATGATTTGGCTTTTCCTACAAATAATGAAGTCATTACTCATTACTATT

GGAAAAA

>FK826445.1 JutEST4441 Jute Plasmid library 20-45 days after sowing Corchorus capsularis cDNA similar to NTCP23-like cysteine proteinase, mRNA sequence

GGTGTGGCAACTTGTGCATCATACCCTGTTGTTGCTTGACCCAGTCGATGAACCGTGTTGGCTCATGTCA

CGGCTTTGTGTTATACGAGGCGTCTAGTTATGGTCTACAAATGTTGTAGTTGTCGATTTCAGAGGTGAGG

GAGGGAGGGGTCATGTTGTAGTTAAAAAGGGTTATGGTCTAGTTGAAACCCATGGGTTCATGTACAGGCA

GGATCCAAATACATGTAAGATGCAAAATTTGCTGTACACAACAAAGGTGTAAATTTTTATGTTGTAATGC

TTGTTGTGTAATTACTTGTCAACATTGGAAACCAATAGGGGTTTTAATAACAATACAAACGAACCTTTTC

GTGT

>FK826444.1 JutEST4440 Jute Plasmid library 20-45 days after sowing Corchorus capsularis cDNA similar to Adenine phosphoribosyltransferase -like protein, mRNA sequence

GGGGAAGCAAATATAATGAATGAAACTTTTTGATCTTGATACGACTTTTCTACCCAATTCAAACTGGGTA

TTTGACTATTTGTCATTTAAACTAAAGTTGCTTGATTTATCTAAATTAGTTTAGTTCCATGTTATGTTCT

TTTTTTTTTTTTCCAATGCAAGATTGTAAATTTGGTTTTGTTATGGTGAAAATGGCAGGTATTATGTTTC

AAGACATTACTACTCTGTTACTGGATCCTAAAGCATTCAAGGACACAGTGGATTTGTTCGTTGAGAGATA

CAAAGGGAAGAACATTTCAGTTGTTGCAGGTAAAGATTATCTCTCATTGCAATTCATAATGTTAATTAAT

TTTCTATACATAGTGTTATTT

>FK826443.1 JutEST4438 Jute Plasmid library 20-45 days after sowing Corchorus capsularis cDNA similar to drought-induced protein SDi-6 - common sunflower, mRNA sequence

GGGATTGAAACATTTAACAATGGCGTCAGAGAAAGACCACAAGCCAACTCGTCCGGGGACCTTTTTTCCA

GCAGCAAGCTGGTTGCGGAGGCCGCAAAATCAGCATTCGGGCAGAAGAGCGGCGACAAGTTCGATAAGGC

GAAGGTTGCCGAAGCCGCTTCTGATCTTTTAGATGCAGCCAAAGACTATGGCAAGCTGGAGGATAAGGGA

ATTGGTCAGTATGTTGACAAGGCACAGAGTTACCTTCACCAATTCAGCTTCCCAATCAACCACTAAGAGA

TACCCGACGCACCGGCTGCCGGCTAAAGATGGTGAGGCAAAGCCAGAGTCGCATCCAAGCGGTGGCCATG

GAGGCAAGGAGTCTGGAAGCGGTGGCG

>FK826442.1 JutEST4437 Jute Plasmid library 20-45 days after sowing Corchorus capsularis cDNA similar to RNA polymerase transcriptional regulation mediator~gene, mRNA sequence

GGGGAATTAAGCGAGAGCGCTGTCTTTCACTGAAGAAGACTCTGAACCCCAAATCATGGCGACACCACCG

GTGCCGCCACCACAGGCGGCGGCATTAGGGAACTTTGAAGCTCCGCCACCGCCGGCGATGCAGCCTCCTG

GAACGGACATGACAGGGATATGTTTCAGAGACCAATTGTGGCTAAATACTTACCCTCTTGATCGGAATTT

GATCTTCGATTACTTTGCTCTGTCTCCGTTCTATGACTGGACCTGCAACAACGAACAGCTCAGAATGCGC

TCTATTCACCCTCTCGACATCTCTCAGCTCTCGAAAATGGCTGGCTTGGAGTACATGCTTAGTGAGGTCA

TGGAGCCACACCTATTTGTTATCCGTAAGCAGAAGAGAGATAGTCCAGAGAAAGTCACACCAATGCTGGC

ATATTATATTTTGGATGGTTCCATTTACCAAGCACCACAGCTTTGCAATGTCTTTGCAGCTCGAGTTGGA

CGTGCTCTCTACTATATATCAAAAGCTTTTACTACTGCTGCTTCAAAATTGGAGAAGATTGGTTATGTTG

ATACTGAAAATGAGGAATGAAATTTCTGAATCAAAGGGTGGTAAAGAGGCAATTGATTTC

>FK826407.1 JEST42121 Jute Plasmid library 20-45 days after sowing Corchorus capsularis cDNA similar to Drm3, mRNA sequence

GGGAGAAAACGAAAAAAGGATTCTTTATTATCGCCGTGGATCCGTTAAACTACGCCGATCTCCTGCTTAG

ATACGCCTTGATCTGTTATTATGGGGCTACTTGACCAGTTGTGGGACGACACCGTTGCGGGTCCCCGACC

CGATAACGGCCTCTGCAAGCTTCTAAAGCACTCCACTTTCACTTTCCGGCCTATCTCCGCCAAGGAATCT

GATGGCGGGACTGTGAGATCGTACGGTGATGAAAC

>FK826406.1 JEST42114 Jute Plasmid library 20-45 days after sowing Corchorus capsularis cDNA similar to Uridylate kinase (UK) (Uridine monophosphate kinase) (UMP kinase) (UMP/CMP kinase), mRNA sequence

GGGGGGAAACTTACAAGGATTCCCTTAGTAACGGCGAGCGAACCGGGAGGAGCCCAGCTTGAGAATCGGT

CGGCCCCGGCGTCCGAATTGTAGTCTGGAGAAGCGTCCTCAGCGGCGGACCGGGCCCAAGTCCCCTGGAA

GGGGGCGCCGGAGAGGGTGAGAGCCCCGTCGTGCCCGGACCCTGTCGCACCACGAGGCGCTGTCTACGAG

TCGGGTTGTTTGGGAATGCAGCCCCAATCGGGCGGTAAATTCCGTCCAAGGCTAAATACGGGCGAGAGAC

CGATAGCAAACAAGTACCGCGAGGG

>FK826405.1 JEST4270 Jute Plasmid library 20-45 days after sowing Corchorus capsularis cDNA similar to Ran-binding protein 1 homolog a (Ran-binding protein siRanBP), mRNA sequence

GGGTCGAATATGACAGTGCAGGAGCACGCAGGGAACGACAAATCGTGCCTATGGCACGCTGCTGACTTTG

CAGATGGTGGATTGAAAGATGAACTTTTCTGCATCCGCTTTGCATCAGTGGAAAATTGCAAAACTTTTAT

GGAAATGTTCCAAGAAGTTGCTGAATCACAAAAACCAAAGGAGGAAGATAAAGATGCATCTGCAGCAGCT

GGATTACTTGAGAAGTTGAGTGTTGAGGAAAAGAAAACAGATGATAAAGCTGAAGAGGAGAAGCCTGTTG

CGGCAAAGGAAGAAAGGGAAACAAAGGATGATACAGAAAAAGCAGATGCTGAGAAGAAAGATGGGGAGGC

AGCTTCCTCAACCTAAGTTGGTTTTGTTTTCCTTGCCATGTGGTTTTGGCGAATATGGGATGATTAGTCA

CCCAACCTTACCTCCCACCCATGTCTTGGCATATTCCTGCAGATGACACTAGATGGGTCGATTTATTGGA

TGGAATGGTTGAGGTCCTTGGTGTCCTTTAGGTTTGAGTGCCTGTTGAGGTTTGTCAACCCTCGGTCGTG

TCAGTCATGTTTGCATGTTGAGAGTTGTCTTTCGCGGTT

>FK826404.1 JEST420045 Jute Plasmid library 20-45 days after sowing Corchorus capsularis cDNA similar to Auxin-repressed protein-like protein ARP1, mRNA sequence

GGGATTACTTGAAAAGAGAAACCGGAGAGGAAAAGAGAAAGAAAAAGAAAAGAAAAATGGTTCTCCTTGA

TAAGCTTTGGGATGACGTAGTAGCCGGGCCTCAACCGGAACGTGGCCTCGGCAGTCTTAGGAAGCTGGTC

ACCGCCCAGCCCTTGACCATCAAAGATGAAGGAGAAGGGAGCAGGTTCCAGAGATCGTTGTCGATGCCGT

CGAGTCCAGGGACACCGACGACTCCGATAACACCAACGACGCCGGTGTCGGCGCGTAGGGATAACGTGTG

GCGGAGCGTGTTCCATCCGGGTAGCAACCTTGCCACCAAGGGAATCGGCGCTGCGGTTTTCGACAAGCCT

TCACAAGCCAACTCTCCTACTGTCTATGACTGGCTTTACAGTGGGGAGACAAGGAGCAAGCATCATCGCT

GATTGGGTGGTGGTGACTGTGACTGACAGACGGGGGAAGCTTAGCAGTATAAGCAGGGGGGATTTGGTTG

AAGCATGTAAATAGGCGTCGTTCTTTCAATCTTCGATGTGCCAACGTTGTCCTTGTTTTGATGGGTATTT

GTGATTAAGTGTCCTAAGTAGTAGAACGTTTTAAGACTTGACTAAGTTCTTATTAAGCCCTTT

>FK826403.1 JEST4223 Jute Plasmid library 20-45 days after sowing Corchorus capsularis cDNA similar to Germin-like protein subfamily 3 member 2 precursor, mRNA sequence

GGGATTTCGTCTTCTCAGGGTTAAAAGCAGCCGGAAACTTCAGTGAAACAGGCCTTGCTACCATTCCGGT

GAATCCATCAGTCTTTCCAGGGATTAACACACTAGGCATGTCATTTGTACGTGCTGACCTCAAAGTTGGC

GGCATAGATCCACCTCATTATCATCCGAGAGCAACTGAAATCGCCTATGTGGTGCAAGGAAGTGTTTATT

CAGGCTTTGTTGATTCAAGCAACCGCGTTTTCGCCAAGGTGATTGAGCAGGGAGAGGTGATGGTGTTCCC

AAGGGGGCTGGTGCACTTCATGATGAATGTTGGTGAGAAGCCTGCTACAATATATGCAAATCTTAACAGC

CAAAATCCAGGACTTCAAAAGATCCCAACTGCTATTTTTGGATCTGGAATCAATGAGGAGCTGTTGGAGA

AGGCTTTTGGCCTGAGTTCTAAGCAGATTGGGACAATGAGGAGAAGATTTGATCCCAAAACAGTGAGGTA

AAAGTTCATGAAAAACCAAAAAGATCGTCAGACAATATATATGAATAGTAA

>FK826402.1 JEST427 Jute Plasmid library 20-45 days after sowing Corchorus capsularis cDNA similar to Rac GTPase activating protein, mRNA sequence

GGTGAAGAAAGAGATTGAAGTTCTTGACCATGGGAACAGAAAGAAGTCTGATAATAAGTGCTCCGTTCCG

GGGCATAAGCTGGAATTAGTCCAGAAGGCAGAGCCATTAATTCTGGAAGAGCCCTTAAAGCCATGGATTA

CCTTGAAAAGGAATATACAAAACGTGAAGTATTTGTCATCTATGGTTGATATAAGTTTCAAGAATAGGTA

TGTTTATGTTGATGTTGGGGCTAGGAGTTATGGTTCTAGTATTGGGAGCTGGTTTAGGAAACAATAT

>FK826401.1 JEST423 Jute Plasmid library 20-45 days after sowing Corchorus capsularis cDNA similar to Always early protein 3, mRNA sequence

GGCTCCTGCAGTTAAATGAAGTCAATGAGCAGGTTTCTTCTGCTCTGTTTTGCTTGAGGCAACGCAACAC

ATATCAAGGGACCTCCTCAGTTAGGTTGCTGAAGCCCATGGTTGGTGGGCATGGTTGTCAGTTGACCTCT

TTTGATCATTCTATGCATAATCTCCAAGAATCTGTATCAAATGTGGCTGAAATTGTAGAAGGTTCAAGGA

TGAAAGCTCGGTCAATGGTAGATGCAGCTATGCAGGCTGTATCATCATTGAGAAAAGGCGTTGAGAGGAT

TGAAGACGCAATAGATGTCGTAAATAACCAACTTTCAGTGGATGATTTAAGTCTGCCTGCTCCAAGGTCG

TCGTCTATTCCAATAGACTCAATCCGCAGTACGGTAACACTTAACGATCATCCATCAGCAACTAGTCATT

CGCCTGATATTAAGTTACAAAATTTGTCTGACCAAGATGATCCTAAAATCCCGTCAGACCTTATTGCGCA

CTGTGTAGCCACCTTGCTCATGATACAGAAATGTACGGAAAGACAGTTTCCGCCCGGAGAAGTTGCCCAG

GTACTAGATTCCGCCGTTACCAGTTTGAAGCCATATTGTTCACAGAATCTTCCGATTTACGCCG

>FK826400.1 JEST4142 Jute Plasmid library 20-45 days after sowing Corchorus capsularis cDNA similar to Glutathione S-transferase GST 22, mRNA sequence

GGAGTGAACAACAAAAGAAAAGTCCTTAGGAAGACCAAGAGGGAATTGAGTACTGAGTAGTAGCTTTTAC

AGTCAACAACAATGGTGGTGAAGGTTTATGGTCCAGCCTATGCTTCCCCAAAGCGTGTGATCCTGTGTCT

TATTGAGAAAGGGGTTGAATTTGAGACTGTCAATGTTGATCTACTCAAAGGAGAAAACAGAGATCCTGAA

TTCCTTAAGATGCAGCCTTTTGGAAGTCTTCCTGTCACTCAAGATGGAGACTTGACTTTATTTGAATCCC

GAGCCACCATGAGGTATTTTGCAGAGAAATACAAGTCACAGGGAACTGATCTGCTAGGGAAGACATTGGA

GGAAAAGGCTCTAGTTGATAATTGGGA

>FK826399.1 JEST4138 Jute Plasmid library 20-45 days after sowing Corchorus capsularis cDNA similar to RNA-binding glycine-rich protein-1, mRNA sequence

GGGATTCGCAGTTCTATTCTCAAAAACAACCTCCTTCATTTCTTAAACCTAGTTTTTTTCTTGTTTTTCT

TCATGGCTTCTGCAGATGTTGAGTTCCGGTGCTTCGTCGGAGGCCTTGCCTGGGCCACCGACGACCGCGC

CCTCGAAAGTGCTTTCAGTCAATACGGCGAGATCGTCGAATCCAAGATCATCAATGATCGTGAGACTGGT

AGATCCAGGGGATTTGGCTTCGTTACCTTCCGTGATGAGAAATCTATGAGGGATGCTATTGAAGGAATGA

ACGGTCAGAATCTCGATGGAAGGAACATTACTGTCAACGAGGCTCAATCTCGCAGGAGCGGCGGCGGCGG

CGGCGGTTTCGGAGGAGGAAACGGTGGTGGCTACAGCCGCGGTGGAGGTGGCGGAGGCTA

>FK826398.1 JutEST416 Jute Plasmid library 20-45 days after sowing Corchorus capsularis cDNA similar to Induced stolon tip protein, mRNA sequence

GGAAATCGATCAAGAAATGATCAAGTGGCCGCTGTGGACAAAAAGGTGGTAAACCGATGTTCCGGTTGCA

GAAAACGTGTGGGGTTGACCGGGTTCCGATGTCGTTGTGGCGAACTTTTTTGTGCGGAGCACCGGTATTC

GAATCGACACGATTGCAGTTATGATTACAAGGCAGCGGGTCGCGAGGCCATCGCTAGGGAAAATCCCGTG

GTTAAAGCAGCTAAGATTATTAGAGTTTGATGTTGCCAAATTAAACTGAATCATATATTGCAAAATTATT

TTTCGACCTTAAAGTTGAGTTAGTCGAGGAGATCCATGCTCCATCTCATCATCTCGTACAAAAAGATGAC

GGCATCTCAGACAAAAGATGGTTTGACAGAGAGAAAAGAGGAGATGAACAACATAGTCCCTCCTTAAGAC

TGATTGTTCATGTCCGATCAATCAATTTATAGTATTTCATTTTTCATTTTACTTGAGTTTTCTTTTCTTT

TTTT

>FK826441.1 JutEST4432 Jute Plasmid library 20-45 days after sowing Corchorus capsularis cDNA similar to Beta-glucosidase precursor, mRNA sequence

GGGAAATGTAGCTATCGGATTTTATCATAAATACAAGGAGGATATAGTATTGATGAAGAAGATTGGGTTG

GATTCCTTCAGATTCTCCATTTCATGGTCTCGAGTTTTACCAAAGGGGAGAGTTAGTGAAGGAGTAAACG

ACCTAGGAGTCAAATTCTACAACGACCTCATCGATGAACTCATTAAGTAATGACATCAAGCCTTTTGTGA

CTTTGCTTCATTTTGATCATCCACAAGCTATTGAAGAAGAATACGGTGCATTCGGAAACTCGCAAGTTTG

TAGATGATTTTGCTGAATATGCGGAATTCTGCTTCAAAACATTTGGTGACCGAGTCAAGTACTGGTGCAC

CTTGAATGAACCAAATGGGTGGATCGTATCCATCATCCAAGCAGGTGGCTATGGATGGGTTGAACAAGGC

GGTAATGCCACTGAGTCGGGGCCCTTTCCCTTGGGCCATCATTTGATTCTTGCCCATGCAGCTGCTGCCA

GAATCTACATGGACAAGTACCAGCCAACTCAAAAGGGAAAGATTGGAATTACCATAAATACCACTTGGTA

CAAACCACTGAATGC

>FK826440.1 JutEST4426 Jute Plasmid library 20-45 days after sowing Corchorus capsularis cDNA similar to Chalcone synthase 2, mRNA sequence

GGGACTTAGTTATTAATTAAATTGAGTATCAAATTAATAAGCTTATAATAACTTATATTACTTTTACTAA

AAATTATCTTTGTATTCAATTGAACTTAAACTTTGTATTTCATATAGTATCCAATGAAAGTTCATTTCCC

CTTTGATGATCTCTTGAC

>FK826439.1 JutEST4420 Jute Plasmid library 20-45 days after sowing Corchorus capsularis cDNA similar to F6F9.8 protein, mRNA sequence

GGGGAAAAACTTGATGATGCTAAAGTTGAGTCTGCACATTCAGTTATTGTGGACTCCTCAGAAGTAGAGC

AGAAGCCACTGAAAAACACATCTGATACGCAGAGAGAACAGGACTCTGTGACTGTTAATCTCTCATACAG

GTCATCCCCAGAAGCCTCTCCAAGGAGCCATTTGACTGTTCCAGAATCCCAGGGAACACCTTCTAGTCAG

GTATCAGTGAAACC

>FK826438.1 JutEST4414 Jute Plasmid library 20-45 days after sowing Corchorus capsularis cDNA similar to FH protein interacting protein FIP1, mRNA sequence

GGCTACCAGCGCTGCTACCACCATGCCGGTTGAGTCCAATCCTTACGTTTCTCCCGCTCCCGCTCCATCT

TCTACTGTCAAGAATACGATGGAAACCGTTAAAGATGTGTTAGGGAAATGGGGTAAGAAGGCAGCAGAGG

ATACTAAGAAGGCGAAAGAGATTGCTGGGAACATGTGGCAACACTTGAAAACTGGGCCAAGTCTTGCTGA

TGTTGCTGTAGGAAGAATTGCACAGGGTACTAAAGTTCTTGCAGAAGGTGGTTATGAGAAGATCTTTCGA

GGAACATTTGACACTGTTCCTGAGGAAAAGCTTCTGAAAACATATGCGTGCTACCTGTCCACCTCGGCTG

GTCCTGTTATGGGAGTGATGTATCTATCAACTGCAAAGCTCGCTTTTTGTAGTGACAATCCACTGTCTTA

CCAAGTTGGTGGTCAGACTCAGTGGAGCTATTATAAGGTGGTTCTACCATTACATCAGCTCAGGGCGGTG

AATCAATCGGCTAGCAAAGCCAACCCAGCTGAGAAATACATCCAGGTTATATCAGTTGACAACCATGAGT

TTTGGTTCATGGGTTTTGTACACTATGATAGTGCTGTTTAAAACCTTCAAGGAGCATTG

>FK826437.1 JutEST4410 Jute Plasmid library 20-45 days after sowing Corchorus capsularis cDNA similar to unspecific monooxygenase, mRNA sequence

GGGGCGATACTGACGTGCAAATCGTTCGTCTGACTTGGGTATAGGGGCGAAAGACTAATCGAACCGTCTA

GTAGCTGGTTCCCTCCGAAGTTTCCCTCAGGATAGCTGGAGCCCTTAGCGAGTTCTATCGGGTAAAGCCA

ATGATTAGAGGCATCGGGGGCGCAACGCCCTCGACCTATTCTCAAACTTTAAATAGGTAGGACGGCGCGG

CTGCTTTGTTGAGCCGTGCCAAGGAATCGAGAGCTCCAAGTGGGCCATTTTTGGTAAGCAGAACTGGCGA

TGCGGGATGAACCGGAAGCCGGGTTACGGTGCCCAACTGCGCGCTAACCTAGAACCACAAAGGGTGTTGG

TCGATTAAGACAGCAGGACGGTGGTCATGGAAGTCGAAATCCGCTAAGGAGTGAATGTTATAGGATTTGT

ATACTCTGGGATTCAAGCATATGATCTAGCTTACCAATTGACCTCCGGGGAAACAGAAACCCCGGGACGC

GCCTCCGGTGTTACTTAGATTTCGTTTTGGGATCAGGTTCTAGCTTATCTTCTTATATCAGCATCATCGT

CAGCTGCTGTCCGAGTTGATGACTGGCAATCAACTGGGGGAAAGACAAGTTCCCGGAGATGGCA

>FK826436.1 JutEST41193 Jute Plasmid library 20-45 days after sowing Corchorus capsularis cDNA similar to Thiazole biosynthetic enzyme, mRNA sequence

TCATCATCCATGGCTGTAACTCTCACTTCCTCAACCAAACTCTACAAGAACCCTTCTGTTTTTGAGTCTT

CCTTCCATGGGGTTCCCATAACTCCACTTTCTTTCAATTTCAAAACAAAATCCTCTCCATGCAATGCATC

GATTTCCATGTCTGCTGCCTCATCATCACCTCCACCCTATGATCTGAATAATTTCAGGTTTGACCCCATC

AAGGAGTCCATCGTGTCTCGTGAGATGACACGTAGGTACATGATGGACATGATCACTTATGCTGACACTG

ATGTGGTCGTGGTTGGTGCTGGCTCCGCCGGTCTTTCTTGTGCCTATGAACTCAGCAAGAACCCTTCTGT

CCAGGTTGCCATTATTGAACAATCTGTTAGCCCTGGAGGTGGTGCCTGGCTTGGTGGACAACTTTTCTGT

GCCATGGTAGTTCGCAAACCAGCTCATCGCTTCCTTGATGAACTTGGTATTGAGTATGATGAACAAGATG

ACTACGTAGTCATCA

>FK826435.1 JutEST41109 Jute Plasmid library 20-45 days after sowing Corchorus capsularis cDNA similar to Proline-rich protein APG-like; GDSL-motif lipase/hydrolase-like protein, mRNA sequence

GTACTAAGCTCTGTCACTACTCCTTTTTCTATTATCTTAATCAAAAAGTGACTCGTATTGTAGAAGCTTG

TATTTGTAGTCAAACCTTCCTTAAAACTCATCGTATATTTCTCATCAGATTATTTACAATTCCTAAGTGA

CATATAGGTTAAAGAAATTGTGATACAGCTACATCTTTAAGTTATAAATTGAGGATAATGAAAAT

>FK826434.1 JutEST43143 Jute Plasmid library 20-45 days after sowing Corchorus capsularis cDNA similar to 14-3-3-like protein D, mRNA sequence

GGGGCAACCAAACCAAATGAGAAGCAAATAAATAAATTCGTCTGCAAAAAAAAGAGAAAAAAGGGGAGAG

AATCTCTCTTTTAGATTTCCAGCAGCAAAATTCTCTCACTATAAATCCCTAAAGGAGAAGTCTGTTTTTG

TCAGAGATCAGCTATGGCTTCTTCTAAAGAACGCGAAACTTTCGTCTACGTTGCCAAACTCGCTGAACAA

GCTGAACGCTACGATGAAATGGTGGAGTCCATGAAGAATGTGGCAAAGCTTGATGTTGAGTTGACTGTGA

GGAGAGGAACTTGTTGTCCGTCGGTTATAAGAATGTGATTGGTGCTCGGAGAGCTTCATGGAGGATTCTT

TCCTCTATAGAGCAAAAGGAAGAAGGTAAAGGAAATGAGGTTAATGCAAAGCGGATCAAGGAGTACAGGC

ACAAAGTTGAATCTGAGCTTTCAAACATATGCAATGATATAATGACTGTGATTGATGAACATCTTATCCC

TTCAGCATCTGCTGGGGAATTGACTGTCTTCTTCTATAAAATGAAAGGAGATTATTATCGATACCTTGCT

GAATTCAAATCTGGCAATGAGAAAAAAGAGGCTGCTGATCATTCTATGAAGGCATATGAGACTGCTACCA

CT

>FK826433.1 JutEST43139 Jute Plasmid library 20-45 days after sowing Corchorus capsularis cDNA similar to WRKY transcription factor, mRNA sequence

GGAATTCAACTTCAACCGTTTCTCCTCCTGTTAGCCAAAACCAAGCAAATCAAGAGCTTGAAACTAAGGT

TTACTATGCAACGCCGATCCAGCAGATTCCACCGCCGGTTTCTTACCATCCTCCTCCTCCTGCTCATCAT

CATCATCACCATCATGATTTTTTGAGTGTGAAAAGTGGGGGTTTAGAGAGGAAAGATTCATCAACAACCA

TCAATTTTTCGTATTCTTCAGCTGGTGGGAATTCGTTTATGTCATCTTTAACAGGGGATACTGACAGCAA

GCAGCCATCTTCGTCATCTGCTTTCCAAATTACTAATCTTTCTCAGGTTTCCTCCGCCGGGAAGCCTCCT

CTGTCTTCTTCTTCCTTTAAAAGGAAGTGTAGCTCTGAAACCTTGGCTTCCGGCAAGTGCAGTGGCTCTT

CTGCTGGTCGCTGCCATTGCTCTAAGAAAAGAAAACTGAGATCAAAAAGAGTTGTAAGGGTTCCAGCAAT

AAGCCTGAAAATGGCTGATATTCCACCAGATGATTATTCTTGGAGAAAGTACGGTCAAAAACCAATCAAG

GGTTCTCCACATCCAAGGGGGATATTATAAGTGCAGTAGCGTGAGAGGATGTCCAGCTCG

>FK826432.1 JutEST43138 Jute Plasmid library 20-45 days after sowing Corchorus capsularis cDNA similar to transcription factor, mRNA sequence

GGGGATCCCCATGGAGGAAGACGAAGAGATCCAATCGCACCGGTCGCCTGGCGCTACCGGATCTCAATCT

CCGTCGTCACCGTCACCGACGTCCAACGGTCGGATAACTGTAACGGTAGCCGCGGTGCCTCCTTGCGAAC

CACCTCTTTCAGAAGAGCCTCAGCAACATGAAAATAAGCTTGCTTTGGCGCTTGTGCCGCCGCAGCCGCA

ACCTTTGCAGACGAAGAACAACGGCGGCGGAGGTGGAAGAGAAGACTGCTGGAGCGAAGGCGCGACGGGG

GTTTTGATCGACGCGTGGGGGGAGAGGTACTTGGAGCTGAGCCGTGGGAACTTAAAGCAGAAGCATTGGA

AAGAAGTGGCGGATATCGTGAGTAGCAGAGAGGACTATATGAAGACTCCCAAGACGGATATTCAGTGTAA

GAATCGAATCGATACGGTGAAGAAGAAGTATAAGTTAGAGAAAACTAAGATGGCGAGTGGGAGGTGGGCC

GAGTAAGTGGGTCTTTTTCGAGAAATTGGATCGACTTATAGGACCCACGGCCAAGGTTTCCGTCAATGCT

GCGGCCGTGGGTGGGTGGGTGGGGAGTCCTGGGGTTTATGTCTA

>FK826431.1 JutEST43137 Jute Plasmid library 20-45 days after sowing Corchorus capsularis cDNA similar to DNA-binding protein 4, mRNA sequence

GGATCCTAAAACTAAAAAAAAAAAAAAAACCAAAACCAAAACGGAAAAAAAATCCGGCAGGGTTGGTCGG

AAAAAGAAAACATGTCGTGGGACAAAAAAAAGAGCAATCGAAGAACTTGCTAAAGGCAAGGACTTAACAA

ATCAGCTGCGAGATCTCCTCAATGGTAGTAATTATTTGGTTGGTAGCGAAGATCTAGTGATGAAAATCTT

CAACTCTTTTTCAAATACACTTTCGATTTTAAGGAACAGTAGTAGCGGAGACTATGACGAAGTTTCTCAG

TTGAATCCGAGGAACAACAACAGTTATGATGGTCGGAAAGATCGAAGAGGATGTTACAAAAGAAGGAAGT

GTGGAGACTCATGGACAAGAGATACCCTTACTTTGTTTGATGATGGGTATGCATGGAGAAAATATGGGCA

AAAAACAATCCTCAATGCCAATCATTCAAGAAACTACTTCAGATGCACACACAAGGATGATCAAGGGTGC

CAAGCAACCAAGCAAGTGCAGCAGATTGAAGATGACCCACCAAAGTACAGGACAATCTATTATGGCCATC

ACACTTGCAAAACTCTGCTCAAAGCTTCCCAACTCATCTTGGATTCCACTACTTCCAATGACTCTT

>FK826430.1 JutEST43134 Jute Plasmid library 20-45 days after sowing Corchorus capsularis cDNA similar to Late embryogenesis-like protein, mRNA sequence

GGGGAATTAGTTACATTGAGATTCCCATCACCTTCAGGCCTAAGGATTTTGGATCTGCACTTTGGGACAT

GATTAGGGGAAAAGGAACAGGTTACACTATGAAAGGTCTAATCCATGTCGACACCCCTTTTGGAGCAATG

AAATTGCCTATCAGTAGGGAGGGTGGCACAACCCGCTTGAAGAAAAACAAGGAAGATGGTGGCGATGATG

ATGATGAGGATTAGAGCAATATTGATGTCGATGTCGATAGGGGTGCAGTGTGGATGGTCGTTTTGAGCTA

TCTTCTGTTTTATCATCGATAGAATAACGGACAAATAGATTGTTGAACTTTAATGTATTATAAGATTGTG

ATATATATT

>FK826429.1 JutEST43132 Jute Plasmid library 20-45 days after sowing Corchorus capsularis cDNA similar to Arabinogalactan protein, complete, mRNA sequence

GGGCATCCGCCTCATGAAAGCCACCCAAGTTGCTAACCAGCTCCTTGGTCAGCTCAATAACTCAAACAAT

GGCATCACCATTTTTGCACCATCCGATAGTGCTTTCTCCAGCCTTTCTGCCGGCACCTTGAACACCTCAC

CGACGAACAAAAAGTTGAGCTGGTTCAGTTCCACATTATCCCAACTTTCCTTTCTTCGTCTCAGTTCCAA

ACTGTTTAGTAACCCCTTTTGAGAACCCCAAGCTGGGGGATAGCGCCCGACGGCAAGTTTCCCTCTCAAT

GTGACCACTTTCGGGAAACTCAGTCAACATAAC

>FK826428.1 JutEST43127 Jute Plasmid library 20-45 days after sowing Corchorus capsularis cDNA similar to Stearoyl-ACP desaturase, complete, mRNA sequence

GGATTCCAAGAAAGGGCAACTTTTATCTCCCATGGTAACACTGCCAGGCTTGCCAAGGAGCATGGGGACA

TTAACTTGGCTCAGATATGTGGAATAATTGCCTCAGACGAAAAACGCCATGAGACAGCCTATACTAAAAT

TGTGGAGAAGCTCTTTGAGATTGATCCTGATGGTACTGTCCTGGCCTTTGCTGACCTGATGAAGAAGAAA

ATCTCCATGCCAGCCCACTTGATGTATGATGGCAAGGATGACAACCTTTTCGACCACTTCTCAGCTGTTG

CACAAAGGCTTGGGGTGTACACTGCCAAGGACTATGCTGATATCCTAGAGTTCCTTGTGGAAAGATGGAA

GGTGAAGGACCTAACTGGACTTTCTGGAGATGGGCGTAAAGCTCAGGATTATGTGTGTGGACTTCCTGCA

AGAATTAGGAGGCTGGAGGAGAGAGCTCAAGGAAGGGCCAAGCAAGGACCCAGTATTCCTTTCAGTTGGA

TTTATGACAGAGAAGTGATGCTGTAGGTGTTTAAGTAAAGAAGTGCTATCAATTTTCGTCTCTTGCAAGC

GTCTTATTTGATTCAAGGAAAAACAGAGATATAGAGAAACATGATGAAGCTGCAACT

>FK826427.1 JutEST43124 Jute Plasmid library 20-45 days after sowing Corchorus capsularis cDNA similar to GTP binding, mRNA sequence

GGGAGTTTGATGATGATGGGTTTGGGAAATTGGAGGAGAAAAATGATGAGGAAGAAGATGCTCCTTCATT

GTCTTTCTCTGATAAAAAGAAGTCATCCAAGTCTTCTAAGAAGAGTGGTAGGACTTTTGGGGGGTCAGCT

TTTGATGCCATTGCAGATGAAGATGATGACGATGGAGAGGTAGTTGAAGAAGAGAATAAAGTGGATGATG

ACGACGATGATGATATGGTCGCTAAAGCTTTGAAAGGGAAAAAAAAGAAGTCCAAGGGGGGAAAGAAAGC

AGGAAACTCGTTTGCAGCAGTTGCACTTGATGAGCTTGATGATGGAACAGGTGAAGAGGAGGATGAGAAG

AAAGAAGAGGAAGAAGATGTTCCTTCATTTCCTTTCTCGGACAAGAAAAAAAAGTCCTCAAAGTCTTCTA

AGAAGGGTGGTAATTCCTTTAGTGCTGCATTCCTCGAGGATGAAAATGATGGAGACGCTCCTGTATCAGA

ATCCACTAGTGCTATGGATGATTATGATGATGCTTCAGCAATTGCATTTTCAGGTAAGAAAAAGAAATCC

TCCCAGAAGAAGGGTGGGACTGAGCTGCAAACTGAAGTTTTAGA

>FK826426.1 JutEST43123 Jute Plasmid library 20-45 days after sowing Corchorus capsularis cDNA similar to Tetratricopeptide-like helical, mRNA sequence

GGGATAAAGGTCAGGTGAGAAAAGCTTTTGAGTGCATGAAGGCTGCCCTATCATTAACCGCGGTTAGTAA

GGTATGGAGACCCAACCTTAGGGTGGTTACAAGCATCTTGGATTGGCTAGGTGATGAAGGCAGTCTTCAA

GAGGTGGAAGATTTCGTGGCCTCACTGAGGACTGTCATGCCAGTTCACAGAAGAATGTACGTTGCCTTGT

TGAAGGCAAATGTGAGAAACGGTAATGGAGTTGATGAACTTCTGAAGCTCATGAAAGCTGATAAAATAGA

TGAAGACAAAGAAACAAAAGCAATATTAAGTATGAAGTCCTGCTAAGTTACAGATAGGTTCCTTACTGTT

GATCCATAGTTGGTCATAATGTATTTTGCATTTGTTCATTTGCTTCAACAATTGATATGGATTTTGTTCA

GACTTTTATCGGCTATCAAAATGGCAGTGGAAAGAATAAAATCAAATTTTAACC

>FK826425.1 JutEST43118 Jute Plasmid library 20-45 days after sowing Corchorus capsularis cDNA similar to Peptide methionine sulfoxide reductase, mRNA sequence

GGGAGAGTCAGGTCCAATACACCCAAAAACCCAACCTCCAACTCATAATAACAATGCTTCGAAGTCTTAC

GCTTGCCACCTCTTCATCTCTTTCTTCTTCCCTCATCTCCAAGCCTTTCTTTTCTCTCTCTCCAAAACTC

CCTCTCTCTTTCTCTCCCATTTCCCTCCCTTTCCCTCAAACCAAACGCTCCATTTCTCTCTACAGGGCCC

CCATGAACATCCTCAAGAACCTCGGCTTCGGAGCCAAGCCGGCCCAGTCAGCTGAAAACACTGCCATCGC

TCAGGGTCCGGACGACGACGTCCCGGCTCCCGGTCAGCAATTTGCTCAGTTCGGCGCGGGTTGCTTTTGG

GGCGTTGAATTGGCTTTCCAGAGAGTTCCGGGGGTCACCAAAACAGAGGTTGGGTATACCCAAGGGTTCA

TACATAATCCGACTTATGAAGACGTGTGTTCCGGGACCACGAACCATAGTGAAGTTGTGAGGGTTCAGTA

CGACCCTAAGGGAGTGTAGCTATGAGAGTTTGCTAGATGCCTTCTGGGCTCGCCATGATCCTACTACACT

GAATCGCCAGGGGAATGATGTTGGAACACAGTACAGATCCGGTATATACTTT

>FK826424.1 JutEST43112 Jute Plasmid library 20-45 days after sowing Corchorus capsularis cDNA similar to Cullin 1C (Fragment), complete, mRNA sequence

GGGATCTCAATCTCCCCGCAGAAATGGTGAAGTGTGTGGAAGTTTTTAAAGGATTTTATGAAACAAAAAC

AAAACACAGAAAGCTCACATGGATATACTCGTTGGGAACCTGTCACATTAATGGCAAATTTGACCAGAAA

ACTATTGAATTGGTTGTCTCCACTTATCAGGCTGCTGTACTGTTGCTCTTTAATGCCTCTGATCGGCTGA

GTTATTCAGAAATCATGGCTCAGTTGAACTTGACCAGTGAAGACTTGAACAGATTACTGCATTCACTATC

ATGTGCCAAGTATAAGATTCTACTCAAAGAGACAAGTACAAAAAGCATTTCCTCAAGTGACAACTTTGAG

TTCAACTCTAAATTTACGGACAAATTGAGGAGGATTAAGATTCCTCTCCCTCCAGTAGATGAAAGGAAGA

AAGTAGTTGAGGATGTCGATAAAGATCGTCGTTATGCCATTGATGCTGCTATTGTGCGTATAATGAAGAG

TAGGAAAGTTTTGGGCCACCAACAGTTAGTTTCAGAGTGTGTTGAGCAGTTGAGTCGCATGTTCAAGCCT

GATATTAAGGCCATCAAGAAGAGGATGGAAGATCTCATCACA

>FK826423.1 JutEST43101 Jute Plasmid library 20-45 days after sowing Corchorus capsularis cDNA similar to Epicotyl-specific tissue protein, mRNA sequence

GGGGCACAAGTTGATATATTTTATTATCAATATAATATAATATAATATTCTATTATATCCCTACATTGCA

GTTCTGCAACTATGAAATCTTTCCCTTCTTTCTTAGGTTTCCTGTCACTTCTCATGCTTGCTGACACCAT

AGCAGCAGCAAGAAAAGATGGTGGGGAATACTGGAGAGTAGCTGTAATGAAAGATGAATCCATTCCAGAA

GCCATCGAAAGCGCCCTCGTTCCGGTTAATGCCGCAGCTGCTTCCTCCTCCGGTGACAAAACCAACTGTC

ACGATTTACCTAGCAACATCGAGATTAAGGAAGAGAAGATTTTTGTTGAGGATTTTGAATTACCAAGGCC

TAATAATTATAACAGTGTTGAAGGAGGGGTGACGAAGGAACGTTCATTTGCTAAAGATTTCGAACCCAGG

CCCAACCTTTCGGCTTATGGTGATGATGGTGATCTTAAAGAAGAAAAAAAATCATTTGCTAAAGATTTCG

AACCGAGGCCCAACCTTTCGGCTTATGGTGGTGATGGTGATCTTGAAGAAGAGGAAAAATCATTTGCTAA

AGATTTCGAACCCAGGCCTAACCTTTCGGCTTATGGTGATGATGGTTATCT

>FK826422.1 JutEST43100 Jute Plasmid library 20-45 days after sowing Corchorus capsularis cDNA similar to Photosystem I subunit XI, mRNA sequence

GGGTTCACCGGTGGATTCTTCTTCGGTGGAATCTCTGGTGTCGTTTGGGCTTACTTCCTTCTCTATGTCC

TTGATCTTCCTTACTACTTTAAGTAGATGACATCTGTAATTGTTACTTTGTTCTGTCACTCCATATTTGT

AACTTGAATGTAAATCTATCTACTTTGGACCATCCTTTTGGAACGAATATCAAGTGATTTGTTACTGTG

>FK826421.1 JutEST4395 Jute Plasmid library 20-45 days after sowing Corchorus capsularis cDNA similar to Homeodomain-leucine zipper protein 57, mRNA sequence

GGAGGAGAAATATAAGCTAAAACAAGAGGTTGTCCAGCTCACAGATAAGTTGCTTGTGAAAGAGAAAGAG

AAGGGGAACTCAGAAGAACCACAGCAAAAGCTTGTTGTTGCTGAACAAGAGTCTAAGGTTTCTATGGCGG

TGGGCTGTAAGCAGGAAGATATGAGCTCAGCAAAAAGTGATATATTCGATTCAGAGAGCCCACATTACAC

AGATGGGGTTGTTCATTCTTCACTCCTGGAGGCTGCAGATTCATCTTATGCATTCGAACCAGAGCAATCT

GATTTGTCACAAGATGAGGAAGATAACTTGAGTAAGGGCCTCTTACATCCTCCTCCAATTCCATACATCT

TCCCAAAGCTTGAAGATGATGGTGATTACTCCGACCCGCCTGCAACTTCATGTAATTTTGGATTCCCTCT

TGAAGATCATGCCTTTTGGTCTTGGGCTTACTAAGCTCTATATTATCTATATCAAAATTCTGGTATATAT

ATATGATATGATCTTTATTGTGGTATTATATTTACTTCCTTGGTTAAGTCATCTTGTAGTATCTTTCT

>FK826420.1 JutEST4392 Jute Plasmid library 20-45 days after sowing Corchorus capsularis cDNA similar to Peroxiredoxin Q, mRNA sequence

GGGAAAATCATTTTTATTAGTAACACAAATTTCTTTCCCCATTTTTGTCACAAAAACAAAAAAATCATGG

CTTCATCATCTCTCACTCTTCCTAACCACACTCTTGCTTCTCTTCTTCCTTCTCACACTCCCAAGTCTCA

ATCTTCTGTTTCTCAATGTCTCCCACTTGCCTCCAAATCATCCCACTCTAGCTTTTTTGGCCTCAAGTTG

TCTTATTCTTCTTCTATTTCCATCCCTTCCTCTTCTTCTTTCAAGACTTCCATTTCTGCTAAGGTCAACA

AGGGTCAAGTGCCTCCAGCATTTACATTGAAAGACCAGGATGGGAAAAATGTGAGCCTTTCCAAGTTTAA

AGGAAAGCCTGTTGTTGTCTATTTCTATCCTGCTGATGAGACCCCTGGCTGCACCAAGCAGGCTTGTGCT

TTCCGAGATTCTTATGAGAAGTTTAAGAAAGCAGGAGCTGAAGTTGTCGGAATAAGTGGTGATGATACAT

CATCACACAAGGCATTTGCTAAGAAATACAGACTCCCATTTACATTACTGAGTGATGACGGAAACAAGGT

GAGGAAAGAGTGGGGAGTGCCATCAGATCTGTTTGG

>FK826419.1 JutEST4385 Jute Plasmid library 20-45 days after sowing Corchorus capsularis cDNA similar to Translation elongation factor 1A-4, mRNA sequence

GGTGCTGCAACAAGATGGATGCCACAACTCCAAAGTACTCCAAGGCAAGGTATGATGAAATTGTTAAGGA

AGTCTCTTCTTACCTGAAGAAGGTTGGTTACAACCCAGACAAAATCCCATTCGTCCCAATCTCTGGTTCT

GAGGGTGATAACATGATTGAGAGGTCAACCAACCTTGACTGGTACAAGGGTCCAACACTCCTTGAGGCTC

TTGACCAGATCAATGAGCCCAAGAGGCCCTCAGACAAGCCACTCCGTCTCCCACTTCAGGATGTCTACAA

GATTGGTGGTATTGGAACTGTCCCTGTTGGTCGTGTTGAGACTGGTGTTCTTAAGCCTGGTATGGTTGTC

ACCTTTGGTCCCTCTGGACTGACCACTGAAGTTAAGTCTGTCGAGATGCACCATGAAGCCCTCCAGGAGG

CTCTCCCCGGTGACAATGTCGGGTTCAATGTTAAGAACGTTGCTGTTAAGGATCTCAAGCGTGGTTACGT

CGCATCTAACTCTAAGGATGATCCTGCCAAGGAAGCTGCCAACTTCACCTCTCAGGTTATCATCATGAAC

CACCCTGGCCAGATCGGAAATGGATACGCCCCGGTGCTCGATTGCCACAC

>FK826418.1 JutEST4371 Jute Plasmid library 20-45 days after sowing Corchorus capsularis cDNA similar to hydrolase, hydrolyzing O-glycosyl compounds, mRNA sequence

GGGAGATGTAGCTTCGGATTTCTATCATAAATACAAGGAGGATATAGTATTGATGAAGAAGATTGGGTTG

GATTCCTTCAGATTCTCCATTTCATGGTCTCGAGTTTTACCAAAGGGGAGAGTTAGTGAAGGAGTAAACG

ACCTAGGAGTCAAATTCTACAACGACCTCATCGATGAACTCATTAAGAATGACATCAAGCCTTTTGTGAC

TTTGCTTCATTTTGATCATCCACAAGCTATTGAAGAAGAATACGGTGCATTCGGAAATCGCAAGTTTGTA

GATGATTTTGCTGAATATGCGGAATTCTGCTTCAAAACATTTGGTGACCGAGTCAAGTACTGGGCAACCT

TGAATGAACCAAATGGGTGGATCGTATCCATCATCCAAGCAGGTGGCTATGGATGGGTTGAACAAGGCGG

TAATGCCACTGAGTCGGGGCCCTTTCCCTTGGGCCATCATTTGATTCTTGCCCATGCAGCTGCTGCCAGA

ATCTACATGGTACAAGTACCAGCCAACTCAAAAGGGAAAGATTGGAATTACCATAAATACCACTTGGTAC

AAACCACTGTATGCCACAAAAGAACAGGAAGAGGCAGCGGGATAGAGCTCTTGATTTCGT

>FK826417.1 JutEST4369 Jute Plasmid library 20-45 days after sowing Corchorus capsularis cDNA similar to Photosystem I reaction center subunit II, chloroplast precursor (Photosystem I 20 kDa subunit) (PSI-D), mRNA sequence

GGGGTCGTCAAGGAGTTGGGCAGAACTTCAGATCGATTGGGAAGAATGTGAGTCCCATTGAGGTTAAGTT

CACTGGGAAACAACCTTATGACTTGTAAGAGATGAACGTCTGAGAGACGGTGCTTAG

>FK826416.1 JutEST4368 Jute Plasmid library 20-45 days after sowing Corchorus capsularis cDNA similar to structural constituent of ribosome, mRNA sequence

GGTCAGGAATGTAAGATTCTTGGCGAACTTCTTGACAGATGCTGGTATTATCGTCAAGAGGAGCTCGACT

GGTATTAGTGCCAAGGCCCAGAGGAAGCTTGCCAGGGAGATCAAGACCGCCAGGGCTTTTGGTCTGATGC

CTTTTACAACAATGGGTACAAAGAGGTTTGTTTTTGGAAGAACTATGGAAGATCAGGATGAAGATTATTC

ATACGAGTCTCTTGATACGAACCCTGAAGAAAACGATCTTTATGGACGCTAAGCTTGGACTAAGATCGGA

CTCTACAATTGTTTCTTGTCCGGGTATCATACAATTTACTGCAAGTGCCATGGCCAGCCTACTACAGCTG

AAATCATCAAACTTCATATATCTAAGCATTTCTGATGTCTACTTGAAATATACACTGCGTTACTATCAGA

TTAGTGACATCAAGGATGTAAGAAGTTACTTCATTCTATCAAGTGAACAATTTTATTTTCGAAAA

>FK826415.1 JutEST4355 Jute Plasmid library 20-45 days after sowing Corchorus capsularis cDNA similar to Transcription factor LIM, mRNA sequence

GGATCAAACTACATTGCACATGAAGGAAAACTTTACTGCAAACATCGCCATATTCAACTCTTCAAGGAGA

AAGGAAACTACAGTCAGCTCGAGAATGATCGCGAGAAAGCTGCCATGACAGAGAATCTTGCATCCATGAA

ATTGCAGCTGAATCATAGTCCATTGCTCTTGTTTTGGGTTTGTCACTGAGGATTTTTCTGTTTCTTCTCT

ACGTACCATATGTTAGTCTGTATCCTCCCCTTTTCCTCTCCATCTTTGAAGTGTGCCATTGGGTTACTAT

AAAGTGGAGTGAGCTTTTGAAGAATAAGACTAATCATGGTTTGTTTCAGCATTTGCTTCAAAGTATTATA

AAAAAAAGAGAGAATATTTTTGTGGATTTCAAAGAAGGAATGTATTTGCTTCACTTTTTGTTCAGGTTTT

TGCTTTGCTGGTTTTTTCCTGATGGAAAATTGCTATGGTATAAGTATTGTCAAAAATCAAAATGATTCCC

AATTGTTTGTACATTGAATATTGAAA

>FK826414.1 JutEST4347 Jute Plasmid library 20-45 days after sowing Corchorus capsularis cDNA similar to Arabinogalactan protein, mRNA sequence

GGCATCCGCCTCATGAAAGCCACCCAAGTTGCTAACCAGCTCCTTGGTCAGCTCAATAACTCAAACAATG

GCATCACCATTTTTGCACCATCCGATAGTGCTTTCTCCAGCCTTTCTGCCGGCACCTTGAACACCCTCAC

CGACGAACAAAAAGTTGAGCTGGTTCAGTTCCACATTATCCCAACTTTCCTTTCTTCGTCTCAGTTCCAA

ACTGTTAGTAACCCTTTGAGAACCCAAGCTGGGGATAGCGCCGACGGCAAGTTTCCTCTCAATGTGACCA

CTTCGGGAAACTCAGTCAACATAACGTCAGGGCTTACCAACACCGGCGTTTCTGGCACCATTTACACCGA

TGGTCAACTTGCTGTTTATCAAATCGATCAGGTGCTTAAGCCTTATCAGATATTTGATCCTAAGCCTCCA

GCTCCGGCACCGGCTCCGGCCAAGTCTAAGAAGAAGGCTGCTGCTATTGCTGAGAGCCCCGATGATACCA

CGGCTGATAACTCTAAGGCTGTCAGCCTCACCGCCATGCAAAATGTAGTGTTTTGGGGAGTTAGTGCTAT

TGTCATTGCACTTTCTTTGTGATCATGAGAAAAAAGAAATTATCAGAAGAAGTTTTGAT

>FK826413.1 JutEST4341 Jute Plasmid library 20-45 days after sowing Corchorus capsularis cDNA similar to 3-hydroxy-3-methylglutaryl-coenzyme A reductase (HMG-CoA reductase), mRNA sequence

GGGCTTGCATCTCAATCAGCATGTTTGAACTTGCTTGGTGTCAAGGGTGCACACAAAGAGTCACCAGGAG

CAAACTCTAGGATGCTTGCCACCATTGTTGCTGGTGCTGTTCTTGCCGGAGAGCTCTCACTTATGTCTGC

ACTTGCCGCCGGGCAGCTAGTTAAGAGCCATATGAAATACAATAGGTCAAGCAAGGATGTTACCACGGCT

GCTTCCTCTTAGAGACCACATAAATGGAAGGCTTTGTTCAAAAAGGCCTGAATTTGGTGTAAAGATGTTA

GTCCCAATGGTGTAAACGTAGCTAAGCTAAGGAGACATCATCATCATCATCATCTCGTTGTAATAAGTTA

AGATCTCTTTGTAATTTCTTTTATTTATTTTGTATTTTGTTCCTTTCTTTCTTTATTTTTTTTAATATTA

AGAAAAAAAAGAAAGAAAGAACCAACCAAGCCATGTAATCTGAGAATTGTGTAAGATGACAGAACTGTTT

AGTAACATTTTTTTTTCTTCTCCTGAAAAAAGA

>FK826412.1 JutEST4336 Jute Plasmid library 20-45 days after sowing Corchorus capsularis cDNA similar to Nodulin-26, mRNA sequence

GGGCCATCGGTTTCATCGTTGGGGCCAACATTTTGGCTGGTGGGCCATCCTCTGGAGCCTCAATGAACCC

AGCCGTTGCTTTCGGACCAGCTGTCGTTGGCTGGAGCTGGGCCAACCACTGGATCTACTGGGCTGGACCT

CTTGTTGGCGGTGGGATTGCTGGAGTCCTGTACGAGTTCATCTTCATTAGCCACACCCATGAGCAACTCC

CCACAACCGAATACTAGAGGGGGGGATTTGATAATAAAAAAAAAAAAATTGTCAATGAAAGGATTTCTCC

GGTTTATTCTAATTTGTATGGTTTTCTTTTCTATTATTTTCCTTTGTAATTTGTGTTTCATTGTGGCCGT

TGATCTTTCATTATCATCCATCATCATGATCATCATGGAAGGTGTTGTGATTGAAAAATTTGCATGCCCC

CTGGTAAAATATTGTGTGGGAATATATTTATGACCAAAAATACCAGGGGATTGATGTAGATGTGAAATGT

TAGCCTTTTTTAATTTCTTTGTTAATTTCTATGGGTTCATTTTCCTCCAAAAAAAA

>FK826411.1 JutEST4333 Jute Plasmid library 20-45 days after sowing Corchorus capsularis cDNA similar to Alpha-expansin 1, complete, mRNA sequence

GGGATTCACGCCGCTGCTTCTGTAGAACAACACTAACAAGTGATTTGGTACTGCAAAAGGAGACCCTTTT

CGAAGACTTTTCACCATGAGAGCCAAGTGGAAGAAGAAGCGTATGAGAAGGTTGAAGAGGAAGAGAAGAA

AGATGAGACAAAGATCTAAGTAGACTAACTTTACTTTAGAGCAATTATGCCCGTTTAGATTTCTTAGTTT

GTTTGTTGTTGATAGATCCTTTATGTTAAAAGGATCGAATTTATCATTCTTGAAACCTGTTTAGTATTGT

GAACACTTGTTAACCACTTTTCTATTTACTGGGAAAGCTTATGGATCTTAATTTTCTTTTGGATCTG

>FK826410.1 JutEST4320 Jute Plasmid library 20-45 days after sowing Corchorus capsularis cDNA similar to Eukaryotic initiation factor 4A-11 (eIF4A-11), mRNA sequence

GGGCATTAGGTGATTACCTTGGTGTTAAGGTTCATGCTTGTGTAGGTGGGACCAGTGTCCGTGAAGATCA

AAGGATTCTCTCCAGTGGAGTTCATGTGGTTGTTGGTACACCTGGTCGTGTGTTTGACATGTTGAGGAGA

CAATCTCTCCGCCCTGATCACATAAAGATGTTTGTGTTGGATGAAGCAGATGAAATGCTCTCCCGAGGTT

TCAAGGATCAGATCTATGATATTTTCCAATTACTGCCACCAAAAATTCAAGTAGGGGTTTTCTCTGCTAC

AATGCCTCCTGAAGCCCTTGAGATCACAAGGAAGTTTATGAATAAACCTGTGAGAATTCTGGTGAAGCGT

GATGAACTTACCCTTGAGGGTATTAAGCAATTTCATGTCAATGTTGAGAAGGAGGAATGGAAGCTTGAGA

CGCTATGTGATCTCTATGAAACCTTGGCAATCACCCAAAGTGTCATTTTTGTGAACACTAGGCGTAAAGT

GGATTGGCTTACAGACAAGATGAGAGGTCGTGACCATACAGTGTCTGCCACCCATGGAGATATGGACCAA

AATAC

>FK826409.1 JEST4311 Jute Plasmid library 20-45 days after sowing Corchorus capsularis cDNA similar to Aquaporin, mRNA sequence

GGACCACCAGTACATTCTCAGAGCTGCAGCCATCAAGGCTTTGGGATCTTTCCGCAGCAACCCCACCAAC

TAAATGATGAAAAACAAAGAAACGACTCATCGAAAACCCAAAAATAGCTTTGTCTTCTTCACTTCTCAAC

CCCCATCTTATTTTCTCCATTTGTTCGTTTGTGTGTATGAGAGAATGATGATGATTATGATGATATGATG

ACCCTTTTTTCTATTTCATTTATCTTATTTATAATTTAAATCTTTCTTTTGTTGTTAGCTCTTGCTCCCT

TGTGTATTATTGTTCATGAGTAGTAGTAGTACTTCTGGTGTTCATTTTTTCC

>FK826408.1 JEST42124 Jute Plasmid library 20-45 days after sowing Corchorus capsularis cDNA similar to ATP synthase F0 subunit 9, mRNA sequence

GGGAAGAAAAATAAGGGTCTAATTTCTGGTTTTTTTTATATAAAATAAAATGAAAAAAAGAAAATCTTAA

AGCGTGAGAAATTCAATTATAAATAATTTAATAAATGCTAGAAGGAGCTAAATTAATTGGTGCAGGAGCA

GCTACAATTGCTTTGGCGGGCGCAGCAGTAGGAATCGGAAATGTATTTAGTTCATTAATACATTCCGTGG

CAAGAAATCCATCCCTAGCAAAACAATTATTCGGATATGCAATTTTAGGCTTTGCTCTAACCGAAGCAAT

TGCCCTCTTTGCATTAATGATGGCATTTTTGATCTTATTCGTATTTTGAGAGTATCGGCTCAACATTACA

TTTTTTTCCATTTCCCGACTCGAATTTCTTTTTTTAGAAAAAAAAAGAAAAA

>EL595867.1 EST00179 Jute Plasmid library Corchorus capsularis cDNA, mRNA sequence

ATGCCAAGATCTTGGCCATGGACATTAACAGGGAAAACTACGAGTTGGGTCTGCCTGTAATCCAAAAAGC

AGGCGTTGCACACAAAATTGAATTCAAAGAGGGCCCTGCTTTGCCTGTTCTTGACAAACTAGTCGAAGAT

GAAAAGAATCATGAATCATATGACTTCATCTTCGTGGATGCTGACAAAGACAACTACATAAACTACATAA

ACTACCACAAGAGGTTAATAGACCTTGTGAAAGTTGGAGGCTTAATCGGCTACGACAACACTCTATGGAA

TGGTTCGGTCGTCGCACCACCTGACGCACCAATGAGGAAGTATGTTCGGTACTATCGGGATTTCGTCTTG

GAAATAAATAAAGCACTCGCTGCCGACCCAAGGATCGAGATTTGCATGCTACCTGTCGAAGATGGTATCA

CAGTATGTAGGCGGATCCAATGAGGGATTCTGGCTATCCAGCTAG

>EE332679.1 EST00175 Jute Plasmid library Corchorus capsularis cDNA, mRNA sequence

AAACAGGGGATGCAGTGGCCCGCGCTGATGCAAGCCCTTGCCCTGAGAGAAGGCGGTCCGCCCTCGTTCC

GGCTAACCGGAATTGGGCCTCCCGCGGCGGACAACACCGATCATCTGCATGAAGTGGGTTGGAAATTGGC

CCAGCTGGCGGAGGCGATTCATGTGGAGTTCGAGTACAGAGGTTTCGTGGCCAACAGTTTGGCAGATCTC

GACGCATCCATGCTTGAGCTTCGACGAGCA

>EE332677.1 EST00173 Jute Plasmid library Corchorus capsularis cDNA, mRNA sequence

TCGTTGCAGCTTTTCCTGGGAGTATAGCATGGGTTACTTCAGCGCCGTAGCCCTGGTACCTGAACATTGG

CTCGGGGCATTTTCTCTACCCCTTCTTNCCCTGAAAAAGCAGGGGACCTTGCGTCCTTGAACCGATAACC

ATCTTTCGGCTAACCTAGCCTCCTCGNCCTCGGGACCAACAAGGGGTAGTACAGGAATATTCACCTGTTG

TCATCGACTA

>EE332676.1 EST00172 Jute Plasmid library Corchorus capsularis cDNA, mRNA sequence

TGTAGGGGCAGAGGGAATTTCCGGTGGAGCGGTGAAATGCGTAGAGATCGGAAAGAACACCAACGGCGAA

AGCACTCTGCTGGGCCGACACTGACACTGAGAGACGAAAGCTAGGGGAGCGAATGGGATTAGATACCCCA

GTAGTCCTAGCCGTAAACGATGGATACTAGGCGCTGTCGGTATCGACCCGTGCAGTGCTGTAGCTAACGG

TAAGTATCCGCCTGGAGCATGGCGCAGGCTGAAACTTAAGGATC

>EE332675.1 EST00171 Jute Plasmid library Corchorus capsularis cDNA, mRNA sequence

AGTCAGCTGATGCTGCAGTCGACTCTAGAGATCGGTACCAGCTCTTCGCGCCGCTGGACCCTACCTCCGA

CTGAGTCGTTTCCAGGGTGGGCAGGCTGTTAAACAGAAAAGATAACTCTTCCCGAGGCCCCCGACGACGT

CTCCGGACTCCCTAACGTTGCCGTCAGCCGCCACGTCCCGGTTCAGGAATTTTAACCCGATTCCCTTTCG

AAGCTAGCGTTGCACGCGCTATCGGACGGGCTTCCCCCGTCTCTTAGGATCGACTAACCCATGTGCAAGT

GCCGTTCACATGGAACCTTTCCCCTCTTCG

>EE332674.1 EST00170 Jute Plasmid library Corchorus capsularis cDNA, mRNA sequence

GCCGTCGCTCGATGGGAATTGCGCTTCAAGGCGGCCTGCGCGGCTCGTCCGCCAGCGAGCTTAGCCTACG

ACACGTGCCTTTGGGGGCCAAGGCCCCTACTGCGGGTCGGCAATCAGGCGACGGGCGCACGCGTCGCTTC

TAGCCCGGATTCTGACTTAGAGGCGTTCAGTCATAATCCACGCACGGTAGCTTCGCGCCACTGGCTTTTC

AACCAAGCGCGATGACCAATTGTGCGAATCAACGTCTTGCAAGG

>EE332673.1 EST00169 Jute Plasmid library Corchorus capsularis cDNA, mRNA sequence

AAATTGCCGGTACTTACCAAGACGGCAATGGGTTTTGTGAAAGCCACGCTGAAAGCTCTAAGTGAAGAGC

CTGCCCCATTGACTGAAGCACAACGGCATGAACTGTCATGGGAAGCTGCTACAGAGCGGTTTCTAAGTGT

ATCTGAGCTCAAACAGGTAGTTGATAAAAAATTTTCGAAAATTTCTCAAAAGAAATTTGCATCAACGTCT

TTAAACCTACGAAAGAATATAGGATGATACATCAGCATATTTTCATCATGTGGCCTTAGGATTT

>EE332678.1 EST00174 Jute Plasmid library Corchorus capsularis cDNA, mRNA sequence

TCTTGAAAATCGGGTCCGGCTTCCCGTCGTCCGAATTGTAGTCTGGAGAAGCGTCCTCAGCGGCGGACCG

GGCCCAAGTCCCCTGAAAGGGGGCGCCGGAGAGGGTGAGTGCCCCGTCGTGCCCGGACCCTGTCGCACCA

CGAGGCCGTGTCTGCGAGTCGGGTTGTTTGGGAATGCAGCCCCAATCGGGCGGTAAATTCCGTCCAAGGC

TAAATACCGCCATGAGACGATAGCGAACAAGTACC

>EE332665.1 EST00161 Jute Plasmid library Corchorus capsularis cDNA, mRNA sequence

TAACAAAATCAATGTTTTGAGACTCCATAGCATGGATTCTGATCATGCCCATCATCATCTCATGGTTGGG

GTAACTAATATTCATCCATCGTCTCATCACATCCACATGGATCCTTCATTGAGGGTGTTTTTCACCTTGA

ATGAGCTTAGAGTTGGGAAGACAATACCCATATATTTCCCTAAGAATGACCCTTCAA

>EE332663.1 EST00159 Jute Plasmid library Corchorus capsularis cDNA, mRNA sequence

TCACAGCTTGCGGTTCATGCAAACCCAAGCCCAAACCCACCCCTAGTCCTCCTCCAGCTCCAAAAGCTTG

CCCTCCGCCACCAAAACCGGCGTCTTGCCCTAAGGACACATTGAAGCTCGGTGTCTGCGCCGACCTCCTC

GGACTTGTCAACGTCATTGTCGGAACACCCCCTTCAAGCAAATGCTGCGCTTTGCTCTCAGGCTTGGCTG

ATGCCGAAGCTGCCCTTTGCCTTTGCACTGCCATTAAAGCCAATGTACTTGGAATCACCCTCAACATTCC

TGTTACTCTTAGCTTACTCCTTAGTGCATGCCAAAAGGAAATTCCTCCAGGCTTCAAGTGTGAATAAGGA

AG

>EE332662.1 EST00158 Jute Plasmid library Corchorus capsularis cDNA, mRNA sequence

ATCTTGGGGTGGGCTTACCCCTTAGATGCTTTCAGCAGTTATCCGCTCCGCACTTGGCTACCCTGCGTGC

GCCGTGGGCACGACGACAGGTACACCAGAGGTACGTTCAACCCGGTCCTCTCGTACTAGGGTCAACTCCT

CTCAAATTTCGACGCCCACGGCAGATAGGGACCAAACTGTCTCGCGACGTTCTGAACCCAGCTCACGTAC

CACTTTAATTGGCGAACAGCCAAACCCTTGGGACCTGCTCCAGCCCCAGGATGTGATGAGCCGACATCGA

GGTGCCAAACAACCCCGTCGATATGAGCTCTTGGGGGTTATCAGCCTGTTATCCCCGGCGTACCTTTTAT

CCGTTGAGCGATGGCCCTTCCA

>EE332661.1 EST00157 Jute Plasmid library Corchorus capsularis cDNA, mRNA sequence

TCAAGAATCTGAATTGTCTCCCTCAGGCATGTTTCGGTTAGCTCTCGAAGACATGGTGATGTTTAATCTT

GAAAGGATCGATTAGTTAGACAACATTTTCACAGTACAAGCAAATTCCTTGAAATGGTGGACCATTATTC

TTAAATCTTAAACATACCCAAAAGCATAAACAGAGGATGCATGAAGTTGAAGCCTCTTACTCATATAGAA

AATCTTTTTCTTAGCACAATAGTTGGTTAGTCCAGCTGAATCCAGGTTCCTTGTGACTGGTTTTTTTTTT

TTACACCATAGGACCACAACAATAACTCTAAAAATAAATTAAGAAAAACGGAATTTAATGAAAAAAAAAT

AAAAGCCAAAAAAGAAATTTTTGGGAGGTGGAAATTAAAAAAAAAAAAAAGAGCAAAGGGGGCCAACCAT

GTGATTGATGATGAAGACTGGAATATCAATCCTCAACCGGATA

>EE332660.1 EST00156 Jute Plasmid library Corchorus capsularis cDNA, mRNA sequence

TATCTCGAGGGAGGCTTCCCGCTTAGATGCTTTAAGCGGTTATCCCGTCCGTACATAGCTACCCTGCTGC

GCCGTTGGCACGACGACAGGTACACCAGAGGTACGTTCAACCCGGTCCTCTCGTACTAGGGTCAACTCCT

CTCAAATTTCGACGCCCACGGCAGATAGGGACCAAACTGTCTCGCGACGTTCTGAACCCAGCTCACGTAC

CACTTTAATTGGCGAACAGCCAAACCCTTGGGACCTGCTCCAGCCCCAGGATGTGATGAGCCGACATCGA

GGTGCCAAACAACCCCGTCGATATGAGCTCTTGGGGGTTATCAGCCTGTTATCCCCGGCGTACCTTTTAT

CCGTTGAGCGATGGCCCTTCCA

>EE332659.1 EST00155 Jute Plasmid library Corchorus capsularis cDNA, mRNA sequence

TATCTCGAGGGAGGCTTCCCGCTTAGATGCTTTAAGCGGTTATCCCGTCCGTACATAGCTACCCTGCTGC

GCCGTTGGCACGACGACAGGTACACCAGAGGTACGTTCAACCCGGTCCTCTCGTACTAGGGTCAACTCCT

CTCAAATTTCGACGCCCACGGCAGATAGGGACCAAACTGTCTCGCGACGTTCTGAACCCAGCTCACGTAC

CACTTTAATTGGCGAACAGCCAAACCCTTGGGACCTGCTCCAGCCCCAGGATGTGATGAGCCGACATCGA

GGTGCCAAACAACCCCGTCGATATGAGCTCTTGGGGGTTATCAGCCTGTTATCCCCGGCGTACCTTTTAT

CCGTTGAGCGATGGCCCTTCCA

>EE332664.1 EST00160 Jute Plasmid library Corchorus capsularis cDNA, mRNA sequence

TCCCCGTAGGAGTCTGGGCCGTGTCTCAATCCCAGTGTGGCTGATCATCCTCTCGGACCAGACTACTGAT

CATCGCCTTGGTAAGCTATTGCCTCACCAACTAGCTAATCAGACGCGAGCCCCTCCTCGGGCGGATTCCT

CCTTTTGCTCCTCAGGCTACGGGGTATTAGCAGCCGTTTCCAGCTGTTGGTCCCCTCCCAAGGGCAGGTT

CTTACGCGTTACTCACCCGACCGCCACTGGAAACACCACTTCCCGTCCGACTTGCATGTGTTAAGCATGC

CGCCAGCGTTCATCCTGAGCCAGGAT

>EE332651.1 EST00147 Jute Plasmid library Corchorus capsularis cDNA, mRNA sequence

AAAAAAACAAAGGACAATACCTCAAAAAGAAAGGAAAAGTAAAAAGGTTTCGACAAATGCTAAGTACAAT

TTAAACAAGCCGAATCCCTACAATGTGGGGAAAACAACTCTGTTCATTTAAGCTGAGGTGAAGCTTGGGG

GCTTGTAGGCAATGAAACTGATGCATTGCACTTGACGCTTGTTGTCGAATCCAATGTTTCTGATGAAGGA

CTCGGGGTATTCCCTCTTGGCCTCCTCANGCTCCTTCAACACCTGGGATGAGTCGGTGCACCCGAACATG

GGANGTTTTCNCATGGTCCAGTAGCGTCCATCGTANTACCCTGGGGGAGCTGGGGTTCTTACGGTANACN

ANTCNGGNCNCCAATCCAATTCNNAACCNGGCNNANCTGNTTCCCGGGGGAAATTTTTTNCNCTCCNAAT

TCCNNNAANANACAGNCCGNANCNTAAANGTNAAACCCGGGGGNCCCAAGGGGGGGCC

>EE332649.1 EST00145 Jute Plasmid library Corchorus capsularis cDNA, mRNA sequence

TGTCAAGGGCCAAGCAACATGCAAGATTTAAGGTGTTACAGTGCTAGTTATGCAAATTCTGTTGTTCAGG

TTCAGCCAAATCAGCTTGGGAACGAGATTAAGATCAAGAAAAGCAAAAGCAGTTTTGGGTCATCTTCAAA

AAGCTGGAGCTTTAATGATCCTGAGTTGCAGAGGAAGAAAAGGGTTGCAAGTTACAAGGTTTATGCAGTT

GAAGGCAAGATGAAAGGGTCTTTCAGGAAGAGTTTTAGGTGGATCAAAGACACTTATACCCAAGTTGTCT

ATGGATGGAGA

>EE332648.1 EST00144 Jute Plasmid library Corchorus capsularis cDNA, mRNA sequence

GTTGTAACGACGGCCAGCTGCATTAATGAATCGGCCAACGCGCGGGGAGAGGCGGTTTGCGTATTGGGCG

CTCTTCCGCTTCCTCGCTCACTGACTCGCTGCGCTCGGTCGTTCGGCTGCGGCGAGCGGTATCAGCTCAC

TCAAAGGCGGTAATACGGTTATCCACAGAATCAGGGGATAACGCAGGAAAGAACATGTGAGCAAAAGGCC

AGCAAAAGGCCAGGAACCGTAAAAAGGCCGCGTTGCTGGCGTTTTTCCATAGGCTCCGCCCCCCTGACGA

GCATCACAAAAATCGACGCTCAAGTCAGAGGTGGCGAAACCCGACAGGACTATAAAGATACCAGGCGTTT

CCCCCTGGAAGCTCCCTCGTGCGCTCTCCTGTTCCGACCCTGCCGCTTACCGGATACCTGTCCGCCTTTC

TCCCTTCNGGAAGNGTGNCGCTTTCTCATANCTCAC

>EE332647.1 EST00143 Jute Plasmid library Corchorus capsularis cDNA, mRNA sequence

AGAGGCGGTTTGCGTATTGGGCGCTCTTCCGCTTCCTCGCTCACTGACTCGCTGCGCTCGGTCGTTCGGC

TGCGGCGAGCGGTATCAGCTCACTCAAAGGCGGTAATACGGTTATCCACAGAATCAGGGGATAACGCAGG

AAAGAACATGTGAGCAAAAGGCCAGCAAAAGGCCAGGAACCGTAAAAAGGCCGCGTTGCTGGCGTTTTTC

CATAGGCTCCGCCCCCCTGACGAGCATCACAAAAATCGACGCTCAAGTCAGAGGTGGCGAAACCCGACAG

GACTATAAAGATACCAGGCGTTTCCCCCTGGAAGCTCCCTCGTGCGCTCTCCTGTTCCGACCCTGCCGCT

TACCGGATACCTGTCCGCCTTTCTCCCTTCGGGAAGCGTGGCGCTTTCTCATAGCTCACGCTGTAGGTAT

CTCAGTTCGGTGTAGGTCGTTCGCTCCAAGCTGGGCTGTGTGCACGAACCCCCCGTTCAGCCCGACCGCT

GCGCCTTATCCGGTAACTATCGTCTTGAGTCCAACCCGGTAAGACACGACTTATCGCCACTGGCAGCAGC

CACTGGTAACAGGATTAGCAGANCGAGGTATGTAGGCGGTGCTACAGAGTTCTTGAAGTGGTGGCCTAAC

TACGGCTACACTAGAAGAACAGTATTTGGTATCTGCGCTCTGCTGAANCCAGTTACNTNCGNAAAANANT

TGGTAGCTCTTGATCCGGCAAAAAACCACCGCTGGTA

>EE332646.1 EST00142 Jute Plasmid library Corchorus capsularis cDNA, mRNA sequence

TGCCTAAGATCAGGCCGAAAGGCGTAGTCGATGGACAACAGGTGAATATTCCTGTACTACCCCTTGTTGG

TCCCGAGGGACGGAGGAGGCTAGGTTAGCCGAAAGATGGTTATCGGTTCAAGGACGCAAGGTGCCCCTGC

TTTTTCAGGGTAAGAAGGGGTAGAGAAAATGCCCCGAGCCAATGTTCGAGTACCAGGCGCTACGGCGCTG

AAGTAACCCATGCTATACTCCCAGGAAAGCTCGAACGACCTTCAACAAAAGGGTACCTGTACCCGAAACC

GACACAGGTGGGTAGGTAGAGAATACCTAGA

>EE332645.1 EST00141 Jute Plasmid library Corchorus capsularis cDNA, mRNA sequence

TCGAGACCTTGAAAGCAGCTTTTTCAAGTGTCAGTAGCGGGACGTTCTGTCAATCGGGGAAGGTTTTTGG

TAACAACACCTGGAGATATCAGAAGTGAGAATGCTGACATGAGTAACGAGAAATCCTGTGAAAAACACGA

TCGCCTGCCAGTGGAAGGCTTTCTGCGTTCAGTCAATCTACGCAGAGTGAATCGGTCCCTAAGGAACCCC

CGAAAGGGCTGCCGTCCGATGGGTACACGAAAGTGACGAAGTTGCTTTGACTACAGAACCATGCCTGTCT

GTTGGAGCGAATTGGATGGATGATCGGGCCGAGGTTCTCGGCAAAGACCTCTTCCCCTCACTCTCCTTTC

CCTAATATTCACA

>EE332650.1 EST00146 Jute Plasmid library Corchorus capsularis cDNA, mRNA sequence

TATCTCGAGGGAGGCTTCCCGCTTAGATGCTTTAAGCGGTTATCCCGTCCGTACATAGCTACCCTGCTGC

GCCGTTGGCACGACGACAGGTACACCAGAGGTACGTTCAACCCGGTCCTCTCGTACTAGGGTCAACTCCT

CTCAAATTTCGACGCCCACGGCAGATAGGGACCAAACTGTCTCGCGACGTTCTGAACCCAGCTCACGTAC

CACTTTAATTGGCGAACAGCCAAACCCTTGGGACCTGCTCCAGCCCCAGGATGTGATGAGCCGACATCGA

GGTGCCAAACAACCCCGTCGATATGAGCTCTTGGGGGTTATCAGCCTGTTATCCCCGGCGTACCTTTTAT

CCGTTGAGCGATGGCCCTTCCA

>EE332637.1 EST00133 Jute Plasmid library Corchorus capsularis cDNA, mRNA sequence

TGGAGCCGTAGCGAAAGCGAGTCTTCATAGGGCAATTGTCACTGCTTATGGACCCGAACCTGGGTGATCT

ATCCATGACCAGGATGAAGCTTGGGTGAAACTAAGTGGAGGTCCGAACCGACTGATGTTGAAGAATCAGC

GGATGAGTTGTGGTTAGGGGTGAAATGCCACTCGAACCCAGAGCTAGCTGGTTCTCCCCGAAATGCGTTG

AGGCGCAGCAGTTGACTGGACATCTAGGGGTAAAGCACTGTTTCGGTGCGGGCCGCGAGAGCGGTACCAA

ATCGAGGCAAACTCTGAATACTAGATATGACCTCAAAATAACAGGGGTCGAGGTCGGCCAGTGAGACGAT

GGGGGATAAGCTTCCTCGTCGAGAGGGAAACAGCCCGGATCACCAGCTAAGGCCCCTACCTCCTTTATCA

CTGAGA

>EE332635.1 EST00131 Jute Plasmid library Corchorus capsularis cDNA, mRNA sequence

TTTTCTCTTCAAAAACAGAGAAAAAAGAGGACCATTCTAAGCACTTAGTTGTGTCACTTGAAAAATGGCT

TCAAAATCTGTAGCATCCATTGCCTTCCTTCTCACTCTCAACCTTCTCTTCTTCACTTTGGTCAGTTCAA

CCTACACCCCTTGCCCACCATCACCCAAAAAGCCCACCCCGACCCCTTCCCCGAAGCCAAAACCCAAACC

CAAGCCCGCCGCCTCTTGCCCTAAAGACACTCTCAAATTGGGATTTGTGCTGATTTGTTGGGTTTGGTTA

ACGTTGTTGTTGGTTCCCCTCCAAAGACTCCTCCTGCATCTCCTCGAAGGTTTGGCTGATCTTGAAGCTG

CTGCTTGCCTTTGCACTGCCATTAAGGCTAATGTTTTGGGAATTAACCTTAATGTTCCCGTTTCATTGTC

TTTGCTCCTTAATTACTGTGGAAAGAAGGTTCCTGCTGGCTACCAGTGTGCTTAATTAAAAGCTCTTTAA

TTTCGTCTTACTTGTAATAATCAATTACTGTCTTGGTACTTTCGGGCTTATGTTTGGTCGTTTTTATAGT

CGTTTTCATGGTGTGATATTGGAAGTTTGGAATCTGGGTACGTACGTATAGTATTATGTCATACTGCAAT

TTATTACTATATTTTTACTTTTAATTTGATCAAGGAATTAGGAAGGCTTCTCACCTTCTAATCCTTTTAT

ATATGCGAAACAAGAATGTATTATTTACATCACTCTGTATCTAANAAANGGAAAAGTTTTANTTTTATCA

>EE332634.1 EST00130 Jute Plasmid library Corchorus capsularis cDNA, mRNA sequence

AAAAAAACAAAGGACAATACCTCAAAAAGAAAGGAAAAGTAAAAAGGTTTCGACAAATGCTAAGTACAAT

TTAAACAAGCCGAATCCCTACAATGTGGGGAAAACAACTCTGTTCATTTAAGCTGAGGTGAAGCTTGGGG

GCTTGTAGGCAATGAAACTGATGCATTGCACTTGACGCTTGTTGTCGAATCCAATGATTCTGATGAAGGA

CTCGGGGTATTCCTTCTTGGCCTCCTCAAGCTCCTTCAACACCTGGGATGAGTCGGTGCACCCGAACATG

GGAAGTTTCCACATGGTCCAGTAGCGTCCATCGTAGTACCCTGGGGAGCTGTGGTTCTTACGGTAGACGA

ATCCGTGCTCCAATTC

>EE332633.1 EST00129 Jute Plasmid library Corchorus capsularis cDNA, mRNA sequence

TCAAACATTAATTAAACATTACATTAGGCCGAGGTATAAAATTAACCCACAAGCCTAGTTATTTAAAACA

TGAACAAATATCACACAATGTCGGAAGTCTTTGTAATAAAGACTCCCTAAAAATATGTAAAAATGGAAAC

CCTATCCCAACTTCAAATCGATCCAAACCACTAAACAAAAATACATTGAATATTTGAAACTACA

>EE332632.1 EST00128 Jute Plasmid library Corchorus capsularis cDNA, mRNA sequence

GTTTAACGACGGCCAGTGCCAAGCTTGCATGCCTGCAGGTCGACTCTAGAGGATCCCCGGGTACCGAGCT

CGCGTAATCATGTCATAGCTGTTTCCTGTGTGAAATTGTTATCCGCTCACAATTCCACACAACATACGAG

CCGGAAGCATAAAGTGTAAAGCCTGGGGTGCCTAATGAGTGAGCTAACTCACATTAATTGCGTTGCGCTC

ACTGCCCGCTTTCCAGTCGGGAAACCTGTCGTGCCAGCTGCATTAATGAATCGGCCAACGCGCGGGGAGA

GGCGGTTTGCGTATTGGGCGCTCTTCCGCTTCCTCGCTCACTGACTCGCTGCGCTCGGTCGTTCGGCTGC

GGCGAGCGGTATCAGCTCACTCAAAGGCGGTAATACGGTTATCCACAGAATCAGGGGATAACGCAGGAAA

GAACATGTGAGCAAAAGGCCAGCAAAAGGCCAGGAACCGTAAAAAGGCCGCGTTGCTGGCGTTTTTCCAT

AGGCTCCGCCCCCCTGACGAGCATCACAAAAATCGACGCTCAAGTCAGAGGTGGCGAAACCCGACAGGAC

TATAAAGATACCAGGCGTTTCCCCCTGGAAGCTCCCTCGTGCGCTCTCCTGTTCCGACCCTGCCGCTTAC

CGGATACCTGTCCGCCTTTCTCCCTTCGGGAAGCGTGGCGCTTTCTCATAGCTCACGCTGTAGGTATCTC

AGTTCGGTGTAGGTCGTTCGCTCCAAGCTGGGCTGTGTGCACGAACCCCCCGTTCAGCCCGACCGCTGCG

CCTTATCCGGTAACTATCGTCTTGAGTCCAACCCGGTAANACACGACTTATCGCCNCTGGCAGCAGCCAC

TGGTNACAGGATTANCAAACCGAGTTTGTAGGCGGNGCTACAANTTCTTGAAGNGGGGCCTANCTACG

>EE332631.1 EST00127 Jute Plasmid library Corchorus capsularis cDNA, mRNA sequence

GAATTGGAGCACGGATTCGTCTACCGTAAGAACCACAGCTCCCCAGGGTACTACGATGGACGCTACTGGA

CCATGTGGAAACTTCCCATGTTCGGGTGCACCGACTCATCCCAGGTGTTGAAGGAGCTTGAGGAGGCCAA

GAAGGAATACCCCGAGTCCTTCATCAGAATCATTGGATTCGACAACAAGCGTCAAGTGCAATGCATCAGT

TTCATTGCCTACAAGCCCCCAAGCTTCACCTCAGCTTAAATGAACAGAGTTGTTTTCCCCACATTGTAGG

GATTCGGCTTGTTTAAATTGTACTTAGCATTTGTCGAAACCTTTTTACTTTTCCTTTCTTTTTGAGGTAT

TGTCCTTTGTTTTTTT

>EE332636.1 EST00132 Jute Plasmid library Corchorus capsularis cDNA, mRNA sequence

ACCAAGTGTTGGATTGTTCACCCACCAATAGGGAACGTGAGCTGGGTTTAGACCGTCGTGAGACAGGTTA

GTTTTACCCTACTGATGGCCGCTTCGCAATAGTAATTCAACCTAGTACGAGAGGAACCGTTGATTCGCAC

AATTGGTCATCGCGCTTGGTTGAAAAGCCAGTGGCGCGAAGCTACCGTGCGCTGGATCATGACTGAACGC

CTCTAAGTCAGAATCCGGGCTAGAAGCGACGCATGCGCCCGCCGCCCGATTGCCGACCCGCAGTTGGGGC

CTTTGGCCCCCAAGGGCACGTGTCGTAGGAGTAGTCCGTGCGGCGGATGAGCCGTGCAGGCCTCCTTGAA

GCGTAATTCCCACCGAGCGGCGGGTAGAATCCTTTA

>EE332623.1 EST00119 Jute Plasmid library Corchorus capsularis cDNA similar to Tobacco chloroplast 23S and 4.5S rRNA genes, mRNA sequence

TCTCAATGCTCTAACGCCCACACCGGATATGGACCGAACTGTCTCACGACGTTCTGAACCCAGCTCACGT

ACCGCTTTAATGGGCGAACAGCCCAACCCTTGGAACATACTACAGCCCCAGGTGGCGAAGAGCCGACATC

GAGGTGCCAAACCTTCCCGTCGATGTGAGCTCTTGGGGAAGATCAGCCTGTTATCCCTAGAGTAACTTTT

ATCCGTTGAGCGACGGCCCTTCAACTCGGCACCGTCGGATCACTAAGGCCGACTTTCGTCCCTGCTCGAC

GGGTGGGTCTTGCAGTCAAGCTCCCTTCTGCCTTTGCACTCGAGGGCCAATCTCCGTCCGGCCCGAGGAA

ACCTTTGCACGCCTCCGTTACCTTTTGGGAGGCCTACGCCCCATAGAAACTGTCTACCTGAGACTGTCCA

>EE332621.1 EST00117 Jute Plasmid library Corchorus capsularis cDNA similar to Ribulose bisphosphate carboxylase small chain, chloroplast precursor (RuBisCO small subunit), mRNA sequence

TGGAGTACTAGCAATGGCTTCCGCAATGATGTCATCCGCCACCGTTGTTAACCGTGCCTCCCCGGCACAG

GCCACCCTGGTCGCCCCATTCGTTGGCCTCAAGTCTGCCTCTGCTTTCCCCACCACCCGCAAGAACAACA

AAGACATCACCTCCATCGTCAGCAATGGTGGCAGAGTTCGATGCATGCAGGTATGGCCACCACTTGGGAA

GAAGAAGTTCGAGACTCTCTCATACCTCCCGGATCTCACCCCCGTGCAATTGGCTAAGGAAGTAGACTAC

CTTCTCCGCTCTGGATGGATTCCTTGCTT

>EE332620.1 EST00116 Jute Plasmid library Corchorus capsularis cDNA similar to Type II keratin E1, mRNA sequence

GTAATCATGTCATAGCTGTTTCCTGTGTGAAATTGTTATCCGCTCACAATTCCACACAACATACGAGCCG

GAAGCATAAAGTGTAAAGCCTGGGGTGCCTAATGAGTGAGCTAACTCACATTAATTGCGTTGCGCTCACT

GCCCGCTTTCCAGTCGGGAAACCTGTCGTGCCAGCTGCATTAATGAATCGGCCAACGCGCGGGGAGAGGC

GGTTTGCGTATTGGGCGCTCTTCCGCTTCCTCGCTCACTGACTCGCTGCGCTCGGTCGTTCGGCTGCGGC

GAGCGGTATCAGCTCACTCAAAGGCGGTAATACGGTTATCCACAGAATCAGGGGATAACGCAGGAAAGAA

CATGTGAGCAAAAGGCCAGCAAAAGGCCAGGAACCGTAAAAAGGCCGCGTTGCTGGCGTTTTTCCATAGG

CTCCGCCCCCCTGACGAGCATCACAAAAATCGACGCTCAAGTCAGAGGTGGCGAAACCCGACAGGACTAT

AAAGATACCAGGCGTTTCCCCCTGGAAGCTCCCTCGTGCGCTCTCCTGTTCCGACCCTGCCGCTTACCGG

ATACCTGTCCGCCTTTCTCCCTTCGGGAAGCGTGGCGCTTTCTCATAGCTCACGCTGTAGGTATCTCAGT

TCGGTGTAGGTCGTTCGCTCCAAGCTGGGCTGTGTGCACGAACCCCCCGTTCAGCCCGACCGCTGCGCCT

TATCCGGTAACTATCGTCTTGANTCCAACCCGGTAA

>EE332619.1 EST00115 Jute Plasmid library Corchorus capsularis cDNA similar to Oryza sativa (japonica cultivar-group) cDNA clone:006-209-G02, full insert sequence, mRNA sequence

GTTGTAACGANGGCCAGTGCCAAGCTTGCATGCCTGCAGGTCGACTCTAGAGGAGAGGCGGTTTGCGTAT

TGGGCGCTCTTCCGCTTCCTCGCTCACTGACTCGCTGCGCTCGGTCGTTCGGCTGCGGCGAGCGGTATCA

GCTCACTCAAAGGCGGTAATACGGTTATCCACAGAATCAGGGGATAACGCAGGAAAGAACATGTGAGCAA

AAGGCCAGCAAAAGGCCAGGAACCGTAAAAAGGCCGCGTTGCTGGCGTTTTTCCATAGGCTCCGCCCCCC

TGACGAGCATCACAAAAATCGACGCTCAAGTCAGAGGTGGCGAAACCCGACAGGACTATAAAGATACCAG

GCGTTTCCCCCTGGAAGCTCCCTCGTGCGCTCTCCTGTTCCGACCCTGCCGCTTACCGGATACCTGTCCG

CCTTTCTCCCTTCGGGAAGCGTGGCGCTTTCTCATAGCTCACGCTGTAGGTATCTCAGTTCGGTGTAGGT

CGTTCGCTCCAAGCTGGGCTGTGTGCACGAACCCCCCGTTCAGCCCGACCGCTGCGCCTTATCCGGTAAC

TATCGTCTTGAGTCCAACCCGGTAAGACACGACTTATCGCCACTGGCAGCAGCCACTGGTAACAGGATTA

GCAGAGCGAGGTATGTAGGCGGTGCTACAGAGTTCTTGAAGTGGTGGCCTAACTACGGCTACACTAGAAG

AACAGTATTTGGTATCTGCGCTCTGCTGAAGCCAGTTACCTTCGGAAAAAGAGTTGGTAGCTCTTGATCC

GGCAAANAAACCACCGCTGGTAGCGGTGGTTTTTTTGTTTGCAAGCAGCAAATTACNCGCAAAAAAAAGG

ATCTCAAAAAAATCTTTNGATNTTTNTACGGGGTCTGACNCTCANTGGAACGAAANTCCCGTTAAGGGN

>EE332618.1 EST00114 Jute Plasmid library Corchorus capsularis cDNA similar to Alnus incana chloroplast 23S ribosomal RNA, mRNA sequence

TATAGGGTCTTTCTGTCCAGGTGCGGGTAGTCCGCATCTTCACAGACATGTCTATTTCACCGAGCCTCTC

TCCGAGACAGTGCCCAGATCGTTACGCCTTTCGTGCGGGTCGGAACTTACCCGACAAGGAATTTCGCTAC

CTTAGGACCGTTATTGTTACGGCCGCCGTTCACCGGGGCTTCGGTCGCCGGCTCCCCTGTCATCAGGTCA

CCAACTTCCTTGACCTTCCGGCACTGGGCAGGCGTCAGCCCCCATACATGGTCTTACGACTTTGCGGAGA

CCTGTGTTTTTGGTAAACAGTCGCCCGGGCCTGGA

>EE332617.1 EST00113 Jute Plasmid library Corchorus capsularis cDNA similar to Idesia polycarpa 18S ribosomal RNA gene, complete sequence, mRNA sequence

TCGGTAATTCCAGCTCTAATAGCGTATACTAAAGTTGTTGTGGTTAAAACGCTCGTAGTTCAGATTCTGG

GGTACTCTGAATACATAAACTCATGTGATTTAACCGTTACGTGGGCAGGGTACCTGAGATCACGTTGGGA

CCGCGGGTTTCTGATCTTAACTGACCGGAGATTCGTACCCGACCGTTTACTTTGAATAAAACAGAGCGCT

CAATGCAAGCTGACGCCTGAATGAAACAGCATGGGATAATGGAATAGGAGCTTTAGTGGGAGTAGGTACA

CCGCTAAGGCTAATGACGAATAGAGCTTATCGGGGGTGTTCGTATTGCACGGCTAGAGGTGAGATTCTTG

GACCCGTA

>EE332622.1 EST00118 Jute Plasmid library Corchorus capsularis cDNA similar to Probable peroxisomal (S)-2-hydroxy-acid oxidase 2 (Glycolate oxidase 2) (GOX 2) (Short chain alpha-hydroxy acid oxidase 2), mRNA sequence

TCTTAGCCCTGGGAGCCTCTGAATATTTATTGGGCGACCTGTGGTGTTCTCATTGGCTGCTGAGGGAGAA

GCTGGGGTGAGGAAAGTACTGTCAATGCTGAGGGAGGAATTTGAGCTAACCATGGCGCTGAGTAGGTGCC

GTTCACTGAAAGAAATCACCCGCAATCACATTGTGACAGATTGGGACACTCCTAAAGGTCTTCCGGCCCC

AAGGTTGTAAATGCATGATTCAGAGTATTAGTATCTGCATGCATTAATTACTACTTTTATGGCAAATCAT

AGATTTGGGTTGATTTTACTTGGTCTTCAACAGACCTGATCTTCTCTTTGAAACACTTTTATGTTTCAAC

AATGAGTTAATACTTTTGTTATGATGATTATCATGGCTGCTGCTGCCTTTCATTGTATATTCTA

>EE332609.1 EST00105 Jute Plasmid library Corchorus capsularis cDNA similar to Putative senescence-associated protein (Pisum sativum), mRNA sequence

TGGATTCTACCCGCCGCTCGGTGGGAATTACGCTTCAAGGAGGCCTGCACGGCTCATCCGCCGCACGGAC

TACTCCTACGACACGTGCCCTGGGGGCCAAAGGCCCCTACTGCGGGTCGGCAATCGGGCGGCGGGCGCAT

GCGTCGCTTCTAGCCCGGATTCTGACTTAGAGGCGTTCAGTCATAATCCAGCGCACGGTAGCTTCGCGCC

ACTGGCTTTTCAACCAAGCGCGATGACCAATTGTGCGAATCAACGGTTCCTCTCGTACTAGGTTGAATTA

CTA

>EE332607.1 EST00103 Jute Plasmid library Corchorus capsularis cDNA similar to Cytochrome P450 like_TBP [Nicotiana tabacum], mRNA sequence

TAAGGATTCTACCCGCCGCTCGGTGGGAATTACGCTTCAAGGAGGCCTGCACGGCTCATCCGCCGCACGG

ACTACTCCTACGACACGTGCCCTTGGGGGCCAAAGGCCCCTACTGCGGGTCGGCAATCGGGCGGCGGGCG

CATGCGTCGCTTCTAGCCCGGATTCTGACTTAGAGGCGTTCAGTCATAATCCAGCGCACGGTAGCTTCGC

GCCACTGGCTTTTCAACCAAGCGCGATGACCAATTGTGCGAATCAACGGTTCCTCTCGTACTAGGTTGAA

TTACTATTGCGACGCGGCCATCAGTAGGGTAAAACTAACA

>EE332606.1 EST00102 Jute Plasmid library Corchorus capsularis cDNA similar to Alpha-hemolysin, mRNA sequence

GGCCAGTGCCAGCTGCATTAATGAATCGGCCAACGCGCGGGGAGAGGCGGTTTGCGTATTGGGCGCTCTT

CCGCTTCCTCGCTCACTGACTCGCTGCGCTCGGTCGTTCGGCTGCGGCGAGCGGTATCAGCTCACTCAAA

GGCGGTAATACGGTTATCCACAGAATCAGGGGATAACGCAGGAAAGAACATGTGAGCAAAAGGCCAGCAA

AAGGCCAGGAACCGTAAAAAGGCCGCGTTGCTGGCGTTTTTCCATAGGCTCCGCCCCCCTGACGAGCATC

ACAAAAATCGACGCTCAAGTCAGAGGTGGCGAAACCCGACAGGACTATAAAGATACCAGGCGTTTCCCCC

TGGAAGCTCCCTCGTGCGCTCTCCTGTTCCGACCCTGCCGCTTACCGGATACCTGTCCGCCTTTCTCCCT

TCGGGAAGCGTGGCGCTTTCTCATAGCTCACGCTGTAGGTATCTCAGTTCGGTGTAGGTCGTTCGCTCCA

AGCTGGGCTGTGTGCACGAACCCCCCGTTCAGCCCGACCGCTGCGCCTTATCCGGTAACTATCGTCTTGA

GTCCAACCCGGTAAGACACGACTTATCGCCACTGGCAGCAGCCACTGGTAACAGGATTAGCAGAGCGAGG

TATGTAGGCGGTGCTACAGAGTTCTTGAAGTGGTGGCCTAACTACGGCTACACTAGAAGAACAGTATTTG

GTATCTGCGCTCTGCTGAAGCCAGTTACCTTCGGAAAAAGAGTNGGTAGCTCTTGATCCGGCAAAAAACC

ACGCTGGTAGCGGTGGTTTTTTTGTTTGCANGCAGCAAATNACNCGCANAAAAAAGGATCTCAAAAAATC

TTTGATNTTTTCTACGGGGTCTGACCCTCANTGGAACNAAACTCACGTTAAGGGATTTGGTCATGAA

>EE332605.1 EST00101 Jute Plasmid library Corchorus capsularis cDNA similar to Solanum nigrum chloroplast tRNA-Ala, tRNA-Ile, 16S rRNA, tRNA-Val, rps12, rps7, ndhB genes, mRNA sequence

TATGGCTGACCGGCGATTACTAGCGATTCCGGCTTCATGCAGGCGAGTTGCAGCCTGCAATCCGAACTGA

GGACGGGTTTTTGGAGTTAGCTCACCCTCGCGGGATCGCGACCCTTTGTCCCGTCCATTGTAGCACGTGT

GTCGCCCAGGGCATAAGGGGCATGATGACTTGACGTCATCCTCACCTTCCTCCGGCTTATCACCGGCAGT

CTGCTCAGGGTTCCAAACTCAACGGTGGCAACTAAACACGAGGGTTGCGCTCGTTGCGGGACTTAACCCA

ACACCTTACGGCACGAGCTGACGACAGCCATGCACCACCTGTGTCCGCGTTCCCGAAGGCACCCCTCTCT

TTCAAGAGGATTCGCGGCATGTCAAGCCCTGGTAAGGTTCA

>EE332604.1 EST00100 Jute Plasmid library Corchorus capsularis cDNA similar to Alnus incana chloroplast 23S ribosomal RNA (23S rRNA) gene, mRNA sequence

TGGAAAGGTCCCCTCAATGCTCTAACGCCCACACCGGATATGGACCGAACTGTCTCACGACGTTCTGAAC

CCAGCTCACGTACCGCTTTAATGGGCGAACTNGCCCAACCCTTGGAACATACTACAGCCCCAGGTGGCGA

AGAGCCGACATCGAGGTGCCAAACCTTCCCGCCGATGTGAGCTCTTGGGGAAGATCAGCCTGTTATCCCT

AGAGTAACTTTTATCCGA

>EE332603.1 EST00099 Jute Plasmid library Corchorus capsularis cDNA similar to Hypothetical protein c17, mRNA sequence

TCACGGTCGCAACGTACCACATCCACACGGTGGCGACTAACCAGTGTTGTTCCGCCGTCGTTCAATCCAT

CGACGCCCCCGTCGAGACGGTTTGGTCGGTGGTTCGTCGTTTCGATAACCCTCAAGCTTACAAACACTTT

TTAAAGAGCTGCCAAGTCATCGTCGGCGACGGCGACGTGGGTACACTCCGGGAGGTCCACGTTATTTCGG

GTCTACCCGCCGCGTGCAGCACGGAGAGACTCGATATCCTCGACGACGAACGCCACGTGCTTAGCTTCAG

CGTGGTTGGCGGCGATCATCGGTTGAATAATTACAGGTCAGTGACGACGCTACACGCTTCGCCGAATGGG

AAAGGGACAGTTGTCGTCGAATCATTCGTCGTTGATATACCGCCCGGTAACACTAGAGACGAGACGTGTA

CTTTTGTTGATACCATTGTACGTTGCAACCTTTAATCTTTGGCTCAAATGGCTGAAAATATGGCTAGAAG

AGATCAAAAATGAAGAAATCAAATGTCTTAATTCTTTTAAAAGAATCCCTTTTTTTTGTTTTCGATATGG

GATCGAAGGAAGGAGCTCAGAAATTAATGGAAGATCAACGAATTAAAGCTTGGCTCAAAAAAAAAA

>EE332608.1 EST00104 Jute Plasmid library Corchorus capsularis cDNA similar to Putative senescence-associated protein (Pisum sativum), mRNA sequence

TGATTCTACCCGCCGCTCGGTGGGAATTACGCTTCAAGGAGGCCTGCACGGCTCATCCGCCGCACGGACT

ACTCCTACGACACGTGCCCTTGGGGGCCAAAGGCCCCTACTGCGGGTCGGCAATCGGGCGGCGGGCGCAT

GCGTCGCTTCTAGCCCGGATTCTGACTTAGAGGCGTTCAGTCATAATCCAGCGCACGGTAGCTTCGCGCC

ACTGGCTTTTCAACCAAGCGCGATGACCAATTGTGCGAATCAACGGTTCCTCTCGTACTAGGTTGAATA

>EE332595.1 EST00091 Jute Plasmid library Corchorus capsularis cDNA similar to Metallothionein-like, mRNA sequence

TATAGTTCATTAAGATGAGACAAGGCCAAAGGCATGGCCATGGCCATTACATCACAGTATATAAGGAGAC

ACAAACATGACAGACAGGACATGAAAACACAGACACCAACAGAAACAGGAAAGACACAAATTATTCAAAC

CCAAAATAAGATTTTAGGTCCCAAATATTGCCCATGTTTCTGCAACATGGGATCATTTGTTTCATTTCTA

TATATATGTTTGCTTAATGACCACCGCATGTGCAGTTTGTGCAGCTGCAGCTAGATCCACACTTGCACTT

TGTCGTTCTCTGCTGGAGGAGCCTCCATCACCACGGTGGCAACGTAGCTCTTCTCAGTCTCAATGATCAT

GGTGTTTCCCTTCTTGGTGCACTGGCTCTTGTCAGCGCAGTCGCAGTCTCCGCACTTGTCCGACATGGTG

AATTGAGGAAGGTTTTTTGAAAGAAGTTGGAAGAGCTTTTGGGTTATTGAAGATTGA

>EE332593.1 EST00089 Jute Plasmid library Corchorus capsularis cDNA similar to Tobacco chloroplast 23S and 4.5S rRNA genes, mRNA sequence

AAGGGTCCTATCAATGCTGTAACCCCCAAACATGGGGGAGATAANACCGATCTGTCTCACGACGTTCTGA

CCTGGGTCACGTACCGCTTTAATGGGCTGAACAGCCCAGACCTTGGAGCATACTGCGGCCCCGAATGGCC

CGGGGCCGACTTCGAGGTGCCTGAGCTTCCGGTCGATGTGAGCTCTTGGGGAAGATCACCCTGTTATCTC

AAGAGCAACTTTTATCCGTTGAGCGACGGCCCTTCGACTCGGCACCGGTCGGATCACTAAGGTCGACTTT

CGTCCCTGCTCGACTGGCGGGTCTTGCAATCAAGCTGCCTTCTGCCTTTGCACTCGAGGGCCAATCTCCG

TGCGGCCCGAGGAAACCTTTGCACGCCTCCGTTACCTTTTGGGAGGCCTAGGCCCCATAGAAACTGCCTA

CCTGAGACTGTCCCTTGGCCCGTAGGTCGTGACACAAGGGGNTAGAGGTNCNTAANNCANGTNNATNGCT

TGTCTACNTNGNGNGAAANTTNCTTANCNCACATCACAATTCCAAACNANTANCNNNCNGCCCGGNAANC

GNAAANNNTATANTCCTGNCGNCGCTCTAATGGAGNCGAGCTAGNTGNNGTTANTTGCCTTNTNGCTCAN

GGANGGCTTTAGAANCNNGAAGNNCTGANANGNCGGNANCANNANTAANGNCGCACNNCNAGGCAGANNN

AATTTGTATANANGGGGGNTNATCCANNNTNNTCCGNTGACNGAATAAAGANNCANGNANTTNANGNNGC

GGANAGCGTNTNANCTNANGCNANGCGGGANATNTGTNAATCTCNN

>EE332592.1 EST00088 Jute Plasmid library Corchorus capsularis cDNA similar to At2g39050/T7F6.22 (Arabidopsis thaliana), mRNA sequence

TGGCAAATATGCAGGCCTTCAGAGAAGCCTGCAGCAGATGCTTTGTTATTCAATTTTAACACAGTACGAT

TACACATACATTATTCATAACAAGTCCATAACATAAAGTAGTAGAAGACCATAATTCAACTCATAGTAAT

AGTAGGCCTCTCTACACTCAACATATAAAGAGAATGCATAGGGACACTACTAAGGGTATATTAGCAAGCG

CGATACAGCTTCTGGAACTCACTCGCGTGCTCCATATCACGAGCATCAATAGTGTTCAATCTTCCATCTC

TGGTTATCTCCCTTATTCCACTCCCACACCCCAACAGTAGATCCATTTTGCAATCTACCGTTATCATCTT

TGAAGGCATCCACATTCAACTTTATGTTATTAACCATTCTTAGAGCAAAGTAACCTTCACCTGTGTCCTT

GCTCTCAGTCCGTAATATGGATTCATCCAGATGGTTTGCCTTGTAAGGAGTCAATTGCAAAGGTTGAGAA

GCTCCATTGGCATGCTTAATGGCTTGACCAGGGCTTTGTTAACCAAAGAAAAGCTGGGGAAACCTGCTGA

ATCTTTCACTTGATTGCTGAATTTCTCATCCTTGTACCATTGCTGGTAGGGATCAGAGTGAGCAGCTGGA

GCAAGAACAACCTTGCCACCTCTTACAGTAAGAAGAAGATGGNGATCAGCTTTGCATAGACTTTGAAAGT

AGGTTAGAGGCNA

>EE332591.1 EST00087 Jute Plasmid library Corchorus capsularis cDNA similar to BURP domain containing protein, mRNA sequence

TAACAAAATCAATGTTTTGAGACTCCATAGCATGGATTCTGATCATGCCCATCATCATCTCATGGTTGGG

GTAACTAATATTCATCCATCGTCTCATCACATCCACATGGATCCTTCATTGAGGGTGTTTTTCACCTTGA

ATGAGCTTAGAGTTGGGAAGACAATACCCATATATTTCCCTAAGAATGACCCTTCAATAGGGAACCCAAT

AACTGTCTCCTTCCCAAGAGGGGATCTTTCTGCTGCTCCGAGTTTCTTGCCCACAGAAGAAGCTGATTCC

ATTCCATTTACTCAAGAGGAACTCCCAAACATCCTTCAACGTTTCTCATTCTCTCGAAA

>EE332590.1 EST00086 Jute Plasmid library Corchorus capsularis cDNA similar to Type II keratin E1, mRNA sequence

GGAATCATGTCATATCTGTTTCCTGTGTGAAATTGTTATCCGCTCAGGATTCCACACGACATACGAGCCG

GAGGCATGGAGTGTGAAGCCTGGGGTGCCTAATGAGTGAGCTAACTCACATTAATTGCGTTGCGCTGACT

GCCCGCTTTCCAGTCGGGAAACCTGGCGTGCCAGCTGCATTAATGAATCGGCCAACGCGCGGCGAGAGGC

GGTTTGCGTAGTGGGCGCTCTTCCGCTTCCTCGCTCACTGACTCGCTGCGCTCGGTCGTTCGGCTGCGAG

CGAGCGGTATCACCTCACTCGAAGGCGGTAATGGGGNTATCCACAGAAGCAGGGNATAACGCAAGAAAGA

ACATGTGAGCAAAAGGCCACCAAAAGGCCAGGAACCGTAGAAAGGCCGCGTTGCTGGGGTTTTTCCATAG

GCTCCGCCCGCCTGACGAGCATCACAAAAATCCACGCTCAAGGCGGAGGTGGGGAAGCCCGACAGGACTA

TATAGATACCGGGCGTTTCGCCCTGGAAGCTCCCGCGTGCGCTCTCCTGTTCCGAGCCTGCCGCTTACCG

GATACCTGTCGGGCTTTCTCCCTTCAGGAAGCGTGGCGCTTTCTCATAGCTCACGCTGTAGGTATCTCAG

TTCGGGGTAGGTC

>EE332589.1 EST00085 Jute Plasmid library Corchorus capsularis cDNA similar to N. plumbaginifolia chloroplast 16SrDNA, trnV and trnI genes for 16S ribosomal RNA, transfer RNA-Val and transfer RNA-Ile (5'exon), mRNA sequence

TCCCCGTAGGAGTCTGGGGCGTGTCTCAATCCCAGTGTGGCTGATCATCCTCTCGGACCAGACTACTGAT

CATCGCCTTGGTAAGCTATTGCCTCACCCACTAGCTAATCAAAAGCGAGCGCCTCCTCGGCCGGATTCCT

CCTTTTGCTCCTCAGGCTACGGGGCTATTATCAGCCGTTTCCAGCTGTTGGTCCCCTCCCAAGGGCAGGT

TCTTACGCGTTACTCACCCGATCCGCCACTGTTAAATACCACTTCGCGCCCGACTTGAATGTGTTAATCG

GCCGCGAGCATTAATCCTGAGCCAGGATATCGGGGGC

>EE332594.1 EST00090 Jute Plasmid library Corchorus capsularis cDNA similar to Type II keratin E1, mRNA sequence

TGCCACAAAGGAAGAAGCTCCAGTTGGTTTCACTCCTCCTGAATTGGACCCAAGCACCCCATCACCCATC

TTCGCTGGAAGCACCGGCGGACTGCTGAGGAAGGCCCAAGTGGAAAAATTCNTAATCATGTCATAGCTGT

TTCCTGTGTGAAATTGTTATCCGCTCACAATTCCACACAACATACGACCCGGAAGCATAAAGTGTAAAGC

CTGGGGTGCCTAATGAGTGAGCTAACTCACATTAATTGCCTTGCGCTCACTGCCCGCTTTCCAGTCGGGA

AACCTGTCGTGCCAGCTGCATTAATGAATCGGCCAACGCGCGNGGAGAGGCGGTTTGCGTATTGGGCGCT

CTTCCGCTTCCTCGCTCACTGACTCGCTGCGCTCGGTCGTTCGGCTGCGGCGAGCGGTATCAGCTCACTC

AAAGGCGGTAATACGGTTATCCACAGAATCAGGGGATAACGCAGGAAAGAACATGTGAGCAAAAGGCCAG

CAAAAGGCCAGGAACCGTAAAAAGGCCGCGTTGCTGGCGTTTTTCCATAGGCTCCGCCCCCCTGACGAGC

ATCACAAAAATCGACGCTCAAGTCAAAAGTGGCGAAACCCGACAGGACTATAAAGATACCAGGCGTTTCC

CCCTGGAACCTCCCTCGTGNGCTCTCCTGTTCGACCCTGCCGCTTACCGGAAACCTGTCCCCCTTTTCCC

TTCGGGAAGGTGGAGCTTTNNN

>EE332581.1 EST00077 Jute Plasmid library Corchorus capsularis cDNA similar to S.alba 18S, 5.8S & 25S rRNA genes, mRNA sequence

ACCAAGTGTTGGATTGTTCACCCACCAATAGGGAACGTGAGCTGGGTTTAGACCGTCGTGAGACAGGTTA

GTTTTACCCTACTGATGGCCGCGTCGCAATAGTAATTCAACCTAGTACGAGAGGAACCGTTGATTCGCAC

AATTGGTCATCCCGCTTGGTTGAAAAGCCAGTGGCGCGAAGCTACCGTGCGCTGGATTATGACTGAACGC

CTCTAAGTCAGAATCCGGGCTAGAAGCGACGCATGCGCCCGCCGCCCGATTGCCGACCCGCAGTAGGGGC

CTTTGGCCCCCAAGGGCACGTGTCGTAGGAGTAGTCCGTGCGGCGGATGAGCCGTGCAGGCCTCCTTGAA

GCGTAATTCCCACCGAGCGGCGGGTAGAATCCA

>EE332579.1 EST00075 Jute Plasmid library Corchorus capsularis cDNA similar to Transferase, transferring glycosyl groups (Arabidopsis thaliana), mRNA sequence

TCACGAATGCTCGAGCTACTTGTAAATGTCTGGGCCTACCACAGTGCACGGCGGATGGTATATGTGAATC

CTGAGACTGGCATGTTGCANGAGTNCCACANGTTCANGAGTCGGAGAGGTAAAATGTGGGTCGGGTGGTT

TTCATACAACACCTTATTGGGTATGGATGAATATTTGGCAGAGGAGGCAAATTCTGACCACCCAAGAAAA

CGATGGTTATGGCCTTCCACAGGCGAGGTTGTCTGGCAAGGTGTGCTTGATAGGGAAAGATATCATCGCA

ACAAACAGAANGAGAAACGGAAGCAGCAAAGTAAGGATAAACTATAGAGGATGAGGCATAAACACCTCCG

GAAGGCATTACGGAAGTATGTGAAACCCCTTCCCGAAGAAATGGAAAATTCAAACTCAACTTTATTAAGA

TGATGAGATTAATTTAATGTGTGTTTTGCATTACAATGGGAGTTTTGATAATTCTTTATTTTTTTCACAA

AGGGGGGGAAAATGTCTTGAATGCCTTAATTTTTGTCAGAAGGGGTGCATTAATTATTTTGTGTACATGG

GAGTAATTTTTTGTATGTAGTTAAGCTAACCGCGTGCATCCTTCTGGTCATTGCCATTTGATGAATGGCT

CAGTGGAATACTACAGCGGCCGCGGATCCGTAGTCATGTCATAACTGGTTCCTGTGTGAAATTGTTATCC

GCTCCGATTCCGCGGACGTGACGAGCCGGAAGCNTAAAGTGTGAAGCCGGGGGTGCCTAATGACGGGCTA

ACTCGGGTTATTGCGTTGCGCCCCTGCCCGTTTTCCGTTG

>EE332578.1 EST00074 Jute Plasmid library Corchorus capsularis cDNA similar to Tobacco chloroplast 23S and 4.5S rRNA genes, mRNA sequence

TCGGTCCTCTCGTACTAGGGAAAGGTCCTCTCAATGCTCTAACGCCCACACCGGATATGGACCGAACTGT

CTCACGACGTTCTGAACCCAGCTCACGTACCGCTTTAATGGGCGAACAGCCCAACCCTTGGAACATACTA

CAGCCCCAGGTGGCGAAGAGCCGACATCGAGGTGCCAAACCTTCCCGTCGATGTGAGCTCTTGGGGAAGA

TCAGCCTGTTATCCCTAGAGTAACTTTTATCCGTTGAGCGAAGGCCCTTCCACTCGGCACCGTCGGATCA

CTAAGGCCGACTTTCGTCCCTGCTCGACGGGTGGGTCTTGCAGTCAAGCTCCCTTCTGCCTTTGCACTCG

AGGGCCAATCTCCGTCCGGCCCGAGGAAACCCTTGCACGCCTCCGTTACCTTTTGGGAGGCCTACGCCCC

ATAGAAACTGTCTACCTGAGACTGTCCCTTGGCCCGTAGGTCCTGACA

>EE332577.1 EST00073 Jute Plasmid library Corchorus capsularis cDNA similar to Ribulose bisphosphate carboxylase small chain, chloroplast precursor (RuBisCO small subunit), mRNA sequence

AAGCAGGGTAATCCGTCGGGAGCGGACAACGTGAGTCTACTTCCTTAGCCATTTGCACGGGGGTGAGATC

CCGGAGGTATGAGAGAGTCTCGAACTTCTTCTTCCGGGGTGGTGGCCCTACCTGCGTGCATGGGGGTCTG

CCACCATTGCTGACGATGGAGGTGATGTCTTTGTTGTTCTTGCGGGTGGTGGGGAAAGCAAAGGCAGACT

TGATGGCTACGAATGGGGGGACCATGGAGGGCTGTGCAGTTGAAGCACGGCTAACCGCAGTGGCTGGATG

ACATCATTGACGAACCCATTGTTTAGTACTCCTTATGCTGAAGGATATCGGCCCCGTAATCATAATCATG

TTATAGCTGTTTCCGGGGTGAATCTGGTATGCCTTTTATTTTCCAACAAAAACCATCCGAAGTTTAATCG

AAAACCTGGGGGGCTGTGGGGGGCCCACAACCCTAATTGGTTGGCGCCTGCCCGTTTCATTGGGAAACGG

CGGGCCCCGCATTAAGACCGGCCCCGCGGGTAAATCGGTTCCAATAGGGCCTTCCCTTTATTTTTTTCCC

CGGCCCGGTTTGGGGGGAAAGGGGTCCCTCCTTAAGGGTTACTTTTCCCAAAAAGGATAAAAGGAAAAAT

TGAAAAAGACCCAAAGGGGGGGCTTAATGGGGGGTGGGTTTTATTA

>EE332576.1 EST00072 Jute Plasmid library Corchorus capsularis cDNA similar to Ribulose bisphosphate carboxylase small chain, chloroplast precursor (RuBisCO small subunit), mRNA sequence

CAAGCAAGGAATGCATCCAGAGCGGAGAAGGTAGTCTACTTCCTTAGCCAATTGAACGGGGGTGAGATCC

GGGAGGTATGAGAGAGTCTCGAACTTCTTCTTCCCAAGTGGTGGCCATACCTGCATGCATCGAACTCTGC

CACCATTGCTGACGATGGAGGTGATGTCTTTGTTGTTCTTGCGGGTGGTGGGGAAAGCAGAGGCAGACTT

GAGGCCAACGAATGGGGCGACCAGGGTGGCCTGTGCCGGGGAGGCACGGTTAACAACGGTGGCGGATGAC

ATCATTGCGGAAGCCATTGCTAGTACTCCTTACTAGTAATGTATATGCTAAAGAGA

>EE332575.1 EST00071 Jute Plasmid library Corchorus capsularis cDNA similar to Eucryphia lucida large subunit 26S ribosomal RNA gene, partial sequence, mRNA sequence

TCTTGGTGGTAGTAGCAAATATTCAAATGAGAACTTTGAAGGCCGAAGAGGGGAAAGGTTCCATGTGAAC

GGCACTTGCACATGGGTTAGTCGATCCTAAGAGACGGGGGAAGCCCGTCCGATAGCGCGCAAGCGCGAGC

TTCGAAAGGGAATCGGGTTAAAATTCCTGAACCGGGACGCGGCGGCTGACGGCAACGTTAGGGGGTCCGG

AGACGTCGGCGGGGGCCCCGGGAAGAGTCATCTTTTCTGTTTAACGGCCTGCCA

>EE332580.1 EST00076 Jute Plasmid library Corchorus capsularis cDNA similar to Ribulose bisphosphate carboxylase small chain, chloroplast precursor (RuBisCO small subunit), mRNA sequence

TCTCGAACTTCTTCTTCCCAAGTGGTGGCCATACCTGCATGCATCGAACTCTGCCACCATTGCTGACGAT

GGAGGTGATGTCTTTGTTGTTCTTGCGGGTGGTGGGGAAAGCAGAGGCAGACTTGAGGCCAACGAATGGG

GCGACCATGGAGGCCTGTGCGGGGGAGGCACGGCTAACGACGGTGGCGGATGACATCATTGAGGAAGCCA

TTGCTAGTACTCCTTATGCTAAAGGACA

>EE332567.1 EST00063 Jute Plasmid library Corchorus capsularis cDNA similar to LRRG00134, mRNA sequence

TCTACCCGCCGCTCGGTGGGAATTACGCTTCAAGGAGGCCTGCACGGCTCATCCGCCGCACGGACTACTC

CTACGACACGTGCCCTTGGGGGCCAAAGGCCCCTACTGCGGGTCGGCAATCGGGCGGCGGGCGCATGCGT

CGCTTCTAGCCCGGATTCTGACTTAGAGGCGTTCAGTCATAATCCAGCGCACGGTAGCTTCGCGCCACTG

GCTTTTCAACCAAGCGCGATGACCAATTGTGCAAATCAACGGTTCCTCTCGTACTAGGTTGAATTACTAT

TGCGACGCGGCCATCAGTAGGGTAAAACTAACCTGTCA

>EE332565.1 EST00061 Jute Plasmid library Corchorus capsularis cDNA similar to Type II keratin E1, mRNA sequence

GTAATCATGTCATAGCTGTTTCCTGTGTGAAATTGTTATCCGCTCACAATTCCACACAACATACGAGCCG

GAAGCATAAAGTGTAAAGCCTGGGGTGCCTAATGAGTGAGCTAACTCACATTAATTGCGTTGCGCTCACT

GCCCGCTTTCCAGTCGGGAAACCTGTCGTGCCAGCTGCATTAATGAATCGGCCAACGCGCGGGGAGAGGC

GGTTTGCGTATTGGGCGCTCTTCCGCTTCCTCGCTCACTGACTCGCTGCGCTCGGTCGTTCGGCTGCGGC

GAGCGGTATCAGCTCACTCAAAGGCGGTAATACGGTTATCCACAGAATCAGGGGATAACGCAGGAAAGAA

CATGTGAGCAAAAGGCCAGCAAAAGGCCAGGAACCGTAAAAAGGCCGCGTTGCTGGCGTTTTTCCATAGG

CTCCGCCCCCCTGACGAGCATCACAAAAATCGACGCTCAAGTCAGAGGTGGCGAAACCCGACAGGACTAT

AAAGATACCAGGCGTTTCCCCCTGGAAGCTCCCTCGTGCGCTCTCCTGTTCCGACCCTGCCGCTTACCGG

ATACCTGTCCGCCTTTCTCCCTTCGGGAAGCGTGGCGCTTTCTCATAGCTCACGCTGTAGGTATCTCAGT

TCGGTGTAGGTCGTTCGCTCCAAGCTGGGCTGTGTGCACGAACCCCCGTTCAGCCCGACGCTGCGCCTTA

TCNGGTANCTATCGTCTTGAGTCCACCCGGTAAANACGACTTATCGCCCTGGCAGCANCCCTGGTANCAG

GATNACAAAACGAGGTATGTAGGCGGTGCTACAAANTNTT

>EE332564.1 EST00060 Jute Plasmid library Corchorus capsularis cDNA similar to Ribulose bisphosphate carboxylase small chain F1, chloroplast precursor (RuBisCO small subunit F1), mRNA sequence

GAATTGGAGCACGGATTCGTCTACCGTAAGAACCACAGCTCCCCAGGGTACTACGATGGACGCTACTGGA

CCATGTGGAAACTTCCCATGTTCGGGTGCACCGACTCATCCCAGGTGTTGAAGGAGCTGAGGAGGCCAAG

AAGGAATACCCCGAGTCCTTCATCAGAATCATTGGATTCGACAACAAGCGTCAAGTGCGATGCATCAGTT

TCATTGCCTACAAGCCCCCAAGCTTCACCTCAGCTTAAATGAACAGAGTTGTTTTCCCCACATTGTAGGG

ATTCGGCTTGTTTAAATTGTACTTAGCATTTGTCGAAACCTTTTTACTTTTCCTTTCTTTTTGAGGTATT

GTCCTTTGTTTTTTT

>EE332563.1 EST00059 Jute Plasmid library Corchorus capsularis cDNA similar to Expressed protein, mRNA sequence

TGTCAAGGGCCAAGCAACATGCAAGATTTAAGGTGTTACAGTGCTAGTTATGCAAATTCTGTTGTTCAGG

TTCAGCCAAATCAGCTTGGGAACGAGATTAAGATCAAGAAAAGCAAAAGCAGTTTTGGGTCATCTTCAAA

AAGCTGGAGCTTTAATGATCCTGAGTTGCAGAGGAAGAAAAGGGTTGCAAGTTACAAGGTTTATGCAGTT

GAAGGCAAGATGAAAGGGTCTTTCAGGAAGAGTTTTAGGTGGATCAAAGACACTTATACCCAAGTTGTCT

ATGGATGGAGA

>EE332562.1 EST00058 Jute Plasmid library Corchorus capsularis cDNA similar to Expressed protein, mRNA sequence

TGTCAAGGGCCAAGCAACATGCAAGATTTAAGGTGTTACAGTGCTAGTTATGCAAATTCTGTTGTTCAGG

TTCAGCCAAATCAGCTTGGGAACGAGATTAAGATCAAGAAAAGCAAAAGCAGTTTTGGGTCATCTTCAAA

AAGCTGGAGCTTTAATGATCCTGAGTTGCAGAGGAAGAAAAGGGTTGCAAGTTACAAGGTTTATGCAGTT

GAAGGCAAGATGAAAGGGTCTTTCAGGAAGAGTTTTAGGTGGATCAAAGACACTTATACCCAAGTTGTCT

ATGGATGGAGA

>EE332561.1 EST00057 Jute Plasmid library Corchorus capsularis cDNA similar to Type II keratin E1, mRNA sequence

GTTGTAACGACGGCCAGTGCCAAGCTTGCATGCCTGCAGGTCGACTCTAGAGGATCCCCGGGTACCGAGC

TCGAATTTTCGTAATCATGTCATAGCTGTTTCCTGTGTGAAATTGTTATCCGCTCACAATTCCACACAAC

ATACGAGCCGGAAGCATAAAGTGTAAAGCCTGGGGTGCCTAATGAGTGAGCTAACTCACATTAATTGCGT

TGCGCTCACTGCCCGCTTTCCAGTCGGGAAACCTGTCGTGCCAGCTGCATTAATGAATCGGCCAACGCGC

GGGGAGAGGCGGTTTGCGTATTGGGCGCTCTTCCGCTTCCTCGCTCACTGACTCGCTGCGCTCGGTCGTT

CGGCTGCGGCGAGCGGTATCAGCTCACTCAAAGGCGGTAATACGGTTATCCACAGAATCAGGGGATAACG

CAGGAAAGAACATGTGAGCAAAAGGCCAGCAAAAGGCCAGGAACCGTAAAAAGGCCGCGTTGCTGGCGTT

TTTCCATAGGCTCCGCCCCCCTGACGAGCATCACAAAAATCGACGCTCAAGTCAGAGGTGGCGAAACCCG

ACAGGACTATAAAGATACCAGGCGTTTCCCCCTGGAAGCTCCCTCGTGCGCTCTCCTGTTCCGACCCTGC

CGCTTACCGGATACCTGTCCGCCTTTCTCCCTTCGGGAAGCGTGGCGCTTTCTCATAGCTCACGCTGTAG

GTATCTCAGTTCGGTGTAGGTCGTTCGCTCCAGGCTGGGCTGTGTGCACGAACCCCCGTTCAGCCCGACC

GCTGCGCCTTATCCGGTAGCTATCGTCTTGAGTCCAGCCGGTAAAAACGACTTATCGCCCTGGCAGCAGC

CCTGGTAGCAGGATTACNAAACGAGGTATGTAGGCGGGGCTCAAAATTATTGAATGGTGGCCTACTAA

>EE332566.1 EST00062 Jute Plasmid library Corchorus capsularis cDNA similar to Oxygen evolving enhancer protein 1 precursor, mRNA sequence

TCTCTTGTGGTCTCGGGGGCAAGCGCTGAAGGAGTTCCAAAGAGGCTAACCTTTGATGAAATCCAGAGCA

AAACGTACATGGAAGTAAAGGGATCAGGAACAGCTAACCAGTGCCCAACCATTGAAGGTGGTGTTGACAA

ATTTGCCTTCAAGCCAGGCAAATACTACGCCAAGAAGTTCTGCTTAGAGCCCACTTCCTTCACTGTCAAG

GCAGAGGGTGTTAACAA

>EE332553.1 EST00049 Jute Plasmid library Corchorus capsularis cDNA similar to Putative senescence-associated protein (Pisum sativum), mRNA sequence

ACCAAGTGTTGGATTGTTCACCCACCAATAGGGAACGTGAGCTGGGTTTAGACCGTCGTGAGACAGGTTA

GTTTTACCCTACTGATGGCNGCGTCNCANTAGTANTTCAACCTAGTACGAGAGGAACCGTTGATTCGCAC

NATTGGTCATCGCGCTTGGCTGAAAAGCCAGTGGCGCGAAGCTACCGTGCGCTGGATTATGACNGAACGC

CTCTAAGTCAGAATCCGGGCTAGAAGCGACGCATGCGCCCGCCGCCCGATTGCCGACCCGCAGTANGGGC

CTTTGGCCCCCAAGGGCACGTGTCGTATGANTANTCCGTGCGGCGNATGAACCNTGCAGGCCTCCTTGAA

NCGTAATTCCCACCNAGCGGCNNGTAGA

>EE332551.1 EST00047 Jute Plasmid library Corchorus capsularis cDNA similar to Photosystem I chain II precursor - spinach, mRNA sequence

TACAGGGCCAGCATCAGGGCTGAAGCTGCGGAAGAGACTGCCGCCGCCGCTGCCACAAAGGAAGAAGCTC

CAGTTGGTTTCACTCCTCCTGAATTGGACCCAAGCACCCCATCACCCATCTTCGCTGGAAGCACCGGCGG

ACTGCTGAGGAAGGCCCAAGTGGAA

>EE332550.1 EST00046 Jute Plasmid library Corchorus capsularis cDNA similar to Photosystem I chain II precursor - spinach, mRNA sequence

GGAAGTCCCCACCAAAGAAGCGCCAGTTGGTTTCACTCCTCCTGAATTGGACCCAAGCACCCCATCACCC

ATCTTCGCTGGAAGCACCGGCGGACTGCTGAGGAAGGCCCAAGTGGAAGGAGTTCTACGTCATCACATGG

GAATCGCCAAAAGAACAGATCTTCGAAATGCCCACCGGCGGCGCCGCCATCATGAGGGAAGGACCGAATC

TGTTGAAATTAGCGAGAAAAGAACAGTGTTT

>EE332549.1 EST00045 Jute Plasmid library Corchorus capsularis cDNA similar to Unknown protein, mRNA sequence

TCACAAACTGAAGGAATGCTGTCATATATCTTGGAAGAAATCATGCAACATTATTTATGGATAATGCAAG

TTGTTTGTAAGCACCCCAAAGACAGACAATCAAGTGTAGCCAAATTTCAGGCTAAGAAATGTACAAAGGC

AATCCCTGCGGATCACGATAACCTCACTTCTGAAAGCCGATGAAGCAACAATGAACCTAGTTCCCTCTAT

AGTATCCTCATTTAATACATTAAAGCCTGCTCTCGAAATTAGGTTAATGCAAGGGATGGAGGATTACCTG

CGGGACATCTTTGCCTGCGAGGCCGTCTAGTTGTCTTCCCCCATCCAGTAAGGCAG

>EE332548.1 EST00044 Jute Plasmid library Corchorus capsularis cDNA similar to S.alba 18S, 5.8S & 25S rRNA genes, mRNA sequence

TCAGGCCTCCTTGAAGCGTAATTCCCACCGAGCGGCGGGTAGATCCAGCGCACGGTAGCTTCGCGCCACT

GGCTTTTCAACCAAGCGCGATGACCAATTGTGCGAATCAACGGTTCCTCTCGTACTAGGTTGAATTACTA

TTGCGACGCGGCCATCAGTAGGGTAAAACTAACCTGTCTGACGACGGTCTAAACCCAGCTCACGTTCCCT

ATTGGTGGGTGAACAATCCAACACTTGGT

>EE332547.1 EST00043 Jute Plasmid library Corchorus capsularis cDNA similar to Ribulose bisphosphate carboxylase small chain, chloroplast precursor (RuBisCO small subunit), mRNA sequence

CAAGCAAGGAATCCATCCAGAGCGGAGAAGGTAATCTACTTCCTTAGCCAATTGCACAGGGCTGAGATCA

GGGTGGTATGAAAGAGTCTCGAACTTCTTCTTTCCAAGTGGTGGCCATACCTGCATGCATCGCACTTTGC

CACCATCGCTTACAATAGAAGTAATGTCCTTGTTGTTCTTGCGGGTGGTGGGGAAAGCAGAGGCAGACTT

GAGA

>EE332552.1 EST00048 Jute Plasmid library Corchorus capsularis cDNA similar to Metallothionein-like, mRNA sequence

TATATTTCATTAAGATGAGACAAGGCCAAAGGCATGGCCATGGCCATTACATCACAGTATATAAGGAGAC

ACAAACNTGACANACAGGACATGAAAACACAGACACCAACAGAAACNGGAAAGACACAAATTATTCNNAC

CCAAAATAAGATTTTAGGTCCCAAATATTGCCCATGTTTCTGCAACATGGGATCATTTGTTTCATTTCTA

TATATATGTTTGCTTAATGACCACCGCNTGTGCAGTTTGTGCAGCTGCAGCTAGATCCACACTTGCACTT

TGTCGTTCTCTGCTGGAGGAGCCTCCATCACCACGGTGGCAACGTAGCTCTTCTCAGTCTCAATGATCAT

GGTGTTTCCCTTCTTGGTGCACTGGCTCTTGTCAGCGCAGTCGCAGTCTCCGCACTTGTCCGANATGGTG

AATTGAGGAAGGTTTTTTGAAAGAAGTTGGAAGAGCTTTTGGGTTATTGAAGATTGA

>EE332539.1 EST00035 Jute Plasmid library Corchorus capsularis cDNA similar to S.alba 18S, 5.8S & 25S rRNA genes, mRNA sequence

TAAGGATTCTACCCGCCGCTCGGTGGGAATTACGCTTCAAGGAGGCCTGCACGGCTCATCCGCCGCACGG

ACTACTCCTACGACACGTGCCCTTGGGGGCCAAAGGCCCCTACTGCGGGTCGGCAATCGGGCGGCGGGCG

CATGTGTCGCTTCTAGCCCGGATTCTGACTTAGAGGCGTTCAGTCATAATCCAGCGCACGGTAGCTTCGC

GCCACTGGCTTTTCAACCAAGCGCGATGACCAATTGTGCGAATCAACGGTTCCTCTCGTACTAGGTTGAA

TTACTATTGCGACGCGGCCATCAGTAGGGTAAAACTAACCTGTCTCACGACGGTCTAAACCCAGCTCACG

TTCCCTATTGGTGGGTGAACAATCCAACACCTG

>EE332537.1 EST00033 Jute Plasmid library Corchorus capsularis cDNA similar to Type II keratin E1, mRNA sequence

TGTCATAGCTGTTTCCTGTGTGAAATTGTTATCCGCTCACAATTCCACACAACATACGAGCCGGAAGCAT

AAAGTGTAAAGCCTGGGGTGCCTAATGAGTGAGCTAACTCACATTAATTGCGTTGCGCTCACTGCCCGCT

TTCCAGTCGGGAAACCTGTCGTGCCAGCTGCATTAATGAATCGGCCAACGCGCGGGGAGAGGCGGTTTGC

GTATTGGGCGCTCTTCCGCTTCCTCGCTCACTGACTCGCTGCGCTCGGTCGTTCGGCTGCGGCGAGCGGT

ATCAGCTCACTCAAAGGCGGTAATACGGTTATCCACAGAATCAGGGGATAACGCAGGAAAGAACATGTGA

GCAAAAGGCCAGCAAAAGGCCAGGAACCGTAAAAAGGCCGCGTTGCTGGCGTTTTTCCATAGGCTCCGCC

CCCCTGACGAGCATCACAAAAATCGACGCTCAAGTCAGAGGTGGCGAAACCCGACAGGACTATAAAGATA

CCAGGCGTTTCCCCCTGGAAGCTCCCTCGTGCGCTCTCCTGTTCCGACCCTGCCGCTTACCGGATACCTG

TCCGCCTTTCTCCCTTCGGGAAGCGTGGCGCTTTCTCATAGCTCACGCTGTAGGTATCTCAGTTCGGTGT

AGGTCGTTCGCTCCAGCTGGGCTGTGTGCACGAACCCCCCGTTCAGCCCGACCGCTGCGCCTTATCNGGT

ANCTATCGTCTTGAGTCCACCCGGTAAAAACGACTTATCGCCCTGGCAGCANCCCTGGTAACAGGATTAN

CAAANCGAGGTTGTAG

>EE332536.1 EST00032 Jute Plasmid library Corchorus capsularis cDNA similar to Type II keratin E1, mRNA sequence

ATGTCATAGCTGTTTCCTGTGTGAAATTGTTATCCGCTCNCAATTCCACACANNATACNAGNCGGAAGCA

TAAAGTGTAAAGCCTGNGGTGCCTAATGAGTGAGCTAACTCACATTANTTGNGTTGCGCTCNCTGCCCGC

TTTCCAGTCGGGAAACCTGTCGTGCCANCTGCATTAATGAATNNGNCAACNCGCGGGGTAGAGGCGGTTT

GCGTATTGGGCGCTCTTCCGCTTCCTCGCTCACTGACTCGCTGCGCTCGGTCGTTCGGCTGCGGCNAGNG

GTATCAGCTCACTCAAAGGCGGTAATACNGTTATCCACAGAATCAGGGGATAACNCANGAAAGAACATGT

GAGCNAAANGCCANNNNAANGCCAGGAACCGTAAAAAGGCCGCGATTGCTGGCGTTTTTCCATANGNTCC

GCCCCCCTGACGAGCATCACANAAATCNANNCTCAAGTCAGAGGTGGCGAAACCCGACNNGACTATAAAG

ATACCNGGCGTTTCCCCCTGGAAGCTCCCTCGTGCGCTCTCCTGTTCCGACNCTGCCNCTTACCGGATAC

NTGTCCGCCTTTCTCNCTTCAGGAAGCGTGGNNCTTTCNCATAGNTCACGCTGTANGTATCTCANTTCGG

NGAACGTCNTT

>EE332535.1 EST00031 Jute Plasmid library Corchorus capsularis cDNA similar to LRRG00134, mRNA sequence

TTTCTACCCGCCGCTCGGTGGGAATTACGCTTCAAGGAGGCCTGCACGGCTCATCCGCCGCACGGACTAC

TCCTACGACACGTGCCCTTGGGGGCCAAAGGCCCCTACTGCGGGTCGGCAATCGGGCGGCGGGCGCATGC

GTCGCTTCTAGCCCGGATTCTGACTTAGAGGCGTTCAGTCATAATCCAGCGCACGGTAGCTTCGCGCCAC

TGGCTTTTCAACCAAGCGCGATGACCAATTGTGCGAATCAACGGTTCCTCTCGTACTAGGTTGAATCACT

ATTGCGACGCGGCCATCAGTAGGGTAAAACTAACCTGTCTCACGACGGTCTAATCCCAGCTCACGTTCCC

TATTGGTGGGTGAACAATCA

>EE332534.1 EST00030 Jute Plasmid library Corchorus capsularis cDNA similar to Predicted protein, mRNA sequence

TTGCAGTTAAAGCACACCAGTTCTTCACAGAGGAAAGCTGGGATACCTTTAAGCAGATTTTTGATACCTA

CATGCTGGAAGCGTCAAAGGAATGGATTGAGACATATGGGGGCAGTTCACTCCTTGAAACTGTGTACAAG

GCTCTTGATGAGGTGGTGAAACTCTCAGAGTGCGAAATTTACAGCTATAATCCAGACTCTGATGCCGATC

CCTTTCTGGAGAAAGGAGCAATATGGTCTTTCAGTTTCTTCTTCTACAATAGAAAGCTTAAGCGAGTAGT

AAGTTTCAGCTTTTCTTGCCTAAGCAATCCGGTGGCCGATGGATTTCTTGCAGACAACTTATGTTACGAG

GAAGATGGTGAGATATTTGATAACATGGACATATGACCACAAACGTGTACATACTGCTTGTACAATGTGT

CAAAACCCTTTGACCATTGTCTGTAGTTACATTACCGTAGTAGGTATCTTGCTCACATGAACTGAGTAGC

TTAGCTTTTCTACTTTGCAGTGCATATGTAGTCGGTAGTTGCTTCCTACATAATAGGAGTATAAATAATG

ATGTCTGTAAGAATGTGTCGAATGTGAGTTCGGATTCATTAAGGTTGAAGACATTCCTTACTACAATGTT

TGTTTCCTATGCTTATACACTTATATATGTTTGAACAAAAAAAAAAATTC

>EE332533.1 EST00029 Jute Plasmid library Corchorus capsularis cDNA similar to Ribulose bisphosphate carboxylase small chain, chloroplast precursor (RuBisCO small subunit), mRNA sequence

CAAGCAAGGAATCCATCCAGAGCGGAGAAGGTAGTCTACTTCCTTAGCCAATTGCACGGGGGTGAGATCC

GGGAGGTATGAGAGAGTCTCGAACTTCTTCTTCCCAAGTGGTGGCCATACCTGCATGCATCGAACTCTGC

CACCATTGCTGACGATGGAGGTGATGTCTTTGTTGTTCTTGCGGGTGGTGGGGAAAGCAGAGGCAGACTT

GAGGCCAACGAATGGGGCGACCAGGGTGGCCTGTGCCGGGGAGGCACGGTTAACAACGGTGGCGGATGAC

ATCATTGCGGAAGCCATTGCTAGTACTCCTTACTAGTAATGTATATGCTAAAGAGTGAGAG

>EE332538.1 EST00034 Jute Plasmid library Corchorus capsularis cDNA similar to Protein transporter (Arabidopsis thaliana), mRNA sequence

TAATTAGGATCCAGAAAAGGACCATTCATTGTAAGCAATGCATACAATTGCAAATATCTCTGATTCACTG

ATGGTAAATCTGTTTCGGAGTGCCATAAGATTTTACTTAAAAGAAGTCTTCAAGAAAACATCCGACATTC

AATTCATAAGAAGAGGTCTTTCACCAACTCAACCAATCACAACGAAGAAGGTCTCTTGAAATATGGAGAG

ACATGGAAATCATAAATGAAAACACGAAACATAATCAACCCAAAAAATTTTCCATTTTCATTTAAAATGC

CTAGAACTGAACAGAGAGAGATAAGCAAACTTGTTTAAATGCTTGATATTGTATTCCCATCCCTTGTGAT

TAATTGTAGTTGTCTTCATGCATAGTTCAACCTGAACATGTCATTCAACACATAGTAGCTCCCCTGTGGT

GTTGGCATCAAATGGAACATCTGGCTGAACTTGAGAGCATGCTGTTCGCCGGCGAGCTGGAGATTACCAC

TGACGAAGACGAGCATGCCACCGGCGGCGCCGGAAGGCA

>EE332525.1 EST00021 Jute Plasmid library Corchorus capsularis cDNA similar to LRRG00134, mRNA sequence

TCAAAGGATTCTACCCGCCGCTCGGTGGGAATTACGCTTCAAGGAGGCCTGCACGGCTCATCCGCCGCAC

GGACTACTCCTACGACACGTGCCCTTGGGGGCCAAAGGCCCCTACTGCGGGTCGGCAATCGGGCGGCGGG

CGCATGCGTCGCTTCTAGCCCGGATTCTGACTTAGAGGCGTTCAGTCATAATCCAGCGCACGGTAGCTTC

GCGCCACTGGCTTTTCAACCAAGCGCGATGACCAATTGTGCGAATCAACGGTTCCTCTCGTACTAGGTTG

AATTACTATTGCGACGCGGCCATCAGTAGGGTAAAACTAACCTGTCTCACGACGGTCTAAACA

>EE332523.1 EST00019 Jute Plasmid library Corchorus capsularis cDNA similar to Proline-rich protein, mRNA sequence

TAAACATTAATTAAACATTACATTAGGCCGAGGTATAAAATTAACCCACAAGCCTAGTTATTTAAAACAT

GAACAAATATCACACAATGTCGGAAGTCTTTGTAATAAAGACTCCCTAAAAATATGTAAAAATGGAAACC

CTATCCCAACTTCAAATCGATCCAAACCACTAAACAAAAATACATTGAATATTTGAAACTACAACACCAA

TTGAAGAAATTATACTAATGATAAGCAAATAAACTTAGCCGGAGAAATTGTTAACGTCGGCGGCCGGCCG

CTTCCTTATTCACACTTGAAGCCTGGAGGAATTTCCTTTTGGCATGCACTAAGGAGTAAGCTAAGAGTAA

CAGGAATGTTGAGGTTGATTCCAAGTACATTGGCTTTAATGGCAGTGCAAAGGCAAAGGGCAGCTTCGGC

ATCAGCCAAGCCTGAGAGCAAAGAGCGGCCGC

>EE332522.1 EST00018 Jute Plasmid library Corchorus capsularis cDNA similar to Ribulose bisphosphate carboxylase small chain F1, chloroplast precursor (RuBisCO small subunit F1), mRNA sequence

GAATTGGAGCACGGATTCGTCTACCGTAAGAACCACAGCTCCCCAGGGTACTACGATGGACGCTACTGGA

CCATGTGGAAACTTCCCATGTTCGGGTGCACCGACTCATCCCAGGTGTTGAAGGAGCTTGAGGAGGCCAA

GAAGGAATACCCCGAGTCCTTCATCAGAATCATTGGATTCGACAACAAGCGTCAAGTGCAATGCATCAGT

TTCATTGCCTACAAGCCCCCAAGCTTCACCTCAGCTTAAATGAACAGAGTTGTTTTCCCCACATTGTAGG

GATTCGGCTTGTTTAAATTGTACTTAGCATTTGTCGAAACCTTTTTACTTTTCCTTTCTTTTTGAGGTAT

TGTCCTTTGTTTTTTT

>EE332521.1 EST00017 Jute Plasmid library Corchorus capsularis cDNA similar to Light harvesting chlorophyll a/b-binding protein precursor, mRNA sequence

TCGCCGGTCTCTCAGCTGACCCCGAGACTTTCGCTAAGAACCGTGAGCTGGAGGTCATCCACTGCAGATG

GGCCATGTTGGGAGCTCTCGGCTGCGTTTTCCCGGAGCTTTTGGCCAGGAACGGCGTCAAGTTCGGCGAA

GCCGTGTGGTTCAAGGCCGGAGCCCAGATCTTCAGCGAAGGAGGACTCGACTACTTGGGAAACCCAAGCT

TGATCCACGCTCAGAGCATCTTGGCCATCTGGGCTTGCCAAGTTATCTTGATGGGAGCCGTCGAGGGTTA

CCGTATTGCCGTGGACCCCTCGGTGAGGTGGTTGACCCATTACCCAGGTGGCAGCTTCGACCCATTGGGT

CTTGCCGATGACCCAGAGGCCTTCGCTGAGCTCAAGGTTAAGGAAATCAAGAACGGCCGATTGGCTATGT

TCTCCATGTTTGGATTCTTCGTCCAAGCCATTGTCACCGGAAAGGGACCATTGGAGAACTTGGCCGACCA

CCTTGCTGACCCAGTCAACAACAACGCCTGGGCCTACGCCACCAACTTTGTCCCCGGAAAGTGAGCTTAG

AAGTTAGAATGAGATGAGAGAGATGTATTTGTCTTTTATGTTTTCTTTGTTTGGTCTGATGTACATTTGT

TGCAAATCAATGACCATAATGTGAATTTTAGTGTGCTCCAAAAAAAAAAA

>EE332520.1 EST00016 Jute Plasmid library Corchorus capsularis cDNA similar to Tomato 25S ribosomal RNA gene, mRNA sequence

TCCCAACTGCGCGCTAACCTAGAACCCACAAAGGGTGTTGGTCGATTAAGACAGCAGGACGGTGGTCATG

GAAGTCGAAATCCGCTAAGGAGTGTGTAACAACTCACCTGCCGAATCAACTAGCCCCGAAAATGGATGGC

GCTTAAGCGCGCGACCTATACCCGGCCGTCGGGGCAAGAGCCAGGCCTCGATGAGTAGGAGGGCGCGGCG

GTCGCCGTAAAACCCGGGGCGTGAGCCCGGGCGGAGCGGCCGTCGGTGCAGATCTTGGTGGTAGTAGCAA

ATATTCAAATGAGAACTTTGAAGGCCGAAGAGGGGAAAGGTTCCA

>EE332519.1 EST00015 Jute Plasmid library Corchorus capsularis cDNA similar to Oryza sativa (japonica cultivar-group) cDNA clone: 006-209-G02, mRNA sequence

CGAATTGGGCGCCCTTCCGCTTCCGGGCTCCCTGACCCGCTCCGCTCGGTCGTTCAAGGGGCGGGGAGCG

GTATCAGCTCACTCAAAGGCCGGAGTACGGTTAACCACAGCAAACCGGGGGTAATGTATTGAAAGATTAA

GTAAACCCCCCGGCCAAAATAAGGCCAAGCATCCGATTAATGGCCAGGGTTTGCCTGGCTTTTTTTCCCT

AGGCCCCCGCCCCCTTGATTGATCCATTGAAATAATATGGAATATAAAGTACAGAGGCCGCGTAACCCGG

GGGTGATTAATAAGGGTTCCCTCCTGGATACCCACCGAGCAGCCTCCTGGGCGCACCCTGGCCCTTA

>EE332524.1 EST00020 Jute Plasmid library Corchorus capsularis cDNA similar to At2g39050/T7F6.22 (Arabidopsis thaliana), mRNA sequence

TCCATGGTTCATGGAAATGTGCCATCTGCAACTCACCATACTTCTCATGCTGATCTTGGTGAACTTGCCT

CTAAACCTACTTTCAAAGTCTATTGCAAAGCTGATCACCATCTTCTTCTTACTGTAAGAGGTGGCAAGGT

TGTTCTTGCTCCAGCTGCTCACTCTGATCCCTACCAGCAATGGTACAAGGATGAGAAATTCAGCAATCAA

GTGAAAGATTCAGCAGGTTTCCCCAGCTTTTCTTTGGTTAACAAAGCCACTGGTCAAGCCATTAAGCATG

CCAATGGAGCTTCTCAACCTTTGCAATTGACTCCTTACAAGGCAAACCATCTGGATGAATCCATATTACG

GACTGAGAGCAAGGACACAGGTGAAGGTTACTTTGCTCTAAGAATGGTTAATAACATAAAGTTGAATGTG

GATGCCTTCAAAGATGATAACGGTAGATTGCAAAATGGATCTACTGTTGGGGTGTGGGAGTGGAATAAGG

GAGATAACCAGAGATGGAAGATTGAACACTATTGATGCTCGTGATATGGAGCACGCGAGTGAGTTCCAGA

AGCTGTATCGCGCTTGCTAATATACCCTTAGTAGTGTCCCTATGCATTCTCTTTATATGTTGAGTGTAGA

GAGGCCCTACTATTACTATGAGTGNATTATGGTCTTCTACTACTTTATGTTATGGACTGGTTATGAANAA

NGTANGNGTAATCGTACTGGGTTNAAATTGAATANCNAAGC

>EE332682.1 EST00178 Jute Plasmid library Corchorus capsularis cDNA, mRNA sequence

TCTTACGGTGGATACCTAGGCACCCAGAGACGAGGAAGGGCGTAGTAAGCGACGAAATGCTTCGGGGAGT

TGAAAATAAGCGTAGATCCGGAGATTCCCGAATAGGTCAACCTTTCGAACTGCTGCTGAATCCATGGGCA

GGCAAGAGACAACCTGGCGAACTGAAACATCTTAGTAGCCGGAGGAAAAGAAAGCAAAAGCGATTCCCGT

AGTAGCGGCGAGCGAAATGGGAGCAGCCTAAACCGTGAAAGCGGGGTTGTGGGAGAGCAATAAAAGA

>EE332681.1 EST00177 Jute Plasmid library Corchorus capsularis cDNA, mRNA sequence

AACGCTCCATCCCACGCTCCTTCGTTCTTTATGACCTCTATCATCTCCTCCTCGGCTCCTCCCTTGCAAC

TAACTATTTCCACAGCCTACCTTACAATCTCCCCTACGTCGCCTGGCCGCTCTACTGGGCCTGCCAAGGC

TGCGTCCTAACGGGCCTCTGGGTCATAGCCCACGAATGCGGTCATCATGCCTTCAGCGACCACTAATGGC

TTGACGACGCCGTCGGCCTTGTCCTACACTCATCTCTCCTCGTTCCCTATTTCTCCTGGAAGTACAGCCA

GATCTCGC

>EE332680.1 EST00176 Jute Plasmid library Corchorus capsularis cDNA, mRNA sequence

TTGATGGGAATTACGCTTCAAGGCGGCCTGCACGGCTCGTCCGCCGCACGAGCTACTCCTACGACACGTG

CCTTTGGGGGCCGAGGGCCCTACTGCTGGTCGGCAAACAGGCGACGGGCACATGCGTCGCTTCTAGCCCG

GATTCTGACTTAGAGGCGTTCAGTCATCCTCCAGCGCACGGTAGCTTCGCGCCACTGGCTTTTCAACCAA

GCGCGAGTACCAATTGTGCGAATCAACGGTTCCTCTCGTACTAGGTGATAC

>EE332672.1 EST00168 Jute Plasmid library Corchorus capsularis cDNA, mRNA sequence

ATGGGAGCGGGTGGCAGAAGCAGGAGTCAGCTCGGCCGCGCCGGCGGCGGCGCAGCCTTCAAGCGCTCGC

CGACGGACAAGCCGCCATTCACGCTCGGTCAGATCAAGAAAGCGATCCCGCCTCACTGCTTCCAGCGCTC

CATCATCAAGTCCTTCTCCTACGTGGTTCATGACCTGGTCCTCATCGCCGCCCTGCTGTACTC

>EE332671.1 EST00167 Jute Plasmid library Corchorus capsularis cDNA, mRNA sequence

CGCTGTATGAACAAATTCAGACTGTGACACTGCGAATGGCTCATTAAATCAGTTATAGTTTGTTTGATGG

TATCTACTACTCGGATAACCGTAGTAATTCTAGAGCTAATACGTGCAACAAACCCCGACTTCTGGAAGGG

ATGCATCTATTAGATAAAAGGTCGACGCGGGCTTTGCCCGTTGCTGCGATGATTCATGATAACTCGACGG

ATCGCACGGCCATCGTGCCGGCGACTCATCATTGAAATTTG

>EE332670.1 EST00166 Jute Plasmid library Corchorus capsularis cDNA, mRNA sequence

GCATGCTACATGCAAGTCGGACGGGAAGTGGTGTTTCCAGTGGCGGACGGGTGAGTAACGCGTAAGAACC

TGCCCTTGGGAGGGGAACAACAGCTGGAATACGGCTGCTAATACCCCGTAGGCTGAGGAGCAAAAGGAGG

AATCCGCCCGAGGAGGGGCTCGCGTCTGATTAGCTAGTTGGTGAGGCAATAGCTTACCAAGGCGATGATC

AGTAGCTGGTCCGAGAGGATGATCAGCCACACTGGGCATGAGACACGGCCCAGACTCCTACGGGAGGCAG

TGGA

>EE332669.1 EST00165 Jute Plasmid library Corchorus capsularis cDNA, mRNA sequence

TACTTAGATGCTTTCAGCAGTTATCCGCTCCGCACTTGGCTACCCAGCGTTTACCGTAGGCACGATAACT

GGTACACCAGAGGTGTGTCCTTCCCGGTCCTCTCGTACTAGGGAAAGGTCCTCTCAATGCTCTAACGCCC

ACACCGGATATGGACCGAACTGTCTCACGACGTTCTTGAACCCAGCTCACGTCACAGCTTTAATGGGCGA

ACAGCCCAACCCTTGGAAATACCTACAGCCCCAGG

>EE332668.1 EST00164 Jute Plasmid library Corchorus capsularis cDNA, mRNA sequence

ATGGGAGCTGGTGGCAGAATGCAAGTCTCTCCTCCCTCCAGCTCCCCCGGAACCAACACCCTCAAACGCG

TCCCTGCTGAGACACCACCATTCACTCTCGGAGACCTCAAGAAAGCAATCCCACCTCACTGCTTCAAACG

CTACACCCTTTCTCCTTCGTTCTTTATCACCTCATCATAGCCTTTCTCCTTTCCTCCCTTGCAACCACCT

ATTTCCACAGCCTCCCTTACAATCTATCGTACCTCGCCTGACCCCTCTACTGGGCCTGCCAAGGCTGCGT

CCTAACAGGCCTCTGGGTCATAGCCCACGAATGTGGTCATCACGCCTTCAGTGACTATCAATGGCTTGAT

GACGCTGTTGGCCTTGTCCTACACACATCTCTACTTGTTCCCTATTTCTCTTGGAAATATAGTCA

>EE332667.1 EST00163 Jute Plasmid library Corchorus capsularis cDNA, mRNA sequence

GTAAGTATGAACAAATTCAGACTGTGAAACTGCGAATGGCTCATTAAATCACTTATAGTTTGTTTGATGG

TATCTACTACTCGGATAACCGTAGTAATTCTAGAGCTAATACGTGCAACAAACCCCGACTTCTGGAAGGG

ATGCATTTATTAGATAAAAGGTCGACGCGGGCTCTGCCCGTTGCTCTGATGATTGATGATAACTCGACGG

ATCGCACGGCCATTGTGCCGGCGACGCATCATTCAAATTTCTGCCCTATCAACTTTCGATGGTAGGATAG

TGGCCTACTATGGTGGTGACGGGTGACGGAGAATTAGGGTTCGATTCCGGAGAGGGAGCCTGAGAAACGG

CTACCACATCCAAGGAAGGCAATTAGCGGCCG

>EE332666.1 EST00162 Jute Plasmid library Corchorus capsularis cDNA, mRNA sequence

CCCATCTTCTGCCCTCGGCATCAACATCCTCCAATGGTGGCATGTCAGCATCAGCTTCAGCTTCACCAGC

ATCCTCATCAATGCTCATCCCAATTTCAACATTCTGTGGATTCTGTTGCCGAAGTGTTGGGGTCATCCAG

GCTGANGCCAGAGGTGAGAATTTCAGTCTCAAAGAGCAGGAGAACAAGGTCCTTGACAGACTTGTCATTC

TTGTCAGCATCAGCTTCTTCTGCTACCTCTAGTTGGGTCTCTTGGGGTAGTACCTTCTCAACTCCTTCTC

TCGACTACGACTGTTCTTACTGTTCAGACAGTTCCTGGAACAGAGGACGAGAAACTCTGACGGGAAGAGT

GGAGACNNNAGTCGGACCTACTGGGGTTGTGGAAGCCGTTGTCTTAGGTGTCTTACAACTTTAACCCTAA

CTCGTAACTACTCCTACGACCACTTCGACTTCGACTACGACTGTACGGTGGTAACCTCCTACAACTACGG

CTCCCAGTCGTTCTACCTCTCGCCAACCTTAGCTCGAGCC

>EE332658.1 EST00154 Jute Plasmid library Corchorus capsularis cDNA, mRNA sequence

TATCTCGAGGGAGGCTTCCCGCTTAGATGCTTTAAGCGGTTATCCCGTCCGTACATAGCTACCCTGCTGC

GCCGTTGGCACGACGACAGGTACACCAGAGGTACGTTCAACCCGGTCCTCTCGTACTAGGGTCAACTCCT

CTCAAATTTCGACGCCCACGGCAGATAGGGACCAAACTGTCTCGCGACGTTCTGAANGCAGCTCACGTAC

CACTTTAATTGGCGAACAGCCAAACCCTTGGGACCTGCTCCAGCCCCAGGATGTGATGAGCCGACATCGA

GGTGCCAAACAACCCCGTCGATATGAGCTCTTGGGGGTTATCAGCCTGTTATCCCCGGCGTACCTTTTAT

CCGTTNAGCGATGGCCCTTCCA

>EE332657.1 EST00153 Jute Plasmid library Corchorus capsularis cDNA, mRNA sequence

TATCTCGAGGGAGGCTTCCCGCTTAGATGCTTTAAGCGGTTATCCCGTCCGTACATAGCTACCCTGCTGC

GCCGTTGGCACGACGACAGGTACACCAGAGGTACGTTCAACCCGGTCCTCTCGTACTAGGGTCAACTCCT

CTCAAATTTCGACGCCCACGGCAGATAGGGACCAAACTGTCTCGCGACGTTCTGAACCCAGCTCACGTAC

CACTTTAATTGGCGAACAGCCAAACCCTTGGGACCTGCTCCAGCCCCAGGATGTGATGAGCCGACATCGA

GGTGCCAAACAACCCCGTCGATATGAGCTCTTGGGGGTTATCAGCCTGTTATCCCCGGCGTACCTTTTAT

CCGTTGAGCGATGGCCCTTCCA

>EE332656.1 EST00152 Jute Plasmid library Corchorus capsularis cDNA, mRNA sequence

TGGAGGGGTCGGCCGGTCCGCTTTACGGTGTGCACCGGTCTACTCTTCTCTTCTGCCGGCGATGCGCTCC

TGGCCTTAATTGGCCGGGTCGCTCCTCCGGCTCTGTTACTTTGAAGAAATTAGAGTGCTCAAAGCAGGCC

TACGCTTGTATACATTAGCATGGGATAACATCATAGGATTTCGGTCCTATTGTGTTGGCCTTCGGGATCG

GAGTAATGATTAACAGGGACAGTCGGGGGCATTCGTATTTCATAGTCAGAGGTGAAATTCTTGGATTTAT

GAAAGACGAACAACTGCGAAAGCATTTGCCAAGGATGA

>EE332655.1 EST00151 Jute Plasmid library Corchorus capsularis cDNA, mRNA sequence

AAGCAAGGAATCCATCCAGAGCGGAGAAGGTAGTCTACTTCCTTAGCCAATTGCACGGGGGTGAGATCCG

GGAGGTATGAGAGAGTCTCGAACTTCTTCTTCCCAAGTGGTGGCCATACCTGCATGCATCGAACTCTGCC

ACCATTGCTGACGATGGAGGTGATGTCTTTGTTGTTCTTGCGGGTGGTGGGGAAAGCAGAGGCAGACTTG

AGGCCAACGAATGGGGCGACCATGGAGGCCTGTGCGGGGGAGGCACGGCTAACGACGGTGGCGGATGACA

TCATTGA

>EE332654.1 EST00150 Jute Plasmid library Corchorus capsularis cDNA, mRNA sequence

TCGCCATTTCAGCTCGAGAGTGATAATAGTGTTTCTGAACCTTATTTAAGGCCAACTTTCTTATCAAGGA

TATCACTCAGTGATATATGGAGACCAGCTACAAGAAAATACTGTGGCAATGATGTTTGGAAAATGTTAAA

AACTTCTGTTTCCATTCTCCTTTCACGGTCAGTCCTCCAGGAGCCAGCATTTCAAGAATTAATTTTACTT

TATACAACGGATTCTGCTGAAGGAGATACTGAAGATAGCTCTGAGGCTCAAGCATTACAATTAGAGATAT

ATGAGAGAATACCTATTCCGGATTTACCAGTCATTTTTCCTCACAAGAAGCTGTCTTTCCGTATCATAGA

CACGGTACGCCTAGATGTAGCCTCAATTTTGGGACTTCTGGCATTCGTCATAAACTACAAATTCGAGAAC

ATTTCATCTTCCCCATCAGCAATAGTTCTTGATGCGGTTGCAATAACTGCTCTCTTAGTGTATGTGTTCC

GTGTGGCCTTGGGTTACAAACAGACTTGGGATAGATATCAGCTATTAGTTAACAAGACCCTATACGAGAA

AACCTTGGCAAGTGGCTTTGGCACAGTTCATTTTCTTTTGGATGCTTCTGAACAGCAGCAATACAAGGAG

GCTATTTTAACCTATGCTATCCTGCTTAACATAGAGAACAGCCAGGTGTCTTGTCGCCNAGAAGTTGGAG

ATAAATGTGAGAGATTTATGTACGACGTTTTCNANCATAAGGTCGAATGCCAGTTGACAAGGCGTTGAAT

ACATTGTTAAAATGGGGA

>EE332653.1 EST00149 Jute Plasmid library Corchorus capsularis cDNA, mRNA sequence

TATCTCGAGGGAGGCTTCCCGCTTAGATGCTTTAAGCGGTTATCCCGTCCGTACATAGCTACCCTGCTGC

GCCGTTGGCACGACGACAGGTACACCAGAGGTACGTTCAACCCGGTCCTCTCGTACTAGGGTCAACTCCT

CTCAAATTTCGACGCCCACGGCAGATAGGGACCAAACTGTCTCGCGACGTTCTGAACCCAGCTCACGTAC

CACTTTAATTGGCGAACAGCCAAACCCTTGGGACCTGCTCCAGCCCCAGGATGTGATGAGCCGACATCGA

GGTGCCAAACAACCCCGTCGATATGAGCTCTTGGGGGTTATCAGCCTGTTATCCCCGGCGTACCTTTTAT

CCGTTGAGCGATGGCCCTTCCA

>EE332652.1 EST00148 Jute Plasmid library Corchorus capsularis cDNA, mRNA sequence

TATCTCGAGGGAGGCTTCCCGCTTAGATGCTTTAAGCGGTTATCCCGTCCGTACATAGCTACCCTGCTGC

GCCGTTGGCACGACGACAGGTACACCAGAGGTACGTTCAACCCGGTCCTCTCGTACTAGGGTCAACTCCT

CTCAAATTTCGACGCCCACGGCAGATAGGGACCAAACTGTCTCGCGACGTTCTGAACCCAGCTCACGTAC

CACTTTAATTGGCGAACAGCCAAACCCTTGGGACCTGCTCCAGCCCCAGGATGTGATGAGCCGACATCGA

GGTGCCAAACAACCCCGTCGATATGAGCTCTTGGGGGTTATCAGCCTGTTATCCCCGGCGTACCTTTTAT

CCGTTGAGCGATGGCCCTTCCA

>EE332644.1 EST00140 Jute Plasmid library Corchorus capsularis cDNA, mRNA sequence

TATAGGTCGCGCGCTTAAGCGCCATCCATTTTCGGGGCTAGTTGATTCGGCAGGTGAGTTGTTACACACT

CCTTAGCGGATTTCGACTTCCATGACCACCGTCCTGCTGTCTTAATCGACCAACACCCTTTGTGGGTTCT

AGGTTAGCGCGCAGTTGGGCACCGTAACCCGGCTTCCGGTCCATCCCGCATCGCCAGTTCTGCTTACCAA

AAATGGCCCACTTGGAGCTCTCGATTCCTTGGCACGGCTCAACAAAGCAGCCGCGCCGTCCTACCTATTT

AAAGTTTGAGAATAGGTCGAGGGCGTTGCGCCCCCGATGCCTCTAATCATTGGCTTTACCCGATAGAACT

CGCTAAGGGCTCCAGCTATCCTGAGGGAAACTTGGGAGGGAACCAGCTACTAGACGGTTCGATTAGTCTT

TCGCCCCTATACCCAAGTCAGACGAACGATTTGCACGTCAGTATCGCTGCGGGCCTCCACCAGAGTTTCC

TCTGGCTTCGCCCCGCTCAGGCATAGTTCACCATCTTTCGGGTCCCGACAGGCATGCTCTCACTCGAACC

CTTCTCAGAAGATCAAGGTCGGTCGGCGGTGCACCCGTGAGGGATCCCACCAATCAGCTTCCTTACGCCT

TACGGGTTTTCTAGCCCGTTGACTCGCACACATGTCAGACTCCTTGGTCCGTGTTACAAGACGGGCCGAA

TGGGGAGCCCACAGGCCGATGCCGGGAGCACGCA

>EE332643.1 EST00139 Jute Plasmid library Corchorus capsularis cDNA, mRNA sequence

GTTGTAANGANGGCCAGCTGCATTAATGAAATCGGCCAACGCGCGGGGAGAGGCGGTTTGCGTATTGGGC

GCTCTTCCGCTTCCTCGCTCACTGACTCGCTGCGCTCGGTCGTTCGGCTGCGGCGAGCGGTATCAGCTCA

CTCAAAGGCGGTAATACGGTTATCCACAGAATCAGGGGATAACGCAGGAAAGAACATGTGAGCAAAAGGC

CAGCAAAAGGCCAGGAACCGTAAAAAGGCCGCGTTGCTGGCGTTTTTCCATAGGCTCCGCCCCCCTGACG

AGCATCACAAAAATCGACGCTCAAGTCAGAGGTGGCGAAACCCGACAGGACTATAAAGATACCAGGCGTT

TCCCCCTGGAAGCTCCCTCGTGCGCTCTCCTGTTCCGACCCTGCCGCTTACCGGATACCTGTCCGCCTTT

CTCCCTTCGGGAAGCGTGGCGCTTTCTCATAGCTCACGCTGTAGGTATCTCAGTTCGGTGTAGGTCGTTC

GCTCCAAGCTGGGCTGTGTGCACGAACCCCCCGTTCAGCCCGACCGCTGCGCCTTATCCGGTAACTATCG

TCTTGATCCAACCCGGTAAGACACGACTTATCGCCACTGGCAGCAGCCACTGGTAACAGGATTAGCAGAG

CGAGGTATGTAGGCGGTGCTACAGAGTTCTTGAAGTGGTGGCCTAACTACGGCTACACTAGAAGAACAGT

ATTTGGTATCTGCGCTCTGCTGAAGCCAGTTACCTTCGGAAAAAGAGTTGGTAGCTCTNGATCCGGCAAA

AAACCACNGCTGGTAGCGGTGGTTTTTTTGTTTGCAAGCAGCAAATTA

>EE332642.1 EST00138 Jute Plasmid library Corchorus capsularis cDNA, mRNA sequence

TCGCGAATCCTCTTGAAAGAGAGGGGTGCCTTCGGGAACGCGGACACAGGTGGTGCATGGCTGTCGTCAG

CTCGTGCCGTAAGGTGTTGGGTTAAGTCCCGCAACGAGCGCAACCCTCGTGTTTAGTTGCCACCGTTGAG

TTTGGAACCCTGAGCAGACTGCCGGTGATAAGCCGGAGGAAGGTGAGGATGACGTCAAGTCATCATGCCC

CTTATGCCCTGGGCGACACACGTGCTACAATGGACGGGACAAAGGGTCGCGATCCCGCGAGGGTGAGCTA

ACTCCAAAAACCCGTCCTCAGTTCGGATTGCAGGCTGCAACTCGCCTGCATGAAGCCGGAATCGCTAGTA

ATCGCCGGTCAGCCATACGGCGGT

>EE332641.1 EST00137 Jute Plasmid library Corchorus capsularis cDNA, mRNA sequence

TCTACCGCCGCTCGGTGGGAATTACGCTTCAAGGAGGCCCGCACGGCTCATCCGCCGCACGGACTACTCC

TACGACACGTGCCCTTGGGGGCCAAAGGCCCCTACTGCGGGTCGGCAATCGGGCGGCGGGCGCATGCGTC

GCTTCTAGCCCGGATTCTGACTTAGAGGCGTTCAGTCATAATCCAGCGCACGGTAGCTTCGCGCCACTGG

CTTTTCAACCAAGCGCGATGACCAATTGTGCGAATCAACGGTTCCTCTCGTACTAGGTTGAATTACTATT

GCGACGCGGCCATCAGTAGGGTAAAACTAACA

>EE332640.1 EST00136 Jute Plasmid library Corchorus capsularis cDNA, mRNA sequence

TTTTTTTTTTTTTTTAAAACACAGACAAAAGCCACAAACGTAGAACTTAAATTTATTTAAGTTTTAAATA

ATGACAATATGAAACAAGAGCATCATATTGTTAAACGATGTGCCAGTCTGCCAGAGCATCACACTAAACC

AAAAGATGTCATCATAATGAAAATAGTAATATCACAAGACAAATATTCAATAGTTGCTGTTTAGTTTGGG

GGTTCTAAGTCAAGACCCTCATTTCTTCCCACTCTTCTTGAGTCCAGCTCCACCAAATGATCCCTTTTGT

TGTGCCTTTGCTCTCAGCTCTTTAAGAGCCTTCTCTTCTTCCTTCTTCTTCTGAAGCAAAGCCTTATCAT

TCTCGTCGAGTTCTTTCTTCTCAGCCTTGGGTTGCTTCAAAGGCTTCAATTTGCCACCTTGCTTGGACGA

CATGGTTAGGATTTGCGACTAAAATTCGAAGTGATCAGGGAAATTTGGGAGACAGAACAAATAAGA

>EE332639.1 EST00135 Jute Plasmid library Corchorus capsularis cDNA, mRNA sequence

GATGAGCACGCTGAGAAACCAAAGACCCTACCAGAGACCACAACAAACGAAAACAAGCTTATTCTCTATG

GACTGTTCAAGCAAGCAACCGTTGGACCTGTGAATACGAGCCGTCCTGGAATGTTCAACATGAGAGACAG

GTACAAGTGGGATGCATGGAAGGCTGTTGAAGGGAAATCAAAGGAGGAAGCTATGAGTGATTATATCACC

AAGGTGAAACAGTTGCTAGAGGAGGCTGCTGCATCTTCTTGATGTCTGACATAACATTGTGTGTTCCCAG

CCTTGCTACATGTTTCTGAAAATGTAAAACTAAAAATAAGTGTTTGTACTTTTATTGTTGCTACCATGTT

ATTCCCGAATTATGCAGGTTTAAACCAGATTTGATTTATCTATCAATAGTATAAACAAGCCCAAATAAGA

TGTTTA

>EE332638.1 EST00134 Jute Plasmid library Corchorus capsularis cDNA, mRNA sequence

AGTGCCAAGCTTGCATGCCTGCAGGTCGACTCTAGAGGATCCCCGGGTGCCTAATGAGTGAGCTAACTCA

CATTAATTGCGTTGCGCTCACTGCCCGCTTTCCAGTCGGGAAACCTGTCGTGCCAGCTGCATTAATGAAT

CGGCCAACGCGCGGGGAGAGGCGGTTTGCGTATTGGGCGCTCTTCCGCTTCCTCGCTCACTGACTCGCTG

CGCTCGGTCGTTCGGCTGCGGCGAGCGGTATCAGCTCACTCAAAGGCGGTAATACGGTTATCCACAGAAT

CAGGGGATAACGCAGGAAAGAACATGTGAGCAAAAGGCCAGCAAAAGGCCAGGAACCGTAAAAAGGCCGC

GTTGCTGGCGTTTTTCCATAGGCTCCGCCCCCCTGACGAGCATCACAAAAATCGACGCTCAAGTCAGAGG

TGGCGAAACCCGACAGGACTATAAAGATACCAGGCGTTTCCCCCTGGAAGCTCCCTCGTGCGCTCTCCTG

TTCCGACCCTGCCGCTTACCGGATACCTGTCCGCCTTTCTCCCTTCGGGAAGCGTGGCGCTTTCTCATAG

CTCACGCTGTAGGTATCTCAGTTCGGTGTAGGTCGTTCGCTCCAAGCTGGGCTGTGTGCACGAACCCCCC

GTTCAGCCCGACCGCTGCGCCTTATCCGGTAACTATCGTCTTGAGTCCAACCCNGGTAAGACACGACTTA

TCGCCACTGGCAGCAGCCACTGGTANCAGGATTAGCANANCGAGGTATGTAGGCGGTGCTACANANTTCT

TGAAGTGGTGGCCTANCTACGGCTACCCTANAAAAAAGTATTTGGTATCTGCCCTTTGCTGAANCCAGTT

ACNTNCGAAAAANANTTGGTAGCTNTNGATCCGGCAAAAAAACACCCTTGGTACGGTGGTTTTTTTTTNG

CANCNNCAAATNCCCCCAAAAAAAGNNTNNAAAAAN

>EE332630.1 EST00126 Jute Plasmid library Corchorus capsularis cDNA similar to Chain A, Structures of Adenylosuccinate Synthetase From Triticum Aestivum And (Arabidopsis thaliana), mRNA sequence

TATGAACTTCTCATCCCTCGCACTCGACCCCAGCCCAGTCGCCACCTCAGCAAGATGGGTCAACCAAAAG

CCCATTCTTCACTTCCACCACCACTCCTCCCGCCGTAACTTCGTTGTCTGCTCCTTGAAGCCCGCCGCCT

CCTCTTCCCTTAGTGTGGCTGAGTCAGCCGCCAGTGAGTCTCTTAAACGAATCGGCTCTCTCAGTCAGGT

TTCCGGCGTCTTGGGGTCTCAGTGGGGTGATTAAGGCAAAGGCAAACTTGTTGATATCTTGGCTGAGCAT

TTCGACATAGTCGCTCGCTGCCAGGGTGGAGCTAATGCGGGACATACTATCTATAACTCAGAGGGAAAGA

AGTTTGCCTTGCATCTTGTTCCTTCTGGCATCCTTAATGAGGACACCCTATGTGTTATTGGAAATGGAGT

AGTAGTCCATTTGCCTGGACTTTTTAAGGAAATTGATGGGTTGGAAGCTAATGGGGTCTCCTGCAAAGGA

CGGATATTGGTATCAGACCGTGCTCATTTGCTCTTCGACTTTCATCAAGTAGTTGATGGCCTCAGGGAAG

CCGAGCTCGCTAAATCGTTCATTGGCACAACTAAAAGGGGAATCGGACCCTGCTACTCTAGCAAGATGAT

CAGGAATGGCATTAGAGTAGGTGACTTGAGGCACATGGACACTTTCCCTCAGAAGCTTGATCTTTTATTA

TCAGATGCAGCTTCAAGATTCCCTGGTTTTAACTATACCCCAAAAATGCTCAAGGAAGAGGTTGAAAACT

ACAAAAAATACGCTGAAAGGTTGGAACCTTTCATGGCTGATACAGGGCACGGGAAGAATGAAGCCATTTC

CAAAAAAAAAA

>EE332629.1 EST00125 Jute Plasmid library Corchorus capsularis cDNA similar to Photosystem II 10 kDa polypeptide, chloroplast precursor (Light-inducible tissue-specific ST-LS1 protein), mRNA sequence

TGAAATGGCTGCCTCAGTTATGGCTTCTTCAATTACCCTCAAGCCTTCTTTCACTCTCAAGAAATCTAGT

GTCAAAGGAGTTCCTTCCCTTGCTAGGTCTCCTAAAGTTTTCAACGTTCTAGCCTCCGGTGGCAAGAAGA

TCAAGACTGACAAGCCTTATGGAATCAATGGAGGCATGGCCTTGAGGGATGGTCTTGATGCCTCTGGAAG

GAAGGGCAAGGGAAAGGGTGTTTACCAATTTGTTGACAAATATGGTGCTAACGTCGATGGCTACAGCCCT

ATCTACGACCAAAAGGACTGGTCTCCCACCGGTGATGTTTATGCTGGCGGTGTAACTGGTTTGGCAATCT

GGGCAGTTACCCTAGCCGGTATTCTTGCAGGTGGTGCTCTTCTTGTTTACAACACAAGTGCTTTGGCACA

GTAGATTTCTATAATGTAATGTAATCCGTGTTGTATAATATAGCCACTCCAACGTTATTGTTCTCATCAT

AATACTTCAAGAACTACTTTGCCTATCTAAAAAAAAAAAA

>EE332628.1 EST00124 Jute Plasmid library Corchorus capsularis cDNA similar to At2g43970/F6E13.10 (Arabidopsis thaliana), mRNA sequence

TGTGATCCTTCTTCACAACAGCCTCAACTTCTTGCTCTCGAGGATGATGATGATGGTTCTGGTAAGTAGG

AGGAGGATGATGATGATGATTCCGCATGGGCATGTGATGGTGAACGGGAACCATGTGAGGCGGCATAGGC

ACGGGGCTATGGACGGGGTAGGTGAAACGCGGCCGGAGGAGGAGGTGGAGAAGGCGGGTAGACGTGCAGC

ATCCCAGGGGGCGGAGGAGGATGGATCATCATCCTGTGACTCTGCTGCTGCTGCTGCTGCTGCTGTGGCT

GCTGAGGAGTAGAGCGAGTAGGCACAAACTCCGGCGCCTGAGCGTTGAGCCGACTGGATGACAAATTCCG

CAATAGTTGGGGATCTGAGGATAAAGATGAGGATTGCGGAGAATCTAGGGTTTCGGATTGGGCCATCGTT

TGGGAGATGTACAGGGTTTTTCTTTTTTTCTTTTTTTCTTTTTCCTTGATTTGAAGAATTAAAATAAAAA

TAAATAGTACAAGTANATGGTAAANATAAATAAAATGGGGGAGATANGGTANCTGCTGCGCGGCGGCCTG

GTTCTTGCGGGAAGGAANAGCCCTA

>EE332627.1 EST00123 Jute Plasmid library Corchorus capsularis cDNA similar to At2g43970/F6E13.10 (Arabidopsis thaliana), mRNA sequence

TAGGGCTCTTCCTTCCCGCAAGAACCAGGCCGCCGCGCAGCAGCTACCCTATCTCCCCCATTTTATTTAT

TTTTACCATCTACTTGTACTATTTATTTTTATTTTAATTCTTCAAATCAAGGAAAAAGAAAAAAAGAAAA

AAAGAAAAACCCTGTACATCTCCCAAACGATGGCCCAATCCGAAACCCTAGATTCTCCGCAATCCTCATC

TTTATCCTCAGATCCCCAACTATTGCGGAATTTGTCATCCAGTCGGCTCAACGCTCAGGCGCCGGAGTTT

GTGCCTACTCGCTCTACTCCTCAGCAGCCACAGCAGCAGCAGCAGCAGCAGCAGAGTCACAGGATGATGA

TCCATCCTCCTCCGCCCCCTGGGATGCTGCACGTCTACCCGCCTTCTCCACCTCCTCCTCCGGCCGCGTT

TCACCTCCCCGTCCATAGCCCCGTGCCTATGCCGCCTCACATGGTTCCCGTTCACCATCACATGCCCATG

CGGAATCATCATCATCATCCTCCTCCTACTTACCAGAACCATCATCATCATCCTCGAGAGCAAGAAGTTG

AGGCTGTTGTGAAGAAGGATCACA

>EE332626.1 EST00122 Jute Plasmid library Corchorus capsularis cDNA similar to Hybrid proline-rich protein, mRNA sequence

TTTTCTCTTCAAAAACAGAGAAAAAAGAGGACCATTCTAAGCACTTAGTTGTGTCACTTGAAAAATGGCT

TCAAAATCTGTAGCATCCATTGCCTTCCTTCTCACTCTCAACCTTCTCTTCTTCACTTTGGTCAGTTCAA

CCTACACCCCTTGCCCACCATCACCCAAAAAGCCCACCCCGACCCCTTCCCCGAAGCCAAAACCCAAACC

CAAGCCCGCCGCCTCTTGCCCTAAAGACACTCTCAAATTGGGAGTTTGTGCTGATTTGTTGGGTTTGGTT

AACGTTGTTGTTGGTTCCCCTCCAAAGACTCCTCCTGCACTCTCCTCGAAGGTTTGGCTGATCTTGAAGC

TGCTGCTTGCCTTTGCACTGCCATTAAGGCTAATGTTTTGGGAATTAACCTTAATGTTCCCGTTTCATTG

TCTTTGCTCCTTAATTACTGTGGAAAGAAGGTTCCTGCTGGCTACCAGTGTGCTTAATTAAAAGCTCTTT

AATTTCGTCTTACTTGTAATAATCAATTACTGTCTTGGTACTTTCGGGCTTATGTTTGGTCGTTTTTATA

GTCGTTTTCATGGTGTGATATTGGAAGTTTGGAATCTGGGTACGTACGTATAGTATTATGTCATACTGCA

ATTTATTACTATATTTTTACTTTTAATTTGATCAAGGAATTAGGAAGGCTNCTCACCTTCTANTCCTTTT

ATATATGCGAAANAAGAATGTATTATTTACATCACTCTGTATCTAANAAAATGAAANGTTTTACTTTTAT

CAGCGGGCCGCGAAATNANTTCACCAAGNGGTGGAATGNTNCNCCCNCNAAANGGGANCGNGANCTGGGN

ATNAACCNCNCNNGAAAANG

>EE332625.1 EST00121 Jute Plasmid library Corchorus capsularis cDNA similar to Bimodular protein, mRNA sequence

TCAAACATTAATTAAACATTACATTAGGCCGAGGTATAAAATTAACCCACAAGCCTAGTTATTTAAAACA

TGAACAAATATCACACAATGTCGGAAGTCTTTGTAATAAAGACTCCCTAAAAATATGTAAAAATGGAAAC

CCTATCCCAACTTCAAATCGATCCAAACCACTAAACAAAAATACATTGAATATTTGAAACTACAG

>EE332624.1 EST00120 Jute Plasmid library Corchorus capsularis cDNA similar to Pyruvate decarboxylase, mRNA sequence

TGTCACCGCACAAGATGTGTCAACAATGCTGCGATGTGGCCAAAAAACCATAATCTTCCTGATAAACAAT

GGTGGTTACACCATTGAAGTTGAAATTCATGATGGACCATACAATGTAATCAAGAACTGGAACTACACTG

GTTTGGTCGATGCAATTCATAATGGTGAAGGCAAGTGCTGGACAGCCAAGGTCTTTTGTGAGGAAGAACT

GATTGAAGCGATTGAGACAGCAACTGGAGCCAAGAAGGATTGCTTATGTTTCATTGAAGTGATAGTTCAC

AAGGATGATACAAGCAAAGAACTGTTGGAGTGGGGTTCTAGGGTCTCTGCTGCTAATAGCCGTCCACCAA

ATCCTCAGTAAAAAAAAGAACTCATGGCGCTGCTTGAGTGATTTCCAGCATTGGACGCTATAAAATCTTT

CTAAGTACATTGAGAATTAATCAGTCTCAGTTTAAGTTTCTTGGATGCAGAGTTAAATCTGCGATTTCAA

CATTTGCATATTCTACTGAGTGCTCGCTGATATTTGTGTAAATTTACTTTCAATGTCAATTCTAGTACTA

ATGAAATGTTTNGCCTTTACNGNCCCGAANTCNTANCATGTCATANCTGTTTCCNGNGGNAANTGTTATC

CNCTCCCANTNCCCNNACATACNANCCGNAACATAAC

>EE332616.1 EST00112 Jute Plasmid library Corchorus capsularis cDNA similar to Solanum nigrum chloroplast tRNA-Ala, tRNA-Ile, 16S rRNA, tRNA-Val, rps12, rps7, ndhB genes, mRNA sequence

TCGCTCCCCTAGCTTTCGTCTCTCAGTGTCAGTGTCGGCCCAGCAGAGTGCTTTCGCCGTTGGTGTTCTT

TCCGATCTCTACGCATTTCACCGCTCCACCGGAAATTCCCTCTGCCCCTACCGTACTCCAGCNTGGTAGT

TTCCACCGCCTGTCCAGGGTTGAGCCCTGGGATTTGACGGCGGACTTAAAAAGCCACCTACAGACGCTTT

ACGCCCAATCATTCCGGATAACGCTTGCATCCTCTGTATTACCGCGGCTGCTGGCACAGAGTTAGCCGAT

GCTTATTCCCCAGATACCGTCATTGCTTCTTCTCCGGGAAAAGAAGTTCACGACCCGTGGGNCTTCTACC

TCCACGCGGCATTGCTCCGTCAGGCTTTCGCCCATTGCGGAAAATTCCCCACTGCTGCCTCCCGTAGGAG

TCTGGGCCGTGTCTCAGTCCCAGTGTGGCTGATCATCCTCTCGGACCAGCTACTGATCATCGNCTTGGTA

AGCTATTGCCTCACCAACTAGNTAATCAGACGCGAGA

>EE332615.1 EST00111 Jute Plasmid library Corchorus capsularis cDNA similar to Type II keratin E1, mRNA sequence

GTTGTAACGACGGCCAGTGCCAAGCTTGCATGCCTGCAGGTCGACTCTAGAGGATCCCCGGGTACCGAGC

TCGATTCGTAATCATGTCATAGCTGTTTCCTGTGTGAAATTGTTATCCGCTCACAATTCCACACAACATA

CGAGCCGGAAGCATAAAGTGTAAAGCCTGGGGTGCCTAATGAGTGAGCTAACTCACATTAATTGCGTTGC

GCTCACTGCCCGCTTTCCAGTCGGGAAACCTGTCGTGCCAGCTGCATTAATGAATCGGCCAACGCGCGGG

GAGAGGCGGTTTGCGTATTGGGCGCTCTTCCGCTTCCTCGCTCACTGACTCGCTGCGCTCGGTCGTTCGG

CTGCGGCGAGCGGTATCAGCTCACTCAAAGGCGGTAATACGGTTATCCACAGAATCAGGGGATAACGCAG

GAAAGAACATGTGAGCAAAAGGCCAGCAAAAGGCCAGGAACCGTAAAAAGGCCGCGTTGCTGGCGTTTTT

CCATAGGCTCCGCCCCCCTGACGAGCATCACAAAAATCGACGCTCAAGTCAGAGGTGGCGAAACCCGACA

GGACTATAAAGATACCAGGCGTTTCCCCCTGGAAGCTCCCTCGTGCGCTCTCCTGTTCCGACCCTGCCGC

TTACCGGATACCTGTCCGCCTTTCTCCCTTCGGGAAGCGTGGCGCTTTCTCATAGCTCACGCTGTAGGTA

TCTCAGTTCGGTGTAGGTCGTTCGCTCCAAGCTGGGCTGTGTGCACGAACCCCCGTTCAGCCCGACCGCT

GCGCCTTATCCGGTAACTATCGTCTTGAGTCCAACCCGGTANANACGACTTATCGCCACTGGCAGCANCC

CTGGTAACAGGATTANCAAANCGAGGTATGTAGGCGGTGCTANAANTTCTTGAANTGGNGCCTANCTACG

GCTACCTAAAAAANANTATTGGNTTCNGCCCN

>EE332614.1 EST00110 Jute Plasmid library Corchorus capsularis cDNA similar to Catalytic/ structural constituent of eye lens (Arabidopsis thaliana), mRNA sequence

TATTTAATGGTGTTTCTTTCGAGAGCAATGGATCGCTGGATGAAATCGTGGAACTGGGCGATATTGTTAG

CTCGCTACAAATGCTGAATCGCCGCTAGTGAAAGGCGAGAAGTTGAANCNNGGGCGCATCTGGATTTGGT

TGGGTCGTTTACGCATACCATGAAGGAATGTGACGATGAGGCAATAAAGAGAGGGAGAGTTTTTGTTGAT

AACGAGGCGGCTTTGATAGAGGCATGGGAGCTGGTGGGTGCTTTCGAGAGAGGGGTGATTGGGAAGCAAG

ATATTGGGGGAAATTTGNCGGAGTTGATTCAAGGANAAAAGATTGGGAGGAA

>EE332613.1 EST00109 Jute Plasmid library Corchorus capsularis cDNA similar to Ribulose bisphosphate carboxylase small chain 2, chloroplast precursor (RuBisCO small subunit 2), mRNA sequence

GAATTGGAGCACGGATTCGTCTACCGTAAGAACCACAGCTCCCCAGGGTACTACGATGGACGCTACTGGA

CCATGTGGAAACTTCCCATGTTCGGGTGCACTGACTCATCCCAGGTGTTGAAGGAGCTTGAGGAGGCCAA

GAAGGAATACCCCGAGTCCTTCATCAGAATCATTGGATTCGACAACAAGCGTCAAGTGCAATGCATCAGT

TTCATTGCCTACAAGCCCCCAAGCTTCACCTCAGCTTAAGAAAATAGAGTTGTTTTCCCCACATTAAGTA

GGGATTCGGCTTCTTTAACTTGTACTTAATTATATATAGCATTGTTCGAAAACATTTTTCTTTCCTTTTG

AGGTATTGTTGTTTGTTGTTTGATTTGGAATGGAAGGGCTGAGAACCATAGTTTCTTGCCATA

>EE332612.1 EST00108 Jute Plasmid library Corchorus capsularis cDNA similar to Cytochrome P450 like_TBP [Nicotiana tabacum], mRNA sequence

TGTTAGTTTTACCCTACTGATGGCCGCGTCGCAATAGTAATTCAACCTAGTACGAGAGGAACCGTTGATT

CGCACAATTGGTCATCGCGCTTGGTTGAAAAGCCAGTGGCGCGAAGCTACCGTGCGCTGGATTATGACTG

AACGCCTCTAAGTCAGAATCCGGGCTAGAAGCGACGCATGCGCCCGCCGCCCGATTGCCGACCCGCAGTA

GGGGCCTTTGGCCCCCAAGGGCACGTGTCGTAGGAGTAGTCCGTGCGGCGGATGAGCCGTGCAGGCCTCC

TTGAAGCGTAATTCCCACCGAGCGGCGGGTAGA

>EE332611.1 EST00107 Jute Plasmid library Corchorus capsularis cDNA similar to Type II keratin E1, mRNA sequence

GTAATCATGTCATAGCTGTTTCCTGTGTGAAATTGTTATCCGCTCACAATTCCACACAACATACGAGCCG

GAAGCATAAAGTGTAAAGCCTGGGGTGCCTAATGAGTGAGCTAACTCACATTAATTGCGTTGCGCTCACT

GCCCGCTTTCCAGTCGGGAAACCTGTCGTGCCAGCTGCATTAATGAATCGGCCAACGCGCGGGGAGAGGC

GGTTTGCGTATTGGGCGCTCTTCCGCTTCCTCGCTCACTGACTCGCTGCGCTCGGTCGTTCGGCTGCGGC

GAGCGGTATCAGCTCACTCAAAGGCGGTAATACGGTTATCCACAGAATCAGGGGATAACGCAGGAAAGAA

CATGTGAGCAAAAGGCCAGCAAAAGGCCAGGAACCGTAAAAAGGCCGCGTTGCTGGCGTTTTTCCATAGG

CTCCGCCCCCCTGACGAGCATCACAAAAATCGACGCTCAAGTCAGAGGTGGCGAAACCCGACAGGACTAT

AAAGATACCAGGCGTTTCCCCCTGGAAGCTCCCTCGTGCGCTCTCCTGTTCCGACCCTGCCGCTTACCGG

ATACCTGTCCGCCTTTCTCCCTTCGGGAAGCGTGGCGCTTTCTCATAGCTCACGCTGTAGGTATCTCAGT

TCGGTGTAGGTCGTTCGCTCCAAGCTGGGCTGTGTGCACGAACCCCCGTTCAGCCCGACCGCTGCGCCTT

ATCCGGTAACTATCGTCTTGAGTCCAA

>EE332610.1 EST00106 Jute Plasmid library Corchorus capsularis cDNA similar to Type II keratin E1, mRNA sequence

GTAATCATGTCATAGCTGTTTCCTGTGTGAAATTGTTATCCGCTCACAATTCCACACAACATACGAGCCG

GAAGCATAAAGTGTAAAGCCTGGGGTGCCTAATGAGTGAGCTAACTCACATTAATTGCGTTGCGCTCACT

GCCCGCTTTCCAGTCGGGAAACCTGTCGTGCCAGCTGCATTAATGAATCGGCCAACGCGCGGGGAGAGGC

GGTTTGCGTATTGGGCGCTCTTCCGCTTCCTCGCTCACTGACTCGCTGCGCTCGGTCGTTCGGCTGCGGC

GAGCGGTATCAGCTCACTCAAAGGCGGTAATACGGTTATCCACAGAATCAGGGGANANCGCAGGAAAAAA

CATGTGANCAAAAGGCC

>EE332602.1 EST00098 Jute Plasmid library Corchorus capsularis cDNA similar to S.alba 18S, 5.8S & 25S rRNA genes, mRNA sequence

TGCAAAGGATTCCACCCGCCGCTCGGTGGGAATTACGCTTCAAGGAGGCCTGCACGGCTCATCCGCCGCA

CGGACTACTCCTACGACACGTGCCCTTGGGGGCCAAAGGCCCCTACTGCGGGTCGGCAATCGGGCGGCGG

GCGCATGCGTCGCTTCTAGCCCGGATTCTGACTTAGAGGCGTTCAGTCATAATCCAGCGCACGGTAGCTT

CGCGCCACTGGCTTTTCAACCAAGCGCGATGACCAATTGTGCGAATCAACGGTTCCTCTCGTACTAGGTT

GAATTACTATTGCGACGCGGCCATCAGTAGGGTAAAACTAACCTGTCTCACGACGGTCTAAACCCAGCTC

ACGTTCCCTATTGGTGGGTGAACAATCCAACACTTGGT

>EE332601.1 EST00097 Jute Plasmid library Corchorus capsularis cDNA similar to Citrus limon cistron for 26S ribosomal RNA, mRNA sequence

TCAAAGGATTCTACCCGCCGCTCGGTGGGAATTACGCTTCAAGGAGGCCTGCACGGCTCATCCGCCGCAC

GGACTACTCCTACGACACGTGCCCTTGGGGGCCAAAGGCCCCTACTGCGGGTCGGCAATCGGGCGGCGGG

CGCATGCGTCGCTTCTAGCCCGGATTCTGACTTAGAGGCGTTCAGTCATAATCCAGCGCACGGTAGCTTC

GCGCCACTGGCTTTTCAACCAAGCGCGATGACCAATTGTGCGAATCAACGGTTCCTCTCGTACTAGGTTG

AATTACTATTGCGACGCGGCCATCAGTAGGGTAAAACTAACCTGTCTCACGACGGTCTAAACCCAGCTCA

CGTTCCCTCGCGCGGGCGCTTCCCCCTAAGGCGGAACGCTCCCCTACCGATGCATTTTTACATCCCACAG

CTTCGGCAGATCGCTTAGCCCCGTTCATCTTCGGCGCAAGAGCGCTCGATCAGTGAGCTATTACGCACTC

TTTCAAGGGTGGCTGCTTCTAGGCAAACCTCCTGGCTGTCTCTGCACCCCTACCTCCTTTATCACTGAGC

GGTCATTTAGGGGGCCTTAGCTGGTGATCCGGGCTGTTTCCCTCTCGACGATGAAGCTTATCCCCCATCG

TCTCACTGGCCGA

>EE332600.1 EST00096 Jute Plasmid library Corchorus capsularis cDNA similar to Unspecific monooxygenase (EC 1.14.14.1) - common tobacco, mRNA sequence

TGTGCGAGTCAACGGGCTAGAAAACCCGTAAGGCGTAAGGAAGCTGATTGGTGGGATCCCTCACGGGTGC

ACCGCCGACCGACCTTGATCTTCTGAGAAGGGTTCGAGTGAGAGCATGCCTGTCGGGACCCGAAAGATGG

TGAACTATGCCTGAGCGGGGCAAGCCAGAGGGAACTCTGGTGGAGGCCCGCAGCGATACTGACGTGCAAA

TCGTTCGTCTGACTTGGGTATAGGGGCGAAAGACTAATCGAACCGTCTAGTAGCTGGTTCCCTCCGAAGT

TTCCCTCAGGATAGCTGGAGCCCTTAGCGAGTTCTATCGGGTAAAGCCAATGATTACAGGCATCGGGGGC

GCAACGCCCTCGACCTATTCTCAAACTTTAAATAGGTAGGACGGCGCGGCTGCTTTGTTGAGCCGTGCCA

AGGAATCGAGAGCTCCAAGTGGGCCATTTTTGGTAAGCAGAACTGGCGATGCGGGATGAACCGGAAGCCG

GGTTACGGTGCCCAACTGCGCGCTAACCTAGAACCCACAAAGGGTGTTGGTCGATTAAGA

>EE332599.1 EST00095 Jute Plasmid library Corchorus capsularis cDNA similar to Tobacco chloroplast 23S and 4.5S rRNA genes, mRNA sequence

TGGGAAAGGTCCTCTCAATGCTCTAACGCCCACACCGGATATGGACCGAACTGTCTCACGACGTTCTGAA

CCCAGGTCACGTACCGCTTTAATGGGCGAACAGCCCAACCCTTGGAACATACTACAGCCCCAGGTGGCGA

AGAGCCGACATCGAGGTGCCAAACCTTCCCGTCGATGTGAGCTCTTGGGGAAGATCAGCCTGTTATCTCT

AGAGTAACTTTTATCCGTTGAGCGACGGCCCTTCCACTCGGCACCGTCGGATCACTAAGGCCGACTTTCG

TCCCTGCTCGACGGGTGGGTCTTGCAGTCAAGCTCCCTTCTGCCTTTGCACTCGAGGGCCAATCTCCGTC

CGGCCCGAGGAAACCTTTGCACGCCTCCGTTACCTTTTGGGAGGCCTACGCCCCATAGAAACCGTCTACC

TGAGACTGTCCCTTGGCCCGTAGGTCCTGACACAAGGTTA

>EE332598.1 EST00094 Jute Plasmid library Corchorus capsularis cDNA similar to Tragopogon dubius large subunit 26S ribosomal RNA gene, partial sequence, mRNA sequence

TCGCTGGCTGAATTTTTCACTTGACATTCAGAGCACTGGGCAGAAATCACATTGCGTGAGCATCCGCAGG

GACCATCGCAATGCTTTGTTTTAATTAAACAGTCGGATTCCCCTTGTCCGTACCAGTTCTGAGTCGGCTG

CTAATTGCCCGGGGAAGGCCCCCGAAGGGACCGTTCCCAGTCCGTCCCCCGGCCGGCACGCGGCGACCCG

CTCTCGCCACGGAAGCAGCTCGAGCAGTCCGCCGACAGCCGACGGGTTCGGGACTGGGACCCCCGTGCCC

AGCCCTCAGAGCCAATCCTTTTCCCGAGGTTACGGATCCATTTTGCCGACTTCCCTTGCCTACGTGGAAT

CGGACACCACTAAACTTAAGCTTCACGATGCATGCATGCATATATGGTTCTAAGTCCATATGAGTGGATG

CATGTTCTGTTAAATGTGCTTTATAGAGTATTATTTATCTTTCTACAGTACCACAATATTCCGAACGATT

TTGTTGAAAGATATATATATTTATCTTTACAATAAATTAATAACCTCTTACAAATCCTAGCTATGTACCT

TCTTGCTACTTTTATATGTCTCGTATTATTCTCTTTCCCCTCTAGATATATANTGCAAAGCTTTGATTTG

TGAAAAAAAAAA

>EE332597.1 EST00093 Jute Plasmid library Corchorus capsularis cDNA similar to S.alba 18S, 5.8S & 25S rRNA genes, mRNA sequence

ACCAAGTGTTGGATTGTTCACCCACCAATAGGGAACGTGAGCTGGGGTTAGACCGTCGTGAGACAGGTTA

GTTTTACCCTACTGATGGCCGCGTCGCAATAGTAATTCAACCTAGTACGAGAGGAACCATTGATTCGCAC

AATTGGTCATCGCGCTTGGTTGAAAAGCCAGTGGCGCGAAGCTACCGTGCGCTGGATTATGACTGAACGC

TTCTAAGTCAGAATCCGGGCTAGAAGCGACGCATGCGCCCGCCGCCCGATTGCCGACCCGCAGTAGGGGC

CTTTGGCCCCCAAGGGCACGTGTCGTAGGAGTAGTCCGTGCGGCGGATGAGCCGTGCAGGCCTCCTTGAA

GCGTAATTCCCACCGAGCGGCGGGTAGAATCCA

>EE332596.1 EST00092 Jute Plasmid library Corchorus capsularis cDNA similar to Tragopogon dubius large subunit 26S ribosomal RNA gene, partial sequence, mRNA sequence

TCGCTCGGTGAATTTTTTCACTTGGACATTCAGAGCACTGGGCAGAAATCCCATTGCGTGAGCATCCGCA

GGGACCATCGCAATGCTTTGTTTTAATTAAACAGTCGGATTCCCCTTGTCCGTACCAGTTCTGAGTCGGC

TGCTAATTGCCCGGGGAAGGCCCCCGAAGGGACCGTTCCCAGTCCGTCCCCCGGCCGGCACGCGGCGACC

CGCTCTCGCCACGGAAGCAGCTCGAGCAGTCCGCCGACAGCCGACGGGTTCGGGACTGGGACCCCCGTGC

CCAGCCCTCAGAGCCAATCCTTTTCCCGAGGTTACGGATCCATTTTGCCGACTTCCCTTGCCTACGTGGA

ATCGGACACCACTAAACTTAAGCTTCACGATGCATGCATGCATATATGGTTCTAAGTCCATATGAGTGGA

TGCATGTTCTGTTAAATGTGCTTTATAGAGTATTATTTATCTTTCTACAGTACCACAATATTCCGAACGA

TTTTGTTGAAAGATATATATATTTATCTTTACAATAAATTAATAACCTCTTACAAATCCTAGCTATGTAC

CTTCTTGCTACTTTTATATGTCTCGTATTATTCTCTTTCCCCTCTAGATATATANTGCAAAGCTTTGATT

TGTGAAAAAAAAAA

>EE332588.1 EST00084 Jute Plasmid library Corchorus capsularis cDNA similar to Proline-rich protein, mRNA sequence

TTCCTTATTCACACTTGAAGCCTGGAGGAATTTCCTTTTGGCATGCACTAAGGAGTAAGCTAAGAGTAAC

AGGAATGTTGAGGGTGATTCCAAGTACATTGGCTTTAATGGCAGTGCAAAGGCAAAGGGCAGCTTCGGCA

TCAGCCAAGCCTGAGAGCAAAGCGCAGCATTTGCTTGAAGGGGGTGTTCCGACAATGACGTTGACAAGTC

CGAGGAGGTCGGCGCAGACACCGAGCTTCAATGTGTCCTTAGGGCAAGACGCCGGTTTTGGTGGCGGAGG

GCAAGCTTTTGGAGCTGGAGGAGGACTAGGGGTGGGTTTGGGCTTGGGTTTGCATGAACCGCAAGCTGTG

A

>EE332587.1 EST00083 Jute Plasmid library Corchorus capsularis cDNA similar to Cell wall-associated hydrolase, mRNA sequence

TATCTCGAGGGAGGCTTCCCGCTTAGATGCTTTAAGCGGTTATCCCGTCCGTACATAGCTACCCTGCTGC

GCCGTTGGCACGACGACAGGTACACCAGAGGTACGTTCAACCCGGTCCTCTCGTACTAGGGTCAACTCCT

CTCAAATTTCGACGCCCACGGCAGATAGGGACCAAACTGTCTCGCGACGTTCTGAACCCAGCTCACGTAC

CACTTTAATTGGCGAACAGCCAAACCCTTGGGACCTGCTCCAGCCCCAGGATGTGATGAGCCGACANCGA

GGTGCCAAACAACCCCGTCGATATGAGCTCTTGGGGGTTATCAGCCTGTTATCCCCGGCGTACCTTTTAT

CCGTTGAGCGATGGCCCTTCCAG

>EE332586.1 EST00082 Jute Plasmid library Corchorus capsularis cDNA similar to Cell wall-associated hydrolase, mRNA sequence

TATCTCGAGGGAGGCTTCCCGCTTAGATGCTTTAAGCGGTTATCCCGTCCGTACATAGCTACCCTGCTGC

GCCGTTGGCACGACGACAGGTACACCAGAGGTACGTTCAACCCGGTCCTCTCGTACTAGGGTCAACTCCT

CTCAAATTTCGACGCCCACGGCAGATAGGGACCAAACTGTCTCGCGACGTTCTGAACCCAGCTCACGTAC

CACTTTAATTGGCGAACAGCCAAACCCTTGGGACCTGCTCCAGCCCCAGGATGTGATGAGCCGACATCGA

GGTGCCAAACAACCCCGTCGATATGAGCTCTTGGGGGTTATCAGCCTGTTATCCCCGGCGTACCTTTTAT

CCGTTGAGCGATGGCCCTTCCA

>EE332585.1 EST00081 Jute Plasmid library Corchorus capsularis cDNA similar to Cell wall-associated hydrolase, mRNA sequence

TATCTCGAGGGAGGCTTCCCGCTTAGATGCTTTAAGCGGTTATCCCGTCCGTACATAGCTACCCTGCTGC

GCCGTTGGCACGACGACAGGTACACCAGAGGTACGTTCAACCCGGTCCTCTCGTACTAGGGTCAACTCCT

CTCAAATTTCGACGCCCACGGCAGATAGGGACCAAACTGTCTCGCGACGTTCTGAACCCAGCTCACGTAC

CACTTTAATTGGCGAACAGCCAAACCCTTGGGACCTGCTCCAGCCCCAGGATGTGATGAGCCGACATCGA

GGTGCCAAACAACCCCGTCGATATGAGCTCTTGGGGGTTATCAGCCTGTTATCCCCGGCGTACCTTTTAT

CCGTTGAGCGATGGCCCTTCCA

>EE332584.1 EST00080 Jute Plasmid library Corchorus capsularis cDNA similar to Cell wall-associated hydrolase, mRNA sequence

TATCTCGAGGGAGGCTTCCCGCTTAGATGCTTTAAGCGGTTATCCCGTCCGTACATAGCTACCCTGCTGC

GCCGTTGGCACGACGACAGGTACACCAGAGGTACGTTCAACCCGGTCCTCTCGTACTAGGGTCAACTCCT

CTCAAATTTCGACGCCCACGGCAGATAGGGACCAAACTGTCTCGCGACGTTCTGAACCCAGCTCACGTAC

CACTTTAATTGGCGAACAGCCAAACCCTTGGGACCTGCTCCAGCCCCAGGATGTGATGAGCCGACATCGA

GGTGCCAAACAACCCCGTCGATATGAGCTCTTGGGGGTTATCAGCCTGTTATCCCCGGCGTACCTTTTAT

CCGTTGAGCGATGGCCCTTCCA

>EE332583.1 EST00079 Jute Plasmid library Corchorus capsularis cDNA similar to Cell wall-associated hydrolase, mRNA sequence

TATCTCGAGGGAGGCTTCCCGCTTAGATGCTTTAAGCGGTTATCCCGTCCGTACATAGCTACCCTGCTGC

GCCGTTGGCACGACGACAGGTACACCAGAGGTACGTTCAACCCGGTCCTCTCGTACTAGGGTCAACTCCT

CTCAAATTTCGACGCCCACGGCAGATAGGGACCAAACTGTCTCGCGACGTTCTGAACCCAGCTCACGTAC

CACTTTAATTGGCGAACAGCCAAACCCTTGGGACCTGCTCCAGCCCCAGGATGTGATGAGCCGACATCGA

GGTGCCAAACAACCCCGTCGATATGAGCTCTTGGGGGTTATCAGCCTGTTATCCCCGGCGTACCTTTTAT

CCGTTGAGCGATGGCCCTTCCA

>EE332582.1 EST00078 Jute Plasmid library Corchorus capsularis cDNA similar to Solanesyl diphosphate synthase 2, mRNA sequence

GATCATTGAGTCTGAGTTTTGTGAGCCTGGATCGCTTGAAGAAGCGATTGAAATAGTTAGAAATCGCGGT

GGGATCAAGAAAGCTCAAGAATTAGCAAAGGAGAAAGCTGAACTTGCGTTAAAGAATTTGAATTGTCTTC

CTAGGAGTGGTTTCAGATCGGCTCTTGAGGATATGGTAATGTTTAATCTTGAAAGGATTGATTAGTTTCA

AAGGGTATTTTCAAGAGTCTACAATGTCTCCCTCAGAGCATGTTTCGGTTAGCTCTCGAAGACATGGTGA

TGTTTAATCTTGAAAGGATCGATTAGTTAGACAACATTTTCACAGTACAAGCAAATTCCTTGAAATGGTG

GACCATTATTCTTAAATCTTAAACATACCCAAAAGCATAAACAGAGGATGCATGAAGTTGAAGCCTCTTA

CTCATATAGAAAATCTTTTTCTTAGCACAATAGTTGGTTAGTCCAGCTGAATCCAGGTTCCTTGTGACTG

GTTTTTTTTTTTTACACCATAGGACCACAACAATAACTCTAAAAATAAATTAAGAAAAACGGAATTTAAT

GAAAAAAAAATAAAAGCCAAAAAAGAAATTTTTGGGAGGTGGAAATTAAAAAAAAAAAAAAGAGCAAAGG

GGGCCAACCATGTGATTGATGATGAAGACTGGAATATCAATCCTCAACCGGATA

>EE332574.1 EST00070 Jute Plasmid library Corchorus capsularis cDNA similar to Transcription regulator (Arabidopsis thaliana), mRNA sequence

TCTTGAAGTTGAACCCAAAGAACAGAAAGAAGAAGAAATAAGGAAGCGGCGACAGAGGAGAAGAGATGGG

AGTAACAAAGGAACAAGTTGAATCTTCGTTGACTTCAAAATTGAATCCTTCTCATCTGGAAGTAATTGAT

ACATCTGGAGGGTGTGGTGCAAGTTTTGCAATTGAGATTGTCTCGGAACAATTCGAAGGGAAGAGGCTTC

TGGAGAGGCATAGAATAGTGAATGGTGCTCTGGAAGAGGAGATGAAGGAAATCCATGCTCTCTCAATAAA

GAAAGCTCTGACTCCGGAGCAGTGGAAACAACAGCAAGAATCTGAAAAATCTAAATCAGATGCTTAGAAG

GACACTAAACTCAATTAGACATTTAAATAAGGGTTCATGTAGCCACTTGAGATATTTGAACTCAGATAAC

CTGGATAGAAAATTGCCAATGCTGGAATACTGAAGCATTGTTTTACA

>EE332573.1 EST00069 Jute Plasmid library Corchorus capsularis cDNA similar to Type II keratin E1, mRNA sequence

GTAATCATGTCATAGCTGTTTCCTGTGTGAAATTGTTATCCGCTCACAATTCCACACAACATACGAGCCG

GAAGCATAAAGTGTAAAGCCTGGGGTGCCTAATGAGTGAGCTAACTCACATTAATTGCGTTGCGCTCACT

GCCCGCTTTCCAGTCGGGAAACCTGTCGTGCCAGCTGCATTAATGAATCGGCCAACGCGCGGGGAGAGGC

GGTTTGCGTATTGGGCGCTCTTCCGCTTCCTCGCTCACTGACTCGCTGCGCTCGGTCGTTCGGCTGCGGC

GAGCGGTATCAGCTCACTCAAAGGCGGTAATACGGTTATCCACAGAATCAGGGGATAACGCAGGAAAGAA

CATGTGAGCAAAAGGCCAGCAAAAGGCCAGGAACCGTAAAAAGGCCGCGTTGCTGGCGTTTTTCCATAGG

CTCCGCCCCCCTGACGAGCATCACAAAAATCGACGCTCAAGTCAGAGGTGGCGAAACCCGACAGGACTAT

AAAGATACCAGGCGTTTCCCCCTGGAAGCTCCCTCGTGCGCTCTCCTGTTCCGACCCTGCCGCTTACCGG

ATACCTGTCCGCCTTTCTCCCTTCGGGAAGCGTGGCGCTTTCTCATAGCTCACGCTGTAGGTATCTCAGT

TCGGTGTAGGTCGTTCGCTCCAAGCTGGGCTGTGTGCACGAACCCCCCGTTCAGCCCGACCGCTGCGCCT

TATCCGGTANCTATCGTCTTGAGTCCACCCGGTAAANCCGACTTATCGCCNCTGGCAGCANCCCTGGTAA

CAGGATTACCAAAACGAGNTTGTA

>EE332572.1 EST00068 Jute Plasmid library Corchorus capsularis cDNA similar to LRRG00134, mRNA sequence

TCTACCCGCCGCTCGGTGGGAATTACGCTTCAAGGAGGCCTGCACGGCTCATCCGCCGCACGGACTACTC

CTACGACACGTGCCCTTGGGGGCCAAAGGCCCCTACTGCGGGTCGGCAATCGGGCGGCGGGCGCATGCGT

CGCTTCTAGCCCGGATTCTGACTTAGAGGCGTTCAGTCATAATCCAGCGCACGGTAGCTTCGCGCCACTG

GCTTTTCAACCAAGCGCGATGACCAATTGTGCAAATCAACGGTTCCTCTCGTACTAGGTTGAATTACTAT

TGCGACGCGGCCATCAGTAGGGTAAAACTAACCTGTCA

>EE332571.1 EST00067 Jute Plasmid library Corchorus capsularis cDNA similar to Type II keratin E1, mRNA sequence

GTAATCATGTCATAGCTGTTTCCTGTGTGAAATTGTTATCCGCTCACAATTCCACACAACATACGAGCCG

GAAGCATAAAGTGTAAAGCCTGGGGTGCCTAATGAGTGAGCTAACTCACATTAATTGCGTTGCGCTCACT

GCCCGCTTTCCAGTCGGGAAACCTGTCGTGCCAGCTGCATTAATGAATCGGCCAACGCGCGGGGAGAGGC

GGTTTGCGTATTGGGCGCTCTTCCGCTTCCTCGCTCACTGACTCGCTGCGCTCGGTCGTTCGGCTGCGGC

GAGCGGTATCAGCTCACTCAAAGGCGGTAATACGGTTATCCACAGAATCAGGGGATAACGCAGGAAAGAA

CATGTGAGCAAAAGGCCAGCAAAAGGCCAGGAACCGTAAAAAGGCCGCGTTGCTGGCGTTTTTCCATAGG

CTCCGCCCCCCTGACGAGCATCACAAAAATCGACGCTCAAGTCAGAGGTGGCGAAACCCGACAGGACTAT

AAAGATACCAGGCGTTTCCCCCTGGAAGCTCCCTCGTGCGCTCTCCTGTTCCGACCCTGCCGCTTACCGG

ATACCTGTCCGCCTTTCTCCCTTTCGGGAAGCGTGGCGCTTTCTCATAGCTCACGCTGTAGGTATCTCAG

TTCGGTGTAGGTCGTTCGCTCCAAGCTGGGCTGTGTGCACGAACCCCCCGTTCAGCCCGACCGCTGCGCC

TTATCCGGTAA

>EE332570.1 EST00066 Jute Plasmid library Corchorus capsularis cDNA similar to Type II keratin E1, mRNA sequence

AGGAATCCGTGCAGAGCGGACAAGGTAATCTACTTCCTTAGCCAATTGCCCAGGGCTGAAATCAGGGAGG

TATGAAAGAATCTCGAACTTCTTCTTTCCGGGTGGTGNCCATACCTGGGTGCGTCGCACTTTGCCACCAT

TGGTTACCATAGAAGTAATGTCCTTGTTGTTCTTGCGGNTGGTGGGGAAAGCACAGGCCCACTTGAGGCC

AACGAATGGGGCCACCATGGAGGCCTGTGCGGGGGAGGCACGGATAACCACTGTGGCCGATGACATCATT

GAGGAACCCATTGACCGGCCGCCAATTCCTAAACATGTCCTAAATGTTTCCTGTGTGAATTTGTTATCCG

CTCACAATTCCACCCAAAATAAGAGCCGGAAAAATAAAATGTAAAGCCTGGGGTGCCTAATGAGTGAGCT

AACTCCCGTTAATTGCCTTGCGCTCACTGCCCGGTTTCCAGTCCGGAAACCTGTGGTGCCAGCTGCATTA

ATGAATCGTCCAACCCGGGCGCAGAGGCGGTTTGCGTATTGGGCGCTCTTCGGCTTCCTTTCTCACTGAC

TCCCTGCGTCTCGGTCGTTCGGCTGCTGGGAGCGGTATCAGCTCACTCAAAGGCGGTTATACTGTTATCC

ACAAAATCAAGGGATTACCCAGGAAATAACATGTGACCAAAGGCCAGCTAAAGGGCAGGAACGGTAAAAG

GCCNCGTTGCTGGAGTTTTTCCTAGGGTCCCCCCCTGACGAGGAAAAAAAAAA

>EE332569.1 EST00065 Jute Plasmid library Corchorus capsularis cDNA similar to Tobacco chloroplast 23S and 4.5S rRNA genes, mRNA sequence

TCCAGCGTTTACCGTGGGCACGATAACTGGTACACCAGAGGTGCGTCCTTCCCGGTCCTCTCGTACTAGG

GAAAGGTCCTCTCAATGCTCTAACGCCCACACCGGATATGGACCGAACTGTCTCACGACGTTCTGAACCC

AGCTCACGTACCGCTTTAATGGGCGAACAGCCCAACCCTTGGAACATACTACAGCCCCAGGTGGCGAAGA

GCCGACATCGAGGTGCCAAACCTTCCCGTCGATA

>EE332568.1 EST00064 Jute Plasmid library Corchorus capsularis cDNA similar to Putative senescence-associated protein (Pisum sativum), mRNA sequence

TGGTTAGTTTTACCCTACTGATGGCCGCGTCGCAATAGTAATTCACCCTANTACGAGAGGAACCGTTGAT

TCGCACAATTGGTCATCGCGCTTGGTTGAAAAGGCAGTGGCGCGAAGCTACCGTGCGCTGGATTATGACT

GAACGCCTCTAATTCAGAATCCGGGCTAGAAGCGACGCATGCGCCCGCCGCCCGATTGGCGACCCAAAAT

AAGGGCCTTTGGCCCCCAAGGGGAGGTGTCGTATGAGGAGCCGGTGCGGCGGATGAGCCGTGNAT

>EE332560.1 EST00056 Jute Plasmid library Corchorus capsularis cDNA similar to Oenothera berteriana mitochondrial gene for 26S ribosomal RNA, mRNA sequence

GCGGCCGCTAAGCACCGCGAGGTGTGAAGCGATCTCGTACTAAACGAAGTTCTGTTCAAAAACCTCGTCT

GAGAACTTGTATAGAGAAGGATTTCCCGCTAGCTGCAGTTCTTCCATACCAACTTAGCTGCCCGGCGCTG

CTATTGGCATAACAACCGGTACACCATAGGTTGGCCCAATCCAGTCCTCTCGTACTAGGGTTGGCTCCTC

GAGCGGCCGC

>EE332559.1 EST00055 Jute Plasmid library Corchorus capsularis cDNA similar to NBS-LRR-like resistance protein (Oryza sativa (indica cultivar-group)), mRNA sequence

GTAATCATGTCATAGCTGTTTCCTGTGTGAAATTGTTATCCGCTCACAATTCCACACAACATACGAGCCG

GAAGCATAAAGTGTAAAGCCTGGGGTGCCTAATGAGTGAGCTAACTCACATTAATTGCGTTGCGCTCACT

GCCCGCTTTCCAGTCGGGAAACCTGTCGTGCCAGCTGCATTAATGAATCGGCCAACGCGCGGGGAGAGGC

GGTTTGCGTATTGGGCGCTCTTCCGCTTCCTCGCTCACTGACTCGCTGCGCTCGGTCGTTCGGCTGCGGC

GAGCGGTATCAGCTCACTCAAAGGCGGTAATACGGTTATCCACAGAATCAGGGGATAACGCAGGAAAGAA

CATGTGAGCAAAAGGCCAGCAAAAGGCCAGGAACCGTAAAAAGGCCGCGTTGCTGGCGTTTTTCCATAGG

CTCCGCCCCCCTGACGAGCATCACAAAAATCGACGCTCAAGTCAGAGGTGGCGAAACCCGACAGGACTAT

AAAGATACCAGGCGTTTCCCCCTGGAAGCTCCCTCGTGCGCTCTCCTGTTCCGACCCTGCCGCTTACCGG

ATACCTGTCCGCCTTTCTCCCTTCGGGAAAGCGTGGCGCTTTCTCATAGCTCACGCTGTAGGTATCTCAG

TTCGGTGTAGGTCGTTCGCTCCAAGCTGGGCTGTGTGCACGAACCCCCGTTCAGCCCGACCGCTGCGCCT

TATCCGGTAGCTATCGTCTTGAGTCCAACCCGGTAAAATACGACTTATCGCCACTGGCAGCAGCCCTGGT

AGCAGGATTACCAAATCGAGGTATGTAGGCC

>EE332558.1 EST00054 Jute Plasmid library Corchorus capsularis cDNA similar to Alpha-hemolysin, mRNA sequence

GTTGTAAGGACGGCCAGCTGCATTAATGAATCGGCCAACGCGCGGGGAGAGGCGGTTTGCGTATTGGGCG

CTCTTCCGCTTCCTCGCTCACTGACTCGCTGCGCTCGGTCGTTCGGCTGCGGCGAGCGGTATCAGCTCAC

TCAAAGGCGGTAATACGGTTATCCACAGAATCAGGGGATAACGCAGGAAAGAACATGTGAGCAAAAGGCC

AGCAAAAGGCCAGGAACCGTAAAAAGGCCGCGTTGCTGGCGTTTTTCCATAGGCTCCGCCCCCCTGACGA

GCATCACAAAAATCGACGCTCAAGTCAGAGGTGGCGAAACCCGACAGGACTATAAAGATACCAGGCGTTT

CCCCCTGGAAGCTCCCTCGTGCGCTCTCCTGTTCCGACCCTGCCGCTTACCGGATACCTGTCCGCCTTTC

TCCCTTCGGGAAGCGTGGCGCTTTCTCATAGCTCACGCTGTAGGTATCTCAGTTCGGTGTAGGTCGTTCG

CTCCAAGCTGGGCTGTGTGCACGAACCCCCCGTTCAGCCCGACCGCTGCGCCTTATCCGGTAACTATCGT

CTTGAGTCCAACCCGGTAAGACACGACTTATCGCCACTGGCAGCAGCCACTGGTAACAGGATTAGCAGAG

CGAGGTATGTAGGCGGTGCTACAGAGTTCTTGAAGTGGTGGCCTAACTACGGCTACACTAGAAGAACAGT

ATTTGGTATCTGCGCTCTGCTGAACCAGTTACCTTCGGAAAAAGAGTTGGTAGCTCTTGATCCGGCAAAA

AACCACCGCTGGTAGCGGTGGTTTTTTTGTTGCAAGCAGCAAATACGCGCANAAAAAAGGATCTCAAAAT

ATCCTTTGATATTTTCTACGGGGTCTGAT

>EE332557.1 EST00053 Jute Plasmid library Corchorus capsularis cDNA similar to LRRG00134, mRNA sequence

TGGATTCTACCCGCCACTCGGTGGGAATTACGCTTCAAGGAGGCCTGCACGGCTCATCCGCCGCACGGAC

TACTCCTACGACACGTGCCCTTGGGGGCCAAAGGCCCCTACTGCGGGTCGGCAATCGGGCGGCGGGCGCA

TGCGTCGCTTCTAGCCCGGATTCTGACTTAGAGGCGTTCAGTCATAATCCAGCGCACGGTAGCTTCGCGC

CACTGGCTTTTCAACCAAGCGCGATGACCAATTGTGCGAATCAACGGTTCCTCTCGTACTAGGTTGAATT

ACTATTGCGACGCGGGCCATCAGTAGGGTAAAACTAACCTGTATCACGACGGTCTAAACCCAGCTCACGA

>EE332556.1 EST00052 Jute Plasmid library Corchorus capsularis cDNA similar to Metallothionein-like, mRNA sequence

TCAATCTTCAATAACCCAAAAGCTCTTCCAACTTCTTTCAAAAAACCTTCCTCAATCCACCATGTCGGAC

AAGTGCGGAAACTGCGACTGCGCTGACAAGAGCCAGTGCACCAAGAAGGGAAACACCATGATCATTGAGA

CTGAGAAGAGCTACGTTGCCACCGTGGTGATGGAGGCTCCTCCAGCAGAGAACGATGGAAAGTGCAAGTG

TGGATCTAGCTGCAGCTGCACAAACTGCACATGCGGTGGTCATTAAGCAAACATATATATAGAAATGAAA

CAAATGATCCCATGTTGCAGAAACATGGGCAATATTTGGGACCTAAAATCTTATTTTGGGTTTGAATAAT

TTGTGTCTTTCCTGTTTCTGTTGGTGTCTGTGTTTTCATGTCCTGTCTGTCATGTTTGTGTCTCCTTATA

TACTGTGATGTAATGGCCATGGCCATGCCTTTGGCCTTGTCTCATCTTA

>EE332555.1 EST00051 Jute Plasmid library Corchorus capsularis cDNA similar to Metallothionein-like, mRNA sequence

TATAGTTCATTAAGATGAGACAAGGCCAAAGGCATGGCCATGGCCATTACATCACAGTATATAAGGAGAC

ACAAACATGACAGACAGGACATGAAAACACAGACACCAACAGAAACAGGAAAGACACAAATTATTCAAAC

CCAAAATAAGATTTTAGGTCCCAAATATTGCCCATGTTTCTGCAACATGGGATCATTTGTTTCATTTCTA

TATATATGTTTGCTTAATGACCACCGCATGTGCAGTTTGTGCAGCTGCAGCTAGATCCACACTTGCACTT

TGTCGTTCTCTGCTGGAGGAGCCTCCATCACCACGGTGGCAACGTAGCTCTTCTCAGTCTCAATGATCAT

GGTGTTTCCCTTCTTGGTGCACTGGCTCTTGTCAGCGCAGTCGCAGTCTCCGCACTTGTCCGACATGGTG

AATTGAGGAAGGTTTTTTGAAAGAAGTTGGAAGAGCTTTTGGGTTATTGAAGATTGA

>EE332554.1 EST00050 Jute Plasmid library Corchorus capsularis cDNA similar to Type II keratin E1, mRNA sequence

GTTGTAACGACGGCCAGTGCCAAGCTTGCATGCCTGCAGGTCGACTCTAGAGGATCCCCGGGTACCGAGC

TCGCGTAATCATGTCATAGCTGTTTCCTGTGTGAAATTGTTATCCGCTCACAATTCCACACAACATACGA

GCCGGAAGCATAAAGTGTAAAGCCTGGGGTGCCTAATGAGTGAGCTAACTCACATTAATTGCGTTGCGCT

CACTGCCCGCTTTCCAGTCGGGAAACCTGTCGTGCCAGCTGCATTAATGAATCGGCCAACGCGCGGGGAG

AGGCGGTTTGCGTATTGGGCGCTCTTCCGCTTCCTCGCTCACTGACTCGCTGCGCTCGGTCGTTCGGCTG

CGGCGAGCGGTATCAGCTCACTCAAAGGCGGTAATACGGTTATCCACAGAATCAGGGGATAACGCAGGAA

AGAACATGTGAGCAAAAGGCCAGCAAAAGGCCAGGAACCGTAAAAAGGCCGCGTTGCTGGCGTTTTTCCA

TAGGCTCCGCCCCCCTGACGAGCATCACAAAAATCGACGCTCAAGTCAGAGGTGGCGAAACCCGACAGGA

CTATAAAGATACCAGGCGTTTCCCCCTGGAAGCTCCCTCGTGCGCTCTCCTGTTCCGACCCTGCCGCTTA

CCGGATACCTGTCCGCCTTTCTCCCTTCGGGAAGCGTGGCGCTTTCTCATAGCTCACGCTGTAGGTATCT

CAGTTCGGTGTAGGTCGTTCGCTCCAAGCTGGGCTGTGTGCACGAACCCCCGTTCAGCCCGACCGCTGCG

CCTTATCCGGTAGCTATCGTCTTGAGTCCAACCCGGTAAAACCGACTTATCGCCCCTGGCAGCAGCCCTG

GTACCAGGATTATCAAAACGAGTTTGTAGGCGGGGCTATAAATTTTTGAAAAGGGGCCTACTACGGCTAC

CC

>EE332546.1 EST00042 Jute Plasmid library Corchorus capsularis cDNA similar to Type II keratin E1, mRNA sequence

GTAATCATGTCATAGCTGTTTCCTGTGTGAAATTGTTATCCGCTCACAATTCCACACAACATACGAGCCG

GAAGCATAAAGTGTAAAGCCTGGGGTGCCTAATGAGTGAGCTAACTCACATTAATTGCGTTGCGCTCACT

GCCCGCTTTCCAGTCGGGAAACCTGTCGTGCCAGCTGCATTAATGAATCGGCCAACGCGCGGGGAGAGGC

GGTTTGCGTATTGGGCGCTCTTCCGCTTCCTCGCTCACTGACTCGCTGCGCTCGGTCGTTCGGCTGCGGC

GAGCGGTATCAGCTCACTCAAAGGCGGTAATACGGTTATCCACAGAATCAGGGGATAACGCAGGAAAGAA

CATGTGAGCAAAAGGCCAGCAAAAGGCCAGGAACCGTAAAAAGGCCGCGTTGCTGGCGTTTTTCCATAGG

CTCCGCCCCCCTGACGAGCATCACAAAAATCGACGCTCAAGTCAGAGGTGGCGAAACCCGACAGGACTAT

AAAGATACCAGGCGTTTCCCCCTGGAAGCTCCCTCGTGCGCTCTCCTGTTCCGACCCTGCCGCTTACCGG

ATACCTGTCCGCCTTTCTCCCTTCGGGAAAGCGTGGCGCTTTCTCATAGCTCACGCTGTAGGTATCTCAG

TTCGGTGTAGGTCGTTCGCTCCANGCTGGGCTGTGTGCACGAACCCCCGTTCAGCCCGACCGCTGCGCCT

TATCCGGTAACTATCGTCTTGAGTCCACCCGGTATAAACGACTTTCGCCCTGGCAGCANCCCTGGTAACN

GGATNANCAAACCGAGTTTGTAGGCGGNGCTAAAAATATTGAATGGNGCCTACTACGCTACCTAAAAAAA

TATTGTT

>EE332545.1 EST00041 Jute Plasmid library Corchorus capsularis cDNA similar to Ribulose bisphosphate carboxylase small chain, chloroplast precursor (RuBisCO small subunit), mRNA sequence

AAGCAAGGAATCCATCCAGAGCGGAGAAGGTAGTCTACTTCCTTAGCCAATTGCACGGGGGTGAGATCCG

GGAGGTATGAGAGAGTCTCGAACTTCTTCTTCCCAAGTGGTGGCCATACCTGCATGCATCGAACTCTGCC

ACCATTGCTGACGATGGAGGTGACGTCTTTGTTGTTCTTGCGGGTGGTGGGGAAAGCAGAGGCAGACTTG

AGGCCAACGAATGGGGCGACCATGGAGGCCTGTGCGGGGGAGGCACGGCTAACGACGGTGGCGAATGACA

TCATTGAGGAAGCCATTGCTAGTACTCCTTATGCTAAAGGGCA

>EE332544.1 EST00040 Jute Plasmid library Corchorus capsularis cDNA similar to Chlorophyll A/B binding protein precursor, mRNA sequence

TAGCACACTAAAATTCACATTATGGTCATTGATTTGCAACAAATGTACATCAGACCAAACAAAGAAAGCA

TAAAAGACAAATACATCTCTCTCATCTCATTCTAACTTCTAAGCCCACTTTCCGGGGACAAAGTTGGTGG

CGTAGGCCCAGGCGTTGTTGTTGACTGGGTCAGCAAGGTGGTCGGCCAAGTTGTCCAATGGTCCCTTTCC

GGTGACAATGGCTTGGACGAAGAATCCAAACATGGAGAACATAGCCAATCGGCCGTTCTTGATTTCCTTA

ACCTTGAGCTCAGCGAAGGCCTCTGGGTCATCGGCAAGACCCAATGGGTCGAAGCTGCCACCTGGGTACA

ATGGGTCAACCACCTCACCGAGGGGTCCACCGGCAATACGGTAACCCTCGACGGCTCCCATCAAGATAAC

TTGGCAAGCCCAGATGGCCAAGATGCTCTGAGCGTGGATCAAGCTTGGGTTTCCCAAGTAGTCGAGTCCT

CCTTCGCTGAAGATCTGGGCTCCGGCCTTGAACCACACGGCTTCGCCGAACTTGACGCCGTTCCTGGCCA

AAAGCTCCGGGAAAACGCAGCCGAGAGCTCCCAACATGGCCCATCTGCAGTGGATGACCTCCAGCTCACG

GTTCTTAGCGAAAGTCTCGGGGTCAGCTGAGANACCGGCGGTGTCCNACCGTAGTCACCGGGGAACTCAC

GGGTCAAGTAGAATGGTGGCTCNCCGAAAATGGGCCNAGTAAAAACTCTGTCTGGGCCGTACATGGGCTG

CCGGAGGANCTGAGGCTNGGNGCCTTCCCATGGAAATAGCAATGCCTGGATTAAAA

>EE332543.1 EST00039 Jute Plasmid library Corchorus capsularis cDNA similar to Type II keratin E1, mRNA sequence

GAGTGTAAATGCTGGGGTGCCTAGTGCGCGAAATAACTCACATTTATTGGGTTGCTGCTCACTGCCCGCT

TTCCAGTCTGAAAAACGGTGGGGCCACCTGCATTGGGGCATCGGCCGCCGCGGGAAAAGAGGCGATTTGC

GTATGGTGCGCCCTTCCGCTTCCTCGCTCACTGACTCGCTGCGCGCGGTCGCTCGGCTGCTTGCGAGCGG

GAGCAGGTCCCTCNAAGGCGGTGAANACGGTTATCCATACAATCCAGGGATTACGCGNTTACAGAACATG

NGATTTTTATGGCGGCAGAAGGCCGGGAACCGGAAAAAGGCCCCGTTGCTGGCGTTTTTCCATAGGCGCC

GCCCCCCTGACGAGCATCACCGAATCGACGCTCAANTCANAGGTGGCTTNAACCCGACTGGACTAGNAAG

ATACCANGCGTTTCCCCCTGNAAGCTCCCTCGTGCGCTCCCGTGTTCCGACTTTGCCGCTTAGCGGATAC

CTGTGCGCCNTTCTCGTTCGGCTGGCGCGTGGCTTTCGCATGACTCACGCTGGAGGTATCTCATTTCGGT

GTAAGTCGTCGCTCCAGTTGGGCCGGGTGCAAAAACCCGGGTTAATACTACCCTGCACTTATCGGTACTA

CNACTGAGTCACCGGTAAACCGACTACGCCCTGGAGCACACTGGTACGGATAAATACTAGTTGAAGCGGC

AAAAATTATGAGGGCGCTNCACGCTCCAAAAAATATTTTGN

>EE332542.1 EST00038 Jute Plasmid library Corchorus capsularis cDNA similar to Type II keratin E1, mRNA sequence

TCGGATAATCATGTCATAGCCTGTTTCCNGTGTGAGATTGTTGTGGGCGCACGATTCCACAGAGCATACG

AGCCGGCAAGCATAAGGTGTAAGGCCTGGGGTGCCTAATGAGTGAGCTAGCTCACATTAGTTGCGTTGCG

CTCACTGCCCGCTTTCCAGTCGGGAAACCTGTCGTGCCAGCTGCATTGAGGAACCGGCCAACGCGCGGGG

AGAGGCGGTTTGCGGACTGGGCGCTCTTCCGGTTCCTGGCTCGGTGACTCGCTGCGCTCGGTCGTTCGGC

TGCGGCGAGGGGTATCGGCTCACTCAAAGGCGGTCATACGGTTATCCGCAGAGTCCGGGGATAACGCTGG

ACAGAACATGGGAGCAAAAGGCCAGGGGGGGCCAGTAGCAGGAAAAGGGCCGCGGTTGCTGGCGTTTTTC

CATAGGCGCCGCCCCCCTGACTGAGCATCACAGAAGTCGACGCTCGAGGCAGGAGGTGGTCGGAGCCCGA

GAGGAACTATAAAGGAGACCGGGGCGCGTACCCCAGGAGGGCTCCGAGGTGCGCTCGCCGGGCTCAAACC

CGGCCGCTTGGGGGAGGCGGGTCCCGCTTTGGGCCTGGGGGCAAGGGGGAGCCTTGCTCTGGGACTGCCT

TTGTGCAGTTCGTTTGGAGTTAGGCGGCTGACCCAGGGGGCGGGATCCCAAATGGCCGGTTAAGCCCAAA

GGTGGTGCTATTCAACGTGATGCTGTTGGGCTGTGGCTGCTCGGGGATAAGCGCTTGCGGCCCTGCTGGG

ACAGGGGGGATAGCTCAGTAAACCAACCGGGTAAGGACGGCTCTGGCGGTGTGGTAAGGGCCCCCGAAGT

GGGGCCAGGTAATAACCTTAGCGCCCTTTTTATT

>EE332541.1 EST00037 Jute Plasmid library Corchorus capsularis cDNA similar to 60S ribosomal protein L7a, mRNA sequence

TGCAAAATCCTTGAGGCTATCAAGGCAAACTTCAACGACAAGTACGATGAGTACAGGAAGAAGTGGGGAG

GTGGTATCATGGGCTCCAAGTCACAGGCCAAAACCAAGGCAAAGGAAAGGCTTCTTGCAAAGGAAGCTGC

ACAAAGAATGACTTAGGATTTGTCCCTTGAGATTCTCCTCTAGTCATGGCTTTTTTTGTTTACTATTCTT

GTTTGATTAAATGTTCTTGTGTAGTCAGCTGCAATCAACTTTTGACTCGTATCACTCTTTTTCCAGACTA

ATGGTTATATTTTTCGTAGTTGAACTTATAAAATACTACTACCCAGTAGATTCTGTTTGCGGTGAAGGTC

TGTGCATTTCTAATGAATTACTTTCAATTAAAAAAAAAAAAAAAAAATTTTTTTTTTTTTTTTATGAATG

AAAGGAGCCGCCATAGATCTCCATATCATATATTTATTATGTAAATTTTAATAGCTCAACAAGGCACAAG

CATACAAAGAGTAGTACATTGTAGTAATGAAAAAAGAAAAGGAAAAGACATGTAGTTAATTATAATTAAG

GAAGAACAAAGATGGTTTCACTGGTGATTATTAATTGGTTGATGTTCGTACGGAGAGAGGAGAAGAGAGT

GAATACTGCAGACTCCAGAGCTCGAAAGATCCTTTANCGAAATCGTAGTACCCCCCTTCANNGACAGAGT

TTGTTCNCCATCCANCCNCNCAATGGAGCGNCCCGAATCGNAACANGNCAANCTGTTCCNGGGGNAATGT

TACCGCNCCATNCCCCACANACGACCGGAGCTAAGNNAAGC

>EE332540.1 EST00036 Jute Plasmid library Corchorus capsularis cDNA similar to Ribulose bisphosphate carboxylase small chain, chloroplast precursor (RuBisCO small subunit), mRNA sequence

AAGCAAGGAATCCATCCAGAGCGGAGAAGGTAGTCTACTTCCTTAGCCAATTGCACGGGGGTGAGATCCG

GGAGGTATGAGAGAGTCTCGAACTTCTTCTTCCCAAGTGGTGGCCATACCTGCATGCATCGAACTCTGCC

ACCATTGCTGACGATGGAGGTGATGTCTTTGTTGTTCTTGCGGGTGGTGGGGAAAGCAGAGGCAGACTTG

AGGCCAACGAATGGGGCGACCAGGGTGGCCTGTGCCGGGGAGGCACGGTTAACAACGGTGGCGGATGACA

TCATTGCGGAAGCCATTGCTAGTACTCCTTACTAGTAATGTATATGCTAAAGAGTGAGA

>EE332532.1 EST00028 Jute Plasmid library Corchorus capsularis cDNA similar to Unknown protein, mRNA sequence

CGTTGATTGGGCCACAAAATGGTGATCGCGGTTGGTTGAAAAGCCTGGGCCCCGAAGGTACCGTTCGCTG

GGTTATGACGGAACGCCTCTAAGTCAGAATCCGGGCTAGAAGCGACGCAGGCGCCCGCCGCCCGATGGCC

GACCCGCAGTAGGGGCGCTTTGGCCCCCCAAGGGCAGGGGTAGTAAGTGAAGCGACCGCGGCGGACAATC

CGGCGTCCTTCTTGGAACGGGATAAACAACCGAAATCTGGGTCAGAAGTTTGAAACGGGGCGTGGTGGCC

CGGGGTGAGGAGACCAGGAGGTAGTTCCCCCTGGAGAGAGGGAAGAAGCCCATCCGTGGGGATTGGAAAA

CCAAAGGAGAAGGGGAGTTCAGCGCTGCGGGGAAATGACCAACATGATGAACAATCCCCAAAGGGGAAAA

AGGGCCCACGGGTGCGCCGGCCTCAATACATGGGAGCTGGGCAAGACAGGATCTCTCCAAAGAGGAACGC

GGGGGAGTTGAGGCGCAGGAGCAAGTACTGTCAGCCCGATCCAGTCAGGGGGGGAGGCCATCGGTATCCC

ACTTGAATGGTTAGGCCTGGTAGGTTGGGAGGAATGGTGAGCCCGGGAACAAAATCACACACGGAAGACA

CCTAGGACCTGGAGTGGGGATGGGGGGATCGCA

>EE332531.1 EST00027 Jute Plasmid library Corchorus capsularis cDNA similar to Alnus incana chloroplast 16S rRNA gene, complete sequence, mRNA sequence

TGTAGGGGCAGAGGGAATTTCCGGTGTAGCGGTGAAATGCGTAGAGATCGGAAAGAACACCAACGGCGAA

AGCACTCTGCTGGGCCGACACTGACACTGAGAGACGAAAGCTAGGGGAGCGAATGGGATTAGATACCCCA

GTAGTCCTAGCCGTAAACGATGGATACTAGGCGCTGTGCGTATCGACCCGTGCAGTGCTGTAGCTAACGC

GTTAAGTATCCCGCCTGGGGAGTATGGTCGCAAGGCTGAAACTTAAAGGAATTGACGAAGAGGGGAAAGG

TTCCATGTGAACGGCACTTGCACATGGGTTAGTCGATCCTAAGAGACGGGGGAAGCCCGTCCGATAGCGC

GCAAGCGCGAGCTTCGAAAGGGAATCGGGTTAAAATTCCTGAACCGGGACGCGGCGGCTGACGGCAACGT

TAGGGAGTCCGGAGACGTCGGCGGGGGCCTCGGGAAGAGTTATCTTTTCTTTTAACGGCCTGCCCACCCT

GGAAACGGCTCAGCCGGAGGTAGGGTCCA

>EE332530.1 EST00026 Jute Plasmid library Corchorus capsularis cDNA similar to Chlorophyll A/B binding protein precursor, mRNA sequence

TCGCCGGTCTCTCAGCTGACCCCGAGACTTTCGCTAAGAACCGTGAGCTGGAGGTCATCCACTGCAGATG

GGCCATGTTGGGAGCTCTCGGCTGCGTTTTCCCGGAGCTTTTGGCCAGGAACGGCGTCAAGTTCGGCGAA

GCCGTGTGGTTCAAGGCCGGAGCCCAGATCTTCAGCGAAGGAGGACTCGACTACTTGGGAAACCCAAGCT

TGATCCACGCTCAGAGCATCTTGGCCATCTGGGCTTGCCAAGTTATCTTGATGGGAGCCGTCGAGGGTTA

CCGTATTGCCGTGGACCCCTCGGTGAGGTGGTTGACCCATTACCCAGGTGGCAGCTTCGACCCATTGGGT

CTTGCCGATGACCCAGAGGCCTTCGCTGAGCTCAAGGTTAAGGAAATCAAGAACGGCCGATTGGCTATGT

TCTCCATGTTTGGATTCTTCGTCCAAGCCATTGTCACCGGAAAGGGACCATTGGAGAACTTGGCCGACCA

CCTTGCTGACCCAGTCAACAACAACGCCTGGGCCTACGCCACCAACTTTGTCCCCGGAAAGTGAGCTTAG

AAGTTAGAATGAGATGAGAGAGATGTATTTGTCTTTTATGTTTTCTTTGTTTGGTCTGATGTACATTTGT

TGCAANTCAATGACCATAATGTGAATTTTAGTGTGCTCCAAAAAAAAAAN

>EE332529.1 EST00025 Jute Plasmid library Corchorus capsularis cDNA similar to Putative senescence-associated protein (Pisum sativum), mRNA sequence

TGGATTCTACCCGCCGCTCGGTGGGAATTACGCTTCAAGGAGGCCTGCACGGCTCATCCGCCGCACGGAC

TACTCCTACGACACGTGCCCTGGGGGCCAAAGGCCCCTACTGCGGGTCGGCAATCGGGCGGCGGGCGCAT

GCGTCGCTTCTAGCCCGGATTCTGACTTAGAGGCGTTCAGTCATAATCCAGCGCACGGTAGCTTCGCGCC

ACTGGCTTTTCAACCAAGCGCGATGACCAATTGTGCGAATCAACGGTTCCTCTCGTACTAGGTTGAATTA

CTAG

>EE332528.1 EST00024 Jute Plasmid library Corchorus capsularis cDNA similar to Chlorophyll a-b binding protein, chloroplast precursor (LHCI type II CAB), mRNA sequence

TCTGACATCCTCAAGCCAGGCTGTGTAAACACTGACCCCATTTTCCCCAACAACAAGCTCACTGGGACCG

ATGTTGGGTACCCGGGTGGGCTCTGGTTCGACCCACTGGGCTGGGGAAGTGGTTCACCTGAGAAAATCAA

GGAATTGTGGACAAAAGAGATCAAGAATGGTAGATTGGCTATGTTGGCTGTGATGGGTGCTTGGTTCCAA

CACATTTACACTGGCACTTGTCCTATTGACAACCTCTTTGCCCACCTCGCCGATCCCGGTCATGCTACCA

TTTTTGCTGCTTTCACCCCCAAGTGAGACACAAACAAGGAGAAAACCCTTAACAAGAATTGAATGATTTA

TATATCTTTGCCTAATTAGCAAGGAGATGTACTTTTGTGTGGGGAGGGAATGATGCTTGTATCAGTAGAA

ATCTTAATGGCTCTGTTTGTATTCTATAGATGCTCCAGTGATGTACAAATACCTCTGTGAAATTAGATCC

CATGAATATTTTTATTCATGTTAACTTGTTCA

>EE332527.1 EST00023 Jute Plasmid library Corchorus capsularis cDNA similar to Chlorophyll A-B binding protein 48, chloroplast precursor (LHCII typeI CAB-48) (LHCP). (Zea mays), mRNA sequence

TATTTCATTAAGACTAGCATTACACAGAGATTTAACATTTCTCCATAAGCAATACTAGCCCCAAACAAAT

TCATTACATCATCACAAAACAACCCTCTCAAATTCACACCCAAATTGGAGGATGATTACATTTTTCAAAG

CATTTTAACATTTTCTCACTTTCCGGGGACAAAGTTGGTGGCATAGGCCCAGGCGTTGTTGTTGACTGGG

TCGGCAAGGTGGTCGGCCAAGTTCTCCAATGGTCCCTTTCCGGTGACGATGGCTTGGACGAAGAATCCAA

ACATGGAGAACATAGCCAATCGGCCGTTCTTGATTTCCTTAACCTTGAGCTCAGCGAAGGCTTCTGGGTC

ATCGGCAAGACCCAATGGGTCGAAGCTACCACCTGGGTACAATGGGTCAACCACCTCACCGAGGGGTCCA

CCGGCAATACGGTAACCCTCGACAGCTCCCATCAAGATAACTCGGCAAGCCCAGATGGCCAAGATGCTCT

GAGCGTGGATCAAGCTTGGGTTTCTCAAGTAGTCGAGTCCTCCTTCGCTGAAGATCTGTGCTCCGGCCTT

GAACCACACGGCTTCGCCGAACTTGACTCCGTTCCTGGCCAAAAGCTCCGGGGAAANCGCAGCCGAGA

>EE332526.1 EST00022 Jute Plasmid library Corchorus capsularis cDNA similar to Humulus lupulus 26S ribosomal RNA gene, partial sequence; 18S ribosomal RNA gene, internal transcribed spacer 1, 5.8S ribosomal RNA gene, internal transcribed spacer 2 and 26S ribosomal RNA gene, complete sequence; and 18S ribosomal RNA gene, partial sequ, mRNA sequence

TGGATTCTACCCGCCGCTCGGTGGGAATTACGCTTCAAGGAGGCCTGCACGGCTCATCCGCCGCACGGAC

TACTCCTACGACACGTGCCCTTGGGGGCCAAAGGCCCCTACTGCGGGTCGGCAATCGGGCGGCGGGCGCA

TGCGTCGCTTCTAGCCCGGATTCTGAGAGAGGCGTTCAGTCATAATCCAGCGCACGGTAGCTTCGCGCCA

CTGGCTTTTCAACCAAGCGCGATGACCAATTGTGCGAATCAACGA

>EE332518.1 EST00014 Jute Plasmid library Corchorus capsularis cDNA similar to Type II keratin E1, mRNA sequence

GACGGCCAGTGCCAAGCTTGCATGCCTGCAGGTCGACTCTAGAGGATCCCCGGGTACCGAGCTCGCGTAA

TCATGTCATAGCTGTTTCCTGTGTGAAATTGTTATCCGCTCACAATTCCACACAACATACGAGCCGGAAG

CATAAAGTGTAAAGCCTGGGGTGCCTAATGAGTGAGCTAACTCACATTAATTGCGTTGCGCTCACTGCCC

GCTTTCCAGTCGGGAAACCTGTCGTGCCAGCTGCATTAATGAATCGGCCAACGCGCGGGGAGAGGCGGTT

TGCGTATTGGGCGCTCTTCCGCTTCCTCGCTCACTGACTCGCTGCGCTCGGTCGTTCGGCTGCGGCGAGC

GGTATCAGCTCACTCAAAGGCGGTAATACGGTTATCCACAGAATCAGGGGATAACGCAGGAAAGAACATG

TGAGCAAAAGGCCAGCAAAAGGCCAGGAACCGTAAAAAGGCCGCGTTGCTGGCGTTTTTCCATAGGCTCC

GCCCCCCTGACGAGCATCACAAAAATCGACGCTCAAGTCAGAGGTGGCGAAACCCGACAGGACTATAAAG

ATACCAGGCGTTTCCCCCTGGAAGCTCCCTCGTGCGCTCTCCTGTTCCGACCCTGCCGCTTACCGGATAC

CTGTCCGCCTTTCTCCCTTCGGGAAGCGTGGCGCTTTCTCATAGCTCACGCTGTAGGTATCTCAGTTCGG

TGTAGGTCGTTCGCTCCAGCTGGGCTGTGTGCACGAACCCCCGTTCANCCGACGCTGCGCCTTATCCGGT

AACTATCGTCTTGAGTCAACCCGGTAAAAACGACTTATCGCCCTGGCAGCACCCCTGGTACAGGATAACC

AAACGAGTATGTAGCGGGCTAAAATTAATGAATGGGGCCTACTCGCTACTTAAAAAAAATTTTGAACTGC

CTGGCTAACAATTACTTCGAAAAATTGTATTTTGTCGCAAAAAACTTTGGACGGGTTTTTTTTCACCCAA

TACCCAAAAAGATAAAAACTTTTTTTTCGGGGGACCCGGAAAACCCTAGTTTGGTGAATAAAAAT

>EE332517.1 EST00013 Jute Plasmid library Corchorus capsularis cDNA similar to Alnus incana chloroplast 23S ribosomal RNA (23S rRNA) gene, mRNA sequence

TAGCCGAAAGCATCACTAGCTTACGCTCTGACCCGAGTAGCATGGGGCACGTGGAATCCCGTGTGAATCA

GCGAGGACCACCTTGCAAGGCTAAATACTCCTGGGTGACCGATAGCGAAGTAGTACCGTGAGGGAAGGGT

GAAAAGAACCCCCGTCGGGGAGTGAAATAGAACATGAAACCGTAAGCTCCCAAGCAGTGGGAGGAGCCCA

GGGCTCTGACCGCGTGCCTGTTGAAGAATGAGCCGGCGACTCATAGGCAGTGGCTTGGTTAAGGGAACCC

ACCGGAGCCGTAGCGAAAGCGAGTCTTCATAGGGCAATTGTCACTGCTTATGGACCCGAACCTGGGTGAT

CTATCCATGACCAGGATGAAGCTTGGGTGAAACTAAGTGGTGGTCCGAACCGACTGATGTTGAAGAATCA

GCGGATGAGTTGTGGTTAGGGGTGAAATGCCACTCGAACCCAGAGCTAGCTGGTTCTCCCCGAAATGCGT

TGAGGAG

>EE332516.1 EST00012 Jute Plasmid library Corchorus capsularis cDNA similar to Alnus incana chloroplast 16S rRNA gene, complete sequence, mRNA sequence

TGTAGGGGCAGAGGGAATTTCCGGTGGAGCGGTGAAATGCGTAGAGATCGGAAAGAACACCAACGGCGAA

AGCACTCTGCTGGGCCGACACTGACACTGAGAGACGAAAGCTAGGGGAGCGAATGGGATTAGATACCCCA

GTAGTCCTAGCCGTAAACGATGGATACTAGGCGCTGTGCGTATCGACCCGTGCAGTGCTGTAGCTAACGC

GTTAAGTATCCCGCCTGGGGAGTATGGTCGCAAGGCTGAAACTTAAAGGAATTGACGAAGAGGGGAGAGG

TTCCATGTGAACGGCACTTGCACATGGGTTAGTCGATCCTAAGAGACGGGGGAAGCCCGTCCGATAGCGC

GCAAGCGCGAGCTTCGAAAGGGAATCGGGTTAAAATTCCTGAACCGGGACGCGGCGGCTGACGGCAACGT

TAGGGAGTCCGGAGACGTCGGCGGGGGCCTCGGGAAGAGTTATCTTTTCTTTTAACGGCCTGCCCACCCT

GGAAACGGCTCAGCCGGAGGTAGGGTCCAG

>EE332515.1 EST00011 Jute Plasmid library Corchorus capsularis cDNA similar to Ribulose bisphosphate carboxylase small chain, chloroplast precursor (RuBisCO small subunit), mRNA sequence

AAGCAAGGAATCCATCCAGAGCGGAGAAGGTAATCTACTTCCTTAGCCAATTGCACAGGGCTGAGATCAG

GGAGGTATGAAAGAGTCTCGAACTTCTTCTTTCCAAGTGGTGGCCATACCTGCATGCATCGCACTTTGCC

ACCATTGCTTACAATAGAAGTAATGTCCTTGTTGTTCTTGCGGGTGGTGGGGAAAGCAGAGGCAGACTTG

AGGCCAACGAATGGGGCGACCATGGAGGCCTGTGCGGGGGAGGCACGGCTAACGACGGTGGCGGATGACA

TCATTGAGGAAGCCATTGCTAGTTAGTACTCCTTACTAATAATGTATAAACA

>EE332514.1 EST00010 Jute Plasmid library Corchorus capsularis cDNA similar to Ribulose bisphosphate carboxylase small chain, chloroplast precursor (RuBisCO small subunit), mRNA sequence

TCTCACTCTTTAGCATATACATTACTAGTAAGGAGTACTAGCAATGGCTTCCGCAATGATGTCATCCGCC

ACCGTTGTTAACCGTGCCTCCCCGGCACAGGCCACCCTGGTCGCCCCATTCGTTGGCCTCAAGTCTGCCT

CTGCTTTCCCCACCACCCGCAAGAACAAAAAAGACATCACCTCCATCGTCAGCAATGGTGGCAGAGTTCG

ATGCATGCAGGTATGGCCACCACTTGGGAAGAAGAAGTTCGAGACTCTCTCATACCTCCCGGATCTCACC

CCCGTGCAATTGGCTAAGGAAGTAGACTACCTTCTCCGCTCTGGATGGATTCCTTGCTTG

>EE332513.1 EST00009 Jute Plasmid library Corchorus capsularis cDNA similar to Ribosomal protein S28, mRNA sequence

TGAAAGAAGCGCACTCGTTGCTCGCCGCTACCTTCATCCTCACTCTCTCTAATCACAGAATGGATTCTAC

GATTAAGCATGCTGTTGTTGTGAAAGTGATGGGAAGGACTGGATCAAGAGGTCAGGTGACTCAAGTCAGA

GTCAAGTTTTTGGATGATCGGAACCGTTACATCATGAGGAACGTCAAGGGACCGGTCAGGGAGGGAGATA

TTCTCACCTTGCTTTAGTCTGAAAGAGAAGCTAGGAGGCTTCGTTGATCGGGTCGGGTCTTTTACTATAT

CTTTGTTGATGGTTTTCTGGTATTAGCTAGTGTTATTTTATCCTGAGTTTTGTTACTTAAATCTCATATC

AAGACTTGGATTTTGGGAGAGGATGGATTTACATGTTTTAAAGCTTAATTATGTTGTGGAAATGAACTAA

GTTGA

>EE332512.1 EST00008 Jute Plasmid library Corchorus capsularis cDNA similar to Ribosomal protein L28-like, mRNA sequence

TGCAGCTCTCATCCCTTAAAAAACCCTAAAAATCAACAGGGAAAGAGAAGATGGCGACAGTTCCAGGACA

ATTGATCTGGGAGGTAGTTAAGAAGAACAACTGCTTTTTGGTTAAGGAGTTTGGCAGAGGAACTGCTGGC

GTTCAGTTCAGCAAAGAGCCTAACAATCTCTACAATATCAACTCCTACAAACACTCTGGGTTGGCAAACA

AGAAAACAGTCACCATTCAACGCGCCGGCAACCAGTCTGTGCTTCTAGCAACAACCAAGACTAAGAAGCA

GAACAAGCCTTCAAGCTTGCTTCACAAATCTGTCATGAGGAAGGAGTTCCCCAGGATGGCCAAGGCAGTC

AAAAACCAGGTGACAGACAACTACTACGGGCCAGACTTAACAAAAGCAGCCCTTGCAAGGTTGAGTGCTG

TGCACAAGAGTCTGAAGGTTGCCAAGTCCGGTGTCAAGAAGAGGAACAGACAAGCATTGAAGAAGTGAGA

GAGCTAGGGGGTTTAAGTTGTAGTGTGCTGCTATTTTCATTGTCCAGACATTTTTGGTTTTACTTTTGCC

TTTTTAAATTCTAAATTGTCGTATTTTCCATCAATGGATTCAGCAGGGTGGTGGATCTGTCTAGTTTTTG

AACAAAGAGATCAGTGTCCTTAATAGTATGTTTTTGGATTATAATATGGTAATCTTGATATTGAGCAAAA

AAAAAAAAAAGCGGCCGAAATTCGTATCAGTCAAACTGTTCCCGGGGAAATGTTTCGCTCAATTCCCCAC

ATCGACCGAAGCTAATGTAGCTGGGGGCTATTTGACTACTCAATATGGTGCCTTTGCCCTCCATCGAACG

TTGCCCTCTTAGACGCACCCGGGAGGGTGGGGTGGCTTCTTCCCCAGGCCGCCGGGTGGGGGAG

>EE332511.1 EST00007 Jute Plasmid library Corchorus capsularis cDNA similar to Ribulose bisphosphate carboxylase small chain 1A, chloroplast precursor (RuBisCO small subunit 1A), mRNA sequence

AATTGGAGCACGGATTCTTCTACCGTAAGAACCACAGCTCCCCAGGGTACTACGATGGACGCTACTGGAC

CATGTGGAAACTTCCCATGTTCGGGTGCCCCGACTCATCCCAGGTGTTGAAGGAGCTTGAGGAGGCCAAG

AAGGAATACCCCGAGTCCTTCATCAGAATCATTGGATTCAACAACAAGCGTCAAGTGCAATCCATCAGTT

TCATTGCCTACAAGCCCCCAAGCTTCACCTCAACTTAAATGAACAGAGTTGTTTTCCCCACATTGTAAGG

TTTCGACAAATGCTAAGTACAATTTAAACAAGCCA

>EE332510.1 EST00006 Jute Plasmid library Corchorus capsularis cDNA similar to Predicted protein, mRNA sequence

TTGCAGTTAAAGCACACCAGTTCTTCACAGAGGAAAGCTGGGATACCTTTAAGCAGATTTTTGATACCTA

CATGCTGGAAGCGTCAAAGGAATGGATTGAGACATATGGGGGCAGTTCACTCCTTGAAACTGTGTACAAG

GCTCTTGATGAGGTGGTGAAACTCTCAGAGTGCGAAATTTACAGCTATAATCCAGACTCTGATGCCGATC

CCTTTCTGGAGAAAGGAGCAATATGGTCTTTCAGTTTCTTCTTCTACAATAGAAAGCTTAAGCGAGTAGT

AAGTTTCAGCTTTTCTTGCCTAAGCAATCCGGTGGCCGATGGATTTCTTGCAGACAACTTATGTTACGAG

GAAGATGGTGAGATATTTGATAACATGGACATATGACCACAAACGTGTACATACTGCTTGTACAATGTGT

CAAAACCCTTTGACCATTGTCTGTAGTTACATTACCGTAGTAGGTATCTTGCTCACATGAACTGAGTAGC

TTAGCTTTTCTACTTTGCAGTGCATATGTAGTCGGTAGTTGCTTCCTACATAATAGGAGTATAAATAATG

ATGTCTGTAAGAATGTGTCGAATGTGAGTTCGGATTCATTAAGGTTGAAGACATTCCTTACTACAATGTT

TGTTTCCTATGCTTATTACACTTATATATGTTTGAACAAAAAAAAAAATTCTACCCGCCCCTCGGGGGGA

ATACACTTCAAGGAGGCTGCAAGTTCATCCGCCGCAGGAATACTCTACAAAACGTGCCCTGGGGGGCAAA

GCCCCTACTGGGGTCGGCAACGGGGGGGGGGGCAGCGTCCCTTTATCCAAATCCCAACAAGCAAACTGTT

CCTGGGGAAATGTTTCCCCCCATTCCCCCACAACACCCGAACAAAATGAAACCGGGGGCCAAGAAGACAA

CCAATAATGNTGCCCCCCCCCTCCCTCGGAACGGCCGCCGCATAAAA

>EE332509.1 EST00005 Jute Plasmid library Corchorus capsularis cDNA similar to High mobility group protein 2 HMG2, mRNA sequence

GAATCAGGGGATAACGCAGGAAAGAACATGTGAGCAAAAGGCCAGCAAAAGGCCAGGAACCGTAAAAAGG

CCGCGTTGCTGGCGTTTTTCCATAGGCTCCGCCCCCCTGACGAGCATCACAAAAATCGACGCTCAAGTCA

GAGGTGGCGAAACCCGACAGGACTATAAAGATACCAGGCGTTTCCCCCTGGAAGCTCCCTCGTGCGCTCT

CCTGTTCCGACCCTGCCGCTTACCGGATACCTGTCCGCCTTTCTCCCTTCGGGAAGCGTGGCGCTTTCTC

ATAGCTCACGCTGTAGGTATCTCAGTTCGGTGTAGGTCGTTCGCTCCAAGCTGGGCTGTGTGCACGAACC

CCCCGTTCAGCCCGACCGCTGCGCCTTATCCGGTAACTATCGTCTTGAGTCCAACCCGGTAAGACACGAC

TTATCGCCACTGGCAGCAGCCACTGGTAACAGGATTAGCAGAGCGAGGTATGTAGGCGGTGCTACAGAGT

TCTTGAAGTGGTGGCCTAACTACGGCTACACTAGAAGAACAGTATTTGGTATCTGCGCTCTGCTGAAGCC

AGTTACCTTCGGAAAAAGAGTTGGTAGCTCTTGATCCGGCAAACAAACCACCGCTGGTAGCGGTGGTTTT

TTTGTTTGCAAGCAGCAGATTACGCGCAGAAAAAAAGGATCTCAAAAAGATCCTTNGATCTTTTCTACGG

GGTCTGACGCTCAGTGGACCGAAAACTCACGTTAAGGGATTTGGGTCATGAAATTATCAAAAAGGATCTT

CCCTTATATCCTTTTAAATAAAAATGAAGTTTTAATCAATCTAAAGTTATTTGATAAACTGGGTTGAAAA

TAAAATTTTTATTCATGGGGCCCTTTTTATCGATGGTTATTTGTTAACCAAATGCCGACCCCCCGGGAAA

AAAAAAAAAGGGGGTTTCCTTGCCCAGTTTAATAACCCGAAA

>EE332508.1 EST00004 Jute Plasmid library Corchorus capsularis cDNA similar to Solanum nigrum chloroplast tRNA-Ala, tRNA-Ile, 16S rRNA, tRNA-Val, rps12, rps7, ndhB genes, mRNA sequence

GGGATGCAGCAGTGGGGAATTTTCCGCAATGGGCGAAAGCCTGACGGAGCAATGCCGCGTGGAGGTAGAA

GGCCCACGGGTCGTGAACTTCTTTTCCCGGAGAAGAAGCAATGACGGTATCTGGGGAATAAGCATCGGCT

AACTCTGTGCCAGCAGCCGCGGTAATACAGAGGATGCAAGCGTTATCCGGAATGATTGGGCGTAAAGCGT

CTGTAGGTGGCTTTTTAAGTCCGCCGTCAAATCCCAGGGCTCAACCCTGGACAGGCGGTGGAAACTACCA

AGCTGGAGTACGGTAGGGGCAGAGGGAATTTCCGGTGGAGCGGTGAAATGCGTAGAGATCGGAAAGAACA

CCAACGGCGAAAGCACTCTGCTGGGCCGACACTGACACTGAGAGACGAAAGCTGGGGGAGCGAATGGGAT

TAGATACCCCAGTAGTCCTAGCCGTAAACGATGGATACTAGGCGCTGTGCGTATCGACCCGTGCAGTGCT

GTAGCTAACGCGTTAAGTATCCCGCCTGGGGAGTACGTTCGCAAGAATGAAACTCAAAGGAATTGACGGG

GGCCCGCACAAGCGGTGGAGCATGTGGTTTAATTCGATGCAAAGCGAAGAACCTTACCAGGGCTTGACAT

GCCGCGAATCCTCTTGAAAGAGAGGGGTGCCTTCCGGAACNCGAAACAGGTGGTGCATGGCTGTCGTCAC

TCGTGCCTAGGTGTTGGTTAATACGGCCCGATTCGTATCTGTCTACTGTTCCGGGTAATTGTATCCCTCA

ATTCCCCACTCAACCGAACTAATGAAACCGGGGCAAGAGGACAAACATATAATTGCNCTGCCTTTTTTGA

ACTTTGCCCTCTTTATCCCCCCGGGAGGTTTTTTGG

>EE332507.1 EST00003 Jute Plasmid library Corchorus capsularis cDNA similar to Unknown protein, mRNA sequence

TCACTCATATATTAGCAGAAAAGAAATATATCTTTTCTTCTCTTTCTTTTTTTTCCTTTTTTTCCCTAAA

AGGAGGTGAATACACATAAGATTTTGTATCATTTACATGCCGCTTAATATGTCCATTTTGACTAAATTAT

CATTATTGCAATCAATTTTAAAAAAAAATTAAAGCTCTGGTGTTATCTAGTTCCACTACATTCTGATGCA

AATCTTTTGGCTTTTATGTATAGGTCTATCCTTGTCCTAAAGGGTCTATAAGACATAGGGAAATGGTTAG

AAACCTTGGTGGAAAAGAGCAGTATGTGGTTCTTTCTCACACTTTTGTTGTATGATGATGTTATTCTGTT

TGGTTTTTGGTTGATCACTTGGAAGTTTTAGATAAGTCATAATAAAACTCCAGCTATCTTGAAGATTTGC

A

>EE332506.1 EST00002 Jute Plasmid library Corchorus capsularis cDNA similar to Predicted protein, mRNA sequence

TTGCAGTTAAAGCACACCAGTTCTTCTACAGAGGAAAGCTGGGATACCTTTAAGCAGATTTTTGATACCT

ACATGCTGGAAGCGTCAAAGGAATGGATTGAGACATATGGGGGCAGTTCACTCCTTGAAACTGTGTACAA

GGCTCTTGATGAGGTGGTGAAACTCTCAGAGTGCGAAATTTACAGCTATAATCCAGACTCTGATGCCGAT

CCCTTTCTGGAGAAAGGAGCAATATGGTCTTTCAGTTTCTTCTTCTACAATAGAAAGCTTAAGCGAGTAG

TAAGTTTCAGCTTTTCTTGCCTAAGCAATCCGGTGGCCGATGGATTTCTTGCAGACAACTTATGTTACGA

GGAAGATGGTGAGATATTTGATAACATGGACATATGACCACAAACGTGTACATACTGCTTGTACAATGTG

TCAAAACCCTTTGACCATTGTCTGTAGTTACATTACCGTAGTAGGTATCTTGCTCACATGAACTGAGTAG

CTTAGCTTTTCTACTTTGCAGTGCATATGTAGTCGGTAGTTGCTTCCTACGTAATAGGAGTATAAATAAT

GATGTCTGTAAGAATGTGTCGAATGTGAGTTCGGATTCATTAAGGTTGAAGACATTCCTTACTACNATGG

TTTGGTTCCTATGCTTATTACACTTATATATGTTTGAACAAAAAAAAAAAATCTACCCGCCGCTC

>EE332505.1 JutEST00001 Jute Plasmid library Corchorus capsularis cDNA similar to Ribulose bisphosphate carboxylase small chain, chloroplast precursor (RuBisCO small subunit), mRNA sequence

TATCCGCCACCGTCGTTAGCCGTGCCTCCCCCGCACAGGCCTCCATGGTCGCCCCATTCGTTGGCCTCAA

GTCTGCCTCTGCTTTCCCCACCACCCGCAAGAACAACAAAGACATCACCTCCGTCGTCAGCAATGGTGGC

AGAGTTCGATGCATGCAGGTATGGCCACCACTTGGGAAGAAGAAGTTCGAGACTCTCTCATACCTCCCGG

ATCTCACCCCCGTGCAATTGGCTAAGGAAGTAGACTACCTTCTCCGCTCTGGATGGATTCCTTGCTTG

>JZ532445.1 JEC18 Library of differentially expressed transcripts in jute upon elicitor and drought stress treatment Corchorus capsularis cDNA, mRNA sequence

CCCGTTCCCAGTCTAGACGAAAACTCCCAATGCTGTGAATCCCCTTGTGCTCGGCTCTTTCCCATTGCTT

ACCATTGATGTCTGGGAGCATGCTTACTACCTTGACTTCCAGAACCGAAGACCAGATTACATAAAGACAT

TCATGACCAATCTTGTGTCTTGGGAAGCTGTAAGTGCCAGACTTGAGGCCGCCAAGGCTGCTTCTGCTTA

AGCAAATTTCTGAACAATTTGACTTCAGTGACAGTG

>JZ532444.1 JEC17 Library of differentially expressed transcripts in jute upon elicitor and drought stress treatment Corchorus capsularis cDNA, mRNA sequence

ACCTATATATGAGGTTCACCCATGGCGCCAACAATGGCCTTGAATCGCTGTTAGGCTATTGGAGCCGATC

AAGGAACAGTTCCCCATCCCCTCATATGCAGACTTTACCTTGGCTGGTGTTGTTGCTGTTGGGTTACTGG

AGGACCCGAAGTTCCTTTCCACCCTGGTAGGAGGACAAACCCGAGCCACCACCAGAAGGGGGCGCTTGCT

GATGCTACCAAGGGCTCTGATCA

>JZ532443.1 JEC16 Library of differentially expressed transcripts in jute upon elicitor and drought stress treatment Corchorus capsularis cDNA, mRNA sequence

ACCGCTCGAGTAAGTAAGGAGGTAGAGTCGGTCAAGAGTTTCCAACTTTGCTTACCAAGCATAGAGAGAT

TAAATGCTTCCATATCACAAAAGCCTAGATCTCCATGACTTTTGTGTAGAGTACTCACAAATTTATAATT

>JZ532442.1 JEC15 Library of differentially expressed transcripts in jute upon elicitor and drought stress treatment Corchorus capsularis cDNA, mRNA sequence

CGCCTTCAATGCCTTCAGATGTCAGGAGTCCTCTTTTCAGACCAAACTTTATTCAACACACCAAGGAGCA

AATCTTGTTAACCCCTATGCACTTAACCAAGCCAAGTTCAACTTTGATTTCCAACAGGCCATGCGCAAGA

TGAGCAATCTTGATGTTAAACTTGGCTCTCAAGGTGAAGTTTGTCGGAGTATTA

>JZ532441.1 JEC14 Library of differentially expressed transcripts in jute upon elicitor and drought stress treatment Corchorus capsularis cDNA, mRNA sequence

GTAATTCGGCCAGTCGACGTGAGAGTGGTGGCCGTTAGCAAGACGAAGCCTATTTCTCTTATAACCCAAG

TGTATGAAGCCGGTCACCGTTGTTTTGGTGAAAATTATGTACAAGAGATTATCCAGAAGGCTCCTGAGCT

TCCTGAGGACATAGA

>JZ532440.1 JEC13 Library of differentially expressed transcripts in jute upon elicitor and drought stress treatment Corchorus capsularis cDNA, mRNA sequence

CGTGCCGTAACTGCCGGTGCCTGGAGATCCTGTGCGCCATCCTCTCGGCGCCCCTCGGCGTCTGCTCCGA

CGGCTGCTGCTCCATGGAGTTCTGGATCAGCGTGCTGCTCACCATCCTCGGCTACCGTCCTCTACGCCGC

CTACGTCATCTGCTCCGTCGACCCCGACCGCGTCCGCCGCCGCGACGACGACTACATCTACTGCCCGCAA

>JZ532439.1 JEC12 Library of differentially expressed transcripts in jute upon elicitor and drought stress treatment Corchorus capsularis cDNA, mRNA sequence

TAATAAGATGGTTAGGGCAAATATAGTGAGCATGGTTCTGTTGATGCATGCCATCGTTGGGTTTCCTTTT

CATGCGAGGGGATTAAGTATGACTTACTACATGATGAGCTGTCCTTTCGCTGAACAGATTGTGAAGAACA

GTGTTAACAATGCTCTTCAAGCCGATC

>JZ532438.1 JEC11 Library of differentially expressed transcripts in jute upon elicitor and drought stress treatment Corchorus capsularis cDNA, mRNA sequence

CCGTTCCCAGGCGGCTCCGATGGCTCATGACACAAAGGACAACCGCCTCAACCCCTGGTTGGGTTTCCGA

GAAATTTTCCACAGGATTGGGAGGCAAGAAAGCTGAGGCAGAAGAAGCGGCTGAATCTGCGACGGGTGGA

TCCGCCTATGACTTTGCGGGTTACGCCTTGGACAATGCATCAGACAAAGCCGGTTTCTGC

>JZ532437.1 JEC10 Library of differentially expressed transcripts in jute upon elicitor and drought stress treatment Corchorus capsularis cDNA, mRNA sequence

AAGGATCCATGGCTGCTTCAGTAACCGCCTACGTCCTAGCCACCTCCATACTCTGATGCTTTGAGCCGCA

TATGAGCACTTTCACTGGAGTTTCATGGGGAAAACATCACAGGGCTTATGTGGCAACCTCAAGAACAGGT

TCTGGATCCGAGCTTGAAGGAAAGCCCTTGGAGCATATCATCCAAAACACTTACAACACGGCGACCTGCT

GGGTCTTCAACAACGCT

>JZ532436.1 JEC9 Library of differentially expressed transcripts in jute upon elicitor and drought stress treatment Corchorus capsularis cDNA, mRNA sequence

CTTCGTGTTATTAGTCATGTAAATAGAGGAAGCAGGCTTTGGTTGGGTACCTTCTCTTCTGCCCAGGAAG

CTGCTCTTGCCTATGATGAAGCTGCTAGAGCTATGTATGGTCCTTGTGCACGCCTCAATTTTCCCGGCAT

CACAGATTATGCTTCTTTTAAGGAATCGTTGAAGGAATCTCCGATGGCCGCATCGTC

>JZ532435.1 JEC8 Library of differentially expressed transcripts in jute upon elicitor and drought stress treatment Corchorus capsularis cDNA, mRNA sequence

GCTTGATTGCCCCATGTTATGAGCCCAGGAAGCCCAACTGGATCTGATGTCAGATTCAAGTACCGATATT

ACCTTCCGTTGAGCTTTCAAGCCTGTGCCTAGACCGGTGCTGTTGTGCTACCGCTTCCTATCGAAACGTC

GTCTCTTCCGATGATCCACCGACTTCGTTAAGCTTGTCACTTCCTGGTGCCG

>JZ532434.1 JEC7 Library of differentially expressed transcripts in jute upon elicitor and drought stress treatment Corchorus capsularis cDNA, mRNA sequence

ATTATCGGAAAGTGCTCCAAGTTTCATCATCGTCATCATCACCATGGGAAAGTGCTACCCTGTTGTGAGT

GAGGAATATCAGAAAGCCATTGAAAAGGCTAAGAGAAAGCTCAGAGGATTCATCGCTGAGAAGAATTGTG

CTCCTCTAATGCTTCGTCTTGCATGGCACTCTGCTGGAACCTTTTGCAAGGATTCCAAGACTGGTGGTC

>JZ532433.1 JEC6 Library of differentially expressed transcripts in jute upon elicitor and drought stress treatment Corchorus capsularis cDNA, mRNA sequence

AAAGTTATGTTACAGTTATTTCTAAGCAAGAAGTTCACCGATTGTGAAACTCGGTATACAATGCTTGAAA

AGACTTGTTGTGCTCTAGCTTGGGCTGCCAAACGCCTGCGTCATTATTTGGTGAATCATACAACTTGGCT

GATATCCAGAATGGATCCGA

>JZ532432.1 JEC5 Library of differentially expressed transcripts in jute upon elicitor and drought stress treatment Corchorus capsularis cDNA, mRNA sequence

CGCAGATATAGTTGCCATGGCTGCTAGAGATGCTGTCTTTTGGGCTGGTGGTCCATATTATGACATACCG

AAAGGAAGGTTTGATGGTAAAAGATCAAAGATAGAAGATACAAGAAATCTACCTTCACCTTTTCTTAATG

CCTCTCAACTCATTCAAACTTTTGGCCAACGTGGTTTCACTCCACAAGATGTTGTTGCTCTCTCTGGAGC

CGT

>JZ532431.1 JEC4 Library of differentially expressed transcripts in jute upon elicitor and drought stress treatment Corchorus capsularis cDNA, mRNA sequence

GCTTTCTGGACATACTCGCTGTGACGGCGCTGCGGTCGGTGCTCCACCGTGTCCGTCAGGCTGCTGAAAG

GTCCGGTCGCCGTGCTGATGACGTGAGAGTGTTGGCCGTTAGCAAGACGAAGCCTATTTCTCTTATTACC

CAAGTGTATGAAGCCGGTCACCGGTGTTTTGGTGAAAATTATGTTCAAGAGATTATCCAGAAGGCTCCTG

AGCTTCCTGAGGAC

>JZ532430.1 JEC3 Library of differentially expressed transcripts in jute upon elicitor and drought stress treatment Corchorus capsularis cDNA, mRNA sequence

TGCGCCATCCTCCTCCCGCCCCTCGGCGTCTGCCTCCGCCACGGCTGCTGCTCCATGGAGTTCTGGATCA

GCGTGCTGCTCACCATCCTCGGCTACCTCCCCGGCGTCCTCTACGCCGCCTACGTCATCTGCTCCGTCGA

CCCCGACCGCGTCCGCCGCCGCGACGACGACTACATCTACGTCGCC

>JZ532429.1 JEC2 Library of differentially expressed transcripts in jute upon elicitor and drought stress treatment Corchorus capsularis cDNA, mRNA sequence

CTCCATCCTGACAAAGGGAAGATATAGTTGCCATGGCTGCTAGAGATGCTGTCTTTTGGGCTGGTGGTCC

ATATTATGACATACCGAAAGGAAGGTTTGATGGTAAAAGATCAAAGATAGAAGATACAAGAAATCTACCT

TCACCTTTTCTTAATGCCTCTCAACTCATTAAACTTTTGGCCAACGTGGTTCACTCCACAAGATGTTGTT

GCTCTCTCTGGAGCACATACCCTAGGAGTTGCACGATGCTCCTCCCAAGGCTAGACTTACCGTCCCAGAC

TCTT

>JZ532428.1 JEC1 Library of differentially expressed transcripts in jute upon elicitor and drought stress treatment Corchorus capsularis cDNA, mRNA sequence

TTTATCAAAATCTAAAGATGAAGCAAGAAAAGCGGCTCCAAGCCGCGGAAAACTATGCTTATGACAAGGC

GGGTTACGTCATGGACTCAGCTTACGACAATGCAGGTTACGCCAAGGATTTTACTGAGAACAAGGCAGAG

TACGCAAAGGATTTCGCTTATGACAAGGCTGGTGATGCAAAGAACATGAAAATTT

>JK743820.1 JutESTSSH3(N78) Jute Plasmid Suppression Subtractive cDNA library Corchorus capsularis cDNA similar to beta-tubulin, mRNA sequence

ATAGTTCTTTCTTCTGCATGCTTTACTGAAATTAACTCGACGATCTTAAAGAAATTAAAACTCAAATGCA

AAGCAAAAACTACAGCATAATGACTAATTCAACTGTTTCACATTCCAAACAAACCAGCAACCGGAAGTTG

TGGTTTTGAATGGCCAATAACCAGTCTCAAATCATTTAGTCATCATATCCTTCTGGTTCCTCTTCCTCAC

CATACTCATCTTCATCAGCTGTTGCATCTTGGTATTGCTGATACTCAGAAACCAAATCATTCATGTTGCT

CTCAGCCTCAGTAAACTCCATTTCATCCATACCTTCCCCGGTATACCAATGCAAGAAAGCCTTCCTCCTA

AACATAGCTGTAACCTGATCACTCACACGCCTGAACATCTTGTGAAAAGATGTTGAGATGCCAAGAATGA

GATGCCATCTTTGAGGACTGTAAG

>JK743819.1 JutESTSSH3(N72) Jute Plasmid Suppression Subtractive cDNA library Corchorus capsularis cDNA similar to cationic peroxidase, mRNA sequence

TGAAGGAAACAGACAGGAGCTTTGGCCTTAGGAATTTCAGATACATTGAAGACATCAAAGAAGCTGTCGA

AAGGGAGTGTCCTGGAGTTGTTTCTTGCGCTGATATCCTTGTGCTCTCTGCAAGAGAAGGAATTGTTTCG

TTGGGAGGCCCTTACATCCCTCTAAAAACTGGAAGAAGAGATGGCAGAAAAAGCAGAGCAGATGTTCTAG

AAGAGTATCTTCCAGACCACAATGAGTCCATGTCTGTCGTCCTTGACAGATTTGCTGCAATCGGAATCGA

CACCCCTGGAGTCGTTGCCCTGCTAGGATCTCACAGTGTCGGTCGAACCCACTGTGTCAAACTGGTGCAC

CGTTT

>JK743818.1 JutESTSSH3(N71) Jute Plasmid Suppression Subtractive cDNA library Corchorus capsularis cDNA similar to cationic peroxidase, mRNA sequence

TGAAGGAAACAGACAGGAGCTTTGGCCTTAGGAATTTCAGATACATTGAAGACATCAAAGAAGCTGTCGA

AAGGGAGTGTCCTGGAGTTGTTTCTTGCGCTGATATCCTTGTGCTCTCTGCAAGAGAAGGAATTGTTTCG

TTGGGAGGCCCTTACATCCCTCTAAAAACTGGAAGAAGAGATGGCAGAAAAAGCAGAGCAGATGTTCTAG

AAGAGTATCTTCCAGACCACAATGAGTCCATGTCTGTCGTCCTTGACAGATTTGCTGCAATCGGAATCGA

CACCCCTGGAGTCGTTGCCCTGCTAGGATCTCACAGTGTCGGTCGAACCCACTGTGTCAAACTGGTGCAC

CGTTT

>JK743817.1 JutESTSSH3(N68) Jute Plasmid Suppression Subtractive cDNA library Corchorus capsularis cDNA similar to SKP1 component-like 1, mRNA sequence

AAAGGCAATTAAAAAGAAAAAGAAACATAGAAAAAAGATATGTTCAAATTCAATTCGATATCCTTCACTC

AAAGGCCCACTGATTCTCCCGACGAACCTCCTCCTCCTCTTCCGGAGTAAAATCATTCTTAATATTGAAA

GTCTTACGGATCTCCTCAGGCGTTTTCCCCTTTATCATATCCGCCACCGTTTGACACGTCAGGTCCAGCA

GGCTCTTGATGTTCAAATAGTTAGCAGCCAGGATGAGATCGAAGAGGGTAGCTTGATCGACCCTGGCGAA

ATCAGCATCCCAAAGCTTAAGCTCATCTTCGACGGAGGCAGAACGATCGTCGGATTTAGGTGCTTCCACA

TGCTTCTTGCA

>JK743816.1 JutESTSSH3(N61) Jute Plasmid Suppression Subtractive cDNA library Corchorus capsularis cDNA similar to 26S ribosomal RNA, mRNA sequence

TGGAAATCAGAATCAAACGAGCTTTTACCCTTTTGTTCCACACGAGATTTCTGTTCTCGTTGAGCTCATC

TTAGGACACCTGCGTTATCTTTTAACAGATGCGCCGCCCCAGCCAAACTCCCCACCTGACAATGTCTTCC

GCCCGGATCGACCGACAGATGCCGGTCTTGGGTCTAAAAAGAGGGGCGTAGCCCCGCTTCCGATTCACGG

AATAAGTAAAATAACGTTAAAAGTAGTGGTATTTCACCGTTGCCGAAGCTCCCACTTATACTACACCTCT

CAAGTCATTTCACAAAGTCGGACTAGAGTCAAGCTCAACAGGGTCTTCTTTCCCCGCTGATTCTGCCAAG

CCCGTTCCCTTGGCTGTGGTTTC

>JK743815.1 JutESTSSH3(N60) Jute Plasmid Suppression Subtractive cDNA library Corchorus capsularis cDNA similar to alpha-tubulin, mRNA sequence

TTACCGCCATCTCTTCCACCCAGAGCAACTCATCAGCGGCAAGGAAGATGCTGCCAACAACTTTGCTCGT

GGTCACTACACCATTGGCAAGGAAATTGTTGATCTCTGCTTGGACCGTATCAGGAAGCTTGCTGATAACT

GCACAGGTCTCCAAGGGTTCCTTGTTTTCAACGCTGTTGGTGGAGGAACTGGATCTGGTCTTGGTTCTCT

TCTGTTGGAGCGTTTGTCTGTTGACTATGGCAAGAAATCCAAGTTGGGTTTCACTGTCTACCCCTCTCCC

CAGGTCTCCACCTCTGTTGTTGAGCCCTACAATAGTGTTCTCTCAACTCACTCCCTATTGGAACACACTG

ATGTGGCTGTTCTCCTCGACAATGAGGCTATCTATGACATCTGCAGGCGTTCTCTTGACATTGAGCGACC

CACCTACACCAACCTTAACCGTCTCGTGTCTCAGGTGATTTCCTCCTTGACTGCCTCACTCAGGTTTGAT

GGTGCATTGAACGTGGATGTGACTGAATTCCAGACCAACTTGGTCCCATACCCAAGAATCCACTTCATGC

TTTCCTCATATGCTCCAGTCATCTCTGCCGAGAAAGCTTACCATGAACAGCTCTCAGTTTCTGAAATCAC

CAACAGTGCCTTTGAGCCTTCATCTATGATGGCTAAGTGTGATCCTCGCCATGGCAA

>JK743814.1 JutESTSSH3(N59) Jute Plasmid Suppression Subtractive cDNA library Corchorus capsularis cDNA similar to 40S ribosomal protein S24, mRNA sequence

TGTTATACTAAATATTTCAAAATTCTCATCTGGTAGAATTTACTCAAAACAAAGACAATCAGAGAATATA

ACTAACCATCGAATAACACCAAAATCCAGTTTCCACAAAAACAGGTTCTAAGAACACCTAAACATTTCAG

ACTAGTAATAATAGTAAAGAAATGCAGTTGAAAACCAGTGGCTCTAAGATAAGCACGAACTTCAATTCTT

TCCCCTCTTTTAGCTTACTTCTTCTTTGCATCACCGGCCTTGGTCTTCTTAACTCCGCGGACCTTCTTAG

ATCTGTTCTTCCTTTCCTTCATTTGTTTCCTTGACTTCTCTACCTTGGTGTCAAGTCCATTCCTGATAAG

CCT

>JK743813.1 JutESTSSH3(N58) Jute Plasmid Suppression Subtractive cDNA library Corchorus capsularis cDNA similar to methionine synthase, mRNA sequence

TTTGGTGAGCAGTTGTCTGGTTTTGCTTTCACTGCTAATGGGTGGGTGCAATCTTATGGTTCTCGCTGTG

TCAAGCCACCAATCATCTATGGTGATGTTTGCCGCCCCAAGCCAATGACCGTCTTCTGGTCATCAACCGC

CCAGAGCATGACCTCTCGCCCAATGAAGGGAATGCTTACTGGCCCAGTTACCATCCTCAACTGGTCTTTT

GTCAGAAATGATCAACCTAGATTTGAGACTTGCTACCAGATCGCCTTGGCCATCAAGGATGAAGTCGAGG

ATCTTGAGAAGGCCGGTATCAATGTTATCCAAATTGATGAGGCTGCTTTGAGAGAAGGTTTGCCTCTTAG

GAAGTCTGAGCATGCTTTCTACCTGAAATGGGCTGTCCACTCCTTCAGGATTACCAACTGCGGTGTCCAG

GACACTACCCAGATCCACACACACATGTGCTACTCCAACTTCAATGATATCATCCACTCAATCATTGACA

TGGATGCCGATGTCATCACCATTGAGAACTCACGTTCAGATGAGAAGCTCCTGTCAGTCTTCCGTGAAGG

AGTCAA

>JK743812.1 JutESTSSH3(N57) Jute Plasmid Suppression Subtractive cDNA library Corchorus capsularis cDNA similar to ATP-dependent subunit of the HslUV protease, mRNA sequence

AGAATAAGTTCATGCGACAATTCCATGAAATGACTCAAATGAAACTGTTTTCTTCCCGTCTGCAGGTGAG

AGACAGGGGGAGAGGGAGAAGGTAGGAAAATAAACCAAAGCAACAGTCAGGAACTTAAAATTATTTTCCT

AAATTCAGTTTTCTCCATGAGAAACACTCTAGTGTCAACGTTTACAAGAACAAGCTTAGTCACATAAATG

ACATATACTAACATAGTTCGAGAGACCAGAGAGATGGGAACTAAATACTAAAACAAAAACAATCTACTTT

AAAATGCAGGAGAGAAAAAAGTTGTCCTTTCATCCAACTGAAAAGGAGAAGACTTATTTCATGCTCTCTT

GGCAAAGTCATTGGTATCATCCTCATCCCGGAAGTTCTCATTCAGTGGATCATTTTGGAAACCTTCTGGT

TCATCTTTGAAGCTACCTTCACCAGTATCTTGATGATCAGCACCACCAAATCCCATATTACCACCTCCAA

ATCCACCTGAACCACCAGTTCCAAAACCATCACTGCCGCCAACCTCCACCAGCAAACATTGAAGACCTCG

GCCGCGACAAGCTA

>JK743811.1 JutESTSSH3(N56) Jute Plasmid Suppression Subtractive cDNA library Corchorus capsularis cDNA similar to 70 kDa heat shock cognate protein 2, mRNA sequence

ATCTTAGCAATGATGGGGTTGCAGATACTCTCCAACTCCTTCATCTTGTCTTCGAATTCATCAGCCTCGG

CGAGCTGGTTGCTGTCGAGCCATTGGATGGCAGCCTCAATTGCATCCTCAATCTTCTTCTTGTCTGCTGG

AGTGAGCTTGGCACCGATCTTCTCATCCTTCACTGTGTTTCTCATGTTGTAAGCGTAGTTCTCGAGTGCA

TTCTTGGCCTCAACCTTCTTTTTGTGCTCCTCATCCTCAGATTT

>JK743810.1 JutESTSSH3(N55) Jute Plasmid Suppression Subtractive cDNA library Corchorus capsularis cDNA, mRNA sequence

TATTTTTCCCTATAAAGCCAAACTAACAACTTCCAATATCCTTTTTCTTCTAAAATTCTTCCCATTTCCT

AAAAAACTTCCAAAATGGTGTAAAAAAGAACCACCCATATCATATCAGTATGTCCACTCCTGGGGTAACC

GAACTTCATGACTCATCCCTTGTTGTGGCCTGAAATTGCGCCTACTCCGAACCCTCAAAGCAGCAGCATC

ACTATCAGAAGTGAAACTTGCATTAAAACGAGGGTGAAGCTGGGGCTGAGCATTAATCTCATCAAGATTC

AAGTTCAAGGCCTGAACAACTCTGGCTGGAAGCAGAACAGTAGAACAACCTGTCTTCTTGCGGGTTTCAG

TAGGGGCACCAACACGACGAGGCAGAAAAACACCAGTTCCAGCACATTCCCGTTTACCAGCTGGGTTACC

CAAGAAAACAGCTCGCATACCCGACCCGTTTTGAGACTGGGTTTGTTGTTGCTGCTGTTGCAGAGGGGGC

CATGCAGATGGAGACAAACCCAAAGAAGGCCTATTGTTGCTATTACTGTTACCATTTCTCCCTCTGTTTT

GGACCACATGGTGGTTGTTCTGATGCTGCTGCTGCTTCTGTCCGCCCCAAACCGATGCATTTTGTTGTTT

CATTAGCTGCTGCTGTTGTTGCTTCAGTTGTTGAAATTGAGTGGTCTGCAACTTCTGGTGATGAGAAAAC

GACTGGTGAGAAGGAGGGTAATAGCTTGAAACATCGAGGTTTGGAGATGGTTTTCTTGCTGGTGGACCCA

AAAGGGCCTCTGTTATTGGAATCCACCGTATGATTCTTCATTCATTCGCATTCGTGCCTACTTCCCCTGC

AGCGCGTATAGCAAGATTCCAATGTTCCCGGCGGCGACCAAACCTCGGTGAATGACAGTTGGGCCTTCTC

GCCGCGAACCACGCCTAATTCTAGATTGCATCGCGAG

>JK743809.1 JutESTSSH3(N54) Jute Plasmid Suppression Subtractive cDNA library Corchorus capsularis cDNA similar to adenine nucleotide translocator, mRNA sequence

TTCAATATATATATCCTTTAATAAAACCAAAACTCCGAAATGGGTGGTGGCATTAGCCGCCAATCTTATA

AGGTTTTTTTCAATCAAAACTTCCATCGGAAAAATAAAAATGGTTTGGAGAGTAACCCCCCCCTAAACTG

AATTATTTAATGAAATTTTACCGAAATAAAATTTCTCGGGAGTCAAAGTTTTTTTCATCACCATCTAAAA

CCTAAACGGAACAAAAAGGGAGAGTGTAGCCCCAAAGCTTAAGCACCACCGGATCCATACTTCTTTCCGA

ACACAATGAGCTGCAACTTGTCATATCCAGCAAGCACACCAGCACCAGCAATAGCACGAAGGATGTTAGC

TCCAGCACCCTTAAATAGAGACTTCGCACCCTCATTTTTCAAGATCTGAGAAAATGCATCAAGGGAGCTC

TTGTATTTGACTGCTTCACCAGAGGTCATCATCATTCTTCTACGAACAGTATCAATTGGGTAGGATGCAA

GGCCAGCACCATTGGTAATGAGCCAACCGAGAGCAAAGCTAGCAAAGAAACTATCCTGCATGTTTCCGGT

CAAAAGCACAGGCTTCAGAGAGTCATACATTCCAAA

>JK743808.1 JutESTSSH3(N53) Jute Plasmid Suppression Subtractive cDNA library Corchorus capsularis cDNA similar to senescence-associated protein, mRNA sequence

GAATACCAACCGTGGAAGCGTGGCCTATCGATCCTTTATACCTTCGGAATTTGAAGCTAGAGGTGTCATA

AAAGTTACCACAGGGATAACTGGCTTGTGGCAGCCTTCCGTTCATAGCGACGTTGCTTTTTGATCCTTTG

TGTCGGCTCTTCCTATCATTGTGAAGCAGAATTCACCATGTGTTGGATTGTTCACCCACCAATACGGAAC

GTGAGCTGGGTTTAGACCGTCGTGAGACAGGTTAGTTTTACCCTACTGATGGCCGCGTCGCAATAGTAAT

TCAGCCTACTACCTCGTCCGCGACCACGCTA

>JK743807.1 JutESTSSH3(N52) Jute Plasmid Suppression Subtractive cDNA library Corchorus capsularis cDNA similar to Alanyl-tRNA synthetase mitochondrial precursor, mRNA sequence

TGAAAACCATTATTGGCATTTTCTTCTGCGCCATAACTTTTGAAACTGCTTCACGGAGAGCAGCAGCATC

TAGACCAACATCAATACGGGAGACACAGAAGGTCTTTCCCTCTGAGGCAGCAACTTCAGCCAATTCAATT

GCTTTCTTGACAGCCTTCTGCATATTTTCTTCAGCAAGCTTTTTTTGTGCCTTCTTCAGTTGATTCTGAA

GTTGAGCAATCTTGGCCCTAATATCAGCTTTCTTGGCAGCAGGAATTGATGCTGCATCCACACGAGTTTT

TAAGGAAGCCACTGTCTTCTCCAACAAGCTTACTTCCATCTTGGATGCATCATCTACTTCTTTCAAAAGC

TGGTCAGCCAATTCCATTGCCTTCAAAGCATCCTCAGTAGTGACAGCAGTGATCCTTCTAACTCCTTTAG

CAATTCCCTCCTCAGATAAAAGAGCAAATGCCTTTGCTTCCCGGGTATTTGTTATGTGT

>JK743806.1 JutESTSSH3(N51) Jute Plasmid Suppression Subtractive cDNA library Corchorus capsularis cDNA similar to Ribosomal protein L19, mRNA sequence

TGCCACAAACATTATCTTCAAATATAACAGCATAGGGGTTTCCAAAATTCCGTAATTTATAGTAATTAAA

ACTTTTAACATCAAAGAAAACAGAGTTTTGCAAACAAACTCCTACAGTGAAAACAAAAGGCATTAACATA

TAAAGTTTATCCATCTTCACTACTTGGTTCACTTCTTGGACTTCTTACCAACCTCGGCCTGTTGGGGTGC

GGGTGTGGGCACAGGAGGTGCTGCCTTCTCACCACCAGGTCCCTGAGCAAGCCGTTCTTCCCTTCTGGCC

TGCTTTCTCTCCCTGCTAGCCTTGTTCTTTGCACGCTTGGCCTCAAACTGGTCGGAAAGAGTCTTCTCTC

TAGCCTTCTCAGCCTTGGACTTGTGAATGCTTTCCATCAAGACACGCTTGTTCTTGAACACATTACCCTT

AACCTTCAT

>JK743805.1 JutESTSSH3(N48B) Jute Plasmid Suppression Subtractive cDNA library Corchorus capsularis cDNA similar to MAP kinase kinase kinase Ste11/SteC, mRNA sequence

AAGCAAACAAAGGCAAAACCAGAAACCGTGCTCGTTCCCTCCCTTAATCTGCATGTAAACGCATCTGAGG

TTGTTCAAGTTCATATTAATTTGGCCCTCCAGCAGGCAACCAAAAAAATCAGCCGAACCTTG

>JK743804.1 JutESTSSH3(N48A) Jute Plasmid Suppression Subtractive cDNA library Corchorus capsularis cDNA similar to Pectin methylesterase, mRNA sequence

TGATTCTTTCCAGAAAAAAAGGAGATGTGCCCATGGGAATATACATGGGGCTTTTTTTCTAATTACTTTC

TGTAAATTTATATATGTATGAAATATATATGAATTAAGTAGCTCTTGGAGATCTTG

>JK743803.1 JutESTSSH3(N47) Jute Plasmid Suppression Subtractive cDNA library Corchorus capsularis cDNA similar to senescence-associated protein, mRNA sequence

AAACATCACTCCATAAACAAAAACAACTTCAACATAATCCACTGAAACAAACTAAAACATGATAGTCATT

GCCGAAACACTTTTCATTCCAAGTTCATAGACAAGACTAGTAATGGAAAGTGTCCTACTACTGCTGTTTG

ATTTCAGCTTTCTTCTTTTGCCAAAGCCTGTCAAGAGCGCTGTCGGCATCACCCTGAGCACGGACATTGC

CGCAAAGGGCG

>JK743802.1 JutESTSSH3(N46) Jute Plasmid Suppression Subtractive cDNA library Corchorus capsularis cDNA similar to senescence-associated protein, mRNA sequence

GAATACGAACCGTGAAAGCGTGGCCTATCGATCCTTTAGACCTTCGGAATTTGAAGCTAGAGGTGTCAGA

AAAGTTACCACAGGGATAACTGGCTTGTGGCAGCCAAGCGTTCATAGCGACGTTGCTTTTTGATCCTTCG

ATGTCGGCTCTTCCTATCATTGTGAAGCAGAATTCACCAAGTGTTGGATTGTTCACCCACCAATAGGGAA

CGTGAGCTGGGATTAGACCGTCGTGAGACAGGTTAGTTTTACCCTACTGATGGCCGCGTCACAATAGTAA

TTCAACCTA

>JK743801.1 JutESTSSH3(N45) Jute Plasmid Suppression Subtractive cDNA library Corchorus capsularis cDNA similar to Ubiquitin-like protein SMT3, mRNA sequence

TACTAAGAACATGATTTCAAACAAATTAGAACACAGCCAAAGCAATTGTTCCACCAGTTTGGTGCAGCAT

AGCATCAATCTCATCACCATCCTCCATTTCCAGCTCATCAGGAGTTTGCTCTGCTCGCAGACGGCGACCA

TCAAACAAGAAGGCAATCGAATTGAGCTCCACAGATTGTCGATCACAATATGCATTCATAAGCTTCTTCA

GCTGTGTGCTTCTTTTGATCCTGAAGAAGACTTCATTCCCATCCTGGCCTTTGACTTTGAGATTGATGTG

AGCGGCGGATTGATCGCCTGGCTTCTTATCCTCTTCTTGCTGCCCTGACATGTTTCTTTTCTCCTCCGCG

ATTCCTTGCTGCTTTAGATGAGAATTCTGTAATTCGA

>JK743800.1 JutESTSSH3(N43) Jute Plasmid Suppression Subtractive cDNA library Corchorus capsularis cDNA, mRNA sequence

CCGGCCCCCCTTTTTTCCCTCTCTCCGGGGGGGGGGGGGGGGCCCCCCCCCCCCCCCCCTCTCCTTCTTT

CCAAACAGGTGGCTGGTGGGGGGGGGACAAAAAAATTACCCCCCAAATTGGTTTCCCCCTTGCGTTGTTT

AGGCGGGGTGGGTCTCCACCTTTTAGACGTAGGGTTCGCCTCCCATTCCGAGTTATTTCCCTGAAAAAAG

GGGGGGTTGTGTGGGGGGGGGTGACCGGGAAACTAGGGAACCTAACAAAAAGGCGGGCGACTGGGGACAG

CGGGGGTATCGGTCGTAAACGGGCCCCGTGGGTAAATGGCCGGCCCCCCCCCCCTTTTTTTCGGGGTTTT

GGGGGGGGGGGGGGCGCCGAGAAACCCCCCCCCTGGAAACCCCCCCCCCCAAATACATTTTTTTTGGGGG

GGAGGCGGGAGAACGGGGGCAATGTTAGGGGAAGCCACAGAGAGGGAAGAGCGCGCGTTGGGGGGGGGGG

GGTGAGGATAGCTTGGACATATTTGTCATAAAACAGCGACACTATCCCCCCGCCAACCGGCGCCGGGGTT

AAGGGGGGGGGAGTAGTGCGGCGGGTAACCGGCCGTATATTTCTCCCGCCACGCCCCCACGCTATTATTC

CTGGTCGTTAGATCGTTTTATATGTGGGCTCACTTGTCGGGGGAGAATGCAGGCACGAAAGAACACACGA

GATGATGATACTTTTCTTTTCTCTATAGATGACGAAAGGGAGGAGGGGTGCGGGACTGTCTAGTGAGTGT

GCCTCCCCCCCGCCGACCTGTTGTCCCTCCCTCGTTTATCTGGGCCGTAGATATGGCCTTCATGAAG

>JK743799.1 JutESTSSH3(N42) Jute Plasmid Suppression Subtractive cDNA library Corchorus capsularis cDNA similar to Chloroplast pigment-binding protein, mRNA sequence

TCAAAGTTCCAAGTGATTCATCCTACTTTCTCTTAAGAGCTGAAGGTGTCAATAATGGTTGTGTGAAGAG

GGTCACTCAAGTGGGTTGCCCAGTTGTTAAGTGGACCTTTGCCTGTTACGGCAGCTTGGACGGCAAAGCC

CAAGAAAGCAACCATGGCGAGGCAGGCGTGCTTGATCTCAGCCAATTGAAGGGTGGCCTTCTTTTCTGGG

TCTGCGGCCAAGCCCAGTGGGTCGAAGAATGATCCTCCTGGGTAAAGCCTCTTCTCTGGGTCAAGCTCTG

CGTTCCTTTGGAATTCAATGTATCCAATCACCAACACTTCAATCCAAATCAAAGTTGAAATGGAGAATGG

AAGAGGTTGGCCCAAGTATGACGATCCCTCAACTAGCTCCACCTTGCCAGCATCTTGCCAGGTAACACCA

GTGAGCCATTCAACAGTAAGAGCACCGAGAGTAGCCAACATGGCCCATCTGCCATGAATAAGCTCACATT

CACGGAACCTCTGAAGACCAAACACCTCGGCGTAAGGCTGGAACGGCGTTGCCTTAACGTCGGCAACCTC

AGTCCTTGTCCCAATGATGTCACCAGCCACATTCTTTGCCAAGTTCTGGTCCAAAGAATCCAGTTCATAT

TGCAA

>JK743798.1 JutESTSSH3(N41A) Jute Plasmid Suppression Subtractive cDNA library Corchorus capsularis cDNA similar to NADP+-isocitrate dehydrogenase, mRNA sequence

TATTAGAAGTCCCAATTGTGTGTTGAGCGTTGAATGGTTATGGATATCTCTGGTTGTGTGAGCTTGCTGC

TATGCAGTGGCTTTTAAATAATTTCTTTAAACTTATCTAGTATTATGCTATGGTTAATGTTGCCATGCTC

TGGCATTGAATAAATTTCAATTTCCAT

>JK743797.1 JutESTSSH3(N40) Jute Plasmid Suppression Subtractive cDNA library Corchorus capsularis cDNA similar to S-adenosylmethionine synthetase, mRNA sequence

TAGCAAATGGGCTTGCCCGCAGGTGCCTTGTTCAGGTTTCCTATGCTATTGGTGTGCCTGAGCCCTTGTC

AGTCTTTGTTGACAGCTTTGGAACTGGAAAGATCCCCGACAAGGAAATCCTCCAAATTGTGAAGGAGAAC

TTCGACTTCAGGCCTGGTATGATCACCATCAACTTGGACCTCAAGAGGGGTGGCAATGGCAGGTTCTTGA

AGACAGCCGCCTATGGACACTTTGGAAGGGACGACCCAGACTTCACCTGGGAAGTTGTCAAGCCCCTCAA

GTGGGAGAAGCCTCAATCTTAAGTTCAACCTTTCTCCATCCCTGCTGCTCAGTGCTTTGCATATCATAGT

GCTGAATAATCTGGATGTTTCCTTGCTGCCGCCTTTTATAACTTCTAAGCAATCCCCCTAATGTTCTAAT

CTCATACATATGATGCCATATTCATTTTCTTTTTTCTTTTTTTTTTCTTATTTTTTCTTTTATTCTGGTT

GTTTTTGAATTGTGTCCTTTTATTATCAGTGTCGGTGATCCAAAAATGTATGAGAATTTTGCTTTATAGA

TGGCACTGAAAAGTAACAAGCTGCCCTCTTTATGATTGCTTGTGAAATTAGCTTTGCAAAGGCAGCAGCA

AATACTTTAGTAGTAATAT

>JK743796.1 JutESTSSH3(N39) Jute Plasmid Suppression Subtractive cDNA library Corchorus capsularis cDNA similar to Calmodulin-7, mRNA sequence

TCTTGAAGAAAGCCGTGGAACGTGAATCCACCAAACTTGCTCTGGCTGCTTGATCTACCTAGTTCGGTTT

TGCTTTCATTTTTGGAACCTAATTTAATCATCCACCAAACTTACCTTATAATTCTTCTCGATTTCCACTT

TCATCTCTTATATATATAGTTTATATCATATGCTCCAAATGGATCATTTCTTCCCCAAGCTAAATGTAAT

TTCGTATGATGTATAATTTAAATC

>JK743795.1 JutESTSSH3(N38B) Jute Plasmid Suppression Subtractive cDNA library Corchorus capsularis cDNA similar to Elongation factor 1-alpha, mRNA sequence

TTTGGAGTTGTGGCATCCATCTTGTTGCAGCAGCAAATCATTTGCTTGACACCAAGGGTAAAAGCAAGCA

AAGCATGCTCACGGGTCTGACCATCCTTAGAGATACCGGCTTCAAAACCTAAGGATGATCCTGCCAAGGA

AGCTGCCAACTTCACCTCTCAGGTTATCATCATGAACCACCCTGGCCAGATCGGAAATGGATACGCCCCG

GTGCTCGATTGCCACACCTCCCACATTGCTGTCAAGTTTGCTGAACTCTTGACAAAGATTGACAGACGAT

CTGGAAAGGAGCTCGAGACGGAGCCTAAGTTCTTGAAGAACGGTGATGCCGGGATGGTTAAGATGATTCC

CACTAAGCCCATGGTTGTGGAGACTTTCTCTGA

>JK743794.1 JutESTSSH3(N38A) Jute Plasmid Suppression Subtractive cDNA library Corchorus capsularis cDNA similar to Elongation factor 1-alpha, mRNA sequence

CAGTCGAGGTTGGTTGACCTCTCAATCATGTTATCACCCTCAAAACCAGAGATTGGGACGAATGGGATTT

TGTCTGGGTTGTAACCAACCTTCTTCAGGTAAGAAGAGACTTCCTTAACAATTTCATCATACCTTGCCTT

GGA

>JK743793.1 JutESTSSH3(N37) Jute Plasmid Suppression Subtractive cDNA library Corchorus capsularis cDNA similar to beta-galactosidase, mRNA sequence

TCAGAGATCTCAGCACAAACACTGGACACTGATCTTTTTACAAGTGAGATTCTCGTGGGATCACCTCCGA

GCTCCTCAAAAATAACCAACAGATTTTGTGTAGGTTTCAACCAAGACCGAGGAACATGATACCATCTTTG

GGTTGGTTGGCCGCATCCAAGCTGACACTTTGGTGGTCGGAATGTCCCAGCATAGCTGCACCCATTGCAG

TCACCATGAGCATATGCAGTCCAATATCTACCAATGCTCTGCCCATTAATCCATATTTGACCTTTTCCCA

TACTCTCCATGTCCAAAGCCAATGGCTCGTCACCCTCAGGTGCATTGAAATAAGCCTTATGCCATCTTAA

TGGCTGCTGTTTCTGTGCAGCCAATGATCCCTCTATCCACTCAACAGAGTAAATATTATTTGGTGAGACA

AGATTCATGGACTCTCCTTTAAGTCCAACCTGGTAGGTCCATTTTTGCCAGGACAGGTCCCATTTTCCTT

GGTTAAGTCCATGAAGTGCAACTGGAC

>JK743792.1 JutESTSSH3(N36) Jute Plasmid Suppression Subtractive cDNA library Corchorus capsularis cDNA, mRNA sequence

TCTAGCTGCTGGAATTCTCCTTCAAGTTCCTCCCACTCTTGATGTGACAATGATACAACATTCCACCCAG

AAGCAGCAATATGACGACGTTTCAAGATGGTATGCCCCAAAGGTGTTCCACTATTCCTTGAGAAATGTGT

TGGTCCATCTATCTCCAAAACAACCTTTTTATCAATCACAGCAGCATCCACGGTATATACATCCACAGAG

TATTCTCTTACCCAATCAAGTCCA

>JK743791.1 JutESTSSH3(N35) Jute Plasmid Suppression Subtractive cDNA library Corchorus capsularis cDNA similar to putative senescence-associated protein, mRNA sequence

TCTGGCTTGTGGCAGCCAAGCGTTCATAGCGACGTTGCTTTTTGATCCTTCGATGTCGGCTCTTCCTATC

ATTGTGAAGCAGAATTCACCAAGTGTTGGATTGTTCACCCACCAATAGGGAACGTGAGCTGGGTTTAGAC

CGTCGTGAGACAGGTTAGTTTTACCCTACTGATGGCCGCGTCGCAATAGTAATTCAACCTA

>JK743790.1 JutESTSSH3(N34) Jute Plasmid Suppression Subtractive cDNA library Corchorus capsularis cDNA similar to Superoxide dismutase, mRNA sequence

AAATGTATTCGTTCTACATTTTATTCACAAAGTCTCTCTGGTTCACATAAAAAGAGTTATTCTCAAGATC

ATTGCATCTCCGACTGCACATATTATTCTGATCACTGGGATAAGGACTTCAGCCTTGCTGACCAATGATA

CCGCAAGCTACTCTGCCACCGGCATTTCCTGTGCTTTTGCTAAGTTCATGGCCACCCTTGCCAAGGTCAT

CAGGATCTCCATGGACAACAACAGCTCTTCCAATAATAGAGTTTGGCCCAGTGAGAGGAATCTGACTGTC

AGTTTTACAGAAACTTGCAGTGCCATCATCACCAACAGTGATATTTCCTAGATCACCAGCATGGCGATTC

TCATCTTCTGGGGCACCATGCTCTTTCCCAGCAGGATTAAAGTGGGGTCCTGTTGACATGCAACCATTTG

TAGTGTCCCCAAGGGCATGAACATGGAAGCCATGGAGTCCAGGCTTAAGACCAGAAACATTCCCGGTCAC

AGTTG

>JK743789.1 JutESTSSH3(N32) Jute Plasmid Suppression Subtractive cDNA library Corchorus capsularis cDNA similar to Major latex-like protein, mRNA sequence

TCGTTCATCACATCCCCTTCGATCACACTAAAAGTGATTGAGTTCTTCGCCTCGTCTGCTGCTTCAATTA

CCTCCTTTGCCACCTTCCTCACTCCATCATGCGTGTAATGCCAGACGACAATGGAACCGACTTTGCCAAA

TTCACCCTCATGCAATTCACATCCGTGGATCTTGTCACCGCAGGTTTTGGAGACGTGGTGGGGTTTTTTA

AAGAACATATCATGGAAATCCTTGGCAGAAGCCTTGATCTCTACATCGGTCTCAAGCTTACCAGTTAGAC

CCGATGACGCCTTTGCAGCAGAGGTGAGGTGAGCATCGATGTCTCTGGAGACATCGGCGACAAACTGCAG

CAGTGTCTCAGGATGATCAACTCCCTTGTGCAGCTTTTCGTATTCCAGAATCCAGTGAACTATGCAGCCG

TTACCATCGCTCTTGGGTGTAGCTTGAATATGGATTACAAAGTTCTT

>JK743788.1 JutESTSSH3(N31) Jute Plasmid Suppression Subtractive cDNA library Corchorus capsularis cDNA similar to Al-induced protein, mRNA sequence

AGTGGAACCTTACCAAATTGATCAGAAGCTACAAACAAGGTGGAGGTTGATTTGTCATAGACAATGAAAG

CAAAGCTCCCACTTAAATGCCCAACCACATGGTTTGGAGGATAAGGTGCTCTGTCTCTAAGAGCCTTGTA

AGCTTCAATAACCAAAATCACTTCATTTGCAGACTTTGCAAGCCCATATTGCTGCTTTAAACTCCCCAAG

TTGTCAAGTGCACCCTCAAACAAGCAGAAGATTTCATCTTTCACAGCAAATGATCTGGGCTGTAGGGCGG

ACTCATTCTGGTGAGAATAAGCGAATTGGACATGATCTCCGATCTGCATGGACACGGCGGAAGAGTTGTT

GTCAAGGAACCTTTTCACCAGAGCATCCGACGTTATCTTAGGCGACGGAGTCCGGCAACCGGCAGCCACC

AG

>JK743787.1 JutESTSSH3(N29) Jute Plasmid Suppression Subtractive cDNA library Corchorus capsularis cDNA similar to Ubiquitin fusion protein, mRNA sequence

AAAGAGAACCAATCCATCATCATATCAAAAATGAAAAACTCACTGTCAAGAACAAACACGATGACATACA

CCAAGAGGTTTAGGCTTAATGAAATTAAATCGAACTCACGATAGTAAAATGTCTGTTATTTAAAACTATA

GTTGTCCAACAAAACCTCAAGAAAACTAATACTAAATGTAGAATTCCTCCCTCAAGATTCACAATCTACT

TGATCTTCTTCTTTGGCCTCAACTGGTTGCTGTGCCCACACTTCTTTTTCCTGCAGTTCACAGCACGGGG

GTGAAGACGAGCA

>JK743786.1 JutESTSSH3(N27) Jute Plasmid Suppression Subtractive cDNA library Corchorus capsularis cDNA similar to SKP1 component-like 1, mRNA sequence

AAAGGCAATTAAAAAGAAAAAGGAACATAGAAAAAAGATATGTTCAAATTCAATTCGATATCCTTCACTC

AAAGGCCCACTGATTCTCCCGACGAACCTCCTCCTCCTCTTCCGGAGTAAAATCATTCTTAATATTGAAA

GTCTTACGGATCTCCTCAGGCGTTTTCCCCTTTATCATATCCGCCACCGTTTGACACGTCAGGTCCAGCA

GGCTCTTGATGTTCAAATAGTTAGCAGCCAGGATGAGATCGAAGAGGGTAGCTTGATCGACCCTGACGAA

ATCAGCATCCCAAAGCTTAAGCTCATCTTCGACGGAGGCAGAACGATCGTCGGATTTAGGTGCTTCCACA

TGCTTCTTGCA

>JK743785.1 JutESTSSH3(N26) Jute Plasmid Suppression Subtractive cDNA library Corchorus capsularis cDNA similar to Vicianin hydrolase, mRNA sequence

TCTCAGCACACAAGTTCGCACACTTGAAACAAGAAATTAAAATTAAAGAAAGCCAACAACAACAACAACA

AACGACGACGACAAACTGTTAATTACGTCCATCTCTTGGTTTGGTTCGGCAGCGGCAGATCAATTAATGG

CTGAGAAACTTCTTGAACCAGTAAGCTGAATATTTCAGGTGTCGTTGCAATTTATGCTTGAAATCAATAT

AGACCAACCCAAATCTAACAGTATATCCAGAATTCCATTCGTAATTATCCAAAAATGACCACATGTAATA

ACCCTTGACATTAGCACCCGCCTTGATCATTCCTTCTAAAAGCGCTTTCAGGTGGCTGTCAAGGGCTGTA

ATCCTAAGGCTATCCTTGATTGCTTCAGTTACCGACAACGAAGCATTGTTTGCTTCAGCTACCCCATTCT

CATTAATGTAAACAACAGGATTGTTGTATTTGTCCTTTATGTGTATCACAAGTTCTTCCAAACCCTTTGG

AACAAGGTAAAGCCAACTCACAACAGTAGGTGTGCCAATTGGAACTCCATTATTTTGAGATGATGTGGAG

ACATTAGCATAATTT

>JK743784.1 JutESTSSH3(N25) Jute Plasmid Suppression Subtractive cDNA library Corchorus capsularis cDNA similar to GDP dissociation inhibitor 1, mRNA sequence

ATCCGTGACGGTCGATTCAAAGTGTGTTGTAGCATCATAACTCGTTGATATAAAACAATTGTCCAAAGAA

GGCTCATTGACTGGTTCATATCCGTCATAGATGTCAAAGAATATCTCATCAACAGGACCAAGAAGGTCAA

TTCCAGGCTTTAGCTCAGTTTCAGGGTGATCCGTCTCGGCCTCCGTAGAAACAAATGCAATAAATTTGCC

CTTAGGAGCAACATTGTGAGAATATGAGCAACAGAAGACATACATGTCCGATCTGCGACCCAACTGCTTT

TGTGGCAGGATAATCTGCACTGAATGAGAGTCATTGGTGTTTGGGATTGGGTGACTCATGATAGCAATTG

CCCTGGCAACCCTGCCAATCTTTCTAACCTTGTTCGTCAAGTAGGAAGGATCACAAACAACTTTCTTACA

CCTGGCAGTTTCACCTTCAGATGTGACGCCGCAAGCTTTACCTTCATCATCAAACTCCACCTTGCACTCA

GGCTTATTCAACATGTATGTTCCACCATATACAGCACTAAGCCGTGCAAAAGCCTGGGGCAGCTCGCCCA

ATCCATACAGTGGATAAATGTAAAGGGATCCACCTTGAAAACGTGCAAGAGACTCTGCATACAGCTTCAT

CCTCTTTACAGTATCCAATGCAGGCTCATCCAGGTAACGATCATCTCTATGAAGGGCCAATGCATGACCA

ATGAAGTCCACGGTGTTATCATCAAGACCATATTTCGCATTTAGCTCCCTAGTGGTCACCGTAGTTCAGG

TCCATTCCCTCATGACTTTAGGATCCCTTTTCGTCATATTCTTGAACATAATATGAGAACTTCCGTGCAC

CGACCGCTTTTCAAATATTCCCATGAGTGGAGATTGGAGTGCCTCCATTGTCCAGTTGGCTTGCACGTAT

GACGTTGGTTTAATGAAACATAGCCTGCCATTCACAGCCTTGGAAGTTACCTCAGGCGGAACCACGCCTA

TCCTGATGCATCGCGAGTTACCGAGTCGATTCCATGTCCGTCGTTGAACACGTCCGTTGACTA

>JK743783.1 JutESTSSH3(N24) Jute Plasmid Suppression Subtractive cDNA library Corchorus capsularis cDNA similar to Ribosomal protein L5, mRNA sequence

CGTTTTGTCGTGCGATTTACCAACAAGGATATCACTGCACAAATAATTTCTGCTAGTATTGCTGGTGATA

TGGTTCTTGCTGCTGCTTACTCACATGAACTTCCTCGTTATGGACTTGAAGTGGGTCTTTCTAACTATGC

TGCTGCTTATTGCACTGGATTACTTTTGGCACGTCGTGTCCTTAAGATGTTTGAAATGGATGATGAATAT

CAGGGCAATGTGGAGGCTACTGGAGAGGATTTCTCAGTTGAGCCCACTGACACAAGGAGGCCATTCCGTG

CTCTCCTTGATGTAGGGCTGATCAGGACAACCACGGGCAATCGTGTGTTTGGTGCTCTTAAGGGAGCATT

GGATGGTGGTTTGGATATTCCTCATAGCGACAAGATGTTACTGCTGCAAAGGACCAACCTGCTGCAGAAC

CAGAGGCGACGGCAACAGCATAATTTCTTACAATACGTTAACCATGGTTTTTTCTTGCCCCATTCTTTTT

TATTTTCCTCAAGATTTTGATTTTTTATTAAATTCCATATTACGATGGTATTTCCTTCACAAGGATTTA

>JK743782.1 JutESTSSH3(N23) Jute Plasmid Suppression Subtractive cDNA library Corchorus capsularis cDNA similar to transcriptional regulator, mRNA sequence

TAAACAATGCTATTAGAAACTTTTATTTTTGACCAATCTCTCCTTGAGCATTGATTATTTGTTATTTTTT

TGCCCCATGCTTTCACCATTTTCCCGGATTCTGATCATGGCCTCTCTCTCTCTTGTATCTCCAAAGCATT

GAAGATCATCATCCAAAAAACAGAGCCCATTGTTAACCCAAAAAAATCTCTTTCATCAAGCAAAAAAGGA

CATTTCAAGCATAAAGAGTCGTCCCATCCTTGCAAAGCAAAACCCAGAAAGCTGTAGACAAAGCGATGGG

AAGAACAAGGAAGACTTTCTTTGAGAGAGATGAATTATATTCAACTCCACCCTCTTCAAGAATGGTTTCT

AGCCTTGGAAAAGATGAAGAGGATGTTCTTCGAGAAACGTGACGGGAAGGCGCCACTGCACCCCAGGTTT

GCTTCTTTGCCACCTCTCTCTTCATCATCACCATCTTTAGAAATATAAGAAATTGATTGAAAAAGCTTGA

TCGAAGTTTGGCTGTTTGGGTTTTTAAGATTGCTTC

>JK743781.1 JutESTSSH3(N21) Jute Plasmid Suppression Subtractive cDNA library Corchorus capsularis cDNA similar to 40S ribosomal protein S24, mRNA sequence

TGTTATACTAAATATTTCAAAATTCTCATCTGGTAGAATTTACTCAAAACAAAGACAATCAGAGAATATA

ACTAACCATCGAATAACACCAAAATCCAGTTTCCACAAAAACAGGTTCTAAGAACACCTAAACATTTCAG

ACTAGTAATAATAGTAAAGAAATGCAGTTGAAAACCAGTGGCTCTAAGATAAGCACGAACTTCAATTCTT

TCCCCTCTTTTAGCTTACTTCTTCTTTGCATCACCGGCCTTGGTCTTCTTAACTCCGCGGACCTTCTTAG

ATCTGTTCTTCCTTTCCTTCATTTGTTTCCTTGACTTCTCTACCTTGGTGTCAAGTCCATTCCTGATAAG

CCT

>JK743780.1 JutESTSSH3(N20) Jute Plasmid Suppression Subtractive cDNA library Corchorus capsularis cDNA similar to RNA-binding protein, mRNA sequence

TCATAGTTACGAACCCAAAGCCACGACTTTGTTCACTCTCTCTATTGTAAATAACCTCAGCAATTTCCAC

AGTTCCAGCTTTGTCAAAGAGCATAGCCAAACTTTGACTGTCAACATCGTAAGGCAAATTGCCAACAAAC

AATTTAGCTTCCTCAGGAGGCTCGGGAATAGACTCCTCTTCTTCCGAATCCCCACCCTGTCCTAAAAACG

CTGCATCCTCACCTTCCTCAGCTTCCCAATCAGACAACTTCGCTTCAGCCCCATCGCTTTCTTCATTTTC

CCACGTTAATTCTCCTTCACTTTCACTTTCAGTTTCTTCCTGAACTTGTTGCTCTTCTCCATCAATGGTT

ATGGTGGTGTCATTTTCTCCCTCTTGCTGGGCCCAATCTGAAGTCTGGGCAACAAAGGCAACCAGAGAAG

GGAAATGGGTTTTTGTTCTGAGTGCCGAAAGGGTTGAAGAATGGGAAGAGGAAAGGAAAAGCTTGATAGG

TTTTGGAGGGAGTGAGAAGAGCATAAGGGAGCCCATGAGCCACCCTGTTTTCTTGAAGGT

>JK743779.1 JutESTSSH3(N18B) Jute Plasmid Suppression Subtractive cDNA library Corchorus capsularis cDNA similar to Cell wall-associated hydrolase, mRNA sequence

ACCTTTGATCCGTTGAGCGAGAGCCCTTCCACACGGGACTCCCGGATCACTATGGCCGACTTTCGTCTCT

GTTCGACCAGTCGGTCTCACAGTCAGGCAGGCTTATACCATTACGCTCACGAGCAGAATCTTAGCTTGAG

CCTACCTTCGCACACCTCCGTTACTCTTTAGGAGGCATCCGCCCCAGATAAACTACCCACCTCGCAGTGT

CCCGCCACCCCCCGAATTCTCGGTTTGGCGGTTAGGCATCCTTAGACGAAAGAGTGGCCTTTCAGGATTG

GTCGTTGTGTGTCACCACCTCCCACCTATCCTACACATTCGATCAGGGTTGTCACTGCGAAGCTATAGTT

AATGTGCACGGGGTC

>JK743778.1 JutESTSSH3(N18A) Jute Plasmid Suppression Subtractive cDNA library Corchorus capsularis cDNA similar to Ultraviolet-B-repressible protein, mRNA sequence

TCAAGCCGCCGATGGAGTTACCCCATCTCTCAAGAACTTCTTGCTTAGCATTGTGGCTGGTGGTGTCGTT

CTTGTTGCCATTGTTGGAGCCGTGATCGGCGTCGCCAACTTCGACCCTGTCAAGCGGAGCTGAGCTAATA

CATATTCTGAGATGCTGTGAATTTTTTTTCTATTAAACCTCTTCATGTATCTGTTTAATGTTTCCAGTAA

TGAACTCTGTCTTTTGCGTGCG

>JK743777.1 JutESTSSH3(N17) Jute Plasmid Suppression Subtractive cDNA library Corchorus capsularis cDNA similar to Peroxidase precursor, mRNA sequence

TAAATTAAATCATTGAAATCCATGCATCATCAAACAACATCAACACCATCCATCCATCCATCCATACATA

CATACATACATACATCCTAACATCCAACCACCAGCCAGGACCATCATCATTCATCAACCTGGGCAAACAA

GAGAAGAGAAACCTAGCTATATGCTAAAGAAAACAGGGGAATTTGATGAGCAAAGTCATCATATCACCAG

ATATAACTGGTCACTACTGCATATTAACAAAGTTGTCTCAAGACTAGATCCAAACAACCATTTTTATTCT

GCCACTCACATTATAGCATTCTAGTGAAGCTTGTTGGCAACATTGCATCGTCTCCTGATTTCACCCTTGT

TACCAGTAAGAGGATTGTTCTCAGAAAGTTTAGTGATGGCCCTGGAAAACTCCTTGAAGAAATAGTCTTG

GCTCTTGGCCATTTTCTTCACATAAGGCCTAGTCCTCTTGTCATAAGCAAGCTGGTGATCCACAATCAAC

AAACCCTTGTTGTCCAATATGTTTCTGTAGTAATTGTTATCAAGAATCATGGGCGTGCCACGGTCGTTTC

TCACATATTGGGACCGCCTTGGGGTCTGGAATTTGGTCAGGGCATTTGTGGAGCATGTGTGGAACATGGT

CAGGGTTTGAGTGCAGGGTCGATCTCTGG

>JK743776.1 JutESTSSH3(N16) Jute Plasmid Suppression Subtractive cDNA library Corchorus capsularis cDNA similar to Cytochrome P450 like, mRNA sequence

TAGAAATCAGAATCAAACGAGCTTTTACCCTTTTGTTCCACACGAGATTTCTGTTCTCGTTGAGCTCATC

TTAGGACACCTGCGTTATCTTTTAACAGATGTGCCGCCCCAGCCAAACTCCCCACCTGACAATGTCTTCC

GCCCGGATCGACCGACAGATGCCGGTCTTGGGTCTAAAAAGAGGGGCGTAGCCCCGCTTCT

>JK743775.1 JutESTSSH3(N15) Jute Plasmid Suppression Subtractive cDNA library Corchorus capsularis cDNA similar to 40S ribosomal protein, mRNA sequence

ATAATTCGTGAGATTTTTATTTATAATATAAAATGGTAAGTTCTACAAAACCTTTTTGTTTATTAATAGT

AATAATTTCTTAATTTGGACAAGAAAGAAATATTACATCATCAAATCTACTACTTCGTATACAAAGATGA

AAGAGGAGCAGAGCAACAAGAAGTTAAGCAGACAAGAAATTAAAAAATCTTCCCTTTTAGAACACAATCG

ATAGCCCCCTTCGATAGAAAACATGTTTGAAACCAACCTCGTCGTCAATCCCATTGAATACTTAGCGGGT

TAGCTTTTGTAATAGTGAA

>JK743774.1 JutESTSSH3(N14) Jute Plasmid Suppression Subtractive cDNA library Corchorus capsularis cDNA similar to Leucine-rich repeat receptor, mRNA sequence

AAAGGCTTTAACTTTCGGAAGCCAGGAATCGGCATCATATCCATGCATGTAAGGTCAAATGTTGCGTGGG

AAAACCTTACTGTGGCATGCATTAAATCATCTTTAGGTGTAGCCTAGGTGATTCCTAAAAACATAATGCA

GCGGTGTTTTGCAACAATCATGGGGTCTGCACATTTGAATCTTTTGACTTGTTCTCTTCAGAAGATGGCC

GGTTCTCTGAGTCTGATTGCCGGACCTCTTCAATCATTCTAACAACTTCATCCATGTTAGGTCGCATGTC

CGGCACCTTTGCCACACATGCCATTGCAATTTGTAACATCTGCACCATTTCTTCTTCAATGTTTTGGAAC

CTCATCAACTCAACATCAAAGACCTCGGCTGTCCATTCCTCCCTCACAACAGATTGAACCCACCTAGGAA

GATCAACCATGTCATCCCGCCCAGGTGATTGGAGTGGTGCTTTCCCAGTCAGCATCTCAAGGAGCAGGAC

ACCAAAGCT

>JK743773.1 JutESTSSH3(N10) Jute Plasmid Suppression Subtractive cDNA library Corchorus capsularis cDNA similar to cytochrome P450-like TBP protein, mRNA sequence

TGGAAATCAGAATCAAACGAGCTTTTACCCTTTTGTTCCACACGAGATTTCTGTTCTCGTTGAGCTCATC

TTAGGACACCTGCGTTATCTTTTAACAGATGTGCCGCCCCAGCCAAACTCCCCACCTGACAATGTCTTCC

ACCCGGATCGACCGACAGATGCCGGTCTTGGGTCTAAAAAGAGGGGCGTAGCCCCGCTTCCGATTCACGG

AATAAGTAAAATAACGTTAAAAGTAGTGGTATTTCACCGTTGCCGAAGCTCCCACTTATACTACACCTCT

CAAGTCATTTCACAAAGTCGGACTAGAGTCAAGCTCAACAGGGTCTTCTTTCCCCGCTGATTCTGCCAAG

CCCGTTCCCTTGGCTGTGGTTTCGCTGGATAGTAGACAGGGACAGTGGGAATCTCGTTAATCCATTCATG

CGCGTCACTAATTAGATGACGAGGCATTTGGCTACCTTAAGAGAGGGAGGATTTCCGATTGAGGCTAGGA

GTCACCAAAAAGATTTGAATGAA

>JK743772.1 JutESTSSH3(N9) Jute Plasmid Suppression Subtractive cDNA library Corchorus capsularis cDNA similar to Cytochrome P450-like protein, mRNA sequence

TGGAAATCAGAATCAAACGAGCTTTTACCCTTTTGTTCCACACGAGATTTCTGTTCTCGTTGAGCTCATC

TTAGGACACCTGCGTTATCTTTTAACAGATGTGCCGCCCCAGCCAAACTCCCCACCTGACAATGTCTTCC

GCCCGGATCGACCGACAGATGCCGGTCTTGGGTCTAAAAAGAGGGGCGTAGCCCCGCTTCCGATTCACGG

AATAAGTAAAATAACGTTAAAAGTAGTGGTATTTCACCGTTGCCGAAGCTCCCACTTATACTACACCTCT

CAAGTCATTTCACAAAGTCGGACTAGAGTCAAGCTCAACAGGGTCTTCTTTCCCCGCTGATTCTGCCGAG

CCCGTCGGCCTGGATGAGGCTAATCTACATGCATTCGCTAGGCACCGAGCTCTCATTCGGTGGCCAGCTC

TATATCACGTGCCCGACTGGGAACACCGTGGCGACTGCCAACTTACTGGCCTTGCAGCACATCCTCGCCT

CGCAGCACATCGTAATTTCGAAGACTGGCGCACCGATCAACATTCCCACACTTATCGCAGCTTGCATGGG

GAAGCGCAGCCTGTAATGCGTTAATGTATTGTGTAAAACTCGAGTATTAATGTTTAGATAAGCGTCTCAT

TTTTTGTAACCATCAGCTCAGTATTCGGCCAAATACCCCTGATAATCGGCAAAGAACCACTTCCGAGATC

AGGATAAAGTGCTCGAGCTAGGTTTGGAGCTGTAGTCCACGTATTGAAAGCACGATGTAACTCCATTCGA

CGAACGGTGGGACTACCCACCGTCTATACAGGCGCGAATGCCGCTACTATCGATGACCATGGACCCCATA

TATCGAGGATTCTTTCGAGCCGCCTCGATCAGCTGTATAGGCAGCTCAAAGTCTGCGCTCATAGACGTAG

TTCCGCGACCCTAAGCTGACCCGCGGATTCCAGCTTAGCCTGCAGACGGCAGACATGACGCGTAGGATCG

CAGCATCGACCTGACAGGTCCTCAC

>JK743771.1 JutESTSSH3(N7) Jute Plasmid Suppression Subtractive cDNA library Corchorus capsularis cDNA similar to putative glycine-rich protein, mRNA sequence

AAGTGCATAAGATGCTTGCGGCAGACTTCTGCTTTTGCTTCACCTGGTTCAACCAATCAACCTAATCAAA

ATGAACGGGAAACTTGGTGGATCAGCCGTTGGGCTTATGGTTGTCCTATGTGTGGCGGAACTTGGGTTCG

AGTTGTATAGATGATGCTTTTGACAGGATATATTTATTAGCTGGGCCTATTTTTATTCTCCTTTTATCCG

TTTATGTGACAAAATATACATCGTAGCTGGTTGCTCTCCGTTGCAGAATCAGTATATATTCGAAGAGTGA

CATACTGTTTTCATGTAGTGGCTGCTGCTCACAATTCCGGGCTACTTGAAGATTGAACCAGTAGAATTCT

ATTTTTCAGAGTGAGTTATCATCATTAATTGTAGAACCCATATCCTTATTATTTCATCTTGTAAATTTTG

TTGATAAATGACAGA

>JK743770.1 JutESTSSH3(N6) Jute Plasmid Suppression Subtractive cDNA library Corchorus capsularis cDNA similar to high mobility group protein, mRNA sequence

ATTTTCCCTTTAGATGACATGAGCATCTAAAAGAATAAACATCGAAACATCAAAAGCATTAACATAAGAT

CGATCAAACACTACCACAGCCTACATTGATTATGCTATCCTTCCTGATGGATCTTCCTACTCATCATCAT

CCTCCTCTCCGCTGCCATCTTCGTCATCCTCATCATTCACCTCAGAAACAGACTTCTCAGACTCCACTTC

ATCATCAGCTTGGGGACCCTCAGCCTGTCTCTTATTGTAAGCTTTCATGTTCTTCTCATACTCAACCTTT

CGCTTCTCTGCCTTAGCAACATAAGGTGCCTTTTCAGCCTCAGACATGCTCTTCCATTTATCTCCTCCAG

CTTTGCCGACAGCAGCAACAGACTTGTTCTTGGGATGATCCTTCTTATATGTCTCGCGGAACTCCTCCAT

GAAAACGAAGAAGGCACTGGCAGGCCTCTTTGGCTTGTTTGGATCCTTGGCTGCCTTCCCCGATTTCCTT

CCTCCAGCTTTGGTGGTTGGTTTCTTATTCACGGAGAGCTTAGCGCTCTTGGTATCTGACTTGGATTTAC

CTCCCTTCATGACGAATCCTGGTTGGTGAACTAG

>JK743769.1 JutESTSSH3(N5) Jute Plasmid Suppression Subtractive cDNA library Corchorus capsularis cDNA similar to Aspartic proteinase nepenthesin-2 precursor, mRNA sequence

AGACAATAGCTTGTTTTTGTTTATAATTCTATCTTTTGCCCTGAATTTTGGCTACAAGCAGCGGAGAATA

TAAAAACAACTATATATATTTGAACAAATCTTCTTCCCTCCACTTATGGAGATGATGATCAAGAGCCTTC

ACTATTATATATATAATCAATATTTATTTTGTAATTAACTCAAACACCTTAGCTACATCCCCCTGATGCA

AACCCAAGTCTCCCTTTAGCACCATCGTAGACCACTTGCAATGTCTTCTGCTGTGTATTCCCAAAAATCC

CCACATCTGTATCATCGCCGTTTGGAGCAAACGCCAGACACACTTGCGAGATACTATTTACGTAAAATAT

TCCCTTTGCGTCGATCGGAACCACGACTGAGCCGCTGAGGAAGAAGCTTATCTTAGGTATGGTTACCGTG

CTGTATCTGCTAAAATCGTAGCAGGTGTCCAGGATTGAGAGTGGTTGGGCCATTGGGTATTGGCTCATCT

GTTTTCTGAATGCAGTTTTCAGCGCATCATAAGCAGTCGGCGGCAATCGCGTAATGACGGTTCCAGAGTC

GATGATCGCGCCGCCGGCGGAGAAAACTGATGCCGAGATTGGTAGTTTTTTGGCCTTCCGACGGTTACTC

CAGTATGTCATGCCGTAAAAGATGTGCCTTTGGGAGATTGTAGACAATGGGGATGAATTTTACGGTTTTC

GAGACCCCGCCGCCGTAGCCAATAGAGAGGAAACCGGTTGAGCTGGCGGATGACGGGCAGCCAATAGGAG

AAGTTATTTTGTA

>JK743768.1 JutESTSSH3(N4B) Jute Plasmid Suppression Subtractive cDNA library Corchorus capsularis cDNA, mRNA sequence

TCCTCTTTCAGCCTCTGAGTCCATGAAGCTCCGTCTCTGGGGCCCGCCTTCGTCGTCAGAAGCGGGATCT

GAGTCAGCGTCGACTTTGTATTCGGGTCCCAGCCTTCCATTTCG

>JK743767.1 JutESTSSH3(N4A) Jute Plasmid Suppression Subtractive cDNA library Corchorus capsularis cDNA, mRNA sequence

GATACCAGCACTTTCCGGTCCAACGTGTCCCTTCCGGATTGGCCGCAGAGATCCGGAACCAATCATTGTC

TTTGGACTTATTCATCTGCGTGTAGGCGATCAAATCCCT

>JK743766.1 JutESTSSH3(N3) Jute Plasmid Suppression Subtractive cDNA library Corchorus capsularis cDNA similar to mipB gene, mRNA sequence

ATCCTCAGAGCTGCAGCCATCAAGGCTTTGGGATCTTTCCGCAGCAACCCCACCAACTAAATGATGAAAA

ACAAAGAAAAGACTCATCGAAAACCCAAAAATAGCTTTGTCTTCTTCACTTCTCAAACCCTCATCTTATT

TTCTCCATTTGTTCGTTTGTGTGTATGAGAGAATGATGATGATTATGATGATATGATGACCCTTTTTCTA

TTTCATTTATCTTATTTATTATAATTTAAATCTCTCTTTTGTTGTTAGCTCTTGCTCCCTTGTGTATTAT

TGTTCATGAGCAGTAATAGTAGTT

>JK743765.1 JutESTSSH3(N2) Jute Plasmid Suppression Subtractive cDNA library Corchorus capsularis cDNA similar to Proline-rich protein, mRNA sequence

GAATGAGAATGATAGACAAAATCAACTTAACATCGACAAACATTAATTAAACATTACATTAGGCCGAGGT

ATAAAATTAACCCACAAGCCTAGTTATTTAAAACATGAACAAATATCACACAATGTCGGAAGTCTTTGTA

ATAAAGACTCCCTAAAAATATGTAAAAATGGAAACCCTATCCCAACTTCAAATCGATCCAAACCACTAAA

CAAAAATACATTGAATATTTGAAACTACAACACCAATTGAAGAAATTATACTAATGATAAGCAAATAAAC

TTAGCCGGAGAAATTGTTAACGTCGGCGGCCGGCCGCTTCCTTATTCACACTTGAAGCCTGGAGGAATTT

CCTTTTGGCATGCACTAAGGAGTAAGCTAAGAGTAACAGGAATGTTGAGGTTGATTCCAA

>JK714342.1 JutESTSSH3(31) Jute Plasmid Suppression Subtractive cDNA library Corchorus capsularis cDNA, mRNA sequence

CAAGCAATAGCAAACTGTTTTTAACTTATTGGATGTAAGCATGGAATGGTATGGTTATCTATATTTGGCA

CAGGGAATTGCAACCCATTAACAGCAAAAGCAACAACAACAAATTGTCATAACATAAACAAAATTACATA

TGAAACAACAACACTGGTTAAAATCTTTATTGTAGCACCAAAAACCCCGACACCAAACTTGACTTACCTA

TTCGTTTCCATGCTTTTCCAAATTTTCTTTAAAAATCCCTTCTTCTAACGAATCTGTCCAAGTCACTTCC

CACTCGAGTTGGATTCATTCAAACAAGTCAAAAAAGGTGAGTGGATTCCTTCAAACAAGTCAGTCAATCA

CCTTCTCGTCTTTGGCCATAGTTACTGCAACCTTCTTGTCTCGTGGACCTTTGGAAACCTTAAGC

>JK714341.1 JutESTSSH3(27) Jute Plasmid Suppression Subtractive cDNA library Corchorus capsularis cDNA similar to Ribosomal protein L32, mRNA sequence

AAGCTTATGCTCATTCATCTTCCTGGCTGCGCAACCTGGCAAGCCTGTTAGTGACCACAACATCAAGCTG

AGCTGCTCGTTCCACAATCTCTTTTCTCTTCTTAGTGGAAACATCATGGGCAATTTCAGCACAGTAAGTC

CTGTTGTGCATCATCGACACTTCAAGCTCTTGGACATTGTGAACAACAAACTTCTTAAACCCATTGGGAA

GATAGTGGCGAGTCTTCTTGTCTGAGCCATAACCAATGTTGGGCATCAAAGTGCACCCCTTGAACTTTCT

TCTCACCCGGGAATCAATACCCTTTGGTCTACGCCAGCTTTCCTTGACGGAGATTTTGCGGTCACTCTGG

GGTCTCTTGAACTTCTTGACCCTCTTCTTCACGATCTTCTTACTTAG

>JK714340.1 JutESTSSH3(4B) Jute Plasmid Suppression Subtractive cDNA library Corchorus capsularis cDNA similar to 60S ribosomal protein L17, mRNA sequence

GATAAGAAAATTGCAGCAAGAATTAACCATAGGAAAAACACCCCATAAACTACATTTAATATTCAAAATC

AAGATCAAATCAAAACACTCCAACTGATATTCGGAAGAAACCATTAACTAAATATAACAAAATCTTTTAT

CAAGAGAACTATGGTTGATCCAACATAGAGAAACAAAATCACTGATATTTGAACTCCTAGACAATAGCAT

TAGCAGCGCTTGCAATCCTAGGCCACAGATCAGCACACTCCTTCCCGATTGGACCGGTAATAGCAGAACC

TTTCATTTTCTCCCCTTTGGATTGACAATCACACCAGCATTATCTTTCAAA

>JK714339.1 JutESTSSH3(51) Jute Plasmid Suppression Subtractive cDNA library Corchorus capsularis cDNA similar to Hypothetical Cys-3-His zinc finger protein, mRNA sequence

ATAGAACAACAGGAGCAGATGGTGGCATAAGGAAAAAGATGATCCGTTTGCTCATAGCTATATTCTAACA

TATTTCATATTCTTGGATTCTTAATATATATATTTGATAGCTTGTAGTCGGCGGTAGAGGAGGTAATATG

TCTTCCCGACCAGTCAAGAAAGAAGCATAAACATACTTGAGCAGGAAGGTTTAAGTCACTGCTCATGTTA

GAATTTTAGGAGCAAAAAGGAGAACCAAGCAACAAAGAAGATTTATTGATTATTTCATATTCTTGATTCT

TTTTCTTCCCCCTTTGGGGATCTTTCTCTTAATGTTTTTAGCAAGAAGTTGAGAGATAAATGAACAGAGA

ACTCTCTCTGATCAAATTGTGTAATCTTAAGAGTTAAGAGGGAAATCCTCCATTAAT

>JK714338.1 JutESTSSH3(50B) Jute Plasmid Suppression Subtractive cDNA library Corchorus capsularis cDNA similar to Acyl-CoA-binding protein, mRNA sequence

TTTTATTGTTGCTACCATGTTATTCCCGAATTATGCAGGTTTAAACCAGATTTGATTTATCTATCAATAG

TATAAACAAGCCCAAATAAGATGTTTGCTCATCTGATGGAAAATTGCTTGAATTCATAAATTAGCCATAA

TTGCTTAGCT

>JK714337.1 JutESTSSH3(50A) Jute Plasmid Suppression Subtractive cDNA library Corchorus capsularis cDNA similar to Acyl-CoA-binding protein, mRNA sequence

AAGTGGGATGCATGGAAGGCTGTTGAAGGGAAATCAAAGGAGGAAGCTATGAGTGATTATATCACCAAGG

TGAAACAGTTGCTAGAGGAGGCTGCTGCATCTTCTTGATGTCTGACATAACATTGTGTGTTCCCAGCCTT

GCTACATGTTTCTGAAAATGTAAAACTAAAAATAAGTGTTT

>JK714336.1 JutESTSSH3(49) Jute Plasmid Suppression Subtractive cDNA library Corchorus capsularis cDNA similar to Expressed protein, mRNA sequence

TCCCGAGAGCCGTCGCTCCGCACCTGCTCTCGGTTTCCCTCGCATTAGGTTCTTCCGTTACCGGCGCTTG

AGATAACCGTTGCCCTGCTTGCCAAACCTCCAAAAAAAGAAACAAGGAAAGAAAAGTTTTTCTATTTCAT

ATTTTCTTCTTTTCTCCGTTGAATTATTGTAAAATATATTACGCCAGATAATTTCAATTCGATGCCGAAT

TCGAGAAACAGCCGTAAGGAACCGGAGAAGAACCGGAGGGGACGGTTGTCTGAAAAGGCATCGTCGTTTC

ACGGTAGAAGCCCTGCGATGGCACCGGCAGAGTTGAGGCGGCCAAAGACGCTGCCCGACCTATTTACAGA

GAGGACTATTTCCATTCCACTTGATGCGAGGCCAAAATTAACGAAGCTGCTATTAAACGTCACCGTCCAA

GGAAGCCTCGGCGCCATACAGGTGCTGATGTCACCGGAAAACACCGTCGGGGACTTAATAAATGCAGCCG

TTCGTCATTATGCGAAGGAAGGTCGCCGGCCGATCTTGCCGCCGACAGACGCCTCGCTTTTCGACCTTCA

TTATTCTCAATTCAGCTTAGAAAGTTTAGATAGGCAAGAGAAGCTGATGGAGTTAGGATCGAGGAATTTC

TTCCTCTGCACAAAAAAGGCGGCGTTGAAGGACGGTGAGAATAAAACGACATCGTGCTCATCAGAAGCCG

AAAATGTTACGAAGAGCGCCATACCTTGGCTCAAATTCATGGATTCCTTGTTTTAATCCTAAAGAACAAG

CATTTTTTCTTTCTTTAGATTTTTGTGTTGTAATATGTAAAAAAATACGATTGAATTAAGGTTAAAAAAT

AGTTGTGGGTTTAGAATATATTATATTAAGGTTTGCAAAATTGGAGGAAAAAAATTGCATGTAGGTTATA

TATTATTTTTTAATTGTGGAAAAAAAAAAAGAGAAGAGTGTTTATGATTATCTTCTCTTCTGTTTAAAGG

GAAAAGGTT

>JK714335.1 JutESTSSH3(48) Jute Plasmid Suppression Subtractive cDNA library Corchorus capsularis cDNA similar to Disulfide-isomerase-like protein, mRNA sequence

GAATACGAACCGTGAAAGCGTGGCCTATCGATCCTTTAGACCTTCGGAATTTGAAGCTAGAGGTGTCAGA

AAAGTTACCACAGGGATAACTGGCTTGTGGCAGCCAAGCGTTCATAGCGACGTTGCTTTTTGATCCTTCG

ATGTCGGCTCTTCCTATCATTGTGAAGCAGAATTCACCAAGTGTTGGATTGTTCACCCACCAATAGGGAA

CGTGAGCTGGGTTTAGACCGTCGTGAGACAGGTTAGTTTTGCCCTACTGATGGCCGCGTCGCAATAGTAA

TTCAACCTA

>JK714334.1 JutESTSSH3(47) Jute Plasmid Suppression Subtractive cDNA library Corchorus capsularis cDNA, mRNA sequence

CGGAAATTGATCGAATCTATGGACCAATGGAGCATAAGGATCATATGGAAAGCGTTATTTATCCACTCCA

AAGTGGGGAAGCCAAGGCGCACACACACAGATGATTAACTTGAAACGAAGTCAATCTTGAGACCACCAAT

CGTAACTTTGCCAGTCCTAGGCACAAGAGTGGCGATGATAGAATCATCACCTTCGACATCCAAATCTTCC

AACAAATCTGTCAATCCCAACTTCAAGCCGGTGGTTATCTCGGCCGAACTTTTTTGCCTGTGCGGCACAC

TGGAGAAGCTCCCTGCAAATTCTGAATCTTGCGGTCCAGTTGGTGTCTCACCATCCTCATCGTTGAT

>JK714333.1 JutESTSSH3(46) Jute Plasmid Suppression Subtractive cDNA library Corchorus capsularis cDNA similar to 26S ribosomal RNA gene partial sequence, mRNA sequence

TCGCGACGGGGTATTGTAAGTGGCAGAGTGGCCTTGCTGCCACGATCCACTGAGATTCAGCCCTTTGTCG

CTTCGATTCGTCTGCAAAGGATTCTACCCGCCGCTCGGTGGGAATTACGCTTCAAGGAGGCCTGCACGGC

TCATCCGCCGCACCTGCCCG

>JK714332.1 JutESTSSH3(44) Jute Plasmid Suppression Subtractive cDNA library Corchorus capsularis cDNA, mRNA sequence

TTTGGCCTAGATATTGGCTTGCATTTGCCCTCCACTGATGAGAGAAGGCATAAGCCATGGCTCGTTCGCG

CTTAACTGCCGCTTCTTCTCTTTGTTGAATTCTGGATAGGATTTCCTCCATAGTTTCAGACCCTCCACAC

CATTCTACCTCAAGCTCTTGAAGCTTAGCCTCAAGTTTCAACTGATTTTCAATCCTCTTTTGCTTTCACC

TTCAAATGTCGCTCCAAGATGTTAACTAACTCCGGAACGTTAACCGGCGACGTATTGCCCAAATTGAAAA

TTCT

>JK714331.1 JutESTSSH3(43) Jute Plasmid Suppression Subtractive cDNA library Corchorus capsularis cDNA similar to Senescence-associated protein putative, mRNA sequence

TAGGTTGAATTACTATTGCGACGCGGCCATCAGTAGGGTAAAACTAACCTGTCTCACGACGGTCTAAACC

CAGCTCACGTTCCCTATTGGTGGGTGAACAATCCAACACTTGGTGAATTCTGCTTCACAATGATAGGAAG

AGCCGACATCGAAGGATCAAAAAGCAACGTCGCTATGAACGCTTGGCTGCCACAAGCCAGTTATCCCTGT

GGTAACTTTTCTGACACCTCTAGCTTCAAATTCCGAAGGTCTAAAGGATCGATAGGCCACGCTTTCACGG

TTCGTATTCGTA

>JK714330.1 JutESTSSH3(42) Jute Plasmid Suppression Subtractive cDNA library Corchorus capsularis cDNA similar to Triosephosphate isomerase, mRNA sequence

TGCATTTTAAACTCTTATGAAACAGATTCTATGCCTCCCCAAACAGCTCTCCTTGCAAGCACCTTAACTA

AGCTTCAAACTAGCACTTAGGCATTCTTCTTCACCTCAGCAGACTTGATAATATCAATGAACTCCGGCTT

CAGTGAAGCTCCCCCAACCAAGAAACCATCAATATCAGGCCGAGCTGCTAGTTCCTTGCAGTTCGCAGCA

GCGACAGACCCTCCATAAATAATCCGGGTTGATGCAGCAACTTCAGCACTGATGTTTGCTTGAAGCCATT

TCCTCAACTCTGAAT

>JK714329.1 JutESTSSH3(40) Jute Plasmid Suppression Subtractive cDNA library Corchorus capsularis cDNA similar to Ras-related GTP-binding protein, mRNA sequence

ACAAACAGGGTAACGTAAGGAAGATAAAACTTTTGATTGTGGAGGGCATTATAAGGGCTACTTAGATACA

GGCAGGTAAACCACCATTGCCTAAGAGCAGCAAGATGCACTTGCAGTCCGTTCTGCAGGTCTATCCATGA

GAACCATTCCAGCAGGATTTTGTGCTGGCTGAACTCGAGGTAATCTTTTAGCTATTTCATAGAAAATATC

ATTCACATTGGATGCAGTCTTAGCAGAAGTCTCCATGAAGAAAAGACCATTCTCCTGAGCATAGGTTTGT

GCTTCCTCAGCTGCCACCTTCC

>JK714328.1 JutESTSSH3(39) Jute Plasmid Suppression Subtractive cDNA library Corchorus capsularis cDNA similar to Profilin-2, mRNA sequence

AGAAACAAAGTAAGATCATAAGCATTCACTATTACAAACACCAAGCCTTTTGTTTCTGGAAAAAAGGTTC

CCCATGACTTTAATACTCGGTAATACTATTATTTAACTTAACTGCACCTAACCTACTACAGGAGTCGGCA

GATGCATGATGATGATTGTATAGAAGCCAAGAAGAATTGCTGCAACAAATAAGGCAAACTGAATAGATCA

AAGGCCCTGGTCAACAAGATAATCACCAAGCCTTTCAACAATCATGTTGCACTGGCCGGGGGTCATTGGC

TCATCATAGATTCCGATGATCATGGCTGCACCGGTCTTTTTAACAGTAACACCTCCAGGTCCCTTCTTCC

CCCGAATAACAGCTCCGGGTTCACCTTGGATCACCATATACTTTGTGCCACCAAG

>JK714327.1 JutESTSSH3(38B) Jute Plasmid Suppression Subtractive cDNA library Corchorus capsularis cDNA similar to Translation elongation factor 1 alpha, mRNA sequence

TTGGAGTTGTGGCATCCATCTTGTTGCAGCAGCAAATCATTTGCTTGACACCAAGGGTAAAAGCAAGCAA

AGCATGCTCACGGGTCTGACCATCCTTAGAGATACCGGCTTCAAAACCTAAGGATGATCCTGCCAAGGAA

GCTGCCAACTTCACCTCTCAGGTTATCATCATGAACCACCCTGGCCAGATCGGAAATGGATACGCCCCGG

TGCTCGATTGCCACACCTCCCACATTGCTGTCAAGTTTGCTGAACTCTTGACAAAGATTGACAGACGATC

TGGAAAGGAGCTCGAGAAGGAGCCTAAGTTCTTGAAGAACGGTGATGCCGGGATGGTTAAGATGATTCCC

ACTAAGCCCATGGTTGTGGAGACTTTCTCTGA

>JK714326.1 JutESTSSH3(38A) Jute Plasmid Suppression Subtractive cDNA library Corchorus capsularis cDNA similar to Elongation factor 1-alpha, mRNA sequence

CAGTCAAGGTTGGTTGACCTCTCAATCATGTTATCACCCTCAAAACCAGAGATTGGGACGAATGGGATTT

TGTCTGGGTTGTAACCAACCTTCTTCAGGTAAGAAGAGACTTCCTTAACAATTTCATCATACCTTGCCTT

GGA

>JK714325.1 JutESTSSH3(36) Jute Plasmid Suppression Subtractive cDNA library Corchorus capsularis cDNA similar to Glutamate decarboxylase putative, mRNA sequence

CTAGTGACCTCCTCTTGAATCTCCCTCTCACTCTTTTTATGCTTCCTTTGCACTTCTTCAGCAACAGCCT

CAACATGTGCAGCTTGAACGGCTGAACGACTTGGGAGCAAGTCTAGTTCCTTCACAACCGTCTCGATATG

AGACACAAGTCGCTCTGCCAGGCTGCGGCTGAAGTCCTCCCTAACAACTATGCGAAGAACTGCAACATGT

TGCGCATCAGCAGGCATTGTATAAGCCGGGATGATCCACCCGAACCTTCTTAAGTTCTCGGCTATCTCGA

AAACAGTGTGTTTACTGCTGTCTTTTAAGGCAAAAGCTACAAGGGGAACACCCACATCCTTGGACACAAC

TTCAAACCGGCCTGTTTTCTCAATCCCTTCTCTCAGAATTCTTGCATTTTCGATGCAGTTCTCCATGATA

TTCTTGTATCCCTCAAAGCCAAGCCGAATTAACTGATAATATTGAGCAATAATTTGACTAGAGCCTTTAG

AGAAGTTTAAGGTAAAAGTGGGCTGATCAGATCCAAGGTAGTTGATGTGAAAGACGAGGTCATCAGGCAA

ATCCTCCTTGGTTCGCCACACAACCCAACC

>JK714324.1 JutESTSSH3(35) Jute Plasmid Suppression Subtractive cDNA library Corchorus capsularis cDNA similar to Disulfide-isomerase-like protein, mRNA sequence

GAATACGAACCGTGAAAGCGTGGCCTATCGATCCTTTAGACCTTCGGAATTTGAAGCTAGAGGTGTCAGA

AAAGTTACCACAGGGATAACTGGCTTGTGGCAGCCAAGCGTTCATAGCGACGTTGCTTTTTGATCCTTCG

ATGTCGGCTCTTCCTATCATTGTGAAGCAGAATTCACCAAGTGTTGGATTGTTCACCCACCAATAGGGAA

CGTGAGCTGGGTTTAGACCGTCGTGAGACAGGTTAGTTTTACCCTACTGATGGCCGCGTCGCAATAGTAA

TTCAACCTA

>JK714323.1 JutESTSSH3(34) Jute Plasmid Suppression Subtractive cDNA library Corchorus capsularis cDNA similar to Triosephosphate isomerase chloroplast precursor, mRNA sequence

ATTAGAGAGAAACAATTTATTTTCCTATCACTTTGGGGCAAAAGAACTCAATAAGTAATCAAAACTCCAA

TGGATAAAACACTCTCATAAGGCTCCATGAAGCTAAGTCGCATGAAGCCTCAATCTGAAATTGCTTATGT

TTGTCCTTTATTTGTTATTTTAATCCGCTTTTTATATTTAAGCAGCAACCTTCTTTGCTGTTACAGAGTT

GACAATTGTAGCAAACTCAGGGCCCTTCAAGGAAGCACCACCGACAAGGAACCCATCAATATCTTCTTTC

TTTGCAAGCTCGACACAGTTGCCTCCGTTAACTGACCCTCCATATATGATACGCGTTTTTGATGCTACTT

CTTCGGAAACATTCTTTTTAAGCCAATCACGAAGAGCTACAT

>JK714322.1 JutESTSSH3(33) Jute Plasmid Suppression Subtractive cDNA library Corchorus capsularis cDNA similar to Auxin-responsive factor TIR1-like protein, mRNA sequence

AACTACTTAAATAAGCAGAAAAGAAATGGCAAGAGCCATGCAGCAGCTATAAAATCCACACAAATTCTGG

TGCATCTTCCCTATGCCCAACCACCGATCGATAGAGATACATCTTCTCTACTTTCTGGTCATCATCGAGG

TTAAATTCCATCTGATCACTATCATTTATGAGCTCCACCTTCAGTCTTGGCATCTTCTCCGCAAGTGTCT

TGCAGGCTCCGAGGGTAACTTCACAGGAGGACATCCAAAGGGATCGCATTGTTTCATACTTTCCCACGTC

CTCTAGAAGTGCTGCATTACCAAAGGGGCAATCCCTGATCTCTAACTTGCGAAGCTTCTTGCAACCATTC

AACACATAAAGCATACCTTTGTCACTCTCGCCAGCAAAAGCAATGGACAACATTTCAAGCTGCTCAGCAT

ACATTCCTATGTAAAGGAAAACCTGATCGGTCAGCAGGCCTGAGAGTGATAACCGCTTGAGACCCTTGCA

AGACTGAACAATGGCCCCAAAACCTTCATCCAATGGCAGATTGGTCACAGGGTCTGGTTTTGTGGGTTCA

AGGATGCACAATCTGAAGCGTATGAAATTTGGG

>JK714321.1 JutESTSSH3(32) Jute Plasmid Suppression Subtractive cDNA library Corchorus capsularis cDNA similar to 29 kDa ribonucleoprotein chloroplast precursor, mRNA sequence

TTAAAAAAGCAAAGAAGCCATGTCTGCAACATCTGCCTCATCTCTTATTCTTCCATCTCTCAAACCCAAA

ACCTTATCTTCATCAAACCCTAGACCCATTTCAGTTTCCCTCTTCTCTGCTTCCTACAAGCTTCATGCTA

AGCCTCTCTTCTGTTCTGCTCTTCAAAGTTTTGACCTTTTAGCCAAAAAGTCTTCTTCTTCTTCTTCTTC

TTCTTCTTCTAAGTTTGTTAGAAACGTTGCCGTTTCTTCTGAGTATGGTCAGGAAGAGGAGCTTTTTGGC

AGTGATGAAGAAGAAGATGCTCCCAGTTTTTCCGATGACTTGAAACTTTTTGTGGGTAATCTTCCTTTTA

CTGTGGATAGTGCCCAGCTTGCTGCTTTGTTTGAAAGTGCTGGAAATGTTGAGATGGTTGAGGTGATTTA

TGACAAGGTTACTGGGAGAAGTAGGGGTTTCGGTTTTGTGACCATGTCCTCAGCCCAGGAAGTTGAAGCT

GCTGCTCAACAGTTCAATGGCTATGAACTGGAAGGAAGGGCATTGAGGGTAAACGCTGGACCTCCTCCTC

CTCGTCGTAGTGAAGAATTTTCTCCTAGAGGGCGAAGAGGCGGAGGCGGTGCTCCAGCTATGGGTTCTTC

AAATCGTGTGTATGTTGGTAACCTTTCATGGGGTGTTGATGATTTAGCACTTGAGACCTTGTTCAGTGAG

CAAGGCAAGGTTGTAGAAGCAAAAGTCGTTTATGATAGGGACAGTGGTAGATCGAGAGGCTTTGGCTTTG

TAACTTACAACTCTGCTGAAGAGGTTGACAGTGCCATTCAATCCTTGGACGGTGTTGATTTGGATGGCAG

ACGAATTCGAGTCTCCGTGGCAGAGTCTTAGCCAAAGCGTCAATTTTTGAGTGTGATTTTGGTGATTGG

>JK714320.1 JutESTSSH3(30) Jute Plasmid Suppression Subtractive cDNA library Corchorus capsularis cDNA similar to 26S ribosomal RNA gene partial sequence, mRNA sequence

TGGAAATCAGAATCAAACGAGCTTTTACCCTTTTGTTCCACACGAGATTTCTGTTCTCGTTGAGCTCATC

TTAGGACACCTGCGTTATCTTTTAACAGATGTGCCGCCCCAGCCAAACTCCCCACCTGACAATGTCTTCC

GCCCGGATCGACCGACAGATGCCGGTCTTGGGTCTAAAAAGAGGGGCGTAGCCCCGCTTCCGATTCACGG

AATAAGTAAAATAACGTTAAAAGTAGTGGTATTTCACCGTTGCCGAAGCTCCCACTTATACTACACCTCT

CAAGTCATTTCACAAAGTCGGACTAGAGTCAAGCTCAACAGGGTCTTCT

>JK714319.1 JutESTSSH3(29) Jute Plasmid Suppression Subtractive cDNA library Corchorus capsularis cDNA similar to Disulfide-isomerase-like protein, mRNA sequence

GAATACGAACCGTGAAAGCGTGGCCTATCGATCCTTTAGACCTTCGGAATTTGAAGCTAGAGGTGTCAGA

AAAGTTACCACAGGGATAACTGGCTTGTGGCAGCCAAGCGTTCATAGCGACGTTGCTTTTTGATCCTTCG

ATGTCGGCTCTTCCTATCATTGTGAAGCAGAATTCACCAAGTGTTGGATTGTTCACCCACCAATAGGGAA

CGTGAGCTGGGTTTAGACCGTCGTGAGACAGGTTAGTTTTACCCTACTGATGGCCGCGTCGCAATAGTAA

TTCAACCTA

>JK714318.1 JutESTSSH3(28) Jute Plasmid Suppression Subtractive cDNA library Corchorus capsularis cDNA similar to Vacuolar ATP synthase 16 kDa proteolipid subunit, mRNA sequence

AAATTTCCCAGTTCTTTCTCCTTCTCTCTCTTCTTCTGTTCGCTGAAGAAATCCACCGATCAAGTTCTCC

CAAATCTCTTCTCTATCATCGGTGAAATTTTTGATTAGCGAAGATGTCTTCCACATTTAGCGGCGATGAA

ACTGCTCCTTTCTTTGGCTT

>JK714317.1 JutESTSSH3(26) Jute Plasmid Suppression Subtractive cDNA library Corchorus capsularis cDNA similar to Cyclin-dependent kinases regulatory subunit, mRNA sequence

TATTATATATATCCAAGATGCAGTAGCAATGACATTATGTCATCAACACCACAACAGGGGAAAAGTAAGA

AACAAGACTAACATGAAATTCTCCCATACATAAGAGAACAACAATAAAAACCAGGAATCCCAGAAAAATA

TCCAAACTGTTTATGTAGAGCATAAACACTAGTAAATATATATTTATGTTTTCATTTCCATTTTAGATAC

TGAGCACTAATTACTTAGCAGGAATAAGTTGCTGAGCCTGATTCTCTTGCTGCTGCTGATAGTTAAGAGG

CCTCCTGAAGAGCATGATATGTGGCTCAGGGCGATGAATGGCATAATGAACCCAACCGCGACTCTGCTGC

ACTCCAATGGCACGCCACTCATTCTCAGAGAGGAGGCGATTCTTAGGGAGGAGTTTGGCCACTTCTGGAG

GAAGAACAACGTGCCTGC

>JK714316.1 JutESTSSH3(25) Jute Plasmid Suppression Subtractive cDNA library Corchorus capsularis cDNA similar to 4-coumarate:CoA ligase, mRNA sequence

ACAAATAAAAATAAGACTCTGAAATCTTCATCCATTTACCAAGAAAAAAGAAAGAAAAACAGAATATTTA

GTCAAATTATATGTCCTCTAACAAATCGAATCCTTTAATCTCATAATTTACATTCATTCCAAACCCACTT

TGTCTATAAGCATAGAATGGAAGAACATATATCAACAGTCCCAATATTGCTTCATAGTAGGGACCCCTAG

CACTCGTGTAATGGTAATATTATTTCAATCTTTTTCGTTGCAGTTTTTGGAATTGGAAACAAAGTTTCAG

TAGTTTCCAGAAGCCAATTTAGCTCTCAATTCCTTTCTCAAAATCTTGCCTGATGGTGCCTTTGGAATTG

CTTCCATGAAGAACACACGGCTTATTCTCTTGTAGAACACCACCTGCTTAAAAATATATTGCTTGATTTC

ATCCTCAGTGATTTCTGATTTCTCTGATTTCACAACAAATGCAACAGGAACTTCTCCAGCTAGCTCATCT

TTCATTGCAACGACAGCAGCATCAATAATCTCAGGGTGGGCAATGAGCATGGCTTCAAGCTCAGCAGGAG

CAACCTGGAAACCCTTATATTTGATC

>JK714315.1 JutESTSSH3(24) Jute Plasmid Suppression Subtractive cDNA library Corchorus capsularis cDNA similar to Protein serine/threonine kinase BNK1-like, mRNA sequence

AAGGTTTTGATCATTTGGTCCCTGATTTAAGTCTACAGCTCTGCGTCCTGTCAAAAGCTCTAGAAGAACC

ACTCCAAATGCATAGACATCACTTTGTAAAGTTAGCTTTCCTGTCGATGTATACTCAGGATCGAAATAGC

CAAATGTGCCTAAAACTCTGGCTGTCACATAAGTCTCTTGACCCTCTGGCATCAGTTTGGCTGGTCCAAA

ATCAGATATCTTTGCTTCAAAGTTGTCATTTAGAAGAACATTGGTGGACTTGAAGTCTCTGTGAACTATT

GGAATTCCAACAGCAGAACTTGAATGTAGATAAGCAAGTCCCCTGGCTGCACCAATTGCCACTTTGAGCC

TTGAAGGCCAATCCATTTTTGTTTCCCCAATACCATTTAAGTGATCTTGC

>JK714314.1 JutESTSSH3(23) Jute Plasmid Suppression Subtractive cDNA library Corchorus capsularis cDNA similar to Ribonucleoprotein A chloroplast precursor, mRNA sequence

TAAAAAAGCAAAGAAGCCATGTCTGCAACATCTGCCTCATCTCTTATTCTTCCATCTCTCAAACCCAAAA

CCTTATCTTCATCAAACCCTAGACCCATTTCAGTTTCCCTCTTCTCTGCTTCCTACAAGCTTCATGCTAA

GCCTCTCTTCTGTTCTGCTCTTCAAAGTTTTGACCTTTTAGTCAAAAAGTCTTCTTCTTCTTCTTCTTCT

TCTTCTAAGTTTGTTAGAAACGTTGCCGTTTCTTCTGAGTATGGCCAGGAAGAGGAGCTTTTTGGCAGTG

ATGAAGAAGAAGATGCTCCCAGTTTTTCCGATGACTTGAAACTTTTTGTGGGTAATCTTCCTTTTACTGT

GGATAGTGCCCAGCTTGCTGCTTTGTTTGAAAGTGCTGGAAATGTTGAGATGGTTGAGGTGATTTATGAC

AAGGTTACTGGGAGAAGTAGGGGTTTCGGTTTTGTGACCATGTCCTCAGCCCAGGAAGTTGAAGCTGCTG

CTCAACAGTTCAATGGCTATGAACTGGAAGGAAGGGCATTGAGGGTAAACGCTGGACCTCCTCCTCGTCG

TAGTGAAGAATTTTCTCCTAGAGGGCGAAGAGGCGGAGGCGGTGCTCCAGCTATGGGTTCTTCAAATCGT

GTGTATGTTGGTAACCTTTCATGGGGTGTTGATGATTTAGCACTTGAGACCTTGTTCAGTGAGCAAGGCA

AGGTTGTAGAAGCAAAAGTCGTTTATGATAGGGACAGTGGTAGATCGAGAGGCTTTGGCTTTGTAACTTA

CAACTCTGCTGAAGAGGTTGACAGTGCCATTCAA

>JK714313.1 JutESTSSH3(22B) Jute Plasmid Suppression Subtractive cDNA library Corchorus capsularis cDNA similar to Ribosomal protein S8, mRNA sequence

CATTGCTTGAAGGGTGCAGCATCCACTTGAACAATGGCGCTCTTTACAAGAGTTTGTGTCCTGACAAGCT

CATTGTTTGAGGCATTGTATACCACATCAAGAATACGGGTCTTGCGGGTGACAGCCTCACTACCCCATGA

ATAGTTGCCTGTATCCAATCTCAATGCTCTCCACTTCACATTCCCTCCCCTAACTCGTATTCTCCTAACA

GTCTTGTTACTTGATAGCTTAGTGTTTGCTGGCTGACGACCAAGTTCATACTTTCTCTTCTTCCTCCAAG

CCTTCTTCTTGCCACCAGTGGCACGCC

>JK714312.1 JutESTSSH3(22A) Jute Plasmid Suppression Subtractive cDNA library Corchorus capsularis cDNA, mRNA sequence

TCCTTTCTCCTCGGTCCACAGAGACAAAATGTAGGACTAGGACTGGCGCCAATAGCTAATCACGGAAGAA

AGGACTCACTGAGCCGGGATCACTAACTAATACTAATCTAATACTAATAGAATAGAATACTAATCTAATA

CTAATTCTAATACTAATAGAATAGAAAAAATAGAATAGAAAATACTAATAGAATAGAAAGAAAAGAACTG

TCTTTTCTGTATACTTTCCCCGGTTCCGTTGCTACCGCGGGCTTTACGCAATCGATCGGATCATATAGAT

ATCCCTTCAACACAACATAGGTCGTCGAAAGGATCTCGGAGACTCACCAAAGCACGAAAGCCAGGATCTT

TCAGAAAATGGATTCCTATTCGAAGAGTGCATAACCGCATGGATAAGCTCACACTAACCCGTCAATTTGG

GATCCAATTCGGGATTTTCCTTGGGGGGTATCAGTAAGGAATTGGAATGTAATAATATCGATTCATATGG

ATACAGAGGAAAAGGTTCTCTATTGATTCAAATGCT

>JK714311.1 JutESTSSH3(21) Jute Plasmid Suppression Subtractive cDNA library Corchorus capsularis cDNA similar to Cellulose synthase catalytic subunit, mRNA sequence

CATTAATACACTCTTGGTAATACACCATGATTTCATACAATTACCTATATTATAAATATATATATTCTGC

TTCTGCACATAACAATAACATGATATGAATCGACATGATACACATTTTATAATAATAAAATATTATAATA

ACGACAAAACAACAAGAGGTAGCAGAAGGAAAAATCCCACCTTGACCATGACCTTGACCAAAAGCAACCA

AACGTTGAGTAGTTGAGATGAGTCTTTCCTGATAATTTTGTCTAACTAACAATTGATTCCACATTGTTCT

ACATCTGGTCCGGTAACTCTGGTGGTGAAGGGATCAATCCGCACCCACAACAATGAGAAGATAGACGCCA

GCAGAATGGACCATACCACAACAATGGTGGGTGTTCGGTTTTGGCGTCCCATCAAACCTTTCAGGAAGGG

GTAGAGATGGATGATGACCCAGAAGGCAAAGAAGAGCTTCCCAAACAAGGGCCCCCATGATTGGTAACCA

CTGTTGATGGCATACGAGATTCCTGCCACAACCCCTACCAAGTTTATGATCAGCAGCGTTGTTGGTGGGA

TCAGAAGGGTTGTCCATTTGACAATGTATAGGTCGGCAAACTCATCATCCACATCATCTCCGCCTGATTT

TGATGTCAAGGTGAAAGAAATCTCAATTCCGGCAATTACTTTAAGTTTC

>JK714310.1 JutESTSSH3(20) Jute Plasmid Suppression Subtractive cDNA library Corchorus capsularis cDNA similar to Phytochrome A, mRNA sequence

TCCACCACCTGCATGTGTTATCCTGAGTTCCAAATGGGCAAGATGAACAGACTGTCCTAGTTGATCTTTT

GTTAAGCTTGCTAAGAGAACAAGCTGGCCTCCATTTGGTGTAAAATTAACTGATATCAACAAGAAATCAG

CTAAGACCTGTTGAAGTCTAACACTATCTCCATATAAGGTCTCAGTCATGACCTCTTCTGCTGTGTCGTT

TACTATTCGGATACCCTTCCCAGTGCTTTTCATCACGACTTGACTAATAGAAGCAACCAATACCTCATTC

AGGGTGAATTCAATCATTTCAAGATCCAAGTAGCCATCAATGATGCTATCAAGATCTGAATCATCAAGTA

TCTTGCTGAGCTGGTGCTGGCACAGA

>JK714309.1 JutESTSSH3(19B) Jute Plasmid Suppression Subtractive cDNA library Corchorus capsularis cDNA similar to 14-3-3 family protein, mRNA sequence

CTGTGATAATCACCCTTCATCTTAAGGTAAAAAACCTTGGACTCGGCAGATGAGGCTGAAGGAATGAGGT

GAGACTCAAGAAGGCTCAAAATGCCATCACAGATCTTGCTGAGCTCGGCTTCAATCTTGCCCCTATAATC

CTTGATGATGGCAACATGATCCTCATTTCCTCTGCTCTCTTCCTTCTGCTCAATGGAAGAGATGATCCTC

CATGAAGCCCTCCTAGCTCCAATCACATTTTTGTAAGCAACAGAAAGAAGGTTCCCCTCCTCAACTGTTA

GCTCCTCAACATCGACAGTTTTTGCAACTTTCTCCATGAACTCCACCATTTCCTCATAGCGTTCAGCCTG

CTCAGCCAACTTGGCCAT

>JK714308.1 JutESTSSH3(19A) Jute Plasmid Suppression Subtractive cDNA library Corchorus capsularis cDNA similar to 14-3-3 family protein, mRNA sequence

TATCCTTGTAAGATTCCTCACCCAATGTATCCAACTCAGAAATAGCCTCATCAAAGGCATTTTTAGCAAG

ATTGCAAGCACGGTCAGGTGAGTTGAGGATTTCATAGTAGAAGACAGAGAAGTTAAGTGCAAGTCCCAGC

CTGATTGGGTGAGTAGGAGCAAGCTCAGCAAGAGCAATATCCTGGGCAGACTTATAAGCCAACAATGTGC

TTTCAGCAGCTTCTTTCCTCTCAGCACCAGTCTTAAACTCAGCAAG

>JK714307.1 JutESTSSH3(18) Jute Plasmid Suppression Subtractive cDNA library Corchorus capsularis cDNA similar to Expressed protein, mRNA sequence

TCCCCTCTCCATCTCCTTCTCCGCCTACCCTTACCTCCATCAACTTATCAATCCCCAACCCTCCACCGTT

CCCACCGCCGTCGAAAGCCATAATTTCAACCCTTCTTTTCAACCCACGAAAACCACCATCTTTTCTCTCC

TCTATTCTCTCTTCGTCTCAATCTTCACTTCCTTAGCCACCGGTTCAATCACTTACAGCGTTTTCCATGG

CTTTTACGGCCAACCCATCAAGCTAAAACCAGCTATCAAGTCAGCTTTCACTTCATTTTTCCGTCTCTTA

TCAACTAGCTTCCTAACGGGGCTTATTCTGTTTGGGTTCTTTTTCCTTCTTGGTTTAATCATTATTTCAT

TAGCCATGGCAACCTTACTTTTAGGGTTCCAAGTTGATTTTTCTTCACCTAATTGTATTTTCCTTTTTAT

ATATTCTATGACTGCTTTCATCTGT

>JK714306.1 JutESTSSH3(17) Jute Plasmid Suppression Subtractive cDNA library Corchorus capsularis cDNA similar to Phytochrome A, mRNA sequence

TCCACCACCTGCATGTGTTATCCTGAGTTCCAAATGGGCAAGATGAACAGACTGTCCTAGTTGATCTTTT

GTTAAGCTTGCTAAGAGAACAAGCTGGCCTCCATTTGGTGTAAAATTAACTGATATCAACAAGAAATCAG

CTAAGACCTGTTGAAGTCTAACACTATCTCCATATAAGGTCTCAGTCATGACCTCTTCTGCTGTGTCGTT

TACTATTCGGATACCCTTCCCAGTGCTTTTCATCACGACTTGACTAATAGAAGCAACCAATACCTCATTC

AGGGTGAATTCAATCATTTCAAGATCCAAGTAGCCATCAATGATACTATCAAGATCTGAATCATCAAGTA

TCTTGCTGAGCTGGTGCTGGCACAGAGC

>JK714305.1 JutESTSSH3(16) Jute Plasmid Suppression Subtractive cDNA library Corchorus capsularis cDNA similar to Disulfide-isomerase-like protein, mRNA sequence

GAATACGAACCGTGAAAGCGTGGCCTATCGATCCTTTAGACCTTCGGAATTTGAAGCTAGAGGTGTCAGA

AAAGTTACCACAGGGATAACTGGCTTGTGGCAGCCAAGCGTTCATAGCGACGTTGCTTTTTTATCCTTCG

ATGTCGGCTCTTCCTATCATTGTGAAGCAGAATTCACCAAGTGTTGGATTGTTCACCCACCAATAGGGAA

CGTGAGCTGGGTTTAGACCGTCGTGAGACAGGTTAGTTTTACCCTACTGATGGCCGCGTCGCAATAGTAA

TTCAACCTA

>JK714304.1 JutESTSSH3(15) Jute Plasmid Suppression Subtractive cDNA library Corchorus capsularis cDNA similar to Adenine phosphoribosyltransferase-like, mRNA sequence

CAGCTTGAAAACATAAACAATGGCATTAAACAGAGAATCTCTCGCATCAACAAACAACAACTTTCAATGA

AAGCTACTGTATAGCCATGGTTCTCATAAAAATACAGGCAACCTCCCTTAAAAGCAGGAATATGACCTTT

GTAGTAACTAGTAAGCACCCACCTCCAAACTTTCTCAATCGAGAAAACCGCGAAAGAAACATCAAAGTAA

CCTAACTACCACAGTTCTTAATTCAATCTAATGTGACCTTAATTACCAAGTTTCACCCTTTTTATTCAAC

CAAGTAATTTTGTTAGGCTTTGGGCAAAGTTCTAGCCATATTGCAAACTCAAAAGGCCAAAGGGACCTGC

TCTTCAACCGCAAAACTTGTGTGATCAAGTGGAGGAGCTTACAAGAACAAACAGTGGTGTCTTTCCTAAA

CGTTCACGACCCTTTAGCTCCGGTAATTCAAAAACACAAGCACACTCCACAACATGCACTCCTACACGCT

CAAGTAGCCTGATTGCAGCACATAAAGTTCCGCCAGTC

>JK714303.1 JutESTSSH3(14) Jute Plasmid Suppression Subtractive cDNA library Corchorus capsularis cDNA similar to Photosystem I reaction center subunit II, mRNA sequence

ACACAGGAGAACAGAGGCAATATATTGGAGAAAAAGGAAGCGATTAACATAAATTAACATTAATTAACAG

AAAACATGTATAATCACATGGATATTACATAATCGTCTTTCTCAATTCTCACTAAGCACCGTCTCTCAAA

CGTTCATCTCTTACAAGTCATAAGGTTGCTTCCCAGTGAACTTAACCTCAATGGGACTCACATTCTTCCC

AATCGATCTGAAGTTCTGCCCAACTCCTTGACGACCAGGGTTCACTTTCTCAGGATAAACTCCATCTTTG

GGGTGAAGGTATTGAACCTCACCATTAGGGAAAACCCTGTAAAACTG

>JK714302.1 JutESTSSH3(13) Jute Plasmid Suppression Subtractive cDNA library Corchorus capsularis cDNA similar to SKP1 component-like 1, mRNA sequence

AAAGGCAATTAAAAAGAAAAAGAAACATAGAAAAAAGATATGTTCAAATTCAATTCGATATCCTTCACTC

AAAGGCCCACTGATTCTCCCGACGAACCTCCTCCTCCTCTTCCGGAGTAAAATCATTCTTAATATTGAAA

GTCTTACGGATCTCCTCAGGCGTTTTCCCCTTTATCATATCCGCCACCGTTTGACACGTCAGGTCCAGCA

GGCTCTTGATGTTCAAATAGTTAGCAGCCAGGATGAGATCGAAGAGGGTAGCTTGATCGACCCTGGCGAA

ATCAGCATCCCAAAGCTTAAGCTCATCTTCGACGGAGGCAGAACGATCGTCGGATTTAGGTGCTTCCACA

TGCTTCTTGCA

>JK714301.1 JutESTSSH3(12) Jute Plasmid Suppression Subtractive cDNA library Corchorus capsularis cDNA similar to Al-induced protein, mRNA sequence

AGTGGAACCTTACCAAATTGATCAGAAGCTACAAACAAGGTGGAGGTTGATTTGTCATAGACAATGAAAG

CAAAGCTCCCACTTAAATGCCCAACCACATGGTTTGGAGGATAAGGTGCTCTGTCTCTAAGAGCCTTGTA

AGCTTCAATAACCAAAATCGCTTCATTTGCAGACTTTGCAAGCCCATATTGCTGCTTTAAACTCCCCAAG

TTGTCAAGTGCACCCTCAAACAAGCAGAAGATTTCATCTTTCACAGCAAATGATCTGGGCTGTAGGGCGG

ACTCATTCTGGTGAGAATAAGCGAATTGGACATGATCTCCGATCTGCATGGACACGGCGGAAGAGTTGTT

GTCAAGGAACCTTTTCACCAGAGCATCCGACGTTATCTTAGGCGACGGAGTCCGGCAACCGGCGGCCACC

AG

>JK714300.1 JutESTSSH3(11) Jute Plasmid Suppression Subtractive cDNA library Corchorus capsularis cDNA similar to Al-induced protein, mRNA sequence

AGTGGAACCTTACCAAATTGATCAGAAGCTACAAACAAGGTGGAGGTTGATTTGTCATAGACAATGAAAG

CAAAGCTCCCACTTAAATGCCCAACCACATGGTTTGGAGGATAAGGTGCTCTGTCTCTAAGAGCCTTGTA

AGCTTCAATAACCAAAATCGCTTCATTTGCAGACTTTGCAAGCCCATATTGCTGCTTTAAACTCCCCAAG

TTGTCAAGTGCACCCTCAAACAAGCAGAAGATTTCATCTTTCACAGCAAATGATCTGGGCTGTAGGGCGG

ACTCATTCTGGTGAGAATAAGCGAATTGGACATGATCTCCGATCTGCATGGACACGGCGGAAGAGTTGTT

GTCAAGGAACCTTTTCACCAGAGCATCCGACGTTATCTTAGGCGACGGAGTCCGGCAACCGGCGGCCACC

AG

>JK714299.1 JutESTSSH3(10) Jute Plasmid Suppression Subtractive cDNA library Corchorus capsularis cDNA, mRNA sequence

TCTAGCTGCTGGAATTCTCCTTCAAGTTCCTCCCACTCTTGATGTGACAATGATACAACATTCCACCCAG

AAGCAGCGATATGACGACGTTTCAAGATGGTATGCCCCAAAGGTGTTCCACTATTCCTTGAGAAATGTGT

TGGTCCATCTATCTCCAAAACAACCTTTTTATCAATCACAGCAGCATCCACGGTATATACATCCACAGAG

TATTCTCTTACCCAATCAAGTCCA

>JK714298.1 JutESTSSH3(9) Jute Plasmid Suppression Subtractive cDNA library Corchorus capsularis cDNA similar to Polyphenol oxidase A1 chloroplast precursor, mRNA sequence

CGGAAATTGATCGAATCTATGGACCAATGGAGCATAAGGATCATATGGAAAGCGTTATTTATCCACTCCA

AAGTGGGGAAGCCAAGGCGCACACACACAGATGATTAACTTGAAACGAAGTCAATCTTGAGACCACCAAT

CGTAACTTTGCCAGTCCTAGGCACAAGAGTGGCGATGATAGAATCATCACCTTCGACATCCAAATCTTCC

AACAAATCTGTCAATCCCAACTTCAAGCCGGTGGTTATCTCGGCCGAACTTTTTTGCCTGTGCGGCACAC

TGGAGAAGCTCCCTGCAAATTCTGAATCTTGCGGTCCAGTTGGTGTCTCACCATCCTCATCGTTGAT

>JK714297.1 JutESTSSH3(8) Jute Plasmid Suppression Subtractive cDNA library Corchorus capsularis cDNA similar to Senescence-associated protein, mRNA sequence

GAATACGAACCGTGAAAGCGTGGCCTATCGATCCTTTAGACCTTCGGAATTTGAAGCTAGAGGTGTCAGA

AAAGTTACCACAGGGATAACTGGCTTGTGGCAGCCAAGCGTTCATAGCGACGTTGCTTTTTGATCCTTCG

ATGTCGGCTCTTCCTATCATTGTGAAGCAGAATTCACCAAGTGTTGGATTGTTCACCCACCAATAGGGAA

CGTGAGCTGGGTTTAGACCGTCGTGAGACAGGTTAGTTTTACCCTACTGATGGCCGCGTCGCAATAGTAA

TTCAACCTA

>JK714296.1 JutESTSSH3(7B) Jute Plasmid Suppression Subtractive cDNA library Corchorus capsularis cDNA similar to Translation elongation factor 1 alpha, mRNA sequence

TTTGGAGTTGTGGCATCCATCTTGTTGCAGCAGCAAATCATTTGCTTGACACCAAGGGTAAAAGCAAGCA

AAGCATGCTCACGGGTCTGACCATCCTTAGAGATACCGGCTTCAAAACCTAAGGATGATCCTGCCAAGGA

AGCTGCCAACTTCACCTCTCAGGTTATCATCATGAACCACCCTGGCCAGATCGGAAATGGATACGCCCCG

GTGCTCGATTACCACACCTCCCACATTGCTGTCAAGTTTGCTGAACTCTTGACAAAGATTGACAGACGAT

CTGGAAAGGAGCTCGAGAAGGAGCCTAAGTTCTTGAAGAACGGTGATGCCGGGATGGTTAAGATGATTCC

CACTAAGCCCATGGTTGTGGAGACTTTCTCTGA

>JK714295.1 JutESTSSH3(7A) Jute Plasmid Suppression Subtractive cDNA library Corchorus capsularis cDNA similar to Elongation factor 1-alpha, mRNA sequence

CAGTCAAGGTTGGTTGACCTCTCAATCATGTTATCACCCTCAAAACCAGAGATTGGGACGAATGGGATTT

TGTCTGGGTTGTAACCAACCTTCTTCAGGTAAGAAGAGACTTCCTTAACAATTTCATCATACCTTGCCTT

GGA

>JK714294.1 JutESTSSH3(6B) Jute Plasmid Suppression Subtractive cDNA library Corchorus capsularis cDNA similar to Ribosomal protein S8, mRNA sequence

CATTGCTTGAAGGGTGCAGCATCCACTTGAACAATGGCGCTCTTTACAAGAGTTTGTGTCCTGACAAGCT

CATTGTTTGAGGCATTGTATACCACATCAAGAATACGGGTCTTGCGGGTGACAGCCTCACTACCCCATGA

ATAGTTGCCTGTATCCAATCTCAATGCTCTCCACTTCACATTCCCTCCCCTAACTCGTATTCTCCTAACA

GTCTTGTTACTTGATAGCTTAGTGTTTGCTGGCTGACGACCAAGTTCATACTTTCTCTTCTTCCTCCAAG

CCTTCTTCTTGCCACCAGTGGCACGCCCTCTTGTGCATGGAATCCCTGTTTTTTTGTCTTTTATGACTTC

CCTGAAAAAAGGGAATATAAAAAATTTTCCTTCAGACAGTCTTTTTAACACCCTGCGCTACTCGAATTTC

GTGCATTAGCCCCAATGAAAGAGAGAGTCTGTCTGAACTTAAAACATTACTGCATAGTCTCAGTCCAAGG

TGTTGTTGACACTGACCATTCAATAGATGTTAAGCCTCTGTATGACAATATAGGCCAA

>JK714293.1 JutESTSSH3(6A) Jute Plasmid Suppression Subtractive cDNA library Corchorus capsularis cDNA, mRNA sequence

TCCTTTCTCCTCGGTCCACAGAGACAAAATGTAGGACTAGGACTGGCGCCAATAGCTAATCACGGAAGAA

AGGACTCACTGAGCCGGGATCACTAACTAATACTAATCTAATACTAATAGAATAGAATACTAATCTAATA

CTAATTCTAATACTAATAGAATAGAAAAAATAGAATAGAAAATACTAATAGAATAGAAAGAAAAGAACTG

TCTTTTCTGTATACTTTCCCCGGTTCCGTTGCTACCGCGGGCTTTACGCAATCGATCGGATCATATAGAT

ATCCCTTCAACACAACATAGGTCGTCGAAAGGATCTCGGAGACTCACCAAAGCACGAAAGCCAGGATCTT

TCAGAAAATGGATTCCTATTCGAAGAGTGCATAACCGCATGGATAAGCTCACACTAACCCGTCAATTTGG

GATCCAATTCGGGATTTTCCTTGGGGGGTATCAGTAAGGAATTGGAATGTAATAATATCGATTCATATGG

ATACAGAGGAAAAGGTTCTCTATTGATTCAAATGCT

>JK714292.1 JutESTSSH3(5B) Jute Plasmid Suppression Subtractive cDNA library Corchorus capsularis cDNA similar to Ribosomal protein S8, mRNA sequence

CATTGCTTGAAGGGTGCAGCATCCACTTGAACAATGGCGCTCTTTACAAGAGTTTGTGTCCTGACAAGCT

CATTGTTTGAGGCATTGTATACCACATCAAGAATACGGGTCTTGCGGGTGACAGCCTCACTACCCCATGA

ATAGTTGCCTGTATCCAATCTCAATGCTCTCCACTTCACATTCCCTCCCCTAACTCGTATTCTCCTAACA

GTCTTGTTACTTGATAGCTTAGTGTTTGCTGGCTGACGACCAAGTTCATACTTTCTCTTCTTCCTCCAAG

CCCTTCTTCTTGCCACCAGTGGCACGCCTCTTGTGCATGAATCCCTGTGTTTTTTCTTTTGACTTTCCTG

AAAAAGGAATATAAAAATTTTCCTTCAGACAGTCTTTTTACACCTGCGCTACTCGAATGTTCGTGCATTT

ATGCCATGAAAGAGAAAGTCTGTTCTGACTTAAAACATACTGCCATTAGTTCTCAGTCAAAGCTGTTTGA

CCACCTGACATCCATAGAAGTAGCCTGAATGCACAATAATGCCAAATTTCCTATTCGTGTAACCCAC

>JK714291.1 JutESTSSH3(5A) Jute Plasmid Suppression Subtractive cDNA library Corchorus capsularis cDNA, mRNA sequence

CCTTTCTCCTCGGTCCACAGAGACAAAATGTAGGACTAGGACTGGCGCCAATAGCTAATCACGGAAGAAA

GGACTCACTGAGCCGGGATCACTAACTAATACTAATCTAATACTAATAGAATAGAATACTAATCTAATAC

TAATTCTAATACTAATAGAATAGAAAAAATAGAATAGAAAATACTAATAGAATAGAAAGAAAAGAACTGT

CTTTTCTGTATACTTTCCCCGGTTCCGTTGCTACCGCGGGCTTTACGCAATCGATCGGATCATATAGATA

TCCCTTCAACACAACATAGGTCGTCGAAAGGATCTCGGAGACTCACCAAAGCACGAAAGCCAGGATCTTT

CAGAAAATGGATTCCTATTCGAAGAGTGCATAACCGCATGGATAAGCTCACACTAACCCGTCAATTTGGG

ATCCAATTCGGGATTTTCCTTGGGGGGTATCAGTAAGGAATTGGAATGTAATAATATCGATTCATATGGA

TACAGAGGAAAAGGTTCTCTATTGATTCAAATGCT

>JK714290.1 JutESTSSH3(4A) Jute Plasmid Suppression Subtractive cDNA library Corchorus capsularis cDNA similar to senescence-associated protein, mRNA sequence

GAATACGAACCGTGAAAGCGTGGCCTATCGATCCTTTAGACCTTCGGAATTTGAAGCTAGAGGTGGCAGA

AAAGTTACCACAGGGATAACTGGCTTGTGGCAGCCAAGCGTTCATAGCGACGTTGCTTTTTGATCCTTCG

ATGTCGGCTCTTCCTATCATTGTGAAGCAGAATTCACCAAGTGTTGGATTGTTCACCCACCAATAGGGAA

CGTGAGCTGGGTTAGACCGTCGTGAGACAGGTTAGTTTTACCCTACTGATGGCCGCGTCGCAATAGTAAT

TCAACCTA

>JK714289.1 JutESTSSH3(3) Jute Plasmid Suppression Subtractive cDNA library Corchorus capsularis cDNA similar to Zinc finger protein, mRNA sequence

ATTGAAGTGGAAGAGCCAAGAGATTACAAAGGCGAAGACCATACAGGCCAGCAGGAAATTTAGAAACCGA

TGGCCTTGCCAGAAGTTTCGGGTTTCTGCTGAGTGAATTGCAATAGGAGCTGTTGCTGCAGCTGTGGAAT

CGTTTGCCTCATTCCACTGCTCCGTCATTTCTGTCTCATTTGCAGCAGCAACGTTCCGAGCAGTTGATCC

ACAAATCTCACAAGTTCTGTTACCCTTAATTTTGAACCATGCCTCAGCACAGTGCTTATGGGCAGCAGCC

AAATCATCCTTACAAGAACATCCCAGTTCTATAGGCACACCAGATTCTTGATTGGTTGGGTCCAAGCTGA

GATGGCAAATCCTGCAATCCCTCTCAACTTTGGCCAAATGTAGCTTGGTTTCACTTTGATTTACCCCAG

>JK714288.1 JutESTSSH3(2) Jute Plasmid Suppression Subtractive cDNA library Corchorus capsularis cDNA similar to Zinc finger protein, mRNA sequence

ATTGAAGTGGAAGAGCCAAGAGATTACAAAGGCGAAGACCATACAGGCCAGCAGGAAATTTAGAAACCGA

TGGCCTTGCCAGAAGTTTCGGGTTTCTGCTGAGTGAATTGCAATAGGAGCTGTTGCTGCAGCTGTGGAAT

CGTTTGCCTCATTCCACTGCTCCGTCATTTCTGTCTCATTTGCAGCAGCAACGTTCCGAGCAGTTGATCC

ACAAATCTCACAAGTTCTGTTACCCTTAATTTTGAACCATGCCTCAGCACAGTGCTTATGGGCAGCAGCC

AAATCATCCTTACAAGAACATCCCAGTTCTATAGGCACACCAGATTCTTGATTGGTTGGGTCCAAGCTGA

GATGGCAAATCCTGCAATCCCTCTCAACTTTGGCCAAATGTAGCTTGGTTTCACTTTGATTTACCCCAG

>JK714287.1 JutESTSSH3(1) Jute Plasmid Suppression Subtractive cDNA library Corchorus capsularis cDNA similar to 40S ribosomal protein, mRNA sequence

AATGCTGAGAAAAGGGGCAAAAGACAGGTCATGATTAGGCCTTCCTCGAAAGTGATTATCAAGTTCCTTT

TGGTCATGCAGAAGCATGGATACATTGGAGAGTTTGAATATGTTGATGACCACAGAGCTGGCAAAATTGT

AGTTGAACTAAACGGTAGACTGAACAAGTGTGGCGTGATCAGTCCTCGATTTGATGTTGGAGTGAAGGAG

ATTGAGGGTTGGACTGCTAGGTTACTTCCTTCAAGACAGTTTGGTTATATCGTGCTCACCACATCTGCTG

GCATCATGGACCATGAAGAAGCCAGAAGAAAGAATGTTGGTGGGAAAGTGCTTGGTTTCTTCTATTGAAG

CACATTTCTGAGATTTTCATCTACTTTTCATTTAGAACTTGTTTCTCTATCCTCTGTTG

>HS412001.1 JBM30 Library of differentially expressed transcripts in jute upon stress Corchorus capsularis cDNA, mRNA sequence

GTGCGATTTTACCCTCGCTTCCACGACAACACCGATAATCTTGCAGTTCCCGTTGATAGGAGTCATAGGC

CATGAAGGATTCAGGAAGCTTTTTTTTTTTATAGTGGGACAGAAGTAGTGCTATTAAAATTGGTACAGTA

CCGGAATTAGCATTAAGCATGTGATATATTTAATGGAAGGTATATAGTTCACACACTAATAAATAATTTA

AAAGTCTTAAATGCCCATGGCGAGCTCCGACCAAATGCAGTTAAACAAAAGTAGTCTTGGAAACACAAAT

CAGAGGCATGCGATGCAACTCTCTGGGTTGCTTATTTATAAGCTACATATATATAATAGATACCCTTCCC

>HS412000.1 JBM29 Library of differentially expressed transcripts in jute upon stress Corchorus capsularis cDNA, mRNA sequence

GTTGTAATGGTTATTTCGATTTACCTCGCTTCCACGACACACCGATAATCTTGCAGTTCCCGTTGATAGG

AGTCATAGGCCATGAAGGATTCAGGAAGCTTCGACTGTACCTTGAATGTGGTGAAAGCTATATTAGGGGA

AGGGTATCTATTATATATATGTAGCTTATAAATAAGCAACCCAGAGAGTTGCATCGCATGCCTCTGATTT

GTGTTTCCAAGACTACTTTTGTTTAACTGCATTTGGTCGGAGCTCGCCATGGGCATTTAAGACTTTTAAA

TTATTTAAAAAAAAAAAGCTTCCTTTCAGGTACTTCTGACCGCCATCTATGACCAGTTTCTTGAATGTTG

CTTCGTTCGCGTCAGTCAGTTTGGCTACAACAAGGCTTCCATTCACTGGCTCGCGTCCAGTATCTACTAA

CACCATATGACCTTCAGGGATGCTTTGACCTACAGGTGAGGTCATGGAATCACCTTCAACCTTCAGCCAG

AATCCATTGCCTAATAAGTTAACGTCACTGTCATACCATTCATCAATGTCCTTGATATCGTAGGGTTCAC

AAGCTTCACACCACGAACCAG

>HS411999.1 JBM28 Library of differentially expressed transcripts in jute upon stress Corchorus capsularis cDNA, mRNA sequence

GATCCCGAATTTTACCCTCGCTTCCACGACACACCGATAATCTTGCAGTTCCCGTTGATAGGAGTCATAG

GCCATGAAGGATTCAGGAAGCTTTTTTTTTTTAGTTCACACACTAATAAATAATTTAAAAGTCTTAAATG

CCCATGGCGAGCTCCGACCAAATGCAGTTAAACAAAAGTAGTCTTGGAAACACAAATCAGAGGCATGCGA

TGCAACTCTCTGGGTTGCTTATTTATAAGCTACATATATATAATAGATACCCTTCCCCTAATATAGCTTT

CACCACATTCAAGGTACAGTCGAAGCTTCCTTTCAGGTACTTCTGACCGCCATCTATGACCAGTTTCTTG

AATGTTGCTTCGTTCGCGTCAGTCAGTTTGGCTACAACAAGGCTTCCATTCACTGGCTCGCGTCCAGTAT

CTACTAACACCATATGACCTTCAGGGATGCTTTGACCTACAGGTGAGGTCATGGAATCACCTTCAACCTT

CAGCCAGAATCCATTGCCTAATAAGTTAACGTCACTGTCATACCATTCATCAATGTCCTTGATATCGTAG

GGTTCACAAGCTTC

>HS411998.1 JBM27 Library of differentially expressed transcripts in jute upon stress Corchorus capsularis cDNA, mRNA sequence

CAACCAAACCAATATTTAACCCCCCCTCCCCGACAACACGCGAAAAAACTGCAGTTCCCGTGAAGGAGTC

TAGGCCTAGAAGGATCAGGAAGCTTTTTTTTTTTGGGACAGAAGTAGTGCTATTAAAATTGGTACAGTAC

CGGAATTAGCATTAAGCATGTGATATATTTAATGGAAGGTATATAGTTCACACACTAATAAATAATTTAA

AAGTCTTAAATGCCCATGGCGAGCTCCGACCAAATGCAGTTAAACAAAAGTAGTCTTGGAAACACAAATC

AGAGGCATGCGATGCAACTCTCTGGGTTGCTTATTTATAAGCTACATATATATAATAGATACCCTTCCCC

TAATATAGCTTTCACCACATTCAAGGTACAGTCGAAGCTTCCTTTCAGGTACTTCTGACCGCCATCTATG

ACCAGTTTCTTGAATGTTGCTTCGTTCGCGTCAGTCAGTTTGGCTACAACAAGGCTTCCATTCACTGGCT

CGCGTCCAGTATCTACTAACACCATATGACCTTCAGGGATGCTTTGACCTACAGGTGAGGTCATGGAATC

ACCTTCAACCTTCAGCCAGAATCCATTGCCTAATAAGTTAACGTCACTGTCATACCATTTCATCAATGTC

CTTGATATCCGTAAGGGTTCAACAAAGCT

>HS411997.1 JBM26 Library of differentially expressed transcripts in jute upon stress Corchorus capsularis cDNA, mRNA sequence

TAACAACGATATTTACCCAGTTTCCCGACAACACCGAATTTGCAGTTCCCTTGAAGGAGTCTAGGCCTGA

AGGTTCAGGAAGCTTCGACTGTACCTGATGTGGTGAAAGCTATATTAGGGGAAGGGTATCTATTATATAT

ATGTAGCTTATAAATAAGCAACCCAGAGAGTTGCATCGCATGCCTCTGATTTGTGTTTCCAAGACTACTT

TTGTTTAACTGCATTTGGTCGGAGCTCGCCATGGGCATTTAAGACTTTTAAATTACTTATTAGTGTGTGA

ACTATATACCTTCCATTAAATATATCACATGCTTAATGCTAATTCCGGTACTGTACCAAAAAAAAAAAGC

TTCCTTTTAGGGACTTTTGAACGCCCTCTATGACCAGTTTCTGATGTGCTTCTTCGCTCATCACTTGGCT

TTGCAAGGCTTCCTTGCTTGGCTTGCGTCGGGTACCCACTACCACCTCTGACATTAAGGGATGCATTCGT

CCTAGGATGCTTTCAAGGAACCAGTGAGGTCATGGCCACATCTTTTGCCTTACAGCCAAAATCCTTGCCA

AATAAATTAACGTCCTCTGTACATACCGTCATAAATGTCCCTTGATAAACGTAAGGGTTCCACAAGCTTC

ACAGAAACGAAACCATTCT

>HS411996.1 JBM25 Library of differentially expressed transcripts in jute upon stress Corchorus capsularis cDNA, mRNA sequence

GCATACGAGTTTTACCCTCGCTTCCACGACAACACCGATAATCTTGCAGTTCCCGTTGATAGGAGTCATA

GGCCATGAAGGATTCAGGAAGCTTGATTGCCAAAGCAGATAAGATTGCCAAGGAATCTGCAGCATAATGA

AATATTTACATAACCTTGTCTCCTTCTGGATAATTGCGAAAATAATCTAGGTGATGTTGGCAATGCCAGC

AACAGCAGGATGTTCATTTTTTGAGATGAAATGTACCAGGATTTTTCAGTTTATGTTTTTGATGGTATGT

TACTCCTGAGTGTGAAAGAGGAATAGAACTAGGGGGCCTGTTTTGCCAAGCCCTCTTTTTTTATTGTTAT

TTAATAAGGGAAATACAAGGCGGACGTGTACATTTTTCCTTCCAAAAAAAAAAAGCTTCCTTTCAGGTAC

TTCTGACCGCCATCTATGACCAGTTTCTTGAATGTTGCTTCGTTCGCGTCAGTCAGTTTGGCTACAACAA

GGCTTCCATTCACTGGCTCGCGTCCAGTATCTACTAACACCATATGACCTTCAGGGATGCTTTGACCTAC

AGGTGAGGTCATGGA

>HS411995.1 JBM23 Library of differentially expressed transcripts in jute upon stress Corchorus capsularis cDNA, mRNA sequence

AACAAAATTTTTACCCTTCGCTTCCCGACAACACCGATAATCTTGCAGTTCCCGTTGATAGGAGTCATAG

GCCACGAAAAGATTCAGGAAGCTTTTTTTTTTTGCAAAATGGGGTGCTATGCAATTACTGCATACTAATC

AATCAAACAAAGGAAGGAGACAGGAGCTACATATGAATAGATGAGAATGATTAGATACATTATTTTTAGC

TTACTTGTGAGGTTCCTCAAAAATAATGAATGTAGTACCACTTTATGCATTTTATCATGCGTGCCAAGTG

CTTCTGCAATCCCTTCACCCTACAGCAAAGTAAATTCGGATTGCCCAGTGATTTTCTTGTGGTCCAGCGG

TAAAAGCTTCCTTTCAGGTACTTCTGACCGCCATCTATGACCAGTTTCTTGAATGTTGCTTCGTTCGCGT

CAGTCAGTTTGGCTACAACAAGGCTTCCATTCACTGGCTCGCGTCCAGTATCTACTAACACCATATGACC

TTCAGGGATGCTTTGACCTACAGGTGAGGTCATGGAATCACCTTCAACCTTCAGCCCAGAATCCATTGCC

TAAATAAGTTAACGTCACTGGTC

>HS411994.1 JBM22 Library of differentially expressed transcripts in jute upon stress Corchorus capsularis cDNA, mRNA sequence

AAATTCCATCTAAATTTTTCCCTTCGCTTCCACGACAAACCGATAATCTTGCAGTTCCCGTTGATAGGAG

TCATACGCCACGAAAAGATTCAGGAAGCTTTTACCGCTATATTGGTTTTACTCCGTCGAAGTATGATTGC

TTACTGCTTCAAATTCAGAGATGAGGGGAGGGATAGCGTATCTCTTATACTAGAAGAAAACTACCCCTTC

GAAAGCTGCATCGCCACCCTATCGCCCTTAGGAAAGGGGAAATAACTAATAGCTCGAACTCAAAGAAAGG

CTCTTAGACTGGCTTAGCTGCAGTCGCTTCCCAAAAAAAAAAAGCTTCCTTTCAGGTACTTCTGACCGCC

ATCTATGACCAGTTTCTTGAATGTTGCTTCGTTCGCGTCAGTCAGTTTGGCTACAACAAGGCTTCCATTC

ACTGGCTCGCGTCCAGTATCTACTAACACCATATGACCTTCAGGGATGCTTTGACCTACAGGTGAGGTCA

TGGAATCACCTTCAACCTTCAGCCAGAATCCATTGCCTAATAAGTTAACGTCACTGTCATACCATTCATC

AATGTCCTTTGATATCGTAAGGTTCACAA

>HS411993.1 JBM21 Library of differentially expressed transcripts in jute upon stress Corchorus capsularis cDNA, mRNA sequence

TGACGGGAGAAAGGAAGAGCCAAATAGTTGAGTTTAGGAAAATCTCAGGAAAAAAGGAAACCAAGAGATT

TCTTACCTCGGAACTGATATATTCTCTTCACCCGATACTGCGTTCGTGATGGATCATAGCCAGAGGATTC

GGAAGCTTTTACCGCTATATTGGTTTTACTCCGTCGAAGTATGATTGCTTACTGCTTCAAATTCAGAGAT

GAGGGGAGGGATAGCGTATCTCTTATACTAGAAGAAAACTACCCCTTCGAAAGCTGCATCGCCACCCTAT

CGCCCTTAGGAAAGGGGAAATAACTAATAGCTCGAACTCAAAGAAAGGCTCTTAGACTGGCTTAGCTGCA

GTCGCTTCCCAAAAAAAAAAAGCTACCTTTCAGGTACTTCTGACCGCCATCTATGACCAGTTTCTTGAAT

GTTGCTTCGTTCGCGTCAGTCAGTTTGGCTACAACAAGGCTTCCATTCACTGGCTCGCGTCCAGTATCTA

CTAACACCATATGACCTTCAGGGATGCTTTGACCTACAGGTGAGGTCATGGAATCACCTTCAACCTTCAG

CCAGAATCCATTGCCTAATAAGTTAACGTCACTGTCATACCATTCATCAATGTCCTTGATATCGTAGGGG

TTCATAAA

>HS411992.1 JBM20 Library of differentially expressed transcripts in jute upon stress Corchorus capsularis cDNA, mRNA sequence

GTTATAAGAGAGGTTCGATTTCCCTCGCTTCCACGACACACCGATAATCTTGCAGTTCCCGTTGATAGGA

GTCATAGGCCATGAAGGATTCAGGAAGCTTGATTGCCCGGAACTGAGGGCCAAGATGTTGCAGATGTTCT

GTTTGGTGACTATGGTTTTACAGGCAAGCTTCCTCGTACTTGGTTTAAGACTGTTGATCAATTACCGATG

AATGTTGGGGATCCACATTATGATCCTCTTTTCCCATTTGGATTTGGCCTTACCACTGAACCTACCGAAA

GATATGAGGATACTTTTCCTATGGAAGTGTAGTTTTCTTTGAAGTCATTTACATCTGTCATTTCCATTTC

TTAGAAGTACCAATTCTTGATAGCATGATATTTATTTACAAGAAATGTAATAAAAGTTCATTGTTACATA

CAATTGTAGTATGACAATACACAGAAAAAATAAATAGAATACTTACCAAAAAAAAAAAGCTTCCTTTCAG

GTACTTCTGACCGCCATCTATGACCAGTTTCTTGAATGTTGCTTCGTTCGCGTCAGTCAGTTTGGCTACA

ACAAGGCTTCCATTCAC

>HS411991.1 JBM19 Library of differentially expressed transcripts in jute upon stress Corchorus capsularis cDNA, mRNA sequence

AGGTGTTTATCGACGATTTCGATTTTACCTCGCTTCCACGACACACCGATAATCTTGCAGTTCCCGTTGA

TAGGAGTCATAGGCCATGAAGGATTCAGGAAGCTTGATTGCCCTTACCCCTGCAACCCAAGTTGCCCAAA

AGTTGATGATTCAGACTGAGTCTCAACTCCCAGACATATAGAAACACTAAATTTTGTGTTATGCGACTCC

CTTTAATTTGTAGTAAATAAAAATGAATGCATTTCTGATTTCTTTTAGAAATAAGTTGTTTGGACGGACT

AGAAGATTACTTCACTCTGTAACTAGTACTGTAGTATTTTACTATACTACATTATTTATTATATTCATGT

AATGTAACTACACAACCACAGTGAAGAAATAATAAATTGAACAAGGGTTGGGTTCGCAGTTTTACTATTA

TCAAAAAAAAAAAGCTTCCTTTCAGGTACTTCTGACCGCCATCTATGACCAGTTTCTTGAATGTTGCTTC

GTTCGCGTCAGTCAGTTTGGCTACAACAAGGCTTCCATTCACTGGCTCGCGTCCAGTATCTACTAACACC

ATATGACCTTCAGGGATGCTTTGA

>HS411990.1 JBM17 Library of differentially expressed transcripts in jute upon stress Corchorus capsularis cDNA, mRNA sequence

GAAATCGACTTTTACCCTCGCTTCCACGACAACACCGATAATCTTGCAGTTCCCGTTGATAGGAGTCATA

GGCCATGAAGGATTCAGGAAGCTTGATTGCCTAGCTAGTGAAATTGCTCCAAGTCTAAAGCAATATTCAA

AGATAATTTCGTCTCTTGGGTATTTCTTCTTGTCTTCATTTCACTATTTAAGTGACTGGGTTACTGTTTA

GCTGTCAAAAAAAAAAAGCTTCCTTTCAGGTACTTCTGACCGCCATCTATGACCAGTTTCTTGAATGTTG

CTTCGTTCGCGTCAGTCAGTTTGGCTACAACAAGGCTTCCATTCACTGGCTCGCGTCCAGTATCTACTAA

CACCATATGACCTTCAGGGATGCTTTGACCTACAGGTGAGGTCATGGAATCACCTTCAACCTTCAGCCAG

AATCCATTGCCTAATAAGTTAACGTCACTGTCATACCATTCATCAATGTCCTTGATATCGTAGGGTTCAC

AAGCTTCACACCACGAACCAGCTCTAACCATGCTAATCAATGGATATTTCCCTTTGGGCTCAACGTGCCC

AACAAATCTAACAT

>HS411989.1 JBM16 Library of differentially expressed transcripts in jute upon stress Corchorus capsularis cDNA, mRNA sequence

ATTATTTAACGTGAGCTCGATTTTACCTCGCTTCCCGACAACACCGATAATCTTGCAGTTCCCGTTGATA

GGAGTCATAGGCCATGAAGGATTCAGGAAGCTTGATTGCCCCATGTTCCGGGGGATGTTTTACTCAATCA

TGGGAGGATAATTTTTATAACCCTGTATGTTTCGTTCACATCGCGACTCTGTATTTCTAGCTTTTGTTAA

CTAACCTGTACTTCTCTTCTGATTTTTTGTTTTAGTATTTTTATAGTTTCGGATCAAAAAAAAAAAGCTT

CCTTTCAGGTACTTCTGACCGCCATCTATGACCAGTTTCTTGAATGTTGCTTCGTTCGCGTCAGTCAGTT

TGGCTACAACAAGGCTTCCATTCACTGGCTCGCGTCCAGTATCTACTAACACCATATGACCTTCAGGGAT

GCTTTGACCTACAGGTGAGGTCATGGAATCACCTTCAACCTTCAGCCAGAATCCATTGCCTAATAAGTTA

ACGTCACTGTCATACCATTCATCAATGTCCTTGATATCGTAGGGTTCACAAGCTTCACACCACGAACCAG

CTCTAACCATGCTAATCAA

>HS411988.1 JBM15 Library of differentially expressed transcripts in jute upon stress Corchorus capsularis cDNA, mRNA sequence

GATATCGGAGTTTTAACCTCGCTTCCACGACAACACCGATAATCTTGCAGTTCCCGTTGATAGGAGTCAT

AGGCCATGAAGGATTCAGGAAGCTTTTTTTTTTTAGTACAAACAATACCACATTACTATTCATAGACACA

CGCCACAGTGGAAAATGGAATAAGAATTTACAGGCTATTCATGCCTAATTATACAGCAAATTAAACTGAT

CCAATCATACAGGCAATCAAGCTTCCTTTCAGGTACTTCTGACCGCCATCTATGACCAGTTTCTTGAATG

TTGCTTCGTTCGCGTCAGTCAGTTTGGCTACCACAAGGGTTCCATTCACTGGCTCGCGTCCAGTATCTAC

TAACACCATATGACCTTCAGGGATGCTTTGACCTACAGGTGAGGTCATGGAATCACCTTCAACCTTCAGC

CACAATCCATTGCCTAATAACTTAACGTCACTGTCATACCATTCATCAATGTCCTTGATATCCTACGGTT

CACAAGCTTCACACCACGAACCAGCTCTAACCATGCTAATCAATGGATATTTCCCTTTGGGCTCAACGTG

CCCAACAAATCTAACATTCGA

>HS411987.1 JBM14 Library of differentially expressed transcripts in jute upon stress Corchorus capsularis cDNA, mRNA sequence

AAAGGGAAGGACATACTCCTTTCAGAAAAAGAAACAGGAAGCTTTTACCGCTCCAAAATGAGGCCTTTTA

TATATGTTGAAGAAAAGCCCATGCGGGGATTGCCTTGTTCATTGCTTCTGCGAGTATTGTGCCCTTTGCC

AAGAATACCGTGAGCTCAAATCTCGTGGCTACGACCTTTCCATCGGCTGGCATGGCAATATGGAGAAGCA

GAACCGGGAAGTCGCCATGACTTCCATCCCACCATCCGTCGAACAAGGCATGACCAGATGATCCAAATGT

CGCCGCTATATTATATATTAAATGTTGAATTCTTAGTTTCAGTGATTAAGATATATCTATGAACATAAAG

ATGCCACTTCGTGTTATTAGTCATGTAATTATTGTTTGTCTCTGTATGGATTGTTCTTTGTTTCCAAAAA

AAAAAAGCTTCCTTTCAGGTACTTCTGACCGCCATCTATGACCAGTTTCTTGAATGTTGCTTCGTTCGCG

TCAGTCAGTTTTGGCTAAAACAAGGCTTCCATTCACTGG

>HS411986.1 JBM13 Library of differentially expressed transcripts in jute upon stress Corchorus capsularis cDNA, mRNA sequence

TAAGGATCGTCTCATCAAACGTACGAGCGAAGAATCAACTTCAGAAATCGGTCTAGTGGCGGTCAGAAGT

TACTGAAAGGAAGCTTTTTTTTTTTGGGAAGCGACTGCAGCTAAGCCAGTCTAAGAGCCTTTTTTGAGTT

CGAGCTATTAGTTATTTCCCCTTTCTAAGGGCGATAGGGTGGCGATGCAGCTTTCGAAGGGGTAGTTTTC

TTCTAGTATAAGAGATACGCTATCCCTCCCCTCTCTCTGATTTGAAGCAGTAAGCAATCATACTTCGACG

GAGTAAAACCAATATAGCGGTAAAAGCTTCCTGAATCCTTCATGGCCTATGATTTCCTTTCAACGGGAAC

TGCAAGATTATCGGTGTTGTCGTGGAAGCGAGGGTAAAATTAAGAATAGATACAAATCATGAAAACAAAA

GGGCCTCATAAAACAACCTATTTTTAAAGGTTAAAAGTCAAAAAAAAATAAAAGATTAAAAAG

>HS411985.1 JBM12 Library of differentially expressed transcripts in jute upon stress Corchorus capsularis cDNA, mRNA sequence

GGGAAAACGCAAGAATTCGATTTTCCCTCGCTTCCCGACACACCGATAATCTTGCAGTTCCCGTTGATAG

GAGTCATAGGCCATGAAGGATTCAGGAAGCTTGTTTGTCTTCTATAAAACTTTGAGTATGTTTGATTATC

AATGAAATCTTTTGCTAATGCTATTTCTGATGTACTCTTTTGACGAATTCTATCTTTGATTTCACTATGA

TTCCATATAATTGCAACCTGATGAATTTTCTTGTCCAAAAAAAAAAAGCTTCCTTTCAGTACTTCTGACC

GCCATCTATGACCAGTTTCTTGAATGTTGCTTCTTTCGCGTCAGTCATTTTGGCTACAACAAGGCTTCCA

TTCACTGGCTCGCGTCCAGTATCTACTAACACCATATGACCTTCAGGGATGCTTTGACCTACAGGTGAGG

TCATGGAATCACCTTCAACCTTCAGCCAGAATCCATTGCCTAATAAGTTAACGTCACTGTCATACCATTC

ATCAATGTCCTTGATATCGTAGGGTTCACAAGCTTCACACCACGAACCAGCTCTAACCATGCTAATCAAT

GGATATTTCCCTTTGGGC

>HS411984.1 JBM11 Library of differentially expressed transcripts in jute upon stress Corchorus capsularis cDNA, mRNA sequence

GGGAGGAAATCGTGGGCACAACGGACGCAGGCGAACGAATCAACTTCAAGAAATCGGTCTAGTGGCGGTC

AGAAGTACCTGAAAGGAAGCTTCGACTGTGGTCAACCCGAGTGAATGCTCATCATTATTGCCTTGCTTGC

AAACGTTAATATTCCTCAGGTGCAAGTGGTCTTGTACATCTTTCTAACAATGATTTTCATTTCTTAATTA

TTATATTAGGAAAGATGGATTTTGTCTTTGAAAGTACTTTGAATATTAGAAGAAGAGTGGACACCATGAT

TTTAGAAGAAATTATGGTATGTTCATTTTAGTTTATATTCAATTACTGATTGAAGTTTGGTAAATGACAT

AATTACATTCAGAATCCTAGTAGAATGGTTTAACTTTTTATTGCCAAAAAAAAAAAGCTTCAAGGATCGG

GCCTTGATGTTACCCGAGAGCTTGGCACCCAGCCTGCGCGAGCAGGGGAATTGATCCGGTGGATGACCTT

TTGAATGACCTTTAATAGATTATATTACTAATTAATTGGGGACCCTAGAGGTCCCCTTTTTTATTTTAAA

AATTTTTTCACAAAACGGTTTACAAGCATAAAGCTTGCTCAATCAATCACCGGGTCCCCCACTCACCAGT

>HS411983.1 JBM10 Library of differentially expressed transcripts in jute upon stress Corchorus capsularis cDNA, mRNA sequence

TAAGGATCGTCTCATCAAACGTACGAGCGAAGAATCAACTTCAGAAATCGGTCTAGTGGCGGTCAGAAGT

TACTGAAAGGAAGCTTTTTTTTTTTGGGAAGCGACTGCAGCTAAGCCAGTCTAAGAGCCTTTTTTGAGTT

CGAGCTATTAGTTATTTCCCCTTTCTAAGGGCGATAGGGTGGCGATGCAGCTTTCGAAGGGGTAGTTTTC

TTCTAGTATAAGAGATACGCTATCCCTCCCCTCTCTCTGATTTGAAGCAGTAAGCAATCATACTTCGACG

GAGTAAAACCAATATAGCGGTAAAAGCTTCCTGAATCCTTCATGGCCTATGATTTCCTTTCAACGGGAAC

TGCAAGATTATCGGTGTTGTCGTGGAAGCGAGGGTAAAATTAAGAATAGATACAAATCATGAAAACAAAA

GGGCCTCATAAAACAACCTATTTTTAAAGGTTAAAAGTCAAAAAAAAATAAAAGATTAAAAAG

>HS411982.1 JBM9 Library of differentially expressed transcripts in jute upon stress Corchorus capsularis cDNA, mRNA sequence

GCATCGAAATTTTACCCTCGCTTCCACGACACACCGATAATCTTGCAGTTCCCGTTGATAGGAGTCATAG

GCCATGAAGGATTCAGGAAGCTTCGACTGTTAACTGCCCACGATGATTCCTGTAAATTTCATGAGATATT

AATTTGAACGACTTTAGAAATGGTACAGAAGCTACATAATGATGAATAAATAGATTTTATAGTTTTCTAA

AAAAAAAAAGCTTCCTTTCAGGTACTTCTGACCGCCATCTATGACCAGTTTCTTGAATGTTGCTTCGTTC

GCGTCAGTCAGTTTGGCTACAACAAGGCTTCCATTCACTGGCTCGCGTCCAGTATCTACTAACACCATAT

GACCTTCAGGGATGCTTTGACCTACAGGTGAGGTCATGGAATCACCTTCAACCTTCAGCCAGAATCCATT

GCCTAATAAGTTAACGTCACTGTCATACCATTCATCAATGTCCTTGATATCGTAGGGTTCACAAGCTTCA

CACCACGAACCAGCTCTAACCATGCTAATCAATGGATATTTCCCTTTGGGCTCAACGTGCCCAACAAATC

TAACATTCGAATCAG

>HS411981.1 JBM8 Library of differentially expressed transcripts in jute upon stress Corchorus capsularis cDNA, mRNA sequence

ATAATCACGATATTTACCCAGTTCCCCGACAACACCGTTATCTGCAGTCTCCCCTTGAAGGAGTCTAGGC

CTGAAGGATCAGGCTTCAATATATTACTACCTGGCCTTCAGGTACTTCTGACCGCCTCTATGACCAGTTT

CTGATGTTGCTTCTTTCGCGCGTCAGTCAGTTTGGCTACAACAAGGCTTCCATTCACTGGCTCGCGTCCA

GTATCTACTAACACCATATGACCTTCAGGGATGCTTTGACCTACAGGTGAGGTCATGGAATCACCTTCAA

CCTTCAGCCAGAATCCATTGCCTAATAAGTTAACGTCACTGTCATACCATTCATCAATGTCCTTGATATC

GTAGGGTTCACAAGCTTCACACCACGAACCAGCTCTAACCATGCTAATCAATGGATATTTCCCTTTGGGC

TCAACGTGCCCAACAAATCTAACATTCGAATCAGAGGTGCCATTGAGCAGCCAGTCAACACTTACGCCAA

GAGCTGACGCAAGTTCTGGTAAAAAGCGTGGTCGCTTAGTTTTACCGTTTTCGAGCTGCTCTATAGACTG

CTGGGTAGTCCCCACCTTTTGAGCAAGTTCAGCCTGGTTAAGTCCAAGCTGAATTCTTTTTTGCTTTTTT

TACCCCTGGAAAGAAT

>HS411980.1 JBM7 Library of differentially expressed transcripts in jute upon stress Corchorus capsularis cDNA, mRNA sequence

ATATCACGGTATTTACCCCGTTCCCCGACAACACCGAATCTGCAGTTCCCTTGAAGGAGTCTAGGCCTGA

AGGTTCAGGAAGCTTTTACCGCTGTGGACTCTAGTTAAGCACTGCATGTTGACTCTTTACTTTTTTTGTT

TTTTCGTTTTTGTTTTGTTTCTGTCGTGTTTCAAAGTACTGTCAGGGCACACTCGGATAATCCAATGTAT

TCTTCTTATTAAGTATCCTGGTGCATCGCAATATTTGGTTTTCCAAAAAAAAAAGCTTCCTTTCAGGTAC

TTCTGACCGCCATCTATGACCAGTTTCTTGAATGTTGCTTCGTTCGCGTCAGTCAGTTTGGCTACAACAA

GGCTTCCATTCACTGGCTCGCGTCCAGTATCTACTAACACCATATGACCTTCAGGGATGCTTTGACCTAC

AGGTGAGGTCATGGAATCACCTTCAACCTTCAGCCAGAATCCATTGCCTAATAAGTTAACGTCACTGTCA

TACCATTCATCAATGTCCTTGATATCGTAGGGTTCACAAGCTTCACACCACGAACCAGCTCTAACCATGC

TAATCAAATGGATATTTCCCTTTGGGGCTCAACGTGCCCAACAAATCTAAACATTCGAATCCAGAAGGTG

GC

>HS411979.1 JBM6 Library of differentially expressed transcripts in jute upon stress Corchorus capsularis cDNA, mRNA sequence

ATAAACACGATATTTACCCAGTTCCCCGACAACACCGAATCTGCAGTTCCCTTGAAGGAGTCTAGGCCTG

AAGGTTCAGGAAGCTTTTTTTTTTTTAAAGAAAACTGGGTTTTTGAGCCTCAAATATACACACATATGAT

TTAATTATCCATCACCGGAGAAAGTAATGGTTTATTTGAAAAGAAAGCTTCCAAGTTGCCCACCGCCAGT

TCAGAGGCAGCAATGAAGGATTCCGGTGTAAAAAACAACTCTATGCGGTAAAAGCTTCCTTTCAGGTACT

TCTGACCGCCATCTATGACCAGTTTCTTGAATGTTGCTTCGTTCGCGTCAGTCAGTTTGGCTACAACAAG

GCTTCCATTCACTGGCTCGCGTCCAGTATCTACTAACACCATATGACCTTCAGGGATGCTTTGACCTACA

GGTGAGGTCATGGAATCACCTTCAACCTTCAGCCAGAATCCATTGCCTAATAAGTTAACGTCACTGTCAT

ACCATTCATCAATGTCCTTGATATCGTAGGGTTCACAAGCTTCACACCACGAACCAGCTCTAAACCATGC

TAATCAATGGATATTTCCCTTTGGGCTCAAACGTGCCCAACAAATACTAAAACATTCGAAATCAGAGGTG

C

>HS411978.1 JBM5 Library of differentially expressed transcripts in jute upon stress Corchorus capsularis cDNA, mRNA sequence

AAAAACAAATTAACTATAAAAATTTTTTTTTGATAAAAAAAAAAAAAAAGTTGGGGCACTTGCATAGGTA

TTACCGAAGCGCTGTCATTGGCCGAGAGTGATTTTTCCCCCCCCCCCCC

>HS411977.1 JBM4 Library of differentially expressed transcripts in jute upon stress Corchorus capsularis cDNA, mRNA sequence

GAAAGACCGTTCGCAAACGTACGAGCGAAGAATCAACTTCAGAAATCGGTCTAGTGGCGGTCAGAAGTTA

CTGAAAGGAAGCTTTTTTTTTTTAGGTAAAGCGTCTTTCTTTTAGCTCTCGAGTTTGCTTTTTTGTGGTA

TTGCAGAGTCTACTCGAAAAATAGGGATAACTTTCCATCCCATATAGCGGTAAAAGCTTCCTGAATCCTT

CATGGCCTATGACTCCTATCAACGGGAACTGCAAGATTATCGGTGTTGTCGTGGAAGCGAGGGTAAAATT

CGTATGATCAATTCTTGAAGACGAAAGGGCCTCGTGATACGCCTATTTTTATAGGTTAATGTCATGATAA

TAATGGTTTCTTAGACGTCAGGTGGCACTTTTCGGGGAAATGTGCGCGGAACCCCTATTTGTTTATTTTT

CTAAATACATTCAAATATGTATCCGCTCATGAGACAATAACCCTGATAAATGCTTCAATAATGGGGATCC

GGTGATTGATTGAGCAAGCTTTATGCTTGTAAACCGTTTTGTGAAAAAAATTTTTAAAATAAAAAAGGGG

ACCTCTAGGGTCCCCCCAATTAATTAGTAAATATAAATCTATTAAAGGTCATTTCAAAAAGGGT

>HS411976.1 JBM3 Library of differentially expressed transcripts in jute upon stress Corchorus capsularis cDNA, mRNA sequence

CCAGATATATCAACCTAATATAAGTTGCTATGCGAGCATCCTTTTAAGTTTCATCGGTATCATTACCCCC

ATGAACAGAAATCCCCCTTACACGGAGGCATCAGTGACCAAACAGGAAAAAA

>HS411975.1 JBM2 Library of differentially expressed transcripts in jute upon stress Corchorus capsularis cDNA, mRNA sequence

AAAAAACCGCCCTTAACATGGCCCGCTTTATCAGAAGCCAGACATTAACGCTTCTGGAGAAACTCAACGA

GCTGGACGCGGATGAACAGGCAGACATCTGTGAATCGCTTCACGACCACGCTGATGAGCTTACCGCAGCT

GCATAATAATCCGCCATGA

>HS411974.1 JBM1 Library of differentially expressed transcripts in jute upon stress Corchorus capsularis cDNA, mRNA sequence

GACTCGTCGCGCACAACGTACGAGCGAAGAATCAACTTCAAGAAATGGTCTAGTGGCGGTCAGAAGTTAC

TGAAAGGAAGCTTTTTTTTTTTATCATAACACTTTTACTGTTCCATAGCATTAGCATAGCAACTAATAAA

TGTCAAAATTTCAATTATCTTCCACAAATAAAGCAGTTTTGGGCGGTAAAAGCTTCCTGAATCCTTCATG

GCCTATGACTCCTATCAACGGGAACTGCAAGATTATCGGTGTTGTCGTGGAAGCGAGGGTAAAATTCGTA

TGATCAATTCTTGAAGACGAAAGGGCCTCGTGATACGCCTATTTTTATAGGTTAATGTCATGATAATAAT

GGTTTCTTAGACGTCAGGTGGCACTTTTCGGGGAAATGTGCGCGGAACCCCTATTTAGTTTAATTTTATA

AAATACATTCAAAAATGTAAACCGCTACATAGAAACAATAACACTGATAAATAGCTTCAATAAATGGGGA

TACGGTGAATAAAATAGAGCAAACAAATAGCATAACAAAAAAAAAAAAAAAAAAAAAAATTTTTAAATAA

AA

>GR463793.1 JuteSSH6H5S178 JuteSSHlibMP2 Corchorus capsularis cDNA clone JuteSSH6HXERICO2 similar to XERICO; protein binding / zinc ion binding [Arabidopsis thaliana] RING zinc finger protein like protein, mRNA sequence

CGAGCGGCCGCCCGGGCAGGTACCGTAGGAAGACAAATCTACACCATGTTTGGGATACGAATATTATTGA

TTCTGCAGTGAAGACATTTTATGGCTCAGATCTCGCAATAATGATTCAAGCTATCCAGAGGAATATTACA

GATTCTTGGTCCAATAACATATCATCGTGGGAATCTTGTGGAAATAATCGCACAGTTTGTCCTAACCTGT

ATGCTTCTGAAAGTGTTAGGTTGGCATGTTTGTATGCATACAGGAATGCCACGCCTGGAAGCACCTTAGA

AGATGAATATTTCCTCTCTCGGTTGCCCATTGTGGAGAAGAGACTTGCTCAAGGTGGGATTCGCCTTGCC

GCCGTACCTCGGCCGCGACCACGC

>GR463792.1 JuteSSH6H5S161 JuteSSHlibMP2 Corchorus capsularis cDNA clone JuteSSH6HXERICO similar to XERICO; protein binding / zinc ion binding [Arabidopsis thaliana] RING zinc finger protein like protein, mRNA sequence

TCGAGTTTTTCAGCAAGATTCGAGCGGCCGCCCGGGCAGGACACATAGCACTCCTTCTGATGGAGCTGGG

AGACTTGAGAGACCCATTTTTTCTGACTTTTACGTGGGATACAGAAAGCAAAAGAGATGTGCAATTTGAG

ATTAAGCTAAAGTTCAGATATGAATAAGGACAGAACGAAGCTACTCTCTTGTTGACCTATATAACCTTAA

AAGCAGAGGGGGAAAACCTTCACATGTCAATTGATGATTGCTGCTAAAAACCCTGGTAATACAAAAATGG

TCAACTTTGCTCACCTCGGCCGCGACCACGC

>GR463791.1 JuteSSH6H5S197 JuteSSHlibMP2 Corchorus capsularis cDNA clone JuteSSH6HAPUM5 similar to APUM5 (arabidopsis pumilio 5) like protein, mRNA sequence

CGAGCGGCCGCCCGGGCAGGTTCGTCATCTAATTAGTGACGCGCATGAATGGATTAACGAGATTCCCACT

GTCCCTGTCTACTATCCAGCGAAACCACAGCCAAGGGAACGGGCTTGGCAGAATCAGCGGGGAAAGAAGA

CCCTGTTGAGCTTGACTCTAGTCCGACTTTGTGAAATGACTTGAGAGGTGTAGTATAAGTGGGAGCTTCG

GCAACGGTGAAATACCACTACTTTTAACGTTATTTTACTTATTCCGTGAATCGGAAGCGGGGCTACGCCC

CTCTTTTTAGACCCAAGACCGGCATCTGTCGGTCGATCCGGGCGGAAGACATTGTCAGGTGGGGAGTTTG

GCTGGGGCGGCACATCTGTTAAAAGATAACGCAGGTGTCCTAAGATGAGCTCAACGAGAACAGAAATCTC

GTGTGGAACAAAAGGGTAAAAGCTCGTTTGATTCTGATTTCCAGTACCTCGGCCGCGACCACGC

>GR463790.1 JuteSSH6H5S167 JuteSSHlibMP2 Corchorus capsularis cDNA clone JuteSSH6HDEGP1A similar to DEGP1 (DEGP protease 1) like protein, mRNA sequence

TTTTTCAGCAAGATTCGAGCGGCCGCCCGGGCAGGTACAAATGAGTCAACATAAATATGCATCAAATGTT

GTTGAGAAGTGCTTGGAGTATGGTGATAGTGCTGAAAAGGAGCTTTTGGTTGAGGAGATAATTGGACAAT

CTGCTGAAAATGACACTTTATTGACTATGATGAAGGACCAATTTGCAAATTATGTGGTCCAGAAGGTTCT

TGAAATTTGCAATGATAGACAGCGGGAAGTATTAATGGATCGTGTAAGAGTTCATCTTAATGCTTTGAAG

AAGTACCTCGGCCGCGACCACGC

>GR463789.1 JuteSSH6H5S162 JuteSSHlibMP2 Corchorus capsularis cDNA clone JuteSSH6HDEGP1 similar to DEGP1 (DEGP protease 1) like protein, mRNA sequence

GCGTGGTCGCGGCCGAGGTACGGGAAGGTGACAAGACCTATTTTAGGGATTAAATTTGCTCCAGATCAAT

CTGTTGAGCAGTTGGGAGTAAGTGGTGTCCTTGTCTTAGATGCTCCTGCAAATGGTCCGGCTGGCAAGGC

GGGTCTGCTACCAACAAAGCGTGATGCCTATGGTAGACTAATTTTGGGTGATATAATAACATCTGTTAAT

GGAAAGAAGGTCACTAGTGGAAGTGACTTGTACCTGCCCGGGCGGCCGCTCG

>GR463788.1 JuteSSH6H5S154 JuteSSHlibMP2 Corchorus capsularis cDNA clone JuteSSH6HACCOX1 similar to ACC oxidase like protein, mRNA sequence

GCGTGGTCGCGGCCGAGGTGGGAAAAAGAGGATGCAAGATAATGCAGTTTTCAAACCTGAGTTCTACGTA

TTTATTTATCTTTGTGCTTGGAAAAGTTAGAAGGATAGATAGATTATCAGAAATAGGTAATTGTCTCGCC

TTGCTTTTCAGGGCCTTCATCGACTTGGGAAGCCTGATATTGCTGTCTATGATGGTTCTTGGACAGAATG

GGGTGCGGAATCTGACACACCCGTCAATACATCATCATCATAAATCTTAATGCAACCCTGTCAGTTCTTA

GCGATACAAATTGAGATCGGCGGGTCTAGTACCTGCCCGGGCGGCCGAGCGTGGTCGCGGCCGAGGTACT

TCTATACTATTAGATAGCATGCCATTGCCACTGCCCTTTTTTTTCTCATCATCACAAGTCAAAGCTCTAG

AACAAACTTTAAACTGTTGCAACTGGGACAGTTGTTTCCACGGCTTTCATGGCTTCAAATCTTGGTTCCT

TGGCCTGGAATTTCAGAGCTGCATACAGCTTCATGTAATCTTCAAACACAAATCTGGGGTAAATTTGCTT

CTTCTACCTGCCCGGGCGGCCGCTCGAATCTTTCTAGAAGATCTCCTACAATATTCTCAGCTGCCATGGG

AAAATCGATGTTCTTCTTTTATTCTCTCAAGATTTTCAGGCTGTATATTAAAACTTATATTAAGAACTAT

GCTAACCACCTCATCAGGAACCGTTGTAGGTGGCGTGGGTTTTCTTGGCAATC

>GR463787.1 JuteSSH6H5S245 JuteSSHlibMP2 Corchorus capsularis cDNA clone JuteSSH6RPL1 similar to 60S acidic ribosomallarge subunit 9 protein like protein, mRNA sequence

GCGTGGTCGCGGCCGAGGTACGCTCACTTTCCCATCAACGCCTCCATCACCAACGGCAACAAGTCCATTG

AGATCCGTAACTTCCTTGGCGAGAAGAAGGTGCGTAAGGTTGATATGTTGGAGGGAGTTTCAATTGTTCG

ATCTGAGAAGGTTAAGGATGAGATTGTTTTGGATGGTAATGATATCGAACTTGTCTCTCGCTCAGCTGCC

TTGATAAACCAGAAATGCCATGTGAAGAACAAGGATATCAGAAAGTTCCTTGATGGTATCTATGTTAGCG

AGAAGGGCAGGATCGCCGAAGAGGAATGAATGATTGTTATCTCTTTGAGTGATAGTTATAGTGCTGCTTT

ACTTTAATGGTGGTAGTTACATGTTTGAACCTTTGTTT

>GR463786.1 JuteSSH6H5S190 JuteSSHlibMP2 Corchorus capsularis cDNA clone JuteSSH6RP1 similar to 60S acidic ribosomal protein like protein, mRNA sequence

CGAGCGGCCGCCCGGGCAGGTACTCCTTCACTTTATCAGCCTGTGGGAATGAGCATTCTGTTACAATAGC

AACAGACAACACATTCTTGTAGGCATTGATGAACATGTGTGGTGCAGCAGCAAGGGTTGGGTATGAGATG

GCAAGTGAGAGTGAGGTAACCATGGAGACACCAACAGCAAATTTCTCAATCAGGTCTTCCTCGGTAAGGT

CAAGAACCTCAGGGCTGAAGACTGAGCCATTGTCATAAACAGACAAGACAATGAGACCGTATGAGAATGG

CCTGATTCCGAGCTTGGCAAGGAGTGCAGCCTCAGAAGACCCAACCTTGTCACCTTCTTGATAAGTTCCA

CAGGGGTGATAATTTCGACAGTACCTCGGCCGCGACCACGC

>GR463785.1 JuteSSH6H5S115 JuteSSHlibMP2 Corchorus capsularis cDNA clone JuteSSH6LECTINP similar to Lectin precursor like protein, mRNA sequence

TCGATTATGGCTTCCTCAAACCCAAATGATCTCATTCCATCTGATATTTCTATCCATTCTCTTACAACAA

TCTTTTTCTTCAGGTGAACTCAACAGAAACCACTTCCTTTTCCATCACCAAGTTTAGCCCAGACCAAAAA

AACCTAATCTTCCAAGGAGATGGCTACACCACAAAAGGGAAGTTGACACTGACCAAGGCAGTAAAGAGCA

CTGTTGGCAGAGCTCTCCATTCCACACCTATCCATATATGGGATAGAGATACAGGTAACGTTGCTAATTT

TGCAACTTCCTTCACTTTTGTCATAGATGCGCCCAGCAGTTACGACGTTGCCGATGGGTTTACGTTCTTC

ATCGCACCGGTGGATACTAAGCCGCAGACTGGCGGGGGATACCTCGGAGTTTTCAATAGCAAAGAATATG

ATAAAACTAGTCAAACTGTTGCTGTGGAGTTTGACACTTTCTATAACGCTGCTTGGGATCCAAGCAATAA

AGAAAGACATATTGGGATTGATGTTAACAGTATCAAATCCGTAAATACTAAGTCGTGGAATTTGCAGAAT

GGTGAACGGGCTAATGTTGTGATAGCTTTTAATGCTGCCACTAATGTGTTAACTGTTACTTTAACTTATC

CTAATTCACTTGAGGAAGAGAATGTAACTTAGTTATACTCTTAATGAAGGTTGGTGCCTTTGAAAAGGAT

GTTTGGTTCCTGAATGGGGGGGAGG

>GR463784.1 JuteSSH6H5S184 JuteSSHlibMP2 Corchorus capsularis cDNA clone JuteSSH6RTN similar to retrotransposon protein like protein, mRNA sequence

GCGTGGTCGCGGCCGAGGTACCAAATCGAGGCAAACTCTGAATACTAGATATGACCTCAAAATAACAGGG

GTCGAGGTCGGCCAGTGAGACGATGGGGGATAAGCTTCATCGTCGAGAGGGAAACAGCCCGGATCACCAG

CTAAGGCCCCTAAATGACCGCTCAGTGATAAAGGAGGTAGGGGTGCAGAGACAGCCAGGAGGTTTGCCAC

CTGCCCGGGCGGCCGCTCG

>GR463783.1 JuteSSH6H5S172 JuteSSHlibMP2 Corchorus capsularis cDNA clone JuteSSH6ATSOC similar to SOC1-like protein, mRNA sequence

CGAGCGGCCGCCCGGGCAGGACACATAGCACTCCTTCTGATGGAGCTGGGAGACTTGAGAGACCCATTTT

TTCTGACTTTTACGTGGGATACAGAAAGCAAAAGAGATGTGCAATTTGAGATTAAGCTAAAGTTCAGATA

TGAATAAGGACAGAACGAAGCTACTCTCTTGTTGACCTATATAACCTTAAAAGCAGAGGGGGAAAACCTT

CACATGTCAATTGATGATTGCTGCTAAAAACCCTGGTAATACAAAAATGGTCAACTTTGCTCACCTCGGC

CGCGACCACGC

>GR463782.1 JuteSSH6H5S183 JuteSSHlibMP2 Corchorus capsularis cDNA clone JuteSSH6ATARFA1E similar to ATARFA1E (ADP-ribosylation factor A1E) like protein, mRNA sequence

GCGTGGTCGCGGCCGAGGTACATGTGCCACCTCCGGGGAAGGGCTGTACGAAGGACTTGACTGGCTCTCC

AACAACATTGCTAACAAAGCTTGAAGGGGAAGGGTCAGATACCCCATACCAGTGACAGTGAGATGTTGGG

CAGCAATATTTGTCAAGAGTTGGAATGTTTTTTCTATGCTATTTTGTGTTTATGGTTTTGTCTTGCTTTG

TAACTGTTGCAAGAATGTATGAGATTGTGTTAGAGATTTTTTACTGTACCTGCCCGGGCGGCCGCTCG

>GR463781.1 JuteSSH6H5S15 JuteSSHlibMP2 Corchorus capsularis cDNA clone JuteSSH6HXET2 similar to XET like protein, mRNA sequence

TTCCACCTATGACTATACTTTTAGGTCTCATGGCCGCGCAGCGGGACCCATCACATTTTCAGAGGCCCAT

TCCTCTATGGCCTCTGGAACCTGATCGAATGAAGAAGAAAGAAATGGGGGAAATAAACCTCCACTGGAGG

TGCAAAAATCCTATCTAAATTATTTTTTAAATTATGATATTGAAAAATAAAATCTTTCGGGAGTTTCTCC

CTAATAATTGCTAAAGCTGCTTCAGCTGAACCTGCATTTAGGGCCTCTGCTGCAGCATCATTAGCTGTCT

GTTGACCTCCTCGAGCAGATCGTCCATCGACCTGAAACTGACCCCAGTCGATGTAATCACCGTCCTTCTC

GATATAGGACTTGACATCTGAGCTGGACTTTGCTCCCTG

>GR463780.1 JuteSSH6H5S95 JuteSSHlibMP2 Corchorus capsularis cDNA clone JuteSSH6HKHSR similar to KH-type splicing regulatory protein like protein, mRNA sequence

ACAGGTTCTTAATTAGCCAGTCAGCCCATGAGCCCTTGGCACGAGTTTACCCAAGTTACCTACGAACCCT

TAAATCCGGTATTTTGGGGGGCCCAGGGATCCCGGATTTTATGAGAAACCCTGGGGGTGCTGCCAGAAAA

GGGAAAAACAGTTGGGGCAAAACACAGAGAATGGGTATTGTTATTCTCCCTTCCCCCCGGGGGCCCGCCC

CAAGGGAGGGGGGGGAGGGGGGGACAGAGGGGGCCAGGGGGGAAAAATACTTTTGTGGTGGGGGGTGGGG

GGGAAAAACACAACCCCCGCGCGGGGGGGGCGCCCCCCCCACCCTCTCTCTCTCTTGGTTATATAAA

>GR463779.1 JuteSSH6H5S30 JuteSSHlibMP2 Corchorus capsularis cDNA clone JuteSSH6HZF1 similar to zinc finger (C2H2 type)-like protein, mRNA sequence

TCCTCTTGATCCGAGGTCACATTTCAGAAGTTTGGGGTGTTTTACGGCTGTGGCCGCGCCGCGCTCCCGG

TGCGAGTGTGCAAACTACTGCGCAGGAGAGGCTGCGGCGAGACCGCCACTGTATTTCGGGGGCGGCCCGG

TGAGGGGCCGATCCCCAACGCCGACCCCCCGGAGGGGTTCGAGGGTTGAAATGACGCTCGGACAGGCATG

CCCGCCAGAATACTGGCGGGCGCAATGTGCGTTCAAAGATTCGATGATTCACTGAATTCTGCAATTCACA

TTACTTATCGCATTTCGCTGCGTTCTTCATCGATGCCAGAACCAAGAGATCCGTTGTTGAAAGTTTTGAT

TCATTTTCAAGACGCCCGCTAGGGTCGCCGAGAAAGGC

>GR463778.1 JuteSSH6H5S181 JuteSSHlibMP2 Corchorus capsularis cDNA clone JuteSSH6HMBD10 similar to MBD10 (methyl-CpG-binding domain 10) like protein, mRNA sequence

GCGTGGTCGCGGCCGAGGTAAAAAAGCCAACTCTTTTTTAGGAAATAGAGATTTTAGAGAGAGAAGGGTT

GTTGATTTTTTCTTCTGTTTTTGGTTTTTTTTGAGAGAAAAATTTTGTTGTTTTTGCAGGTGAAAGAAAA

TGGAGAGTAAAGAGGAAGTTATTTCTGTTGAGCTTCCAGCACCTGCTTCCTGGAAGAAACTGTTTCTTCC

CAAGAGAGTCGGTTCACCTAGGAAGACTGAGATCATGTTTGTTGCACCAACTGGGGAGGAGATTAATAAC

AAAAAACAGTTGGAGCAGTATCTGAAATCCCACCCTGGCAATCCTCCAGTAACAGAGTTTGATTGGGGTA

CCTGCCCGGGCGGCCGCTCG

>GR463777.1 JuteSSH6H5S243 JuteSSHlibMP2 Corchorus capsularis cDNA clone JuteSSH6HRCD1 similar to RCD1 (radical-induced cell death1) like protein, mRNA sequence

GCGTGGTCGCGGCCGAGGTACAGATATGGACCAAATTGCCATTCATTATGAACTATTCAGGAACAAGAAG

ATAAGTCGTCCTGAATTTGTCAAAAAGTTGAGATTGATTGTTGGAGATGATTTGTTGAGAACAACAATAA

CATCTCTGCAACGCAAGGTGCCATCTGCGAATGAAGTGGAGGAGGCAAAGCAGAAGATCATCAAGTGTCC

TGGCAATGTCTGAATTACCCTTATGAAGATTTTGGTATGATGGATGGTTAGTCCTTTGATACGGAGTGAT

GTTCTACCAATGTGTCGACATGAATATGGATTATGGTTGGAAGGTACCTGCCCGGGCGGCCG

>GR463776.1 JuteSSH6H5S156 JuteSSHlibMP2 Corchorus capsularis cDNA clone JuteSSH6HSAP3 similar to senescence-associated protein like protein, mRNA sequence

GCGTGGTCGCGGCCGAGGTGGAAGACATTGTCAGGTGGGGAGTTTGGCTGGGGCGGCACATCTGTTAAAA

GATAACGCAGGTGTCCTAAGATGAGCTCAACGAGAACAGAAATCTCGTGTGGAACAAAAGGGTAAAAGCT

CGTTTGATTCTGATTTCCAGTACCTGCCCGGGCGGCCGCTCG

>GR463775.1 JuteSSH6H5S147 JuteSSHlibMP2 Corchorus capsularis cDNA clone JuteSSH6HSAP2 similar to senescence-associated protein like protein, mRNA sequence

CCCGGGCAGGTACTAGGTTGAATTACTATTGCGACGCGGCCATCAGTAGGGTAAAACTAACCTGTCTCAC

GACGGTCTAAACCCAGCTCACGTTCCCTATTGGTGGGTGAACAATCCAACACTTGGTGAATTCTGCTTCA

CAATGATAGGAAGAGCCGACATCGAAGGATCAAAAAGCAACGTCGCTATGAACGCTTGGCTGCCACAAGC

CAGTTATCCCTGTGGTAACTTTTCTGACACCTCTAGCTCCAAATTCCGAAGGTCTAAAGGATCGATAGGC

CACGCTTTCACGGTTCGTATTCGTACCTGCCCGGGCGGCCGCTCG

>GR463774.1 JuteSSH6H5S142 JuteSSHlibMP2 Corchorus capsularis cDNA clone JuteSSH6HSAP1 similar to senescence-associated protein like protein, mRNA sequence

TTTTCAGCAAGATTCGAGCGGCCGCCCGGGCAGGTCAAGGGAACGGGCTTGGCAGAATCAGCGGGGAAAG

AAGACCCTGTTGAGCTTGACTCTAGTCCGACTTTGTGAAATGACTTGAGAGGTGTAGTATAAGTGGGAGC

TTCGGCAACGGTGAAATACCACTACTTTTAACGTTATTTTACTTATTCCGTGAATCGGAAGCGGGGCTAC

GCCCCTCTTTTTAGACCCAAGACCGGCATCTGTCGGTCGATCCGGGCGGAAGACATTGTCAGGTGGGGAG

TTTGGCTGGGGCGGCACATCTGTTAAAAGATAACGCAGGTGTCCTAAGATGAGCTCAACGAGAACAGAAA

TCTCGTGTGGAACAAAAGGGTAAAGGCTCGTTTGATTCTGATTTCCAGTACCTCGGCCGCGACCACGC

>GR463773.1 JuteSSH6H5S239 JuteSSHlibMP2 Corchorus capsularis cDNA clone JuteSSH6HACY1 similar to Aminoacylase ACY1 like protein, mRNA sequence

CGAGCGGCCGCCCGGGCAGGTGGAGTATCAAGGGAAAAGTAAAAGAAAGGACCTACGGGATACTAAACTA

AAGTCGAGCTTGGTCAGAATTTGCTAGGAATGTGTCAGGTGAATGGGATGGATTTGGGGCAGACTTTTCA

ATTGAAGGGAGTCCAACTGAACTTCCTGAGTCTGTTGTCCCTGAAGCTTACAGGGAATGGGAGGTTAAGG

TGTATGATTGGCAGACCCAGTGCCAACCCTTGCTCAACCTGGGGAGAATGTTATGATTTACAAAAACATA

AAATTGCTTCCAACTGTTGGGTGTGAAGCAGATGCTGCAACACAGTACCTCGGCCGCGACCACGC

>GR463772.1 JuteSSH6H5S195 JuteSSHlibMP2 Corchorus capsularis cDNA clone JuteSSH6HSGST1 similar to Glutathione S transferase like protein, mRNA sequence

CGAGCGGCCGCCCGGGCAGGTACAAAAGGAAAAACTGCTATGATGTAGAAACATATATCTGAGAAAGTAG

ATACCTACTTTTTGAGAATTATTTCATCTTTAGACTTTAGAGATGTGAGAGGAGAGGTGTCTATGCCAAC

AGCAAGCATCACTGAACCACACTCCTAAATATACAACAGCTTACAGTTTGGGATGCACAGTCTATGAAGA

AAACCTTGCTCGGTCATGCGGAGAAGAATAGTCAATATTTGGGCCAGCAGGAATTATCCCAAATGGATTT

ATAGTAGGATGGCTTCCATAGTAATGTCGCTTGATGTGTTGCATGTTCACTG

>GR463771.1 JuteSSH6H5S146 JuteSSHlibMP2 Corchorus capsularis cDNA clone JuteSSH6HXET1 similar to XET like protein, mRNA sequence

GGCCGGCCGGGCAGGTACACTATTCTTTGGACTGATTCTCAGATTATATTTTATGTTGACAATATTCCCA

TTAGAGAGTTTAAGAGAACAGCTTCAATGGGTGGAGATTTCCCAGCTAAGCCAATGTCTTTGTATGCTAC

AATTTGGGATGGTTCAGATTGGGCTACTAATGGTGGCAAATATAGAGTTAATTACAAATATGCTCCTTAC

GTTGCCGAATTCTCCGACCTGATCCTCCACGGCTGCGCGGTTGATCCGATTGAATTTTCATCAAAGAAGT

GTGACACCACTCAAAGTTTGGAAGAACTCTCTACTACTATCACACCATCACAAAGAATGAAGATGGATAG

CTTTAGGAGGAAGCACATGACATACTCATATTGCTATGACCAAACCAGGTACCTGCCCGGGCGGCCGCTC

G

>GR463770.1 JuteSSH6H5S120 JuteSSHlibMP2 Corchorus capsularis cDNA clone JuteSSH6HKUNITZ4 similar to Kunitz-type trypsin inhibitor-like 2 protein precursor like protein, mRNA sequence

CGACCTGCAGGCGGCCGCGAATTCACTAGTGATTCTATGCATCTGCAGCTTGCTTAGAACTCGAAGTTCC

GCCCAACTCAGAATGAAAAGACCATGAATGAACTTCATGTGCTGCAAATTCTGCTCCAGTGGTAGCTGAG

AAACCAATCCTCACCCACTCAGGAACAACATCTTTCAAAGGCACAACTTCATTAAGAGTATAACTAGTTA

CATTCTCTTCCTCAAGTGAATTAGGATAAGTTAAAGTAACAGTTAACACATTAGTGGCAGCATTAAAAGC

TATCACAACATTAGCCCGTTCACCATTCTGCAAATTCCACGACTTAGTATTTACGGATTTGATACTGTTA

ACATCAATCCCAATATGTCTTTCTTTATTGCTTGGATCCCAAGCAGCGTTATAGAAAGTGTCAAACTCCA

CAGCAACAGTTTGACTAGTTTTATCATATTCTTTGCTATTGAAAACTCCGAGGTATCCCCCGCCAGTCTG

CGGCTTAGTATCCACCGGTGCGATGAAGAACGTAAACCCATCGGCAACGTCGTAACTGCTGGGCGCATCT

ATGACAAAAGTGAAGGAAGTTGCAAAATTAGCAACGTTACCTGTATCTCTATCCCATATATGGATAGGTG

TGGAATGGAGAGCTCTGCCAACAGTGCTCTTTACTGCCTTGTTCAGTGTCAACTTCCCTTTTGTGGTGTA

GGCCATCTCCTTGGGAAAAATTAAGGTTTTTTTTGGGTCTG

>GR463769.1 JuteSSH6H5S187 JuteSSHlibMP2 Corchorus capsularis cDNA clone JuteSSH6HLRR2 similar to leucine-rich receptor protein kinase-like protein, mRNA sequence

GCGTGGTCGCGGCCGAGGTACTTTGGTTGCAGTAAAGCAACTATGGAGAGGTGATGGTCTGAAGGTTTTG

GCAGCAGAGATGGAAATTTTGGGGAAAATCAGGCACAGAAATATACTGAAGCTCTATGCTTCTCTTACAA

GAGGTGGTTCAAGCTTTTTAGTGTTTGAGTACCTGCCCGGGCGGCCGCTCG

>GR463768.1 JuteSSH6H5S117 JuteSSHlibMP2 Corchorus capsularis cDNA clone JuteSSH6HKUNITZ2 similar to Kunitz-type trypsin inhibitor-like 2 protein precursor like protein, mRNA sequence

TCGATTCCTCCTATTTGCAATTCTAACCAATCCTTTATTTGCATTCTCCAATAACAATGCTATTGAGCAA

GTGTTGGATACAAATGGTAACCCCCTTATCCCTGGTGATGAATACTACATTTTTCCAGCAAGTGATAACC

CTAAAACTGGAGGACTAACCCTAAACAAAATTAGTGATGCAGAGTGTCCTGTAACTGTCCTACAAAATAA

TGCGACAAGAGGTTTACCAGTTAAATTCACCTTATCAGGAAGTAATAACACTGGTAATAATATCTTGACT

AATACTGATCTTGAAATTGAGTTCACTAAGAAGCCAAATTGCGTTGAATCATCAAAATGGATTGTGTCTG

TTGATGATTTTACTCCTCAAGGTTGTGTTGGTATTGGTGGTCCTGAAAATCATCTTGGTTTAGAAATACT

CAATGGCAAATTTTTAATTGTGAGACATGCTTCTGGATATGTTTATAGGTTTGGATTTTGTTTGGATGTG

AGTGGTGATTGTGGTTTG

>GR463767.1 JuteSSH6H5S118 JuteSSHlibMP2 Corchorus capsularis cDNA clone JuteSSH6HKUNITZ3 similar to Kunitz-type trypsin inhibitor-like 2 protein precursor like protein, mRNA sequence

ACGACCTGCAGGCGGCCGCGAATTCACTAGTGATTTATGAAATCCATTGTATTCCTCCTATTTGCAATTC

TAACCAATCCTTTATTTGCATTCTCCAATAACAATGCTATTGAGCAAGTGTTGGATACAAATGGTAACCC

CCTTATCCCTGGTGATGAATACTACATTTTTCCAGCAAGTGATAACCCTAAAACTGGAGGACTAACCCTA

AACAAAATTAGTGATGCAGAGTGTCCTGTAACTGTCCTACAAAATAATGCGACAAGAGGTTTACCAGTTA

AATTCACCTTATCAGGAAGTAATAACACTGGTAATAATATCTTGACTAATACTGATCTTGAAATTGAGTT

CACTAAGAAGCCAAATTGCGTTGAATCATCAAAATGGATTGTGTTTGTTGATGATTTTACTCCTCAAGGT

TGTGTTGGTATTGGTGGTCCTGAAAATCATCTTGGTTTAGAAATACTCAATGGCAAATTTTTAATTGTGA

GACATGCTTCTGGATATGTTTATAGGTTTGGATTTTGTTTGGATGTGAGTGGT

>GR463766.1 JuteSSH6H5S108 JuteSSHlibMP2 Corchorus capsularis cDNA clone JuteSSH6HKUNITZ1 similar to Kunitz-type trypsin inhibitor-like 2 protein precursor like protein, mRNA sequence

TCGATTTAAACTGACGCATCAAATCCACCTTTAAGAGGCATAATACGTCCAGATTTAACAGAAGCAACAT

CAACAAATACAACATTATAAGAATTAAATATAGTTAAAATTAAACGTGATCCACCTTCTCTCGAATCAAA

AGTATTCAATCCAAGCAAACCACAATCACCACTCACATCCAAACAAAATCCAAACCTATAAACATATCCA

GAAGCATGTCTCACAATTAAAAATTTGCCATTGAGTATTTCTAAACCAAGATGATTTTCAGGACCACCAA

TACCAACACAACCTTGAGGAGTAAAATCATCAACAAACACAATCCATTTTGATGATTCAACGCAATTTGG

CTTCTTAGTGAACTCAATTTCAAGATCAGTATTAGTCAAGATATTATTACCAGTGTTATTACTTCCTGAT

AAGGTGAATTTAACTGGTAAACCTCTTGTCGCATTATTTTGTAGGACAGTTACAGGACACTCTGCATCAC

TAATTTTGTTTAGGGTTAGTCCTCCAGTTTTAGGGTTATCACTTGCTGGAAAAATGTAGTATTCATCACC

AGGGATAAGGGGGTTACCATTTGTATCCAACACTTGCTCAATAGCATTGTTATTGGAGAATGCAAATAAA

GGATTGGTTAGAATTGCAAATAGGAGGAATACAATGGGATTTCATAAATCCCTAGTGAATTCGCCGGGCC

GCCTGCAGGCTCGACCCATATGGGGAGAACCTCCCCCAAC

>GR463765.1 JuteSSH6H5S164 JuteSSHlibMP2 Corchorus capsularis cDNA clone JuteSSH6HLRR similar to leucine-rich receptor protein kinase-like protein, mRNA sequence

AGATAGCGTGGTCGCGGCCGAGGTACTTTGGTTGCAGTAAAGCAACTATGGAGAGGTGATGGTCTGAAGG

TTTTGGCAGCAGAGATGGAAATTTTGGGGAAAATCAGGCACAGAAATATACTGAAGCTCTATGCTTCTCT

TACAAGAGGTGGTTCAAGCTTTTTAGTGTTTGAGTACCTGCCCGGGCGGCCGCTCG

>GR463764.1 JuteSSHRP85 JuteSSHlibMP1 Corchorus capsularis cDNA clone JuteSSHCpn10 similar to Cpn10 Chaperonin 10 kd subunit like protein, mRNA sequence

CGAGCGGCCGCCCGGGCAGGTACTTCGGATGAACAAGGTAACAGGAAGCCGTTATCAGTTGCACCAGGAA

ACACAGTCCTTTACTCCAAGTATGCCGGGAATGACTTCAAGGACACTGATGGGAGTAACTACATTGCTCT

GAGAGCTTCTGAAGTAATGGCAGTGCTCTCTTAGTTGTCTAATAAGATATGCTTTAGGTGTACCTCGGCC

GCGACCACGC

>GR463763.1 JuteSSHPS30 JuteSSHlibMP1 Corchorus capsularis cDNA clone JuteSSHGP10 similar to glutathione peroxidase like protein, mRNA sequence

AGCGTGGTCGCGGCCGAGGTACTCGCTTTAAAGCTGAATATCCCATATTTGATAAGGTTGAGGTGAACGG

TGCTAACACCGCTCCCATATACAAATTCTTGAAGTCCAGCAAAGGCGGACTTTTCGGGGACAGCATCAAG

TGGAACTTCTCCAAGTTCTTGGTCGATAAGGATGGCAATGTTGTTGACCGCTATGCTCCCACAACTTCTC

CTCTTAGCATTGAGAAGGATATAAAGAAGTTGCTGGCCTGATTACTGGAGGCTACTTCCATCAGCTCAGA

GTGAATAAAGTAGATAAATTTCACGAATAAAAACTAGTGTTTCAGACATTTCTCTGCTTTTGTTCCCCAT

CTGATCTGCAAAGATGGGGTTGAACTGTATTGGAAATTAGAACTTGTATTGTACCTGCCCGGGCGGCCGC

TCGA

>GR463762.1 JuteSSHPS31 JuteSSHlibMP1 Corchorus capsularis cDNA clone JuteSSHSLZ similar to secretory laccase like protein, mRNA sequence

AGCGTGGTCGCGGCCGAGGTACGGCCAGGAGCAATGGTGAGAGAATCGGCGAAAGACCTTGTGGTGTAAG

CCCCATCTTGTCCCACAACTGTGAACATGTGGTATTCCACGCCAAACTCCATTGTCTCATTCAAAGCCGC

ATTTACAATTCGAAGGTGATAAACACTTTACCTGCCCGGGCGGCCGCTCGA

>GR463761.1 JuteSSHPS32 JuteSSHlibMP1 Corchorus capsularis cDNA clone JuteSSHXET10 similar to Xyloglucan endotransglucosylase like protein, mRNA sequence

TCGAGCGGCCGCCCGGGCAGGTCAAATTAGATGGAACCATAAGGAAATTGTGTTTTTGGTTGATGATACG

CCAATTAGAGTTTACAAGAACAATGAAGCAAAAGGTATCCCATACCCAAAATCCCAACCCATGGGAGTCT

ATTCAACACTTTGGGAAGCCGATGATTGGGCGACAAGGGGTGGATTAGAGAAAATTGATTGGAGCAAAGC

TCCTTTCTTAGCTTATTACAAGGATTTCGACATCGAAGGTTGTGCTGTTCCTGGACCAGCTAATTGTGCC

TCAAACCCTAGGAATTGGTGGGAAGGCACTGCTTATCAAGGCCTTAATGCCTTGGAGGCTAGAAGATACA

GGTGGGTTCGTATGAACCATATGATCTATGATTATTGCACCGACAAGTCCCGGTACCTCGGCCGCGACCA

CGCT

>GR463760.1 JuteSSHPS34 JuteSSHlibMP1 Corchorus capsularis cDNA clone JuteSSHSACC8 similar to acetyl-CoA carboxylase like protein, mRNA sequence

AGCGTGGTCGCGGCCGAGGTTTTTTTTTTTCCATATATAATTCCAATACAAACTTCATTATTCTTTCCCT

ACTTACATATATTTTTTTCAAACTCTTTACAAATAATCAAATATGGCCAAGAAGGATGGCTGGAAAACAA

AGGAAAACCCCAGGTCATAGTTTCTGGACTTTTGCCCCAAACCCATCATTTTGAGCCGGCCAAGGCCGGA

CGATTTCCGGCTAACCATATATATTATATCTAATACTCTATCAGGTGGGACCAGCTAATTTTCCCCTCTA

AAAATATCCGGATTAGGCCGAGGAGTCTGCACCCCCCTTTCCTTCTCAGGAAGCTCCTTCTAGCTGACAA

AATCCCCCCTATGAGTAGTTGGGGAATTGTTGCAGGAGTTCCGGGCCAAATATGGTTCCTTGATCAATAT

GCATGAAATTATCCCATTTTAAAGGTAGGCGGCAAAGGAATGAGATATATCAAACCTTACTGGCTTACCG

ACTTCTCAATCAGCCAAGCCCCTTACGAAGTTCGTTAACTATTTCTGCCCTGCTGGATGGCCCCTTCTGG

CTTAAAAGGGCAGCAAGACCTTGAGGAAAAGCTGGCATATCTGAAGCTGATTTGCCAATATTGGTCAGCT

GAAGCAGTACCTGCCCGGGCGGCCGCTCG

>GR463759.1 JuteSSHPS35 JuteSSHlibMP1 Corchorus capsularis cDNA clone JuteSSHSAP5 similar to senescence-associated protein like protein, mRNA sequence

AGCGTGGTCGCGGCCGAGGTACGAATACGAACCGTGAAAGCGTGGCCTATCGATCCTTTAGACCTTCGGA

ATTTGAAGCTAGAGGTGTCAGAAAAGTTACCACAGGGATAACTGGCTTGTGGCAGCCAAGCGTTCATAGC

GACGTTGCTTTTTGATCCTTCGATGTCGGCTCTTCCTATCATTGTGAAGCAGAATTCACCAAGTGTTGGA

TTGTTCACCCACCAATAGGGAACGTGAGCTGGGTTTAGACCGTCGTGAGACAGGTTAGTTTTACCCTACT

GATGGCCGCGTCGCAATAGTAATTCAACCTAGTACCTGCCCGGGCGGCCGCTCGA

>GR463758.1 JuteSSHPS36 JuteSSHlibMP1 Corchorus capsularis cDNA clone JuteSSHL16W similar to ribosomal protein L16 like protein, mRNA sequence

TCGAGCGGCCGCCCGGGCAGGTACGTATATTTCCAGACAAACCAGTTACGGTAAGACCTACAGAAACACG

TATGGGTTCGGGGAAAGGATCTCCCGAGTATTGGGTAGCTGTTGTTAAACCGGGCAGAATACTTTATGAA

ATGGGCGGAGTAGCCGAAAATATAGCCAGAAAAGCTATTTCAATAGCGGCGTCAAAATGCCTATAAAAAC

TCAATTCATTATTTCAGGATAGGAATATAGAACCAAAGGAAAGAAGTCTTGGGAATAAAAAAAACAATTG

CGGGTTTCCTTTTTTTTGACAAACAATATTTCTTTTTTTTCTCCGTCCTTTGTTGCATTGAAAGAAGAGA

TTAAAAAAATGATTCAACCTCAGACCCATTTGAATGTAGCAGATAATAGCGGGGCCCGAGAATTGATGTG

TATTCGAGTCATAGGAGCTAGTAATCGCCGATATGCTCATATTGGTGACGTTATTGTTGCTGTGATCAAG

GAAGCAGTACCTCGGCCGCGACCACGCT

>GR463757.1 JuteSSHPS37 JuteSSHlibMP1 Corchorus capsularis cDNA clone JuteSSHSL9 similar to secretory laccase like protein, mRNA sequence

GAGCGTGGTCGCGGCCGAGGTCGGCCATCCTGGAGAAACATATGGCTGCAGCAACGATTCAACATACGAG

CAGCAAGTAGATTACCAAAGTGTTTATCACCTTCGAATTGTAAATGCGGCTTTGAATGAGACAATGGAGT

TTGGCGTGGAATACCACATGTTCACAGTTGTGGGACAAGATGGGGCTTACACCACAAGGTCTTTCGCCGA

TTCTCTCACCATTGCTCCTGGCCGTACCTGCCCGGGCGGCCGCTCGA

>GR463756.1 JuteSSHPC24 JuteSSHlibMP1 Corchorus capsularis cDNA clone JuteSSHHsp901 similar to hsp90 protein family (Stretch 13-112) like protein, mRNA sequence

AGCGTGGTCGCGGCCGAGGTCCATAAGGCTGCTTCTGACGTTCGTGATGAGTTTGTATCTGTTACTGAGA

AGTTAATGGATGAATTGGCACAATGCTACAATGTGCTCCCCCAACTTGATATTAATAACACTATAGACCA

CCGCCCCGAAGGGGACGAAAAATGGTTTTTAGAGAACGAGAAGACGGTTACGCAGTTTTGCCGCAAGCTG

GCTGCTGAACGCCCTCTTAAGGATATTCGCGATGAGTATAATTACCCCAAAAAGAAAGGTATTAAGGATG

AGTGTTCAAGATTGCTGGAGGACCTGCCCGGGCGGCCGCTCGA

>GR463755.1 JuteSSHRP115 JuteSSHlibMP1 Corchorus capsularis cDNA clone JuteSSHNT similar to Nitrite transporter like protein, mRNA sequence

GCGTGGTCGCGGCCGAGGTACACAAGATTAAAAACTTGCAAAGCTGTTATCAACCAATAAAAATACTCCA

ATTTTCCCTTGTTCAAGTTATTGTCCGGAAGCCAATTGGATCCATCAGGCCCAGCACTGAATTTATGCAC

CGATGAAACCAACAAAGTACCTGCCCGGGCGGCCGCTCG

>GR463754.1 JuteSSHRP52 JuteSSHlibMP1 Corchorus capsularis cDNA clone JuteSSHCS similar to Cysteine synthase like protein, mRNA sequence

CGTGGTCGCGGCCGAGGTACTCCAGTCCCAATACCAGAGACAAGGATATCTACCTTCCCTCCAGTGCCTG

TCCATATCTCTGGTCCAGTTGTCTCATAATGGATCTTTGGGTTGGCAGGGTTTTCAAATTGCTGAAGAAT

GTAGGAATTTGGAGTCTTGTCTCGAATTTCCTCAGCTTTCTGAACAGCTCCTTTCATGCCCCGAGCTGGA

TCTGTGAGAACCAGCTCAGCTCCAAAAGCTCGGAGGACCATTCTCCTTTCCAAACTCATTGAAGAGGGCA

TGGTAATTATCAGTTTGTAACCCTTAGCAGCAGCCATGAATGCCAATCCTATTCCGGTATTACCACTGGT

TGGCTCAATACCTGCCCGGGCGGCCGCTCG

>GR463753.1 JuteSSHRP23 JuteSSHlibMP1 Corchorus capsularis cDNA clone JuteSSHSTPP similar to Serine/threonine protoin phosphatase like protein, mRNA sequence

GCGTGGTCGCGGCCGAGGTACCTCTCAATTCTTGCTTTAACACTTTATTCAATAGTTGGCTCATATAGTA

AGATAAAAGTCAGTCACAGAGACATGTCTCATAAAGAAGCCTGGACTACGGCAATATTCGTCGAATTGCC

AAGTCGAATCACCAGCTACTCCAGGTAGAAAACTACATTACATGTGAGCATTGAAGGGTGGGCATGTGTT

GAAATATTGCTGTTGCAGCATCCGTGACTAAAGTCGTTGCTCCAATGTTGATCACACACAAATTCTGATG

GGGACACGAAGCTGAAGCAGATTAGGCTGAGAAAGAAGACTTGGCGAGTATTAATACAAAGCTAACGGAT

CTCTCCTTCATAAAGCTGAACCTGCTTTCCTCCTCTGCCATCCAACAACCTTCTTATTATTATCTTCACC

ATCAGTAACTCCATCAGAAGATGCCATGTTGTTTTCATGCTCTTTTACTTCTTCCATGTACCTGCCCGGG

CGGCCGCTCG

>GR463752.1 JuteSSHRP80 JuteSSHlibMP1 Corchorus capsularis cDNA clone JuteSSHGT5 similar to O-glucosyltransferase like protein, mRNA sequence

CGAGCGGCCGCCCGGGCAGGTACTTAATTTAGTTTTTCCTAAGCTCATTTACGAAGGACTTAAGGTTCTT

GTCGGAAGAACCACCTTCCATGGCAGCTTCTCTTGAAAATTCTTTCCATTTCTTAACATTCCTCTTGATT

TCTTCTCCTCTTTCCCCATCTCCCATCACCATTTCCAAACATCTCTTAATCTCGCAACCTACGACTAATC

CTTGTTCATTCTTCGTAACCCCTCACCCCTGTCTTCCACACATCTTGGGACAAGCTTCGAGTTGGGTCCC

TTGATCTGTCCATTTGAGGGATTTGTCACTTGTCGAACCTCGGCCGCGACCCCGC

>GR463751.1 JuteSSHRP57 JuteSSHlibMP1 Corchorus capsularis cDNA clone JuteSSHGT4 similar to O-glucosyltransferase like protein, mRNA sequence

CGAGCGGCCGCCCGGGCAGGTACACATGAGTGGATATATAGGAATCCAAATCTGCCGAACCACTCATGTT

ATGATCTTCTACATCCTAGGTCTTCCCGTTCCGTCATCTGGCTTATGTTCTTCATGTAGCATTCAGACCG

AATGACTCTATGAAATTACGTCGATACTTCCACATATTATGGGTAACGTAGGAGACATCTCTATTTTTCC

CCGGGGGAATCTTTAGAATTACCACTGCTTAGCTTTCAATTCGCCTCTGACCATCAAATGAAATGTGGAT

AACCCGTCCTCCTCTCTTTGAAACAAGGGGCGCTTCTGGTTCTGTCGGTGCTTGAAACAATTTTGTCTTC

TCCATATTACTATATCTCTAGAGTCAATAATTTTATATGAGGAACTACTGAACTCAATCACTTGCTGCCG

TTACTCTTCAGTTTTCTGTTGAGGTCTATCCTGTAGAGGTACCTCGGCCGCGACCACGC

>GR463750.1 JuteSSHRP62 JuteSSHlibMP1 Corchorus capsularis cDNA clone JuteSSHGT2 similar to O-glucosyltransferase like protein, mRNA sequence

CGAGCGGCCGCCCGGGCAGGTACTTAATTTAGTTTTTCCTAAGCTCATTTACGAAGGACTTAAGGTTCTT

GTCGGAAGAACCACCTTCCATGGCAGCTTCTCTTGGAAATTCTTTCCATTTCTTAACATTCCTCTTGATT

TCTTCTCCTCTTTCCCCATCTCCCATCACCATTTCCAAACATCTCTTAATCTCGCAACCTTCGACTAATC

CTTGTTCATTCTTCGTATCCCTCACCCCTGTCTTCCACACATCTTGGACAAGCTTCGAGTTGGTTCCTTG

ATCTGTCCATTGAGGAATTGTCACTGTCGACCTCGGCCGCGACCACGC

>GR463749.1 JuteSSHRP32 JuteSSHlibMP1 Corchorus capsularis cDNA clone JuteSSHGT3 similar to O-glucosyltransferase like protein, mRNA sequence

CGAGCGGCCGCCCGGGCAGGTACTTAATTTAGTTTTTCCTAAGCTCATTTACGAAGGACTTAAGGTTCTT

GTCGGAAGAACCACCTTCCATGGCAGCTTCTCTTGAAAATTCTTTCCATTTCTTAACATTCCTCTTGATT

TCTTCTCCTCTTTCCCCATCTCCCATCACCATTTCCAAACATCTCTTAATCTCGCAACCTACGACTAATC

CTTGTTCATTCTTCGTAACCCTCACCCCTGTCTTCCACACATCTTGGACAAGCTTCGAGTTGGTTCCTTG

ATCTGTCCATTGAGGAATTGTCACTGTCGACCTCGGCCGCGACCACGC

>GR463748.1 JuteSSHRP45 JuteSSHlibMP1 Corchorus capsularis cDNA clone JuteSSHGT1 similar to O-glucosyltransferase like protein, mRNA sequence

CGGCCGCCCGGGCAGGTACTTAATTTAGTTTTTCCTAAGCTCATTTACGAAGGACTTAAGGTTCTTGTCG

GAAGAACCACCTTCCATGGCAGCTTCTCTTGAAAATTCTTTCCATTTCTTAACATTCCTCTTGATTTCTT

CTCCTCTTTCCCCATCTCCCATCACCATTTCCAAACATCTCTTAATCTCGCAACCTTCGACTAATCCTTG

TTCATTCTTCGTAACCCTCACCCCTGTCTTCCACACATCTTGGACAAGCTTCGAGCTGGTTCCTTGATCT

GTCCATTGAGGAATTGTCACTGTCGACCTCGGCCGCGACCACGC

>GR463747.1 JuteSSHRP18 JuteSSHlibMP1 Corchorus capsularis cDNA clone JuteSSHSUS2 similar to SUS2(ABONORMAL SUSPENSOR 2)like protein, mRNA sequence

CGAGCGGCCGCCCGGGCAGGTACCAAGCTTCACCCCGTATTTCATGCTAACCGTATGCTTCACTCCCATG

AAATTGTAATTCCATGGTACCATTGTCGGGAACCATATAAAATCCGAGGAACCGATCACCTCGGCCGCGA

TCATCTGCACCTTCTCATAATGAGTTGGACCGTAACCATGAGGATTGCTTCCTGTATCTTTATTAACCCG

TCCCCATTCATAACCACTTGGAGTAAGCTTGTATGCTGTTAACGAGCAGGAACCTGGAGTGAAACTGCAT

GTCAAGATAATGCACTTCTCTCCATCCCACTGCTTGTTATTCTCTAAAATCCGGGCATGTGAGGTAACAT

CCTGTGGCGATAATTGAGGAAGCTCATTTGGTTGTGTGTGCATCCAACCAAAGGCTCGAGATCATTCAAG

AAATCATGTTCAGGTTGAGCAGATGGAAGATGAACTTGCTGATGAGTTCCCCACTGCGGTACCTCGGCCG

CGACCACCAC

>GR463746.1 JuteSSHRP14 JuteSSHlibMP1 Corchorus capsularis cDNA clone JuteSSHCMO similar to Carbon monoxide Oxygenase(ACYB-2)like protein, mRNA sequence

CGAGCGGCCGCCCGGGCAGGTACAACAAAGGCTCCATATAGTATGGTGACAATGGCTGTGAAATTCACTA

GGTAGGCCTCGGCACCATATTTGTCAACACCAGAGACCTCAAGGAAAGTGAGCTTCTCCAGGAACCCTAT

GGCAGCAGTGGCTACGGCCAATATGTATACAAACATTCCCACCAGAACGTGCCAGGGAAGCGATTCGCTT

CTTAATCCCCCTGATCCTCCTGGGTAGAAGAATACTAGAAACCCATATATCCACTGAAGGCCATAAATGA

CAGTGACTCCAATCCCAAGTCAAGAATGCAAACTGTAGAGATTGGCAATGTTGCTCTCATTATGGTACCT

CGGCCGCGACC

>GR463745.1 JuteSSHRPC34 JuteSSHlibMP1 Corchorus capsularis cDNA clone JuteSSHACtO similar to Acetyl CoA oxidase like protein, mRNA sequence

ATGCTTGAATTGGCATCACCTCACAGTTTCAGGAGATGTGACATTTGGAAAAAATGTTTCATTAAAGGGA

ACGGTTATCATCATTGCAAATCATGGTGACAGAATTGATATCCCA

>GR463744.1 JuteSSHRP67 JuteSSHlibMP1 Corchorus capsularis cDNA clone JuteSSHACo similar to Acyl CoA oxidase like protein, mRNA sequence

GCGTGGTCGCGGCCGAGGTACAAGATACAATGCATACACATTGCAAAGGATCTCCAACTGTCTTTTCACT

CCCTTTCCTGGTATGTCTTGTTGCAATTTCTCGATAAACTTGGAAACAACAATTAACTGACAATGAGCAA

CTGCAGCCTCAACCAAATTAGGTGAGAGTTCAGCAAAACCCTCCTCAGGATTTGAAAATTTGCTAAGGCT

TTGAGCACAGGCAACAGACATCCGAAGAGCCCTTGCTTCAAATGCCTCCACTATTACACTGGGGTTTAGC

CAATCCTCAGCCCTTTGAACATCACAACTGCATTGCATGAGGTGCTCTGCTCGTCCCATATAGGCTGTTG

TCCCAACAGGTTTCTTTCCAGACCCCAGCTGGGAAACGGTCTTCATTAAAAACCTTGCAACCTGTAAAAG

CAGCACTACATTGTCTCCTTCATAAGTACCTGCCCGGGCGGCCGCTCG

>GR463743.1 JuteSSHPC13 JuteSSHlibMP1 Corchorus capsularis cDNA clone JuteSSHACC1 similar to 1-aminocyclopropane-1-carboxylic acid oxidase like protein, mRNA sequence

TCGAGCGGCCGAGCGTGGTCGCGGCCGAGGTACGGATACCGGAAAGGCTGGATACCTCGGTTATTAGAGG

ATTTTGGAGATGGAGGTGCTTTTCCAGAGATCCATGTGGCCCAGTATCCACTGGATATGGGACGAAAGAA

AAAAATGTCGAATGCGCTGGCCATTCAGGTGGATTCTGAAGGAAAAATTAAATATGATGCAATTGCTCGA

CAAGGACAGTCAAAAGACAAGGTCATTTATAGCAAATACACTGACCTGGTTCCAAAGGAGGTTATGAATG

CAGATGATCCAGACCTGCAAAGGCCCGATGAAGAAGCTATTAAAGAGATAACAGAAAAGACAAGAGTAGC

CTTAGAAAAATCTGTATCACAGAAGGTCGCCGCAGCCATGCCAGTTCGAGCAGCTGACAAATTGGCTCCT

GCTCAGTATATCCGATACACACCATCTCAGCAAGGAGTGGCATTCAACTCTGGAGCTAAACAGAGGGTTA

TTCGGATGGTAGAAATGCAGAAAGATCCAATGGAGCCTCCAAGGTTCAAGATTAATAAGAAAATTCCCCG

GGGGACCACCTTCTCCTCCTGCGCCTGTCATGCATTCTCCTAGCCGAAAGATGACTGTAAAGGAACAACA

AGAGTGGGAAGATTCCTCCTTGTATTTCTAACTGGAAAAATGCAAAGGGTTATACAATTCCATTAGACAA

ACGTCTGGCTGCTGATGGAAGAGGACTACAGACAGTACCTGCCCGGGCGGCCGCTC

>GR463742.1 JuteSSHPC12 JuteSSHlibMP1 Corchorus capsularis cDNA clone JuteSSHSHMT1 similar to Serine hydroxymethyltransferase (Stretch 7-45 amino acid)like protein, mRNA sequence

AGCGTGGTCGCGGCCGAGGTCCTGTTGATGCTAAAGGTGAGCCGCTTAAAGCTACCATTTATATGGCTGT

TGGTTTCTATGTGGCTAAATACGTTAACAAAAAGTCAGATATGGACCTTGCTGCTAAAGGTCTAGGAGCT

AAAGAATGGAACAACTCACTAAAAACCAAGCTGTCGCTACTTCCCAAGAAGCTGTTCAGAATCAGAATGA

GCCGCAACTTCGGGATGAAAATGCTCACAATGACAAATCTGTCCACGGAGTGCTTAATCCAACTTACCAA

GCTGGGTTACGACGCGACGCCGTTCAACCAGATATTGAAGCAGAACGCAAAAAGAGAGATGAGATTGAGG

CTGGGAAAAGTTACTGTAGCCGACGTTTTGGCGGCGCAACCTGTGACGACAAATCTGCTCAAATTTATGC

GCGCTTCGATAAAAATGATTGGCGTATCCAACCTGCAGAGTTTTATCGCTTCCATGACGCAGAAGTTAAC

ACTTTCGGATATTTCTGATGAGTCGAAAAATTATCTTGATAAAGCAGGAATTACTACTGCTTGTTTACGA

ATTAAATCGAAGTGGACTGCTGGCGGAAAATGAGAAAATTCGACCTATCCTTGCGCAGCTCGAGAAGCTC

TTACTTTTGCGACCTTTCGCCATCAACTAACGATTCTGTCAAAAACTGACGCGTTGGATGAGGAGAAGTG

GCT

>GR463741.1 JuteSSHPC35 JuteSSHlibMP1 Corchorus capsularis cDNA clone JuteSSHXET5 similar to Xyloglucan endo-transglycosylase (XET)like protein, mRNA sequence

TCGAGCGGCCGCCCGGGCAGGTACCGAGGAAGGGGATGCCTCACAGCAAGAGGGCTGCACCTTAGGTTCT

GACCCCATCTGCCTCAGTGAGAGCCAGGTTTCTGAGGAACAAGAAGAGATGGGAGGGCAAAGCAGCGCGG

CCCAGGCCACGGCCAGTGTGAATGCAGAGGAGATCAAGGTAGCCCGTATTCATGAGTGTCAGTGGGTGGT

GGAGGATGCTCCAAACCCGGATGTCCTGCTGTCACACAAAGATGACGTGAAGGAGGGAGAAGGTGGTCAG

GAGAGTTTCCCAGAGCTGCCCTCAGAGGAGTGAAAGGGACAATTTGGCTGAAGTCTTTCTCTGAAAAAAG

CCAAAGGGTTATAGGGGTACCTCGGCCGCGACCACGCT

>GR463740.1 JuteSSHPC32 JuteSSHlibMP1 Corchorus capsularis cDNA clone JuteSSHXET4 similar to Xyloglucan endo-transglycosylase (XET)like protein, mRNA sequence

AGCGTGGTCGCGGCCGAGGCACTTAAATTCTCCTGGGGTCTCCTGCTTTCTCTTAACCATTTGCTTAATA

TATATCTACCTAAAGGAGACTTCTGAATTGTAAATGAACTTAAAAATAGAATGTGGATGCAAAATATCAC

ATAAGACATCATGATAACATTTGAAGAAAAAATAAAACTGTAGACCCTAACAGTTGTGATATTTGGTGGT

TTCATGTGGTAATGTAATTTTCTGTTTAATTACAGTACCTGCCCGGGCGGCCGCTCGA

>GR463739.1 JuteSSHPC22 JuteSSHlibMP1 Corchorus capsularis cDNA clone JuteSSHXET3 similar to Xyloglucan endo-transglycosylase (XET)like protein, mRNA sequence

GGCCGCCCGGGCAGGTACTGGATGACCGCTTGGTGTTCACAGTCTTCCCTGCACCAGATTCTCCAGTGAT

CAGGATTGACTGATTCTCTCGGTCAGTCAGCATGAACTGATAGGCGTTGTCAGAGATGGAGAAGATGTGG

GGCGGGGCCTCCTGGCGCTTTTTGCCTCGGTAGGCTGTCACCACCTCGGGCTTATACACAGGCAGCCACT

TGTAGGGGTTGACAGTGACACAGAAGAGACCTGAATAGGTGTAGATCATCCAGGCTGCATAACGTTC

>GR463738.1 JuteSSHRP19 JuteSSHlibMP1 Corchorus capsularis cDNA clone JuteSSHXET2 similar to Xyloglucan endo-transglycosylase (XET) like protein, mRNA sequence

GCGTGGTCGCGGCCGAGGTACGGGACTTGTCGGTGCAATAATCATAGATCATATGGTTCATACGAACCCA

CCTGTATCTTCTAGCCTCCAAGGCATTAAGGCCTTGATAAGCAGTGCCTTCCCACCAATTCCTAGGGTTT

GAGGCACAATTAGCTGGTCCAGGAACAGCACAACCTTCGATGTCGAAATCCTTGTAATAAGCTAAGAAAG

GAGCTTTGCTCCAATCAATTTTCTCTAATCCACCCCTTGTCGCCCAATCATCGGCTTCCCAAAGTGTTGA

ATAGACTCCCATGGGTTGGGATTTTGGGTATGGGATACCTTTTGCTTCATTGTTCTTGTAAACTCTAATT

GGCGTATCATCAACCAAAAACACAATTTCCACCTGCCCGGGCGGCCGCTCG

>GR463737.1 JuteSSHRP9 JuteSSHlibMP1 Corchorus capsularis cDNA clone JuteSSHXET1 similar to Xyloglucan endo-transglycosylase (XET)like protein, mRNA sequence

GCGTGGTCGCGGCCGAGGTACGGGACTTGTCGGTGCAATAATCATAGATCATATGGTTCATACGAACCCA

CCTGTATCTTCTAGCCTCCAAGGCATTAAGGCCTTGATAAGCAGTGCCTTCCCACCAATTCCTAGGGTTT

GAGGCACAATTAGCTGGTCCAGGAACAGCACAACCTTCGATGTCGAAATCCTTGTAATAAGCTAAGAAAG

GAGCTTTGCTCCAATCAATTTTCCCTAATCCACCCCTTGTCGCCCAATCATCGGCTTCCCAAAGTGTTGA

ATAGACTCCCATGGGTTGGGATTTTGGGTATGGGATACCTTTTGCTTCATTGTTCTTGTAAACTCTAATT

GGCGTATCATCAACCAAAAACACAATTTCCTTATGGTTCCAACCTGCCCGGGCGGCCGCTCG

>GR463736.1 JuteSSHPC23 JuteSSHlibMP1 Corchorus capsularis cDNA clone JuteSSHPTP similar to Phosphatidylinositol transfer-like protein III, mRNA sequence

AGCGTGGTCGCGGCCGAGGTCCTGTCTCATCATGGAAGGCGCTGAATTTACGGAAAACATTATTAAATGG

CGTCGAGCGTCCGGTTAAAGCCGCTGAATTGTTCGCGTTTACCTTGCGTGTACGCGCAGGAAACACTGAC

GTTCTTACTGACGCAGAAGAAAACGTGGCGTCAAAAATTACGTGCAGAAGGAGTGATGTAATGTCTAAAG

GTAAAAAACGTTCTGGCGCTCGCCCTGGTCGTCCGCAGCCGTTGCGAGGTACTAAAGGCAAAGCGTAAAG

GCGCTCGTCTTTGGTATGTAGGTGGTCAACAATTTTAATTGCAGGGGCTTCGGACCTGCCCGGGCGGCCG

CTCGA

>GR463735.1 JuteSSHRP7 JuteSSHlibMP1 Corchorus capsularis cDNA clone JuteSSHATCP1a similar to Hypersensitive reaction associated Ca+-binding ATCP1 like protein, mRNA sequence

GCGTGGTCGCGGCCGAGGTACAATTCGGCATTTCCAATGGAAACATAATAAAAAGTGAAATGTTCAAAAC

AAAAGGGACTTTAAACTCAAAACTTAAACTAAAAAGGCAATTTAATGAGCAAAGTCAACAGCTAAAATCT

TAAGCAAACCATCAAAAGAAACGCCTTGATTTTCATCGCCGCCACCCAATCTGATCATAGCCTTAATATC

TTCATCGGTAGCAGAAAAACCAGCCCAGTTCATGTAACTTTTCAAGTCTTGATGGCTCAATCTCCCATCT

CCATCTTTGTCCATAACCTTAAAAGCATCTGCCATAACCCCAAAACCCGACGCCGCCGAACTCTGTTTCC

CTCGAGACCTGCCCGGGCGGCCGCTCG

>GR463734.1 JuteSSHRP10 JuteSSHlibMP1 Corchorus capsularis cDNA clone JuteSSHATCP1 similar to Hypersensitive reaction associated Ca+-binding ATCP1 like protein, mRNA sequence

GCGTGGTCGCGGCCGAGGTCCAATTCGGCATTTCCAACGGTAAACATAATAAAAAGTGAAATGTTCAAAA

CAAAAGGACTTTAAACTCAAAACTTAAACTAAAAAGGCAATTTAATGAGCAAAGTCAACAGCTAAAATCT

TAAGCAGACCATCAAAAGAAACGCCTTGATTTTCATCGCCGCCACCCAATCTGATCATAGCCTTAATATC

TTCATCGGTAGCAGAAAAACCAGCCCAGTTCATGTAACTTTTCAAGTCTTGATGGCTCAATCTCCCATCT

CCATCTTTATCCATAACCTTAAAAGCATCTTCCATAACCCCAAAACCCGACGCCGCCGAGCTCTGTTTCC

CTCGAGACCTGCCCGGGCGGCCGCTCG

>GR463733.1 JuteSSHRP72 JuteSSHlibMP1 Corchorus capsularis cDNA clone JuteSSHUBQC similar to Ubiquitin-conjugating enzyme like protein, mRNA sequence

CGAGCGGCCGCCCGGGCAGGTACCTCGGGAGGTTGGACCAGCTTTCGGTTATGAGGGGCTGCCATCTCCT

TTTTCAGCTGTGTCAGTATATCCTCCATGGTATACTCCCTCTGCCAATTTGCAAGTAGCCCAAACTTCTT

TGGTTCCACCACTCCAGTCTCATGGTTCACACAAGTCATGTTAACCCTTGAATGAAAACGGACACTCGGT

GGCTTCTCAGGATAGTCTTTATCACAGAACAGCTCCAGCTGATAGATTCGACCTTCATGTACCTCGGCCG

CGACCACGC

>GR463732.1 JuteSSHRP79 JuteSSHlibMP1 Corchorus capsularis cDNA clone JuteSSHSTPK similar to Ser/ThrProtein kinase (PRK) like protein, mRNA sequence

GCGTGGTCGCGGCCGAGGTACAAAAACCAGAGACCTTATATGTGAAATCTTGTAGAACTGTCATAGTAAG

TCCATCAAACTTCTGTAACAGATTAAGCCGGTGAACGGGCTGGATTTTTCCCAGAAGGCAGTTGCTTTTA

AATACCATCATAGTAACCTCCAAATGTTATTTGAGCTCTGGTGATTGGAACTCCTCTGGCTGCAGAAATT

GCTTCATTGAGTTTGGTTGCATGATGAACCACTTGCTTCATTCTCTTGTCATCAGTGTAACGCTGGAACA

GAAGCTTCTGATTCTTCTGATGCTGTCTGTGTCTTGTTGACTCCCAGTATGATGATACAAAAAGTAGCAG

CAACAACACCAAAGAACAGATTGCTCCCATTCCCAGTTAAACCCCCATTCCCAAAATTCAGGCCCACCAG

ATTCACCTGCCCGGGGGGCCGCTCC

>GR463731.1 JuteSSHRP13 JuteSSHlibMP1 Corchorus capsularis cDNA clone JuteSSHECT2a similar to ECT2 [Arabidopsis thaliana] like protein, mRNA sequence

CGAGCGGCCGCCCGGGCAGGTACTGTGATCTAAAATTCTAAATCATCAGATCAAAATTTCTGGAAACTCC

TCTTCATATTCTACAAGACCATGAACCGTCGCAAGGGGATTGGCAGGAACTAGCAACCATTGGCAATCCC

ATTCGATAAAACCTTCTTGTCTGGAACTACAATTGGATTCGCTGAACCATTCTCTGAAAGCTTTAGATCC

CCATTGGAATTATCAGTTAAGCTTGGTTCCTTAACCAAATCTGATCCAGCTTCCAAACAGTTCTGTGTTC

CATTTGCTACATCTTTCTTCTCATCTACAGGCTTTCCTTCCCATACCTGCTTTTGAAATTGCTGTTTAGC

CTTTTTATCCTGGATTGCCTTCTGGCGGACCTCATAAAACTCAAAATCATCCAAAATGCAAGTTTTGCTG

GAATGTTCCTTGAAAATTTTGATCAACTTCAGTCCTTGCTCAAGCTTGATCTCTTGTGTACCCCTGCTAT

TCGTGACAGGTTTATTCTCATTGTTCTCTAAGGTGATGTGCTTCAAAGAATTGTTTGGTACCTCGGCCGC

GACCACGC

>GR463730.1 JuteSSHPC4 JuteSSHlibMP1 Corchorus capsularis cDNA clone JuteSSHECT2 similar to ECT2 [Arabidopsis thaliana] like protein, mRNA sequence

ATGCTTGAATTGGATCACCTCACAGTTTCAGGAGATGTGACATTTGGAAAAAATGTTTCATTAAAGGGAA

CGGTTATCATCATTGCAAATCATGGTGACAGAATTGATATCCCACCTGGAGCAGTATTAGAGAACAAGAT

TGTGTCTGGAAACCTTCGCATCTTGGACCACTGAAATGAAAAATACTGCGGACACTTAAATAAT

>GR463729.1 JuteSSHPC25 JuteSSHlibMP1 Corchorus capsularis cDNA clone JuteSSHLip similar to Lipocalin-like domain (Stretch 7-91 amino acid) containing protein, mRNA sequence

CCATGGTGAAGCTGAGCAAAGAGGCCAAGCAGAGACTACAGCAGCTCTTCAAGGGGAGCCAGTTTGCCAT

TCGCTGGGGCTTTATCCCTCTTGTGATTTACCTGGGATTTAAGAGGGGTGCAGATCCCGGAATGCCTGAA

CCAACTGTTTTGAGCCTACTTTGGGGATAAAGGATTATTTGGTCTTCTGGATTTGGAGGCAATCAGCGGA

CAGCATGGAAGATGTGTGCTCTGGCTCGGATAAGAGATGGGACATCATTCAGTCACTAGTTGGATGGCAC

AAGGCTCTTCACAGACGCA

>GR463728.1 JuteSSHRP76 JuteSSHlibMP1 Corchorus capsularis cDNA clone JuteSSHGST4 similar to Glutathione S-transferase N terminal domain like protein, mRNA sequence

GCGTGGTCGCGGCCGAGGTACGCTGCTTGGAAAGAATCTCCTCATATTTGTCCATGGTTTCATATAATTG

TTTCACAGCCTCATCATAAGGCCCTTGCTTTCTAGCAAAGCCGCATCTGTAAACACCATTATTGATTCCA

CTGTATATCCATTCATTTGTCTCATCAATTTGAGCTCTCAAATGAGGAGGATATAGGTCCAAAGCAGGAT

TCTCTGCTATATCATTGAATTCAGTATTAAGCATGCGGATTATCTCTGCGCTCTCATTACTAACATTAGT

CTTGAATTTCTTGTCCCAAAGGACAGGAACGGTGTACCTGCCCGGGCGGCCGCTCG

>GR463727.1 JuteSSHRP26 JuteSSHlibMP1 Corchorus capsularis cDNA clone JuteSSHGST2 similar to Glutathione S-transferase N terminal domain like protein, mRNA sequence

GCGTGGTCGCGGCCGAGGTACGCTGCTTGGAAAGAATCTCCTCACATTTGTCCATGGTTTCATATAATTG

TTTCACAGCCTCATCATAAGGCCCTTGCTTTCTAGCAAAGCCGCATCTGTAAACACCATTATTGATTCCA

CTGTATATCCATTCATTTGTCTCATCAATTTGAGCTCTCAAATGAGGAGGATATAGGTCCAAAGCAGGAT

TCTCTGCTATATCATTGAATTCAGTATTAAGCATGCGGATTATCTCTGCGCTCTCATTACTAACAATAGT

CTTGAATTTCTTGTCCCAAAGGACAGGAACGGTGTACCTGCCCGGGCGGCCGCTCG

>GR463726.1 JuteSSHRP21 JuteSSHlibMP1 Corchorus capsularis cDNA clone JuteSSHGST1 similar to Glutathione S-transferase N terminal domain like protein, mRNA sequence

CGAGCGGCCGCCCGGGCAGGTCAGCAACATTACAATGGCTGCTAACAACTTTGAGAATTTGCCTCCTCCC

TTGGATGCTACTGCTGAGCAACCCCCTCTGTTTGATGGAACTACCAGGTTGTATACTTCTTATTCTTGCC

CCTTTGCCCAGCGAGTGTGGATCACAAGGAATTATAAGGGGTTGCAAGACAAGATCAAATTGGTTCCTTT

GAACCTTCCAAACAGACCTGCTTGGTACAAGGAAAAAGTCTACCCTGAGAACAAGGTGCCTGCCTTAGAA

CACAACGGGAAAATCATCGGGGAGAGTCTTGATCTAATTAAGTACCTCGGCCGCGACCACGCT

>GR463725.1 JuteSSHPC30 JuteSSHlibMP1 Corchorus capsularis cDNA clone JuteSSHGP similar to Glutathione peroxidase (Stretch 4-34 amino acid) like protein, mRNA sequence

TCGAGCGGCCGCCCGGGCAGGTACTACTTGGTTCCCGATATGGATGATGAAGAAGGAGAAGGAGAAGAAG

ATGATGATGATGATGAAGAGGAGGAAGGATTAGAAGATATTGACGAAGAAGGGGATGAGGATGAAGGTGA

AGAAGATGAAGATGATGATGAAGGGGAGGAAGGAGAGGAGGATGAAGGAGAAGATGACTAAATAGAACAC

TGATGGATTCCAACCTTCCTTTTTTTTAAATTTTCTCCAGTCCCTGGGAGCAAGTTGCAGTCTTTTTTTT

TTTTTTTTTTTTTCCCCTCTTGTGCTCCGTCGCCCTGTTCTTGAGGTCTCTTTTCTCTACTCCCTGGTTC

TCAATTTATTTGGGGGG

>GR463724.1 JuteSSHPC29 JuteSSHlibMP1 Corchorus capsularis cDNA clone JuteSSHGP3 similar to Glutathione peroxidase (Stretch 4-34 amino acid) like protein, mRNA sequence

TCGAGCGGCCGCCCGGGCAGGTACCTTATTCCCTTCCTGGATAATTATGTAGCAAAACCAAGTTGGGTGT

GAACAGATTCCGCTGCAGAGGCTATAGATCTTATCTCTCACCATCTGCACGCACTAAAAAACTCACTCCG

CGTGCAATTGCAGCACTTAAGTATCTTGGTCAGTGTGAACCAGCTATGATTTCTACTTCTCTTAGCATGT

ATTTTTTTTTGGCATCAACCAAGATTTGGTGAAGAAAGGTTTTCTTTTTTGCTTTTACTTCTTTTGGTGC

CTGGTAAGTTGATCTGTGCAAACGCTTTAAAGTTTGCTAAATTAACAAATACTTGTTTAAAAATTCAGAT

AAAACAATTATATTCCAATAATGCAGATTACTACAATCATTTTAGAATTGTTGAAGGAGTCTGGGGAGGT

ATAAAAGGGCTTTGTAAAAAACGAGTTTGTTTTTCTTTAGGTGCATTGGCCAAAACATTAAACCAGCTTT

AGACTGTAGTTGCTACACTCCTTAGAGGGGTCATACCAACTGCAGTACCTCGGCCGCGACCACGCT

>GR463723.1 JuteSSHRP33 JuteSSHlibMP1 Corchorus capsularis cDNA clone JuteSSHHsp903 similar to hsp90 protein family (Stretch 13-112) like protein, mRNA sequence

GCGTGGTCGCGGCCGAGGTACGCTTGCTTGCTAGCATCTGATAGCGTTTGAGATTGCATGATTCTCTCCA

TATTAGCACTCCATCCGAACTTAGATGTAACAACCACACAAGGAGTGTCGTCCAAACGGTTGCTTATCTT

TACATCATCGACATTCTCACCGGCAAGTGAACCCTTCCACCACTTGGTTAGTTCCTTGAATGATTCCTTG

AGTTCTTTGTTCTTTGAGTCTTTCCCAATTTTCAGACCCTCCTTGGATACATTTTGGAACTGCTTTCCTT

CATAATCCATCAGGTATTGCATCAGGTACCTGCCCGGGCGGCCGCTCG

>GR463722.1 JuteSSHPS1 JuteSSHlibMP1 Corchorus capsularis cDNA clone JuteSSHSAP10 similar to senescence-associated protein like protein, mRNA sequence

AGCGTGGTCGCGGCCGAGGTACGAATACGAACCGTGAAAGCGTGGCCTATCGATCCTTTAGACCTTTGGA

ATTTGAAGCTAGAGGTGTCAGAAAAGTTACCACAGGGATAACTGGCTTGTGGCAGCCAAGCGTTCACAGC

GACGTTGCTTTTTGATCCTTCGATGTCGGCTCTTCCTATCATTGTGAAGCAGAATTCACCAAGTGTTGGA

TTGTTCACCCACCAATAGGGAACGTGAGCTGGGTTTAGACCGTCGTGAGACAGGTTAGTTTTACCCTACT

GATGGCCGCGTCGCAATAGTAATTCAACCTAGTACCTGCCCGGGCGGCCGCTCGA

>GR463721.1 JuteSSHPS2 JuteSSHlibMP1 Corchorus capsularis cDNA clone JuteSSHCHP similar to conserved hypothetical protein [Arabidopsis thaliana] like protein, mRNA sequence

AGCGTGGTCGCGGCCGAGGTACCTTCTCTGAGTTCAGTATAGCTTCTTTCTTGTCCCTCAATTTCAAATT

TTATTTTCCTGCAGGGCCATGAGGGCAGGTTTTCTGTTTTTGTACATGCTTCCAAAGAAAGGCCAGTGCA

CACTAGCCGATATTTTGTCGGTCGAGACATCCACAGTGAAAAGGTGGTTTGGGGAAAAATTTCCATGGTT

GATGCTGAAAGGAGACTTTTGGCGCATGCATTGGTAGACCCTGATAACCTGCAGTTTGTGTTGCTGTCAG

ATAGTTGAGTACCTGCCCGGGCGGCCGCTCGA

>GR463720.1 JuteSSHPS3 JuteSSHlibMP1 Corchorus capsularis cDNA clone JuteSSHPPI9 similar to peptidylprolyl isomerase[Arabidopsis thaliana] like protein, mRNA sequence

AGCGTGGTCGCGGCCGAGGTACTGTTTTCGTGAAGAAGGGTTACAACGAAGGCGAAGAACCATTTGAGTT

CATTATAGATGAAGAGCAAGTCTGTGATGGTCTTGATAGAACTGCGATGACGATGAAAAAGGGTGAAGTT

GCAGAGCTGACTATTGCACCTGAGTATGCTTTTGGGTCGTCTGGATCACAGCAGGAGTTGGCTTTTGTGC

CTGCTGACTCAACTGTATTCTACGAAGTTGAGATGGTATCTTTCACCAAGGAAAAAGAATCATGGGACAT

GAACACCAAAGAAAAAATTGAAGCGGCTGGTAAAAAGAAAGAAGAAGGAAATGCATTGTTTAAGGTTGGC

AAGTATGTGAGAGCTTCTAAGAGATATGAGAAGGCTGTTAAGTACCTGCCCGGGCGGCCGCTCGA

>GR463719.1 JuteSSHPS4 JuteSSHlibMP1 Corchorus capsularis cDNA clone JuteSSHECT2I similar to ECT2 [Arabidopsis thaliana] like protein, mRNA sequence

AGCGTGGTCGCGGCCGAGGTCTTCAGCTTTTCTTCAGATTTTGGTGATTGATTGTTGGATTCGGCAATCC

CATGGCATCCAAAAGGATCCTCAAAGAGATCAAGCTTGAGCAAGGACTGAAGTTGATCAAAATTTTCAAG

GAACATTCCAGCAAAACTTGCATTTTGGATGATTTTGAGTTTTATGAGGTCCGCCAGAAGGCAATCCAGG

ATAAAAAGGCCAAACAGCAATTTCAAAAGCAGGTATGGGAAGGAAAGCCTGTAGATGAGAAGAAAGATGT

AGCAAATGGAACACAGAACTGTTTGGAAGCTGGATCAGATTTGGTTAAGGAACCAAGCTTAACTGATAAT

TCCAATGGGGATCTAAAGCTTTCAGAGAATGGTTCAGCGAATCCAATTGTAGTTCCAGACAAGAAGGTTT

TATCGAATGGGATTGCCAATGGTTGCTAGTTCCTGCCAATCCCCTTGCGACGGTTCATGGTCTTGTAGAA

TATGAAGAGGAGTTTCCAGAAATTTTGATCTGATGATTTAGAATTTTAGATCACAGTACCTGCCCGGGCG

GCCGCTCGA

>GR463718.1 JuteSSHPS5 JuteSSHlibMP1 Corchorus capsularis cDNA clone JuteSSHCRCK3 similar to CRCK3 (calmodulin-binding receptor-like cytoplasmic kinase 3); kinase [Arabidopsis thaliana] like protein, mRNA sequence

TCGAGCGGCCGCCCGGGCAGGTACTCATTCGGGATTTTACTCATAGAAATTCTGACTGGCCGCCGTCCTG

TGGAGTTAAGGAGACCTGTTGAAGAGCGAGTAACACTGCGATGGGCATTCCATAAGTATAATGATGGACA

TGCAGTGCAATTGGTGGATCCTGCCATGGAAGAAGCTGTAGATGCAGAGGTACTGGTCAAAATATTTGCT

TTGGCATTTCAATGTGCAGCACCCGTAAGAAATGACCGACCGGACATGAAATCTGTGGGAGAACAGTTAT

GGGCGATAAGAGCTGACTACCTTAAGAGTTCAAGGAAAGTGTAGAATGATTGATTGTAGTCTCTCCCCTT

GACTCAATTGTTTTTTATCCCCTTTTCTTTCTTATATAAATTTCCTCTGCTTCTTGCTGTTGCTCACTTT

CTTTTCTCTTGTTCCCTTTTATTTTTCTCTCATTTCAAGCTTATTGTAAAAAGCAAAATTGGTCTTAACA

AAAATATGGTGCATTTTTGCCTTGTGCTCCTGTTTCATGTCTGTAATAGTTTCATTCTGGCAAAGCATCA

AAAGTACCTCGGCCGCGACCACGCT

>GR463717.1 JuteSSHPS6 JuteSSHlibMP1 Corchorus capsularis cDNA clone JuteSSHYCF2 similar to Ycf2 [Gossypium hirsutum] like protein, mRNA sequence

AGCGTGGTCGCGGCCGAGGTACAACAGCCCTTCCTATCTGATAGAAAAAGATCCCATGATCCTGAACCGA

TCTTACCTGGGATCGCAAATCCCAAGTTTGTCTATGAAGAGCAGATCTAATTGTATTAGTGTCTATAATT

GATTTCTTCTGTGTAATACTAATCGATAGGGCCTCATTGGTAAGTGCTACAAGATCTCGTGCATTGGAAC

CCACGGTTATGGACCCGAATCCATTAGTGTGGAACATTTTCTTTTCCAAGTGAAATCCCCTAGTATATGA

AAGAGTGAAAAAGTGCTTTCGTTGTTGTGGAATAAGAAGCCTTCGTATCTTAATACATGTATTTAATTTA

TTCGGGGCTATTAGAGTGGGATCCACTTTTTTGGGAATATGAGTCGAAGCAATAACAAGAATATTTCTAG

TGGAACATCTTTCACAATCCTTGGAGAGATAGTTCACTAATAGACCGAGGGAAAAGTACCTGCCCGGGCG

GCCGCTCGA

>GR463716.1 JuteSSHRP12 JuteSSHlibMP1 Corchorus capsularis cDNA clone JuteSSHGP2 similar to Glutathione peroxidase (Stretch 4-34 amino acid) like protein, mRNA sequence

GCGTGGTCGCGGCCGAGGTCGAAATCTTAGTTCAGTTGCTATTTCTGCTTCTCTTCTTTTTGCAAAGCGT

TTTTCACCGAGTTCCAAGCCAACCCTTTCCAGTTTTCCGCAGATTTCGCCAGTTTCTTTGGTTTCATCCT

CGAAGGAAACAGGGTTTTCAAGGTCTTTCTTGGGTAGTTTAAGATTTGATCATTCCATGGCTAGCCAATC

AGCCCAGACTTCAGTTCATGACTTCACCGTTAAGGATGCAAGAGGAAAGGACGTTGATCTAAGCACTTAC

AAAGGCAAGGTCCTCTTGATTGTCAATGTTGCCTCACAATGTGGCTTGACAAATTCCAACTACACTGAGC

TGAGTCAATTGTATGAGAAATATAAAGATCAAGGTTTTGAGATTCTGGCATTTCCTTGTAACCAGTTTGG

AGGTCAGGAGCCAGGGAACAATGAACAGATCTTAGAATTTGCTTGTACCTGCCCGGGCGGCCGCTCG

>GR463715.1 JuteSSHPS8 JuteSSHlibMP1 Corchorus capsularis cDNA clone JuteSSHSAP3 similar to senescence-associated protein like protein, mRNA sequence

TCGAGCGGCCGCCCGGGCAGGTACGAATACGAACCGTGAAAGCGTGGCCTATCGATCCTTTAGACCTTCG

GAATTTGAAGCTAGAGGTGTCAGAAAAGTTACCACAGGGATAACTGGCTTGTGGCAGCCAAGCGTTCATA

GCGACGTTGCTTTTTGATCCTTCGATGTCGGCTCTTCCTATCATTGTGAAGCAGAATTCACCAAGTGTTG

GATTGTTCACCCACCAATAGGGAACGTGAGCTGGGATTAGACCGTCGTGAGACAGGTTAGTTTTACCCTA

CTGATGGCCGCGTCGCAATAGTAATTCAACCTAGTACCTGCCCGGGCGGCCGCTCGACGTGGTCGCGGCC

GAGGTGGACGATAGGGTGCTTCAGATCAGCGGAGAGAGGAACGTTGAGAAGGAAGACAAGAACGACACCT

GGCACCGCGTTGAGCGTAGCAGCGGCAAATTCATGAGGAGGTTCAGGTTGCCGGAGAATGTGAAGATGGA

TCAAGTTAAGGCTTCAATGGAAAATGGGGTTCTTACTGTTACTGTTCCTAAGGTGGAAGTGAAGAAACCT

GATGTCAAATTCATTGAGATTCTGGGTTAAAATCTTCAAACACTTCATTTGCTCATTAGCTAAACTATAT

AGTGCTCTGTCTGCCTGGGTCCTGTAAGAAAGTTTGAACAATCATCATGTGTACCTGCCCGGGGGGCCGC

TCGA

>GR463714.1 JuteSSHPS9 JuteSSHlibMP1 Corchorus capsularis cDNA clone JuteSSHXET13 similar to Xyloglucan endotransglucosylase like protein, mRNA sequence

CGCAGCGGCCGCCCGGGCAGGTGGAAATTGTGTTTTTGGTTGATGATACGCCAATTAGAGTTTACAAGAA

CAATGAAGCAAAAGGTATCCCATACCCAAAATCCCAACCCATGGGAGTCTATTCAACACTTTGGGAAGCC

GATGATTGGGCGACAAGGGGTGGATTAGAGAAAATTGATTGGAGCAAAGCTCCTTTCTTAGCTTATTACA

AGGATTTCGACATCGAAGGTTGTGCTGTTCCTGGACCAGCTAATTGTGCCTCAAACCCTAGGAATTGGTG

GGAAGGCACTGCTTATCAAGGCCTTAATGCCTTGGAGGCTAGAAGATACAGGTGGGTTCGTATGAACCAT

ATGATCTATGATTGTTGCACCGACAAGTCCCGGTACCTGCCCGGGCGGCCGCTCGA

>GR463713.1 JuteSSHPS10 JuteSSHlibMP1 Corchorus capsularis cDNA clone JuteSSHBIP9 similar to BiP [Glycine max] like protein, mRNA sequence

AGCGTGGTCGCGGCCGAGGTACTTTATTAAGTTGATTAAGAAGAAGCATGAGAAAGGATATCAGCAAGGA

CAACAGGGCTATTGAGAAGCTTAGGAGGGAAGCTGAGCGAGCCAAGAGAGCTCTCAGCAGTCAGCACCAA

GTCCGTGTGGAGATTGAATCACTTTTCGATGGTATTGATTTCTCTGAGCCACTGACCAGGGCTCGCTTTG

AGGAGTTGAACAATGACCTCTTCAGAAAGACCATGGGACCTGTGAAGAAAGCAATGGATGATGCTGGTTT

GGAGAAGCACCAGATTGATGAGATTGTTCTAGTTGGTGGAAGTACCTGCCCGGGCGGCCGCTCGA

>GR463712.1 JuteSSHPS11 JuteSSHlibMP1 Corchorus capsularis cDNA clone JuteSSHPL9 similar to pectinacetylesterase like protein, mRNA sequence

TCGAGCGGCCGCCCGGGCAGGTACGTGGTGCAGACTATGCAAACGCCAATCTTTTTCATAAATGCAGCTT

ATGATTCCTGGCAGATTAAGAACGTTTTGGCTCCCACTGCTGTTGATTCTGGGAAGGCGTGGAAAAGCTG

CAAGCTTGATCTGACGAAGTGTACCTCGGCCGCGACCACGCTTACTTGGATCATGGTATAAGCAAGAGTT

GACAGAAGCTAATGCTCCTGGGAAAGCTGATGCTTTCACAATCAACGGCCATCCTGGAGAAACATATGGC

TGCAGCAACGATTCAACATTCGAGCAGCAGGTAGATTACCAAAGTGCTTATCACCTTCGAATTGTAAATG

CGGCTTTGAATGAGACAATGGAGTTTGGCGTGGAATACCACATGTTCACAGTTGTGGGACAAGATGGGGC

TTACACCACAAGGTCTTTCGCCGATTCTCTCACCATTGCTCCTGGCCGTACCTGCCCGGGCGGCCGCTCG

AATCTTTCTAGAAGATCTCCTACAATATTCTCAGCTGCCATGGAAAATCGATGTTACTTCTTTTA

>GR463711.1 JuteSSHPS12 JuteSSHlibMP1 Corchorus capsularis cDNA clone JuteSSHSHMT9 similar to serine hydroxymethyltransferaselike protein, mRNA sequence

AGCGTGGTCGCGGCCGAGGTGGAGATGTGTCTGCTATGGTTCCTGGTGGTATCAGGATGGGAACACCTGC

TCTTACATCTAGGGGATTTGTTGAGGAGGACTTTGTTAAGGTAGCAGAATACTTTGATGCTGCTGTGAAG

TTAGCATTGAAAGTCAAGGCTGAAGCACCAGGCACAAAGTTGAAAGGTTTCGTTGCAACATTGCCCAAGT

TTCAATCTGAGATTGCAAAACTCCGCCACGATGTTGAGGAGTACCTGCCCGGGCGGCCGCTCGA

>GR463710.1 JuteSSHPS13 JuteSSHlibMP1 Corchorus capsularis cDNA clone JuteSSHACCO9 similar to 1-aminocyclopropane-1-carboxylate oxidase like protein, mRNA sequence

AGCGTGGTCGCGGCCGAGGTGTGTGGAACATAGGGTTATTGCTCAAACTGACGGAGCTCGCATGTCAATT

GCTTCATTCTACAATCCTGGAAGTGAGGCTGTAATTTACCCTGCACCATCTCTGATTGAGAAAGAAGCAG

AGGAGAAGAAGCAAATTTACCCCAGATTTGTGTTTGAAGATTACATGAAGCTGTATGCAGCTCTGAAATT

CCAGGCCAAGGAACCAAGATTTGAAGCCATGAAAGCCGTGGAAACAACTGTCCCAGTTGCAACAGTTTAA

AGTTTGTTCTAGAGCTTTGACTAGTGATGATGAGAAAAAAAAAGGGCAGTGGCAATGGCATGCTATCTAA

TAGTATAGAAGTACCTGCCCGGGCGGCCGCTCGA

>GR463709.1 JuteSSHPS15 JuteSSHlibMP1 Corchorus capsularis cDNA clone JuteSSHHSP708 similar to heat shock protein 70 like protein, mRNA sequence

AGCGTGGTCGCGGCCGAGGTACTTTATTAAGTTGATTAAGAAGAAGCATGGAAAGGATATCAGCAAGGAC

AACAGGGCTATTGGGAAGCTTAGGAGGGAAGCTGAGCGAGCCAAGAGAGCTCTCAGCAGTCAGCACCAAG

TCCGTGTGGAGATTGAATCACTTTTCGATGGTATTGATTTCTCTGAGCCACTGACCAGGGCTCGCTTTGA

GGAGTTGAACAATGACCTCTTCAGAAAGACCATGGGACCTGTGAAGAAAGCAATGGATGATGCTGGTTTG

GAGAAGCACCAGATTGATGAGATTGTTCTAGTTGGTGGAAGTACCTGCCCGGGCGGCCGCTCGA

>GR463708.1 JuteSSHPS14 JuteSSHlibMP1 Corchorus capsularis cDNA clone JuteSSHCSP similar to chromosome segregation protein SMC like protein, mRNA sequence

TCGAGCGGCCGCCCGGGCAGGTTTTTTTTTTTTGTGAATCACCTCCAGAAGTAGAAGACAATCTATTCAT

TATAATAGTTTCCATATGATACGAACGAAACATCATAATATCGTTTTTACTGGAAAAGGAGAATCATAAA

TTCAAAAGCGGGGAGTTTTCATCTAACCAATACAAAGGAATAAGTAACAAAAGATGGATATTAGTAAATC

ATCTCAATCCAGCATCTAATACAGGAGATATATTTACATGATCAAGCAAACAAAATGGATCAAAGTTTTG

CTGGAAAGAGACAAAACAACTTGTTAATATTTGCAATAGGTGAACCCAAGCATTACCAGCAATAAGCTCA

CATCCAAGGTCAACCAACTTCATGTTTAATTTCCTAGCATTGAAAATATAGAGAATAAATAAACATAAGA

TAAAGATGAAAGATAAATTCTTGAATAAGAGGATGAGTAAACTGACAAAATGTATGTGAAAAAGTAATGA

GAACCGTACCTCGGCCGCGACCACGCT

>GR463707.1 JuteSSHPS16 JuteSSHlibMP1 Corchorus capsularis cDNA clone JuteSSHAC09 similar to acyl-CoA oxidaselike protein, mRNA sequence

CCGGCGCGCAAGGCAGCTACTTATGTTTGAACAATGTAGTGCTGCGTTTAACGTTGCAGGATTAAATGAA

CACCGTTTCCCAGCTGGGGTCTGGAAAGAAACCTGTTGGGACAACAGCCTATATGGGACGAGCAGAGCAC

CTCATGCAATGCAGTTGTGATGTTCAAAGGGCTGATGATTGCCTAACCCCAGTGTAATAGTGGAGGCA

>GR463706.1 JuteSSHPS17 JuteSSHlibMP1 Corchorus capsularis cDNA clone JuteSSHEFP9 similar to exonuclease family protein like protein, mRNA sequence

AGCGTGGTCGCGGCCGAGGTACATACACCCCTCATATAGCTGGAAATTCAGTTTACATGGATTTTCTATT

TCTAAAGAAATATATGCCACATTTGGCTAGTCTTTTCTCTCATATAGTCGTTGATGTAAGCAGTGTGAGG

GCTCTTTGCATCCGCTGGTATCCTAAAGATCAAAAGAAAGCCCCTGCCAAAGAAAAGAAACACAGAGCCA

TGGATGATATCAGGGAGAGCATATTAGAACTCAAATACTTCAAGGAGACGATATTTAAAGGGAAGTCCAA

AAGGTGAATGCCAATGCCCAATTGCTTTGGGAGACAGTGATGGTCATTTTTTAGAACAATTAGGTCACTG

GACAAGCTGTTTGCTGATTTGGTACCTGCCCGGGCGGCCGCTCGA

>GR463705.1 JuteSSHRP54 JuteSSHlibMP1 Corchorus capsularis cDNA clone JuteSSHGP1 similar to Hsp20/alpha crystallin family like protein, mRNA sequence

GTGGTCGCGGCCGAGGTACATATAATTCATTCATACAAGTAAAAGTATCATAAATTCATATTAAGACATG

AAAATATAAATAGGCATAGGAAGCAAATGAATGTAATTAAAGTCGGCTGGTGATGTTAATATTTCGCAGT

TTAGATGGGTTGGGCGTAGTATCTTTTGGAACAAGAAGCTTGAGCACACCATTTTCCAGGTGGGCCTTGA

TCTGCTCAACCTTGACATTTGGTGGCAATTCTATATGGCGAGAGAATTTGAATTCACCCAATTTCCCGGT

GCCCCTCTCGGCTACATGCCAAACACTGTCTTTTGCATTGGCGTAGCTCTCGTCTTTACCCTCTCCTTTT

ATATGCAAAATGTTTCCATCTTCTATCTGCACTTTTATGTCCTCTTTGCCGTATCCTGGGACGTTGAGCT

TGAAGATGTGGGCAGTTGGGGATTCTACCCAATCCATGAGAGCCGTTGATGATCCAGTTCACCTGCCCAG

GCGGCCGCTCG

>GR463704.1 JuteSSHPS18 JuteSSHlibMP1 Corchorus capsularis cDNA clone JuteSSHGT9 similar to glutathione transferase like protein, mRNA sequence

CGAGCGGCCGCCCGGGCAGGTACACCGTTCCTGTCCTTTGGGACAAGAAATTCAAGACTATTGTTAGTAA

TGAGAGCGCAGAGATAATCCGCATGCTTAATACTGAATTCAATGATATAGCAGAGAATCCTGCTTTGGAC

CTATATCCTCCTCATTTGAGAGCTCAAATTGATGAGACAAATGAATGGATATACAGTGGAATCAATAATG

GTGTTTACAGATGCGGCTTTGCTAGAAAGCAAGGGCCTTATGATGAGGCTGTGAAACAATTATATGAAAC

CATGGACAAATGTGAGGAGATTCTTTCCAAGCAGCGGTACCTCGGCCGCGACCACGCT

>GR463703.1 JuteSSHPS20 JuteSSHlibMP1 Corchorus capsularis cDNA clone JuteSSHECT10 similar to ECT2 [Arabidopsis thaliana] like protein, mRNA sequence

AGCGTGGTCGCGGCCGAGGTACAAACAATTCTTTGAAGCACATCACCTTAGAGAACAATGAGAATAAACC

TGTCACGAATAGCAGGGATACACAAGAGATCAAGCTTGAGCAAGGACTGAAGTTGATCAAAATTTTCAAG

GAACATTCCAGCAAAACTTGCATTTTGGATGATTTTGAGTTTTATGAGGTCCGCCAGAAGGCAATCCAGG

ATAAAAAGGCTAAACAGCAATTTCAAAAGCAGGTATGGGAAGGAAAGCCTGTAGATGAGAAGAAAGATGT

AGCAAATGGAACACAGAACTGTTTGGAAGCTGGATCAGATTTGGTTAAGGAACCAAGCTTAACTGATAAT

TCCAATGGGGATCTAAAGCTTTCAGAGAATGGTTCAGCGAATCCAATTGTAGTTCCAGACAAGAAGGTTT

TATCGAATGGGATTGCCAATGGTTGCTAGTTCCTGCCAATCCCCTTGCGACGGTTCATGGTCTTGTAGAA

TATGAAGAGGAGTTTCCAGAAATTTTGATCTGATGATTTAGAATTTTAGATCACAGTACCTGCCCGGGCG

GCCGCTCGA

>GR463702.1 JuteSSHPS21 JuteSSHlibMP1 Corchorus capsularis cDNA clone JuteSSHRL6T similar to ribosomal protein L16 like protein, mRNA sequence

TCGAGCGGCCGCCCGGGCAGGTACGTATATTTCCAGACAAACCAGTTACGGTAAGACCTACAGAAACACG

TATGGGTTCGGGGAAAGGATCTCCCGAGTATTGGGTAGCTGTTGTTAAACCGGGCAGAATACTTTATGAA

ATGGGCGGAGTAGCCGAAAATATAGCCAGAAAAGCTATTTCAATAGCGGCGTCTAAATGCCTATAAAAAC

TCAATTCATTATTTCAGGATAGGAATATAGAACCAAAGGAAAGAAGTCTTGGGAATAAAAAAAACAATTG

CGGGTTTCCTTTTTTTGACAAACAATATTTCTTTTTTTCTCCGTCCTTTGTTGCATTGAAAGAAGAGATT

AAAAAAATGATTCAACCTCAGACCCATTTGAATGTAGCAGATAATAGCGGGGCCCGAGAATTGATGTGTA

TTCGAGTCATAGGAGCTAGTAATCGCCGATATGCTCATATTGGTGACGTTATTGTTGCTGTGATCAAGGA

AGCAGTACCTCGGCCGCGACCACGCT

>GR463701.1 JuteSSHPS22 JuteSSHlibMP1 Corchorus capsularis cDNA clone JuteSSHXET12 similar to Xyloglucan endotransglucosylase like protein, mRNA sequence

TCGAGCGGCCGCCCGGGCAGGTAAGATTTCCATGACTACCAAATTAGATGGAACCATAAGGAAATTGTGT

TTTTGGTTGATGATACGCCAATTGGAGTTTACAAGAACAATGAAGCAAAAGGTATCCCATACCCAAAATC

CCAACCCATGGGAGTCTATTCAACACTTTGGGAAGCCGATGATTGGGCGACAAGGGGTGGATTAGAGAAA

ATTGATTGGAGCAAAGCTCCTTTCTTAGCTTATTACAAGGATTTCGACATCGAAGGTTGTGCTGTTCCTG

GACCAGCTAATTGTGCCTCAAACCCTAGGAATTGGTGGGAAGGCACTGCTTATCAAGGCCTTAATGCCTT

GGAGGCTAGAAGATACAGGTGGGTTCGTATGAACCATATGATCTATGATTATTGCACCGACAAGTCCCGG

TACCTCGGCCGCGACCACGCT

>GR463700.1 JuteSSHPS23 JuteSSHlibMP1 Corchorus capsularis cDNA clone JuteSSHPITP8 similar to phosphatidylinositol transfer-like protein III like protein, mRNA sequence

GGCGTGGTCGCGGCCGAGGTATTAGAGCTAGGATTTTGGTGACCGTTATGGCTTTCTTAATGACACTTTA

CACCCTGTTCCGCTCATTTGCATGCCATATAACAAGGAAGCTTCCCAACACAGTGTCTGACCATTCTCAA

AATGCTACTCAACCCACCCGTGCTGCAAATCAAGAGGAAGAGTTTCACCCCCCTTCTCCAACTCCTCCTT

TTAGTCAGTCAGATCTTCTGTCTACTGTGCTCAAGAGGTTGGGTGAGCTTGAGGAGAAAGTTGATACACT

ACAAGCAAAGCCATCCGAGATGCCTTATGAGAAAGAGGAATTGTTGAATGCTGCTGTATGCCGTGTTGAT

GCCCTAGAAGCAGAGCTAATCGCTACTAAGAAGGCTCTTCATGAAGCTTTGAGGAGGCAAGAAGAACTGC

TTGCTTACATAGACAGCCAAGAGGAAGCCAAACTGCGGAAAAAGAAGTTTTGCTGGTGATATGCTTTGCC

GATGGAGAGACTACATTTGAGAGTACTGCCCGGGCGGCCGCTCGA

>GR463699.1 JuteSSHPS24 JuteSSHlibMP1 Corchorus capsularis cDNA clone JuteSSHSHD1 similar to SHEPHERD [Arabidopsis thaliana] like protein, mRNA sequence

TCGAGCGGCCGCCCGGGCAGGTACCTGATGCAATACCTGATGGATTATGAAGGAAAGCAGTTCCAAAATG

TATCCAAGGAGGGTCTGAAAATTGGGAAAGACTCAAAGAACAAAGAACTCAAGGAATCATTCAAGGAACT

AACCAAGTGGTGGAAGGGTTCACTTGCCGGTGAGAATGTCGATGATGTAAAGATAAGCAACCGTTTGGAC

AACACTCCTTGTGTGGTTGTTACATCTAAGTTCGGATGGAGTGCTAATATGGAGAGAATCATGCAATCTC

AAACGCTATCAGATGCTAGCAAGCAAGCGTACCTCGGCCGCGACCACGCT

>GR463698.1 JuteSSHPS25 JuteSSHlibMP1 Corchorus capsularis cDNA clone JuteSSHLIP3 similar to temperature-induced lipocalin like protein, mRNA sequence

AGCGTGGTCGCGGCCGAGGTACGAGATTGCTTCATTCCCATCAAGGTTTCAGCCTAAGGATGGGAAGGAC

ACCAGGGCCACCTATACCTTGAATGAAGATGGAACCGTTAAAGTTCTTAATGAGACTTTCGTCGATGGGA

AACGTTCCTATATTGAAGGCACTGCTTACAAGGCCGATCCCAAGAGTGATGAGGCAAAGCTCAAAGTGAA

ATTCTATGCTCCTCCATTTTTGCCCATCATTCCTGTTGTTGGGGATTACTGGGTTTTGTACCTGCCCGGG

CGGCCGCTCGA

>GR463697.1 JuteSSHPS26 JuteSSHlibMP1 Corchorus capsularis cDNA clone JuteSSHHPA1 similar to histidinol-phosphate aminotransferase like protein, mRNA sequence

AGCGTGGTCGCGGCCGAGGTCATTGCTGAGGTTGTTGAGCGAGAAAAACCTAAATGCATATTTCTAACTT

CTCCAAATAATCCAGACGGAAGTATAATTAGTGAAGAAGTTCTCTTGAAGATCCTTGACATGCCCATATT

GGTGGTTCTGGATGAAGCTTATATCGAGTTTTCAGGAGTTGAATCTAAAATGCAATGGGTTAAAAAGCAT

GAAAATCTAATTGTTCTTCGAACATTTAGCAAAAGAGCTGGTTTAGCTGGACTCCGTGTTGGATATGGAG

CATTCCCTTTGAGCATAATTGAGTATCTGTGGAGAGCAAAGCAACCATACAACGTTTCTGTTGCTGCTGA

AGTTGCTGCATGTGCTGCATTGCAGAACCCAAAGTATCTAGAGATGGTGAAAGAGTCTCTGGTACCTGCC

CGGGCGGCCGCTCGA

>GR463696.1 JuteSSHPS27 JuteSSHlibMP1 Corchorus capsularis cDNA clone JuteSSHRPLQ similar to ribosomal protein L16 like protein, mRNA sequence

TCGAGCGGCCGCCCGGGCAGGTACGTATATTTCCAGACAAACCAGTTACGGTAAGACCTACAGAAACACG

TATGGGTTCGGGGAAAGGATCTCCCGAGTATTGGGTAGCTGTTGTTAAACCGGGCAGAATACTTTATGAA

ATGGGCGGAGTAGCCGAAAATATAGCCAGAAAAGCTATTTCAATAGCGGCGTCAAAATGCCTATAAAAAC

TCAATTCATTATTTCAGGATAGGAATATAGAACCAAAGGAAAGAAGTCTTGGGAATAAAAAAAACAATTG

CGGGTTTCCTTTTTTTTGACAAACAATATTTCTTTTTTTTCTCCGTCCTTTGTTGCATTGAAAGAAGAGA

TTAAAAAAATGATTCAACCTCAGACCCATTTGAATGTAGCAGATAATAGCGGGGCCCGAGAATTGATGTG

TATTCGAGTCATAGGAGCTAGTAATCGCCGATATGCTCATATTGGTGACGTTATTGTTGCTGTGATCAAG

GAAGCAGTACCTCGGCCGCGACCACGCT

>GR463695.1 JuteSSHPS29 JuteSSHlibMP1 Corchorus capsularis cDNA clone JuteSSHGP11 similar to glutathione peroxidase like protein, mRNA sequence

AGCGTGGTCGCGGCCGAGGTACTCGCTTTAAAGCTGAATATTCCATATTTGATAAGGTTGAGGTGAACGG

TGCTAACACCGCTCCCATATACAAATTCTTGAAGTCCAGCAAAGGCGGACTTTTCGGGGACAGCATCAAG

TGGAACTTCTCCAAGTTCTTGGTCGATAAGGATGGCAATGTTGTTGACCGCTATGCTCCCACAACTTCTC

CTCTTAGCATTGAGAAGGATATAAAGAAGTTGCTGGCCTGATTACTGGAGGCTACTTCCATCAGCTCGGA

GTGAATAAAGTAGATAAATTTCACGAATAAAAACTAGTGTTTCAGACATTTCTCTGCTTTTGTTCCCCAT

CTGATCTGCAAAGATGGGGTTGAACTGTATTGGAAATTAGAACTTGTATTGTACCTGCCCGGGCGGCCGC

TCGA

>GR463694.1 JuteSSHPC86 JuteSSHlibMP1 Corchorus capsularis cDNA clone JuteSSHL13AD similar to ribosomal protein L13 like protein, mRNA sequence

AGCGTGGTCGCGGCCGAGGTACTTGAATACTTCTGTTTCCCAGTGTTGCTTGCTGGACATTTTAGTGCCT

TGGACTTCTATTGCTTCTGCCATTAGCATCAACTTACCAGACCCCAGATCAATAAAGGGCATGTGGAAGG

AAATCGTAGGTCCATGTGACCCCAGCAGTCCAGCAGTGGTTATGCCAAAGGGAAATTGAAAAAGTATTTT

TTTAAGTCATTCAACAACTTTGTCTAGAGCAGGTGTAAGATGAGTAGGGTGGGAAGTTAGGTTGGCATCA

GTGGTTAAAAACAGAAAGTTCTGTTTCGGGAATAGTGAGGAGGGGGTGTTGTAACAAAATTGGACAACTT

AAAAGAATGGTGTGTGCTGGGTGAAAGACAAAGACTAAAGAATGAGGAAACAAATGTGATGCCTGGCCAG

TGACTGTCATATAAACCTTTCTTATTTGAGCTAGGCTTGAACAGACGTGACCTAGAAGAAACTGAACATA

AAGAGAAGGGGGTGGGGGGCTAGTTTTCAAGTTGGGGAACCTGATAGTGAAAAGTCACAGATGGAGAAAA

TTGCTCTCAGAAAAACTGTTTGGATTGCTTTCCTCTTGTTGCACATGTACCTGCCCGGGCGGCCGCTCGA

>GR463693.1 JuteSSHPC79 JuteSSHlibMP1 Corchorus capsularis cDNA clone JuteSSHDDPG6 similar to Dolichyl-diphosphooligosaccharide--protein glycosyltransferase subunit 1 precursor like protein, mRNA sequence

AGCGTGGTCGCGGCCGAGGTACTCATAGCTTGGGAGGTTGTAGCCAACGATGTAATGGGTCTTCCACCCG

CCAAAGAGAGGGAAGCGAGGCCGGATTTCCATCTCTACAGAGTCATCCAAAATAAGGAGGTGGCTGGTAG

AAACATTGCCAATCTCATCCCGGTAATAAACATCCTGGGCAGCAGCAGGAAGGGTGGTCTTAAAAGAACG

GATGGAGGATATTCCACTATCTGGCTGTCTCTGGTAATCATAGCGTGAGAAAGGCCCCTTAAGCACAGCT

CCTGTGTGCTTTAAGTCCACATTTTCTTCCACAGCAATATTACCCCAGTGAGAGACTTCAATGACTCGGG

TCATGCTGGTGATGGTCAGGAAAGGGCTGTTGTTCTCATAATGTACCTGCCCGGGCGGCCGCTCGA

>GR463692.1 JuteSSHPC63 JuteSSHlibMP1 Corchorus capsularis cDNA clone JuteSSHMRP2 similar to multidrug resistance protein like protein, mRNA sequence

AGCGTGGTCGCGGCCGAGGTGTAACAGAGACCAGGTTTATCCTCCCAGTTGAAACAGCTAAAAAACTGGA

ATGAATACATGAGACAAGAGCTCTTAGATGTTGGACATCAGGCAATAAAGGACACTGATCCTTGAGGATT

AGGAAACAAATGAGTTTAGCCCTACGGTCTCCCCAGCTGACTGCCTGGAGAGCGTTTCCAGACCACAGCA

CAGGGAAGGAAAACTGAGGTGGGACCAGGAGATTCCCTAAGTTGAAGAGACAGAGCTGGAAATCCAGGGA

AGCCATGGCAGGCAGGGTTCACGTTGCAGAGCACAGAGACACAGTTGAGAGATCACAGAGCAAAGCCCAG

AGGTATGCAGAGAATCCTTGAACATGCAGTTGAGTTCTGCTCAGAGAATACAGACAAGGAAACTACCCTA

GGGCTAGGAAAGAACCACCCAAAAGGAGAAGAGGGAATCATACCTGGAAACACACAGGGCCAGGAATAGT

GTCTATTCCTGACAGGTGAATAAACCTCAGAATTCACAGGGCATTGTTTAGCGTACCTGCCCGGGCGGCC

GCTCGA

>GR463691.1 JuteSSHPC49 JuteSSHlibMP1 Corchorus capsularis cDNA clone JuteSSHPTS similar to phenylalanyl-tRNA synthetase beta subunit like protein, mRNA sequence

TCGAGCGGCCGCCCGGGCAGGTACATAGAACGATAAGGAGCTGTATTCTACCTGCTACACATTGTGCATT

TTACACCTCCAGCTAGGACAGAAACAAGCCACAAGCGGGGAATACCCGCCCAAATTAATACTGGAAAGTT

TACAAAACCTTCACTTCTCATAAGGGTGATGCTACCAGCGGCTTGCTTCACCTGGAATTAGAGTAACTGA

TTTTTAAAGACAACTGAGAATATTAACTCGCTTCAAACAATAAAATAGTCATTTTGCCGACATTTCATAA

AATCGCCAGTTTTAAGCTCAGTTCAGAGGCTTTTAAAAAAAATATTCTAGACCTAGTAAAGTGCCTCCAT

TCATAAAATGAAACATCCAAGATTGAGACAATTTCATACTGATGCGCATCTGAAATAGGTCGACGTATAC

AGTAAATAAACAGCGCCTCTAAGTTGCCATATTTCAGTTAGTGGAATACACACCCAGCCTTCGCCGTTCT

GGATGTGCAAACTAGAAACTGAAGACAAATGGGGATTTCAGCGTCTAGTCCCAATCGTTCAGCACGGGAA

GTACCTCGGCCGCGACCACGCT

>GR463690.1 JuteSSHPC56 JuteSSHlibMP1 Corchorus capsularis cDNA clone JuteSSHMRP1 similar to multidrug resistance protein like protein, mRNA sequence

AGCGTGGTCGCGGCCGAGGTGTAACAGAGACCAGGTTTATCCTCCCAGTTGAAACAGCTAAAAAACTGGA

ATGAATACATGAGACAAGAGCTCTTAGATGTTGGACATCAGGCAATAAAGGACACTGATCCTTGAGGATT

AGGAAACAAATGAGTTTAGCCCTACGGTCTCCCCAGCTGACTGCCTGGAGAGCGTTTCCAGACCACAGCA

CAGGGAAGGAAAACTGAGGTGGGACCAGGAGATTCCCTAAGTTGAAGAGACAGAGCTGGAAATCCAGGGA

AGCCATGGCAGGCAGGGTTCACGTTGCAGAGCACAGAGACACAGTTGAGAGATCACAGAGCAAAGCCCAG

AGGTATGCAGAGAATCCTTGAACATGCAGTTGAGTTCTGCTCAGAGAATACAGACAAGGAAACTACCCTA

GGGCTAGGAAAGAACCACCCAAAAGGAGAAGAGGGAATCATACCTGGAAACACACAGGGCCAGGAATAGT

GTCTATTCCTGACAGGTGAATAAACCTCAGAATTCACAGGGCATTGTTTAGCGTACCTGCCCGGGCGGCC

GCTCGA

>GR463689.1 JuteSSHPC2 JuteSSHlibMP1 Corchorus capsularis cDNA clone JuteSSHIFO similar to indolepyruvate ferredoxin oxidoreductase, alpha subunit like protein, mRNA sequence

AGCGTGGTCGCGGCCGAGGTACATACGGGGAAACTGAAGCCAGGAGAAGAAGAGGGAGAAACTGTCCAGG

GCCTCACAGGGAGGCATTCATGGAGCTGGAGAAACCAGGGATCCTGACCGCAGGGCCCATCAATTGCTAC

ATAGAGATACACATGGAAAGAGGGTTTAACAGACCCTCTCTAGGACACAACTGTTTCTGCTTTGAGCAAT

AATTTCTAACCTGAGGCCAACATGTCCCACTGCCCCTTGGGTTGGCTGGGGTTGTTTCCTGAGCCAAGCA

TCCAATATCATGCCCCACTCAATGGCCTAGAAGCTGCCTGTACCTGCCCGGGCGGCCGCTCGA

>GR463688.1 JuteSSHRP5 JuteSSHlibMP1 Corchorus capsularis cDNA clone JuteSSHHsp702 similar to Hsp70 (Stretch 6-103 amino acid) like protein, mRNA sequence

CGAGCGGCCGCCCGGGCAGGTACTTCCACCAACTAGAACAATCTTATCAATCTGGTGCTTCTCCAAACCA

GCATCATCCATTGCTTTCTTCACAGGTCCCATGGTCTTTCTGAAGAGGTCATTGTTCAACTCCTCAAAGC

GAGCCCTGGTCAGTGGCTCAGAGAAATCAATACCATCGAAAAGTGATTCAATCTCCACACGGACTTGGTG

CTGACTGCTGAGAGCTCTCTTGGCTCGCTCAGCTTCCCTCCTAAGCTTCCCAATAGCCCTGTTGTCCTTG

CTGATATCCTTTCCATGCTTCTTCTTAATCAACTTAATAAAGTACCTCGGCCGCGACCAC

>GR463687.1 JuteSSHRP113 JuteSSHlibMP1 Corchorus capsularis cDNA clone JuteSSHSDDS2 similar to Dehydrodolichyl diphosphate synthase like protein, mRNA sequence

CGAGCGGCCGCCCGGGCAGGTACAAAGAAGAGCTCTGTATAGGCCAACTGCCACAGTAAAAAGTTACTGA

CTCTAAGTTCATTACTTGTTCTAATTAACAAATCCGGATATGGATACTCGGTGCAGTTTGTTTCTAGTTC

TTGTTCAATTAGGCTTTCGTTGATGTCTTCGATTTGGATTTCGCCGTCTTTTGCTTTCTCAGCAATGCTT

CTGCATGCTTGTACCTCGGCCGCGACCACGC

>GR463686.1 JuteSSHRP108 JuteSSHlibMP1 Corchorus capsularis cDNA clone JuteSSHSDR2 similar to plant disease resistance response protein family like protein, mRNA sequence

GCGTGGTCGCGGCCGAGGTACTTTGGTGGTGAACTAGTATTAATTAAATTATTATTAAAGCTGTGAGAAT

AAATAAAAGAATGTCCTATATATATAATATTTATGCAAAGCAAAAGTCATTGAGCTAATGACTTGTGTTT

TTCTTTTGTTTTCTAGCTATCTTTCTTGGCAATATTTGATTATTTGTAATGAGTTTGAAATATATATATG

TATGTAGTGAAAGAATAATGAAGTTTGTATTGTAATTATATATTGATCTCTTTAGCCTTTCCCTTGCCGA

AAAAAAAAAAAAAAAAAAAAAAAAAAGCTTGTACCTGCCCGGGCGGCCGCTCG

>GR463685.1 JuteSSHRP98 JuteSSHlibMP1 Corchorus capsularis cDNA clone JuteSSHSlac2 similar to Secretory laccase (Lac1)like protein, mRNA sequence

CGAGCGGCCGCCCGGGCAGGTACTTATGAACAAGTGATACCATCAATGTCCCAATATAGTTCCCTATCGC

TATTGTAATCCAGAACAATGCTGCAGCTGTGCTTCTTATGCTTTCTGGTGACTGATCATACAGAAATTCC

AAGTGCCCCACTGACATGAAAACTTCTGCTACCCCATGGAGGCAAAATTGAGGAACTAACCAGAACACGC

TGATGGGAATTACTGCTTTTGGATCATCCAACAAGTTATGATTTGCTGCAAACCTCGGCCGCGACCACGC

>GR463684.1 JuteSSHRP97 JuteSSHlibMP1 Corchorus capsularis cDNA clone JuteSSHSPOT similar to Arabidopsis thaliana proton-dependent oligopeptide transport (POT) family protein, mRNA sequence

CGAGCGGCCGCCCGGGCAGGTACTTATGAACAAGTGATACCATCAATGTCCCAATATAGTTCCCTATCGC

TATTGTAATCCAGAACAATGCTGCAGCTGTGCTTCTTATGCTTTCTGGTGACTGATCATACAGAAATTCC

AAGTGCCCCACTGACATGAAAACTTCTGCTACCCCATGGAGGCAAAATTGAGGAACTAACCAGAACACGC

TGATGGGAATTACTGCTTTTGGATCATCCAACAAGTTATGATTTGCTGCAAACCTCGGCCGCGACCACGC

>GR463683.1 JuteSSHRP92 JuteSSHlibMP1 Corchorus capsularis cDNA clone JuteSSHSAP1 similar to Senescence-associated protein, mRNA sequence

GCGTGGTCGCGGCCGAGGTCAAGCTTTTTTTTTTTTTTTTTTTTTTTTTTTTTTTTTCCATAAATTCATA

AAATGTTTTGATTTAAAAAAAATTTAACAGATACTACAAGAATTTTCCTAGAGATCTCAAGAACTATTCA

CATGATGGAAATTTAACGGGCGCCTTTCGATCACCTAACTGGTCAAAGATTGTAGCTAATCCGTCAAAAT

CTCACCGCCAAAATCATGCTACTCCAACACGTGCAAACTTCAAAGCCCCAATCTCATTGGTCAAGTCTCT

ACTCTGCAGTCTGCACATTTGCTGCCAAGCCAATCCATAGCCATTCAACCAAAGTACCTGCCCGGGCGGC

CGCTCG

>GR463682.1 JuteSSHRP89 JuteSSHlibMP1 Corchorus capsularis cDNA clone JuteSSHDR similar to plant disease resistance response protein family like protein, mRNA sequence

GCGTGGTCGCGGCCGAGGTCAAGCTTTTTTTTTTTTTTTTTTTTTTTTTTTTTTTTTCCATAAATTCATA

AAATGTTTTGATTTAAAAAAAATTTAACAGATACTACAAGAATTTTCCTAGAGATCTCAAGAACTATTCA

CATGATGGAAATTTAACGGGCGCCTTTCGATCACCTAACTGGTCAAAGATTGTAGCTAATCCGTCAAAAT

CTCACCGCCAAAATCATGCTACTCCAACACGTGCAAACTTCAAAGCCCCAATCTCATTGGTCAAGTCTCT

ACTCTGCAGTCTGCACATTTGCTGCCAAGCCAATCCATAGCCATTCAACCAAAGTACCTGCCCGGGCGGC

CGCTCG

>GR463681.1 JuteSSHRP87 JuteSSHlibMP1 Corchorus capsularis cDNA clone JuteSSHGT6 similar to glutathione S-transferase like protein, mRNA sequence

GCGTGGTCGCGGCCGAGGTCAGCAACATTACAATGGCTGCTAACAACTTTGAGAATTTGCCTCCTCCCTT

GGATGCTACTGCTGAGCAACCCCCTCTGTTTGATGGAACTACCAGGTTGTATACTTCTTATTCTTGCCCC

TTTGCCCAGCGAGTGTGGATCACAAGGAATTATAAGGGGTTGCAAGACAAGATCAAATTGGTTCCTTTGA

ACCTTCCGAACAGACCTGCTTGGTACCTGCCCGGGCGGCCGCTCG

>GR463680.1 JuteSSHRP86 JuteSSHlibMP1 Corchorus capsularis cDNA clone JuteSSHL3 similar to ribosomal protein L14 like protein, mRNA sequence

CGAGCGGCCGCCCGGGCAGGTACTAGGGTTTATGCTAATCGATTGATATTGTGATTGTGCTTGAATTGAG

TTTTTGTGCAGCAACTTTGGAGAAGAATTTCGACATAAGGAACCTGCATTAAGTGGTTTGGTTTGGTTTA

AGATCACTTTTCCTTTGATTCACAAATAACTTTGCTACTAATTGTTTGTTTCTCAAGTAGGTCACGAGAA

GAAAGGAGTGCTCTAGGAGTTGCTCTTATTAGGCCTATTGAAGCTAGTAAAAGGCAAATCTTGTGAGATC

ATTAAGAGAGTAGTGGACCGCCAATCCATAGGACTTGAGTACCTCGGCCGCGACCACG

>GR463679.1 JuteSSHRP75 JuteSSHlibMP1 Corchorus capsularis cDNA clone JuteSSHPS2 similar to pseudouridine synthase like protein, mRNA sequence

CGAGCGGCCGCCCGGGCAGGTACTAGGGTTTATGCTAATCGATTGATATTGTGATTGTGCTTGAATTGAG

TTTTTGTGCAGCAACTTTGGAGAAGAATTTCGACATAAGGAACCTGCATTAAGTGGTTTGGTTTGGTTTA

AGATCACTTTTCCTTTGATTCACAAATAACTTTGCTACTAATTGTTTGTTTCTCAAGTAGGTCACGAGAA

GAAAGGAGTGCTCTAGGAGTTGCTCTTATTAGGCCTATTGAAGCTAGTAAAAGGCAAATCTTGTGAGATC

ATTAAGAGAGTAGTGGACCGCCAATCCATAGGACTTGAGTACCTCGGCCGCGACCACG

>GR463678.1 JuteSSHPC15 JuteSSHlibMP1 Corchorus capsularis cDNA clone JuteSSHHsp701 similar to Hsp70 (Stretch 6-103 amino acid) like protein, mRNA sequence

AGCGTGGTCGCGGCCGAGGTCCTGTCTCATCATGGAAGGCGCTGAATTTACGGAAAACATTATTAATGGC

GTCGAGCGTCCGGTTAAAGCCGCTGAATTGTTCGCGTTTACCTTGCGTGTACGCGCAGGAAACACTGACG

TTCCTACTGACGCAGAAGAAAACGTGCGTCAAAAATTACGTGCAGAAGGAGTGATGTAATGTCTAAAGGT

AAAAAACGTTCTGGCGCTCGCCCTGGTCGTCCGCAGCCGTTGCGAGGTACTAAAGGCAAGCGTAAAGGCG

CTCGTCTTTGGTATGTAGGTGGTCAACAATTTTAATTGCAGGGGCTTCGGACCTGCCCGGGCGGCCGCTC

GA

>GR463677.1 JuteSSHRP74 JuteSSHlibMP1 Corchorus capsularis cDNA clone JuteSSHNBSLRR similar to disease resistance protein (TIR-NBS-LRR class),[Arabidopsis thaliana] like protein, mRNA sequence

CGAGCGGCCGCCCGGGCAGGTACACTAATACAGATTCCGGCACTAAATTTGAGATAACAAAAGAACTACT

ACAATGAAAGAACTCACTGAAAGTTTTCTGAAGTATCTAAAAACTAAACTTTTCGCTCAAAACATTCCTT

CAAGTTCAAATGTCTGCTTTAAGAAAGTTGTCTGATAGAGATACCTTCTCCCCAGCTTGAATGGAGGAAG

ACCCTATATGGACATGTACCTCGGCCGCGACCACGC

>GR463676.1 JuteSSHRP69 JuteSSHlibMP1 Corchorus capsularis cDNA clone JuteSSHHsp20 similar to Heat shock protein Hsp20 like protein, mRNA sequence

CGAGCGGCCGCCCGGGCAGGTACACATGAAGACTTGTCAAACTTTACTACAGGACACAGGAAGACAGAGC

ACTATATAGTTTTAGCTAATGAGCAAATGAAGTGTTTGAAGATTTTTAACCAGAAATCTCAATGGATTTG

ACATCAGGTTTCTTCACCTCCACCTTAGGAACAGTAACAGTAAGAACCCCATTTTCCATTGAAGCCTTAA

CTTGATCCATCTTCACATTCTCCGGCAACCTGAACCTCCTCATGAATTTGCCGCTGCTACGCTCAACGCG

GTGCCAGGTGTCGTTCTTGTCTTCCTTCTCAACGTTCCTCTCTCCGCTGATCTGAAGCACCCTATCGTCC

ACCTCGGCCGCGACCACGGTCGAGCGGCCGCCCGGGCAGGTACTAGGTTGAATTACTATTGCGACGCGGC

CATCAGTAGGGTAAAACTAACCTGTCTCACGACGGTCTAATCCCAGCTCACGTTCCCTATTGGTGGGTGA

ACAATCCAGCACTTGGTGAATTCTGCTTCACAATGATAGGAAGAGCCGACATCGAAGGGATCAAAAAGCA

ACGTCGCTATGAACGCTTGGCTGCCACAAGCCAGTTATCCCTGTGGTAACTTTTCTGACACCTCTAGCTT

CAAATTCCCGAAGGTCTAAAGGATCGATAGGCCACGCTTTCACGGTTCGTATTCGGACCTGCCCGGGGCG

GCAGCTTCAAATCTTGCTTGAAAACTCG

>GR463675.1 JuteSSHRP60 JuteSSHlibMP1 Corchorus capsularis cDNA clone JuteSSHDNAJ20 similar to Chaperone protein dnaJ 20 like protein, mRNA sequence

CGAGCGGCCGCCCGGGCAGGTACTTGCTTTTTGAGCAATGCTAACCCAACGGCCATCTATAATTAGGGTA

GAAAGTTATATATACATATGATCAACTGGAAAAACCTAACCAAAAGACTATTCATAGTTGTATGCACCGA

GGAAAATGTAAGTTGTTTACTATTATTCATTAGACAATCCATCCCTTTGTCTCCGCATTCGTGCTCCCCA

TGACATGTTTCCCCTGGCATCCTTGTTCATGCTTCTTCTCTTCAGCTTTGATAGCTGAGATTGCCAACGA

TTTTTCCACTCACCGCGCTCATCCAGACCTCGGCCGCGACCACGCT

>GR463674.1 JuteSSHRP68 JuteSSHlibMP1 Corchorus capsularis cDNA clone JuteSSHBIP similar to BIP like protein, mRNA sequence

CGAGCGGCCGCCCGGGCAGGTACTTCCACCAACTAGAACAATCTTATCAATCTGGTGCTTCTCCAAACCA

GCATCATCCATTGCTTTCTTCACAGGTCCCATGGTCTTTCTGAAGAGGTCATTGTTCAACTCCTCAAAGC

GAGCCCTGGTCAGTGGCTCAGAGAAATCAATACCATCGAAAAGTGATTCAATCTCCACACGGACTTGGTG

CTGACTGCTGAGAGCTCTCTTGGCTCGCTCAGCTTCCCTCCTAAGCTTCCCAATAGCCCTGTTGTCCTTG

CTGATATCCTTTCCATGCTTCTTCTTAATCAACTTAATAAAGTACCTCGGCCGCGACCACG

>GR463673.1 JuteSSHRP64 JuteSSHlibMP1 Corchorus capsularis cDNA clone JuteSSHGP4 similar to glutathione peroxidase like protein, mRNA sequence

CGAGCGGCCGCCCGGGCAGGTGTTCTCCGCAAAATCTCATTCGACGAAATCTTAGTTCAGTTGCTATTTC

TGCTTCTCTTCTTTTTGCAAAGCGTTTTTCACCGAGTTCCAAGCCAACCCTTTCCAGTTTTCCGCAGATT

TCGCCAGTTTCTTTGGTTTCATCCTCGAAGGAAACAGGGTTTTCAAGGTCTTTCTTGGGTAGTTTAAGAT

TTGATCATTCCATGGCTAGCCAATCAGCCCAGACTTCAGTTCATGACTTCACCGTTAAGGATGCAAGAGG

AAAGGACGTTGATCTAAGCACTTACAAAGGCAAGGTCCTCTTGATTGTCAATGTTGCCTCACAATGTGGC

TTGACAAATTCCAACTACACTGAGCTGAGTCAATTGTATGAGAAATATAAAGATCAAGGTTTTGAGATTC

TGGCATTTCCTTGTAACCAGTTTGGAGGTCAGGAGCCAGGGAACAATGAACAGATCTTAGAATTTGCTTG

TACCTCGGCCGCGACCACGC

>GR463672.1 JuteSSHRP58 JuteSSHlibMP1 Corchorus capsularis cDNA clone JuteSSHGT2 similar to O-glucosyltransferase 3 like protein, mRNA sequence

CGAGCGGCCGCCCGGGCAGGTACTTAATTTAGTTTTTCCTAAGCTCATTTACGAAGGACTTAAGGTTCTT

GTCGGAAGAACCACCTTCCATGGCAGCTTCTCTTGAAAATTCTTTCCATTTCTTAACATTCCTCTTGATT

TCTTCTCCTCTTTCCCCATCTCCCATCACCATTTCCAAACATCTCTTAATCTCGCAACCTACGACTAATC

CTTGTTCATTCTTCGTAACCCTCACCCCTGTCTTCCACACATCTTGGACAAGCTTCGAGTTGGTTCCTTG

ATCTGTCCATTGAGGAATTGTCACTGTCGACCTCGGCCGCGACCACGC

>GR463671.1 JuteSSHRP56 JuteSSHlibMP1 Corchorus capsularis cDNA clone JuteSSHDNACRR2 similar to Cystein rich repeat(C_tripleX) containing like protein, mRNA sequence

CGAGCGGCCGCCCGGGCAGGTACATATGGTAGATTTTCTGCATTTTTCTAATTTCCACGCCTCTTTCATC

TTCCCAAGAGCTAACTCAGATTTCCCTCCTGTTTTGAGATCGAATCTGGACCGCATCTGATCTGGCACTT

TCGGTCACAACGCTGCATAAGCCTTGTGTTGCTTTGAACGCAGCCAGTAGAACAAGCATCATAGCAATCA

ACCTGCGACTCAGCGTTTTCCATTTGTGAAGTGAAAATGCAGAAAATAAGGACAGCAACCATCATTTTCT

TCATCATCGCCATTTCTTCGATCTGCTTTGAGGAATACAATACCTCGGCCGCGACCACGCT

>GR463670.1 JuteSSHRP50 JuteSSHlibMP1 Corchorus capsularis cDNA clone JuteSSHDNAJ20 similar to Chaperone protein dnaJ 20 like protein, mRNA sequence

CGAGCGGCCGCCCGGGCAGGTACTTGCTTTTTGAGCAATGCTAACCCAACGGCCATCTATAATTAGGGTA

GAAAGTTATATATACATATGATCAACTGGAAAAACCTAACCAAAAGACTATTCATAGTTGTATGCACCGA

GGAAAATGTAAGTTGTTTACTATTATTCATTAGACAATCCATCCCTTTGTCTCCGCATTCGTGCTCCCCA

TGACATGTTTCCCCTGGCATCCTTGTTCATGCTTCTTCTCTTCAGCTCTGATAGCTGAGATTGCCAACGA

TTTTTCCACTCACCGCGCTCATCCAGACCTCGGCCGCGACCACGC

>GR463669.1 JuteSSHRP48 JuteSSHlibMP1 Corchorus capsularis cDNA clone JuteSSHE1 similar to exonuclease family protein [Arabidopsis thaliana] like protein, mRNA sequence

CGAGCGGCCGCCCGGGCAGGTACAAATCAGCAAACAGCTTGTCCAGTGACCTAATTGTTCTAAAAAATGA

CCATCACTGTCTCCCAAAGCAATTGGGCATTGGCATTCACCTTTTGGACTTCCCTTTAAATATCGTCTCC

TTGAAGTATTTGAGTTCTAATATGCTCTCCCTGATATCATCCATGGCTCTGTGTTTCTTTTCTTTGACAG

GGGCTTTCTTTTGATCTTTAGGATACCAGCGGATGCAAAGAGCCCTCACACTGCTTACATCAACGACTAT

ATGAGAGAAAAGACTAGCCAAATGTGGCATATATTCCTTTAGAAATAGAAAATCCATGCAAACTGAATTT

CCAGCTATATGAGGGGTGTATGTACCTCGGCCGCGACCC

>GR463668.1 JuteSSHRP43 JuteSSHlibMP1 Corchorus capsularis cDNA clone JuteSSHrp2 similar to 30S ribosomal protein, mRNA sequence

CGAGCGGCCGCCCGGGCAGGTACAAATATCTAAATTTGATATTTTTAGATACTTTTTCAGACCTATTGCC

GATACTAAGTAGCAGTCAAAAATTTGTATCCATTTTTCATGATATTATGCATGGATCAGATATATCATGG

CGTATTCTTCAGAAAAAATTGTGCCTTCCACAACGGAATCTGAGGAATCTGATAAGTGAGATTTCGAGTA

AGTGTTTACAGAATTTTCTTCTGTCCGAAGAAATGATTCATCGAAATAATGAGTCACCATTGATATCGAC

ATATCTGAGATCGCCAAATGTTCAGGAGTTCCTCTATTCAATCCTTTTCCTTCTTCTTGTTGCTGGATAT

CTCGTTTGTACCTCGGCCGCGACCACGC

>GR463667.1 JuteSSHPC10 JuteSSHlibMP1 Corchorus capsularis cDNA clone JuteSSHAT2 similar to Hsp70 (Stretch 6-103 amino acid) like protein, mRNA sequence

TCGAGCGGCCGCCCGGGCAGGTACAAGATACACCAGGAAAAGTAGTTGGGCCAATAAGATTCACCAATAT

TACTGGGGAGAAGGTGACTCTGTGGTGGGATGCCCCACTCAATGACGGTTGTGCTCCCATAACCCACTAC

ATCATTGAAAAACGGGAAACCAGCAGACTTGCCTGGGCACTAATTGAGGATAAATGTGAAGCCCAAAGTT

ACACTGCCATTAAACTAATAAACGGCAATGAATACCAATTCCGTGTTTCTGCAGTTAACAAGTTTGGTGT

TGGCAGGCCACTTGATTCTGATCCAGTGGTTGCTCAAATACAATATACTGTTCCTGATGCCCCTGGCATT

CCAGAACCTAGCAACATAACAGGCAACAGCATTACCCTGACATGGGCAAGGCCAGAATCAGATGGTGGCA

GTGAAATTCAACAGTATATCCTTGAAAGAAGAGAAAAGAAAAGCACAAGATGGGTAAAAGTGATCAGCAA

ACGACCAATCTCTGAAACAAGATTCAAAGTCACTGGTCTGACAGAAGGCAATGAGTATGAATTCCATGTC

ATGGCTGAAAATGCTGCAGGAGTTGGACCTGCAAGTGGCATCTCAAGACTCATTAAATGTAGAGAGCCCG

TCAACCCAC

>GR463666.1 JuteSSHRP42 JuteSSHlibMP1 Corchorus capsularis cDNA clone JuteSSHYCF2 similar to Ycf2 like protein, mRNA sequence

CGAGCGGCCGCCCGGGCAGGTACAAATATCTAAATTTGATATTTTTAGATACTTTTTCAGACCTATTGCC

GATACTAAGTAGCAGTCAAAAATTTGTATCCATTTTTCATGATATTATGCATGGATCAGATATATCATGG

CGTATTCTTCAGAAAAAATTGTGCCTTCCACAACGGAATCTGAGGAATCTGATAAGTGAGATTTCGAGTA

AGTGTTTACAGAATTTTCTTCTGTCCGAAGAAATGATTCATCGAAATAATGAGTCACCATTGATATCGAC

ATATCTGAGATCGCCAAATGTTCAGGAGTTCCTCTATTCAATCCTTTTCCTTCTTCTTGTTGCTGGATAT

CTCGTTTGTACCTCGGCCGCGACCACGC

>GR463665.1 JuteSSHRP41 JuteSSHlibMP1 Corchorus capsularis cDNA clone JuteSSHRP2 similar to chloroplast-specific ribosomal protein, mRNA sequence

GCGTGGTCGCGGCCGAGGTACTACCACAGGCTCTCTAACTTTCAATCTATTGAAGCCCTTCATGTGTCGA

CCATGGTCTGATACTTTCTCCTTAATCTTCCTTAACTTTCTTTGTAGAATTGAGGATACCAAACCTATAC

TCGCATATACAGTCTCAGCATCTTCCTCTGCACGAACCACTCCATGTTTCTTCGTAAATAAGACCTGCCC

GGGCGGCCGCTCG

>GR463664.1 JuteSSHRP40 JuteSSHlibMP1 Corchorus capsularis cDNA clone JuteSSHRP similar to chloroplast-specific ribosomal protein, mRNA sequence

GCGTGGTCGCGGCCGAGGTACTACCACAGGCTCTCTAACTTTCAATCTATTGAAGCCCTTCATGTGTCGA

CCATGGTCTGATACTTTCTCCTTAATCTTCCTTAACTTTCTTTGTAGAATTGAGGATACCAAACCTATAC

TCGCATATACAGTCTCAGCATCTTCCTCTGCACGAACCACTCCATGTTTCTTCGTAAATAAGACCTGCCC

GGGCGGCCGCTCG

>GR463663.1 JuteSSHRP39 JuteSSHlibMP1 Corchorus capsularis cDNA clone JuteSSHL2A similar to Arabidopsis thaliana ribosomal protein L2 like protein, mRNA sequence

CGAGCGGCCGCCCGGGCAGGCACCACATGAGTGGATGTATAGGAATCCAAATCTGCCGAACCACTCATGT

TATGATCTTCTACATCCTAGGTCTTCCCGTTCCGTCATCTGGCTTATGTTCTTCATGTAGCATTCAGACC

GAATGACTCTATGAAATTACGTCGATACTTCCACATATTATGGGTAACGTAGGAGACATCTCTATTTTTC

CCCGGGGGAATCCTTAGAATTACCACTGCTTAGCTTTCAATTCGCCTCTGACCATCAAATGAAATGTGAA

TAACCCGTCCTCCTCTCTTTGGAACAAGGGGCGCTTCTGGTTCTGTCGGTGCTTGAAACAATTTTGTCTT

CTCCATATTACTATATCTCTAGAGTCAATAATTTTATATGAGGAACTACTGAACTCAATCACTTGCTGCC

GTTACTCTTCAGTTTTCTGTTGAGGTCTATCCTGTAGAGGTACCTCGGCCGCGACCACGC

>GR463662.1 JuteSSHRP37 JuteSSHlibMP1 Corchorus capsularis cDNA clone JuteSSHLACCCASE similar to secretory laccase [Gossypium arboreum] like protein, mRNA sequence

CGAGCGCGCCGGGCAGTACGGCCAGGAGCATGGTGAGAGATCGGCGAAGACCTTGTGGTGTAAGCCCCAT

CTTGTCCCACAACTGTGAACATGTGGTATTCCACGCCAAACTCCATTGTCTCATTCAAAGCCGCATTTAC

AATTCGAAGGTGATAAACACTTTGGTAATCTACTTGCTGCTCGTATGTTGAATTGTTGCTGCAGCCATAT

GTTTCTCCAGGATGGCCGACCTCGGCCGCGACC

>GR463661.1 JuteSSHRP34 JuteSSHlibMP1 Corchorus capsularis cDNA clone JuteSSHCRP30S similar to Lycopersicon esculentum chloroplast-specific ribosomal protein like protein, mRNA sequence

GCGTGGTCGCGGCCGAGGTACTACCACAGGCTCTCTAACTTTCAATCTATTGAAGCCCTTCATGTGTCGA

CCATGGTCTGATACTTTCTCCTTAATCTTCCTTAACTTTCTTTGTAGAATTGAGGATACCAAACCTATAC

TCGCATATACAGTCTCAGCATCTTCCTCTGCACGAACCACTCCATGTTTCTTCGTAAATAAGACCTGCCC

GGGCGGCCGCTCG

>GR463660.1 JuteSSHRP31 JuteSSHlibMP1 Corchorus capsularis cDNA clone JuteSSHCRPL2 similar to ribosomal protein L2 (rpl2) gene like protein, mRNA sequence

CGAGCGGCCGCCCGGGCAGGTACACATGAGTGGATATATAGGAATCCAAATCTGCCGAACCACTCATGTT

ATGATCTTCTACATCCTAGGTCTTCCCGTTCCGTCATCTGGCTTATGTTCTTCATGTAGCATTCAGACCG

AATGACTCTATGAAATTACGTCGATACTTCCACATATTATGGGTAACGTAGGAGACATCTCTATTTTTCC

CCGGGGGAATCTTTAGAATTACCACTGCTTAGCTTTCAATTCGCCTCTGACCATCAAATGAAATGTGAAT

AACCCGTCCTCCTCTCTTTGAAACAAGGGGCGCTTCTGGTTCTGTCGGTGCTTGAAACAATTTTGTCTTC

TCCATATTACTATATCTCTAGAGTCAATAATTTTATATGAGGAACTACTGAACTCAATCACTTGCTGCCG

TTACTCTTCAGTTTTCTGTTGAGGTCTATCCTGTAGAGGTACCTCGGCCGCGACCACGC

>GR463659.1 JuteSSHRP30 JuteSSHlibMP1 Corchorus capsularis cDNA clone JuteSSHCRR similar to Cysteine rich repeat containing(55-92aa) like protein, mRNA sequence

CGAGCGGCCGCCCGGGCAGGTGCATATGGTAGATTTTCTGCATTTTTCTAATTTCCACGCCTCTTTCATC

TTCCCAAGAGCTAACTCAGATTTCCCTCCTGTTTTGAGATCGAATCTGGACCGCATCTGATCTGGCACTT

TCGGTCACAACGCTGCATAAGCCTTGTGTTGCTTTGAACGCAGCCAGTAGAACAAGCATCATAGCAATCA

ACCTGCGACTCAGCGTTTTCCATTTGTGAAGTGAAAATGCGGAAAATAAGGACAGCAACCATCATTTTCT

TCATCATCGCCATTTCTTCGATCTGCTTTGAGGAATACAATACCTCGGCCGCGACCACGC

>GR463658.1 JuteSSHRP29 JuteSSHlibMP1 Corchorus capsularis cDNA clone JuteSSHGT5 similar to glutathione transferase like protein, mRNA sequence

GCGTGGTCGCGGCCGAGGTACGCTGCTTGGAAAGAATCTCCTCATATTTGTCCATGGTTTCATATAATTG

TTTCACAGCCTCATCATAAGGCCCTTGCTTTCTAGCAAAGCCGCATCTGTAAACACCATTATTGATTCCA

CTGTATATCCATTCATTTGTCTCATCAATTTGAGCTCTCAAATGAGGAGGATATAGGTCCAAAGCAGGAT

TCTCTGCTATATCATTGAATTCAGTATTAAGCATGCGGATTATCTCTGCGCTCTCATTACTAACATTAGT

CTTGAATTTCTTGTCCCAAAGGACAGGAACGGTGTACCTGCCCGGGCGGCCGCTCG

>GR463657.1 JuteSSHRP28B JuteSSHlibMP1 Corchorus capsularis cDNA clone JuteSSHSOUL similar to SOUL heme-binding family protein [Arabidopsis thaliana] like protein, mRNA sequence

CGAGCGGCCGCCCGGGCAGGTACAAGAATCCTTGACCTTCAGTTTCTCATCTCAGACAAAGTCTTAAAAC

TTAAAAAGACAACAACTCATCTATTCTTCCATGTCAAACAACAGCCAAATTTCATTCACCCTGTCACTAT

ATTCAAACGGTGAATTGTACCTCGGCCGCGACCACGC

>GR463656.1 JuteSSHPC26 JuteSSHlibMP1 Corchorus capsularis cDNA clone JuteSSHAT1 similar to Aminotransferase (Stretch 7-91amino acid) like protein, mRNA sequence

ACGTGGTCGCGGCCGAGGTCCTCCACTATGAAATCGCGTAGAGGCTTTGCTATTCAGCGTTTGATGAATG

CAATGCGACAGGCTCATGCTGATGGTTGGTTTATCGTTTTTGACACTCTCACGTTGGCTGACGACCGATT

AGAGGCGTTTTATGATAATCCCAATGCTTTGCGTGACTATTTTCGTGATATTGGTCGTATGGTTCTTGCT

GCCGAGGGTCGCAAGGCTAATGATTCACACGCCGACTGCTATCAGTATTTTTGTGTGCCTGAGTATGGTA

CAGCTAATGGACCTGCCCGGGCGGCCGCTCGA

>GR463655.1 JuteSSHRP17 JuteSSHlibMP1 Corchorus capsularis cDNA clone JuteSSHPp1 similar to protein phosphatase 1, regulatory (inhibitor) subunit 1C like protein, mRNA sequence

GTATGGTAGTAGTGACGACGTGGATGAAATCTATCCTGTCTACTGTCGTGGTGCCGGTGTTGAGTTGCTG

TACTGCGTGGATGAGTGGGGGCTGATGTATGGGGAGCGAGAGCATGCGCGGAGAGTGTGAGTACGTGTGT

ATGAGTAGGACAGTACTGATGCTGTATATAGTAACGATATATATATCAGATAGAGAAATACACAAAGTAT

GTAGATACGATGGCATTGGAGAGTGGTACGTGTTGATATTTCTCTTCGTGAGTACTCGATTGAAGTAATC

TTATGACGGACATACTATATGACTACTAATCTATTGATTCTTGATCTTGTTGTGCCCCGCCGCAGTCCCC

CTTATGGGAAAATATATTGGTAGAGGGAT

>GR463654.1 JuteSSHRP3 JuteSSHlibMP1 Corchorus capsularis cDNA clone JuteSSHPS similar to 23S rRNA pseudouridine synthase like protein, mRNA sequence

GCGTGGTCGCGGCCGAGGTTTTTTTTTTTTCATATATAATTACAATACAAACTTCATTATTCTTTCACTA

CATACATATATATATTTCAAACTCATTACAAATAATCAAATATTGCCAAGAAAGATAGCTAGAAAACAAA

AGAAAAACACAAGTCATTAGCTCAATGACTTTTGCTTTGCATAAATATTATATATATAGGACATTCTTTT

ATTTATTCTCACAGCTTTAATAATAATTTAATTAATACTAGTTCACCACCAAAGTACCTGCCCGGGCGGC

CGCTCGA

>GR463653.1 JuteSSHRP4 JuteSSHlibMP1 Corchorus capsularis cDNA clone JuteSSHATCP1C similar to hypersensitive reaction associated Ca2+-binding protein(ATCP1) like protein, mRNA sequence

GTGTGTCGGGCGAATACATTCGGCTTTCAATGTACATAGTAAAAAGTGCAATGTTCAAAACAAAAGGACT

TTAAACTCAAAACTTAAACTAAAAAGGCAATTTAATGCGCAAAGTCAACAGCTGAAAATCTTAAGCAAAC

CATCAAAAGAAACGCCTTGATTTTCATCGCCGCCACCCAATCTGATCATAGCCTTAATATCTT

>GR463652.1 JuteSSHRP1 JuteSSHlibMP1 Corchorus capsularis cDNA clone JuteSSHATCP1B similar to hypersensitive reaction associated Ca2+-binding protein(ATCP1) like protein, mRNA sequence

GCGTGGTCGCGGCCGAGGTACAATTCGGCATTTCCAATGGAAACATAATAAAAAGTGAAATGTTCAAAAC

AAAAGGACTTTAAACTCAAAACTTAAACTAAAAAGGCAATTTAATGAGCAAAGTCAACAGCTAAAATCTT

AAGCAAACCATCAAAAGAAACGCCTTGATTTTCATCGCCGCCACCCAATCTGATCATAGCCTTAATATCT

TCATCGGTAGCAGAAAAACCAGCCCAGTTCATGTAACTTTTCAAGTCTTGATGGCTCAATCTCCCATCTC

CATCTTTATCCGTAACCTTAAAAGCATCTTCCATAACCCCAAAACCCGACGCCGCCGAGCTCTGTTTCCC

TCGAGACCTGCCCGGGCGGCCGCTCG

>GR463651.1 JuteSSHPC3 JuteSSHlibMP1 Corchorus capsularis cDNA clone JuteSSHppi3 similar to FKBP-type peptidyl-prolyl cis-trans isomerase like protein, mRNA sequence

AGCGTGGTCGCGGCCGAGGTACAGACCTGGTTTGGGGGAGAACTGCCAAGGAACTGGAGGGGCAGGGCTG

TGGCCCCAGAGGTTGGAGGATTTGGGGGGAGAGTTTCTTCTTTGGCAAGGGAGAGGGTGGCACAAAGCCC

AGGCAGAGGGGTCAGCTCGGGGTGAGACCATGACTAGCAGTACCTGCCCGGGCGGCCGCTCGA

>GR463650.1 JuteSSHRP88 JuteSSHlibMP1 Corchorus capsularis cDNA clone JuteSSHppi2 similar to FKBP-type peptidyl-prolyl cis-trans isomerase like protein, mRNA sequence

CGAGCGGCCGCCCGGGCAGGTACTTAACAGCCTTCTCATATCTCTTAGAAGCTCTCACATACTTGCCAAC

CTTAAACAATGCATTTCCTTCTTCTTTCTTTTACCAGCCGCTTCAATTTTTTCTTTGGTGTTCATGTCCC

ATGATTCTTTTTCCTTGGTGAAAGATACCATCTCAACTTCGTAGAATACAGTCGAGTCAGCAGGCACAAA

AGCCAACTCCTGCTGTGATCCAGACGACCCAAAAGCATACTCAGGTGCAATAGTCAGCTCTGCAACTTCA

CCCTTTTTCATCGTCATCGCAGTTCTATCAAGACCATCACAGACTTGCTCTTCATCTATAATGAACTCAA

ATGGTTCTTCGCCTTCGTTGTAACCCTTCTTCACGAAAACAGTACCTCGGCCGCGACCACGC

>GR463649.1 JuteSSHRP28A JuteSSHlibMP1 Corchorus capsularis cDNA clone JuteSSHppi1 similar to FKBP-type peptidyl-prolyl cis-trans isomerase like protein, mRNA sequence

CGAGCGGCCGCCCGGGCAGGTACTTAACAGCCTTCTCATATCTCTTAGAAGCTCTCACATACTTGCCAAC

CTTAAACAATGCATTTCCTTCTTCTTTCTTTTTACCAGCCGCTTCAATTTTTTCTTTGGTGTTCATGTCC

CATGATTCTTTTTCCTTGGTGAAAGATACCATCTCAACTTCGTAGAATACAGTTGAGTCAGCAGGCACAA

AAGCCAACTCCTGCTGTGATCCAGACGACCCAAAAGCATACTCAGGTGCAATAGTCAGCTCTGCAACTTC

ACCCTTTTTCATCGTCATCGCAGTTCTATCAAGACCATCACAGACTTGCTCTTCATCTATAATGAACTCA

AATGGTTCTTCGCCTTCGTTGTAACCCTTCTTCACGAAAACAGTACCTCGGCCGCGACCACGCT
