## Supplementary File 3 for "*In silico* Identification and Functional Characterization of Conserved miRNAs in Fibre Biogenesis Crop *Corchorus capsularis*"

**Supplementary File 3:** Total of 3350 miRNA after redundancy screening

>aqc-miR477b MIMAT0012586

CUCUCCCUCAAGGGCUUCUA

>aqc-miR167 MIMAT0012560

UCAAGCUGCCAGCAUGAUCUA

>aqc-miR530 MIMAT0012593

UGCAUUUGCACCUGCAUCUC

>aqc-miR398a MIMAT0012578

UGUGUUCUCAGGUCACCCCUU

>aqc-miR482a MIMAT0012589

UCUUGCCGACUCCUCCCAUACC

>aqc-miR166d MIMAT0012559

UCGGACCAGGCUUCAUUCCUC

>aqc-miR169c MIMAT0012562

CAGCCAAGGAUGACUUGCCGG

>aqc-miR160a MIMAT0012553

UGCCUGGCUCCCUGGAUGCCA

>aqc-miR477c MIMAT0012587

CUCUCCCUCAAGUUCUUCUA

>aqc-miR168 MIMAT0012561

UGGCUUAGUGCAGCUCGGGGA

>aqc-miR529 MIMAT0012592

AGAAGAGAGAGAGCACAACCC

>aqc-miR171d MIMAT0012569

UGAUUGAGCCGUGCCAAUAUC

>aqc-miR477d MIMAT0012588

CUCUUCUUCAAAGGCUUCUA

>aqc-miR171c MIMAT0012568

UAAUUGAACCGCACUAAUAUC

>aqc-miR172b MIMAT0012572

GGAAUCUUGAUGAUGCUGCAU

>aqc-miR159 MIMAT0012552

UUUGGACUGAAGGGAGCUCUA

>aqc-miR171e MIMAT0012570

UGAAUGAACCGAGCCAACAUC

>aqc-miR395a MIMAT0012575

CUGAAGGGUUUGGAGGAACUC

>aqc-miR171f MIMAT0012565

UAAUUGAGCCGUGCCAAUAUC

>aqc-miR169a MIMAT0012563

UAGCCAAGGAUGACUUGCCUA

>aqc-miR396b MIMAT0012577

UUCCACAGCUUUCUUGAACUU

>aqc-miR398b MIMAT0012579

UGUGUUCUCAGGUCGCCCCUG

>aqc-miR399 MIMAT0012580

UGCCAAAGGAGAGUUGCCCUA

>aqc-miR319 MIMAT0012573

UUGGACUGAAGGGAGCUCCCU

>aqc-miR408 MIMAT0012581

UGCUCUGCCUCAUCCUUGUCU

>aqc-miR535 MIMAT0012594

UGACAACGAGAGAGAGCACGCG

>aqc-miR477f MIMAT0012583

CUCUCCUUCAAAGGCUUCUA

>ama-miR396-3p MIMAT0031155

AAGCUCAAGAAAGCUGUGGGA

>ama-miR156 MIMAT0031153

CUGACAGAAGAGAGUGAGCAC

>bcy-miR529 MIMAT0020967

GAAGAAGAGAGAUGGUAGAG

>bcy-miR156 MIMAT0020964

UUUGACAGAAGAUAGAGAGCAC

>ccl-miR167b MIMAT0014083

UGAAGCUGCCAGCAUGAUCUGA

>crt-miR166a MIMAT0014084

UCGGACCAGGCUUCAUUCCCGU

>crt-miR166b MIMAT0014085

UCGGACCAGGCUUCAUUCCCUU

>csi-miR3952-5p MIMAT0037415

GCUGUAGAUAGGCCCUUCAAC

>csi-miR2275a-3p MIMAT0048951

UUUAGUUUCCUCCAAUAUCUUA

>csi-miR399f-5p MIMAT0048935

GGGCAAUUCUCCUUUGGCAGA

>csi-miR171f-5p MIMAT0048910

UAUUGGCCUGGUUCACUCAGA

>csi-miR396c MIMAT0018493

UUCAAGAAAUCUGUGGGAAG

>csi-miR403a-5p MIMAT0037405

CAUGUUUCGGGUUUGUGCGUG

>csi-miR3947-5p MIMAT0018487

UUAUUUCAGUAGACGACGUCACA

>csi-miR399e-3p MIMAT0048934

UGCCAAAGGAGAUUUGCCCGG

>csi-miR166k-3p MIMAT0048883

UCUUGGACCAGGCUUCAUUCC

>csi-miR9560-3p MIMAT0048980

UCAUAUUUGCUCCACCACCUGUGG

>csi-miR390a-3p MIMAT0037374

CGCUAUCCAUCCUGAGUUUCA

>csi-miR399c-5p MIMAT0037401

GUGCAGUCCUCCUUUGGCGUG

>csi-miR482b-5p MIMAT0037411

AAUGGGAGGCUUGGCAAGAAG

>csi-miR156e-3p MIMAT0048856

GCUCUCUGUGCUUCUGUCAUCA

>csi-miR536-5p MIMAT0048967

CGCACCCCAGCGUGGAACCAUC

>csi-miR482g-5p MIMAT0048987

AGUGGGAGCGUGGGGUAAGAAG

>csi-miR166d-5p MIMAT0048874

CUGUUGUCUGGUUCAAGG

>csi-miR171b-5p MIMAT0037396

UGUGAUAUUGGUUCGGCUCAUC

>csi-miR535a MIMAT0018484

UGACAAUGAGAGAGAGCACAC

>csi-miR399e-5p MIMAT0048933

GGGCAAUUCUCUAUUGGCAGU

>csi-miR156a-3p MIMAT0037391

UGCUCGCUCCUCUUCUGUCAGC

>csi-miR3952b-3p MIMAT0048962

UGAAGGGCCUUUCUAGAGCAC

>csi-miR477c-3p MIMAT0037408

GAAGUCCUUGGGGUUGAGUGA

>csi-miR159a-5p MIMAT0037392

GAGCUCCUUGAAGUCCAAAAG

>csi-miR169a MIMAT0014074

GAGCCAAGAAUGACUUGCCGA

>csi-miR408-3p MIMAT0018472

AUGCACUGCCUCUUCCCUGGC

>csi-miR156i-5p MIMAT0048990

UUGACAGAAGAUAGAUAGCAC

>csi-miR3627b-3p MIMAT0048955

UUGUCGCAGGAGCGUUGGCACC

>csi-miR167c-3p MIMAT0037394

AGGUCAUCUUGCAGCUUCAAU

>csi-miR482e-5p MIMAT0048973

GGUCAUGGGAGGAUUGGCGA

>csi-miR827 MIMAT0018485

UUAGAUGACCAUCAACAAACA

>csi-miR12108-3p MIMAT0048976

AUUUUAAAACGUGCGGAUGUCUGC

>csi-miR162-3p MIMAT0014088

UCGAUAAACCUCUGCAUCCAG

>csi-miR167d-3p MIMAT0048885

AUCGGAUCAUGUGGUAGCUUCACC

>csi-miR12109-5p MIMAT0048977

UGCUUGGUGCAUAAAACUACAGAA

>csi-miR172a-5p MIMAT0017387

GCAGCGUCCUCAAGAUUCACA

>csi-miR858-3p MIMAT0048949

CUCGUUGUCUGUUCGACCUUG

>csi-miR390a-5p MIMAT0014092

AAGCUCAGGAGGGAUAGCGCC

>csi-miR2111-5p MIMAT0049002

UAAUCUGCAUCCUGAGGUUUG

>csi-miR482a-3p MIMAT0018480

UCUUCCCUAUGCCUCCCAUUCC

>csi-miR166h-3p MIMAT0048880

UUUGGACCAGGCUUCCUUCCUC

>csi-miR164b-3p MIMAT0048869

CACGCGCUCCCCUUCUCCAAC

>csi-miR3951a-3p MIMAT0037414

UUUCUCUUAUCGUUAUCUGU

>csi-miR393b-5p MIMAT0048919

UUCCAAAGGGAUCGCAUUGAUC

>csi-miR408-5p MIMAT0037406

ACGGGGAACAGGCAGAGCAUG

>csi-miR3952b-5p MIMAT0048961

GCUGUAGAAAGGCCCCUCAAC

>csi-miR482d-3p MIMAT0048944

UCCCUACUCCACCCAUGCCAUA

>csi-miR171e-3p MIMAT0048909

UUGAGCCGCGUCAAUAUCUCC

>csi-miR482c-3p MIMAT0018495

UUCCCUAGUCCCCCUAUUCCUA

>csi-miR166g-3p MIMAT0048879

UCUCGGACCAGGCUUCAUUCC

>csi-miR156e-5p MIMAT0048855

GUGACAGAAGAUAGAGAGCGC

>csi-miR3950 MIMAT0018491

UUUUUCGGCAACAUGAUUUCU

>csi-miR167b-3p MIMAT0037395

AGAUCAUGCGGCAGUUUCACC

>csi-miR477a-5p MIMAT0018475

ACCUCCCUCGAAGGCUUCCAA

>csi-miR12107-3p MIMAT0048972

UCAUUCGCGCUCUCAUCAUUA

>csi-miR3948 MIMAT0018488

UGGAGUGGGAGUGGGAGUAGGGUG

>csi-miR393b-3p MIMAT0048920

UCAUGCGAUCCCUUCGGAAUU

>csi-miR160b-5p MIMAT0048864

UGCCUGGCUCCCUGUAUGCCG

>csi-miR159c-3p MIMAT0048863

CUUGGACUGAAGGGAGCUCCC

>csi-miR12110-5p MIMAT0048981

CUGGGGAGUUGCACCCGGAGU

>csi-miR172c-5p MIMAT0037397

GGAGCACCAUCAAGAUUCACA

>csi-miR12111-5p MIMAT0048983

GAAAAACUUAGGUGGGAUAGUGUG

>csi-miR169e-3p MIMAT0048894

CAGGCAGUCUCCUUGGCUAAG

>csi-miR530a-3p MIMAT0037412

AGGUGCAGCUGUAUAUGCAGG

>csi-miR166c-5p MIMAT0037393

GGGAAUGUUGUCUGGUUCGAG

>csi-miR172b-5p MIMAT0048915

GCAGCAUCAUCAAGAUUCACA

>csi-miR395b-5p MIMAT0048924

GUUCCCCUGAACACUUCAUUG

>csi-miR156f-3p MIMAT0048858

GCUCUCUCUUCUCCUGUCAUUG

>csi-miR1515b-5p MIMAT0048959

UCAUUUUUGCGUGCAAUGAUCC

>csi-miR156g-3p MIMAT0048860

GCUCUCUAUUCUUCUGUCAUC

>csi-miR159c-5p MIMAT0048862

GAGCUUUCUUCGGUCCACUU

>csi-miR3627a-5p MIMAT0048954

UUGUCGCCGGAGAGAUAGCACC

>csi-miR172c-3p MIMAT0018458

UGGAAUCUUGAUGAUGCUGCAG

>csi-miR2275b-5p MIMAT0048952

AGAAUUGGAUGGAACUAAACA

>csi-miR164a-3p MIMAT0037371

CAUGUGCCCUUCUUCCCCAUC

>csi-miR156h-5p MIMAT0048989

ACAGAAGAUAGAGAACACAUA

>csi-miR160b-3p MIMAT0048865

GCGUACGAGGAGCCAAGCAUA

>csi-miR482f-3p MIMAT0018473

UUUUCCCACACCUCCCAUCCC

>csi-miR166b-5p MIMAT0048872

GGAAUGUUGUUUGGCUCGAGGG

>csi-miR399d-3p MIMAT0018468

UGCCAAAGGAGAGUUGCCCUG

>csi-miR857 MIMAT0018486

UUUUGAAUGUUGAAUGGUGGCUAU

>csi-miR160a-3p MIMAT0037370

GCGUAUGAGGAGCCAUGCAUA

>csi-miR535d-3p MIMAT0048999

GUGCUCUCUACCAUUGUCAUA

>csi-miR160c-3p MIMAT0048867

GCGUGCGAGGAGCCAUGCAUG

>csi-miR391-5p MIMAT0048957

UGCAGGUGAGAUGAUACCGUCA

>csi-miR403b-5p MIMAT0048937

AGUUUGUGCGUGAAUCUAACC

>csi-miR169n-3p MIMAT0048905

GGCAAGUUGCCCUUGGCUACA

>csi-miR171b-3p MIMAT0018455

CGAGCCGAAUCAAUAUCACUC

>csi-miR167e-3p MIMAT0048887

GGUCAUGCUCUGACAGCCUCACU

>csi-miR477e-3p MIMAT0048942

UGAGGUUCUUGGGGAGAGUAG

>csi-miR393c-3p MIMAT0048922

AUCAUGCUAUCCCUUUGGAUU

>csi-miR482c-5p MIMAT0037416

GGAAUUGGGUGCUAGGGAAGG

>csi-miR3951b-5p MIMAT0049003

CAGGUAAAGAUGAGAGAAAAA

>csi-miR156b-3p MIMAT0048851

GCUCACUGCUCUUUCUGUCAGCU

>csi-miR166e-5p MIMAT0017386

GGAAUGUUGUCUGGCUCGAGG

>csi-miR3954b-3p MIMAT0048964

ACCGUGUUUCUCUGCCCAAUC

>csi-miR399b-5p MIMAT0037404

AGGGCUUCUCUCCUUUGGCAG

>csi-miR12110-3p MIMAT0048982

CCCGAGUCACAGCUCCCUGGAC

>csi-miR12108-5p MIMAT0048975

AGACGUUCGCACUUUUUAAAAUGC

>csi-miR3954b-5p MIMAT0048963

UUGGACAGAGAAAUCACGGUCA

>csi-miR477c-5p MIMAT0018474

ACUCUCCCUCAAGGGCUUCGC

>csi-miR477b-3p MIMAT0037410

UGAAGCCCUUAGGGCAGAGUGA

>csi-miR12111-3p MIMAT0048984

GACUGUCCCAUCUAAGUUUUUCUC

>csi-miR12109-3p MIMAT0048978

CUGCAGAUUUGUGCAUUAAGCUGC

>csi-miR3627c-5p MIMAT0048956

UCGCAGGGGAGAUGGGACCAAC

>csi-miR12106-5p MIMAT0048969

GUUUAGGACAGCUGCUGCCAAA

>csi-miR3951a-5p MIMAT0018492

UAGAUAAAGAUGAGAGAAAAA

>csi-miR12106-3p MIMAT0048970

UGGGGCAGCUGUCCUAAACGG

>csi-miR482e-3p MIMAT0048974

UUGCCAACUCCUCCCAUGCCGA

>csi-miR160c-5p MIMAT0048866

UGCCUGGCUCCCUGUAUGCUU

>csi-miR399c-3p MIMAT0018467

UGCCAAAGGAGAAUUGCCCUG

>csi-miR530b-5p MIMAT0018483

UGCAUUUGCACCUGCAUCUUG

>csi-miR398a-5p MIMAT0037372

AGAACAGAGGGUGGCGUUGGCU

>csi-miR530a-5p MIMAT0018482

UGCAUUUGCACCUGCACCUUG

>csi-miR397-3p MIMAT0037400

ACCAGCGCUGCACUCGAUCAU

>csi-miR166a-3p MIMAT0014090

UCGGACCAGGCUUCAUUCCCCC

>csi-miR393a MIMAT0018460

AUCCAAAGGGAUCGCAUUGAUC

>csi-miR159d MIMAT0018459

UUUGGACUGAAGGGAGCUCCU

>csi-miR3951b-3p MIMAT0049004

UUUCUCUUAUCUUUUAUCUGUG

>csi-miR536-3p MIMAT0048968

UGGUGCCACGCUGUGUGCGUC

>csi-miR399d-5p MIMAT0037402

GGGCACCUCUUACUUGGCAUG

>csi-miR477b-5p MIMAT0018476

CUCUCCCUCAAGGGCUUCUCU

>csi-miR396a-3p MIMAT0037398

AUUCAAGAAAGCUGUGGAAAA

>csi-miR162-5p MIMAT0017388

UGGAGGCAGCGGUUCAUCGAUC

>csi-miR12105-5p MIMAT0048965

UUUCUUGAGACAAAACAUGCAUC

>csi-miR156j-3p MIMAT0048992

AUCCAACGAAGCAGGAGCUGC

>csi-miR399f-3p MIMAT0048936

CGCCAAAGGAGAAUUGCCCUG

>csi-miR396b-3p MIMAT0037399

GUUCAAUAAAGCUGUGGGAAG

>csi-miR169l-3p MIMAT0048902

AGGCAGUCUCCUUGGCUAAC

>csi-miR477e-5p MIMAT0048941

ACUCUCCCUCAAGGGCUUCUGA

>csi-miR172d-5p MIMAT0048917

GCGGCAUCAUCAAGAUUCACA

>csi-miR9560-5p MIMAT0048979

ACAGGAGGUGGAACAAAUAUGAAA

>csi-miR477d-3p MIMAT0048940

UGAGGCCGUUGGGGAGAGUGG

>csi-miR477d-5p MIMAT0048939

ACUCUCCCUCAAGGGCUUCUGG

>csi-miR396f-3p MIMAT0048931

GCUCAAGAAAGCUGUGGGAGA

>csi-miR530b-3p MIMAT0037413

AGGUGCAGUUGCAAGUGCAGA

>csi-miR477a-3p MIMAT0037409

GGAAACCCUAGGGGGAGGUCG

>csi-miR1515b-3p MIMAT0048960

AUCAUUCACGCAAAAAUGAUU

>csi-miR159b-5p MIMAT0048861

AGCUGCCGACUCAUUCAUUCA

>csi-miR399a-5p MIMAT0037403

GGGCAACAUCUCCAUUGGCAGG

>csi-miR166f-5p MIMAT0048876

GGACUGUUGUCUGGCUCGAUG

>csi-miR12107-5p MIMAT0048971

CUGAUGAGAGAGCGAAUGAUA

>csi-miR482b-3p MIMAT0018481

UCUUGCCCACCCCUCCCAUUCC

>csi-miR156f-5p MIMAT0048857

AUGACAGAAGAGAGAGAGUAC

>csi-miR166g-5p MIMAT0048878

GAAUGCUGUCUGGUUCGAGAC

>csi-miR482d-5p MIMAT0048943

UGGUAUGGGUGAGUAGGGAAG

>csi-miR3949 MIMAT0018490

UGAUGUUGAGGCAAAAAUGUAG

>csi-miR167a-3p MIMAT0037417

AGAUCAUCUGGCAGUUUCACC

>csi-miR12105-3p MIMAT0048966

UUGUUUUGGGUGAAACGGGUGU

>csi-miR396e-3p MIMAT0048929

GCUCAAGAAUGCCGUGGGAAA

>csi-miR171c-5p MIMAT0048906

UGUUGGAACGGCUCAAUCAAA

>csi-miR159a-3p MIMAT0018450

UUUGGAUUGAAGGGAGCUCUA

>csi-miR156g-5p MIMAT0048859

UUGACGGAAGAUAGAGAGCAC

>csi-miR156j-5p MIMAT0048991

AGCUGCUGACUCGUUGGUUCA

>csi-miR166j-3p MIMAT0048882

UUGGACCAGGCUUCAUUCCUC

>csi-miR391-3p MIMAT0048958

CCGGAAUCAUUUCUCCCGCGUG

>csi-miR3946 MIMAT0018449

UUGUAGAGAAAGAGAAGAGAGCAC

>csi-miR482f-5p MIMAT0037407

GAUGGGUGAGUUGGGAAAAUA

>csi-miR3953 MIMAT0018496

UUGAGUUCUGCAAGCCGUCGA

>dpr-miR397 MIMAT0023537

CCAUUGAGUGCAGCGUUGAUG

>dpr-miR408 MIMAT0023538

CUGCACUGCCUCUUCCCUGGC

>dpr-miR156a MIMAT0023526

UGACAGAAGAGAGGGAGCAC

>dpr-miR160 MIMAT0023528

UGCCUGGCUCCUUGUAUGCCA

>eun-miR482a-5p MIMAT0041082

CAUGGGUUGUUUGGUGAGAGG

>eun-miR530-5p MIMAT0041088

UCUGCAUUUGCACCUGCACCU

>eun-miR10214a-3p MIMAT0041105

UAGAUAUAGUGGAUUUUCGAU

>eun-miR397a-3p MIMAT0041079

CGGUUUCGACAGCGCUGCACU

>eun-miR10220-3p MIMAT0041117

UCGUGAAGGAAGAAUGUGCAAU

>eun-miR395-5p MIMAT0041072

UCCCCUAGAGUUCUCCUGAACA

>eun-miR159-3p MIMAT0041052

CUUGCAUAUGCCAGGAGCUUC

>eun-miR10214b-3p MIMAT0041106

CAGAGAUAUAGUGGAUUUACG

>eun-miR530-3p MIMAT0041089

AGGUGCGGGUGCAGGUGCAGA

>eun-miR10216-5p MIMAT0041108

UCUCUGUUGAUCUGAUAAAUA

>eun-miR396b-3p MIMAT0041077

GUUCAAGCUAGCUGUGGGAAG

>eun-miR167b-3p MIMAT0041062

UCAGGUCAUCUUGCAGCUUCA

>eun-miR10217-3p MIMAT0041111

UAAGUGGUGAUCUGACUCUAA

>eun-miR397b-3p MIMAT0041081

UCAACGCUGCACUCAAUGAUG

>eun-miR167d-5p MIMAT0041065

UGAAGCUGCCACAUGAUCUGA

>eun-miR535a-3p MIMAT0041091

GUGCUCUCUAUCGCUGUCAUA

>eun-miR10219-5p MIMAT0041114

UUCAAGUCUAACAACCUCAGCU

>eun-miR167c-3p MIMAT0041064

AUCAGAUCAUGUGGCAGCUUCACC

>eun-miR10218-3p MIMAT0041113

AUCUGUGGAAGAGACUCGACU

>eun-miR10222-3p MIMAT0041121

UUCAUGGACAGCCAGCUUAUU

>eun-miR10223-3p MIMAT0041123

UCUCGUUCCGCUUCAUCUGAA

>eun-miR10211a-3p MIMAT0041097

AUCCAGAAAUUGGCAGCCGUU

>eun-miR10213-3p MIMAT0041103

UGGAUCAAUAGAACGAGCAGGUGA

>eun-miR482a-3p MIMAT0041083

UCUUGCCAAUACCACCCAUGCC

>eun-miR395-3p MIMAT0041073

AUGAAGUGUUUGGGGGAACUC

>eun-miR482b-3p MIMAT0041085

UUUCCUAUUCCUCCCAUUCCAU

>eun-miR10215-3p MIMAT0041107

GAGAAUGAUGAGUUAAAUGGA

>eun-miR10221-5p MIMAT0041118

GCUCGAGGUCAGUUUGUCGCC

>eun-miR167c-5p MIMAT0041063

UGAAGCUGCCAGCGUGAUCUCA

>eun-miR10223-5p MIMAT0041122

UGAAGCAGAUCAAGAACCCAG

>eun-miR482c-3p MIMAT0041087

UUCCCAAGGCCGCCCAUUCCGA

>eun-miR10213-5p MIMAT0041102

UACUCGUUCCGUUGAUCCAUC

>eun-miR172c-5p MIMAT0041070

GUAGCAUCAUCAAGAUUCACA

>eun-miR10214a-5p MIMAT0041104

UCGUAAAUCCACUAUAUCUCU

>eun-miR397b-5p MIMAT0041080

UGCAGCGCUGUCGAAACCGAU

>eun-miR10211b-5p MIMAT0041098

CAAUUUCUGGAUUUCAGUUCG

>eun-miR10212-3p MIMAT0041101

CGAUCUAAUCAAUCAUUUUUCGGG

>eun-miR10220-5p MIMAT0041116

UGCUGUUCUUCCGUUCACGAAU

>eun-miR10218-5p MIMAT0041112

UCGAGCCCCUCCCACAGAUUG

>eun-miR10212-5p MIMAT0041100

CGAAAAAUGAUUGGUUGUAUCGCU

>eun-miR482b-5p MIMAT0041084

GAAAUGGGAGGGUGGGAAAGA

>eun-miR10216-3p MIMAT0041109

UUUGUCGGAUGAACAGGGAAU

>eun-miR167b-5p MIMAT0041061

UGAAGCUGCCAGCAUGAUCUGG

>eun-miR396a-5p MIMAT0041074

UUCCACGGCUUUCUUGAACUG

>eun-miR827-5p MIMAT0041094

CUUUGUUGAUGGCCAUCUAAUC

>eun-miR482c-5p MIMAT0041086

GAGAUUCGAGCUACCGGAAGUUGUG

>eun-miR10222-5p MIMAT0041120

CAUGUAACAAGUUGGCUGUCA

>eun-miR10224-3p MIMAT0041124

CAAUGAACGCAUUUGCAGGUG

>eun-miR535b-3p MIMAT0041093

UGCUCUCUACCGUUGUCAUG

>eun-miR159-5p MIMAT0041051

AGCUGCUGGUCUAUGGAUCCC

>eun-miR10211a-5p MIMAT0041096

UCGGCUGUCAAUUUCUGGAUU

>eun-miR10217-5p MIMAT0041110

UAGGGUCAGAUCGCUACUUAG

>eun-miR827-3p MIMAT0041095

UUAGAUGACCAUCAGCGAACA

>eun-miR10221-3p MIMAT0041119

CGGCAAACUGGACCUCGAGAUC

>eun-miR167a-3p MIMAT0041060

AGAUCAUCUGGCAGUUUCAAC

>eun-miR10219-3p MIMAT0041115

UGGAGGUUGUUUGGCUUGAGCU

>eun-miR160-3p MIMAT0041054

GCGUAUGAGGAGCCAAGCAUA

>eun-miR10211b-3p MIMAT0041099

UCGAACUGAAAUCCAGAAAUU

>eun-miR172a-5p MIMAT0041066

CAGGUGUAGCAUCAUCAAGAU

>fve-miR11287 MIMAT0044568

UCAGGGAUUGUUUCAUAGACC

>fve-miR11299 MIMAT0044582

CAAAUAGGGUUGGCUGAUACU

>fve-miR11314 MIMAT0044598

AGAGUUGUGGAUGCUAUGAAU

>fve-miR11289 MIMAT0044571

UGCUUCAAGUCUGGCCAAUACU

>fve-miR482a MIMAT0044557

UCUUUCCAAUUCCUCCCAUGCC

>fve-miR11284 MIMAT0044473

UUCGUGAUCUGCGAAAGGCUC

>fve-miR11297 MIMAT0044580

CAGACAAGAUCGAUCUCGCCU

>fve-miR11300 MIMAT0044583

CAACAUCACUGUUCUCUUCCU

>fve-miR477a MIMAT0044556

ACUCUCCCUCAAGGGCUUCUC

>fve-miR408 MIMAT0044555

UGCACUGCCUCUUCCCUGGCU

>fve-miR535a MIMAT0044563

UGACGAUGAGAGAGAGCACGC

>fve-miR845 MIMAT0044566

AACCGGCUCUGAUACCAAUUG

>fve-miR482b MIMAT0044558

UCUUUCCUAGUCCUGCCAUUCC

>fve-miR11294 MIMAT0044577

UUCACCUGGACCAUAACUGACC

>fve-miR11291 MIMAT0044574

UUGCGGUCUUGUCUCUUCCAAU

>fve-miR11295 MIMAT0044578

CUCAUUCAAUUUCGGUAUUCAG

>fve-miR11305 MIMAT0044588

UUUUGGUCCGAAUCCGAGCUCC

>fve-miR159c MIMAT0044490

AUUGGAUUGAAGGGAGCUCCC

>fve-miR11301 MIMAT0044584

UCAGAGUUGUAAUAUAUUGAU

>fve-miR11286 MIMAT0044567

UUGGAGAGAGAGUAGACAAUG

>fve-miR2109 MIMAT0044531

UGCGAGUGUCUUCACCUCUGAA

>fve-miR11309 MIMAT0044592

UUUGUUUGGCAUGCAGUUGGC

>fve-miR156j MIMAT0044486

UUGACGGAAGAGAGCGAGCAC

>fve-miR166d-5p MIMAT0044502

GGGAAUGUCGUCUGGUUCGA

>fve-miR11312 MIMAT0044595

UGGAGCUGUUGGGGAGAGUUA

>fve-miR11288e MIMAT0044601

UGAAGUGGGAUUUGGCGAAUU

>fve-miR164a-3p MIMAT0044496

CACGUGCUCCCCUUCUCCAAC

>fve-miR396b-3p MIMAT0044545

GCUCAAGAAAGCUGUGGGACA

>fve-miR11304 MIMAT0044587

UAGUGUUUCAGCUUGACAAG

>fve-miR11308 MIMAT0044591

UAAGUUAGGAUUCUAGUUACC

>fve-miR171b MIMAT0044525

CGAGCCGAACCAAUAUCACUC

>fve-miR11288a MIMAT0044573

UGGGAUUUGGCGAAUUGUGGU

>fve-miR11288c-5p MIMAT0044569

UUCGUUCGGAUUCCAAUUCAAA

>fve-miR11310 MIMAT0044593

AGGGAUCACAACUCCAGUGCU

>fve-miR482c MIMAT0044559

UCUUUCCUAUUCCUCCCAUCCC

>fve-miR159a-5p MIMAT0044487

GAGCUCCUUGAAGUCCAAUAG

>fve-miR171f-5p MIMAT0044526

CGAUGUUGGUGAGGUUCAAUC

>fve-miR11288c-3p MIMAT0044570

UGAAUUGGGAUUUGUCGAAUU

>fve-miR3627b MIMAT0044538

UCGCAGGAGAGAUGGCACUACC

>fve-miR11292 MIMAT0044575

UUGUAGUUCAGCGCCUCCGCC

>fve-miR11298 MIMAT0044581

UUGAGGGGCUUAACGAUUACC

>fve-miR169f MIMAT0044517

UGAGCCAAGAAUGACUUGCUG

>fve-miR11293 MIMAT0044576

UUCUUCCUCAGGAACCUCCACC

>fve-miR164c MIMAT0044498

UGGAGAAGCAGGGCACAUGCU

>fve-miR11290 MIMAT0044572

UCGUCGACAUCAAAGGGCACC

>fve-miR11302 MIMAT0044585

AGGACCGCCAUCACGUUUUGG

>fve-miR482d MIMAT0044560

UUCCCUAUUCCACCUAUUCCCC

>fve-miR11285 MIMAT0044474

UCUAUUCAAAGAGAUGACUGUU

>fve-miR11313 MIMAT0044597

GAAAAGAAUGGACUCUCCGGGG

>fve-miR11311 MIMAT0044594

UGAGAAUGAUGUGGAUCUCAGC

>fve-miR171f-3p MIMAT0044527

UUGAGCCGCGCCAAUAUCACU

>fve-miR11315 MIMAT0044602

CUAGUCAUUGGUCAUAGCAUC

>fve-miR399b MIMAT0044554

AGCCAAAGGAGAAUUGCCCUG

>fve-miR393b MIMAT0044541

UCCAAAGGGAUCGCAUUGAUCU

>fve-miR11296 MIMAT0044579

UUUUUGAUGGCUGGAAUCCAGU

>fve-miR11306 MIMAT0044589

CUACCGAAGAACUUUGCAAAAG

>fve-miR11303 MIMAT0044586

UCAAACAUCACUGCAGCUGUA

>fve-miR156h MIMAT0044484

UGACAGAAGAGAGUGAGCUC

>fve-miR169e MIMAT0044516

UGAGCCAAGGAUGACUUGCCU

>fve-miR11288b MIMAT0044599

UGGGAUUGGGCGAAUUUUGGU

>fve-miR171c-5p MIMAT0044522

AGAUAUUGGUGCGGUUCAAUC

>fve-miR167d MIMAT0044506

UGAAGCUGCCAGCAUGAUCUCA

>fve-miR1511 MIMAT0044475

ACCUAGCUCUGAUACCAUGUG

>fve-miR530 MIMAT0044562

UGCAUUUGCACCUGCACCUCU

>fve-miR11283 MIMAT0044472

AGGCUUUGUAGAGGAUGGAAU

>fve-miR11307 MIMAT0044590

UAAGCGACGGACUCCAAUCGC

>fve-miR5225 MIMAT0044561

CUGUCGUAGGAGAGAUGGCGCC

>fve-miR477b MIMAT0044596

CGCGCACCCGUUCAUCUUCGC

>fve-miR2111b-3p MIMAT0044535

GCCCUUGGGAUGCGGAUUACC

>fve-miR11288d MIMAT0044600

UCCAUCGUUUUGAGACACAGG

>fve-miR396e MIMAT0044551

UUCCACAGGCUUUCUUGAACU

>fve-miR159b MIMAT0044489

AUUGGAUUGAAGGGAGCUCUC

>gar-miR2947 MIMAT0014330

UAUACCGUGCCCAUGACUGUAG

>ghr-miR7492b MIMAT0029135

CUAUAGAACAUGAUCUUUAGCGG

>ghr-miR7507 MIMAT0029154

AAGGUAGUGAAGUAGGCAAUUGGG

>ghr-miR399e MIMAT0025840

UGCCAAAGGAGAUUUGCCCCG

>ghr-miR482b MIMAT0014340

UCUUGCCUACUCCACCCAUGCC

>ghr-miR7508 MIMAT0029155

CAAGAAAAGAAGUCGGGAGAG

>ghr-miR7498 MIMAT0029145

AUGGUGACACAUGGUAGUCUCACA

>ghr-miR7497 MIMAT0029144

ACAUGUGGACUGUCAUAUGGGUU

>ghr-miR399d MIMAT0014350

UGCCAAAGGAGAUUUGCCCUG

>ghr-miR2950 MIMAT0014348

UGGUGUGCAGGGGGUGGAAUA

>ghr-miR399a MIMAT0005820

CGCCAAUGGAGAUUUGUCCGG

>ghr-miR7490 MIMAT0029132

AGUCUAGAAAACUUCACUGACGGU

>ghr-miR3476-3p MIMAT0015654

AGCCAACAACAUCAGUUCUAA

>ghr-miR2949a-5p MIMAT0014345

ACUUUUGAACUGGAUUUGCCGA

>ghr-miR7495b MIMAT0029141

UUACUUUAGAUGUCUCCUUCA

>ghr-miR7494 MIMAT0029139

AGCUUGUGGACUAGUUUUAACAA

>ghr-miR3476-5p MIMAT0015653

UGAACUGGGUUUGUUGGCUGC

>ghr-miR169b MIMAT0029157

CAGCCAAGGAUGAUUUGCCGG

>ghr-miR172 MIMAT0014333

AGAAUCCUGAUGAUGCUGCAG

>ghr-miR7504a MIMAT0029151

UAUGAAACUGUGAUUCCACGUCAU

>ghr-miR7504b MIMAT0029159

AGGAGGAAAAAUCUGAUUUGUCAU

>ghr-miR7505 MIMAT0029152

UUCAGAAACCAUCCCUUCCUU

>ghr-miR7510b MIMAT0029163

AAGAACAUGAUCUUUAGCGGCGU

>ghr-miR7513 MIMAT0029162

AAUCAGCCAGGAAUCGUUUGA

>ghr-miR7484a MIMAT0029124

UUUGUAUAUUAGAUCAAAGAGCAA

>ghr-miR479 MIMAT0014338

CGUGAUAUUGGUUCGGCUCAUC

>ghr-miR2949b MIMAT0014346

UCUUUUGAACUGGAUUUGCCGA

>ghr-miR7500 MIMAT0029147

AUCGAGUUAUUCGAGUUAAUCGAG

>ghr-miR7486b MIMAT0029128

AAGGAAGCGCUUUGUCCACGUGGA

>ghr-miR7510a MIMAT0029158

AAGGUCAUGAUCUUUAGCGGCGUU

>ghr-miR7496a MIMAT0029142

AUGACCAAAUUGAUAGAAUGUGUA

>ghr-miR7506 MIMAT0029153

AUGUCUGGGACAUGGCGUUGGCAC

>ghr-miR7501 MIMAT0029148

AUAUCUGAUUCUGACACGAAAAAA

>ghr-miR7493 MIMAT0029138

AAUAUUUUAAUAAUUCAAUCGUCA

>ghr-miR7503 MIMAT0029150

AGAUCGAUGGCUGAACAAGUUAGA

>ghr-miR156c MIMAT0005808

UGUCAGAAGAGAGUGAGCAC

>ghr-miR7485 MIMAT0029126

AAAGACAUCUUUGAAUUCUUGGAG

>ghr-miR7487 MIMAT0029129

AUACUCUUAUAGGACACUUGUUAA

>ghr-miR482a MIMAT0014339

UCUUUCCUACUCCUCCCAUACC

>ghr-miR2948-5p MIMAT0014344

UGUGGGAGAGUUGGGCAAGAAU

>ghr-miR7499 MIMAT0029146

AUAUAAUUUUCGGUUAAUUCGGUU

>ghr-miR7502 MIMAT0029149

UUUUUAACAGUAGAAAUGAAUGAA

>ghr-miR7514 MIMAT0029164

AUAAAGUGAUAAGUGAGAUCGUCU

>ghr-miR399c MIMAT0014349

UGCCAAAGGAGAGUUGGCCUU

>ghr-miR2949a-3p MIMAT0015374

UGCAAAUCCAGUCAAAAGUUA

>ghr-miR7511 MIMAT0029160

AGAAGUUUUGCAUGUGUAGCUGAG

>ghr-miR7491 MIMAT0029133

UGGGAUCUUCGAGAGGAUUGAGCC

>ghr-miR7489 MIMAT0029131

AUUGUUGCCAAUACAGGAGAACGU

>ghr-miR7509 MIMAT0029156

UCAAAAGCACUUUUUGACAGCAAU

>ghr-miR7488 MIMAT0029130

UUUUGAGUACAGGGGACAAAA

>ghr-miR160 MIMAT0029137

UAUGAGGAGCCAUGCAUGUAU

>ghr-miR7512 MIMAT0029161

UGCUACUUGUAGUUAUGCAUG

>gra-miR8787 MIMAT0034241

UUUUCUUUUAAUUGGACGAGAUA

>gra-miR8659a MIMAT0034028

UUUGGAAAGUUAUAAAAUGGUCAU

>gra-miR8681 MIMAT0034052

ACAUUGUUGAGGGUCUAAUCGG

>gra-miR7494b MIMAT0034062

AGAGGGAGAAGCAGAAGAGAAUA

>gra-miR8735 MIMAT0034147

GGGGACAAUACCUUCGAUUGUUGG

>gra-miR7486g MIMAT0034122

CGCUUUGCUGACGUGGAAACAAAU

>gra-miR8675b MIMAT0034084

AGGUGAUGAUGUGGUACAAUCUCA

>gra-miR8693 MIMAT0034073

AGGAUGAAAAUAUUGAUGUAGCAU

>gra-miR8677 MIMAT0034048

AAUGAAUCUAGGUUCUCUCUU

>gra-miR8690 MIMAT0034063

AGAGUGACUACUUCGUAACAAAAC

>gra-miR8778 MIMAT0034260

UUUCCAUAUUAGGGUUUGAACUUU

>gra-miR8680 MIMAT0034051

ACAGUGGAGGUAUUGUGCCUG

>gra-miR8784 MIMAT0034236

UUUGUCGACAUGUCAGGAAAGCGC

>gra-miR8742a MIMAT0034158

UAUCUUAUUCAUCUUGGACUG

>gra-miR7504g MIMAT0034069

AGGAGGAAAUCUGAUUUGUCAUUC

>gra-miR8785 MIMAT0034237

UUUUACAGCAGCUACAUCCAU

>gra-miR7484f MIMAT0034157

UAUCUAACGUGUAGGGACUAA

>gra-miR8771b MIMAT0034144

GGAUUGUCGUUAGGGAGGUAA

>gra-miR8669 MIMAT0034040

AAAGACGAAAGACAAAAUCUCAAU

>gra-miR8747 MIMAT0034171

UCCCAGUUGUAGUUGGUCGUUCGG

>gra-miR8786b MIMAT0034239

UUUUAGUGAUGUGGCAGAAAGAUG

>gra-miR8745 MIMAT0034166

UCAACGGAGUUGGGAGACAAA

>gra-miR7492d MIMAT0034016

UGGGCUUAGAUUUUUUGCGGCGUU

>gra-miR7493b MIMAT0034256

UUGGAUUGUUAAAAGGUUAAUUGU

UAGCACUGAAGAUGAUGAUGG

>gra-miR8651 MIMAT0034010

UUAAUGGAUCAUAACAGCAGGUAU

>gra-miR7504h MIMAT0034070

AGGAGGAAUAAGUCUGAUUUGUCA

>gra-miR7504c MIMAT0034027

AAGAGAAAAAAUCGGAUUUAUCAU

>gra-miR8783 MIMAT0034233

UUUGUACGUGGCGGGAGAUAU

AUUGGAGUGAAGGGAGCUCGA

>lus-miR398d MIMAT0031859

UGUGUUCUCAGGUCACCCCUC

>lus-miR156d MIMAT0027180

UGACAGAAGAAAGAGAGCAC

>lus-miR171h MIMAT0027219

AUGAGCCGAACCAAUAUCACU

>lus-miR171i MIMAT0027224

UUGAGCCGUGCCAAUAUCACG

>lus-miR169d MIMAT0027128

UAGCCAAGGAUGACUUGCCCA

>lus-miR395e MIMAT0027208

CUGAAGUGUUUGGAGGAACUC

>lus-miR159b MIMAT0027233

UUUGGAUUGAAGGGAGCUCUC

>lus-miR169l MIMAT0027215

CAGCCAAGGAUGACUUGCCGA

>lus-miR159c MIMAT0027235

UUUGGAUUGAAGGGAGCUCUU

>lus-miR171g MIMAT0027193

UGAUUGAGCCGCGUCAAUAUC

>lus-miR398f MIMAT0031861

GGUGUUCUCAGGUCGCCCCUG

>lus-miR397a MIMAT0027170

AUUGAGUGCAGCGUUGAUGAA

>lus-miR171a MIMAT0027122

AGAUUGAGCCGCGCCAAUAUC

>lus-miR166f MIMAT0027221

UCGGACCAGGCUUCAUUCCUU

>lus-miR166b MIMAT0027172

UCGGACCAGGCUUCAUCCCCC

>lus-miR399g MIMAT0027188

UGCCAAAGGAGAUUUGCCCAG

>lus-miR828a MIMAT0027220

UCUUGCUCAAAUGAGUGUUCCA

>lus-miR172j MIMAT0027234

GCAGCAUCAUCAAGAUUCCCA

>lus-miR396d MIMAT0027237

UCCCACAGCUUUAUUGAACUG

>mdm-miR2118b MIMAT0026035

CUACCGAUGCCACUAAGUCCCA

>mdm-miR159c MIMAT0026053

GAAUUCCUUCUCCUCUCCUUU

>mdm-miR156ad MIMAT0025896

UGACAGAAGAAAGUGAGCAC

>mdm-miR7121e MIMAT0026044

UCCUCUUGGUGAUCGCCCUGC

>mdm-miR395i-5p MIMAT0037471

GUUCCCUUGACCACUUCAUUG

>mdm-miR171i MIMAT0025946

UGAGCCGAACCAAUAUCACUC

>mdm-miR828b MIMAT0026030

UCUUGCUCAAAUGAGUAUUCCA

>mdm-miR11012b MIMAT0043615

CCGCUGCCUAUGCAAUGCACCC

>mdm-miR171m MIMAT0025950

UUGAGCCGUGCCAAUAUCACA

>mdm-miR10999b MIMAT0043592

GGGCGUGAUAUUCACACACCU

>mdm-miR169f MIMAT0026068

UGAAGAGAAGAGCGUUGUUUGG

>mdm-miR482a-3p MIMAT0011165

UUCCCAAGCCCGCCCAUUCCUA

>mdm-miR395k MIMAT0043586

GUUUCCUCAAACACUUCAUU

>mdm-miR3627d MIMAT0043534

UCCAUCCUCCUGUGACAUGAA

>mdm-miR319e MIMAT0043567

GAGCUUUCUUCAGUCCACUC

>mdm-miR408c MIMAT0026032

ACAGGGAAGAGGUAGAGCAUG

>mdm-miR319d MIMAT0043552

AACUGCCGACUCAUUCACUCA

>mdm-miR10989c MIMAT0043557

CAAAGCUUUUAAUAUCAGUCGA

>mdm-miR10980b MIMAT0043531

CACCUGGGACUUGCAGCCAUG

>mdm-miR393f MIMAT0026063

AUCAUGCGAUCCCUUCGGACG

>mdm-miR399g MIMAT0026007

UGCCAAAGGAGAUUUGCUCGG

>mdm-miR10993f MIMAT0043590

ACAUGUGGUGUACCAUCCUGU

>mdm-miR11001 MIMAT0043599

UACAAAGAUGGACUCCACCCUA

>mdm-miR10991c MIMAT0043563

CGAGCCAUUGAAAUUCGAUCC

>mdm-miR10984a-5p MIMAT0043547

GGUAAUUGACUGUGAAAUCGU

>mdm-miR169c MIMAT0025936

UAGCCAAGGAUGACUUGCCCG

>mdm-miR156s MIMAT0025885

CUGACAGAAGAUAGAGAGCAC

>mdm-miR7124a MIMAT0026054

CACCAAUAUCAACUUUAUUUG

>mdm-miR11002b MIMAT0043601

GAGGAUGAGCUUCGGCGGUGA

>mdm-miR10986 MIMAT0043551

UGGCACCAAAGUCACCACCCG

>mdm-miR530a MIMAT0043542

UGCAUUUGCACCUGCACUUGU

>mdm-miR399k MIMAT0043550

UGCCAAAGGAGAGUUGCCCUU

>mdm-miR10978b MIMAT0043528

GUUGGGAAUCGAAGCAUCACGA

>mdm-miR10997 MIMAT0043581

UAACCUUAUUUGAUUUCACGA

>mdm-miR535b MIMAT0026025

UGACAAGGAGAGAGAGCACGC

>mdm-miR10990 MIMAT0043560

CCAAGGAAAAUUUUAUGACGA

>mdm-miR396a MIMAT0025989

UUCCACAGCUUUCUUGAACAG

>mdm-miR10982c MIMAT0043537

CGGAAUGAAGCUUACGAGAAUG

>mdm-miR10981a MIMAT0043532

UGACCAACAUAUAUGGGCCGU

>mdm-miR169b MIMAT0025935

UAGCCAAGGAUGAUUUGCCUGC

>mdm-miR11007 MIMAT0043607

CCGUCAAUUAUUUUGGUACGU

>mdm-miR827 MIMAT0026028

UUAGAUGACCAUCAACGAACA

>mdm-miR10994-3p MIMAT0043573

UGCUUUUUUCUUGACCAUAGC

>mdm-miR11000 MIMAT0043598

GUGUUCCAAAGAAAUCCGGAGU

>mdm-miR10993b MIMAT0043571

AUCCCACCAUUUAUAUAGCGA

>mdm-miR7127a MIMAT0026064

AUACUCAUCGAAUUUGUCAUA

>mdm-miR11009 MIMAT0043611

AUGCACAACAAUAUGAGGGUGU

>mdm-miR10984b-3p MIMAT0043610

CUCACGUACGCUGUCCCGAGAA

>mdm-miR11014 MIMAT0043623

CGACCAUUCAUGAAAACUGCC

>mdm-miR11011a MIMAT0043613

AAGUUCAUUCAAACACCAUGU

>mdm-miR171q MIMAT0043605

GGAUAUUGGUCCGGUUCAAUA

>mdm-miR11020 MIMAT0043631

GACAUUACAACGGUUACACGG

>mdm-miR10996b MIMAT0043622

UCACCAUUGCAUCUCAUGUUCC

>mdm-miR5225a MIMAT0026056

UCUGUCGAAGGUGAGAUGGUGC

>mdm-miR397b MIMAT0025997

UUGAGUGCAGCGUUGAUGAAA

>mdm-miR10979 MIMAT0043529

CUUGCCGAUAGAUUUGGGGAG

>mdm-miR319h MIMAT0043595

GAGCUCUUCUUCAGUCCAGUCC

>mdm-miR482b MIMAT0026022

UCUUUCCUAUCCCUCCCAUUCC

>mdm-miR11017 MIMAT0043627

CUCUAAUCUGCCCAACAACUU

>mdm-miR10983 MIMAT0043546

CAGAGCAAAACAGUCGUGGAA

>mdm-miR11018 MIMAT0043628

CCAACAUCAAAAGUAAAAGGAA

>mdm-miR11015 MIMAT0043624

UGGGUAAAAUGACCCAACCUG

>mdm-miR10981d MIMAT0043594

AGCCGUUUAAUCAAAAUCCAA

>mdm-miR11011b MIMAT0043630

UAAGUUCAUCCAAACACCAUA

>mdm-miR319c-3p MIMAT0026058

AUCCAACGAAGCAGGAGCUGA

>mdm-miR482d MIMAT0026039

AAUGGAAGGGUAGGAAAGAAG

>mdm-miR11006 MIMAT0043606

CAAUGGGGAGGAGUCAUUCGUA

>mdm-miR10987 MIMAT0043553

CCAUAUGUCCCUCCAUAUACU

>mdm-miR477b MIMAT0026020

ACUCUCCCUCAAGGGCUUCGAC

>mdm-miR164a MIMAT0025907

UGGAGAAGCAGGGCACAUGCC

>mdm-miR11005 MIMAT0043604

CUACUAUAUGGUCGUACACAUC

>mdm-miR858 MIMAT0026070

UUCGUUGUCUGUUCGACCUGA

>mdm-miR172m MIMAT0025964

AGAAUCUUGAUGAUGCUGCAG

>mdm-miR7121c MIMAT0026042

UCCUCUUGGUGAUCGCCCUGU

>mdm-miR11004 MIMAT0043603

GUAUUCUUUCAUCUUCUACUA

>mdm-miR167j MIMAT0025931

UGAAGCUGCCAGCAUGAUCUUA

>mdm-miR7125 MIMAT0026059

CGAACUUAUUGCAACUAGCUU

>mdm-miR171o MIMAT0026066

UGGGAUGUUGGUAUGGUUCAA

>mdm-miR2111b MIMAT0026015

UAAUCUGCAUCCUGAGGUUUA

>mdm-miR10996a MIMAT0043580

ACACCAUCGCAUCUCAUGUUCC

>mdm-miR482a-5p MIMAT0011164

AGGAAUGGGCUGUUUGGGAAGA

>mdm-miR11013 MIMAT0043621

CAUGGGAAUUUUAAAGUCACCU

>mdm-miR7120a-3p MIMAT0037472

CAGUCUGACAAUAUAACGUGC

>mdm-miR395l MIMAT0043626

CUGAAGUGUUUGGGGGAACCC

>mdm-miR535d MIMAT0026027

UGACGACGAGAGAGAGCACGC

>mdm-miR7120b-5p MIMAT0026038

UGUUAUAUUGUCAGAUUGUCA

>mdm-miR7126-5p MIMAT0026060

AAAGUAUCAAGGAGCGCAAAG

>mdm-miR7128 MIMAT0026069

AUCAUUAACACUUAAUAACGA

>mdm-miR11010 MIMAT0043612

AGUAACUAUAGCUGUUUUCUA

>mdm-miR1511 MIMAT0026188

ACCUAGCUCUGAUACCAUGAA

>mdm-miR477a MIMAT0026021

ACUCUCCCUCAAGAGCUUCUC

>mdm-miR11003 MIMAT0043602

ACACAAUAUACGAUGAACAGA

>mdm-miR7122a MIMAT0026048

UUAUACAGAGAAAUCACGGUCG

>mdm-miR10988 MIMAT0043554

AGAGAAAAUUCAUUCCAACGC

>mdm-miR391 MIMAT0026019

UACGCAGGAGAGAUGACGCCG

>mdm-miR7126-3p MIMAT0037475

UUGCGUUCCACUGAUUCUUUCG

>mdm-miR11016 MIMAT0043625

UCCAUAAUUUUUCCAGAUCAA

>mdm-miR10992 MIMAT0043566

CAUAACAAAUUAUUACUCAGU

>mdm-miR5225c MIMAT0026052

UCUGUCGUGGGUGAGAUGGUGC

>mdm-miR7123a MIMAT0026050

AAGAGCGGGAUGUGUAAAAGG

>mdm-miR11002c-5p MIMAT0043596

CCCAGUCACCUCCGAAGCUCA

>mdm-miR482c MIMAT0026023

UCUUUCCUAACCCUCCCAUUCC

>mdm-miR10995 MIMAT0043579

CAAGCUUCCUCUUCAUACUCGU

>mdm-miR10994-5p MIMAT0043572

UAUGGUUAAGAAACAAGCAGA

>mdm-miR11008 MIMAT0043608

GUGACCGCACAAAAUAGAAGA

>mdm-miR10984a-3p MIMAT0043548

AGUCAAUUACCUCAUAAACUC

>mdm-miR10985 MIMAT0043549

CCACUCGUAGUGAAACAGUUG

>mdm-miR11019 MIMAT0043629

CCAGAUGAUCAUAAUCUCCUGA

>mdm-miR10998 MIMAT0043585

CUUGGGAUUCAGUCUAGGACUU

>nta-miR156f MIMAT0024634

UGACAGAAGAGAAUGAGCAC

>nta-miR6021 MIMAT0023588

UUGGAAGAGGCUGCUAUUGGA

>nta-miR6151c MIMAT0024744

UGAAUGUGAGGCAUUGGAUUGA

>nta-miR1446 MIMAT0024763

UGAACUCUCUCCCUCAAUGGCU

>nta-miR482c MIMAT0024727

UUUCCAAUUCCACCCAUUCCUA

>nta-miR168b MIMAT0024663

UCGCUUGGUGCAGGUCGGGAC

>nta-miR399e MIMAT0024717

CGCCAAAGGAGAGCUGCCCUG

>nta-miR6151i MIMAT0024750

UGAGUGUGAGGCGUUGGAUUGA

>nta-miR171a MIMAT0024686

UAUUGGUGCGGUUCAAUGAGA

>nta-miR6158a MIMAT0024766

AAGUUCGAUUUGUACGAAGGGC

>nta-miR6155 MIMAT0024757

UAAGGUUGCCUUGCUCUUGCA

>nta-miR6148b MIMAT0024738

UGUGUUAAUCGUUUGUUCUCA

>nta-miR6154b MIMAT0024756

UGGGUCUCCUGGAGAAAGGUC

>nta-miR6147 MIMAT0024736

UGACAUCUUCAAAACCCACUA

>nta-miR6025c MIMAT0024730

UCAAUUGAGAUGACAUCUAGU

>nta-miR6157 MIMAT0024760

UGGUAGACGUAGGAUUUGAAGA

>nta-miR6161b MIMAT0024775

GCUGGACCGGUAUACUUUGCUGAC

>nta-miR6152a MIMAT0024751

UAUUGUAUUCGACUGUAUUCACGG

>nta-miR6020a-3p MIMAT0023585

AGAUACUCAGCAAAACAUUUAC

>nta-miR6024 MIMAT0023595

UUUUAGCCAGAGUUGUUUUCCC

>nta-miR6145c MIMAT0024758

CAGUGCACAUAUAACAGUAA

>nta-miR6025a MIMAT0023606

UACCAACAAUUGAGAUAACAUC

>nta-miR6025e MIMAT0024733

UGCCAAUUAUAGAGAUGACAUC

>nta-miR6145d MIMAT0024764

AUUGUUACAUGUAACACUGGC

>nta-miR6146b MIMAT0024735

UUUGUCCAAUGAAAUACUUAUC

>nta-miR6148a MIMAT0024737

UACGUCGAUCGAUUGUUCUUA

>nta-miR6163 MIMAT0024779

UGGAAGUACUGCCUAAGUUUGA

>nta-miR6019a MIMAT0023583

UACAGGUGACUUGUAAAUGUUU

>nta-miR482d MIMAT0024776

UUCCCGACUCCCCCCAUACCAC

>nta-miR6161d MIMAT0024773

UGAACUCCAGCAUAUUAUACU

>nta-miR6149b MIMAT0024740

UUGAUACGCACCUGAAUCGGC

>nta-miR479a MIMAT0024723

CGUGAUAUUGGUUUGGCUCAUC

>nta-miR6150 MIMAT0024741

AGAUUUGUUUGAUCGUCUUGGC

>nta-miR6144 MIMAT0024729

UGGCAACUUCUUCAUCAUGCC

>nta-miR6160 MIMAT0024772

GCAUAUAUGGGCCAACUGUGUAAC

>nta-miR6020b MIMAT0023587

AAAUGUUCUUCGAGUAUCUUC

>nta-miR5303c MIMAT0024770

ACGGGUGCGGCUACAUUUUGG

>nta-miR172b MIMAT0024690

AGAAUCAUGAUGAUGCUGCAU

>nta-miR6145f MIMAT0024771

AUCGUAACAUAUAGCACUAGC

>nta-miR6156 MIMAT0024759

UUGAAGAUGUUCUAUUUCUGU

>nta-miR6162 MIMAT0024777

AGAAAAAUGGUAGCCAUUGGA

>nta-miR6164a MIMAT0024780

UCACAUAAAUUGAAACGGAGG

>nta-miR6145b MIMAT0024753

UUAUCAUACGUAGCACUAGCC

>nta-miR5303a MIMAT0024761

AAAAUGUGGCCGGAUACGUGU

>nta-miR482b-3p MIMAT0023605

UCUUGCCAAUGCCAUCCAUUCC

>nta-miR6145a MIMAT0024731

CAUUUUCACAUGUAGCACUGAC

>nta-miR160d MIMAT0024643

UGCCUGGCUCCCUGCAUGCCA

>nta-miR477b MIMAT0024722

UCUCUCCCUCAAGGGCUUCUC

>nta-miR6025b MIMAT0023607

UGCCAACUAUUGAGAUGACAUC

>nta-miR479b MIMAT0024724

CGUGAUAUUUGUUUGGCUCAUC

>nta-miR6161c MIMAT0024778

AAUAUACUGGAGUUCGGUGCACCU

>nta-miR6153 MIMAT0024754

UAGGACCAUAUUCACUAUUUG

>nta-miR6146a MIMAT0024734

UUUGUCCAAUGAAACACUUAUC

>nta-miR482b-5p MIMAT0023604

AGUGGGUGGAGUGGUAAGAUA

>nta-miR6151f MIMAT0024747

UGAGUGUGAGGCAUUGGAUUGA

>nta-miR827 MIMAT0024725

UUAGAUGAACAUCAACAAACA

>nta-miR6159 MIMAT0024769

UAGCAUAGAAUUCUCGCACCUA

>nta-miR390a MIMAT0024701

AAGCUCAGGAGGGAUAGCACC

>nta-miR6025d MIMAT0024732

AACAAUUGAGAUAACAUCUAGG

>nta-miR6020a-5p MIMAT0023584

AAAUGUUUUUCGAGUAUCUUC

>nta-miR1919 MIMAT0024726

GAGCGAGUCAUCUGUGACAGG

>nta-miR6145e MIMAT0024765

AUUGUUACAUGUAGCACUGGC

>pla-miR11609 MIMAT0045462

UAGCUUGGUGUGAGGUCAACUU

>pla-miR11603 MIMAT0045456

UUAUAAUUAGGUUGAGCGGAC

>pla-miR11600 MIMAT0045453

AACGUUCCUCGAUUUCGCGAU

>pla-miR11612 MIMAT0045465

AAGACGGUCCAAAACGCCCAC

>pla-miR11598 MIMAT0045451

UCGUUCAAAGUAGGUUGUCAA

>pla-miR11608 MIMAT0045461

UCGCUUAGGGGUUGUUGAAGCGC

>pla-miR11601 MIMAT0045454

UGCUCUAAAAGAUCGUAGUUC

>pla-miR11610 MIMAT0045463

ACAGAUAUGGUAGGGGGCACA

>pla-miR11606 MIMAT0045459

UGGUGGACUCCAAUUCGCAUA

>pla-miR11599 MIMAT0045452

UUCAACCGUGGUAGAUGUUAA

>pla-miR11607 MIMAT0045460

GAUCACUCGGUUGUCUGACACAC

>pla-miR11604 MIMAT0045457

UCCAGAGGGAGAACGUGGCGA

>pla-miR11611 MIMAT0045464

UAGGCAACCGUGGUAAAAUGUC

>pla-miR11602 MIMAT0045455

UCUAACGGAACGCUAUUGGAUC

>pla-miR11605 MIMAT0045458

UUGAGGCGGCAUAUUCUCAAU

>peu-miR2916 MIMAT0012884

UGGGGACUCGAAGACGAUCAUAU

>peu-miR2912b MIMAT0012880

UCUAGAACUCGAGAUAUGGGC

>peu-miR2913 MIMAT0012881

GAGGUCGGGGAUUGCAAGGAG

>ptc-miR475d-3p MIMAT0002070

UUACAGAGUCCAUUGAUUAAG

>ptc-miR1447 MIMAT0006009

CAGAAUUGCAGUGCCUUGAUU

>ptc-miR482b-3p MIMAT0025271

UUACCAAUACCUCUCAUGCCAA

>ptc-miR169o MIMAT0001971

AAGCCAAGGAUGACUUGCCUG

>ptc-miR169b-3p MIMAT0022898

GGCAGGUUGUUCUUGGCUAC

>ptc-miR7834 MIMAT0030406

UAAUAAAAUCUCGACUAUUAU

>ptc-miR397c MIMAT0002040

UCAAUGAGUGGAGCUUUGAUG

>ptc-miR6434 MIMAT0025237

CUAUGUAUGCUUGACCAAUCA

>ptc-miR156l MIMAT0025268

UUGACAGAAGAUGGAGAGCAC

>ptc-miR481b MIMAT0002099

AGGACCUCACUUAACAGCUUAAGC

>ptc-miR169u-3p MIMAT0022900

GGCAGUCUCCUUUGGCUAUCC

>ptc-miR167h-3p MIMAT0022895

AGAUCAUGUGGCAGUUUCACC

>ptc-miR6467 MIMAT0025196

AUAAGCUGUCGAGCGUUUUGU

>ptc-miR482a.2 MIMAT0006785

UCUUGCCUACUCCUCCCAUU

>ptc-miR477e-5p MIMAT0002062

ACUCUCCCUCAAGGCUUCCA

>ptc-miR6442 MIMAT0025245

CUGAACGGCUUGAAGGGCACG

>ptc-miR160e-3p MIMAT0022892

GCAUGAGGGGAGUCGAGCAGG

>ptc-miR478l MIMAT0002086

UAACGUGUCUCCUAUUUUUAGGGA

>ptc-miR478f MIMAT0002081

UGACAUGUCUUCUAUUUUUAGGGA

>ptc-miR6430 MIMAT0025231

UGAUGAUUAAUUGACUGCAAA

>ptc-miR171l-5p MIMAT0002095

UGUGAUAUUGGUCCGGCUCAUC

>ptc-miR6437b MIMAT0025225

UCACGGACGGCGGCUCCAAGCA

>ptc-miR6428 MIMAT0025229

UCUGGCAACUCAUUAGACUCAU

>ptc-miR6478 MIMAT0025220

CCGACCUUAGCUCAGUUGGUG

>ptc-miR6421-3p MIMAT0025181

UAGAGCAGAUUGUAAGGGAAG

>ptc-miR6422 MIMAT0025236

UGUGAUAAUGAAGGCUAUGGU

>ptc-miR482d-3p MIMAT0025235

UUGCCGACCCCACCCAUGCCAA

>ptc-miR399e MIMAT0002055

CGCCAAAGGAGAGUUGCCCUC

>ptc-miR1444c MIMAT0006001

UUCACAUUCGGUCAACGUUC

>ptc-miR403c-5p MIMAT0022932

UUUGUGCGUGGAUCUGAGGCC

>ptc-miR6440c MIMAT0025200

GAGUUUGAUCGAUUUCGAGUU

>ptc-miR6459b MIMAT0030396

UCUCAAGCCCAAAUUCGAUC

>ptc-miR7817b MIMAT0030379

UCUCUUCUGUUCCUGAACGGU

>ptc-miR6429 MIMAT0025230

UAGUAGAAAUGCAUUGACUAG

>ptc-miR6438b MIMAT0025197

UUGUACACAGAAUAGGUGAAAU

>ptc-miR6461 MIMAT0025187

UAGCUAGCAAGUUCAUGGAUC

>ptc-miR169n-3p MIMAT0022899

GCAAGCAUCCUUGGUUCUCC

>ptc-miR169ac MIMAT0001954

UAGCCAAGGACGACUUGCCCA

>ptc-miR6474 MIMAT0025213

UGUUCAGAUCAGUAGAUAGCA

>ptc-miR7821 MIMAT0030402

AGAUGGGCAUCGGCAUUGUGA

>ptc-miR6423 MIMAT0025252

CCGCUGUCGCCACUAUCUUCCU

>ptc-miR398c-5p MIMAT0022912

GGAGCGACCUGAAAUCACAUG

>ptc-miR6476a MIMAT0025217

UCAGUGGAGAUGAAACAUGA

>ptc-miR477d-3p MIMAT0025276

UGGACUCCUUUGGGGAGAUGG

>ptc-miR6425e MIMAT0025219

UUGUCUUCCAUGGAAUAGGCAG

>ptc-miR7815 MIMAT0030373

CUCUUCAAAUAAAUCGUGGGA

>ptc-miR7812 MIMAT0030371

CUGUUAUGAAUUGAUGGAGUG

>ptc-miR6447 MIMAT0025250

UUGACGAAAUGUGACGACUAC

>ptc-miR1448 MIMAT0006010

CUUUCCAACGCCUCCCAUAC

>ptc-miR7829 MIMAT0030404

ACACAGAAACUCCAAGCCCAC

>ptc-miR478a MIMAT0002076

UGACGUGUCUUCUAUUUUUAGGGA

>ptc-miR7819 MIMAT0030401

UCUUGAGAACAUGAUGAAUCG

>ptc-miR477a-3p MIMAT0022918

GGAUGCCUUUGGGGGAGAUUG

>ptc-miR481d MIMAT0002102

AGGACCUCACCUAACAGCUUAAGC

>ptc-miR475a-3p MIMAT0002067

UUACAGUGCCCAUUGAUUAAG

>ptc-miR7835 MIMAT0030407

GAUGGGAUUUUUCGGGAAGUG

>ptc-miR6463 MIMAT0025189

UGGAUGAUCAUGUUGGCAACC

>ptc-miR1450 MIMAT0006012

UUCAAUGGCUCGGUCAGGUUAC

>ptc-miR474b MIMAT0002065

CAAAAGUUGUUGGGUUUGGCUGGG

>ptc-miR482c-5p MIMAT0025277

UAUGGGAGAGGCGGGAAUGACU

>ptc-miR477f MIMAT0002063

GCUCUCCCUCAGGGCUUCCA

>ptc-miR480 MIMAT0002096

ACUACUACAUCAUUGACGUUGAAC

>ptc-miR6457b MIMAT0025203

UUAGUUUGGCAGCCUCUUCUC

>ptc-miR394b-3p MIMAT0006784

CUGUUGGUCUCUCUUUGUAA

>ptc-miR6436 MIMAT0025239

CCAGACUCAAUAGCAGGACCCA

>ptc-miR7817a MIMAT0030377

UUUGGUUAUUGUCUCGAGACA

>ptc-miR6427-3p MIMAT0025228

GUGGGAAUGAACAUUAUGAGA

>ptc-miR1449 MIMAT0006011

UGAGGUGCACGUAAGAUAACUC

>ptc-miR408-5p MIMAT0022913

CGGGGAACAGGCAGAGCAUGG

>ptc-miR6462c-3p MIMAT0025206

UCUCUUAUGCAUUUUUGUCCC

>ptc-miR169z MIMAT0001982

CAGCCAAGAAUGAUUUGCCGG

>ptc-miR1444e MIMAT0030386

CGAACGUUGACCGAAUGUGAA

>ptc-miR7830 MIMAT0030389

UGAUCUAGAGAACCGUUGCU

>ptc-miR6468-5p MIMAT0027337

GUUUUCCCUGAAUCACUCCCA

>ptc-miR6421-5p MIMAT0025180

UCCCUUACAAUCUACUCUUUC

>ptc-miR6472 MIMAT0025211

UAGUGAAUUCUAGGUCUCAAUC

>ptc-miR475d-5p MIMAT0022917

AAUGGCCAUUGUAAGAGUAGA

>ptc-miR477c MIMAT0025257

GGAAACCUUUUGUGGGGGUUUG

>ptc-miR6457a MIMAT0025267

UAAUCUCUCUGCAGAAUGCUG

>ptc-miR6451 MIMAT0025255

AUGUCAGAUCAUGUUAGGUAU

>ptc-miR476b MIMAT0002072

UAGUAAUUCUUCUUUGCAAAA

>ptc-miR390d-3p MIMAT0022907

CGCUAUCCAUCCUGAGUUUUA

>ptc-miR7828 MIMAT0030388

GAUGACAUGGACACCAAAAUC

>ptc-miR7839 MIMAT0030395

AGUGGCAUUGGAGGUAUCCC

>ptc-miR6456 MIMAT0025266

UUGAGUCCUUCCAUUAGAUCC

>ptc-miR159d MIMAT0001904

CUUGGAUUGAAGGGAGCUCCU

>ptc-miR7824 MIMAT0030383

UUGAGAAAAGUCAAUCGGACC

>ptc-miR482b-5p MIMAT0027338

GGCAUGAGGUGUUUGGCAAGA

>ptc-miR6445b MIMAT0025256

UUCAUUCCUCUUCCUAAAAUGG

>ptc-miR474a MIMAT0002064

CAAAAGUUGCUGGGUUUGGCUGGG

>ptc-miR6470 MIMAT0025209

CUCUGAUAUCAUAUUAAAAAA

>ptc-miR7822 MIMAT0030381

UUUGAAAUUGAACAAAUGGUA

>ptc-miR477b MIMAT0002075

AUCUCCCUCAGAGGCUUCCAA

>ptc-miR399j MIMAT0002053

UGCCAAAGGAGAUUUGUCCGG

>ptc-miR472a MIMAT0002060

UUUUCCCUACUCCACCCAUCCC

>ptc-miR399h MIMAT0002051

UGCCAAAGGAGAGUUUCCCUG

>ptc-miR7814 MIMAT0030372

UAGAUUGUUUUUAUGCUUUGA

>ptc-miR6448 MIMAT0025251

UAGGCACAGAAUUAACAAGGC

>ptc-miR169s MIMAT0001975

UCAGCCAAGGAUGACUUGCCG

>ptc-miR475c MIMAT0002069

UUACAAUGUCCAUUGAUUAAG

>ptc-miR472b MIMAT0002061

UUUUCCCAACUCCACCCAUCCC

>ptc-miR7818 MIMAT0030378

UUUCUUAUCGAUCACUAGACG

>ptc-miR7840 MIMAT0030397

CAAGGAGUAAUUAGUGACAUC

>ptc-miR6475 MIMAT0025216

UCUUGAGAAGUAAAGAACGAC

>ptc-miR6449 MIMAT0025253

CAUGAUUCUGAAUAACGGUUU

>ptc-miR6431 MIMAT0025232

UUAUGUGGCAUAAAAGAAUCAA

>ptc-miR6460 MIMAT0025186

UGAUAUGUGGCAUUCAAUCGA

>ptc-miR478e MIMAT0002080

UGACGAGUCUUCUAUUUUUAGGGA

>ptc-miR171k MIMAT0003943

GGAUUGAGCCGCGCCAAUAUC

>ptc-miR6473 MIMAT0025212

UCCACAAUCCCAUCAAGACUU

>ptc-miR7836 MIMAT0030408

UGGGUGGGAGGUGUGGUAGCU

>ptc-miR169q MIMAT0001973

UAGCCAAGGACGACUUGCCUG

>ptc-miR482c-3p MIMAT0025278

UCUUUCCGAGUCCUCCCAUACC

>ptc-miR478p MIMAT0002090

UAACGUGUCUUCUAUUUUUAGGGA

>ptc-miR6444 MIMAT0025247

UAGGAAAAUUAGAGAUUAUGU

>ptc-miR172i MIMAT0002001

AGAAUCCUGAUGAUGCUGCAA

>ptc-miR7831 MIMAT0030390

UACAUGUAGAGACCACCAAAC

>ptc-miR6458 MIMAT0025182

ACGCUCAAAUAUAUAAGGUGUU

>ptc-miR7838 MIMAT0030394

ACAUGUGCUGGGUAGGAGGAA

>ptc-miR399d MIMAT0002047

UGCCAAAGAAGAUUUGCCCCG

>ptc-miR6465 MIMAT0025193

UUAAAGCAGGGGGUCUAGUAGU

>ptc-miR7827 MIMAT0030403

GCUAGGACCAAGUUUUUUGGA

>ptc-miR6462c-5p MIMAT0025205

AAGGGACAAAAAUGGCAUAAGA

>ptc-miR6480 MIMAT0025223

UUGCUGAAACGAUUGAACUAU

>ptc-miR6455 MIMAT0025265

UCAAAUAGCAUCCUCAACAUU

>ptc-miR7823 MIMAT0030382

UUGCAUGCAUGAACUUGAAAU

>ptc-miR6450b MIMAT0025261

CGAACACAGGACUCAAGGCUA

>ptc-miR6432 MIMAT0025233

CGGCUCUAGAGAAAAAGAUGG

>ptc-miR7842 MIMAT0030399

UAAAUCAAGCCCGGUACUUUU

>ptc-miR6462e MIMAT0030375

UCUUAUGCGUUUUUGUCUCU

>ptc-miR477d-5p MIMAT0025275

AUCUCCCUCAAAGGCUUCCUCU

>ptc-miR478d MIMAT0002079

UGACAUGUCUUCUAUUUUUAGUAA

>ptc-miR536 MIMAT0025260

CGCUCGCCAGCGUUGCACCACC

>ptc-miR6479 MIMAT0025222

AUCAAUGAGAAUACUACUGCA

>ptc-miR7826 MIMAT0030387

UUACCAAGUUUCAAAUUCUCA

>ptc-miR169y MIMAT0001981

UAGCCAUGGAUGAAUUGCCUG

>ptc-miR6439b MIMAT0025258

AGCAGAAGCCAUCACUAGCGC

>ptc-miR171a-5p MIMAT0022901

GGAUAUUGGUACGGUUCAAUC

>ptc-miR166p MIMAT0001939

UCGGACCAGGCUCCAUUCCUU

>ptc-miR3627a MIMAT0025263

UCUGUCGCUGGAAAGAUGGUAC

>ptc-miR167h-5p MIMAT0001948

UGAAGCUGCCAACAUGAUCUG

>ptc-miR1444a MIMAT0005999

UCCACAUUCGGUCAAUGUUC

>ptc-miR6452 MIMAT0025262

UGCAAAGGUAACUGAGACAAU

>ptc-miR6425b-3p MIMAT0025199

UCCAUGGAAGAUAAUGACUCG

>ptc-miR828b-3p MIMAT0025281

UCAUUCGAGCAAGAAAUAUUA

>ptc-miR169aa MIMAT0001952

GAGCCAAGAAUGACUUGUCGG

>ptc-miR7813 MIMAT0030400

UGGUAAUGCAAGUGUUGCUAA

>ptc-miR6459a-5p MIMAT0025184

AGCUCAAGCACAAAUUCGAUC

>ptc-miR6441 MIMAT0025244

AAGUUGACGGAAGGACUACAU

>ptc-miR171g-5p MIMAT0022902

UGUUGGGAUGGCUCAAUCAUG

>ptc-miR169u-5p MIMAT0001977

UAGCCAAGGACGACUUGCCUA

>ptc-miR6426b MIMAT0025259

GUGGAGACAUGGAAGUGAAGA

>ptc-miR6469 MIMAT0025202

UGGCAGAAAAGGAUUCGUUUA

>ptc-miR6450a MIMAT0025254

CUUUGUCAGGACUCAAGGCUA

>ptc-miR6443 MIMAT0025246

GUAUGAUCAUAGAUGAUGGAG

>ptc-miR171h-5p MIMAT0022903

UGUUGGGAUGGCUCAAUCAUA

>ptc-miR7833 MIMAT0030392

UAAUUAGAACUCAUACUAGAC

>ptc-miR6464 MIMAT0025192

UGAUUGCUUGUUGGAUAUUAU

>ptc-miR6427-5p MIMAT0025227

UCGUAAUGCUUCAUUCUCACAA

>ptc-miR3627b MIMAT0030391

UGUCGCAGGAGAGAUGGCGCUA

>ptc-miR7832 MIMAT0030405

UAGUUCCCAACCUACACCACA

>ptc-miR319i MIMAT0002010

UUGGGCUGAAGGGAGCUCCC

>ptc-miR7841 MIMAT0030398

GGGGGUUGCUGUCAAGCAUAA

>ptc-miR396g-5p MIMAT0002037

UUCCACGGCUUUCUUGAACUU

>ptc-miR477e-3p MIMAT0022914

UGAGGCCUUUGGGGGAGAGUGG

>ptc-miR169ag MIMAT0025269

AAGCCAAGGGUGACUUGCCUGA

>ptc-miR7816 MIMAT0030374

AAUGUUGUUAUUAACACUGUA

>ptc-miR6477 MIMAT0025218

UGAACAGUAGACGUGAAUUAU

>ptc-miR6454 MIMAT0025264

CUUGUAACCUGAGUAGAGGCA

>ptc-miR159e MIMAT0001905

CUUGGGGUGAAGGGAGCUCCU

>ptc-miR482d-5p MIMAT0025234

GGACAUGGGUUGGUUUGCAAGA

>ptc-miR474c MIMAT0002066

CAAAAGCUGUUGGGUUUGGCUGGG

>ptc-miR6466-3p MIMAT0025195

UAUCAAUCAUCAAAUGUUCGU

>ptc-miR7825 MIMAT0030384

UUGAAGAAAGGUAGACAGAUAG

>ptc-miR6446 MIMAT0025249

UUGCUGGGUCCUGAUGAUGGA

>ptc-miR169ab MIMAT0001953

CAGCCAAGGAAGACUUGCCC

>ptc-miR6439a MIMAT0025242

CCAAAUGAAGAAGCAGAAGCC

>ptc-miR6471 MIMAT0025210

UUUGGGAUCAUCAGGACAGCC

>ptc-miR6435 MIMAT0025238

UGAAUAAUGGAGACACUCUAG

>ptc-miR1445 MIMAT0006002

UCCCUUGUAGACUAGAAAAA

>ptc-miR6459a-3p MIMAT0025185

UCGAAUUUGGGCUUGAGAUUG

>ptc-miR827 MIMAT0006003

UUAGAUGACCAUCAACGAAAA

>ptc-miR396g-3p MIMAT0022911

CUCAAGAAAGCCGUGGGAAAA

>ptc-miR6468-3p MIMAT0025201

GGAGUGAUUCAGGGAACCCAU

>ptc-miR476a MIMAT0002071

UAGUAAUCCUUCUUUGCAAAG

>ptc-miR169x MIMAT0001980

UAGCCAAGGAUGACUUGCUCG

>ptc-miR399c MIMAT0002054

UGCCAAAGGAGAUUUGCUCAC

>ptc-miR6466-5p MIMAT0025194

UCUGGUAUGAGCAUUUGAUGA

>ptc-miR7820 MIMAT0030380

AUCAUAUAGGUUGAUCCUCGU

>ppe-miR169f MIMAT0031459

UAGCCAAGGAUGACUUGCCUGC

>ppe-miR8122-5p MIMAT0031523

UUCCACAGAUCUUUCCUCAUU

>ppe-miR399h MIMAT0031504

UGCCAAAGAAGAGUUGCCCUA

>ppe-miR8130-3p MIMAT0031557

CCCUUCCAGUAAGGCACCCCC

>ppe-miR3627-5p MIMAT0031534

UCGCAGGAGAGAUGGCACUGUC

>ppe-miR6295 MIMAT0027331

GAGGACAGAAGAUGAUUCAGC

>ppe-miR6287 MIMAT0027319

CAAGAAGUGGAAGUUUUGGGC

>ppe-miR6273 MIMAT0027299

AAUGCAGCAUGAUUUUUUUUU

>ppe-miR396a MIMAT0031494

UUCCACAGCUUUCUUGAACGU

>ppe-miR169l MIMAT0031465

GAGCCAAGGAUGAAUUGCCGG

>ppe-miR319b MIMAT0027303

UAGCUGCCGAGUCAUUCAUCCA

>ppe-miR6288c-5p MIMAT0031565

CUUGUUAUUUUUUAAUUGAUU

>ppe-miR393a MIMAT0031474

CAUCCAAAGGGAUCGCAUUGA

>ppe-miR398b MIMAT0027328

CGUGUUCUCAGGUCGCCCCUG

>ppe-miR6288b-3p MIMAT0031561

UCAAUUAGAAAAUGAUAAGUG

>ppe-miR8123-3p MIMAT0031526

UUGUGCCAUUGCUCAAGC

>ppe-miR6267c-5p MIMAT0031542

AUUGCUGAUCACCUCUCUAAU

>ppe-miR6265 MIMAT0027284

UUGAACUUUGACCCGAUUCGCAU

>ppe-miR8129-5p MIMAT0031554

GAUAUCCGCACAUUAUUAUUG

>ppe-miR6292 MIMAT0027327

UAUCUUUUAAUCGUUAGAUCA

>ppe-miR8124-5p MIMAT0031531

ACUUGGUAUCUUGGUGCCGGU

>ppe-miR6293 MIMAT0027329

UAAGAGGCUGAUGACUAAAAC

>ppe-miR482d-3p MIMAT0027315

CCUCCCAUGCCACGCAUUUCUA

>ppe-miR482a-5p MIMAT0031165

GGGUGAGAGGUUGCCGGAAAGA

>ppe-miR8126-3p MIMAT0031539

UUCAGUAUUUUGACUCAGAA

>ppe-miR8127-5p MIMAT0031546

CAACUGUGGACAUACCCUUUG

>ppe-miR395a-3p MIMAT0031478

CUGAAGUGUUUGGGGGGACCC

>ppe-miR6276 MIMAT0027305

AAAGGCUCAUACAAAUAUUCC

>ppe-miR6261 MIMAT0027277

AAGUGAUUAUAUGGAGAAGCAC

>ppe-miR6269 MIMAT0027293

UGUGAAUAGUGAUUGCCAUGG

>ppe-miR8124-3p MIMAT0031532

UGGCACCAAUGAUACCAAGUUU

>ppe-miR482c-3p MIMAT0031169

UUCCCAAGCCCGCCCAUUCCAA

>ppe-miR6282 MIMAT0027312

GUUGAUCGAUGUGGGAUGUUACA

>ppe-miR8130-5p MIMAT0031556

GGGUUCCUUGUUGGAAGGACU

>ppe-miR3627-3p MIMAT0031535

UGGUGUCAUCCCUCCUGUGACC

>ppe-miR6274b-5p MIMAT0031552

AUUUCGACUAAUAACACAAUG

>ppe-miR6272 MIMAT0027298

UAGCUGUAAAUGAGUGUUUUU

>ppe-miR6257 MIMAT0027270

UCUUAACUGUUGGAUUAGGCU

>ppe-miR6259 MIMAT0027275

UAGAAAAAUACGGGCGAUAAA

>ppe-miR6270 MIMAT0027295

UUCUGGUAUUGGAAUUUCAUU

>ppe-miR477b-3p MIMAT0027288

GUUGGGGGCUCUUUUGGGACG

>ppe-miR1511-3p MIMAT0031522

ACCUGGCUCUGAUACCAUAAC

>ppe-miR8133-5p MIMAT0031563

UCCUGUGCGAACGUCCAGAAG

>ppe-miR477-3p MIMAT0031541

CGAAGCCUUUGGGGAGAGUAA

>ppe-miR8126-5p MIMAT0031538

UCUGAGUCAGAUUACUGAAUA

>ppe-miR7122a-5p MIMAT0031527

UUAUACAAUGAAAUCACGGCCG

>ppe-miR6284 MIMAT0027314

UUUGGACCAUGGAUGAAGAUU

>ppe-miR477b-5p MIMAT0031171

UCCCUCAAGGGCUCCCAAUAUU

>ppe-miR482d-5p MIMAT0031174

GAGAUGGGUGGCUGGGAAGGA

>ppe-miR6297b MIMAT0027336

GAUGUAUUGUCGUCGCGCAAAGU

>ppe-miR6267b MIMAT0027297

AUUAGAGAGGCGGUAAACAAU

>ppe-miR172a-5p MIMAT0031468

GUAGCAUCAUCAAGAUUCACG

>ppe-miR8128-3p MIMAT0031551

UCGUGGGGAGAGAUCUAAUCG

>ppe-miR8128-5p MIMAT0031550

AUUAGACCUCUCCCGACGAAA

>ppe-miR5225-3p MIMAT0031549

UCAUCUCUCCUCGACUGAA

>ppe-miR6260 MIMAT0027276

UGGAGUGAGAGAAUGGGAGGU

>ppe-miR8123-5p MIMAT0031525

UGAGCAAUGGCACACAGCCCU

>ppe-miR6266a MIMAT0027290

UAAAUGCAGGGGCAAAAUGAU

>ppe-miR6274a MIMAT0027301

UAUUUUGCUAUCUUCGGGCAAUA

>ppe-miR482e MIMAT0027318

UUGCCUAUUCCUCCCAUGCCAA

>ppe-miR8132 MIMAT0031562

UCCAACGAUGGGUGACCACAA

>ppe-miR6288a MIMAT0027320

GAAAAUGACAAGUGGCUAGUU

>ppe-miR6263 MIMAT0027280

AAGUGGACAAAAGGGGAGUGG

>ppe-miR6291c-5p MIMAT0031544

CCACAUUUAUAGAUUACCUUG

>ppe-miR6294 MIMAT0027330

UGGUGUAGGCUAAUCACAAUC

>ppe-miR395a-5p MIMAT0031477

GUUCCCUCAAACACUUCAUU

>ppe-miR7125-5p MIMAT0031536

GCUAGGUGCAACAAGUUCAAU

>ppe-miR399a MIMAT0031497

CGCCAAAGGAGAGUUGCCCUU

>ppe-miR482a-3p MIMAT0027272

UUUCCGAAACCUCCCAUUCCAA

>ppe-miR6280 MIMAT0027310

UUGGCAGUAAGAUUUUUGGUG

>ppe-miR6296 MIMAT0027332

UAAGGCCCUUAGAUGAGACCC

>ppe-miR8122-3p MIMAT0031524

UGAAGGAAGAUUUGUGGAAAG

>ppe-miR6275 MIMAT0027302

AGUGGAAGUAGCAAGGGGAAGC

>ppe-miR6285 MIMAT0027316

UAGUGAAGUUUGAAUUAGGGCU

>ppe-miR6271 MIMAT0027296

UCAAGAUUGAGAGAUAUAAUG

>ppe-miR6297a MIMAT0027335

AAUAAUUUUUCGUCGCGCAAAAU

>ppe-miR6281 MIMAT0027311

GUUAGAGAUAGAGAGAGUGAG

>ppe-miR482b-3p MIMAT0027273

CUUCCCAAACCUCCCAUUCCUA

>ppe-miR6258 MIMAT0027271

UUCCAGCUGUAAAGAUCAAGA

>ppe-miR6278 MIMAT0027307

UGAACCUUGUGUACAAAUUGGC

>ppe-miR5225-5p MIMAT0031548

UCUGUCGUAGGAGAGAUGGCGC

>ppe-miR169d MIMAT0031458

UGAGCCAAGGAUGACUUGCCA

>ppe-miR477-5p MIMAT0031540

ACUCUCCCUCAAAGGCUUCUAG

>ppe-miR8133-3p MIMAT0031564

UAACUUCCGAACGUCCGCAUA

>ppe-miR6291c-3p MIMAT0031545

CAAGGUAGUUUAUAAAUGUGG

>ppe-miR6288c-3p MIMAT0031566

AACCAAUUAGAAAAUAACAAGUGG

>ppe-miR7122a-3p MIMAT0031528

GCCGUGUUUCUUUGUAUAAAG

>ppe-miR6286 MIMAT0027317

UUUGAACCAUUGGAUCGUAGUUA

>ppe-miR8127-3p MIMAT0031547

UUCAAAGGGUACAUCCACAGU

>ppe-miR6262 MIMAT0027278

UCUUUAGAAAGUUAGAAUUGU

>ppe-miR482f MIMAT0031512

UCUUUCCUACUCCACCCAUUCC

>ppe-miR6290 MIMAT0027325

UGAAUGAGUUCAGAGAUCGUGUA

>ppe-miR6291a MIMAT0027326

CUUACCACAUUUUUAUACCAU

>ppe-miR6283 MIMAT0027313

CAAAAGGGGAGUGGGAAAAUC

>ppe-miR828-3p MIMAT0027282

UCAUUUCAGCAAGCAGCGUUA

>ppe-miR6264 MIMAT0027281

AUGCCUAUGGACACGUGUCAA

>ppe-miR1511-5p MIMAT0031521

CGUGGUAUCAGAGUCAUGUUA

>ppe-miR6279 MIMAT0027308

UAGACAAGAAUUCCAGAGACC

>ppe-miR8131-3p MIMAT0031559

AAUCAACUCAGCUUAGCUGAACUG

>ppe-miR171d-5p MIMAT0031466

UGUGAUAUUGGUUCGGUUCAUA

>ppe-miR171e MIMAT0027289

UUAUUGAACCGGACCAAUAUC

>ppe-miR6277 MIMAT0027306

UGUGUGUGGAAAGAGCGAGAC

>ppe-miR7122b-3p MIMAT0031530

CCGUGUUUCCUUGUAUAAAG

>ppe-miR482c-5p MIMAT0027283

GGAAUGGGCUGUUUGGGAUG

>ppe-miR8125 MIMAT0031533

CAGGAAAGAAUGUGAUGAGUA

>ppe-miR8129-3p MIMAT0031555

AUAAUAAUGUCCGGAUGUCAA

>ppe-miR7122b-5p MIMAT0031529

UUAUACAAUGAAAUCACGGUCG

>ppe-miR8131-5p MIMAT0031558

AUUUCAGCUAAGUUGAGUUGU

>ppe-miR398a-5p MIMAT0031173

GGAGCGACCUGGGAUCACAUG

>ppe-miR6274b-3p MIMAT0031553

UUGUGUUAUUGGCCGAAAAUAG

>ppe-miR6289 MIMAT0027322

UCCUUUGAAUGGUUAGGCUCA

>ppe-miR482b-5p MIMAT0031166

GGAAUGGGAGGAUUGGGAAAA

>ppe-miR399b MIMAT0031498

UCUGCCAAAGGAGAAUUGCCC

>ppe-miR6267c-3p MIMAT0031543

UAGAGAGAUGGUCAGCAAUGU

>ppe-miR6267a MIMAT0027291

UAGAGAGGUGGUACAAUUGUG

>rgl-miR7801 MIMAT0032242

UACGAGAUGAAACACAGUUUG

>rgl-miR5142 MIMAT0020653

AUAUUGAUUGAUAAGUGAU

>rgl-miR5141 MIMAT0020652

AGACCCGACGCGACUGACAGAUAA

>rgl-miR7803b-3p MIMAT0032253

UACACGUGUCAAUCAUCUAU

>rgl-miR7797a MIMAT0032238

AAGACGGAAUCAAACCUCAA

>rgl-miR5140 MIMAT0020651

GCUGGUGAAGAUUUGGUG

>rgl-miR5577 MIMAT0023002

AGAAGCUGAGAAUCACUUUUU

>rgl-miR5574 MIMAT0022999

UUUAAUCAGAAUCUAAAAGCA

>rgl-miR7807b-5p MIMAT0032258

UAACUAUAUGAAAAUCUCAAUU

>rgl-miR7808 MIMAT0032254

AAGGAUGCUCGAUUCAGAAGAA

>rgl-miR7798 MIMAT0032239

AGGGAGUGUUUGCAAAAACU

>rgl-miR7797b MIMAT0032260

UUUGAUUUCGUCUUACAUUUUUC

>rgl-miR7804-5p MIMAT0032245

AGGGGUGUUCAUCGAAUCGAAUU

>rgl-miR5575 MIMAT0023000

UGGAUUUUGAACGUUUCGGUG

>rgl-miR7811 MIMAT0032257

UGAAUGGAGAUACGGAAUGAAGC

>rgl-miR5573 MIMAT0022998

GAGUAGUCGACCUGAGAUGGA

>rgl-miR7802 MIMAT0032243

AGGGAGUGUUUGCAAUCACUAAA

>rgl-miR7972 MIMAT0032261

UUGUCAGGCUUGUUAUUCUCC

>rgl-miR5578 MIMAT0023003

CAUGUGGCAUGAUGGCAGAAAGU

>rgl-miR7809 MIMAT0032255

UCCCAUUGCAUCAGCGGACACA

>rgl-miR7805-3p MIMAT0032248

UAUUCAUUUACACCAAAUUUGG

>rgl-miR7803a MIMAT0032244

UACGGAUAAUUGACACGUGUAUA

>rgl-miR7804-3p MIMAT0032246

UUUAAUCGAAUGAACAUUUUAAA

>rgl-miR5139 MIMAT0020650

AAACCUGGCUCUGAUACCA

>rgl-miR7807a-3p MIMAT0032251

UUGGGAUUUGCAUACAGUUAC

>rgl-miR7803b-5p MIMAT0032252

GGAUGAUUGCCACGUGUAUA

>rgl-miR7799 MIMAT0032240

AGUGGAAUAGGAGAUCUCAA

>rgl-miR164 MIMAT0023004

UGGGGAAGAAGAGCACAUGAA

>rgl-miR7807b-3p MIMAT0032259

UUGAGAUUUUCAUAUAGUUACU

>rgl-miR7810 MIMAT0032256

AGAGGAAGAGUUUUCUGGCUC

>rgl-miR7800 MIMAT0032241

UAUUUUUGUGUCGUUAUGGUC

>rgl-miR5576 MIMAT0023001

AGAAGUUGGCAUUUGCAAACACU

>rgl-miR5138 MIMAT0020649

AAAAAUCGUUAGGCGCUA

>rgl-miR7806 MIMAT0032249

UAGAAGAUGUCCACAUGAGCA

>rgl-miR5137 MIMAT0020648

AGCGGAGAAGACGAUGGGCU

>rgl-miR7805-5p MIMAT0032247

AAAUUUGGUGUAGUGAAUAGU

>smi-miR12112 MIMAT0049006

CGAUCUUGAUACCACCAAUGG

>ssl-miR399 MIMAT0022520

UGCCAAAGGAGAAUUGCCCGG

>ssl-miR156 MIMAT0022506

UGACAGAAGAGAGUGAGCACA

>ssl-miR395 MIMAT0022516

GGGAAAUGUUUGGGGAAACUU

>ssl-miR948 MIMAT0022522

UGUGGCUGUGUGGGUUCCGG

>ssl-miR1078 MIMAT0022523

CUUGAUUGAUUCAAUUGUGAU

>sly-miR6026 MIMAT0023610

UUCUUGGCUAGAGUUGUAUUGC

>sly-miR9472-5p MIMAT0035449

UUUCAGUAGACGUUGUGAAUA

>sly-miR7981e MIMAT0042015

AAGUGUGUCUCUGAGAUUUCGGAU

>sly-miR9477-5p MIMAT0035471

UAUCCGUUGUUCCCUUUUCCUACC

>sly-miR9473-3p MIMAT0035462

AAACGAGUUCAGAUUUACAGC

>sly-miR9470-5p MIMAT0035439

UGAAAUCCAUGAGCCUAAACU

>sly-miR9469-5p MIMAT0035435

CCACAUAAGAAGACCGAAUUC

>sly-miR169e-5p MIMAT0035475

UAGCCAAGGAUGACUUGCCUUU

>sly-miR482e-5p MIMAT0020765

UGUGGGUGGGGUGGAAAGAUU

>sly-miR10539 MIMAT0042033

CUUGGAACCACAGUUACCACC

>sly-miR9475-3p MIMAT0035466

CUACAAUGUAGAGAUCGUUUU

>sly-miR482b MIMAT0023602

UCUUGCCUACACCGCCCAUGCC

>sly-miR7981b MIMAT0042030

ACCCCUUUUUAGCUUACGUGGCAC

>sly-miR394-3p MIMAT0035438

AGGUGGGCAUACUGUCAACA

>sly-miR10537 MIMAT0042025

AUUUACCCCAAGUUCGUUGUC

>sly-miR5300 MIMAT0020764

UCCCCAGUCCAGGCAUUCCAAC

>sly-miR391 MIMAT0022688

ACGCAGGAGAGAUGAUGCUGGA

>sly-miR9471a-5p MIMAT0035447

CAGGUGCUCACUCAGCUAAUA

>sly-miR1916 MIMAT0007908

AUUUCACUUAGACACCUCAA

>sly-miR398a MIMAT0042014

UAUGUUCUCAGGUCGCCCCUG

>sly-miR167b-5p MIMAT0035457

UAAAGCUGCCAGCAUGAUCUGG

>sly-miR5304 MIMAT0020768

UCAAUGCUACAUACUCAUCCC

>sly-miR164b-3p MIMAT0033976

CACGUGUUCUCCUUCUCCAAC

>sly-miR1919a MIMAT0007911

ACGAGAGUCAUCUGUGACAGG

>sly-miR9469-3p MIMAT0035436

AUUCGGUCUUCUUAUGUGGAC

>sly-miR10534 MIMAT0042020

CAAAAUACCCUUGUCAUCCAA

>sly-miR10540 MIMAT0042034

AUAAUAACUAUUAGUUGAAUG

>sly-miR156d-3p MIMAT0035452

GCUCACUGCUCUAUCUGUCACC

>sly-miR168a-3p MIMAT0031130

CCUGCCUUGCAUCAACUGAAU

>sly-miR6023 MIMAT0023592

UUCCAUGAAAGAGUUUUUGGAU

>sly-miR9477-3p MIMAT0035472

UUGGGAAAGGGAACAACUGAUAGU

>sly-miR9479-3p MIMAT0035478

GAGAAUGGUAGAGGGUCGGACC

>sly-miR9476-5p MIMAT0035469

UCUAGUCCUGCAUCUUUUUUU

>sly-miR1918 MIMAT0007910

UGUUGGUGAGAGUUCGAUUCUC

>sly-miR482d-5p MIMAT0035459

GGAGUGGGUGGGAUGGAAAAA

>sly-miR10529 MIMAT0042009

AACGAGUGAGACUUGCUCAGUUGG

>sly-miR403-3p MIMAT0035434

CUAGAUUCACGCACAAGCUCG

>sly-miR171e MIMAT0035482

UUGAGCCGCGUCAAUAUCUCU

>sly-miR5302b-5p MIMAT0035445

UGAAAUGCUAUAGUUGGAAAGU

>sly-miR395b MIMAT0007927

CUGAAGUGUUUGGGGGAACUCC

>sly-miR10533 MIMAT0042019

UCUUAUGAAUUCUAGGUCUUCU

>sly-miR9476-3p MIMAT0035470

AAAAAGAUGCAGGACUAGACC

>sly-miR156e-5p MIMAT0035453

UGAUAGAAGAGAGUGAGCAC

>sly-miR530 MIMAT0042011

AGGUGUAGGUGUUCAUGCAGA

>sly-miR6025 MIMAT0042023

UACCAAUAAUUGAGAUAACAUC

>sly-miR167b-3p MIMAT0035458

AGGUCAUCUAGCAGCUUCAAU

>sly-miR10535a MIMAT0042022

UUGGCAUAAGUUUGUGAAAGCCGG

>sly-miR10542 MIMAT0042037

UAGAAGAAUCAUAUAUACCCCUA

>sly-miR397-3p MIMAT0037335

UCAACGCUAAACUCGAUCAUG

>sly-miR6027-5p MIMAT0032133

AUGGGUAGCACAAGGAUUAAUG

>sly-miR5303 MIMAT0020767

UUUUUGAAGAGUUCGAGCAAC

>sly-miR159b MIMAT0042036

UUGGAAAGAAGGGAGCUCUAC

>sly-miR9473-5p MIMAT0035461

UGGCUGUAAAUCUAAACUCGU

>sly-miR482c MIMAT0023603

UCUUGCCAAUACCGCCCAUUCC

>sly-miR10536 MIMAT0042024

AGACAUGUUCUAAUCGUCAGCUUC

>sly-miR166c-5p MIMAT0035443

GGGAUGUUGUCUGGCUCGACA

>sly-miR403-5p MIMAT0035433

CGUUUGUGCGUGAAUCUAACA

>sly-miR482d-3p MIMAT0035460

UUUCCUAUUCCACCCAUGCCAA

>sly-miR7981c MIMAT0042021

UCGAAAUCUCAGAGACACACUUAU

>sly-miR393 MIMAT0042012

AUCAUGCGAUCUCUUCGGAAU

>sly-miR319d MIMAT0042007

AGGAAACUGUUUAGUCCAACC

>sly-miR164a-3p MIMAT0033974

CAUGUGCCUGUUUUCCCCAUC

>sly-miR7981a MIMAT0042029

ACCCCUUUUCGGCCUACGUGGCAC

>sly-miR9471b-3p MIMAT0035442

UUGGCUGAGUGAGCAUCACUG

>sly-miR319c-5p MIMAT0035431

AGAGCUUCCUUCAGCCCACUC

>sly-miR10530 MIMAT0042013

ACGUCCCUUCCCCAUCGUUCAACA

>sly-miR10528 MIMAT0042006

AAUGCAAUGUCAUAUACCAUC

>sly-miR1919c-5p MIMAT0032040

UGUCGCAGAUGACUUUCGCCC

>sly-miR10541 MIMAT0042035

AGUCACUUUGAUGAUUGUCAAACA

>sly-miR5302b-3p MIMAT0035446

UUUUCAACUAUAGCAUUAUUUU

>sly-miR6024 MIMAT0023594

UUUUAGCAAGAGUUGUUUUACC

>sly-miR9472-3p MIMAT0035450

UUCACAAUCUCUGCUGAAAAA

>sly-miR9479-5p MIMAT0035477

UCCAGUCCUCUACCCUUCUCCA

>sly-miR6027-3p MIMAT0023611

UGAAUCCUUCGGCUAUCCAUAA

>sly-miR9471b-5p MIMAT0035441

GAGGUGCUCACUCAGCUAAUA

>sly-miR9474-5p MIMAT0035463

UGUAGAAGUCAUGAAUAAAAUG

>sly-miR9474-3p MIMAT0035464

UUUUGUUCGCAGAUACUACAGU

>sly-miR10538 MIMAT0042032

AGAUUGAUAUACGUUACUCACAGU

>sly-miR169e-3p MIMAT0035476

UGGCAAGCAUCUUUGGCGACU

>sly-miR477-3p MIMAT0035484

AGUUCUUGUAGGGUGAGACAAC

>sly-miR1917 MIMAT0007909

AUUAAUAAAGAGUGCUAAAGU

>sly-miR408 MIMAT0042005

ACGGGGACGAGCCAGAGCAUG

>sly-miR399b MIMAT0042016

GGGCUACUCUCUAUUGGCAUG

>sly-miR9478-5p MIMAT0035473

GCUUAAAUAUGUAGAUCGAACU

>sly-miR10532 MIMAT0042018

AAGUGUGUCUCUGAGAUUUCGGGC

>sly-miR9471a-3p MIMAT0035448

UUGGCUGAGUGAGCAUCACGG

>sly-miR9475-5p MIMAT0035465

AACGAUCUCUACAUUGUAGGC

>sly-miR477-5p MIMAT0035483

UGUCUCUCCCUCAAGGGCUCC

>sly-miR10535b MIMAT0042027

UUGGCAUGAGUUUGUGAAAGCCGG

>sly-miR171b-5p MIMAT0037334

AUAUUGGUGCGGUUCAAUUAG

>sly-miR7981f MIMAT0042026

AAGUGUGUCUCUGGAAUUUCGGGC

>sly-miR9478-3p MIMAT0035474

UUCGAUGACAUAUUUGAGCCU

>sly-miR169f MIMAT0042028

UAGGCGUUGUCUGAGGCUAAC

>sly-miR6022 MIMAT0023590

UGGAAGGGAGAAUAUCCAGGA

>sly-miR5302a MIMAT0020766

AAACGAGGUUUGUUACUUUGG

>sly-miR9470-3p MIMAT0035440

UUUGGCUCAUGGAUUUUAGC

>sly-miR10531 MIMAT0042017

UGGGGUCCUAGUAGAGUCGGUUC

>stu-miR391-3p MIMAT0031275

GCAUCAUACUCCUGCAUAUU

>stu-miR8032a-5p MIMAT0030936

UGGUCGGCAUGACUCCCGAGGU

>stu-miR8008a MIMAT0030901

AUUUCCAGAAAAGCGACGGACAGU

>stu-miR8005b-5p MIMAT0031191

ACUCUAAAUUUUAAAUUCUAAAUC

>stu-miR167d-3p MIMAT0031310

GAUCAUGUGGUUGCUUCACC

>stu-miR166d-5p MIMAT0031254

AGAAUGUCGUCUGGUUCGAGA

>stu-miR8032a-3p MIMAT0031208

AGUGUGAGUCGGUGUGAUUAGG

>stu-miR1886b MIMAT0030870

AUGGUAUCGUGAGAUGAAAUCAGC

>stu-miR1886i-5p MIMAT0030925

AUGAGAUGAAAUUAGCGUUUGGAU

>stu-miR7999-3p MIMAT0030889

ACGACCCGUAGAACUGCCCACGAC

>stu-miR8030-3p MIMAT0031207

UUAAAACCAAAUCAACCCAAAU

>stu-miR1886i-3p MIMAT0031202

UUUACGUUGAUUUCAUCUCAUGA

>stu-miR167b-3p MIMAT0031306

GAUCAUGUGGCAGCAUCACC

>stu-miR482e-5p MIMAT0031163

AGUGGGUGGUGUGGUAAGAUU

>stu-miR6026-5p MIMAT0031164

AAUACAACUAUUGCCAAGACAA

>stu-miR7983-5p MIMAT0031183

UAAAGUCUUUAGCGACAUUGGUUC

>stu-miR8038b-3p MIMAT0031217

GUUCAACUUGCUCACUUGGAG

>stu-miR7997c MIMAT0030887

AUAUUGCUCGGACUCUUCAAAAAU

>stu-miR8000 MIMAT0030890

ACACCGAAGAACUGACACCGAAGA

>stu-miR8019-3p MIMAT0030916

AAAAGAAUGACCUGGUUUGACUUG

>stu-miR8006-5p MIMAT0030898

UAGUUUUUGGACGACAGGGGCACC

>stu-miR169e-3p MIMAT0031368

GCAAGUUAUCCUGGCUAUC

>stu-miR169c-3p MIMAT0031364

GCAGUCUCCUUGGCUACC

>stu-miR169b-3p MIMAT0031362

GCAGUCUCCUUGGCUACU

>stu-miR7984c-5p MIMAT0030861

ACGAUACCAAACUUUAUGAAGGAC

>stu-miR8004 MIMAT0030894

AGGGGUUGUGUAUGUGUUUGGCCU

>stu-miR482a-5p MIMAT0031161

GGAAUUGGUGGAUUGGAAAGC

>stu-miR390-3p MIMAT0031344

CGCUAUCCAUCUUGAGUUUUA

>stu-miR7992-5p MIMAT0030868

UUUGACAAUGCACAUCUAGACACU

>stu-miR8040-3p MIMAT0030953

CUUAUAAUUGUAAUUAUGAUC

>stu-miR408a-3p MIMAT0031338

UGCACAGCCUCUUCCCUGGUU

>stu-miR8032g-3p MIMAT0031213

AGUGUGAGUCGGUGCGAUUAGG

>stu-miR7990a MIMAT0030864

UUCAAAUGAUCGUAACUUUGGCCU

>stu-miR399i-5p MIMAT0031260

GGGCUACACUCUAUUGGCAUG

>stu-miR164-3p MIMAT0030973

CAUGUGCUCUAGCUCUCCAGC

>stu-miR399g-3p MIMAT0031233

CGCCAAAGGGGAGCUGCCCUA

>stu-miR7980b-5p MIMAT0031205

GUCCAAACACUGAUUCCAUCUCAU

>stu-miR8045 MIMAT0030959

AUUGAUAGUUGAGGUGUGUUU

>stu-miR8005c MIMAT0030897

UUUAGAGUUUAAGGUUUAGAGUUU

>stu-miR8022 MIMAT0030921

UUUAAAUGAGAAUUUUGGACUAUU

>stu-miR7987 MIMAT0030860

ACACUAGUUGAUCUUAUUGAUGAC

>stu-miR5303f MIMAT0030904

AUUUUUGGAGAAUCUGACACGGGU

>stu-miR7980b-3p MIMAT0030930

GAGAUGGAAUCAGUGUUUGGACAU

>stu-miR477a-5p MIMAT0031326

CCUCUCCCUCAAGGGCUUCUC

>stu-miR8033-5p MIMAT0030943

UUCCAAAGCUGCAGAAAUGAGU

>stu-miR8002-3p MIMAT0031190

AUUCCAUUAUUAUCAAGAAAAAAG

>stu-miR7993d MIMAT0030878

AUAUUUUAUGUGGUUAACUUAACU

>stu-miR8041b-3p MIMAT0030955

AUGAUGUAUAGCAAAGAGCCU

>stu-miR477b-5p MIMAT0031328

ACUCUCCCUCAAAGGCUUCUG

>stu-miR8048-3p MIMAT0030972

AGAUGGACAUGCUAAUGAACA

>stu-miR399d-3p MIMAT0031230

UGCCAAAGGAGAGCUGCCCUG

>stu-miR399j-5p MIMAT0031312

GGGCUACUCUCUAUUGGCAUA

>stu-miR8051-5p MIMAT0030978

UAGUAUGGUAGAAAGAUUCA

>stu-miR8024a-5p MIMAT0031201

UUGAAGAAUUUAAAGACUUCAACU

>stu-miR8041a-5p MIMAT0031221

GUGCUUUGCUAUUUUCAUUG

>stu-miR5303b MIMAT0030857

UUUUGGAGAAUCCGACACGCACCC

>stu-miR482b-5p MIMAT0031160

GGAGUGGGUGGCAUGGUAAGA

>stu-miR6024-5p MIMAT0031159

AGAAACAACACUUGCUAAAAGA

>stu-miR172e-5p MIMAT0031294

GCAACAUCAUCAAGAUUCACA

>stu-miR8001b-3p MIMAT0031200

GGAUUUUCAUACUAAUUCCUAGAA

>stu-miR7984c-3p MIMAT0031185

CCUUCAUAAAGUUUGGUAUCGUAA

>stu-miR8046-3p MIMAT0031225

CGCUGAAAUUUCGAUCAUAAU

>stu-miR7994b-3p MIMAT0030880

AUAUUAUACUUGGGCAUAAACUCC

>stu-miR7122-5p MIMAT0031224

UUAUACAGAGAAACCGCUGUCG

>stu-miR391-5p MIMAT0031274

UACGCAGGAGAGAUGAUGCUG

>stu-miR8018 MIMAT0030915

ACGAACCGUAGAUCCCAUCCGUGG

>stu-miR8007a-5p MIMAT0030899

AUGUGGCACUUUUCGGAUUUUGAG

>stu-miR6149-3p MIMAT0031226

UGAUUCAGGUUUGUAUGCAAAC

>stu-miR7996c MIMAT0030884

AUGUGGUACAUAUGAAAUUUGAAA

>stu-miR8021 MIMAT0030920

AUUCAAGGCUCAAACUCGAGACCU

>stu-miR482d-5p MIMAT0031162

CGUGAGUGGUGGGGUAAGAUA

>stu-miR8033-3p MIMAT0031214

UCAAUUCUGCAGCUUUAGGAGU

>stu-miR7995 MIMAT0030881

UUACACGUAGACAAGUUGACCAUU

>stu-miR7984d-5p MIMAT0030928

AUCCGAACUUUGACCGAAAUUGCU

>stu-miR7992-3p MIMAT0031186

UGUCUAGAUGUGCAUUUCAAAGU

>stu-miR8011a-5p MIMAT0031195

UUGUGUGAGGUUUCUUUUUGUUUC

>stu-miR156d-3p MIMAT0031300

GCUCUCUAUGCUUCUGUCAUCA

>stu-miR7991a MIMAT0030865

AGGAGGUCGGAAUUUUUAAUGAAU

>stu-miR8001b-5p MIMAT0030917

AUGGGGAUUAGUAUGAAAAUUUGC

>stu-miR7982b MIMAT0030850

AAGUUGGAUGAUAAUAAUAUAUAU

>stu-miR7988 MIMAT0030862

AACGGAAAAGGGCCAAAAAUACCC

>stu-miR397-5p MIMAT0031341

AUUGAGUGCAGCGUUGAUGAC

>stu-miR7122-3p MIMAT0030960

ACAGCGUUUCUCUGUAUAACC

>stu-miR7980a MIMAT0030847

AUGAGAUGAAGUCAAUGUUUGGAC

>stu-miR8032d-3p MIMAT0031210

AGUGUGAGUUGGUGCGAUUAGG

>stu-miR7998 MIMAT0030888

ACGGACCGUAGAUCAAUCCACAGU

>stu-miR408a-5p MIMAT0031337

ACAGGGACGAGGCAGCGCAUG

>stu-miR8011a-3p MIMAT0030907

AAUAAAAAGAAGCCUCACACAACU

>stu-miR7999-5p MIMAT0031189

CUGGGUCACUUCUACGGGUCCUUC

>stu-miR8008b MIMAT0030931

AAACCCAGAAAAGCGACGGACAGU

>stu-miR8048-5p MIMAT0031234

CUCAUUAGCAUCUCCAUCUUG

>stu-miR8019-5p MIMAT0031199

AGGGAAGCAGGUCAUUCUUUAUG

>stu-miR8025-3p MIMAT0031203

UUUAAUUGCAUGCCAAGUGUGUGG

>stu-miR827-5p MIMAT0031272

UUUGUUGAUGGUCAUCUAUUC

>stu-miR5303j MIMAT0031241

AAUAUUUUUGAAGAGUCUGAGCAA

>stu-miR1886g-5p MIMAT0030900

GAGAUGAGAUCAAUGUUUGGACAU

>stu-miR8050-3p MIMAT0030976

UGACUUGAGAUUCCUACUUGG

>stu-miR5303g MIMAT0031242

AUAUUUUUGAAGAGUCUGAGCAAC

>stu-miR8006-3p MIMAT0031192

UGCCCUGCCGUCCAAAAAAUAGA

>stu-miR8050-5p MIMAT0031238

AAGUAGGAAUCAAGGUCAAU

>stu-miR477a-3p MIMAT0031327

GAAGCUCUAGCAGGGAGAGCCA

>stu-miR8015-5p MIMAT0030912

UAUUGGAUAUUGAAAAUGAAACUU

>stu-miR7981-5p MIMAT0031182

GUUAAUUAAACUAUGGUCCUAUUA

>stu-miR8014-5p MIMAT0031196

AUUGUUUCAUAUUGUAUUGUAUUU

>stu-miR8038b-5p MIMAT0030949

CCUUGUGAGUAAGUUGAAUCUC

>stu-miR8044-5p MIMAT0031223

UUUCAAAUAUGGUUGGAGAUG

>stu-miR8037 MIMAT0030947

AUAAUUUGGAGGAAUAGGAACC

>stu-miR8049-3p MIMAT0031237

CAUGUCUACAUGAGCCUGAUA

>stu-miR6149-5p MIMAT0030963

UUGCAACACACCUGAAUCGUC

>stu-miR7986 MIMAT0030859

AGUUUAAAACGUUACUGUCGGUAA

>stu-miR7984d-3p MIMAT0031204

CAAUUUCGGUCAAAGUUCGGAUAU

>stu-miR8028-5p MIMAT0031206

UCCUUAUGCUACAAUUGUGAACAA

>stu-miR7984b-3p MIMAT0031184

GGUCUUUCAUAAAAUUUGGUAUCG

>stu-miR3627-3p MIMAT0031246

AAGUGCCUCUGUCUUUCGACA

>stu-miR8007a-3p MIMAT0031193

CGAAAAAUGAAAAGUGCCACAUAA

>stu-miR8024a-3p MIMAT0030923

UUGGAGGAUUUGAAGAUUUCAACU

>stu-miR397-3p MIMAT0031342

CAUCAACGCUACACUCAAUCA

>stu-miR6023 MIMAT0023591

UUCCAUGAAAGUGUUUUUGGAU

>stu-miR8007b-3p MIMAT0031197

CGAAAAAUGAAAAGUACCACAUAA

>stu-miR171d-5p MIMAT0031301

AGAUAUUGGUGCGGUUCAAUU

>stu-miR8047 MIMAT0030962

CCAUUUUUUCGAAAUUAGACC

>stu-miR8001a MIMAT0030891

UCCUGGGGAUUAGUAUGAAAAUUC

>stu-miR7981-3p MIMAT0030848

AUAGGACUUUAGUUUAGUUAAGGU

>stu-miR7985 MIMAT0030854

CGGGCUUGCCUAGAACGGGUUACC

>stu-miR8015-3p MIMAT0031198

GUUUCAUUUUCAAGGUCCAAUAGC

>stu-miR7984a MIMAT0030852

AUACCGAACUUUGGAAAUGACCUU

>stu-miR7993b-5p MIMAT0031187

UUAAGUUAACCACAUAAAAUAUGU

>stu-miR5303e MIMAT0030903

UUUUGGAGAAUCUGACACGGGUGU

>stu-miR8020 MIMAT0030919

AAUUUCAUUGAGUAUGUUGUUGUU

>stu-miR8039 MIMAT0030951

UUUCCUAUCUGAACUAUCACC

>stu-miR8023 MIMAT0030922

UUUGGCACAAUUUCAUUGGCAACC

>stu-miR7997a MIMAT0030885

AUGCUGCUCGGACUCUUCAAA

>stu-miR8017 MIMAT0030914

AUCCAAGUGAAGUGUAUCGUCUCA

>stu-miR7994b-5p MIMAT0031188

AGUUUAUGCCCAAGUAUAUAAUAUAU

>stu-miR156f-3p MIMAT0031348

CUCACUUCUCUUUCUGUCAAUC

>stu-miR7990b MIMAT0030918

GAAUUUUCAAAUGAUCGUAACUUU

>stu-miR7989 MIMAT0030863

ACAAAUAAGUCCAUUACCUGAACC

>stu-miR1886h MIMAT0030905

AUUUUACGUUGAUUUCAUCUCAUG

>stu-miR8025-5p MIMAT0030926

ACAUACUCGACAUGCAAUUAAAUU

>stu-miR169d-3p MIMAT0031366

GCAGGUCAUCUUUAGCUAACU

>stu-miR8014-3p MIMAT0030910

AUGAAUACAAUGUUUGGAUAAAUU

>stu-miR384-3p MIMAT0031374

AGGGGGCCAAAGUGCCAAAC

>stu-miR8028-3p MIMAT0030932

GUUCAUAAUUAUAGUAUAAGGAUG

>stu-miR167c-3p MIMAT0031304

GGUCAUGCUCGGACAGCCUCACU

>stu-miR1886g-3p MIMAT0031194

UUUCAUAUUGAUUUCAUCUCAU

>stu-miR8034 MIMAT0030944

UAUGACAAACACUGCAAAAACU

>stu-miR398b-5p MIMAT0031333

GAGUGUGCCUUAGAACACAGGU

>stu-miR5303h MIMAT0031243

AACAUUUUUGAAGAGUCUGAGCAA

>stu-miR7979 MIMAT0030846

AGGUACAUGAACUCUAACGAGGCA

>stu-miR408b-5p MIMAT0031339

ACGGGGACGAGACAGAGCAUG

>stu-miR172b-5p MIMAT0031286

GCAGCACCAUCAAGAUUCACA

>stu-miR8029 MIMAT0030933

AGCCAUUUUUCUUUGUUUUGGAGC

>stu-miR8040-5p MIMAT0031220

UCAUAAUUACAAUUAUAAGCC

>stu-miR3627-5p MIMAT0031245

UCGCAGGAGAGAUGGCACUUAG

>stu-miR398b-3p MIMAT0031334

UUGUGUUCUCAGGUCACCCCU

>stu-miR8046-5p MIMAT0030961

UAUGAUCGAAGUUUCAAUGAC

>stu-miR8043 MIMAT0030957

UGAUAUAAUUGGACUUUGGCC

>stu-miR8044-3p MIMAT0030958

UCUCCAGCGAUAUUUGAAACU

>stu-miR398a-5p MIMAT0031331

GGGUUGAUUUGAGAACAUAUG

>stu-miR8027 MIMAT0030929

AUCUCGAGAUAAGUUAUUCUGGAC

>stu-miR482b-3p MIMAT0023597

UUACCGAUUCCCCCCAUUCCAA

>stu-miR6024-3p MIMAT0023593

UUUUAGCAAGAGUUGUUUUCCC

>stu-miR7983-3p MIMAT0030851

ACUAAUGCCGGUAAAGACUUUAAC

>stu-miR8049-5p MIMAT0030975

CAAGGCUCAUGCAGACAUGCA

>stu-miR171c-3p MIMAT0031259

UGAUUGAGCCGUGUCAAUAUC

>stu-miR319a-5p MIMAT0031277

AGAGCUUUCUUCGGUCCACAC

>stu-miR172a-5p MIMAT0031290

GUAGCAUAAUCAAGAUUCACA

>stu-miR8036-3p MIMAT0030946

UAUGUCUUUCCGAUGCCUCCCA

>stu-miR477b-3p MIMAT0031329

GAGGUCUUUCGAGUGAGAGUGA

>stu-miR172c-5p MIMAT0031288

AGCAUCUUCAAGAUUCACA

>stu-miR8010 MIMAT0030906

AUAGGACCCUAGUUAAAUUUAGGU

>stu-miR169f-3p MIMAT0031370

GCAAGCAUCCUUGGCGACU

>stu-miR393-5p MIMAT0031324

UCCAAAGGGAUCGCAUUGAUCC

>stu-miR8011b-3p MIMAT0030950

UUCGUGAGACAAAAAGAAGCCU

>stu-miR8036-5p MIMAT0031215

GGAGGAAUCGAAAGAUAUAAG

>stu-miR319-3p MIMAT0030952

UUGGACUGAAGGGUUCCCUUC

>stu-miR8051-3p MIMAT0031240

UAUUUCUUCUACCAUACUAUU

>stu-miR8003 MIMAT0030893

AUUUCGGUAUACAAAUGGGAUGAC

>stu-miR8012 MIMAT0030908

AUGACUUUAAGUCGCGUCUGGCCC

>stu-miR8042 MIMAT0030956

AUUAGACUGAAGUGCUGAUCU

>stu-miR8011b-5p MIMAT0031218

ACUCAUUUUUGUCUCACAAAAA

>stu-miR8013 MIMAT0030909

AGAAGAAAAUCGCUCCGUCAGAAG

>stu-miR167a-3p MIMAT0031308

GAUCAUGUGGCAGCCUCACC

>stu-miR8002-5p MIMAT0030892

UUUUUCGUGAUAAUAAUGGAAUCA

>stu-miR8007b-5p MIMAT0030911

AUGUGACACUUUUUGAAUUUCGAG

>stu-miR171b-5p MIMAT0031247

AGAUAUUGAUGUGGCUCAAUC

>stu-miR8035 MIMAT0030945

UCCAUCUUCAAUAUCACUUUCU

>stu-miR5304-3p MIMAT0031236

AGAUGAGUAUGGUGCAUUGGA

>stu-miR396-3p MIMAT0031336

GUCCAAGAAAGCUGUGGGAAA

>stu-miR8031 MIMAT0030935

UUAGACACCUCAACUAAGACUUG

>stu-miR8030-5p MIMAT0030934

UUGGGUUGGUUUGGUCUCGGGUU

>stu-miR8016 MIMAT0030913

AUUUUUGAAUGGAAGGCCCAUGUG

>stu-miR8026 MIMAT0030927

AUGUAGAGAAUAUGUGGUAACCCU

>stu-miR5304-5p MIMAT0030974

CAAUGCAACAUACUCAUCACC

>tcc-miR399a MIMAT0020435

CGCCAAAGGAGAGUUGCCCUG

>tcc-miR169n MIMAT0020404

UGAGUCAAGAAUGACUUGCCG

>tcc-miR399f MIMAT0020440

UGCCAGAGGAGAUUUGCCCUG

>tcc-miR169i MIMAT0020399

UAGCCAAGGAUGAGUUGCCUG

>tcc-miR169f MIMAT0020396

AAGCCAAGAAUGACUUGCCUG

>tcc-miR535 MIMAT0020448

UGACAACGAUAGAGAGCACGC

>tcc-miR169g MIMAT0020397

UAGCCAGGGAUGACUUGCCUA

>tcc-miR399c MIMAT0020437

UGCCAAUGGAGAUUUGCCCAG

>tcc-miR172d MIMAT0020416

AGAAUCCUGAUGAUGCUGCAU

>vvi-miR3631c MIMAT0018033

CAUGUUGACAUCAUCCAAUAUA

>vvi-miR169r MIMAT0005687

UGAGUCAAGGAUGACUUGCCG

>vvi-miR169y MIMAT0005677

UAGCGAAGGAUGACUUGCCUA

>vvi-miR171g MIMAT0006555

UUGAGCCGAACCAAUAUCACC

>vvi-miR3629a-3p MIMAT0018022

GGCUGCUGAGAAAAUGUAGGA

>vvi-miR3637-3p MIMAT0018048

UUUCGACAAGACACAAUGCAUAAA

>vvi-miR3636-5p MIMAT0018045

UCGGUUUGCUUCUUUGAUAGAUUC

>vvi-miR159a MIMAT0005648

CUUGGAGUGAAGGGAGCUCUC

>vvi-miR3640-3p MIMAT0018054

AUCGAAAAGGCAUCAUCAAUCAGG

>vvi-miR3633a-5p MIMAT0018037

GGAAUGGAUGGUUAGGAGAG

>vvi-miR3632-3p MIMAT0018036

UUUCCCAGACCCCCAAUACCAA

>vvi-miR3624-5p MIMAT0018011

UAGUAUGCUGCUGUCUUUAGA

>vvi-miR3634-3p MIMAT0018040

UUUCCGACUCGCACUCAUGCCGU

>vvi-miR2950-3p MIMAT0018010

UGGUGUGCACGGGAUGGAAUA

>vvi-miR3631a-3p MIMAT0018030

UAUAUUGGAUGAUGUCAACAA

>vvi-miR3632-5p MIMAT0018035

GGAUUGGGGGCCGAUGGAAAGG

>vvi-miR3638-3p MIMAT0018050

GCAACAAGCAUGAAAAGGCACACC

>vvi-miR3624-3p MIMAT0018012

UCAGGGCAGCAGCAUACUACU

>vvi-miR3625-5p MIMAT0018013

UUCCAGCAGUCAUCUCCAAGG

>vvi-miR845c MIMAT0006581

AGGCUCUGAUACCAAUUGAUG

>vvi-miR399f MIMAT0006567

UGCCGAAGGAGAUUUGUCCUG

>vvi-miR169b MIMAT0006545

UGAGCCAAGGAUGGCUUGCCG

>vvi-miR3626-5p MIMAT0018015

GGUAGUCGCUGUGAAAUUGAA

>vvi-miR3633b-3p MIMAT0018042

GUUCCCAUGCCAUCCAUUCCUA

>vvi-miR172a MIMAT0005699

UGAAUCUUGAUGAUGCUACAU

>vvi-miR3636-3p MIMAT0018046

GUCUGUCGGAGAAGCAAGUCGGAG

>vvi-miR169i MIMAT0006547

GAGCCAAGGAUGACUGGCCGU

>vvi-miR3631b-3p MIMAT0018032

UGUUGGAUGAUGUCAAUAAGU

>vvi-miR3630-3p MIMAT0018028

UUUGGGAAUCUCUCUGAUGCAC

>vvi-miR3633b-5p MIMAT0018041

GGAAUGGGUGGCUGGGAUCUA

>vvi-miR477b-3p MIMAT0018026

CGAAGUCUUUGGGGAGAGUGG

>vvi-miR3627-5p MIMAT0018017

UUGUCGCAGGAGAGACGGCACU

>vvi-miR3639-3p MIMAT0018052

GAGCUUUUGGCUUCUCAGAAGUCA

>vvi-miR399d MIMAT0006566

UGCCAAAGGAGAUUUGCUCGU

>vvi-miR479 MIMAT0005734

UGUGGUAUUGGUUCGGCUCAUC

>vvi-miR3629a-5p MIMAT0018021

CGCAUUUUCUCAGCAGCCAAG

>vvi-miR171h MIMAT0005697

UGGUUGAGCCGCGCCAAUAUC

>vvi-miR2111-5p MIMAT0016364

UAAUCUGCAUCCUGAGGUCUA

>vvi-miR845e MIMAT0006583

UGGCUCUGAUACCAAUUGAUG

>vvi-miR845b MIMAT0006580

UAGCUCUGAUACCAAUUGAUA

>vvi-miR3627-3p MIMAT0018018

UCGCCGCUCUCCUGUGACAAG

>vvi-miR172d MIMAT0005702

UGAGAAUCUUGAUGAUGCUGCAU

>vvi-miR2950-5p MIMAT0018009

UUCCAUCUCUUGCACACUGGA

>vvi-miR396a MIMAT0005724

UUCCACAGCUUUCUUGAACUA

>vvi-miR160b MIMAT0005652

UGCCUGGCUCCCUGAAUGCCAUC

>vvi-miR156e MIMAT0005644

UGACAGAGGAGAGUGAGCAC

>vvi-miR169l MIMAT0006548

GAGCCAAGGAUGACUUGCCGU

>vvi-miR3623-5p MIMAT0018007

UCACAAGUUCAUCCAAGCACCA

>vvi-miR477a MIMAT0006575

AUCUCCCUCAAAGGCUUCCAA

>vvi-miR394c MIMAT0006558

UUGGCAUUCUGUCCACCUCCAU

>vvi-miR395n MIMAT0006559

CUGAAGAGUCUGGAGGAACUC

>vvi-miR169t MIMAT0005689

CGAGUCAAGGAUGACUUGCCG

>vvi-miR3638-5p MIMAT0018049

UGUGCCUUUUCGCGCUUGUUGCUA

>vvi-miR2111-3p MIMAT0016365

GUCCUCUGGUUGCAGAUUACU

>vvi-miR3628-5p MIMAT0018019

AUGCGAGAGCCGUGCUUAGUA

>vvi-miR169o MIMAT0006550

GAGCCAAGGAUGACUUGCCGC

>vvi-miR3630-5p MIMAT0018027

UGCAAGUGACGAUAUCAGACA

>vvi-miR3635-3p MIMAT0018044

AUUAUGUCCCACACAUGCCUC

>vvi-miR319g MIMAT0005706

UUGGACUGAAGGGAGCUCCCA

>vvi-miR399g MIMAT0005731

UGCCAAAGGAGAUUUGCCCCU

>vvi-miR3623-3p MIMAT0018008

UGGUGCUUGGACGAAUUUGCUA

>vvi-miR172b MIMAT0005700

UGAAUCUUGAUGAUGCUACAC

>vvi-miR3633a-3p MIMAT0018038

UUCCUAUACCACCCAUUCCCUA

>vvi-miR156h MIMAT0006544

UGACAGAAGAGAGAGAGCAU

>vvi-miR3639-5p MIMAT0018051

AUUGACUUCUGAAAGGCUAAAAGC

>vvi-miR3640-5p MIMAT0018053

ACCUGAUUGGUGAUGCUUUUUUGG

>vvi-miR3634-5p MIMAT0018039

GGCAUAUGUGUGACGGAAAGA

>vvi-miR482 MIMAT0006576

UCUUUCCUACUCCUCCCAUUCC

>vvi-miR3637-5p MIMAT0018047

AUUUAUGUAUUGUGUUUUGUCGGA

>vvi-miR3635-5p MIMAT0018043

GGCAUGUGUGGGGCAUAAUAG

>vvi-miR3625-3p MIMAT0018014

CGGGAGAUGACUACUGGAAGC

>vvi-miR169v MIMAT0006552

AAGCCAAGGAUGAAUUGCCGG

>vvi-miR3628-3p MIMAT0018020

CUAAGCACAGCUCUCGCAUCC

>vvi-miR3626-3p MIMAT0018016

CUUCAAUUUCACAGCGACCAC

>vvi-miR477b-5p MIMAT0018025

ACUCUUUCUCAAGGGCUUCUAG

GAAGUUUCAAGUGUAAAAAAGUGG

>cca-miR6118-5p MIMAT0024565

UGGAAUUGGGUGCUUCGGAAGA

>cca-miR6105a MIMAT0024538

ACGAAAACAUGUUGGUCUCACGUG

>cca-miR395a MIMAT0024526

UUGAAGUGUUUGGGGGGACUC

>cca-miR6108c MIMAT0024555

AAGCGUAAGAAGAGAUCUGAACC

>cca-miR6106-5p MIMAT0024539

UUGCAAGUAUCCGGAUUUAAA

>cca-miR6116-5p MIMAT0024562

CAUGCUUGUGAUCAAAUGAUG

>cca-miR6109 MIMAT0024546

AUGGACGUGUUAUUCAUCAUGAAU

>cca-miR6108e-5p MIMAT0024557

GUAAGAAGAGAUCUCCACCCUUGG

>cca-miR6104 MIMAT0024537

AUACGACAAAUAGAACAAAUAAAC

>cca-miR6114-5p MIMAT0024553

UGAAAGGAAUCAUGAACGUGA

>cca-miR6111-3p MIMAT0024550

CAUGCAUGGUGAUAUAAAUAGC

>cca-miR6114-3p MIMAT0024554

UCACGUCCAUUGUCCCUUUCA

>cca-miR6110-3p MIMAT0024548

AUCCACGUCAUCAAAUGUUACAAG

>cca-miR6113 MIMAT0024552

UCUGAAACUCAAGAACACGUUG

>cca-miR169b MIMAT0024519

UAGCCAAGGAUGACUUGCUAC

>cca-miR396c MIMAT0024561

UUCAAGAAAGCUGUGGGAAAA

>cca-miR6103-3p MIMAT0024536

CAAGAAGUUGUCUUAGGCAUG

>cca-miR6118-3p MIMAT0024566

UUCCGAGGCCACCCAUUCCAAC

>cca-miR6115 MIMAT0024560

UCUGGACGGUAUGCACAUGUGCAU

>cca-miR6107 MIMAT0024541

AAAGGGGACAAUAUCUGGUACGGU

>cca-miR6117 MIMAT0024564

GGUUAGGUUGAUCGGGUUGAAGAC

>cca-miR6111-5p MIMAT0024549

UCUUUAUGUCACGAUGUAUGAC

>cca-miR396a-3p MIMAT0024530

GUUCAAGAAAGCUGUGGGAAA

>cca-miR6116-3p MIMAT0024563

UCAUUUGAUCACAAGCAUGAG

>cca-miR6103-5p MIMAT0024535

UGUCUAAGACAACUCCUUGGA

>han-miR160a MIMAT0025498

UGCUCGGCUCCCUGUAUGCCA

>han-miR3440 MIMAT0025524

UGGGUUGGUCAAGGGAAACGC

>han-miR3630-3p MIMAT0025526

UGUGGGAAUCUCUCUGAUGCUU

>han-miR3630-5p MIMAT0025525

GCAAGUGAUGAUAGACCAGACA

>hci-miR156a MIMAT0025490

UGACAGAAGAGAGUGAGUAC

>hci-miR164a MIMAT0025502

UGGAGAAGCAGGGCACGUGAA

UUUAGGUCGAGCUUCAUUGGA

>ath-miR447c-3p MIMAT0002115

UUGGGGACGACAUCUUUUGUUG

>ath-miR5648-5p MIMAT0022412

UUUGGAAAUAUUUGGCUUGACU

>ath-miR827 MIMAT0004243

UUAGAUGACCAUCAACAAACU

>ath-miR5017-3p MIMAT0020521

UUAUACCAAAUUAAUAGCAAA

>ath-miR405d MIMAT0001008

AUGAGUUGGGUCUAACCCAUAACU

>ath-miR163 MIMAT0000184

UUGAAGAGGACUUGGAACUUCGAU

>ath-miR823 MIMAT0004240

UGGGUGGUGAUCAUAUAAGAU

>ath-miR447a.2-3p MIMAT0020345

UAUGGAAGAAAUUGUAGUAUU

>ath-miR5660 MIMAT0022434

CAGGUGGUUAGUGCAAUGGAA

>ath-miR840-3p MIMAT0032021

UUGUUUAGGUCCCUUAGUUUC

>ath-miR8183 MIMAT0032782

UUUAGUUGACGGAAUUGUGGC

>ath-miR854d MIMAT0004283

GAUGAGGAUAGGGAGGAGGAG

>ath-miR5654-5p MIMAT0022426

AUAAAUCCCAACAUCUUCCA

>ath-miR840-5p MIMAT0004262

ACACUGAAGGACCUAAACUAAC

>ath-miR398c-5p MIMAT0031912

AGGGUUGAUAUGAGAACACAC

>ath-miR3932b-5p MIMAT0032113

UUUGACGUGCUCGAUCUGCUC

>ath-miR402 MIMAT0001003

UUCGAGGCCUAUUAAACCUCUG

>ath-miR5629 MIMAT0022388

UUAGGGUAGUUAACGGAAGUUA

>ath-miR5012 MIMAT0020516

UUUUACUGCUACUUGUGUUCC

>ath-miR832-3p MIMAT0004251

UUGAUUCCCAAUCCAAGCAAG

>ath-miR5029 MIMAT0020535

AAUGAGAGAGAACACUGCAAA

>ath-miR8167d MIMAT0037259

AGAUGUGGAGAUCGUGGGGAUG

>ath-miR8176 MIMAT0032775

GGCCGGUGGUCGCGAGAGGGA

>ath-miR158a-5p MIMAT0031873

CUUUGUCUACAAUUUUGGAAA

>ath-miR171b-5p MIMAT0031899

AGAUAUUAGUGCGGUUCAAUC

>ath-miR781b MIMAT0022420

UUAGAGUUUUCUGGAUACUUA

>ath-miR8121 MIMAT0032262

AAAGUAUAAUGGUUUAGUGGUUUG

>ath-miR830-3p MIMAT0004248

UAACUAUUUUGAGAAGAAGUG

>ath-miR8181 MIMAT0032780

UGGGGGUGGGGGGGUGACAG

>ath-miR167c-3p MIMAT0031917

UAGGUCAUGCUGGUAGUUUCACC

>ath-miR396b-3p MIMAT0031909

GCUCAAGAAAGCUGUGGGAAA

>ath-miR3933 MIMAT0018348

AGAAGCAAAAUGACGACUCGG

>ath-miR8166 MIMAT0032762

AGAGAGUGUAGAAAGUUUCUCA

>ath-miR859 MIMAT0004303

UCUCUCUGUUGUGAAGUCAAA

>ath-miR842 MIMAT0004264

UCAUGGUCAGAUCCGUCAUCC

>ath-miR4240 MIMAT0017948

UGACUAGACCCGUAACAUUAC

>ath-miR5664 MIMAT0022442

AUAGUCAAUUUUAUCGGUCUG

>ath-miR851-5p MIMAT0004273

UCUCGGUUCGCGAUCCACAAG

>ath-miR829-3p.1 MIMAT0004245

AGCUCUGAUACCAAAUGAUGGAAU

>ath-miR8178 MIMAT0032777

UAACAGAGUAAUUGUACAGUG

>ath-miR2936 MIMAT0014140

CUUGAGAGAGAGAACACAGACG

>ath-miR5635d MIMAT0022405

UGUUAAGGAGUGUUAACGGUG

>ath-miR2938 MIMAT0014142

GAUCUUUUGAGAGGGUUCCAG

>ath-miR779.1 MIMAT0003938

UUCUGCUAUGUUGCUGCUCAU

>ath-miR406 MIMAT0001009

UAGAAUGCUAUUGUAAUCCAG

>ath-miR8169 MIMAT0032767

AUAGACAGAGUCACUCACAGA

>ath-miR5016 MIMAT0020520

UUCUUGUGGAUUCCUUGGAAA

>ath-miR775 MIMAT0003934

UUCGAUGUCUAGCAGUGCCA

>ath-miR865-3p MIMAT0004314

UUUUUCCUCAAAUUUAUCCAA

>ath-miR5025 MIMAT0020531

ACUGUAUAUAUGUAAGUGACA

>ath-miR852 MIMAT0004275

AAGAUAAGCGCCUUAGUUCUG

>ath-miR837-3p MIMAT0004259

AAACGAACAAAAAACUGAUGG

>ath-miR169a-3p MIMAT0031886

GGCAAGUUGUCCUUGGCUAC

>ath-miR864-5p MIMAT0004311

UCAGGUAUGAUUGACUUCAAA

>ath-miR419 MIMAT0001327

UUAUGAAUGCUGAGGAUGUUG

>ath-miR773b-5p MIMAT0017735

GGCAAUAACUUGAGCAAACA

>ath-miR8180 MIMAT0032779

UGCGGUGCGGGAGAAGUGC

>ath-miR5648-3p MIMAT0022413

AUCUGAAGAAAAUAGCGGCAU

>ath-miR2934-3p MIMAT0020364

CAUCCAAGGUGUUUGUAGAAA

>ath-miR774a MIMAT0003933

UUGGUUACCCAUAUGGCCAUC

>ath-miR167c-5p MIMAT0001018

UAAGCUGCCAGCAUGAUCUUG

>ath-miR168b-3p MIMAT0031885

CCCGUCUUGUAUCAACUGAAU

>ath-miR824-3p MIMAT0032024

CCUUCUCAUCGAUGGUCUAGA

>ath-miR165a-5p MIMAT0031879

GGAAUGUUGUCUGGAUCGAGG

>ath-miR836 MIMAT0004257

UCCUGUGUUUCCUUUGAUGCGUGG

>ath-miR8184 MIMAT0032783

UUUGGUCUGAUUACGAAUGUA

>ath-miR5645c MIMAT0022414

AACCUAUUUAACGACAUGACU

>ath-miR1886.1 MIMAT0007853

UGAGAGAAGUGAGAUGAAAUC

>ath-miR5019 MIMAT0020523

UGUUGGGAAAGAAAAACUCUU

>ath-miR774b-3p MIMAT0017740

CAUCCAUAUUUUCAUCUCGAA

>ath-miR426 MIMAT0001337

UUUUGGAAAUUUGUCCUUACG

>ath-miR5015 MIMAT0020519

UUGGUGUUAUGUGUAGUCUUC

>ath-miR8165 MIMAT0032761

AAUGGAGGCAAGUGUGAAGGA

>ath-miR826b MIMAT0035543

UGGUUUUGGACACGUGAAAAU

>ath-miR394b-3p MIMAT0031907

AGGUGGGCAUACUGCCAAUAG

>ath-miR8171 MIMAT0032770

AUAGGUGGGCCAGUGGUAGGA

>ath-miR5022 MIMAT0020527

GUCAUGGGGUAUGAUCGAAUG

>ath-miR161.1 MIMAT0000181

UGAAAGUGACUACAUCGGGGU

>ath-miR4221 MIMAT0017942

UUUUCCUCUGUUGAAUUCUUGC

>ath-miR845b MIMAT0004317

UCGCUCUGAUACCAAAUUGAUG

>ath-miR164c-3p MIMAT0031916

CACGUGUUCUACUACUCCAAC

>ath-miR415 MIMAT0001323

AACAGAGCAGAAACAGAACAU

>ath-miR393b-3p MIMAT0031906

AUCAUGCGAUCUCUUUGGAUU

>ath-miR5650 MIMAT0022416

UUGUUUUGGAUCUUAGAUACA

>ath-miR159b-5p MIMAT0031889

GAGCUCCUUGAAGUUCAAUGG

>ath-miR1887 MIMAT0007854

UACUAAGUAGAGUCUAAGAGA

>ath-miR5636 MIMAT0022396

CGUAGUUGCAGAGCUUGACGG

>ath-miR870-3p MIMAT0004322

UAAUUUGGUGUUUCUUCGAUC

>ath-miR8175 MIMAT0032774

GAUCCCCGGCAACGGCGCCA

>ath-miR864-3p MIMAT0004312

UAAAGUCAAUAAUACCUUGAAG

>ath-miR841a-3p MIMAT0032022

AUUUCUAGUGGGUCGUAUUCA

>ath-miR5639-5p MIMAT0022400

UAGUCCACUGUGGUCUAAGGC

>ath-miR159c MIMAT0001015

UUUGGAUUGAAGGGAGCUCCU

>ath-miR8179 MIMAT0032778

UGACUGCAUUAACUUGAUCGU

>ath-miR782 MIMAT0003941

ACAAACACCUUGGAUGUUCUU

>ath-miR8173 MIMAT0032772

AUGUGCUGAUUCGAGGUGGGA

>ath-miR156b-3p MIMAT0031866

UGCUCACCUCUCUUUCUGUCAGU

>ath-miR407 MIMAT0001010

UUUAAAUCAUAUACUUUUGGU

>ath-miR826a MIMAT0004242

UAGUCCGGUUUUGGAUACGUG

>ath-miR391-3p MIMAT0031904

ACGGUAUCUCUCCUACGUAGC

>ath-miR418 MIMAT0001326

UAAUGUGAUGAUGAACUGACC

>ath-miR863-3p MIMAT0004310

UUGAGAGCAACAAGACAUAAU

>ath-miR3440b-5p MIMAT0017945

UUUUCUUGGCCCAUCCACUUC

>ath-miR869.1 MIMAT0004320

AUUGGUUCAAUUCUGGUGUUG

>ath-miR5644 MIMAT0022406

GUGGGUUGCGGAUAACGGUA

>ath-miR5020b MIMAT0020529

AUGGCAUGAAAGAAGGUGAGA

>ath-miR832-5p MIMAT0004250

UGCUGGGAUCGGGAAUCGAAA

>ath-miR164b-3p MIMAT0031878

CAUGUGCCCAUCUUCACCAUC

>ath-miR156f-3p MIMAT0031869

GCUCACUCUCUAUCCGUCACC

>ath-miR838 MIMAT0004260

UUUUCUUCUACUUCUUGCACA

>ath-miR835-3p MIMAT0004256

UGGAGAAGAUACGCAAGAAAG

>ath-miR156g MIMAT0001012

CGACAGAAGAGAGUGAGCAC

>ath-miR414 MIMAT0001322

UCAUCUUCAUCAUCAUCGUCA

>ath-miR8182 MIMAT0032781

UUGUGUUGCGUUUCUGUUGAUU

>ath-miR398b-3p MIMAT0000949

UGUGUUCUCAGGUCACCCCUG

>ath-miR5646 MIMAT0022410

GUUCGAGGCACGUUGGGAGG

>ath-miR865-5p MIMAT0004313

AUGAAUUUGGAUCUAAUUGAG

>ath-miR398a-5p MIMAT0031910

AAGGAGUGGCAUGUGAACACA

>ath-miR5027 MIMAT0020533

ACCGGUUGGAACUUGCCUUAA

>ath-miR2937 MIMAT0014141

AUAAGAGCUGUUGAAGGAGUC

>ath-miR778 MIMAT0003937

UGGCUUGGUUUAUGUACACCG

>ath-miR2111a-3p MIMAT0011151

GUCCUCGGGAUGCGGAUUACC

>ath-miR5651 MIMAT0022422

UUGUGCGGUUCAAAUAGUAAC

>ath-miR408-5p MIMAT0031915

ACAGGGAACAAGCAGAGCAUG

>ath-miR166e-5p MIMAT0031882

GGAAUGUUGUCUGGCACGAGG

>ath-miR395f MIMAT0000943

CUGAAGUGUUUGGGGGGACUC

>ath-miR773a MIMAT0003932

UUUGCUUCCAGCUUUUGUCUC

>ath-miR158b MIMAT0001014

CCCCAAAUGUAGACAAAGCA

>ath-miR866-5p MIMAT0004315

UCAAGGAACGGAUUUUGUUAA

>ath-miR5637 MIMAT0022397

AAUGCGCAACUCUAUAUUUCC

>ath-miR1886.2 MIMAT0012129

UGAGAUGAAAUCUUUGAUUGG

>ath-miR829-3p.2 MIMAT0004246

CAAAUUAAAGCUUCAAGGUAG

>ath-miR853 MIMAT0004276

UCCCCUCUUUAGCUUGGAGAAG

>ath-miR5633 MIMAT0022393

UAUGAUCAUCAGAAAACAGUG

>ath-miR862-3p MIMAT0004308

AUAUGCUGGAUCUACUUGAAG

>ath-miR400 MIMAT0001001

UAUGAGAGUAUUAUAAGUCAC

>ath-miR866-3p MIMAT0004316

ACAAAAUCCGUCUUUGAAGA

>ath-miR5652 MIMAT0022424

UUGAAUGUGAAUGAAUCGGGC

>ath-miR10515 MIMAT0041984

ACCCCGAUGGUUAUCCUCACC

>ath-miR5645b MIMAT0022409

AUUUGAGUCAUGUCGUUAAG

>ath-miR5026 MIMAT0020532

ACUCAUAAGAUCGUGACACGU

>ath-miR5663-3p MIMAT0032129

UGAGAAUGCAAAUCCUUAGCU

>ath-miR862-5p MIMAT0004307

UCCAAUAGGUCGAGCAUGUGC

>ath-miR169f-3p MIMAT0031898

GCAAGUUGACCUUGGCUCUGC

>ath-miR4245 MIMAT0023523

ACAAAGUUUUAUACUGACAAU

>ath-miR173-3p MIMAT0022843

UGAUUCUCUGUGUAAGCGAAA

>ath-miR1888a MIMAT0007855

UAAGUUAAGAUUUGUGAAGAA

>ath-miR4243 MIMAT0017949

UUGAAAUUGUAGAUUUCGUAC

>ath-miR5023 MIMAT0020528

AUUGGUAGUGGAUAAGGGGGC

>ath-miR777 MIMAT0003936

UACGCAUUGAGUUUCGUUGCUU

>ath-miR855 MIMAT0004279

AGCAAAAGCUAAGGAAAAGGAA

>ath-miR5655 MIMAT0022427

AAGUAGACACAUAAGAAGGAG

>ath-miR5018 MIMAT0020522

UUAAAGCUCCACCAUGAGUCCAAU

>ath-miR780.1 MIMAT0004218

UCUAGCAGCUGUUGAGCAGGU

>ath-miR3434-3p MIMAT0017738

UCAGAGUAUCAGCCAUGUGA

>ath-miR868-5p MIMAT0020355

UCAUGUCGUAAUAGUAGUCAC

>ath-miR5656 MIMAT0022429

ACUGAAGUAGAGAUUGGGUUU

UUUGAUUCCAGCUUUUGUCUC

>ath-miR1886.3 MIMAT0013773

AAUUAAAGAUUUCAUCUUACU

>ath-miR831-5p MIMAT0032020

AGAAGCGUACAAGGAGAUGAGG

>ath-miR844-5p MIMAT0004266

UGGUAAGAUUGCUUAUAAGCU

>ath-miR5014a-3p MIMAT0020518

UUGUACAAAUUUAAGUGUACG

>ath-miR856 MIMAT0004300

UAAUCCUACCAAUAACUUCAGC

>ath-miR5996 MIMAT0023519

UGACAUCCAGAUAGAAGCUUUG

>ath-miR848 MIMAT0004270

UGACAUGGGACUGCCUAAGCUA

>ath-miR8177 MIMAT0032776

GUGUGAUGAUGUGUCAUUUAUA

>ath-miR5640 MIMAT0022401

UGAGAGAAGGAAUUAGAUUCA

>ath-miR846-5p MIMAT0032023

CAUUCAAGGACUUCUAUUCAG

>ath-miR5639-3p MIMAT0032128

UUUAGCCUCAGACCACGGUGGACU

>ath-miR825 MIMAT0004241

UUCUCAAGAAGGUGCAUGAAC

>ath-miR828 MIMAT0004244

UCUUGCUUAAAUGAGUAUUCCA

>ath-miR822-3p MIMAT0032018

UGUGCAAAUGCUUUCUACAGG

>ath-miR839-5p MIMAT0004261

UACCAACCUUUCAUCGUUCCC

>ath-miR5024-5p MIMAT0020530

AUGACAAGGCCAAGAUAUAACA

>ath-miR399e MIMAT0000955

UGCCAAAGGAGAUUUGCCUCG

>ath-miR447c-5p MIMAT0020346

CCCCUUACAAUGUCGAGUAAA

>ath-miR4227 MIMAT0017943

UCACUGGUACCAAUCAUUCCA

>ath-miR833a-5p MIMAT0004252

UGUUUGUUGUACUCGGUCUAGU

>ath-miR8170-3p MIMAT0032769

UUGCUUAAAGAUUUUCUAUGU

>ath-miR822-5p MIMAT0004239

UGCGGGAAGCAUUUGCACAUG

>ath-miR390b-3p MIMAT0031903

CGCUAUCCAUCCUGAGUUCC

>ath-miR851-3p MIMAT0004274

UGGGUGGCAAACAAAGACGAC

>ath-miR837-5p MIMAT0004258

AUCAGUUUCUUGUUCGUUUCA

>ath-miR844-3p MIMAT0004267

UUAUAAGCCAUCUUACUAGUU

>ath-miR5028 MIMAT0020534

AAUUGGGUUUAUGCUAGAGUU

>ath-miR835-5p MIMAT0004255

UUCUUGCAUAUGUUCUUUAUC

>ath-miR164c-5p MIMAT0001017

UGGAGAAGCAGGGCACGUGCG

>ath-miR156c-3p MIMAT0031867

GCUCACUGCUCUAUCUGUCAGA

>ath-miR5632-3p MIMAT0022392

UUGGAUUUAUAGUUGGAUAAG

>ath-miR4228-3p MIMAT0032109

UCGGAUGCGAAACGGUGGUGU

>ath-miR5013 MIMAT0020517

UUUGUGACAUCUAGGUGCUUU

>ath-miR774b-5p MIMAT0017739

UGAGAUGAAGAUAUGGGUGAU

>ath-miR404 MIMAT0001005

AUUAACGCUGGCGGUUGCGGCAGC

>ath-miR5661 MIMAT0022438

AGAGGUACAUCAUGUAGUCUG

>ath-miR867 MIMAT0004318

UUGAACAUGGUUUAUUAGGAA

>ath-miR472-5p MIMAT0032014

AUGGUCGAAGUAGGCAAAAUC

>ath-miR849 MIMAT0004271

UAACUAAACAUUGGUGUAGUA

>bna-miR397b MIMAT0005601

UCAUUGAGUGCAGCGUUGAUGU

>bna-miR167b MIMAT0005627

UGAAGCUGCCAGCAUGAUCUAA

>bna-miR6030 MIMAT0023650

UCCACCCAUACCAUACAGACCC

>bna-miR168b MIMAT0023628

UCGCUUGGUGCAGGUCGAGAA

>bna-miR1140 MIMAT0005637

ACAGCCUAAACCAAUCGGAGC

>bna-miR161 MIMAT0005634

UCAAUGCACUGAAAGUGACUA

>bna-miR2111b-3p MIMAT0011781

AUCCUCGGGAUACAGAUUACC

>bna-miR171g MIMAT0004446

UGAUUGAGCCGCGCCAAUAUCU

>bna-miR6028 MIMAT0023648

UGGAGAGUAAGGACAUUCAGA

>bna-miR6036 MIMAT0023656

AUAGUACUAGUACUUGCAUGAUCA

>bna-miR6035 MIMAT0023655

UGGAGUAGAAAAUGCAGUCGU

>bna-miR162a MIMAT0023621

UCGAUAAACCUGUGCAUCCAG

>bna-miR860 MIMAT0023645

UCAAUACAUUGGACUACAUAU

>bna-miR6029 MIMAT0023649

UGGGGUUGUGAUUUCAGGCUU

>bna-miR6033 MIMAT0023653

UGAACCAGAUAGAGUGGGACU

>bna-miR6034 MIMAT0023654

UCUGAUGUAUAUAGCUUUGGG

>bna-miR2111c MIMAT0023647

UAAUCUGCAUCCUGGGGUUUA

>bna-miR6032 MIMAT0023652

UGGAGCAUCAACAGAUCUCGG

>bna-miR6031 MIMAT0023651

AAGAGGUUCGGAGCGGUUUGAAGC

>bol-miR9408 MIMAT0035363

GUUUCAUCUUAGAGAAUGUUGUC

>bol-miR9409 MIMAT0035364

UUUUGUUCAUGACUGCAUUUUC

>bol-miR9411 MIMAT0035366

UACUGGACGACUUACACGGAAG

>bol-miR9410 MIMAT0035365

UACUUAAUUAUAAGUCGUCUGG

>bol-miR398a-5p MIMAT0010173

GGAGUGUCAUGAGAACACGGA

>bra-miR9565-3p MIMAT0035690

CUGAAGCUAGUGAAAGAGAGA

>bra-miR9553-5p MIMAT0035659

UACAAAGCUGAAGCUAAUUAUG

>bra-miR5717 MIMAT0023014

GUUUGGAUUGUUUGCCUUGGC

>bra-miR9567-3p MIMAT0035694

AAACUAUAUGUGUUGCUUAGA

>bra-miR9563b-5p MIMAT0035695

AAGAACUCGUCUCUUAACUUUUAA

>bra-miR9563b-3p MIMAT0035696

AAAUUAAGAGAUGAAUUCUUAC

>bra-miR9564-3p MIMAT0035688

CGAGCUGUGUAAUCGUUUUGUU

>bra-miR400-3p MIMAT0035857

GACUUAUAAUGAUCUCAUGAA

>bra-miR5718 MIMAT0023016

UCAGAACCAAACACAGAACAAG

>bra-miR9555b-5p MIMAT0035679

UGUAAUUGCGGGGUUCUAAGC

>bra-miR5716 MIMAT0023013

UUGGAUAAUUGAAGAUAUAAA

>bra-miR156a-3p MIMAT0035817

GCUUACUCUCUCUCUGUCACC

>bra-miR408-5p MIMAT0035683

GGGAGCCAGGGAAGAGGCAGU

>bra-miR172c-5p MIMAT0035872

GCAUCAUCAUCAAGAUUCAGA

>bra-miR9564-5p MIMAT0035687

ACAAAACGAUUACACAGCUCGGUC

>bra-miR9558-5p MIMAT0035669

AGAGAUGUCUGGCUUGCAACA

>bra-miR9554-5p MIMAT0035661

GAAUGAUACUUGGAUAUAAUC

>bra-miR9558-3p MIMAT0035670

UUGCAAGCCAGACAUUUCCUUU

>bra-miR1885a MIMAT0009213

CAUCAAUGAAAGGUAUGAUUCC

>bra-miR9552b-3p MIMAT0035656

UGACCGAGUAGACCGAUAGUC

>bra-miR5714 MIMAT0023011

AGACUCUACGACAUCAAGAAAC

>bra-miR9568-3p MIMAT0035698

UCAUCGUAAGAGAUCUGCAUU

>bra-miR9563a-5p MIMAT0035685

ACCCGUCUCUUAAUUUUUAAC

>bra-miR9562-5p MIMAT0035681

ACUAUGCAAUUGUGAACAAAC

>bra-miR9556-5p MIMAT0035665

GUCAAUUGGUGAUAGUAGUUC

>bra-miR9560a-5p MIMAT0035673

ACAGGUGGUGGAACAAAUAUGAGU

>bra-miR9565-5p MIMAT0035689

UCUCGUUCUCUCGUUUCAGCU

>bra-miR9567-5p MIMAT0035693

UAAACAACACAUAUAGUUUGC

>bra-miR5721 MIMAT0023019

AAAAAUGGAGUGAGAAAUGGA

>bra-miR161-3p MIMAT0035879

GUCACUUUCAAUGCGUUGAUC

>bra-miR9556-3p MIMAT0035666

UCUACUUUCACCAAUUGGCCU

>bra-miR9408-5p MIMAT0035657

CAACAGUCUCAGGAUGGAAAA

>bra-miR172b-5p MIMAT0010167

GCAGCACCAUUAAGAUUCACA

>bra-miR9568-5p MIMAT0035697

UGCGGAUAUCUUAGGAUGAGGU

>bra-miR2111b-3p MIMAT0016371

AUCCUCGGGAUACGGAUUACC

>bra-miR164e-5p MIMAT0035870

UGGAGAAGCAGGGCACGUGCAA

>bra-miR158-5p MIMAT0035701

CUUUGUCUAUCGUUUGGAAAAG

>bra-miR9552b-5p MIMAT0035655

CUAUCGGUCUACUUGGUCAGC

>bra-miR390-3p MIMAT0035841

CGCUGUCCAUCCUGAGUUUCA

>bra-miR9555b-3p MIMAT0035680

UUAGAAACUUGCAAUUAUAUA

>bra-miR408-3p MIMAT0035684

UGCUUGUUCCCUGUCUCUCUC

>bra-miR9566-5p MIMAT0035691

UUGUUGACAAAUACUUAGGCUC

>bra-miR9562-3p MIMAT0035682

UUAUUCACAACUGCAUAAUUC

>bra-miR5654b MIMAT0023008

AUAAAUCCCAAGCAUCAUCCA

>bra-miR5723 MIMAT0023022

AAUGUGCUGCAAUAUCUCUGC

>bra-miR9552a-5p MIMAT0035653

CUAUCGGUCUACUCGGUCAGC

>bra-miR5711 MIMAT0023006

UGUUUUGUGGGUUUCUACCGA

>bra-miR9555a-5p MIMAT0035663

UUCUAAGCUUUACGGGAAACC

>bra-miR9569-3p MIMAT0035700

ACACAGGAACAAUACUAACUCAUU

>bra-miR395a-5p MIMAT0035850

GUUCCUCUGAGCACUUCAUUG

>bra-miR9553-3p MIMAT0035660

UAAUCAGCUCCAGCUAUGUACA

>bra-miR5719 MIMAT0023017

UUGUGAUGAUAAUACGACUUC

>bra-miR9552a-3p MIMAT0035654

UGACCAAGUAGACCGAUAGUC

>bra-miR9561-5p MIMAT0035677

UGAGUCUCUCACCAGUCUUUCAC

>bra-miR860-5p MIMAT0035860

AUGUAGUCCAAUCUAUUGAAG

>bra-miR5713 MIMAT0023010

AGGCUUAGAAGAACGUUUGUU

>bra-miR6032-3p MIMAT0035863

UCUGCUGGUCGUUCCAUGUUAA

>bra-miR9408-3p MIMAT0035658

UUUCAUCUUAGAGAAUGUUGUU

>bra-miR5725 MIMAT0023024

AUUUGGCACAAUCUGAUCUGC

>bra-miR9559-3p MIMAT0035672

ACAAUGAACGAAAUCCAAAUC

>bra-miR9560b-3p MIMAT0035676

UCAUAUUAGUUCUACCUCCUGCUG

>bra-miR5726 MIMAT0023025

CAAAGGUUGCUUGAAUAAGGU

>bra-miR156f-3p MIMAT0035827

UGCUCACUGCUCUUUCUGUCAGA

>bra-miR9561-3p MIMAT0035678

GAGAGACUCUGAAAGACUCACC

>bra-miR319-5p MIMAT0035838

AGAGCUUCCUUGAGUCCAUUC

>bra-miR403-5p MIMAT0035858

UGUUUUGUGCGUGAAUCUAAUU

>bra-miR9560a-3p MIMAT0035674

UCAUAUUAGUUCUACCUCCUGUUG

>bra-miR2111-3p MIMAT0035875

GACCUCAGGAUGCGGAUUACC

>bra-miR5715 MIMAT0023012

ACGUGAUAAGCCUCUGAAGAA

>bra-miR172d-5p MIMAT0035876

GCAGCAUCAUUAAGAUUCACA

>bra-miR5712 MIMAT0023009

AAUAUUAAUAUAAUUGGUGAG

>bra-miR9557-5p MIMAT0035667

UUUUGCGUUUCAACUCGGUCC

>bra-miR9555a-3p MIMAT0035664

UUUCCCGUAAAGCUUAGAACC

>bra-miR9557-3p MIMAT0035668

GCUGAGUUGGAACGCAAAAUC

>bra-miR1885b MIMAT0023015

UACAUCUUCUCCGCGGAAGCUC

>bra-miR398-5p MIMAT0035854

GGGUCGACAUGAGAACACAUG

>bra-miR5720 MIMAT0023018

UUGUGAUUUGGUUGGAAUAUC

>bra-miR9554-3p MIMAT0035662

UCAUAUCCAAGUAUCAUUCCU

>bra-miR9559-5p MIMAT0035671

UUUGGAUUUUGGUCAUUGUUG

>bra-miR9563a-3p MIMAT0035686

UAAAAGUUAAGAGACAAGUUA

>bra-miR5724 MIMAT0023023

AACCGCCGGUUUGAUAAUAGC

>bra-miR9569-5p MIMAT0035699

UGAGUUAUCAUUGGUCUUGUG

>bra-miR164e-3p MIMAT0035871

CACGUGCUCCCCUCCUCCAAC

>bra-miR5722 MIMAT0023020

UGAAAUAGAGUCAUGUGGAACG

>bra-miR158-3p MIMAT0035702

UUUCCAAAUGUAGACAAAGCA

>bra-miR168a-3p MIMAT0035837

CCCGCCUUGUAUCAAGUGAAU

>bra-miR6032-5p MIMAT0035862

AACAUGGAGCAUCAACAGAUC

>bra-miR9566-3p MIMAT0035692

GAGCCUAAGUAUUUGUCAACAAUG

>cas-miR162b MIMAT0045372

AUCGAUGAACCGCUGCCUCC

>cas-miR399b MIMAT0045353

GGGCAAGAUCACCAUUGGCAGA

>cas-miR167b MIMAT0045307

GGUCAUGCUGUGACAGCCUCACU

>cas-miR827a MIMAT0045359

UUAGAUGACCAUCAACAAACG

>cas-miR160b-3p MIMAT0045292

GCGUACAGAGUAGUCAAGCAUG

>cas-miR156k-3p MIMAT0045281

UGCUUACUCUCUAUCUGUCACC

>cas-miR395c-5p MIMAT0045342

GUUCCCUUUAACGCUUCAUUG

>cas-miR159c-5 MIMAT0045263

AGAAGAAGAGGAAGAGCUCCU

>cas-miR159b-5p MIMAT0045287

AGCUGCUAAGCUAUGGAUCCC

>cas-miR157d MIMAT0045285

CAGAAGAUAGAGAGCACUGA

>cas-miR860 MIMAT0045364

UCAAUAGAUUGGACUAUAUAU

>cas-miR166d MIMAT0045302

UCGGACCAGGCUUCAUUCCCCU

>cas-miR395d-5p MIMAT0045344

GUUCUCCCGAACACUUCAUUG

>cas-miR172a-5p MIMAT0045325

GUGGCAUCAUCAAGAUUCACA

>cas-miR403 MIMAT0045355

UUAGAUUCACGCACAAACUCGU

>cas-miR158a MIMAT0045286

CUGUGCUUCUUUGUCUACAAU

>cas-miR319b MIMAT0045332

AGAGCUUUCUUCGGUCCACUC

>cas-miR857 MIMAT0045362

AAACUUUCACCAUACAAAAUAA

>cas-miR827b MIMAT0045262

UAAGAUGACCAUCAACAAAC

>cas-miR158b MIMAT0045261

UCCGAAAUGUAGACAAAGCA

>cas-miR169d MIMAT0045313

GCAAGUUGACCUUGGCUCUGU

>cas-miR3435 MIMAT0045367

GGCAUAAUUCUGUCAAUUCC

>cas-miR394 MIMAT0045339

UUUGGCAUUCUGUCCACCUCC

>cas-miR157a MIMAT0045282

GCUCUCUAGCCUUCUGUCAUCA

>cas-miR169g-3p MIMAT0045318

GGCAGUCUCCUUGGCUAUU

>cas-miR165b MIMAT0045297

GAAUGUUGUUUGGAUCGAGG

>cas-miR156d-3p MIMAT0045271

GCUCACUCUCUUUCUGUCAUA

>cas-miR156i MIMAT0045278

CAGAAGAGAGAGAGCACAAU

>cas-miR11592 MIMAT0045371

GAACCGAACCGAACCGAAA

>cas-miR167c MIMAT0045308

AGGUCAUGCUGGUAGCUUCAC

>cpa-miR8152 MIMAT0031849

UUGGACUGCUAGGUGGCCCAU

>cpa-miR8140 MIMAT0031837

CUUUUCAAGACUUCAGCUUCA

>cpa-miR8144 MIMAT0031841

AACAGUAGAACGAGUUAGAAAGGA

>cpa-miR5211 MIMAT0031851

GUGCCGUCGCACUGUGACAAG

>cpa-miR8146 MIMAT0031843

AGGAAGACGGUGAGUAGAAGCCAA

>cpa-miR8139b MIMAT0031833

UCCUGGCUGAGGACGGGUGUUGAA

>cpa-miR319 MIMAT0031782

AUUGGACUGAAGGGAGCUCC

>cpa-miR8153 MIMAT0031850

UGCACUGUAGAGCCGUAUUCGGAC

>cpa-miR172b MIMAT0031811

GGGAAUCUUGAUGAUGCUGCA

>cpa-miR8137 MIMAT0031830

UUCGCCAGCCAUUCACAAAAU

>cpa-miR8138 MIMAT0031831

UAAAGAUGGUAACAAAGGAUAA

>cpa-miR8147 MIMAT0031844

ACUGAUACUUGAUGAAUUUGCAUG

>cpa-miR164e MIMAT0031795

UGGAGAAGGGGAGCACGUGCA

>cpa-miR8155 MIMAT0031853

UAACCUGGCUCUGAUACCA

>cpa-miR8151 MIMAT0031848

UGGAUACCAGUAGACAGAUA

>cpa-miR8150 MIMAT0031847

AAAACCUGAGUCAGAUGAUGAGCG

>cpa-miR398 MIMAT0031826

GGGACGACAUGAGAUCACACG

>cpa-miR8148 MIMAT0031845

UGACUGGGUCUGCUGACGUGGCAU

>cpa-miR8154 MIMAT0031852

CAGAGGAGGAGAUGAAGAGGGA

>cpa-miR8142 MIMAT0031839

UGAGGUAAGUAGACAGUAAAGGUU

>cpa-miR8136 MIMAT0031829

UAAAGUGGAAUUGGGAUAAUA

>cpa-miR8145 MIMAT0031842

AUAAACAGAUAGAAUGACAGCCUU

>cpa-miR477 MIMAT0031828

AUUGGAGGACUUUGGGGGAGC

>cpa-miR8134 MIMAT0031825

UGAGAAUUAUGCGGAGGAUGU

>cpa-miR8135 MIMAT0031827

AGGAUUUUGCAGGGUUGAU

>cpa-miR8141 MIMAT0031838

UGGAUACUAGUAGGCUGGUU

>cpa-miR8149 MIMAT0031846

AGCGAAGGGGACGCCUGAAGACUC

>cpa-miR8143 MIMAT0031840

GAGAGAUGGUGGACAGAUCAGGUA

>cme-miR172f MIMAT0026099

UGAAUCUUGAUGAUGCCGCAC

>cme-miR7129 MIMAT0026183

AGUCAAAUCUAAACGAUCGUGUAU

>cme-miR164a MIMAT0022749

UGGAGAAGCAGGGCACGUGCU

>cme-miR169m MIMAT0026092

UAGCCAAAAAUGACUUGCCUGC

>cme-miR166i MIMAT0026091

UCGGACCAGGCUUCAUUCUC

>cme-miR169t MIMAT0026187

UGAGCCAAGAAUGACUUGCCGGC

>cme-miR156j MIMAT0026186

GUUGACAGAAGAGAGUGAGCAC

>cme-miR854 MIMAT0026100

GAUGAGGAUAGUGAGGAGGAG

>cme-miR477a MIMAT0026089

ACCUCCCUCAAAGGCUUCCAA

>cme-miR169q MIMAT0026073

UGAGCCAAAGAUGACUUGCCU

>cme-miR156e MIMAT0026156

UUGACAGAAGAUAGAGGGCAC

>cme-miR399g MIMAT0026185

AGGGCUUCUCUCCAUUGGCAGG

>cme-miR845 MIMAT0026084

UCGCUCUGAUACCAAUAUGAUG

>cme-miR7130 MIMAT0026184

GUUUGGAAUGUGCGAGAUGUGUGC

>cme-miR171i MIMAT0026085

UUGAGCCGUUCCAAUAUCACA

>cme-miR1863 MIMAT0026101

AGCUCUGAUACCAUGUUAGAUUUG

>cme-miR399f MIMAT0026087

UGCCAAAGGAGAAUUGCAC

>cme-miR164b MIMAT0026088

UGGAGAGGCAGGGCACAUGCU

>cme-miR395b MIMAT0026157

UUGAAGUGUUUGGGGGAACUC

>cme-miR858 MIMAT0026083

UCUCGUUGUCUGUUCGACCUU

>cme-miR530b MIMAT0026104

UGCAUUUGCACCUACACCUU

>cme-miR159b MIMAT0026168

AUUGGAUUGAAGGGAGCUCCU

>cst-miR11334 MIMAT0044624

AAUUACUAUAAUAACACCUUCACAU

>cst-miR11332 MIMAT0044622

CUUGUGGGUAAUAGGCUUUCCUUCCUUG

>cst-miR11333 MIMAT0044623

ACUUCGCGGUAAGUCUAACCCUAAUU

>cst-miR11331 MIMAT0044621

CGUUGAGUAUUAUCUUUCAGUUC

>cst-miR11335 MIMAT0044625

GUUGGGAGGAUGGAGCGGUU

>hbr-miR6485 MIMAT0025295

UAGGAUGUAGAAGAGCAUAA

>hbr-miR6168 MIMAT0024786

UCUUGGUGCGGUGGACUGCUG

>hbr-miR2118 MIMAT0024784

GAAAUGGGUGGAUGGGAGUGA

>hbr-miR6175 MIMAT0024796

UGGUGGCUUUUAGGGCUCAAG

>hbr-miR9386 MIMAT0035235

UUUGCAGUUCGAAAGUGGAAGC

>hbr-miR6172 MIMAT0024792

UGGACCGUGUUUAAGUAAGAA

>hbr-miR482a MIMAT0024787

AGAUGGGUGGCUGGGCAAGAAG

>hbr-miR476 MIMAT0025290

UAAUCCUUCUUUGCAAAGUC

>hbr-miR9387 MIMAT0035236

GAACACAAUUAUAGGAAUC

>hbr-miR6174 MIMAT0024795

AUGCUAAGCUGUCAGACUUUG

>hbr-miR6483 MIMAT0025292

UAUUGUAGAAAUUUUCAGGAUC

>hbr-miR6482 MIMAT0025291

CAGGAACUGGUAUCAACCCAGC

>hbr-miR408b MIMAT0025289

ACUGGGAACAGGCAGAGCAUGA

>hbr-miR396a MIMAT0024791

CACAGCUUUCUUGAACUUUCU

>hbr-miR408a MIMAT0025288

AAGACUGGGAACAGGCAGAGCA

>hbr-miR482b MIMAT0035237

GAAUGGGCGGUUUGGGAAAGA

>hbr-miR6169 MIMAT0024788

UAGUAUUUCUCUUUUCUCUCU

>hbr-miR6173 MIMAT0024793

AGCCGUAAACGAUGGAUACU

>hbr-miR398 MIMAT0024794

GGAGCGACCUGAGAUCACAUG

>hbr-miR6170 MIMAT0024789

CAAGAAACAGAAGAGAGGGAU

>hbr-miR6166 MIMAT0024783

AUUGGAUAACGGGAUUACAUGGU

>hbr-miR6171 MIMAT0024790

AUGUGGAUUGCUGAAGGCUUU

>hbr-miR6484 MIMAT0025293

UAAUGGGCUCUGCAUAGAUGG

>hbr-miR6167 MIMAT0024785

UACCCAGGUGGAAGCUUUGA

>mes-miR1446 MIMAT0029305

UUCUGAACUCUCUCCCUCAU

>mes-miR11891 MIMAT0045987

CAUAAAUUGAACUAUAGACC

>mes-miR482d MIMAT0045978

UUCCCGACACCACCCAUUCCAU

>mes-miR399h MIMAT0045972

GGGCACCUCUCGCUUGGCAGG

>mes-miR1446b MIMAT0045982

UGAACUCUCCCCCUCAACGGCU

>mes-miR159a-5p MIMAT0037459

AGCUGCUGAGCUAUGGAUCCC

>mes-miR482b MIMAT0045976

UUCCCAAUGUCGCCCAUUCCGA

>mes-miR169l MIMAT0029220

CAGCCAAGAAUGACUUGCCGG

>mes-miR160g MIMAT0029185

UGCCUGGCUCCCUGUAUGCCAUC

>mes-miR2275 MIMAT0029308

UUUGGUUUCCUCCAAUAUCUUA

>mes-miR477i MIMAT0029289

ACUCUCCCUCAAGGGCUUCCG

>mes-miR535b MIMAT0029301

UGACAACGAGAGAGAGCACGG

>mes-miR319h MIMAT0029260

CUUGGACUGAAGGGAGCUCCU

>mes-miR535d MIMAT0045981

UUGACGACGAGAGAGAGCACG

>mes-miR169f MIMAT0029214

UAGCCAAGGAUGACUUGCCGG

>mes-miR482 MIMAT0029297

UCUUCCCUACUCCACCCAUUCC

>mes-miR159d MIMAT0029178

AUUGGAGUGAAGGGAGCUCUG

>mes-miR166i MIMAT0029200

UUGGACCAGGCUUCAUUCCCC

>mes-miR169ad MIMAT0045986

UCACAGGCUCUUAUUUUUCAUG

>mes-miR3627 MIMAT0045983

UUGUCGCAGGAGCGGUGGCACC

>mes-miR397 MIMAT0029279

UUUGAGUGCAGCGUUGAUGA

>mes-miR172c MIMAT0029249

UGAAUCUUGAUGAUGCUACGC

>mes-miR530b MIMAT0029299

UGCAUUUGCACCUGCACCUUA

>mes-miR2118 MIMAT0045975

GUUCCCAUGCCACCCAUUUCUA

>mes-miR11892 MIMAT0045988

UUGUCAUCUCAACCUUGUGUC

>mes-miR482c MIMAT0045977

UUUUCCCAAGACCUCCCAUACC

>mes-miR535c MIMAT0045980

UUGACGACGAGAGAGAGCACA

>mes-miR160h MIMAT0029186

UGCCUGGCUCCCUGUAUGCCAUU

>mes-miR171d MIMAT0029240

AUGAGCCGUGCCAAUAUCACG

>mes-miR477j MIMAT0045973

ACUCUCCCUAAAGGCUUCAAC

>mes-miR482e MIMAT0045979

UCUUACCUACACCGCCCAUGCC

>mes-miR535a MIMAT0029300

UGACAACGAGAGAGAGCACGU

>mes-miR477h MIMAT0029288

ACUCUCCCUCAAGGGCUUCAG

>aau-miR160 MIMAT0022672

UGGCAUACAGGGAGCCAGGCA

>aau-miR168 MIMAT0022678

AUUCAGUUGAUGCAAGGCGGGAUC

>aau-miR2086 MIMAT0022676

GACAUGAAUGCAGAACUGGAA

>ahy-miR3511-3p MIMAT0016335

UGUUACUAUGGCAUCUGGUAA

>ahy-miR3517 MIMAT0016343

CUGACCACUGUGAUCCCGGAA

>ahy-miR3511-5p MIMAT0016334

GCCAGGGCCAUGAAUGCAGA

>ahy-miR3510 MIMAT0016333

UUAUACCAUCUUGCGAGACUGA

>ahy-miR3518 MIMAT0016344

UGACCUUUGGGGAUAUUCGUG

>ahy-miR3519 MIMAT0016345

UCAAUCAAUGACAGCAUUUCA

>ahy-miR3513-5p MIMAT0016337

UUAAUUUCUGAGUUUGUCAUC

>ahy-miR3520-3p MIMAT0016347

AAGGGAGACGUUUGAAUUAUC

>ahy-miR3514-5p MIMAT0016339

AGGAUUCUGUAUUAACGGUGGA

>ahy-miR3513-3p MIMAT0016338

UUGAUAAGAUAGAAAUUGUAU

>ahy-miR3512 MIMAT0016336

CGCAAAUGAUGACAAAUAGA

>ahy-miR3514-3p MIMAT0016340

UCACCGUUAAUACAGAAUCCUU

>ahy-miR3515 MIMAT0016341

AAUGUAGAAAAUGAACGGUAU

>ahy-miR3509-3p MIMAT0016332

AUCUAACGACUCUCAGAUAUCA

>ahy-miR3508 MIMAT0016330

UAGAGGGUCCCCAUGUUCUCA

>ahy-miR3520-5p MIMAT0016346

AGGUGAUGGUGAAUAUCUUAUC

>ahy-miR3509-5p MIMAT0016331

AUACUUGAGAGCCGUUAGAUGA

>ahy-miR3516 MIMAT0016342

GCUGGGUGAUAUUGACAGAAG

>ahy-miR3521 MIMAT0016348

UGGUGAGUCGUAUACAUACUG

>ahy-miR160-3p MIMAT0016323

GCAUGAAGGGAGUCACGCAGG

>gma-miR10447 MIMAT0041727

UGUUUCCAUGUUGUUGAGUGAC

>gma-miR9732 MIMAT0036351

CAAGGGUAUGAUGUGCAAUCU

>gma-miR5038a MIMAT0021063

UGAGAAUUUGGCCUCUGUCCA

>gma-miR5672 MIMAT0022454

CAUGGUAGUGGAAGAAAUGGA

>gma-miR10406b MIMAT0041685

CAGUCGAGUUAUUGUAUAAUUCAC

>gma-miR10186d MIMAT0041653

UUUUGGGAAUUAAAAUGUAAACUU

>gma-miR5030b MIMAT0040957

UUCCGGAAGAACAAAGCUACC

>gma-miR10419 MIMAT0041665

CAAAAAACACAUAUGAAACUG

>gma-miR1520f-3p MIMAT0018244

CAAUCAGAACAUGACACAUGACAA

>gma-miR9739 MIMAT0036359

UUUGAAUGUCCAGAUACGUAC

>gma-miR319p MIMAT0036379

UUUUGGACUGAAGGGAGCUCC

>gma-miR1520l MIMAT0018292

AAUCAGAACAUGACACGUGAUAGU

>gma-miR2606a MIMAT0023226

AAAAGCACUUAAGGAACGGUA

>gma-miR403b MIMAT0021614

UUAGAUUCACGCACAAACUUG

>gma-miR5371-5p MIMAT0021608

UAGGAAUUAGUCACUCAGAUC

>gma-miR482b-5p MIMAT0018286

UAUGGGGGGAUUGGGAAGGAAU

>gma-miR169w MIMAT0041676

CAAGGAUGACUUGCCGGCAUU

>gma-miR10430 MIMAT0041695

UCGGUCUGACCUUUUUAAAAGCCU

>gma-miR1532 MIMAT0007394

AACACGCUAAGCGAGAGGAGCUC

>gma-miR408c-5p MIMAT0022993

CAGGGGAACAGGCAGAGCAUG

>gma-miR4994-3p MIMAT0032119

UGAUAUCCUUGAGCUAAUACA

>gma-miR391-5p MIMAT0018280

UACGCAGGAGAGAUGACGCUGU

>gma-miR1524 MIMAT0007385

CGAGUCCGAGGAAGGAACUCC

>gma-miR10424d MIMAT0041722

UUUUCUAAUUUAUUAGGGACU

>gma-miR9760 MIMAT0036395

UGGAUGAUGUAGUUUUGAUUG

>gma-miR4358 MIMAT0018250

CAGUGCAUGACUAUAUCGCCAG

>gma-miR4398 MIMAT0018314

UGUCAGCGGAGUGAGAAGACGAAA

>gma-miR393b MIMAT0023170

UUUGGGAUCAUGCUAUCCCUU

>gma-miR1520h MIMAT0018255

AACGUCCAAUCAGAACGUGACAUG

>gma-miR171b-5p MIMAT0017336

ACGGCGUGAUAUUGGUACGGCUC

>gma-miR395e MIMAT0023244

UGAAGUGUUUGGGGGAACUUU

>gma-miR4353 MIMAT0018242

CAAGUCGUAGCCGGUGUUAUUACU

>gma-miR4996 MIMAT0021014

UAGAAGCUCCCCAUGUUCUC

>gma-miR10405d MIMAT0041704

UUGUUUCUUAUAAAAAGGACC

>gma-miR4379 MIMAT0018285

UAGAGUGUAUACUGUGAGAGGCCU

>gma-miR10426 MIMAT0041682

UAGAAAAGAAUCAAUGUAGAAAGU

>gma-miR10441 MIMAT0041714

CAACCCUGAGAACAAUGAAAUCGU

>gma-miR399m MIMAT0041681

GGGCUCCUCUCUCCUGGCAUG

>gma-miR2118b-5p MIMAT0022982

GGAGAUGGGAGGGUCGGUAAAG

>gma-miR1513c MIMAT0021674

UAUGAGAGAAAGCCAUGAC

>gma-miR2108a MIMAT0010084

UUAAUGUGUUGUGUUUGUCGG

>gma-miR4366 MIMAT0018265

CUACUUAGUAGAGAUUUGUUGG

>gma-miR1512a-5p MIMAT0007370

UAACUGAAAAUUCUUAAAGUAU

>gma-miR5679 MIMAT0022467

UUGGUGACCCAGAAGAAGUUGA

>gma-miR4389 MIMAT0018302

UCGGUCGGACCGAUCCAAUCGGAA

>gma-miR159d MIMAT0018340

AGCUGCUUAGCUAUGGAUCCC

>gma-miR9748 MIMAT0036375

GAAGGAAGUGUAGAGGGAUGAC

>gma-miR5784 MIMAT0023190

AAUUAGCUAAUGGUUAGCUAA

>gma-miR4411 MIMAT0018331

UUAUUGUAACUAAUUUGUCGGU

>gma-miR4401a MIMAT0018319

ACAACGUCUUUGAAAGUAGGCAUU

>gma-miR10438 MIMAT0041710

UUGGACUAAGGUUUUUUGGCAC

>gma-miR4997 MIMAT0021015

GAUCGUCAAGCGCGAAGAUGAGG

>gma-miR4412-3p MIMAT0018337

AGUGGCGUAGAUCCCCACAAC

>gma-miR4404 MIMAT0018323

AUUCGUGGAAGACUGGCGGAUCAA

>gma-miR530c MIMAT0023197

UGCAUUUGCACCUGCACUUUA

>gma-miR171i-5p MIMAT0021056

AUAAGAAAGCAAUGCUCAAA

>gma-miR4415b-3p MIMAT0021676

UUGAUUCUCAUCACAACAUGG

>gma-miR1521a MIMAT0007382

CUGUUAAUGGAAAAUGUUGA

>gma-miR5035-3p MIMAT0032121

UGUUUAGAAGCUCAUAGAAUAGAU

>gma-miR4390 MIMAT0018303

UCGUACUCGUCGGGUAUCGGGUAU

>gma-miR9723 MIMAT0036338

CAAAGGAGAUUUGGACAACUC

>gma-miR5043 MIMAT0021074

UGUCCCCUUCUCUGCACCACC

>gma-miR10422 MIMAT0041669

UUCUGAUUAGAGAGCAACACC

>gma-miR1516d MIMAT0024927

GUACUUGUGGCUUGUAUCCAA

>gma-miR4344 MIMAT0018232

AAGUAGACAUUCUAAGACGUUGCU

>gma-miR319i MIMAT0021663

UUGGACUGAAGGGGAGCUCCUUC

>gma-miR1520m MIMAT0018293

AAUCAGAACAUGACAUGUGACAAU

>gma-miR4363 MIMAT0018259

CGAUUACCAGAAGGCUUAUUAG

>gma-miR2111a MIMAT0022457

GUCCUUGGGAUGCAGAUUACG

>gma-miR4383 MIMAT0018290

UAUUGGAUCUCAGUUGAACCGGUC

>gma-miR4370 MIMAT0018271

AGUAGACUCGUCCGAUUUUGCGUA

>gma-miR9752 MIMAT0036381

UGCUUCUUCUUUUCCCUGUUU

>gma-miR5773 MIMAT0023175

UUUUUAAAAGGUUCAGUUAGGU

>gma-miR5671a MIMAT0022453

CAUGGAAGUGAAUCGGGUGAC

>gma-miR4357 MIMAT0018249

CAGUCGUGUGAUUGUACGGUUCAU

>gma-miR5770a MIMAT0023167

UUAGGACUAUGGUUUGGACGA

>gma-miR5764 MIMAT0023161

UCCAUUCGCGGACAUGAUGGAU

>gma-miR159f-5p MIMAT0021642

GAGUUCCCUGCACUCCAAGUC

>gma-miR10409 MIMAT0041646

CCUCUGAAGAUAUUAAAAGCCU

>gma-miR1520c MIMAT0007395

UUCAAUAAGAACGUGACACGUGA

>gma-miR4414b MIMAT0041688

AGCUGCUGACUCGUUGGUUCG

>gma-miR10418 MIMAT0041661

GAAGUAAUCCUAGGAACUCCCU

>gma-miR5044 MIMAT0021075

GUAGUGGAUGCCUAGAGGUCCA

>gma-miR9745 MIMAT0036365

AGAAUUAAAUUUGGACCGUAUAAC

>gma-miR9751 MIMAT0036380

GUAAUUUUAAACCUAAACCCUAAA

>gma-miR9740 MIMAT0036360

UGUAGGUUCCAGUGAGGGAAA

>gma-miR5780b MIMAT0036341

UCUGAGUCCAUGAUAUAUUAAA

>gma-miR1516c MIMAT0023178

AAUGUCUGGGCUUAGCGAGGCGGU

>gma-miR4415b-5p MIMAT0022997

AAGUUGUGAUGGGAAUCAAUGGCA

>gma-miR171r MIMAT0023229

CGAGCCGAAUCAAUACCACUC

>gma-miR4377 MIMAT0018281

UACGUCAUCGCUGAAUGGAAGACG

>gma-miR4410 MIMAT0018329

UAUGUUGAUCCGUAUGAGUCGUAC

>gma-miR9738 MIMAT0036358

UGAAACAUGAUGUGGACUCUUC

>gma-miR169i-3p MIMAT0021068

CCGGUGCCAUCCCGUCUCAUA

>gma-miR4374c MIMAT0041680

CAACACCGUCUUUGAAGCCUGG

>gma-miR169l-3p MIMAT0022994

CGGGCAAGUUGUUUUUGGCUAC

>gma-miR159c MIMAT0007352

AUUGGAGUGAAGGGAGCUCCG

>gma-miR1525 MIMAT0007386

UGGGUUAAUUAAGUUUUUAGU

>gma-miR159f-3p MIMAT0021643

AUUGGAGUGAAGGGAGCUCCA

>gma-miR4368b MIMAT0018269

AAGGACGGUACUUACGUAAGCAAC

>gma-miR4365 MIMAT0018264

AAGAACUUCUUCCGCGAGAUCGCA

>gma-miR10198 MIMAT0040965

AAGGAACUCGAAAUUCAUAGU

>gma-miR156s MIMAT0023216

UGACAGAAGAGAGUGAGCACU

>gma-miR5369 MIMAT0021603

UGAGAAAAGGAGGAUGUCA

>gma-miR10437 MIMAT0041709

UAUUGUUCAAACAUAUACUGUU

>gma-miR5786 MIMAT0023222

UGUCGCAGGAUAGAGGGCACU

>gma-miR1507c-5p MIMAT0021005

GAGGUGUUUGGGAUGAGAGAA

>gma-miR9750-5p MIMAT0039317

AGGCGCGUAUGGACUUACGACC

>gma-miR5042-3p MIMAT0032123

UGGGGCUUGAUCCAAGAUAGG

>gma-miR4354 MIMAT0018246

CAAUUGGAUCGGUCCAACCGGC

>gma-miR9750-3p MIMAT0036378

UGUAAGUCCAUAUGUGCUCUCU

>gma-miR1521b MIMAT0021065

GACUGUCACGUGUCAUAAUCAUA

>gma-miR1515a MIMAT0007374

UCAUUUUGCGUGCAAUGAUCUG

>gma-miR9746f MIMAT0036371

AAAGUGUUUGAAUCUCAAUUAGAU

>gma-miR10407b MIMAT0041706

AGUUAACGGAUGAAUGAAUUUGUC

>gma-miR5370 MIMAT0021607

CUAAAGAUUGUCCAAAAGGAA

>gma-miR9724 MIMAT0036339

UAGAGAUAGUGUCAAAAUAGAA

>gma-miR9758 MIMAT0036392

UGUUUAGUCAUGCAAGUUUAG

>gma-miR4362 MIMAT0018258

CCUUAGGACAGACGUCAUGUAG

>gma-miR5378 MIMAT0021619

CAUCUGAAGGAUAGAACACAUA

>gma-miR482d-3p MIMAT0021672

UCUUCCCUACACCUCCCAUACC

>gma-miR4416a MIMAT0018344

ACGGGUCGCUCUCACCUAGG

>gma-miR10431 MIMAT0041697

UUAGAAUAAACAUGUGUAGAAAUU

>gma-miR4396 MIMAT0018311

UGUAGUUUCUAAGACGAUGCUGAC

>gma-miR4351 MIMAT0018240

AUUGGGAUUCAGUUGGAGUUGG

>gma-miR9766 MIMAT0036403

UUUGAAGGGAAGGAAUGAAAC

>gma-miR4405b MIMAT0041662

UUCCGAAUAACCGACUUAGAAAUC

>gma-miR4392b MIMAT0040970

UCUGCGAAAAUGUGAUUUCGGA

>gma-miR5762 MIMAT0023158

UCAUAGGAGGAAUCAACUGGC

>gma-miR10435 MIMAT0041703

UUCCAUUUCAACGUCGGGUUAGAA

>gma-miR10411 MIMAT0041648

UAAUUUUAGACUGAUAUUUUGCCU

>gma-miR9756 MIMAT0036387

UGAGAACUUUAUCCAAACAGAG

>gma-miR5037b MIMAT0021605

AACCCUCAAAGGCUUCCUAG

>gma-miR169h MIMAT0021062

GGCGAGACAUCUUGGCUCAUU

>gma-miR1522 MIMAT0007383

UUUAUUGCUUAAAAUGAAAU

>gma-miR10190 MIMAT0040955

UCCUGAUGACUAUUAUGAGCU

>gma-miR5041-3p MIMAT0032122

GUUGAGCAAGUUGAAGAUGAA

>gma-miR4368a MIMAT0018267

AAGACGGUACUUACCUCAGUAACA

>gma-miR1528 MIMAT0007390

AUAGAUUAGAUCAAUAUAUUAGU

>gma-miR4378b MIMAT0018284

UAGAACUGUCUUAGAAUGUGCUAC

>gma-miR9767 MIMAT0036404

AUGGAAUGGUUACUUAUGAAAAGA

>gma-miR5674b MIMAT0023248

UAAUUGUGUUGUACAUUAUCA

>gma-miR1509b MIMAT0011201

UUAAUCAAGGAAAUCACGGUU

>gma-miR4399 MIMAT0018315

UUAACGAAAAAGGACUAACGAC

>gma-miR5676 MIMAT0022462

UCGACACCAUAUGUAGAGGCAG

>gma-miR171p MIMAT0023208

UUGAGCCGCGUCAAUAUCUUA

>gma-miR1520r MIMAT0018313

UGUCACAUCCUGGUUGGACAUGAA

>gma-miR4397-3p MIMAT0018312

UGUCAAAGAUGUGGCGAAUACU

>gma-miR5559 MIMAT0023168

UACUUGGUGAAUUGUUGGAUC

>gma-miR4342 MIMAT0018230

AAUCGACUUAGAAUGUAGGAUGGU

>gma-miR9749 MIMAT0036376

UUAGCUUCUUUCACCUUUCCC

>gma-miR9728 MIMAT0036346

CGCAGAACUGAAACAAGUUGA

>gma-miR1520o MIMAT0018296

UCAUCGUCCAAUCAGAAUGUGACA

>gma-miR1519 MIMAT0007378

UAAGUGUUGCAAAAUAGUCAUU

>gma-miR1508a MIMAT0007366

UCUAGAAAGGGAAAUAGCAGUUG

>gma-miR169c MIMAT0007357

AAGCCAAGGAUGACUUGCCGA

>gma-miR1520i MIMAT0018257

AACGUGACACGUGACGGUCAACAU

>gma-miR1514a-3p MIMAT0032031

AUGCCUAUUUUAAAAUGAAAA

>gma-miR171h MIMAT0021054

AUUGAGACGAGCCGAAUCAAU

>gma-miR5774b MIMAT0023193

GCUGGCGUCGACACGUGGCAU

>gma-miR5379 MIMAT0021620

AUGAAAAUCAUUCAUUAUGAUAUC

>gma-miR10201 MIMAT0040969

UACCGGUAGUAAAUGGAUGCCU

>gma-miR10448 MIMAT0041729

UUAGGCUUUGUGGGACGUGAUU

>gma-miR10445 MIMAT0041724

AAGGACCAAAUUUAUGAAUUAAAU

>gma-miR9724b MIMAT0041687

UAUCUUGGCACUAUCUCUAAC

>gma-miR169u MIMAT0023249

CAGCCAAGGAUGACUUGCCGU

>gma-miR5368 MIMAT0021602

GGACAGUCUCAGGUAGACA

>gma-miR4997b MIMAT0041691

UCGCGGCAGAAGAAAUCCUGAU

>gma-miR5380b MIMAT0021622

GAAAAUGAAUGAUGAGGAUGGGGA

>gma-miR4348b MIMAT0036344

UCUUUUGAAUUUGACUAUUAG

>gma-miR172h-5p MIMAT0021657

GCAGCAGCAUCAAGAUUCACA

>gma-miR4346 MIMAT0018235

GAAAGACCAAACGAGAAGCUGCAU

>gma-miR4403 MIMAT0018321

ACGGACACCGAACACGACACGGAC

>gma-miR5376 MIMAT0021617

UGAAGAUUUGAAGAAUUUGGGA

>gma-miR10191 MIMAT0040956

GGCGACUGCGGCUCCUCCGCCG

>gma-miR169r MIMAT0023217

UGAGCCAGGAUGGCUUGCCGGC

>gma-miR5034 MIMAT0021055

GGUACCCUUUCAGAUAGUCUCA

>gma-miR862a MIMAT0021004

UGCUGGAUGUCUUUGAAGGAAU

>gma-miR9762 MIMAT0036399

UACAGAUCUUUGGAAACAGGC

>gma-miR169s-5p MIMAT0023218

AAGCCAAGGAUGACUUGCCGG

>gma-miR9730 MIMAT0036349

CGAUUGCUGUCAUAACUGCUGC

>gma-miR166i-5p MIMAT0021645

GGAAUGUCGUCUGGUUCGAG

>gma-miR5371-3p MIMAT0021609

UCUCAGUGACUAAUUUCUAGA

>gma-miR9736 MIMAT0036356

UGAAAGACAAACAAAGGUGGG

>gma-miR1513a-5p MIMAT0007371

UGAGAGAAAGCCAUGACUUAC

>gma-miR4387d MIMAT0018301

AUGUCACUGAUUAGGCAUGAUGAU

>gma-miR4348a-5p MIMAT0018237

AAACUUGUAAGAUGGUGACAUU

>gma-miR4349-3p MIMAT0037390

UCACUUAUAUACUCUUUCUUGGCC

>gma-miR10425 MIMAT0041675

UUACAACCGUCUUUGAAAUCGCCU

>gma-miR5380c MIMAT0036391

AUGAAUGGUGAAGAUGAAGAG

>gma-miR1520f-5p MIMAT0032111

AUUGUCACGUGUCAUGUUCUGAUU

>gma-miR4402 MIMAT0018320

ACAUAUUAUGGGUCUCAGACGGAC

>gma-miR10413b MIMAT0041659

UUUUGGUAGAGAACGAAACCCU

>gma-miR395h MIMAT0024920

AUGAAGUGUUUGGGAGAACUC

>gma-miR4348c MIMAT0036389

UGUUAAACUUGCAAGAUGACA

>gma-miR1516a-5p MIMAT0020937

CAAGUUAUAAGCUCUUUUGAGAG

>gma-miR5669 MIMAT0022451

CAAUGUAGUGUGGUAAGUGGUC

>gma-miR9761 MIMAT0036397

UGAAGGUCUAGGAUAUUUUGU

>gma-miR5677 MIMAT0022465

UUUGGUCUUUAAUCAAGCUGA

>gma-miR1523b MIMAT0021051

UCAUCGCUCCUGAGCUCACA

>gma-miR5779 MIMAT0023184

CAAGUCCAAAGUAGGAAUGUUGCA

>gma-miR398b-5p MIMAT0037314

GAGUGGAUCUGAGAACACAAGG

>gma-miR4995 MIMAT0021013

AGGCAGUGGCUUGGUUAAGGG

>gma-miR396j MIMAT0023214

AUUCAAGAUAGCUGUGGAAAA

>gma-miR4378a MIMAT0018283

AUAGGACUGUCUUAGAAUGGUGUA

>gma-miR169j-5p MIMAT0021649

UAGCCAAGAAUGACUUGCCGG

>gma-miR10193f MIMAT0041725

UAAAAAAACCAAUGUUAACUGU

>gma-miR4994-5p MIMAT0021012

GGUUAGCUCAAGGAUCUCAC

>gma-miR171a MIMAT0007358

UGAGCCGUGCCAAUAUCACGA

>gma-miR9755 MIMAT0036386

UGAUCCAGGAACUUUUCAUCU

>gma-miR1527 MIMAT0007389

UAACUCAACCUUACAAAACC

>gma-miR1533 MIMAT0007396

AUAAUAAAAAUAAUAAUGA

>gma-miR4341 MIMAT0018229

UGUGUUGAAAGUUUAACAUGACGG

>gma-miR10412 MIMAT0041649

GAACGUUCUGGAGAACUCUAGAGU

>gma-miR4409 MIMAT0018328

UAACAAGUGGGUUUGUUGACUG

>gma-miR10199 MIMAT0040966

AGCAAUGUUGAGCUUGGGCCU

>gma-miR482a-3p MIMAT0007364

UCUUCCCAAUUCCGCCCAUUCCUA

>gma-miR9737 MIMAT0036357

UUGUGGCUGAAAUCACUGUUGC

>gma-miR167h MIMAT0021064

AUCAUGCUGGCAGCUUCAACUGGU

>gma-miR5040 MIMAT0021067

AUGAUAUAUAACAAGCAUGAG

>gma-miR4393a MIMAT0018308

UGAGAAAAGGACGGCAGAAAAGCC

>gma-miR156p MIMAT0023196

UUGACAGAAGAAAGGGAGCAC

>gma-miR10442 MIMAT0041715

CUACAUGCUGCACUUGGAUCC

>gma-miR169t MIMAT0023233

UAGCCAAGGAUGGACUUGCCUA

>gma-miR9729 MIMAT0036347

GUAAUGAGUAGAAACAUUUAGAAG

>gma-miR5785 MIMAT0023191

UAGUGUUGUCCUGUCGAACACGGA

>gma-miR169n-5p MIMAT0021654

CAGCCAAGGGUGAUUUGCCGG

>gma-miR4345 MIMAT0018233

UAAGACGGAACUUACAAAGAUU

>gma-miR5678 MIMAT0022466

UUCCAUGAUAAGAUCUUUGAC

>gma-miR4374a MIMAT0018278

UAAGACGGUCGUGAUGUCAGCA

>gma-miR391a-3p MIMAT0020951

AGCAUCAUAUCUCCUGCAUAG

>gma-miR5761b MIMAT0023173

UUUUGUGUCGUGAAGCUUUUG

>gma-miR9747 MIMAT0036374

CAUGCGAUGAUUUAAAUACUUU

>gma-miR169j-3p MIMAT0021650

UUUCGACGAGUUGUUCUUGGC

>gma-miR5033 MIMAT0021053

GGCUGUACAAAAGGAAACUAC

>gma-miR9735 MIMAT0036355

UACGGCUUAAGUUCAACUUUGGAG

>gma-miR5765 MIMAT0023162

CGAAACGUUGAGGUAUAUGUGGAC

>gma-miR4393b MIMAT0018317

UUGAAAAGGGACAGCAGAGAAGCC

>gma-miR10192 MIMAT0040958

UAAACGUUUGAUCCCUUGUAU

>gma-miR5772 MIMAT0023174

AGAAUGUGAGUUAGAGUGAGCAUC

>gma-miR10189 MIMAT0040954

AAGAACACUAGAACCAUCUCC

>gma-miR4371a MIMAT0018273

AAGUGAUGACAUGACAAGCGAAGU

>gma-miR5771 MIMAT0023171

AUCUCAAGUGGAUUGCUUAAGGAC

>gma-miR396g MIMAT0021071

UUCUUGAACUUCUUAUGCAUC

>gma-miR1520a MIMAT0007381

UAGAACAUGAUACAUGACAGUCA

>gma-miR10407d MIMAT0041655

UUAACGGCUGGAUAUAUUUGUUGC

>gma-miR4416c-3p MIMAT0037453

ACGGGUCGCUCUCACCUGGAG

>gma-miR9734 MIMAT0036354

UCGUGAAUGAGAUUUGUGUUGCUU

>gma-miR10407c MIMAT0041650

UUAACAGUCGGAUAUAUUUAUCGC

>gma-miR4387b MIMAT0018272

AAGGUGUGAUGGCAUGACACUCUG

>gma-miR1520q MIMAT0018332

AUUGACCAAUCAGAACAUGACACA

>gma-miR4350 MIMAT0018239

UCAAAUGAUUUUGUGUCGUUGG

>gma-miR397b-3p MIMAT0022991

UAUUGACGCUGCACUCAAUCA

>gma-miR1510a-5p MIMAT0017338

AGGGAUAGGUAAAACAAUGACUGC

>gma-miR4352a MIMAT0018241

AUUUCUAGGACAUACUACGACGGU

>gma-miR4391 MIMAT0018304

UCUCGGCAAAGAACUAAGAAGAAG

>gma-miR4400 MIMAT0018316

UUCGGAAAAAUUCUGGAAGACGUC

>gma-miR167k MIMAT0036388

UGAAGCUGCCAGCCUGAUCUUA

>gma-miR4372b MIMAT0021073

UAAUAAAAUCGUGACAUGUAAC

>gma-miR5037d MIMAT0022455

CGGGAGCCUAUGAAGGUUAAC

>gma-miR5375 MIMAT0021616

ACUAUAGAAGUACUUGUGGAGC

>gma-miR5781 MIMAT0023186

CUGAAACUGAGACUGCAUCUGG

>gma-miR5032 MIMAT0021052

AGAGCCACUUUUGGGUUCCCUAU

>gma-miR5667-5p MIMAT0032130

AAUCCAUUCAGAUCUGUUUCG

>gma-miR5768 MIMAT0023165

AAGUGCAAUACUGAUCUUCGGAAC

>gma-miR4382 MIMAT0018289

UAUGUUAACUGAUUUCAUGGAU

>gma-miR408b-5p MIMAT0021630

CUGGGAACAGGCAGGGCACG

>gma-miR9733 MIMAT0036353

UAACAGACUUAGAUUAACAGA

>gma-miR319f MIMAT0020994

UUGGACUGAAGGGGCCUCUU

>gma-miR4371c MIMAT0018268

GACGUGACAGACGGAAUAUCACAU

>gma-miR4375 MIMAT0018279

UACCACUAGUGGUCGCGCCUGGCA

>gma-miR4380b MIMAT0018287

UAUGGUCAUACGGAUUGUUGAU

>gma-miR4388 MIMAT0018300

AAUCUUAGGGACCAAAUUGACAGC

>gma-miR4364b MIMAT0018234

UAACAACAGCGGAAGAACCUUCUU

>gma-miR10188 MIMAT0040953

UCUCUGAUUUUGCAGAUAAGGACU

>gma-miR5675 MIMAT0022460

UAGAGACGACAACAAUGGAAA

>gma-miR10427a MIMAT0041684

AAGAGAAUUGGGCUAGAGCUGC

>gma-miR10193a MIMAT0040960

UCUGAAACCGAUGUUAACUACU

>gma-miR9754 MIMAT0036385

UGACCACAUUAGACUGAAAGUA

>gma-miR4408 MIMAT0018327

UAACAACAUUGGAUGAGGGUUGGA

>gma-miR10443 MIMAT0041721

UCGACUGAGAACAACAUAAUUCAU

>gma-miR1536 MIMAT0007380

AAGCAGAGACAAAUGUGUUUA

>gma-miR4394 MIMAT0018309

AAUGGACUAAAGAGAAAGGGGCCG

>gma-miR9764 MIMAT0036401

UGGAUACACUCUUACUUUCUC

>gma-miR4359b MIMAT0018253

AACGCGUGAUAUGUUAACAUCGGU

>gma-miR1520k MIMAT0018291

AAUCAGAACAUGACACAUGACAGU

>gma-miR9741 MIMAT0036361

AAUGUGUUGAACUGAGUGAAGACA

>gma-miR4343b MIMAT0018306

UCUUACAGAUCAAGUUGAUUCGGA

>gma-miR4348d MIMAT0041640

UUUCGGUGUCGGUGAAUUGCC

>gma-miR10410a MIMAT0041647

UUUCCACAUCGUUGCUGACAUAAG

>gma-miR10195 MIMAT0040962

UAUGAUUUUGUGGAUCAAAGGA

>gma-miR5775 MIMAT0023179

AUAAGCUCUUUUGAGAGCUUC

>gma-miR5763a MIMAT0023159

UGAACUAUACAAAGACGGUUA

>gma-miR10416 MIMAT0041658

AAGAUUUGACUGUUGGAACUGCCU

>gma-miR5776 MIMAT0023181

AACUUGGGCUGAGCUUAGGUG

>gma-miR1520g MIMAT0018245

CAAUCAGAACAUGACACGUGACAA

>gma-miR5037a MIMAT0021060

GCCUCAAAGGCUUCCACUACUG

>gma-miR156ab MIMAT0024887

AUUUAAGUGAUGGGAGCUCCG

>gma-miR4352b MIMAT0018276

UAAAAUGUAGACAUUCUAAGACGG

>gma-miR5668 MIMAT0022450

AGCAAUGGAAUUAUAGACUGC

>gma-miR172d MIMAT0011199

GGAAUCUUGAUGAUGCUGCAGCAG

>gma-miR10436 MIMAT0041707

UCUUGGCAACACAACUUUUAACAU

>gma-miR4387e MIMAT0022464

UGUUAGUGAUAAGGCGUGAUG

>gma-miR5031 MIMAT0021050

UUAAUGAUUAACAUCUAAUUU

>gma-miR4385 MIMAT0018297

AAUCGAUGUAGAAAAGUGAUUGGU

>gma-miR5782 MIMAT0023187

UAGCUGGUAGGAGAAGUUCAG

>gma-miR4348a-3p MIMAT0037389

UAUCACAAUGUCAUCAUCUUG

>gma-miR10414 MIMAT0041652

UCUGAGAAGAACCAUUGGUACC

>gma-miR1512c MIMAT0022458

UAACUGAACAUUCUUAGAGCAU

>gma-miR156r MIMAT0023212

CUGACAGAAGAUAGAGAGCAU

>gma-miR1509a MIMAT0007367

UUAAUCAAGGAAAUCACGGUCG

>gma-miR9765 MIMAT0036402

UAAUACAGAAUUCGGAGACAAC

>gma-miR1530 MIMAT0007392

UUUUCACAUAAAUUAAAAUAU

>gma-miR1507a MIMAT0007365

UCUCAUUCCAUACAUCGUCUGA

>gma-miR4364a MIMAT0018260

CGCGAGAUCGCACGGAAGAAGGUU

>gma-miR1512a-3p MIMAT0032030

GCUUUAAGAAUUUCAGUUAUG

>gma-miR10440 MIMAT0041713

UUGGGACAAUACUUUAGAUAU

>gma-miR390b-3p MIMAT0020935

UACUUGGCGCUAUCUAUCUUGA

>gma-miR10423 MIMAT0041670

AUGAAAGCACCAAAUUGAUAGGACU

>gma-miR9759 MIMAT0036394

UAUAAGCAAGUAGAAUUUAAU

>gma-miR5667-3p MIMAT0022449

AAACAGAUCUAAAUGGAUUCC

>gma-miR10200 MIMAT0040967

AGGUUUUAAAGAAAUUAAAUG

>gma-miR319n MIMAT0023207

UUUGGACCGAAGGGAGCCCCU

>gma-miR1446 MIMAT0040968

UUCUGAACUCUCUCCCUCAAU

>gma-miR10427b MIMAT0041694

CAAGAGAAUUGGGCUAGAGCUG

>gma-miR1520n MIMAT0018295

UCAAUCAGAACAUGACACGUGACA

>gma-miR4401b MIMAT0022461

UCAAAGACGUUGCUGAGGUAA

>gma-miR171d MIMAT0020988

UGAUUGAGUCGUGUCAAUAUC

>gma-miR9726 MIMAT0036342

UAUAGGCAUUAUUUUUUUCUUC

>gma-miR156g MIMAT0018339

ACAGAAGAUAGAGAGCACAG

>gma-miR5225 MIMAT0023169

CCUGUCGUAGGAGAGAUGACGC

>gma-miR2107 MIMAT0010083

CAAACCUCCGUAGCCUGUAUC

>gma-miR4993 MIMAT0021011

GAGCGGCGGCGGUGGAGGAUG

>gma-miR10408 MIMAT0041644

CGAGGACUCCUUUAAAUAGACU

>gma-miR9727 MIMAT0036345

UGAAGUUACUCUGAGCACUGAG

>gma-miR10444 MIMAT0041723

GACAUACAUGUGAGUUCGGAUGUA

>gma-miR10415 MIMAT0041656

AUCUUAGAAUGUGAGUUUGGAUCU

>gma-miR5039 MIMAT0021066

CCCUUUUUUAAUCGUUGCAUG

>gma-miR9763 MIMAT0036400

ACCCAUCCAACUCUGAAGAUA

>gma-miR4413b MIMAT0021675

UAAGAGAAUUGUAAGUCACU

>gma-miR9742 MIMAT0036362

UGUGUUGUUUGUUUUGUAGCA

>gma-miR1513a-3p MIMAT0020936

UUUAAAUGUGUAUAAGUCAUGGU

>gma-miR4359a MIMAT0018251

AACGAAGUGACUCUAACAUCGGUU

>gma-miR1520j MIMAT0018263

AAGAACGUGACACAUGACAAUCAA

>gma-miR9722 MIMAT0036337

UAAUAGAGGGAAGAAGAUGAA

>gma-miR4369 MIMAT0018270

GGAUCAAGCUGAUCCGGAAGUGGA

>gma-miR5780c MIMAT0036352

UUUAAUAUAUCAGGGACUUGGA

>gma-miR1529 MIMAT0007391

UUAAAGGAAACAAUUAAUCGUUA

>gma-miR4413a MIMAT0018338

AAGAGAAUUGUAAGUCACUG

>gma-miR396e MIMAT0018345

UUCCACAGCUUUCUUGAACUGU

>gma-miR9753 MIMAT0036384

ACAAUUUGGGACUUAGGGCUACAA

>gma-miR9746i MIMAT0036383

AAAGUGUUUGAAUUUCAAUUAGAU

>gma-miR10446 MIMAT0041726

UCAACUCAAGUCGAGCUCAAGCCU

>gma-miR10194 MIMAT0040961

UGUUGAAGGUGUGUAUGACAAG

>gma-miR4349-5p MIMAT0018238

UAUUGGCUAGAGAUAAGACAAAGA

>gma-miR5770c MIMAT0040959

UAGGACUAUGGUUUGGAUUU

>gma-miR5764b MIMAT0041701

UGAUGGAUCUUUGCUUGGUACC

>gma-miR5770b MIMAT0023189

UUAGGACUAUGGUUUGGACAA

>gma-miR10420 MIMAT0041666

CCUGAACCAUCAUUUUUUU

>gma-miR482a-5p MIMAT0017337

AGAAUUUGUGGGAAUGGGCUGA

>gma-miR5767 MIMAT0023164

UGGAGGACCUUUGAAGGUGCA

>gma-miR5035-5p MIMAT0021058

CUUCUAAACAUUUUUUCCCUUA

>gma-miR396f MIMAT0021069

AGCUUUCUUGAACUUCUUAUGCCUA

>gma-miR10433 MIMAT0041700

CUGCGGAUCAAGUUGAUUGUUACG

>gma-miR9743 MIMAT0036363

AGAGAGUGCUUUGAAGAAAAUGCC

>gma-miR5783 MIMAT0023188

GACGACGACGGGGAGGACGCGC

>gma-miR5670b MIMAT0036393

CACAUCAUACCAUAUUUGCUUC

>gma-miR6299 MIMAT0024928

AUUUAAAAUUAUUGAUUUGUCA

>gma-miR10417 MIMAT0041660

GAGGCUGUUUCCAGAUUUGAACCC

>gma-miR1518 MIMAT0007377

UGUGUUGUAAAGUGAAUAUCA

>gma-miR4412-5p MIMAT0020954

UGUUGCGGGUAUCUUUGCCUC

>gma-miR5377 MIMAT0021618

CUGAAGGAUCGAUGUAGAAUGCU

>gma-miR4998 MIMAT0021016

AGUUUCGUGACUACAACUUCUG

>gma-miR4356 MIMAT0018248

CAGGACUGUCUUAGAAAGCCAGGC

>gma-miR4416b MIMAT0023177

UGGGUGAGAGAAACGCGUAUC

>gma-miR4407 MIMAT0018326

CAGAGGAAGCAGCACUUGUACC

>gma-miR4397-5p MIMAT0020953

CAUCGUUGACGCUGACUGUACG

>gma-miR5373 MIMAT0021611

UCUCUUGAUUCUAGAUGAUGU

>gma-miR1516b MIMAT0021061

AGCUUCUCUACAGAAAAUAUA

>gma-miR10187 MIMAT0040952

CGAGGAUAAUUGUUGGAACCA

>gma-miR2108b MIMAT0010085

UUAAUGUGUUGUGUUUGUGAG

>gma-miR6300 MIMAT0024929

GUCGUUGUAGUAUAGUGG

>gma-miR2119 MIMAT0011167

UCAAAGGGAGUUGUAGGGGAA

>gma-miR1523a MIMAT0007384

AUGGGAUAAAUGUGAGCUCA

>gma-miR5778 MIMAT0023183

CGACGAACUCUUCGUCGGCAUC

>gma-miR4395 MIMAT0018310

UGGAUAGGAGUAUGGGCUUGAG

>gma-miR171m MIMAT0023200

UUGAGCCGCGUCAAUAUCUCA

>gma-miR4360 MIMAT0018252

CAGUUGACGUACGUACGGAUUGAC

>gma-miR4347 MIMAT0018236

AAGCUUCUUACGGAUCAAGUUGAU

>gma-miR5374-3p MIMAT0032125

UUCAAAUGUCAGAUUAUAAAA

>gma-miR1520b MIMAT0007387

GUGACAGUCAUCAUUUAAUAAGA

>gma-miR5377b MIMAT0041693

GACCGAUGUAGAAUGUUCAACAUU

>gma-miR4414a-5p MIMAT0018342

AGCUGCUGACUCGUUGGCUC

>gma-miR4373 MIMAT0018277

AAGUUGACGUACGUACGGAUUGAC

>gma-miR10439 MIMAT0041711

CAGACAGAUAAUGCUAGAGCC

>gma-miR4387a MIMAT0018299

AACAAGACGUGAUGACGUGACACU

>gma-miR394b-3p MIMAT0018336

AGGUGGGCAUACUGUCAACU

>gma-miR5037c MIMAT0021606

AGUGGAACUUUGAGGCCUGC

>gma-miR5030 MIMAT0021049

AGAACAAUUUGUGUUUUACCGG

>gma-miR5670a MIMAT0022452

CAUCAUACCAUAUUUGCUUCAU

>gma-miR1526 MIMAT0007388

CCGGAAGAGGAAAAUUAAGCAA

>gma-miR10197 MIMAT0040964

AUUUUAGAACGACUUUUUCCU

>gma-miR4355 MIMAT0018247

CACUGUUGUGCUGGGUGUACCA

>gma-miR1534 MIMAT0007397

UAUUUUGGGUAAAUAGUCAU

>gma-miR5036 MIMAT0021059

AGAGGCCCUUGGGGAGGAGUAA

>gma-miR169i-5p MIMAT0022983

UGAGCCGGGAUGGCUUGCCGGCA

>gma-miR4381 MIMAT0018288

UAUGUGACGGUAAACGGUGACAAG

>gma-miR1531-5p MIMAT0032033

AUAUGGACGAAGAGAUAGGUAAAU

>gma-miR5780d MIMAT0036396

UGUUUUGAGUUUCUGAUAAAUU

>gma-miR5042-5p MIMAT0021072

UAUCUUGGAUCACAGCCCCAUU

>gma-miR4361 MIMAT0018256

CCGGAAGAGACUUACGGAUCAACU

>gma-miR5673 MIMAT0022456

CGUGGAAUCUCGCGGAAGACAU

>gma-miR1516a-3p MIMAT0007375

CAAAAGAGCUUAUGGCUUGUA

>gma-miR4372a MIMAT0018275

UAAAAUCGUGACAUGUGACGGUCA

>gma-miR159a-5p MIMAT0020919

GAGCUCCUUGAAGUCCAAUUG

>gma-miR10432 MIMAT0041698

UUUUCCAGAUCAUAUCGCCGGUUC

>gma-miR1520p MIMAT0018322

AUGUUGUUAUUGGAUGAUGACGGU

>gma-miR9757 MIMAT0036390

CAACCCUCCUCAGUUAGAUCUC

>gma-miR1514a-5p MIMAT0007372

UUCAUUUUUAAAAUAGGCAUU

>gma-miR5769 MIMAT0023166

UGAGGGAAAUGAAGACGACGA

>gma-miR4343a MIMAT0018231

AAAAAACUUACGGAUCAAGUUGAU

>gma-miR4416c-5p MIMAT0023172

CUGGGUGAGAGAAACACGUAU

>gma-miR4340 MIMAT0018228

UGCAGAGAUAGGGACGCGCUUA

>gma-miR10196 MIMAT0040963

UGAUUGUGGGAGAGCAUUUCAU

>gma-miR4406 MIMAT0018325

AUUGAUUCUGAGAGAACCGGUGUA

>gma-miR1507c-3p MIMAT0021006

CCUCAUUCCAAACAUCAUCU

>gma-miR10434 MIMAT0041702

CUACUGAUUAUAUAUCUGAUACCAG

>gma-miR4371b MIMAT0018274

AAGUGAUGACGUGGUAGACGGAGU

>gma-miR1517 MIMAT0007376

AGUCUUGGUCAAUGUCGUUCGAAA

>gma-miR169n-3p MIMAT0032126

UGCCGGCAAGUUUCUCUUGGC

>gma-miR5041-5p MIMAT0021070

UUUCAUCUUCAACUUGCUCAA

>gma-miR4380a MIMAT0018261

CGGAUUGUUGAUCCGUAUGUGCAU

>gma-miR1510b-5p MIMAT0020939

AGGGAUAGGUAAAACAACUACU

>gma-miR5374-5p MIMAT0021615

UUAUAGUCUGACAUCUGGAAU

>gma-miR4367 MIMAT0018266

CUGAACCCUAGCGAAGUAAAUC

>gma-miR4374b MIMAT0018282

UACUUUCAAAGACGUUGUUGAG

>gma-miR4415a-5p MIMAT0018343

AAGUUGUGAUGAGAAUCAAUG

>gma-miR5766 MIMAT0023163

UUGAGGCUGAGAAGAGGCAAG

>gma-miR9731 MIMAT0036350

AUACAUAUCGUGUUGCCAAGC

>gma-miR169s-3p MIMAT0037454

CGGCAAGUAAUCUUUGGCUGC

>gma-miR10428 MIMAT0041690

UGAGGACAAAACACGUUGAAAAGU

>gma-miR9725 MIMAT0036340

UUAAUUUUUUUGGAUCAGCAU

>gma-miR5372 MIMAT0021610

UUGUUCGAUAAAACUGUUGUG

>gma-miR1510a-3p MIMAT0007368

UUGUUGUUUUACCUAUUCCACCC

>gma-miR171o-5p MIMAT0032131

AGAUAUUGGUACGGUUCAAUC

>gma-miR5777 MIMAT0023182

CUAGCAAUAAUGUUGGAUGCAC

>gma-miR1535b MIMAT0021009

CUUGUUUGUGGUGAUGUCUAG

>gma-miR10429 MIMAT0041692

UGAGGACAAAACAUGUUGAAAAGU

>gma-miR156f MIMAT0018318

UUGACAGAAGAGAGAGAGCACA

>gma-miR172k MIMAT0023201

UGAAUCUUGAUGAUGCUGCAU

>gma-miR394a-3p MIMAT0018341

AGCUCUGUUGGCUACACUUU

>gma-miR4405 MIMAT0018324

AUUCUAAGACGGUUAUCUGGGACC

>gma-miR1531-3p MIMAT0007393

UCGUCCAUAUGGGAAGACUUGUC

>gma-miR4384 MIMAT0018294

AAUCAGACACUGCAUUCAAAGACG

>gma-miR5780a MIMAT0023185

AUCACUUAGCUGACGGUAGGGAC

>gma-miR2109-3p MIMAT0032101

GGAGGCGUAGAUACUCACACC

>gma-miR10421 MIMAT0041667

CAGAAUGAGACCUUGAGCGUGG

>gma-miR1514b-5p MIMAT0007373

UUCAUUUUUAAAAUAGACAUU

>gma-miR1512b MIMAT0021673

UAACUGGAAAUUCUUAAAGCAU

>gma-miR4387c MIMAT0018254

AGCGUGAUGACGUGACACUCCGUC

>gma-miR1520e MIMAT0018243

CAAUAAGAACGUGACAUAUGACAG

>gma-miR9744 MIMAT0036364

AAUGGAUAUGAGCUGCAUACA

>gma-miR4386 MIMAT0018298

UCGAAGGUUCUGGAGAGGACUGCA

>gma-miR5671b MIMAT0036348

UACCCGAAUUUGCUUCCAUGAU

>gma-miR4992 MIMAT0021010

AUUCUAAGAUGGUUUUUGUUAG

>gma-miR169d MIMAT0018330

UGAGCCAAGGAUGACUUGCCGGU

>gma-miR482c-5p MIMAT0021001

AUUUGUGGGAAUGGGCUGAUUGG

>gma-miR396d MIMAT0018262

AAGAAAGCUGUGGGAGAAUAUGGC

>gso-miR2218 MIMAT0016361

UUGCCGAUUCCACCCAUUCCUA

>gso-miR2109 MIMAT0016363

UGCGAGUGUCUUCGCCUCUGA

>gso-miR3522b MIMAT0016360

UGAGACCAAAUGAGCAGCUGAC

>lja-miR7541 MIMAT0029368

UGCAUUCUCUUUUGGUGGCCC

UCAGAAGGACUAAAAUCAAACUCG

>lja-miR11068a-3p MIMAT0043704

ACUUUACCAUUGGAGGUGC

>lja-miR11079-3p MIMAT0043725

GUGGUGUCCAAUAUCUCCUUGG

>lja-miR11140-3p MIMAT0043914

UUUCCGAGCUUAACGCAUAAUUGC

>lja-miR11073-5p MIMAT0043712

>lja-miR11103b-5p MIMAT0043853

UUCUCCAGUGGACAUUUGUGG

>lja-miR11088a-3p MIMAT0043770

UCUCAACCCAACCCAUAUUUUGGC

>lja-miR11133-5p MIMAT0043896

UUCCUUAAGGCUAACAGCUUGG

>lja-miR11102-3p MIMAT0043799

CUAUAUGAAUUUGAUGAAGGC

>lja-miR11077b-5p MIMAT0043798

CGAUGGAUGAAUUAAGACUAC

>lja-miR11114-3p MIMAT0043837

UGGACAGAUAAAUAGGAACGG

>lja-miR5287b-3p MIMAT0043938

UUAUAUAUGUGAUCGGAGGGUGU

>lja-miR11138-3p MIMAT0043910

AAAGCUUAAGCUGUUGGGUAAGG

CGGAGGAUUAGGUAAAACAAC

>mtr-miR2680a MIMAT0013510

UCCUCGGUACCUAUGUUGAU

>mtr-miR2655l MIMAT0013451

CGUUUAGGUCCCUUAACUUUA

>mtr-miR5299 MIMAT0021373

UUCAUUGGUAUUGUAAAGCGACAU

>mtr-miR2605 MIMAT0013309

ACUUAGUUUAUAUGACCUAC

>mtr-miR166d MIMAT0011068

UCGGGCCAGGCUUCAUCCCCC

>mtr-miR399k MIMAT0011078

UGCCAAAGAAGAUUUGCCCUG

>mtr-miR2656b MIMAT0013456

AAGUUGCAUAAUCGAGUUGG

>mtr-miR5291c MIMAT0021356

GUUUGAUGGAUGGAUUGGAUGGAU

>mtr-miR5215 MIMAT0021150

AGGAGGAUGAGCUACCUGCUU

>mtr-miR5232 MIMAT0021217

UACAUGUCGCUCUCACCUGAA

>mtr-miR2636 MIMAT0013402

UUUGGUUAGUGUGCUGAAUAU

>mtr-miR2602b MIMAT0013306

UGGCAGUGAUUGCCACGUCAU

>mtr-miR1507-3p MIMAT0010031

CCUCGUUCCAUACAUCAUCUAG

>mtr-miR2617a MIMAT0013334

UGUAGUGUAGCAUGCCCGUU

>mtr-miR2647c MIMAT0013419

AUUCACGGGGACGAACCUCCU

>mtr-miR5212-5p MIMAT0021145

UGGAUUUCGUAUUUCUUUGGUA

>mtr-miR2669b MIMAT0029977

AAAGUUCAGUCUUCAUAGUAUC

>mtr-miR5263 MIMAT0021263

UGACUAAAACUAGUAACGGGG

>mtr-miR2644 MIMAT0013412

CACUUCAGAUUGAUGGUGUGU

>mtr-miR2646b MIMAT0013416

CAUGACAUUUAGUGAUGAUGU

>mtr-miR2614 MIMAT0013328

CGGUUCGACUCGUUAGGUUC

>mtr-miR5225c MIMAT0023143

UCGCAGGAGAGAUGACACCUUC

>mtr-miR5283 MIMAT0021327

CGUGCGUAUCGGGAUGUAUCGGAA

>mtr-miR2590e MIMAT0013265

AUCUAAAGGUGAUUAUUGUGCC

>mtr-miR395h MIMAT0003861

AUGAAGUGUUUGGGGGAACUU

>mtr-miR5298c MIMAT0021371

UGAUGGAGAUGAUAUGAAGAUGAA

>mtr-miR2679a MIMAT0013507

CUUUUCACUUUCGAACGGGUG

>mtr-miR2592bm-3p MIMAT0022214

GGAAAACAUGAAUGUCGGGUG

>mtr-miR5295c MIMAT0021365

UCGGCUCUGGGAAUGAAAAGAGGC

>mtr-miR5237 MIMAT0021227

UUCAAAAGAUUUAGUUGGGAU

>mtr-miR7696a-3p MIMAT0029987

UUGAAUUAUGAGAACUUGAAG

>mtr-miR399t-5p MIMAT0030005

GGGUGAGUUCUCCAUUGGCAGGU

>mtr-miR398a-5p MIMAT0022935

GGAGUGACACUGAGAACACAAG

>mtr-miR5294a MIMAT0021360

GCUAAACGGAAUGAGGGUAGUCAU

>mtr-miR5740 MIMAT0023117

UGAACAGAAAGAACAUUUGGC

>mtr-miR5563-3p MIMAT0022232

ACUGAGUUGCCUAAUGUCGUU

>mtr-miR166e-5p MIMAT0026726

GGAAUGUUGGCUGGCUCGAGG

>mtr-miR2600a MIMAT0013303

ACAUUAGCCAAUCACAAUGCC

>mtr-miR5298d MIMAT0021372

UGGAGAUGAUAUGAAGAUGAAAAA

>mtr-miR2629h MIMAT0021310

GCAGAAGAUCCUCGGCAGUUAACU

>mtr-miR5249 MIMAT0021245

ACUUAGGGGGCAGUUUUGUAG

>mtr-miR2635 MIMAT0013401

AUUAUUGUCAACGUGACUAG

>mtr-miR5233 MIMAT0021218

GAGGAGGAUGGCCGUCUGGAC

>mtr-miR5562-5p MIMAT0022229

UGUGGAGUCUUUUGCAUGAAG

>mtr-miR7699-3p MIMAT0029999

UGUAAUCAAUGCAUUAAAUGC

>mtr-miR2588b MIMAT0013259

UAACACUGUGCAACUAAGUCC

>mtr-miR5284a MIMAT0021335

GAGGGACCAAAAGUGGAAGAAUCU

>mtr-miR2592s-5p MIMAT0026925

ACAACAGGACUCAAGCAUUUC

>mtr-miR2659g MIMAT0029970

CCAUGGGUGCGACUUGGUAAG

>mtr-miR5280 MIMAT0021319

UAAUUAGAAACGGGCCGUGAUGGG

>mtr-miR2111g-3p MIMAT0013288

AGCCUCGGAGUGCGGAUUAUC

>mtr-miR2610a MIMAT0013318

AGAUUGAGACUUGUAUGGCUU

>mtr-miR5759 MIMAT0023146

AAGGGGGUGAAAAGAUUCAAA

>mtr-miR2615a MIMAT0013329

CCUGAUCGCAUUUUAAAAGGC

>mtr-miR5271a MIMAT0021297

CGGAUAAUUGUGGUUACUAACGGU

>mtr-miR5224a MIMAT0021182

UCGAGGACAUGAGGGACGUUAU

>mtr-miR399j MIMAT0011077

CGCCAAAGAAGAUUUGCCCCG

>mtr-miR2670c MIMAT0013480

CAAGAAGGUUGCUCACUAUUU

>mtr-miR5206a MIMAT0021137

AUGGGAUCCUGUUGGUGGGUUAC

>mtr-miR2645 MIMAT0013414

UUUCUAGAGAUGAGCAUAUAU

>mtr-miR171c MIMAT0011083

UGAUUGAGCCGUGCCAAUAUU

>mtr-miR5560-5p MIMAT0022223

CUCAUUCACUCAGCCGGUACA

>mtr-miR408-5p MIMAT0022237

ACAGGGAACAUGCAGAGCAUG

>mtr-miR5221 MIMAT0021178

AGGAGAGAUGGUGUUUUGACUU

>mtr-miR5741a MIMAT0023118

UAGGGACUAAAUUGAUGGUUU

>mtr-miR2088-3p MIMAT0010039

UCCAAUGUAAUCUAGGUCUA

>mtr-miR5284h MIMAT0021342

GAGGGAUCAAAAGUGGAGGAAUCU

>mtr-miR2676f MIMAT0013503

CAUUGUUUGGAUAAUAAUUUG

>mtr-miR2634 MIMAT0013400

UUUAUUCUCAGUUUGUUGCUC

>mtr-miR5556-3p MIMAT0022212

UGAUGACGGAAGAAAUCCAAA

>mtr-miR5250 MIMAT0021246

UGAGAAUGUUAGAUACGGAAC

>mtr-miR5297 MIMAT0021368

AUCGGGAAGUAUCGGAUAAUUAUU

>mtr-miR2671j MIMAT0013482

UUAAAAGUUUCGUUUCGGUCC

>mtr-miR5254 MIMAT0021252

AGGAGGUGGAAGCAUUUGUGA

>mtr-miR5269b MIMAT0021294

AAAGUGGUGGGACAUACAUUGAUU

>mtr-miR5745a MIMAT0023125

CGUGACAACACAUCAUUGGAUGCA

>mtr-miR5270b MIMAT0021296

GAGGAGGAGUAGUUUUAGGUCAUU

>mtr-miR5286a MIMAT0021346

CAGGACAAACUGGAGGCAAGGGAC

>mtr-miR2643a MIMAT0013411

UUUGGGAUCAGAAAUUAGAGA

>mtr-miR5747 MIMAT0023127

AAAAGAAUACUCAUACAUAACAUU

>mtr-miR5222 MIMAT0021179

UUACAGGAGAAGAAUGUAUGGC

>mtr-miR2590j MIMAT0021334

AGAAUGACAUGGCAGAAUAAUCAC

>mtr-miR5559-3p MIMAT0022222

UCUAAUUAUUCACCAAGUAAA

>mtr-miR482-3p MIMAT0027102

CUUACCUACACCUCCCAUGCC

>mtr-miR2599 MIMAT0013302

UGGGUACAAGGAAUCUACUUU

>mtr-miR2670g MIMAT0021210

AGUGGUCUGUUAGGUUGGGGA

>mtr-miR395b MIMAT0001649

AUGAAGUAUUUGGGGGAACUC

>mtr-miR7696c-3p MIMAT0029991

UUUUGAAUUAUGAGAACUUGA

>mtr-miR2592bl-5p MIMAT0022205

UGGCAAGUUUGAAUUUACCUCA

>mtr-miR2607 MIMAT0013313

AUGUGAUUAUGUGAUAAGUGU

>mtr-miR2592bm-5p MIMAT0022213

CUCGGCAUUCAUGUUUUUCCUU

>mtr-miR5288 MIMAT0021349

CAGCAUUGAAGAACAUAGGGAUUA

>mtr-miR5208a MIMAT0021139

AACAUGGAUGUUGUGAGUUUGUU

>mtr-miR2613 MIMAT0013327

CGGUCGCCGGUGGUCAAUGGU

>mtr-miR5246 MIMAT0021242

UUGCAGACAGCUUUGAAGGUU

>mtr-miR169d-3p MIMAT0022933

GGCAGGUCAUCCUUCGGCUAUA

>mtr-miR5216a MIMAT0021173

UUAGGAGUGAAAAACGGUGGAA

>mtr-miR2087-3p MIMAT0010037

CUGCAGUCGGUUUCUUACUUC

>mtr-miR5563-5p MIMAT0022231

UGAUAUCAGGCAACUCGGUCC

>mtr-miR5287b MIMAT0021348

UGCUUAUAAUAGUGAUCGGAGGGU

>mtr-miR396c MIMAT0023119

AUUCAAGAAGGUCGUGGAAAA

>mtr-miR2639 MIMAT0013406

UAGUCGGCUUACGUCACCUUG

>mtr-miR2606c MIMAT0021312

AGUUAAGAACCAUACAAAAAACAC

>mtr-miR168a MIMAT0011089

UUGCUUGGUGCUGGUCGGGAA

>mtr-miR2675 MIMAT0013496

CGAGGCAUAUUUGCAGGGAUU

>mtr-miR5743a MIMAT0023121

UGAGAACUGUUUUCCGCACCUU

>mtr-miR2199 MIMAT0011319

UGAUACACUAGCACGGAUCAC

>mtr-miR5756 MIMAT0023140

CCGACCGGAUUCUCAGACGGG

>mtr-miR2625 MIMAT0013346

CCAUCGUGCCACGUUACGAUCC

>mtr-miR2592bp-3p MIMAT0030010

CGGCAGAACUCCAGCAUUCGA

>mtr-miR5256 MIMAT0021254

UAAUGGAUUAUGUAAGAUUAA

>mtr-miR5230 MIMAT0021215

CAAAUCUUGAAUCGAUUGGCA

>mtr-miR5269a MIMAT0021293

AAAGUGGUGGAACAUACAUUGAUU

>mtr-miR2587e MIMAT0013256

UUGACCGUUCAUAUGAACCCUG

>mtr-miR5244 MIMAT0021240

UAUCUCAUGAAGAUUGUUGGU

>mtr-miR169i MIMAT0029963

UGAGCCAAAGAUGACUUGCCGG

>mtr-miR5225a MIMAT0021183

UCAGUCGCAGGAGAGAUGACAC

>mtr-miR156j MIMAT0021272

UGACAGAAGAGGGUGAGCAC

>mtr-miR2620 MIMAT0013340

UUCUGAUAGACACCGGCUCUGC

>mtr-miR2601 MIMAT0013304

UAUUUGGUAUCGCUUUGGUCCC

>mtr-miR2642 MIMAT0013410

AUGAGUUUCAUCAAAUCAUGU

>mtr-miR5286b MIMAT0021352

ACAAACUGGAGGCAAGGGACAGGA

>mtr-miR5213-3p MIMAT0021147

CAGAGUGCAGAUACACGCAUC

>mtr-miR5296 MIMAT0021367

AUUUUGUGUGGGUGUAAGAGGUGU

>mtr-miR5757 MIMAT0023141

UAGAGAUUUGUUUAACAGCCA

>mtr-miR5214-3p MIMAT0021148

UGAUAGAGCUAGACCAUCGGAG

>mtr-miR7699-5p MIMAT0029998

AUUUAAUGCAUUGAUUACACA

>mtr-miR7700-5p MIMAT0030000

GUGGAGUGUGGGACAGCUUGC

>mtr-miR2670e MIMAT0021208

UCUCAACAGGACGGAUCACUA

>mtr-miR171a MIMAT0001655

UGAUUGAGUCGUGCCAAUAUC

>mtr-miR5255 MIMAT0021253

UGACUUGAUAGAGGACAUGGG

>mtr-miR5260 MIMAT0021260

UUUGUAUUGUUGACAUGGCUU

>mtr-miR5206b MIMAT0023152

GUGGGAUCCGUUGAUGGGUUAC

>mtr-miR5284b MIMAT0021336

GAGGGAUCAAAAGUGGAAGAAUCU

>mtr-miR5272f MIMAT0021307

GAAUUGAUUAUGUUUGGAUACACU

>mtr-miR5287a MIMAT0021347

UGCUUAUAUUAGUGACCGGAGGAU

>mtr-miR5247 MIMAT0021243

GCAGGAGCAAGCAUCUGAUGA

>mtr-miR5293 MIMAT0021359

GAUGAAGAAGUGGAAGGAAGAAGA

>mtr-miR169f MIMAT0011069

AAGCCAAGGAUGACUUGCCUA

>mtr-miR2660 MIMAT0013466

UAAGACAUCAGCUAUAAGCUA

>mtr-miR5231 MIMAT0021216

UUAUGCAAGUAGAUAAGCUCA

>mtr-miR2608 MIMAT0013314

GUUGUACAUAUAUCACUACUCU

>mtr-miR7698-5p MIMAT0029996

UUUUCAUCAAAGUUUUCUGGA

>mtr-miR2626 MIMAT0013347

AACGUCGGGAUUUAGGGUGUU

>mtr-miR2633 MIMAT0013399

UGACAUUUUGCUCCAGAUUCA

>mtr-miR397-3p MIMAT0029985

UCUACGCUACACUCAAUUAUG

>mtr-miR7697-5p MIMAT0029994

UCCGAGAUAUUUGAUCGGAGG

>mtr-miR5290 MIMAT0021353

AAUUUGGAGAGAGAUAGACACAUA

>mtr-miR5275 MIMAT0021313

AGCUGGAGUCACAUGCUUGAAUUU

>mtr-miR5240 MIMAT0021234

UUGAAAAAAUUGUGGAUUUGA

>mtr-miR5562-3p MIMAT0022230

AUGUGGAGAAGGCUGCAAC

>mtr-miR5554a-5p MIMAT0022199

UGUGCAUCUUGAACAAUGGUAU

>mtr-miR2631 MIMAT0013395

UGACACGCCACGUGGCACACU

>mtr-miR2592bl-3p MIMAT0022206

GAGUAAUUCAAACUUGUUAAA

>mtr-miR169l-3p MIMAT0030002

GGCAAGUUUUUCCUUGGCUAUA

>mtr-miR7698-3p MIMAT0029997

CAGACAACUUUGAUGAAAAGC

>mtr-miR5274b-5p MIMAT0022209

AUAUGACGGAGUGUAAAUGCC

>mtr-miR7701-3p MIMAT0030016

UUAGGUUCAUUCAAUUAAUGA

>mtr-miR160f MIMAT0021268

GCGUGAAGGGAGUCAAGCAGG

>mtr-miR2592bo-5p MIMAT0030007

CGGCCAGGACUCAAGCAUUUCG

>mtr-miR172d-5p MIMAT0029978

AGUGGAGCAUCAUCAAGAUUCACA

>mtr-miR319b-5p MIMAT0026727

GAGCUUUCUUUAGUCCACUCA

>mtr-miR2592br-5p MIMAT0030013

GACUAGGACUCAAGUAUUUCG

>mtr-miR5210 MIMAT0021143

UAAAUGUGUUGGAAUUAAGGUU

>mtr-miR2627 MIMAT0013348

UUUCGGUAGUUAACUGCUGAGG

>mtr-miR5561-5p MIMAT0022227

CAUUUGGAGAGACAUAGACAA

>mtr-miR2597 MIMAT0013287

UUUGGUACUUCGUCGAUUUGA

>mtr-miR2658 MIMAT0013459

AUGUGACCUUGUAUAUGAUC

>mtr-miR2600b MIMAT0021328

AAGCAUUGUGGCAUUGUGAUUGGU

>mtr-miR4414a-3p MIMAT0021207

AUCCAACGAUGCGGGAGCUGC

>mtr-miR2638b MIMAT0013405

AUGAUUAAUAUUUGCAGUGGC

>mtr-miR5258 MIMAT0021258

UCAAGUGACAAGGAAGAUCUU

>mtr-miR2641 MIMAT0013409

GUUUGAUCCUUUACGUUUAU

>mtr-miR2612 MIMAT0013325

UGAUAGUGUCAACUAGUACAG

>mtr-miR2640 MIMAT0013407

UUCCUUGCCGGAGCUGGACUAC

>mtr-miR5274a MIMAT0021311

CGUUCUACAAUAUGACGGAGUGUA

>mtr-miR2087-5p MIMAT0010036

GAAGUAAAGAACCGGCUGCAG

>mtr-miR5226 MIMAT0021184

UUUGUACAACUUGGAGGAUUCA

>mtr-miR319a-5p MIMAT0026556

AGAGCUUCCUUCAGUCCACUC

>mtr-miR2637 MIMAT0013403

AAAUACUUCCUCUGAUCACUG

>mtr-miR5252 MIMAT0021248

UGAGAGCUCACUGAAGUCUGC

>mtr-miR5229a MIMAT0021213

UUAGCAGGAAGAGUGACUAUG

>mtr-miR5760 MIMAT0023149

UGCUUUAAGGAUAUUUGUUAAGGA

>mtr-miR5285b MIMAT0021344

UGGGACUUUGGGUAGAAUUAGGCG

>mtr-miR2606a MIMAT0013310

UACAAUUCCUUAGGUGCUUUU

>mtr-miR2590g MIMAT0021309

AAAUGAGACUGAAAUCUAAAGGUG

>mtr-miR5277 MIMAT0021316

AGGUUGUUUCUUGAAGUGCAAGGC

>mtr-miR2619b-3p MIMAT0022204

CCAAAGAAUCAAUACAUAGGG

>mtr-miR2621 MIMAT0013341

AGCUUGGGCUAGGAAUUUGUGC

>mtr-miR2673a MIMAT0013493

CCUCUUCCUCUUCCUCUUCCAC

>mtr-miR2089-5p MIMAT0010040

UUACCUAUUCCACCAAUUCCAU

>mtr-miR2118 MIMAT0011318

UUACCGAUUCCACCCAUUCCUA

>mtr-miR2672 MIMAT0013492

UUAAUCGACCAAGUGGGUACUA

>mtr-miR156g-3p MIMAT0026728

GCUCUCUAGACUUCUGUCAUC

>mtr-miR5219 MIMAT0021177

UCAUGGAAUCUCAGCUGCUGCA

>mtr-miR2585c MIMAT0013322

CAGGAUUAGCGAUUACAGGGAC

>mtr-miR5557-3p MIMAT0022216

UGCUUCCUUAGUACUUGUUGA

>mtr-miR2665 MIMAT0013472

UGAUUUCAGGUCAAGAAUUGA

>mtr-miR2667a MIMAT0013474

UCCUUGAUCUGACGGCUACC

>mtr-miR5265 MIMAT0021274

AAGUGAUGUUGGAAUGGUUA

>mtr-miR5561-3p MIMAT0022228

GUCUAUCUCUCUCUAAAUGGA

>mtr-miR5205d MIMAT0021136

CUUAUAAUUAGGGACGGAGGUAGU

>mtr-miR5752b MIMAT0023134

CAUUGUUUGGUUUAGUACAAA

>mtr-miR5211 MIMAT0021144

UCGCAGGAGUGAUGGGACCGGC

>mtr-miR482-5p MIMAT0021232

GGCAUGGGAUAGUAGGGAAGA

>mtr-miR5266 MIMAT0021275

CUGGGGGACUGUCUGGGGCG

>mtr-miR2598 MIMAT0013301

CUAAGGGUGAUUAUUCUGCCA

>mtr-miR2611 MIMAT0013320

UAUUUGUCAGUGUUUGAUGAA

>mtr-miR5217 MIMAT0021175

AGGUCAUUUUGAACGGUCGGAU

>mtr-miR2591 MIMAT0013267

GGAACUUCUACGGUACACCUGC

>mtr-miR5748 MIMAT0023128

ACAAAGACAUUGGAAGGCUUA

>mtr-miR319d-5p MIMAT0029982

AGAGCUCUCUUCAGUCCACUC

>mtr-miR2111e-3p MIMAT0026927

AGCCUUGGGAUGCUGAUUAUC

>mtr-miR156i-3p MIMAT0026733

UGCUCACUUCUCUUUCUGUCAUC

>mtr-miR7696c-5p MIMAT0029990

AAGUUCUCAUAAUUCAAAAAG

>mtr-miR2643b-5p MIMAT0021249

UCUAAUCUCUGUUCCCAAUUA

>mtr-miR5259 MIMAT0021259

CAAGGGGUAUUGCGGAGGAUA

>mtr-miR7701-5p MIMAT0030015

AUUAAAUGAAUGAAUCUAAAA

>mtr-miR319c-5p MIMAT0029980

GGAGUUCCUUGCAGCCCAAAG

>mtr-miR2603 MIMAT0013307

UUUGGUAUUGGUCCCUGCACUU

>mtr-miR2674 MIMAT0013495

CACUCGCUUUGGAAGUCAUGG

>mtr-miR2622 MIMAT0013342

UUUGUGUGCCAUCGUGAACUUA

>mtr-miR2088-5p MIMAT0010038

AGGCCUAGAUUACAUUGGAC

>mtr-miR2650 MIMAT0013422

AACUUAAAUAUGUUUUCAGUCC

>mtr-miR5213-5p MIMAT0021146

UACGUGUGUCUUCACCUCUGAA

>mtr-miR5245 MIMAT0021241

CAUCGUAGAACACAGGCAGUA

>mtr-miR7696d-5p MIMAT0029992

AAGUUCUCAUAAUUCAAAACG

>mtr-miR2662 MIMAT0013469

GAGUAAAAAUGUGAACCGAAU

>mtr-miR5755 MIMAT0023137

CUUUAACGCGGGAAAGACACA

>mtr-miR5292a MIMAT0021357

AUUCAGAUGAUAGCAACAAAGAGC

>mtr-miR5276 MIMAT0021314

AGGGGGAGCACCUUGCUGGGGCAU

>mtr-miR5253 MIMAT0021251

GAUGAAAAUGAUUAUGUUGGA

>mtr-miR5745b MIMAT0023145

UUUAAUUUAUAUACAUCGUCA

>mtr-miR5204 MIMAT0021132

GCUGGAAGGUUUUGUAGGAAC

>mtr-miR156d-3p MIMAT0026725

UGCUCACUCAUCUUUCUGUCAAA

>mtr-miR5751 MIMAT0023131

UUGAUUUGAUCAGAUGGUUUU

>mtr-miR2119 MIMAT0011168

UCAAAGGGAGGUGUGGAGUAG

>mtr-miR2628 MIMAT0013362

CAUGAAAGAAUGAUGAGUAA

>mtr-miR2666 MIMAT0013473

CGAAAGUGAGGAUAUCAAGGA

>mtr-miR2589 MIMAT0013260

GGCAUCCACGUGUGCUUCACCG

>mtr-miR5264 MIMAT0021273

UUGAUCAAGGACUUUGCAUC

>mtr-miR5205b MIMAT0021134

CUUAUAAUUAGGGACGGAGGGAGU

>mtr-miR2648 MIMAT0013420

UAGCCAAUGGGAAUAACAGAU

>mtr-miR5218 MIMAT0021176

UGAGACUUGGUAGUAAGAUGAU

>mtr-miR395k MIMAT0003869

UUGAAGCGUUUGGGGGAACUC

>mtr-miR5208d MIMAT0021315

CAUAUUAGUCAUAUUUGUAGGCAU

>mtr-miR2619a MIMAT0013339

ACAUAGGAGGCUGUUUUGUAU

>mtr-miR5207 MIMAT0021138

CAUUAAUGUGGGUUUGGACGGUU

>mtr-miR1510a-5p MIMAT0010032

UUGUCUUACCCAUUCCUCCCA

>mtr-miR5273 MIMAT0021308

UAGGGGCUGUAGUUUGAGAAGAGG

>mtr-miR2595 MIMAT0013285

UACAUUUUCUUCUUUAUGUCU

>mtr-miR5279 MIMAT0021318

CGGAACCACUCGGAUGACUCGGUU

>mtr-miR2623 MIMAT0013343

UCGGCUGUACUGUCCUUCAUG

>mtr-miR2651 MIMAT0013423

UUUGAUUGGUAUGCCUGCAUU

>mtr-miR5234 MIMAT0021219

UUUUGUUGUGGAUGGCAGAAG

>mtr-miR5292b MIMAT0021358

GAUUCAGAUGAUAGCAACAAAGAG

>mtr-miR5268b MIMAT0021292

CCAGAGUGGAAUGAAGAUAUGGUU

>mtr-miR5235b MIMAT0021221

AUAAGGUCAAUGAUUGGCGUG

>mtr-miR5742 MIMAT0023120

CCACAUCAAUGGUCGUUGGAU

>mtr-miR5238 MIMAT0021228

UGUAGAAAAAACAAAGGGCAA

>mtr-miR2661 MIMAT0013468

UAGGUUUGAGAAAAUGGGCAG

>mtr-miR5228 MIMAT0021212

UCUGGUGUACAACUUGAUGGA

>mtr-miR5298a MIMAT0021369

UGGAUAUGAUAUGAAGAUGAAGAA

>mtr-miR5224b MIMAT0021185

CGGAAGAGGAUUGUCGAGGACA

>mtr-miR5212-3p MIMAT0027100

CCAAGGAAAUAGAUAUCCAGC

>mtr-miR2586a MIMAT0013251

CGAGGAGUGUCCGUGCUUCAU

>mtr-miR5555-3p MIMAT0022202

AAGUCGUAUUACACUCUUAGA

>mtr-miR5754 MIMAT0023136

UAUUGCACUCAUCUUCCAUGGC

>mtr-miR5560-3p MIMAT0022224

UGCCGGCUCAAUGAAUGCGGAG

>mtr-miR5558-3p MIMAT0022220

UAGAUUUAGAAUUAGAAAAGC

>mtr-miR5242 MIMAT0021238

UUGUAGAAACAAGCGAUGUCA

>mtr-miR5750 MIMAT0023130

AAGAGAGAUAGAUCAGAAUUGACA

>mtr-miR5744 MIMAT0023124

UAGGUAUUUUAAGGAGCACGUU

>mtr-miR5251 MIMAT0021247

AGUAGAUCUAGUUGGUGUCUU

>mtr-miR5281a MIMAT0021320

CUCUUGUAAAUAGGAUCGGAGGGA

>mtr-miR7697-3p MIMAT0029995

UCCGAUCAAAUAGAUCGGAUA

>mtr-miR5248 MIMAT0021244

UUUUUAGUUGGCAUGCAUUCA

>mtr-miR2624 MIMAT0013344

CGAAAGACGAGGUUGCCGGCU

>mtr-miR2616 MIMAT0013332

AUUGGGUUUGGUUCGGGCGGAU

>mtr-miR5555-5p MIMAT0022201

UAAGAGUAUAAUAUGACUUUG

>mtr-miR2089-3p MIMAT0010041

AGGAUUGGUGUAAUAGGUAAAA

>mtr-miR2592a-5p MIMAT0020945

CCCGGCAUUCAUGUUUUCCU

>mtr-miR2592s-3p MIMAT0013279

AAAUGCUUGAGUCAUGUUGUU

>mtr-miR1509a-5p MIMAT0010034

UUAAUCUAGGAAAAUACGGUG

>mtr-miR2649 MIMAT0013421

AAAGGUGCCAAUUAUGAGUGU

>mtr-miR2592a-3p MIMAT0020946

GAAAAACAUGAAUGUCGAGCG

>mtr-miR2604 MIMAT0013308

UAAUUUUUAUGUGGGAGUGUU

>mtr-miR530 MIMAT0023156

UGCAUUUGCACCUGCACUUUC

>mtr-miR2678 MIMAT0013506

UGAAAUUGUUGCGAGUGUCUU

>mtr-miR4414b MIMAT0021211

UGUGAAUGAUGCGGGAGCUAA

>mtr-miR5205a MIMAT0021133

CAUACAAUUUGGGACGGAGGGAG

>mtr-miR5278 MIMAT0021317

GAAAUUAUCUGCAGGAAAUGUGAA

>mtr-miR2655b MIMAT0013441

CGUUUUGGUCCCUUAACUUUA

>mtr-miR2111m-3p MIMAT0026928

GUCCUCGGGAUACAGAUUACU

>mtr-miR5243 MIMAT0021239

UGGGCAGAGAAAUCGUGAGGC

>mtr-miR5746 MIMAT0023126

UGGAUUCAACUCUAAGUGUCGGUU

>mtr-miR2111a-3p MIMAT0022240

AGCCUUGGAAUGCAGAUUAUC

>mtr-miR5227 MIMAT0021186

UGAAGAGAAGAAGAUUGAUGAA

>mtr-miR169j MIMAT0013321

UGAGCCAGGAUGACUUGCCGG

>mtr-miR5557-5p MIMAT0022215

AACAAGUACUAAGGAAGCACA

>mtr-miR5209 MIMAT0021142

CGAGGAGGCGGUAUUGUUUGAA

>mtr-miR5239 MIMAT0021233

UGGGAGAAAAGAUAGAAUGUG

>mtr-miR166b MIMAT0011061

UCGGACCAGGCUUCAUUCCUA

>mtr-miR167b-3p MIMAT0022234

GAUCAUGUUGGAGCUUCACC

>mtr-miR2657 MIMAT0013457

UGUUAUUUCAUCGAUUUUGUUG

>mtr-miR2596 MIMAT0013286

UCUAUUUCAUUGUUCCACACA

>mtr-miR1510b-3p MIMAT0010029

ACAUGGUCGGUAUCCCUGGAA

>mtr-miR156b-3p MIMAT0026723

UGCUCACUCUCUAUCUGUCACC

>mtr-miR2668 MIMAT0013475

UUCAUCCUUGCAAUUAGGGGUC

>mtr-miR5753 MIMAT0023135

AUUUUGAUUGGUGCCAACUAA

>mtr-miR2111l MIMAT0021255

AUCCUUGGAAUGCAGAUUAUC

>mtr-miR2600e MIMAT0021331

AAGCAUUGUGGCAUUGUGAUUGGC

>pvu-miR482-5p MIMAT0011172

GGAAUGGGCUGAUUGGGAAGCA

>pvu-miR159a.2 MIMAT0011170

CUUCCAUAUCUGGGGAGCUUC

>pvu-miR1514a MIMAT0011171

UUCAUUUUGAAAAUAGGCAUUG
