## Supplementary File 5 for "*In silico* Identification and Functional Characterization of Conserved miRNAs in Fibre Biogenesis Crop *Corchorus capsularis*"

**Supplementary File 5:** Potential 621 ESTs of jute genome for predicting putative miRNA

>lcl|Query_168562 FK826528.1 JutEST4850 Jute Plasmid library 20-45 days after sowing Corchorus capsularis cDNA similar to Leucine-rich repeat protein, mRNA sequence

GACGCATCCATGCTTGAGCTTCGACGAGCA

>lcl|Query_240391 FK826413.1 JutEST4341 Jute Plasmid library 20-45 days after sowing Corchorus capsularis cDNA similar to 3-hydroxy-3-methylglutaryl-coenzyme A reductase (HMG-CoA reductase), mRNA sequence

CCGCTAAGGCTAATGACGAATAGAGCTTATCGGGGGTGTTCGTATTGCACGGCTAGAGGTGAGATTCTTG

GACCCGTA

>lcl|Query_65408 FK826570.1 JutEST43102 Jute Plasmid library 20-45 days after sowing Corchorus capsularis cDNA similar to 60S ribosomal protein L22-2, mRNA sequence

GGGGCTAAAACATTTTGCTATCTTCAAAACGTAGAAGAGGCAGCGAAGAAGCAGTCAGGTATTGCGAAGA

TGAGTCGCGGAGCAGCAGCTGCACCCAAGGGGAAGAAGAAGGGAGCGACGTTCACCATCGACTGCGCGAA

GCCGGTGGAGGATAAGATCATGGACATCGCCTCATTGGAAAAGTTCCTCCAAGAAAGAATTAAGGTCGGT

GGTAAAGCCGGTGCTCTCGGTGATTCCGTCACTATCTCCCGCGACAAGAACAAGATCACCGTCACTTCTG

ACTCCAACTTCTCCAAGCGGTATCTTAAATACTTGACTAAGAAGTACTTGAAGAAGCACAATGTCCGGGA

TTGGCTTCGGGTGATTGCTTCCAACAAGGACCGTTCCGTTTATGAACTCCGCTACTTCAACATTGCTGAG

AATGAGGGGGGAGGAGGAAGAGTAAAGAATCAGTTCAGTATTTGCTCCAGTTACCAATCTGTTTTATTAG

TTGTGAGGACGTTGTTAATCTATTTCAGGTGTCAAATATTTGGACGTGTTTGGGTTGTTTTCTGACCCTT

CTTACTTAATGTTCAATGGAAATTTGTTGCG

>lcl|Query_195035 GR463700.1 JuteSSHPS23 JuteSSHlibMP1 Corchorus capsularis cDNA clone JuteSSHPITP8 similar to phosphatidylinositol transfer-like protein III like protein, mRNA sequence

GGCGTGGTCGCGGCCGAGGTATTAGAGCTAGGATTTTGGTGACCGTTATGGCTTTCTTAATGACACTTTA

CACCCTGTTCCGCTCATTTGCATGCCATATAACAAGGAAGCTTCCCAACACAGTGTCTGACCATTCTCAA

AATGCTACTCAACCCACCCGTGCTGCAAATCAAGAGGAAGAGTTTCACCCCCCTTCTCCAACTCCTCCTT

TTAGTCAGTCAGATCTTCTGTCTACTGTGCTCAAGAGGTTGGGTGAGCTTGAGGAGAAAGTTGATACACT

ACAAGCAAAGCCATCCGAGATGCCTTATGAGAAAGAGGAATTGTTGAATGCTGCTGTATGCCGTGTTGAT

GCCCTAGAAGCAGAGCTAATCGCTACTAAGAAGGCTCTTCATGAAGCTTTGAGGAGGCAAGAAGAACTGC

TTGCTTACATAGACAGCCAAGAGGAAGCCAAACTGCGGAAAAAGAAGTTTTGCTGGTGATATGCTTTGCC

GATGGAGAGACTACATTTGAGAGTACTGCCCGGGCGGCCGCTCGA

>lcl|Query_88522 FK826468.1 JutEST4487 Jute Plasmid library 20-45 days after sowing Corchorus capsularis cDNA similar to GMFP7 (Fragment), complete, mRNA sequence

GGGACAGGATAACAAATAAAGAGAATAGTTTCACATAACGACACAGGCGGAGGGGTTTTCGTCGCTGAAA

GCAGTGGTGTAGCGGACCTCAGTGGAGCTGGCACGTGCGAGATGGGAATAGTGAGGATCATCGGTGTTGC

GGACATAACCGGAGTCAAACAACAAAGAAAAAATACAACCTGTTCTGAAAAACAGGTTAAAGAAAGGAAA

TGGAAGAGTCATCGCTTGATGGAAGAACGGAGGAAGAC

>lcl|Query_186401 FK826450.1 JutEST4458 Jute Plasmid library 20-45 days after sowing Corchorus capsularis cDNA similar to Spermatid-specific protein T2, mRNA sequence

TATAGCTCCATGATTGAGCTTTCTCTTTGTTTTTCTCTTCCGTTTCTTCTCTCCATCTTTATAAATTTTA

CTGCTCGGATGAGTTTAAACTATTC

>lcl|Query_46677 JK714334.1 JutESTSSH3(47) Jute Plasmid Suppression Subtractive cDNA library Corchorus capsularis cDNA, mRNA sequence

GCTTCTTTGCCACCTCTCTCTTCATCATCACCATCTTTAGAAATATAAGAAATTGATTGAAAAAGCTTGA

TCGAAGTTTGGCTGTTTGGGTTTTTAAGATTGCTTC

>lcl|Query_90977 HS411995.1 JBM23 Library of differentially expressed transcripts in jute upon stress Corchorus capsularis cDNA, mRNA sequence

AACAAAATTTTTACCCTTCGCTTCCCGACAACACCGATAATCTTGCAGTTCCCGTTGATAGGAGTCATAG

GCCACGAAAAGATTCAGGAAGCTTTTTTTTTTTGCAAAATGGGGTGCTATGCAATTACTGCATACTAATC

AATCAAACAAAGGAAGGAGACAGGAGCTACATATGAATAGATGAGAATGATTAGATACATTATTTTTAGC

TTACTTGTGAGGTTCCTCAAAAATAATGAATGTAGTACCACTTTATGCATTTTATCATGCGTGCCAAGTG

CTTCTGCAATCCCTTCACCCTACAGCAAAGTAAATTCGGATTGCCCAGTGATTTTCTTGTGGTCCAGCGG

TAAAAGCTTCCTTTCAGGTACTTCTGACCGCCATCTATGACCAGTTTCTTGAATGTTGCTTCGTTCGCGT

CAGTCAGTTTGGCTACAACAAGGCTTCCATTCACTGGCTCGCGTCCAGTATCTACTAACACCATATGACC

TTCAGGGATGCTTTGACCTACAGGTGAGGTCATGGAATCACCTTCAACCTTCAGCCCAGAATCCATTGCC

TAAATAAGTTAACGTCACTGGTC

>lcl|Query_107358 FK826556.1 JutEST44130 Jute Plasmid library 20-45 days after sowing Corchorus capsularis cDNA similar to Lipid transfer protein precursor, mRNA sequence

GGATGGCTAGCGCTATTAAGATTGCATGTGTTGTGGTTGTGTGCATGGCGTTGGTGAGTGCACCAGTAGC

TCAGGGAGCCATTTCATGTGGTCAAGTCGCCAAAGCCCTGGGTCCATGCATTCCTCACTTGACTGGTAAG

GGTCCTTTGGCTCCTGGGTGCTGCAGTGGGGTGAAATCTCTCAACGGCGCCGCCAAAACCACCCCTGACC

GCCAAGCTGCTTGCAACTGCCTTAAAAGTAACGCTGGCTCCATCTCTGGCATCAACTATGGCCTCGCTGC

TGGACTCCCAGGCAAATGTGGTGTCAGCATTCCTTACCAGATCAGCCCCTCCACTGACTGCAGCAAGGTT

AAGTGAAAATGCGGAGATTGGAACGGTTCTCAATGTGAAGGATATAAAGAGAGTTTTAGTCTTTTAGATG

ATACATAATAAAATGGATGTAAAAATCCATGTGGAAGCACCCCTTCGTAGGAGAGTTATCCTATAAGTTT

TAGAGTTGTACAAGGTCGCTTTATATGCGACTATGCAAAATTTCATTTCATATATAAAAAATTTATTACA

ATTATAATACCCC

>lcl|Query_117979 JK743790.1 JutESTSSH3(N34) Jute Plasmid Suppression Subtractive cDNA library Corchorus capsularis cDNA similar to Superoxide dismutase, mRNA sequence

AAATGTATTCGTTCTACATTTTATTCACAAAGTCTCTCTGGTTCACATAAAAAGAGTTATTCTCAAGATC

ATTGCATCTCCGACTGCACATATTATTCTGATCACTGGGATAAGGACTTCAGCCTTGCTGACCAATGATA

CCGCAAGCTACTCTGCCACCGGCATTTCCTGTGCTTTTGCTAAGTTCATGGCCACCCTTGCCAAGGTCAT

CAGGATCTCCATGGACAACAACAGCTCTTCCAATAATAGAGTTTGGCCCAGTGAGAGGAATCTGACTGTC

AGTTTTACAGAAACTTGCAGTGCCATCATCACCAACAGTGATATTTCCTAGATCACCAGCATGGCGATTC

TCATCTTCTGGGGCACCATGCTCTTTCCCAGCAGGATTAAAGTGGGGTCCTGTTGACATGCAACCATTTG

TAGTGTCCCCAAGGGCATGAACATGGAAGCCATGGAGTCCAGGCTTAAGACCAGAAACATTCCCGGTCAC

AGTTG

>lcl|Query_85197 GH985225.1 JutEST5227 Jute Plasmid library 20-45 days after sowing Corchorus capsularis cDNA similar to Malate dehydrogenase, mRNA sequence

TGACGAAGACGAGCATGCCACCGGCGGCGCCGGAAGGCA

>lcl|Query_200154 EE332656.1 EST00152 Jute Plasmid library Corchorus capsularis cDNA, mRNA sequence

TGGAGGGGTCGGCCGGTCCGCTTTACGGTGTGCACCGGTCTACTCTTCTCTTCTGCCGGCGATGCGCTCC

TGGCCTTAATTGGCCGGGTCGCTCCTCCGGCTCTGTTACTTTGAAGAAATTAGAGTGCTCAAAGCAGGCC

TACGCTTGTATACATTAGCATGGGATAACATCATAGGATTTCGGTCCTATTGTGTTGGCCTTCGGGATCG

GAGTAATGATTAACAGGGACAGTCGGGGGCATTCGTATTTCATAGTCAGAGGTGAAATTCTTGGATTTAT

GAAAGACGAACAACTGCGAAAGCATTTGCCAAGGATGA

>lcl|Query_13081 GH985262.1 JutEST5289 Jute Plasmid library 20-45 days after sowing Corchorus capsularis cDNA similar to Fructose-bisphosphate aldolase, mRNA sequence

GAAGCCTAACATGGTTACCCCTGGTTCTGATTCACCAAAGGTTGCCCCTGAGGTTATTGCCGAGTACACT

GTCCGTGCCTTGCAGAGAACCGTCCCTCCTGCAGTCCCAGCTGTTGTTTTCTTGTCCGGTGGACAAAGTG

AGGAGGAGGCCACTGTTAACTTGAATGCCATGAACAAGCTCCAGACCAAGAAGCCATGGGCACTCTCATT

CTCATTCGGCCGTGCCCTTCAACAGAGCACACTCAAGGCCTGGGGTGGAAAGGCAGAGAATGTGAAGAAA

GCCCAAGAGGCATTCCTTGTTAGGTGCAAGGCTAACTCTGAGGCCACTCTTGGTACCTACAAGGGTGATG

CTAAGCTTGGTGAGGGTGCCGCTGAAAGCCTCCATGTCAAGGACTACAAATACTAAGAAGATAGGCTTTT

GTGGTCTCTTTTTAATGTCTATTTTGCAGAGTACATTGCGGTATGAATATTTTTAGGCTTTTAGAACTTT

TTTCCTTTTTGGGTTTGTGTCGTTTCTCATGGAATTTAGAGGGTGAAAATCAGACTTAATAAGCTTGGGG

AGCCCTTAGTGGCTCGTCTGAGTTTGGACATGTTTTGTTTATTTTCAGTTTAAAAACAATATTCTCTGCT

TTTTTT

>lcl|Query_231938 FK826451.1 JutEST4476 Jute Plasmid library 20-45 days after sowing Corchorus capsularis cDNA similar to Superoxide dismutase [Cu-Zn], mRNA sequence

GGGGATTGTCATTATCTCAGTCATTTCTGTTTTATCTTGTACTCTCTACATTCATTCAAGGGGTGCACTG

AGATCACATACGAAAATGGTGAAGGCCGTTGCCGTCCTTAGCAGCAGTGAAGGTGTTAGCGGCACCATCT

TCTTCACTCAAGATGGAGACGCACCAACCACAGTTACTGGAAATGTCTCTGGCCTAAAACCCGGACTTCA

TGGCTTCCATGTCCACGCCCTTGGAG

>lcl|Query_160959 FK826448.1 JutEST4452 Jute Plasmid library 20-45 days after sowing Corchorus capsularis cDNA similar to Dynein light chain, mRNA sequence

TTTCCCAAAAATGAGCACTGAAGAAGCAAAGAGAAGCCTCAACGGAGCTCTGACGGTGAAACCCAATTCA

GATGACCGGAAGCCTTCTATGGCGCCAATCGCGGCGTCGGCTGCTCCAACCGGCAAGAAAGTCATCATCA

AAAGCGCCGACATGAAAGATGACATGCAAAAAGAAGCTATCGATATAGCCATCTCTGCATTTGAGAAGAA

CAATGTAGAGAAAGATGTAGCAGAGCACATAAAGAAAGAATTCGATAAGAGACATGGACCCACTTGGCAT

TGCATCGTTGGTCGCAATTTTGGATCATATGTAACTCATGAGACAAACCACTTTGTTTACTTCTATTTAG

ATCAGAAAGCAGTTTTGCTCTTCAAATCTGGTTGATCATGTTGTTATGTTGATGAAGACTGCTGATCATG

AGGATTATGGAAGGCTTTTCACTTTTATTAATTATGCTACTATAAATAGCAACTGTATGTATTATCAATT

GAAAAACCTGAGTACGTATGTGGAATTCTCTCCTAAATTTCTTGTTT

>lcl|Query_187699 FK826593.1 JutEST4794 Jute Plasmid library 20-45 days after sowing Corchorus capsularis cDNA, mRNA sequence

ATCTTCCCTTTAACTAATCATTAAAAAGTGCATTTCTCACAACCACTTTCACCAAATATCTCCCCACTGC

CAACACCTCACTGCCACCCATGTCCGCCGCCTCTGCCGCCACAATTTCCACCCCTACCCTCGACTCCGAG

TCCCCTTCCCTCCTCCAATCCATCACCTCCCATGGCGGCTACGCCTATGCCCGCATGGCCACTTTAGCCG

CTTCCGGCGACCAACGCGCTGCCGAGGCGGCGCGTGAGATGGCCTGGGAGCAGCTTCATTCTGGCCCTTG

GCACTCCGTTCTCCCCGTCTGGCGTGACGCTTACTCCATGGCTTGCCTCTACGTTGCCAAGTTCCATTTC

TCCAATGGTGAGTTTAGAGATGCTCTTAGGGCTTTGGATATGGGTATCATCATGGGCGGTCCTTTGCTTA

GGAAAGACTTGGACTCGGCGATCGAAGTCGTTTCGGCCGCTAAAGCAAGAGTTGATCAGAACGGTGGAGC

TGCTGCTGGTAATGAGGGTTCTCAAAAAGCTGAGTCTCGATTGCTTGTTTCTCCGGAGGAGATTGATAAG

TCCGAGTTACTACGGCTTTTACCAATAAGGTCCTTGTCTTCTAAGATTGTGGGGAAGAGGTCTGCCCTGT

CTTTGGAGGGATTTCTA

>lcl|Query_102048 JK743780.1 JutESTSSH3(N20) Jute Plasmid Suppression Subtractive cDNA library Corchorus capsularis cDNA similar to RNA-binding protein, mRNA sequence

TCATAGTTACGAACCCAAAGCCACGACTTTGTTCACTCTCTCTATTGTAAATAACCTCAGCAATTTCCAC

AGTTCCAGCTTTGTCAAAGAGCATAGCCAAACTTTGACTGTCAACATCGTAAGGCAAATTGCCAACAAAC

AATTTAGCTTCCTCAGGAGGCTCGGGAATAGACTCCTCTTCTTCCGAATCCCCACCCTGTCCTAAAAACG

CTGCATCCTCACCTTCCTCAGCTTCCCAATCAGACAACTTCGCTTCAGCCCCATCGCTTTCTTCATTTTC

CCACGTTAATTCTCCTTCACTTTCACTTTCAGTTTCTTCCTGAACTTGTTGCTCTTCTCCATCAATGGTT

ATGGTGGTGTCATTTTCTCCCTCTTGCTGGGCCCAATCTGAAGTCTGGGCAACAAAGGCAACCAGAGAAG

GGAAATGGGTTTTTGTTCTGAGTGCCGAAAGGGTTGAAGAATGGGAAGAGGAAAGGAAAAGCTTGATAGG

TTTTGGAGGGAGTGAGAAGAGCATAAGGGAGCCCATGAGCCACCCTGTTTTCTTGAAGGT

>lcl|Query_51871 GH985254.1 JutEST5273 Jute Plasmid library 20-45 days after sowing Corchorus capsularis cDNA similar to Unknown, mRNA sequence

AGACGTCGGCGGGGGCCCCGGGAAGAGTCATCTTTTCTGTTTAACGGCCTGCCA

>lcl|Query_179160 FK826617.1 JutEST4018 Jute Plasmid library 20-45 days after sowing Corchorus capsularis cDNA similar to Solanesyl diphosphate synthase 2, mRNA sequence

GATCATTGAGTCTGAGTTTTGTGAGCCTGGATCGCTTGAAGAAGCGATTGAAATAGTTAGAAATCGCGGT

GGGATCAAGAAAGCTCAAGAATTAGCAAAGGAGAAAGCTGAACTTGCGTTAAAGAATTTGAATTGTCTTC

CTAGGAGTGGTTTCAGATCGGCTCTTGAGGATATGGTAATGTTTAATCTTGAAAGGATTGATTAGTTTCA

AAGGGTATTTTCAAGAGTCTACAATGTCTCCCTCAGAGCATGTTTCGGTTAGCTCTCGAAGACATGGTGA

TGTTTAATCTTGAAAGGATCGATTAGTTAGACAACATTTTCACAGTACAAGCAAATTCCTTGAAATGGTG

GACCATTATTCTTAAATCTTAAACATACCCAAAAGCATAAACAGAGGATGCATGAAGTTGAAGCCTCTTA

CTCATATAGAAAATCTTTTTCTTAGCACAATAGTTGGTTAGTCCAGCTGAATCCAGGTTCCTTGTGACTG

GTTTTTTTTTTTTACACCATAGGACCACAACAATAACTCTAAAAATAAATTAAGAAAAACGGAATTTAAT

GAAAAAAAAATAAAAGCCAAAAAAGAAATTTTTGGGAGGTGGAAATTAAAAAAAAAAAAAAGAGCAAAGG

GGGCCAACCATGTGATTGATGATGAAGACTGGAATATCAATCCTCAACCGGATA

>lcl|Query_179412 EE332592.1 EST00088 Jute Plasmid library Corchorus capsularis cDNA similar to At2g39050/T7F6.22 (Arabidopsis thaliana), mRNA sequence

TGGCAAATATGCAGGCCTTCAGAGAAGCCTGCAGCAGATGCTTTGTTATTCAATTTTAACACAGTACGAT

TACACATACATTATTCATAACAAGTCCATAACATAAAGTAGTAGAAGACCATAATTCAACTCATAGTAAT

AGTAGGCCTCTCTACACTCAACATATAAAGAGAATGCATAGGGACACTACTAAGGGTATATTAGCAAGCG

CGATACAGCTTCTGGAACTCACTCGCGTGCTCCATATCACGAGCATCAATAGTGTTCAATCTTCCATCTC

TGGTTATCTCCCTTATTCCACTCCCACACCCCAACAGTAGATCCATTTTGCAATCTACCGTTATCATCTT

TGAAGGCATCCACATTCAACTTTATGTTATTAACCATTCTTAGAGCAAAGTAACCTTCACCTGTGTCCTT

GCTCTCAGTCCGTAATATGGATTCATCCAGATGGTTTGCCTTGTAAGGAGTCAATTGCAAAGGTTGAGAA

GCTCCATTGGCATGCTTAATGGCTTGACCAGGGCTTTGTTAACCAAAGAAAAGCTGGGGAAACCTGCTGA

ATCTTTCACTTGATTGCTGAATTTCTCATCCTTGTACCATTGCTGGTAGGGATCAGAGTGAGCAGCTGGA

GCAAGAACAACCTTGCCACCTCTTACAGTAAGAAGAAGATGGNGATCAGCTTTGCATAGACTTTGAAAGT

AGGTTAGAGGCNA

>lcl|Query_179149 GH985248.1 JutEST5256 Jute Plasmid library 20-45 days after sowing Corchorus capsularis cDNA similar to Fiber protein Fb19, mRNA sequence

CCACCTGACATCCATAGAAGTAGCCTGAATGCACAATAATGCCAAATTTCCTATTCGTGTAACCCAC

>lcl|Query_179408 EE332603.1 EST00099 Jute Plasmid library Corchorus capsularis cDNA similar to Hypothetical protein c17, mRNA sequence

TTAAACCTACGAAAGAATATAGGATGATACATCAGCATATTTTCATCATGTGGCCTTAGGATTT

>lcl|Query_39412 FK826462.1 JutEST44122 Jute Plasmid library 20-45 days after sowing Corchorus capsularis cDNA similar to Putative calmodulin-binding protein (Fragment), mRNA sequence

GGGGCCAGGTAATAACCTTAGCGCCCTTTTTATT

>lcl|Query_232132 GH985239.1 JutEST5242 Jute Plasmid library 20-45 days after sowing Corchorus capsularis cDNA similar to T-complex protein 1, alpha subunit, mRNA sequence

GCAGGCGGAGGTGCTGTTGAAGCTGCTTTGTCTGTGTACTTGGAGTATCTAGCCACGACTCTAGGATCAC

GAGAACAGTGGGCCATTGCGGAGTTTGCGGAATCTCTATTGATTATACCAAAGGTGCTTGCGGTCAATGC

TGCCAAGGATGCCACTGAATTAGTTGCAAAACTTCGGGCTTATCACCACACCGCTCAAACCAAGGCTGAT

AAGAAACATTTTTCAAGCATGGGTCTAGACCTTTCCAACGGTACTGTTCGTAACAACTTAGAAGCTGGAG

TCATTGAACCCGCAATGAGCAAAGTGAAGATCATTCAGTTTGCTACTGAAGCAGCCATAACAATTCTTCG

CATTGATGATATGATCAAGCTTGTCAAAGATGAAAGTCAGAATGAAGAAAGGGATAAGGGGGGTTTGGTT

TCAACGTTAGTGATGTTATCTCTGTATCTTAATTTTGGGGAAGGGATTGGGATTCCATTTGGGTTAGATT

>lcl|Query_232392 EE332579.1 EST00075 Jute Plasmid library Corchorus capsularis cDNA similar to Transferase, transferring glycosyl groups (Arabidopsis thaliana), mRNA sequence

TACGGTGAATAAAATAGAGCAAACAAATAGCATAACAAAAAAAAAAAAAAAAAAAAAAATTTTTAAATAA

AA

>lcl|Query_177978 GR463673.1 JuteSSHRP64 JuteSSHlibMP1 Corchorus capsularis cDNA clone JuteSSHGP4 similar to glutathione peroxidase like protein, mRNA sequence

CGAGCGGCCGCCCGGGCAGGTGTTCTCCGCAAAATCTCATTCGACGAAATCTTAGTTCAGTTGCTATTTC

TGCTTCTCTTCTTTTTGCAAAGCGTTTTTCACCGAGTTCCAAGCCAACCCTTTCCAGTTTTCCGCAGATT

TCGCCAGTTTCTTTGGTTTCATCCTCGAAGGAAACAGGGTTTTCAAGGTCTTTCTTGGGTAGTTTAAGAT

TTGATCATTCCATGGCTAGCCAATCAGCCCAGACTTCAGTTCATGACTTCACCGTTAAGGATGCAAGAGG

AAAGGACGTTGATCTAAGCACTTACAAAGGCAAGGTCCTCTTGATTGTCAATGTTGCCTCACAATGTGGC

TTGACAAATTCCAACTACACTGAGCTGAGTCAATTGTATGAGAAATATAAAGATCAAGGTTTTGAGATTC

TGGCATTTCCTTGTAACCAGTTTGGAGGTCAGGAGCCAGGGAACAATGAACAGATCTTAGAATTTGCTTG

TACCTCGGCCGCGACCACGC

>lcl|Query_177970 GR463683.1 JuteSSHRP92 JuteSSHlibMP1 Corchorus capsularis cDNA clone JuteSSHSAP1 similar to Senescence-associated protein, mRNA sequence

GCGTGGTCGCGGCCGAGGTCAAGCTTTTTTTTTTTTTTTTTTTTTTTTTTTTTTTTTCCATAAATTCATA

AAATGTTTTGATTTAAAAAAAATTTAACAGATACTACAAGAATTTTCCTAGAGATCTCAAGAACTATTCA

CATGATGGAAATTTAACGGGCGCCTTTCGATCACCTAACTGGTCAAAGATTGTAGCTAATCCGTCAAAAT

CTCACCGCCAAAATCATGCTACTCCAACACGTGCAAACTTCAAAGCCCCAATCTCATTGGTCAAGTCTCT

ACTCTGCAGTCTGCACATTTGCTGCCAAGCCAATCCATAGCCATTCAACCAAAGTACCTGCCCGGGCGGC

CGCTCG

>lcl|Query_177650 EE332670.1 EST00166 Jute Plasmid library Corchorus capsularis cDNA, mRNA sequence

GCATGCTACATGCAAGTCGGACGGGAAGTGGTGTTTCCAGTGGCGGACGGGTGAGTAACGCGTAAGAACC

TGCCCTTGGGAGGGGAACAACAGCTGGAATACGGCTGCTAATACCCCGTAGGCTGAGGAGCAAAAGGAGG

AATCCGCCCGAGGAGGGGCTCGCGTCTGATTAGCTAGTTGGTGAGGCAATAGCTTACCAAGGCGATGATC

AGTAGCTGGTCCGAGAGGATGATCAGCCACACTGGGCATGAGACACGGCCCAGACTCCTACGGGAGGCAG

TGGA

>lcl|Query_177789 JK743771.1 JutESTSSH3(N7) Jute Plasmid Suppression Subtractive cDNA library Corchorus capsularis cDNA similar to putative glycine-rich protein, mRNA sequence

AAGTGCATAAGATGCTTGCGGCAGACTTCTGCTTTTGCTTCACCTGGTTCAACCAATCAACCTAATCAAA

ATGAACGGGAAACTTGGTGGATCAGCCGTTGGGCTTATGGTTGTCCTATGTGTGGCGGAACTTGGGTTCG

AGTTGTATAGATGATGCTTTTGACAGGATATATTTATTAGCTGGGCCTATTTTTATTCTCCTTTTATCCG

TTTATGTGACAAAATATACATCGTAGCTGGTTGCTCTCCGTTGCAGAATCAGTATATATTCGAAGAGTGA

CATACTGTTTTCATGTAGTGGCTGCTGCTCACAATTCCGGGCTACTTGAAGATTGAACCAGTAGAATTCT

ATTTTTCAGAGTGAGTTATCATCATTAATTGTAGAACCCATATCCTTATTATTTCATCTTGTAAATTTTG

TTGATAAATGACAGA

>lcl|Query_177392 FK826582.1 JutEST4467 Jute Plasmid library 20-45 days after sowing Corchorus capsularis cDNA similar to unknown protein, mRNA sequence

GGTGAGACAACTTTGTTAATATGCAGTAGTGACCCAGTTATTCTGGTGATATGATGACTTTGCTCATCAA

ATTCCCCTGTTTTCTTTAGCATATAGCTAGGTTTCTCTTCCTTGTTTGCCCTTCGGTTATCTCCACGCCA

ATGGATTCGTTTAGTACTCAGAAACCTAAATCAAGAAAGATGAAGAAGAGGCTTGCGAAAAAGTCTCGAG

TTGAGATAACAGGTTGATGAATGATGATGGTCCTGGCTGGTGGTTGGATGTTAGGATGTATG

>lcl|Query_177928 GR463729.1 JuteSSHPC25 JuteSSHlibMP1 Corchorus capsularis cDNA clone JuteSSHLip similar to Lipocalin-like domain (Stretch 7-91 amino acid) containing protein, mRNA sequence

CCATGGTGAAGCTGAGCAAAGAGGCCAAGCAGAGACTACAGCAGCTCTTCAAGGGGAGCCAGTTTGCCAT

TCGCTGGGGCTTTATCCCTCTTGTGATTTACCTGGGATTTAAGAGGGGTGCAGATCCCGGAATGCCTGAA

CCAACTGTTTTGAGCCTACTTTGGGGATAAAGGATTATTTGGTCTTCTGGATTTGGAGGCAATCAGCGGA

CAGCATGGAAGATGTGTGCTCTGGCTCGGATAAGAGATGGGACATCATTCAGTCACTAGTTGGATGGCAC

AAGGCTCTTCACAGACGCA

>lcl|Query_177791 JK743769.1 JutESTSSH3(N5) Jute Plasmid Suppression Subtractive cDNA library Corchorus capsularis cDNA similar to Aspartic proteinase nepenthesin-2 precursor, mRNA sequence

GGGTGAAGGTATTGAACCTCACCATTAGGGAAAACCCTGTAAAACTG

>lcl|Query_205497 GR463788.1 JuteSSH6H5S154 JuteSSHlibMP2 Corchorus capsularis cDNA clone JuteSSH6HACCOX1 similar to ACC oxidase like protein, mRNA sequence

GGATTGGCTTACAGACAAGATGAGAGGTCGTGACCATACAGTGTCTGCCACCCATGGAGATATGGACCAA

AATAC

>lcl|Query_60294 JK743789.1 JutESTSSH3(N32) Jute Plasmid Suppression Subtractive cDNA library Corchorus capsularis cDNA similar to Major latex-like protein, mRNA sequence

TCGTTCATCACATCCCCTTCGATCACACTAAAAGTGATTGAGTTCTTCGCCTCGTCTGCTGCTTCAATTA

CCTCCTTTGCCACCTTCCTCACTCCATCATGCGTGTAATGCCAGACGACAATGGAACCGACTTTGCCAAA

TTCACCCTCATGCAATTCACATCCGTGGATCTTGTCACCGCAGGTTTTGGAGACGTGGTGGGGTTTTTTA

AAGAACATATCATGGAAATCCTTGGCAGAAGCCTTGATCTCTACATCGGTCTCAAGCTTACCAGTTAGAC

CCGATGACGCCTTTGCAGCAGAGGTGAGGTGAGCATCGATGTCTCTGGAGACATCGGCGACAAACTGCAG

CAGTGTCTCAGGATGATCAACTCCCTTGTGCAGCTTTTCGTATTCCAGAATCCAGTGAACTATGCAGCCG

TTACCATCGCTCTTGGGTGTAGCTTGAATATGGATTACAAAGTTCTT

>lcl|Query_11715 EE332634.1 EST00130 Jute Plasmid library Corchorus capsularis cDNA, mRNA sequence

AAAAAAACAAAGGACAATACCTCAAAAAGAAAGGAAAAGTAAAAAGGTTTCGACAAATGCTAAGTACAAT

TTAAACAAGCCGAATCCCTACAATGTGGGGAAAACAACTCTGTTCATTTAAGCTGAGGTGAAGCTTGGGG

GCTTGTAGGCAATGAAACTGATGCATTGCACTTGACGCTTGTTGTCGAATCCAATGATTCTGATGAAGGA

CTCGGGGTATTCCTTCTTGGCCTCCTCAAGCTCCTTCAACACCTGGGATGAGTCGGTGCACCCGAACATG

GGAAGTTTCCACATGGTCCAGTAGCGTCCATCGTAGTACCCTGGGGAGCTGTGGTTCTTACGGTAGACGA

ATCCGTGCTCCAATTC

>lcl|Query_11968 HS411996.1 JBM25 Library of differentially expressed transcripts in jute upon stress Corchorus capsularis cDNA, mRNA sequence

GGCCATCTCCTTGGGAAAAATTAAGGTTTTTTTTGGGTCTG

>lcl|Query_11957 JK714296.1 JutESTSSH3(7B) Jute Plasmid Suppression Subtractive cDNA library Corchorus capsularis cDNA similar to Translation elongation factor 1 alpha, mRNA sequence

ACAGCCCAACCCTTGGAAATACCTACAGCCCCAGG

>lcl|Query_190983 FK826452.1 JutEST4479 Jute Plasmid library 20-45 days after sowing Corchorus capsularis cDNA similar to DNA binding, mRNA sequence

GGGACCATCATCAGTGCTTGAGTTGGATACTTCTGCATCTGTTTCGGTCTCAAAATCAAATGACAATGTT

TTTTATTTTATGTGGTCAAAAAAAAAACTAGAAAAATG

>lcl|Query_245382 GH985200.1 JutEST5179 Jute Plasmid library 20-45 days after sowing Corchorus capsularis cDNA similar to Actin-depolymerizing factor, mRNA sequence

GGGTAGATGGCATTCAGGTAGAGCTGCAGGCAACTGATCCAACTGAAATGGGTCTTGATGTCATTAAGAG

CCGAGCCAGCTAAATGGAAACCTGGAAATTTCCAAGAGGCGTGTGATGGAGTCTATATCAAAGCTTTCTG

CTTTTTGCTTAATATGAAATGTTAATAGGTTGTGAGGGGGGCGGGTATCGACCTGACTGAATGATAAGAT

TGGTTTGATCGTTCAAGTTTTATATTGAAGTCTTTATCTAGAACAAGTTTAGTGATTTTAAATTTATATT

TATGATTTGTTATGCGTGGAGATTAACC

>lcl|Query_216655 FK826559.1 JutEST41120 Jute Plasmid library 20-45 days after sowing Corchorus capsularis cDNA similar to T30E16.31, mRNA sequence

GCCGCGAACCACGCCTAATTCTAGATTGCATCGCGAG

>lcl|Query_207471 JK714312.1 JutESTSSH3(22A) Jute Plasmid Suppression Subtractive cDNA library Corchorus capsularis cDNA, mRNA sequence

TCCTTTCTCCTCGGTCCACAGAGACAAAATGTAGGACTAGGACTGGCGCCAATAGCTAATCACGGAAGAA

CAGGGATTGATGTAGATGTGAAATGTTAGCTTTTTTTAATTTCTTTGTTAATTTCTATGGTTTCATTTTC

CTTCATA

>lcl|Query_158683 EE332513.1 EST00009 Jute Plasmid library Corchorus capsularis cDNA similar to Ribosomal protein S28, mRNA sequence

TGAAAGAAGCGCACTCGTTGCTCGCCGCTACCTTCATCCTCACTCTCTCTAATCACAGAATGGATTCTAC

GATTAAGCATGCTGTTGTTGTGAAAGTGATGGGAAGGACTGGATCAAGAGGTCAGGTGACTCAAGTCAGA

GTCAAGTTTTTGGATGATCGGAACCGTTACATCATGAGGAACGTCAAGGGACCGGTCAGGGAGGGAGATA

TTCTCACCTTGCTTTAGTCTGAAAGAGAAGCTAGGAGGCTTCGTTGATCGGGTCGGGTCTTTTACTATAT

CTTTGTTGATGGTTTTCTGGTATTAGCTAGTGTTATTTTATCCTGAGTTTTGTTACTTAAATCTCATATC

AAGACTTGGATTTTGGGAGAGGATGGATTTACATGTTTTAAAGCTTAATTATGTTGTGGAAATGAACTAA

GTTGA

>lcl|Query_225528 EE332543.1 EST00039 Jute Plasmid library Corchorus capsularis cDNA similar to Type II keratin E1, mRNA sequence

GAGTGTAAATGCTGGGGTGCCTAGTGCGCGAAATAACTCACATTTATTGGGTTGCTGCTCACTGCCCGCT

TTCCAGTCTGAAAAACGGTGGGGCCACCTGCATTGGGGCATCGGCCGCCGCGGGAAAAGAGGCGATTTGC

GTATGGTGCGCCCTTCCGCTTCCTCGCTCACTGACTCGCTGCGCGCGGTCGCTCGGCTGCTTGCGAGCGG

GAGCAGGTCCCTCNAAGGCGGTGAANACGGTTATCCATACAATCCAGGGATTACGCGNTTACAGAACATG

NGATTTTTATGGCGGCAGAAGGCCGGGAACCGGAAAAAGGCCCCGTTGCTGGCGTTTTTCCATAGGCGCC

GCCCCCCTGACGAGCATCACCGAATCGACGCTCAANTCANAGGTGGCTTNAACCCGACTGGACTAGNAAG

ATACCANGCGTTTCCCCCTGNAAGCTCCCTCGTGCGCTCCCGTGTTCCGACTTTGCCGCTTAGCGGATAC

CTGTGCGCCNTTCTCGTTCGGCTGGCGCGTGGCTTTCGCATGACTCACGCTGGAGGTATCTCATTTCGGT

GTAAGTCGTCGCTCCAGTTGGGCCGGGTGCAAAAACCCGGGTTAATACTACCCTGCACTTATCGGTACTA

CNACTGAGTCACCGGTAAACCGACTACGCCCTGGAGCACACTGGTACGGATAAATACTAGTTGAAGCGGC

AAAAATTATGAGGGCGCTNCACGCTCCAAAAAATATTTTGN

>lcl|Query_225318 FK826484.1 JutEST4634 Jute Plasmid library 20-45 days after sowing Corchorus capsularis cDNA similar to catalytic/ protein phosphatase type 2C, mRNA sequence

TGGATGCCGATGTCATCACCATTGAGAACTCACGTTCAGATGAGAAGCTCCTGTCAGTCTTCCGTGAAGG

AGTCAA

>lcl|Query_18282 FK826408.1 JEST42124 Jute Plasmid library 20-45 days after sowing Corchorus capsularis cDNA similar to ATP synthase F0 subunit 9, mRNA sequence

AACTTGGTTATTATCCTCTTTTGGTGCAATTCTAAATTAGTAAATTAGGGATATTATAATTTT

>lcl|Query_228629 FK826430.1 JutEST43134 Jute Plasmid library 20-45 days after sowing Corchorus capsularis cDNA similar to Late embryogenesis-like protein, mRNA sequence

AACAAGGTAAAGCCAACTCACAACAGTAGGTGTGCCAATTGGAACTCCATTATTTTGAGATGATGTGGAG

ACATTAGCATAATTT

>lcl|Query_41008 FK826558.1 JutEST41107 Jute Plasmid library 20-45 days after sowing Corchorus capsularis cDNA similar to Ferredoxin, mRNA sequence

GGGGAAACAAAGGCAAATTCTCTAAACGCCTCTGATTCTCTCTCTTCCATTCCCAAAAATGGCGATTTCA

ACACTGCAAAAACTTTCCTCTCAAATCCACCGTCTCCCGTCACTATCAGCCCACATCCTCACCCGATCCT

CCGCCACCTCTGCCTCCCCCTCCGCCTCCACGTCCTCCAAGAAAGTTGCCGACAGGATCGTCAAGCTCTT

CGCCATTGACCCAGAAGGGCAGAAGCGTGAGGTAGTGGGCTTAACCGGCCAAACCTTACTTCGGGCCTTG

ACTAACAGTGGGTTGATTGACCCTGCCTCTCACCGCCTTGAGGAGATCG

>lcl|Query_151664 GR463733.1 JuteSSHRP72 JuteSSHlibMP1 Corchorus capsularis cDNA clone JuteSSHUBQC similar to Ubiquitin-conjugating enzyme like protein, mRNA sequence

CGAGCGGCCGCCCGGGCAGGTACCTCGGGAGGTTGGACCAGCTTTCGGTTATGAGGGGCTGCCATCTCCT

TTTTCAGCTGTGTCAGTATATCCTCCATGGTATACTCCCTCTGCCAATTTGCAAGTAGCCCAAACTTCTT

TGGTTCCACCACTCCAGTCTCATGGTTCACACAAGTCATGTTAACCCTTGAATGAAAACGGACACTCGGT

GGCTTCTCAGGATAGTCTTTATCACAGAACAGCTCCAGCTGATAGATTCGACCTTCATGTACCTCGGCCG

CGACCACGC

>lcl|Query_151571 JK714301.1 JutESTSSH3(12) Jute Plasmid Suppression Subtractive cDNA library Corchorus capsularis cDNA similar to Al-induced protein, mRNA sequence

GGGTGATTAATAAGGGTTCCCTCCTGGATACCCACCGAGCAGCCTCCTGGGCGCACCCTGGCCCTTA

>lcl|Query_245041 FK826553.1 JutEST45139 Jute Plasmid library 20-45 days after sowing Corchorus capsularis cDNA similar to Ubiquitin ligase SINAT2, mRNA sequence

AGCAGCACTGGAGCTGCCAAGGCCGTCGGCAAAGTTTTGCCAGCACTGAATGGCAAGCTGA

>lcl|Query_112270 JK743820.1 JutESTSSH3(N78) Jute Plasmid Suppression Subtractive cDNA library Corchorus capsularis cDNA similar to beta-tubulin, mRNA sequence

ATAGTTCTTTCTTCTGCATGCTTTACTGAAATTAACTCGACGATCTTAAAGAAATTAAAACTCAAATGCA

AAGCAAAAACTACAGCATAATGACTAATTCAACTGTTTCACATTCCAAACAAACCAGCAACCGGAAGTTG

TGGTTTTGAATGGCCAATAACCAGTCTCAAATCATTTAGTCATCATATCCTTCTGGTTCCTCTTCCTCAC

CATACTCATCTTCATCAGCTGTTGCATCTTGGTATTGCTGATACTCAGAAACCAAATCATTCATGTTGCT

CTCAGCCTCAGTAAACTCCATTTCATCCATACCTTCCCCGGTATACCAATGCAAGAAAGCCTTCCTCCTA

AACATAGCTGTAACCTGATCACTCACACGCCTGAACATCTTGTGAAAAGATGTTGAGATGCCAAGAATGA

GATGCCATCTTTGAGGACTGTAAG

>lcl|Query_112282 JK743807.1 JutESTSSH3(N52) Jute Plasmid Suppression Subtractive cDNA library Corchorus capsularis cDNA similar to Alanyl-tRNA synthetase mitochondrial precursor, mRNA sequence

TAAATACCGCCATGAGACGATAGCGAACAAGTACC

>lcl|Query_82557 JK743779.1 JutESTSSH3(N18B) Jute Plasmid Suppression Subtractive cDNA library Corchorus capsularis cDNA similar to Cell wall-associated hydrolase, mRNA sequence

TGTGAAGGATATACTAATGATATCCCAATATATATATACTTTCTTTGGAAATTGTATAATAAGGGCAATG

AAATCTTTTGTCCATTTTCATGTTAGT

>lcl|Query_204866 GH985158.1 JEST5114 Jute Plasmid library 20-45 days after sowing Corchorus capsularis cDNA similar to Actin 7, mRNA sequence

TTAGTGACATCAAGGATGTAAGAAGTTACTTCATTCTATCAAGTGAACAATTTTATTTTCGAAAA

>lcl|Query_149507 EE332644.1 EST00140 Jute Plasmid library Corchorus capsularis cDNA, mRNA sequence

TATAGGTCGCGCGCTTAAGCGCCATCCATTTTCGGGGCTAGTTGATTCGGCAGGTGAGTTGTTACACACT

CCTTAGCGGATTTCGACTTCCATGACCACCGTCCTGCTGTCTTAATCGACCAACACCCTTTGTGGGTTCT

AGGTTAGCGCGCAGTTGGGCACCGTAACCCGGCTTCCGGTCCATCCCGCATCGCCAGTTCTGCTTACCAA

AAATGGCCCACTTGGAGCTCTCGATTCCTTGGCACGGCTCAACAAAGCAGCCGCGCCGTCCTACCTATTT

AAAGTTTGAGAATAGGTCGAGGGCGTTGCGCCCCCGATGCCTCTAATCATTGGCTTTACCCGATAGAACT

CGCTAAGGGCTCCAGCTATCCTGAGGGAAACTTGGGAGGGAACCAGCTACTAGACGGTTCGATTAGTCTT

TCGCCCCTATACCCAAGTCAGACGAACGATTTGCACGTCAGTATCGCTGCGGGCCTCCACCAGAGTTTCC

TCTGGCTTCGCCCCGCTCAGGCATAGTTCACCATCTTTCGGGTCCCGACAGGCATGCTCTCACTCGAACC

CTTCTCAGAAGATCAAGGTCGGTCGGCGGTGCACCCGTGAGGGATCCCACCAATCAGCTTCCTTACGCCT

TACGGGTTTTCTAGCCCGTTGACTCGCACACATGTCAGACTCCTTGGTCCGTGTTACAAGACGGGCCGAA

TGGGGAGCCCACAGGCCGATGCCGGGAGCACGCA

>lcl|Query_149370 FK826453.1 JutEST4483 Jute Plasmid library 20-45 days after sowing Corchorus capsularis cDNA similar to NOI protein, mRNA sequence

GGGATTTTTTTTTCTCTCTGAACCCTCCCCTTCTCTTTCTCTCTCTTCAAACAAAAGTTTCTCTCAAGAA

TGACAGATCTTGTTCAGGGGTTGCTGGTGAGGTCTGTTCTCGAATCGTGGATGTGTTCTTTTCAAAAAAA

>lcl|Query_53538 FK826438.1 JutEST4414 Jute Plasmid library 20-45 days after sowing Corchorus capsularis cDNA similar to FH protein interacting protein FIP1, mRNA sequence

GATCAACCATGTCATCCCGCCCAGGTGATTGGAGTGGTGCTTTCCCAGTCAGCATCTCAAGGAGCAGGAC

ACCAAAGCT

>lcl|Query_213754 FK826436.1 JutEST41193 Jute Plasmid library 20-45 days after sowing Corchorus capsularis cDNA similar to Thiazole biosynthetic enzyme, mRNA sequence

TCATCATCCATGGCTGTAACTCTCACTTCCTCAACCAAACTCTACAAGAACCCTTCTGTTTTTGAGTCTT

CCTTCCATGGGGTTCCCATAACTCCACTTTCTTTCAATTTCAAAACAAAATCCTCTCCATGCAATGCATC

GATTTCCATGTCTGCTGCCTCATCATCACCTCCACCCTATGATCTGAATAATTTCAGGTTTGACCCCATC

AAGGAGTCCATCGTGTCTCGTGAGATGACACGTAGGTACATGATGGACATGATCACTTATGCTGACACTG

ATGTGGTCGTGGTTGGTGCTGGCTCCGCCGGTCTTTCTTGTGCCTATGAACTCAGCAAGAACCCTTCTGT

CCAGGTTGCCATTATTGAACAATCTGTTAGCCCTGGAGGTGGTGCCTGGCTTGGTGGACAACTTTTCTGT

GCCATGGTAGTTCGCAAACCAGCTCATCGCTTCCTTGATGAACTTGGTATTGAGTATGATGAACAAGATG

ACTACGTAGTCATCA

>lcl|Query_239773 GR463722.1 JuteSSHPS1 JuteSSHlibMP1 Corchorus capsularis cDNA clone JuteSSHSAP10 similar to senescence-associated protein like protein, mRNA sequence

ATCCACCTGAACCACCAGTTCCAAAACCATCACTGCCGCCAACCTCCACCAGCAAACATTGAAGACCTCG

GCCGCGACAAGCTA

>lcl|Query_101407 GR463667.1 JuteSSHPC10 JuteSSHlibMP1 Corchorus capsularis cDNA clone JuteSSHAT2 similar to Hsp70 (Stretch 6-103 amino acid) like protein, mRNA sequence

TCGAGCGGCCGCCCGGGCAGGTACAAGATACACCAGGAAAAGTAGTTGGGCCAATAAGATTCACCAATAT

TACTGGGGAGAAGGTGACTCTGTGGTGGGATGCCCCACTCAATGACGGTTGTGCTCCCATAACCCACTAC

ATCATTGAAAAACGGGAAACCAGCAGACTTGCCTGGGCACTAATTGAGGATAAATGTGAAGCCCAAAGTT

ACACTGCCATTAAACTAATAAACGGCAATGAATACCAATTCCGTGTTTCTGCAGTTAACAAGTTTGGTGT

TGGCAGGCCACTTGATTCTGATCCAGTGGTTGCTCAAATACAATATACTGTTCCTGATGCCCCTGGCATT

CCAGAACCTAGCAACATAACAGGCAACAGCATTACCCTGACATGGGCAAGGCCAGAATCAGATGGTGGCA

GTGAAATTCAACAGTATATCCTTGAAAGAAGAGAAAAGAAAAGCACAAGATGGGTAAAAGTGATCAGCAA

ACGACCAATCTCTGAAACAAGATTCAAAGTCACTGGTCTGACAGAAGGCAATGAGTATGAATTCCATGTC

ATGGCTGAAAATGCTGCAGGAGTTGGACCTGCAAGTGGCATCTCAAGACTCATTAAATGTAGAGAGCCCG

TCAACCCAC

>lcl|Query_100783 FK826616.1 JutEST4017 Jute Plasmid library 20-45 days after sowing Corchorus capsularis cDNA similar to Glycolate oxidase 2, mRNA sequence

TAGAATATACAATGAAGGCAGCAGCAGCCATAAGATAATCATCATACAAAAGTATTAACTCATTGTTGAA

ACATAAAATTTTCAAAAGATTTGAAGATCCAGGTCTAGTTGAACGACCAAGTAAAATCAACCCAACATGC

TATGATTTGCCAAAAGTAGATAATTAAGCATGCAGATACTAGATACTCTGAGATCATGCATTTACAACCT

TGGGGCCGGAAGACCTTTAGGAGTGTCCCAATCTGTCACAATGTGATTGCGGGTGATTTCTTTCAGTGAA

CGGCACCTACTCAGCGCCATGGTTAGCTCAAATTCCTCCCTCAGCATTGACAGTACTTTCCTCACCCCAG

CTTCTCCCTCAGCAGCCAATGAGAACACCACAGGTCGCCCAATAAATATTCAGAGGCTCCCAGGGCTAAG

A

>lcl|Query_101076 EE332667.1 EST00163 Jute Plasmid library Corchorus capsularis cDNA, mRNA sequence

CTGTTTTTGTCAATGTCTGCGTGTTAAAATTGTAGTTGGATTGCTTTTAAAAGCAATCTATAATTTACAG

AATTTACTGGA

>lcl|Query_76463 GR463695.1 JuteSSHPS29 JuteSSHlibMP1 Corchorus capsularis cDNA clone JuteSSHGP11 similar to glutathione peroxidase like protein, mRNA sequence

CTGCATGCTTGTACCTCGGCCGCGACCACGC

>lcl|Query_20746 GH985172.1 JutEST5132 Jute Plasmid library 20-45 days after sowing Corchorus capsularis cDNA similar to Inositol-1,4,5-triphosphate-5-phosphatase, mRNA sequence

TGCAAAACAGAAATTTCATTTCTATAATTTATTCATTGTTATAAGGGCGCAAA

>lcl|Query_199335 JZ532438.1 JEC11 Library of differentially expressed transcripts in jute upon elicitor and drought stress treatment Corchorus capsularis cDNA, mRNA sequence

TGTACTTTGAGAGCTATTCTACCTAGTTTGCTTTTCTTTATATTTCTATGGTGTTTTCAAAACTA

>lcl|Query_104055 EE332509.1 EST00005 Jute Plasmid library Corchorus capsularis cDNA similar to High mobility group protein 2 HMG2, mRNA sequence

TGAAGATGTGGGCAGTTGGGGATTCTACCCAATCCATGAGAGCCGTTGATGATCCAGTTCACCTGCCCAG

GCGGCCGCTCG

>lcl|Query_178907 FK826602.1 JutEST4002 Jute Plasmid library 20-45 days after sowing Corchorus capsularis cDNA similar to Hybrid proline-rich protein, mRNA sequence

ATATTGGGGGAAATTTGNCGGAGTTGATTCAAGGANAAAAGATTGGGAGGAA

>lcl|Query_185435 GR463691.1 JuteSSHPC49 JuteSSHlibMP1 Corchorus capsularis cDNA clone JuteSSHPTS similar to phenylalanyl-tRNA synthetase beta subunit like protein, mRNA sequence

GCTTTCCATAAACTTGTTGGATTATGGTAATCTATCTGTGCTACTTCCAACTTGTCTTTGACTAAATGCA

GAACATTCGATTGTAACTAATAAAGTTTAAAGCTGT

>lcl|Query_106136 HS411984.1 JBM11 Library of differentially expressed transcripts in jute upon stress Corchorus capsularis cDNA, mRNA sequence

GGGAGGAAATCGTGGGCACAACGGACGCAGGCGAACGAATCAACTTCAAGAAATCGGTCTAGTGGCGGTC

AGAAGTACCTGAAAGGAAGCTTCGACTGTGGTCAACCCGAGTGAATGCTCATCATTATTGCCTTGCTTGC

AAACGTTAATATTCCTCAGGTGCAAGTGGTCTTGTACATCTTTCTAACAATGATTTTCATTTCTTAATTA

TTATATTAGGAAAGATGGATTTTGTCTTTGAAAGTACTTTGAATATTAGAAGAAGAGTGGACACCATGAT

TTTAGAAGAAATTATGGTATGTTCATTTTAGTTTATATTCAATTACTGATTGAAGTTTGGTAAATGACAT

AATTACATTCAGAATCCTAGTAGAATGGTTTAACTTTTTATTGCCAAAAAAAAAAAGCTTCAAGGATCGG

GCCTTGATGTTACCCGAGAGCTTGGCACCCAGCCTGCGCGAGCAGGGGAATTGATCCGGTGGATGACCTT

TTGAATGACCTTTAATAGATTATATTACTAATTAATTGGGGACCCTAGAGGTCCCCTTTTTTATTTTAAA

AATTTTTTCACAAAACGGTTTACAAGCATAAAGCTTGCTCAATCAATCACCGGGTCCCCCACTCACCAGT

>lcl|Query_106234 GR463703.1 JuteSSHPS20 JuteSSHlibMP1 Corchorus capsularis cDNA clone JuteSSHECT10 similar to ECT2 [Arabidopsis thaliana] like protein, mRNA sequence

AGCGTGGTCGCGGCCGAGGTACAAACAATTCTTTGAAGCACATCACCTTAGAGAACAATGAGAATAAACC

TGTCACGAATAGCAGGGATACACAAGAGATCAAGCTTGAGCAAGGACTGAAGTTGATCAAAATTTTCAAG

GAACATTCCAGCAAAACTTGCATTTTGGATGATTTTGAGTTTTATGAGGTCCGCCAGAAGGCAATCCAGG

ATAAAAAGGCTAAACAGCAATTTCAAAAGCAGGTATGGGAAGGAAAGCCTGTAGATGAGAAGAAAGATGT

AGCAAATGGAACACAGAACTGTTTGGAAGCTGGATCAGATTTGGTTAAGGAACCAAGCTTAACTGATAAT

TCCAATGGGGATCTAAAGCTTTCAGAGAATGGTTCAGCGAATCCAATTGTAGTTCCAGACAAGAAGGTTT

TATCGAATGGGATTGCCAATGGTTGCTAGTTCCTGCCAATCCCCTTGCGACGGTTCATGGTCTTGTAGAA

TATGAAGAGGAGTTTCCAGAAATTTTGATCTGATGATTTAGAATTTTAGATCACAGTACCTGCCCGGGCG

GCCGCTCGA

>lcl|Query_106068 JK743772.1 JutESTSSH3(N9) Jute Plasmid Suppression Subtractive cDNA library Corchorus capsularis cDNA similar to Cytochrome P450-like protein, mRNA sequence

GCTCTCTCTGGAGCACATACCCTAGGAGTTGCACGATGCTCCTCCCAAGGCTAGACTTACCGTCCCAGAC

TCTT

>lcl|Query_150693 FK826519.1 JutEST484 Jute Plasmid library 20-45 days after sowing Corchorus capsularis cDNA similar to Serine/threonine kinase, mRNA sequence

GAGTCAGCGTCGACTTTGTATTCGGGTCCCAGCCTTCCATTTCG

>lcl|Query_208548 JK743803.1 JutESTSSH3(N47) Jute Plasmid Suppression Subtractive cDNA library Corchorus capsularis cDNA similar to senescence-associated protein, mRNA sequence

TTTGGTTCCTATGCTTATTACACTTATATATGTTTGAACAAAAAAAAAAAATCTACCCGCCGCTC

>lcl|Query_47678 JK743793.1 JutESTSSH3(N37) Jute Plasmid Suppression Subtractive cDNA library Corchorus capsularis cDNA similar to beta-galactosidase, mRNA sequence

TCAGAGATCTCAGCACAAACACTGGACACTGATCTTTTTACAAGTGAGATTCTCGTGGGATCACCTCCGA

GCTCCTCAAAAATAACCAACAGATTTTGTGTAGGTTTCAACCAAGACCGAGGAACATGATACCATCTTTG

GGTTGGTTGGCCGCATCCAAGCTGACACTTTGGTGGTCGGAATGTCCCAGCATAGCTGCACCCATTGCAG

TCACCATGAGCATATGCAGTCCAATATCTACCAATGCTCTGCCCATTAATCCATATTTGACCTTTTCCCA

TACTCTCCATGTCCAAAGCCAATGGCTCGTCACCCTCAGGTGCATTGAAATAAGCCTTATGCCATCTTAA

TGGCTGCTGTTTCTGTGCAGCCAATGATCCCTCTATCCACTCAACAGAGTAAATATTATTTGGTGAGACA

AGATTCATGGACTCTCCTTTAAGTCCAACCTGGTAGGTCCATTTTTGCCAGGACAGGTCCCATTTTCCTT

GGTTAAGTCCATGAAGTGCAACTGGAC

>lcl|Query_47174 GH985212.1 JutEST522 Jute Plasmid library 20-45 days after sowing Corchorus capsularis cDNA similar to Putative uncharacterized protein, mRNA sequence

AGTCTGGAAGCGGTGGTGGGTATGGTGACTACGTGAAGATGGCTGAAGGGTTTCTTGGGCAAGTACTGAA

GCTAATTAGCTTTGCTTGTTAAACCTGTGTGAGGTGTTGTTATGTACCATAATTTACAATATTTCCTCTC

TTCTATGCTCTGTATGTTGCTTATAATTCTGTACCCAATCTTCTGTCTAAGTAATTGCAGCAAATTTGGT

ACATTACTGTTAACTTTATGTATCAATTAATCCAAGATAGTCGCTAGTTGC

>lcl|Query_47812 GR463755.1 JuteSSHRP115 JuteSSHlibMP1 Corchorus capsularis cDNA clone JuteSSHNT similar to Nitrite transporter like protein, mRNA sequence

AGTAGCGGCGAGCGAAATGGGAGCAGCCTAAACCGTGAAAGCGGGGTTGTGGGAGAGCAATAAAAGA

>lcl|Query_107368 JK714307.1 JutESTSSH3(18) Jute Plasmid Suppression Subtractive cDNA library Corchorus capsularis cDNA similar to Expressed protein, mRNA sequence

ATAGAAACTGTCTACCTGAGACTGTCCCTTGGCCCGTAGGTCCTGACA

>lcl|Query_92990 GR463711.1 JuteSSHPS12 JuteSSHlibMP1 Corchorus capsularis cDNA clone JuteSSHSHMT9 similar to serine hydroxymethyltransferaselike protein, mRNA sequence

TTACTCTTCAGTTTTCTGTTGAGGTCTATCCTGTAGAGGTACCTCGGCCGCGACCACGC

>lcl|Query_75641 GR463756.1 JuteSSHPC24 JuteSSHlibMP1 Corchorus capsularis cDNA clone JuteSSHHsp901 similar to hsp90 protein family (Stretch 13-112) like protein, mRNA sequence

GCGCGTGCTGTCGCGTGACGTGGCTACTCTGGGTTTACCTTGTCGTCATGTTCCTCCTC

>lcl|Query_44230 FK826431.1 JutEST43137 Jute Plasmid library 20-45 days after sowing Corchorus capsularis cDNA similar to DNA-binding protein 4, mRNA sequence

GGATCCTAAAACTAAAAAAAAAAAAAAAACCAAAACCAAAACGGAAAAAAAATCCGGCAGGGTTGGTCGG

AAAAAGAAAACATGTCGTGGGACAAAAAAAAGAGCAATCGAAGAACTTGCTAAAGGCAAGGACTTAACAA

ATCAGCTGCGAGATCTCCTCAATGGTAGTAATTATTTGGTTGGTAGCGAAGATCTAGTGATGAAAATCTT

CAACTCTTTTTCAAATACACTTTCGATTTTAAGGAACAGTAGTAGCGGAGACTATGACGAAGTTTCTCAG

TTGAATCCGAGGAACAACAACAGTTATGATGGTCGGAAAGATCGAAGAGGATGTTACAAAAGAAGGAAGT

GTGGAGACTCATGGACAAGAGATACCCTTACTTTGTTTGATGATGGGTATGCATGGAGAAAATATGGGCA

AAAAACAATCCTCAATGCCAATCATTCAAGAAACTACTTCAGATGCACACACAAGGATGATCAAGGGTGC

CAAGCAACCAAGCAAGTGCAGCAGATTGAAGATGACCCACCAAAGTACAGGACAATCTATTATGGCCATC

ACACTTGCAAAACTCTGCTCAAAGCTTCCCAACTCATCTTGGATTCCACTACTTCCAATGACTCTT

>lcl|Query_44548 GR463791.1 JuteSSH6H5S197 JuteSSHlibMP2 Corchorus capsularis cDNA clone JuteSSH6HAPUM5 similar to APUM5 (arabidopsis pumilio 5) like protein, mRNA sequence

CGAGCGGCCGCCCGGGCAGGTTCGTCATCTAATTAGTGACGCGCATGAATGGATTAACGAGATTCCCACT

GTCCCTGTCTACTATCCAGCGAAACCACAGCCAAGGGAACGGGCTTGGCAGAATCAGCGGGGAAAGAAGA

CCCTGTTGAGCTTGACTCTAGTCCGACTTTGTGAAATGACTTGAGAGGTGTAGTATAAGTGGGAGCTTCG

GCAACGGTGAAATACCACTACTTTTAACGTTATTTTACTTATTCCGTGAATCGGAAGCGGGGCTACGCCC

CTCTTTTTAGACCCAAGACCGGCATCTGTCGGTCGATCCGGGCGGAAGACATTGTCAGGTGGGGAGTTTG

GCTGGGGCGGCACATCTGTTAAAAGATAACGCAGGTGTCCTAAGATGAGCTCAACGAGAACAGAAATCTC

GTGTGGAACAAAAGGGTAAAAGCTCGTTTGATTCTGATTTCCAGTACCTCGGCCGCGACCACGC

>lcl|Query_212559 EE332505.1 JutEST00001 Jute Plasmid library Corchorus capsularis cDNA similar to Ribulose bisphosphate carboxylase small chain, chloroplast precursor (RuBisCO small subunit), mRNA sequence

TATCCGCCACCGTCGTTAGCCGTGCCTCCCCCGCACAGGCCTCCATGGTCGCCCCATTCGTTGGCCTCAA

GTCTGCCTCTGCTTTCCCCACCACCCGCAAGAACAACAAAGACATCACCTCCGTCGTCAGCAATGGTGGC

AGAGTTCGATGCATGCAGGTATGGCCACCACTTGGGAAGAAGAAGTTCGAGACTCTCTCATACCTCCCGG

ATCTCACCCCCGTGCAATTGGCTAAGGAAGTAGACTACCTTCTCCGCTCTGGATGGATTCCTTGCTTG

>lcl|Query_212234 FK826576.1 JutEST4874 Jute Plasmid library 20-45 days after sowing Corchorus capsularis cDNA similar to 40S ribosomal protein S23, mRNA sequence

GGGGAACAACCTCTCTCAGCCAACATGGGCAAGACACGTGGTATGGGAGCTGGTCGCAAGCTGAGGACCC

ACCGCAGGAACCAGAGGTGGGCTGATAAGTCATACAAGAAATCTAACCTTGGAAATGAATGGAAGAAGCC

ATTTGCCGGTTCTTCCCACGCCAAGGGCATTGTCCTTGAGAAGATCGGTATTGAGGCTAAGCAGCCTAAC

TCTGCTATCCGTAAGTGTGCTAGAGTTCAATTGATCAAGAACGGAAAGAAGATCGCCGCTTTCGTTCCAA

ATGATGGTTGCTTAAACTTCATTGAAGAGAATGATGAGGTGTTGATTGCTGGGATTGGACGTAAGGGTCA

CGCCGTCGGAGATATTCCCGGAGTTCGATTCAAGGTCGTCAAGGTCTCTGGTGTTTCACTCTTGGCACTC

TTCAAGGAGAAGAAGGAGAAGCCTAGGTCGTAAGTTTTAGAAGAGGGATGTTCAGTTTTCTTATGGTCCT

TTTATAGTTCTACTTAAACCTGAGCTTTTTATTGCTTCTTAGCTATTATGGACAATGTTTTGTTTATTTT

CTGGTTATTGGATTGAATTTGTCATGTCATCTTGATCAAGAAAGATTTTGGTGCTTTATTGCAAAAAA

>lcl|Query_212633 JK714341.1 JutESTSSH3(27) Jute Plasmid Suppression Subtractive cDNA library Corchorus capsularis cDNA similar to Ribosomal protein L32, mRNA sequence

AAGCTTATGCTCATTCATCTTCCTGGCTGCGCAACCTGGCAAGCCTGTTAGTGACCACAACATCAAGCTG

AGCTGCTCGTTCCACAATCTCTTTTCTCTTCTTAGTGGAAACATCATGGGCAATTTCAGCACAGTAAGTC

CTGTTGTGCATCATCGACACTTCAAGCTCTTGGACATTGTGAACAACAAACTTCTTAAACCCATTGGGAA

GATAGTGGCGAGTCTTCTTGTCTGAGCCATAACCAATGTTGGGCATCAAAGTGCACCCCTTGAACTTTCT

TCTCACCCGGGAATCAATACCCTTTGGTCTACGCCAGCTTTCCTTGACGGAGATTTTGCGGTCACTCTGG

GGTCTCTTGAACTTCTTGACCCTCTTCTTCACGATCTTCTTACTTAG

>lcl|Query_212340 FK826470.1 JutEST4514 Jute Plasmid library 20-45 days after sowing Corchorus capsularis cDNA similar to Caffeic acid 3-O-methyltransferase, mRNA sequence

GGTGATTGTATCATGTTGGCTCACAACCCTGGTGGGAAAGAAAGGACAGAGAAGGAATTTGAAGCCTTGG

CAAAGGGGGCAGGTTTTCAAGGTTTCCAAGTAAAATGTTGTGCTTTTGGCACTTACATCATGGAGTTCCT

CAAAACTGTTTAAATGATCTCTGGCTAAAAACTTCATGCTTGACCATTTCTTTTGGATTCTGAGCTTTGT

GATTGTTTCAAAGGTGGCTTTTCCGGAAAATATATAGTTCTTTCTTTCTTGTCACAGGAAACAATAAGAA

TATGAAATTTCATGCTCCTTGTGTAAAATAAGTAACCTGTTTTCTATATTGATAATTGCCCTTTCTTAAT

GTAACTATTACAGAATATACATATGTTGATTGGTTTCTCTACAAGGCCTAATGCCTTAATCATGGAAGAA

GCTGATTGGTATAAATACAAATTTCATTTTCTGCAAA

>lcl|Query_212128 GH985184.1 JutEST5152 Jute Plasmid library 20-45 days after sowing Corchorus capsularis cDNA similar to MipC, mRNA sequence

GATCCAAAGCATAGAAAACCCAAACTCAAACAGCTCCTTTTGCTTTTCCTAGCAGAAAAAAACAGTCTTT

CATTATCATCCATCATCATGATCATCATGGAAGGTGTTGTGATTGAAAAATTTGCATGCCCCCTGGTAAA

ATATTGTGTGGGAATATATTTATGACCAAAAATAC

>lcl|Query_212505 EE332624.1 EST00120 Jute Plasmid library Corchorus capsularis cDNA similar to Pyruvate decarboxylase, mRNA sequence

TGTCACCGCACAAGATGTGTCAACAATGCTGCGATGTGGCCAAAAAACCATAATCTTCCTGATAAACAAT

GGTGGTTACACCATTGAAGTTGAAATTCATGATGGACCATACAATGTAATCAAGAACTGGAACTACACTG

GTTTGGTCGATGCAATTCATAATGGTGAAGGCAAGTGCTGGACAGCCAAGGTCTTTTGTGAGGAAGAACT

GATTGAAGCGATTGAGACAGCAACTGGAGCCAAGAAGGATTGCTTATGTTTCATTGAAGTGATAGTTCAC

AAGGATGATACAAGCAAAGAACTGTTGGAGTGGGGTTCTAGGGTCTCTGCTGCTAATAGCCGTCCACCAA

ATCCTCAGTAAAAAAAAGAACTCATGGCGCTGCTTGAGTGATTTCCAGCATTGGACGCTATAAAATCTTT

CTAAGTACATTGAGAATTAATCAGTCTCAGTTTAAGTTTCTTGGATGCAGAGTTAAATCTGCGATTTCAA

CATTTGCATATTCTACTGAGTGCTCGCTGATATTTGTGTAAATTTACTTTCAATGTCAATTCTAGTACTA

ATGAAATGTTTNGCCTTTACNGNCCCGAANTCNTANCATGTCATANCTGTTTCCNGNGGNAANTGTTATC

CNCTCCCANTNCCCNNACATACNANCCGNAACATAAC

>lcl|Query_212447 EE332591.1 EST00087 Jute Plasmid library Corchorus capsularis cDNA similar to BURP domain containing protein, mRNA sequence

TAACAAAATCAATGTTTTGAGACTCCATAGCATGGATTCTGATCATGCCCATCATCATCTCATGGTTGGG

GTAACTAATATTCATCCATCGTCTCATCACATCCACATGGATCCTTCATTGAGGGTGTTTTTCACCTTGA

ATGAGCTTAGAGTTGGGAAGACAATACCCATATATTTCCCTAAGAATGACCCTTCAATAGGGAACCCAAT

AACTGTCTCCTTCCCAAGAGGGGATCTTTCTGCTGCTCCGAGTTTCTTGCCCACAGAAGAAGCTGATTCC

ATTCCATTTACTCAAGAGGAACTCCCAAACATCCTTCAACGTTTCTCATTCTCTCGAAA

>lcl|Query_63546 FK826613.1 JutEST4013 Jute Plasmid library 20-45 days after sowing Corchorus capsularis cDNA similar to Fatty acid desaturase 2, mRNA sequence

TTTTTCCACTCACCGCGCTCATCCAGACCTCGGCCGCGACCACGC

>lcl|Query_69709 FK826555.1 JutEST4274 Jute Plasmid library 20-45 days after sowing Corchorus capsularis cDNA similar to Lipid transfer protein isoform 1, mRNA sequence

CTGGCTTTTCAACCAAGCGCGATGACCAATTGTGCGAATCAACGA

>lcl|Query_172841 GR463706.1 JuteSSHPS17 JuteSSHlibMP1 Corchorus capsularis cDNA clone JuteSSHEFP9 similar to exonuclease family protein like protein, mRNA sequence

GAAAGTGAGTCGGTTTTTCTAATGAAATGAACGGAAAACTCCTTTCTCTTTG

>lcl|Query_172431 FK826398.1 JutEST416 Jute Plasmid library 20-45 days after sowing Corchorus capsularis cDNA similar to Induced stolon tip protein, mRNA sequence

CTTACAGTAATTTGTATTTGCTGCGTAATTAACAATTATGTGGTGGCCCTCGTTCATGAGCATTAAAATT

GTTTATGTTGGTTTG

>lcl|Query_51828 FK826552.1 JutEST4287 Jute Plasmid library 20-45 days after sowing Corchorus capsularis cDNA similar to ubiquitin-protein ligase, mRNA sequence

TGTTTAAAATTTGTGTTTTTTGTTGGAACAATATTGGTAGTTGCTATTATCTTAAATGGTACTGAGATGC

TAAGTTTATGTTTAC

>lcl|Query_217125 FK826415.1 JutEST4355 Jute Plasmid library 20-45 days after sowing Corchorus capsularis cDNA similar to Transcription factor LIM, mRNA sequence

TTAATAACAACTCAACTGAAATGTTCCTAAACTTTGAATGTCAAAACAGTTTATAATGTTTCACCTGGAA

GTTTTGCATTCGAAAAA

>lcl|Query_217126 FK826414.1 JutEST4347 Jute Plasmid library 20-45 days after sowing Corchorus capsularis cDNA similar to Arabinogalactan protein, mRNA sequence

ATCTGTCCATTGAGGAATTGTCACTGTCGACCTCGGCCGCGACCACGC

>lcl|Query_169063 GR463710.1 JuteSSHPS13 JuteSSHlibMP1 Corchorus capsularis cDNA clone JuteSSHACCO9 similar to 1-aminocyclopropane-1-carboxylate oxidase like protein, mRNA sequence

TAAATGCTTCCATATCACAAAAGCCTAGATCTCCATGACTTTTGTGTAGAGTACTCACAAATTTATAATT

>lcl|Query_239655 EE332530.1 EST00026 Jute Plasmid library Corchorus capsularis cDNA similar to Chlorophyll A/B binding protein precursor, mRNA sequence

GTTTGGTTCCTGAATGGGGGGGAGG

>lcl|Query_51225 EE332515.1 EST00011 Jute Plasmid library Corchorus capsularis cDNA similar to Ribulose bisphosphate carboxylase small chain, chloroplast precursor (RuBisCO small subunit), mRNA sequence

GATGTAAATAATTTAGGAATAGGCGATCGCTGTTTTTTACAACTATTTAAAGTTCAAACTATTATT

>lcl|Query_144910 GR463782.1 JuteSSH6H5S183 JuteSSHlibMP2 Corchorus capsularis cDNA clone JuteSSH6ATARFA1E similar to ATARFA1E (ADP-ribosylation factor A1E) like protein, mRNA sequence

CGCATCTAACTCTAAGGATGATCCTGCCAAGGAAGCTGCCAACTTCACCTCTCAGGTTATCATCATGAAC

CACCCTGGCCAGATCGGAAATGGATACGCCCCGGTGCTCGATTGCCACAC

>lcl|Query_130741 EE332677.1 EST00173 Jute Plasmid library Corchorus capsularis cDNA, mRNA sequence

TCGTTGCAGCTTTTCCTGGGAGTATAGCATGGGTTACTTCAGCGCCGTAGCCCTGGTACCTGAACATTGG

CTCGGGGCATTTTCTCTACCCCTTCTTNCCCTGAAAAAGCAGGGGACCTTGCGTCCTTGAACCGATAACC

ATCTTTCGGCTAACCTAGCCTCCTCGNCCTCGGGACCAACAAGGGGTAGTACAGGAATATTCACCTGTTG

TCATCGACTA

>lcl|Query_39239 JK714308.1 JutESTSSH3(19A) Jute Plasmid Suppression Subtractive cDNA library Corchorus capsularis cDNA similar to 14-3-3 family protein, mRNA sequence

TATCCTTGTAAGATTCCTCACCCAATGTATCCAACTCAGAAATAGCCTCATCAAAGGCATTTTTAGCAAG

ATTGCAAGCACGGTCAGGTGAGTTGAGGATTTCATAGTAGAAGACAGAGAAGTTAAGTGCAAGTCCCAGC

CTGATTGGGTGAGTAGGAGCAAGCTCAGCAAGAGCAATATCCTGGGCAGACTTATAAGCCAACAATGTGC

TTTCAGCAGCTTCTTTCCTCTCAGCACCAGTCTTAAACTCAGCAAG

>lcl|Query_41670 GH985182.1 JutEST5149 Jute Plasmid library 20-45 days after sowing Corchorus capsularis cDNA similar to oxysterol binding protein, mRNA sequence

GAACACCTTCAATGATCATCAATATTATCTCAATTGAAACTAAAATAAATAAAGAGTCTCTCATTCTTCC

AGCTTGGCCTTTCTCTCACCAAACCATCTTGGAGGATCGAACCCAGAGCCACCAAGAGGATTGCAGCGAC

AAAGACGCCAAGTTGTGAGAACAGTGCCTTTAAGAACCCCATATTTCTTGTAAGCTTCCATTGAATACTC

ACTACAAGTCGGAACATAACGGCAGCTTTTTGGTAACAAAGGCGATATCTCTCTCTTGTAGAATCTCAGC

ATTGCTAAAGCTGCTTTGACT

>lcl|Query_166260 HS411998.1 JBM27 Library of differentially expressed transcripts in jute upon stress Corchorus capsularis cDNA, mRNA sequence

CAACCAAACCAATATTTAACCCCCCCTCCCCGACAACACGCGAAAAAACTGCAGTTCCCGTGAAGGAGTC

TAGGCCTAGAAGGATCAGGAAGCTTTTTTTTTTTGGGACAGAAGTAGTGCTATTAAAATTGGTACAGTAC

CGGAATTAGCATTAAGCATGTGATATATTTAATGGAAGGTATATAGTTCACACACTAATAAATAATTTAA

AAGTCTTAAATGCCCATGGCGAGCTCCGACCAAATGCAGTTAAACAAAAGTAGTCTTGGAAACACAAATC

AGAGGCATGCGATGCAACTCTCTGGGTTGCTTATTTATAAGCTACATATATATAATAGATACCCTTCCCC

TAATATAGCTTTCACCACATTCAAGGTACAGTCGAAGCTTCCTTTCAGGTACTTCTGACCGCCATCTATG

ACCAGTTTCTTGAATGTTGCTTCGTTCGCGTCAGTCAGTTTGGCTACAACAAGGCTTCCATTCACTGGCT

CGCGTCCAGTATCTACTAACACCATATGACCTTCAGGGATGCTTTGACCTACAGGTGAGGTCATGGAATC

ACCTTCAACCTTCAGCCAGAATCCATTGCCTAATAAGTTAACGTCACTGTCATACCATTTCATCAATGTC

CTTGATATCCGTAAGGGTTCAACAAAGCT

>lcl|Query_75326 GH985186.1 JutEST5154 Jute Plasmid library 20-45 days after sowing Corchorus capsularis cDNA similar to Endo-xyloglucan transferase, mRNA sequence

ACATGGAAAGGGTGACAGGGAACAAAGGGTTAACCTTTGGTTCGATCCATCTCCAGATTTCCATACTTAC

CAAATTAGATGGAACCATAAGGAAATTGTGTTTTTGGTTGATGATACGCCAATTAGGGTTTACAAGAACA

ATGAAGCAAAAGGTATCCCATACCCAAAACTCCAACCAATGGGAGTCTATTCAACACTTTGGGAAGCCGA

TGATTGGGCGACAAGGGGTGGTTTAGAGAAAATTGATTGGACCAAAGCTCCTTTCTTAGCTTATTACAAG

GATTTCGACATCGAAGGTTGTGCTGT

>lcl|Query_170284 JZ532435.1 JEC8 Library of differentially expressed transcripts in jute upon elicitor and drought stress treatment Corchorus capsularis cDNA, mRNA sequence

GCTTGATTGCCCCATGTTATGAGCCCAGGAAGCCCAACTGGATCTGATGTCAGATTCAAGTACCGATATT

ACCTTCCGTTGAGCTTTCAAGCCTGTGCCTAGACCGGTGCTGTTGTGCTACCGCTTCCTATCGAAACGTC

GTCTCTTCCGATGATCCACCGACTTCGTTAAGCTTGTCACTTCCTGGTGCCG

>lcl|Query_105126 EE332568.1 EST00064 Jute Plasmid library Corchorus capsularis cDNA similar to Putative senescence-associated protein (Pisum sativum), mRNA sequence

TGGTTAGTTTTACCCTACTGATGGCCGCGTCGCAATAGTAATTCACCCTANTACGAGAGGAACCGTTGAT

TCGCACAATTGGTCATCGCGCTTGGTTGAAAAGGCAGTGGCGCGAAGCTACCGTGCGCTGGATTATGACT

GAACGCCTCTAATTCAGAATCCGGGCTAGAAGCGACGCATGCGCCCGCCGCCCGATTGGCGACCCAAAAT

AAGGGCCTTTGGCCCCCAAGGGGAGGTGTCGTATGAGGAGCCGGTGCGGCGGATGAGCCGTGNAT

>lcl|Query_161782 GH985230.1 JutEST5233 Jute Plasmid library 20-45 days after sowing Corchorus capsularis cDNA similar to Putative uncharacterized protein, mRNA sequence

ATGAACAGAAATCCCCCTTACACGGAGGCATCAGTGACCAAACAGGAAAAAA

>lcl|Query_213185 GH985195.1 JutEST5172 Jute Plasmid library 20-45 days after sowing Corchorus capsularis cDNA similar to 60S ribosomal protein L27, mRNA sequence

GTCGACCTAGGGTTTTTAGCGAGGAGAGAAACAGAGCGCTGCAGCCGCCGTCGCCAATGGTGAAGTTCTT

GAAGCCCAACAAGGCCGTCATTGTCCTCCAAGGCAGATATGCCGGACGCAAAGCCGTGATCGTCAGGTCC

TTCGACGATGGTACACGTGATCGCCCGTACGGCCACTGTTTGGTTGCAGGGATTAAGAAGTACCCAAGCA

AGGTTATCCGCAAGGACTCCGCCAAAAAGACTGCCAAGAAATCCCGGGTCAAGTGCTTCATCAAGCTAGT

GAATTACCAGCACCTGATGCCCACACGTTACACGCTGGATGTGGACTTGAAAGATGTGGTGACCGCTGAT

TGTTCTTCAACAAAGGACAAGAAAGTTGCAGCTTGTAAGGCTACTAAGGAACGTTTTGAGGAGAGGTTCA

AGACCGGGAAGAACAGGTGGTTTTTCACCAAGCTTAGGTTCTGAGCTTTTGTTGGTTTGTCAGTTGTTGA

GGTATTATAAGTCGTTTTGAATAGGAAAAATTTGAGCTGTCATGTAGTGTCTTTTAACAGTTTTGATTTT

AAGATAATCGAACTGCTTCCATGAGATATTACTTCTGGGCATGATAGTAATGCTGATTTGCTCATTTCTG

TTTGAGAGTTATGTTC

>lcl|Query_186647 FK826539.1 JutEST4323 Jute Plasmid library 20-45 days after sowing Corchorus capsularis cDNA similar to Metallothionein 3b, mRNA sequence

GGGGGGCACATTTTCACTGTAAACAGAAAGCTCAATACAGGAACATATCCACCATCAACGAGCAACAATG

GCAACCTTATCTAGCCTTAACCTTGTAACTCCAAGAGTCGTCGCCAACGTGCCAGACTCCACGAGGGTTG

GCCCTGTGAAAGTCCCATGCCTTAACCAGCCCTGGAAAGGGTATCTCAGTTGGGGTCAAGGCGGATGCAG

GTGTTCAGACCTGTGAGAGCCGCTCCCGACCAAATATCGGAGAAGGTGGAAAAGAGCATCAAGGAAGCCG

AGGAGGCATGCTCAGGTGACCCAACGAGCTCAGAGTGCGTGGCGGCTTGGGACGAGGTGGAGGAGCTCAG

CGCCGCCGCCAGCCACGCCAGGGACAGGCAGAAAGCAAACAACGACCCTTTGGAGAACTACTGCAAAGAG

AACCCTGAGACAGATGAGTGCCGCACTTATGATAATTGAAAATCGCTTTCTTTTTCTTTAATTAGTTTAT

GATGATAAAGGTTTTAGAACCTTCGATGATGAATATGGCGGGGGATTTATGTGGGGGGG

>lcl|Query_121685 JK743773.1 JutESTSSH3(N10) Jute Plasmid Suppression Subtractive cDNA library Corchorus capsularis cDNA similar to cytochrome P450-like TBP protein, mRNA sequence

TGGAAATCAGAATCAAACGAGCTTTTACCCTTTTGTTCCACACGAGATTTCTGTTCTCGTTGAGCTCATC

TTAGGACACCTGCGTTATCTTTTAACAGATGTGCCGCCCCAGCCAAACTCCCCACCTGACAATGTCTTCC

ACCCGGATCGACCGACAGATGCCGGTCTTGGGTCTAAAAAGAGGGGCGTAGCCCCGCTTCCGATTCACGG

AATAAGTAAAATAACGTTAAAAGTAGTGGTATTTCACCGTTGCCGAAGCTCCCACTTATACTACACCTCT

CAAGTCATTTCACAAAGTCGGACTAGAGTCAAGCTCAACAGGGTCTTCTTTCCCCGCTGATTCTGCCAAG

CCCGTTCCCTTGGCTGTGGTTTCGCTGGATAGTAGACAGGGACAGTGGGAATCTCGTTAATCCATTCATG

CGCGTCACTAATTAGATGACGAGGCATTTGGCTACCTTAAGAGAGGGAGGATTTCCGATTGAGGCTAGGA

GTCACCAAAAAGATTTGAATGAA

>lcl|Query_124450 EE332672.1 EST00168 Jute Plasmid library Corchorus capsularis cDNA, mRNA sequence

ATGGGAGCGGGTGGCAGAAGCAGGAGTCAGCTCGGCCGCGCCGGCGGCGGCGCAGCCTTCAAGCGCTCGC

CGACGGACAAGCCGCCATTCACGCTCGGTCAGATCAAGAAAGCGATCCCGCCTCACTGCTTCCAGCGCTC

CATCATCAAGTCCTTCTCCTACGTGGTTCATGACCTGGTCCTCATCGCCGCCCTGCTGTACTC

>lcl|Query_210945 GH985258.1 JutEST5277 Jute Plasmid library 20-45 days after sowing Corchorus capsularis cDNA similar to ATP-dependent protease catalytic subunit, mRNA sequence

GATCTTAACTGTCGTGGGCCCATTTCAGCGTCAAAAACTTTTTTCCTTTTCACACCAGAAACTATTAACA

ATATTTATGTTATGAAAGAGTACAAAAAAAAGAATCTTTTTTCTTTCTTTTTTTGAAAAAAAAAAACTTT

GGGATTGCTGAATAAATATATAAAGCAACGGAGCCATCATAGTATTTTTTATCCGAAAAAAAAGAAGGGT

GGTAATTGGATTATTTATTGATCTGGGTCTAATAGACTCTATATTTTCTCTTTTTTCGATGAAAGAAAAA

AAGAGAATTCATTCCATTGTAAAGCCGTATGCAATGCGCAAAAAATGCCTGTACGGTTGTTCATCAGGGT

AATGATCCATCAACCTACTAGTTCTTTTTATGAGGACCGAACGGGAGAGTTTTCGCTGGACACGGGAGAA

CTCATGAGAATTCGCCAAAAGATCACAAAAGTTTATGCACAAAGAACGGGCAAGCC

>lcl|Query_81759 FK826545.1 JutEST4181 Jute Plasmid library 20-45 days after sowing Corchorus capsularis cDNA similar to 36kDA porin II, mRNA sequence

ACTTTGCCCTCTTTATCCCCCCGGGAGGTTTTTTGG

>lcl|Query_66118 JZ532439.1 JEC12 Library of differentially expressed transcripts in jute upon elicitor and drought stress treatment Corchorus capsularis cDNA, mRNA sequence

ACTGGCTTTTCAACCAAGCGCGATGACCAATTGTGCGAATCAACGGTTCCTCTCGTACTAGGTTGAATTA

CTAG

>lcl|Query_205389 GR463732.1 JuteSSHRP79 JuteSSHlibMP1 Corchorus capsularis cDNA clone JuteSSHSTPK similar to Ser/ThrProtein kinase (PRK) like protein, mRNA sequence

GCGTGGTCGCGGCCGAGGTACAAAAACCAGAGACCTTATATGTGAAATCTTGTAGAACTGTCATAGTAAG

TCCATCAAACTTCTGTAACAGATTAAGCCGGTGAACGGGCTGGATTTTTCCCAGAAGGCAGTTGCTTTTA

AATACCATCATAGTAACCTCCAAATGTTATTTGAGCTCTGGTGATTGGAACTCCTCTGGCTGCAGAAATT

GCTTCATTGAGTTTGGTTGCATGATGAACCACTTGCTTCATTCTCTTGTCATCAGTGTAACGCTGGAACA

GAAGCTTCTGATTCTTCTGATGCTGTCTGTGTCTTGTTGACTCCCAGTATGATGATACAAAAAGTAGCAG

CAACAACACCAAAGAACAGATTGCTCCCATTCCCAGTTAAACCCCCATTCCCAAAATTCAGGCCCACCAG

ATTCACCTGCCCGGGGGGCCGCTCC

>lcl|Query_155017 EE332613.1 EST00109 Jute Plasmid library Corchorus capsularis cDNA similar to Ribulose bisphosphate carboxylase small chain 2, chloroplast precursor (RuBisCO small subunit 2), mRNA sequence

GAATTGGAGCACGGATTCGTCTACCGTAAGAACCACAGCTCCCCAGGGTACTACGATGGACGCTACTGGA

CCATGTGGAAACTTCCCATGTTCGGGTGCACTGACTCATCCCAGGTGTTGAAGGAGCTTGAGGAGGCCAA

GAAGGAATACCCCGAGTCCTTCATCAGAATCATTGGATTCGACAACAAGCGTCAAGTGCAATGCATCAGT

TTCATTGCCTACAAGCCCCCAAGCTTCACCTCAGCTTAAGAAAATAGAGTTGTTTTCCCCACATTAAGTA

GGGATTCGGCTTCTTTAACTTGTACTTAATTATATATAGCATTGTTCGAAAACATTTTTCTTTCCTTTTG

AGGTATTGTTGTTTGTTGTTTGATTTGGAATGGAAGGGCTGAGAACCATAGTTTCTTGCCATA

>lcl|Query_173117 GR463764.1 JuteSSHRP85 JuteSSHlibMP1 Corchorus capsularis cDNA clone JuteSSHCpn10 similar to Cpn10 Chaperonin 10 kd subunit like protein, mRNA sequence

CGAGCGGCCGCCCGGGCAGGTACTTCGGATGAACAAGGTAACAGGAAGCCGTTATCAGTTGCACCAGGAA

ACACAGTCCTTTACTCCAAGTATGCCGGGAATGACTTCAAGGACACTGATGGGAGTAACTACATTGCTCT

GAGAGCTTCTGAAGTAATGGCAGTGCTCTCTTAGTTGTCTAATAAGATATGCTTTAGGTGTACCTCGGCC

GCGACCACGC

>lcl|Query_212957 EE332552.1 EST00048 Jute Plasmid library Corchorus capsularis cDNA similar to Metallothionein-like, mRNA sequence

TATATTTCATTAAGATGAGACAAGGCCAAAGGCATGGCCATGGCCATTACATCACAGTATATAAGGAGAC

ACAAACNTGACANACAGGACATGAAAACACAGACACCAACAGAAACNGGAAAGACACAAATTATTCNNAC

CCAAAATAAGATTTTAGGTCCCAAATATTGCCCATGTTTCTGCAACATGGGATCATTTGTTTCATTTCTA

TATATATGTTTGCTTAATGACCACCGCNTGTGCAGTTTGTGCAGCTGCAGCTAGATCCACACTTGCACTT

TGTCGTTCTCTGCTGGAGGAGCCTCCATCACCACGGTGGCAACGTAGCTCTTCTCAGTCTCAATGATCAT

GGTGTTTCCCTTCTTGGTGCACTGGCTCTTGTCAGCGCAGTCGCAGTCTCCGCACTTGTCCGANATGGTG

AATTGAGGAAGGTTTTTTGAAAGAAGTTGGAAGAGCTTTTGGGTTATTGAAGATTGA

>lcl|Query_237453 FK826604.1 JutEST4004 Jute Plasmid library 20-45 days after sowing Corchorus capsularis cDNA similar to Peptidyl-prolyl cis-trans isomerase, mRNA sequence

ATAGTATTTCCATATGAAATCCACAGACTGCTAATAGATCGTTCTTGATAAGACATTTGAATAAATTAAG

CATAAATTTATTTGAGTGGTTC

>lcl|Query_107341 FK826507.1 JutEST4741 Jute Plasmid library 20-45 days after sowing Corchorus capsularis cDNA similar to Wound-induced protein 1, mRNA sequence

TGGACCATTATTCTTAAATCTTAAACATACCCAAAAGCATAAACAGAGGATGCATGAAGTTGAAGCCTCT

TACTCATATAGAAAA

>lcl|Query_149944 GH985156.1 JEST5110 Jute Plasmid library 20-45 days after sowing Corchorus capsularis cDNA similar to Epsin, mRNA sequence

GGGGCCTCTTCCTTATCCATAACATTTTAGTACACGGTCCGTTCATCCTGAAAGATCAGCTCTCTGTATA

TCCCTGTTTTGGTGGTCGGAATTATCTGAAACTTTCTGCTTTTCGAGATGATTCTGATTCAGAAACGCTT

GAATTATCAACGTGGGTTCGATGGTACGCAGGATATCTTGAACAGAATTTATCGTCTTCAAGAGTCCTAG

GTTATCACCTTTATTCCTCATCGTGTGCTAATAAGAACAAGATTTGGTCATTGT

>lcl|Query_155853 GR463663.1 JuteSSHRP39 JuteSSHlibMP1 Corchorus capsularis cDNA clone JuteSSHL2A similar to Arabidopsis thaliana ribosomal protein L2 like protein, mRNA sequence

CTTTGGCCCCCAAGGGCACGTGTCGTAGGAGTAGTCCGTGCGGCGGATGAGCCGTGCAGGCCTCCTTGAA

GCGTAATTCCCACCGAGCGGCGGGTAGAATCCA

>lcl|Query_103197 GH985181.1 JutEST5147 Jute Plasmid library 20-45 days after sowing Corchorus capsularis cDNA, mRNA sequence

GGAAAAAACATCTACGGTTAAGCATCTTTTAACTTTGCCGCAGTTCAACAGCAGTGTGACTGCACTCTCT

TGGGCTGGTCTTGATCGAAAGAGAAATCATGGGCTTCTCGCAGTTGGAATGGAGAGCGGACTCCTTGAAC

TTTGGAGCCTTGATGTTGAAAGAACACCAGATAAAAGCACCCTGACACCAGCTGTTTCTGCTGCTCTAAT

CGTGCGGCTAGATCCTTTCATGTGCCATGTTTCTGCAATTAACCGGTTGGCCTGGCAGAACCGAGAGAAT

GATGAAAATTGCACAAGCCTCAAGCTTGCTTCTTGTGGAGCTGATCATTGTGTGAGGTTGTATGAGGTAA

TTGTTGACTAAGTCTAATCTGATAATGATTTTGTATTATAGATACATCAGTATGTCATTCTTCAAATCAA

ATACTACAACATATGTGATTCATATATATATATCTAGATTTTAACTTTCATCATTC

>lcl|Query_103349 FK826527.1 JutEST48136 Jute Plasmid library 20-45 days after sowing Corchorus capsularis cDNA similar to Hydroxyproline-rich glycoprotein DZ-HRGP precursor, mRNA sequence

GTGAGACGAGGTGAGTCGTTGGATTCGAAGATTAAGAGCAGTAACGTGGAAATGAAGGCGGGTTGGTTGT

TACGGGTACGGATAGATTGGGTCCGGACCCCAAAATGGTTCCAAAACAGATCAAGATCGTGGATATTAGG

TCTCCGGTCACCGGGAATTCCGACGTTTACGCCGGATCGGCCTTTTCAGTATCTCCGCCCGCCGAGTTCT

CTTCCTTTGCCGTCGTTCTCAAAGAAGAAGCAGGTTTCGATCGATGATTCAGCTACTAGGGATCTCAGGC

GTCTACTCCGGCTCGACTTCTGAGACCCGAATAGAATCTTTCGGGTCCGGGTTGCCTTTTTCTCTCCATG

TTAATCTCTCTCCTGATCCGGATTTATATTCGGATCCCAGAAAAAGTCTCCGTCTCTTATCTCCTCTCTC

CCTCTCTATCTATCGATCGATCAGTTGTCTCTAATTAGAACTCTCCGGAAATTTTCGAAATCCAGTTTCC

TCCGTCATATTTTTTTCAAAACGGTG

RID parameter must be supplied

>lcl|Query_210262 GR463662.1 JuteSSHRP37 JuteSSHlibMP1 Corchorus capsularis cDNA clone JuteSSHLACCCASE similar to secretory laccase [Gossypium arboreum] like protein, mRNA sequence

CGAGCGCGCCGGGCAGTACGGCCAGGAGCATGGTGAGAGATCGGCGAAGACCTTGTGGTGTAAGCCCCAT

CTTGTCCCACAACTGTGAACATGTGGTATTCCACGCCAAACTCCATTGTCTCATTCAAAGCCGCATTTAC

AATTCGAAGGTGATAAACACTTTGGTAATCTACTTGCTGCTCGTATGTTGAATTGTTGCTGCAGCCATAT

GTTTCTCCAGGATGGCCGACCTCGGCCGCGACC

>lcl|Query_27778 EE332649.1 EST00145 Jute Plasmid library Corchorus capsularis cDNA, mRNA sequence

TGTCAAGGGCCAAGCAACATGCAAGATTTAAGGTGTTACAGTGCTAGTTATGCAAATTCTGTTGTTCAGG

TTCAGCCAAATCAGCTTGGGAACGAGATTAAGATCAAGAAAAGCAAAAGCAGTTTTGGGTCATCTTCAAA

AAGCTGGAGCTTTAATGATCCTGAGTTGCAGAGGAAGAAAAGGGTTGCAAGTTACAAGGTTTATGCAGTT

GAAGGCAAGATGAAAGGGTCTTTCAGGAAGAGTTTTAGGTGGATCAAAGACACTTATACCCAAGTTGTCT

ATGGATGGAGA

>lcl|Query_27617 FK826551.1 JutEST48119 Jute Plasmid library 20-45 days after sowing Corchorus capsularis cDNA similar to Specific tissue protein, mRNA sequence

GGGGCAGTGCAAAAGTTATTGCATTTTTTAAACCATGAAATCTTCTATTGCATTGTTTCTCACTTTTCTT

GCTTCTGCAGCATGCAGACAGCAAGAAAAGAGGTGGGGAATATGCTGGAGGGCACTATAGAGCTGAAACG

ATGAATCAACCTATGCCGGAAGCAATCCAAGCGCTTGTTCGGGTTAATGCCGCAGCTGCCCTCCCCGGTG

ACAAAACCGACTGTCATACACCAAGTAACGTGGAGACTAAAGAAGAGAAGATTATTGTGAATGAGGAATT

TGAACCAAGGCCAAACATATCACCTTGAGTTGAAGAGGGTAAGATCATTGTTGCTGATGATTTTGAACCA

AGGCCCAATATTTCTGCTTACCATGATGATGGTGATGTAAAGGGGGAGAAGAAATCATCTTGTGTTGAAG

ATTTTGAACCAAGGCCCAACATTTCTGCTTACGGTGGCGGATGATGCTGATCTTAAGGGGGGGAAAGAAT

CATTTGCTAATGATTTTGAACCAAGGCCCAACATTTCTACTTACCATGATTAATTGGAGGTGAAATTAAA

ACAGAAATAAATAAAAGAGAGAAACCAATTTCTTTGATTTCTCTTAATTAGAAAAAATCATTATGGCCTC

GTGGCCTCTATGTATGGCATGGTTTGTTTT

>lcl|Query_234610 GR463753.1 JuteSSHRP23 JuteSSHlibMP1 Corchorus capsularis cDNA clone JuteSSHSTPP similar to Serine/threonine protoin phosphatase like protein, mRNA sequence

GCGTGGTCGCGGCCGAGGTACCTCTCAATTCTTGCTTTAACACTTTATTCAATAGTTGGCTCATATAGTA

AGATAAAAGTCAGTCACAGAGACATGTCTCATAAAGAAGCCTGGACTACGGCAATATTCGTCGAATTGCC

AAGTCGAATCACCAGCTACTCCAGGTAGAAAACTACATTACATGTGAGCATTGAAGGGTGGGCATGTGTT

GAAATATTGCTGTTGCAGCATCCGTGACTAAAGTCGTTGCTCCAATGTTGATCACACACAAATTCTGATG

GGGACACGAAGCTGAAGCAGATTAGGCTGAGAAAGAAGACTTGGCGAGTATTAATACAAAGCTAACGGAT

CTCTCCTTCATAAAGCTGAACCTGCTTTCCTCCTCTGCCATCCAACAACCTTCTTATTATTATCTTCACC

ATCAGTAACTCCATCAGAAGATGCCATGTTGTTTTCATGCTCTTTTACTTCTTCCATGTACCTGCCCGGG

CGGCCGCTCG

RID parameter must be supplied

>lcl|Query_163914 HS411990.1 JBM17 Library of differentially expressed transcripts in jute upon stress Corchorus capsularis cDNA, mRNA sequence

GAAATCGACTTTTACCCTCGCTTCCACGACAACACCGATAATCTTGCAGTTCCCGTTGATAGGAGTCATA

GGCCATGAAGGATTCAGGAAGCTTGATTGCCTAGCTAGTGAAATTGCTCCAAGTCTAAAGCAATATTCAA

AGATAATTTCGTCTCTTGGGTATTTCTTCTTGTCTTCATTTCACTATTTAAGTGACTGGGTTACTGTTTA

GCTGTCAAAAAAAAAAAGCTTCCTTTCAGGTACTTCTGACCGCCATCTATGACCAGTTTCTTGAATGTTG

CTTCGTTCGCGTCAGTCAGTTTGGCTACAACAAGGCTTCCATTCACTGGCTCGCGTCCAGTATCTACTAA

CACCATATGACCTTCAGGGATGCTTTGACCTACAGGTGAGGTCATGGAATCACCTTCAACCTTCAGCCAG

AATCCATTGCCTAATAAGTTAACGTCACTGTCATACCATTCATCAATGTCCTTGATATCGTAGGGTTCAC

AAGCTTCACACCACGAACCAGCTCTAACCATGCTAATCAATGGATATTTCCCTTTGGGCTCAACGTGCCC

AACAAATCTAACAT

>lcl|Query_34177 JK743784.1 JutESTSSH3(N25) Jute Plasmid Suppression Subtractive cDNA library Corchorus capsularis cDNA similar to GDP dissociation inhibitor 1, mRNA sequence

ATCCGTGACGGTCGATTCAAAGTGTGTTGTAGCATCATAACTCGTTGATATAAAACAATTGTCCAAAGAA

GGCTCATTGACTGGTTCATATCCGTCATAGATGTCAAAGAATATCTCATCAACAGGACCAAGAAGGTCAA

TTCCAGGCTTTAGCTCAGTTTCAGGGTGATCCGTCTCGGCCTCCGTAGAAACAAATGCAATAAATTTGCC

CTTAGGAGCAACATTGTGAGAATATGAGCAACAGAAGACATACATGTCCGATCTGCGACCCAACTGCTTT

TGTGGCAGGATAATCTGCACTGAATGAGAGTCATTGGTGTTTGGGATTGGGTGACTCATGATAGCAATTG

CCCTGGCAACCCTGCCAATCTTTCTAACCTTGTTCGTCAAGTAGGAAGGATCACAAACAACTTTCTTACA

CCTGGCAGTTTCACCTTCAGATGTGACGCCGCAAGCTTTACCTTCATCATCAAACTCCACCTTGCACTCA

GGCTTATTCAACATGTATGTTCCACCATATACAGCACTAAGCCGTGCAAAAGCCTGGGGCAGCTCGCCCA

ATCCATACAGTGGATAAATGTAAAGGGATCCACCTTGAAAACGTGCAAGAGACTCTGCATACAGCTTCAT

CCTCTTTACAGTATCCAATGCAGGCTCATCCAGGTAACGATCATCTCTATGAAGGGCCAATGCATGACCA

ATGAAGTCCACGGTGTTATCATCAAGACCATATTTCGCATTTAGCTCCCTAGTGGTCACCGTAGTTCAGG

TCCATTCCCTCATGACTTTAGGATCCCTTTTCGTCATATTCTTGAACATAATATGAGAACTTCCGTGCAC

CGACCGCTTTTCAAATATTCCCATGAGTGGAGATTGGAGTGCCTCCATTGTCCAGTTGGCTTGCACGTAT

GACGTTGGTTTAATGAAACATAGCCTGCCATTCACAGCCTTGGAAGTTACCTCAGGCGGAACCACGCCTA

TCCTGATGCATCGCGAGTTACCGAGTCGATTCCATGTCCGTCGTTGAACACGTCCGTTGACTA

RID parameter must be supplied

>lcl|Query_230379 JZ532428.1 JEC1 Library of differentially expressed transcripts in jute upon elicitor and drought stress treatment Corchorus capsularis cDNA, mRNA sequence

GCATTAGAATTTGGACACAAGACAGTAGCTGGTGGTGTTCATGACAGCTTCTTGTTTAAACCTGATGTCC

TAGGCCCTGCAGGTTTGACTCCTCTCCACATTGCACTGGTAAAGATGGTTCTGAAGAAGTGTTGGATGCA

TTA

>lcl|Query_236627 EE332675.1 EST00171 Jute Plasmid library Corchorus capsularis cDNA, mRNA sequence

CGTGAGCTGGGTTAGACCGTCGTGAGACAGGTTAGTTTTACCCTACTGATGGCCGCGTCGCAATAGTAAT

TCAACCTA

>lcl|Query_134151 FK826437.1 JutEST4410 Jute Plasmid library 20-45 days after sowing Corchorus capsularis cDNA similar to unspecific monooxygenase, mRNA sequence

TTCTTGGCCTCAACCTTCTTTTTGTGCTCCTCATCCTCAGATTT

>lcl|Query_205284 FK826544.1 JutEST4384 Jute Plasmid library 20-45 days after sowing Corchorus capsularis cDNA similar to cytochrome c oxidase, Vc subunit, mRNA sequence

GGATTGGATTATCACTTGGTCTTTTTGCTGGTGGGCTCTGGAAGATGCATCACTGGAATGAGCAAAGGAA

AGTGAGGGCATTCTATGACATGCTTGAGAAAGGTGATATCTCTGTTGTCGTAGAAGAATAGATGTTCATG

TCTTGTGCAATTGTGGACAGTCTTGGCAGTATTTTGATATTTTGTGTTTCCTTCATTTGGCTCTTAGATT

CAAAATGAGCAAACTCAGTTTGGAGTTATTAAGAATAATTTTATTGTATTAAGATCCAGATCACATATCT

GCTTAACAATCTCAATGGTTTTGTGATATTGTGTCCTTAAACTTTTTTCCTTCTGTTTTTGTGGCATCTC

TTGAGATG

>lcl|Query_50334 FK826518.1 JutEST47139 Jute Plasmid library 20-45 days after sowing Corchorus capsularis cDNA similar to Aux/IAA protein, mRNA sequence

AAATTATAAAACCATCAAGGCCTAGTGTTTGTGAGTAACCTTTTTACTATTGAATCAAGTTTTAGAGTCT

CTGGACAACTCTCATGTTCCCACGCAGGATCTCTAGTTTGTAAAGTGCATTGTTTGTTAAAAACTTTCAT

CGGTGTGTGTTTGTGTGATGAAAGACCTCGACATTATCTGTAACCCCGACATGATTGTATTAACTCTATT

GTGCGGCTGCTTGTGAGTGTTGTATTTCTTCTGCTTAAGTATTATTCTGCAAATCATTTTAGTTCCTATT

CTGCTAACTATAACCCCAATTCCGAAAACCGGTCGGGGGCTTTGAAAATGCATATGTACTGAATCATATT

AGTAGTAGTTTTATAATGTGTTTATGAGTTCCAAAAAAA

>lcl|Query_30729 GH985246.1 JutEST5253 Jute Plasmid library 20-45 days after sowing Corchorus capsularis cDNA similar to Unknown, mRNA sequence

GCACAAAACAGGGTTTCCTGGACATGGACTCTGAATCAAGCCTGGAGAATATCATCAAGAAAAATGGCTG

GTGGACAAGAAGCAACTGGGCATTCCTGAACGAACCACCTGTTCTTGAACGCCCTACCAACAGTTACAAA

GCCCAGTTCCACATAGCCAACCTTGCTGCTTCAAAGCTTAATAGGGATTCTGGAATCAGTGCTTAAATCC

TCAGCTATCTTAATCTTAATCCTACCTTGTGCTAGAAATATATAGCATAAAACTTGTTCTGCCTAAAATC

TGTATATATAATGATGTTCTACAAAGTTTGTAAGAACATTACCCTGATGTGTCGGCTGAGTGTGTTTAGT

CATGAGTGTTGTCCTGATCTGGCGTCTGTACAAAACTAGTAGTAAAATGAGCTGTGTAATTTGCAGGTGG

ATGATATTTCGTAAGCTGTCTGCATTATATAAAGATTTGTTTGAGATTTTTAT

>lcl|Query_2066 GH985247.1 JutEST5254 Jute Plasmid library 20-45 days after sowing Corchorus capsularis cDNA similar to Eukaryotic translation initiation factor 5A isoform VI, mRNA sequence

GCTCCTCCTTATCCTCTCGCTATCTTCCTCGAAAAGCTTTGAAATTTCTTAGTGTCTGGTCAATCGCGGC

CATGTCTGACGAGGAGCAGCACTTCGAGTCCAAAGCTGACGCGGGAGCTTCGAAAACTTACCCTCAACAA

GCCGGTACCATCCGCAAGAGCGGTTACATTGCCATCAAGAATCGTCCTTGCAAGGTTGTGGAAGTGTCAA

CTTCCAAAACTGGCAAGCACGGTCACGCTAAGTGTCACTTTGTTGGAATTGATATCTTCAATGCTAAAAA

GCTTGAAGATATTGTTCCCTCTTCTCACAACTGTGATGTTCCTCATGTCAACCGTGTTGATTACCAGCTG

ATTGATATCTCTGAGGATGGATTTGTGAGTCTGCTCACTGAGGATGGTAACACTAAGGATGACCTCAGGC

TTCCAACCGATGAGGCTCTTCTTAAGACGATTAAGGGATGGATTTGCTGA

>lcl|Query_143906 EE332645.1 EST00141 Jute Plasmid library Corchorus capsularis cDNA, mRNA sequence

TCGAGACCTTGAAAGCAGCTTTTTCAAGTGTCAGTAGCGGGACGTTCTGTCAATCGGGGAAGGTTTTTGG

TAACAACACCTGGAGATATCAGAAGTGAGAATGCTGACATGAGTAACGAGAAATCCTGTGAAAAACACGA

TCGCCTGCCAGTGGAAGGCTTTCTGCGTTCAGTCAATCTACGCAGAGTGAATCGGTCCCTAAGGAACCCC

CGAAAGGGCTGCCGTCCGATGGGTACACGAAAGTGACGAAGTTGCTTTGACTACAGAACCATGCCTGTCT

GTTGGAGCGAATTGGATGGATGATCGGGCCGAGGTTCTCGGCAAAGACCTCTTCCCCTCACTCTCCTTTC

CCTAATATTCACA

>lcl|Query_37765 EE332520.1 EST00016 Jute Plasmid library Corchorus capsularis cDNA similar to Tomato 25S ribosomal RNA gene, mRNA sequence

ACGAATTCGAGTCTCCGTGGCAGAGTCTTAGCCAAAGCGTCAATTTTTGAGTGTGATTTTGGTGATTGG

>lcl|Query_170257 GR463704.1 JuteSSHPS18 JuteSSHlibMP1 Corchorus capsularis cDNA clone JuteSSHGT9 similar to glutathione transferase like protein, mRNA sequence

CGAGCGGCCGCCCGGGCAGGTACACCGTTCCTGTCCTTTGGGACAAGAAATTCAAGACTATTGTTAGTAA

TGAGAGCGCAGAGATAATCCGCATGCTTAATACTGAATTCAATGATATAGCAGAGAATCCTGCTTTGGAC

CTATATCCTCCTCATTTGAGAGCTCAAATTGATGAGACAAATGAATGGATATACAGTGGAATCAATAATG

GTGTTTACAGATGCGGCTTTGCTAGAAAGCAAGGGCCTTATGATGAGGCTGTGAAACAATTATATGAAAC

CATGGACAAATGTGAGGAGATTCTTTCCAAGCAGCGGTACCTCGGCCGCGACCACGCT

>lcl|Query_221535 GH985187.1 JutEST5158 Jute Plasmid library 20-45 days after sowing Corchorus capsularis cDNA similar to 10 kDa photosystem II polypeptide, mRNA sequence

GTCTCCTAAAGTTTTCAAGGTTCTAGCCTCCGGTGGCAAGAAGATCAAGACTGACAAGCCTTATGGAATC

AATGGAGGCATGGCCTTGAGGTATGGTCTTGATGCCTCTGGAAGGAAGGGCAAGGGAAAGGGTGTTTACC

AATTTGTTGACAAATATGGTGCTAACGTCGATGGCTACAGCCCTATCTACGACCAAAAGGACTGGTCCCC

ATCCGGTGATGTTTATGCTGGCGGTGTAACTGGTTTGGCAATCTGGGCAGTTACCCTAGCCGGTATTCTT

GCAGGTGGTGCTCTTCTTGTTTACAACACAAGTGCTTTGGCACAGTAGATTTCTACTAATGTAATGTACC

GTGTTGTATAATATAGCCACTCCAAGTTATTGTTCTCATACATAATCACTTCAAGCTACTTTGCCTATCT

ATATGTCATTTTATTGTTTAAGCCTGGACCAA

>lcl|Query_239809 EE332671.1 EST00167 Jute Plasmid library Corchorus capsularis cDNA, mRNA sequence

CGCTGTATGAACAAATTCAGACTGTGACACTGCGAATGGCTCATTAAATCAGTTATAGTTTGTTTGATGG

TATCTACTACTCGGATAACCGTAGTAATTCTAGAGCTAATACGTGCAACAAACCCCGACTTCTGGAAGGG

ATGCATCTATTAGATAAAAGGTCGACGCGGGCTTTGCCCGTTGCTGCGATGATTCATGATAACTCGACGG

ATCGCACGGCCATCGTGCCGGCGACTCATCATTGAAATTTG

>lcl|Query_206535 EE332550.1 EST00046 Jute Plasmid library Corchorus capsularis cDNA similar to Photosystem I chain II precursor - spinach, mRNA sequence

GGAAGTCCCCACCAAAGAAGCGCCAGTTGGTTTCACTCCTCCTGAATTGGACCCAAGCACCCCATCACCC

ATCTTCGCTGGAAGCACCGGCGGACTGCTGAGGAAGGCCCAAGTGGAAGGAGTTCTACGTCATCACATGG

GAATCGCCAAAAGAACAGATCTTCGAAATGCCCACCGGCGGCGCCGCCATCATGAGGGAAGGACCGAATC

TGTTGAAATTAGCGAGAAAAGAACAGTGTTT

>lcl|Query_206659 JK743808.1 JutESTSSH3(N53) Jute Plasmid Suppression Subtractive cDNA library Corchorus capsularis cDNA similar to senescence-associated protein, mRNA sequence

GAATCTACATGGACAAGTACCAGCCAACTCAAAAGGGAAAGATTGGAATTACCATAAATACCACTTGGTA

CAAACCACTGAATGC

>lcl|Query_185046 FK826508.1 JutEST4742 Jute Plasmid library 20-45 days after sowing Corchorus capsularis cDNA similar to Wound-induced protein 1, mRNA sequence

ACCAGTTCTTTCTGGTTTTCGTTTATCCAAATCTGAAACAACTCTCCTTAATTAAGGAACCCAAGAACTC

TTTTCATCTTTCTTTTGTGCCTATCGATCGGTTTTCTTGGTTCGTTGTCTCTTTCAAGAACTGGCTAACT

CGCAGGAAAAAATTCTGGAGGAAACCAACAAGAACCAAGAGGTGGTTCTTGCCCTCTATGAGGCCTTAAA

GGCTCGAGACGTAGAAACCGTTCACCAAATCCTCGCACCAGACCTAGAGTGGTGGTTCCATGGTCCGCCC

ACCCACCAGTTTTTGATGCGCCTTCTCACTGGAGCTTCAGCCGACGATTCCTTCCGTTTCGACGTTGATC

CTAACTCCGTCACCACCTTTGGGTCCACCGTAATCGTCGAAGGCTGCGATCACAGCCGCTCAATCTCCTG

GGTTCACGCCTGGGACTGTTTCTGATGGGATAATCACACAAGTGAGGGAGTACTTCAACACTTCTCTCAC

CGTCACTCGCCT

>lcl|Query_58997 EE332640.1 EST00136 Jute Plasmid library Corchorus capsularis cDNA, mRNA sequence

TTTTTTTTTTTTTTTAAAACACAGACAAAAGCCACAAACGTAGAACTTAAATTTATTTAAGTTTTAAATA

ATGACAATATGAAACAAGAGCATCATATTGTTAAACGATGTGCCAGTCTGCCAGAGCATCACACTAAACC

AAAAGATGTCATCATAATGAAAATAGTAATATCACAAGACAAATATTCAATAGTTGCTGTTTAGTTTGGG

GGTTCTAAGTCAAGACCCTCATTTCTTCCCACTCTTCTTGAGTCCAGCTCCACCAAATGATCCCTTTTGT

TGTGCCTTTGCTCTCAGCTCTTTAAGAGCCTTCTCTTCTTCCTTCTTCTTCTGAAGCAAAGCCTTATCAT

TCTCGTCGAGTTCTTTCTTCTCAGCCTTGGGTTGCTTCAAAGGCTTCAATTTGCCACCTTGCTTGGACGA

CATGGTTAGGATTTGCGACTAAAATTCGAAGTGATCAGGGAAATTTGGGAGACAGAACAAATAAGAs

>lcl|Query_161507 HS411989.1 JBM16 Library of differentially expressed transcripts in jute upon stress Corchorus capsularis cDNA, mRNA sequence

ATTATTTAACGTGAGCTCGATTTTACCTCGCTTCCCGACAACACCGATAATCTTGCAGTTCCCGTTGATA

GGAGTCATAGGCCATGAAGGATTCAGGAAGCTTGATTGCCCCATGTTCCGGGGGATGTTTTACTCAATCA

TGGGAGGATAATTTTTATAACCCTGTATGTTTCGTTCACATCGCGACTCTGTATTTCTAGCTTTTGTTAA

CTAACCTGTACTTCTCTTCTGATTTTTTGTTTTAGTATTTTTATAGTTTCGGATCAAAAAAAAAAAGCTT

CCTTTCAGGTACTTCTGACCGCCATCTATGACCAGTTTCTTGAATGTTGCTTCGTTCGCGTCAGTCAGTT

TGGCTACAACAAGGCTTCCATTCACTGGCTCGCGTCCAGTATCTACTAACACCATATGACCTTCAGGGAT

GCTTTGACCTACAGGTGAGGTCATGGAATCACCTTCAACCTTCAGCCAGAATCCATTGCCTAATAAGTTA

ACGTCACTGTCATACCATTCATCAATGTCCTTGATATCGTAGGGTTCACAAGCTTCACACCACGAACCAG

CTCTAACCATGCTAATCAA
