## Supplementary File 6 for "*In silico* Identification and Functional Characterization of Conserved miRNAs in Fibre Biogenesis Crop *Corchorus capsularis*"

**Supplementary File 6:** Total of 36 Putative non-coding EST sequences of jute genome

>lcl|Query_82742 GR463689.1 JuteSSHPC2 JuteSSHlibMP1 Corchorus capsularis cDNA clone JuteSSHIFO similar to indolepyruvate ferredoxin oxidoreductase, alpha subunit like protein, mRNA sequence

AGCGTGGTCGCGGCCGAGGTACATACGGGGAAACTGAAGCCAGGAGAAGAAGAGGGAGAAACTGTCCAGGGCCTCACAGGGAGGCATTCATGGAGCTGGAGAAACCAGGGATCCTGACCGCAGGGCCCATCAATTGCTACATAGAGATACACATGGAAAGAGGGTTTAACAGACCCTCTCTAGGACACAACTGTTTCTGCTTTGAGCAATAATTTCTAACCTGAGGCCAACATGTCCCACTGCCCCTTGGGTTGGCTGGGGTTGTTTCCTGAGCCAAGCATCCAATATCATGCCCCACTCAATGGCCTAGAAGCTGCCTGTACCTGCCCGGGCGGCCGCTCGA

>lcl|Query_149086 GH985271.1 JutEST527 Jute Plasmid library 20-45 days after sowing Corchorus capsularis cDNA, mRNA sequence

GCAAGGAAGAAGGAACAGTTAACTTACTTCCTATAGATTGTTGTCGAATGGAGGCTTCTTATGTAATCATCAACAGAATTAATCTCTATGAAGAGATGTGAAAAACTTTGAGCCAAAAGCCCCTTCAACGAGTAATTCAATCGTCCTGTTCCAGGCATGCACGGGGTAGCATCAAGACGGAAAGCTCTGGCACGGGTCCTTGTACAAAGAAATGCTGACCAATACCACAGGGCTCCTTGGTAGAGTGTGTGTGAAAAATAGGTATTAAAAGTATATATCTGGCATTTGTACATCATGCTTAAGAGAGGGTTCTTTGGCAAAGGGTACTCCATTCCATGAACCATTTGCCACCAATCTTTCGGTTACTTTTAGTCTGTGTTCTGTGAATGTGTGGTATTTGGAAATGCCTAGTTTTGACTTTCAAAACATGGGAGGAGTTTGTATTGGATTTGTTTAGAGATTCTGTAGTTCAGTCTATAATTTATTTATAATATAATATTTTTAGTTACA

>lcl|Query_149615 JK743796.1 JutESTSSH3(N39) Jute Plasmid Suppression Subtractive cDNA library Corchorus capsularis cDNA similar to Calmodulin-7, mRNA sequence

TCTTGAAGAAAGCCGTGGAACGTGAATCCACCAAACTTGCTCTGGCTGCTTGATCTACCTAGTTCGGTTTTGCTTTCATTTTTGGAACCTAATTTAATCATCCACCAAACTTACCTTATAATTCTTCTCGATTTCCACTTTCATCTCTTATATATATAGTTTATATCATATGCTCCAAATGGATCATTTCTTCCCCAAGCTAAATGTAATTTCGTATGATGTATAATTTAAATC

>lcl|Query_126885 GR463724.1 JuteSSHPC29 JuteSSHlibMP1 Corchorus capsularis cDNA clone JuteSSHGP3 similar to Glutathione peroxidase (Stretch 4-34 amino acid) like protein, mRNA sequence

TCGAGCGGCCGCCCGGGCAGGTACCTTATTCCCTTCCTGGATAATTATGTAGCAAAACCAAGTTGGGTGTGAACAGATTCCGCTGCAGAGGCTATAGATCTTATCTCTCACCATCTGCACGCACTAAAAAACTCACTCCGCGTGCAATTGCAGCACTTAAGTATCTTGGTCAGTGTGAACCAGCTATGATTTCTACTTCTCTTAGCATGTATTTTTTTTTGGCATCAACCAAGATTTGGTGAAGAAAGGTTTTCTTTTTTGCTTTTACTTCTTTTGGTGCCTGGTAAGTTGATCTGTGCAAACGCTTTAAAGTTTGCTAAATTAACAAATACTTGTTTAAAAATTCAGATAAAACAATTATATTCCAATAATGCAGATTACTACAATCATTTTAGAATTGTTGAAGGAGTCTGGGGAGGTATAAAAGGGCTTTGTAAAAAACGAGTTTGTTTTTCTTTAGGTGCATTGGCCAAAACATTAAACCAGCTTTAGACTGTAGTTGCTACACTCCTTAGAGGGGTCATACCAACTGCAGTACCTCGGCCGCGACCACGCT

>lcl|Query_126709 JK743804.1 JutESTSSH3(N48A) Jute Plasmid Suppression Subtractive cDNA library Corchorus capsularis cDNA similar to Pectin methylesterase, mRNA sequence

TGATTCTTTCCAGAAAAAAAGGAGATGTGCCCATGGGAATATACATGGGGCTTTTTTTCTAATTACTTTCTGTAAATTTATATATGTATGAAATATATATGAATTAAGTAGCTCTTGGAGATCTTG

>lcl|Query_124223 GR463708.1 JuteSSHPS14 JuteSSHlibMP1 Corchorus capsularis cDNA clone JuteSSHCSP similar to chromosome segregation protein SMC like protein, mRNA sequence

TCGAGCGGCCGCCCGGGCAGGTTTTTTTTTTTTGTGAATCACCTCCAGAAGTAGAAGACAATCTATTCATTATAATAGTTTCCATATGATACGAACGAAACATCATAATATCGTTTTTACTGGAAAAGGAGAATCATAAATTCAAAAGCGGGGAGTTTTCATCTAACCAATACAAAGGAATAAGTAACAAAAGATGGATATTAGTAAATCATCTCAATCCAGCATCTAATACAGGAGATATATTTACATGATCAAGCAAACAAAATGGATCAAAGTTTTGCTGGAAAGAGACAAAACAACTTGTTAATATTTGCAATAGGTGAACCCAAGCATTACCAGCAATAAGCTCACATCCAAGGTCAACCAACTTCATGTTTAATTTCCTAGCATTGAAAATATAGAGAATAAATAAACATAAGATAAAGATGAAAGATAAATTCTTGAATAAGAGGATGAGTAAACTGACAAAATGTATGTGAAAAAGTAATGAGAACCGTACCTCGGCCGCGACCACGCT

>lcl|Query_72459 JK743775.1 JutESTSSH3(N15) Jute Plasmid Suppression Subtractive cDNA library Corchorus capsularis cDNA similar to 40S ribosomal protein, mRNA sequence

ATAATTCGTGAGATTTTTATTTATAATATAAAATGGTAAGTTCTACAAAACCTTTTTGTTTATTAATAGTAATAATTTCTTAATTTGGACAAGAAAGAAATATTACATCATCAAATCTACTACTTCGTATACAAAGATGAAAGAGGAGCAGAGCAACAAGAAGTTAAGCAGACAAGAAATTAAAAAATCTTCCCTTTTAGAACACAATCGATAGCCCCCTTCGATAGAAAACATGTTTGAAACCAACCTCGTCGTCAATCCCATTGAATACTTAGCGGGTTAGCTTTTGTAATAGTGAA

>lcl|Query_107385 HS411997.1 JBM26 Library of differentially expressed transcripts in jute upon stress Corchorus capsularis cDNA, mRNA sequence

TAACAACGATATTTACCCAGTTTCCCGACAACACCGAATTTGCAGTTCCCTTGAAGGAGTCTAGGCCTGAAGGTTCAGGAAGCTTCGACTGTACCTGATGTGGTGAAAGCTATATTAGGGGAAGGGTATCTATTATATATATGTAGCTTATAAATAAGCAACCCAGAGAGTTGCATCGCATGCCTCTGATTTGTGTTTCCAAGACTACTTTTGTTTAACTGCATTTGGTCGGAGCTCGCCATGGGCATTTAAGACTTTTAAATTACTTATTAGTGTGTGAACTATATACCTTCCATTAAATATATCACATGCTTAATGCTAATTCCGGTACTGTACCAAAAAAAAAAAGCTTCCTTTTAGGGACTTTTGAACGCCCTCTATGACCAGTTTCTGATGTGCTTCTTCGCTCATCACTTGGCTTTGCAAGGCTTCCTTGCTTGGCTTGCGTCGGGTACCCACTACCACCTCTGACATTAAGGGATGCATTCGTCCTAGGATGCTTTCAAGGAACCAGTGAGGTCATGGCCACATCTTTTGCCTTACAGCCAAAATCCTTGCCAAATAAATTAACGTCCTCTGTACATACCGTCATAAATGTCCCTTGATAAACGTAAGGGTTCCACAAGCTTCACAGAAACGAAACCATTCT

>lcl|Query_106996 FK826520.1 JutEST489 Jute Plasmid library 20-45 days after sowing Corchorus capsularis cDNA similar to Ubiquitin Carboxyl-terminal Hydrolase-like zinc finger protein, mRNA sequence

TATTAAAAATAATTATTATTAATAGATCGGTAATCCAATATTTGTATTACATCTCAATTTACCGGTAGAATATTTTATGATAAACAATAATTTCAGAACAGTACAATAGTATAGAGTAAGATCAGAAAATGTACTATATTAATACACAAACTACTCCATATGCTAATATTATGGAAAACATAATTACAAGTAGATCATCAGATGCATAATTATGCCAAATATATAGCCTTAAAGCAGCTGACTGGCGGTGGGGATTACCGATGCCGCGATTTGTAATTAATGTGGATGATGATGATGGTGAGGACCTGGTGGAGGAGCGTGGGGAGCTGGGGCAGGATGAGGATGCTGCGGTGGACCTGGATGCGGTGCATGATGTGGTGGGGGGTGGTGGGGATCATGGTGGGGCTGTGGTGGTGGTGGTGGATTGTGCGGTGGTGGGTGGTGGTGGTCATGGTGGTGTGGTGGTGGTGGATTATGCATCTTCAATTTTCTTTTTTTTGCTATGTTAGGCTTTGTCTAGATATGGTGTGATTTAGCGAAGAT

>lcl|Query_92844 JK714338.1 JutESTSSH3(50B) Jute Plasmid Suppression Subtractive cDNA library Corchorus capsularis cDNA similar to Acyl-CoA-binding protein, mRNA sequence

TTTTATTGTTGCTACCATGTTATTCCCGAATTATGCAGGTTTAAACCAGATTTGATTTATCTATCAATAGTATAAACAAGCCCAAATAAGATGTTTGCTCATCTGATGGAAAATTGCTTGAATTCATAAATTAGCCATAATTGCTTAGCT

>lcl|Query_172361 FK826503.1 JutEST4720 Jute Plasmid library 20-45 days after sowing Corchorus capsularis cDNA similar to Glycerol-3-phosphate acyltransferase, mRNA sequence

GGCTATGGAAGTTTTGCTCCTGTCATTTTAATCTCGGTTGCTTGTTTCATTAATTGGTCAAGTAAGTGACTTTTGTAGTCAGGGCCATATATCATATATGTTGTTTAGCATTGACTTTAAGAATTGTAATCGATGCAAATGGATCGAGTGTCTATCACTCCATGAAGTTCTAATTGATAATGCAACTTTGTTTTTAAGGTGC

>lcl|Query_172287 FK826577.1 JutEST4892 Jute Plasmid library 20-45 days after sowing Corchorus capsularis cDNA similar to 60S ribosomal protein L41, mRNA sequence

ATTCACGCCGCTGCTTCTGTAGAAACAACATCTAACAAGTAGATTTGGTACTGCAGGAGACCCTTTTCGAAGACTTTTCACCATGAGAGCCAAGTGGAAGAAGAAGCGTATGAGAAGGTTGAAGAGGAAGAGAAGAAAGATGAGACAAAGATCTAAGTAGACTACTTTACTTTAGAGCTTATGCCCGTTTAGATTTCTGTTTGTTTGTTTTGATAAGATCCTTTTGTTAAAAGGATCGAACTTTATCTTCTTGTTTTCTGGGTTTAGTTTGTAACTTTTGTTGAACCAATTTTCATTTACTGGGAAAGTGTTATTTGAAGTTTAATATGCTTTTGGATCTAAAAAA

>lcl|Query_172689 JK714339.1 JutESTSSH3(51) Jute Plasmid Suppression Subtractive cDNA library Corchorus capsularis cDNA similar to Hypothetical Cys-3-His zinc finger protein, mRNA sequence

ATAGAACAACAGGAGCAGATGGTGGCATAAGGAAAAAGATGATCCGTTTGCTCATAGCTATATTCTAACATATTTCATATTCTTGGATTCTTAATATATATATTTGATAGCTTGTAGTCGGCGGTAGAGGAGGTAATATGTCTTCCCGACCAGTCAAGAAAGAAGCATAAACATACTTGAGCAGGAAGGTTTAAGTCACTGCTCATGTTAGAATTTTAGGAGCAAAAAGGAGAACCAAGCAACAAAGAAGATTTATTGATTATTTCATATTCTTGATTCTTTTTCTTCCCCCTTTGGGGATCTTTCTCTTAATGTTTTTAGCAAGAAGTTGAGAGATAAATGAACAGAGAACTCTCTCTGATCAAATTGTGTAATCTTAAGAGTTAAGAGGGAAATCCTCCATTAAT

>lcl|Query_52380 GR463677.1 JuteSSHRP74 JuteSSHlibMP1 Corchorus capsularis cDNA clone JuteSSHNBSLRR similar to disease resistance protein (TIR-NBS-LRR class),[Arabidopsis thaliana] like protein, mRNA sequence

CGAGCGGCCGCCCGGGCAGGTACACTAATACAGATTCCGGCACTAAATTTGAGATAACAAAAGAACTACTACAATGAAAGAACTCACTGAAAGTTTTCTGAAGTATCTAAAAACTAAACTTTTCGCTCAAAACATTCCTTCAAGTTCAAATGTCTGCTTTAAGAAAGTTGTCTGATAGAGATACCTTCTCCCCAGCTTGAATGGAGGAAGACCCTATATGGACATGTACCTCGGCCGCGACCACGC

>lcl|Query_168562 FK826528.1 JutEST4850 Jute Plasmid library 20-45 days after sowing Corchoruscapsularis cDNA similar to Leucine-rich repeat protein, mRNA sequence

GAGTACCAACCAAAGTGGGAGTCTCAATATAGTCAGTGTGAGTATAAGGCTATAAGCCAGAGGTATTTTACAGACAACATTAAACCATACAAGGGTACATACCCGAAGTAATCTTGTTACTACCATTACATCAAACACTGGTTTGTCTTAAAACATCATAGGAGTTTTTGGTATATATGCTAATAAATTCTGATTTACAGATACTTATACGAGATTGCATTTCCCCACACTGACTATATTGAGACTCCCACTTTGGTTGGTACTCTTTAAGGACACTGACTATATTGAGACTCCCACTTTGGTTGGTACTCTTTAAGG

>lcl|Query_177970 GR463683.1 JuteSSHRP92 JuteSSHlibMP1 Corchoruscapsularis cDNA clone JuteSSHSAP1 similar to Senescence-associated protein, mRNA sequence

GCGTGGTCGCGGCCGAGGTCAAGCTTTTTTTTTTTTTTTTTTTTTTTTTTTTTTTTTCCATAAATTCATAAAATGTTTTGATTTAAAAAAAATTTAACAGATACTACAAGAATTTTCCTAGAGATCTCAAGAACTATTCACATGATGGAAATTTAACGGGCGCCTTTCGATCACCTAACTGGTCAAAGATTGTAGCTAATCCGTCAAAATCTCACCGCCAAAATCATGCTACTCCAACACGTGCAAACTTCAAAGCCCCAATCTCATTGGTCAAGTCTCTACTCTGCAGTCTGCACATTTGCTGCCAAGCCAATCCATAGCCATTCAACCAAAGTACCTGCCCGGGCGGCCGCTCG

>lcl|Query_177392 FK826582.1 JutEST4467 Jute Plasmid library 20-45 days after sowing Corchoruscapsularis cDNA similar to unknown protein, mRNA sequence

GGTGAGACAACTTTGTTAATATGCAGTAGTGACCCAGTTATTCTGGTGATATGATGACTTTGCTCATCAAATTCCCCTGTTTTCTTTAGCATATAGCTAGGTTTCTCTTCCTTGTTTGCCCTTCGGTTATCTCCACGCCAATGGATTCGTTTAGTACTCAGAAACCTAAATCAAGAAAGATGAAGAAGAGGCTTGCGAAAAAGTCTCGAGTTGAGATAACAGGTTGATGAATGATGATGGTCCTGGCTGGTGGTTGGATGTTAGGATGTATG

>lcl|Query_60400 GR463780.1 JuteSSH6H5S95 JuteSSHlibMP2 Corchoruscapsularis cDNA clone JuteSSH6HKHSR similar to KH-type splicing regulatory protein like protein, mRNA sequence

ACAGGTTCTTAATTAGCCAGTCAGCCCATGAGCCCTTGGCACGAGTTTACCCAAGTTACCTACGAACCCTTAAATCCGGTATTTTGGGGGGCCCAGGGATCCCGGATTTTATGAGAAACCCTGGGGGTGCTGCCAGAAAAGGGAAAAACAGTTGGGGCAAAACACAGAGAATGGGTATTGTTATTCTCCCTTCCCCCCGGGGGCCCGCCCCAAGGGAGGGGGGGGAGGGGGGGACAGAGGGGGCCAGGGGGGAAAAATACTTTTGTGGTGGGGGGTGGGGGGGAAAAACACAACCCCCGCGCGGGGGGGGCGCCCCCCCCACCCTCTCTCTCTCTTGGTTATATAAA

>lcl|Query_12092 GR463686.1 JuteSSHRP108 JuteSSHlibMP1 Corchoruscapsularis cDNA clone JuteSSHSDR2 similar to plant disease resistance response protein family like protein, mRNA sequence

GCGTGGTCGCGGCCGAGGTACTTTGGTGGTGAACTAGTATTAATTAAATTATTATTAAAGCTGTGAGAATAAATAAAAGAATGTCCTATATATATAATATTTATGCAAAGCAAAAGTCATTGAGCTAATGACTTGTGTTTTTCTTTTGTTTTCTAGCTATCTTTCTTGGCAATATTTGATTATTTGTAATGAGTTTGAAATATATATATGTATGTAGTGAAAGAATAATGAAGTTTGTATTGTAATTATATATTGATCTCTTTAGCCTTTCCCTTGCCGAAAAAAAAAAAAAAAAAAAAAAAAAAAGCTTGTACCTGCCCGGGCGGCCGCTCG

>lcl|Query_207005 GH985240.1 JutEST5244 Jute Plasmid library 20-45 days after sowing Corchoruscapsularis cDNA, mRNA sequence

GTTGAGGAACGAGTTTGGAAGTTGCTGATGGGATGGGAATAAGTATAATCTCAACATTTGTTCCTCCCGGTTGGCTATGGCCCCTGTACAGTATACTTTGCCCTGCCACGCAATGTAGTTGATGTAAATTACGGCTCACCTGCAACTAAAATTTCCCATCTTACCATTCCAGCTTCGATCCTGTGAGGCTGACCTTGTGATATAGAGGTGAGTTGGTAGTAGCAAAGAAGGTAAAAGTAATAGTGAAGCTCTTATGGCAATGTATACATCAAAGTTGTAGCATAGAGGTAGATATCATGCATATCCTTGTTTTGTAAAGCATCTTTCATGATGGACCCTTTGGTGGGTCAGTTGGGTCTAATCAAGCAATCCGACGAAATTTCGACATCAATTCTCATTTCCCTGTACAGAAAAATGCAGAACACATGTAGAAAAGAATAGCATTTTCCAGTTTCCGCATATTATCAATATTATGCCCTCA

>lcl|Query_151484 FK826464.1 JutEST44132 Jute Plasmid library 20-45 days after sowing Corchoruscapsularis cDNA similar to Glycosyl transferase family 2, mRNA sequence

GGTGAATTGATTCCAATCATCTGAGATGATCATAATCTCAGTGCATCAAATTGATGGCACAATTATTTTCTCCCTATACAAATGATGGCACAATTGGGAATTTATCTTGTCTACATGAGCCTTCCTGACGTATTTATCTAGTTATAGTTGTATAAATAGGCCTGTTAATATAGGGCAACATTAGCTTAGGTAGAATCAGAAATGCCATTGCAGTCAACCAGATTACCAGAAGACAAAAGTTTTGTATCTAAATATTTATATTACTGTCATTTTTAACTGTTATAGAAAGGTATATATATAAGTAAATTTT

>lcl|Query_230492 FK826548.1 JutEST4284 Jute Plasmid library 20-45 days after sowing Corchoruscapsularis cDNA similar to Acyl carrier protein, mRNA sequence

CGGATACTTCTCTTTCTTTTTCGCTCTTTATTCCTCTCACCATCACTAGAAGACAATCAAGAATCAGAAAACATATGAACAAAAACAACAATGAAGCAAAAAATAGATGTGATAGTAAAAAAGAACATTGACTGAGAAAGTAGCAAAGACAAATTTTTAATACTGCATATTCAGACAAAAGCCTCTTACCAGATACACTTAAGAACAAAAGTACTATACCTAGGTTGATACATTGGGGATCCGGCTTCTTCCTGTTTTTGATTCTCTCTTCC

>lcl|Query_151255 FK826459.1 JutEST44107 Jute Plasmid library 20-45 days after sowing Corchoruscapsularis cDNA similar to AXS2 (UDP-D-APIOSE/UDP-D-XYLOSE SYNTHASE 2), mRNA sequence

GGCTGTTGCCTCTTAAGAAGCTGAAGACGTTCATGACTACTCGTAAGAAAGGGATATATCAAGATGTGCACCTTTTTAGAAAAGTGTTTCAATTCTTTTTAAGGCAGTCTTAAATATATCTTAATTGTTGATTCCTTATACACTAAAATTTTTGTCTTCCCTGTGTCGGTAGTCAATAGCTCGATGTAGCTCAGGATGATGTGTTCTCAGTATGGATATTATGTGGAAACAAATAGATTCTACTTAAGGGTGGTATTTGTGATAGTATTATAAATATTCCAATGGAGCTTTTGGGCTACAGCATGTTGGGGGTATATTATATAGAGGAAAGGAAAGCTGTTA

>lcl|Query_80878 GR463692.1 JuteSSHPC63 JuteSSHlibMP1 Corchorus capsularis cDNA clone JuteSSHMRP2 similar to multidrug resistance protein like protein, mRNA sequence

AGCGTGGTCGCGGCCGAGGTGTAACAGAGACCAGGTTTATCCTCCCAGTTGAAACAGCTAAAAAACTGGAATGAATACATGAGACAAGAGCTCTTAGATGTTGGACATCAGGCAATAAAGGACACTGATCCTTGAGGATTAGGAAACAAATGAGTTTAGCCCTACGGTCTCCCCAGCTGACTGCCTGGAGAGCGTTTCCAGACCACAGCACAGGGAAGGAAAACTGAGGTGGGACCAGGAGATTCCCTAAGTTGAAGAGACAGAGCTGGAAATCCAGGGAAGCCATGGCAGGCAGGGTTCACGTTGCAGAGCACAGAGACACAGTTGAGAGATCACAGAGCAAAGCCCAGAGGTATGCAGAGAATCCTTGAACATGCAGTTGAGTTCTGCTCAGAGAATACAGACAAGGAAACTACCCTAGGGCTAGGAAAGAACCACCCAAAAGGAGAAGAGGGAATCATACCTGGAAACACACAGGGCCAGGAATAGTGTCTATTCCTGACAGGTGAATAAACCTCAGAATTCACAGGGCATTGTTTAGCGTACCTGCCCGGGCGGCCGCTCGA

>lcl|Query_11971 JK743800.1 JutESTSSH3(N43) Jute Plasmid Suppression Subtractive cDNA library Corchorus capsularis cDNA, mRNA sequence

CCGGCCCCCCTTTTTTCCCTCTCTCCGGGGGGGGGGGGGGGGCCCCCCCCCCCCCCCCCTCTCCTTCTTTCCAAACAGGTGGCTGGTGGGGGGGGGACAAAAAAATTACCCCCCAAATTGGTTTCCCCCTTGCGTTGTTTAGGCGGGGTGGGTCTCCACCTTTTAGACGTAGGGTTCGCCTCCCATTCCGAGTTATTTCCCTGAAAAAAGGGGGGGTTGTGTGGGGGGGGGTGACCGGGAAACTAGGGAACCTAACAAAAAGGCGGGCGACTGGGGACAGCGGGGGTATCGGTCGTAAACGGGCCCCGTGGGTAAATGGCCGGCCCCCCCCCCCTTTTTTTCGGGGTTTTGGGGGGGGGGGGGGCGCCGAGAAACCCCCCCCCTGGAAACCCCCCCCCCCAAATACATTTTTTTTGGGGGGGAGGCGGGAGAACGGGGGCAATGTTAGGGGAAGCCACAGAGAGGGAAGAGCGCGCGTTGGGGGGGGGGGGGTGAGGATAGCTTGGACATATTTGTCATAAAACAGCGACACTATCCCCCCGCCAACCGGCGCCGGGGTTAAGGGGGGGGGAGTAGTGCGGCGGGTAACCGGCCGTATATTTCTCCCGCCACGCCCCCACGCTATTATTCCTGGTCGTTAGATCGTTTTATATGTGGGCTCACTTGTCGGGGGAGAATGCAGGCACGAAAGAACACACGAGATGATGATACTTTTCTTTTCTCTATAGATGACGAAAGGGAGGAGGGGTGCGGGACTGTCTAGTGAGTGTGCCTCCCCCCCGCCGACCTGTTGTCCCTCCCTCGTTTATCTGGGCCGTAGATATGGCCTTCATGAAG

>lcl|Query_148715 FK826411.1 JutEST4333 Jute Plasmid library 20-45 days after sowing Corchorus capsularis cDNA similar to Alpha-expansin 1, complete, mRNA sequence

GGGATTCACGCCGCTGCTTCTGTAGAACAACACTAACAAGTGATTTGGTACTGCAAAAGGAGACCCTTTTCGAAGACTTTTCACCATGAGAGCCAAGTGGAAGAAGAAGCGTATGAGAAGGTTGAAGAGGAAGAGAAGAAAGATGAGACAAAGATCTAAGTAGACTAACTTTACTTTAGAGCAATTATGCCCGTTTAGATTTCTTAGTTTGTTTGTTGTTGATAGATCCTTTATGTTAAAAGGATCGAATTTATCATTCTTGAAACCTGTTTAGTATTGTGAACACTTGTTAACCACTTTTCTATTTACTGGGAAAGCTTATGGATCTTAATTTTCTTTTGGATCTG

>lcl|Query_24568 FK826536.1 JutEST4016 Jute Plasmid library 20-45 days after sowing Corchorus capsularis cDNA similar to Bimodular protein, mRNA sequence

CTGTAGTTTCAAATATTAATGTATTTTTGTTTAGTGTTTGGTCGATTTGAAGTTGGGATCTAGGGTTTCCAGTTGGTTTTACATATTTTATAGGAGTCTTTTTACAAAGACTTCCGACAATTGTCGTTGTAATATTTGTTCATGTTTAATAACTAGGCTTGTGGGTTAATTTACCTCGGCCTAATGTAATGTTTAATTAATGTTGTTTGTTCTCAATTAATGTCAAGTTTGATTTTGCTTGA

>lcl|Query_49587 FK826573.1 JutEST45123 Jute Plasmid library 20-45 days after sowing Corchorus capsularis cDNA similar to ribosomal protein 41, mRNA sequence

ATTCTCCAAGCGAATCGTCTCCTTTCTAGCGAAATCGCCTGAAGAATCTCGAGAAAGAAAGCGCTAACCCAAGATGAGAGCCAAGTGGAAGAAGAAGCGTATGAGGAGGCTGAAGAGGAAGCGCCGAAAGATGAGACAGAGATCCAAGTAGGCGCATGGACACAGTCAAGATGAGCTTTACTGTGACATTCCCTTGTCTTTGCCACCGATCCAGCGTTTGTTGCATGAAGTTGTGAAGATTTGAGGATGTTTTTTCGGGGAAGCTTATCTTTTATGTCTTCTAAATCTAGGTTTAAGTTTCAAGACTATATGAGATGTTCTCTAGTACACTACTATAAATCTTGTTGGTATTTCTGTTTTTGAGATGTGTAGACTATTATCCTTAACATTGCATCCTAAGAGTTTGATATTGGAAACCTTATGGTCTTTCTAGT

>lcl|Query_73943 GH985201.1 JutEST5181 Jute Plasmid library 20-45 days after sowing Corchorus capsularis cDNA similar to S-adenosylmethionine synthetase 2, mRNA sequence

GGGGTATCAGTGTCGGTGATGTGAAAAATGTATGAGAATTTTGCTTTAATAGATGGCACTGAAAGTAACAAGTGCCTCTTTAGATTTCTTGTGAAATTAGCTTTGCAAAGGCAGCAGCAATATACTTATGTGTAATTGTACTTTGGATTTGGGATAATCAAATTTATTTTAC

>lcl|Query_156057 FK826505.1 JutEST4737 Jute Plasmid library 20-45 days after sowing Corchorus capsularis cDNA similar to ATP binding, mRNA sequence

GGGTTCAGTCCGTTCTGGACTCTTTTTTCATGAAATAAATTAATTTCTGTCTCAAGCTGCTTCTTCTTCTTCTTCTCCCCGGTTTTCCTTCTCTTAAATCTGCCGATTTGAATCTCGCAACGACGCCGTTTTAGTTATAATCTCCAGAAAGATTTTATAGATTCCGTCAATCAAAAACCATTTCCGACGAGAA

>lcl|Query_128819 GR463655.1 JuteSSHRP17 JuteSSHlibMP1 Corchorus capsularis cDNA clone JuteSSHPp1 similar to protein phosphatase 1, regulatory (inhibitor) subunit 1C like protein, mRNA sequence

GTATGGTAGTAGTGACGACGTGGATGAAATCTATCCTGTCTACTGTCGTGGTGCCGGTGTTGAGTTGCTGTACTGCGTGGATGAGTGGGGGCTGATGTATGGGGAGCGAGAGCATGCGCGGAGAGTGTGAGTACGTGTGTATGAGTAGGACAGTACTGATGCTGTATATAGTAACGATATATATATCAGATAGAGAAATACACAAAGTATGTAGATACGATGGCATTGGAGAGTGGTACGTGTTGATATTTCTCTTCGTGAGTACTCGATTGAAGTAATCTTATGACGGACATACTATATGACTACTAATCTATTGATTCTTGATCTTGTTGTGCCCCGCCGCAGTCCCCCTTATGGGAAAATATATTGGTAGAGGGAT

>lcl|Query_226921 FK826435.1 JutEST41109 Jute Plasmid library 20-45 days after sowing Corchorus capsularis cDNA similar to Proline-rich protein APG-like; GDSL-motif lipase/hydrolase-like protein, mRNA sequence

GTACTAAGCTCTGTCACTACTCCTTTTTCTATTATCTTAATCAAAAAGTGACTCGTATTGTAGAAGCTTGTATTTGTAGTCAAACCTTCCTTAAAACTCATCGTATATTTCTCATCAGATTATTTACAATTCCTAAGTGACATATAGGTTAAAGAAATTGTGATACAGCTACATCTTTAAGTTATAAATTGAGGATAATGAAAAT

>lcl|Query_79168 JK743805.1 JutESTSSH3(N48B) Jute Plasmid Suppression Subtractive cDNA library Corchorus capsularis cDNA similar to MAP kinase kinase kinase Ste11/SteC, mRNA sequence

AAGCAAACAAAGGCAAAACCAGAAACCGTGCTCGTTCCCTCCCTTAATCTGCATGTAAACGCATCTGAGGTTGTTCAAGTTCATATTAATTTGGCCCTCCAGCAGGCAACCAAAAAAATCAGCCGAACCTTG

>lcl|Query_37800 GR463654.1 JuteSSHRP3 JuteSSHlibMP1 Corchorus capsularis cDNA clone JuteSSHPS similar to 23S rRNA pseudouridine synthase like protein, mRNA sequence

GCGTGGTCGCGGCCGAGGTTTTTTTTTTTTCATATATAATTACAATACAAACTTCATTATTCTTTCACTACATACATATATATATTTCAAACTCATTACAAATAATCAAATATTGCCAAGAAAGATAGCTAGAAAACAAAAGAAAAACACAAGTCATTAGCTCAATGACTTTTGCTTTGCATAAATATTATATATATAGGACATTCTTTTATTTATTCTCACAGCTTTAATAATAATTTAATTAATACTAGTTCACCACCAAAGTACCTGCCCGGGCGGCCGCTCGA

>lcl|Query_50334 FK826518.1 JutEST47139 Jute Plasmid library 20-45 days after sowing Corchorus capsularis cDNA similar to Aux/IAA protein, mRNA sequence

AAATTATAAAACCATCAAGGCCTAGTGTTTGTGAGTAACCTTTTTACTATTGAATCAAGTTTTAGAGTCTCTGGACAACTCTCATGTTCCCACGCAGGATCTCTAGTTTGTAAAGTGCATTGTTTGTTAAAAACTTTCATCGGTGTGTGTTTGTGTGATGAAAGACCTCGACATTATCTGTAACCCCGACATGATTGTATTAACTCTATTGTGCGGCTGCTTGTGAGTGTTGTATTTCTTCTGCTTAAGTATTATTCTGCAAATCATTTTAGTTCCTATTCTGCTAACTATAACCCCAATTCCGAAAACCGGTCGGGGGCTTTGAAAATGCATATGTACTGAATCATATTAGTAGTAGTTTTATAATGTGTTTATGAGTTCCAAAAAAA

>lcl|Query_7089 FK826440.1 JutEST4426 Jute Plasmid library 20-45 days after sowing Corchorus capsularis cDNA similar to Chalcone synthase 2, mRNA sequence

GGGACTTAGTTATTAATTAAATTGAGTATCAAATTAATAAGCTTATAATAACTTATATTACTTTTACTAAAAATTATCTTTGTATTCAATTGAACTTAAACTTTGTATTTCATATAGTATCCAATGAAAGTTCATTTCCCCTTTGATGATCTCTTGAC
